# Plasticity and adaptation to high light intensity amplify the advantage of amphistomatous leaves

**DOI:** 10.1101/2025.08.19.671015

**Authors:** Christopher D. Muir, Wei Shen Lim, Dachuan Wang

## Abstract

The presence of stomata on both leaf surfaces (amphistomy) increases photosynthesis by reducing the distance for CO_2_ diffusion between stomata and chloroplasts. Paradoxically, most leaves are hypostomatous (stomata on lower surface), despite the photosynthetic advantage of amphistomy. Across 29 diverse populations of “wild tomatoes”, leaves developed under high light intensity benefit more from amphistomy in terms of CO_2_ assimilated for a given stomatal conductance than plants developed under low light. Furthermore, populations native to open habitats benefit more from amphistomy than those from more closed habitats. Thus, plasticity and adaptation together may explain why amphistomatous leaves are prevalent in sunny, open habitats, including many crops. Contrary to common assumptions, amphistomy can save water because hypostomatous leaves evaporate 10-65% more to achieve the same photosynthetic rate.

## Main text

Stomata are microscopic pores on the surfaces of leaves and other photosynthetic organs formed by a pair of guard cells. They are essential for balancing carbon gained per unit water lost and were an essential innovation in vascular plants, permitting them to grow tall on land by enabling access to CO_2_ for photosynthetic carbon assimilation while preventing hydraulic failure in variable environments (*1*–*3*). Optimal stomatal function depends on both dynamic changes in aperture on the scale of minutes to hours, as well as static anatomy determined by developmental plasticity and constitutive genetic differences (*4–6*). Understanding how stomata respond to environmental change over daily, developmental, and evolutionary time is important for studying adaptation (*1*, *7*, *8*), inferring paleoclimate from fossil cuticles (*9–11*), predicting responses to climate change (*12–15*), and improving crops (*16*). Stomatal function contributes to global carbon and water cycles (*17*) and therefore to predicting future climate (*18*).

Despite extensive theoretical and empirical progress understanding stomata function and anatomy from molecular to ecosystem levels, the adaptive significance of amphistomatous leaves remains an important unsolved problem in leaf structure-function relationships (*19–26*). Amphistomatous leaves develop abaxial and adaxial stomata whose aperture can be independently regulated (*27–31*) to control gas exchange through each surface. All else being equal, simultaneous gas exchange through stomata on both surfaces increases CO_2_ supply to chloroplasts by providing a second parallel pathway through leaf intercellular airspaces, enhancing photosynthesis (*20*, *32*). The extent to which amphistomy increases CO_2_ supply depends on resistance to diffusion in intercellular airspaces. This resistance can be low in thin, porous, amphistomatous leaves (*28*, *33*), but may be more substantial in thick, dense, hypostomatous leaves (*34*). We refer to the intercellular airspace conductance (*g*_ias_), the inverse of resistance. Amphistomatous leaves also lose more water through evaporation because of a second boundary layer conductance (*35*), but the additional carbon gain should be enough to offset this cost in most realistic scenarios (*36*).

The paradoxical fact is that, despite the photosynthetic benefit, most leaves are not amphistomatous. Many vertically oriented and/or isobilateral leaves are amphistomatous (*25*). But among dorsiventral leaves, it is primarily herbaceous plants in open, high light habitats that tend to have amphistomatous leaves (*22*, *39–44*). Most other leaves, except those from aquatic habitats, are hypostomatous, producing stomata only on the lower, abaxial surface. Even resupinate leaves develop stomata on the lower, albeit adaxial surface (*45*), suggesting that leaf orientation (lower vs. upper) rather than leaf polarity (abaxial vs. adaxial) is causal. Stomatal density ratio, defined as the ratio of upper to total stomatal density, is a quantitative metric of stomatal distribution. The covariation between stomatal density ratio and light habitat is both qualitative and quantitative. A higher proportion of sun leaves are amphistomatous (*39*) and the proportion of stomata on the upper, adaxial surface increases with light (*42*, *43*). Resolving why high light intensity favors amphistomatous dorsivental leaves is an important first step toward understanding variation in stomatal density ratio and leaf structure-function relationships more generally.

The overarching hypothesis is that leaves with greater stomatal density ratio are more common in open, sunny habitats because they increase photosynthesis most in those circumstances. An amphistomatous leaf increases photosynthetic carbon gain compared to an otherwise identical hypostomatous leaf by increasing conductance through the leaf intercellular airspaces and boundary layers. We quantify this benefit as the amphistomy advantage (AA), the log-response ratio of photosynthesis in an amphistomatous leaf compared to an otherwise identical pseudohypostomatous leaf (*20*, *46*). Why would AA be greater in sun than shade? We consider three nonmutually exclusive hypotheses that we classify as ‘acclimation’, ‘plasticity’, and ‘constitutive’ (Fig. 1).

**Figure 1:**
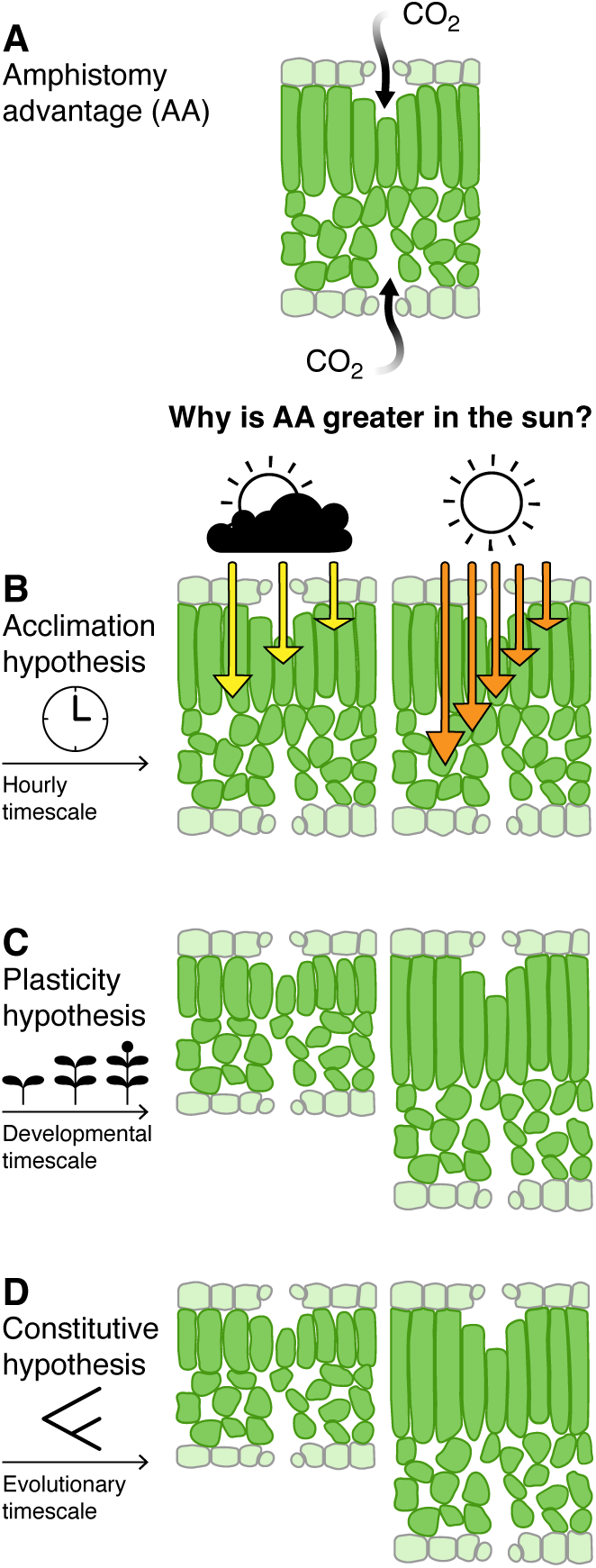
Conceptual outline of three nonmutually exclusive hypotheses explaining why amphistomy advantage (AA) might be greater for leaves in sunny, open habitats. The acclimation hypothesis predicts that AA is greater under high measurement light intensity (PPFD = 2000 µmol m^−2^ s^−1^) than low measurement light intensity (PPFD = 150 µmol m^−2^ s^−1^), regardless of growth light intensity or native plant area index (PAI). The plasticity hypothesis predicts that AA is greater for plants grown in sun than shade, regardless of measurement light intensity or native PAI. The constitutive hypothesis predicts that AA is greater for plants adapted to sunny habitats (lower native PAI) than shaded habitats, regardless of measurement light intensity or growth light intensity.

### Acclimation hypothesis

Photosynthetic induction to high light intensity typically involves increases in total leaf stomatal conductance (increased CO_2_ supply), the concentration of active Rubisco, and electron transport capacity (increased CO_2_ demand). If the acclimation hypothesis is correct, we predict that AA_2000_ > AA_150_ for all populations regardless of native habitat or growth environment. AA_2000_ is the AA measured under high light intensity (PPFD = 2000 µmol m^−2^ s^−1^); AA_150_ is the AA measured under low light intensity (PPFD = 150 µmol m^−2^ s^−1^).

**Table 1:** Three nonmutually exclusive hypotheses and directional predictions explaining why amphistomy advantage (AA) might be greater for leaves in sunny, open habitats. For each hypothesis, we make predictions for how measurement light intensity (PPFD = 150 µmol m^−2^ s^−1^ vs. 2000 µmol m^−2^ s^−1^), growth light intensity (sun vs. shade), and native plant area index (PAI) would affect AA. PPFD: photosynthetic photon flux density.

| Hypothesis | Measurement light intensity | Growth light intensity | Native PAI |
| --- | --- | --- | --- |
| acclimation | $AA_{2000} > AA_{150}$ | $AA_{\text{sun}} = AA_{\text{shade}}$ | $\text{cor}(\text{PAI}, AA) = 0$ |
| plastic | $AA_{2000} = AA_{150}$ | $AA_{\text{sun}} > AA_{\text{shade}}$ | $\text{cor}(\text{PAI}, AA) = 0$ |
| constitutive | $AA_{2000} = AA_{150}$ | $AA_{\text{sun}} = AA_{\text{shade}}$ | $\text{cor}(\text{PAI}, AA) < 0$ |

### Plasticity hypothesis

Individuals of the same genotype often develop dramatically different leaves in sun and shade conditions (*47*). Plastic responses are likely adaptations to optimize photosynthesis at different light intensities in variable environments (*48*). Plastic changes in leaf anatomy and biochemistry could modulate AA as a byproduct. Thicker or less porous leaves, both of which are associated with high leaf mass per area (LMA), will have lower *g*_ias_; leaves with increased total stomatal density and photosynthetic capacity have greater potential CO_2_ supply and demand. Under the plasticity hypothesis, we predict that AA_sun_ > AA_shade_ for all populations and light intensities. AA_sun_ is the AA measured on sun leaves; AA_shade_ is the AA measured on shade leaves.

### Constitutive hypothesis

In environments that are relatively constant or where environmental change cannot be anticipated by a reliable cue, natural selection will favor constitutive expression of optimal phenotypes. We therefore predict genotypes from more sunny, open habitats will have consistently greater AA under all measurement and growth light intensities. For herbaceous plants, light intensity is largely a function of the tree canopy (*49*). Herbs growing in the open will regularly experience high light intensity; herbs growing under a forest canopy will often experience low light intensity.

The primary directional predictions for each hypothesis are summarized in Table 1; detailed predictions for results that would indicate simultaneous support for multiple hypotheses are in Table S3.

We tested these hypotheses by comparing AA among amphistomatous wild tomato species (Table S1; Figure S2; (*50*)) from different native light habitats, grown under simulated sun and shade light treatments, and measured under contrasting light intensities (low and high). We measured AA on 572 individual plants from 29 populations (average of 9.86 replicates per light treatment) using a recently developed method (*46*). With this method, we directly compare the photosynthetic rate of an untreated amphistomatous leaf to that of the same leaf with gas exchange blocked through the adaxial (upper) surface by transparent plastic, which we refer to as ‘pseudohypostomy’. To compare amphi- and pseudohypostomatous leaves at identical whole-leaf stomatal conductance (*g*_sw_), we measure *A* over a range of *g*_sw_, inducing stomatal opening and closure by modulating humidity (see supplementary text for further details). We estimated ‘amphistomy advantage’ (AA) *sensu* (*20*), but with modifications previously described in (*46*) and here (Supplementary Materials). The native light intensity was represented by plant area index (PAI m^2^ m^−2^), estimated using a global gridded data set derived from the Global Ecosystem Dynamics Investigation [GEDI; (*51*)] and georeferenced accession collection information from the Tomato Genetics Resource Center (Table S1). The growth light intensities were PPFD = 761 µmol m^−2^ s^−1^ (sun treatment) and 115 µmol m^−2^ s^−1^ (shade treatment) while all other environment conditions were nearly identical (see Materials and Methods). The high and low measurement light intensities were PPFD = 2000 µmol m^−2^ s^−1^ (97.8:2.24 red:blue) and PPFD = 150 µmol m^−2^ s^−1^ (87.0:13.0 red:blue), respectively.

Consistent with biophysical theory of CO_2_ diffusion within leaves, AA > 0 for all populations (Fig. 2A). Bayesian phylogenetic mixed effects models that allowed AA to vary between measurement light intensities, growth light intensities, and among populations outperformed simpler models based on information criteria (Table S6). Measured under high light intensity, AA was consistently greater for sun plants. The average AA among populations in the shade treatment was 0.041 (range: 0.007–0.113; 19 of 29 populations significant); however, the same populations grown at high light intensity showed a mean AA of 0.052 (range: 0.020–0.120; 20 of29 populations significant). Contrary to the predictions of the acclimation hypothesis, AA was greater in all populations under low measurement light intensity for both sun and shade grown plants. The overall average AA of shade and sun grown plants measured under low light intensity was 0.064 (range: 0.022–0.137; 28 of29 populations significant) and 0.100 (range: 0.049–0.206; 27 of29 populations significant), respectively. There was a modest tendency for populations from more open habitats (lower PAI) to exhibit greater AA and the slope was significantly different than 0 in 3 of the 4 treatment combinations (Fig. 2B).

**Figure 2:**
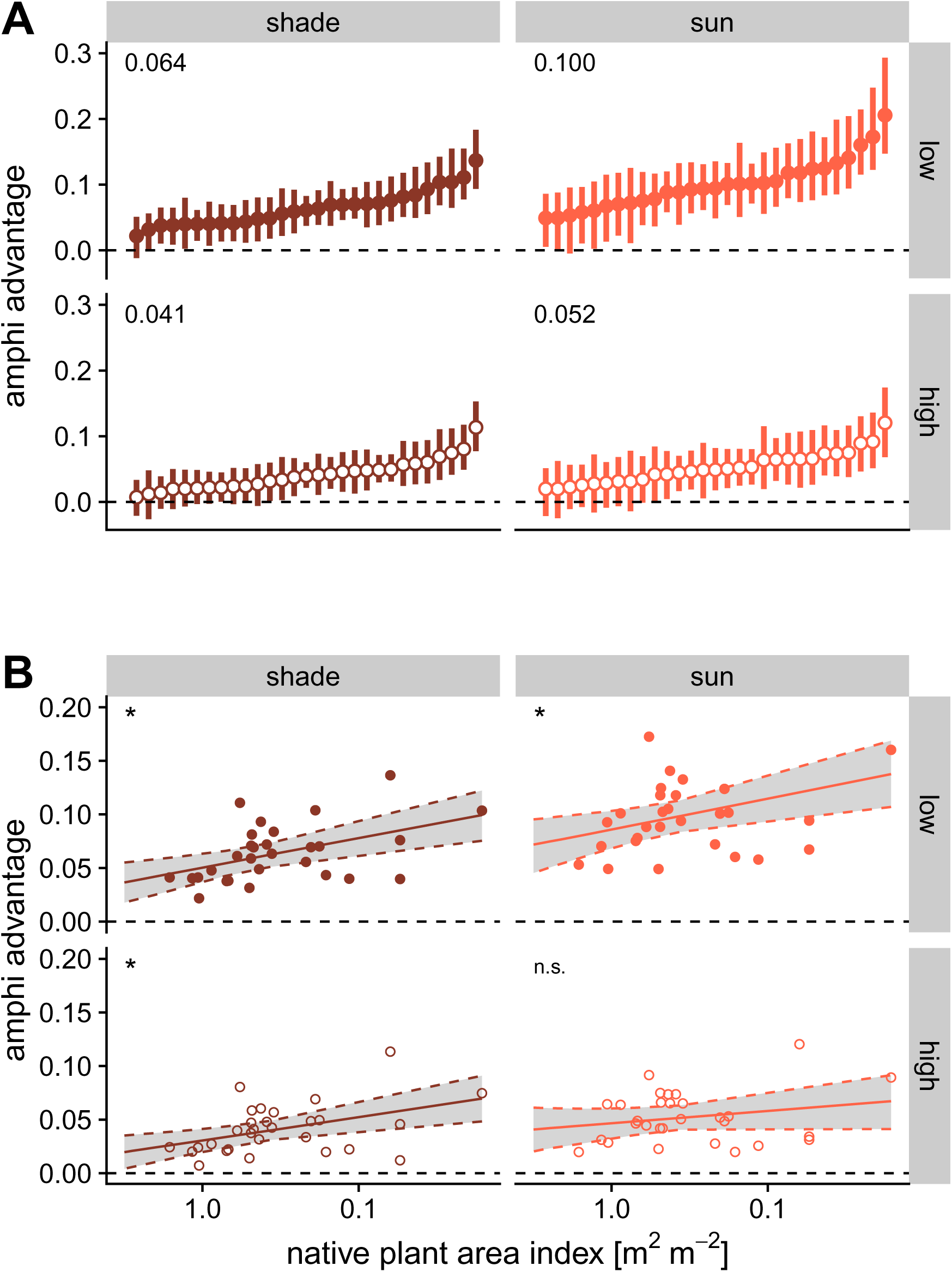
Amphistomy advantage (AA) is greater in wild tomatoes from open habitats and is further amplified by developmental plasticity under sunny growth conditions. AA *(y-* axis) is the log-response ratio of photosynthesis in an amphistomatous leaf compared to an otherwise identical pseudohypostomatous leaf. In both panels, estimates are shown for plants grown under simulated shade (brown) or sun (orange) and measured under high light intensity (PPFD = 2000 µmolm^−2^ s^−1^; open circles) and low light intensity (PPFD = 150 µmolm^−2^ s^−1^; solid circles). The dashed line at zero indicates no difference in photo-synthesis between amphi- and pseudohypostomatous leaves. (**A**) The points are the posterior median AA for each population, with error bars showing 95% confidence intervals. Within each facet, the populations are arranged by AA estimated in that growth and measurement condition. The average AA among populations for each treatment group is written in the top left corner. (**B**) The same estimates of AA in each combination of growth and measurement light intensity plotted against native plant area index (PAI; r-axis, log-scale). The confidence intervals were omitted for visual clarity. The median relationship between native PAI (r-axis; log-scale) and AA is shown as a solid line with the shaded region between dashed lines show-ing the 95% confidence ribbon. The * in the top left corner of each facet indicates the slope of the linear regression is significantly different from zero; n.s. indicates not significant.

The pattern of AA across wild tomatoes strongly supports the plasticity hypothesis, contradicts the acclimation hypothesis, and provides modest support for the constitutive hypothesis. Plastic changes in leaf thickness and/or packing density, summarized by the bulk leaf mass per area (LMA), may mediate the effect of growth light intensity on AA. LMA increased in sun grown plants in all populations by an average of 123% [95% CI: 42.9 to 256%], qualitatively similar to plastic responses in many species (*47*). While LMA is weakly, albeit significantly (Table S5), associated with individual-level AA (Fig. 3), the effect of growth light intensity on AA is still predictive based on model comparison using information criteria (Table S6). However, the direct effect of the sun treatment on AA was weaker when LMA was included in the model (Table S4 vs. Table S5), suggestive of a mediating role. Many anatomical traits underlie LMA (*52*) and future research will be needed to identify which particular traits, such as leaf thickness or mesophyll porosity, are responsible for mediating AA. The fact that the AA of sun plants was greater under low measurement light intensity supports a long-standing hypothesis that resistance to CO_2_ diffusion is greater in the upper than lower portions of the leaf interior (*53*). At low light intensity, photosynthesis is weighted toward the upper palisade where most light is intercepted. If resistance to CO_2_ diffusion is high in the upper palisade, amphistomy may, unexpectedly, be particularly beneficial for sun leaves experiencing intermittent shade or cloud cover. Our study is limited in testing this because we could not directly measure the stomatal conductance ratio and intercellular resistance on each surface. Future experiments measuring AA with a dual sided chamber (*31*, *33*) can overcome these limitations.

**Figure 3:**
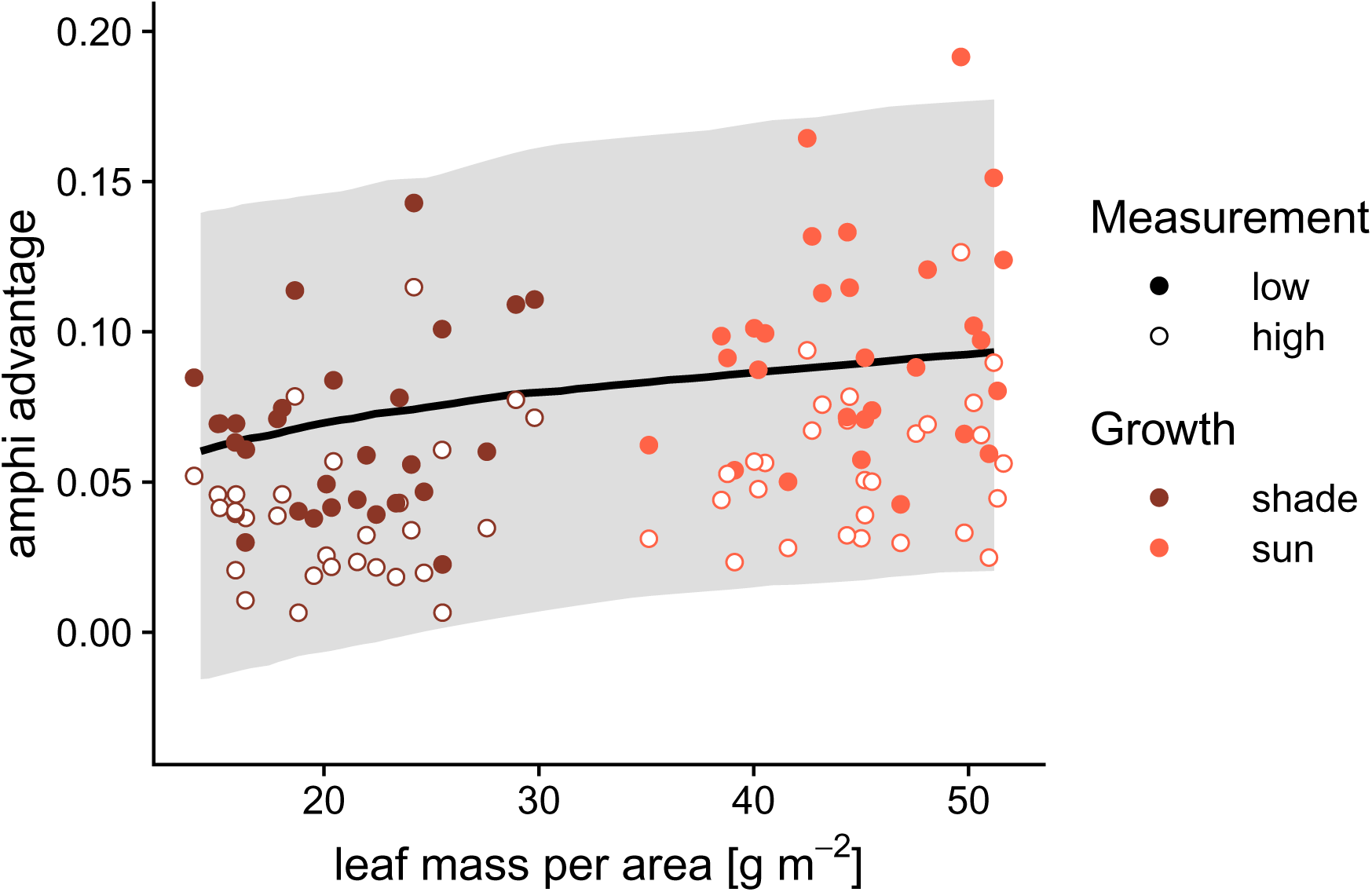
Plastic changes in leaf mass per area (LMA) partially mediate the effect of sun-to shade-grown plants on amphistomy advantage (AA) Each point is the estimated LMA (re-axis) and AA (y-axis) for each population grown under high light intensity (PPFD = 2000 µmol m^−2^ s^−1^) and low light intensity (PPFD = 150 µmol m^−2^ s^−1^). The points are colored by growth light treatment. The solid line is the posterior median of the linear regres-sion of individual-level LMA on AA_2000_ with the shaded region showing the 95% confidence ribbon; the dashed line is the same for AA_150_. Confidence intervals for each population esti-mate were omitted for visual clarity.

We conclude that both developmental plasticity and adaptation to open habitats, but not acclimation, may explain the long-standing observation that amphistomatous leaves are more common in sunny habitats, at least among herbaceous plants including crop relatives. This result changes our understanding by showing that high light intensity *per se* does not increase the benefit of amphistomy. Instead, the benefit increases as a byproduct of anatomical and biochemical changes caused by plasticity and adaptation to higher light intensity. To gain more precise understanding of when amphistomy is most beneficial will require further research on its leaf anatomical and biochemical basis.

The magnitude of AA we document across populations is also noteworthy, as it implies that hypostomatous leaves predominant in mesic to wet forests globally are giving up ‘free’ carbon that would require little to no additional water loss. The cost of hypostomy can be quantified as the difference in total *g*_sw_ that an amphistomatous leaf would require to achieve the same photosynthetic rate. This cost can be locally approximated on a log-ratio scale as *AA/ε_g_* (Supporting Information), where *ε_g_* is the elasticity of *A* to *g*_sw_. For AA = 0.05, the water cost of hypostomy would be 0.5 (64.9%) at low elasticities *(ε_g_* = 0.1) and 0.1 (10.5%) at high elasticities *(ε_g_* = 0.5). For plants to expend this much extra water implies a large fitness cost of upper stomata in certain ecological contexts. As this is the first quantitative comparison ofAA across species, estimates from a broader range of species and more stressful environments will be required to fully understand the ecological causes and consequences of stomatal distributions on leaf surfaces.

## Acknowledgements

Sam McKlin and Tom Buckley helped with protocol development. Justin Alter, Max Gatlin, Joana Kim, Jenna Matsuyama, Brandon Najarian, and Kai Yasuda contributed to data collection. Kate McCulloh provided feedback on an earlier version of this manuscript. Sarah Friedrich helped with graphics. We used Copilot and ChatGPT to assist with coding.

## Author contributions

Conceptualization: C.D.M.; Methodology: C.D.M., W.S.L.; Investigation: C.D.M., W.S.L., D.W.; Visualization: C.D.M.; Funding acquisition: C.D.M.; Writing – original draft: C.D.M.; Writing – review & editing: C.D.M., W.S.L., D.W.

## Supplementary Materials

### Materials and Methods

#### Populations

We compared AA among 29 ecologically diverse populations of wild tomato, including representatives of all described species of *Solanum* sect. *Lycopersicon* and sect. *Lycopersicoides* (*50*) and the cultivated tomato *S. lycopersicum* var. *lycopersicum* (Table S1). These populations were selected in a previous study because they encompass the breadth of climatic variation in the wild tomato clade (*54*). In one case, we substituted a congeric population of the species (*S. galapagense*) because of difficult growing the focal population. We also added an a population of *S. pennellii*. Due to constraints on growth space and time, we spread out measurements over 61.1 weeks from August 29, 2022 to October 31, 2023. Replicates within population were evenly spread out over this period to prevent confounding of temporal variation in growth conditions with variation among populations.

**Table S1:**
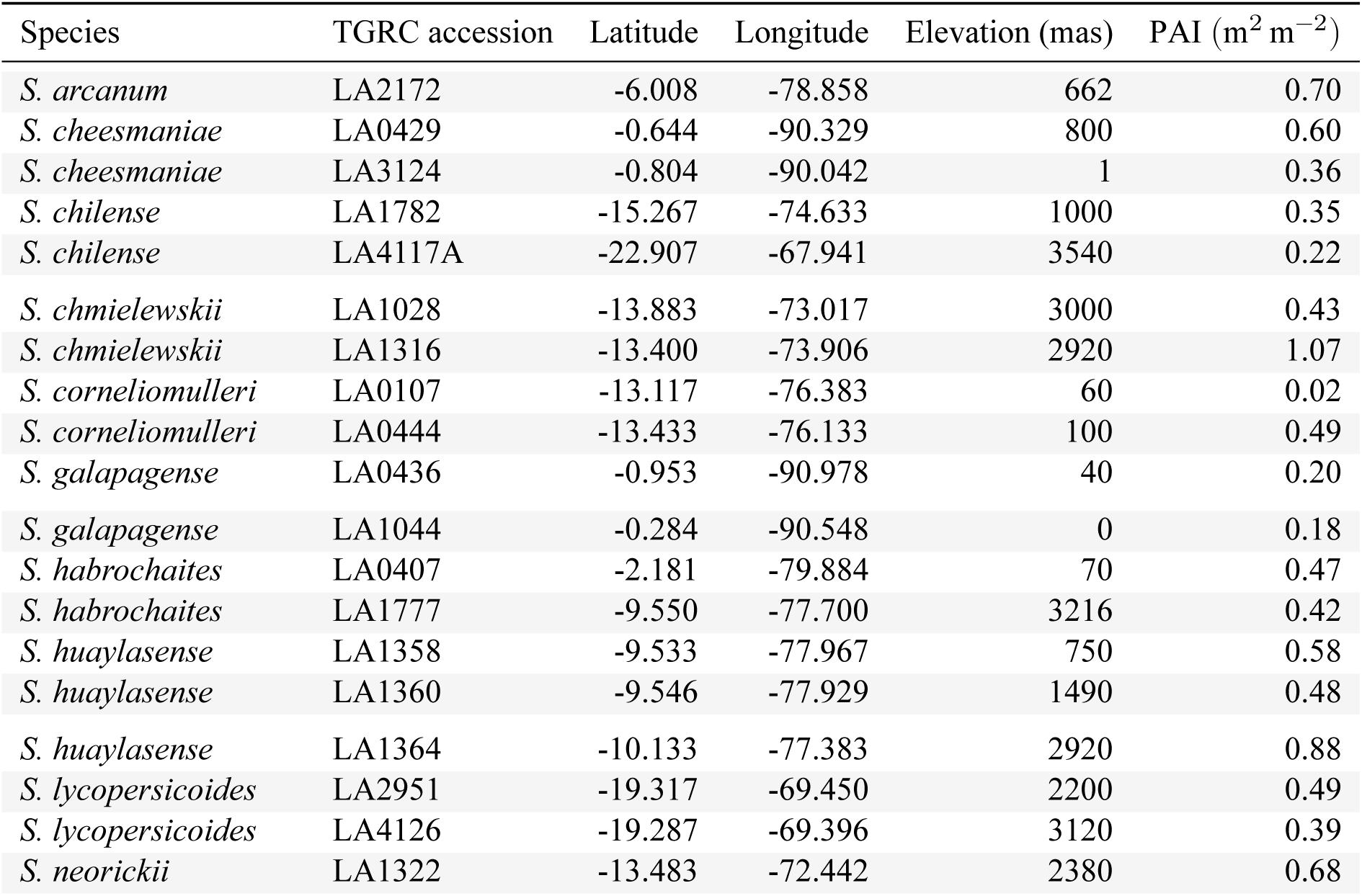
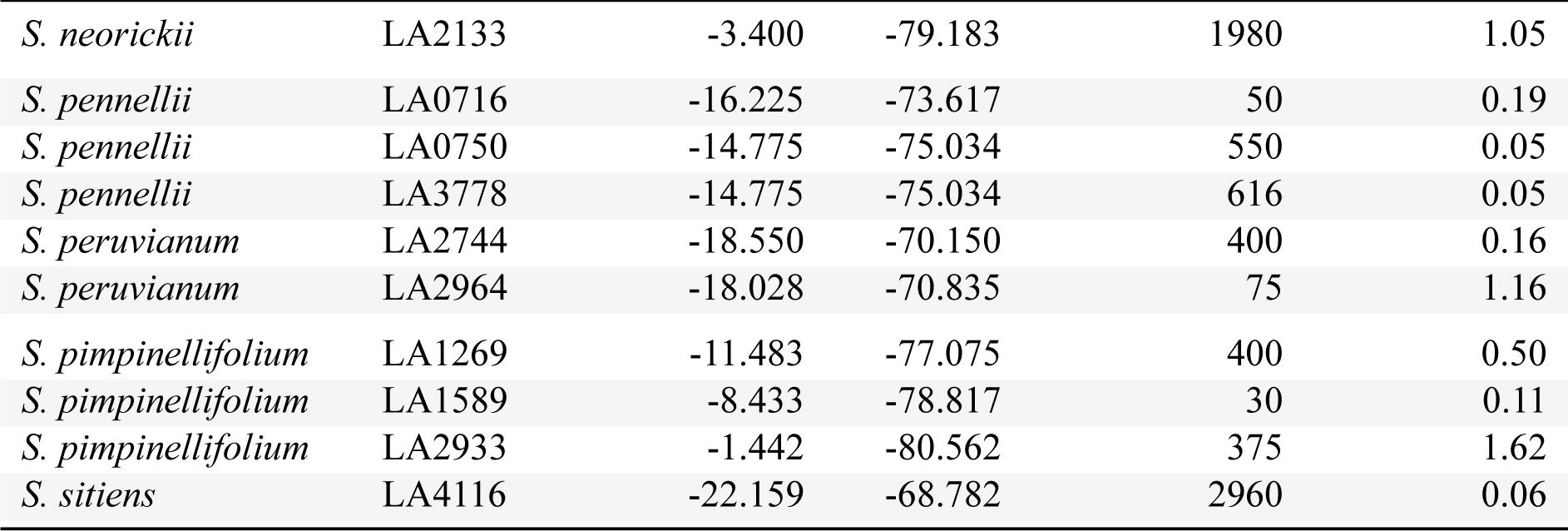
Accession information of *Solanum* populations used in this study. The species name, accession number, collection latitude, longitude, elevation, and plant area index (PAI) from the Global Ecosystem Dynamics Investigation (see ‘Climate data’). TGRC: Tomato Genetics Resource Center.

| Species | TGRC accession | Latitude | Longitude | Elevation (mas) | PAI (m<sup>2</sup> m<sup>-2</sup>) |
| --- | --- | --- | --- | --- | --- |
| *S. arcanum* | LA2172 | -6.008 | -78.858 | 662 | 0.70 |
| *S. cheesmaniae* | LA0429 | -0.644 | -90.329 | 800 | 0.60 |
| *S. cheesmaniae* | LA3124 | -0.804 | -90.042 | 1 | 0.36 |
| *S. chilense* | LA1782 | -15.267 | -74.633 | 1000 | 0.35 |
| *S. chilense* | LA4117A | -22.907 | -67.941 | 3540 | 0.22 |
| *S. chmielewskii* | LA1028 | -13.883 | -73.017 | 3000 | 0.43 |
| *S. chmielewskii* | LA1316 | -13.400 | -73.906 | 2920 | 1.07 |
| *S. corneliomulleri* | LA0107 | -13.117 | -76.383 | 60 | 0.02 |
| *S. corneliomulleri* | LA0444 | -13.433 | -76.133 | 100 | 0.49 |
| *S. galapagense* | LA0436 | -0.953 | -90.978 | 40 | 0.20 |
| *S. galapagense* | LA1044 | -0.284 | -90.548 | 0 | 0.18 |
| *S. habrochaites* | LA0407 | -2.181 | -79.884 | 70 | 0.47 |
| *S. habrochaites* | LA1777 | -9.550 | -77.700 | 3216 | 0.42 |
| *S. huaylasense* | LA1358 | -9.533 | -77.967 | 750 | 0.58 |
| *S. huaylasense* | LA1360 | -9.546 | -77.929 | 1490 | 0.48 |
| *S. huaylasense* | LA1364 | -10.133 | -77.383 | 2920 | 0.88 |
| *S. lycopersicoides* | LA2951 | -19.317 | -69.450 | 2200 | 0.49 |
| *S. lycopersicoides* | LA4126 | -19.287 | -69.396 | 3120 | 0.39 |
| *S. neorickii* | LA1322 | -13.483 | -72.442 | 2380 | 0.68 |

| Species | TGRC accession | Latitude | Longitude | Elevation (mas) | PAI ($\text{m}^2 \text{m}^{-2}$) |
| --- | --- | --- | --- | --- | --- |
| *S. neorickii* | LA2133 | -3.400 | -79.183 | 1980 | 1.05 |
| *S. pennellii* | LA0716 | -16.225 | -73.617 | 50 | 0.19 |
| *S. pennellii* | LA0750 | -14.775 | -75.034 | 550 | 0.05 |
| *S. pennellii* | LA3778 | -14.775 | -75.034 | 616 | 0.05 |
| *S. peruvianum* | LA2744 | -18.550 | -70.150 | 400 | 0.16 |
| *S. peruvianum* | LA2964 | -18.028 | -70.835 | 75 | 1.16 |
| *S. pimpinellifolium* | LA1269 | -11.483 | -77.075 | 400 | 0.50 |
| *S. pimpinellifolium* | LA1589 | -8.433 | -78.817 | 30 | 0.11 |
| *S. pimpinellifolium* | LA2933 | -1.442 | -80.562 | 375 | 1.62 |
| *S. sitiens* | LA4116 | -22.159 | -68.782 | 2960 | 0.06 |

#### Plant growth conditions

We recorded PPFD using full spectrum quantum sensors (SQ-500-SS, Apogee Instruments, Logan, Utah, USA); we recorded temperature, RH, and [CO_2_] using an EE894 sensor (E+E Elektronik, Engerwitzdorf, Austria) protected by a radiation shield. All environmental measurements were taken every 10 minutes from the middle of plant racks at approximately the same height as the leaves we measured. We measured leaf temperature of focal leaves prior to measurement using an infrared radiometer (SI-111-SS, Apogee Instruments, Logan, Utah, USA).

#### Germination and seedling stage

Seeds provided by the Tomato Genetics Resource Center germinated on moist paper in plastic boxes after soaking for 30-60 minutes in a 50% (volume per volume) solution of household bleach and water, followed by a thorough rinse. We transferred seedlings to cell-pack flats containing Pro-Mix BX potting mix (Premier Tech, Rivière-du-Loup, Quebec, Canada) once cotyledons fully emerged, typically within 1-2 weeks of sowing. We grew seeds and seedlings for both sun and shade treatments under the same environmental conditions (12:12 h, 24.3:21.7 °C, 49.6:58.4 RH day:night cycle). LED light provided PPFD = 267 µmol m^−2^ s^−1^ (Fluence RAZRx, Austin, Texas, USA) during the germination and seedling stages.

#### Light treatments

Seedlings were randomly assigned in alternating order within population to the sun or shade treatment during transplanting. After seedlings established in cell-pack flats for ≈ 2 weeks, we transplanted them to 3.78 L plastic pots containing 60% Pro-Mix BX potting mix, 20% coral sand (Pro-Pak, Honolulu, Hawai‘i, USA), and 20% cinders (Niu Nursery, Honolulu, Hawai‘i, USA). Percentage composition is on a volume basis. The soil mixture contained slow release NPK fertilizer following manufacturer instructions (Osmocote Smart-Release Plant Food Flower & Vegetable, The Scotts Company, Marysville, Ohio, USA). We determined pot field capacity one week after transplanting using a scale (Ohaus V12P15 Valor 1000, Parsippany, New Jersey, USA) and watered to field capacity three times per week to prevent drought stress.

We assigned sun and shade treatment to lower and upper racks of a 1.22 m × 2.44 m shelving unit in a climate-controlled growth room. We assigned the sun treatment to the lower rack to limit diffuse light from reaching the shade treatment. The average daytime PPFD was 761 µmol m^−2^ s^−1^ and 115 µmol m^−2^ s^−1^ for sun and shade treatments, respectively. To isolate the effect of light intensity from quality, we used the same LED model with the the same spectrum (Fluence SPYDR 2i, Austin, Texas, USAS), but dimmed the lights in the shade treatment. To maintain homogeneous environmental conditions other than light, we mixed air within the growth room using an air circulator (Vornado 693DC, Andover, Kansas, USA) and within racks using a miniature oscillating air circulator (Vornado Atom 1, Andover, Kansas, USA). Despite these efforts, the air in the sun treatment was on average 2.56 °C warmer and the average RH was consequently 5.75% lower. However, because of evaporative cooling, the leaves in the sun treatment were only 0.886 °C warmer on average *(n* = 699 leaves).

#### Leaf trait measurements

We selected a fully expanded, unshaded leaf at least six leaves above the cotyledons during early vegetative growth. This typically meant that plants had grown in light treatments for ≈ 4 weeks, ensuring they had time to sense and respond developmentally to the light intensity of the treatment rather than the seedling conditions (*55*). Shade plants grew slower than sun plants, hence leaves at the same developmental stage were measured on chronologically older plants in the shade treatment. In some sun plants, we had to use leaves higher on the stem because short internodes made lower leaves inaccessible with the gas exchange equipment. We measured terminal leaflets in 82.6% of cases, but used the lateral leaflet closest to the terminal leaflet when it was damaged or difficult to clamp into the gas exchange chamber. When a leaflet was damaged during gas exchange measurements, we collected anatomical data from the nearest leaflet on the same leaf (1.58% of leaves).

#### Amphistomy advantage

We estimated ‘amphistomy advantage’ (AA) *sensu* (*20*), but with modifications previously described in (*46*). AA is calculated as the log-response ratio of *A* compared at the same total *g*_sw_:

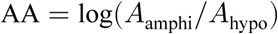

We measured the photosynthetic rate of an untreated amphistomatous leaf (*A*<u>_am_</u>_phi_) over a range of *g*_sw_ values. We refer to this as an *A-g*_sw_ curve. We compared the *A*-*g*_sw_ curve of the untreated leaf to the photosynthetic rate of pseudohypostomatous leaf (*A*_hypo_), which is the same leaf but with gas exchange through the upper surface blocked by a neutral density plastic (propafilm).

We measured *A-g*_sw_ curves using a portable infrared gas analyzer (LI-6800PF, LI-COR Biosciences, Lincoln, Nebraska, USA). Light-acclimated plants were placed under LEDs dimmed to match their light treatment during gas exchange measurements. We estimated the photosynthetic rate (A) and stomatal conductance to CO_2_ (*g*_sw_) at ambient CO_2_ *(C_a_* = 415 µmol mol^−1^) and *T*_leaf_ = 25.0 °C. The irradiance of the light source in the pseudohypo leaf was higher because the propafilm reduces transmission. To compensate for reduced transmission, we increased incident PPFD for pseudohypo leaves by a factor 1/0.91, the inverse of the measured transmissivity of the propafilm. We also set the stomatal conductance ratio, for purposes of calculating boundary layer conductance, to 0 for pseudohypo leaves following manufacturer directions.

We collected four *A-g*_sw_ curves per leaf, an amphi (untreated) curve and a pseudohypo (treated) curve at high light-intensity (PPFD = 2000 µmol m^−2^ s^−1^; 97.8:2.24 red:blue) and low light-intensity (PPFD = 150 µmolm^−2^ s^−1^; 87.0:13.0 red:blue). We always measured high light-intensity curves first because photosynthetic downregulation is faster than upregulation in these species. To control for order effects, we alternated between starting with amphi or pseudohypo leaf measurements. Unlike (*46*), preliminary experiments with *Solanum* indicated a strong order effect in that A declined in the second curve. Therefore, we made measurements over two days. On the first day, we measured high and low light-intensity curves for either amphi or pseudohypo leaves; on the second day, we measured high and low light-intensity curves on the other leaf type.

In all cases, we acclimated the focal leaf to high light (PPFD = 2000 µmol m^−2^ s^−1^) and high relative humidity (RH = 70%) until A and *g*_sw_ reach their maximum. After that, we decreased RH to ≈ 10% to induce rapid stomatal closure without biochemical downregulation. Hence, *A*<u>_am_</u>_phi_ and *A*_hypo_ were both measured at low chamber humidity after the leaf had acclimated to high humidity. All other environmental conditions in the leaf chamber remained the same. We logged data until *g*_sw_ reached its nadir. We then acclimated the leaf to low light (PPFD = 150 µmol m^−2^ s^−1^) and RH = 70% before inducing stomatal closure with low RH and logging data as described above.

#### Stomatal anatomy

We estimated the stomatal density and size on ab- and adaxial leaf surfaces from all leaves, using guard cell length as a proxy for stomatal size since it is proportional to maximum conductance (*56*). We made surface impressions of leaf lamina from the same area used for gas exchange measurements using a-silicone impression material (Zhermack elite HD+, light body, fast set, Rovigo, Italy). We applied clear nail polish to make positive replicas of the impression. After nail polish dried, we mounted replicas on a microscope slide using transparent tape (*57*). We digitized a portion of each leaf surface replica using a brightfield microscope (Leica DM2000, Wetzlar, Germany). We counted and measured guard cell length on all stomata using the FIJI implementation of ImageJ2 version 2.3.0 (*58*), then divided the count by the visible leaf area (0.890 mm^2^) to estimate stomatal density. For each surface we calculated the anatomical maximum stomatal conductance (*g*_max_) at a reference leaf temperature of 25 °C following (*56*) as:

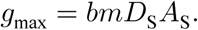

The biophysical and morphological constants *b* and *m* are:

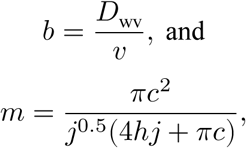

where *D*_wv_ is the diffusion coefficient of water vapor in air and v is by the kinematic viscosity of dry air. We assumed *D*_wv_ = 2.49 × 10_-5_ m^2^ s^−1^ and v = 2.24 × 10^−2^ m_3_ mol^−1^ (*59*). For kidney-shaped guard cells like those in wild tomatoes, *c = h = j =* 0.5. The *g*_max.ratio_ is the *g*_max_ of the adaxial surface divided by the sum total *g*_max_ of both surfaces.

#### Leaf mass per area

Leaf mass per area (LMA) is the dry mass divided by the leaflet area. We scanned fresh leaflets on a flat bed scanner (Epson V600, Los Alamitos, California. USA) and measured leaflet area from digital images using the FIJI implementation of ImageJ2 version 2.3.0 (*58*). We dried leaves for 72 hours at 74 °C in a food dehydrator (Cosori CP267-FD, Vesync Co., Anaheim, California, USA) and weighed using a benchtop analytical balance (Ohaus PR64 Analytical Balance, Parsippany, New Jersey, USA). In 10.5% of leaves we measured LMA on the adjacent leaflet because the focal leaflet was damaged or wilted while making surface impressions and we could not reliably estimate area. LMA data are missing from 2.97% of individuals because the area or mass was not recorded at all or recorded incorrectly.

#### Cleaning *A-g*_sw_ curves

The raw data set consisted of 2,370 *A-g*_sw_ curves with an average of 63.2 points per curve. Manual curation of a data set this size in a principled, consistent manner is not feasible. Therefore, we automated data cleaning using custom *R* scripts. Cleaning is divided into six sequential steps (Table S2).

#### Remove unreliable and unusable data points

##### Rationale

Unreliable data points consisted of those where chamber [CO_2_] was unstable and therefore measurements are not biologically meaningful. Unusable data points were those where A < 0 because the logarithm of a negative number is undefined.

##### Procedure

We retained data points where 410 < *C_a_* < 420 µmol mol^−1^ and A > 0.

**Table S2:** Six sequential steps for cleaning *A-g*_sw_ curves. The rationale and procedure for each step are described in the text. The rightmost columns summarize the number of curves and mean number of points per curve remaining after each step. For reference, there are four possible *A-g*_sw_ curves per replicate: all combinations of leaf type (amphi or pseudohypo) and light intensity (high or low).

| Step: description | Number of curves | Number of points per curve |
| --- | --- | --- |
| 1. remove unreliable and unusable data points | 2,361 | 63.0 |
| 2. remove hysteretic portion of $A$-$g_{sw}$ curves at low $g_{sw}$ | 2,360 | 59.2 |
| 3. remove outliers within each $A$-$g_{sw}$ curve | 2,360 | 58.7 |
| 4. remove replicates with no overlap between amphi and pseudohypo $A$-$g_{sw}$ curves | 2,268 | 58.4 |
| 5. thin redundant data points within each $A$-$g_{sw}$ curve | 2,268 | 28.1 |
| 6. trim extreme AA values | 2,214 | 28.1 |

#### Remove hysteretic portion of *A-g*_sw_ curves at low *g*_sw_

##### Rationale

In most *A-g*_sw_ curves, we observed a hysteretic response at low *g*_sw_. After *g*_sw_ and *A* declined simultaneously, *A* increased slightly as *g*_sw_ continued to decline or stabilize, indicating some leaf acclimation to low RH. We removed this portion of the curve to focus curve-fitting on the primary domain where *A* increases monotonically with *g*_sw_.

##### Procedure

For each curve, we removed data points after *g*_sw_ had reached its minimum unless there were fewer than 10 data points remaining.

#### Remove outliers within each *A-g*_sw_ curve

##### Rationale

Individual outliers within *A-g*_sw_ curves, usually caused by transitory changes in chamber conditions, exert undue leverage on parameter estimates and cause bias and/or low precision in param-eter estimates.

##### Procedure

We fit provisional quadratic regressions for each curve using ordinary least squares with the lm() function in *R*. We sequentially removed data points with an absolute external studentized residual *> 3* until none remained.

#### Thin redundant data points within each *A-g*_sw_ curve

##### Rationale

Data points closely spaced along the *A-g*_sw_ curve provide redundant information and may be highly correlated (i.e. pseudoreplication). This occurred because data was logged at a constant temporal interval, but the rate at which *g*_sw_ declined was not constant. Thinning reduces parameter estimation bias toward densely sampled regions of the curve which may not be the most biologically informative.

##### Procedure

We retained the maxima and minima *g*_sw_ for each curve and thinned all but one point per thinning interval of 0.05 log(mol m^−2^ s^−1^), retaining the point nearest the midpoint of the interval.

#### Remove replicates with no overlap between amphi and pseudohypo *A-g*_sw_ curves

##### Rationale

We could not estimate AA for replicates where amphi and pseudohypo *A-g*_sw_ curves did not overlap.

##### Procedure

We removed replicates where the range of *g*_sw_ values for amphi and pseudohypo *A-g*_sw_ curves did not overlap.

#### Trim extreme AA values

##### Rationale

Extreme AA values were likely due to measurement error or leaf damage. Since amphi and pseudohypo *A-g*_sw_ curves are measured on consecutive days, a poor calibration or a damaged leaf could cause a large difference in *A* between days, which would appear as an extreme AA value.

##### Procedure

We provisionally estimated AA for each replicate by integrating over the range of *g*_sw_ val-ues where amphi and pseudohypo *A-g*_sw_ curves overlap. In this procedure, curve parameters were provisionally estimated using ordinary least squares with the lm() function in *R*. We then used point estimates ofAA for each replicate as the response variable in a linear model with light treatment, light intensity, population, and all interactions as explanatory variables. This model was also fit using ordinary least squares with the lm() function in *R*. We classified extreme AA values as those with an absolute internal studentized residual *> 3*. Because these values likely indicate significant measurement error or leaf damage, we removed *A-g*_sw_ curves at both light intensities if either was classified as extreme.

#### Bayesian data analysis with *Stan*

Our analysis pipeline consisted of four main steps:

1. Estimate *A-g*_sw_ curve parameters
2. Estimate AA for both measurement light intensities using *A-g*_sw_ curve parameters
3. Estimate the effects of light intensity, light treatment, and population on AA (acclimation and plasticity hypotheses)
4. Estimate the effects of native light habitat on population-level AA (constitutive hypothesis)

All models were fit using a Bayesian HMC sampling in the probabilistic programming language *Stan* (*60*) using the *R* package **brms** version 2.22.0 (*61*). We used *CmdStan* version 2.36.0 and **cmdstanr** version 0.9.0 (*62*) to interface with *R* version 4.5.1 (*63*). We sampled the posterior distribution from a single chain with a minimum of 1000 iterations after 1000 warmup iterations. If necessary, we refit models with more iterations to decrease the Gelman-Rubin convergence statistic *(R)* (*64*) to less than 1.01 and the bulk effective sample size (ESS) to greater than 400 for all model parameters, and there were fewer than 10 divergent transitions.

#### *A*–*g*_sw_ curve parameters

We modeled log (A) as a quadratic function of log (*g*_sw_) for each leaf, measured at low and high light intensity, in both amphistomatous (untreated) and pseudohypostomatous (propafilm treatment) states.

#### AA for each light intensity within leaf using *A-g*_sw_ curve parameters

We estimated AA for each light intensity by integrating the difference in log (A) between the amphi and pseudohypo A-*g*_sw_ curves over the range of *g*_sw_ values where the curves overlap (from min(log(*g*_sw_)) tomax(log(*g*_sw_))).

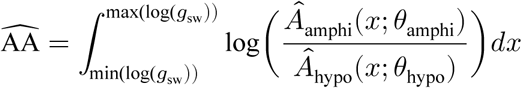

where *θ*_amphi_ and *θ*_hypo_ are the quadratic parameters of the amphi and pseudohypo *A*-*g*_sw_ curves, re-spectively (Figure S1). We calculated 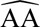 from each draw of the posterior for the paired amphi and pseudohypo *A-g*_sw_ curves to obtain the posterior distribution of 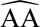 for each leaf at each light inten-sity. We then summarized the posterior distribution of 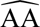for each leaf at each light intensity by the posterior median and standard deviation, which became our point estimate of, and uncertainty in, 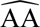 for the phylogenetic mixed effects model described below.

#### Phylogenetic mixed effects models

We fit Bayesian mixed effects models with phylogenetically structured random effects to test predictions of competing hypotheses about why amphistomy advantage (AA) might be greater for leaves in sunny, open habitats. We fit all models combinations of measurement light intensity, growth light intensity, and their interaction as both fixed effects and random effects among populations. In other words, we test whether there were main effects of light intensity treatments and/or whether populations varied in their response to light intensity treatments. In all models, we included population as a phylogenetically structured random effect. We accounted for uncertainty in 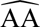 using the standard deviation of the posterior distribution of each 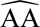 estimate (see above). We crossed fixed and random effect structures with fixed effects of measurement light intensity and/or growth light intensity on the distributional parameter, σ, the residual variance in AA. We used a robust regression approach by assuming ŕ-distributed residuals, which makes estimates less susceptible to extreme values. In total, we fit 100 models, each with a different combination of fixed, random, and distributional effects. We used the RAxML whole-transcriptome concatenated phylogeny based on 2,745 100-kb genomic windows from (*54*). Two of our populations were not in this tree. We used accession LA1044 in place of LA3909, two populations of *S. galapagense*; LA0750 was added as sister to LA0716, two closely related populations of *S. pennellii*. The node separating LA0750 from LA0716 was placed half-way between the next deepest node. We used the leave-one-out cross-validation information criterion (LOOIC) to compare the fit of models (Table S6) using the *R* package **loo** version 2.8.0 (*65*) to calculate LOOIC values. We selected the model with the lowest LOOIC value (Table S6) to generate posterior predictions for hypothesis testing. Parameter estimates from the selected model are shown in Table S4.

**Figure S1:**
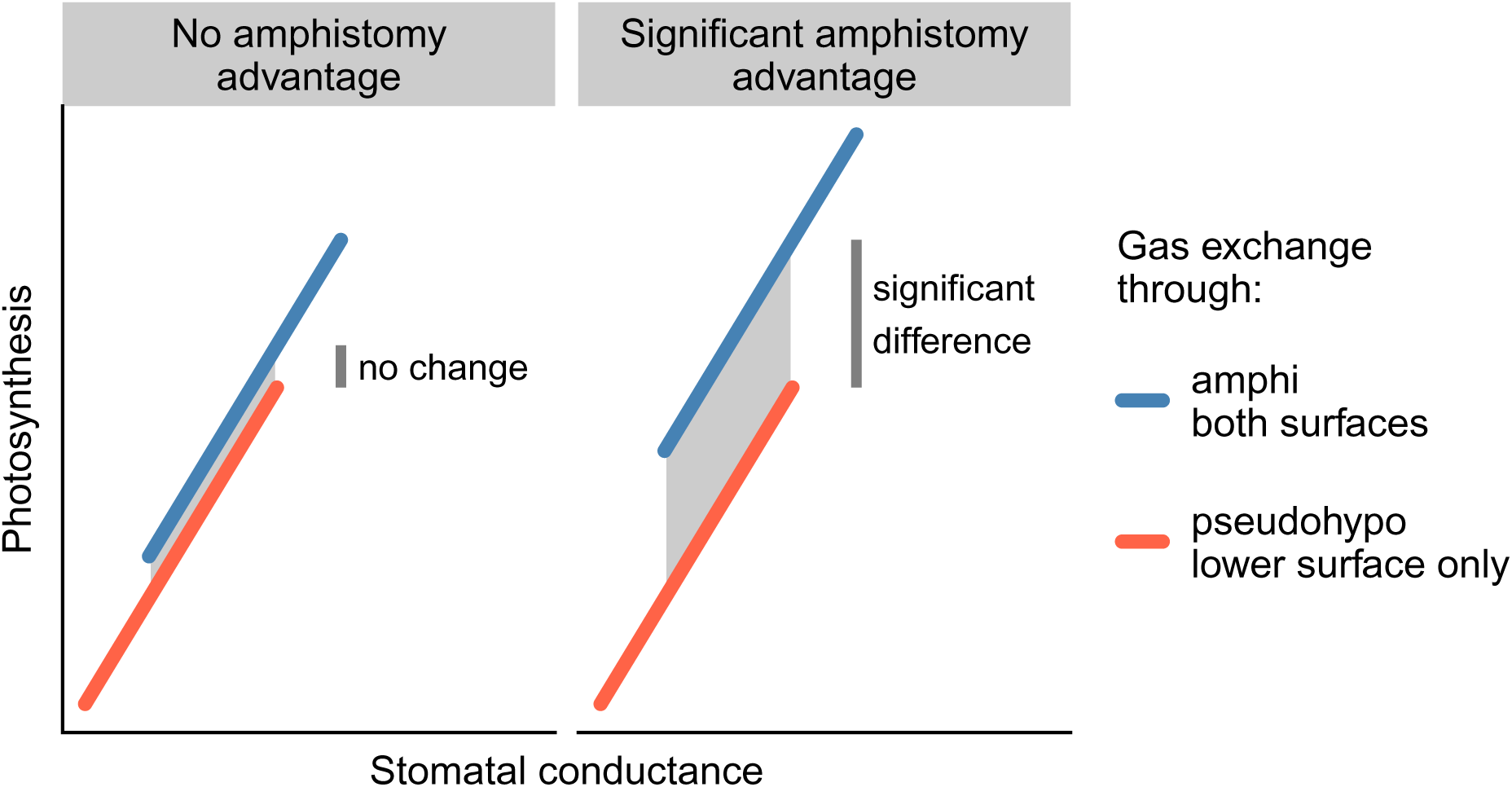
Example amphistomy advantage (AA) calculation from *A-g*_sw_ curves in an amphistomatous leaf with gas exchange through both surfaces (blue) and the same leaf with gas exchange blocked on the upper surface only (orange). We integrate over the gray region between the curves to estimate AA. The AA is low (left facet) when the photosynthetic rate of an amphistomatous leaf is similar to a pseudohypostomatous leaf at the same total stomatal conductance (r-axes); large AA (right facet) is indicated when an amphistomatous leaf has a higher photosynthetic rate than a pseudohypostomatous leaf.

To test whether LMA mediated the relationship between growth light intensity and AA, we repeated a similar model comparison procedure but included log LMA as an additional fixed effect in the model, as well its interaction with measurement and growth light intensity. The model included a direct effect of growth light intensity on log LMA and random phylogeneticlly structured effects of population on log LMA and effects of sun treatment on log LMA. This enabled us to disentangle whether log LMA effects AA directly or indirectly because both traits respond to growth light intensity. We integrated over missing LMA data using the mi() function in **brms**. Because the evidence from the first set of models strongly supported including effects of growth and measurement light intensity on the residual variance in AA, we included these in terms in all models. In total, we fit 75 models, each with a different combination of fixed and random effect structures. We selected the model with the lowest LOOIC value (Table S7) to generate posterior predictions for hypothesis testing. Note that LOOIC was calculated only from pointwise likelihood values of the submodel estimating effects of light intensity, light treatments, and population on AA. Parameter estimates from the selected model are shown in Table S5.

#### Posterior predictions

The acclimation, plasticity, and constitutive hypotheses make different predictions about the relationship between AA and light intensity, light treatment, and native PAI among populations. Since these hypotheses are not mutually exclusive, we describe how we assessed support for one, two, or all three hypotheses simultaneously in Table S3. In general, the acclimation hypothesis was supported if AA was greater at high light intensity than low light intensity. The plastic hypothesis was supported if AA was greater in sun leaves than shade leaves. The constitutive hypothesis was supported if population-level AA increased with native PPFD. In interactive models, we only consider positively reinforcing interactions between high light intensity, sun leaves, and native PPFD because these are the only interactions which could explain why amphistomatous leaves are advantageous in high light habitats.

From the model selected using LOOIC, we predicted the posterior distribution of the expected AA for each population at each measurement light intensity and growth light intensity. If the 95% confidence intervals for AA did not overlap zero, we considered AA to be significantly different from zero. We also calculated posterior distribution for the average AA among populations in each condition. We tested a linear effect of native light habitat on population-level AA in each treatment condition, integrating over the entire posterior distribution to calculate a posterior distribution of the slope. If the 95% confidence interval for the slope did not overlap zero, we considered the slope to be significant.

**Table S3:** Predictions of competing hypotheses about the relationship between AA and light intensity, light treatment, and native plant area index (PAI) among populations. The middle column lists specific directional predictions about parameter values. The rightmost column describes the predictions in words and explains how population-level AA is calculated in the relevant model.

| Hypothesis | Prediction(s) | Description |
| --- | --- | --- |
| Acclimation | $AA_{2000} > 0$ | AA at high light intensity is greater than that at low light intensity |
| Plasticity | $AA_{\text{sun}} > 0$ | AA in sun leaves is greater than that in shade leaves |
| Constitutive | $\beta_{\text{PAI},AA} < 0$ | Population-level AA ($AA_{\text{pop}}$) decreases with native PAI<br>$AA_{\text{pop}} = \beta_{AA,0} + \vec{\beta}_{AA,\text{pop}}$ |
| Acclimation × Plasticity | $AA_{2000,\text{sun}} > 0$ | AA is highest at high light intensity in sun leaves |
| Acclimation × Constitutive | $AA_{2000} > 0$<br>$\beta_{\text{PAI},AA} < 0$ | AA at high light intensity is greater than that at low light intensity<br>Population-level AA at high light intensity ($AA_{\text{pop},2000}$) decreases with native PAI<br>$AA_{\text{pop},2000} = \beta_{AA,0} + \vec{\beta}_{AA,\text{pop}} + \vec{\beta}_{AA,2000,\text{pop}}$ |
| Plasticity × Constitutive | $AA_{\text{sun}} > 0$<br>$\beta_{\text{PAI},AA} < 0$ | AA in sun leaves is greater than that in shade leaves<br>Population-level AA in sun leaves ($AA_{\text{pop},\text{sun}}$) decreases with native PAI<br>$AA_{\text{pop},\text{sun}} = \beta_{AA,0} + \vec{\beta}_{AA,\text{pop}} + \vec{\beta}_{AA,\text{sun},\text{pop}}$ |
| Acclimation × Plasticity × Constitutive | $AA_{2000,\text{sun}} > 0$<br>$\beta_{\text{PAI},AA} > 0$ | AA is highest at high light intensity in sun leaves<br>Population-level AA at high light intensity in sun leaves ($AA_{\text{pop},2000,\text{sun}}$) decreases with native PAI<br>$AA_{\text{pop},2000,\text{sun}} = \beta_{AA,0} + \vec{\beta}_{AA,\text{pop}} + \vec{\beta}_{AA,2000,\text{pop}} + \vec{\beta}_{AA,\text{sun},\text{pop}}$ |

#### The water cost of hypostomy

Amphistomy advantage implies that hypostomatous leaves would need to lose more water by having higher *g*_sw_ to achieve the same photosynthetic rate as an otherwise identical amphistomatous leaf. We estimated this cost of hypostomy approximately. For simplicity, we focus on a sufficiently small region of the *A-g*_sw_ curve such that we can approximate the relationship between *g*_sw_ and *A* as log-log linear. We also assume that the slope is the same in both amphistomatous and hypostomatous leaves, which was often approximately the case in our experiment (Figure S3). With these assumptions, the approximate water cost of hypostomy can be derived from the equations:

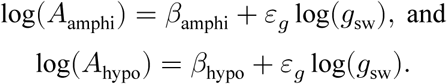

Solving for the difference in log *g*_sw_ at a given log *A*_hypo_ is:

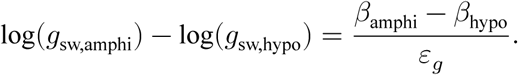

Noting that because the curves are parallel, AA = log(*A*_amphi_/*A*_hypo_) = *β*_amphi_ — *β*_hypo_, this reveals that the negative log-ratio of the *g*_sw_ is related to AA divided by the slope:

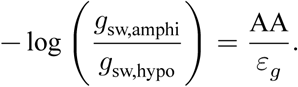

We note that the slope in this case is the elasticity of *A* to *g*_sw_ because both are log-transformed. This quantity will be proportional the water cost of hypostomy assuming that VPD and other environmental variables are held constant and the boundary layer resistance is negligible.

**Table S4:** Parameter estimates and 95% confidence intervals (CIs) from the posterior distribution of the selected model. Models potentially include fixed, random, and distributional effects of growth light intensity (shade and sun), measurement light intensity (low and high), and their interaction on amphistomy advantage (AA). The distributional effects refer to the effect of factors on residual variation in AA, denoted σ, on a log-link scale. For phylogenetically structured random effects, we report the estimated standard deviation (SD).

| Parameter | Estimate [95% CI] |
| --- | --- |
| *Fixed effects* |  |
| AA intercept (shade, low light) | 0.08 [0.01, 0.16] |
| effect of sun on AA | 0.04 [-0.02, 0.12] |
| effect of high light on AA | -0.03 [-0.06, 0] |
| effect of sun × high light interaction on AA | -0.03 [-0.1, 0.03] |
| *Distributional parameters* |  |
| log $\sigma$ intercept (shade, low light) | -2.51 [-2.61, -2.41] |
| effect of sun on log $\sigma$ | 0.23 [0.12, 0.33] |
| effect of high light on log $\sigma$ | -0.19 [-0.3, -0.09] |
| *Random effect SDs* |  |
| AA among populations | 0.06 [0.04, 0.1] |
| effect of sun on AA among populations | 0.05 [0, 0.1] |
| effect of high light on AA among populations | 0.02 [0, 0.05] |
| effect of sun × high light interaction on AA among populations | 0.04 [0, 0.1] |

**Table S5:**
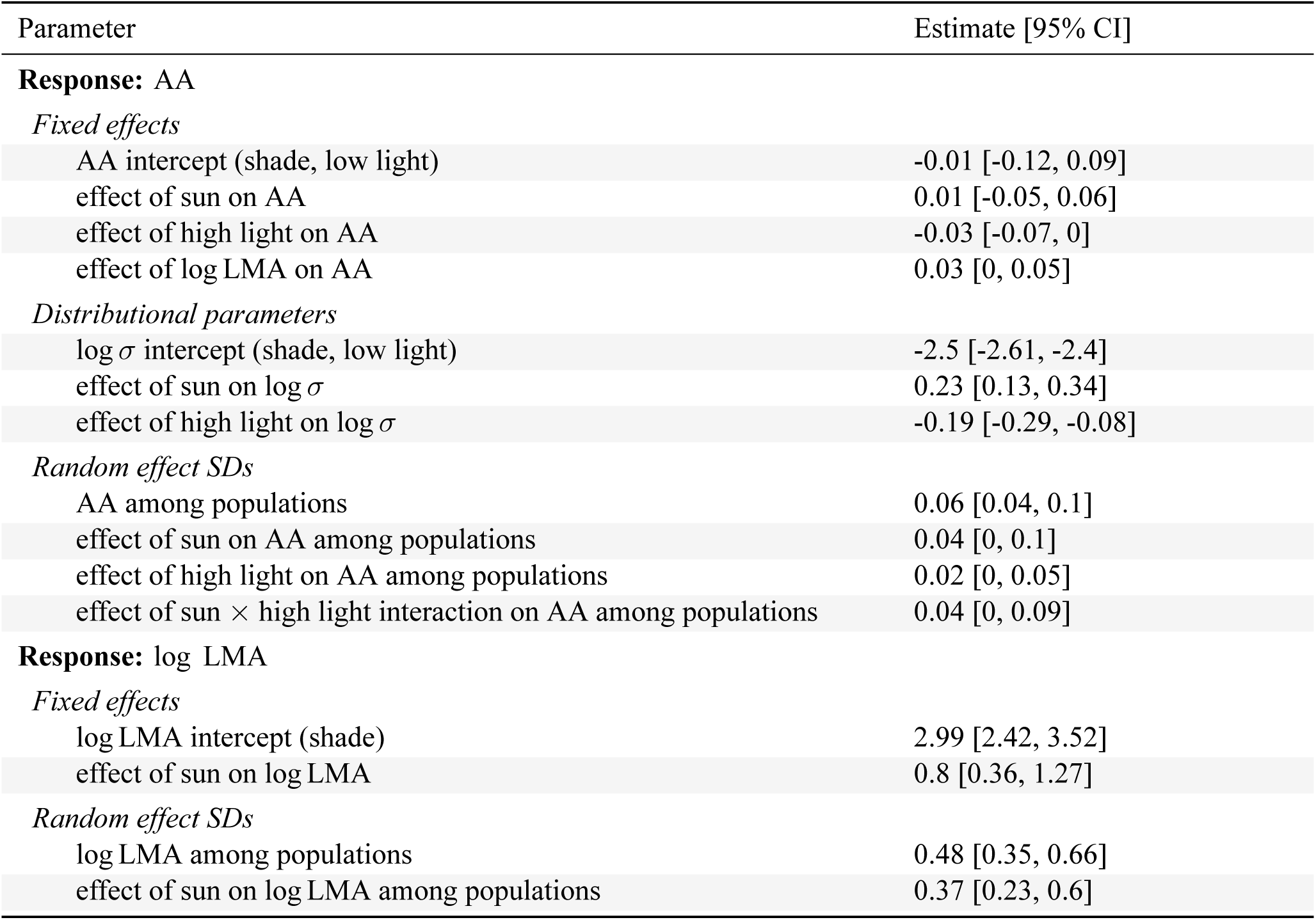
Parameter estimates and 95% confidence intervals (CIs) from the posterior distribution of the selected model with leaf mass per area (LMA) as potential mediator variable. Models potentially include fixed and random effects of growth light intensity (shade and sun), measurement light intensity (low and high), log LMA, and their interactions on amphistomy advantage (AA). LMA can also be affected by growth light intensity and population. For phylogenetically structured random effects, we report the estimated standard deviation (SD).

**Figure S2:**
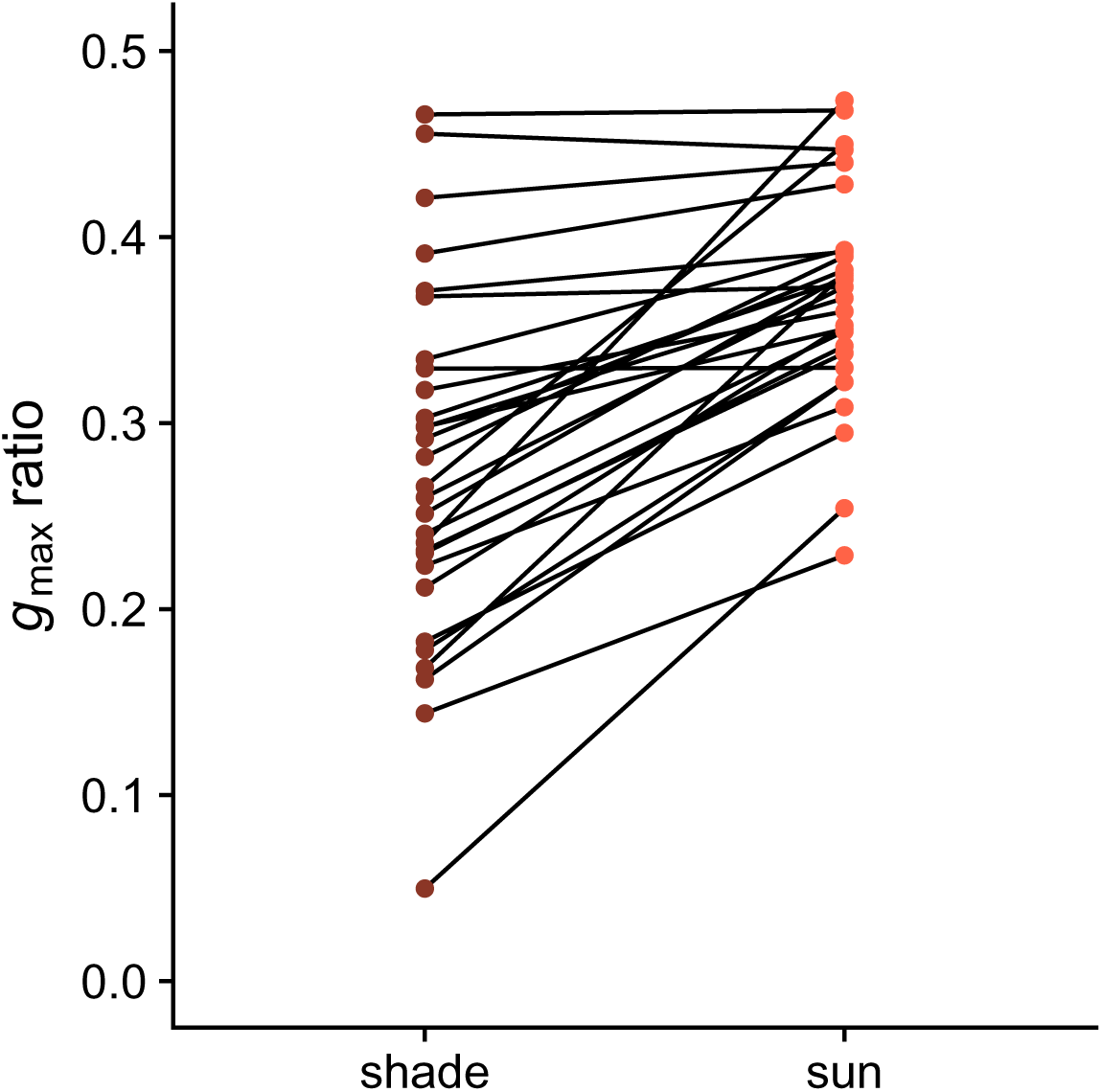
Developmental plasticity in anatomical maximum stomatal conductance ratio (*g*_max,ratio_; *y-* axis) to growth light intensity (re-axis) among 29 wild tomato populations. All populations are amphistomatous, but vary in the stomatal density and size on each surface, which determines *g*_max,ratio_. Each point is the average per population in that treatment; black lines connect the same population across treatments.

**Table S6:** Leave-one-out cross-validation information criterion (LOOIC) for models of AA as a function of light intensity, light treatment, and population. Only the top models (ΔLOOIC < 2) are shown. The model with the lowest LOOIC was selected for generating posterior predictions to test hypotheses. The LOOIC column lists the LOOIC value for each model; the ΔLOOIC column lists the difference between the LOOIC of the corresponding model and the best model (lowest LOOIC). Checkmarks in each row indicate whether the corresponding model included affects of measurement light intensity (“Meas.”) or growth light intensity (“Growth”) as fixed effects, random effects, or distributional effect on the residual variance. We also test for fixed and random interaction effects (“Inter.”) between measurement and growth light intensity. All models included phylogenetically structured random effects of population.

| LOOIC | $\Delta\text{LOOIC}$ | Fixed | | | Random | | | Distributional | |
| --- | --- | --- | --- | --- | --- | --- | --- | --- | --- |
|  |  | Meas. | Growth | Inter. | Meas. | Growth | Inter. | Meas. | Growth |
| -1882.56 | 0.00 | ✓ | ✓ | ✓ | ✓ | ✓ | ✓ | ✓ | ✓ |
| -1881.70 | 0.86 |  |  |  | ✓ | ✓ | ✓ | ✓ | ✓ |
| -1881.00 | 1.56 | ✓ |  |  | ✓ | ✓ | ✓ | ✓ | ✓ |
| -1880.91 | 1.65 | ✓ | ✓ |  | ✓ | ✓ | ✓ | ✓ | ✓ |
| -1880.63 | 1.93 | ✓ | ✓ | ✓ | ✓ | ✓ |  | ✓ | ✓ |

**Table S7:**
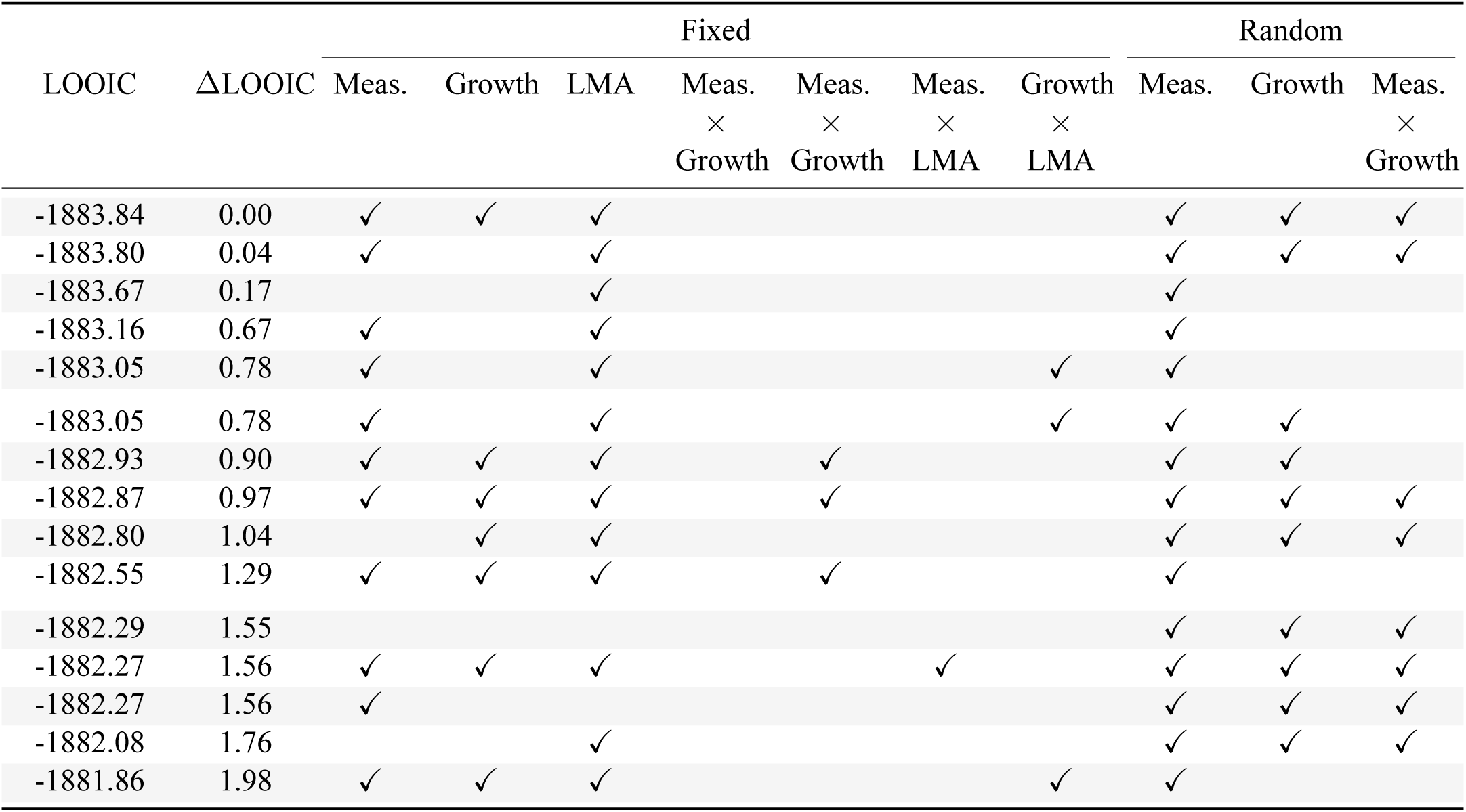
Leave-one-out cross-validation information criterion (LOOIC) for models of AA as a function of light intensity, light treatment, leaf mass per area (LMA), and population. Only the top models (Δ LOOIC < 2) are shown. The model with the lowest LOOIC was selected for generating posterior predictions to test hypotheses. The LOOIC column lists the LOOIC value for each model; the ΔLOOIC column lists the difference between the LOOIC of the corresponding model and the best model (lowest LOOIC). Checkmarks in each row indicate whether the corresponding model included affects of measurement light intensity (“Meas.”), growth light intensity (“Growth”), or LMA as fixed or random effects. We test for fixed interaction effects between measurement and growth light intensity and LMA. We also test for random interaction effects between measurement and growth light intensity. All models included phylogenetically structured random effects of population.

Figure S3: (Figures in separate file) The *A-g*_sw_ curves, fitted lines, and AA estimates for every individual plant. The title provides the Tomato Genetics Resource Center accession number, replicate letter, and species names. The subtitle indicates the growth light intensity (sun or shade). Points are raw data, lines are fitted curves, and shaded regions are 95% confidence ribbons for the fitted curves. The Bayesian correlation coefficient, *r^2^*, is shown to the right of each curve, along with the estimate of AA *±* two standard errors for each pair of amphi (orange) and pseudohypo (blue) curves.

## References

1. J. A. Raven, Selection pressures on stomatal evolution. New Phytologist 153, 371–386 (2002).

2. S. A. M. McAdam, J. G. Duckett, F. C. Sussmilch, S. Pressel, K. S. Renzaglia, R. Hedrich, T. J. Brodribb, A. Merced, Stomata: The holey grail of plant evolution. American Journal of Botany 108, 366–371 (2021).

3. J. W. Clark, B. J. Harris, A. J. Hetherington, N. Hurtado-Castano, R. A. Brench, S. Casson, T. A. Williams, J. E. Gray, A. M. Hetherington, The origin and evolution of stomata. Current Biology 32, R539–R553 (2022).

4. A. M. Hetherington, F. I. Woodward, The role of stomata in sensing and driving environmental change. Nature 424, 901–908 (2003).

5. H. J. de Boer, C. A. Price, F. Wagner□Cremer, S. C. Dekker, P. J. Franks, E. J. Veneklaas, Optimal allocation of leaf epidermal area for gas exchange. New Phytologist 210, 1219–1228 (2016).

6. E. L. Harrison, L. Arce Cubas, J. E. Gray, C. Hepworth, The influence of stomatal morphology and distribution on photosynthetic gas exchange. The Plant Journal 101, 768–779 (2020).

7. T. N. Buckley, K. A. Mott, Modelling stomatal conductance in response to environmental fac-tors: Modelling stomatal conductance. Plant, Cell & Environment 36, 1691–1699 (2013).

8. M. Haworth, C. Elliott-Kingston, J. C. McElwain, Co-ordination of physiological and morpho-logical responses of stomata to elevated [CO2] in vascular plants. Oecologia 171, 71–82 (2013).

9. P. J. Franks, D. L. Royer, D. J. Beerling, P. K. Van de Water, D. J. Cantrill, M. M. Barbour, J. A. Berry, New constraints on atmospheric CO_2_ concentration for the Phanerozoic. Geophysical Research Letters 41, 4685–4694 (2014).

10. J. C. McElwain, M. Steinthorsdottir, Paleoecology, ploidy, paleoatmospheric composition, and developmental biology: A review of the multiple uses of fossil stomata. Plant Physiology 174, 650–664 (2017).

11. The Cenozoic CO Proxy Integration Project (CenCOPIP) Consortium*†, B. Hönisch, D. L. Royer, D. O. Breecker, P. J. Polissar, G. J. Bowen, M. J. Henehan, Y. Cui, M. Steinthorsdottir, J. C. McElwain, M. J. Kohn, A. Pearson, S. R. Phelps, K. T. Uno, A. Ridgwell, E. Anagnostou, J. Austermann, M. P. S. Badger, R. S. Barclay, P. K. Bijl, T. B. Chalk, C. R. Scotese, E. De La Vega, R. M. DeConto, K. A. Dyez, V. Ferrini, P. J. Franks, C. F. Giulivi, M. Gutjahr, D. T. Harper, L. L. Haynes, M. Huber, K. E. Snell, B. A. Keisling, W. Konrad, T. K. Lowenstein, A. Malinverno, M. Guillermic, L. M. Mejía, J. N. Milligan, J. J. Morton, L. Nordt, R. Whiteford, A. Roth-Nebelsick, J. K. C. Rugenstein, M. F. Schaller, N. D. Sheldon, S. Sosdian, E. B. Wilkes, C. R. Witkowski, Y. G. Zhang, L. Anderson, D. J. Beerling, C. Bolton, T. E. Cerling, J. M. Cotton, J. Da, D. D. Ekart, G. L. Foster, D. R. Greenwood, E. G. Hyland, E. A. Jagniecki, J. P. Jasper, J. B. Kowalczyk, L. Kunzmann, W. M. Kürschner, C. E. Lawrence, C. H. Lear, M. A. Martínez-Botí, D. P. Maxbauer, P. Montagna, B. D. A. Naafs, J. W. B. Rae, M. Raitzsch, G. J. Retallack, S. J. Ring, O. Seki, J. Sepúlveda, A. Sinha, T. F. Tesfamichael, A. Tripati, J. Van Der Burgh, J. Yu, J. C. Zachos, L. Zhang, Toward a Cenozoic history of atmospheric CO_2_. Science 382, eadi5177 (2023).

12. F. I. Woodward, Stomatal numbers are sensitive to increases in CO2 from pre-industrial levels. Nature 327, 617–618 (1987).

13. X. Liang, D. Wang, Q. Ye, J. Zhang, M. Liu, H. Liu, K. Yu, Y. Wang, E. Hou, B. Zhong, L. Xu, T. Lv, S. Peng, H. Lu, P. Sicard, A. Anav, D. S. Ellsworth, Stomatal responses of terrestrial plants to global change. Nature Communications 14, 2188 (2023).

14. L. C. Chua, O. S. Lau, Stomatal development in the changing climate. Development 151, dev202681 (2024).

15. P. L. M. Lang, J. M. Erberich, L. Lopez, C. L. Weiβ, G. Amador, H. F. Fung, S. M. Latorre, J. R. Lasky, H. A. Burbano, M. Expósito-Alonso, D. C. Bergmann, Century-long timelines of herbarium genomes predict plant stomatal response to climate change. Nature Ecology & Evolution 8, 1641–1653 (2024).

16. T. A. Hofmann, W. Atkinson, M. Fan, A. J. Simkin, P. Jindal, T. Lawson, Impact of climate-driven changes in temperature on stomatal anatomy and physiology. Philosophical Transac-tions of the Royal Society B: Biological Sciences 380, 20240244 (2025).

17. J. A. Berry, D. J. Beerling, P. J. Franks, Stomata: Key players in the earth system, past and present. Current Opinion in Plant Biology 13, 232–239 (2010).

18. P. J. Franks, J. A. Berry, D. L. Lombardozzi, G. B. Bonan, Stomatal Function across Temporal and Spatial Scales: Deep-Time Trends, Land-Atmosphere Coupling and Global Models. Plant Physiology 174, 583–602 (2017).

19. P. J. Grubb, “Leaf structure and function” in The Encyclopedia of Ignorance, R. Duncan, M. Weston-Smith, Eds. (Pergamon, Oxford, 1977)vol. 2, pp. 317–330.

20. D. F. Parkhurst, The adaptive significance of stomatal occurrence on one or both surfaces of leaves. The Journal of Ecology 66, 367–383 (1978).

21. K. A. Mott, A. C. Gibson, J. W. O’Leary, The adaptive significance of amphistomatic leaves. Plant, Cell & Environment 5, 455–460 (1982).

22. A. C. Gibson, Structure-Function Relations of Warm Desert Plants (Springer Berlin / Heidel-berg, Berlin, Heidelberg, 1996; http://public.eblib.com/choice/PublicFullRecord.aspx?p=6495 247).

23. W. K. Smith, T. C. Vogelmann, E. H. DeLucia, D. T. Bell, K. A. Shepherd, Leaf Form and Photosynthesis. BioScience 47, 785–793 (1997).

24. R. Oguchi, Y. Onoda, I. Terashima, D. Tholen, “Leaf Anatomy and Function” in The Leaf: A Platform for Performing Photosynthesis, W. W. Adams III, I. Terashima, Eds. (Springer Inter-national Publishing, Cham, 2018; 10.1007/978-3-319-93594-2_5)*Advances in Photosynthesis and Respiration*, pp. 97–139.

25. P. L. Drake, H. J. de Boer, S. J. Schymanski, E. J. Veneklaas, Two sides to every leaf: Water and CO_2_ transport in hypostomatous and amphistomatous leaves. New Phytologist 222, 1179–1187 (2019).

26. P. J. Grubb, “Leaf structure and function” in Unsolved Problems in Ecology, A. Dobson, D. Tilman, R. D. Holt, Eds. (Princeton University Press, Princeton, 2020), pp. 124–144.

27. J. Pospíŝilová, J. Solárová, Environmental and biological control of diffusive conductances of adaxial and abaxial leaf epidermes. Photosynthetica 14, 90–127 (1980).

28. K. A. Mott, J. W. O’Leary, Stomatal Behavior and CO□ Exchange Characteristics in Amphis-tomatous Leaves. Plant Physiology 74, 47–51 (1984).

29. P. B. Reich, A. W. Schoettle, R. G. Amundson, Effects of low concentrations of O3, leaf age and water stress on leaf diffusive conductance and water use efficiency in soybean. Physiologia Plantarum 63, 58–64 (1985).

30. K. A. Mott, Z. G. Cardon, J. A. Berry, Asymmetric patchy stomatal closure for the two surfaces of *Xanthium strumarium* L. Leaves at low humidity. Plant, Cell & Environment 16, 25–34 (1993).

31. S. Wall, S. Vialet□Chabrand, P. Davey, J. Van Rie, A. Galle, J. Cockram, T. Lawson, Stomata on the abaxial and adaxial leaf surfaces contribute differently to leaf gas exchange and photo-synthesis in wheat. New Phytologist 235, 1743–1756 (2022).

32. V. P. Gutschick, Photosynthesis model for C_3_ leaves incorporating CO_2_ transport, propagation of radiation, and biochemistry 2. Ecological and agricultural utility. Photosynthetica 18, 569–595 (1984).

33. D. A. Márquez, H. Stuart□Williams, L. A. Cernusak, G. D. Farquhar, Assessing the <span style=”font-variant:small-caps;”> CO_2_ </span> concentration at the surface of photosynthetic mesophyll cells. New Phytologist 238, 1446–1460 (2023).

34. D. F. Parkhurst, K. A. Mott, Intercellular diffusion limits to CO_2_ uptake in leaves: Studies in air and helox. Plant Physiology 94, 1024–1032 (1990).

35. J. R. Foster, W. K. Smith, Influence of stomatal distribution on transpiration in low-wind envi-ronments. Plant, Cell and Environment 9, 751–759 (1986).

36. C. D. Muir, Is amphistomy an adaptation to high light? Optimality models of stomatal traits along light gradients. Integrative and Comparative Biology 59, 571–584 (2019).

37. J. G. Wood, The physiology of xerophytism in Australian plants: The stomatal frequencies, tran-spiration and osmotic pressures of sclerophyll and tomentose-succulent leaved plants. Journal of Ecology 22, 69–87 (1934).

38. J. T. Howell, Concerning stomata on leaves in *a*rctostaphylos. The Wasmann Collector 6, 57–65 (1945).

39. E. J. Salisbury, I. On the causes and ecological significance of stomatal frequency, with special reference to the woodland flora. Philosophical Transactions of the Royal Society of London. Series B, Containing Papers of a Biological Character 216, 1–65 (1928).

40. H. J. Peat, A. H. Fitter, A comparative study of the distribution and density of stomata in the British flora. Biological Journal of the Linnean Society 52, 377–393 (1994).

41. G. J. Jordan, R. J. Carpenter, T. J. Brodribb, Using fossil leaves as evidence for open vegetation. Palaeogeography, Palaeoclimatology, Palaeoecology 395, 168–175 (2014).

42. S. F. Bucher, K. Auerswald, C. Grün-Wenzel, S. I. Higgins, J. Garcia Jorge, C. Römermann, Stomatal traits relate to habitat preferences of herbaceous species in a temperate climate. Flora 229, 107–115 (2017).

43. C. D. Muir, Light and growth form interact to shape stomatal ratio among British angiosperms. New Phytologist 218, 242–252 (2018).

44. G. Triplett, A. S. David, Stomatal distribution andpost□fire recovery: Intra□ and interspecific variation in plants of the pyrogenic Florida scrub. American Journal of Botany, e70050 (2025).

45. O. B. Lyshede, Comparative and functional leaf anatomy of selected Alstroemeriaceae of mainly Chilean origin. Botanical Journal of the Linnean Society 140, 261–272 (2002).

46. G. Triplett, T. N. Buckley, C. D. Muir, Amphistomy increases leaf photosynthesis more in coastal than montane plants of Hawaiian ‘ilima (*Sida fallax*). American Journal of Botany 111, e16284 (2024).

47. H. Poorter, Ü. Niinemets, N. Ntagkas, A. Siebenkäs, M. Mäenpää, S. Matsubara, T. L. Pons, A meta□analysis of plant responses to light intensity for 70 traits ranging from molecules to whole plant performance. New Phytologist 223, 1073–1105 (2019).

48. T. J. Givnish, R. A. Montgomery, Common-garden studies on adaptive radiation of photosyn-thetic physiology among Hawaiian lobeliads. Proceedings of the Royal Society B: Biological Sciences 281, 20132944–20132944 (2014).

49. B. M. J. Engelbrecht, H. M. Herz, Evaluation of different methods to estimate understorey light conditions in tropical forests. Journal of Tropical Ecology 17, 207–224 (2001).

50. I. E. Peralta, D. M. Spooner, S. Knapp, Taxonomy of wild tomatoes and their relatives (*solanum* sect. Lycopersicoides, sect. Juglandfolia, sect. Lycopersicon; Solanaceae). 84 (2008).

51. P. Burns, C. R. Hakkenberg, S. J. Goetz, Multi-resolution gridded maps of vegetation structure from GEDI. Scientific Data 11, 881 (2024).

52. G. P. John, C. Scoffoni, T. N. Buckley, R. Villar, H. Poorter, L. Sack, The anatomical and compositional basis of leaf mass per area. Ecology Letters 20, 412–425 (2017).

53. H. G. Jones, R. O. Slatyer, Effects of Intercellular Resistances on Estimates of the Intracellular Resistance to Co2 Uptake by Plant Leaves. Australian Journal of Biological Sciences 25, 443 (1972).

54. J. B. Pease, D. C. Haak, M. W. Hahn, L. C. Moyle, Phylogenomics reveals three sources of adaptive variation during a rapid radiation. PLOS Biology 14, e1002379 (2016).

55. P.-G. Schoch, C. Zinsou, M. Sibi, Dependence of the stomatal index on environmental factors during stomatal differentiation in leaves of *Vigna sinensis* L.: 1. Effect of light intensity. Jour-nal of Experimental Botany 31, 1211–1216 (1980).

56. L. Sack, T. N. Buckley, The developmental basis of stomatal density and flux. Plant Physiology 171, 2358–2363 (2016).

57. K. A. Mott, O. Michaelson, Amphistomy as an adaptation to high light intensity in *Ambrosia cordifolia* (Compositae). American Journal of Botany 78, 76–79 (1991).

58. J. Schindelin, I. Arganda-Carreras, E. Frise, V. Kaynig, M. Longair, T. Pietzsch, S. Preibisch, C. Rueden, S. Saalfeld, B. Schmid, J.-Y. Tinevez, D. J. White, V. Hartenstein, K. Eliceiri, P. Tomancak, A. Cardona, Fiji: An open-source platform for biological-image analysis. Nature Methods 9, 676–682 (2012).

59. J. L. Monteith, M. H. Unsworth, Principles of Environmental Physics: Plants, Animals, and the Atmosphere (Elsevier/Academic Press, Amsterdam; Boston, 4th ed., 2013).

60. Stan Development Team, Stan Modeling Language Users Guide and Reference Manual (2025; https://mc-stan.org).

61. P.-C. Bürkner, **Brms** : An *r* Package for Bayesian Multilevel Models Using *stan*. Journal of Statistical Software 80 (2017).

62. J. Gabry, R. Češnovar, A. Johnson, S. Bronder, Cmdstanr: R Interface to ‘CmdStan’ (2025; https://mc-stan.org/cmdstanr, https://discourse.mc-stan.org).

63. R Core Team, R: A Language and Environment for Statistical Computing (R Foundation for Statistical Computing, Vienna, Austria, 2025; http://www.R-project.org/).

64. A. Gelman, D. B. Rubin, Inference from iterative simulation using multiple sequences. Statisti-cal Science 7, 457–472 (1992).

65. A. Vehtari, A. Gelman, J. Gabry, Practical Bayesian model evaluation using leave-one-out cross-validation and WAIC. Statistics and Computing 27, 1413–1432 (2017).

