## Supplementary material for "Plasticity and adaptation to high light intensity amplify the advantage of amphistomatous leaves": Figure S3

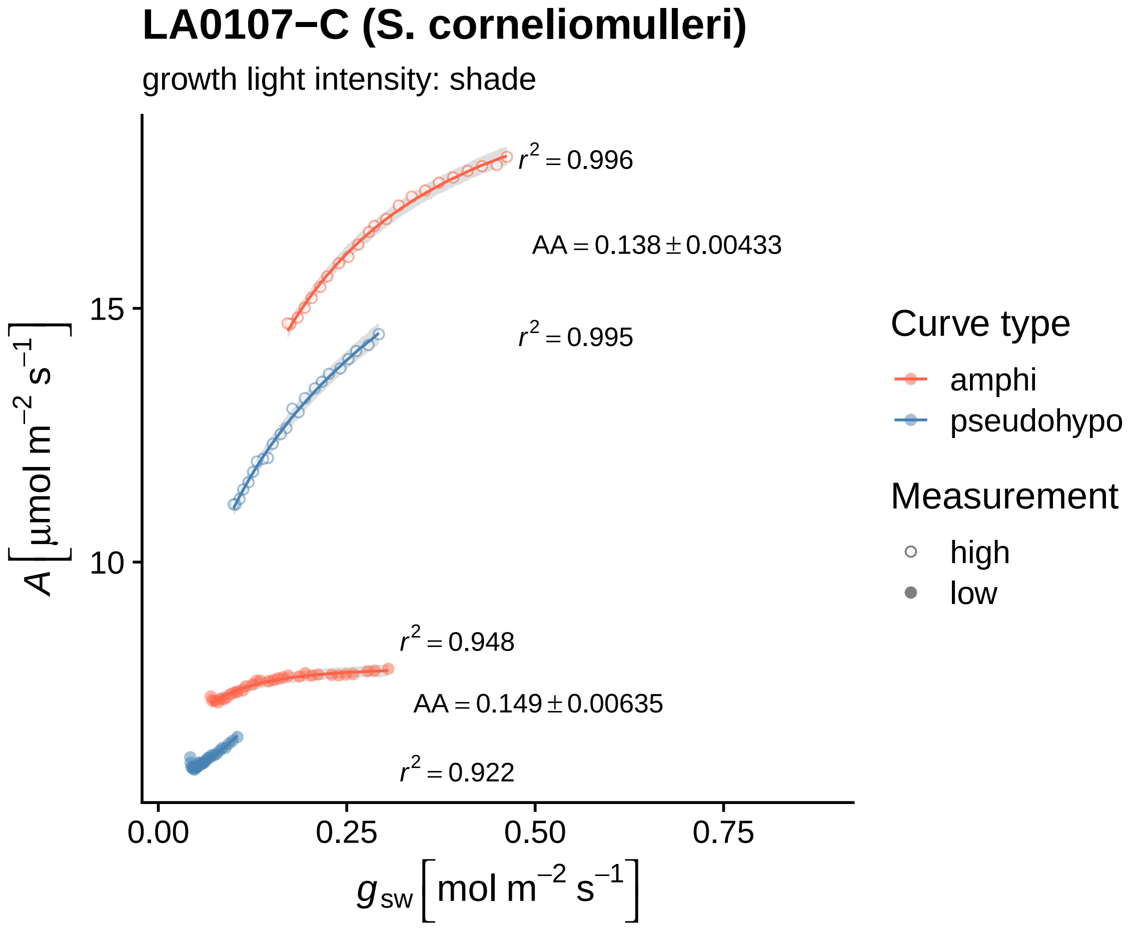
LA0107−C (S. corneliomulleri)
growth light intensity: shade
$r^2 = 0.996$
AA = 0.138 ± 0.00433
$r^2 = 0.995$
$r^2 = 0.948$
AA = 0.149 ± 0.00635
$r^2 = 0.922$
15
10
0.00
0.25
0.50
0.75
$A\ [\mu mol\ m^{-2}\ s^{-1}]$
$g_{sw}\ [mol\ m^{-2}\ s^{-1}]$
Curve type
amphi
pseudohypo
Measurement
high
low

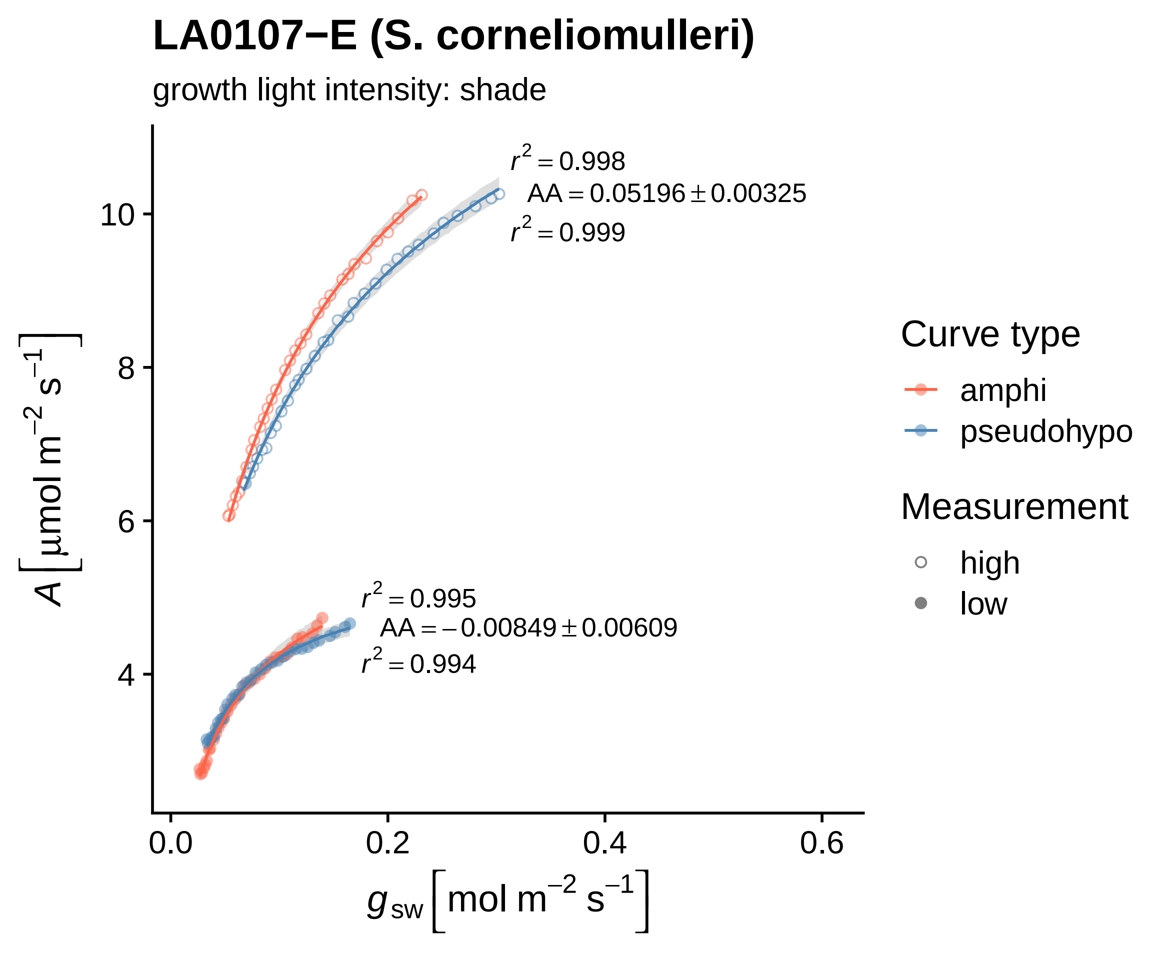

LA0107−E (S. corneliomulleri)
growth light intensity: shade
$r^2 = 0.998$
AA = 0.05196 ± 0.00325
$r^2 = 0.999$
$r^2 = 0.995$
AA = −0.00849 ± 0.00609
$r^2 = 0.994$
$A\left[\mu mol\ m^{-2}\ s^{-1}\right]$
10
8
6
4
0.0
0.2
0.4
0.6
$g_{sw}\left[mol\ m^{-2}\ s^{-1}\right]$
Curve type
amphi
pseudohypo
Measurement
high
low

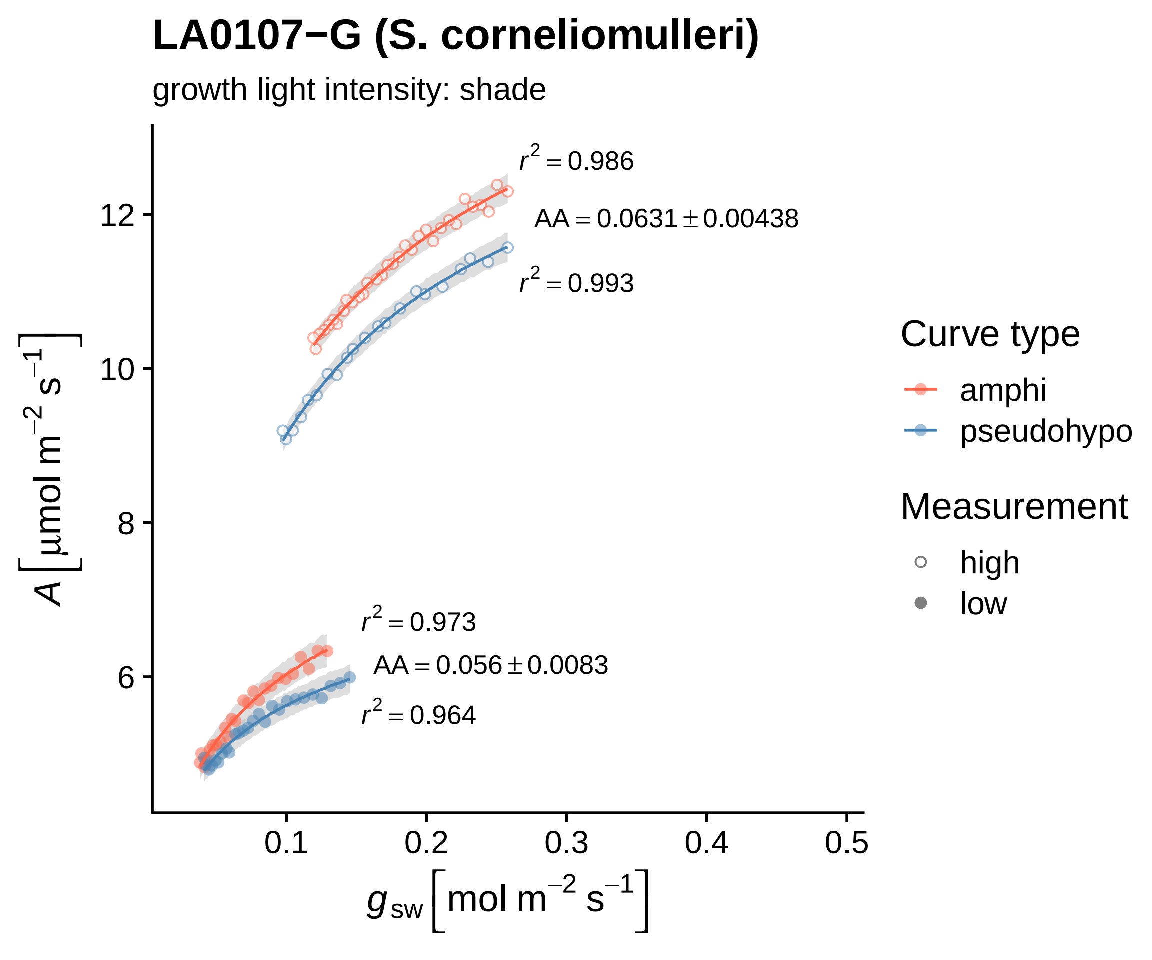
LA0107−G (S. corneliomulleri)
growth light intensity: shade
$r^2 = 0.986$
$AA = 0.0631 \pm 0.00438$
$r^2 = 0.993$
$r^2 = 0.973$
$AA = 0.056 \pm 0.0083$
$r^2 = 0.964$
$A\ [\mu mol\ m^{-2}\ s^{-1}]$
12
10
8
6
0.1
0.2
0.3
0.4
0.5
$g_{sw}\ [mol\ m^{-2}\ s^{-1}]$
Curve type
amphi
pseudohypo
Measurement
high
low

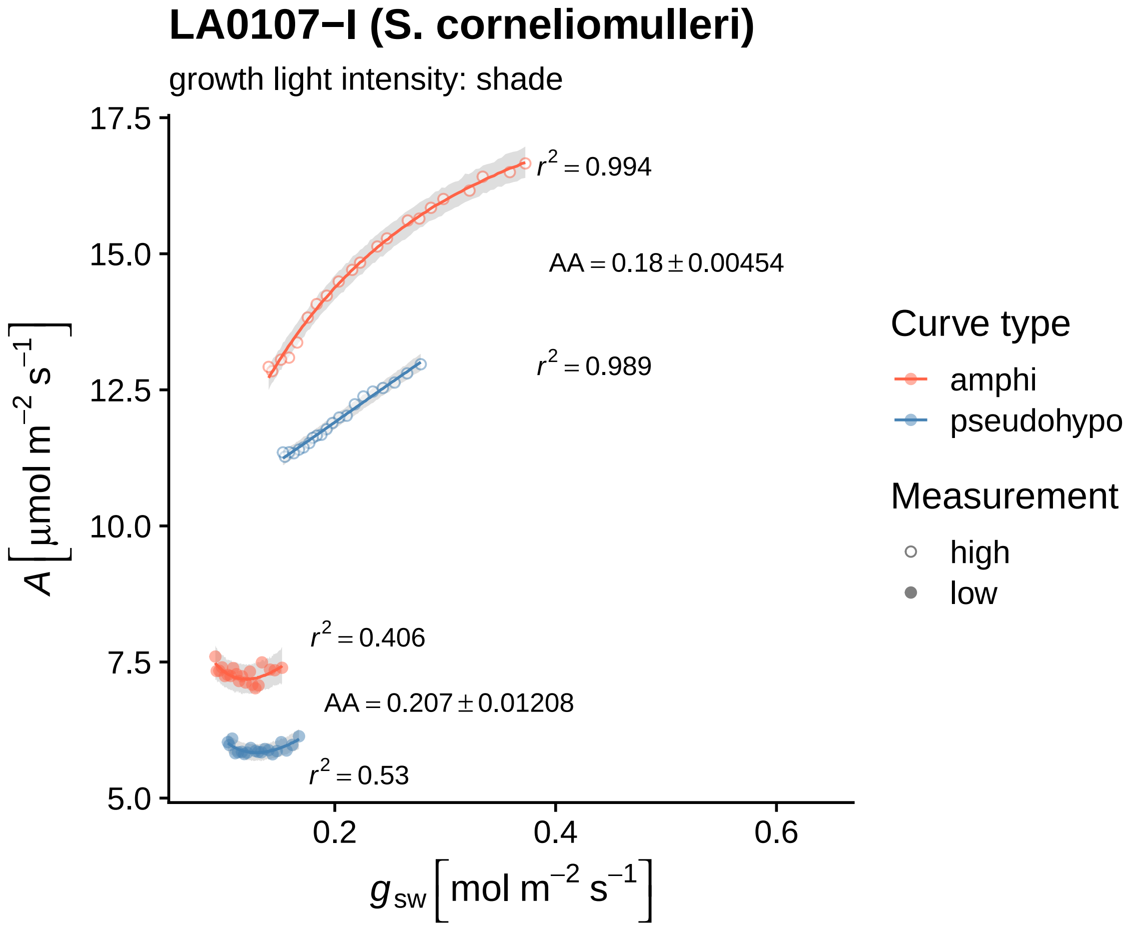

LA0107−I (S. corneliomulleri)
growth light intensity: shade
17.5
15.0
12.5
10.0
7.5
5.0
0.2
0.4
0.6
$A\left[\mu mol\ m^{-2}\ s^{-1}\right]$
$g_{sw}\left[mol\ m^{-2}\ s^{-1}\right]$
$r^2 = 0.994$
$AA = 0.18 \pm 0.00454$
$r^2 = 0.989$
$r^2 = 0.406$
$AA = 0.207 \pm 0.01208$
$r^2 = 0.53$
Curve type
amphi
pseudohypo
Measurement
high
low

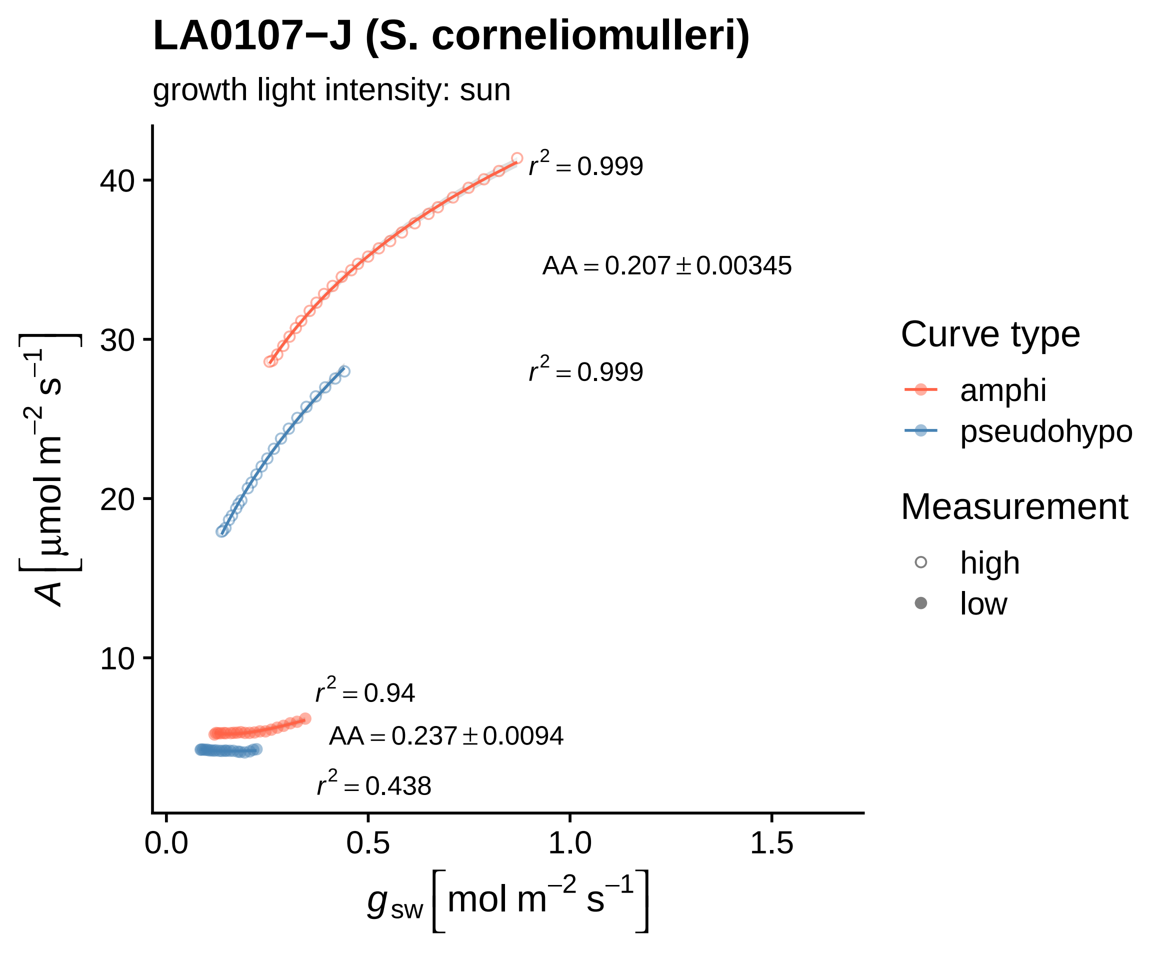

LA0107−J (S. corneliomulleri)
growth light intensity: sun
$r^2 = 0.999$
AA = 0.207 ± 0.00345
$r^2 = 0.999$
$r^2 = 0.94$
AA = 0.237 ± 0.0094
$r^2 = 0.438$
40
30
20
10
0.0
0.5
1.0
1.5
$A\ [\mu mol\ m^{-2}\ s^{-1}]$
$g_{sw}\ [mol\ m^{-2}\ s^{-1}]$
Curve type
amphi
pseudohypo
Measurement
high
low

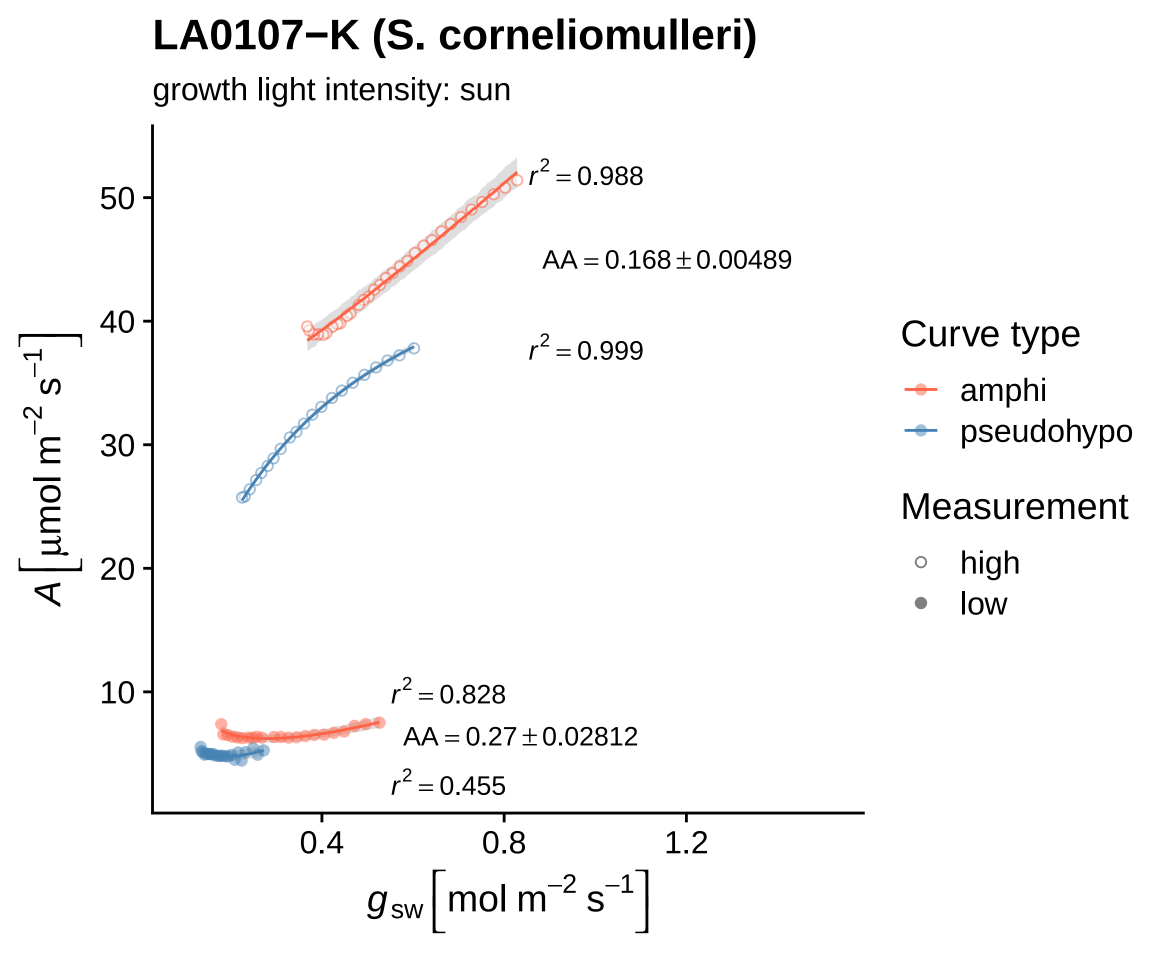
LA0107−K (S. corneliomulleri)
growth light intensity: sun
r^2 = 0.988
AA = 0.168 ± 0.00489
r^2 = 0.999
r^2 = 0.828
AA = 0.27 ± 0.02812
r^2 = 0.455
A [μmol m^−2 s^−1]
g_sw [mol m^−2 s^−1]
10
20
30
40
50
0.4
0.8
1.2
Curve type
amphi
pseudohypo
Measurement
high
low

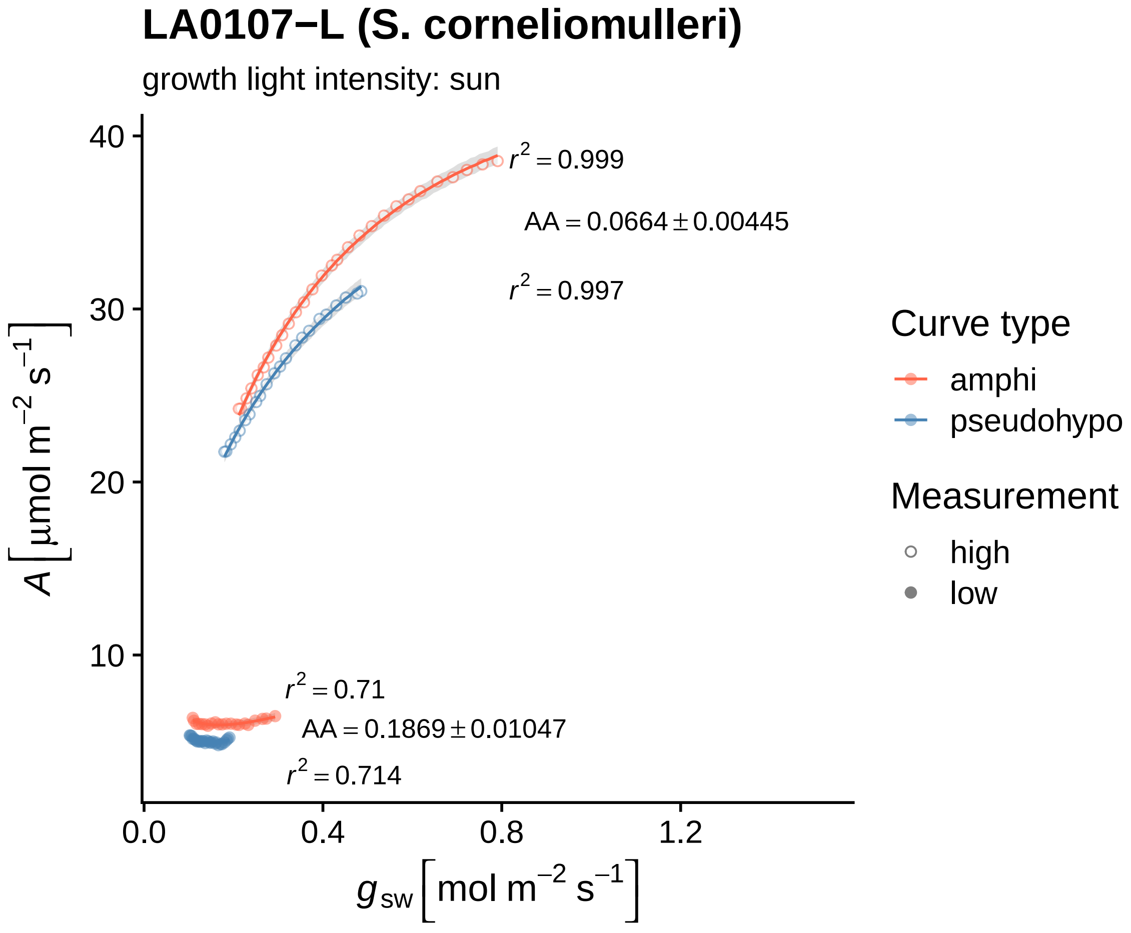
LA0107−L (S. corneliomulleri)
growth light intensity: sun
$r^2 = 0.999$
$AA = 0.0664 \pm 0.00445$
$r^2 = 0.997$
$r^2 = 0.71$
$AA = 0.1869 \pm 0.01047$
$r^2 = 0.714$
40
30
20
10
0.0
0.4
0.8
1.2
$A\ [\mu mol\ m^{-2}\ s^{-1}]$
$g_{sw}\ [mol\ m^{-2}\ s^{-1}]$
Curve type
amphi
pseudohypo
Measurement
high
low

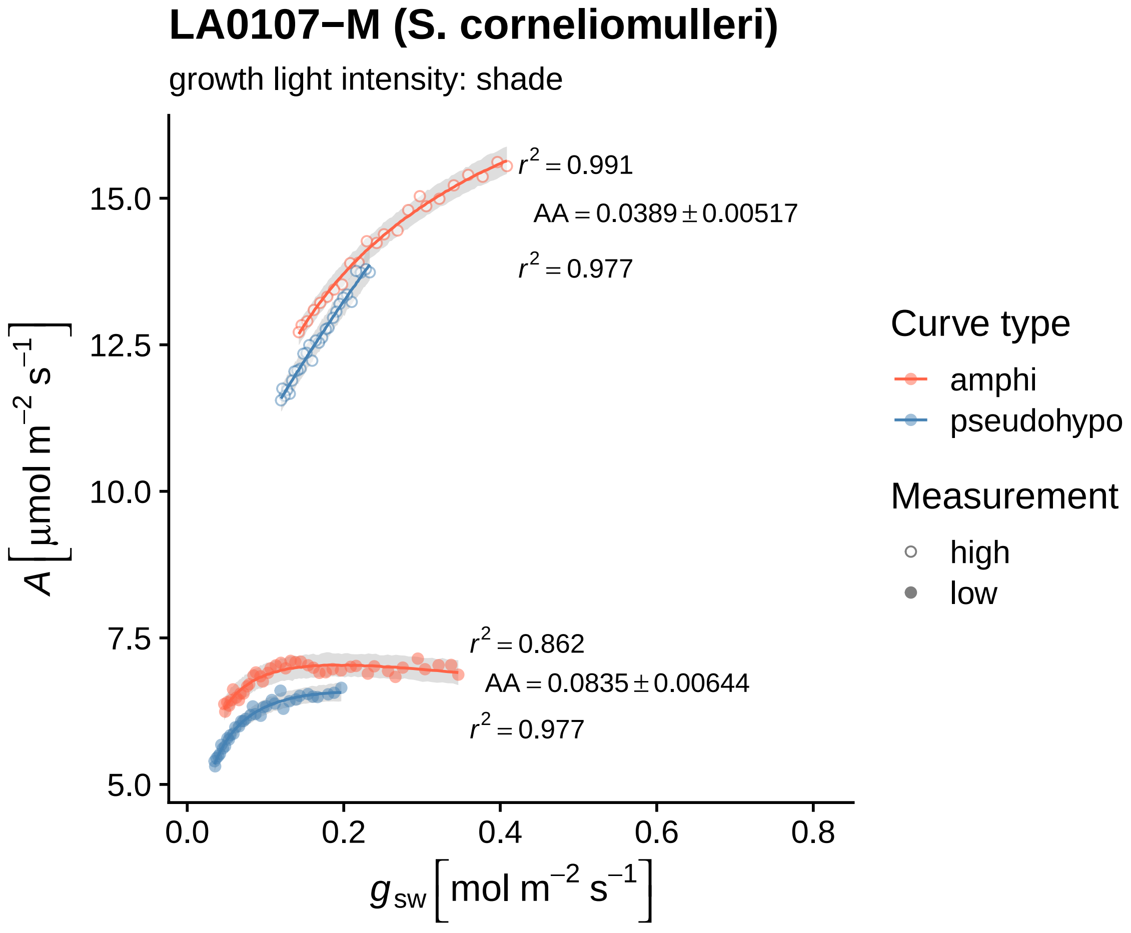
LA0107−M (S. corneliomulleri)
growth light intensity: shade
$r^2 = 0.991$
$AA = 0.0389 \pm 0.00517$
$r^2 = 0.977$
$r^2 = 0.862$
$AA = 0.0835 \pm 0.00644$
$r^2 = 0.977$
$A\ [\mu mol\ m^{-2}\ s^{-1}]$
$g_{sw}\ [mol\ m^{-2}\ s^{-1}]$
15.0
12.5
10.0
7.5
5.0
0.0
0.2
0.4
0.6
0.8
Curve type
amphi
pseudohypo
Measurement
high
low

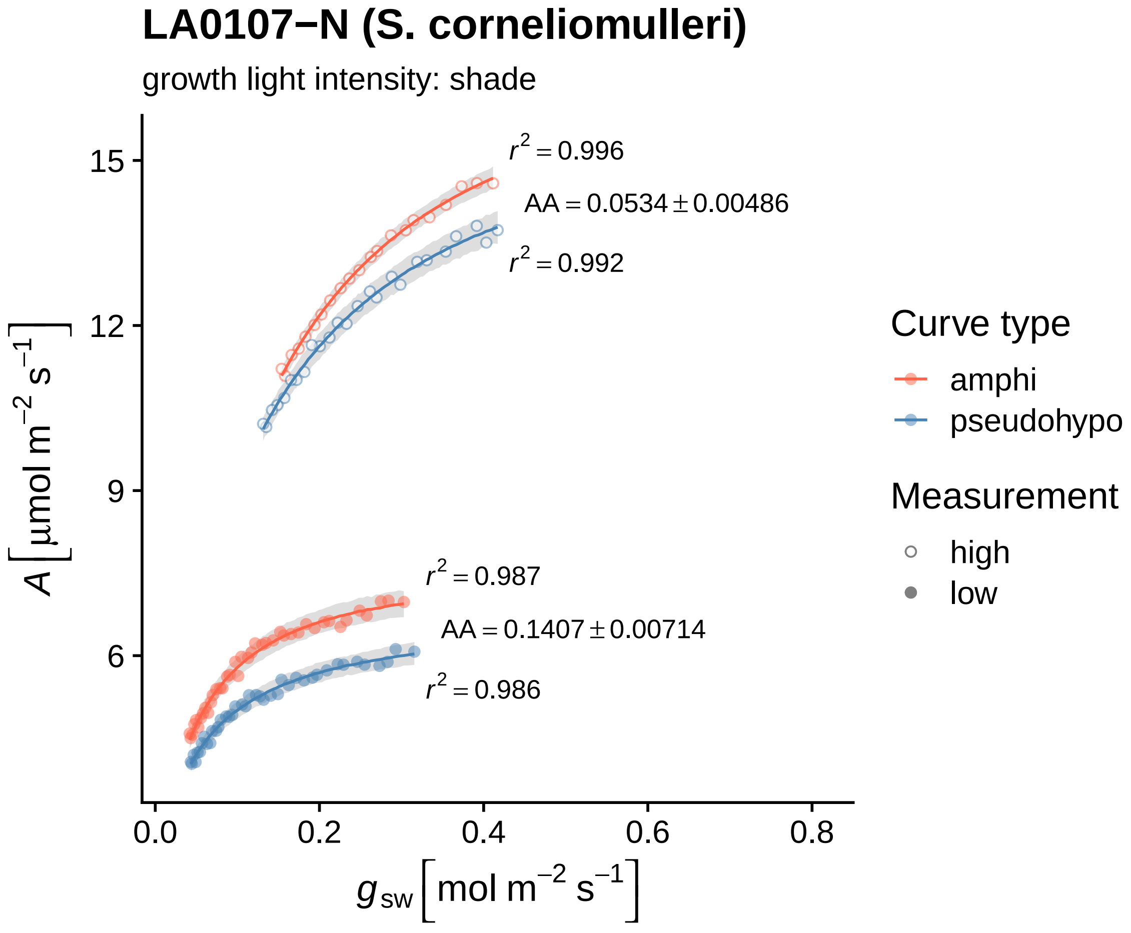

LA0107−N (S. corneliomulleri)
growth light intensity: shade
r² = 0.996
AA = 0.0534 ± 0.00486
r² = 0.992
r² = 0.987
AA = 0.1407 ± 0.00714
r² = 0.986
15
12
9
6
A [µmol m⁻² s⁻¹]
0.0
0.2
0.4
0.6
0.8
g_sw [mol m⁻² s⁻¹]
Curve type
amphi
pseudohypo
Measurement
high
low

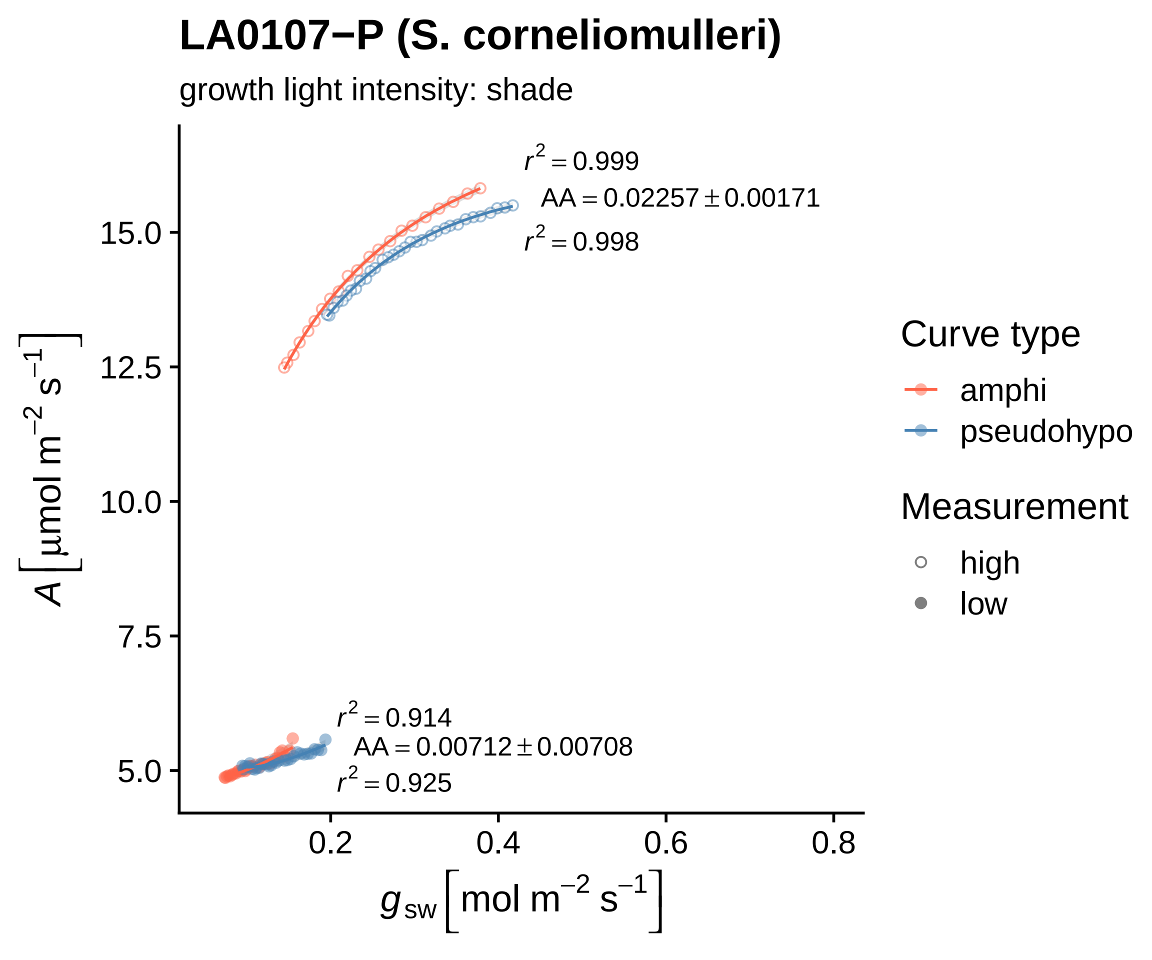

LA0107−P (S. corneliomulleri)
growth light intensity: shade
r² = 0.999
AA = 0.02257 ± 0.00171
r² = 0.998
r² = 0.914
AA = 0.00712 ± 0.00708
r² = 0.925
15.0
12.5
10.0
7.5
5.0
0.2
0.4
0.6
0.8
A [μmol m⁻² s⁻¹]
g sw [mol m⁻² s⁻¹]
Curve type
amphi
pseudohypo
Measurement
high
low

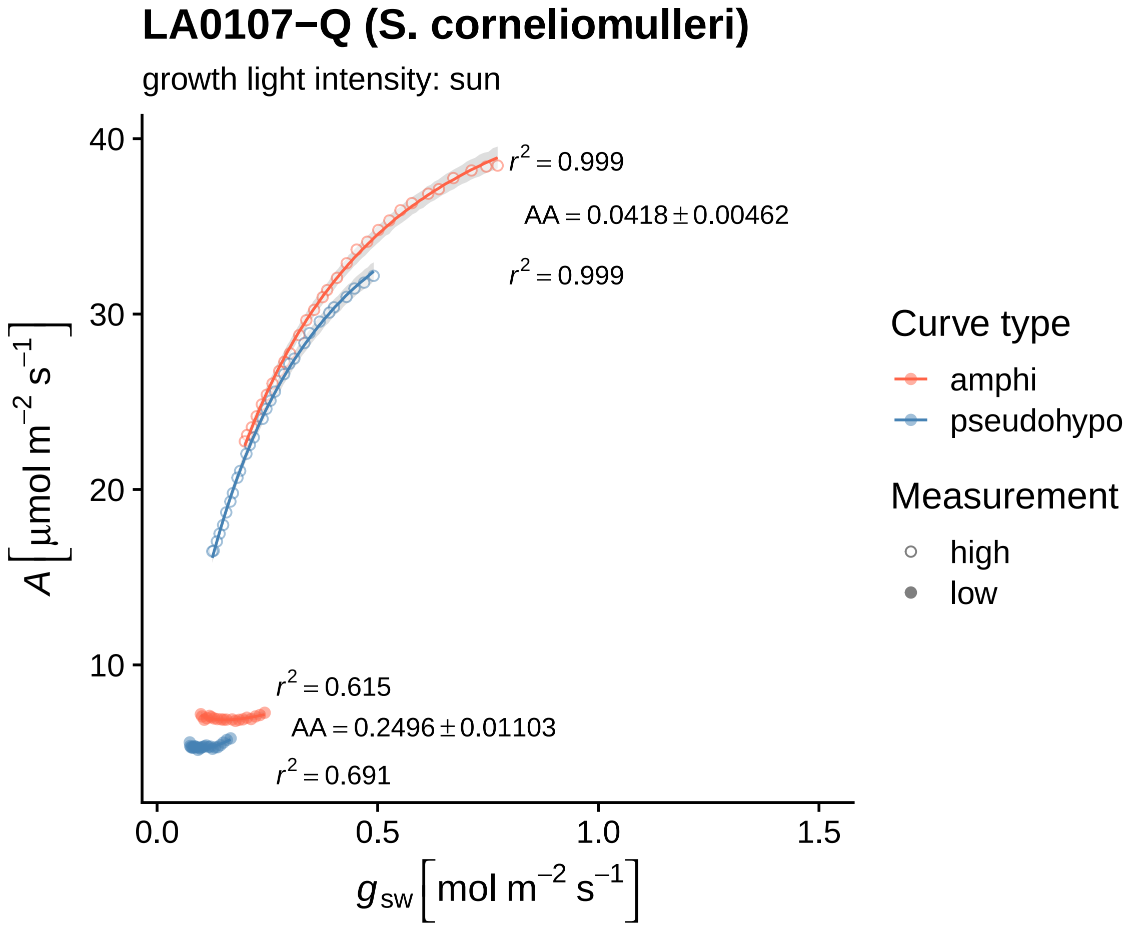

LA0107−Q (S. corneliomulleri)
growth light intensity: sun
$r^2 = 0.999$
$AA = 0.0418 \pm 0.00462$
$r^2 = 0.999$
$r^2 = 0.615$
$AA = 0.2496 \pm 0.01103$
$r^2 = 0.691$
40
30
20
10
0.0
0.5
1.0
1.5
$A\ [\mu mol\ m^{-2}\ s^{-1}]$
$g_{sw}\ [mol\ m^{-2}\ s^{-1}]$
Curve type
amphi
pseudohypo
Measurement
high
low

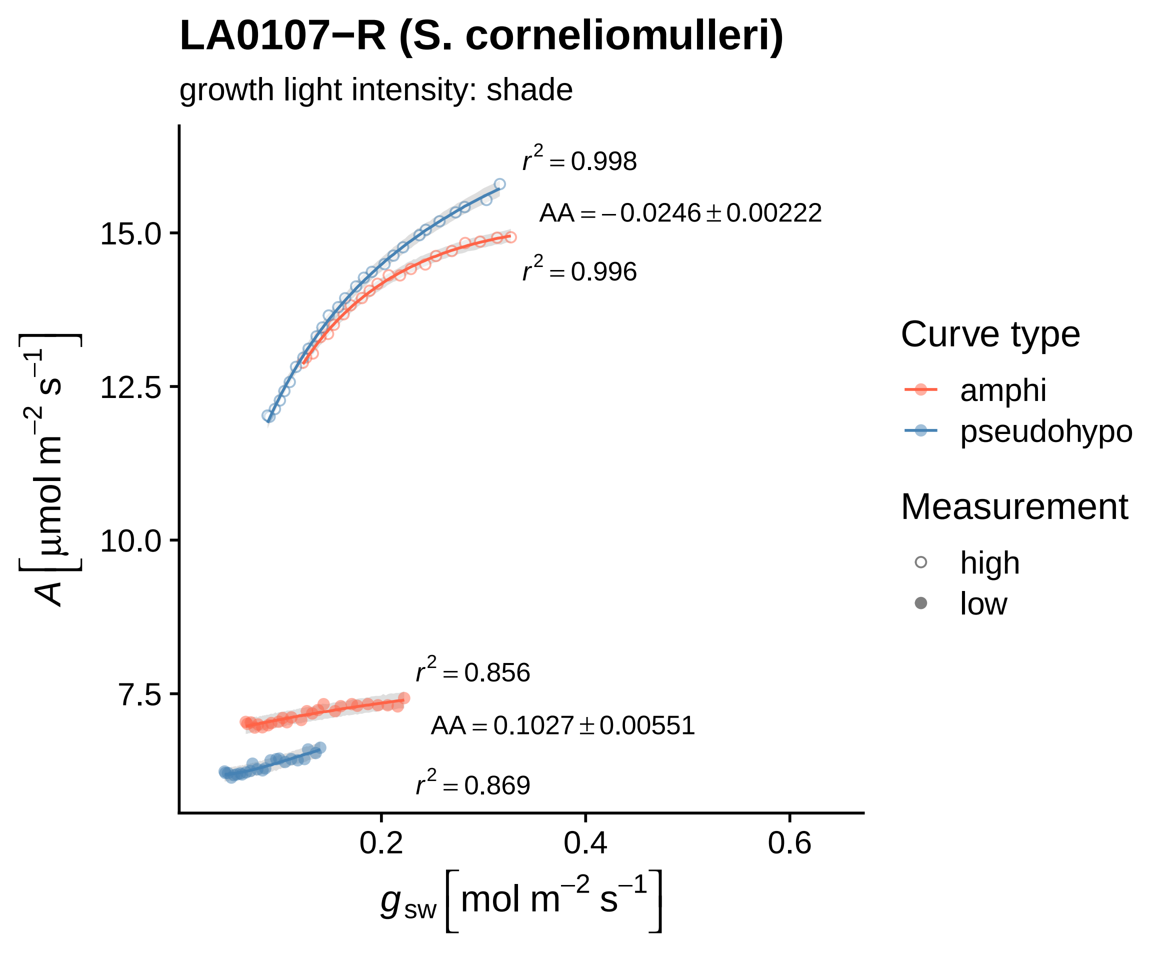

LA0107−R (S. corneliomulleri)
growth light intensity: shade
r² = 0.998
AA = −0.0246 ± 0.00222
r² = 0.996
r² = 0.856
AA = 0.1027 ± 0.00551
r² = 0.869
15.0
12.5
10.0
7.5
0.2
0.4
0.6
A [μmol m⁻² s⁻¹]
g_sw [mol m⁻² s⁻¹]
Curve type
amphi
pseudohypo
Measurement
high
low

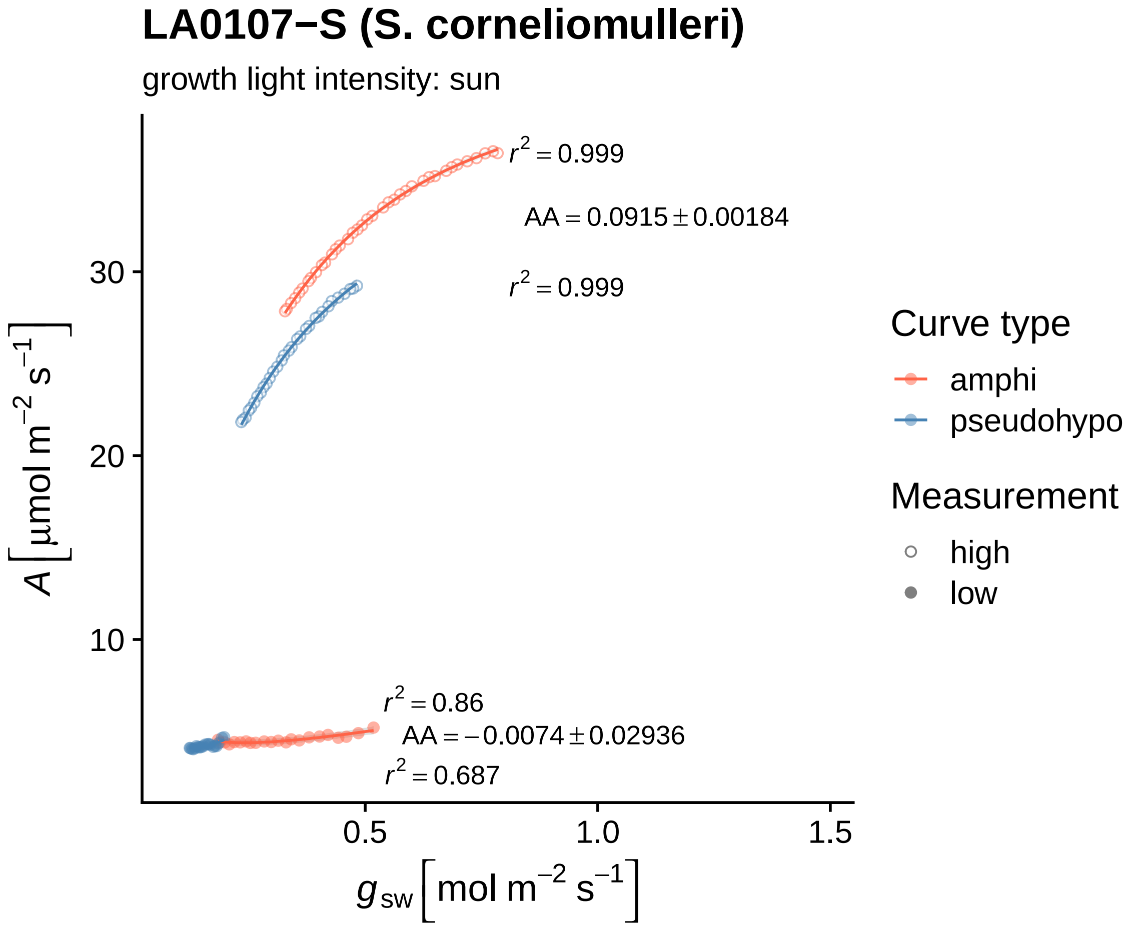
LA0107−S (S. corneliomulleri)
growth light intensity: sun
$r^2 = 0.999$
$AA = 0.0915 \pm 0.00184$
$r^2 = 0.999$
$r^2 = 0.86$
$AA = -0.0074 \pm 0.02936$
$r^2 = 0.687$
30
20
10
0.5
1.0
1.5
$A\ [\mu mol\ m^{-2}\ s^{-1}]$
$g_{sw}\ [mol\ m^{-2}\ s^{-1}]$
Curve type
amphi
pseudohypo
Measurement
high
low

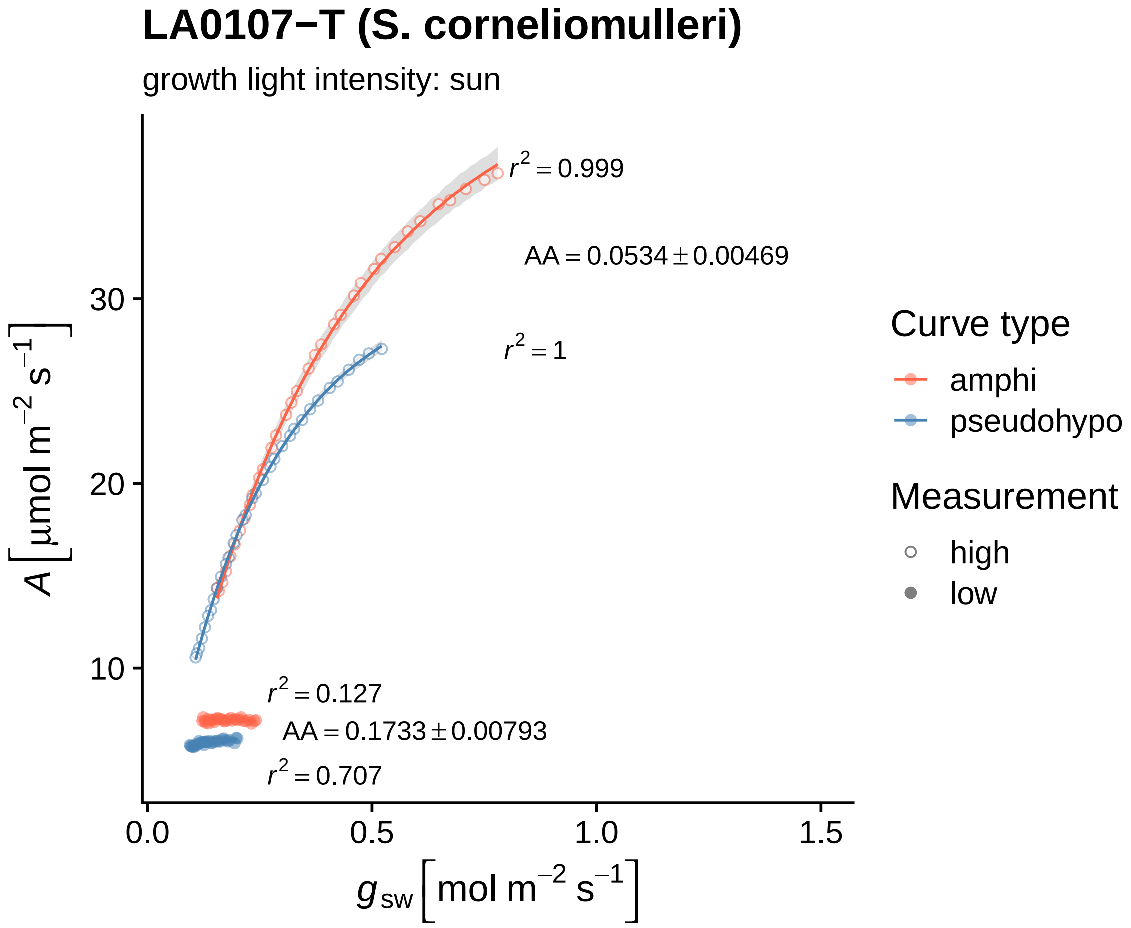
LA0107−T (S. corneliomulleri)
growth light intensity: sun
$r^2 = 0.999$
$AA = 0.0534 \pm 0.00469$
$r^2 = 1$
$r^2 = 0.127$
$AA = 0.1733 \pm 0.00793$
$r^2 = 0.707$
$A$ [µmol m$^{-2}$ s$^{-1}$]
10
20
30
0.0
0.5
1.0
1.5
$g_{sw}$ [mol m$^{-2}$ s$^{-1}$]
Curve type
amphi
pseudohypo
Measurement
high
low

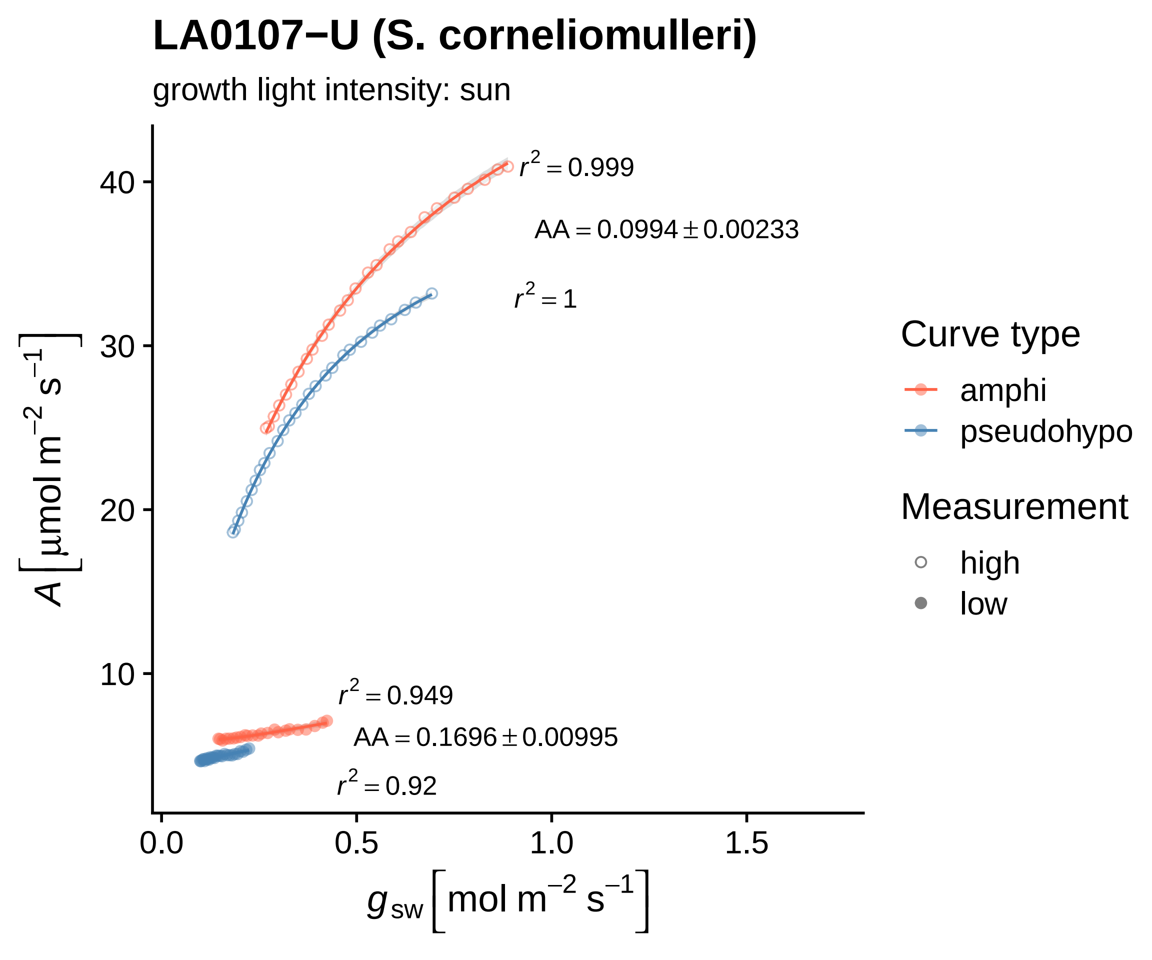
LA0107−U (S. corneliomulleri)
growth light intensity: sun
$r^2 = 0.999$
$AA = 0.0994 \pm 0.00233$
$r^2 = 1$
$r^2 = 0.949$
$AA = 0.1696 \pm 0.00995$
$r^2 = 0.92$
40
30
20
10
0.0
0.5
1.0
1.5
$A\left[\mu mol\ m^{-2}\ s^{-1}\right]$
$g_{sw}\left[mol\ m^{-2}\ s^{-1}\right]$
Curve type
amphi
pseudohypo
Measurement
high
low

### LA0107−V (S. corneliomulleri)

growth light intensity: shade

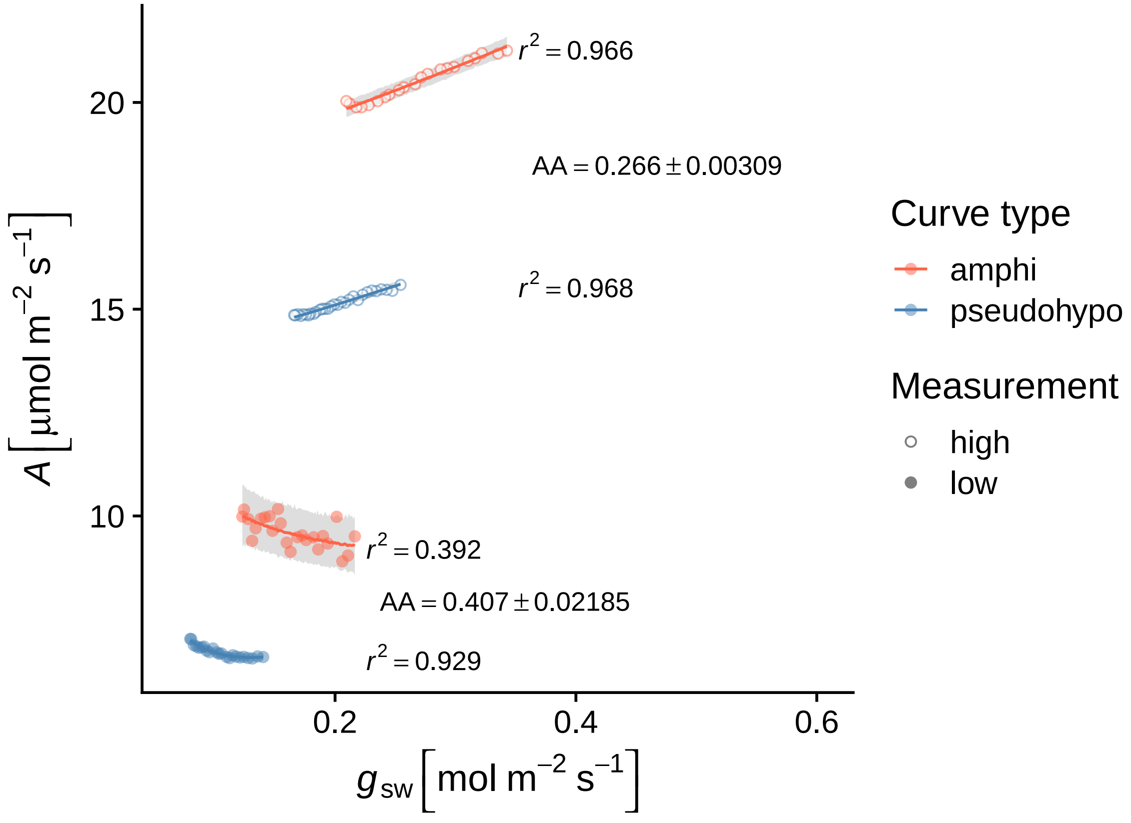

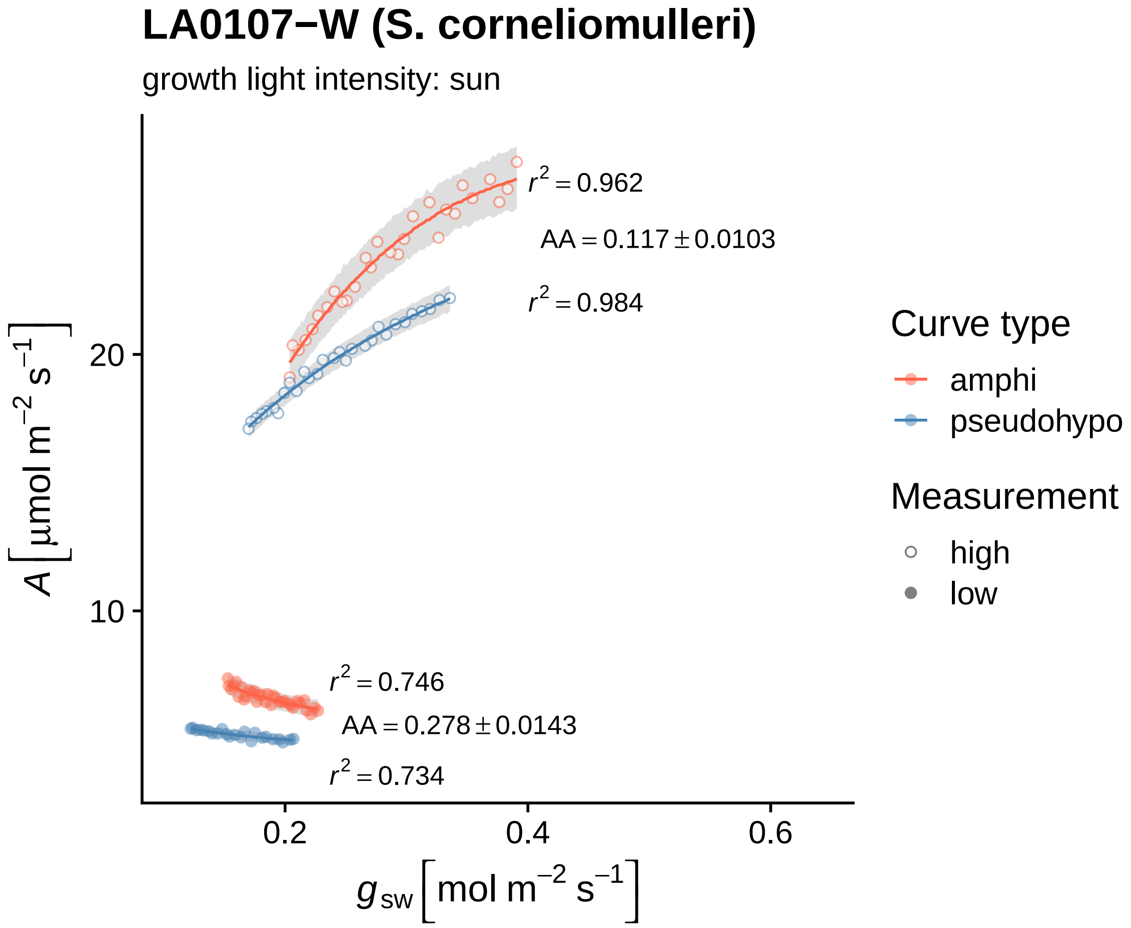

LA0107−W (S. corneliomulleri)
growth light intensity: sun
$r^2 = 0.962$
$AA = 0.117 \pm 0.0103$
$r^2 = 0.984$
$r^2 = 0.746$
$AA = 0.278 \pm 0.0143$
$r^2 = 0.734$
$A\,[\mu mol\ m^{-2}\ s^{-1}]$
20
10
0.2
0.4
0.6
$g_{sw}\,[mol\ m^{-2}\ s^{-1}]$
Curve type
amphi
pseudohypo
Measurement
high
low

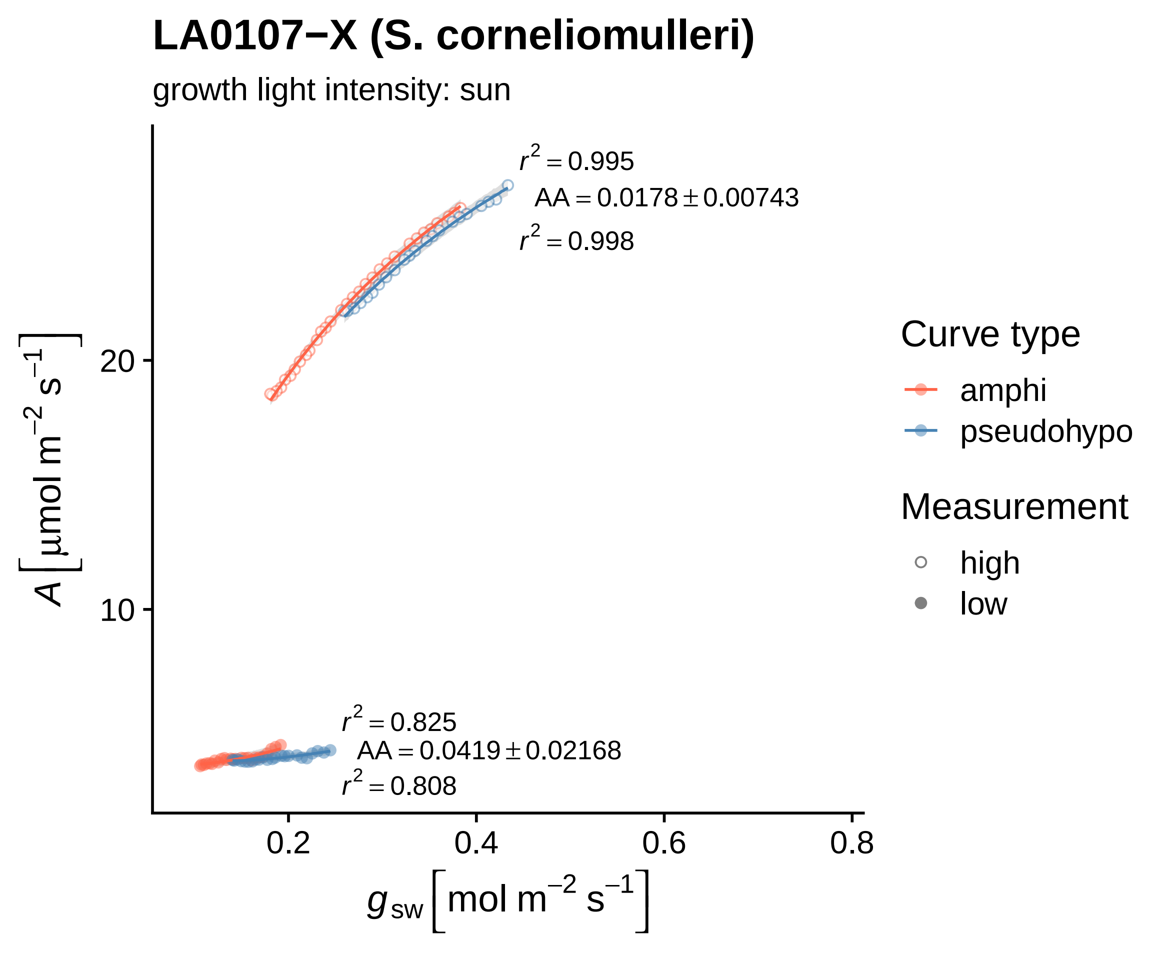

LA0107−X (S. corneliomulleri)
growth light intensity: sun
$r^2 = 0.995$
$AA = 0.0178 \pm 0.00743$
$r^2 = 0.998$
$r^2 = 0.825$
$AA = 0.0419 \pm 0.02168$
$r^2 = 0.808$
20
10
0.2
0.4
0.6
0.8
$A\,[\mu mol\ m^{-2}\ s^{-1}]$
$g_{sw}\,[mol\ m^{-2}\ s^{-1}]$
Curve type
amphi
pseudohypo
Measurement
high
low

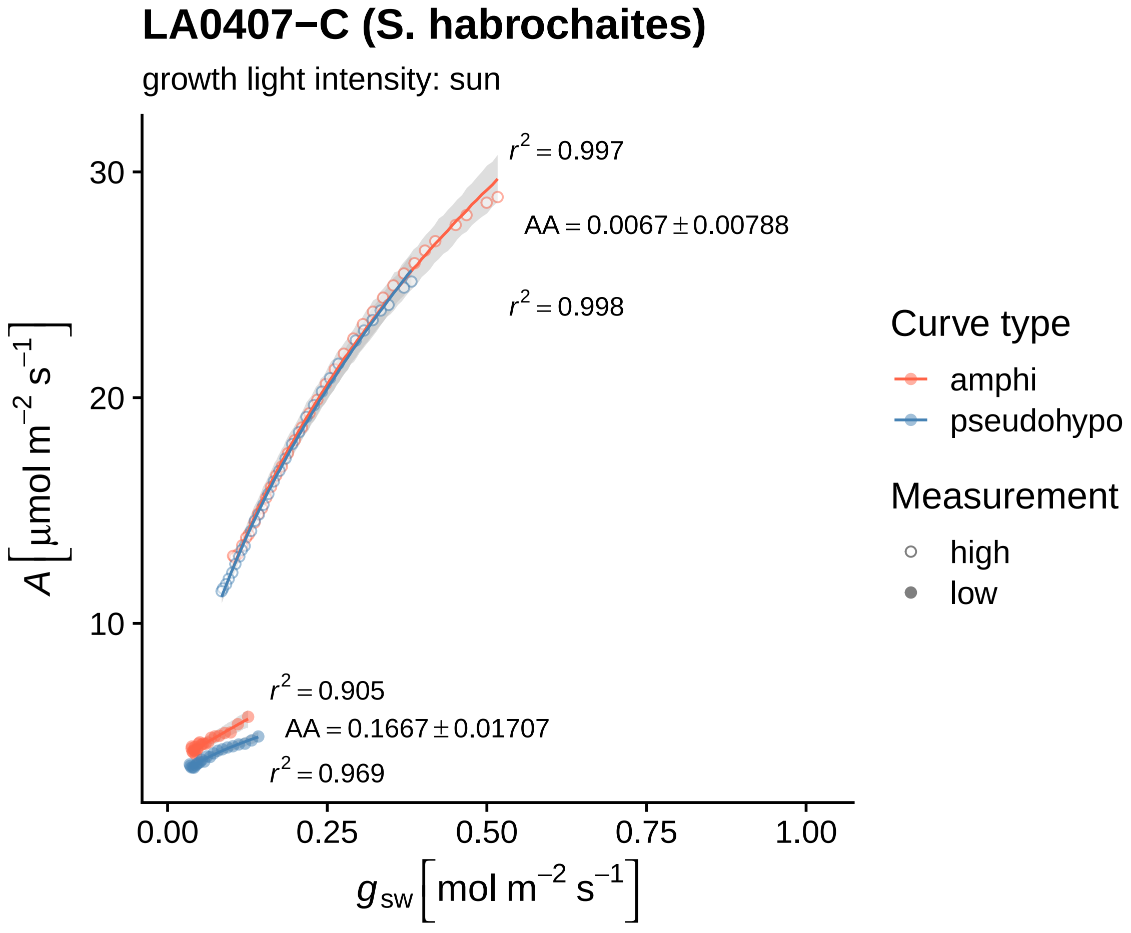

LA0407−C (S. habrochaites)
growth light intensity: sun
$r^2 = 0.997$
$AA = 0.0067 \pm 0.00788$
$r^2 = 0.998$
$r^2 = 0.905$
$AA = 0.1667 \pm 0.01707$
$r^2 = 0.969$
30
20
10
0.00
0.25
0.50
0.75
1.00
$A \left[ \mu mol \; m^{-2} \; s^{-1} \right]$
$g_{sw} \left[ mol \; m^{-2} \; s^{-1} \right]$
Curve type
amphi
pseudohypo
Measurement
high
low

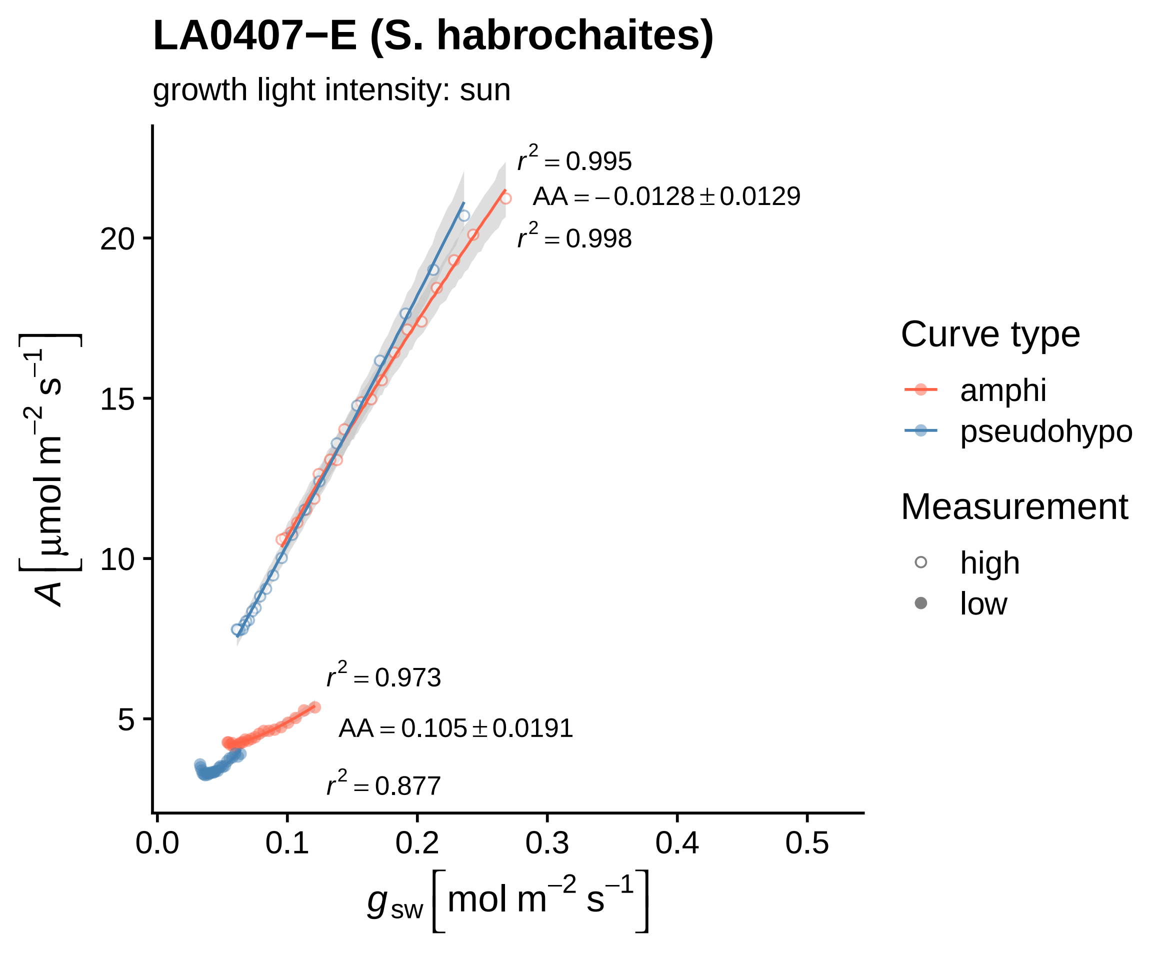
LA0407−E (S. habrochaites)
growth light intensity: sun
r² = 0.995
AA = −0.0128 ± 0.0129
r² = 0.998
r² = 0.973
AA = 0.105 ± 0.0191
r² = 0.877
20
15
10
5
0.0
0.1
0.2
0.3
0.4
0.5
A [µmol m⁻² s⁻¹]
g sw [mol m⁻² s⁻¹]
Curve type
amphi
pseudohypo
Measurement
high
low

#### LA0407−H (S. habrochaites)

growth light intensity: shade

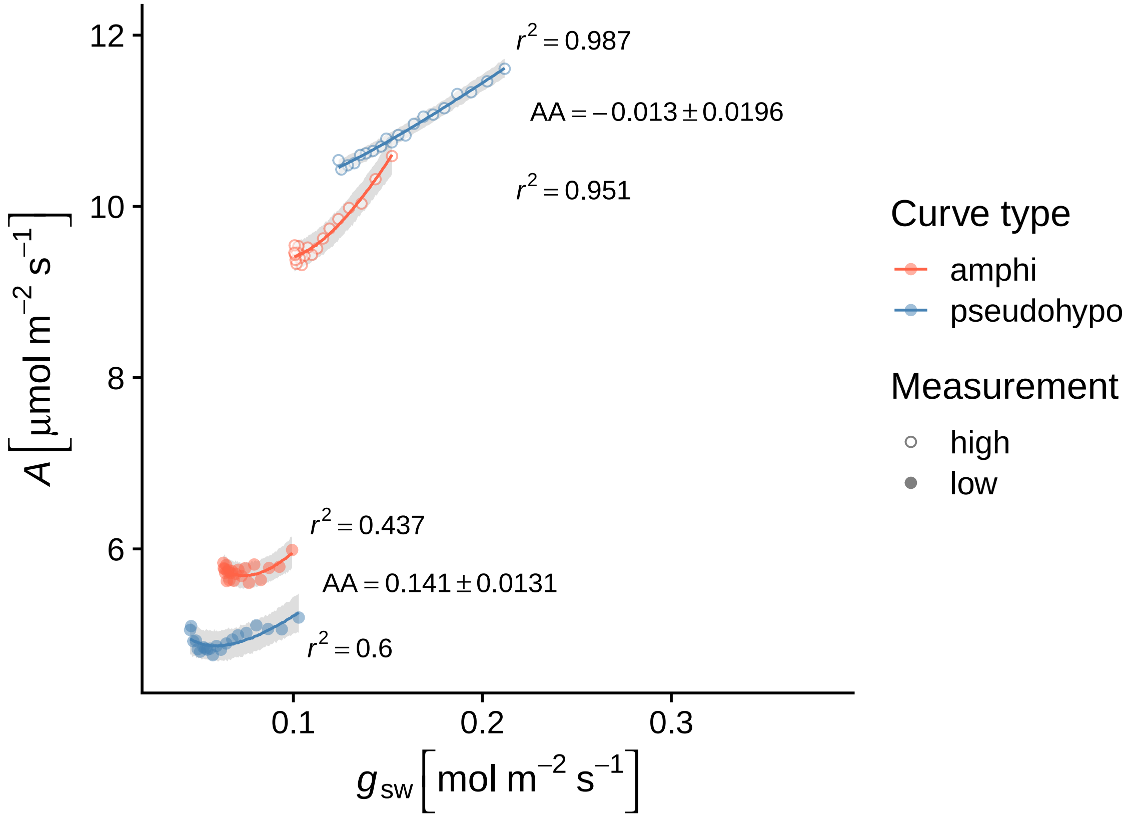

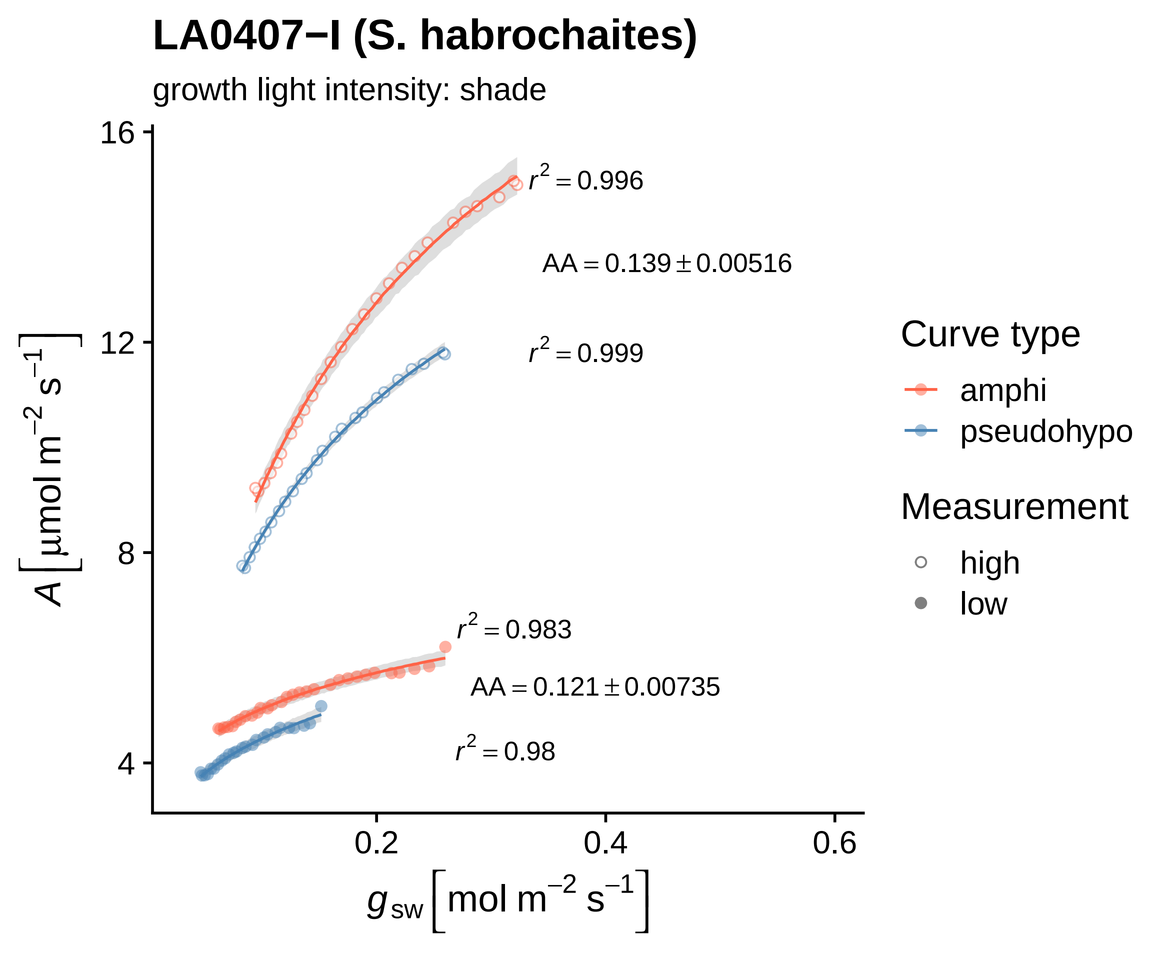

LA0407−I (S. habrochaites)
growth light intensity: shade
16
12
8
4
0.2
0.4
0.6
$A \left[\mu mol\ m^{-2}\ s^{-1}\right]$
$g_{sw} \left[mol\ m^{-2}\ s^{-1}\right]$
$r^2 = 0.996$
$AA = 0.139 \pm 0.00516$
$r^2 = 0.999$
$r^2 = 0.983$
$AA = 0.121 \pm 0.00735$
$r^2 = 0.98$
Curve type
amphi
pseudohypo
Measurement
high
low

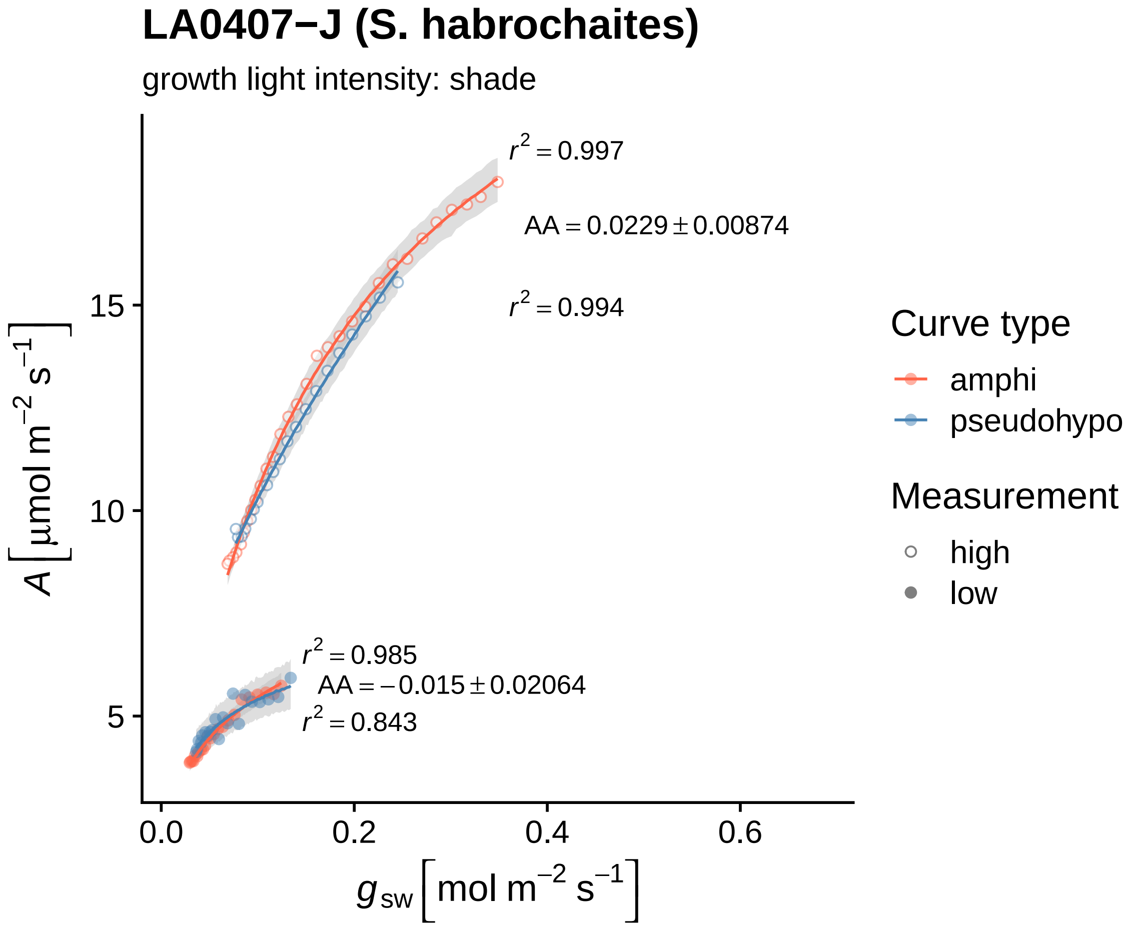

LA0407−J (S. habrochaites)
growth light intensity: shade
$r^2 = 0.997$
$AA = 0.0229 \pm 0.00874$
$r^2 = 0.994$
$r^2 = 0.985$
$AA = -0.015 \pm 0.02064$
$r^2 = 0.843$
15
10
5
0.0
0.2
0.4
0.6
$A\left[\mu mol\ m^{-2}\ s^{-1}\right]$
$g_{sw}\left[mol\ m^{-2}\ s^{-1}\right]$
Curve type
amphi
pseudohypo
Measurement
high
low

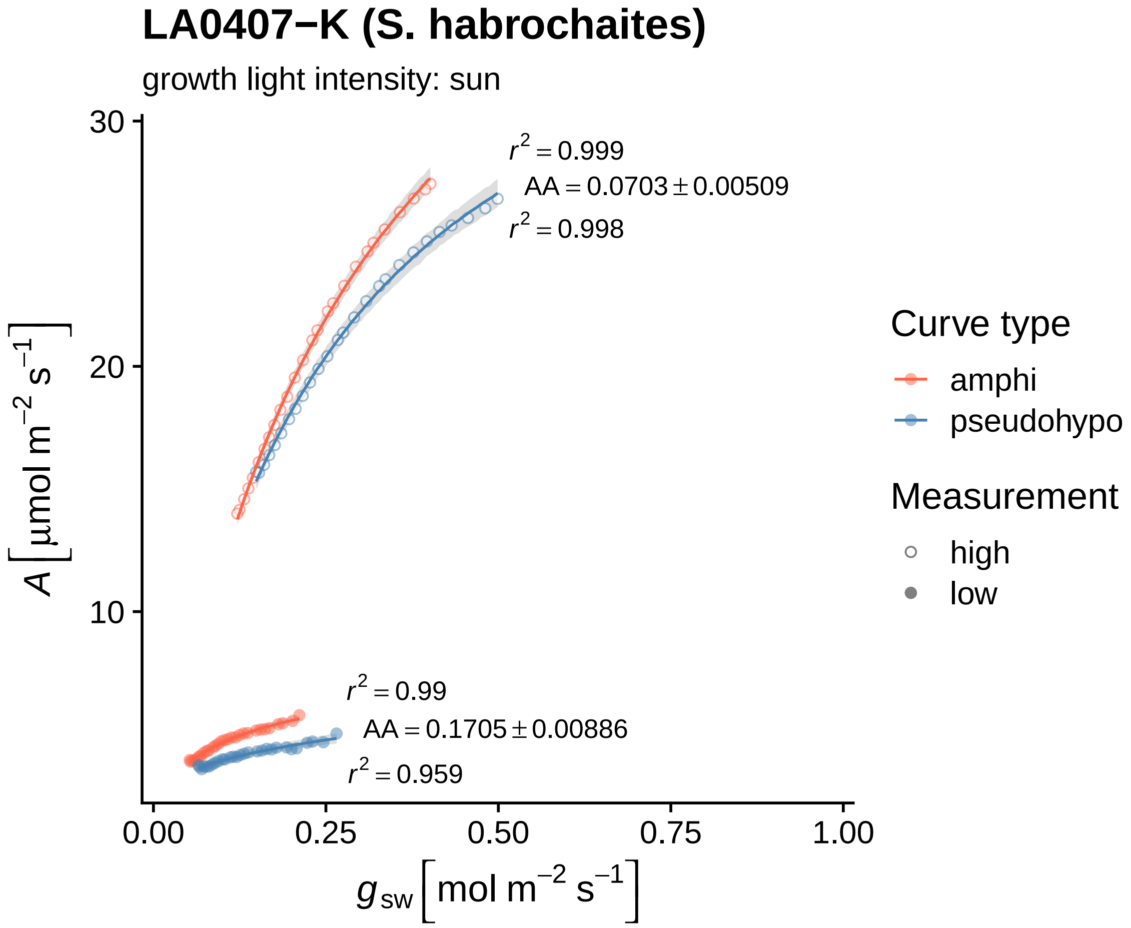
LA0407−K (S. habrochaites)
growth light intensity: sun
$r^2 = 0.999$
$AA = 0.0703 \pm 0.00509$
$r^2 = 0.998$
$r^2 = 0.99$
$AA = 0.1705 \pm 0.00886$
$r^2 = 0.959$
$A\left[\mu mol\ m^{-2}\ s^{-1}\right]$
$g_{sw}\left[mol\ m^{-2}\ s^{-1}\right]$
30
20
10
0.00
0.25
0.50
0.75
1.00
Curve type
amphi
pseudohypo
Measurement
high
low

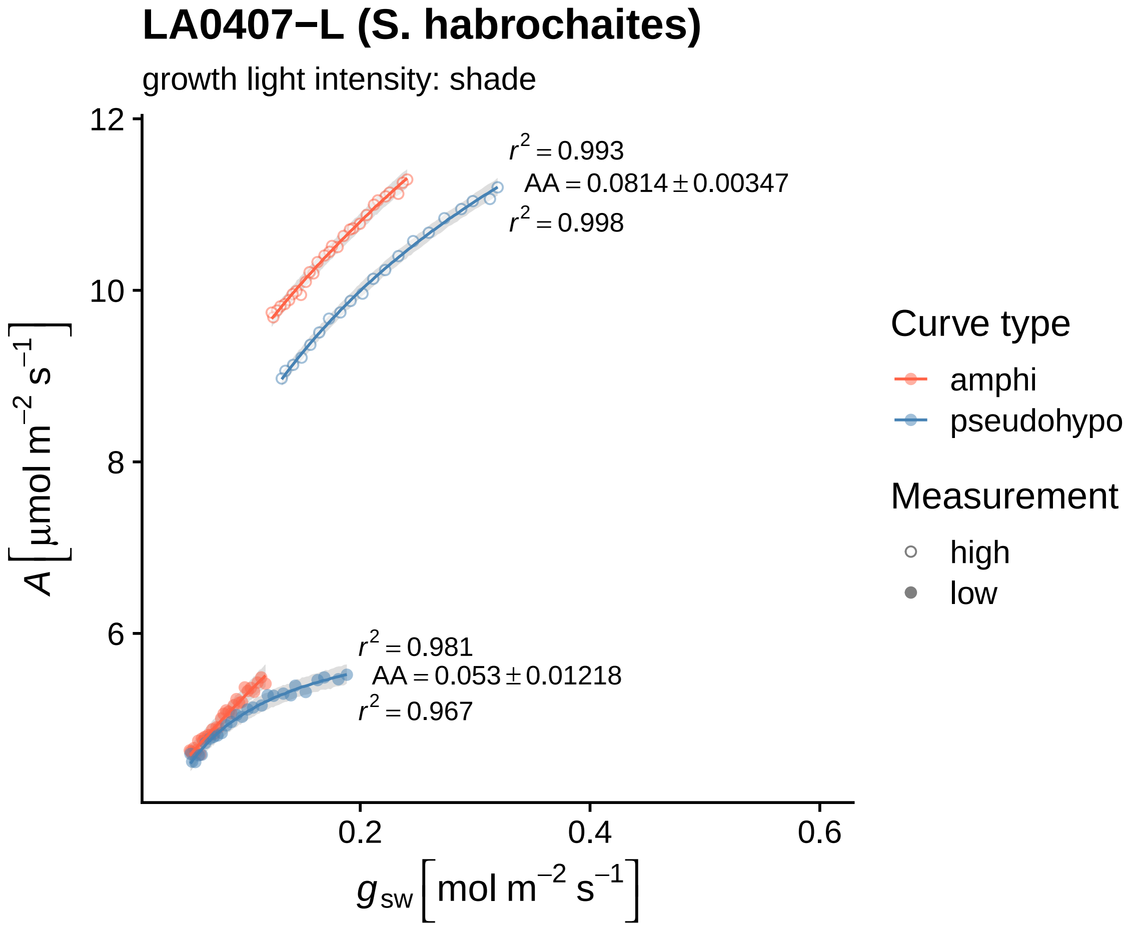

LA0407−L (S. habrochaites)
growth light intensity: shade
$r^2 = 0.993$
$AA = 0.0814 \pm 0.00347$
$r^2 = 0.998$
$r^2 = 0.981$
$AA = 0.053 \pm 0.01218$
$r^2 = 0.967$
12
10
8
6
0.2
0.4
0.6
$A\ [\mu mol\ m^{-2}\ s^{-1}]$
$g_{sw}\ [mol\ m^{-2}\ s^{-1}]$
Curve type
amphi
pseudohypo
Measurement
high
low

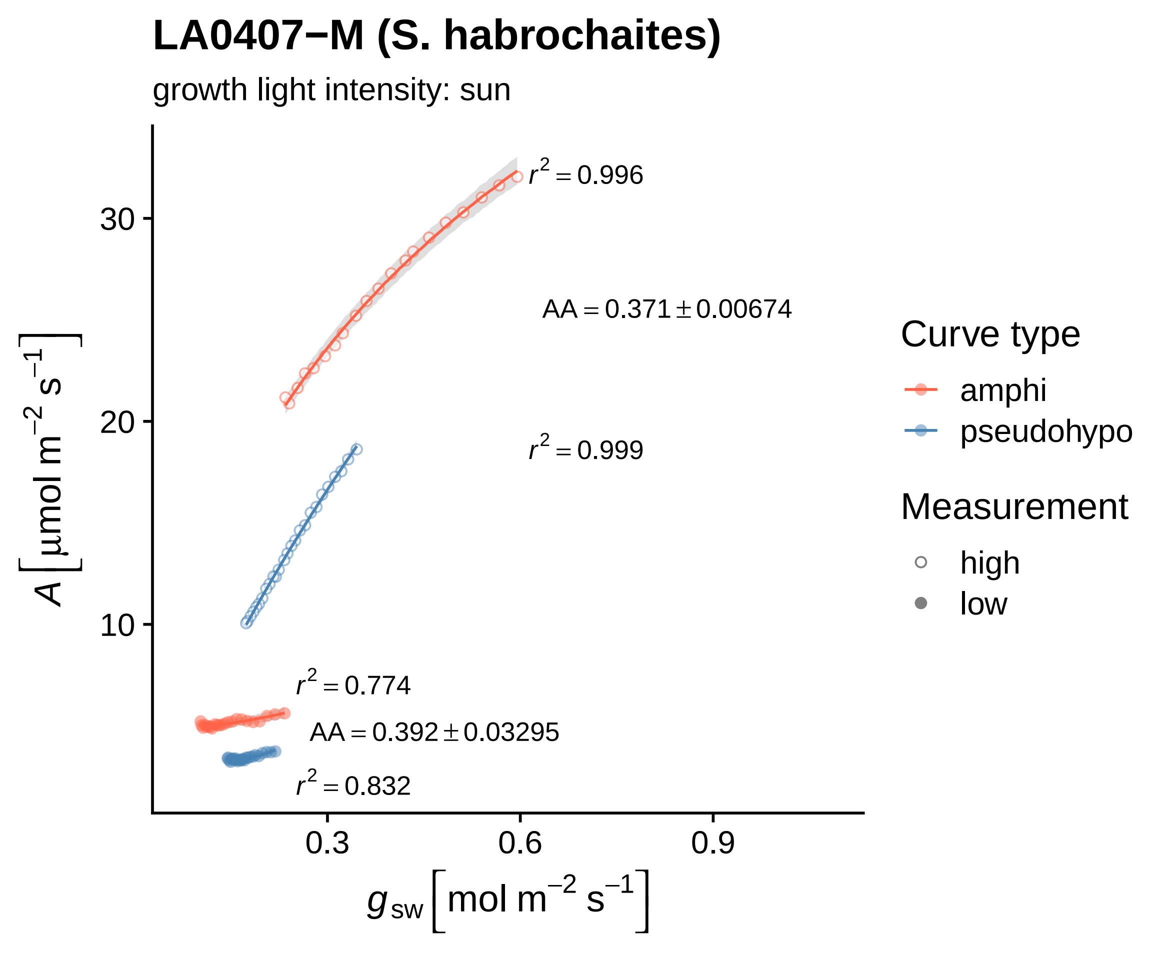

LA0407−M (S. habrochaites)
growth light intensity: sun
$r^2 = 0.996$
$AA = 0.371 \pm 0.00674$
$r^2 = 0.999$
$r^2 = 0.774$
$AA = 0.392 \pm 0.03295$
$r^2 = 0.832$
30
20
10
0.3
0.6
0.9
$A\left[\mu mol\ m^{-2}\ s^{-1}\right]$
$g_{sw}\left[mol\ m^{-2}\ s^{-1}\right]$
Curve type
amphi
pseudohypo
Measurement
high
low

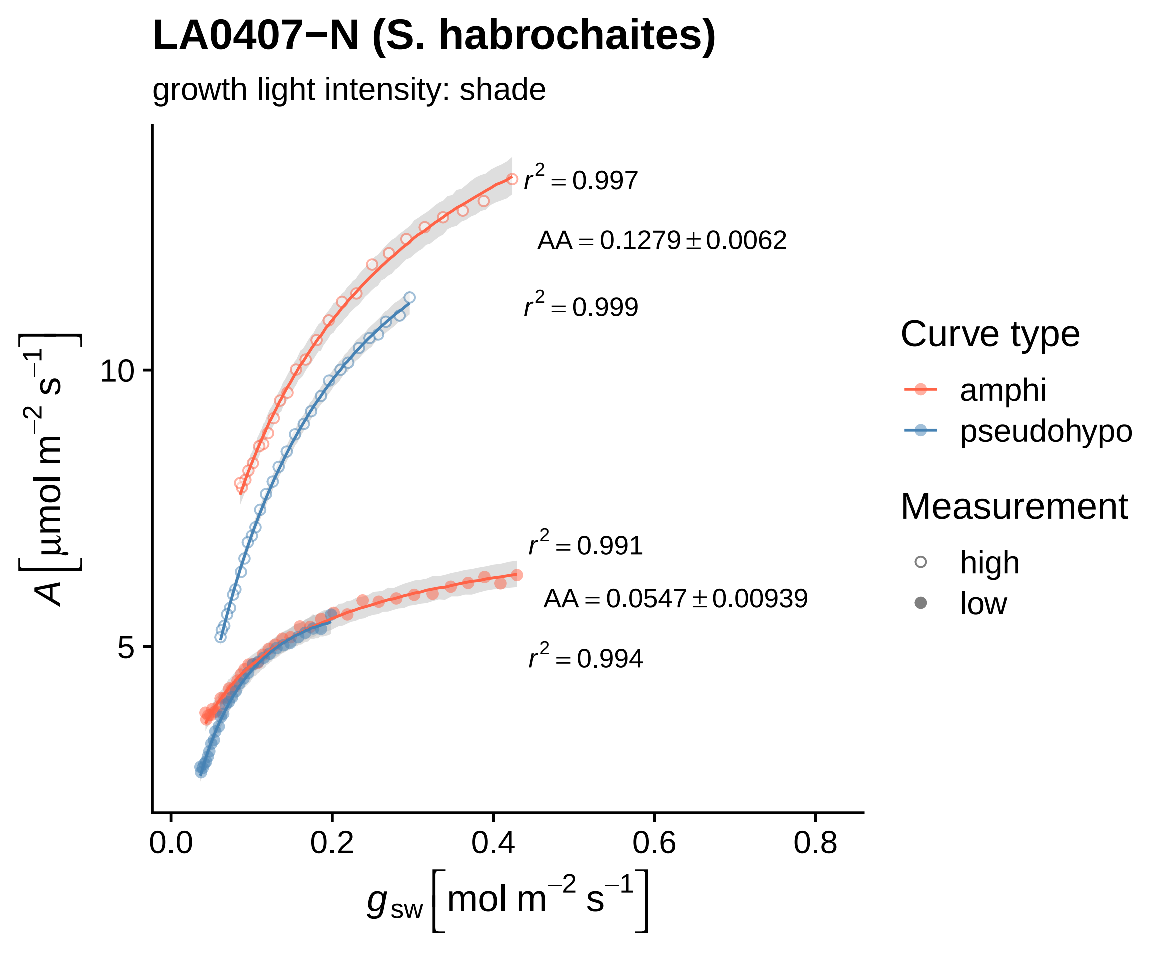

LA0407−N (S. habrochaites)
growth light intensity: shade
$r^2 = 0.997$
$AA = 0.1279 \pm 0.0062$
$r^2 = 0.999$
$r^2 = 0.991$
$AA = 0.0547 \pm 0.00939$
$r^2 = 0.994$
$A\ [\mu mol\ m^{-2}\ s^{-1}]$
10
5
0.0
0.2
0.4
0.6
0.8
$g_{sw}\ [mol\ m^{-2}\ s^{-1}]$
Curve type
amphi
pseudohypo
Measurement
high
low

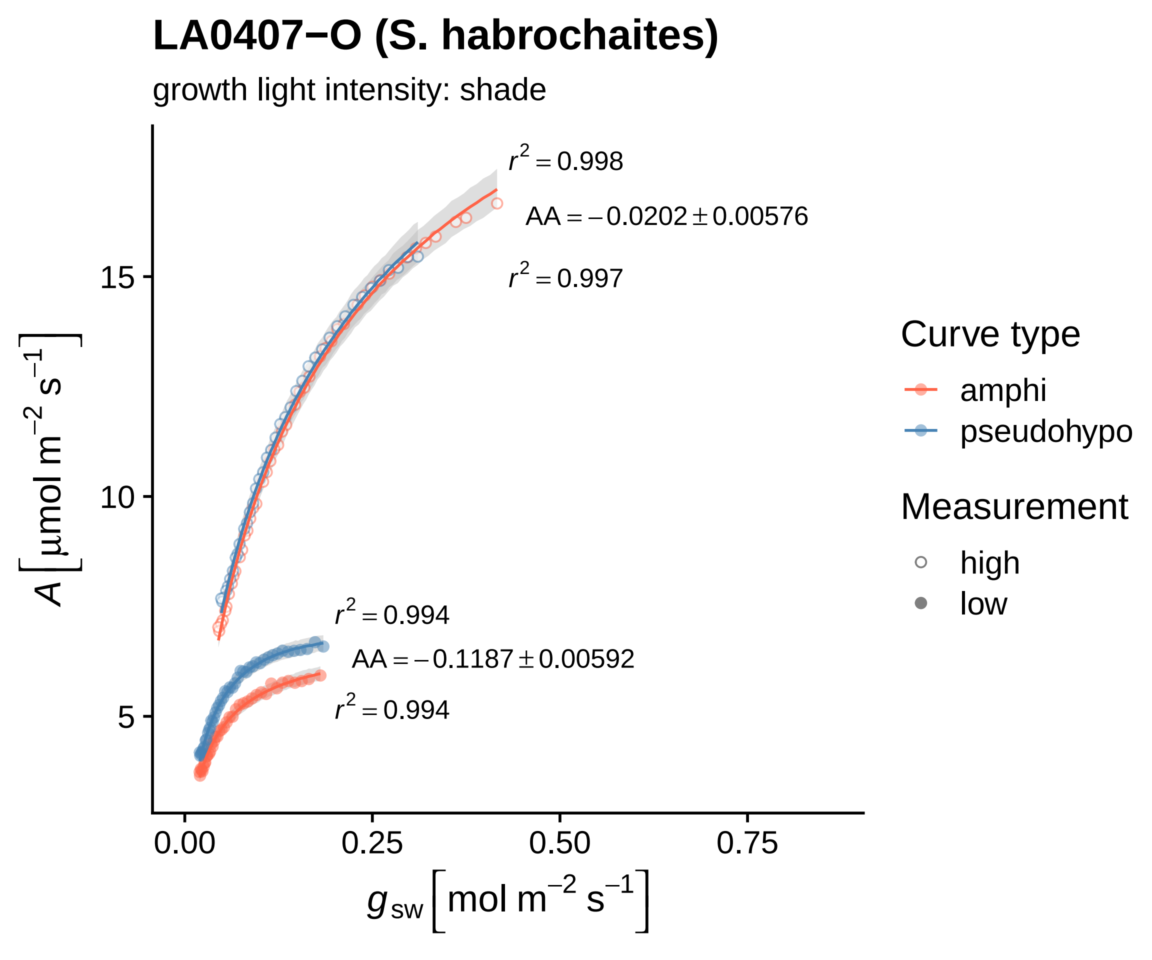

LA0407−O (S. habrochaites)
growth light intensity: shade
$r^2 = 0.998$
$AA = -0.0202 \pm 0.00576$
$r^2 = 0.997$
$r^2 = 0.994$
$AA = -0.1187 \pm 0.00592$
$r^2 = 0.994$
15
10
5
0.00
0.25
0.50
0.75
$A\left[\mu mol\ m^{-2}\ s^{-1}\right]$
$g_{sw}\left[mol\ m^{-2}\ s^{-1}\right]$
Curve type
amphi
pseudohypo
Measurement
high
low

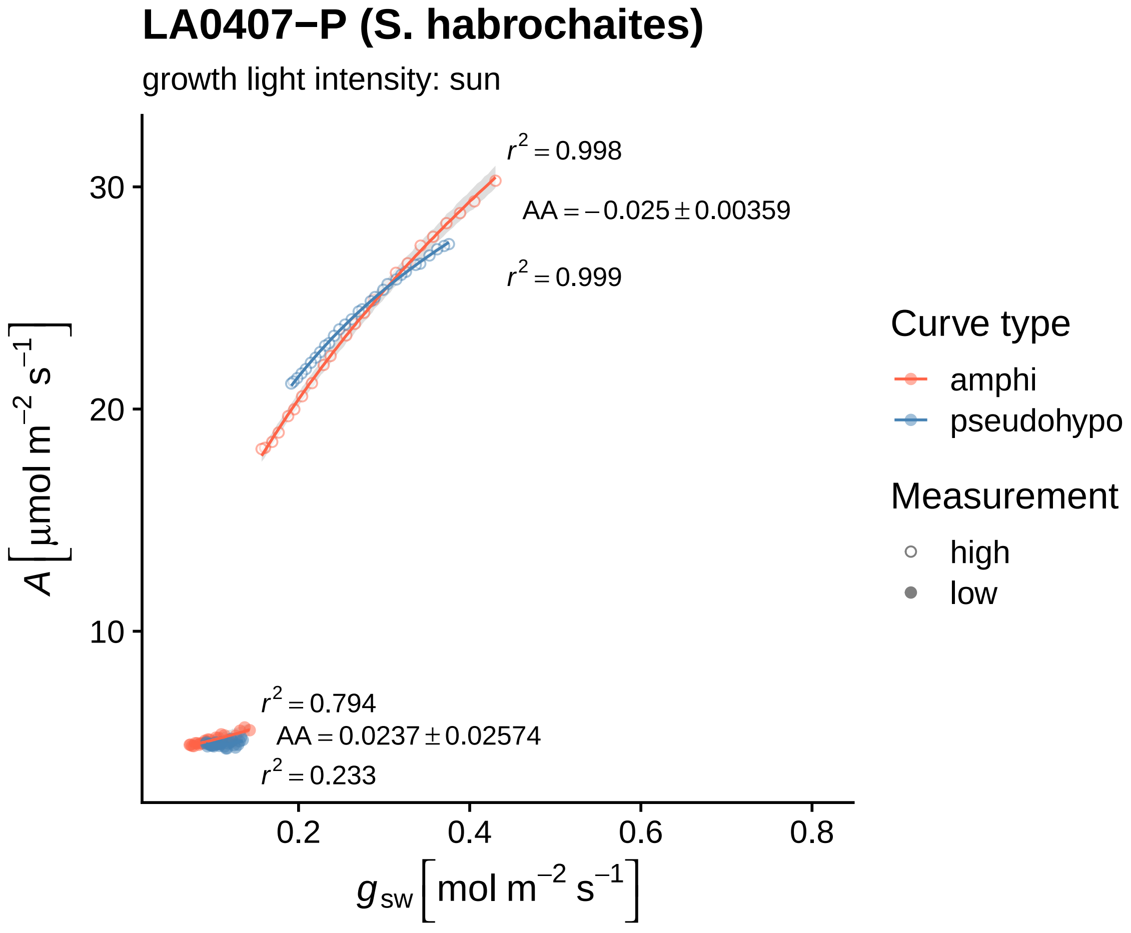

LA0407−P (S. habrochaites)
growth light intensity: sun
$r^2 = 0.998$
AA = −0.025 ± 0.00359
$r^2 = 0.999$
$r^2 = 0.794$
AA = 0.0237 ± 0.02574
$r^2 = 0.233$
30
20
10
0.2
0.4
0.6
0.8
$A\ [\mu mol\ m^{-2}\ s^{-1}]$
$g_{sw}\ [mol\ m^{-2}\ s^{-1}]$
Curve type
amphi
pseudohypo
Measurement
high
low

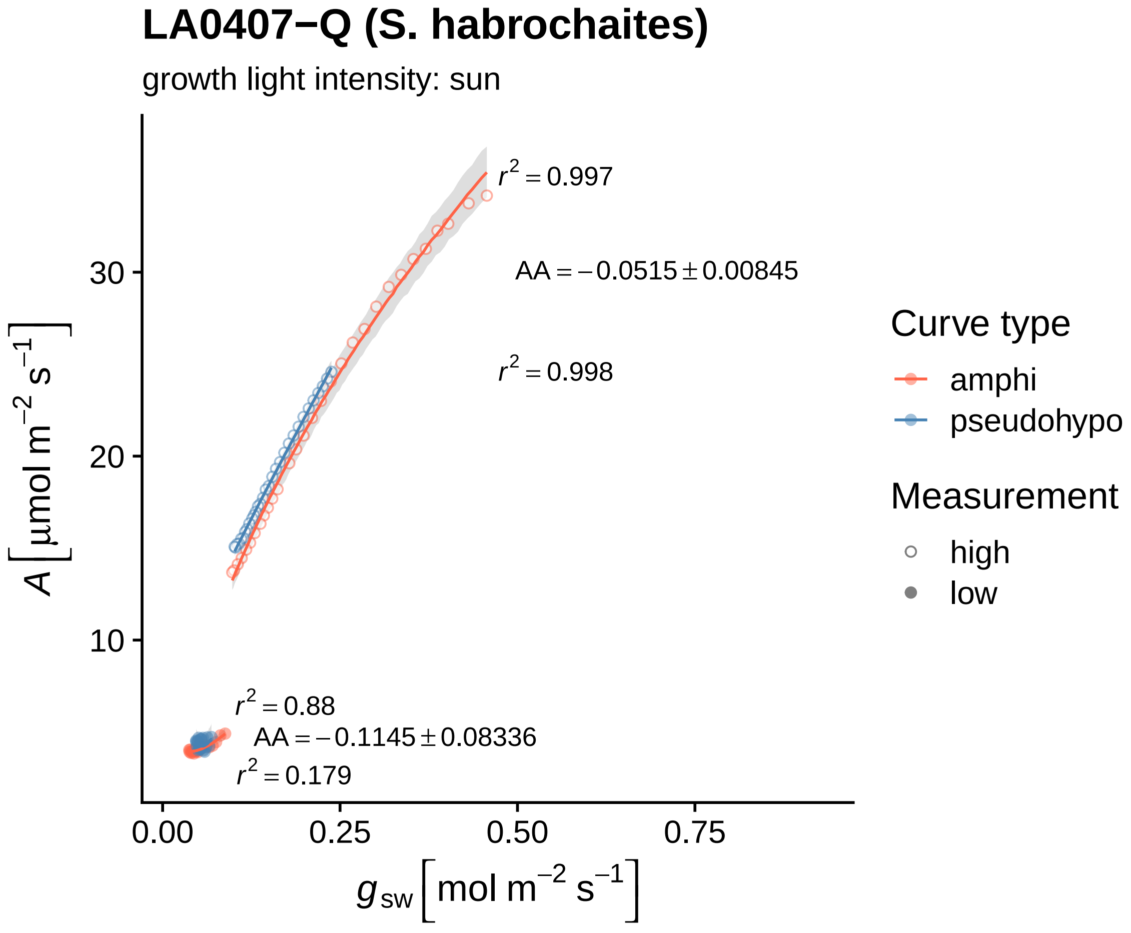

LA0407−Q (S. habrochaites)
growth light intensity: sun
$r^2 = 0.997$
$AA = -0.0515 \pm 0.00845$
$r^2 = 0.998$
$r^2 = 0.88$
$AA = -0.1145 \pm 0.08336$
$r^2 = 0.179$
30
20
10
0.00
0.25
0.50
0.75
$A\ [\mu mol\ m^{-2}\ s^{-1}]$
$g_{sw}\ [mol\ m^{-2}\ s^{-1}]$
Curve type
amphi
pseudohypo
Measurement
high
low

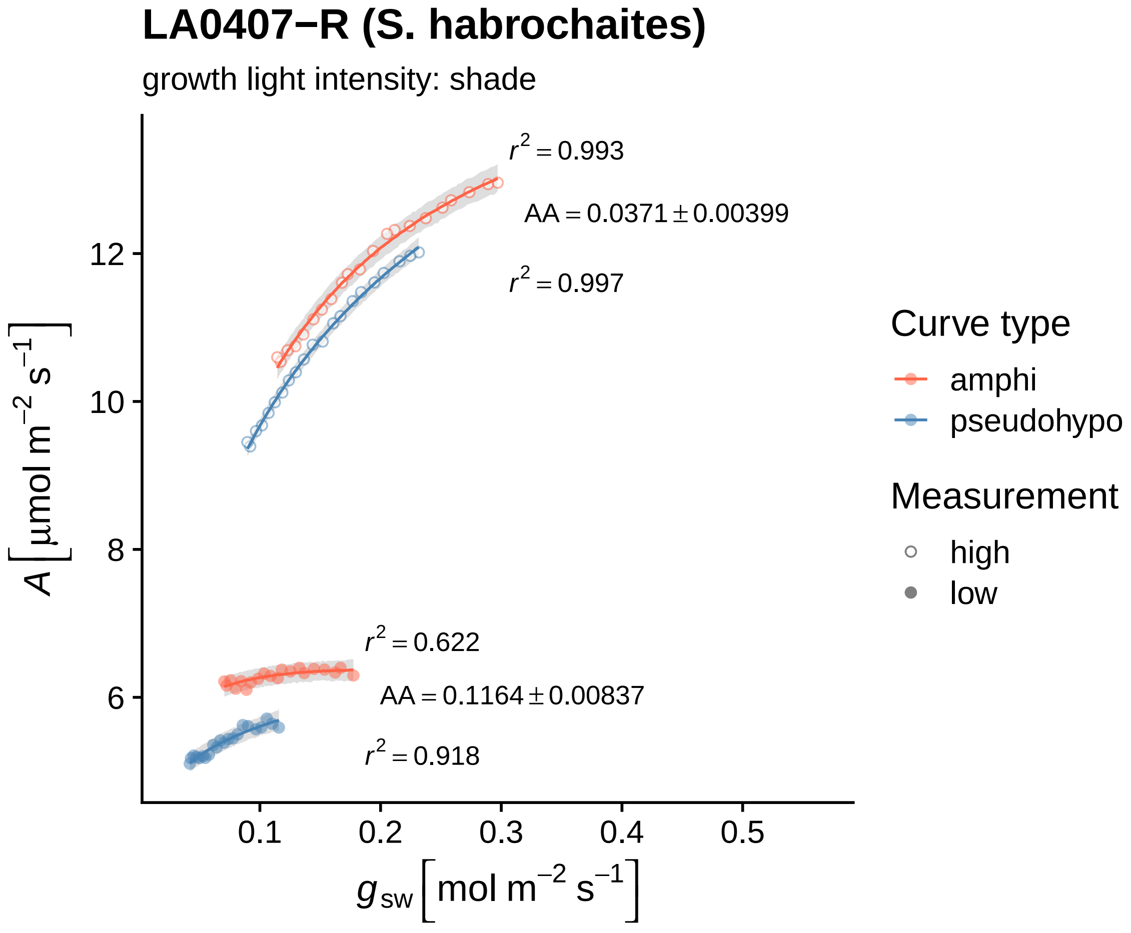

LA0407−R (S. habrochaites)
growth light intensity: shade
$r^2 = 0.993$
$AA = 0.0371 \pm 0.00399$
$r^2 = 0.997$
$r^2 = 0.622$
$AA = 0.1164 \pm 0.00837$
$r^2 = 0.918$
$A\left[\mu mol\ m^{-2}\ s^{-1}\right]$
$g_{sw}\left[mol\ m^{-2}\ s^{-1}\right]$
12
10
8
6
0.1
0.2
0.3
0.4
0.5
Curve type
amphi
pseudohypo
Measurement
high
low

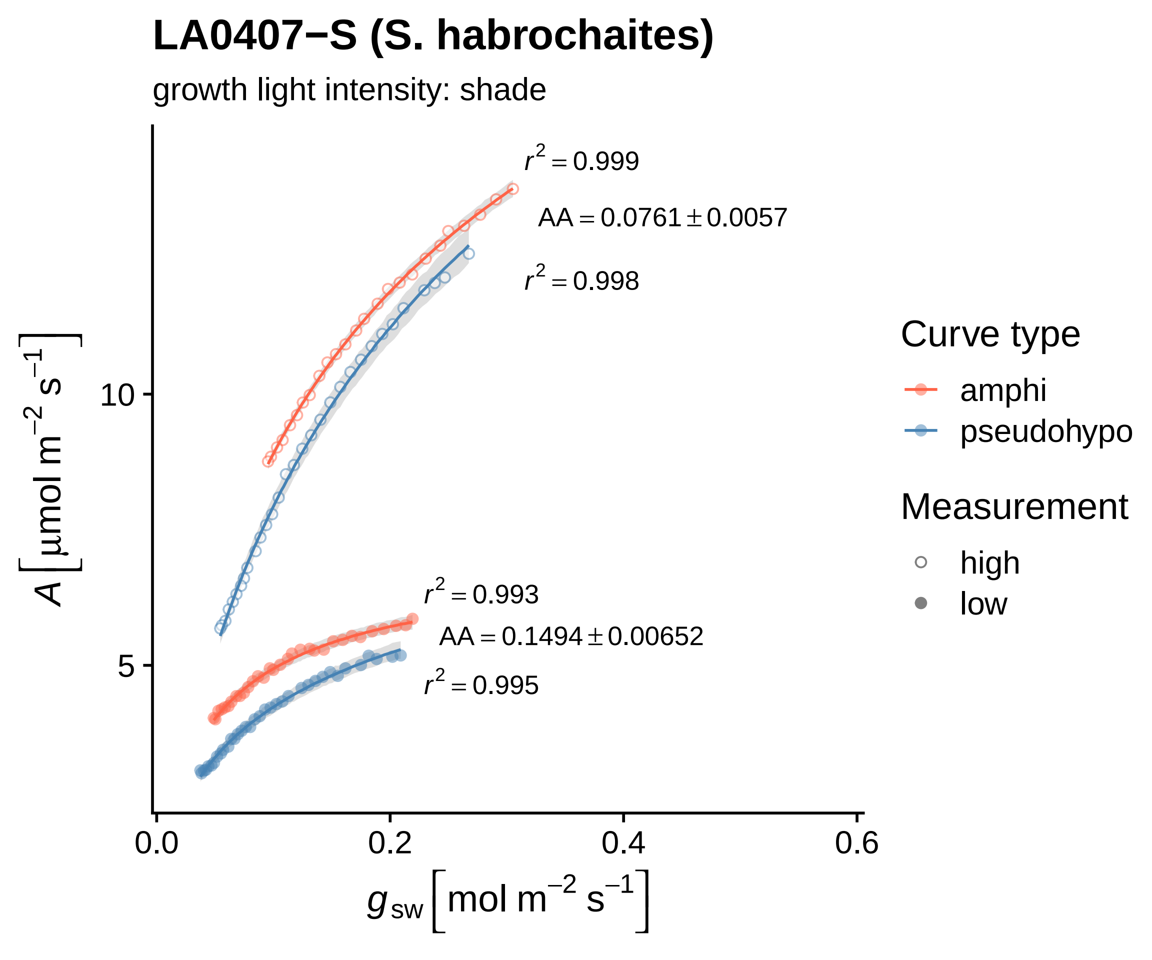

LA0407−S (S. habrochaites)
growth light intensity: shade
$r^2 = 0.999$
AA = 0.0761 ± 0.0057
$r^2 = 0.998$
$r^2 = 0.993$
AA = 0.1494 ± 0.00652
$r^2 = 0.995$
$A\ [\mu mol\ m^{-2}\ s^{-1}]$
10
5
0.0
0.2
0.4
0.6
$g_{sw}\ [mol\ m^{-2}\ s^{-1}]$
Curve type
amphi
pseudohypo
Measurement
high
low

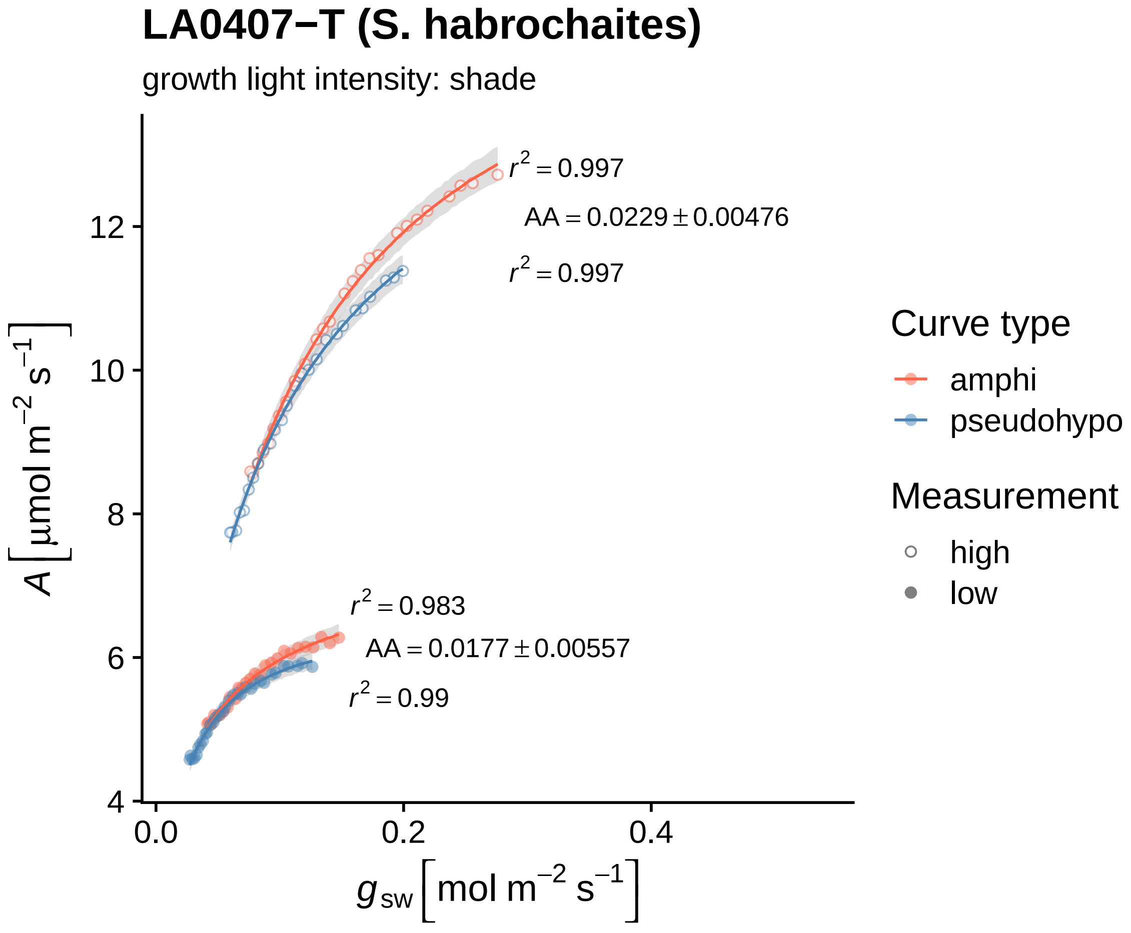

LA0407−T (S. habrochaites)
growth light intensity: shade
$r^2 = 0.997$
$AA = 0.0229 \pm 0.00476$
$r^2 = 0.997$
$r^2 = 0.983$
$AA = 0.0177 \pm 0.00557$
$r^2 = 0.99$
4
6
8
10
12
0.0
0.2
0.4
$A\left[\mu mol\ m^{-2}\ s^{-1}\right]$
$g_{sw}\left[mol\ m^{-2}\ s^{-1}\right]$
Curve type
amphi
pseudohypo
Measurement
high
low

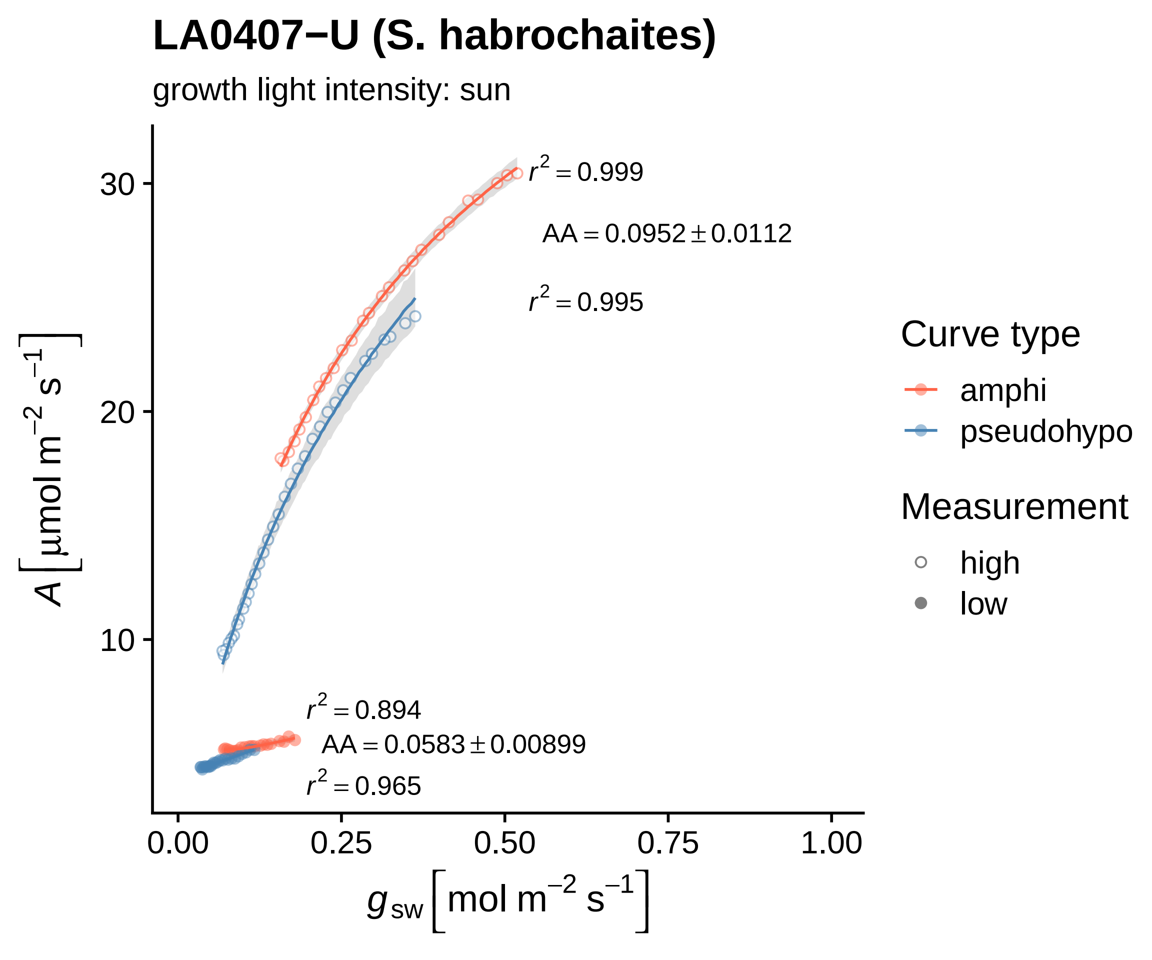
LA0407−U (S. habrochaites)
growth light intensity: sun
A [μmol m−2 s−1]
gsw [mol m−2 s−1]
r2 = 0.999
AA = 0.0952 ± 0.0112
r2 = 0.995
r2 = 0.894
AA = 0.0583 ± 0.00899
r2 = 0.965
30
20
10
0.00
0.25
0.50
0.75
1.00
Curve type
amphi
pseudohypo
Measurement
high
low

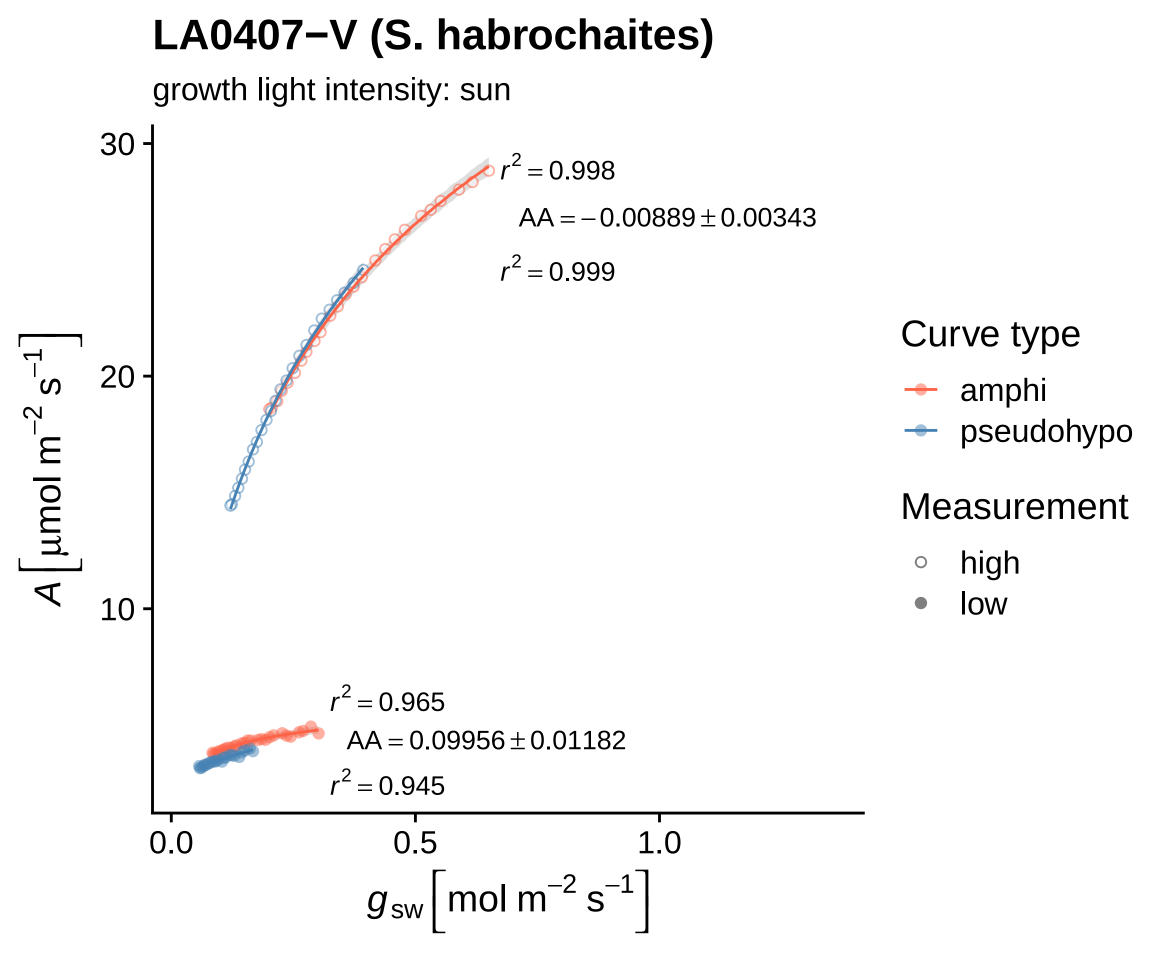

LA0407−V (S. habrochaites)
growth light intensity: sun
$r^2 = 0.998$
$AA = -0.00889 \pm 0.00343$
$r^2 = 0.999$
$r^2 = 0.965$
$AA = 0.09956 \pm 0.01182$
$r^2 = 0.945$
30
20
10
0.0
0.5
1.0
$A\ [\mu mol\ m^{-2}\ s^{-1}]$
$g_{sw}\ [mol\ m^{-2}\ s^{-1}]$
Curve type
amphi
pseudohypo
Measurement
high
low

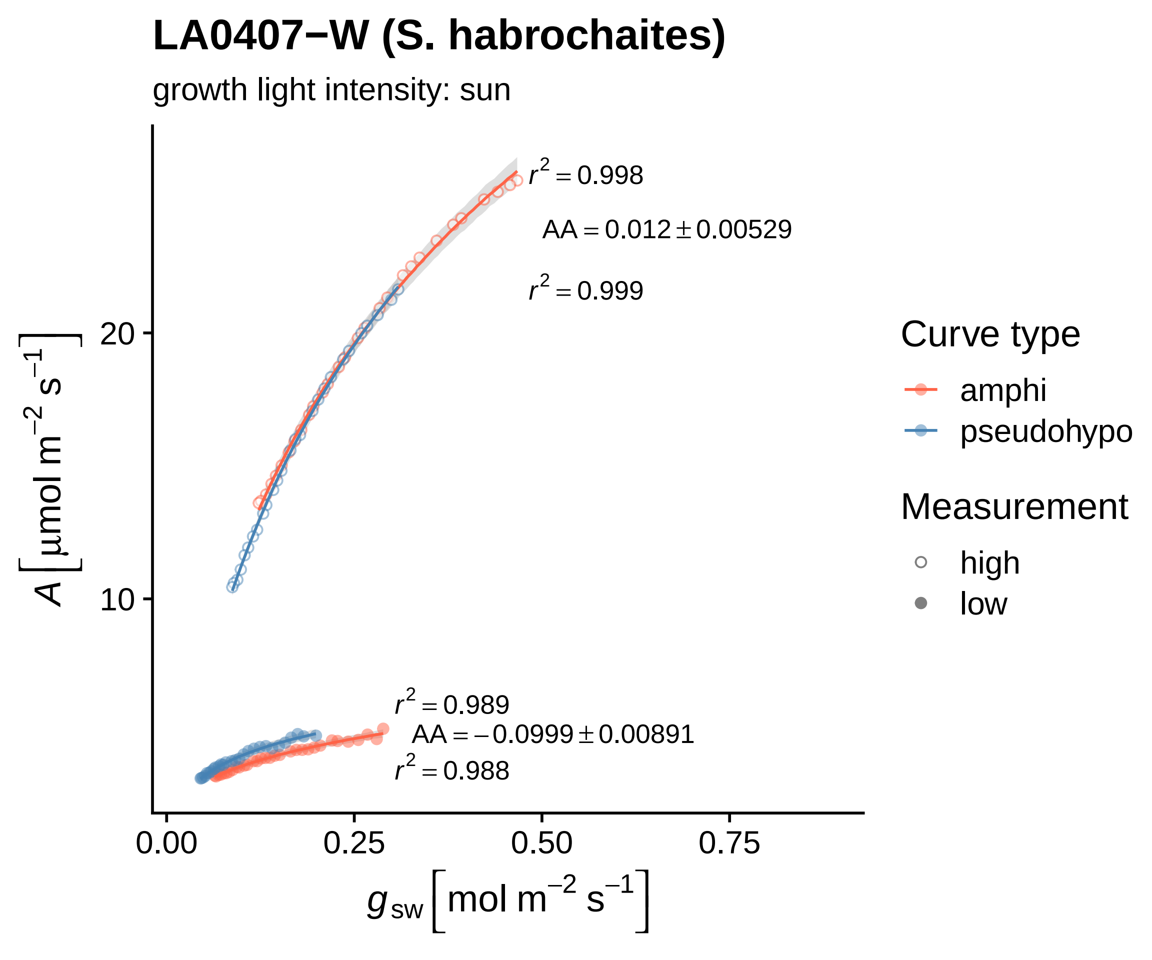

LA0407−W (S. habrochaites)
growth light intensity: sun
$r^2 = 0.998$
$AA = 0.012 \pm 0.00529$
$r^2 = 0.999$
$r^2 = 0.989$
$AA = -0.0999 \pm 0.00891$
$r^2 = 0.988$
20
10
0.00
0.25
0.50
0.75
$A\ [\mu mol\ m^{-2}\ s^{-1}]$
$g_{sw}\ [mol\ m^{-2}\ s^{-1}]$
Curve type
amphi
pseudohypo
Measurement
high
low

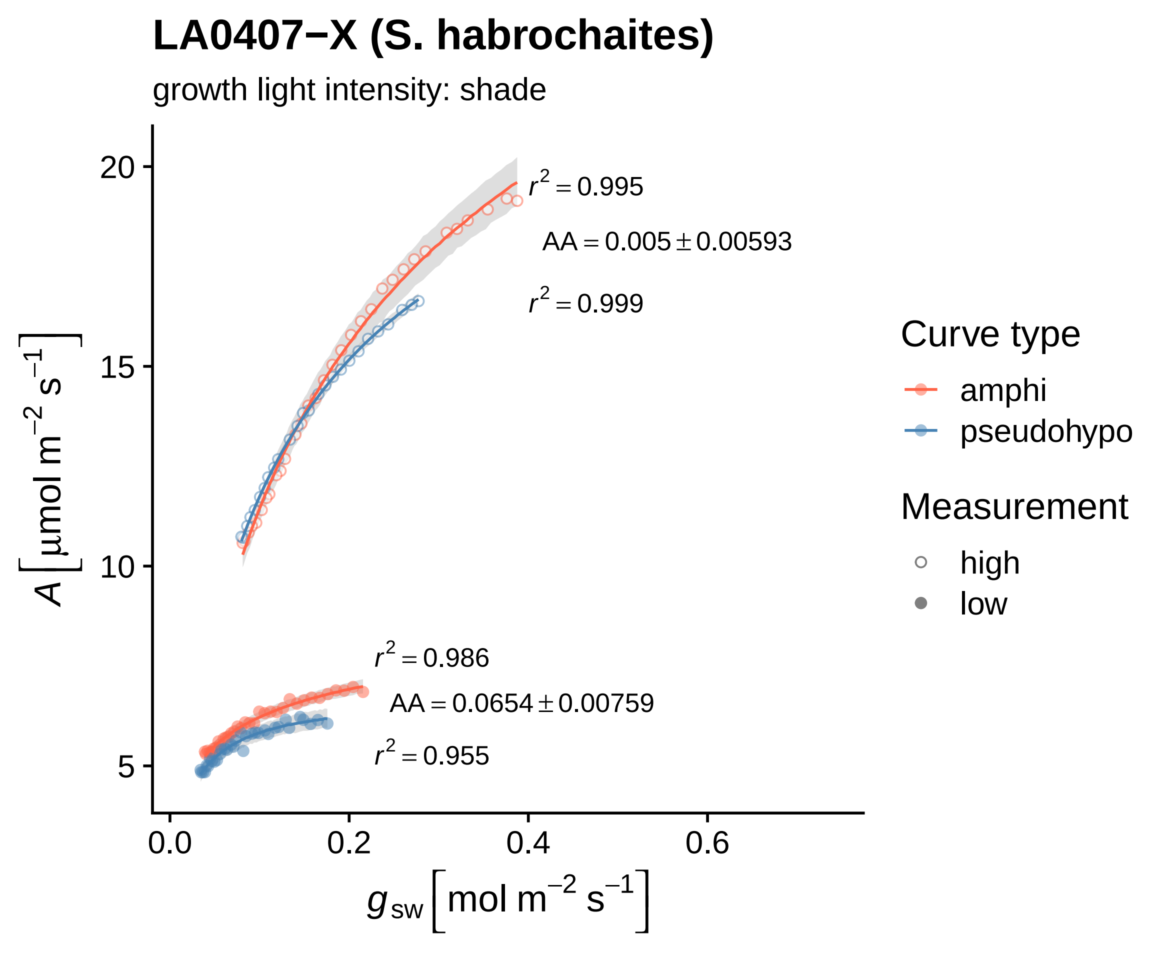
LA0407−X (S. habrochaites)
growth light intensity: shade
$r^2 = 0.995$
$AA = 0.005 \pm 0.00593$
$r^2 = 0.999$
$r^2 = 0.986$
$AA = 0.0654 \pm 0.00759$
$r^2 = 0.955$
20
15
10
5
0.0
0.2
0.4
0.6
$A\ [\mu mol\ m^{-2}\ s^{-1}]$
$g_{sw}\ [mol\ m^{-2}\ s^{-1}]$
Curve type
amphi
pseudohypo
Measurement
high
low

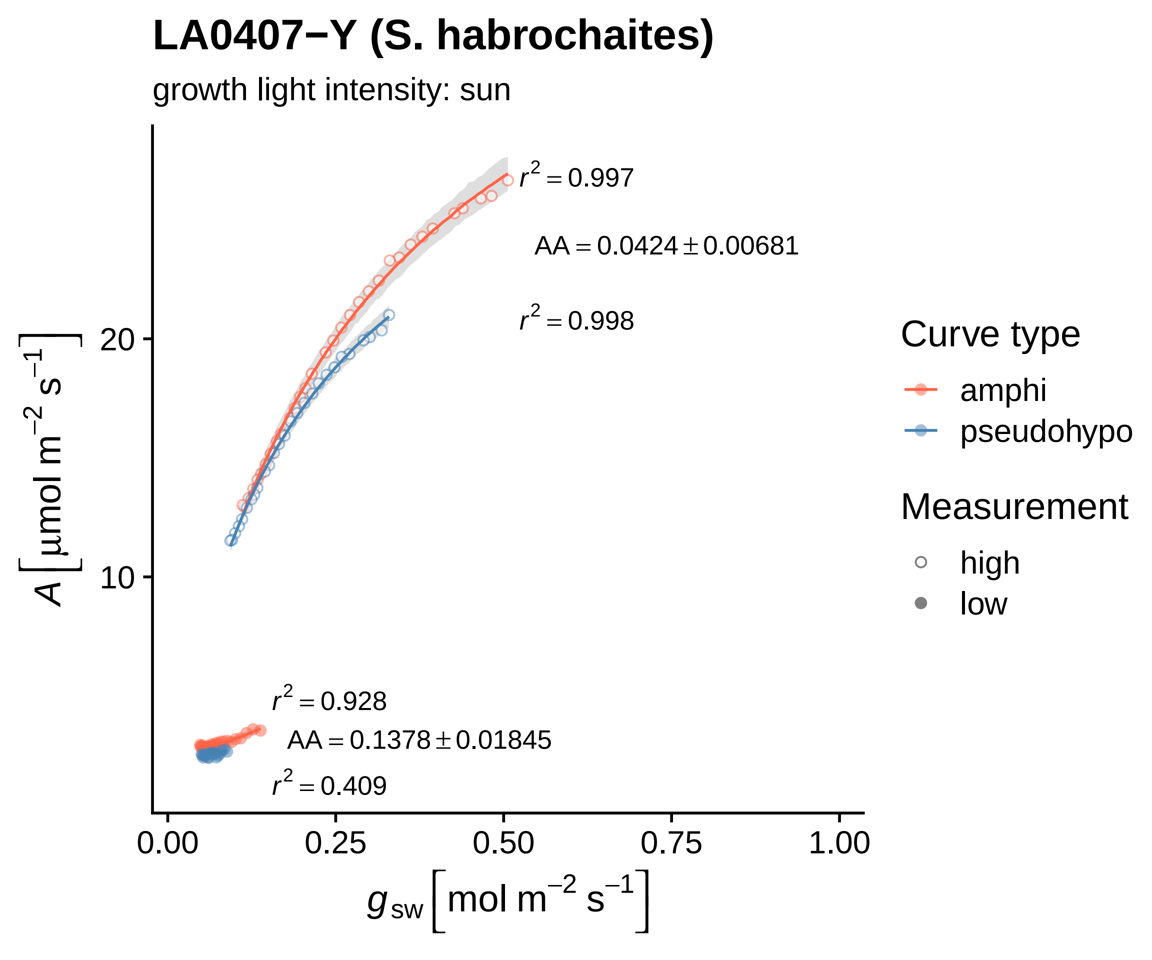

LA0407−Y (S. habrochaites)
growth light intensity: sun
$r^2 = 0.997$
$AA = 0.0424 \pm 0.00681$
$r^2 = 0.998$
$r^2 = 0.928$
$AA = 0.1378 \pm 0.01845$
$r^2 = 0.409$
20
10
0.00
0.25
0.50
0.75
1.00
$A\,[\mu mol\ m^{-2}\ s^{-1}]$
$g_{sw}\,[mol\ m^{-2}\ s^{-1}]$
Curve type
amphi
pseudohypo
Measurement
high
low

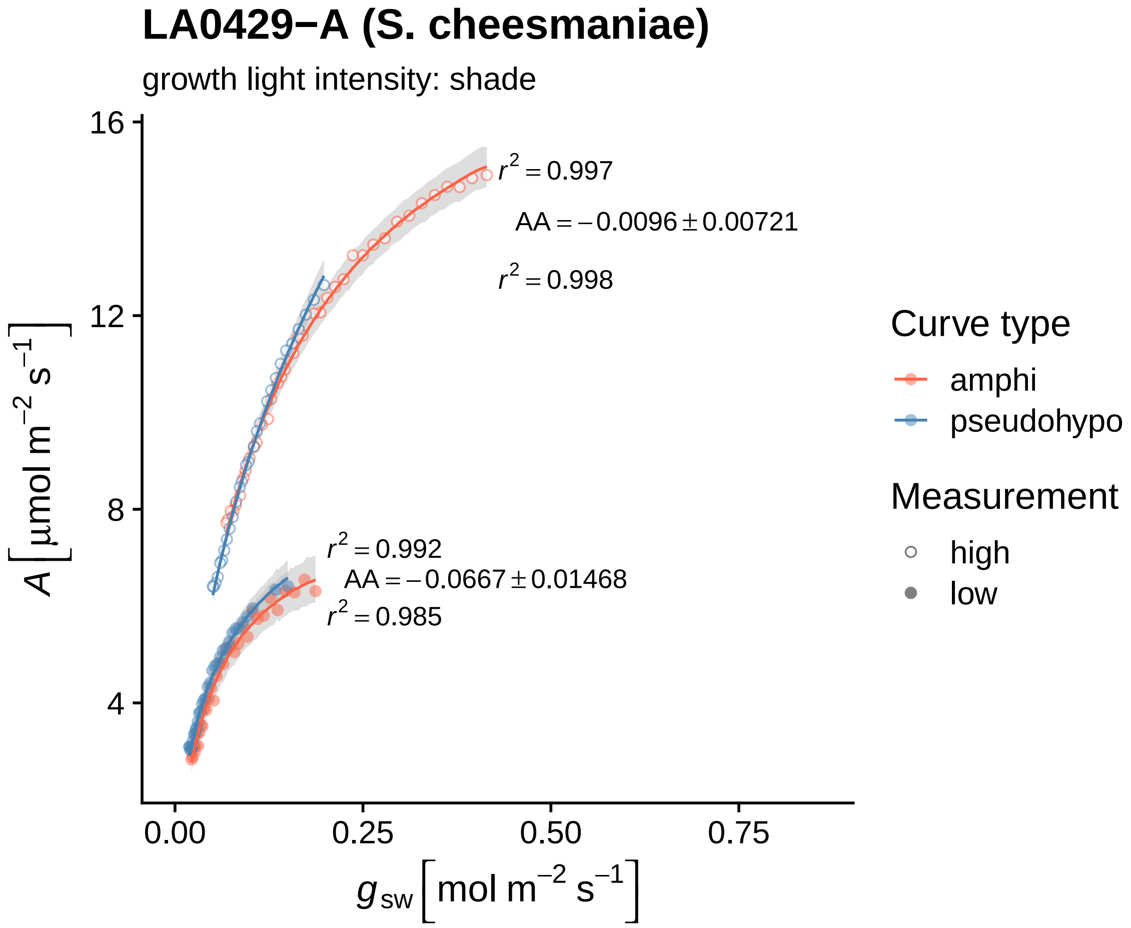

LA0429−A (S. cheesmaniae)
growth light intensity: shade
r² = 0.997
AA = −0.0096 ± 0.00721
r² = 0.998
r² = 0.992
AA = −0.0667 ± 0.01468
r² = 0.985
16
12
8
4
0.00
0.25
0.50
0.75
A [μmol m⁻² s⁻¹]
g_sw [mol m⁻² s⁻¹]
Curve type
amphi
pseudohypo
Measurement
high
low

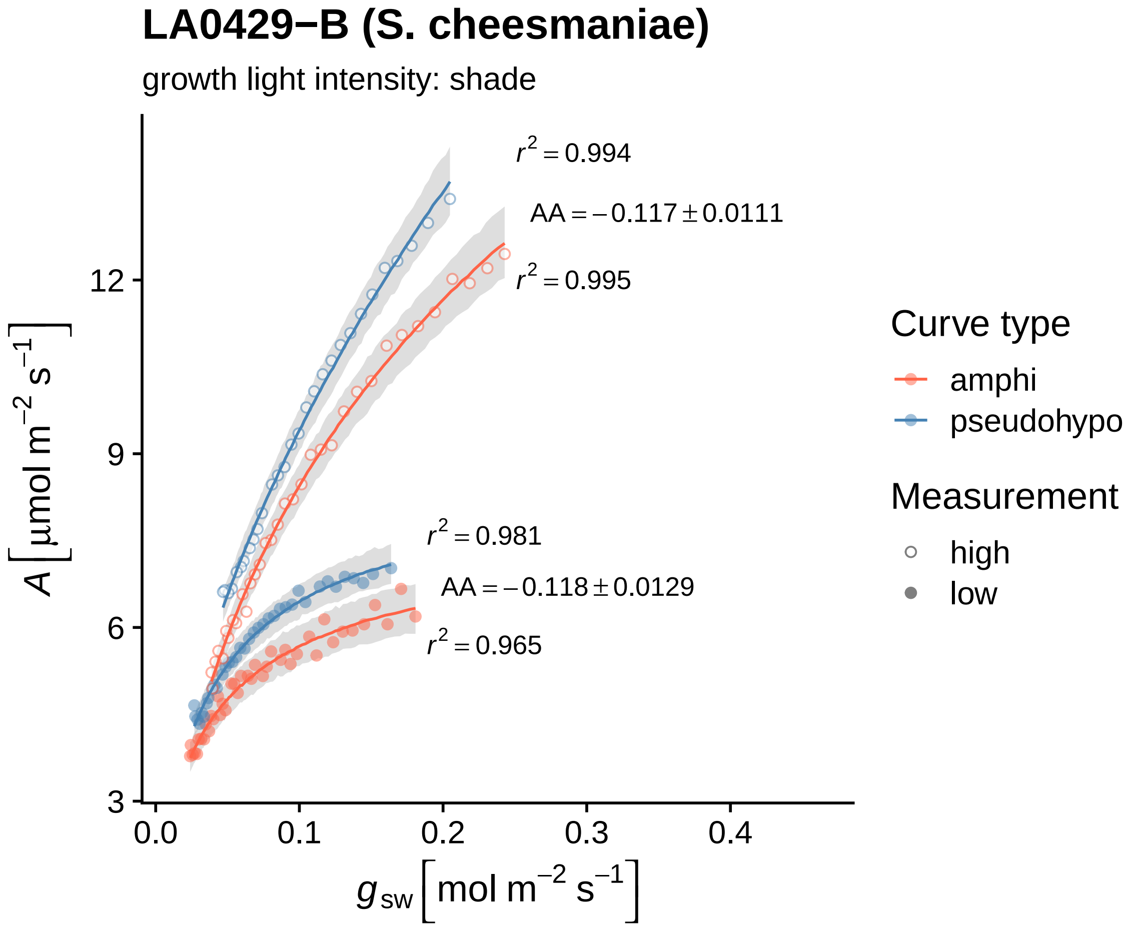

LA0429−B (S. cheesmaniae)
growth light intensity: shade
r² = 0.994
AA = −0.117 ± 0.0111
r² = 0.995
r² = 0.981
AA = −0.118 ± 0.0129
r² = 0.965
12
9
6
3
A [µmol m⁻² s⁻¹]
0.0
0.1
0.2
0.3
0.4
g_sw [mol m⁻² s⁻¹]
Curve type
amphi
pseudohypo
Measurement
high
low

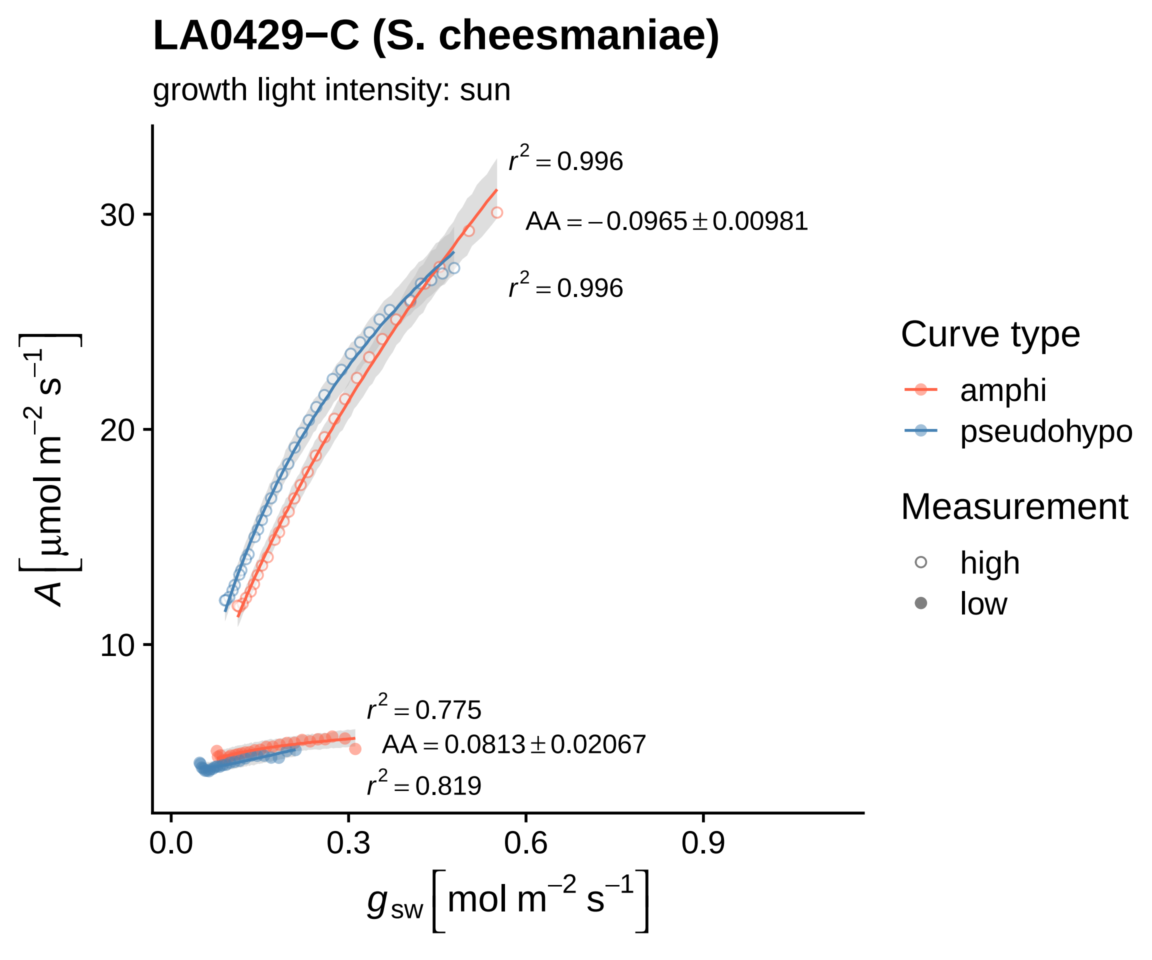
LA0429−C (S. cheesmaniae)
growth light intensity: sun
$r^2 = 0.996$
$AA = -0.0965 \pm 0.00981$
$r^2 = 0.996$
$r^2 = 0.775$
$AA = 0.0813 \pm 0.02067$
$r^2 = 0.819$
$A\left[\mu mol\ m^{-2}\ s^{-1}\right]$
$g_{sw}\left[mol\ m^{-2}\ s^{-1}\right]$
30
20
10
0.0
0.3
0.6
0.9
Curve type
amphi
pseudohypo
Measurement
high
low

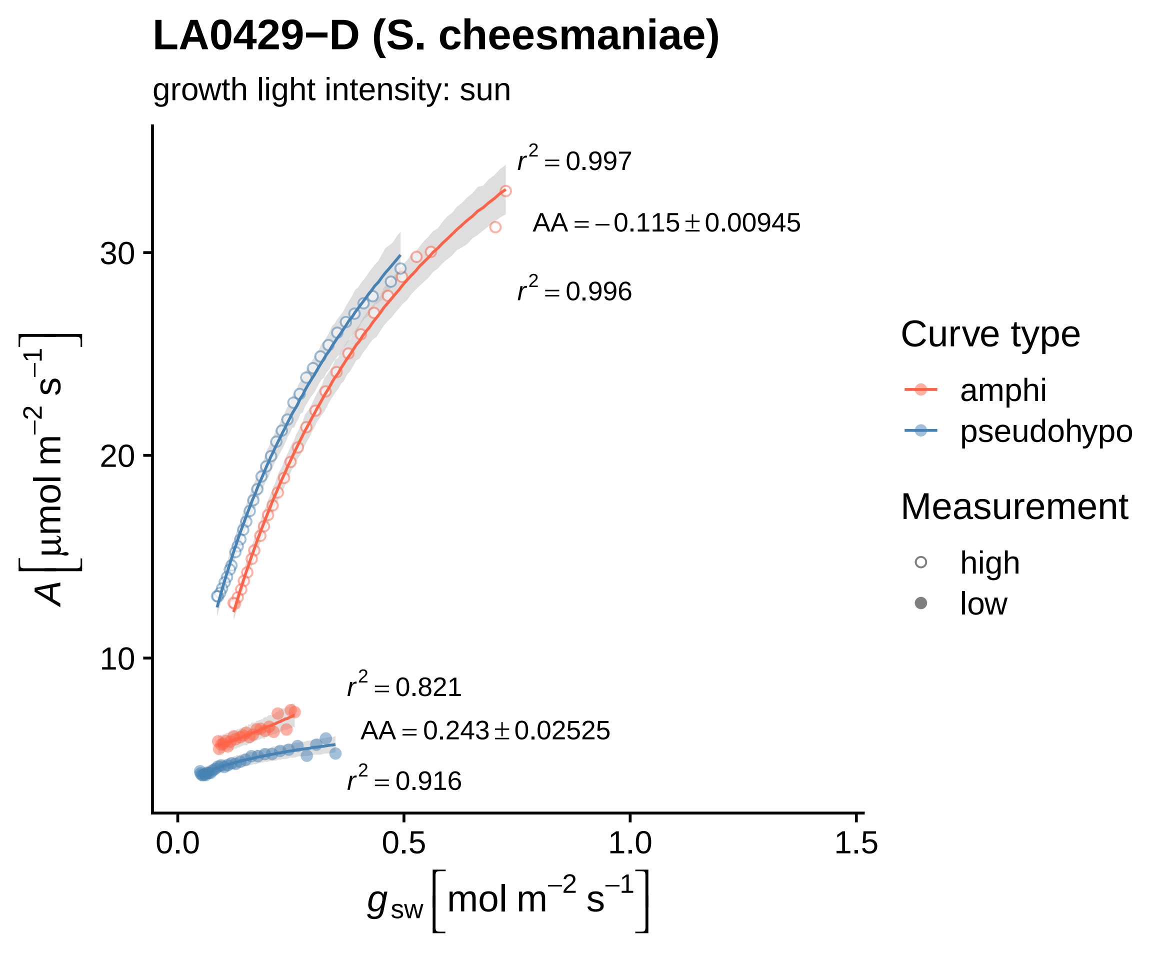

LA0429−D (S. cheesmaniae)
growth light intensity: sun
$r^2 = 0.997$
$AA = -0.115 \pm 0.00945$
$r^2 = 0.996$
$r^2 = 0.821$
$AA = 0.243 \pm 0.02525$
$r^2 = 0.916$
30
20
10
0.0
0.5
1.0
1.5
$A\left[\mu mol\ m^{-2}\ s^{-1}\right]$
$g_{sw}\left[mol\ m^{-2}\ s^{-1}\right]$
Curve type
amphi
pseudohypo
Measurement
high
low

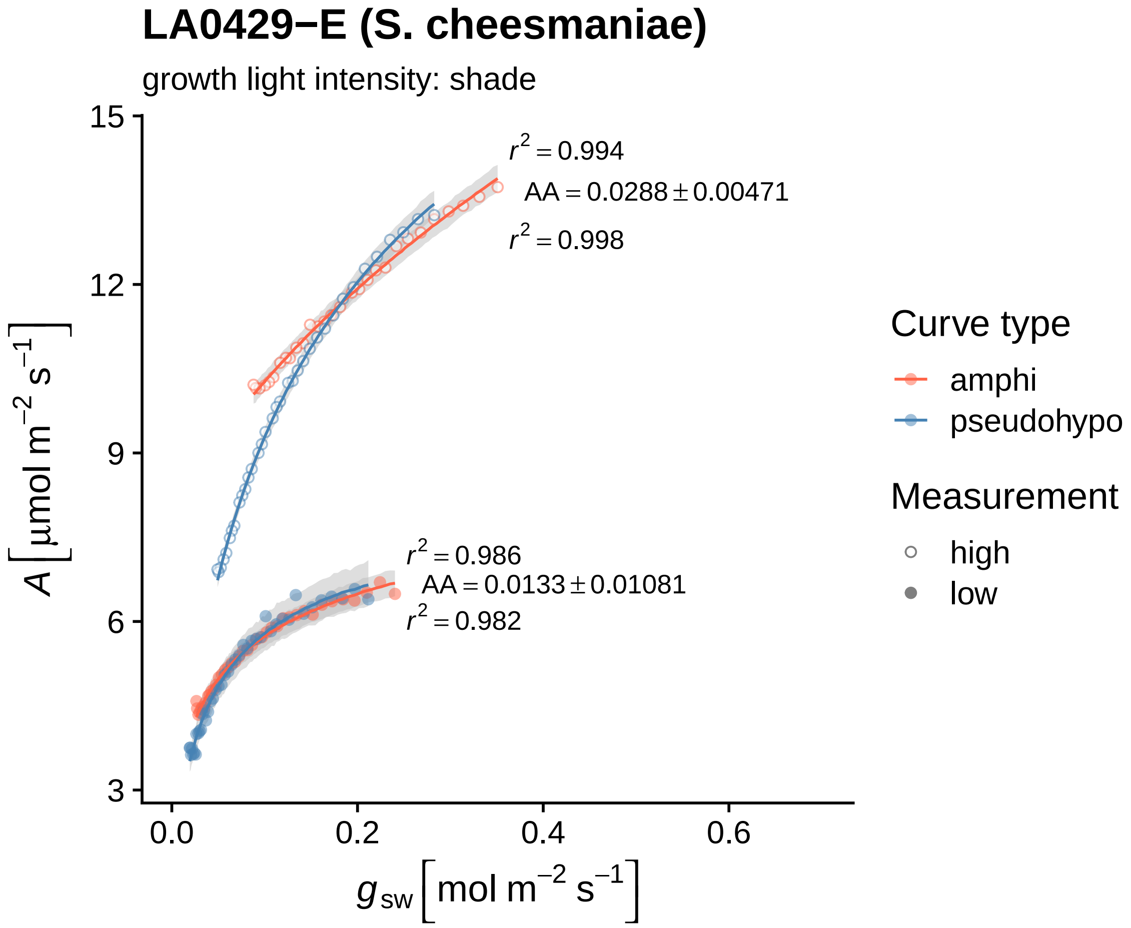
LA0429−E (S. cheesmaniae)
growth light intensity: shade
$r^2 = 0.994$
$AA = 0.0288 \pm 0.00471$
$r^2 = 0.998$
$r^2 = 0.986$
$AA = 0.0133 \pm 0.01081$
$r^2 = 0.982$
15
12
9
6
3
0.0
0.2
0.4
0.6
$A \left[\mu mol\ m^{-2}\ s^{-1}\right]$
$g_{sw} \left[mol\ m^{-2}\ s^{-1}\right]$
Curve type
amphi
pseudohypo
Measurement
high
low

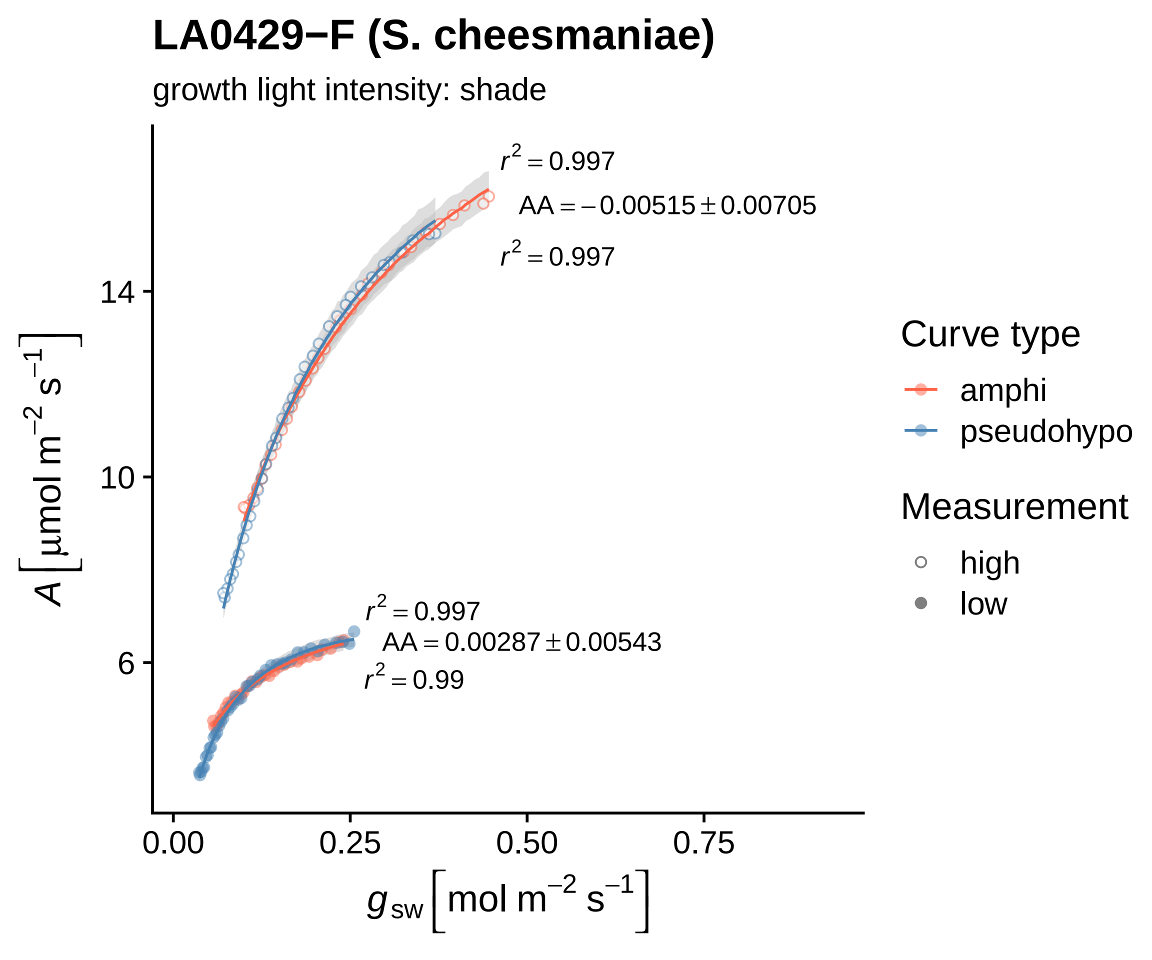

LA0429−F (S. cheesmaniae)
growth light intensity: shade
$r^2 = 0.997$
$AA = -0.00515 \pm 0.00705$
$r^2 = 0.997$
$r^2 = 0.997$
$AA = 0.00287 \pm 0.00543$
$r^2 = 0.99$
14
10
6
0.00
0.25
0.50
0.75
$A\ [\mu mol\ m^{-2}\ s^{-1}]$
$g_{sw}\ [mol\ m^{-2}\ s^{-1}]$
Curve type
amphi
pseudohypo
Measurement
high
low

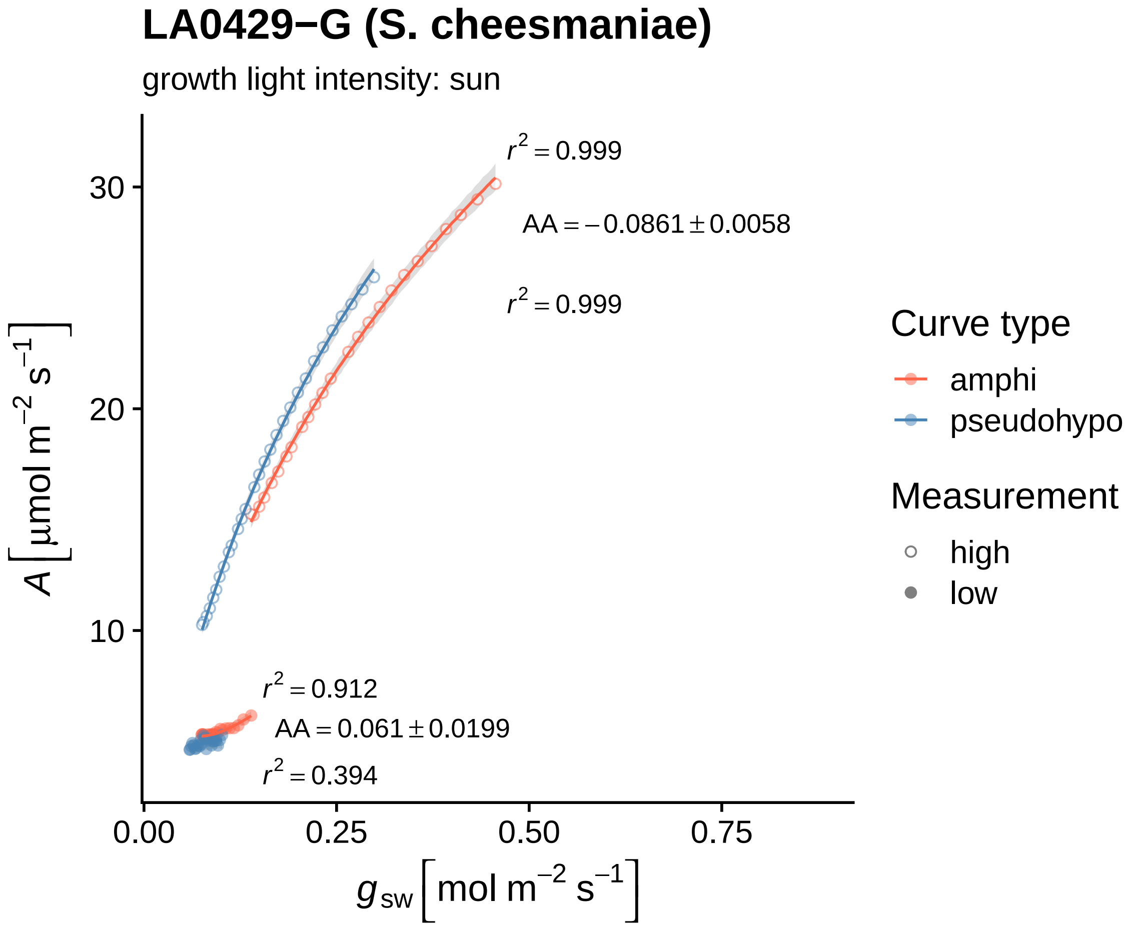

LA0429−G (S. cheesmaniae)
growth light intensity: sun
$r^2 = 0.999$
AA = −0.0861 ± 0.0058
$r^2 = 0.999$
$r^2 = 0.912$
AA = 0.061 ± 0.0199
$r^2 = 0.394$
30
20
10
0.00
0.25
0.50
0.75
A [µmol m⁻² s⁻¹]
g sw [mol m⁻² s⁻¹]
Curve type
amphi
pseudohypo
Measurement
high
low

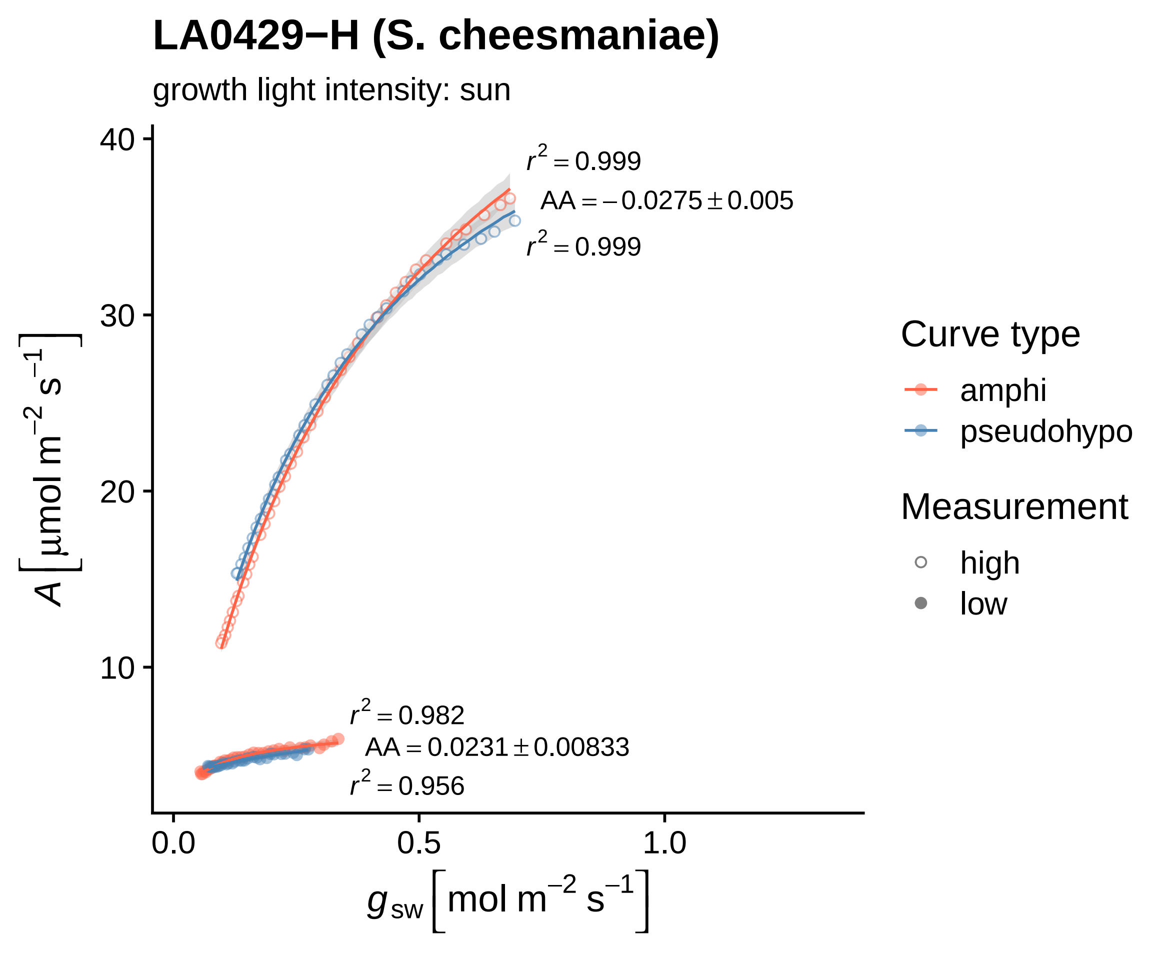

LA0429−H (S. cheesmaniae)
growth light intensity: sun
$r^2 = 0.999$
$AA = -0.0275 \pm 0.005$
$r^2 = 0.999$
$r^2 = 0.982$
$AA = 0.0231 \pm 0.00833$
$r^2 = 0.956$
40
30
20
10
0.0
0.5
1.0
$A\left[\mu mol\ m^{-2}\ s^{-1}\right]$
$g_{sw}\left[mol\ m^{-2}\ s^{-1}\right]$
Curve type
amphi
pseudohypo
Measurement
high
low

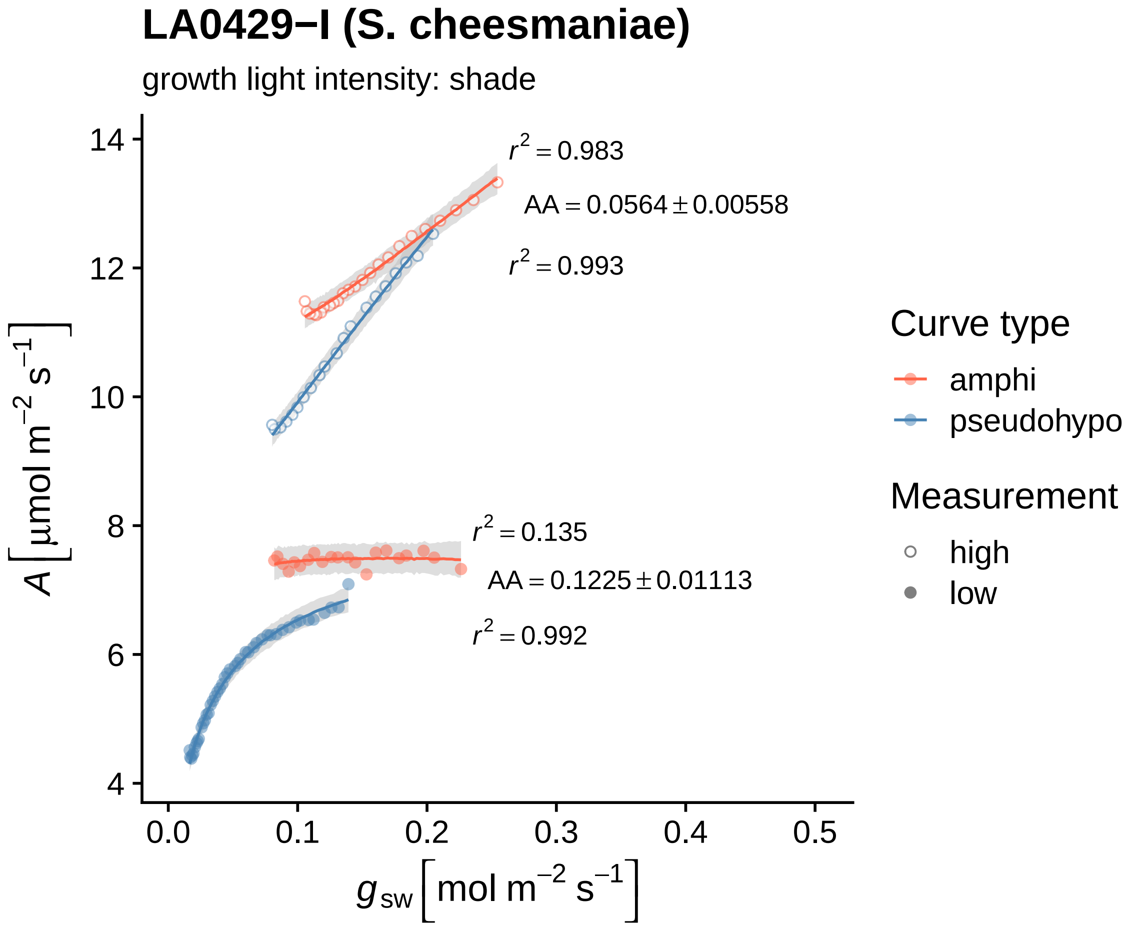

LA0429−I (S. cheesmaniae)
growth light intensity: shade
$r^2 = 0.983$
$AA = 0.0564 \pm 0.00558$
$r^2 = 0.993$
$r^2 = 0.135$
$AA = 0.1225 \pm 0.01113$
$r^2 = 0.992$
$A\left[\mu mol\ m^{-2}\ s^{-1}\right]$
$g_{sw}\left[mol\ m^{-2}\ s^{-1}\right]$
14
12
10
8
6
4
0.0
0.1
0.2
0.3
0.4
0.5
Curve type
amphi
pseudohypo
Measurement
high
low

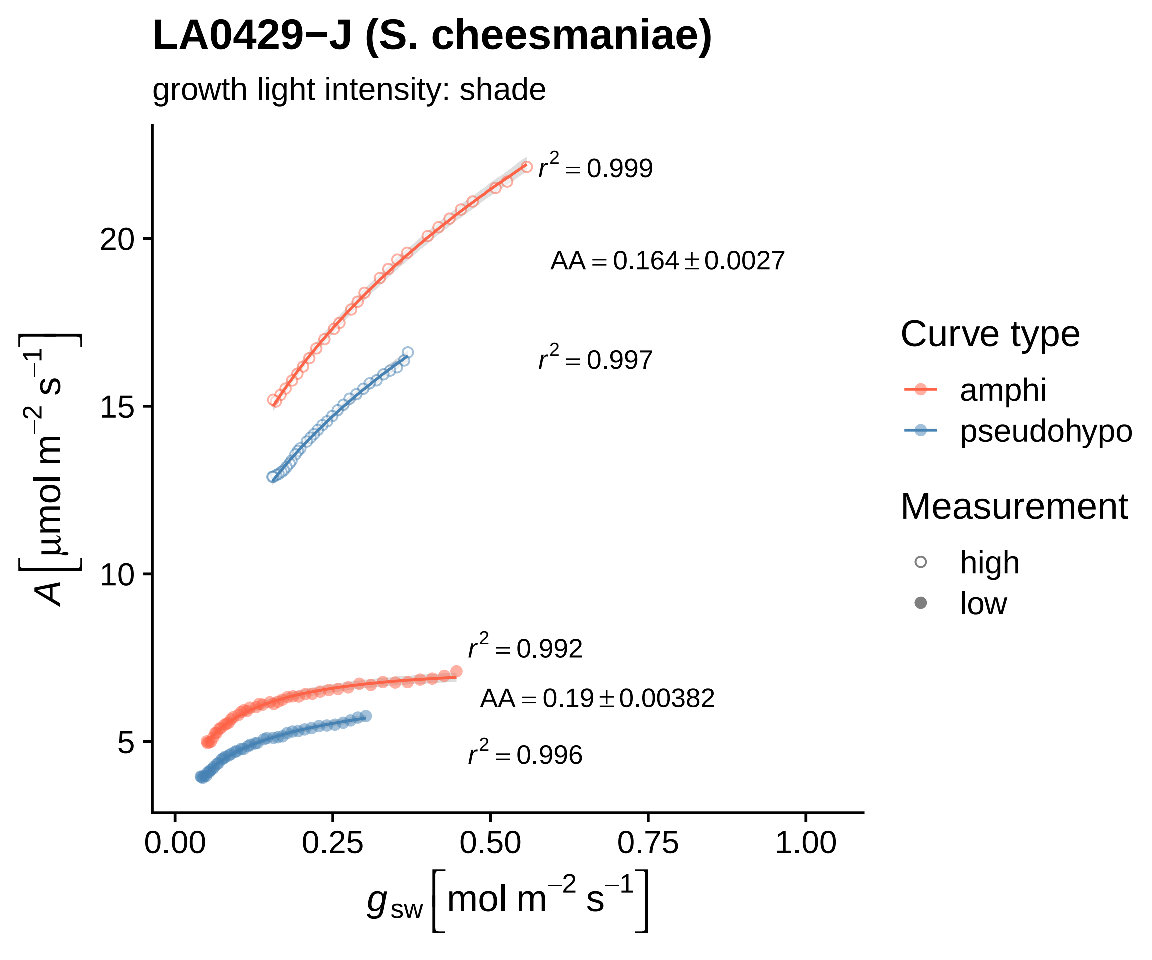
LA0429−J (S. cheesmaniae)
growth light intensity: shade
r^2 = 0.999
AA = 0.164 ± 0.0027
r^2 = 0.997
r^2 = 0.992
AA = 0.19 ± 0.00382
r^2 = 0.996
A [μmol m^−2 s^−1]
g_sw [mol m^−2 s^−1]
20
15
10
5
0.00
0.25
0.50
0.75
1.00
Curve type
amphi
pseudohypo
Measurement
high
low

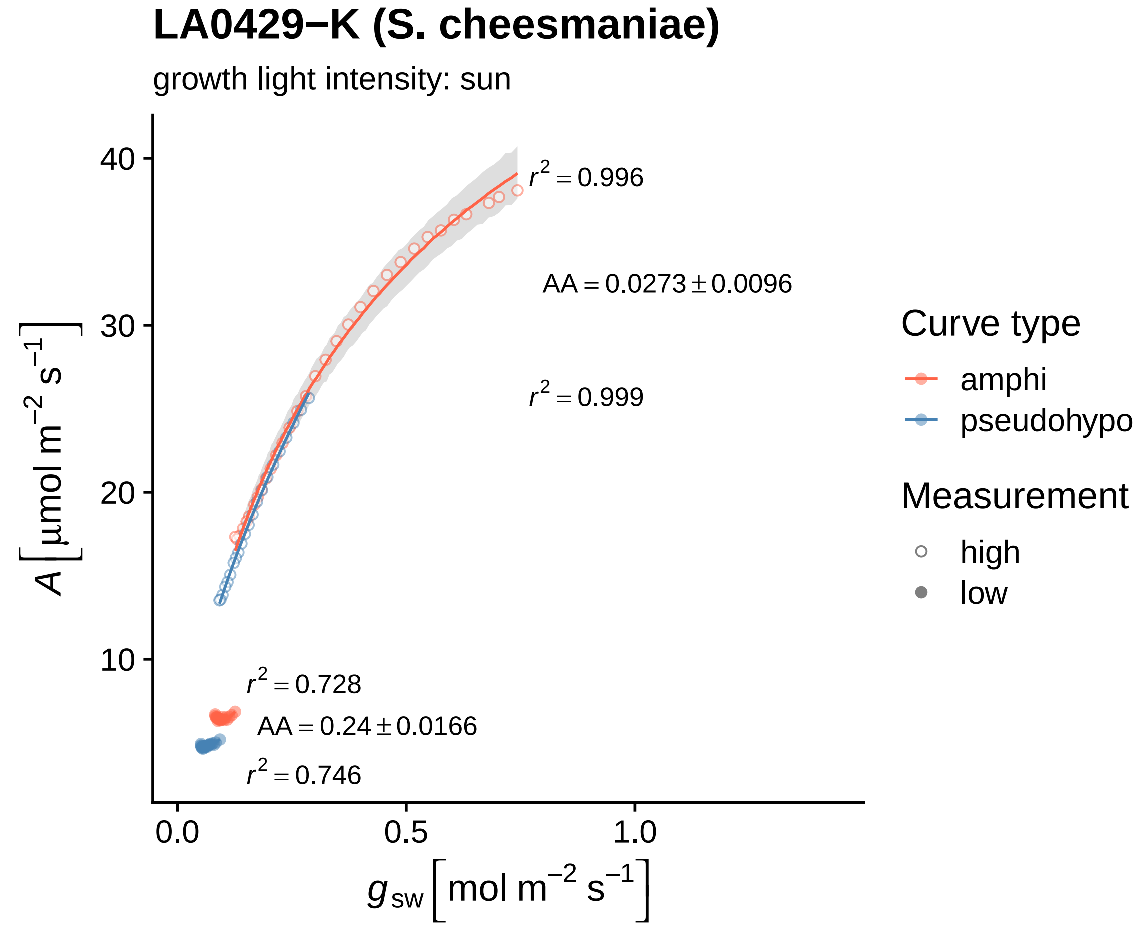
LA0429−K (S. cheesmaniae)
growth light intensity: sun
A [μmol m−2 s−1]
40
30
20
10
r2 = 0.996
AA = 0.0273 ± 0.0096
r2 = 0.999
r2 = 0.728
AA = 0.24 ± 0.0166
r2 = 0.746
0.0
0.5
1.0
gsw [mol m−2 s−1]
Curve type
amphi
pseudohypo
Measurement
high
low

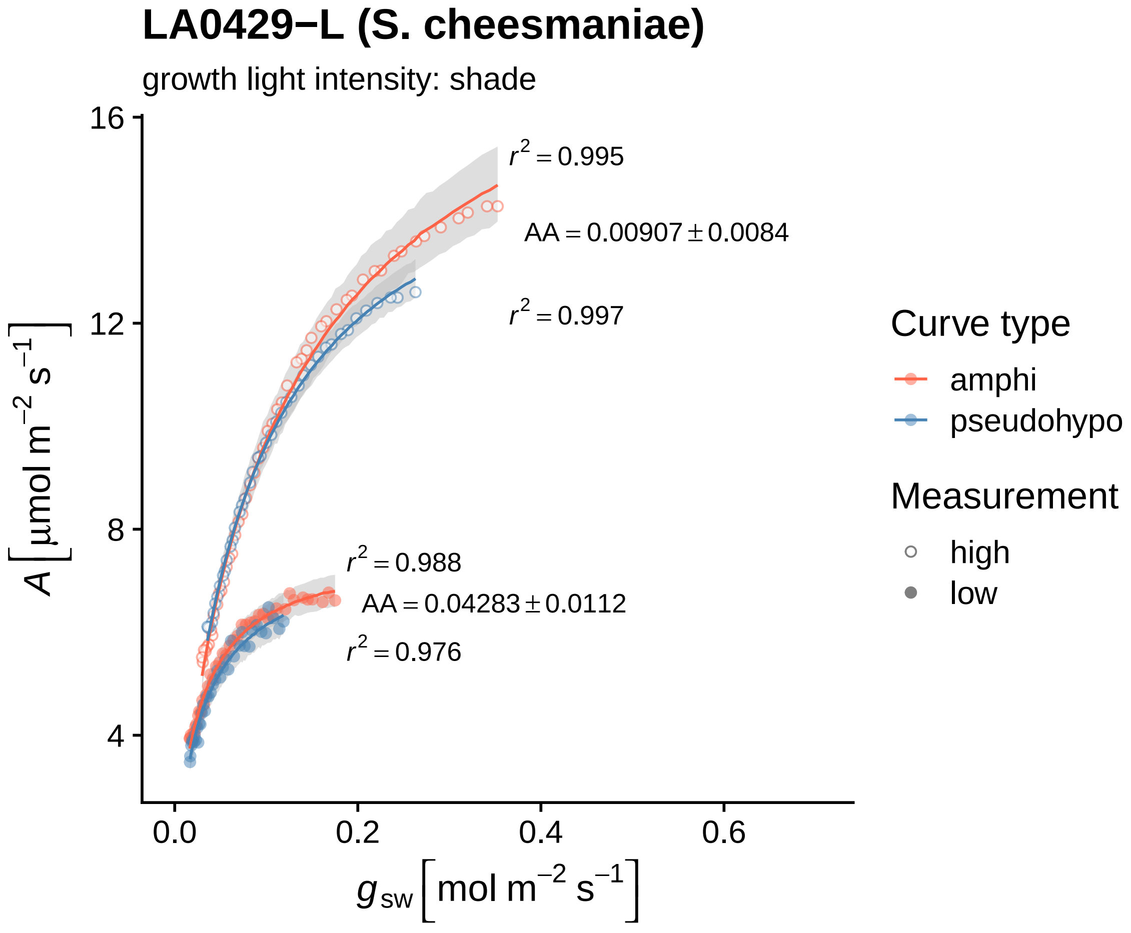
LA0429−L (S. cheesmaniae)
growth light intensity: shade
16
12
8
4
0.0
0.2
0.4
0.6
A [µmol m−2 s−1]
gsw [mol m−2 s−1]
r2 = 0.995
AA = 0.00907 ± 0.0084
r2 = 0.997
r2 = 0.988
AA = 0.04283 ± 0.0112
r2 = 0.976
Curve type
amphi
pseudohypo
Measurement
high
low

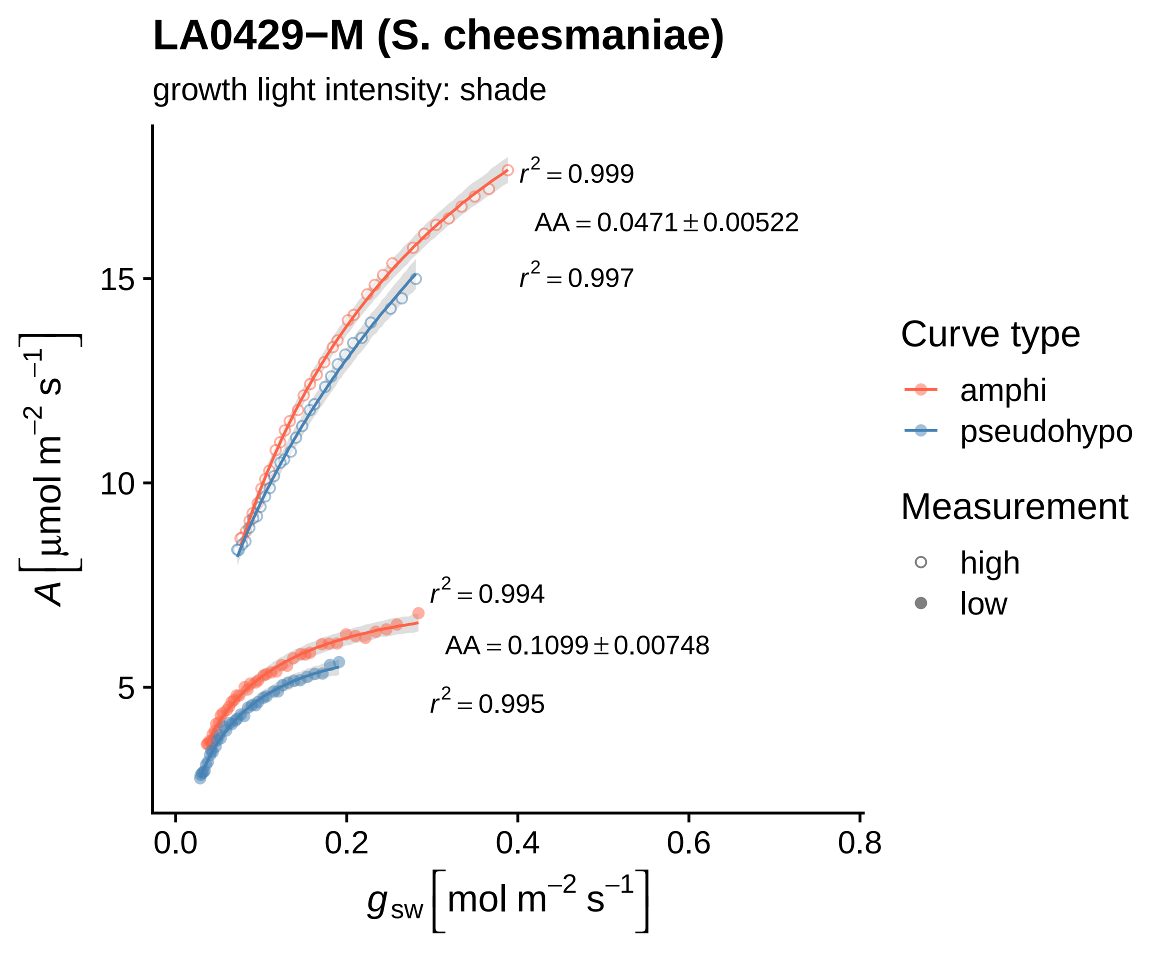

LA0429−M (S. cheesmaniae)
growth light intensity: shade
$r^2 = 0.999$
$AA = 0.0471 \pm 0.00522$
$r^2 = 0.997$
$r^2 = 0.994$
$AA = 0.1099 \pm 0.00748$
$r^2 = 0.995$
$A\ [\mu mol\ m^{-2}\ s^{-1}]$
$g_{sw}\ [mol\ m^{-2}\ s^{-1}]$
5
10
15
0.0
0.2
0.4
0.6
0.8
Curve type
amphi
pseudohypo
Measurement
high
low

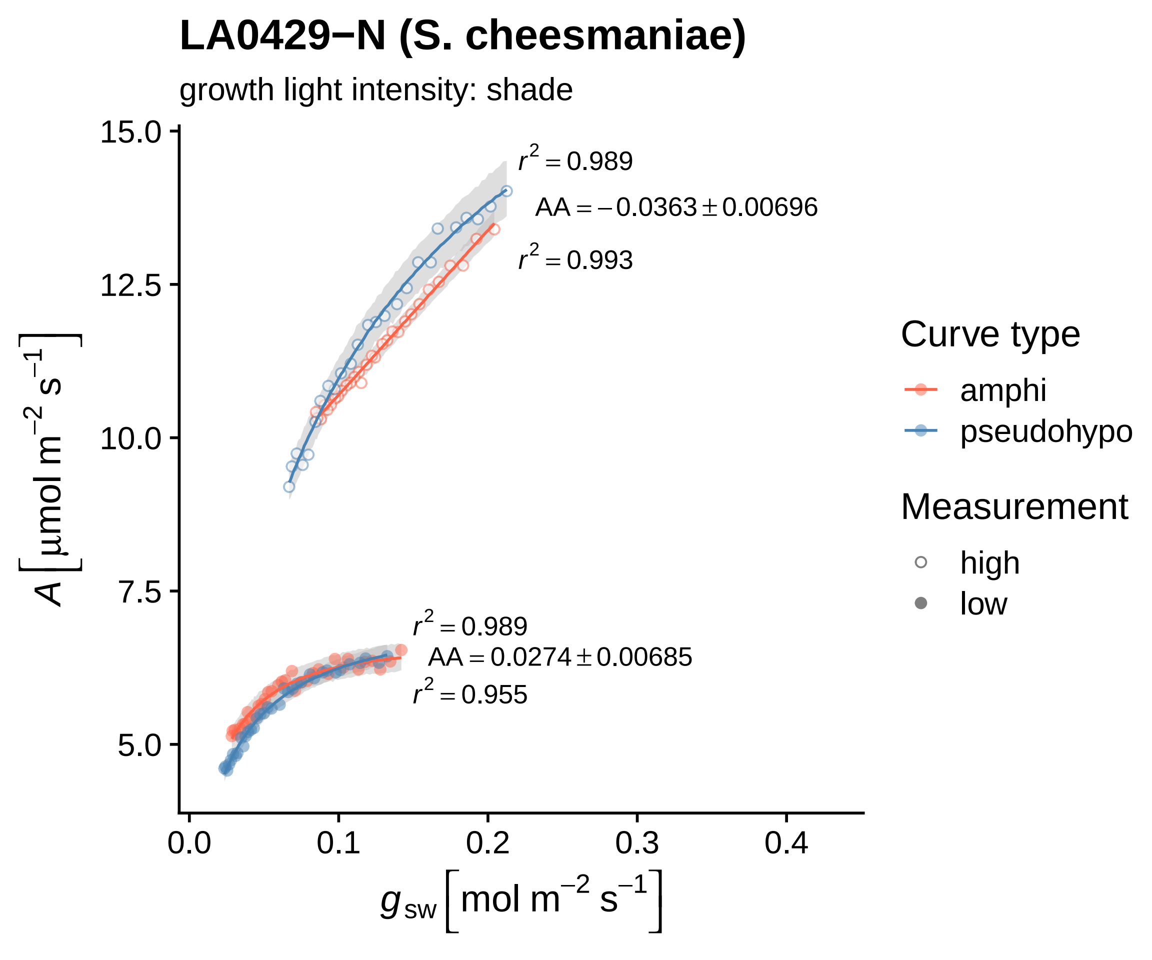

LA0429−N (S. cheesmaniae)
growth light intensity: shade
15.0
12.5
10.0
7.5
5.0
0.0
0.1
0.2
0.3
0.4
r2 = 0.989
AA = −0.0363 ± 0.00696
r2 = 0.993
r2 = 0.989
AA = 0.0274 ± 0.00685
r2 = 0.955
A [μmol m−2 s−1]
gsw [mol m−2 s−1]
Curve type
amphi
pseudohypo
Measurement
high
low

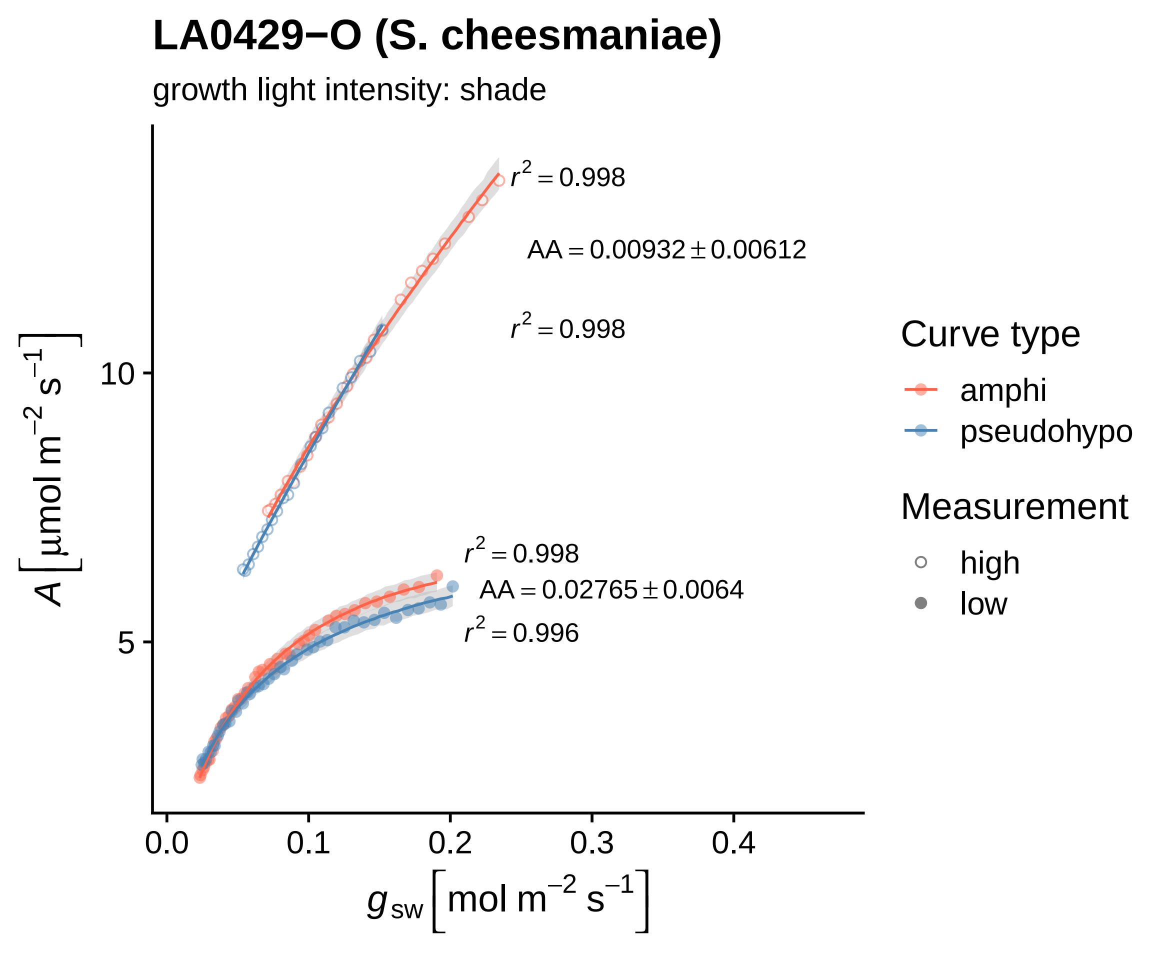
LA0429−O (S. cheesmaniae)
growth light intensity: shade
$r^2 = 0.998$
AA = 0.00932 ± 0.00612
$r^2 = 0.998$
$r^2 = 0.998$
AA = 0.02765 ± 0.0064
$r^2 = 0.996$
10
5
0.0
0.1
0.2
0.3
0.4
$A\,[\mu mol\ m^{-2}\ s^{-1}]$
$g_{sw}\,[mol\ m^{-2}\ s^{-1}]$
Curve type
amphi
pseudohypo
Measurement
high
low

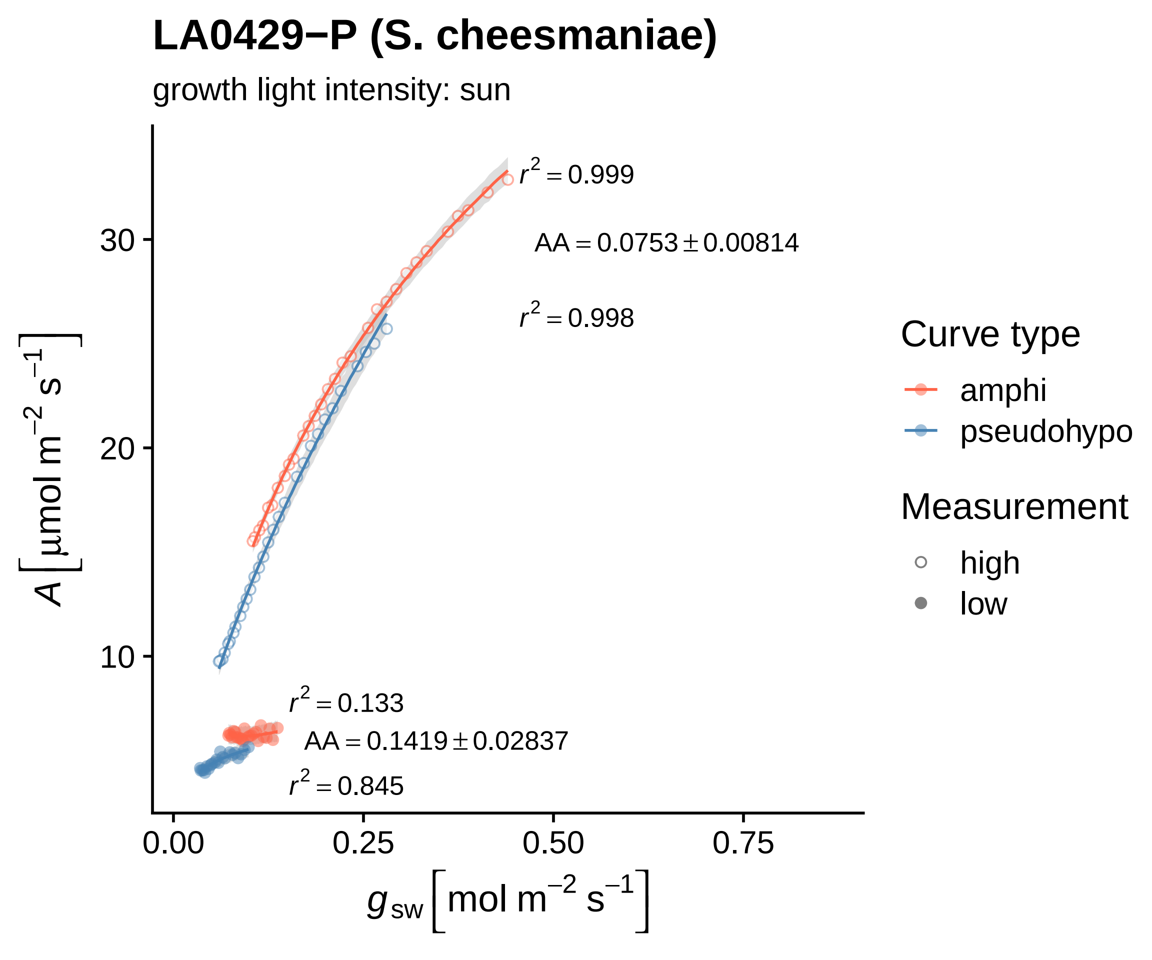
LA0429−P (S. cheesmaniae)
growth light intensity: sun
$r^2 = 0.999$
$AA = 0.0753 \pm 0.00814$
$r^2 = 0.998$
$r^2 = 0.133$
$AA = 0.1419 \pm 0.02837$
$r^2 = 0.845$
30
20
10
0.00
0.25
0.50
0.75
$A\left[\mu mol\ m^{-2}\ s^{-1}\right]$
$g_{sw}\left[mol\ m^{-2}\ s^{-1}\right]$
Curve type
amphi
pseudohypo
Measurement
high
low

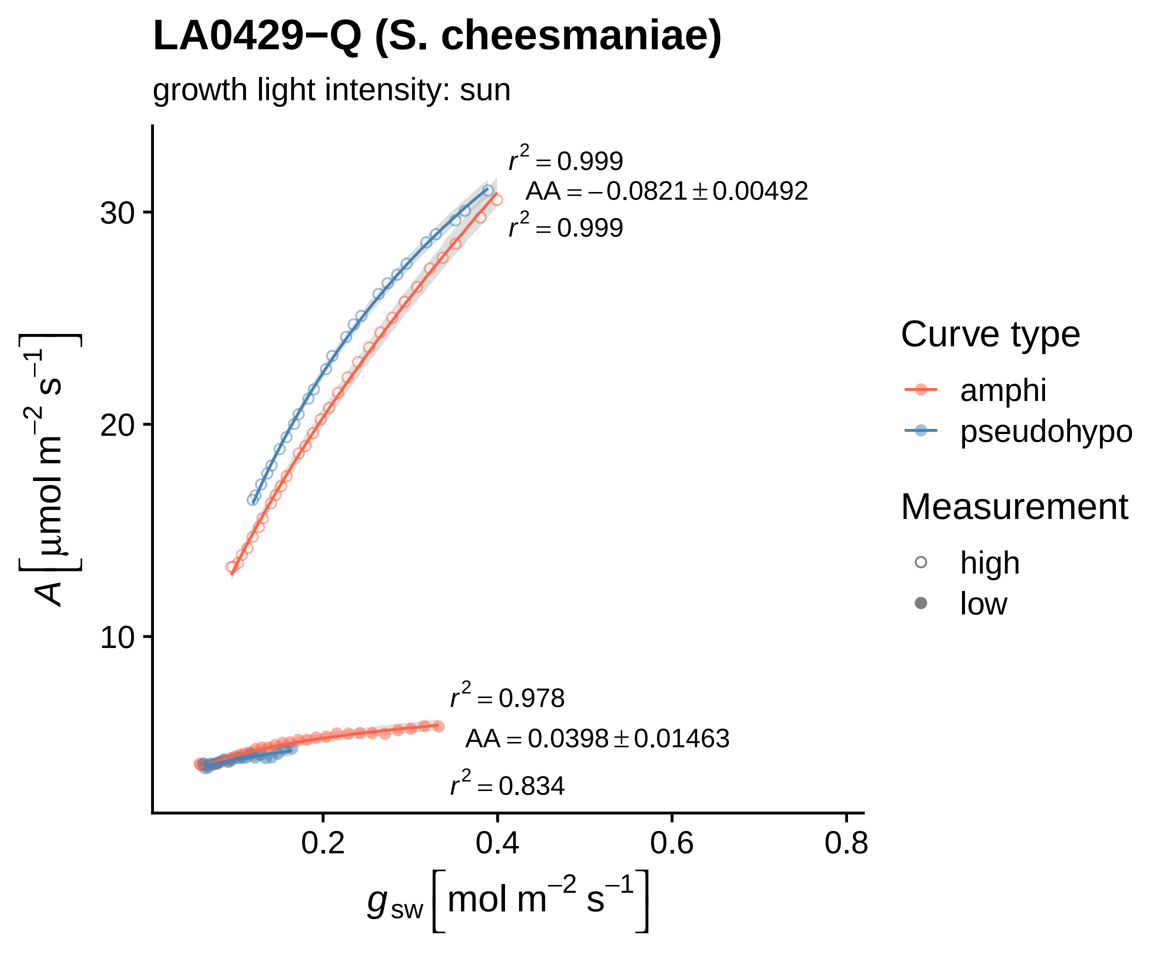
LA0429−Q (S. cheesmaniae)
growth light intensity: sun
$r^2 = 0.999$
$AA = -0.0821 \pm 0.00492$
$r^2 = 0.999$
$r^2 = 0.978$
$AA = 0.0398 \pm 0.01463$
$r^2 = 0.834$
30
20
10
0.2
0.4
0.6
0.8
$A\ [\mu mol\ m^{-2}\ s^{-1}]$
$g_{sw}\ [mol\ m^{-2}\ s^{-1}]$
Curve type
amphi
pseudohypo
Measurement
high
low

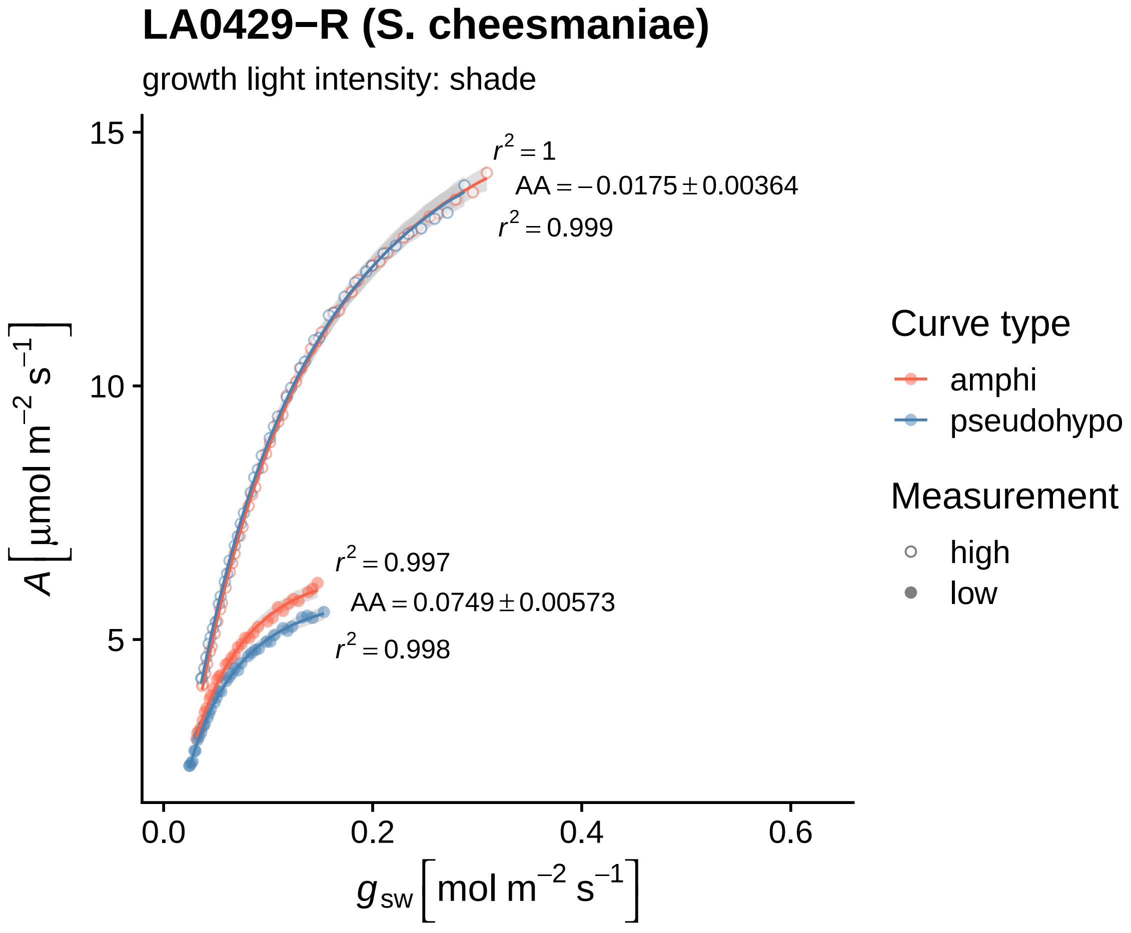
LA0429−R (S. cheesmaniae)
growth light intensity: shade
$r^2 = 1$
$AA = -0.0175 \pm 0.00364$
$r^2 = 0.999$
$r^2 = 0.997$
$AA = 0.0749 \pm 0.00573$
$r^2 = 0.998$
15
10
5
0.0
0.2
0.4
0.6
$A\ [\mu mol\ m^{-2}\ s^{-1}]$
$g_{sw}\ [mol\ m^{-2}\ s^{-1}]$
Curve type
amphi
pseudohypo
Measurement
high
low

### LA0429−S (S. cheesmaniae)

growth light intensity: sun

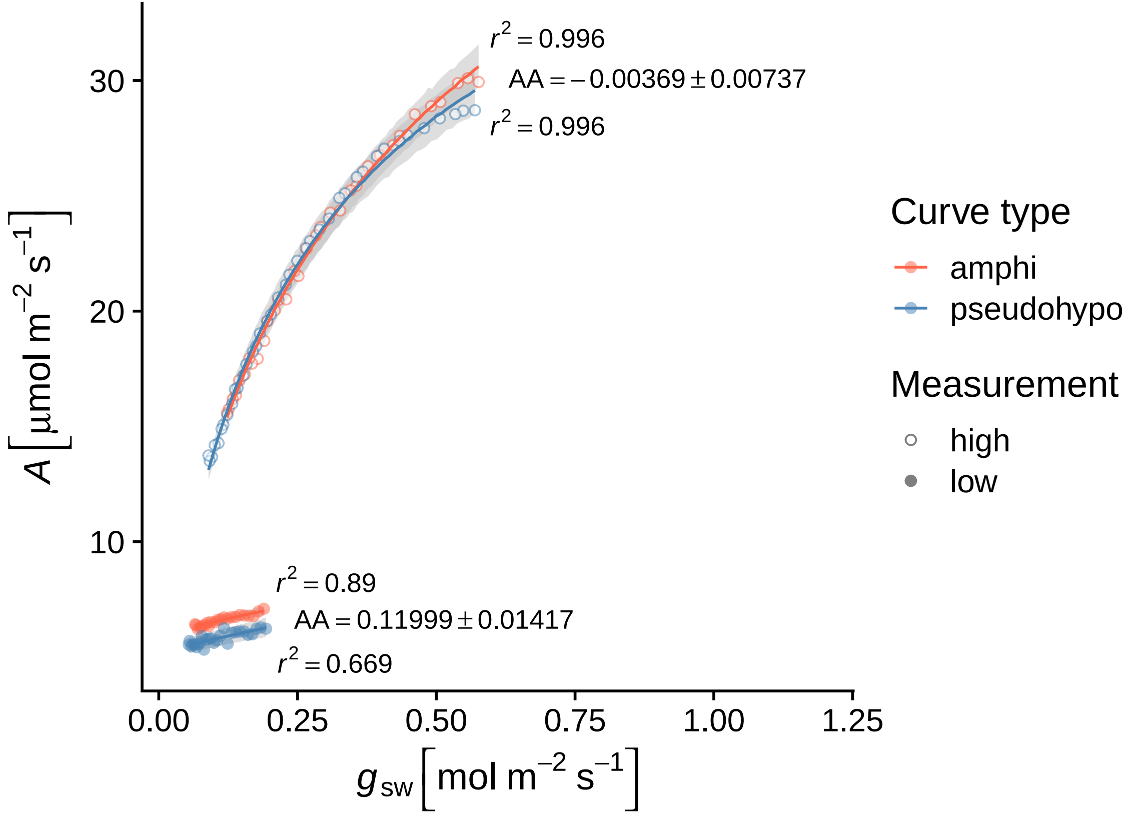

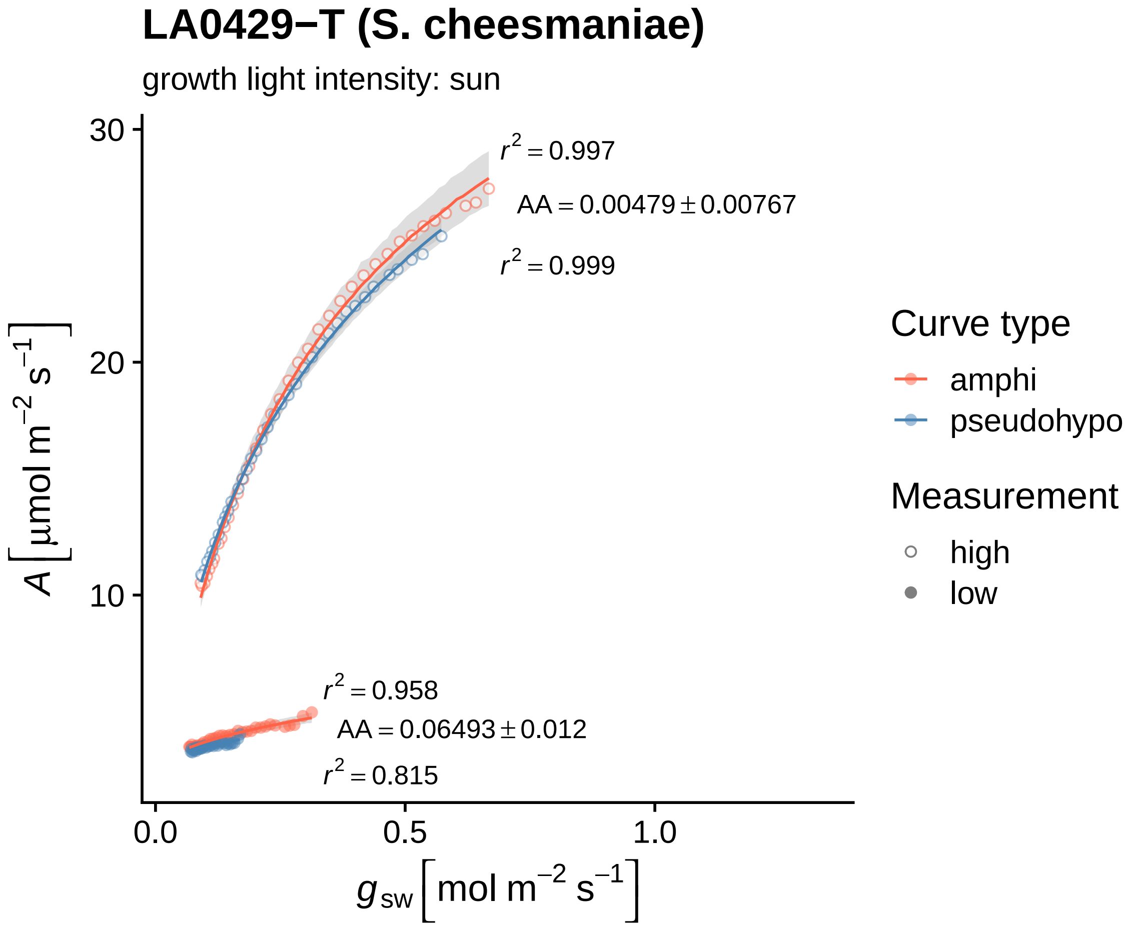
LA0429−T (S. cheesmaniae)
growth light intensity: sun
$r^2 = 0.997$
$AA = 0.00479 \pm 0.00767$
$r^2 = 0.999$
$r^2 = 0.958$
$AA = 0.06493 \pm 0.012$
$r^2 = 0.815$
30
20
10
0.0
0.5
1.0
$A\ [\mu mol\ m^{-2}\ s^{-1}]$
$g_{sw}\ [mol\ m^{-2}\ s^{-1}]$
Curve type
amphi
pseudohypo
Measurement
high
low

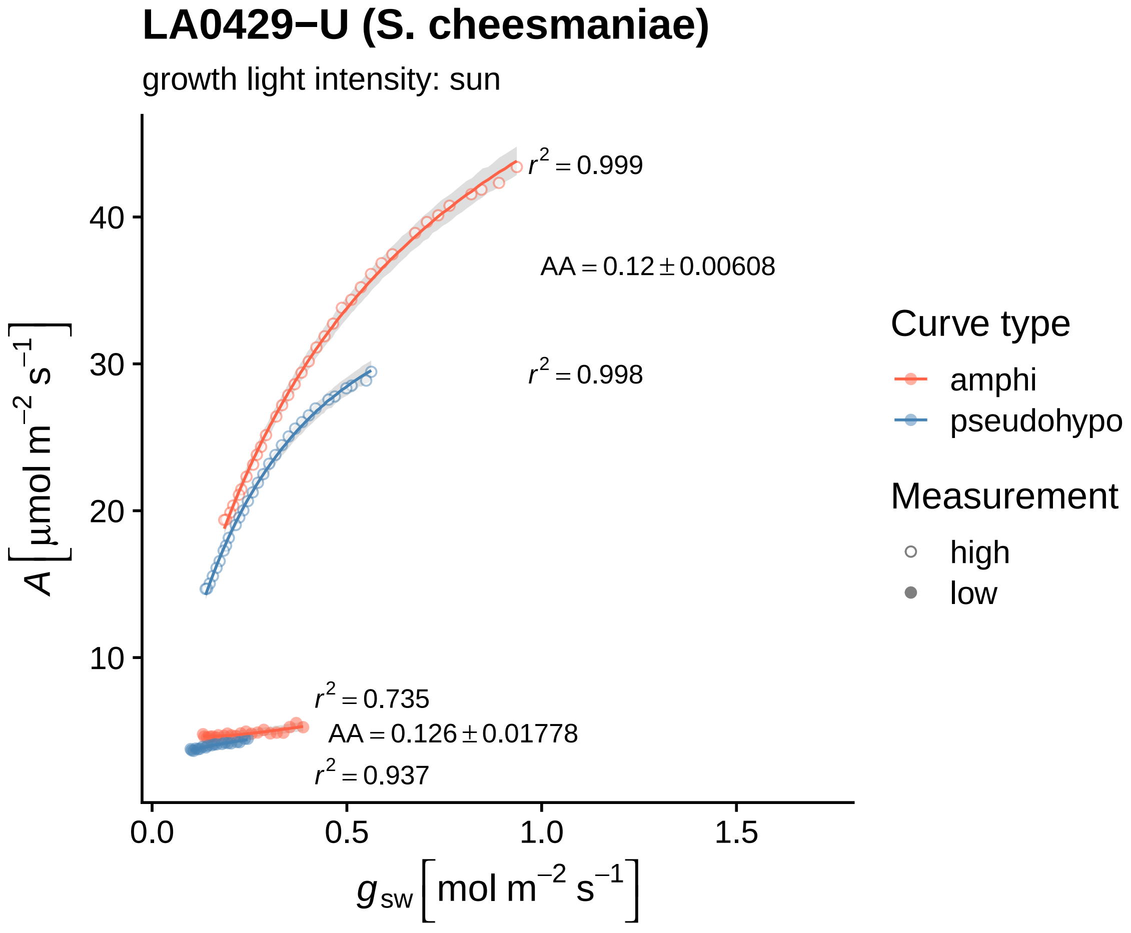

LA0429−U (S. cheesmaniae)
growth light intensity: sun
$r^2 = 0.999$
$AA = 0.12 \pm 0.00608$
$r^2 = 0.998$
$r^2 = 0.735$
$AA = 0.126 \pm 0.01778$
$r^2 = 0.937$
40
30
20
10
0.0
0.5
1.0
1.5
$A\left[\mu mol\ m^{-2}\ s^{-1}\right]$
$g_{sw}\left[mol\ m^{-2}\ s^{-1}\right]$
Curve type
amphi
pseudohypo
Measurement
high
low

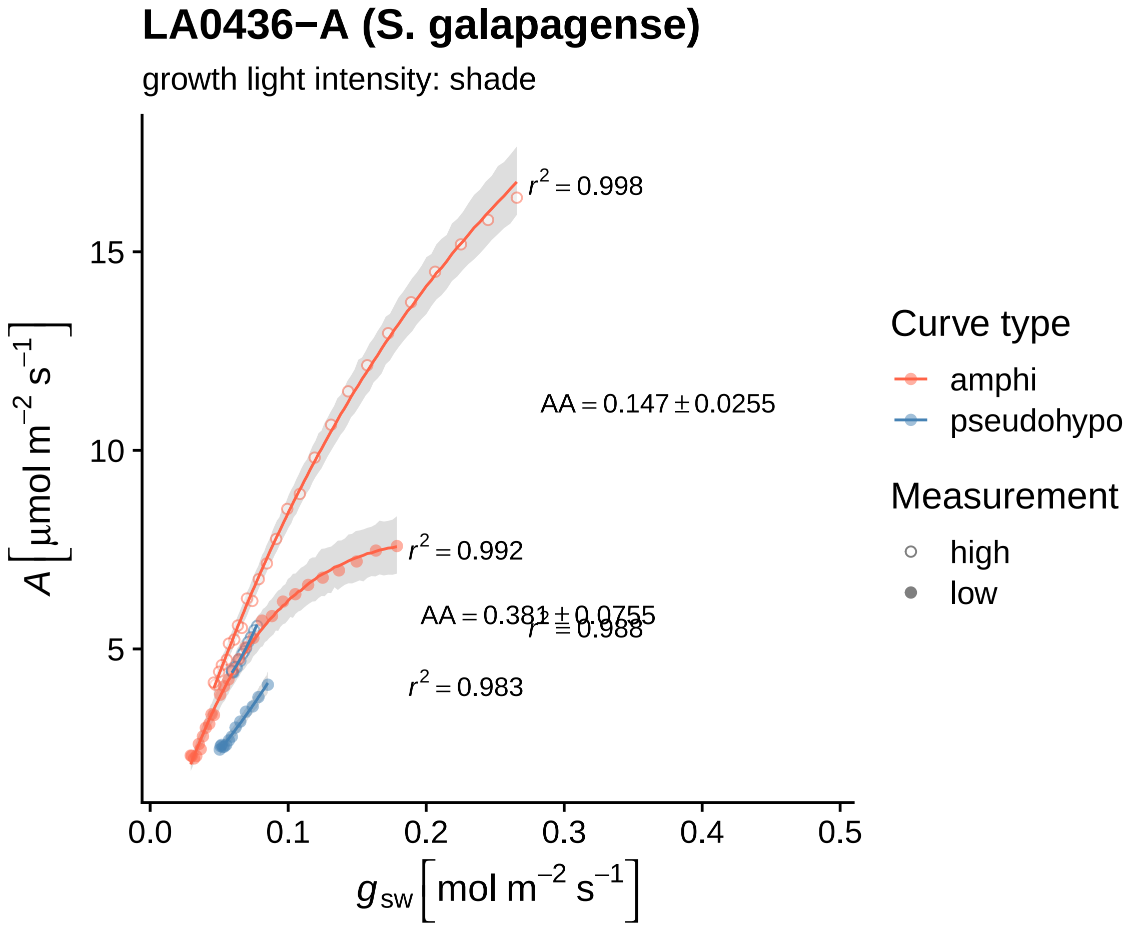

LA0436−A (S. galapagense)
growth light intensity: shade
$r^2 = 0.998$
$AA = 0.147 \pm 0.0255$
$r^2 = 0.992$
$AA = 0.381 \pm 0.0755$
$r^2 = 0.988$
$r^2 = 0.983$
15
10
5
0.0
0.1
0.2
0.3
0.4
0.5
$A\left[\mu mol\ m^{-2}\ s^{-1}\right]$
$g_{sw}\left[mol\ m^{-2}\ s^{-1}\right]$
Curve type
amphi
pseudohypo
Measurement
high
low

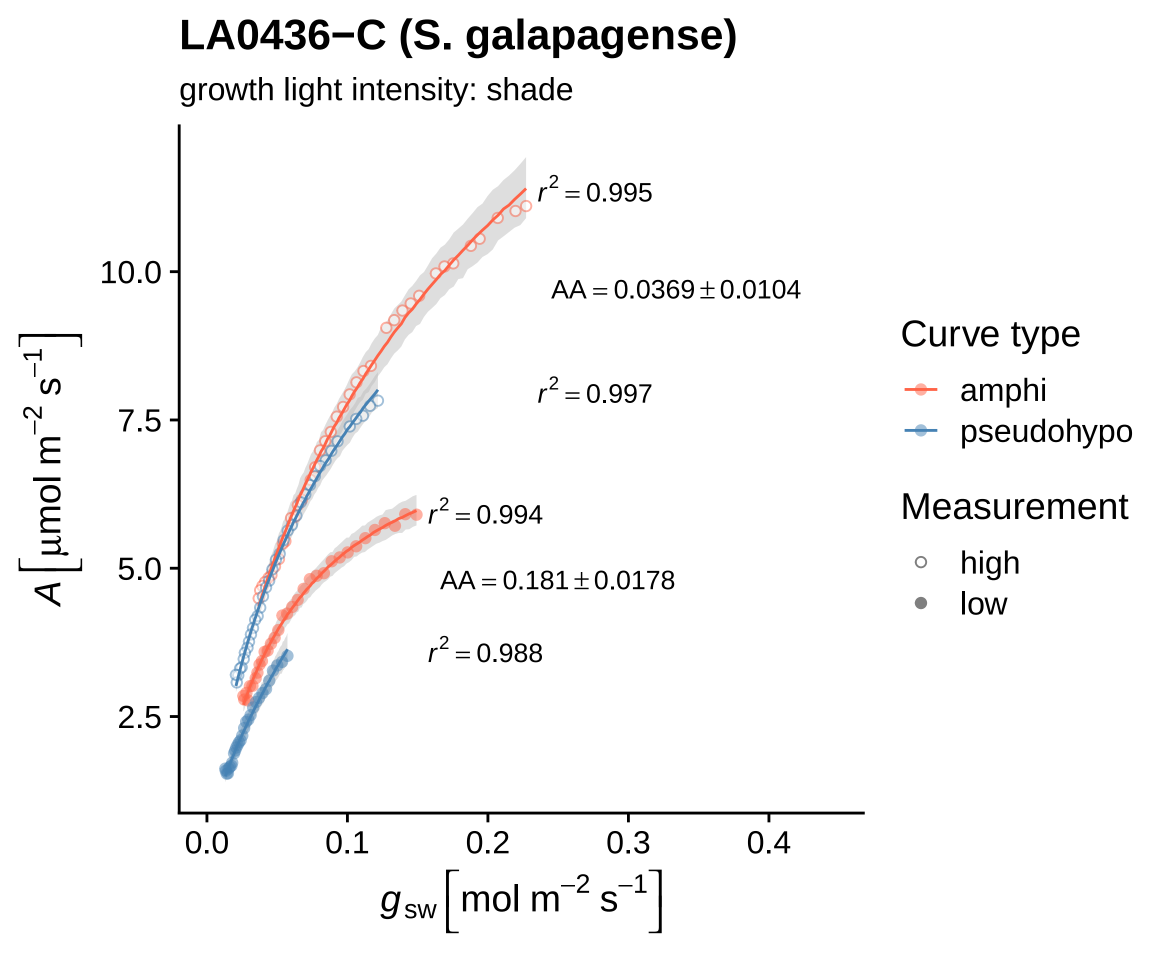
LA0436−C (S. galapagense)
growth light intensity: shade
A [μmol m−2 s−1]
10.0
7.5
5.0
2.5
0.0
0.1
0.2
0.3
0.4
gsw [mol m−2 s−1]
r2 = 0.995
AA = 0.0369 ± 0.0104
r2 = 0.997
r2 = 0.994
AA = 0.181 ± 0.0178
r2 = 0.988
Curve type
amphi
pseudohypo
Measurement
high
low

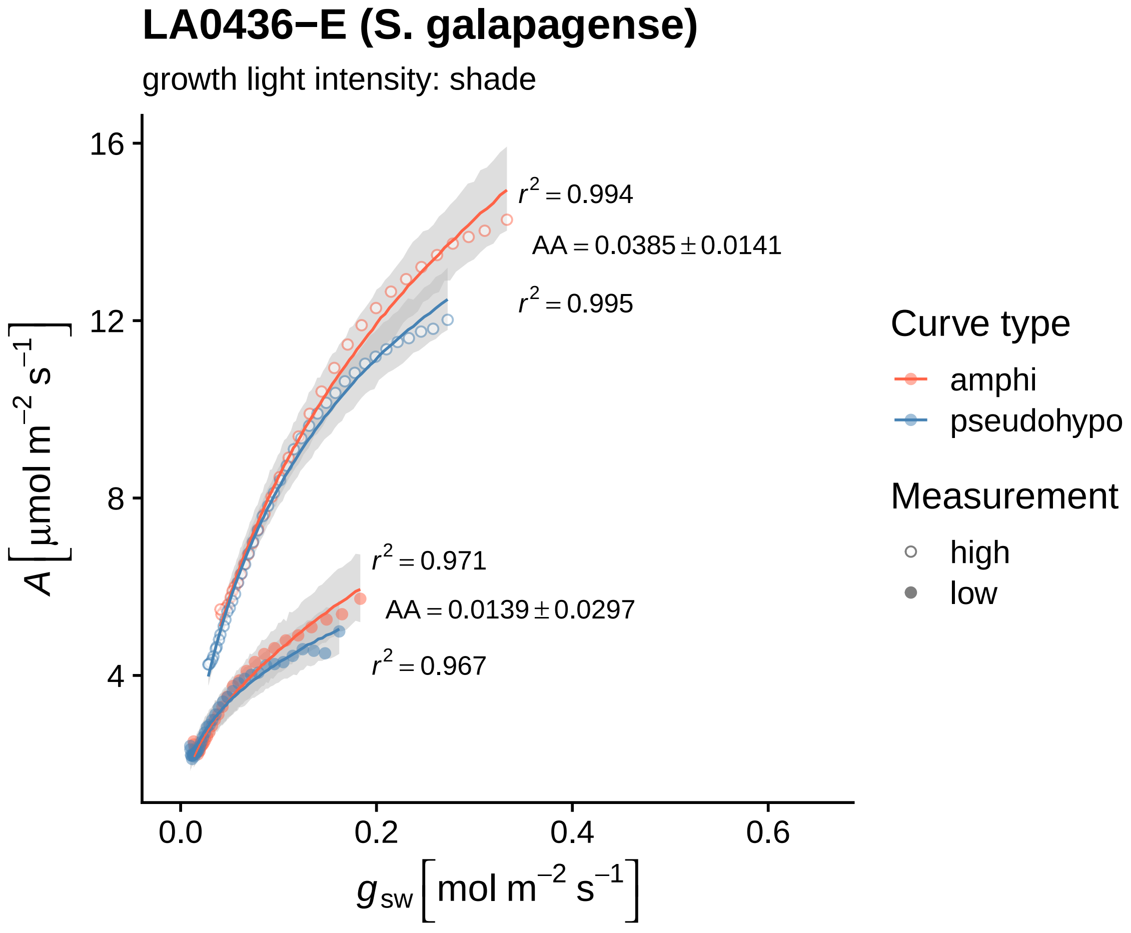
LA0436−E (S. galapagense)
growth light intensity: shade
$r^2 = 0.994$
$AA = 0.0385 \pm 0.0141$
$r^2 = 0.995$
$r^2 = 0.971$
$AA = 0.0139 \pm 0.0297$
$r^2 = 0.967$
16
12
8
4
0.0
0.2
0.4
0.6
$A\ [\mu mol\ m^{-2}\ s^{-1}]$
$g_{sw}\ [mol\ m^{-2}\ s^{-1}]$
Curve type
amphi
pseudohypo
Measurement
high
low

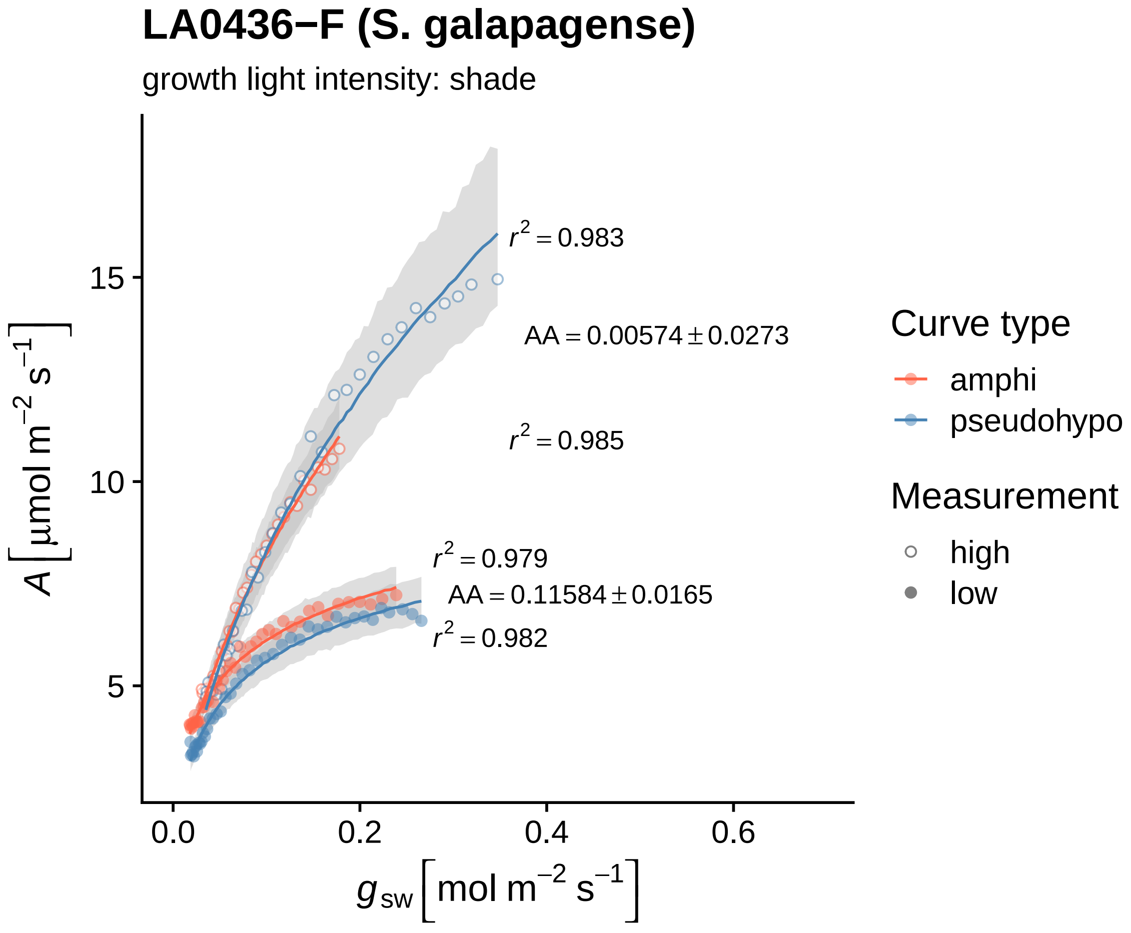

LA0436−F (S. galapagense)
growth light intensity: shade
A [μmol m−2 s−1]
15
10
5
0.0
0.2
0.4
0.6
gsw [mol m−2 s−1]
r2 = 0.983
AA = 0.00574 ± 0.0273
r2 = 0.985
r2 = 0.979
AA = 0.11584 ± 0.0165
r2 = 0.982
Curve type
amphi
pseudohypo
Measurement
high
low

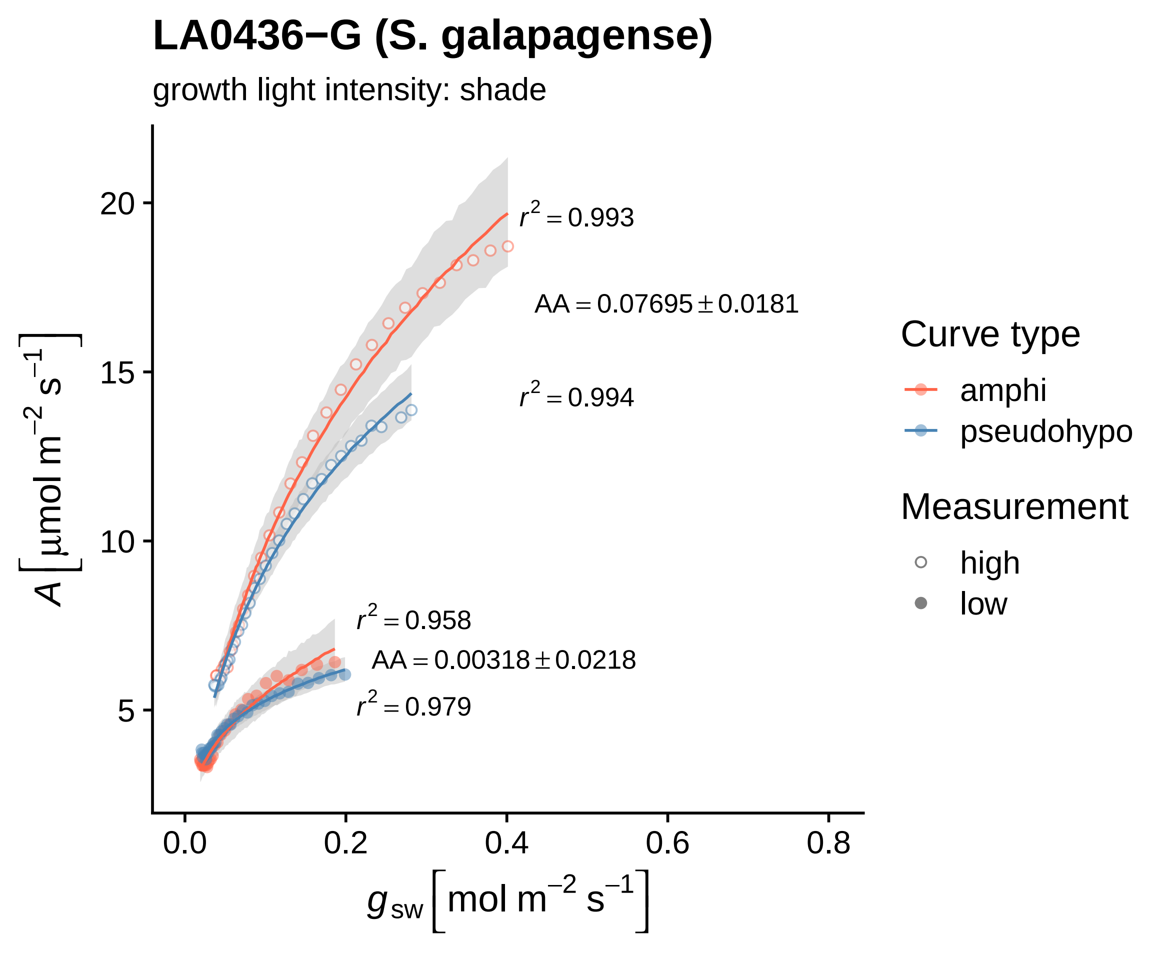

LA0436−G (S. galapagense)
growth light intensity: shade
$r^2 = 0.993$
$AA = 0.07695 \pm 0.0181$
$r^2 = 0.994$
$r^2 = 0.958$
$AA = 0.00318 \pm 0.0218$
$r^2 = 0.979$
20
15
10
5
0.0
0.2
0.4
0.6
0.8
$A\ [\mu mol\ m^{-2}\ s^{-1}]$
$g_{sw}\ [mol\ m^{-2}\ s^{-1}]$
Curve type
amphi
pseudohypo
Measurement
high
low

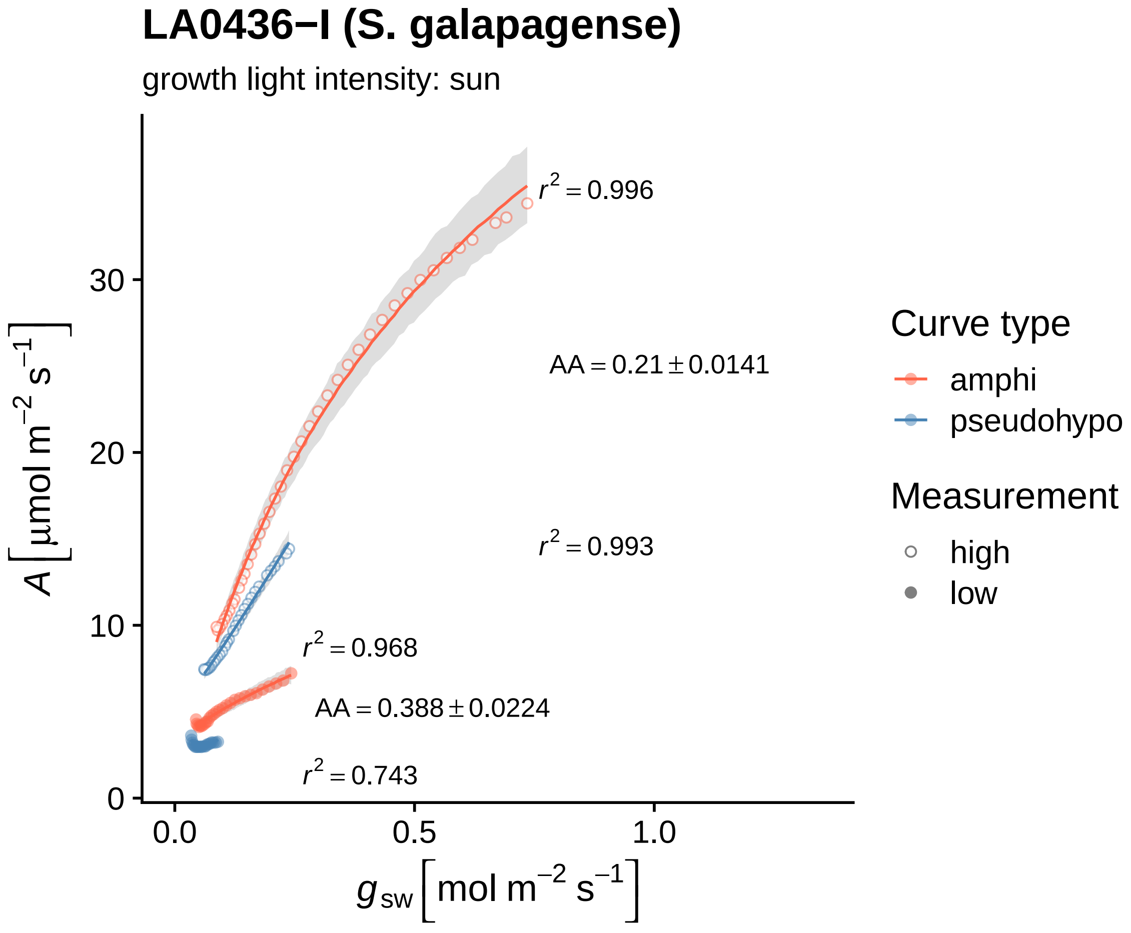

LA0436−I (S. galapagense)
growth light intensity: sun
$r^2 = 0.996$
$AA = 0.21 \pm 0.0141$
$r^2 = 0.993$
$r^2 = 0.968$
$AA = 0.388 \pm 0.0224$
$r^2 = 0.743$
30
20
10
0
0.0
0.5
1.0
$A\ [\mu mol\ m^{-2}\ s^{-1}]$
$g_{sw}\ [mol\ m^{-2}\ s^{-1}]$
Curve type
amphi
pseudohypo
Measurement
high
low

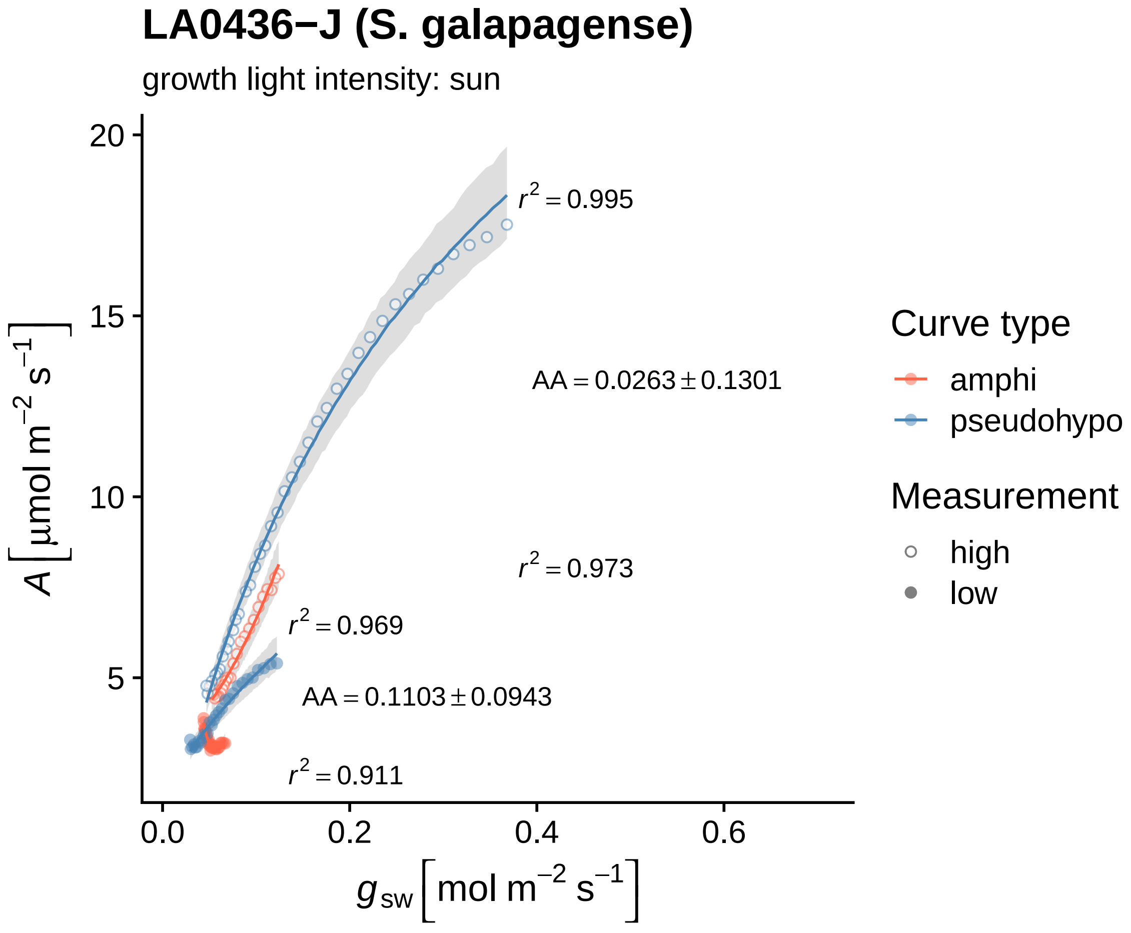

LA0436−J (S. galapagense)
growth light intensity: sun
A [μmol m−2 s−1]
20
15
10
5
0.0
0.2
0.4
0.6
gsw [mol m−2 s−1]
r2 = 0.995
AA = 0.0263 ± 0.1301
r2 = 0.973
r2 = 0.969
AA = 0.1103 ± 0.0943
r2 = 0.911
Curve type
amphi
pseudohypo
Measurement
high
low

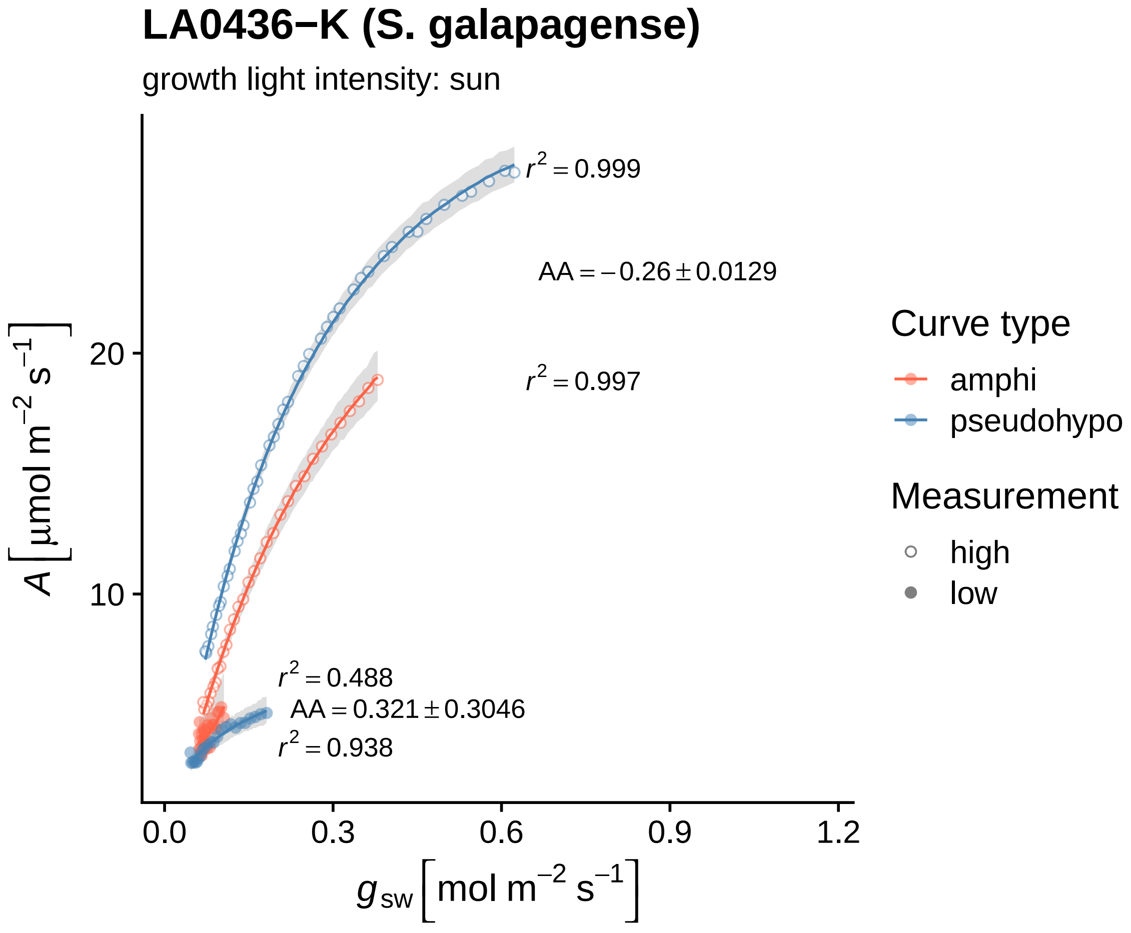

LA0436−K (S. galapagense)
growth light intensity: sun
$r^2 = 0.999$
$AA = -0.26 \pm 0.0129$
$r^2 = 0.997$
$r^2 = 0.488$
$AA = 0.321 \pm 0.3046$
$r^2 = 0.938$
$A\left[\mu mol\ m^{-2}\ s^{-1}\right]$
10
20
0.0
0.3
0.6
0.9
1.2
$g_{sw}\left[mol\ m^{-2}\ s^{-1}\right]$
Curve type
amphi
pseudohypo
Measurement
high
low

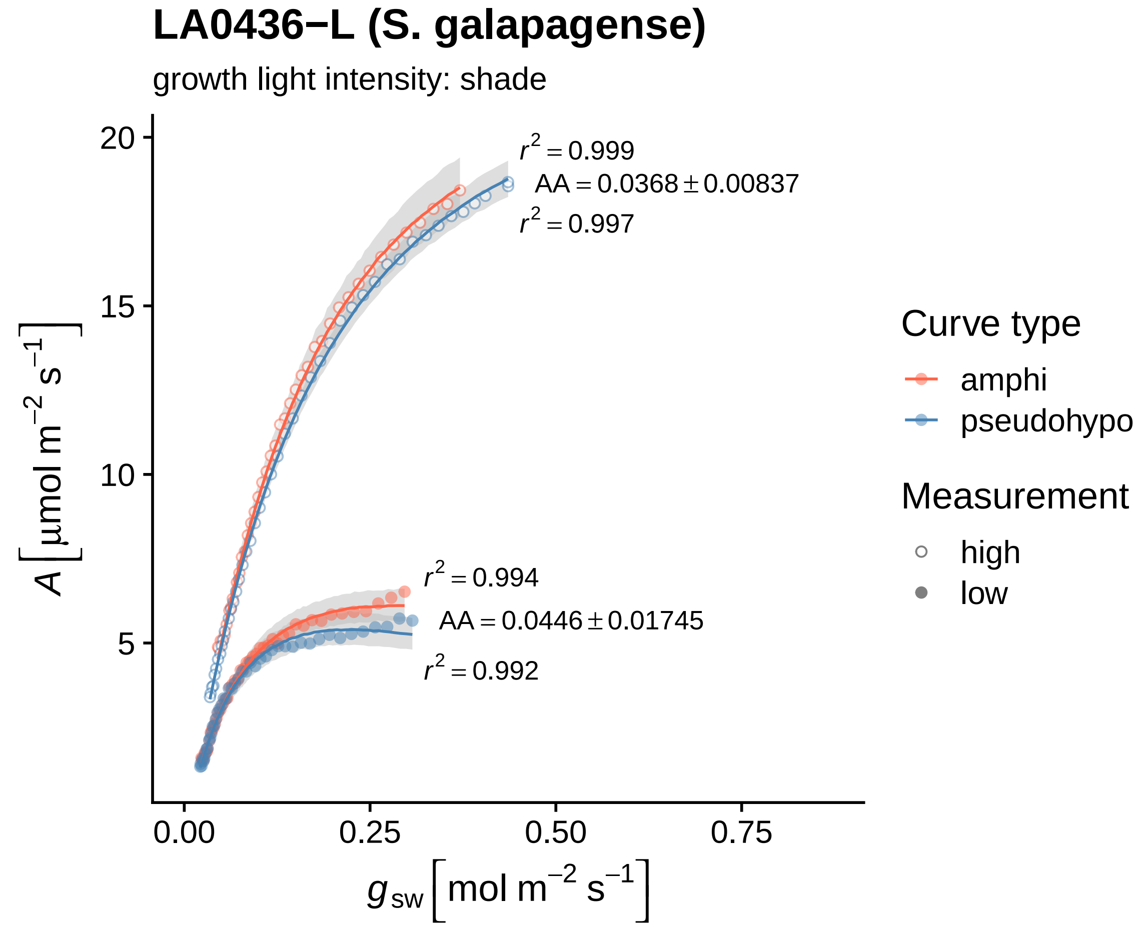

LA0436−L (S. galapagense)
growth light intensity: shade
$r^2 = 0.999$
AA = 0.0368 ± 0.00837
$r^2 = 0.997$
$r^2 = 0.994$
AA = 0.0446 ± 0.01745
$r^2 = 0.992$
20
15
10
5
0.00
0.25
0.50
0.75
$A\,[\mu mol\ m^{-2}\ s^{-1}]$
$g_{sw}\,[mol\ m^{-2}\ s^{-1}]$
Curve type
amphi
pseudohypo
Measurement
high
low

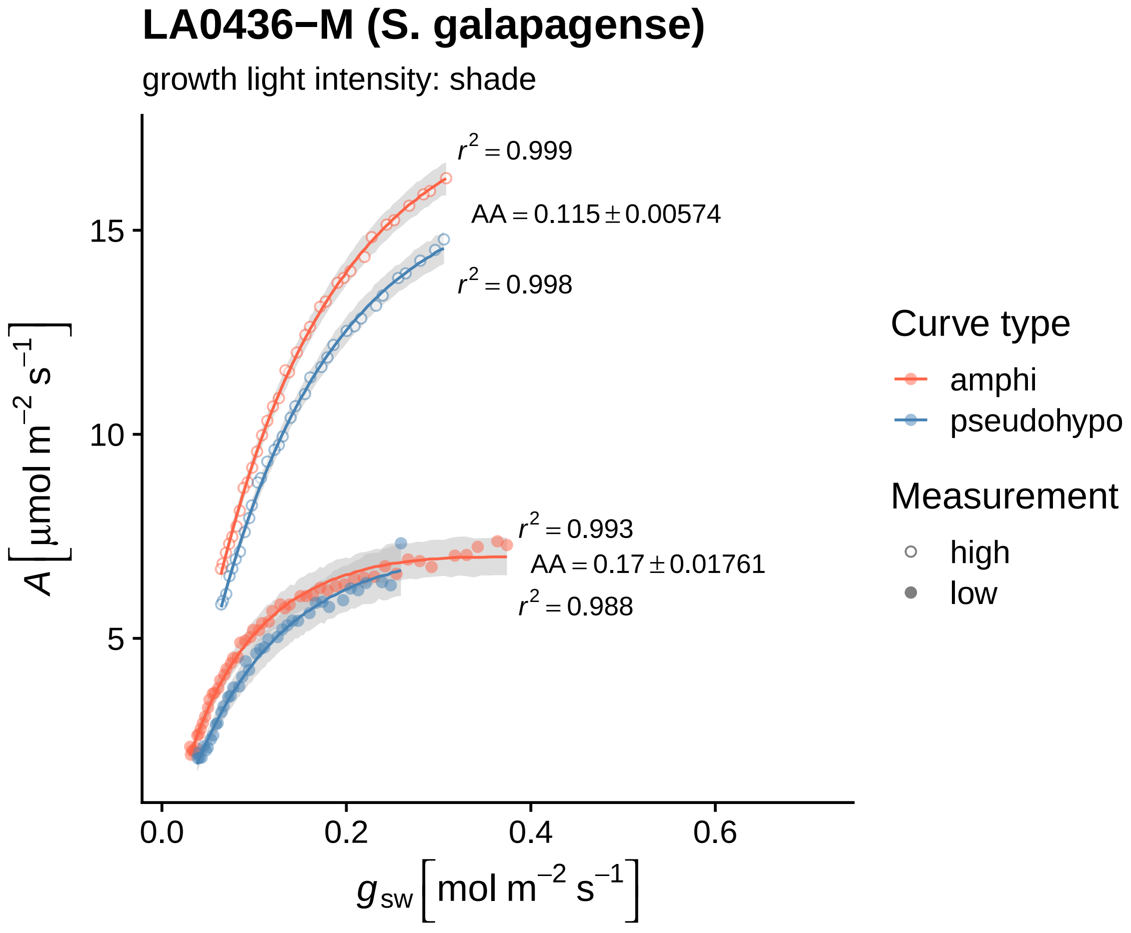

LA0436−M (S. galapagense)
growth light intensity: shade
$r^2 = 0.999$
$AA = 0.115 \pm 0.00574$
$r^2 = 0.998$
$r^2 = 0.993$
$AA = 0.17 \pm 0.01761$
$r^2 = 0.988$
$A\left[\mu mol\ m^{-2}\ s^{-1}\right]$
15
10
5
0.0
0.2
0.4
0.6
$g_{sw}\left[mol\ m^{-2}\ s^{-1}\right]$
Curve type
amphi
pseudohypo
Measurement
high
low

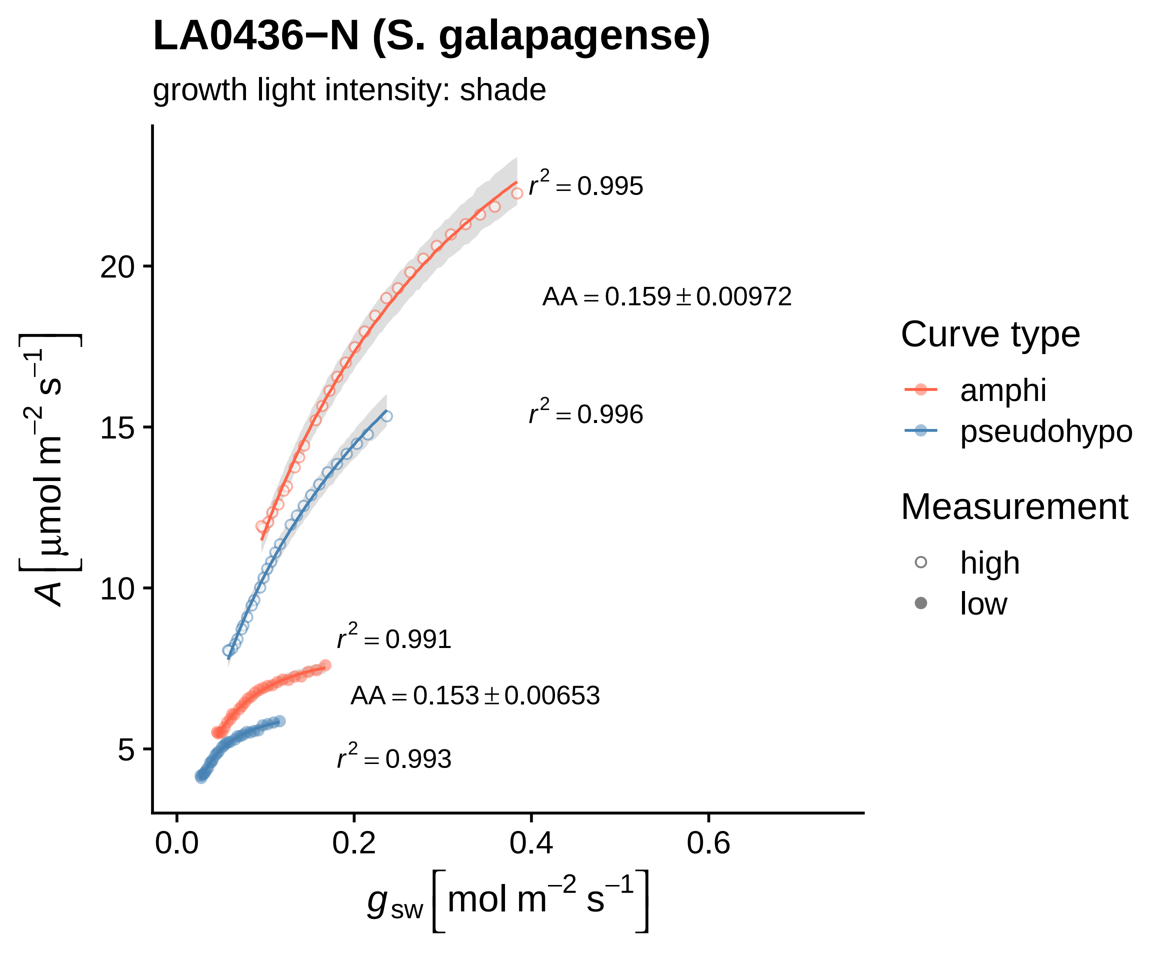

LA0436−N (S. galapagense)
growth light intensity: shade
$r^2 = 0.995$
$AA = 0.159 \pm 0.00972$
$r^2 = 0.996$
$r^2 = 0.991$
$AA = 0.153 \pm 0.00653$
$r^2 = 0.993$
20
15
10
5
0.0
0.2
0.4
0.6
$A\left[\mu mol\ m^{-2}\ s^{-1}\right]$
$g_{sw}\left[mol\ m^{-2}\ s^{-1}\right]$
Curve type
amphi
pseudohypo
Measurement
high
low

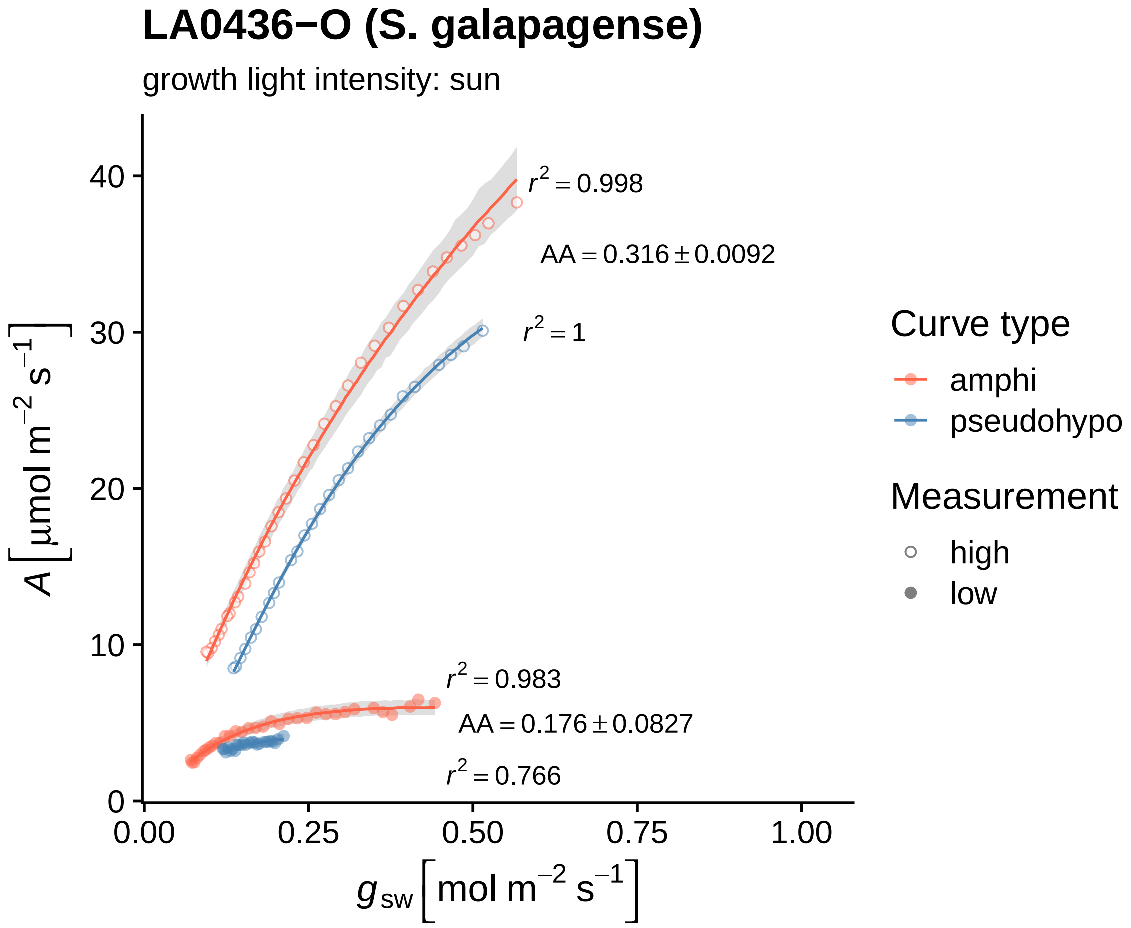
LA0436−O (S. galapagense)
growth light intensity: sun
$r^2 = 0.998$
$AA = 0.316 \pm 0.0092$
$r^2 = 1$
$r^2 = 0.983$
$AA = 0.176 \pm 0.0827$
$r^2 = 0.766$
0
10
20
30
40
0.00
0.25
0.50
0.75
1.00
$A\ [\mu mol\ m^{-2}\ s^{-1}]$
$g_{sw}\ [mol\ m^{-2}\ s^{-1}]$
Curve type
amphi
pseudohypo
Measurement
high
low

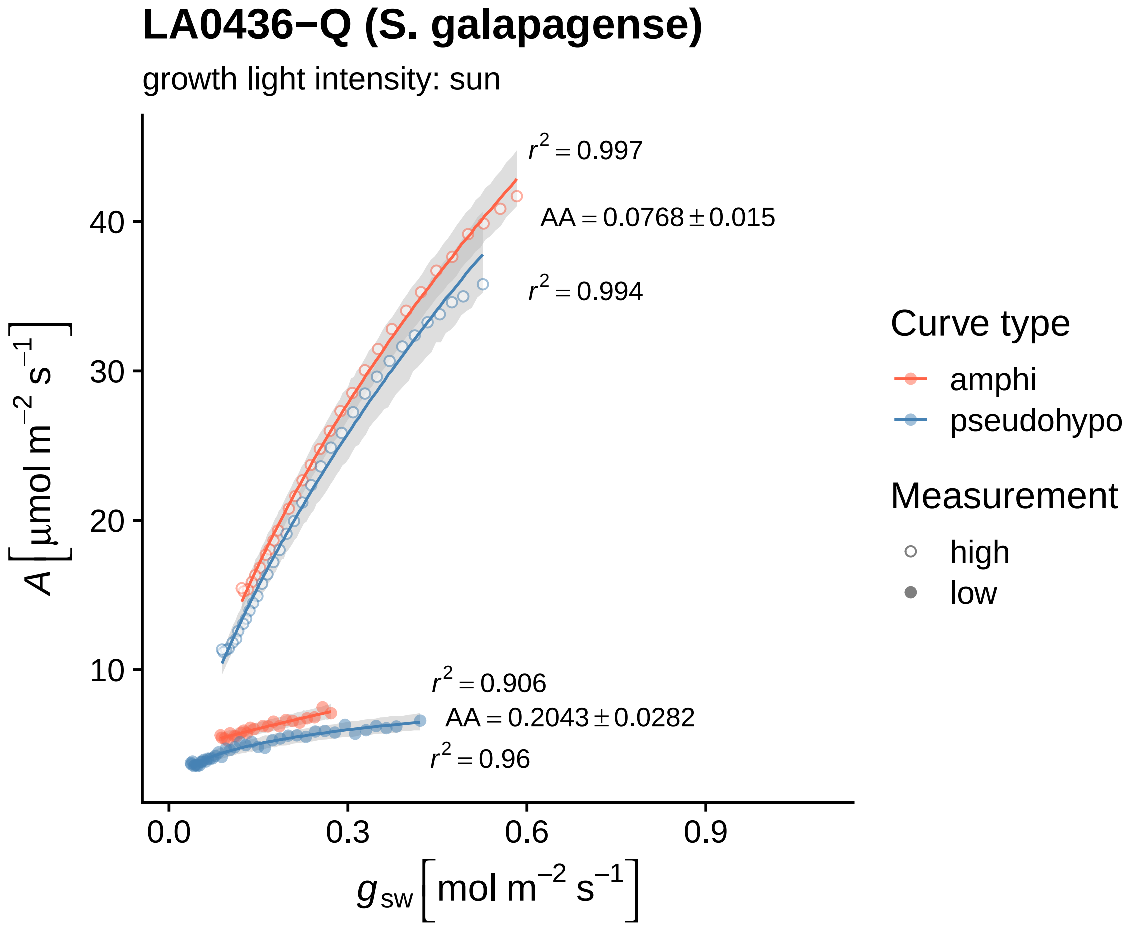
LA0436−Q (S. galapagense)
growth light intensity: sun
r² = 0.997
AA = 0.0768 ± 0.015
r² = 0.994
r² = 0.906
AA = 0.2043 ± 0.0282
r² = 0.96
A [μmol m⁻² s⁻¹]
g_sw [mol m⁻² s⁻¹]
10
20
30
40
0.0
0.3
0.6
0.9
Curve type
amphi
pseudohypo
Measurement
high
low

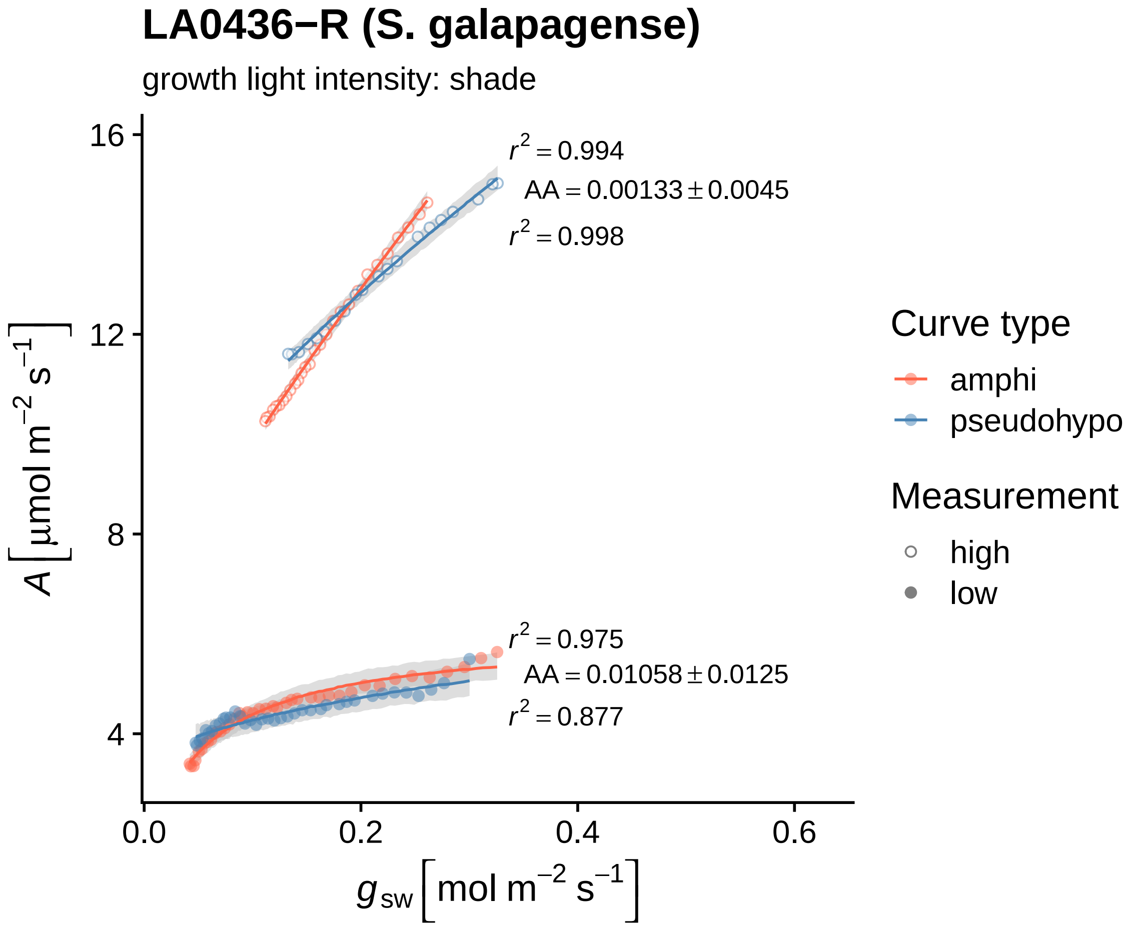
LA0436−R (S. galapagense)
growth light intensity: shade
$r^2 = 0.994$
$AA = 0.00133 \pm 0.0045$
$r^2 = 0.998$
$r^2 = 0.975$
$AA = 0.01058 \pm 0.0125$
$r^2 = 0.877$
16
12
8
4
0.0
0.2
0.4
0.6
$A\left[\mu\text{mol m}^{-2}\text{ s}^{-1}\right]$
$g_{sw}\left[\text{mol m}^{-2}\text{ s}^{-1}\right]$
Curve type
amphi
pseudohypo
Measurement
high
low

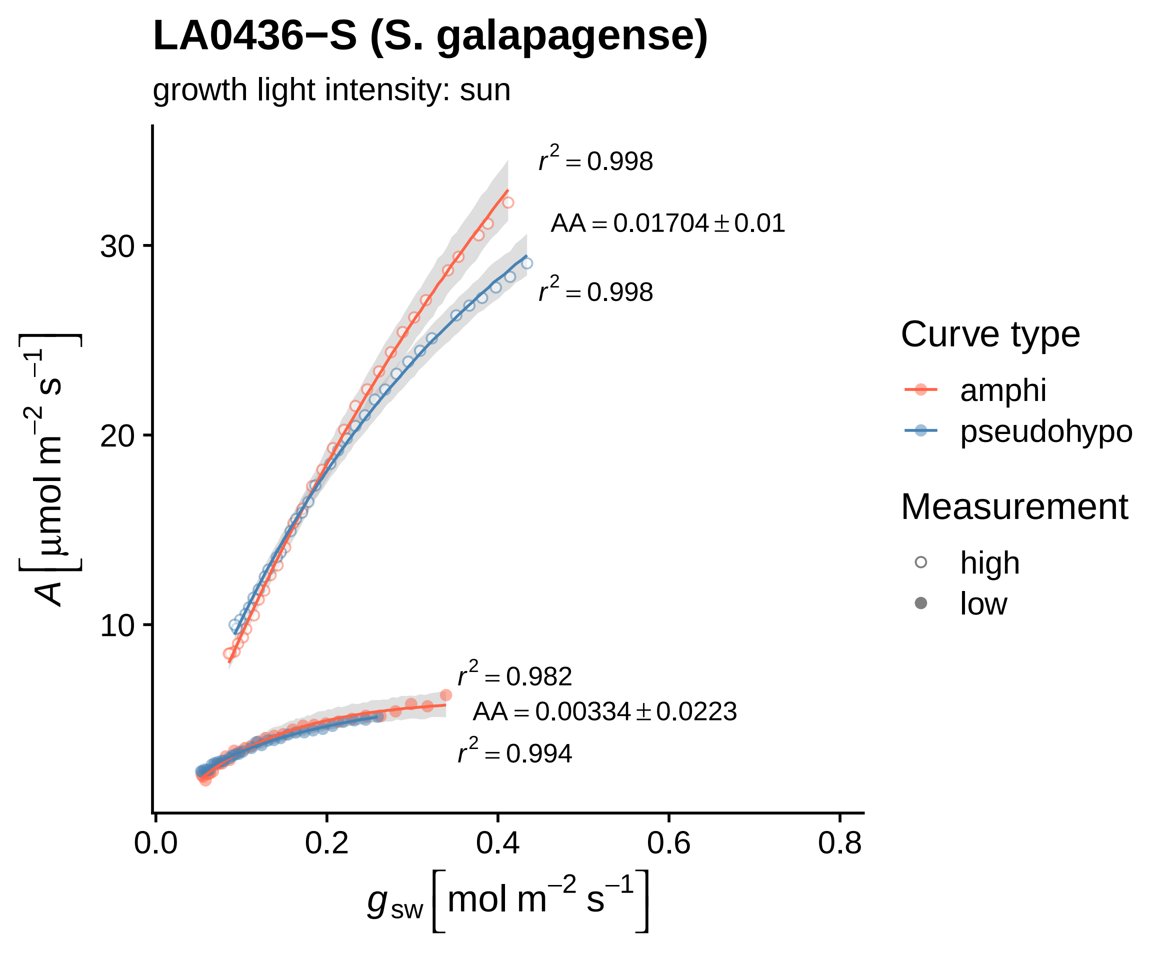

LA0436−S (S. galapagense)
growth light intensity: sun
$r^2 = 0.998$
$AA = 0.01704 \pm 0.01$
$r^2 = 0.998$
$r^2 = 0.982$
$AA = 0.00334 \pm 0.0223$
$r^2 = 0.994$
30
20
10
0.0
0.2
0.4
0.6
0.8
$A\left[\mu mol\ m^{-2}\ s^{-1}\right]$
$g_{sw}\left[mol\ m^{-2}\ s^{-1}\right]$
Curve type
amphi
pseudohypo
Measurement
high
low

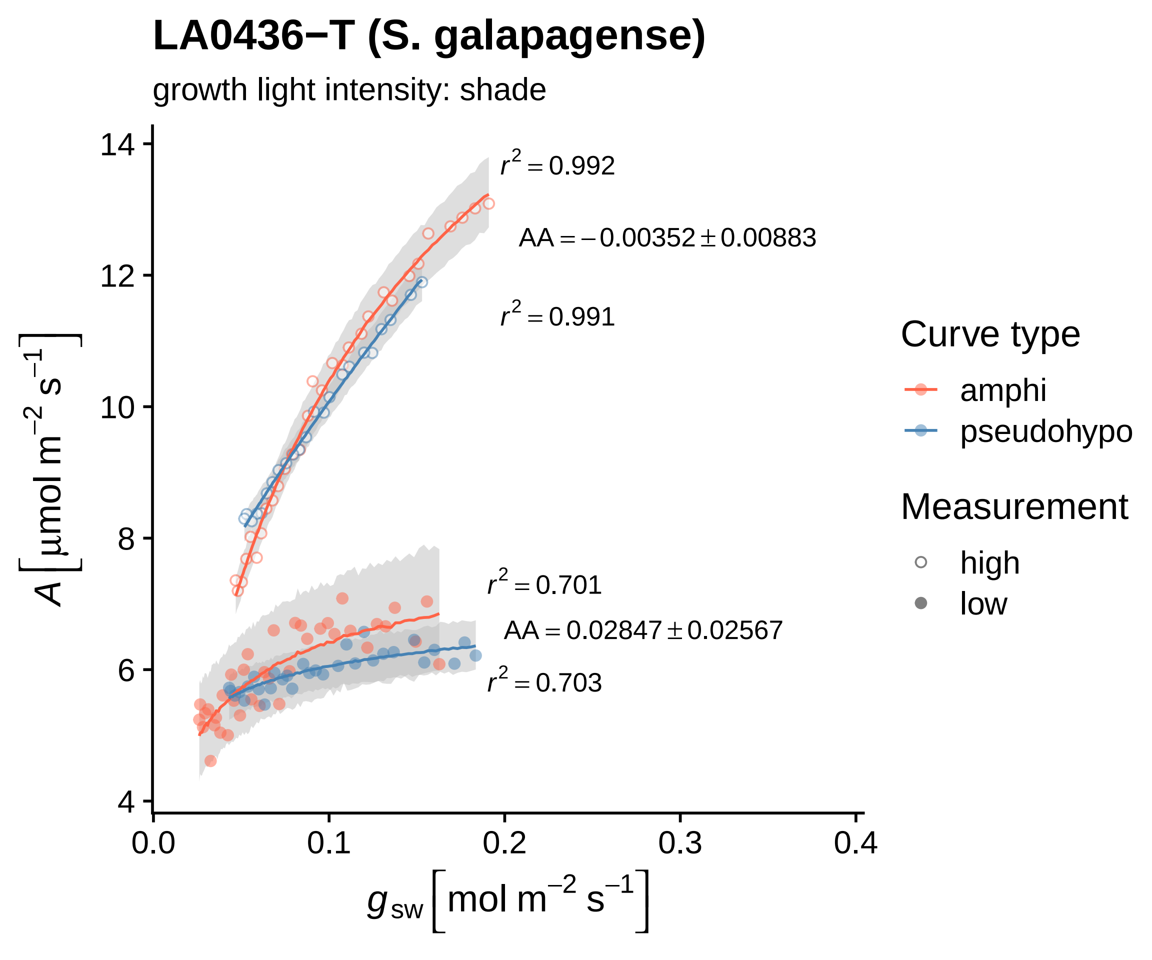

LA0436−T (S. galapagense)
growth light intensity: shade
$r^2 = 0.992$
$AA = -0.00352 \pm 0.00883$
$r^2 = 0.991$
$r^2 = 0.701$
$AA = 0.02847 \pm 0.02567$
$r^2 = 0.703$
$A\left[\mu mol\ m^{-2}\ s^{-1}\right]$
$g_{sw}\left[mol\ m^{-2}\ s^{-1}\right]$
14
12
10
8
6
4
0.0
0.1
0.2
0.3
0.4
Curve type
amphi
pseudohypo
Measurement
high
low

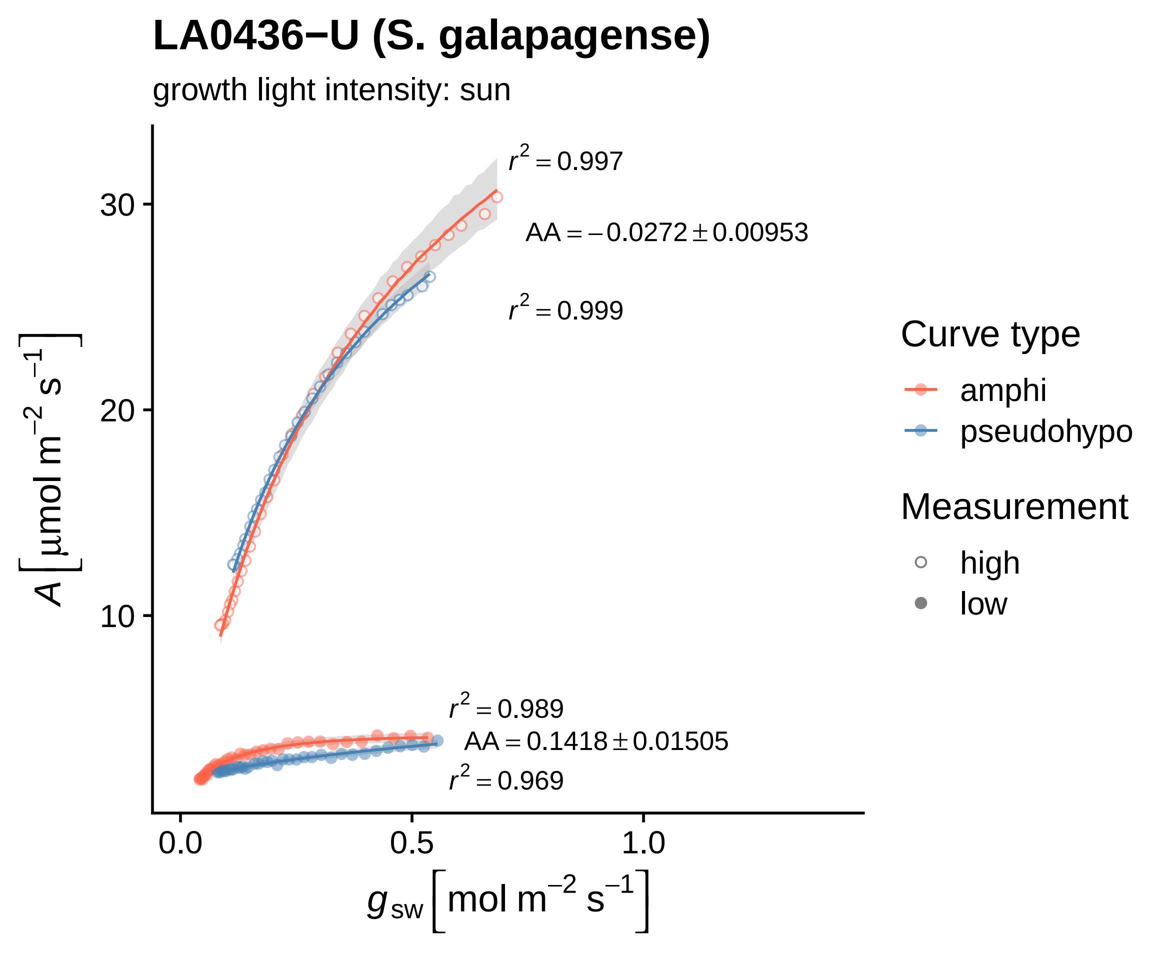
LA0436−U (S. galapagense)
growth light intensity: sun
$r^2 = 0.997$
$AA = -0.0272 \pm 0.00953$
$r^2 = 0.999$
$r^2 = 0.989$
$AA = 0.1418 \pm 0.01505$
$r^2 = 0.969$
30
20
10
0.0
0.5
1.0
$A\left[\mu mol\ m^{-2}\ s^{-1}\right]$
$g_{sw}\left[mol\ m^{-2}\ s^{-1}\right]$
Curve type
amphi
pseudohypo
Measurement
high
low

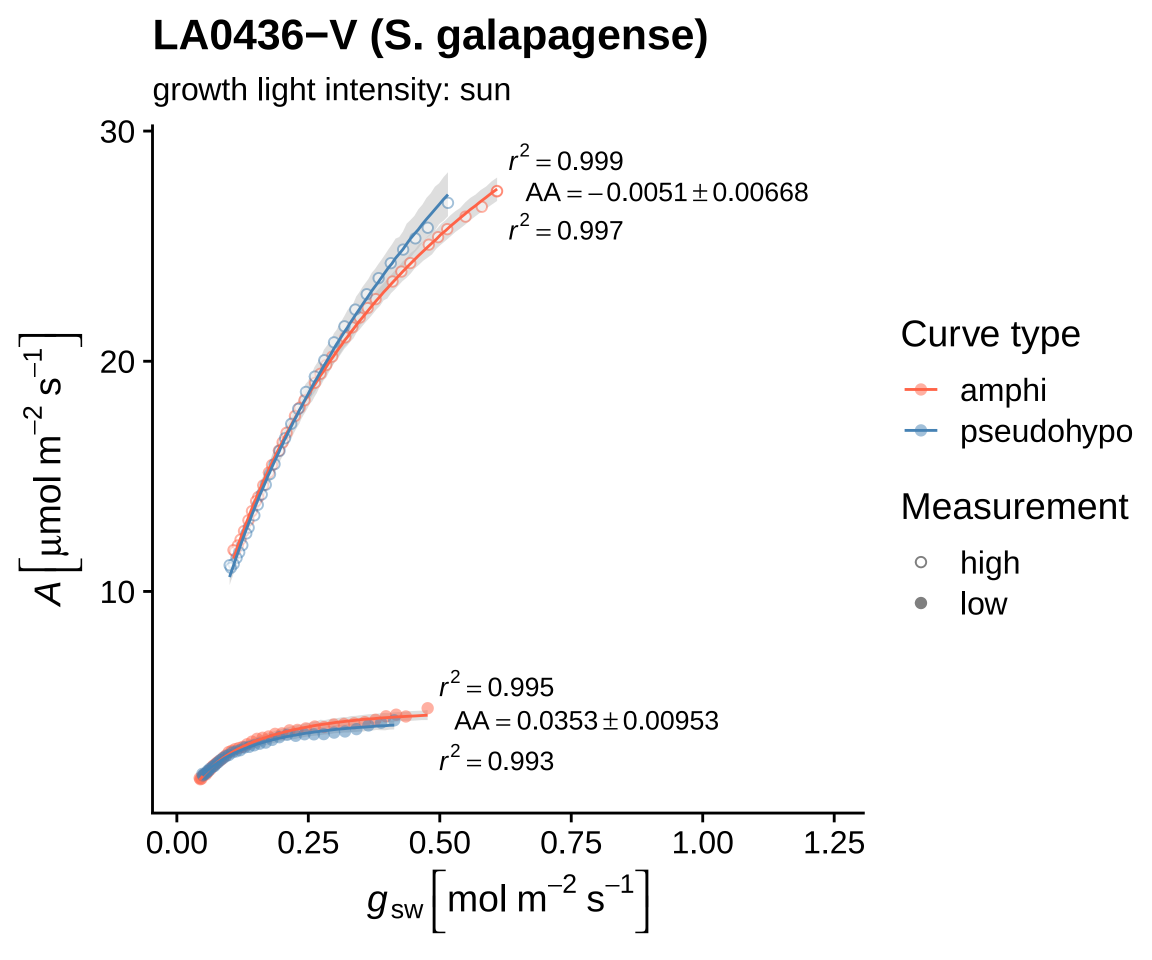

LA0436−V (S. galapagense)
growth light intensity: sun
$r^2 = 0.999$
$AA = -0.0051 \pm 0.00668$
$r^2 = 0.997$
$r^2 = 0.995$
$AA = 0.0353 \pm 0.00953$
$r^2 = 0.993$
30
20
10
0.00
0.25
0.50
0.75
1.00
1.25
$A\ [\mu mol\ m^{-2}\ s^{-1}]$
$g_{sw}\ [mol\ m^{-2}\ s^{-1}]$
Curve type
amphi
pseudohypo
Measurement
high
low

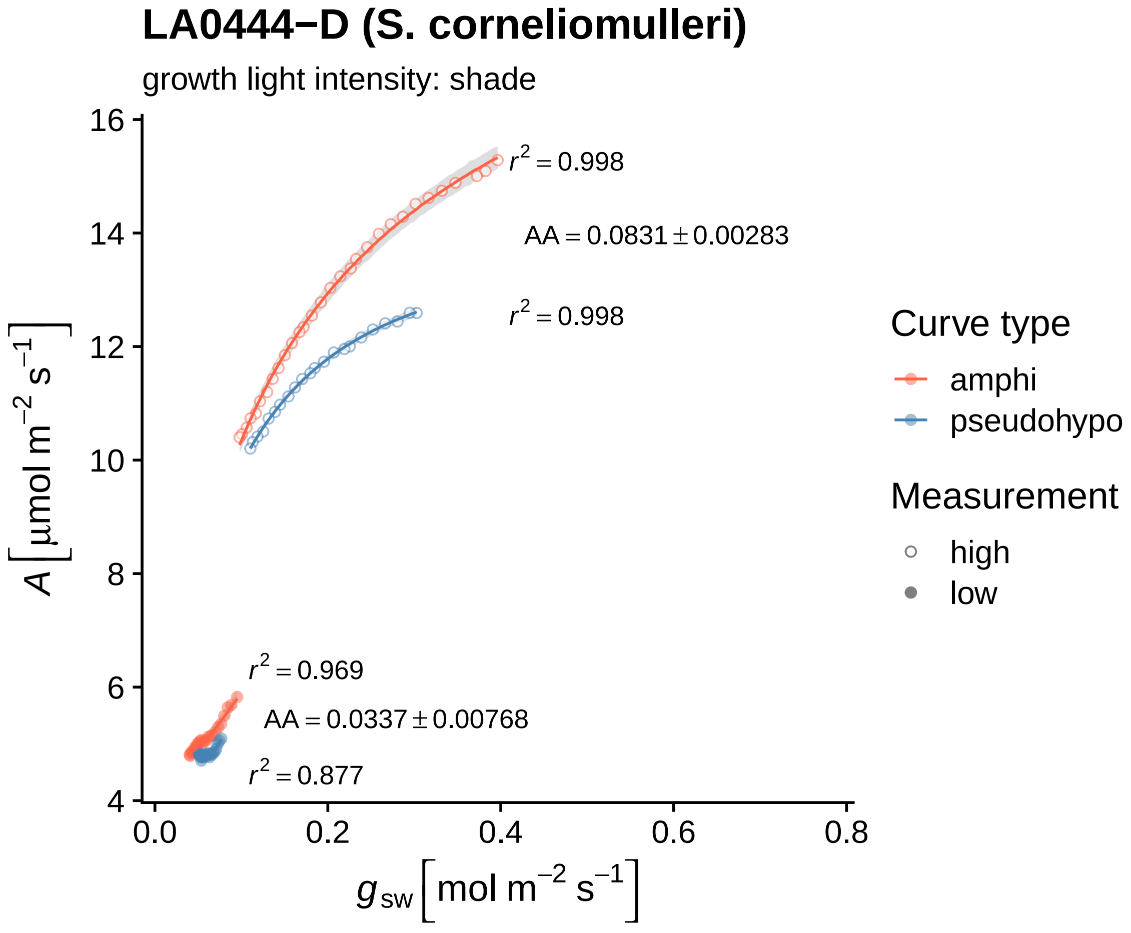
LA0444−D (S. corneliomulleri)
growth light intensity: shade
$r^2 = 0.998$
$AA = 0.0831 \pm 0.00283$
$r^2 = 0.998$
$r^2 = 0.969$
$AA = 0.0337 \pm 0.00768$
$r^2 = 0.877$
$A\ [\mu mol\ m^{-2}\ s^{-1}]$
$g_{sw}\ [mol\ m^{-2}\ s^{-1}]$
16
14
12
10
8
6
4
0.0
0.2
0.4
0.6
0.8
Curve type
amphi
pseudohypo
Measurement
high
low

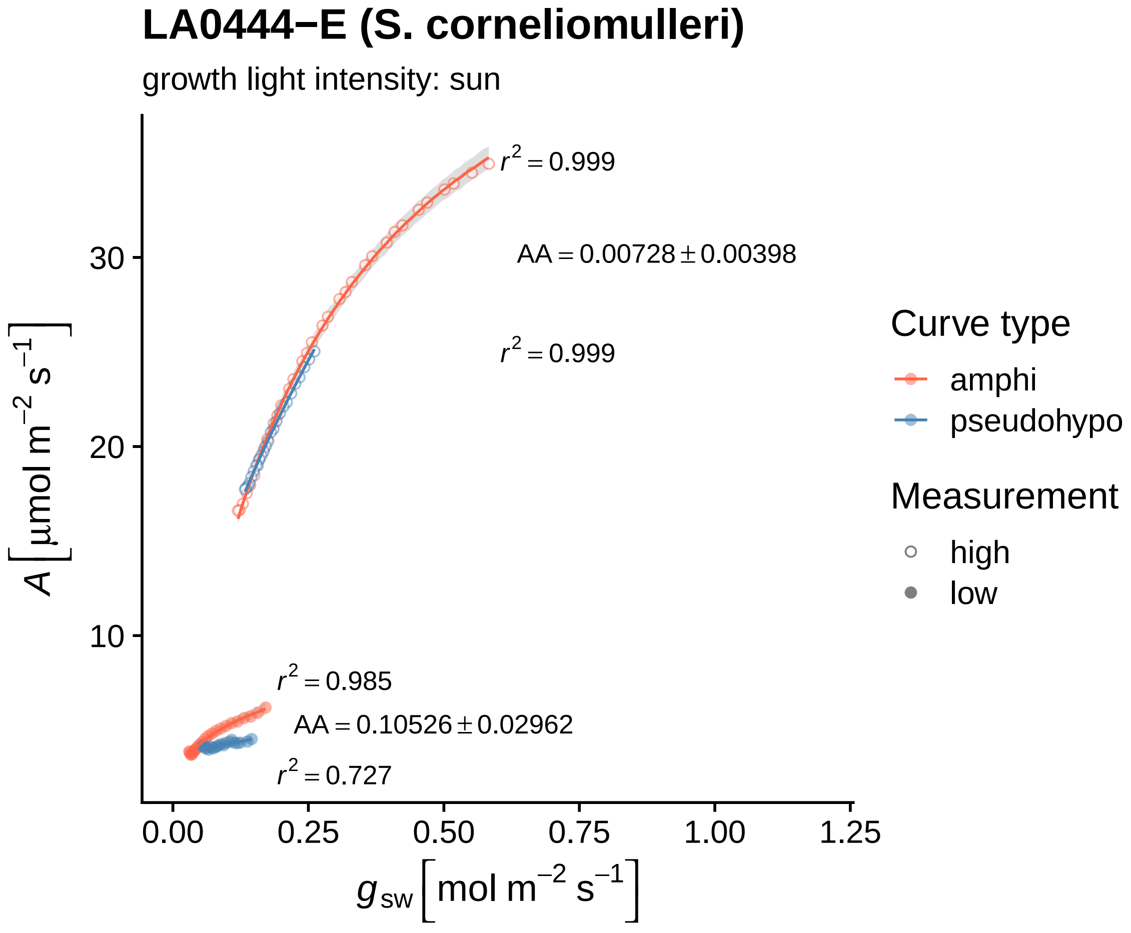

LA0444−E (S. corneliomulleri)
growth light intensity: sun
$r^2 = 0.999$
$AA = 0.00728 \pm 0.00398$
$r^2 = 0.999$
$r^2 = 0.985$
$AA = 0.10526 \pm 0.02962$
$r^2 = 0.727$
30
20
10
0.00
0.25
0.50
0.75
1.00
1.25
$A\ [\mu mol\ m^{-2}\ s^{-1}]$
$g_{sw}\ [mol\ m^{-2}\ s^{-1}]$
Curve type
amphi
pseudohypo
Measurement
high
low

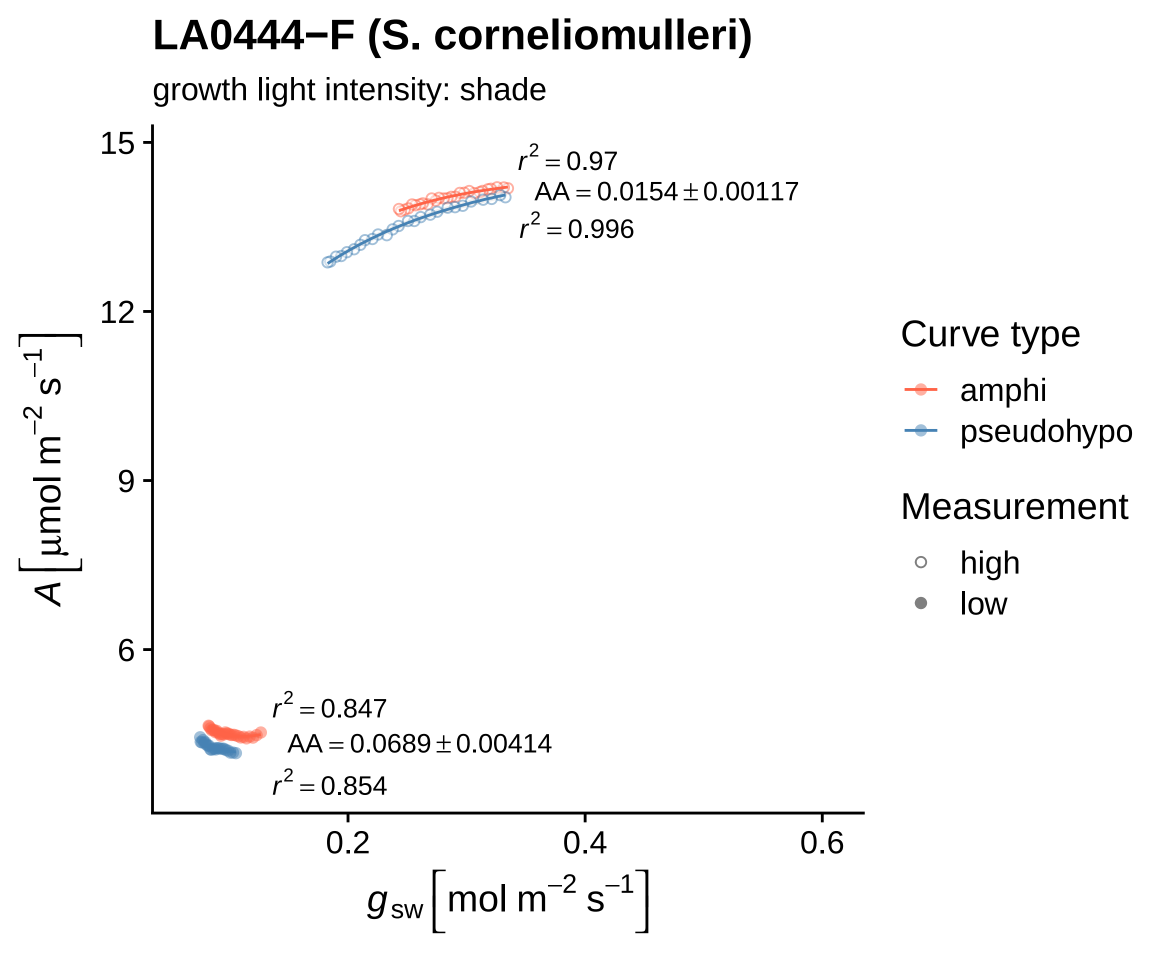

LA0444−F (S. corneliomulleri)
growth light intensity: shade
15
12
9
6
$A\left[\mu mol\ m^{-2}\ s^{-1}\right]$
0.2
0.4
0.6
$g_{sw}\left[mol\ m^{-2}\ s^{-1}\right]$
$r^2 = 0.97$
$AA = 0.0154 \pm 0.00117$
$r^2 = 0.996$
$r^2 = 0.847$
$AA = 0.0689 \pm 0.00414$
$r^2 = 0.854$
Curve type
amphi
pseudohypo
Measurement
high
low

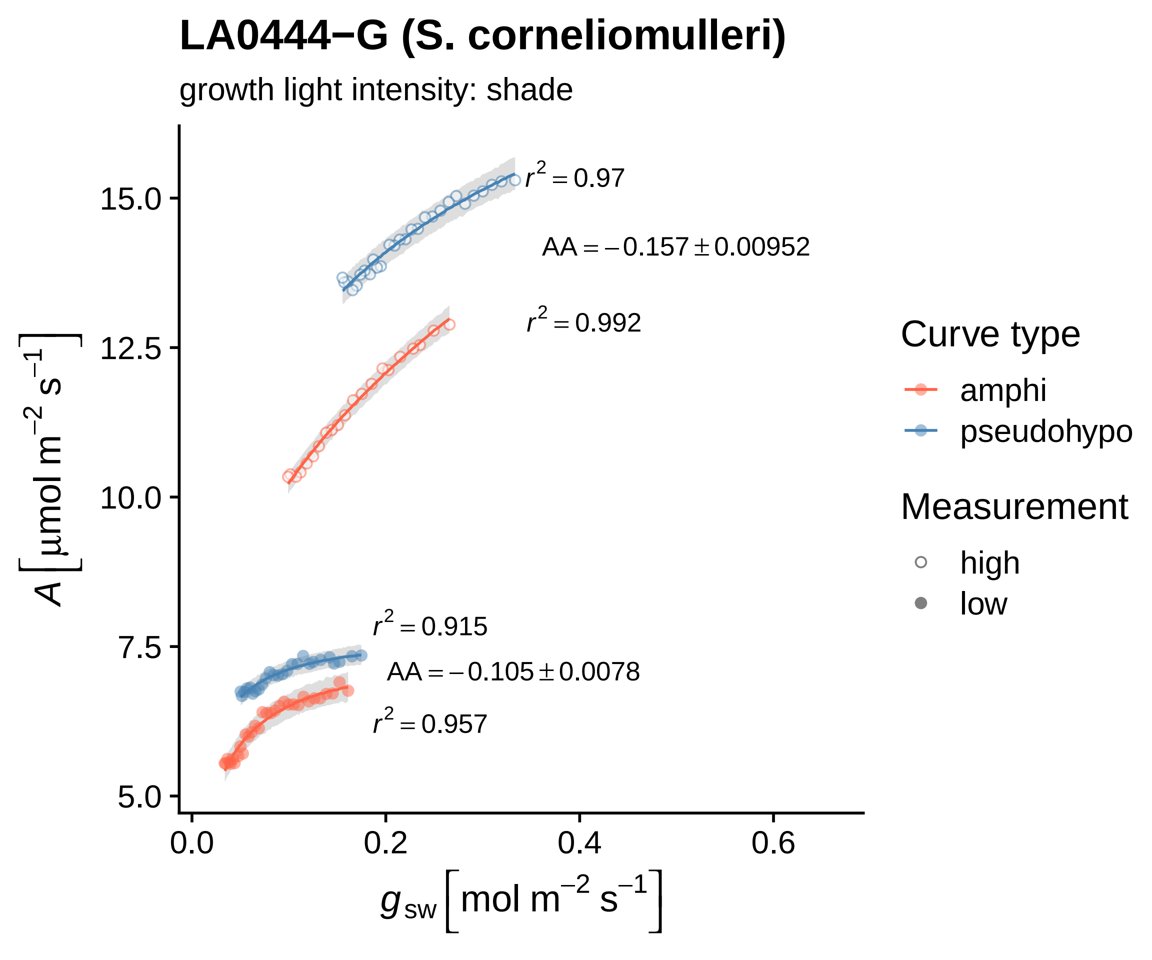

LA0444−G (S. corneliomulleri)
growth light intensity: shade
r² = 0.97
AA = −0.157 ± 0.00952
r² = 0.992
r² = 0.915
AA = −0.105 ± 0.0078
r² = 0.957
15.0
12.5
10.0
7.5
5.0
0.0
0.2
0.4
0.6
A [μmol m⁻² s⁻¹]
g_sw [mol m⁻² s⁻¹]
Curve type
amphi
pseudohypo
Measurement
high
low

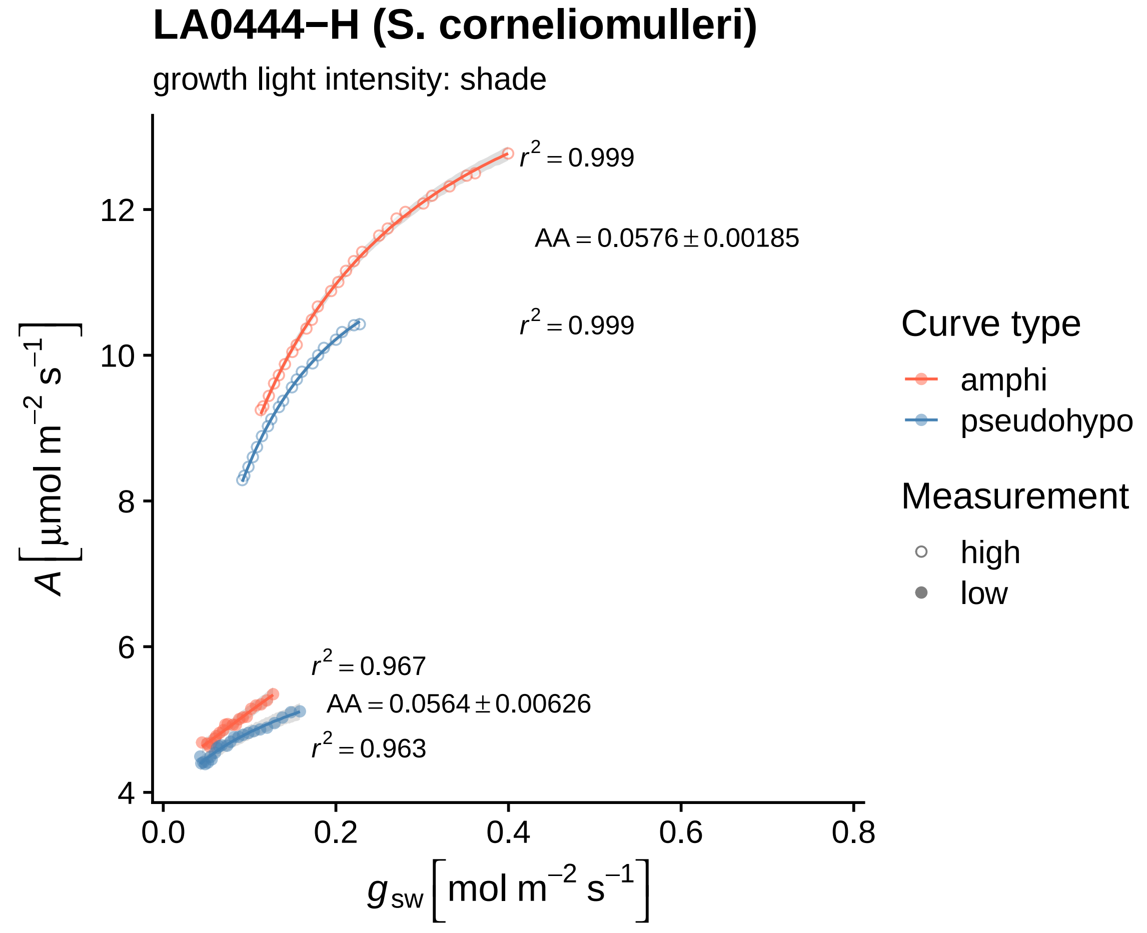

LA0444−H (S. corneliomulleri)
growth light intensity: shade
$r^2 = 0.999$
AA = 0.0576 ± 0.00185
$r^2 = 0.999$
$r^2 = 0.967$
AA = 0.0564 ± 0.00626
$r^2 = 0.963$
A [µmol m⁻² s⁻¹]
12
10
8
6
4
0.0
0.2
0.4
0.6
0.8
$g_{sw}$ [mol m⁻² s⁻¹]
Curve type
amphi
pseudohypo
Measurement
high
low

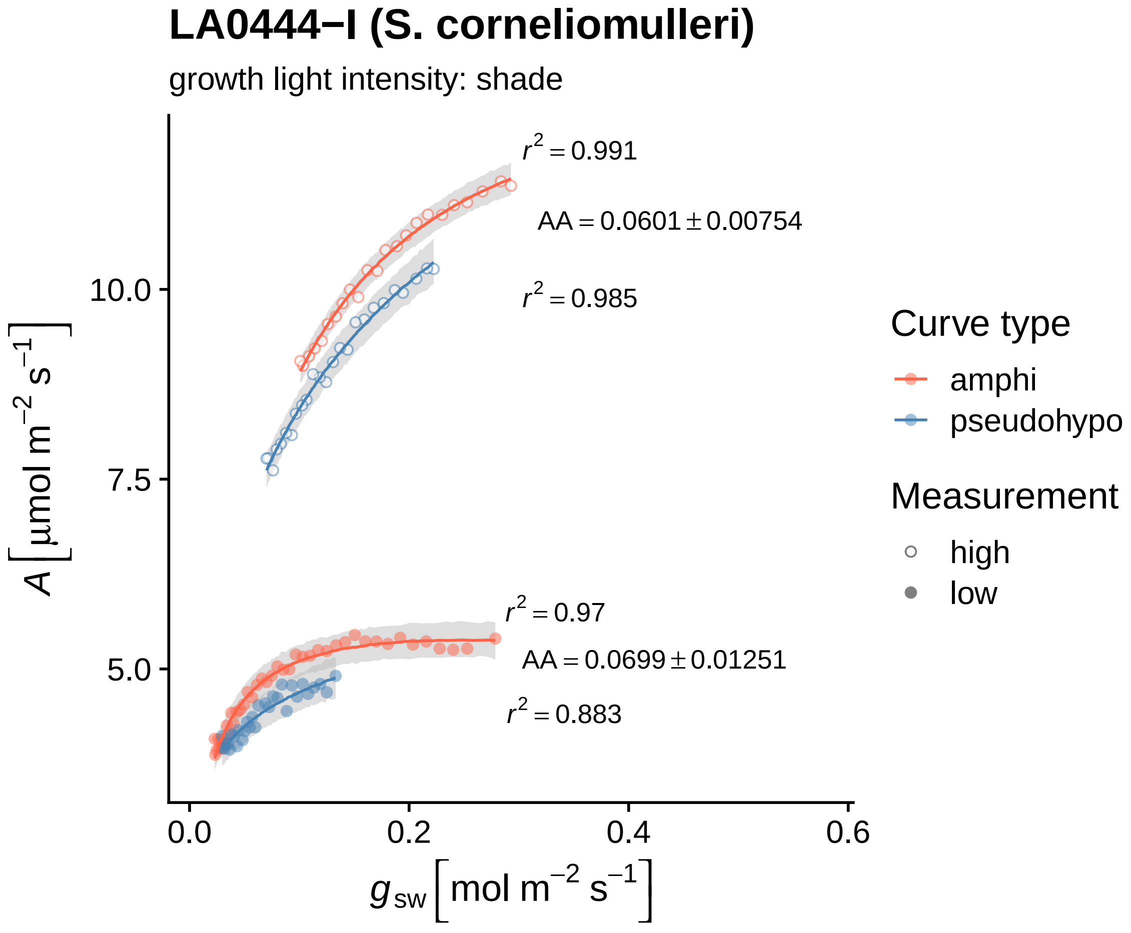
LA0444−I (S. corneliomulleri)
growth light intensity: shade
$r^2 = 0.991$
$AA = 0.0601 \pm 0.00754$
$r^2 = 0.985$
$r^2 = 0.97$
$AA = 0.0699 \pm 0.01251$
$r^2 = 0.883$
10.0
7.5
5.0
0.0
0.2
0.4
0.6
$A\ [\mu mol\ m^{-2}\ s^{-1}]$
$g_{sw}\ [mol\ m^{-2}\ s^{-1}]$
Curve type
amphi
pseudohypo
Measurement
high
low

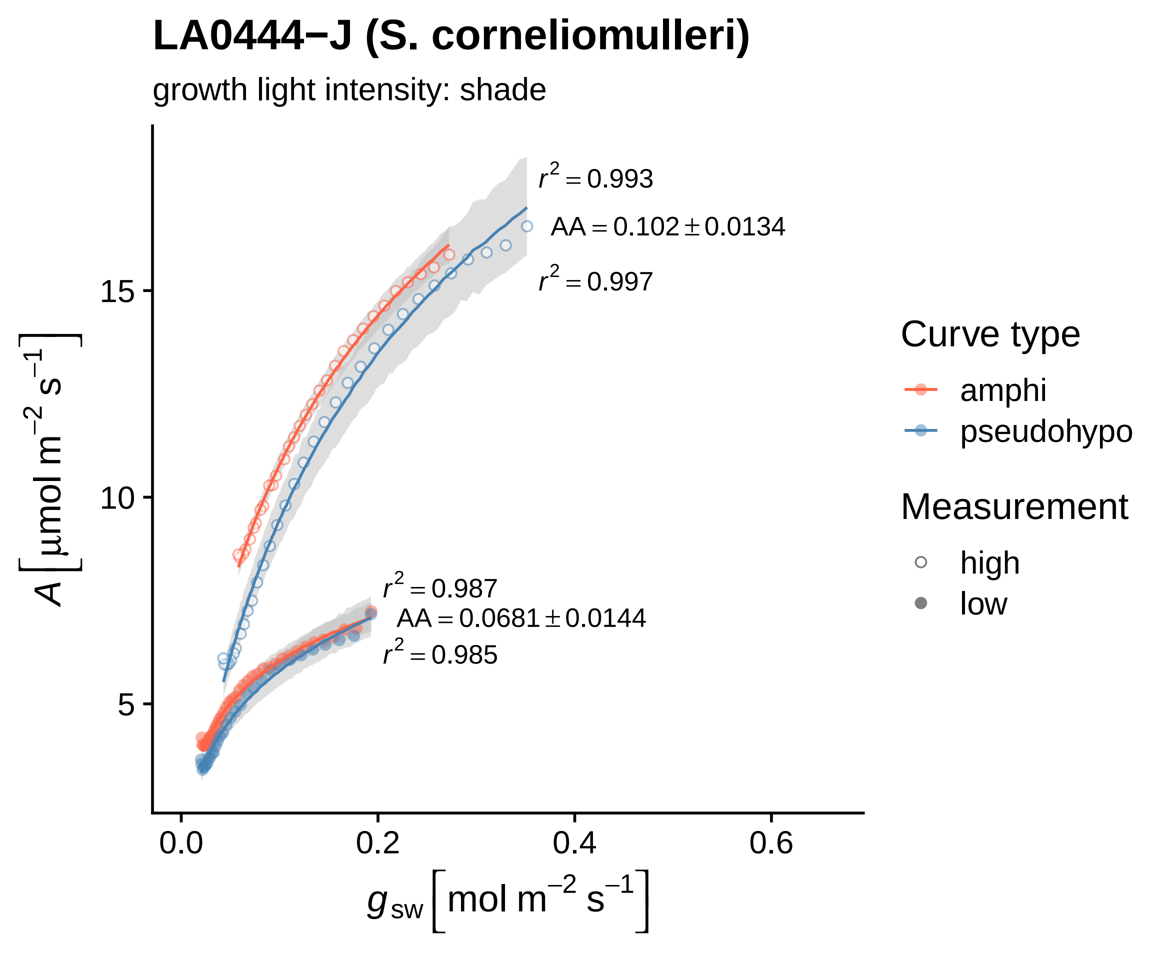
LA0444−J (S. corneliomulleri)
growth light intensity: shade
$r^2 = 0.993$
AA = 0.102 ± 0.0134
$r^2 = 0.997$
$r^2 = 0.987$
AA = 0.0681 ± 0.0144
$r^2 = 0.985$
15
10
5
0.0
0.2
0.4
0.6
$A\ [\mu mol\ m^{-2}\ s^{-1}]$
$g_{sw}\ [mol\ m^{-2}\ s^{-1}]$
Curve type
amphi
pseudohypo
Measurement
high
low

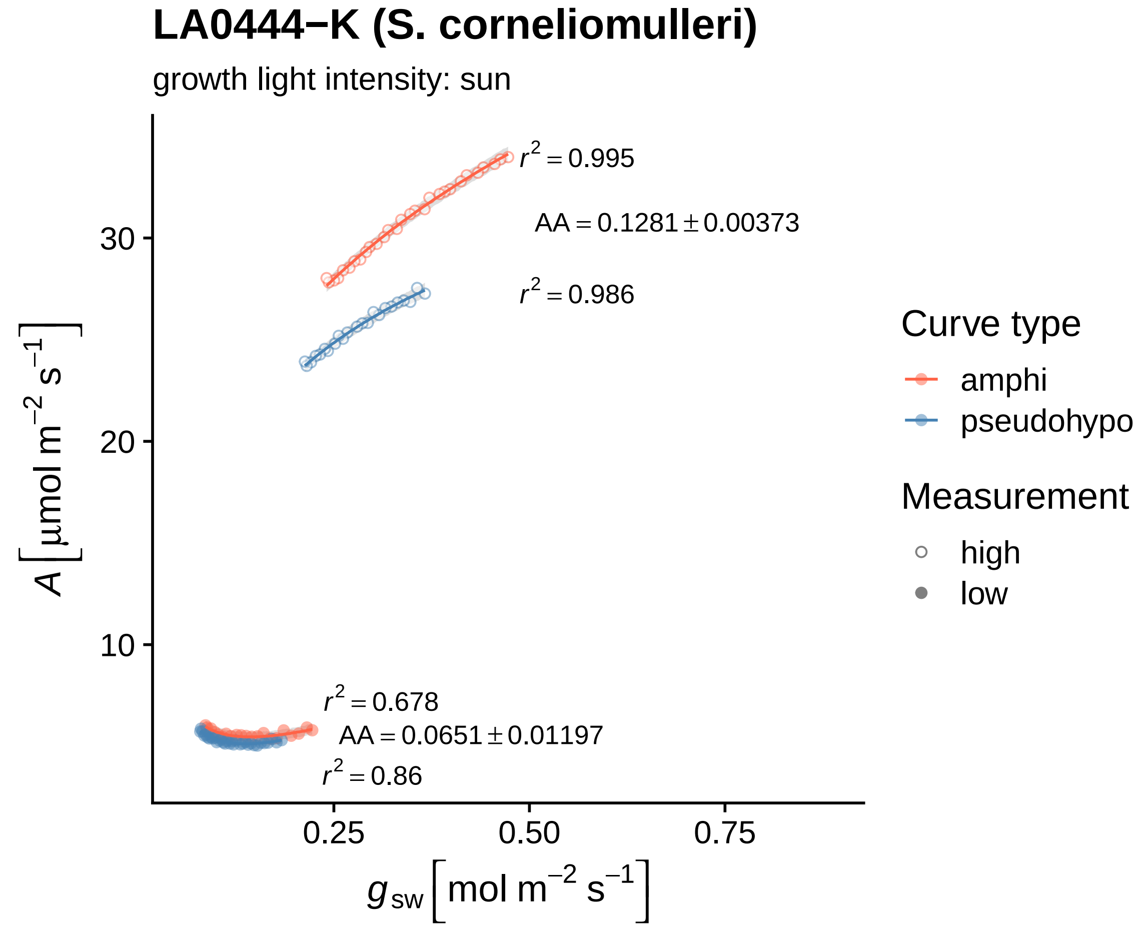

LA0444−K (S. corneliomulleri)
growth light intensity: sun
r^2 = 0.995
AA = 0.1281 ± 0.00373
r^2 = 0.986
r^2 = 0.678
AA = 0.0651 ± 0.01197
r^2 = 0.86
30
20
10
0.25
0.50
0.75
A [µmol m^−2 s^−1]
g_sw [mol m^−2 s^−1]
Curve type
amphi
pseudohypo
Measurement
high
low

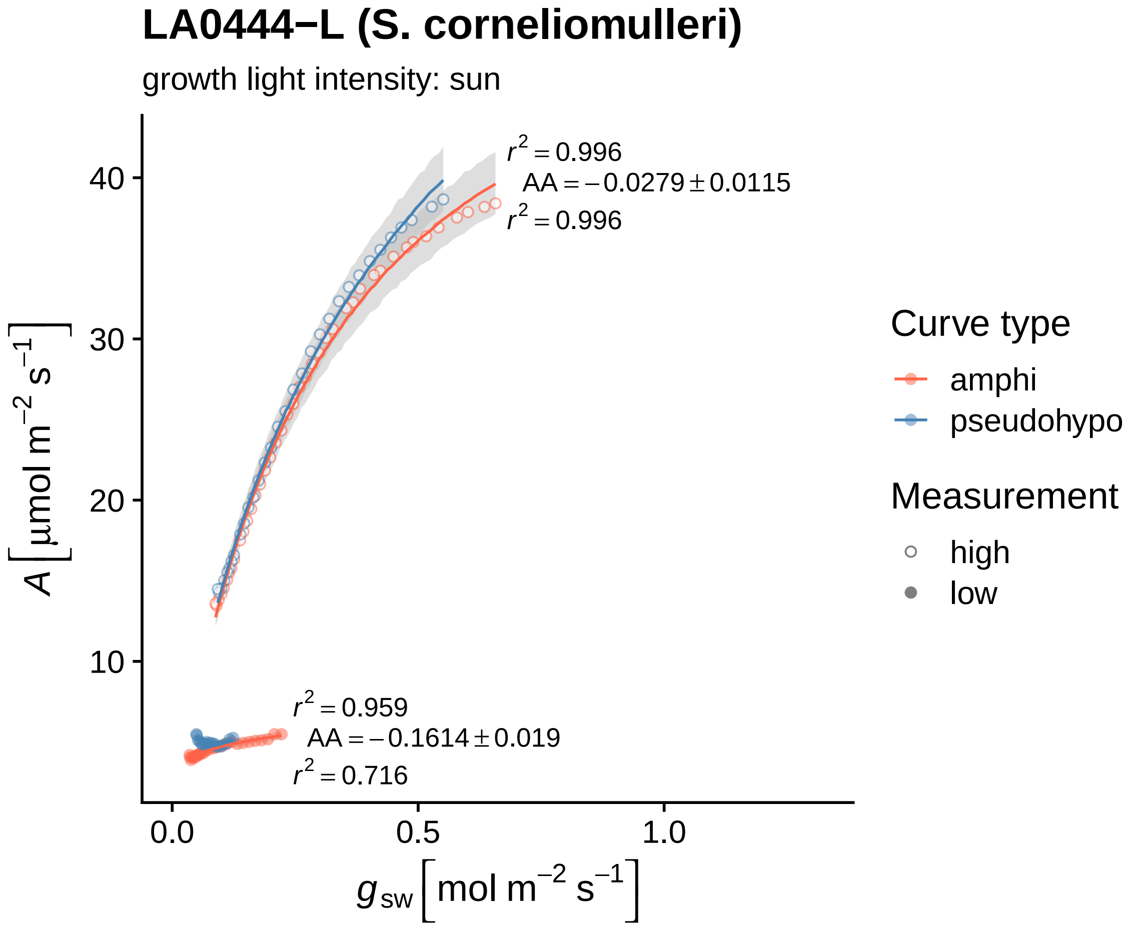
LA0444−L (S. corneliomulleri)
growth light intensity: sun
$r^2 = 0.996$
$AA = -0.0279 \pm 0.0115$
$r^2 = 0.996$
$r^2 = 0.959$
$AA = -0.1614 \pm 0.019$
$r^2 = 0.716$
40
30
20
10
0.0
0.5
1.0
$A\ [\mu mol\ m^{-2}\ s^{-1}]$
$g_{sw}\ [mol\ m^{-2}\ s^{-1}]$
Curve type
amphi
pseudohypo
Measurement
high
low

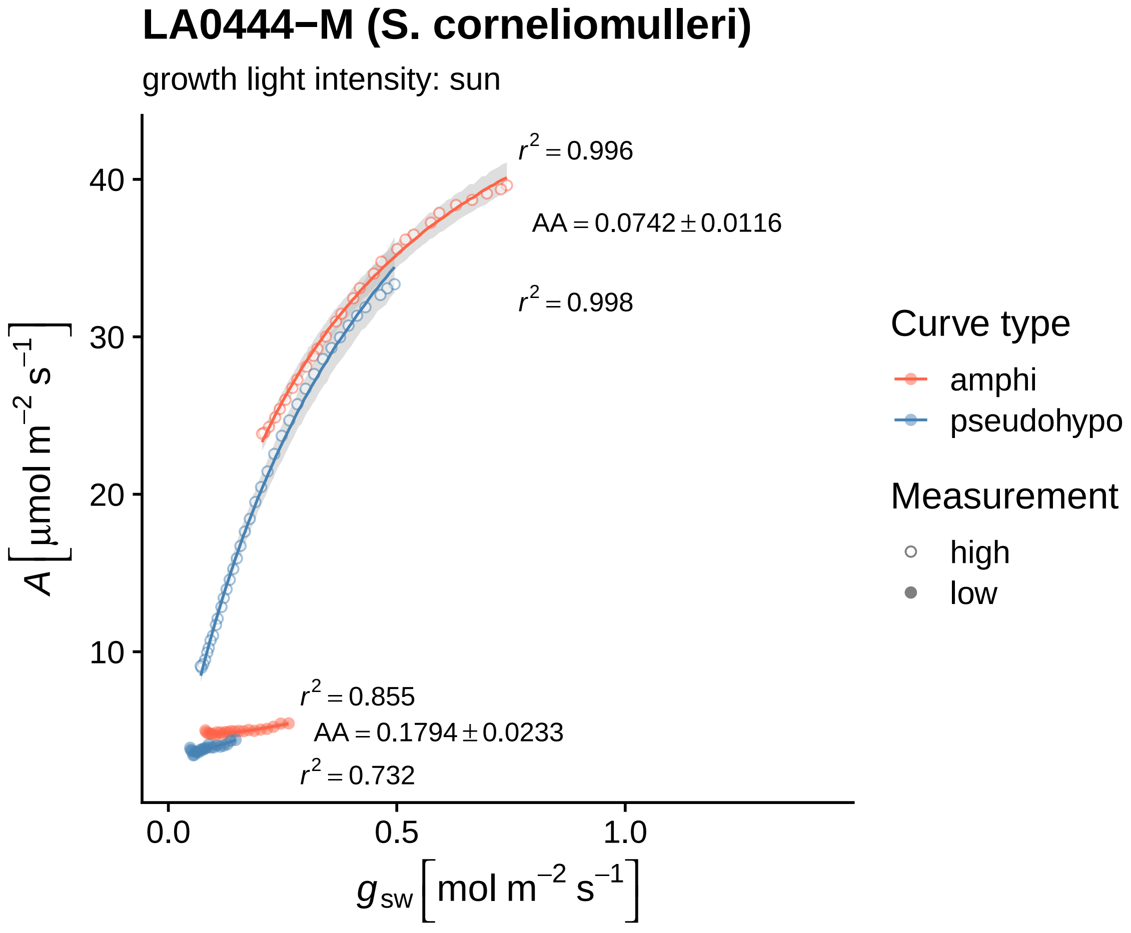
LA0444−M (S. corneliomulleri)
growth light intensity: sun
$r^2 = 0.996$
$AA = 0.0742 \pm 0.0116$
$r^2 = 0.998$
$r^2 = 0.855$
$AA = 0.1794 \pm 0.0233$
$r^2 = 0.732$
40
30
20
10
0.0
0.5
1.0
$A\ [\mu mol\ m^{-2}\ s^{-1}]$
$g_{sw}\ [mol\ m^{-2}\ s^{-1}]$
Curve type
amphi
pseudohypo
Measurement
high
low

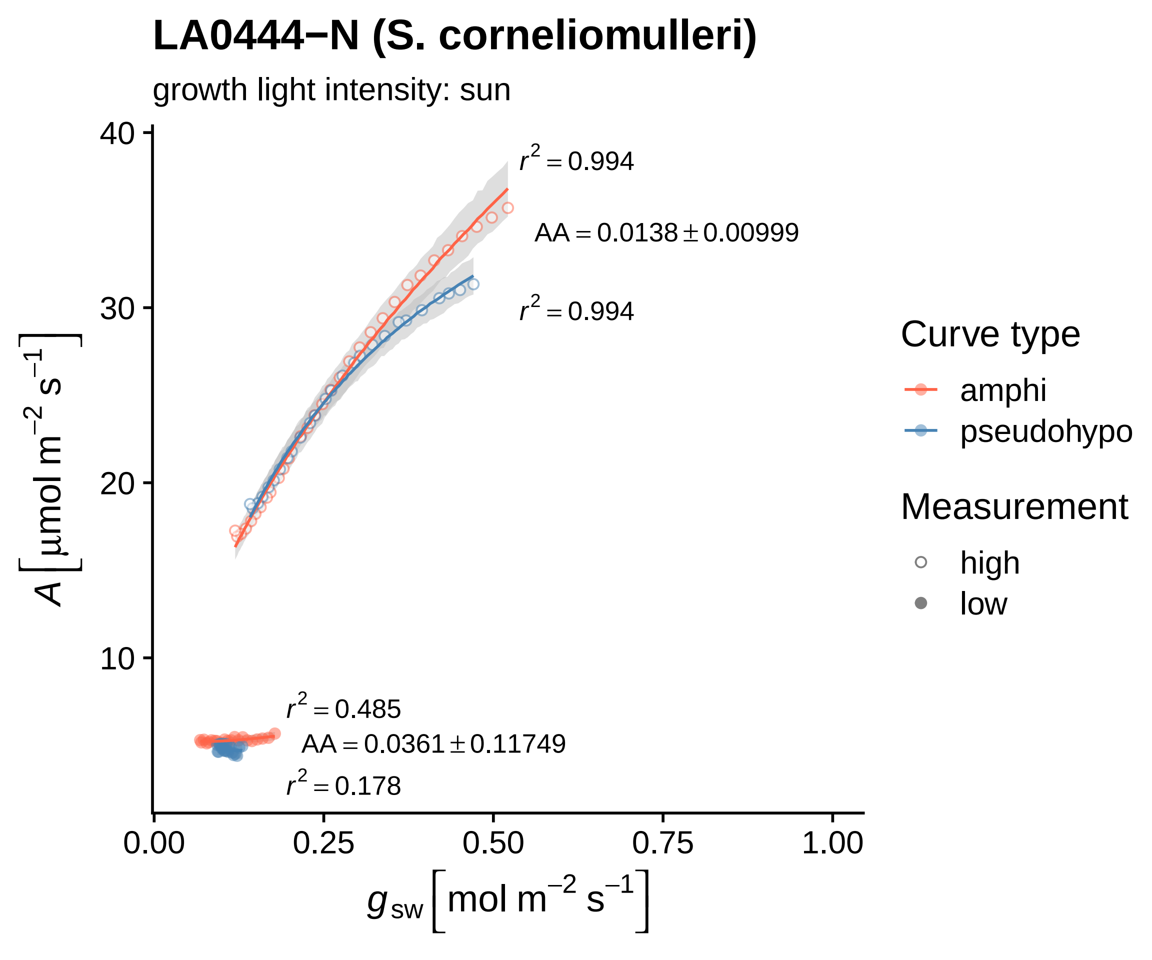

LA0444−N (S. corneliomulleri)
growth light intensity: sun
$r^2 = 0.994$
AA = 0.0138 ± 0.00999
$r^2 = 0.994$
$r^2 = 0.485$
AA = 0.0361 ± 0.11749
$r^2 = 0.178$
40
30
20
10
0.00
0.25
0.50
0.75
1.00
$A\ [\mu mol\ m^{-2}\ s^{-1}]$
$g_{sw}\ [mol\ m^{-2}\ s^{-1}]$
Curve type
amphi
pseudohypo
Measurement
high
low

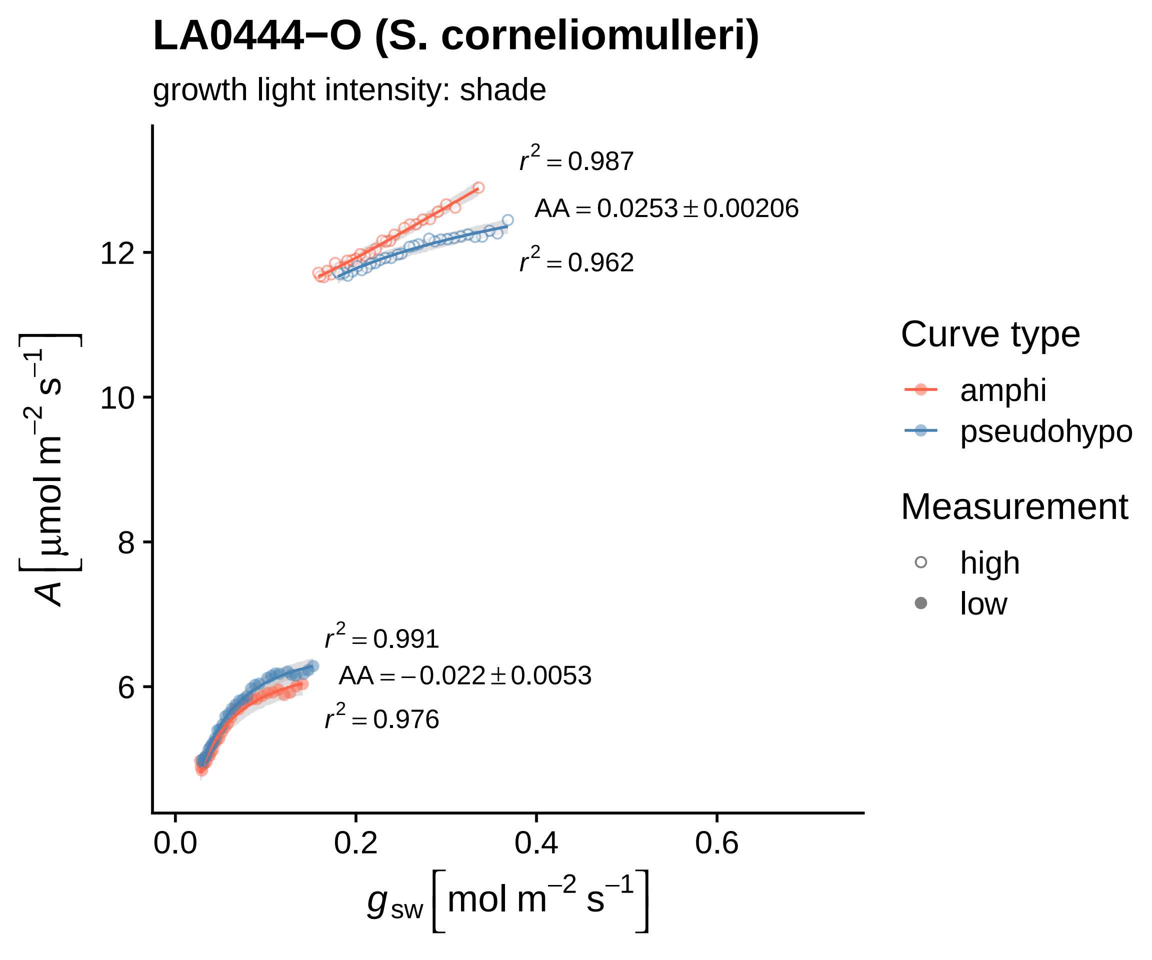
LA0444−O (S. corneliomulleri)
growth light intensity: shade
$r^2 = 0.987$
$AA = 0.0253 \pm 0.00206$
$r^2 = 0.962$
$r^2 = 0.991$
$AA = -0.022 \pm 0.0053$
$r^2 = 0.976$
$A\left[\mu mol\ m^{-2}\ s^{-1}\right]$
$g_{sw}\left[mol\ m^{-2}\ s^{-1}\right]$
12
10
8
6
0.0
0.2
0.4
0.6
Curve type
amphi
pseudohypo
Measurement
high
low

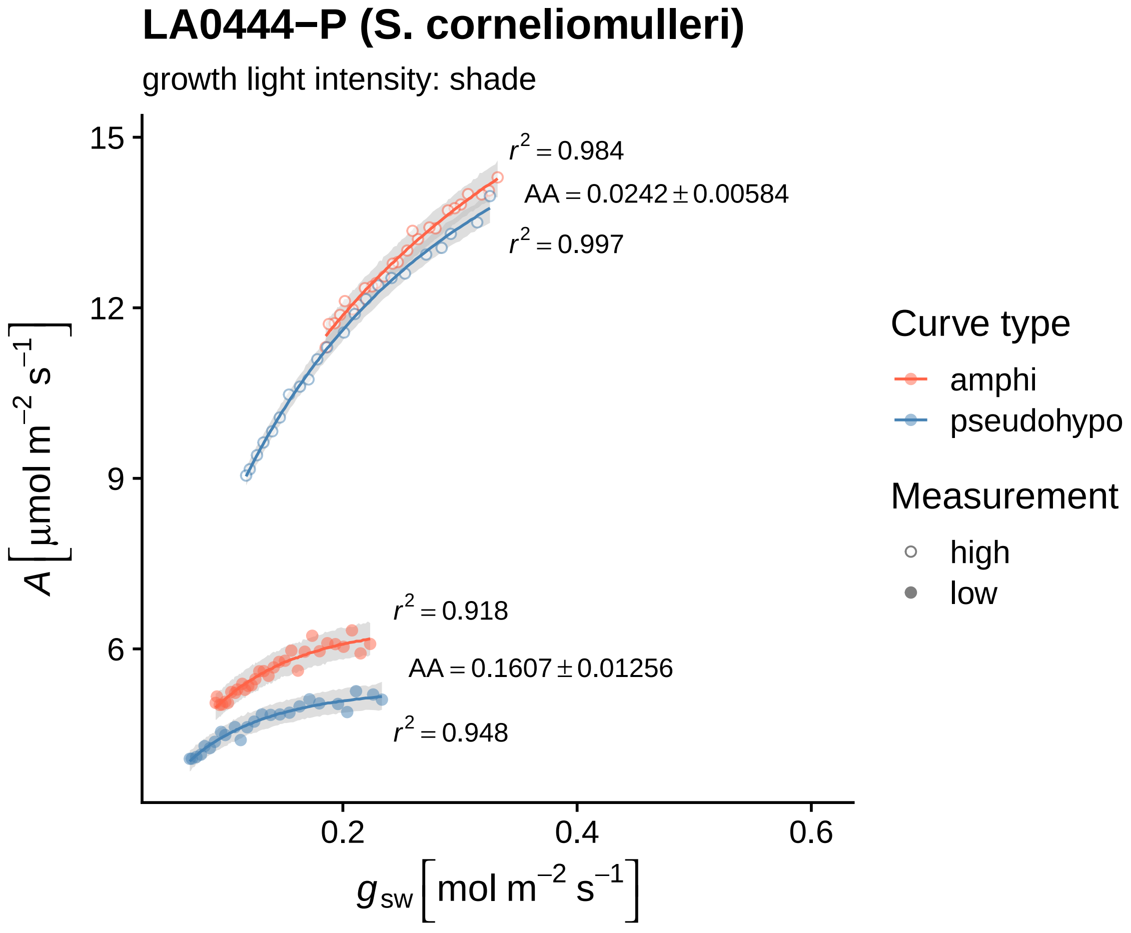

LA0444−P (S. corneliomulleri)
growth light intensity: shade
$r^2 = 0.984$
$AA = 0.0242 \pm 0.00584$
$r^2 = 0.997$
$r^2 = 0.918$
$AA = 0.1607 \pm 0.01256$
$r^2 = 0.948$
15
12
9
6
0.2
0.4
0.6
$A\ [\mu mol\ m^{-2}\ s^{-1}]$
$g_{sw}\ [mol\ m^{-2}\ s^{-1}]$
Curve type
amphi
pseudohypo
Measurement
high
low

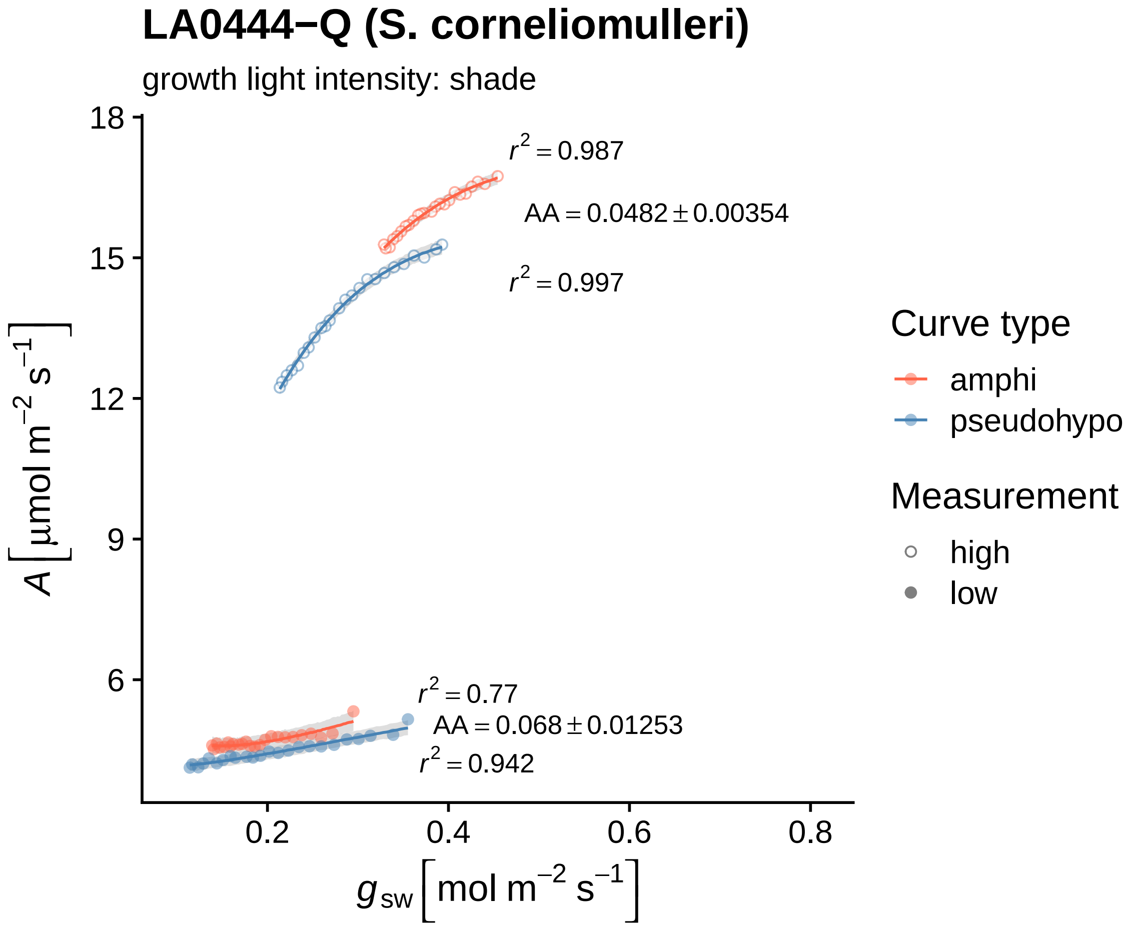

LA0444−Q (S. corneliomulleri)
growth light intensity: shade
18
15
12
9
6
0.2
0.4
0.6
0.8
$r^2 = 0.987$
$AA = 0.0482 \pm 0.00354$
$r^2 = 0.997$
$r^2 = 0.77$
$AA = 0.068 \pm 0.01253$
$r^2 = 0.942$
$A\ [\mu mol\ m^{-2}\ s^{-1}]$
$g_{sw}\ [mol\ m^{-2}\ s^{-1}]$
Curve type
amphi
pseudohypo
Measurement
high
low

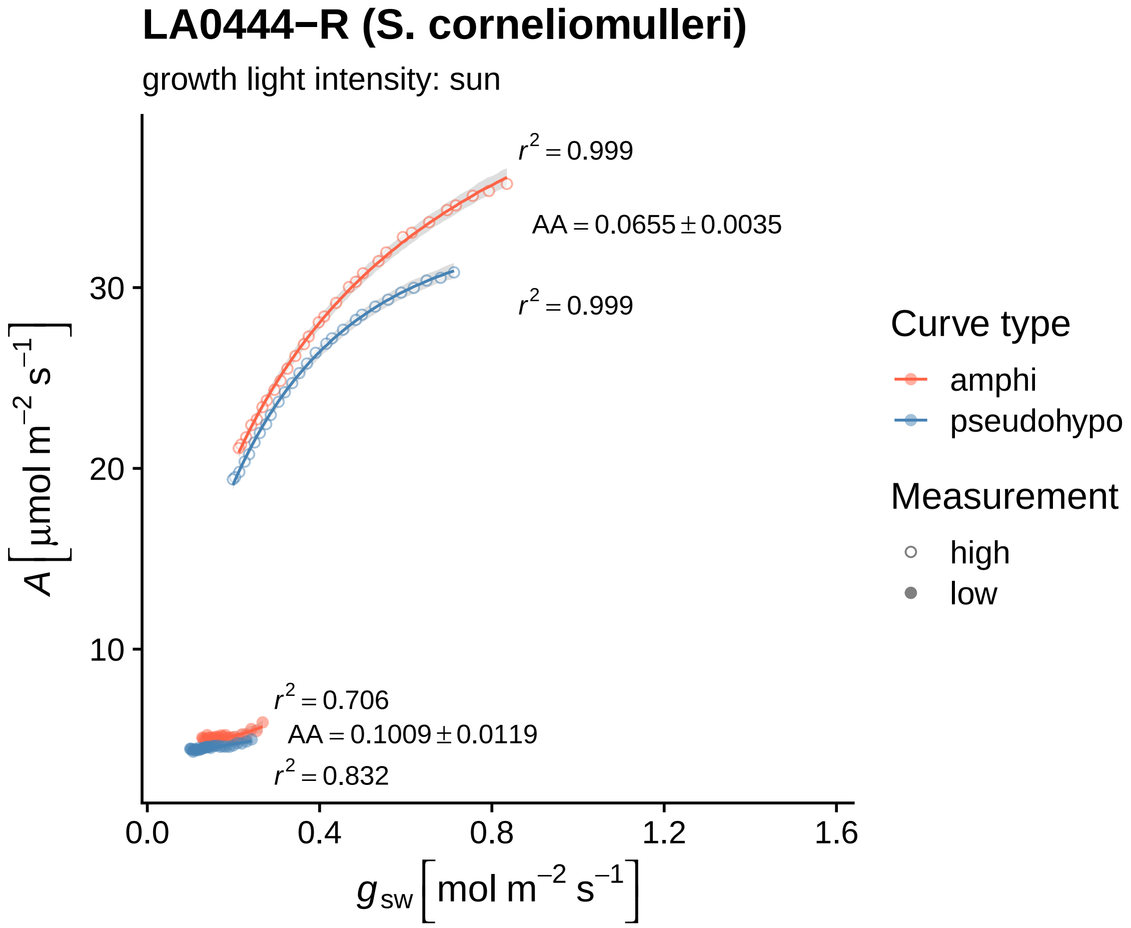

LA0444−R (S. corneliomulleri)
growth light intensity: sun
$r^2 = 0.999$
AA = 0.0655 ± 0.0035
$r^2 = 0.999$
$r^2 = 0.706$
AA = 0.1009 ± 0.0119
$r^2 = 0.832$
30
20
10
0.0
0.4
0.8
1.2
1.6
$A\ [\mu mol\ m^{-2}\ s^{-1}]$
$g_{sw}\ [mol\ m^{-2}\ s^{-1}]$
Curve type
amphi
pseudohypo
Measurement
high
low

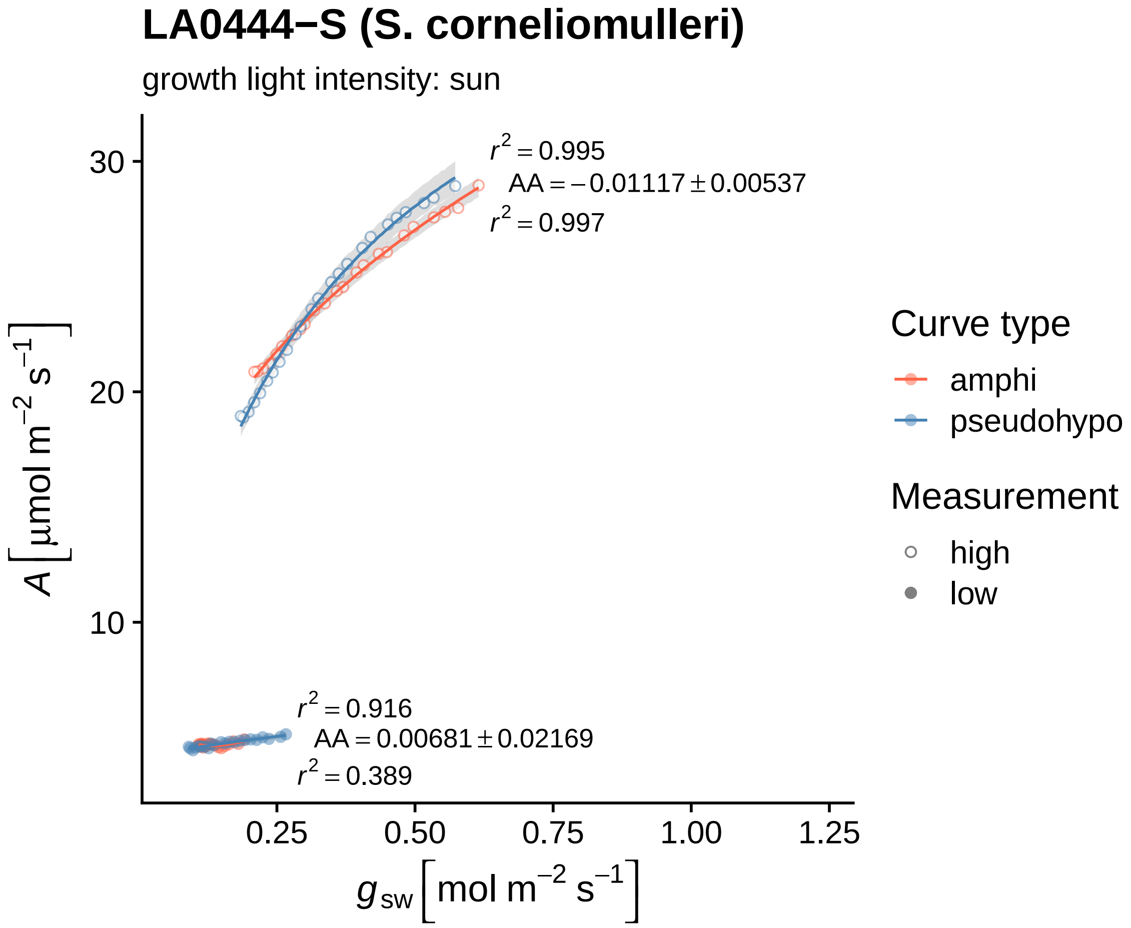

LA0444−S (S. corneliomulleri)
growth light intensity: sun
$r^2 = 0.995$
$AA = -0.01117 \pm 0.00537$
$r^2 = 0.997$
$r^2 = 0.916$
$AA = 0.00681 \pm 0.02169$
$r^2 = 0.389$
30
20
10
0.25
0.50
0.75
1.00
1.25
$A\ [\mu mol\ m^{-2}\ s^{-1}]$
$g_{sw}\ [mol\ m^{-2}\ s^{-1}]$
Curve type
amphi
pseudohypo
Measurement
high
low

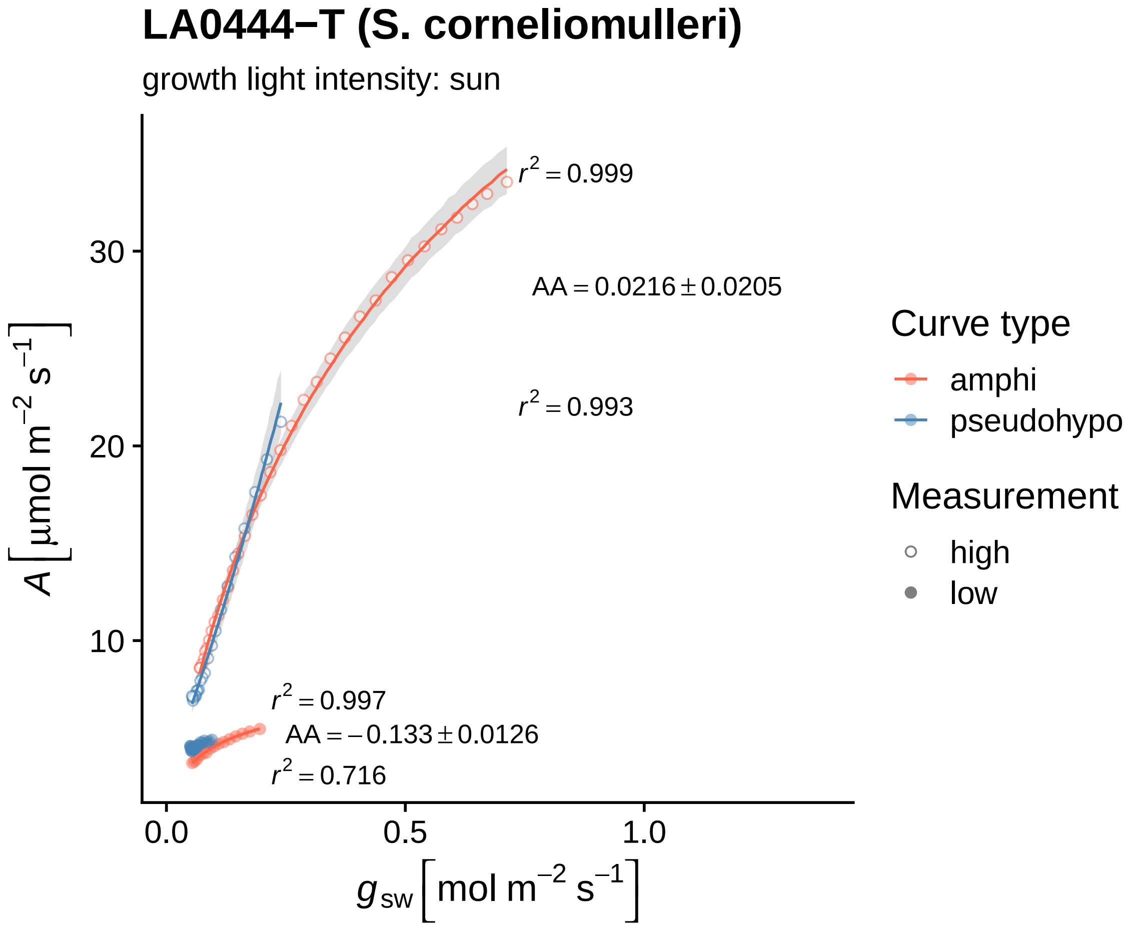
LA0444−T (S. corneliomulleri)
growth light intensity: sun
$A\,[\mu mol\ m^{-2}\ s^{-1}]$
$g_{sw}\,[mol\ m^{-2}\ s^{-1}]$
30
20
10
0.0
0.5
1.0
$r^2 = 0.999$
$AA = 0.0216 \pm 0.0205$
$r^2 = 0.993$
$r^2 = 0.997$
$AA = -0.133 \pm 0.0126$
$r^2 = 0.716$
Curve type
amphi
pseudohypo
Measurement
high
low

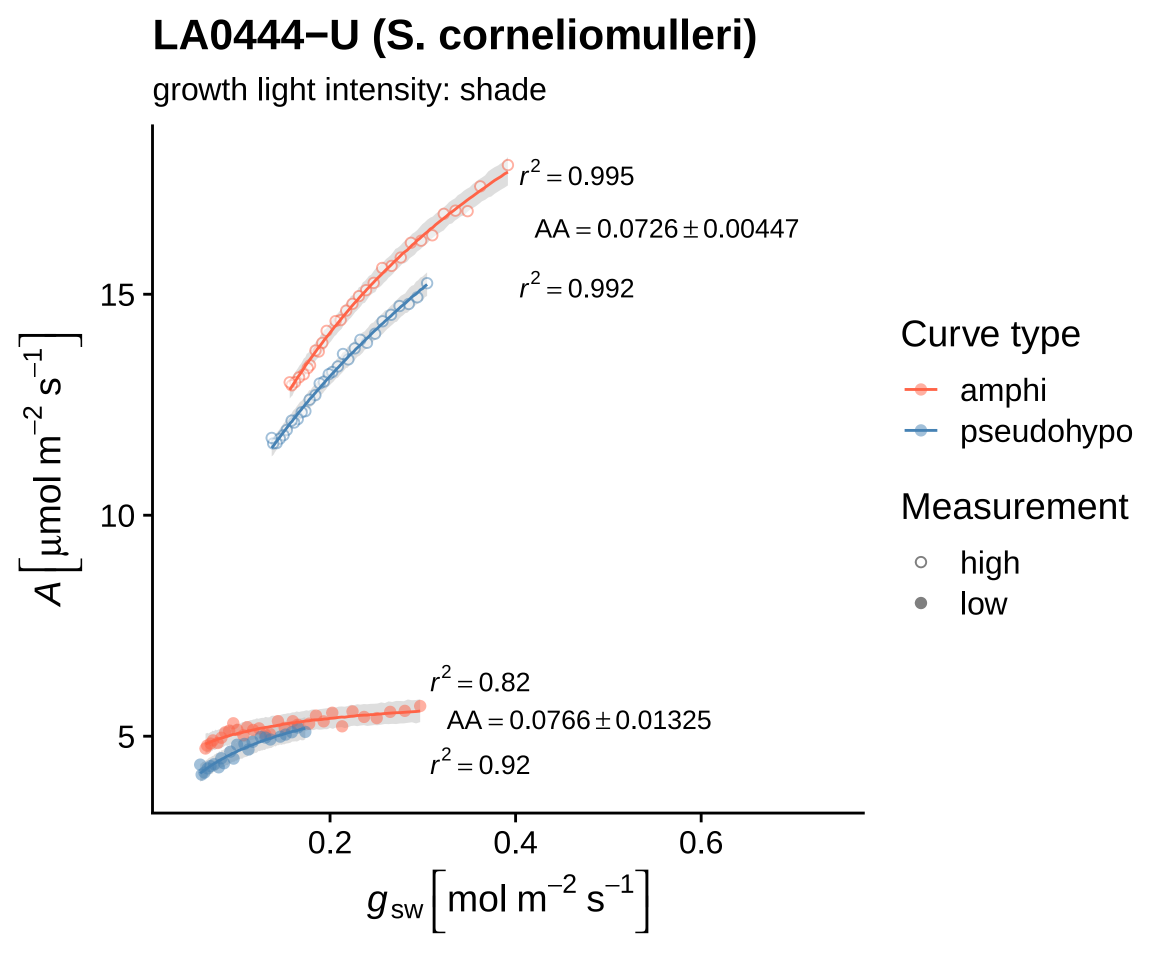

LA0444−U (S. corneliomulleri)
growth light intensity: shade
$r^2 = 0.995$
$AA = 0.0726 \pm 0.00447$
$r^2 = 0.992$
$r^2 = 0.82$
$AA = 0.0766 \pm 0.01325$
$r^2 = 0.92$
15
10
5
0.2
0.4
0.6
$A\left[\mu mol\ m^{-2}\ s^{-1}\right]$
$g_{sw}\left[mol\ m^{-2}\ s^{-1}\right]$
Curve type
amphi
pseudohypo
Measurement
high
low

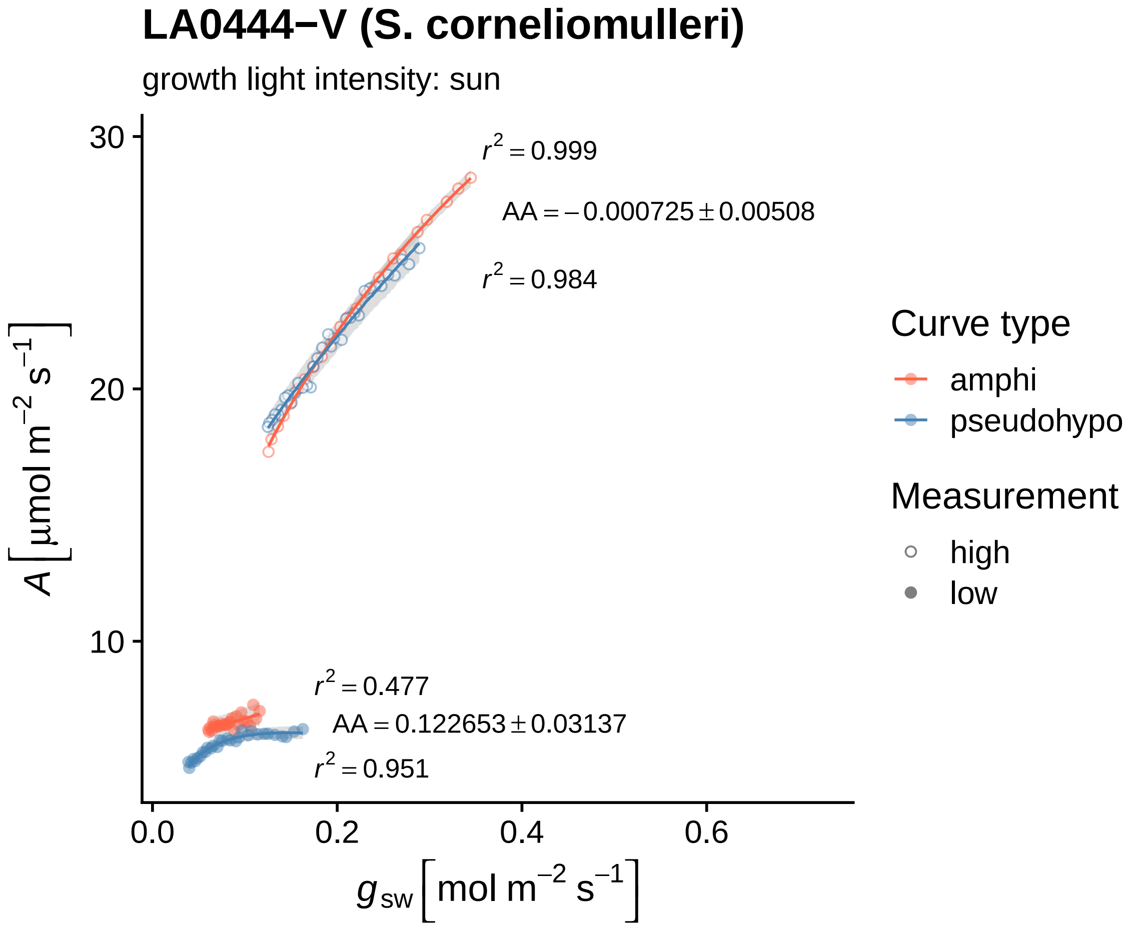
LA0444−V (S. corneliomulleri)
growth light intensity: sun
$r^2 = 0.999$
$AA = -0.000725 \pm 0.00508$
$r^2 = 0.984$
$r^2 = 0.477$
$AA = 0.122653 \pm 0.03137$
$r^2 = 0.951$
$A\ [\mu mol\ m^{-2}\ s^{-1}]$
$g_{sw}\ [mol\ m^{-2}\ s^{-1}]$
30
20
10
0.0
0.2
0.4
0.6
Curve type
amphi
pseudohypo
Measurement
high
low

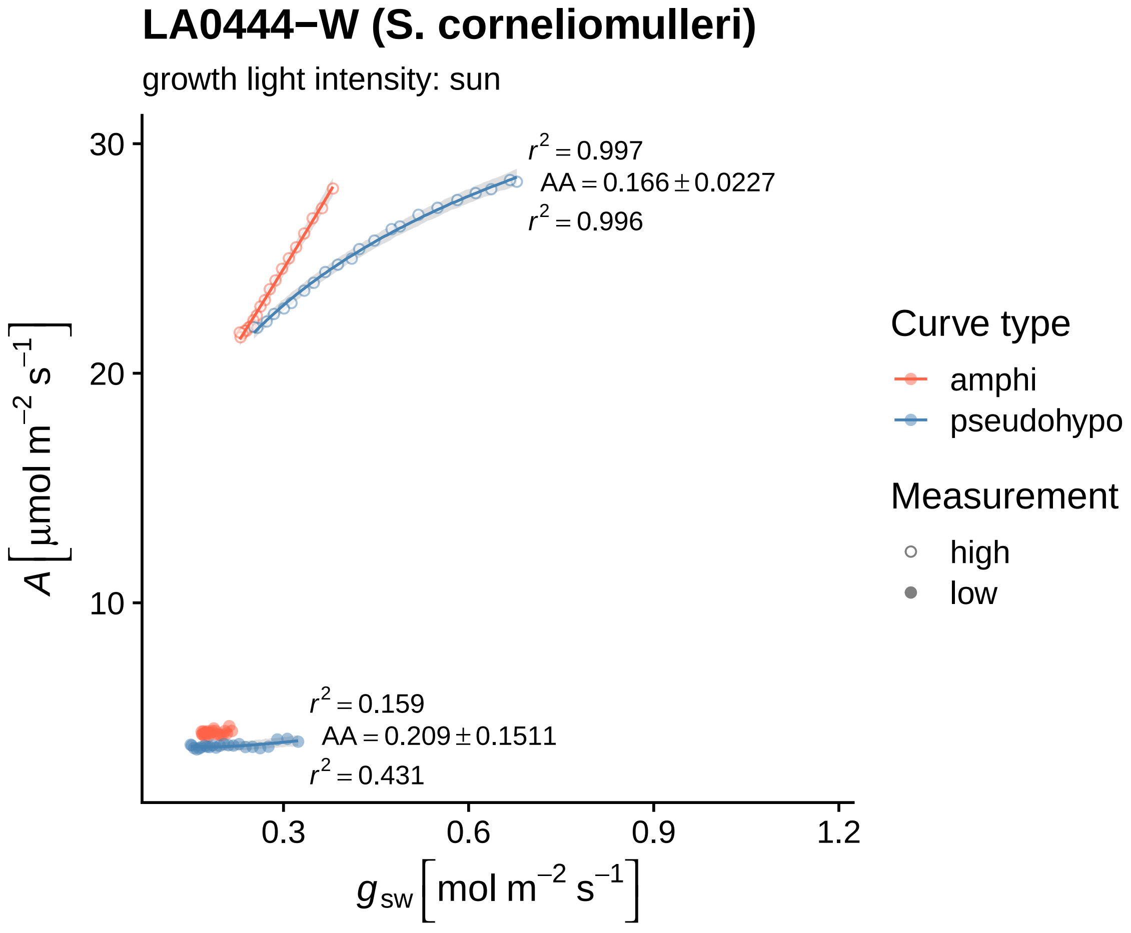

LA0444−W (S. corneliomulleri)
growth light intensity: sun
r2 = 0.997
AA = 0.166 ± 0.0227
r2 = 0.996
r2 = 0.159
AA = 0.209 ± 0.1511
r2 = 0.431
30
20
10
0.3
0.6
0.9
1.2
A [µmol m−2 s−1]
gsw [mol m−2 s−1]
Curve type
amphi
pseudohypo
Measurement
high
low

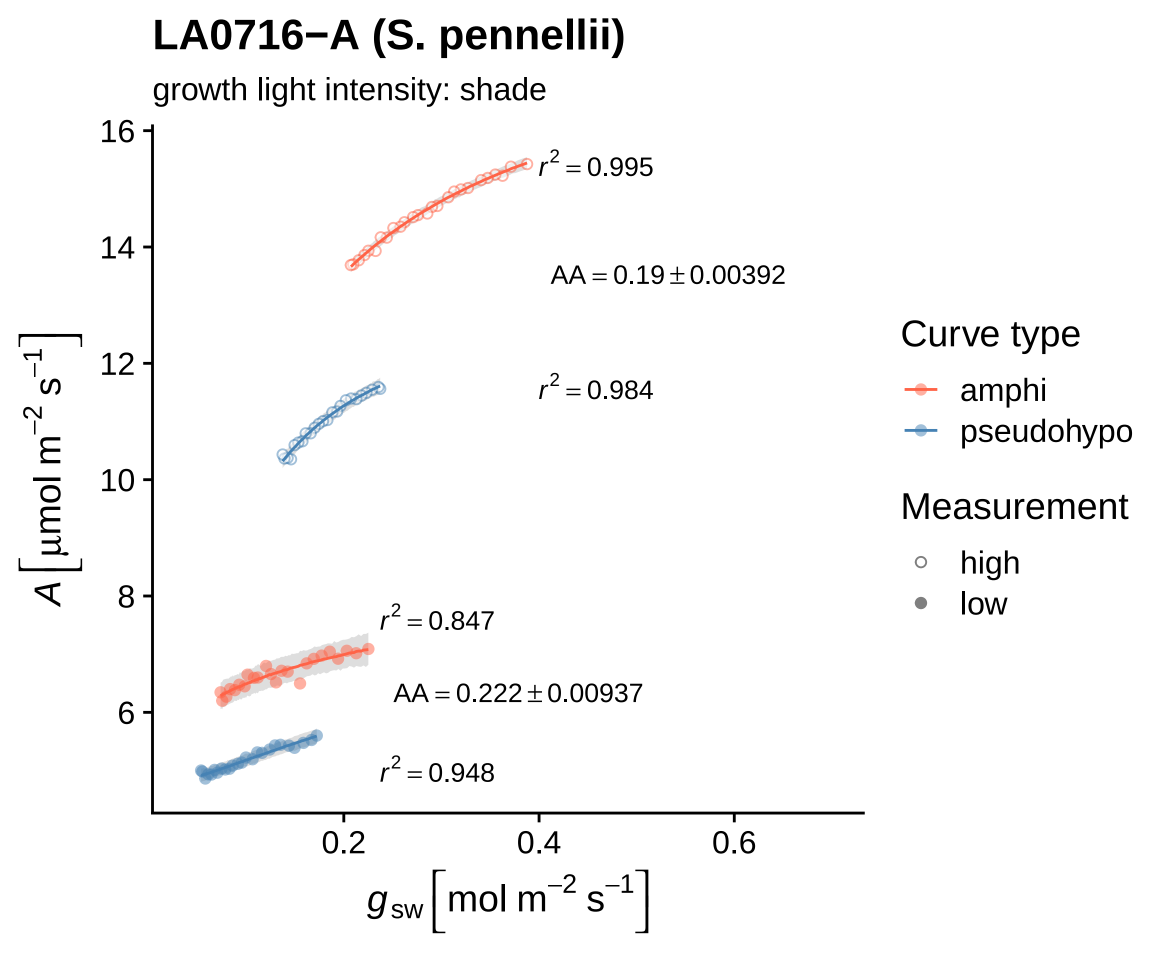

LA0716−A (S. pennellii)
growth light intensity: shade
$r^2 = 0.995$
$AA = 0.19 \pm 0.00392$
$r^2 = 0.984$
$r^2 = 0.847$
$AA = 0.222 \pm 0.00937$
$r^2 = 0.948$
$A\left[\mu mol\ m^{-2}\ s^{-1}\right]$
16
14
12
10
8
6
0.2
0.4
0.6
$g_{sw}\left[mol\ m^{-2}\ s^{-1}\right]$
Curve type
amphi
pseudohypo
Measurement
high
low

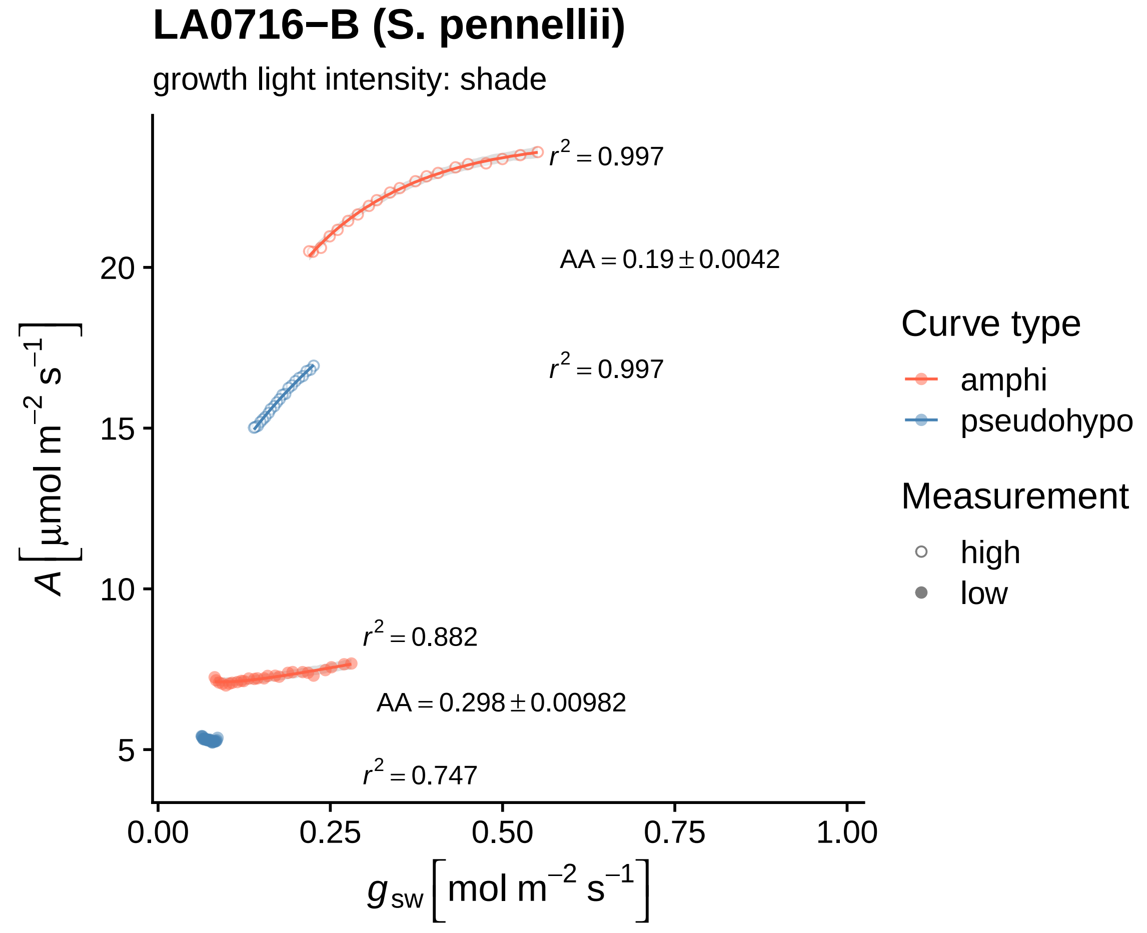

LA0716−B (S. pennellii)
growth light intensity: shade
$r^2 = 0.997$
AA = 0.19 ± 0.0042
$r^2 = 0.997$
$r^2 = 0.882$
AA = 0.298 ± 0.00982
$r^2 = 0.747$
20
15
10
5
0.00
0.25
0.50
0.75
1.00
$A$ [μmol m$^{-2}$ s$^{-1}$]
$g_{sw}$ [mol m$^{-2}$ s$^{-1}$]
Curve type
amphi
pseudohypo
Measurement
high
low

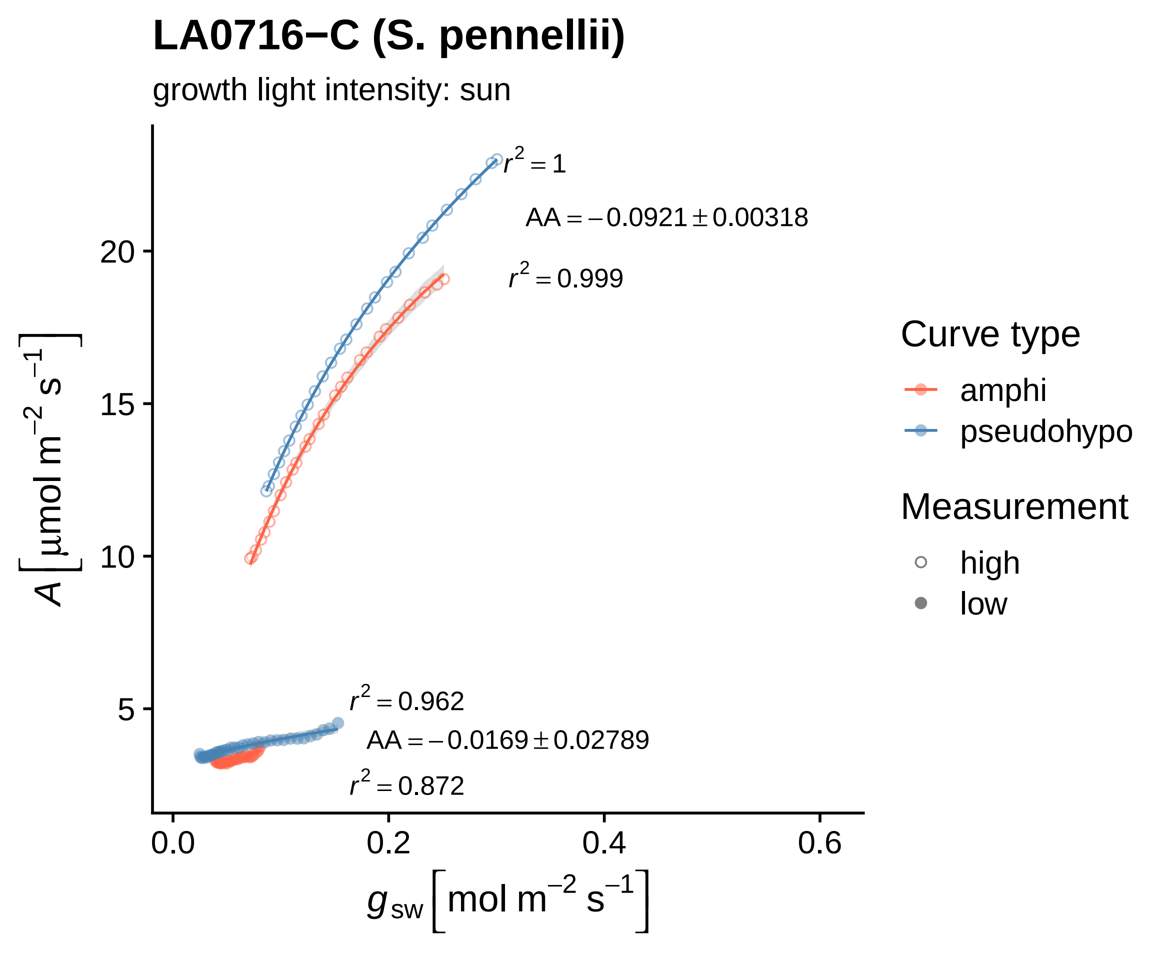
LA0716−C (S. pennellii)
growth light intensity: sun
$r^2 = 1$
AA = −0.0921 ± 0.00318
$r^2 = 0.999$
$r^2 = 0.962$
AA = −0.0169 ± 0.02789
$r^2 = 0.872$
A [µmol m−2 s−1]
20
15
10
5
0.0
0.2
0.4
0.6
gsw [mol m−2 s−1]
Curve type
amphi
pseudohypo
Measurement
high
low

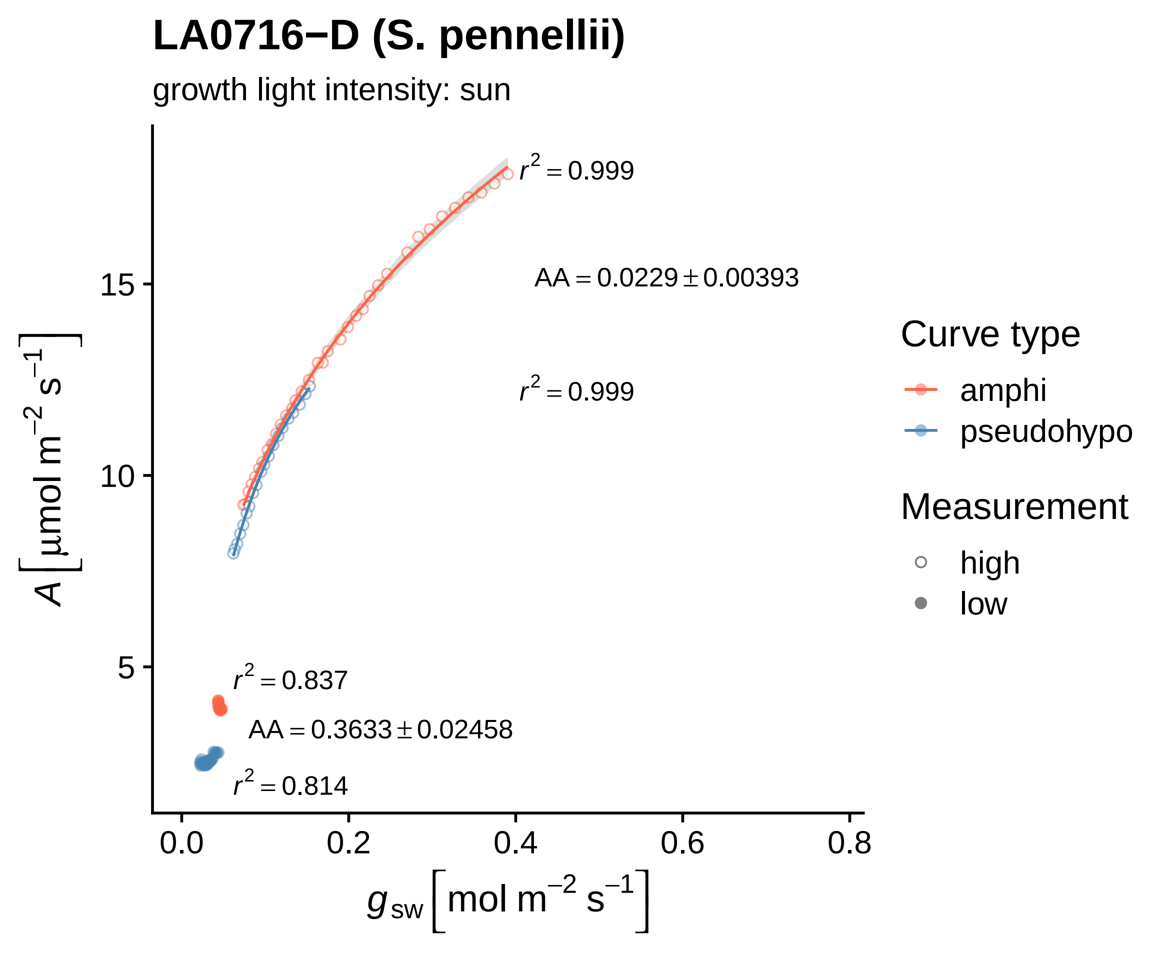

LA0716−D (S. pennellii)
growth light intensity: sun
r^2 = 0.999
AA = 0.0229 ± 0.00393
r^2 = 0.999
r^2 = 0.837
AA = 0.3633 ± 0.02458
r^2 = 0.814
15
10
5
0.0
0.2
0.4
0.6
0.8
A [μmol m^−2 s^−1]
g_sw [mol m^−2 s^−1]
Curve type
amphi
pseudohypo
Measurement
high
low

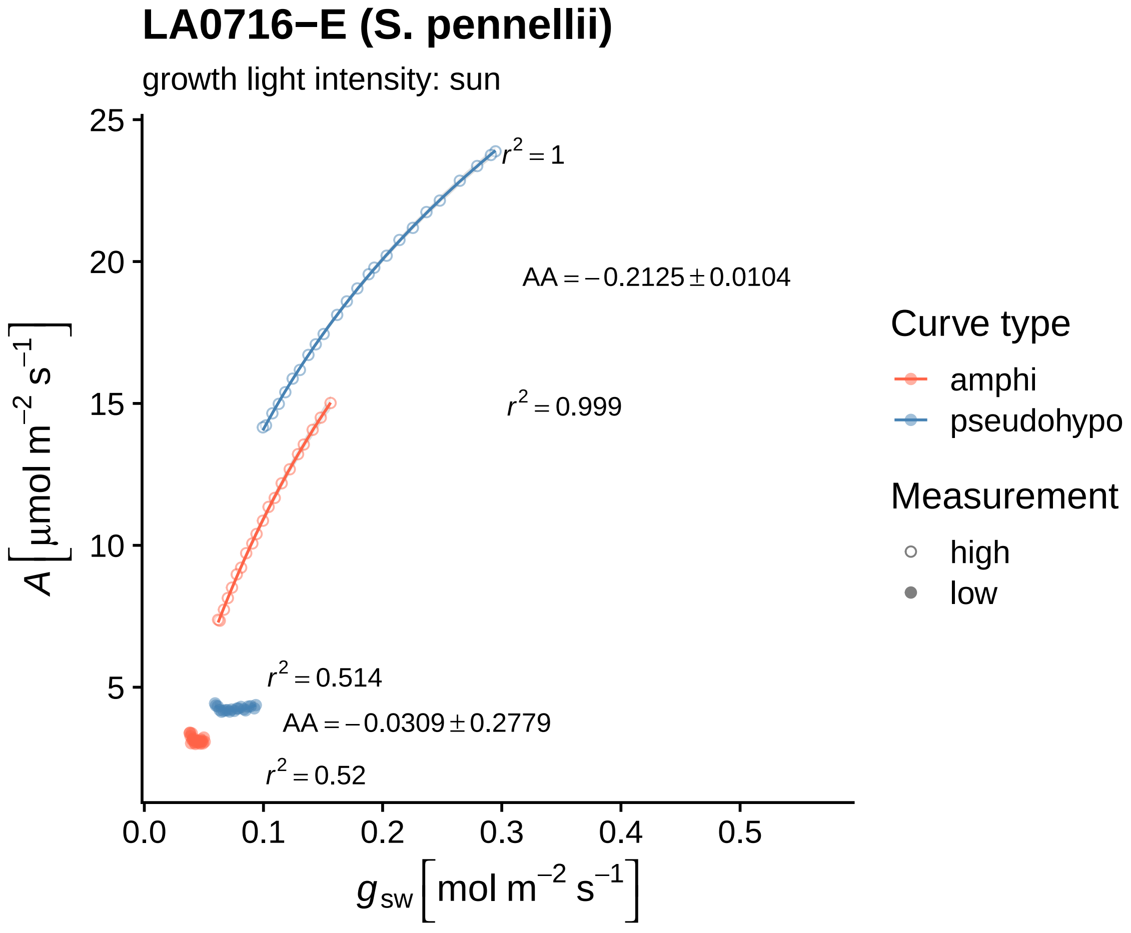

LA0716−E (S. pennellii)
growth light intensity: sun
$r^2 = 1$
$AA = -0.2125 \pm 0.0104$
$r^2 = 0.999$
$r^2 = 0.514$
$AA = -0.0309 \pm 0.2779$
$r^2 = 0.52$
25
20
15
10
5
0.0
0.1
0.2
0.3
0.4
0.5
$A\left[\mu mol\ m^{-2}\ s^{-1}\right]$
$g_{sw}\left[mol\ m^{-2}\ s^{-1}\right]$
Curve type
amphi
pseudohypo
Measurement
high
low

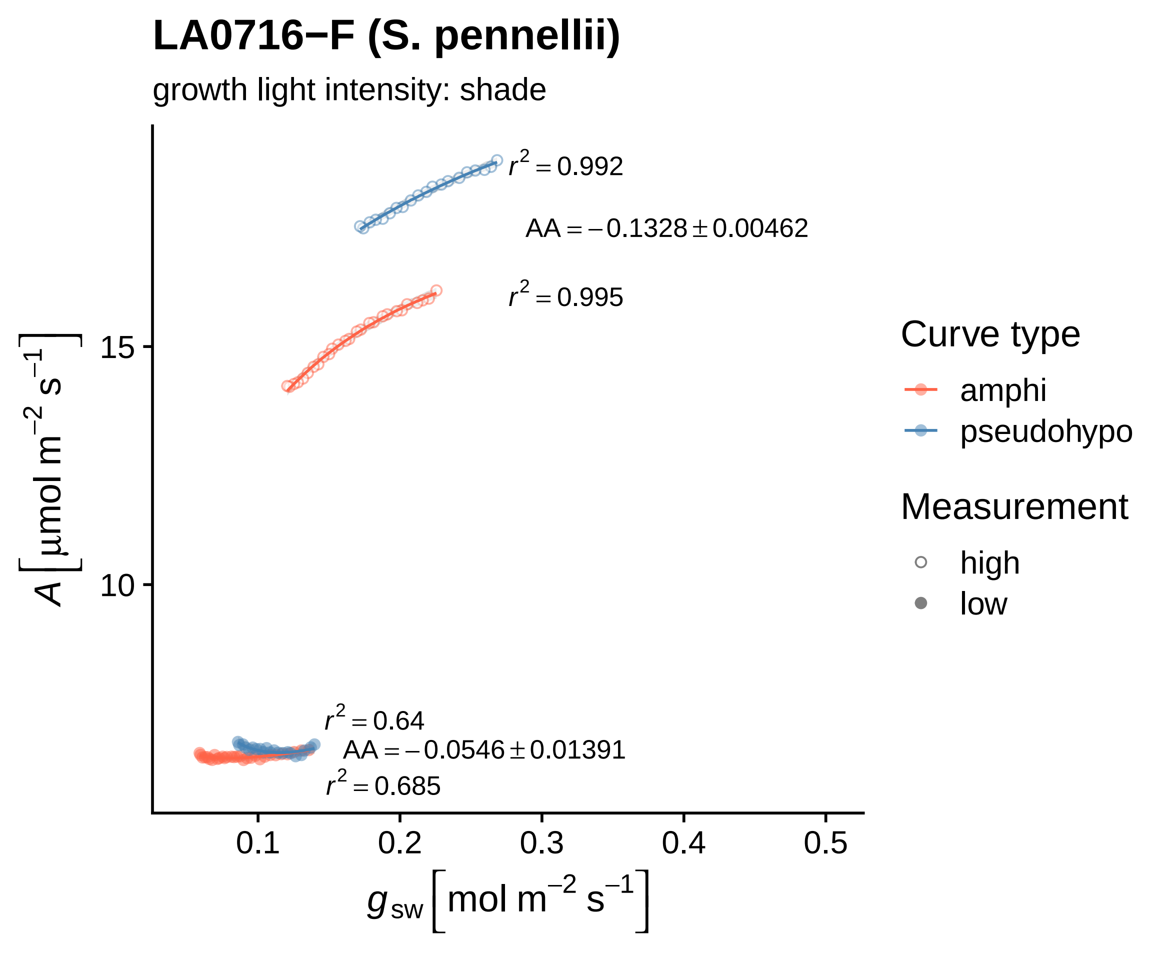

LA0716−F (S. pennellii)
growth light intensity: shade
$r^2 = 0.992$
$AA = -0.1328 \pm 0.00462$
$r^2 = 0.995$
$r^2 = 0.64$
$AA = -0.0546 \pm 0.01391$
$r^2 = 0.685$
15
10
0.1
0.2
0.3
0.4
0.5
$A\ [\mu mol\ m^{-2}\ s^{-1}]$
$g_{sw}\ [mol\ m^{-2}\ s^{-1}]$
Curve type
amphi
pseudohypo
Measurement
high
low

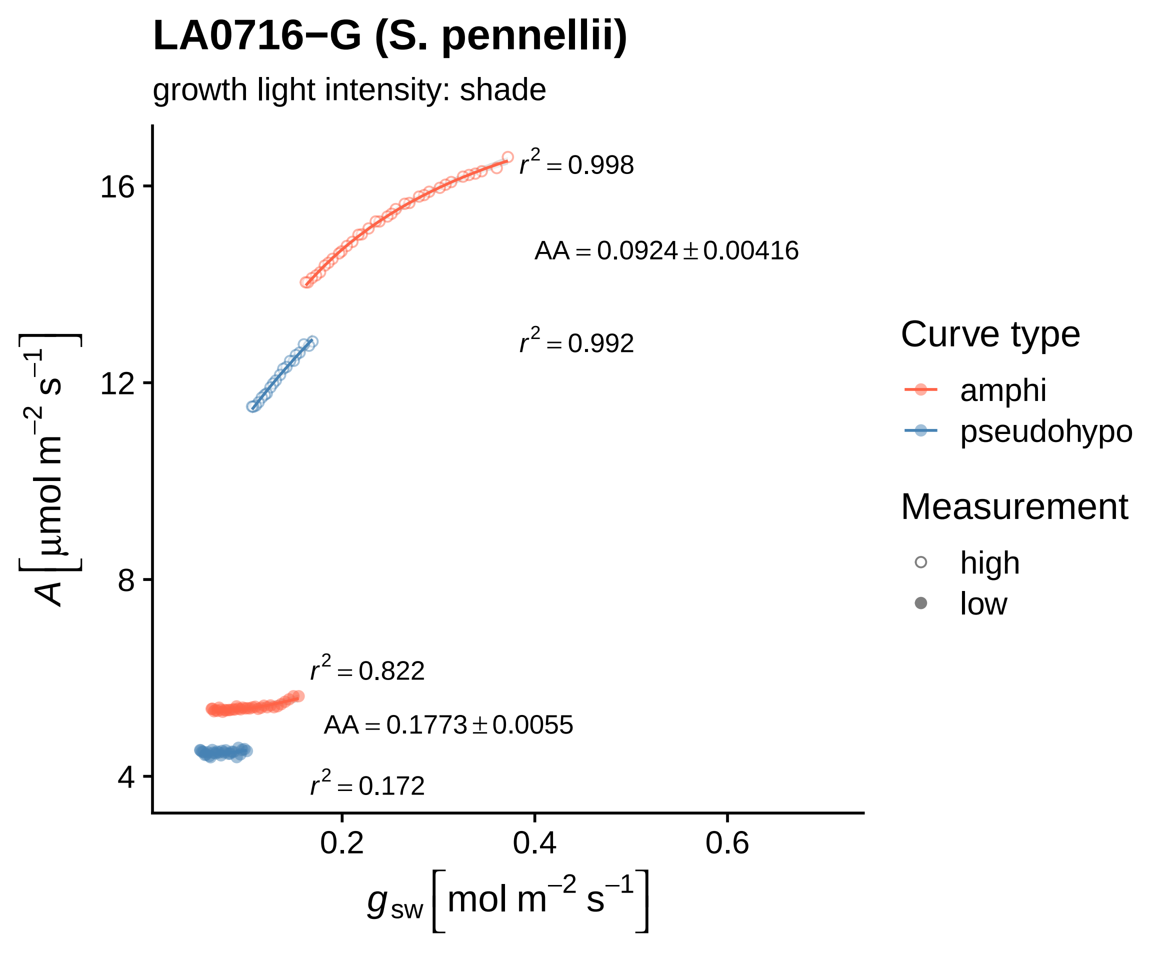

LA0716−G (S. pennellii)
growth light intensity: shade
$r^2 = 0.998$
$AA = 0.0924 \pm 0.00416$
$r^2 = 0.992$
$r^2 = 0.822$
$AA = 0.1773 \pm 0.0055$
$r^2 = 0.172$
16
12
8
4
0.2
0.4
0.6
$A\left[\mu mol\ m^{-2}\ s^{-1}\right]$
$g_{sw}\left[mol\ m^{-2}\ s^{-1}\right]$
Curve type
amphi
pseudohypo
Measurement
high
low

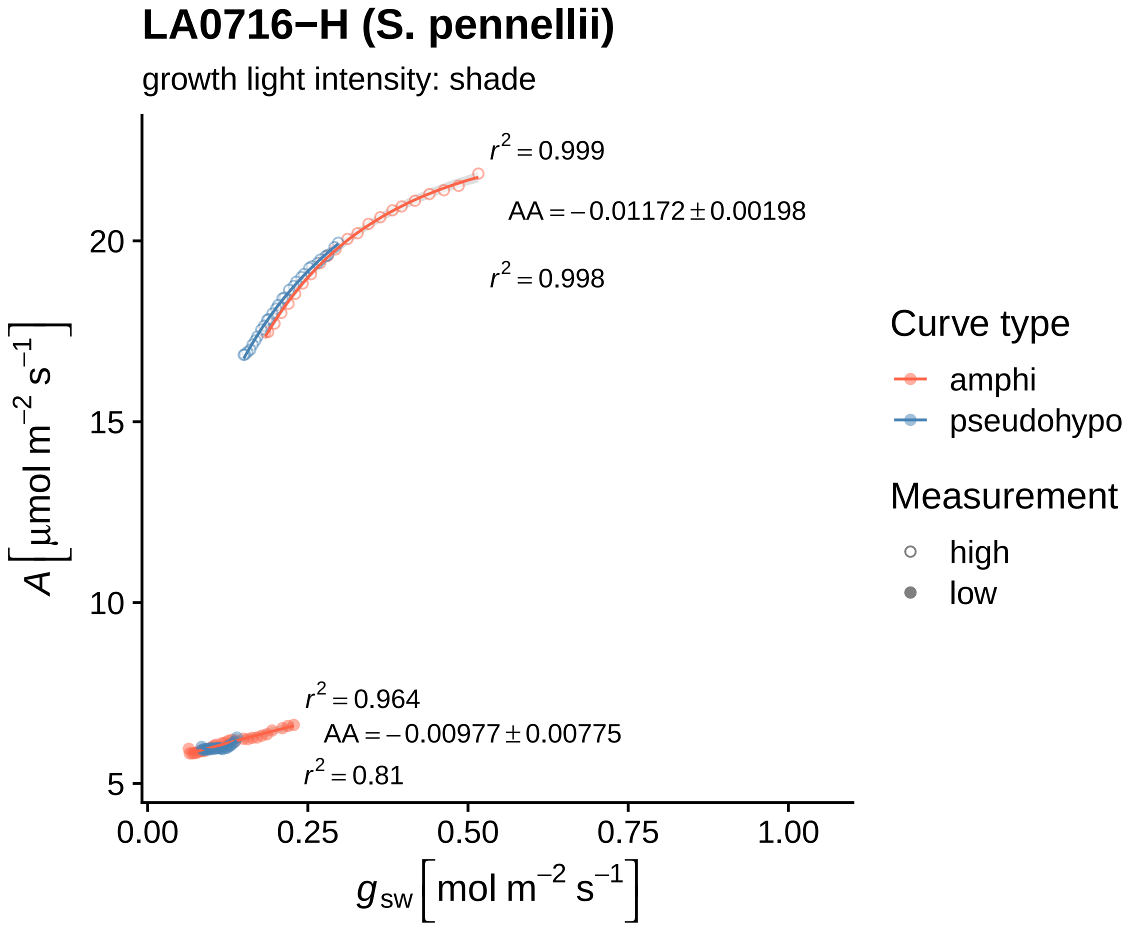
LA0716−H (S. pennellii)
growth light intensity: shade
$r^2 = 0.999$
$AA = -0.01172 \pm 0.00198$
$r^2 = 0.998$
$r^2 = 0.964$
$AA = -0.00977 \pm 0.00775$
$r^2 = 0.81$
$A\ [\mu mol\ m^{-2}\ s^{-1}]$
20
15
10
5
0.00
0.25
0.50
0.75
1.00
$g_{sw}\ [mol\ m^{-2}\ s^{-1}]$
Curve type
amphi
pseudohypo
Measurement
high
low

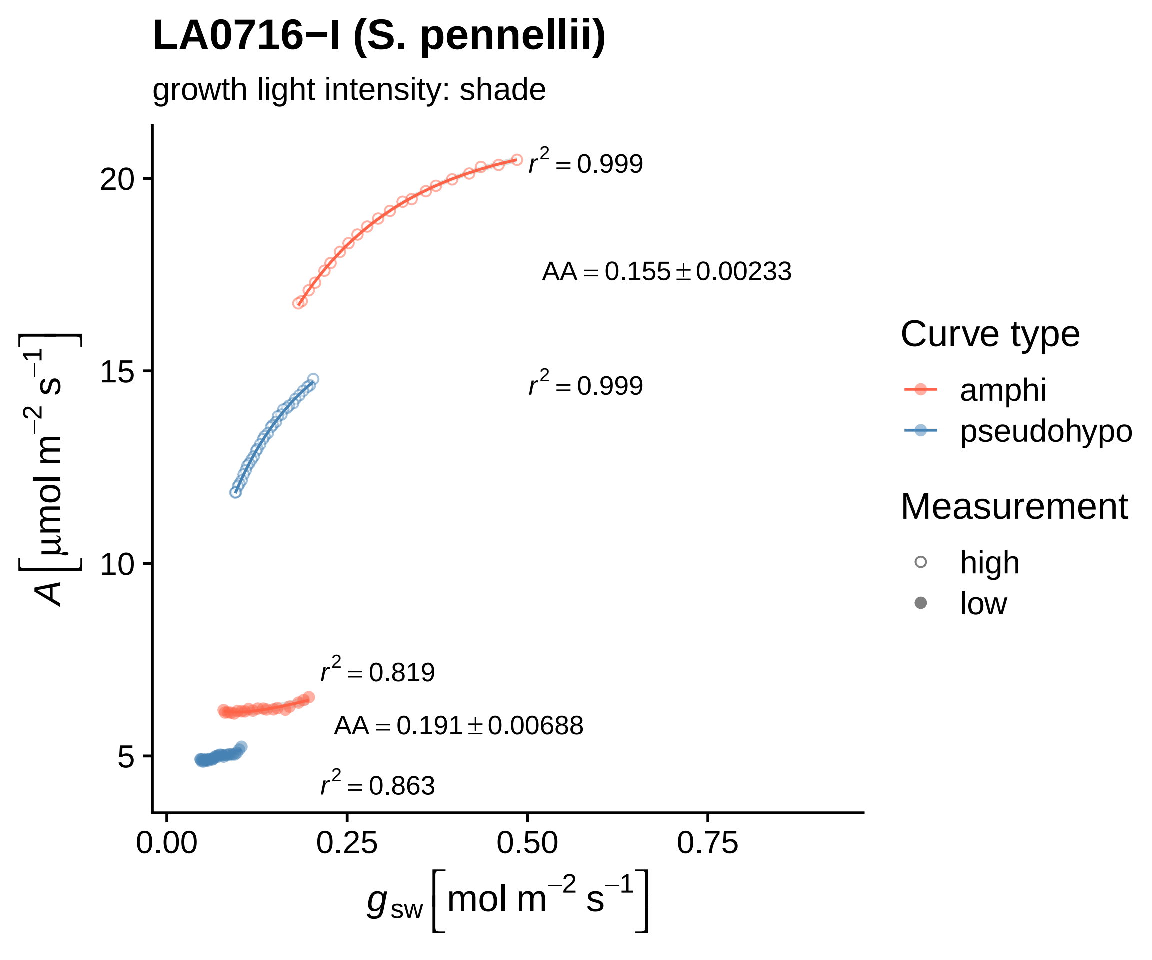

LA0716−I (S. pennellii)
growth light intensity: shade
$r^2 = 0.999$
$AA = 0.155 \pm 0.00233$
$r^2 = 0.999$
$r^2 = 0.819$
$AA = 0.191 \pm 0.00688$
$r^2 = 0.863$
20
15
10
5
0.00
0.25
0.50
0.75
$A\ [\mu mol\ m^{-2}\ s^{-1}]$
$g_{sw}\ [mol\ m^{-2}\ s^{-1}]$
Curve type
amphi
pseudohypo
Measurement
high
low

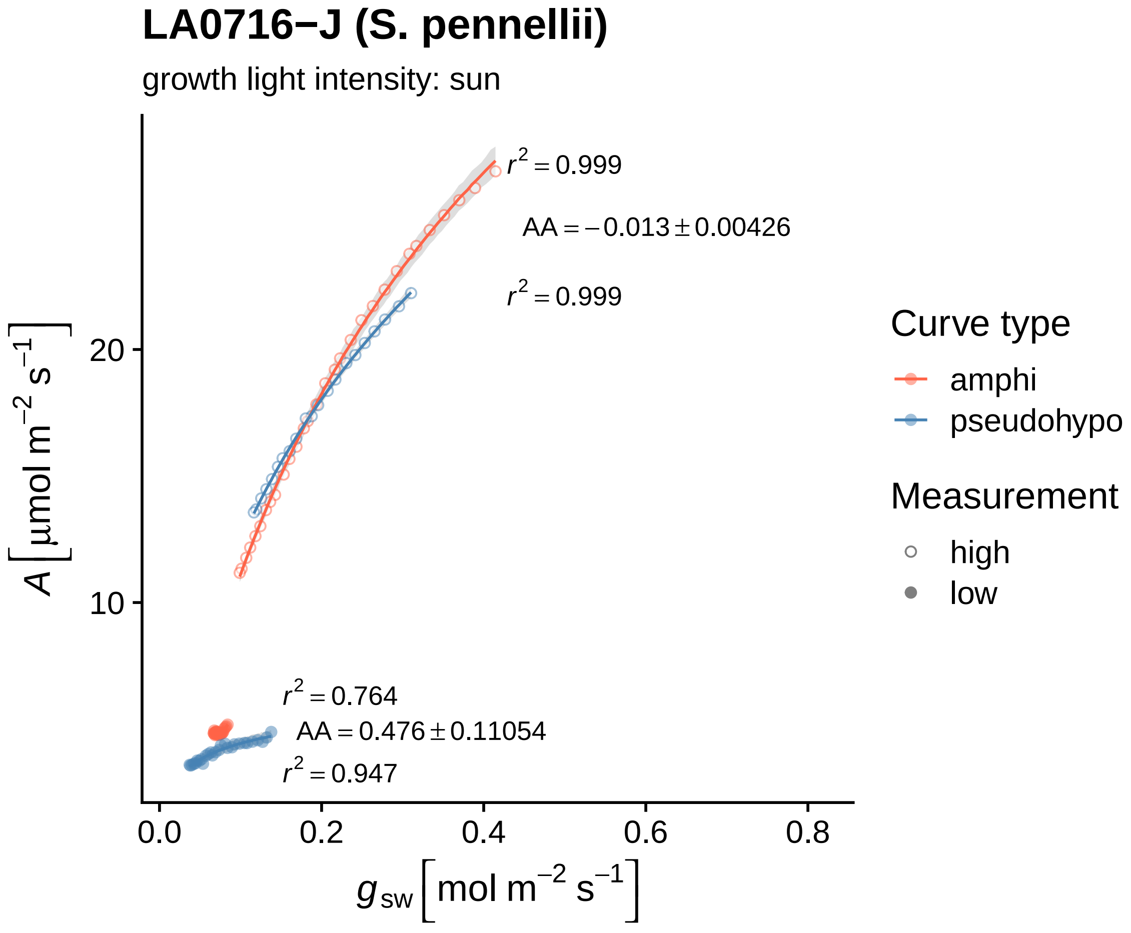
LA0716−J (S. pennellii)
growth light intensity: sun
$r^2 = 0.999$
$AA = -0.013 \pm 0.00426$
$r^2 = 0.999$
$r^2 = 0.764$
$AA = 0.476 \pm 0.11054$
$r^2 = 0.947$
20
10
0.0
0.2
0.4
0.6
0.8
$A\,[\mu mol\ m^{-2}\ s^{-1}]$
$g_{sw}\,[mol\ m^{-2}\ s^{-1}]$
Curve type
amphi
pseudohypo
Measurement
high
low

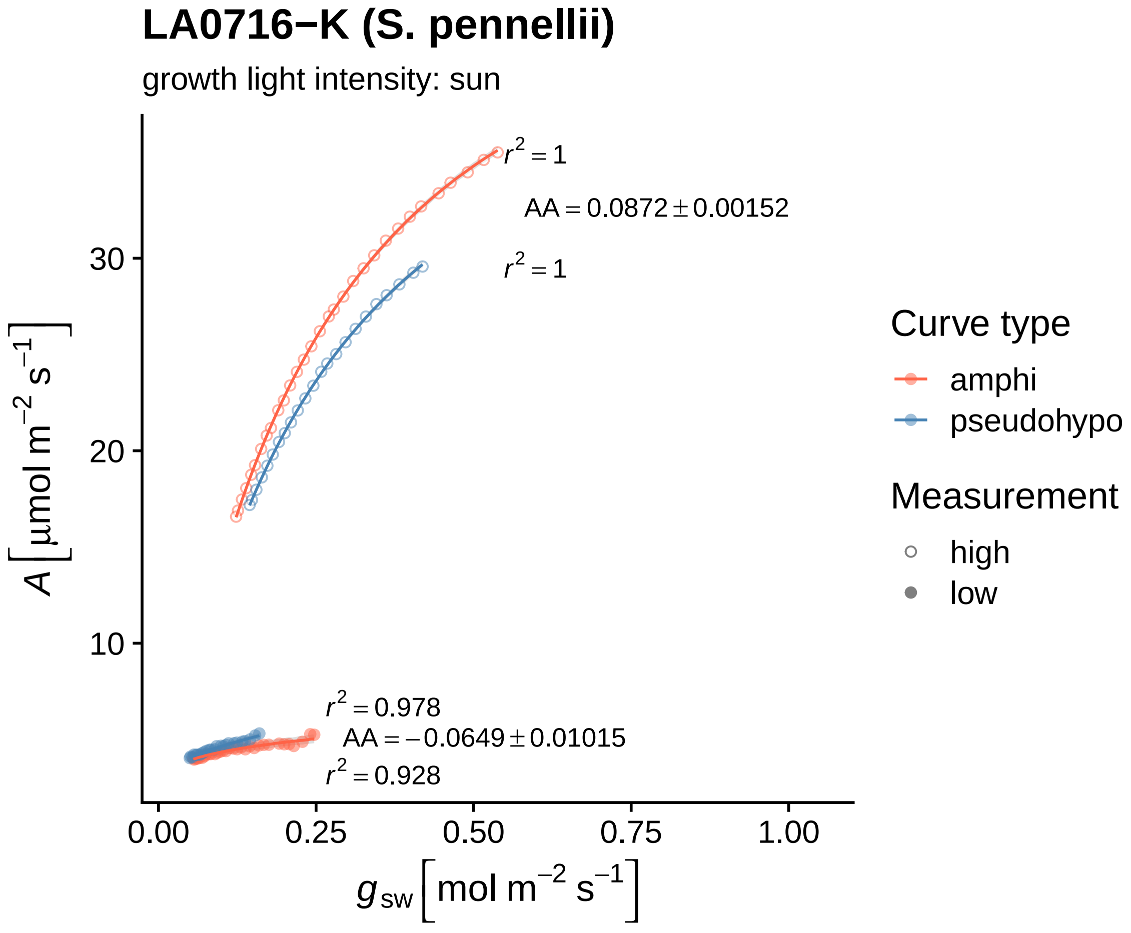

LA0716−K (S. pennellii)
growth light intensity: sun
$r^2 = 1$
AA = 0.0872 ± 0.00152
$r^2 = 1$
$r^2 = 0.978$
AA = −0.0649 ± 0.01015
$r^2 = 0.928$
30
20
10
0.00
0.25
0.50
0.75
1.00
$A\ [\mu mol\ m^{-2}\ s^{-1}]$
$g_{sw}\ [mol\ m^{-2}\ s^{-1}]$
Curve type
amphi
pseudohypo
Measurement
high
low

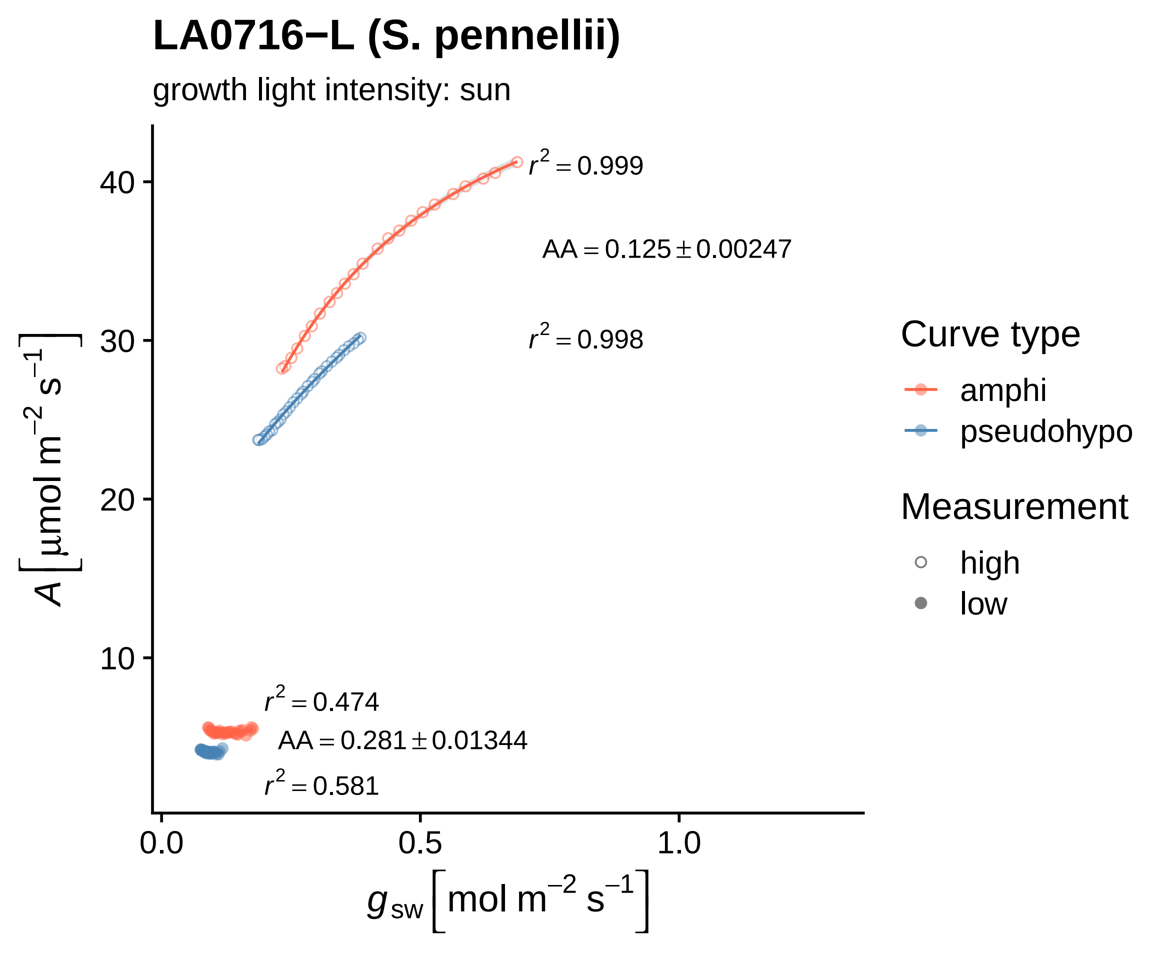
LA0716−L (S. pennellii)
growth light intensity: sun
$r^2 = 0.999$
$AA = 0.125 \pm 0.00247$
$r^2 = 0.998$
$r^2 = 0.474$
$AA = 0.281 \pm 0.01344$
$r^2 = 0.581$
40
30
20
10
0.0
0.5
1.0
$A\left[\mu mol\ m^{-2}\ s^{-1}\right]$
$g_{sw}\left[mol\ m^{-2}\ s^{-1}\right]$
Curve type
amphi
pseudohypo
Measurement
high
low

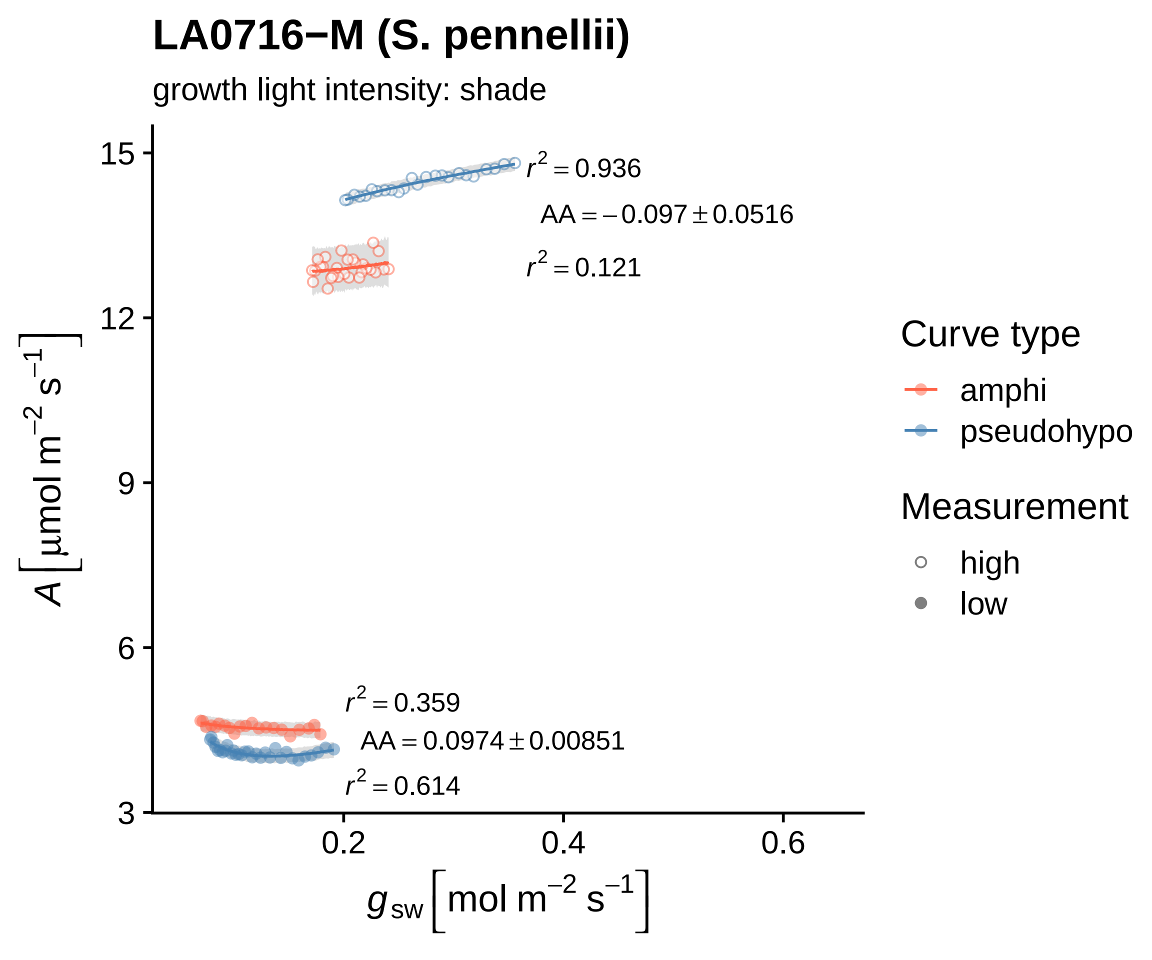

LA0716−M (S. pennellii)
growth light intensity: shade
$r^2 = 0.936$
$AA = -0.097 \pm 0.0516$
$r^2 = 0.121$
$r^2 = 0.359$
$AA = 0.0974 \pm 0.00851$
$r^2 = 0.614$
15
12
9
6
3
0.2
0.4
0.6
$A\left[\mu mol\ m^{-2}\ s^{-1}\right]$
$g_{sw}\left[mol\ m^{-2}\ s^{-1}\right]$
Curve type
amphi
pseudohypo
Measurement
high
low

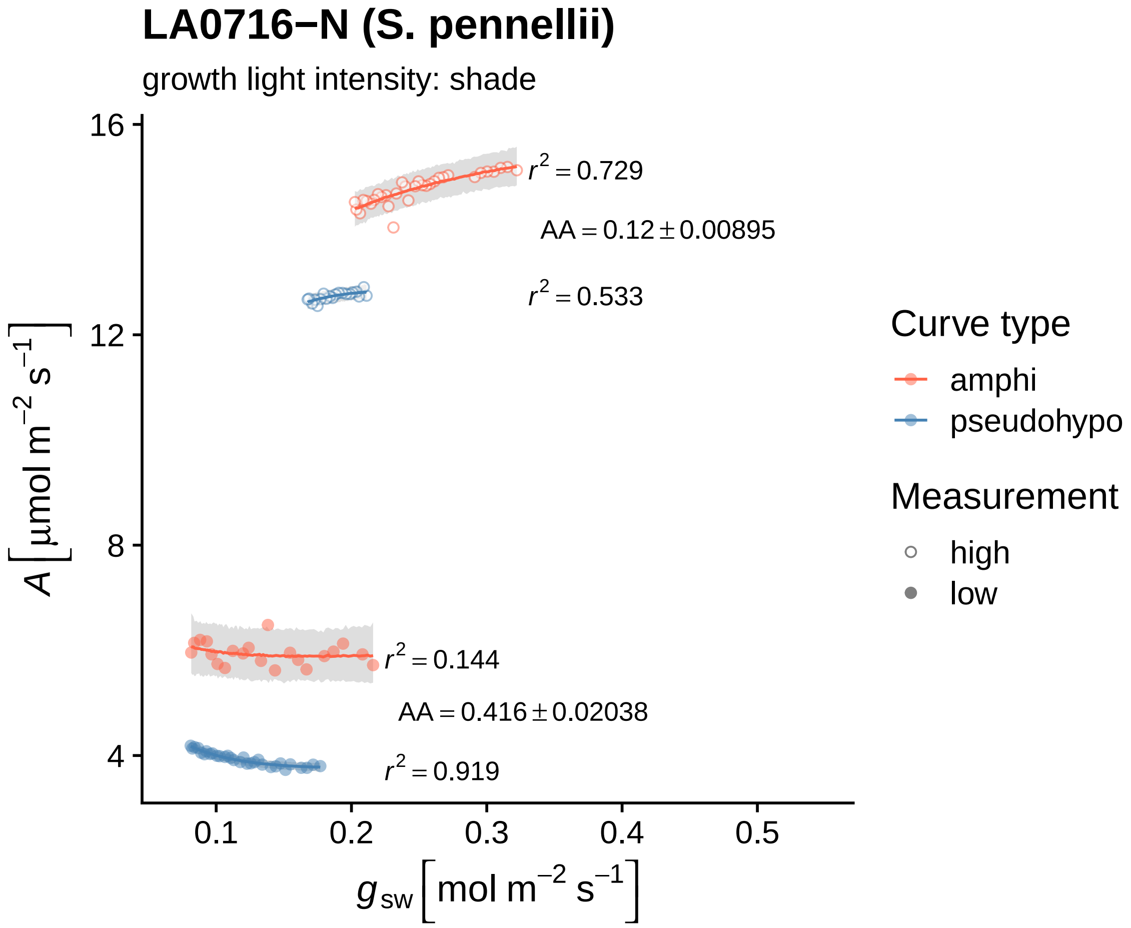

LA0716−N (S. pennellii)
growth light intensity: shade
$r^2 = 0.729$
$AA = 0.12 \pm 0.00895$
$r^2 = 0.533$
$r^2 = 0.144$
$AA = 0.416 \pm 0.02038$
$r^2 = 0.919$
16
12
8
4
0.1
0.2
0.3
0.4
0.5
$A\ [\mu mol\ m^{-2}\ s^{-1}]$
$g_{sw}\ [mol\ m^{-2}\ s^{-1}]$
Curve type
amphi
pseudohypo
Measurement
high
low

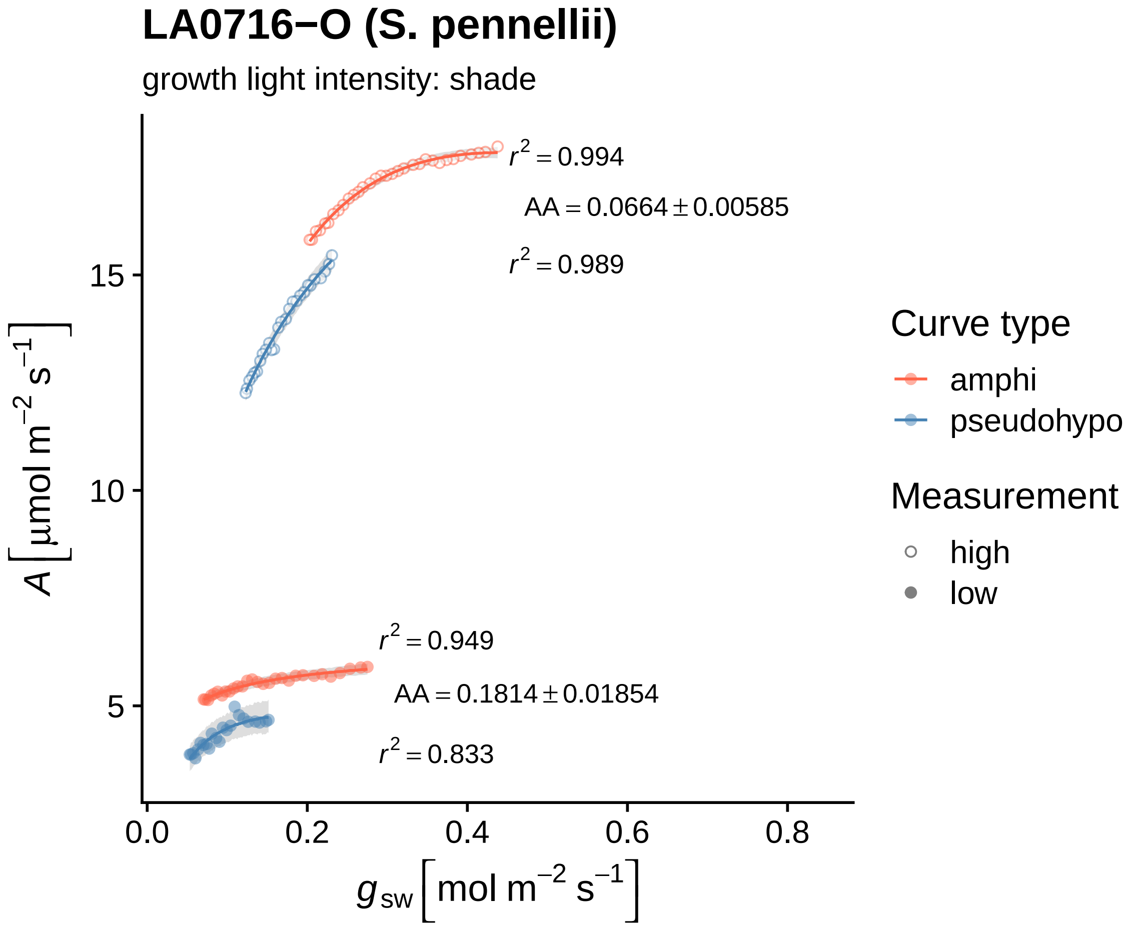

LA0716−O (S. pennellii)
growth light intensity: shade
r^2 = 0.994
AA = 0.0664 ± 0.00585
r^2 = 0.989
r^2 = 0.949
AA = 0.1814 ± 0.01854
r^2 = 0.833
15
10
5
0.0
0.2
0.4
0.6
0.8
A [µmol m^−2 s^−1]
g_sw [mol m^−2 s^−1]
Curve type
amphi
pseudohypo
Measurement
high
low

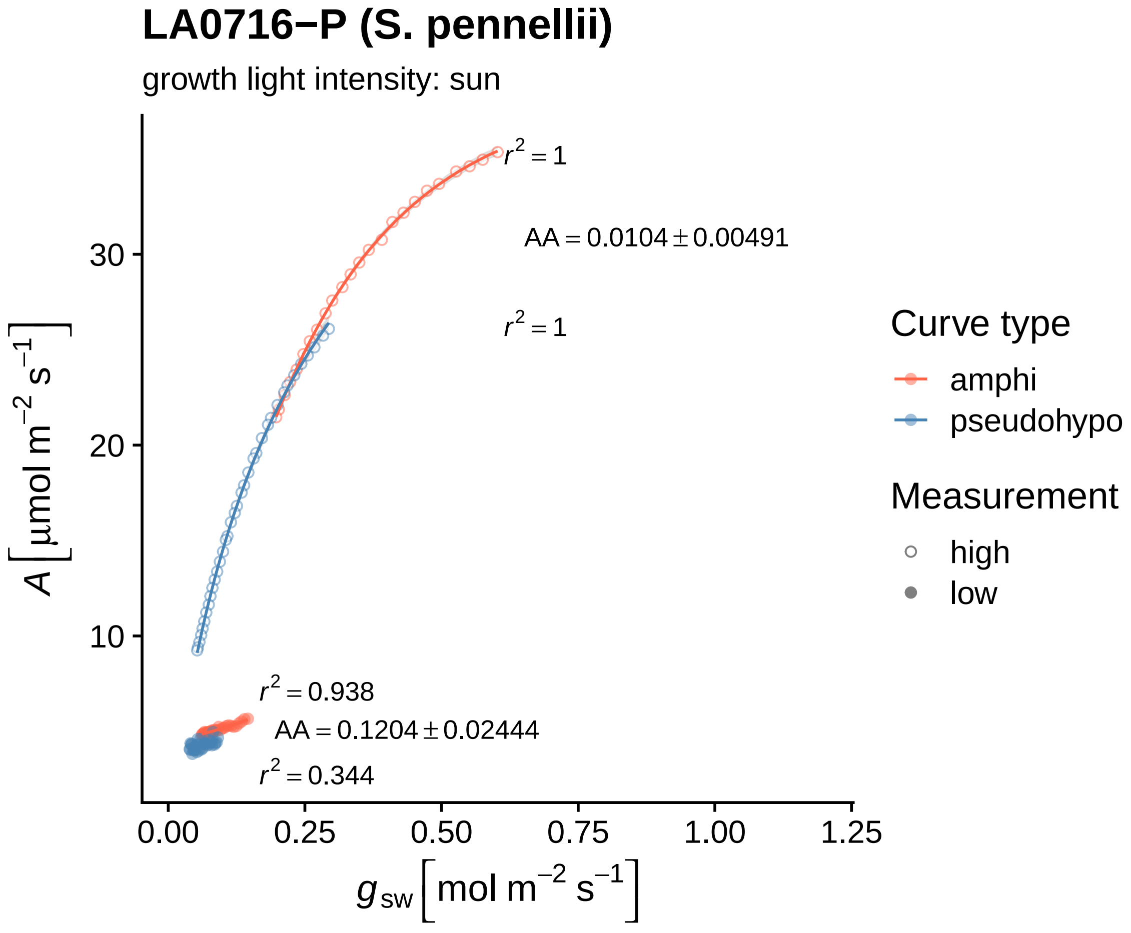

LA0716−P (S. pennellii)
growth light intensity: sun
r² = 1
AA = 0.0104 ± 0.00491
r² = 1
r² = 0.938
AA = 0.1204 ± 0.02444
r² = 0.344
A [μmol m⁻² s⁻¹]
10
20
30
0.00
0.25
0.50
0.75
1.00
1.25
g sw [mol m⁻² s⁻¹]
Curve type
amphi
pseudohypo
Measurement
high
low

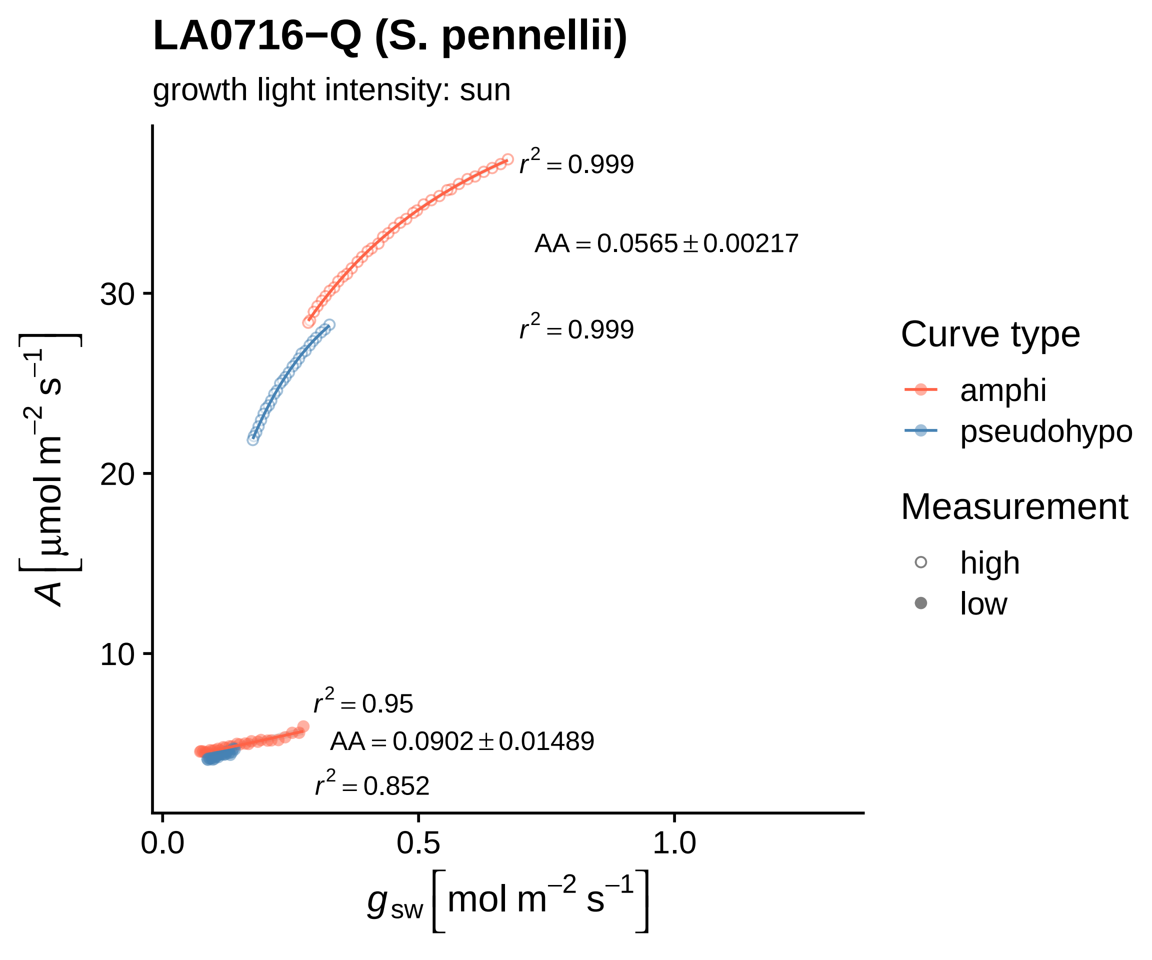

LA0716−Q (S. pennellii)
growth light intensity: sun
$r^2 = 0.999$
$AA = 0.0565 \pm 0.00217$
$r^2 = 0.999$
$r^2 = 0.95$
$AA = 0.0902 \pm 0.01489$
$r^2 = 0.852$
30
20
10
0.0
0.5
1.0
$A\ [\mu mol\ m^{-2}\ s^{-1}]$
$g_{sw}\ [mol\ m^{-2}\ s^{-1}]$
Curve type
amphi
pseudohypo
Measurement
high
low

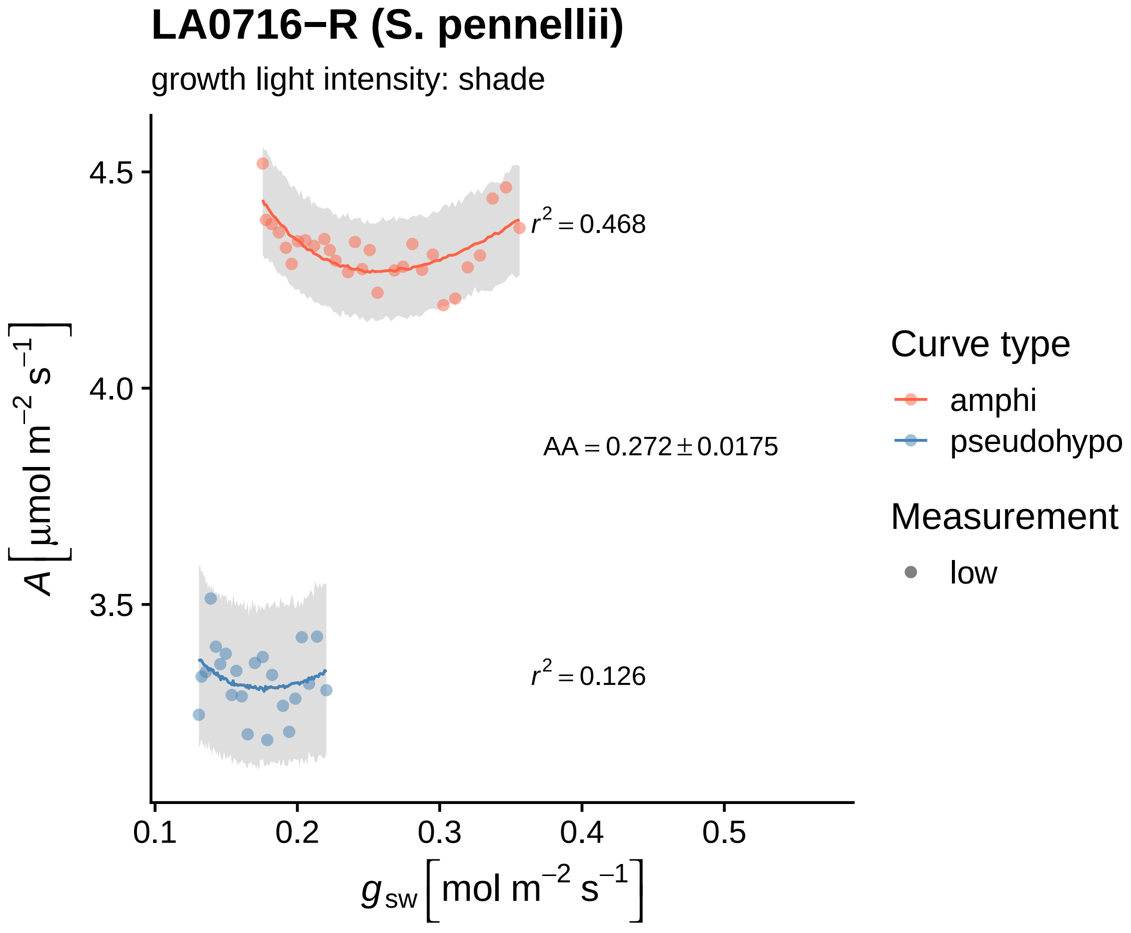

LA0716−R (S. pennellii)
growth light intensity: shade
4.5
4.0
3.5
$r^2 = 0.468$
$AA = 0.272 \pm 0.0175$
$r^2 = 0.126$
$A\left[\mu mol\ m^{-2}\ s^{-1}\right]$
0.1
0.2
0.3
0.4
0.5
$g_{sw}\left[mol\ m^{-2}\ s^{-1}\right]$
Curve type
amphi
pseudohypo
Measurement
low

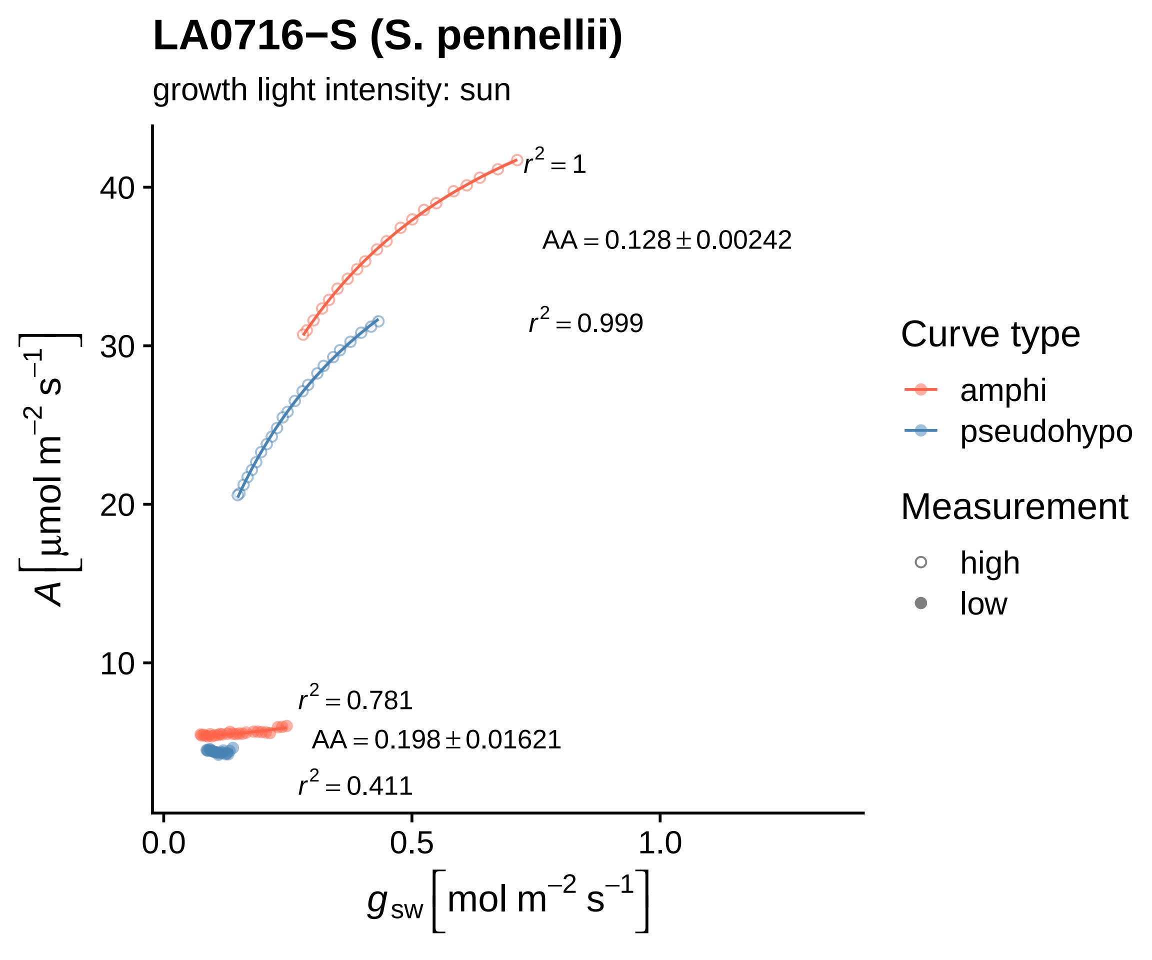
LA0716−S (S. pennellii)
growth light intensity: sun
$r^2 = 1$
AA = 0.128 ± 0.00242
$r^2 = 0.999$
$r^2 = 0.781$
AA = 0.198 ± 0.01621
$r^2 = 0.411$
10
20
30
40
0.0
0.5
1.0
$A\ [\mu mol\ m^{-2}\ s^{-1}]$
$g_{sw}\ [mol\ m^{-2}\ s^{-1}]$
Curve type
amphi
pseudohypo
Measurement
high
low

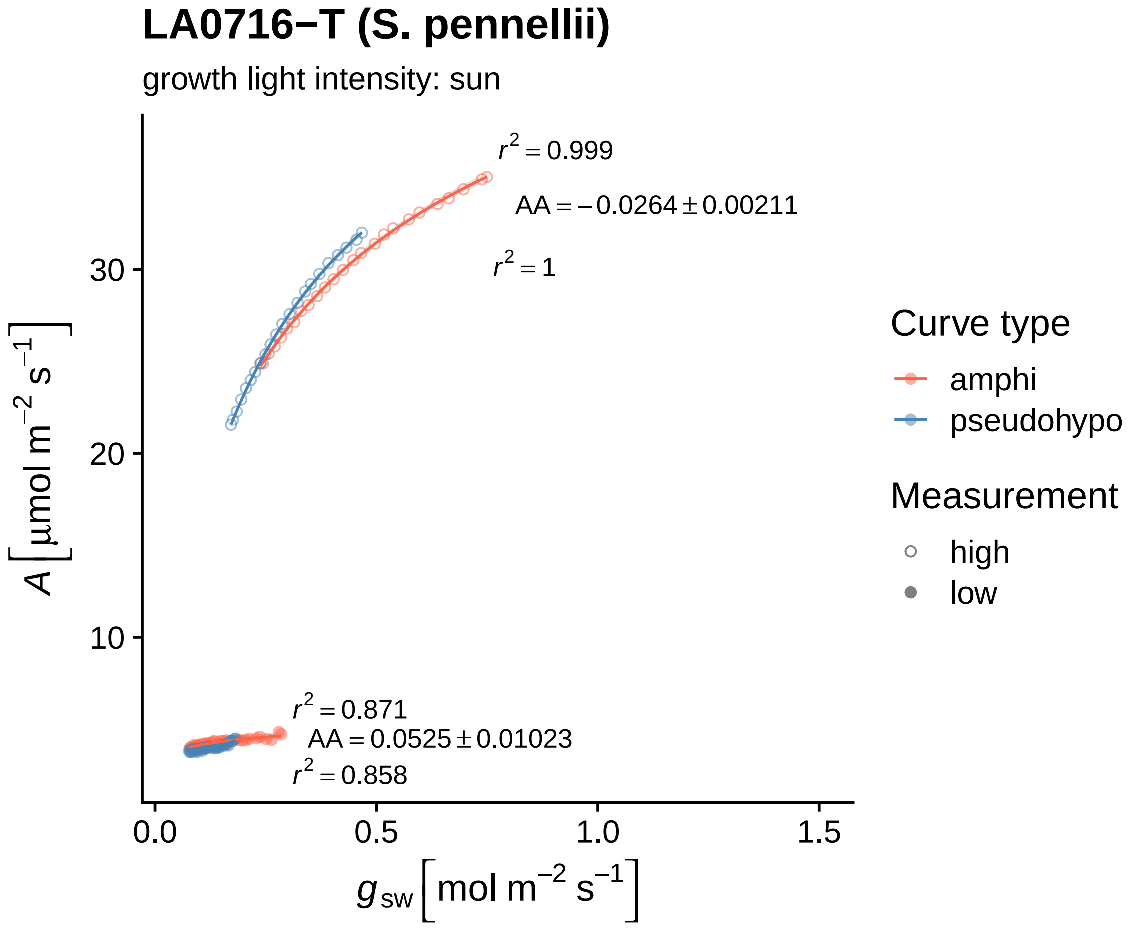
LA0716−T (S. pennellii)
growth light intensity: sun
r² = 0.999
AA = −0.0264 ± 0.00211
r² = 1
r² = 0.871
AA = 0.0525 ± 0.01023
r² = 0.858
30
20
10
0.0
0.5
1.0
1.5
A [μmol m⁻² s⁻¹]
g sw [mol m⁻² s⁻¹]
Curve type
amphi
pseudohypo
Measurement
high
low

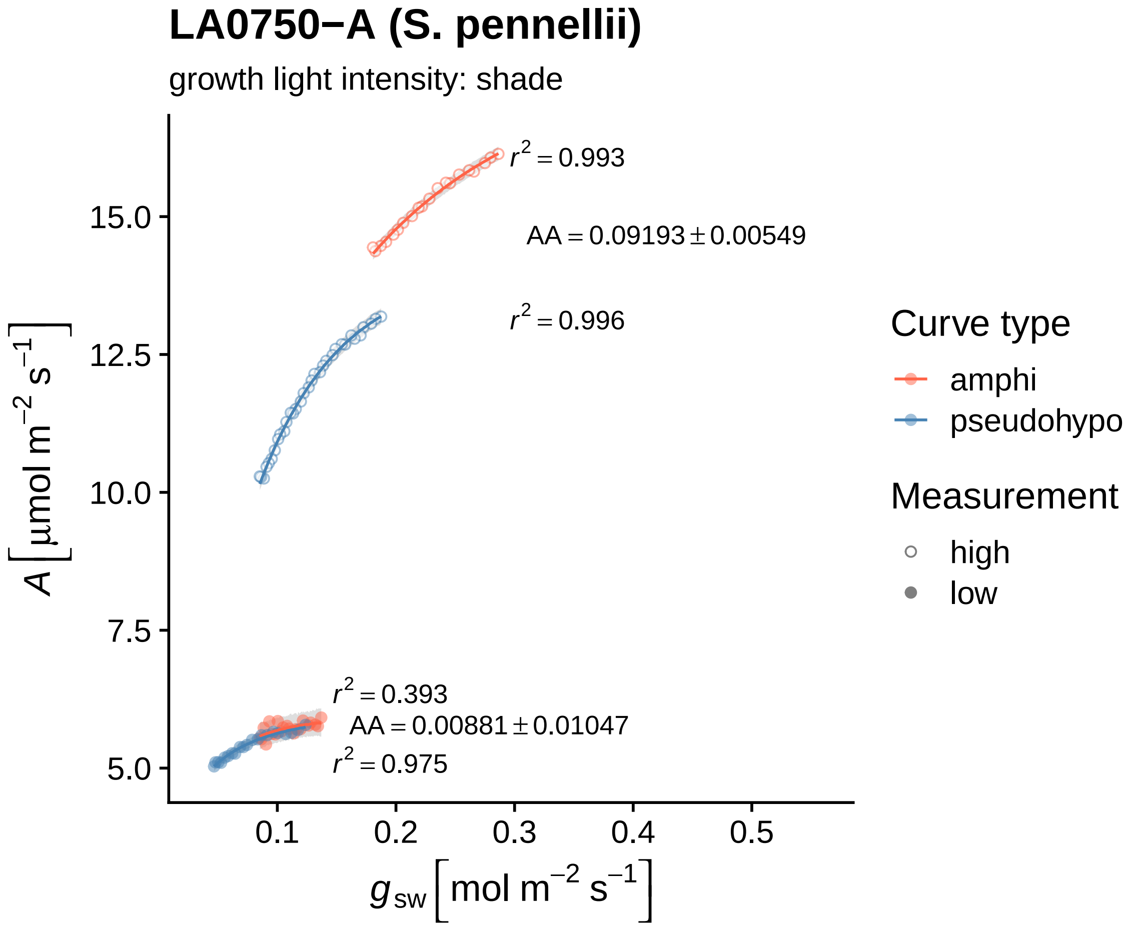

LA0750−A (S. pennellii)
growth light intensity: shade
$r^2 = 0.993$
$AA = 0.09193 \pm 0.00549$
$r^2 = 0.996$
$r^2 = 0.393$
$AA = 0.00881 \pm 0.01047$
$r^2 = 0.975$
15.0
12.5
10.0
7.5
5.0
0.1
0.2
0.3
0.4
0.5
$A\left[\mu mol\ m^{-2}\ s^{-1}\right]$
$g_{sw}\left[mol\ m^{-2}\ s^{-1}\right]$
Curve type
amphi
pseudohypo
Measurement
high
low

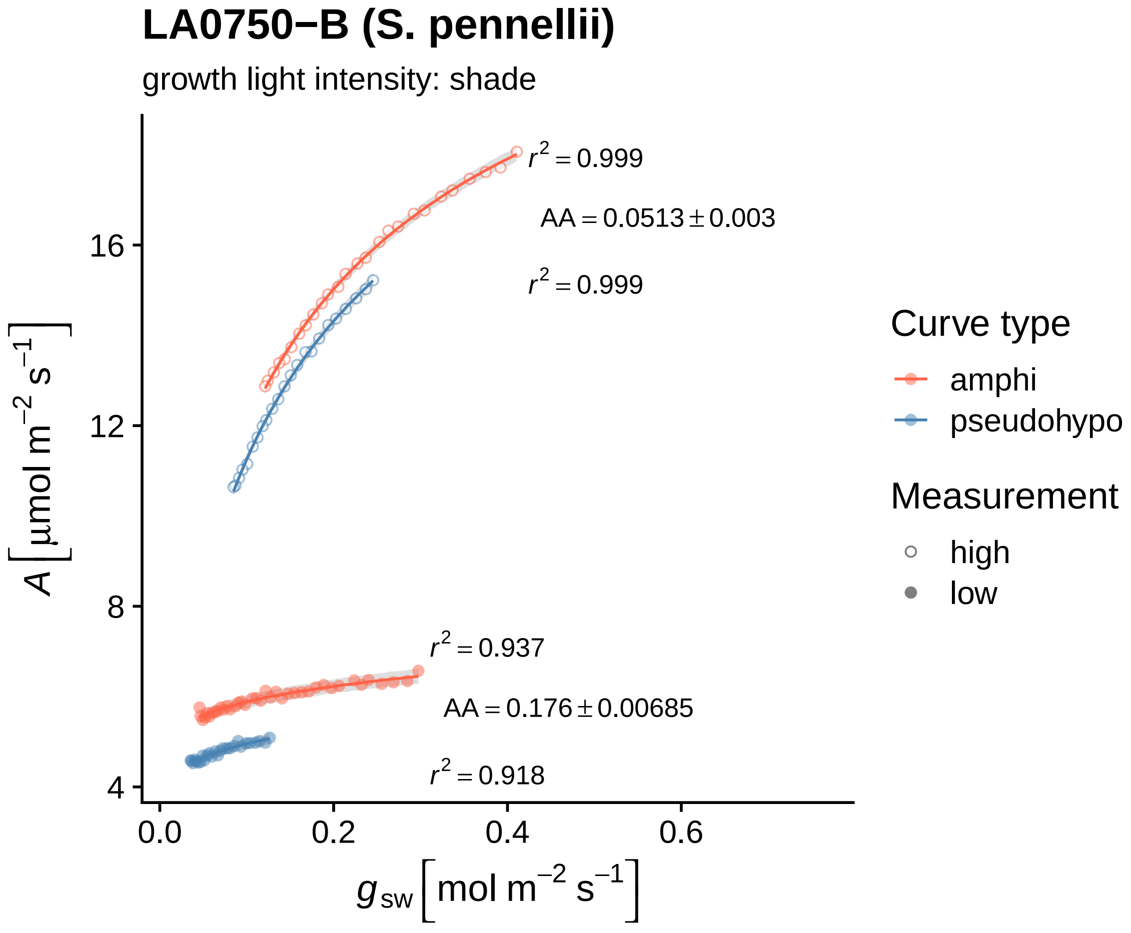

LA0750−B (S. pennellii)
growth light intensity: shade
$r^2 = 0.999$
$AA = 0.0513 \pm 0.003$
$r^2 = 0.999$
$r^2 = 0.937$
$AA = 0.176 \pm 0.00685$
$r^2 = 0.918$
$A\,[\mu mol\ m^{-2}\ s^{-1}]$
$g_{sw}\,[mol\ m^{-2}\ s^{-1}]$
16
12
8
4
0.0
0.2
0.4
0.6
Curve type
amphi
pseudohypo
Measurement
high
low

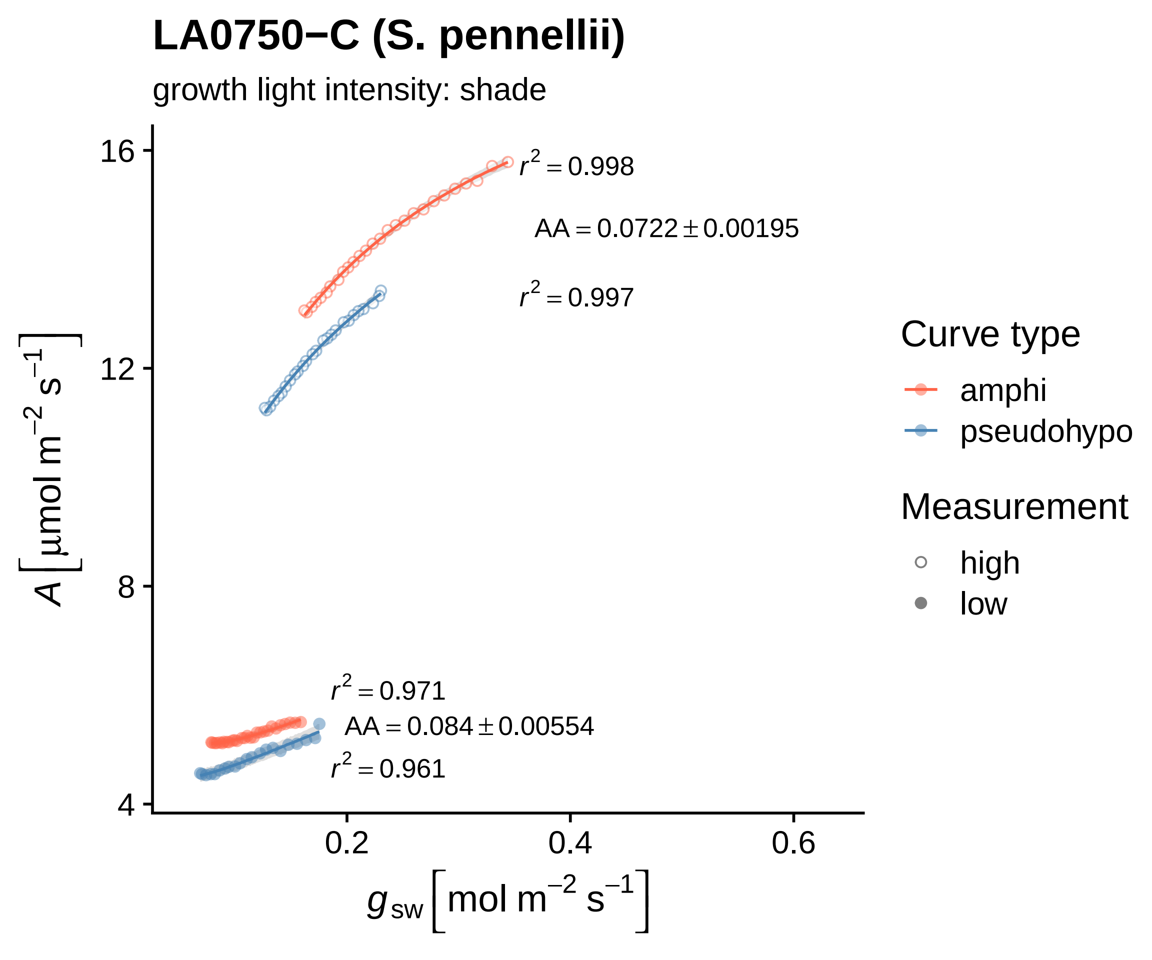

LA0750−C (S. pennellii)
growth light intensity: shade
16
12
8
4
0.2
0.4
0.6
$A\left[\mu mol\ m^{-2}\ s^{-1}\right]$
$g_{sw}\left[mol\ m^{-2}\ s^{-1}\right]$
$r^2 = 0.998$
$AA = 0.0722 \pm 0.00195$
$r^2 = 0.997$
$r^2 = 0.971$
$AA = 0.084 \pm 0.00554$
$r^2 = 0.961$
Curve type
amphi
pseudohypo
Measurement
high
low

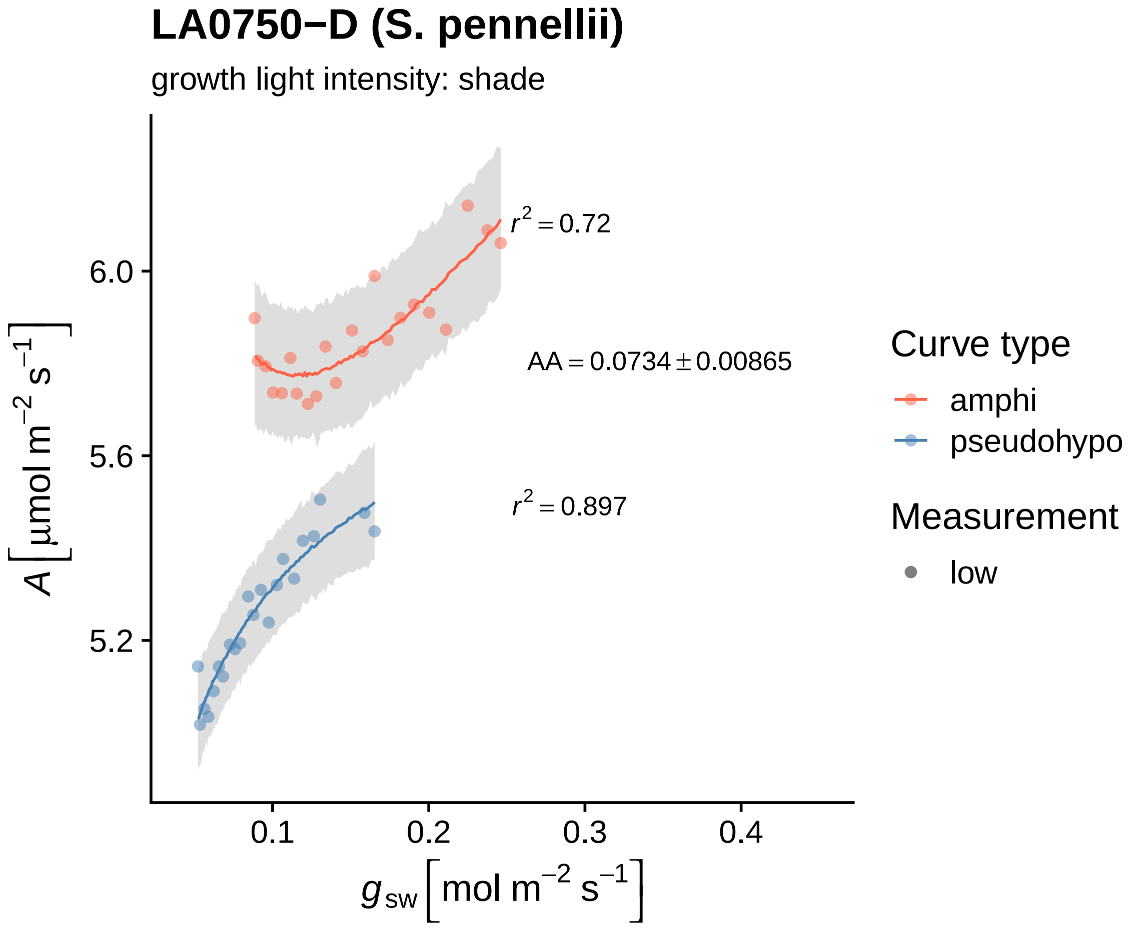

LA0750−D (S. pennellii)
growth light intensity: shade
$r^2 = 0.72$
AA = 0.0734 ± 0.00865
$r^2 = 0.897$
6.0
5.6
5.2
0.1
0.2
0.3
0.4
$A\ [\mu mol\ m^{-2}\ s^{-1}]$
$g_{sw}\ [mol\ m^{-2}\ s^{-1}]$
Curve type
amphi
pseudohypo
Measurement
low

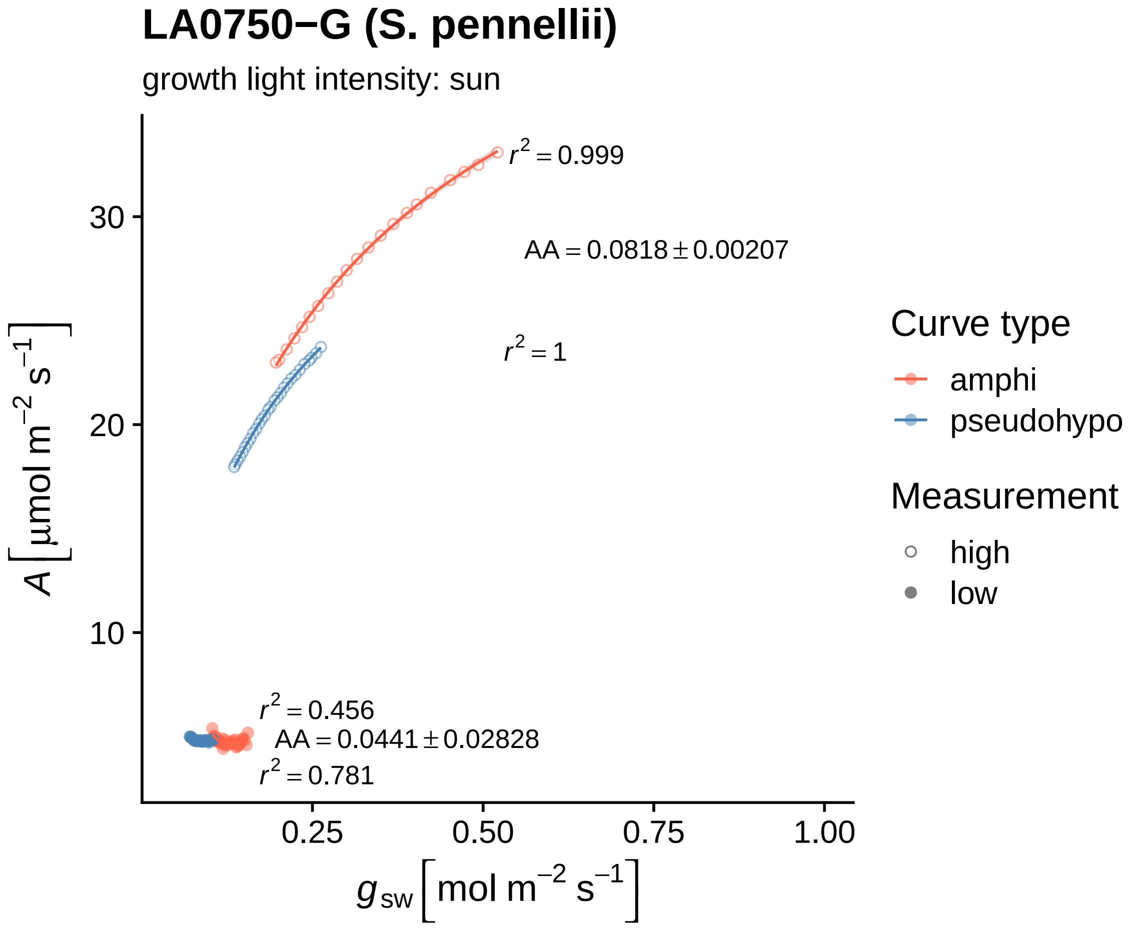

LA0750−G (S. pennellii)
growth light intensity: sun
$r^2 = 0.999$
$AA = 0.0818 \pm 0.00207$
$r^2 = 1$
$r^2 = 0.456$
$AA = 0.0441 \pm 0.02828$
$r^2 = 0.781$
30
20
10
0.25
0.50
0.75
1.00
$A\ [\mu mol\ m^{-2}\ s^{-1}]$
$g_{sw}\ [mol\ m^{-2}\ s^{-1}]$
Curve type
amphi
pseudohypo
Measurement
high
low

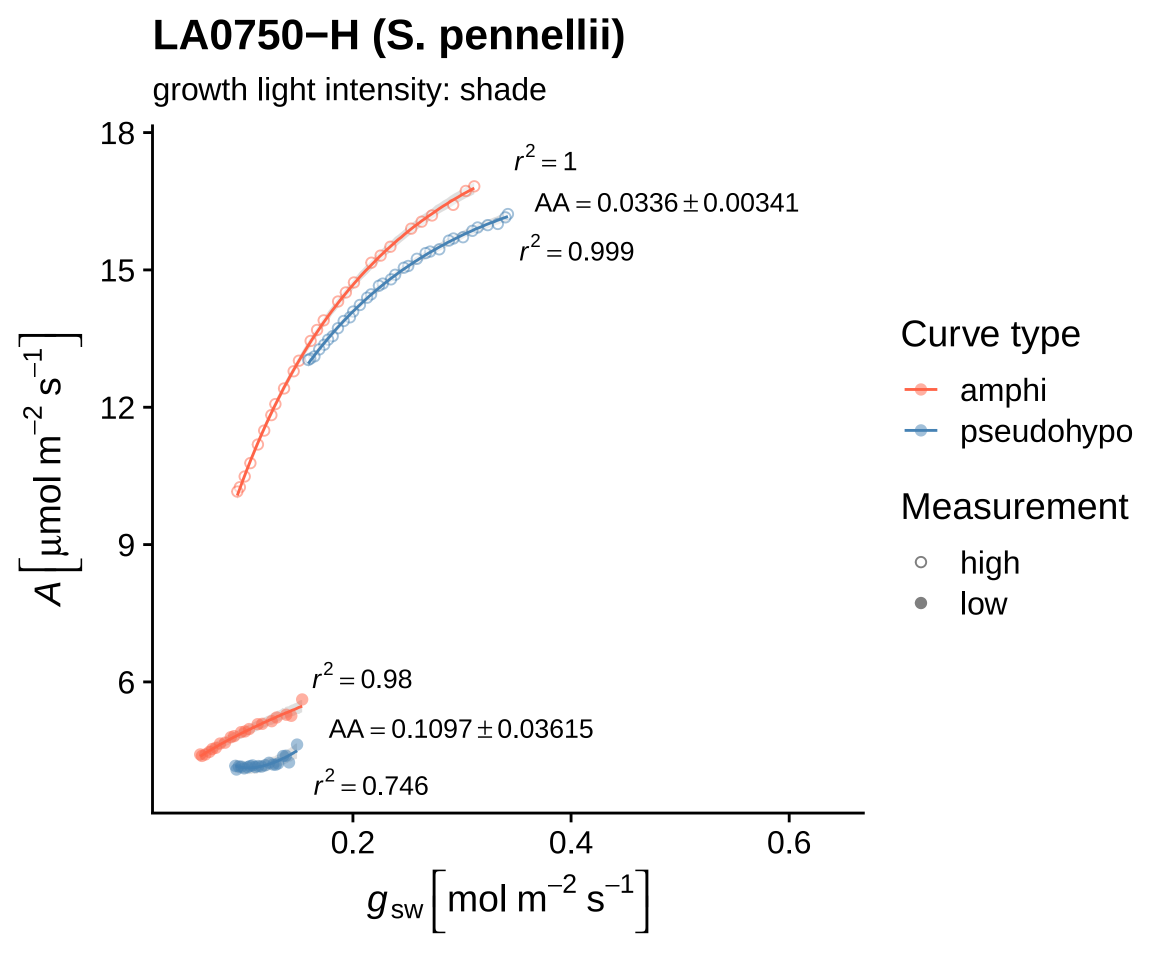
LA0750−H (S. pennellii)
growth light intensity: shade
$r^2 = 1$
$AA = 0.0336 \pm 0.00341$
$r^2 = 0.999$
$r^2 = 0.98$
$AA = 0.1097 \pm 0.03615$
$r^2 = 0.746$
$A\ [\mu mol\ m^{-2}\ s^{-1}]$
$g_{sw}\ [mol\ m^{-2}\ s^{-1}]$
18
15
12
9
6
0.2
0.4
0.6
Curve type
amphi
pseudohypo
Measurement
high
low

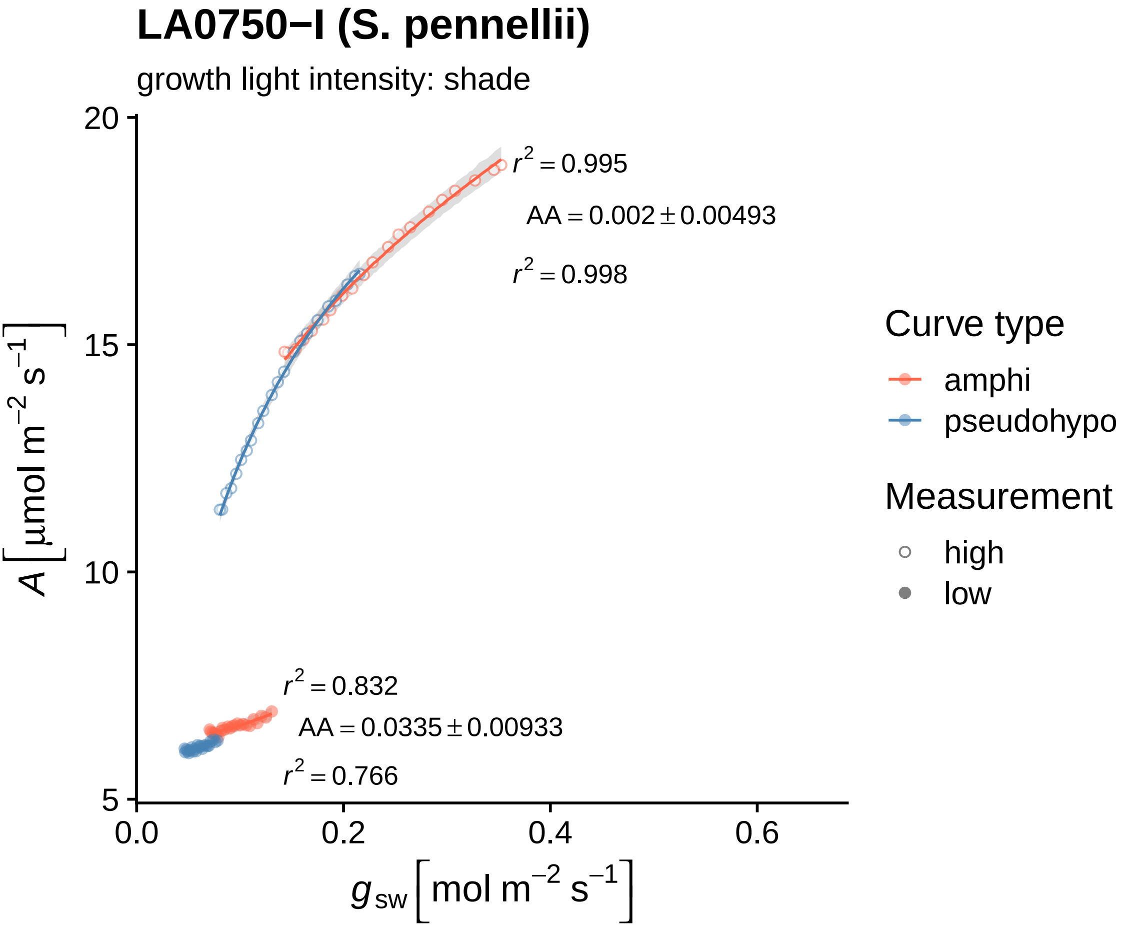
LA0750−I (S. pennellii)
growth light intensity: shade
$r^2 = 0.995$
$AA = 0.002 \pm 0.00493$
$r^2 = 0.998$
$r^2 = 0.832$
$AA = 0.0335 \pm 0.00933$
$r^2 = 0.766$
$A\left[\mu mol\ m^{-2}\ s^{-1}\right]$
$g_{sw}\left[mol\ m^{-2}\ s^{-1}\right]$
20
15
10
5
0.0
0.2
0.4
0.6
Curve type
amphi
pseudohypo
Measurement
high
low

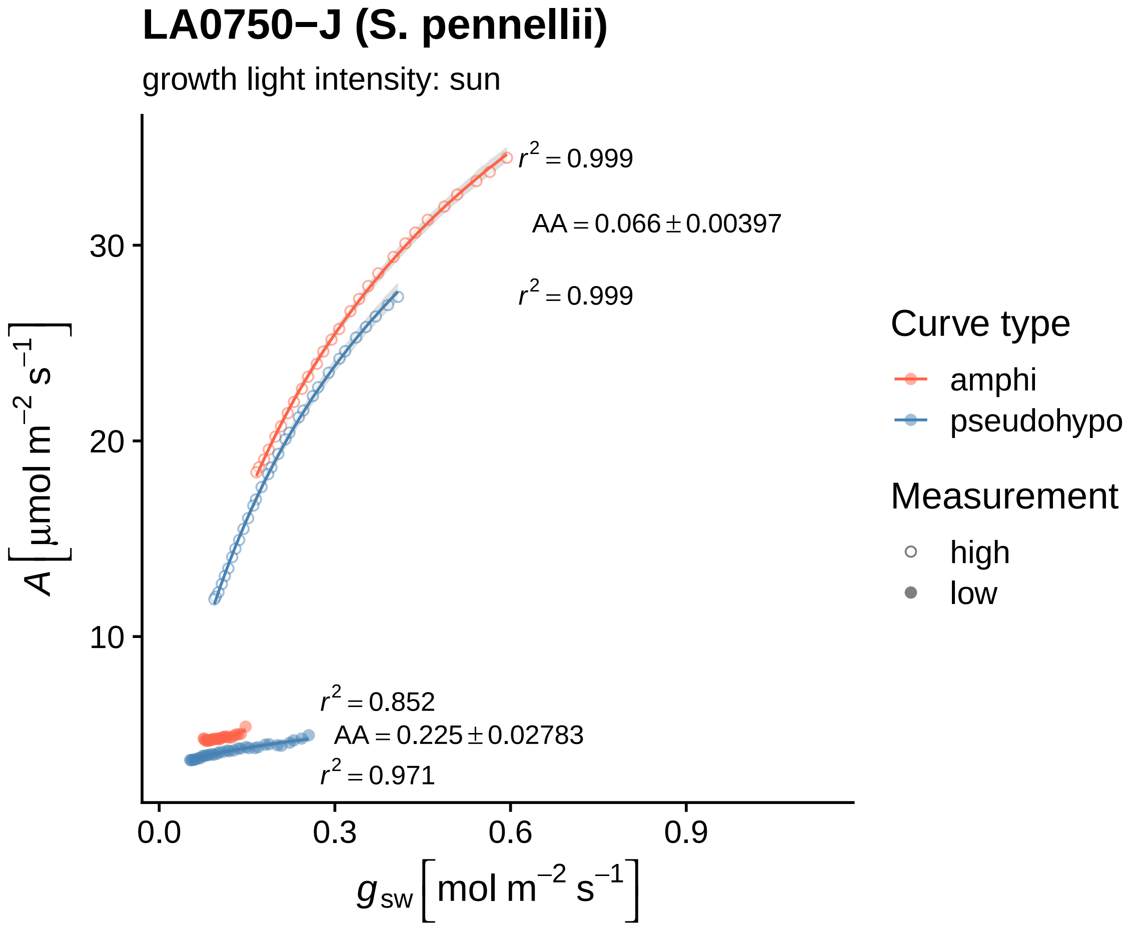
LA0750−J (S. pennellii)
growth light intensity: sun
$r^2 = 0.999$
$AA = 0.066 \pm 0.00397$
$r^2 = 0.999$
$r^2 = 0.852$
$AA = 0.225 \pm 0.02783$
$r^2 = 0.971$
30
20
10
0.0
0.3
0.6
0.9
$A\left[\mu mol\ m^{-2}\ s^{-1}\right]$
$g_{sw}\left[mol\ m^{-2}\ s^{-1}\right]$
Curve type
amphi
pseudohypo
Measurement
high
low

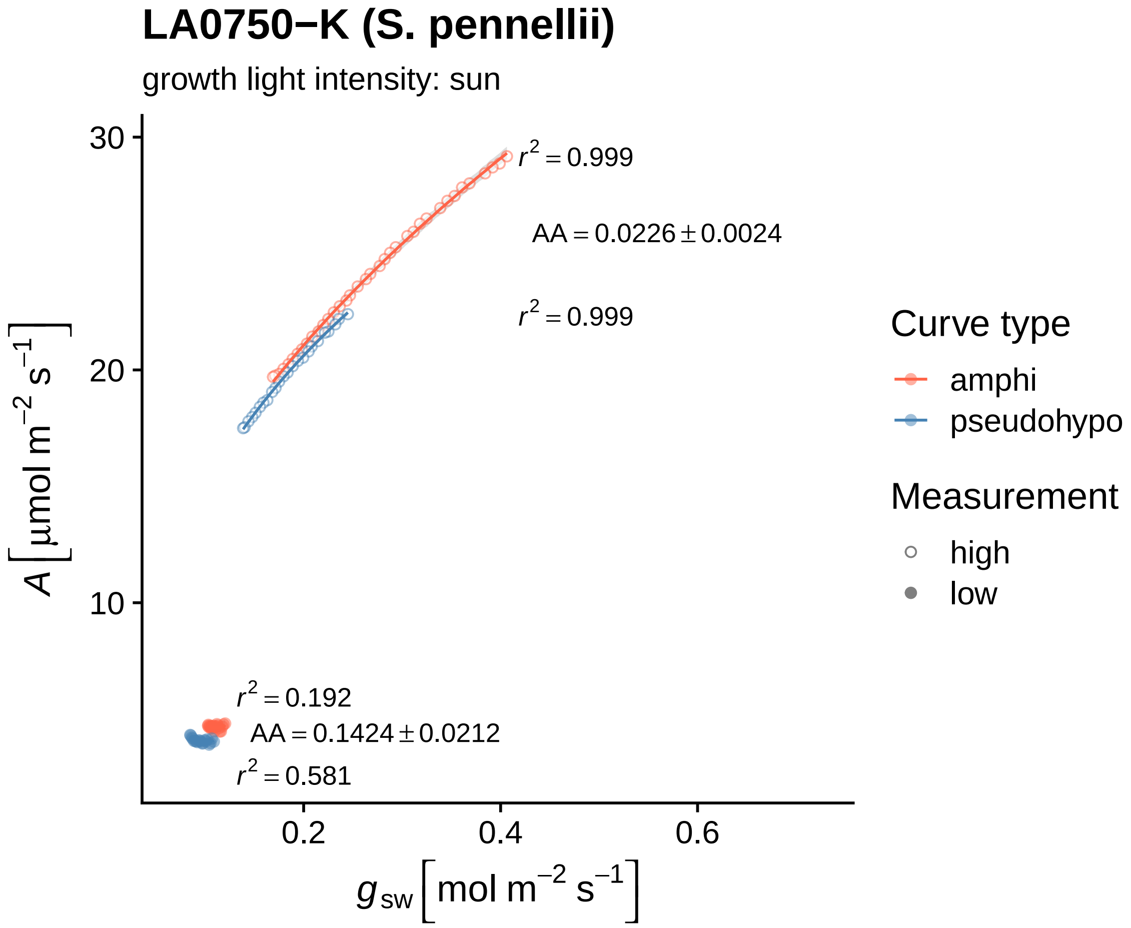
LA0750−K (S. pennellii)
growth light intensity: sun
$r^2 = 0.999$
$AA = 0.0226 \pm 0.0024$
$r^2 = 0.999$
$r^2 = 0.192$
$AA = 0.1424 \pm 0.0212$
$r^2 = 0.581$
30
20
10
0.2
0.4
0.6
$A\left[\mu mol\ m^{-2}\ s^{-1}\right]$
$g_{sw}\left[mol\ m^{-2}\ s^{-1}\right]$
Curve type
amphi
pseudohypo
Measurement
high
low

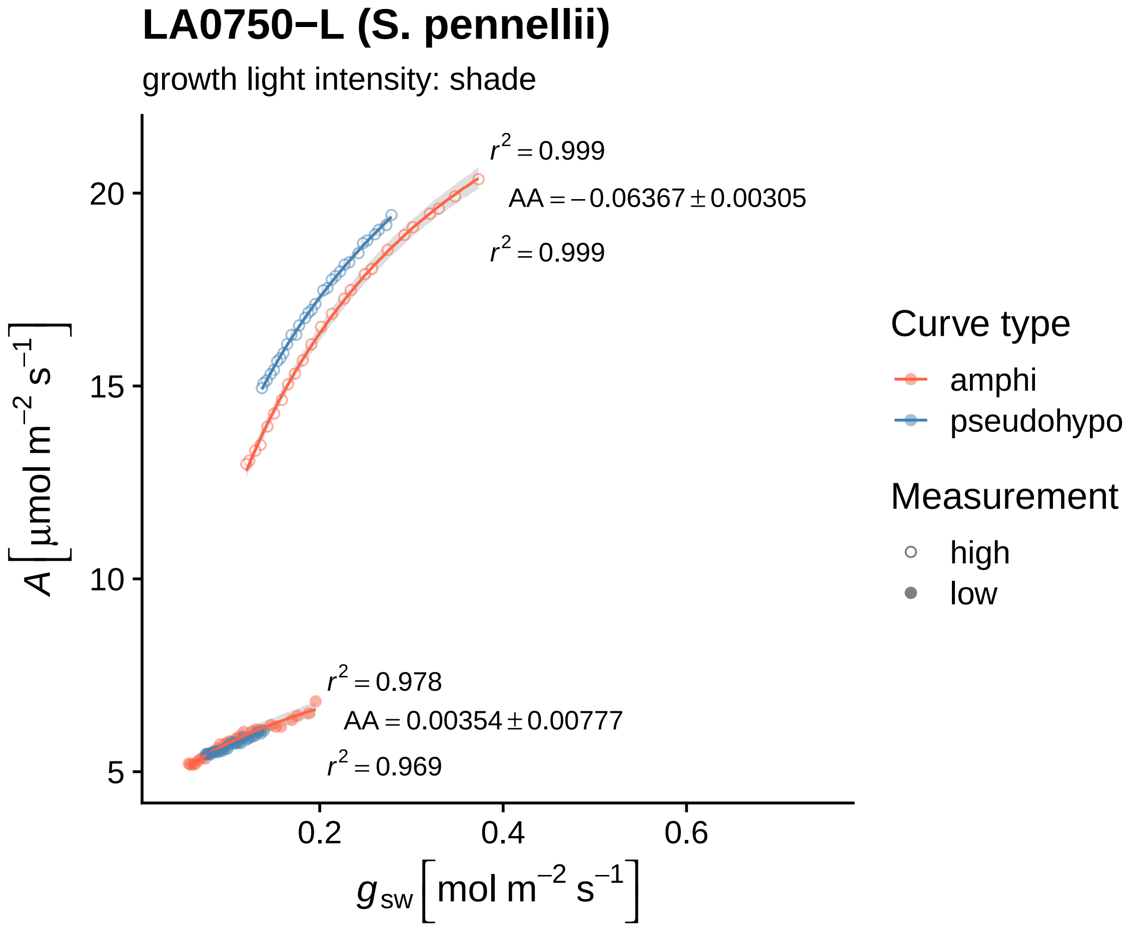

LA0750−L (S. pennellii)
growth light intensity: shade
$r^2 = 0.999$
$AA = -0.06367 \pm 0.00305$
$r^2 = 0.999$
$r^2 = 0.978$
$AA = 0.00354 \pm 0.00777$
$r^2 = 0.969$
20
15
10
5
0.2
0.4
0.6
$A\,[\mu mol\ m^{-2}\ s^{-1}]$
$g_{sw}\,[mol\ m^{-2}\ s^{-1}]$
Curve type
amphi
pseudohypo
Measurement
high
low

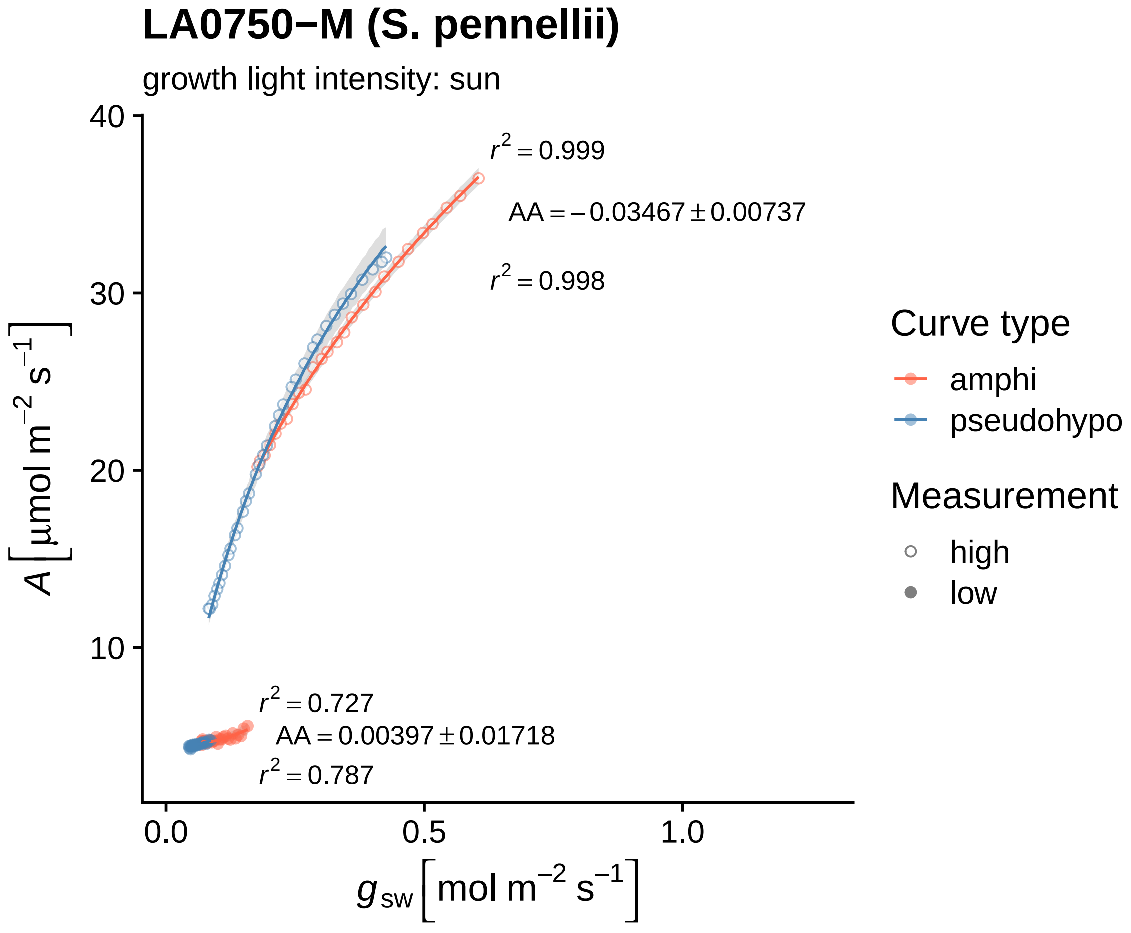

LA0750−M (S. pennellii)
growth light intensity: sun
$r^2 = 0.999$
$AA = -0.03467 \pm 0.00737$
$r^2 = 0.998$
$r^2 = 0.727$
$AA = 0.00397 \pm 0.01718$
$r^2 = 0.787$
40
30
20
10
0.0
0.5
1.0
$A\ [\mu mol\ m^{-2}\ s^{-1}]$
$g_{sw}\ [mol\ m^{-2}\ s^{-1}]$
Curve type
amphi
pseudohypo
Measurement
high
low

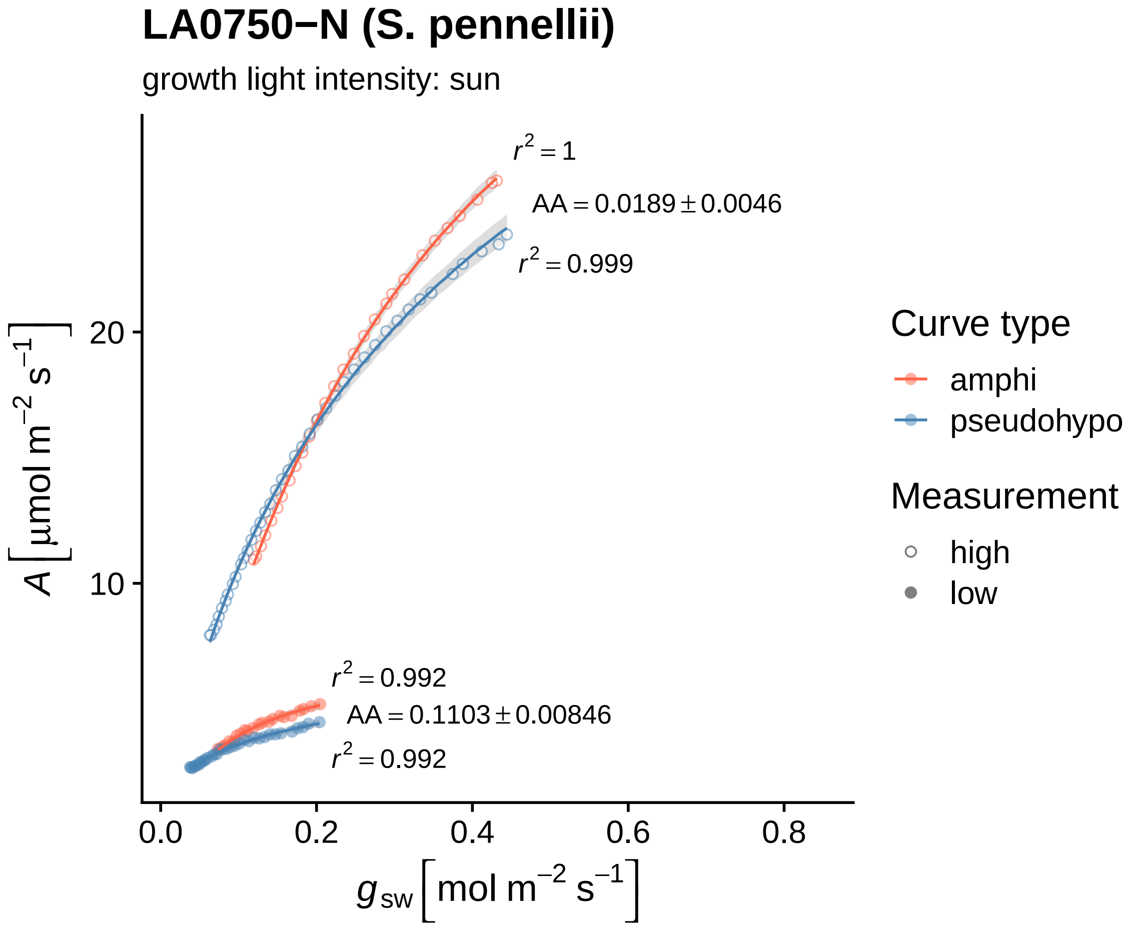
LA0750−N (S. pennellii)
growth light intensity: sun
$r^2 = 1$
$AA = 0.0189 \pm 0.0046$
$r^2 = 0.999$
$r^2 = 0.992$
$AA = 0.1103 \pm 0.00846$
$r^2 = 0.992$
20
10
0.0
0.2
0.4
0.6
0.8
$A\ [\mu mol\ m^{-2}\ s^{-1}]$
$g_{sw}\ [mol\ m^{-2}\ s^{-1}]$
Curve type
amphi
pseudohypo
Measurement
high
low

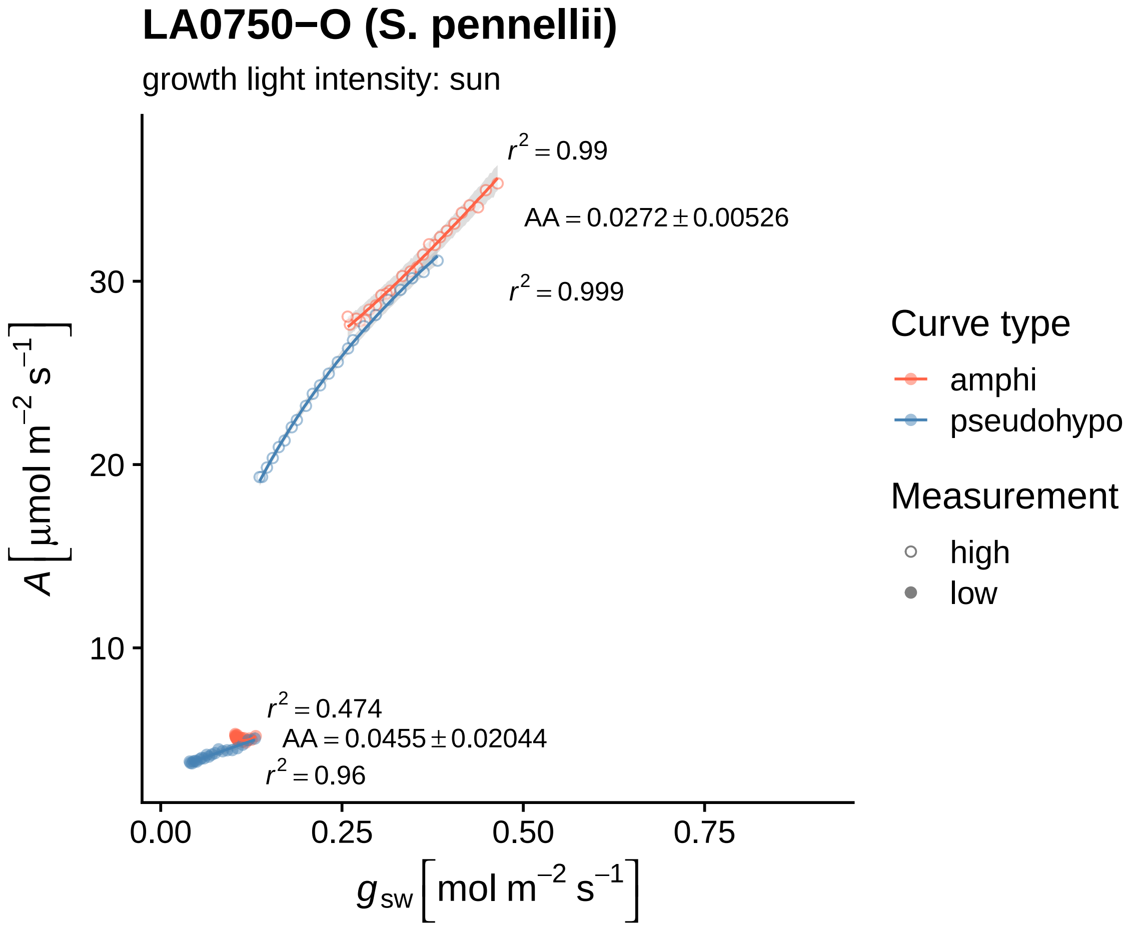

LA0750−O (S. pennellii)
growth light intensity: sun
$r^2 = 0.99$
$AA = 0.0272 \pm 0.00526$
$r^2 = 0.999$
$r^2 = 0.474$
$AA = 0.0455 \pm 0.02044$
$r^2 = 0.96$
30
20
10
0.00
0.25
0.50
0.75
$A\ [\mu mol\ m^{-2}\ s^{-1}]$
$g_{sw}\ [mol\ m^{-2}\ s^{-1}]$
Curve type
amphi
pseudohypo
Measurement
high
low

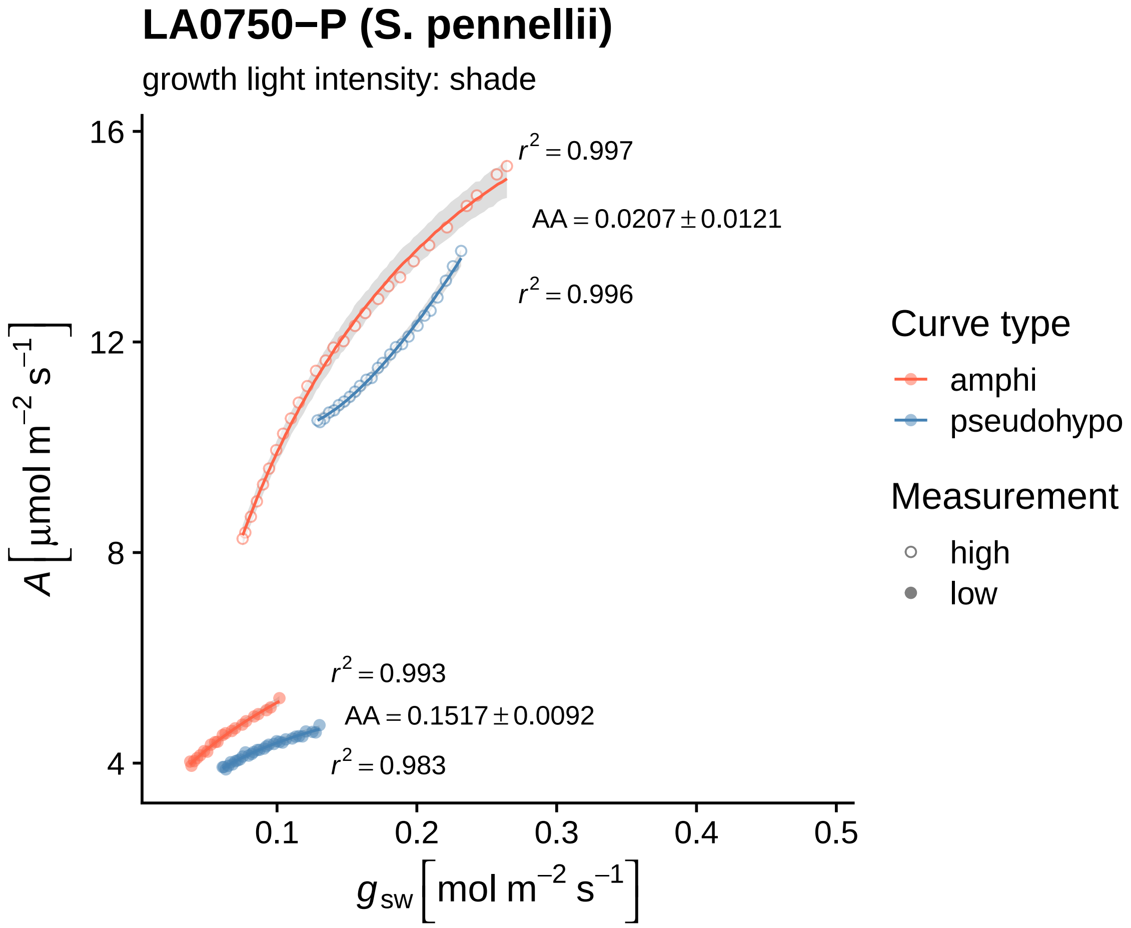

LA0750−P (S. pennellii)
growth light intensity: shade
$r^2 = 0.997$
$AA = 0.0207 \pm 0.0121$
$r^2 = 0.996$
$r^2 = 0.993$
$AA = 0.1517 \pm 0.0092$
$r^2 = 0.983$
16
12
8
4
0.1
0.2
0.3
0.4
0.5
$A\ [\mu mol\ m^{-2}\ s^{-1}]$
$g_{sw}\ [mol\ m^{-2}\ s^{-1}]$
Curve type
amphi
pseudohypo
Measurement
high
low

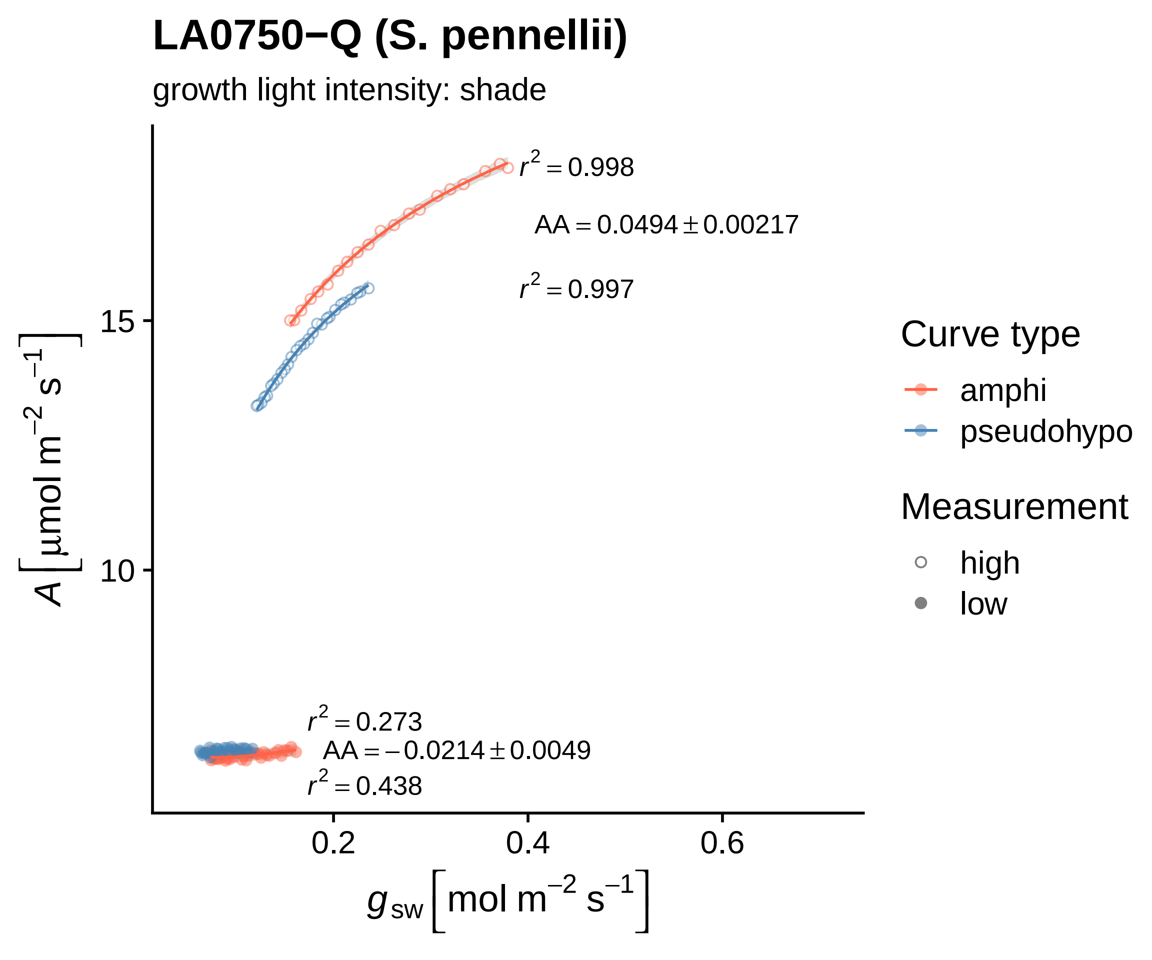
LA0750−Q (S. pennellii)
growth light intensity: shade
$r^2 = 0.998$
$AA = 0.0494 \pm 0.00217$
$r^2 = 0.997$
$r^2 = 0.273$
$AA = -0.0214 \pm 0.0049$
$r^2 = 0.438$
15
10
0.2
0.4
0.6
$A\,[\mu mol\ m^{-2}\ s^{-1}]$
$g_{sw}\,[mol\ m^{-2}\ s^{-1}]$
Curve type
amphi
pseudohypo
Measurement
high
low

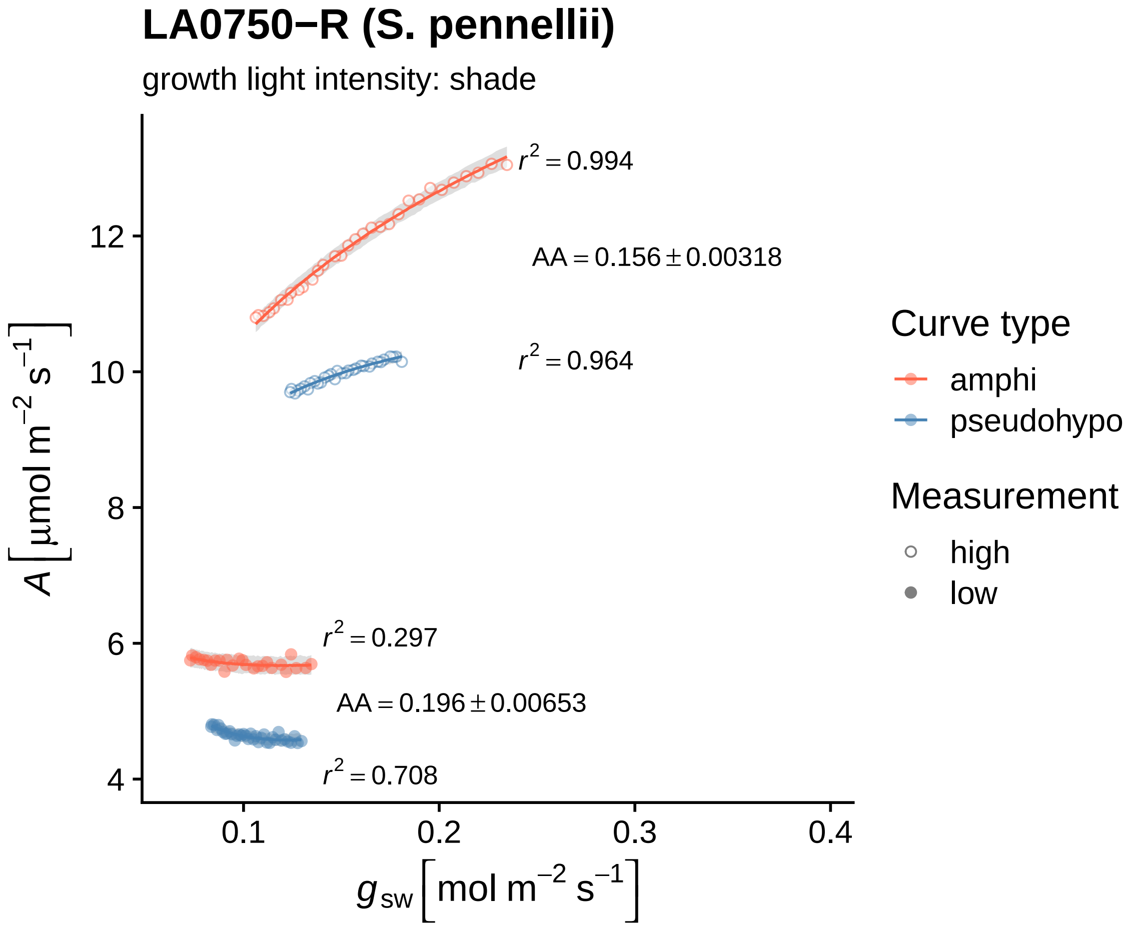

LA0750−R (S. pennellii)
growth light intensity: shade
$r^2 = 0.994$
$AA = 0.156 \pm 0.00318$
$r^2 = 0.964$
$r^2 = 0.297$
$AA = 0.196 \pm 0.00653$
$r^2 = 0.708$
$A\left[\mu mol\ m^{-2}\ s^{-1}\right]$
12
10
8
6
4
0.1
0.2
0.3
0.4
$g_{sw}\left[mol\ m^{-2}\ s^{-1}\right]$
Curve type
amphi
pseudohypo
Measurement
high
low

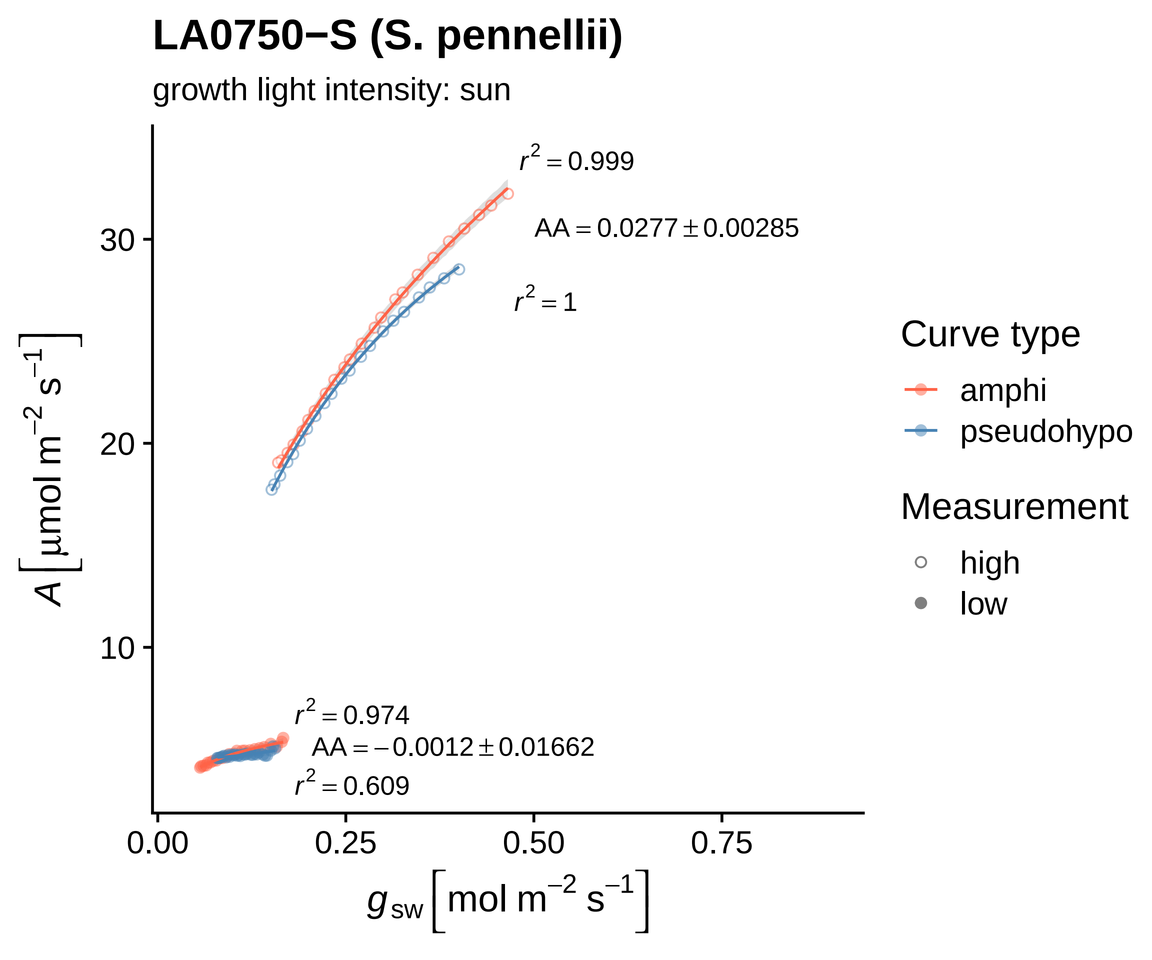

LA0750−S (S. pennellii)
growth light intensity: sun
$r^2 = 0.999$
$AA = 0.0277 \pm 0.00285$
$r^2 = 1$
$r^2 = 0.974$
$AA = -0.0012 \pm 0.01662$
$r^2 = 0.609$
30
20
10
0.00
0.25
0.50
0.75
$A\left[\mu mol\ m^{-2}\ s^{-1}\right]$
$g_{sw}\left[mol\ m^{-2}\ s^{-1}\right]$
Curve type
amphi
pseudohypo
Measurement
high
low

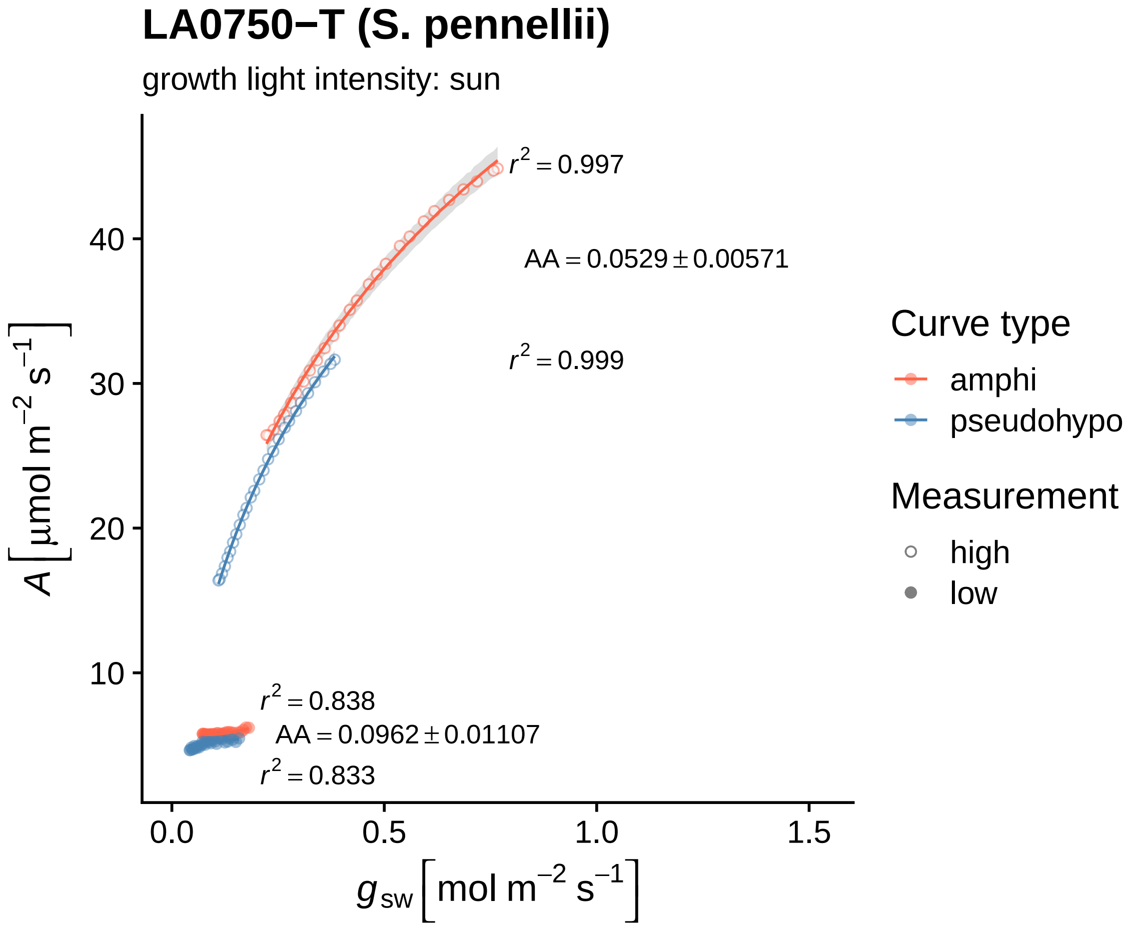

LA0750−T (S. pennellii)
growth light intensity: sun
r² = 0.997
AA = 0.0529 ± 0.00571
r² = 0.999
r² = 0.838
AA = 0.0962 ± 0.01107
r² = 0.833
A [μmol m⁻² s⁻¹]
40
30
20
10
0.0
0.5
1.0
1.5
g_sw [mol m⁻² s⁻¹]
Curve type
amphi
pseudohypo
Measurement
high
low

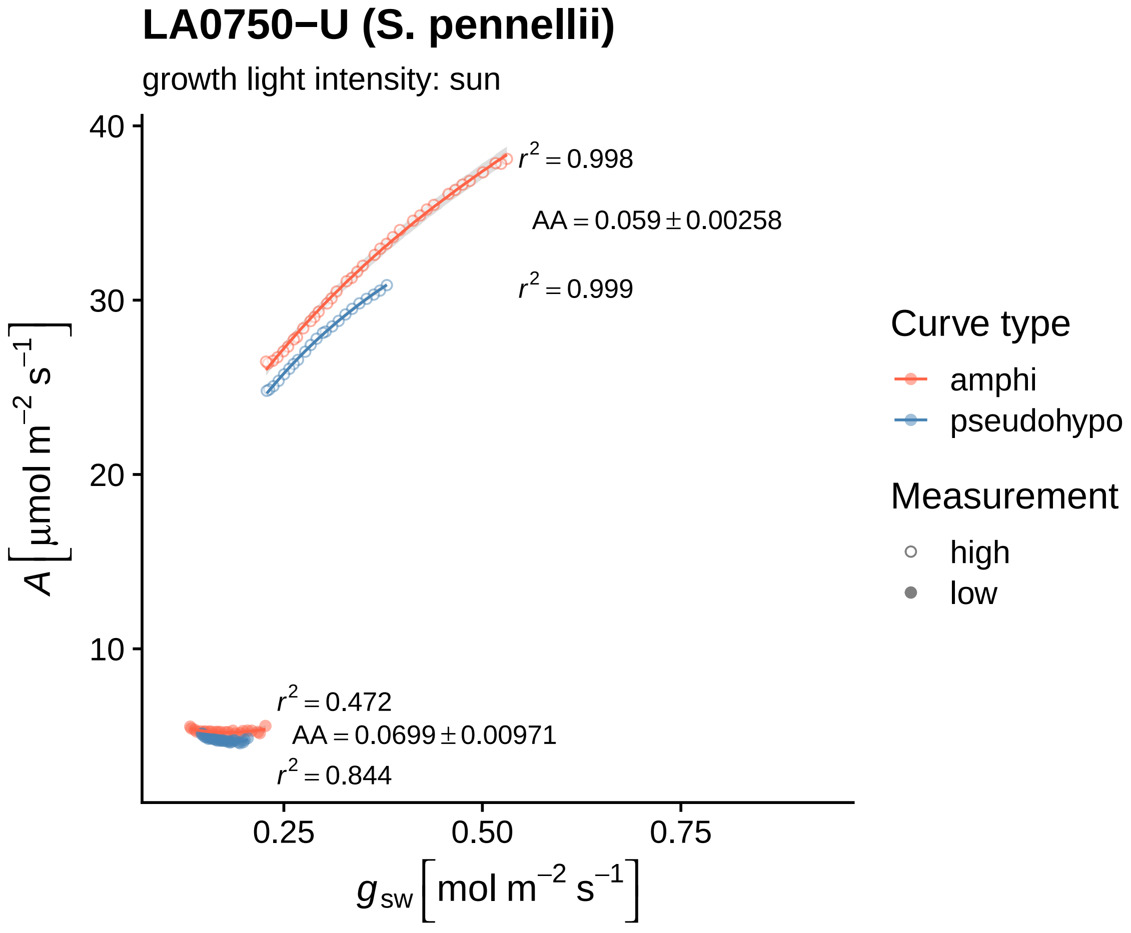

LA0750−U (S. pennellii)
growth light intensity: sun
$r^2 = 0.998$
$AA = 0.059 \pm 0.00258$
$r^2 = 0.999$
$r^2 = 0.472$
$AA = 0.0699 \pm 0.00971$
$r^2 = 0.844$
40
30
20
10
0.25
0.50
0.75
$A\,[\mu mol\ m^{-2}\ s^{-1}]$
$g_{sw}\,[mol\ m^{-2}\ s^{-1}]$
Curve type
amphi
pseudohypo
Measurement
high
low

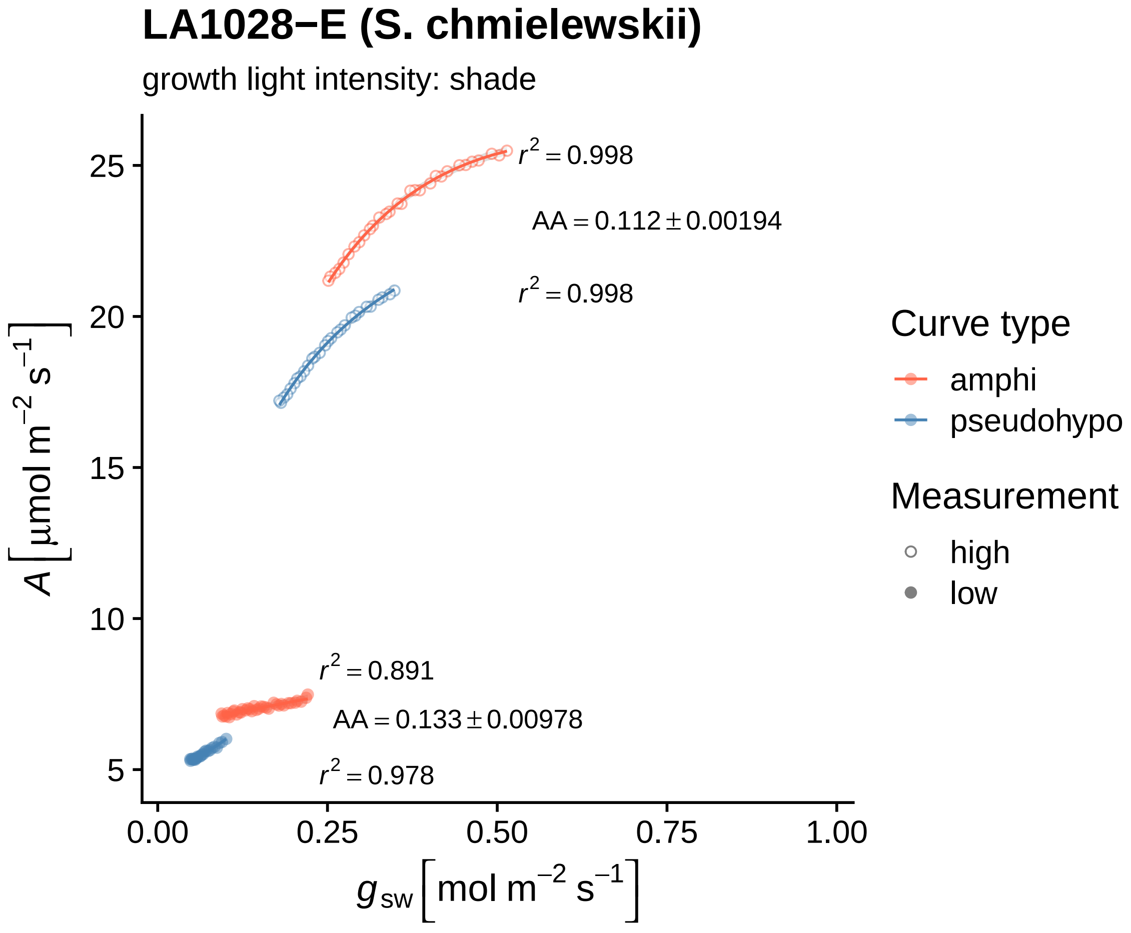
LA1028−E (S. chmielewskii)
growth light intensity: shade
$r^2 = 0.998$
AA = 0.112 ± 0.00194
$r^2 = 0.998$
$r^2 = 0.891$
AA = 0.133 ± 0.00978
$r^2 = 0.978$
25
20
15
10
5
0.00
0.25
0.50
0.75
1.00
$A\ [\mu mol\ m^{-2}\ s^{-1}]$
$g_{sw}\ [mol\ m^{-2}\ s^{-1}]$
Curve type
amphi
pseudohypo
Measurement
high
low

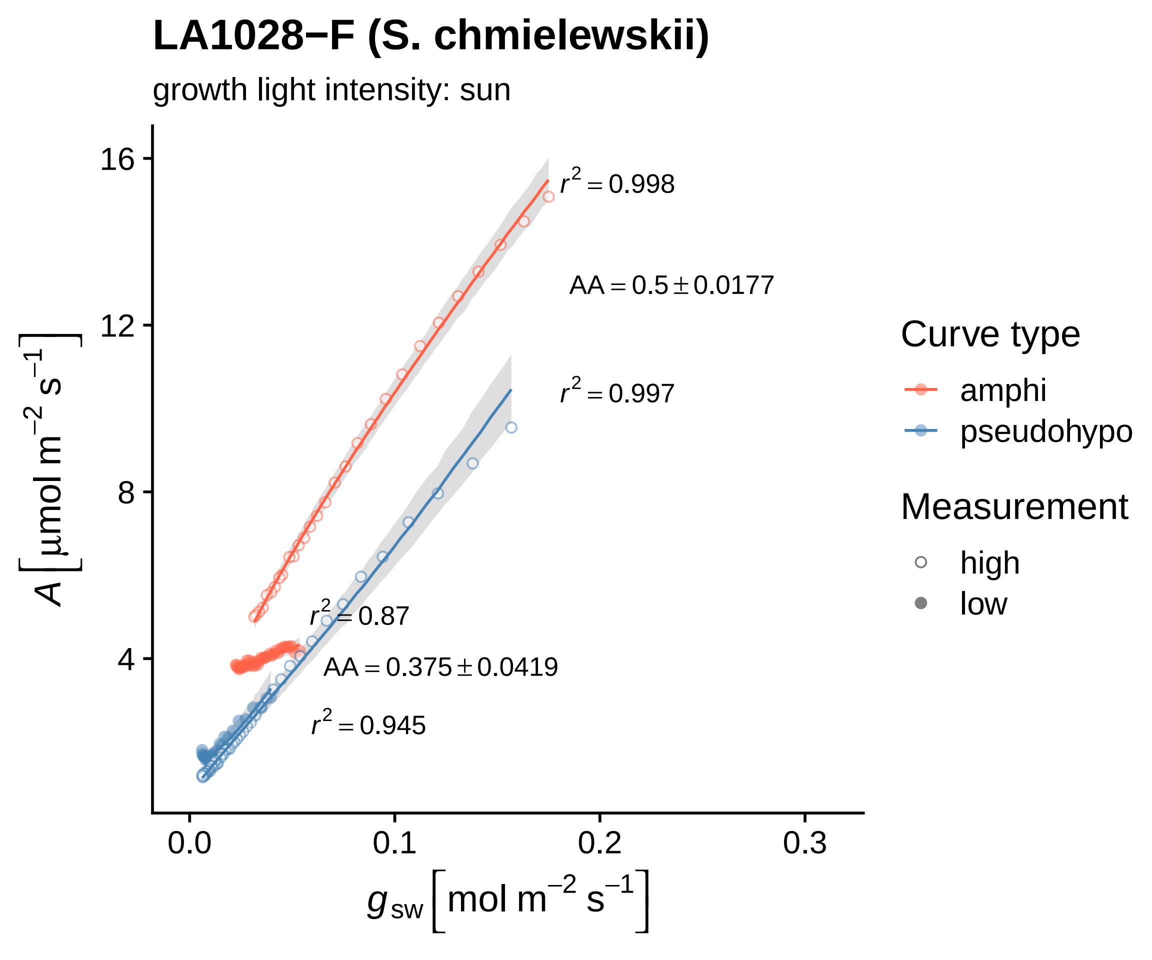

LA1028−F (S. chmielewskii)
growth light intensity: sun
$r^2 = 0.998$
$AA = 0.5 \pm 0.0177$
$r^2 = 0.997$
$r^2 = 0.87$
$AA = 0.375 \pm 0.0419$
$r^2 = 0.945$
16
12
8
4
0.0
0.1
0.2
0.3
$A\left[\mu mol\ m^{-2}\ s^{-1}\right]$
$g_{sw}\left[mol\ m^{-2}\ s^{-1}\right]$
Curve type
amphi
pseudohypo
Measurement
high
low

### LA1028−G (S. chmielewskii)

growth light intensity: shade

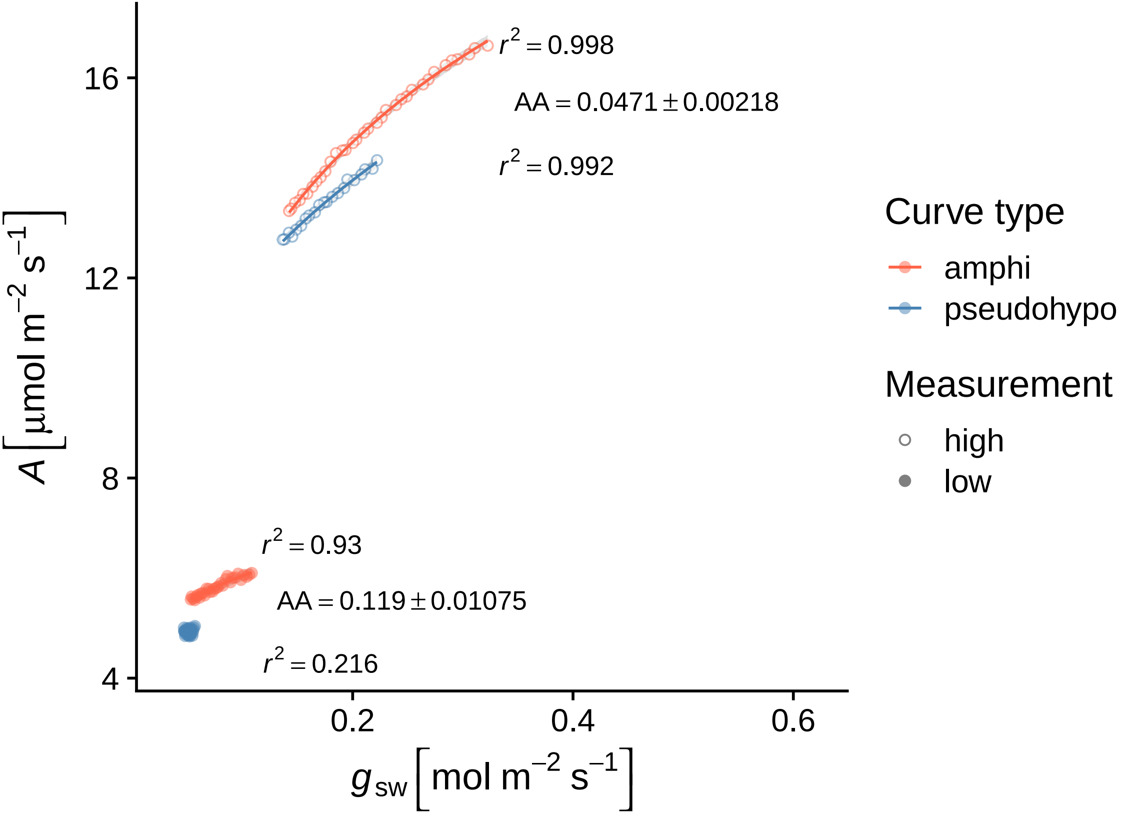

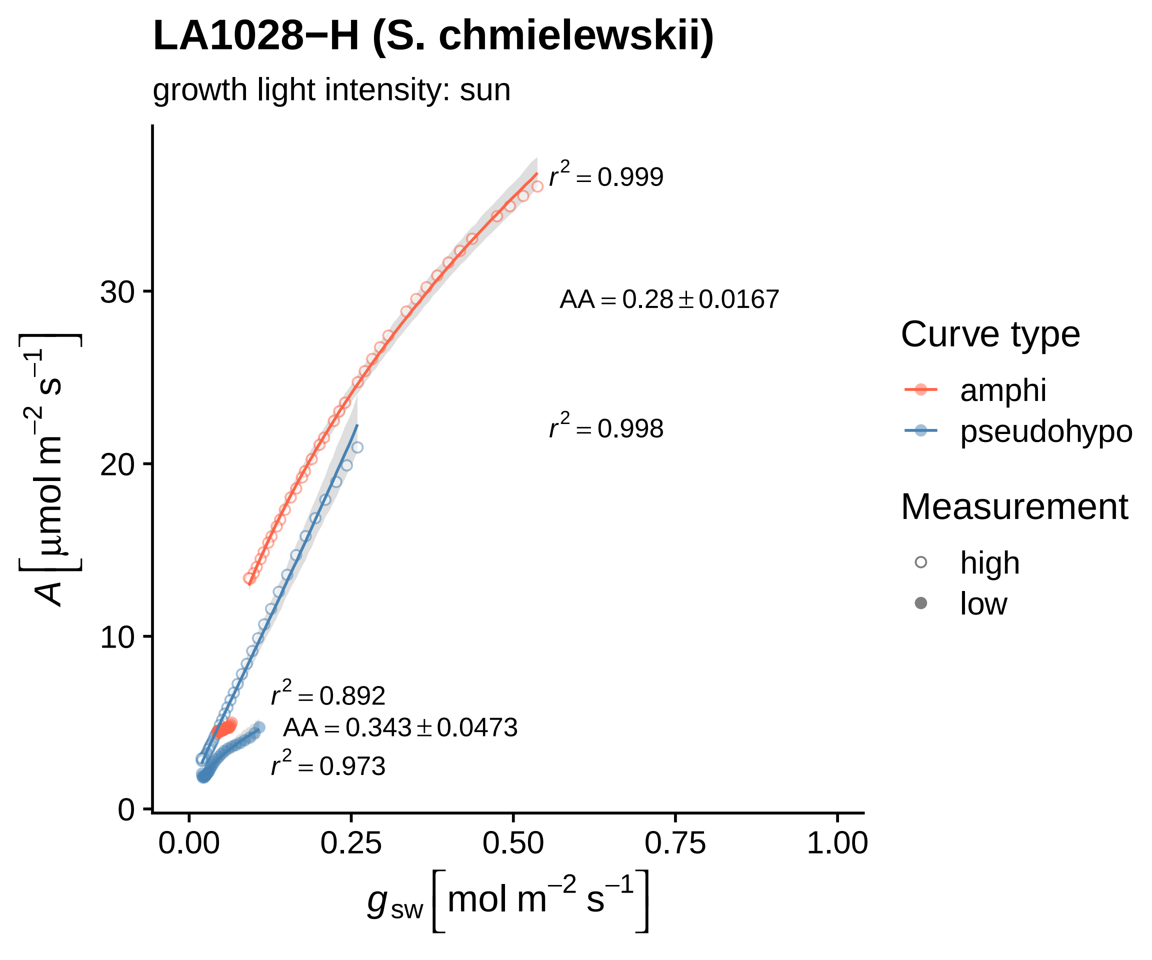
LA1028−H (S. chmielewskii)
growth light intensity: sun
r² = 0.999
AA = 0.28 ± 0.0167
r² = 0.998
r² = 0.892
AA = 0.343 ± 0.0473
r² = 0.973
A [μmol m⁻² s⁻¹]
0
10
20
30
0.00
0.25
0.50
0.75
1.00
g_sw [mol m⁻² s⁻¹]
Curve type
amphi
pseudohypo
Measurement
high
low

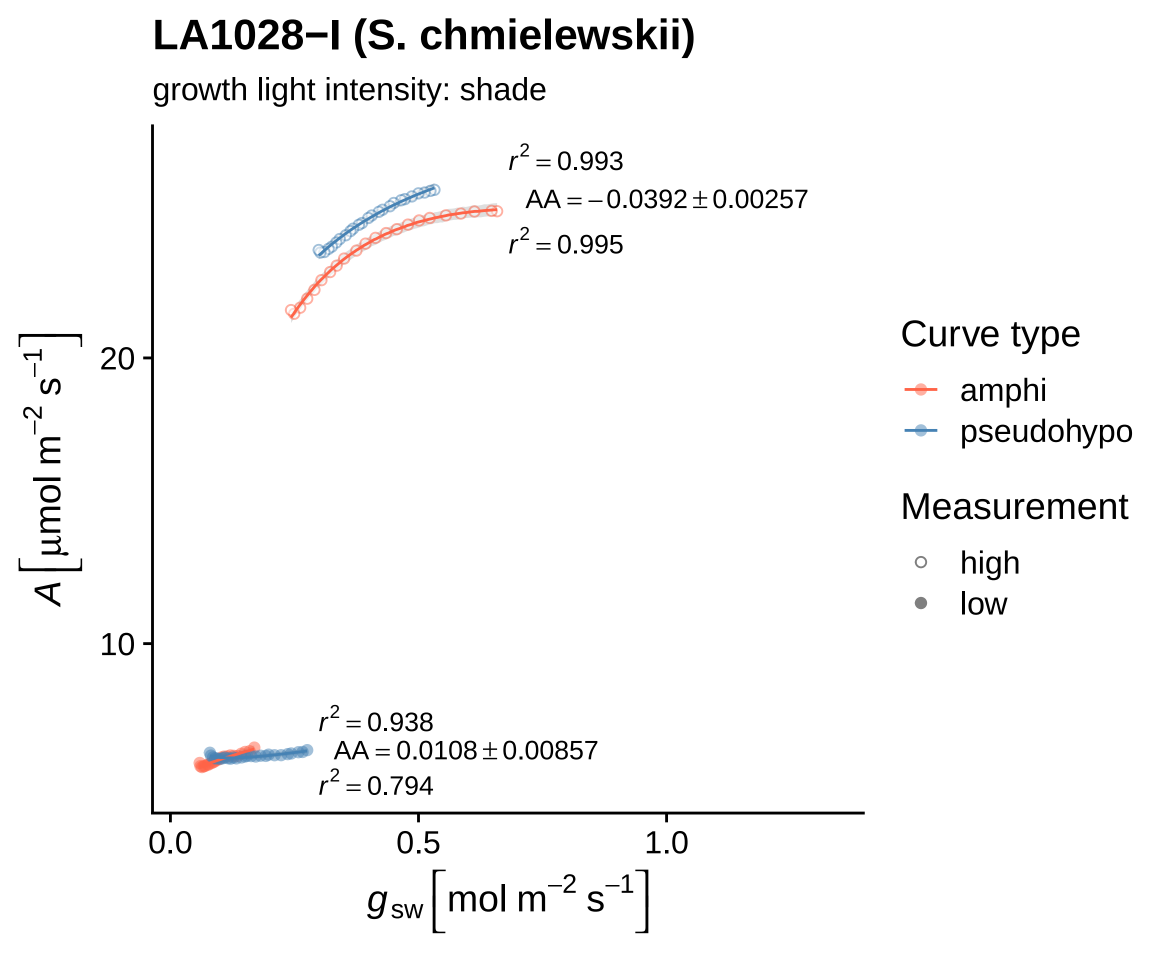

LA1028−I (S. chmielewskii)
growth light intensity: shade
r² = 0.993
AA = −0.0392 ± 0.00257
r² = 0.995
r² = 0.938
AA = 0.0108 ± 0.00857
r² = 0.794
A [µmol m⁻² s⁻¹]
10
20
0.0
0.5
1.0
g_sw [mol m⁻² s⁻¹]
Curve type
amphi
pseudohypo
Measurement
high
low

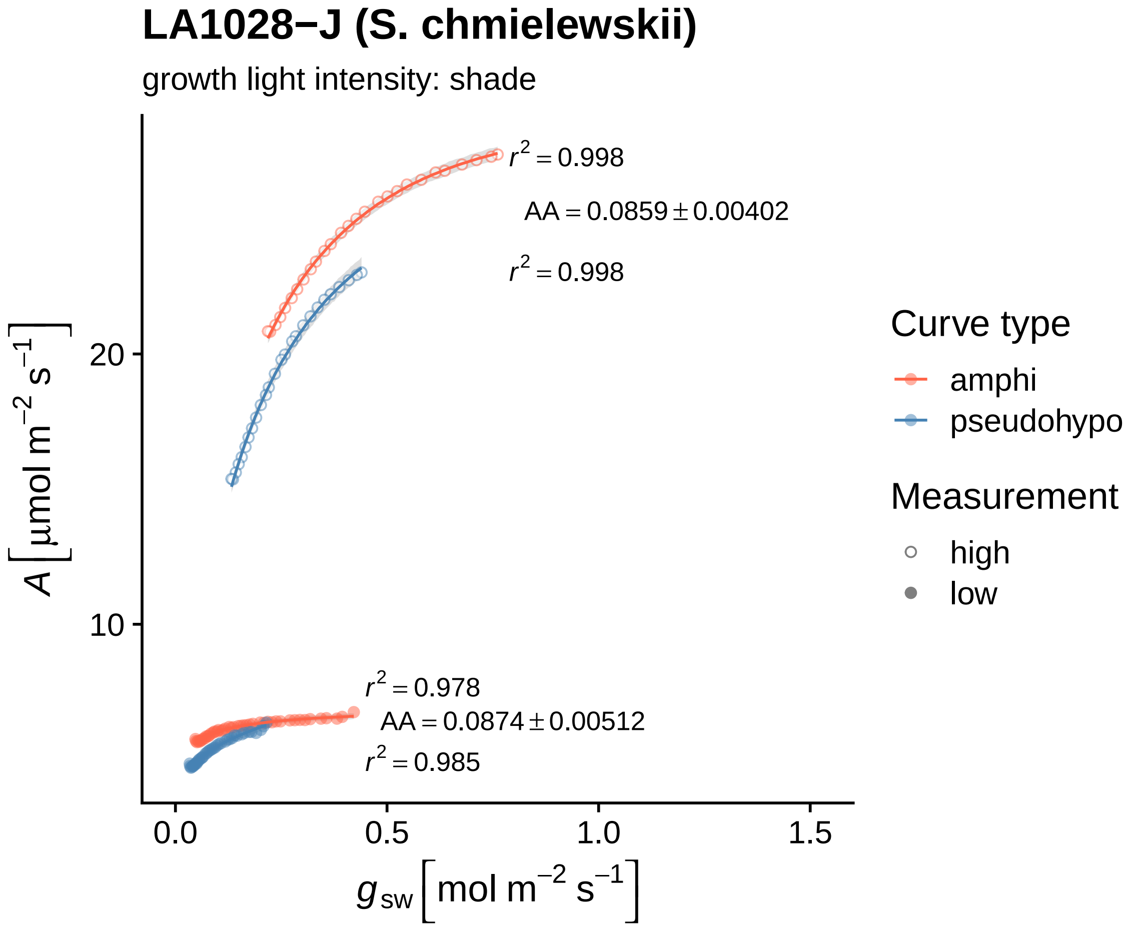
LA1028−J (S. chmielewskii)
growth light intensity: shade
r² = 0.998
AA = 0.0859 ± 0.00402
r² = 0.998
r² = 0.978
AA = 0.0874 ± 0.00512
r² = 0.985
A [μmol m⁻² s⁻¹]
20
10
0.0
0.5
1.0
1.5
g_sw [mol m⁻² s⁻¹]
Curve type
amphi
pseudohypo
Measurement
high
low

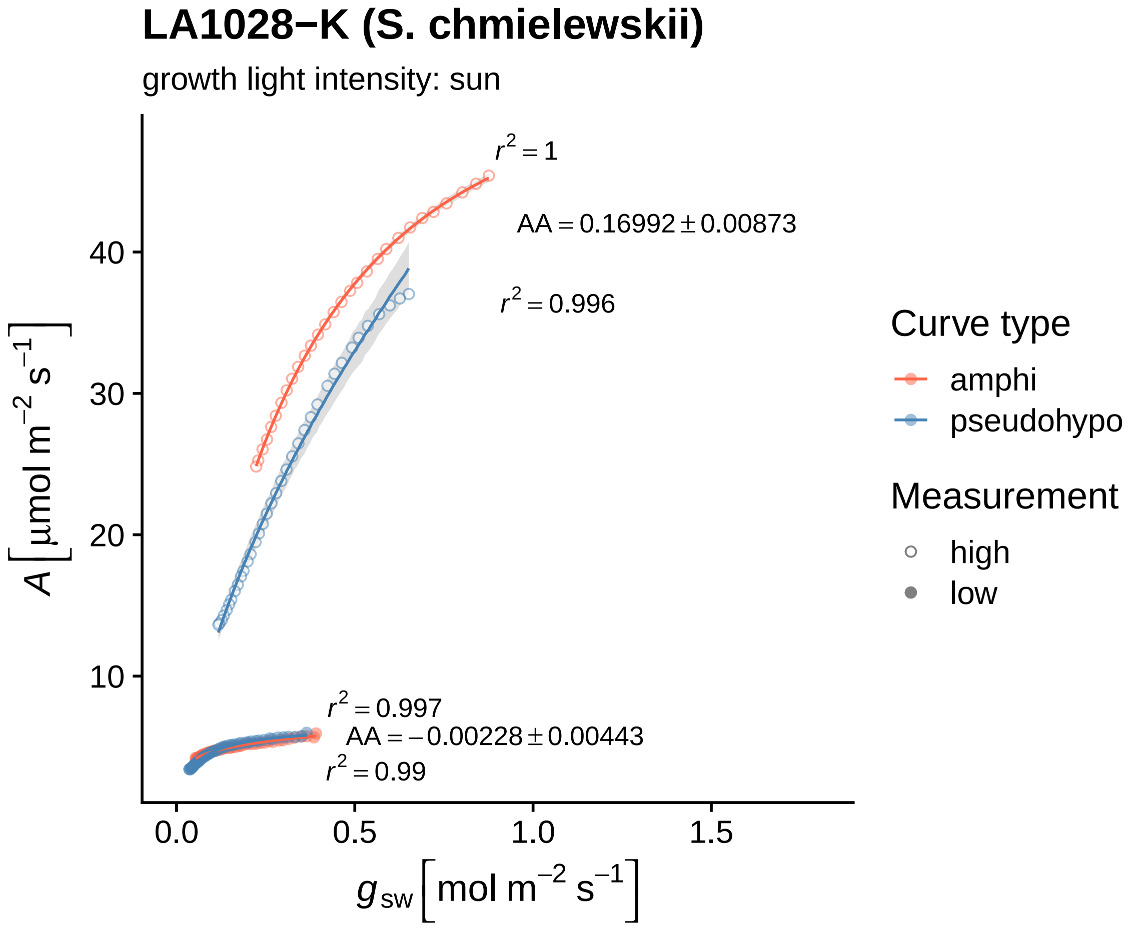

LA1028−K (S. chmielewskii)
growth light intensity: sun
r² = 1
AA = 0.16992 ± 0.00873
r² = 0.996
r² = 0.997
AA = −0.00228 ± 0.00443
r² = 0.99
A [µmol m⁻² s⁻¹]
10
20
30
40
0.0
0.5
1.0
1.5
g sw [mol m⁻² s⁻¹]
Curve type
amphi
pseudohypo
Measurement
high
low

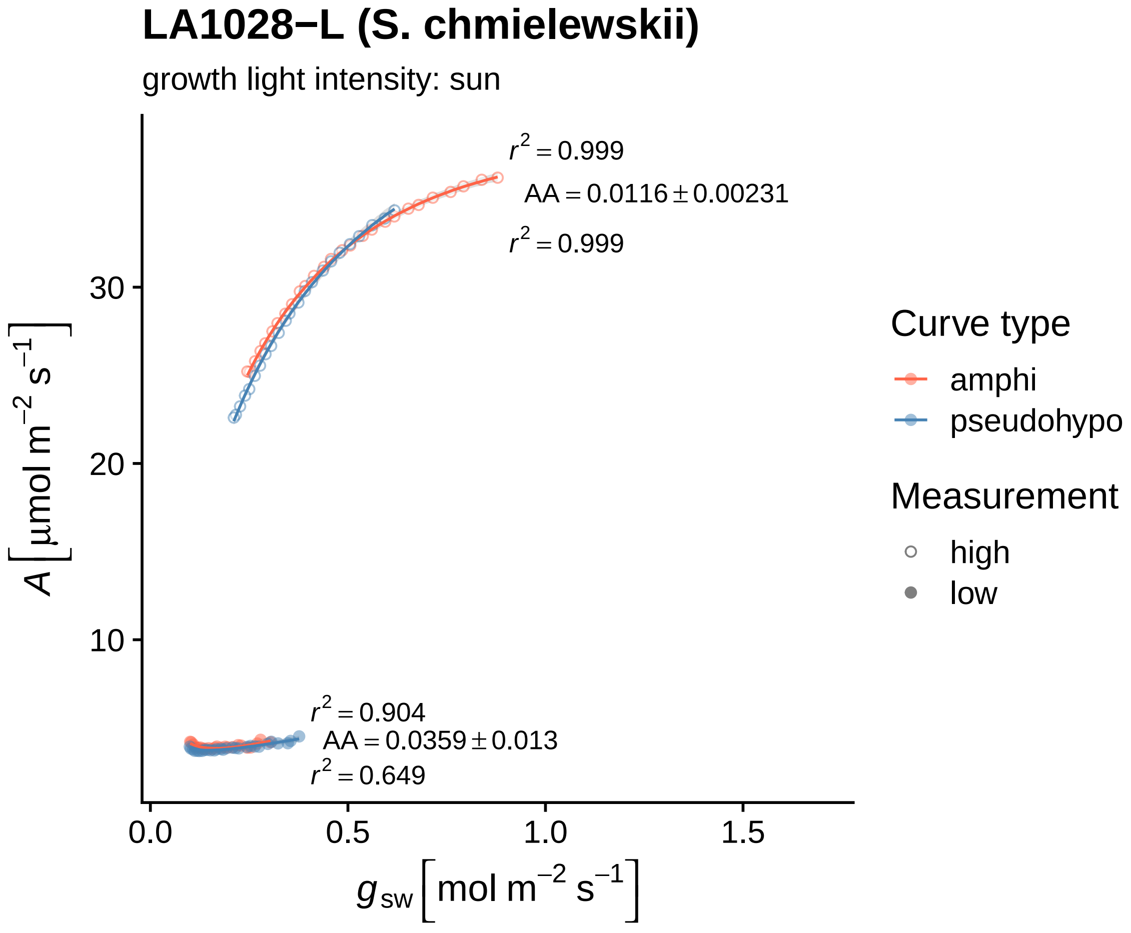

LA1028−L (S. chmielewskii)
growth light intensity: sun
$r^2 = 0.999$
$AA = 0.0116 \pm 0.00231$
$r^2 = 0.999$
$r^2 = 0.904$
$AA = 0.0359 \pm 0.013$
$r^2 = 0.649$
30
20
10
0.0
0.5
1.0
1.5
$A\ [\mu mol\ m^{-2}\ s^{-1}]$
$g_{sw}\ [mol\ m^{-2}\ s^{-1}]$
Curve type
amphi
pseudohypo
Measurement
high
low

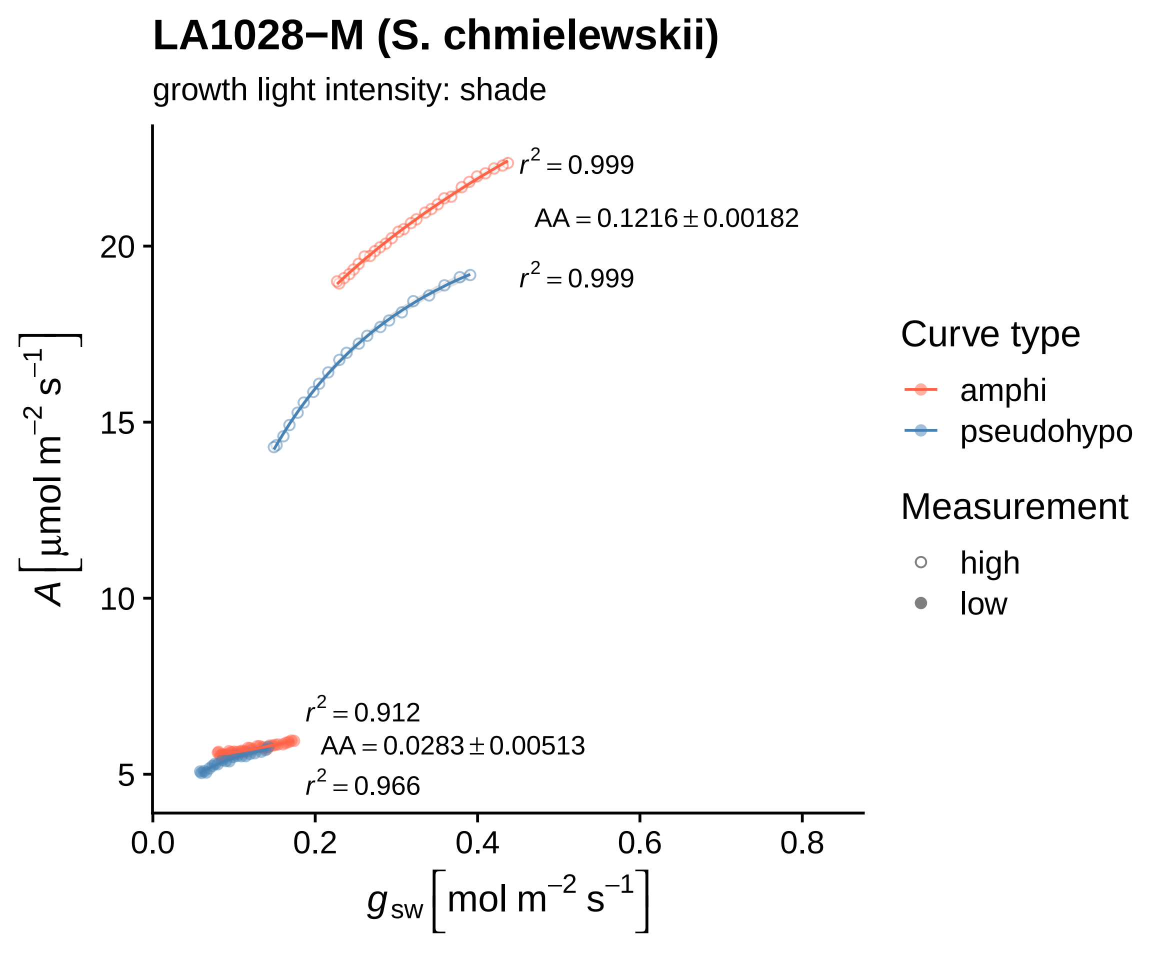

LA1028−M (S. chmielewskii)
growth light intensity: shade
$r^2 = 0.999$
AA = 0.1216 ± 0.00182
$r^2 = 0.999$
$r^2 = 0.912$
AA = 0.0283 ± 0.00513
$r^2 = 0.966$
20
15
10
5
0.0
0.2
0.4
0.6
0.8
$A\ [\mu mol\ m^{-2}\ s^{-1}]$
$g_{sw}\ [mol\ m^{-2}\ s^{-1}]$
Curve type
amphi
pseudohypo
Measurement
high
low

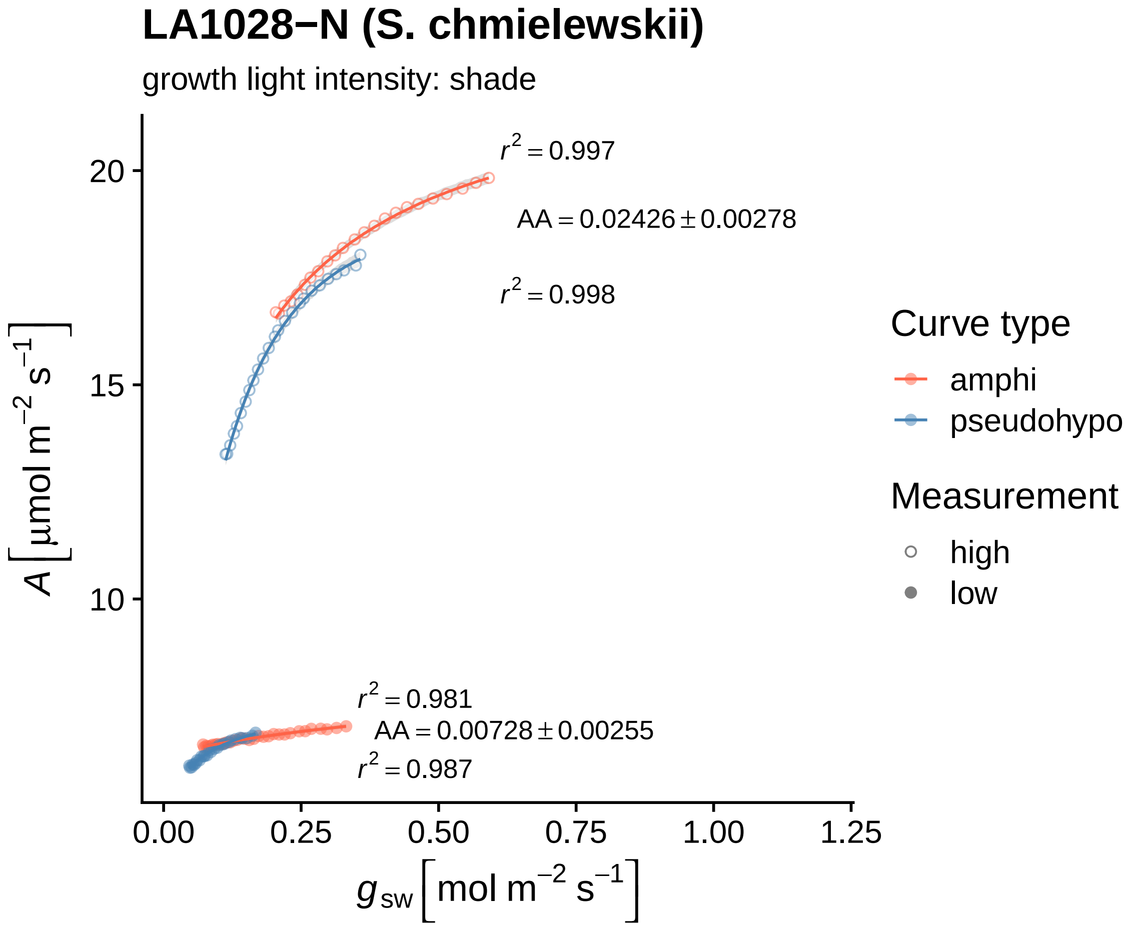

LA1028−N (S. chmielewskii)
growth light intensity: shade
r² = 0.997
AA = 0.02426 ± 0.00278
r² = 0.998
r² = 0.981
AA = 0.00728 ± 0.00255
r² = 0.987
20
15
10
A [μmol m⁻² s⁻¹]
0.00
0.25
0.50
0.75
1.00
1.25
g_sw [mol m⁻² s⁻¹]
Curve type
amphi
pseudohypo
Measurement
high
low

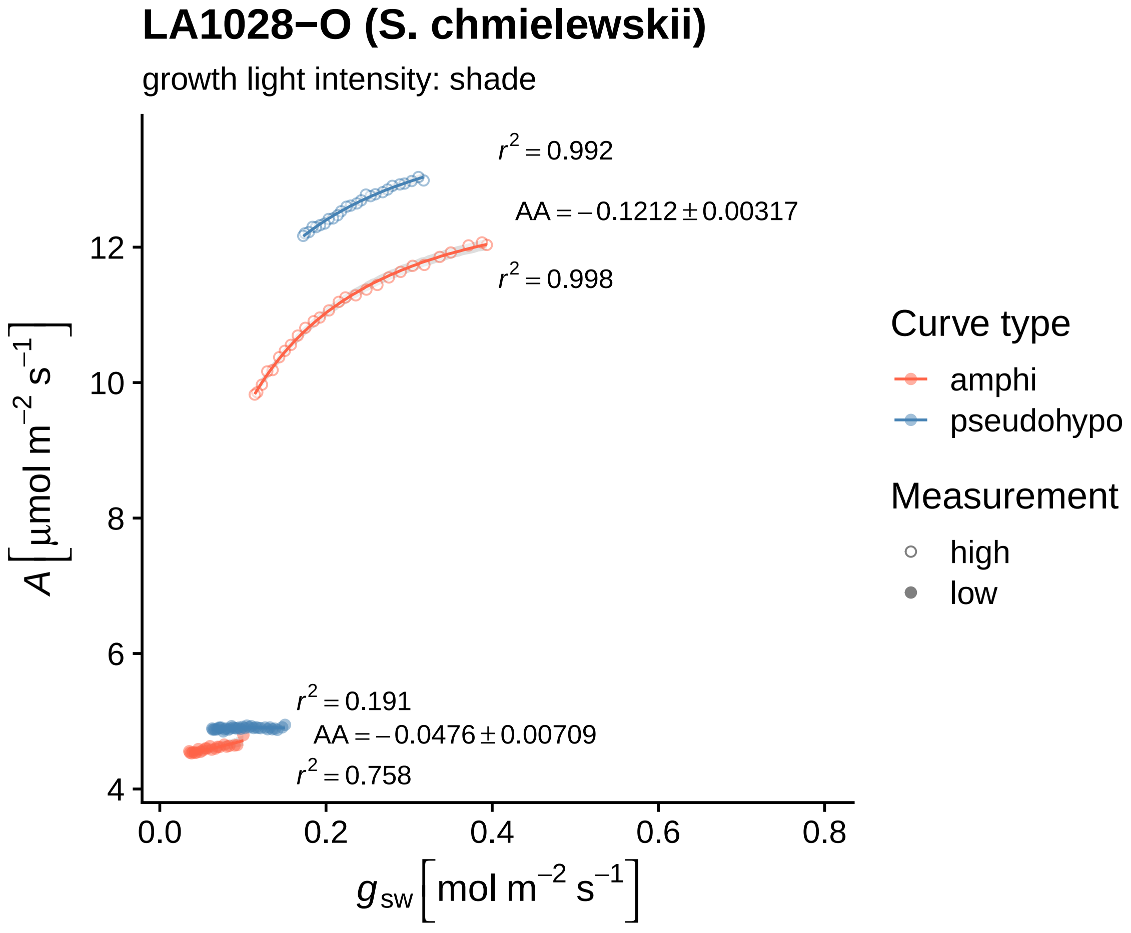
LA1028−O (S. chmielewskii)
growth light intensity: shade
$r^2 = 0.992$
$AA = -0.1212 \pm 0.00317$
$r^2 = 0.998$
$r^2 = 0.191$
$AA = -0.0476 \pm 0.00709$
$r^2 = 0.758$
$A\ [\mu mol\ m^{-2}\ s^{-1}]$
12
10
8
6
4
0.0
0.2
0.4
0.6
0.8
$g_{sw}\ [mol\ m^{-2}\ s^{-1}]$
Curve type
amphi
pseudohypo
Measurement
high
low

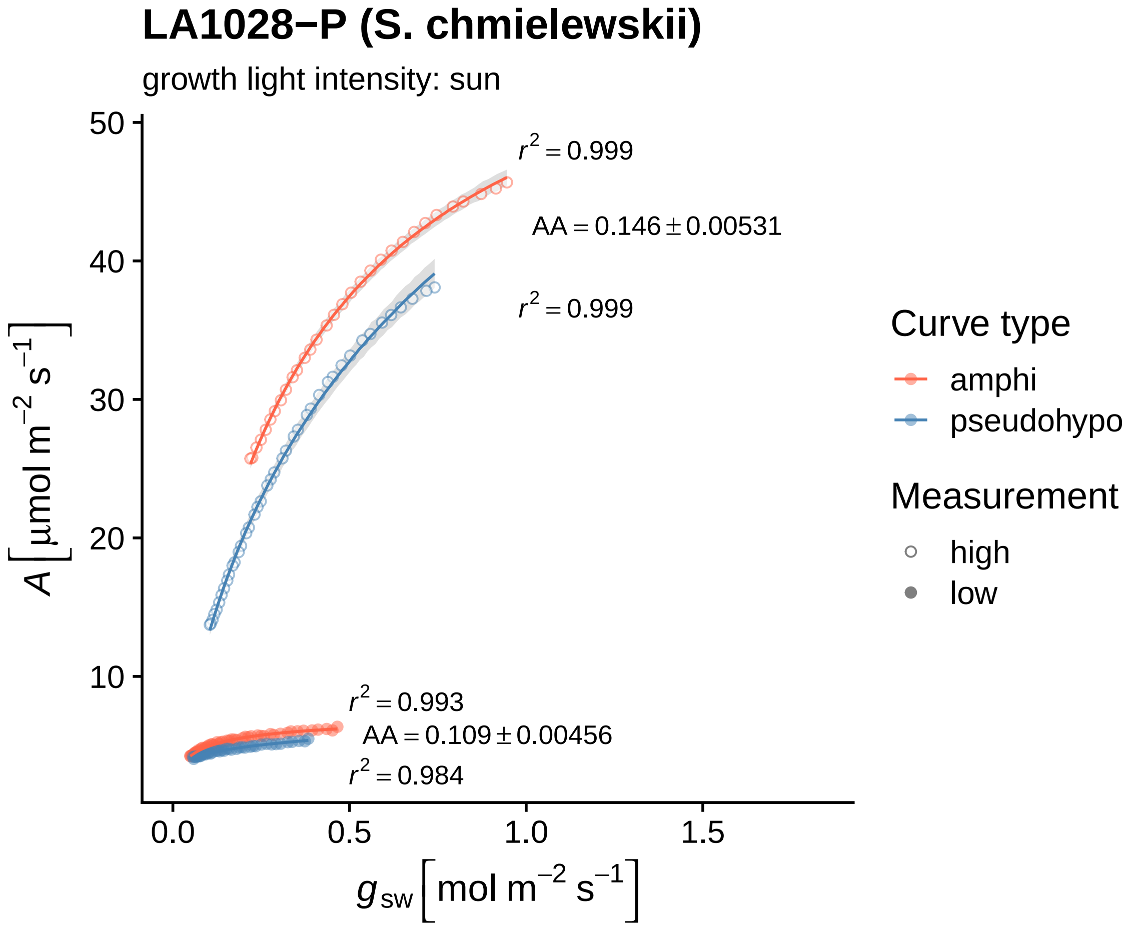
LA1028−P (S. chmielewskii)
growth light intensity: sun
$r^2 = 0.999$
$AA = 0.146 \pm 0.00531$
$r^2 = 0.999$
$r^2 = 0.993$
$AA = 0.109 \pm 0.00456$
$r^2 = 0.984$
$A\left[\mu mol\ m^{-2}\ s^{-1}\right]$
$g_{sw}\left[mol\ m^{-2}\ s^{-1}\right]$
50
40
30
20
10
0.0
0.5
1.0
1.5
Curve type
amphi
pseudohypo
Measurement
high
low

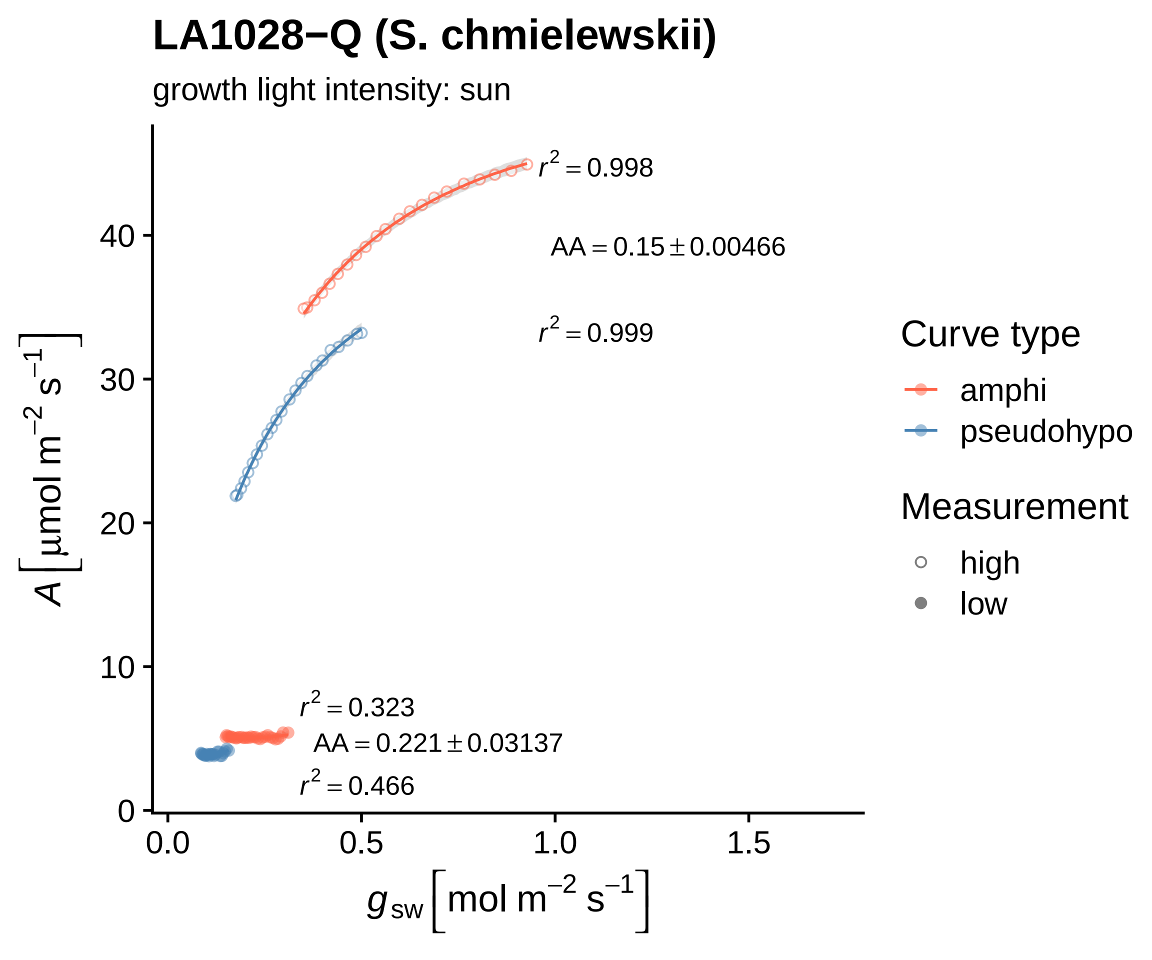

LA1028−Q (S. chmielewskii)
growth light intensity: sun
$r^2 = 0.998$
$AA = 0.15 \pm 0.00466$
$r^2 = 0.999$
$r^2 = 0.323$
$AA = 0.221 \pm 0.03137$
$r^2 = 0.466$
$A\ [\mu mol\ m^{-2}\ s^{-1}]$
$g_{sw}\ [mol\ m^{-2}\ s^{-1}]$
0
10
20
30
40
0.0
0.5
1.0
1.5
Curve type
amphi
pseudohypo
Measurement
high
low

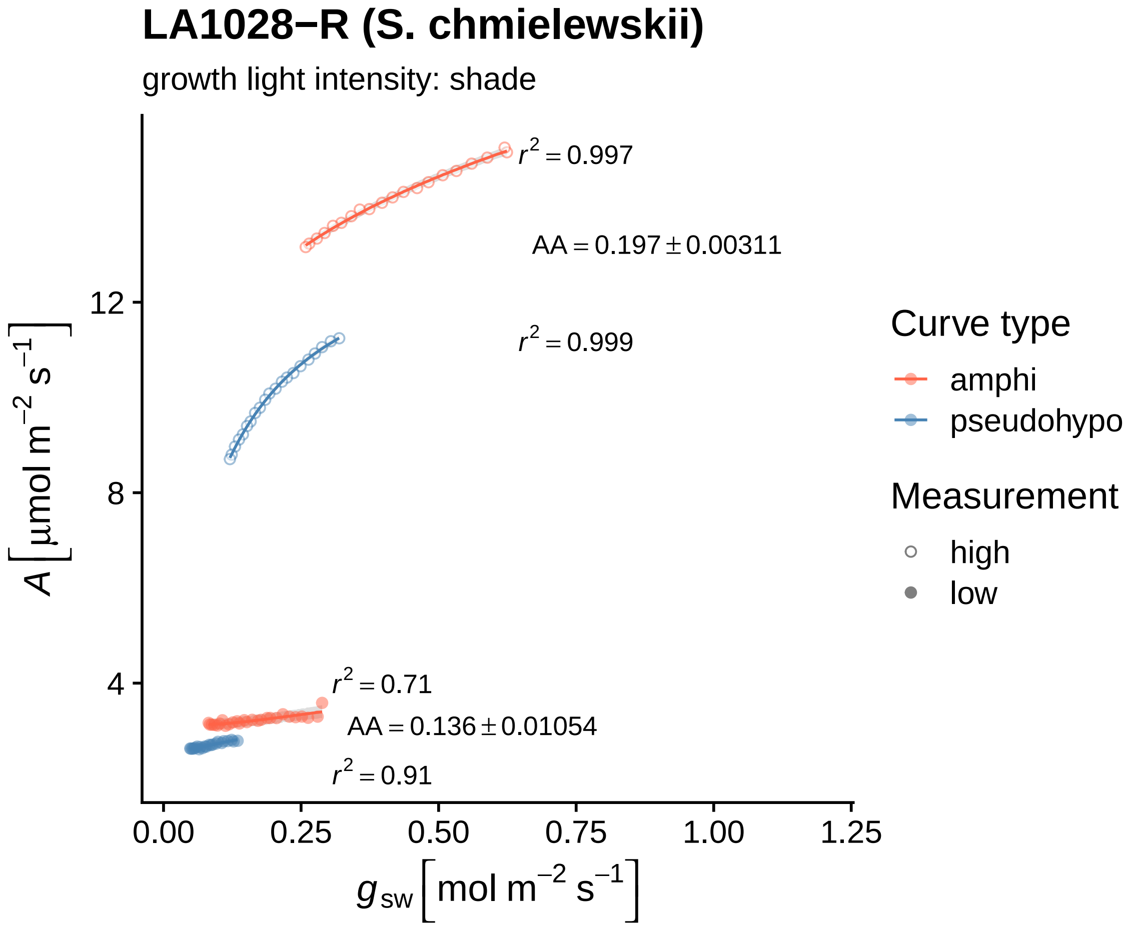
LA1028−R (S. chmielewskii)
growth light intensity: shade
$r^2 = 0.997$
$AA = 0.197 \pm 0.00311$
$r^2 = 0.999$
$r^2 = 0.71$
$AA = 0.136 \pm 0.01054$
$r^2 = 0.91$
$A\,[\mu mol\ m^{-2}\ s^{-1}]$
12
8
4
0.00
0.25
0.50
0.75
1.00
1.25
$g_{sw}\,[mol\ m^{-2}\ s^{-1}]$
Curve type
amphi
pseudohypo
Measurement
high
low

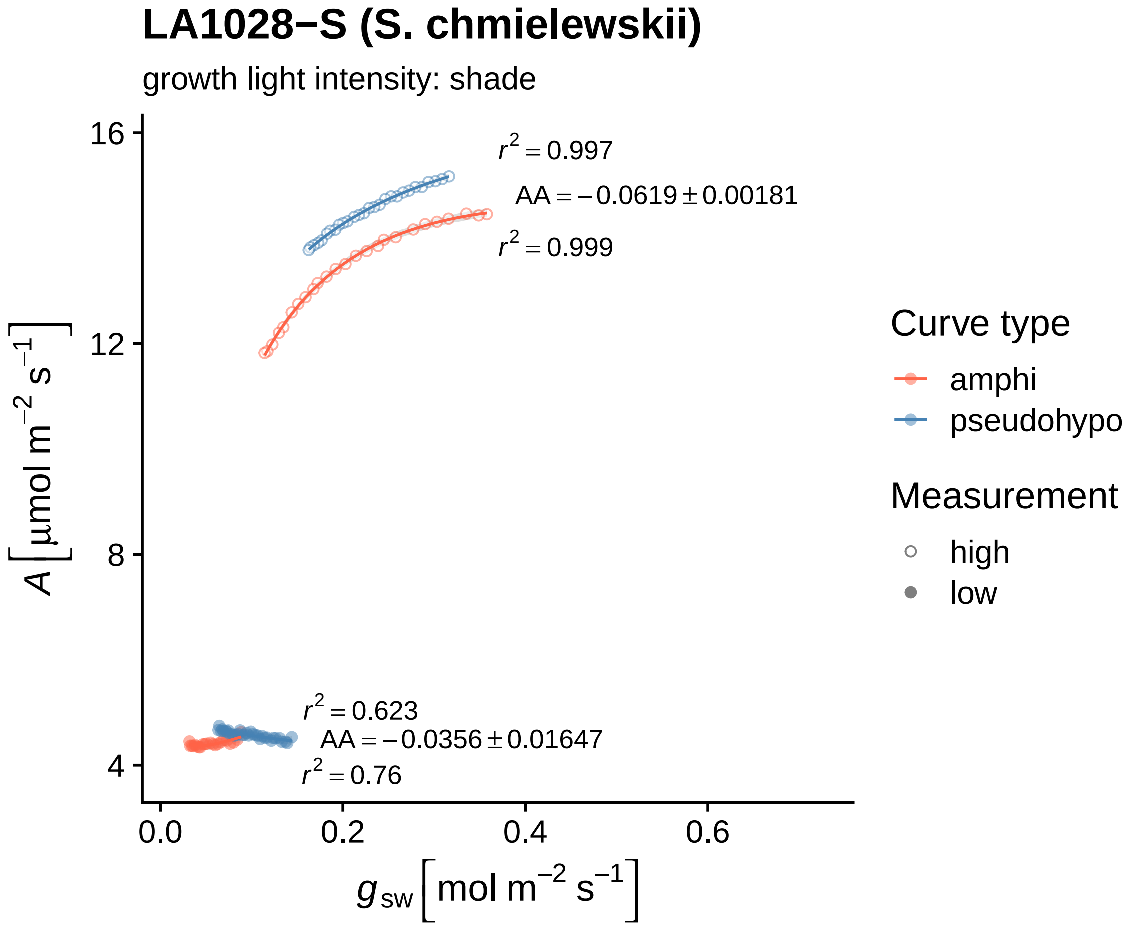
LA1028−S (S. chmielewskii)
growth light intensity: shade
r² = 0.997
AA = −0.0619 ± 0.00181
r² = 0.999
r² = 0.623
AA = −0.0356 ± 0.01647
r² = 0.76
16
12
8
4
0.0
0.2
0.4
0.6
A [µmol m⁻² s⁻¹]
g_sw [mol m⁻² s⁻¹]
Curve type
amphi
pseudohypo
Measurement
high
low

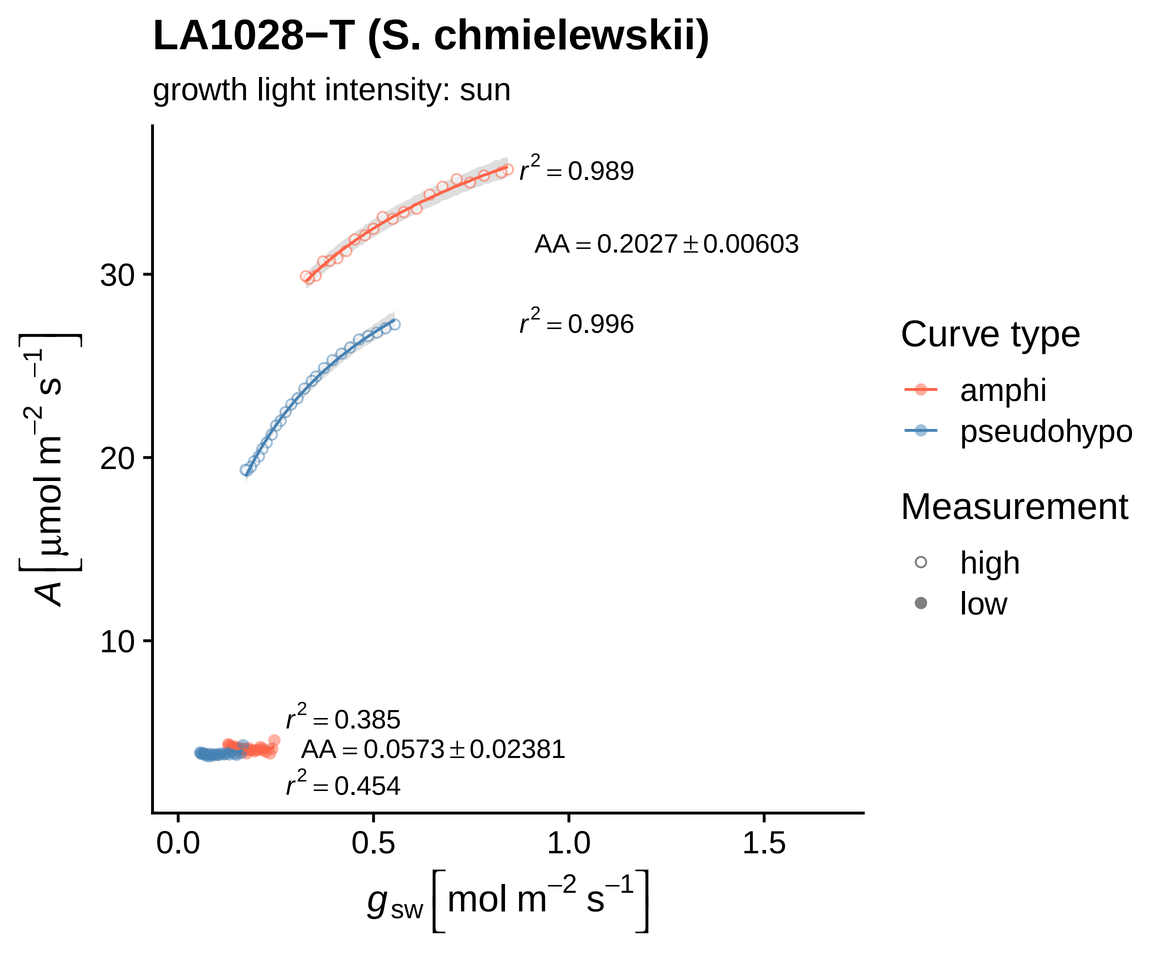

LA1028−T (S. chmielewskii)
growth light intensity: sun
$r^2 = 0.989$
$AA = 0.2027 \pm 0.00603$
$r^2 = 0.996$
$r^2 = 0.385$
$AA = 0.0573 \pm 0.02381$
$r^2 = 0.454$
30
20
10
0.0
0.5
1.0
1.5
$A\ [\mu mol\ m^{-2}\ s^{-1}]$
$g_{sw}\ [mol\ m^{-2}\ s^{-1}]$
Curve type
amphi
pseudohypo
Measurement
high
low

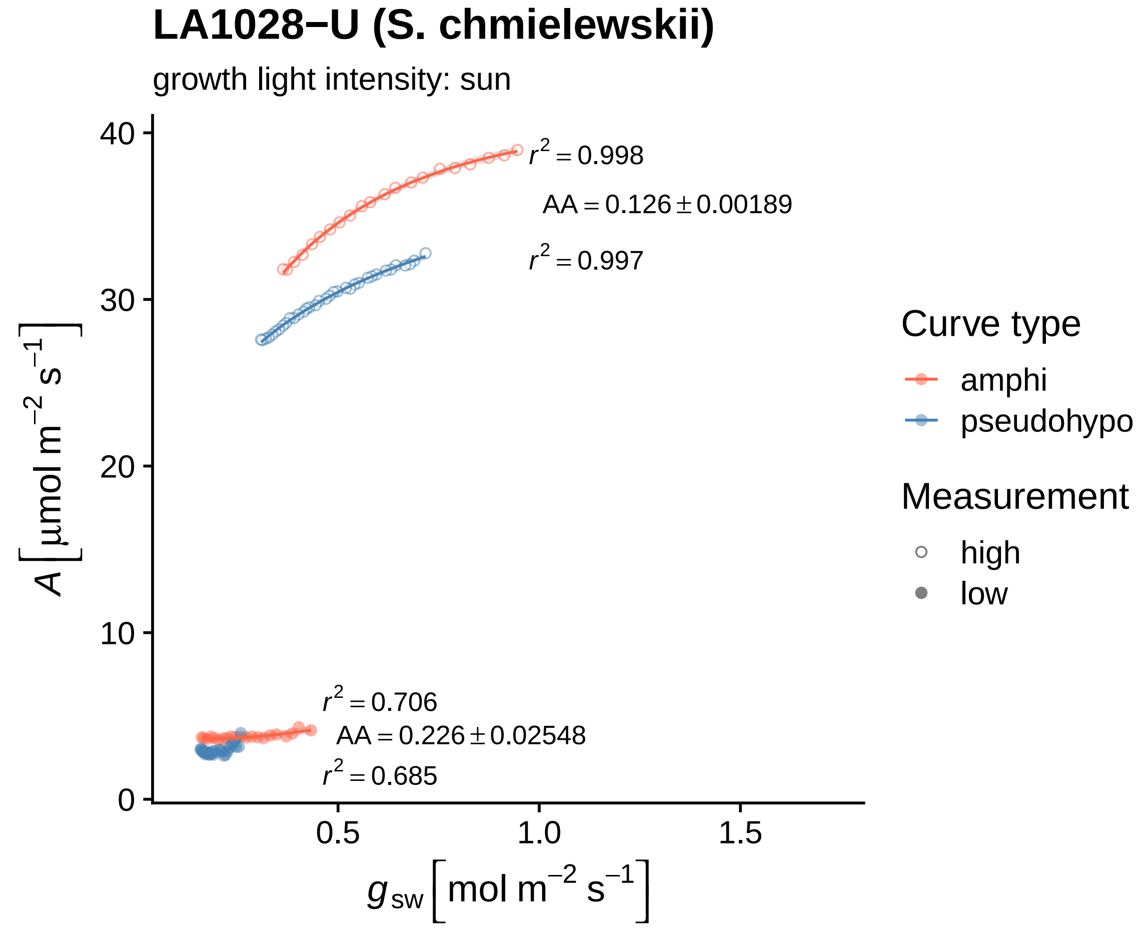
LA1028−U (S. chmielewskii)
growth light intensity: sun
$r^2 = 0.998$
$AA = 0.126 \pm 0.00189$
$r^2 = 0.997$
$r^2 = 0.706$
$AA = 0.226 \pm 0.02548$
$r^2 = 0.685$
40
30
20
10
0
0.5
1.0
1.5
$A \left[\mu mol\ m^{-2}\ s^{-1}\right]$
$g_{sw} \left[mol\ m^{-2}\ s^{-1}\right]$
Curve type
amphi
pseudohypo
Measurement
high
low

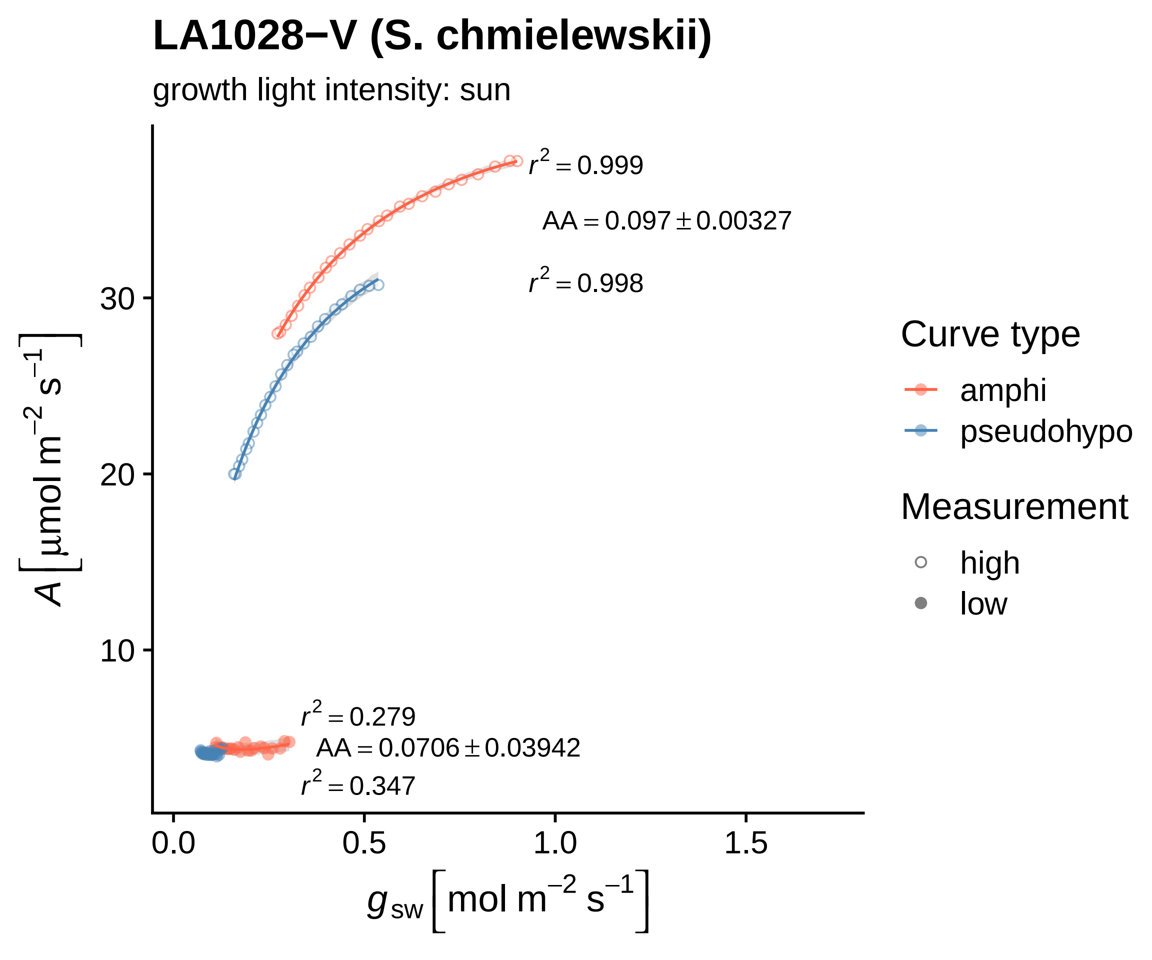
LA1028−V (S. chmielewskii)
growth light intensity: sun
r² = 0.999
AA = 0.097 ± 0.00327
r² = 0.998
r² = 0.279
AA = 0.0706 ± 0.03942
r² = 0.347
A [μmol m⁻² s⁻¹]
g_sw [mol m⁻² s⁻¹]
0.0
0.5
1.0
1.5
10
20
30
Curve type
amphi
pseudohypo
Measurement
high
low

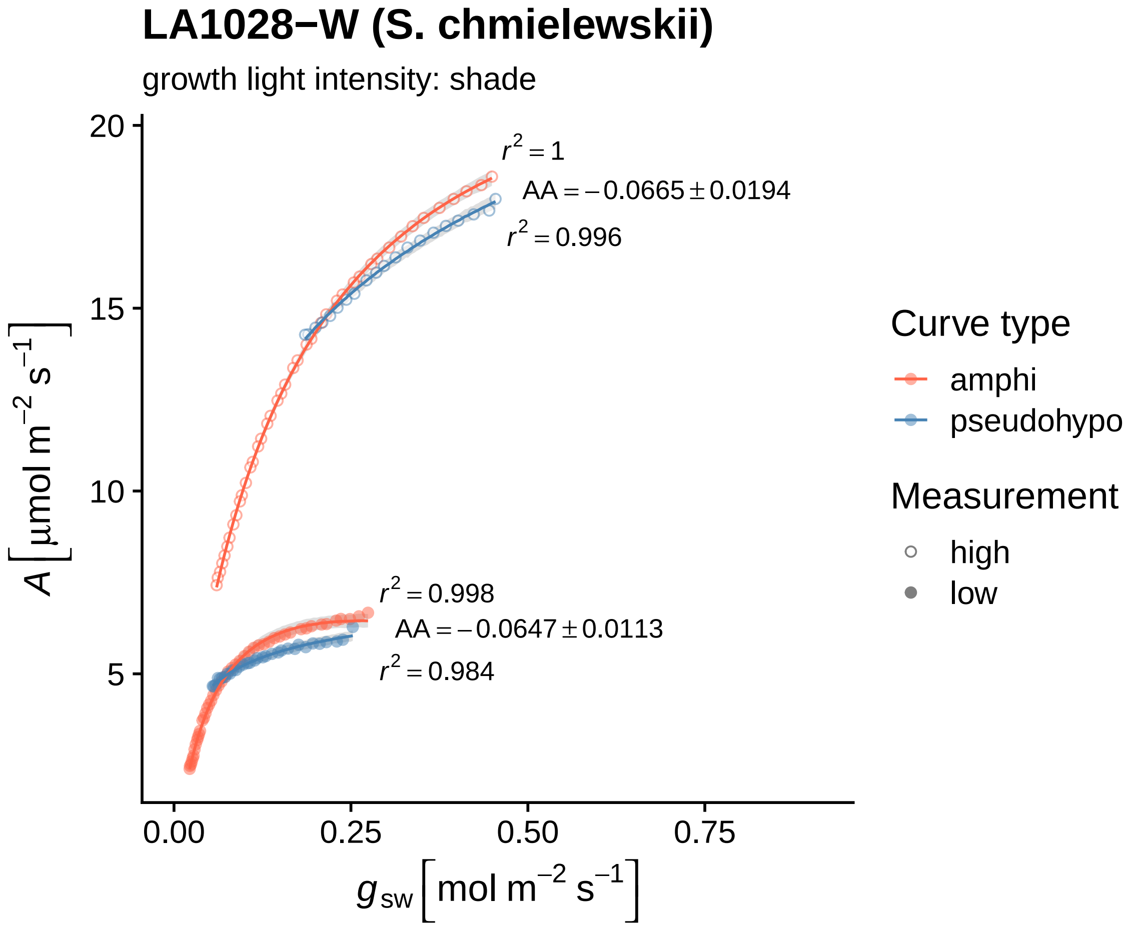

LA1028−W (S. chmielewskii)
growth light intensity: shade
r^2 = 1
AA = −0.0665 ± 0.0194
r^2 = 0.996
r^2 = 0.998
AA = −0.0647 ± 0.0113
r^2 = 0.984
20
15
10
5
0.00
0.25
0.50
0.75
A [μmol m^−2 s^−1]
g_sw [mol m^−2 s^−1]
Curve type
amphi
pseudohypo
Measurement
high
low

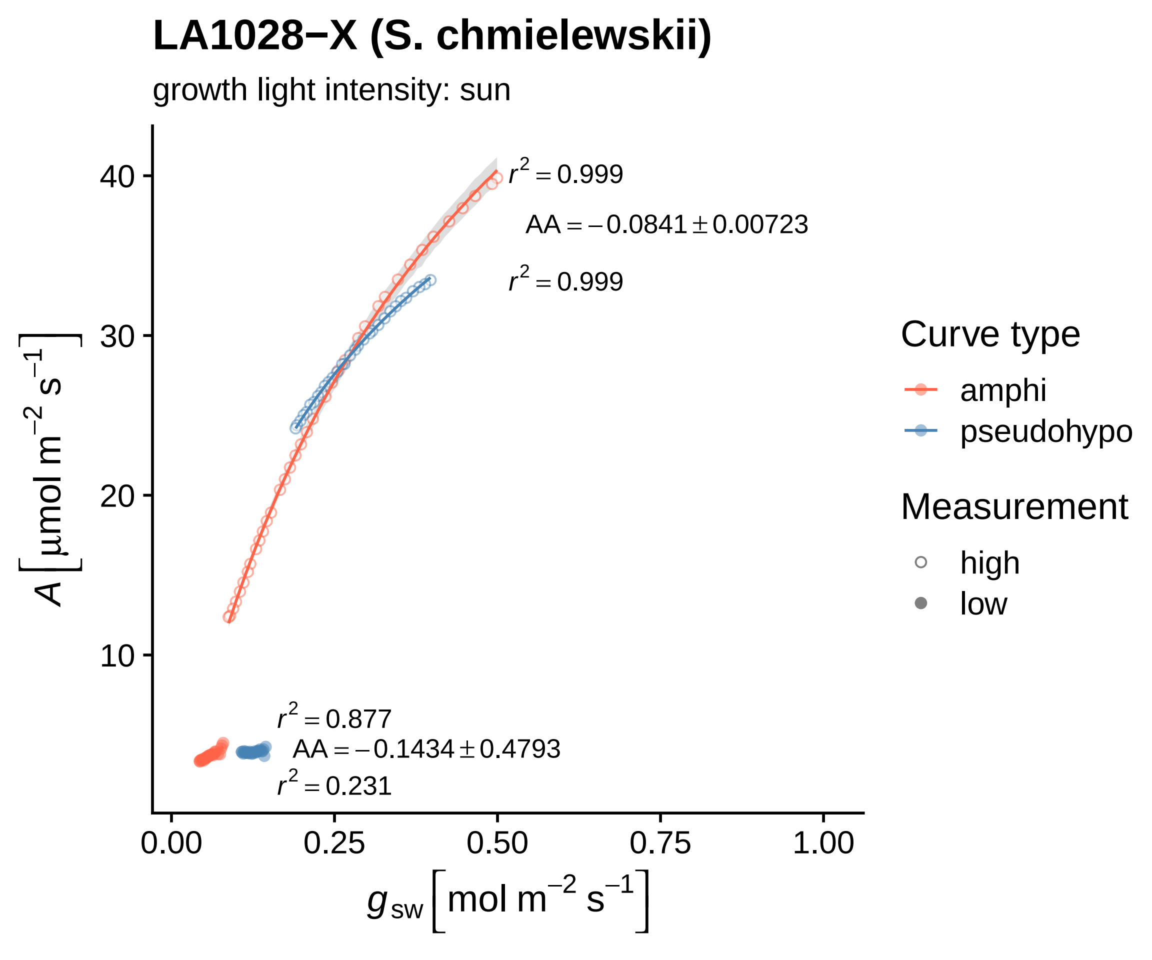
LA1028−X (S. chmielewskii)
growth light intensity: sun
$r^2 = 0.999$
$AA = -0.0841 \pm 0.00723$
$r^2 = 0.999$
$r^2 = 0.877$
$AA = -0.1434 \pm 0.4793$
$r^2 = 0.231$
40
30
20
10
0.00
0.25
0.50
0.75
1.00
$A\ [\mu mol\ m^{-2}\ s^{-1}]$
$g_{sw}\ [mol\ m^{-2}\ s^{-1}]$
Curve type
amphi
pseudohypo
Measurement
high
low

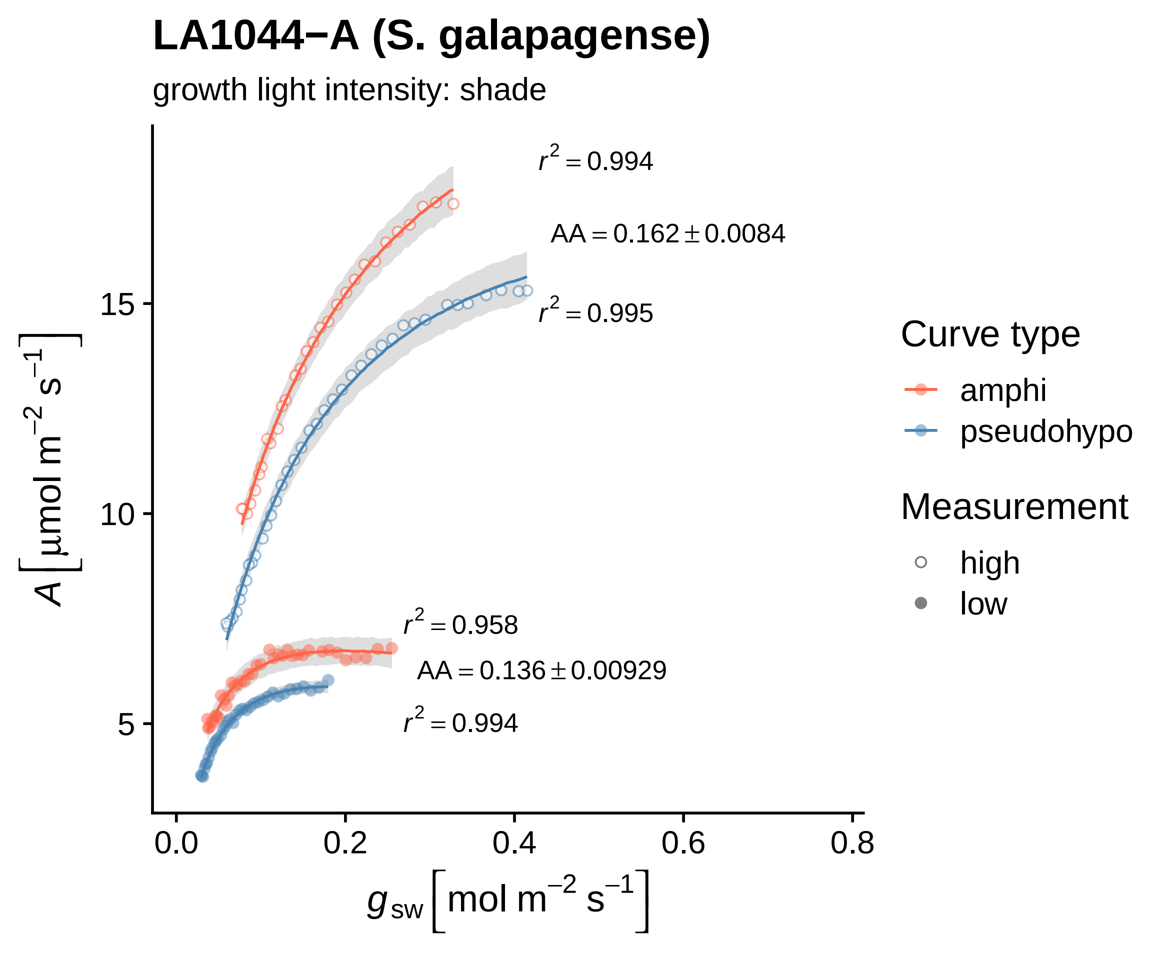

LA1044−A (S. galapagense)
growth light intensity: shade
$r^2 = 0.994$
$AA = 0.162 \pm 0.0084$
$r^2 = 0.995$
$r^2 = 0.958$
$AA = 0.136 \pm 0.00929$
$r^2 = 0.994$
15
10
5
0.0
0.2
0.4
0.6
0.8
$A\left[\mu mol\ m^{-2}\ s^{-1}\right]$
$g_{sw}\left[mol\ m^{-2}\ s^{-1}\right]$
Curve type
amphi
pseudohypo
Measurement
high
low

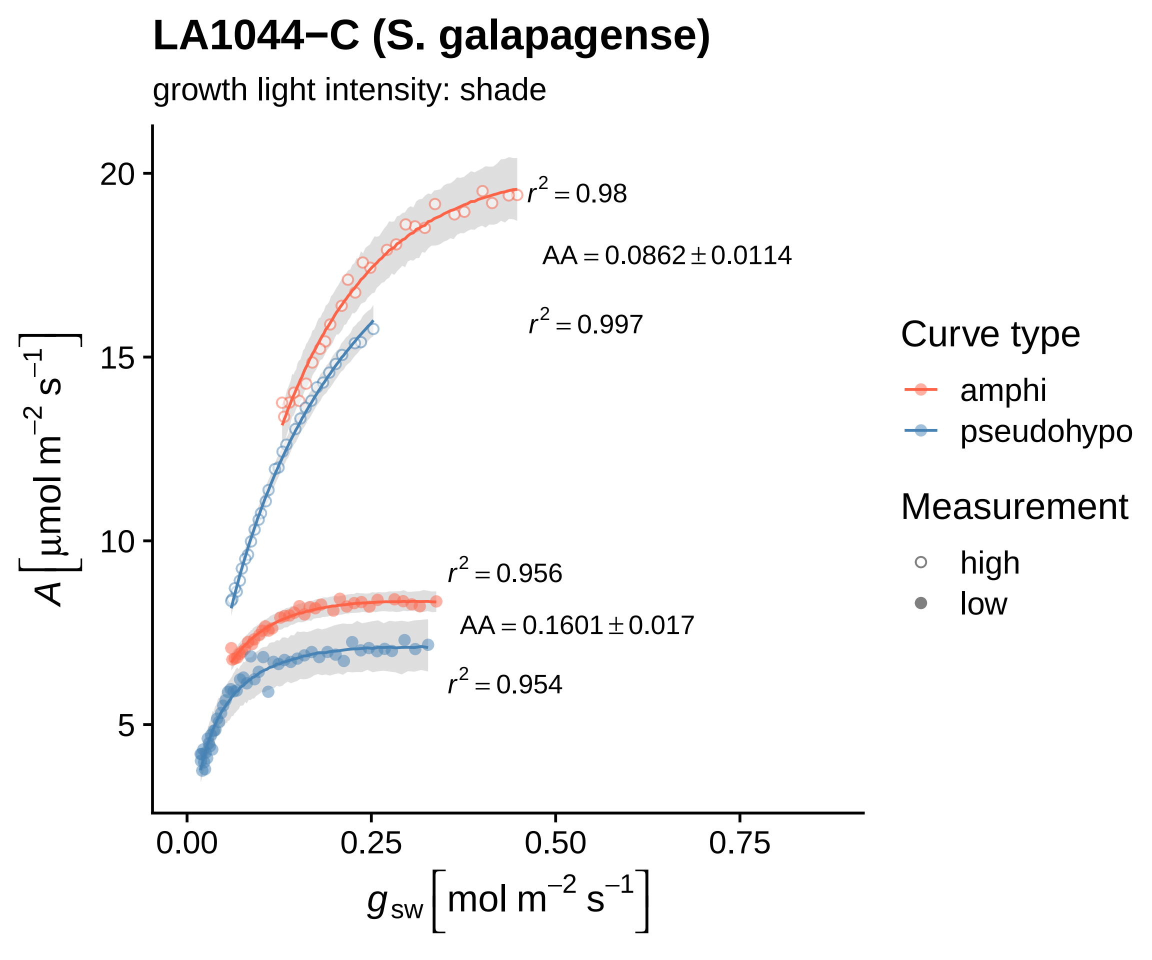
LA1044−C (S. galapagense)
growth light intensity: shade
$r^2 = 0.98$
$AA = 0.0862 \pm 0.0114$
$r^2 = 0.997$
$r^2 = 0.956$
$AA = 0.1601 \pm 0.017$
$r^2 = 0.954$
$A\left[\mu mol\ m^{-2}\ s^{-1}\right]$
20
15
10
5
0.00
0.25
0.50
0.75
$g_{sw}\left[mol\ m^{-2}\ s^{-1}\right]$
Curve type
amphi
pseudohypo
Measurement
high
low

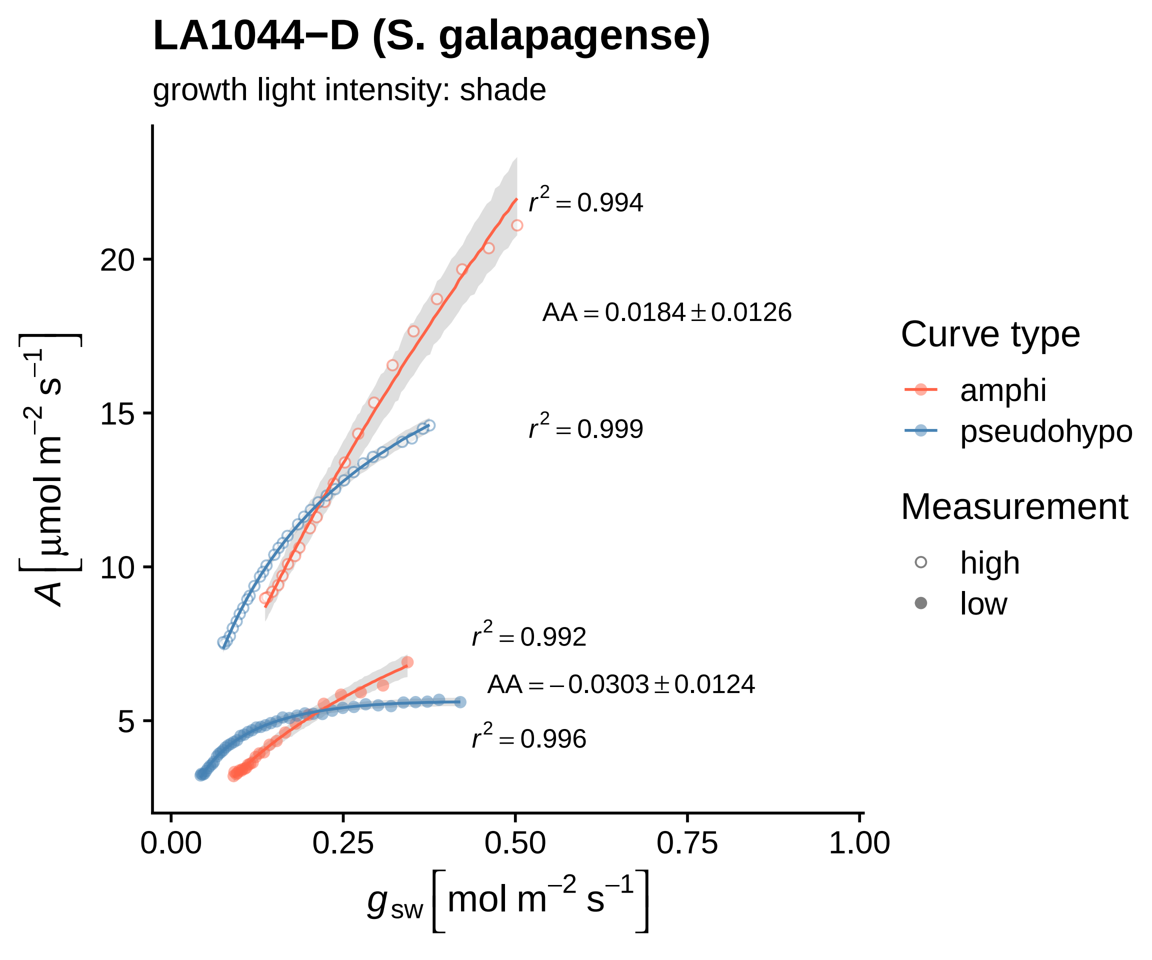

LA1044−D (S. galapagense)
growth light intensity: shade
$r^2 = 0.994$
$AA = 0.0184 \pm 0.0126$
$r^2 = 0.999$
$r^2 = 0.992$
$AA = -0.0303 \pm 0.0124$
$r^2 = 0.996$
20
15
10
5
0.00
0.25
0.50
0.75
1.00
$A\ [\mu mol\ m^{-2}\ s^{-1}]$
$g_{sw}\ [mol\ m^{-2}\ s^{-1}]$
Curve type
amphi
pseudohypo
Measurement
high
low

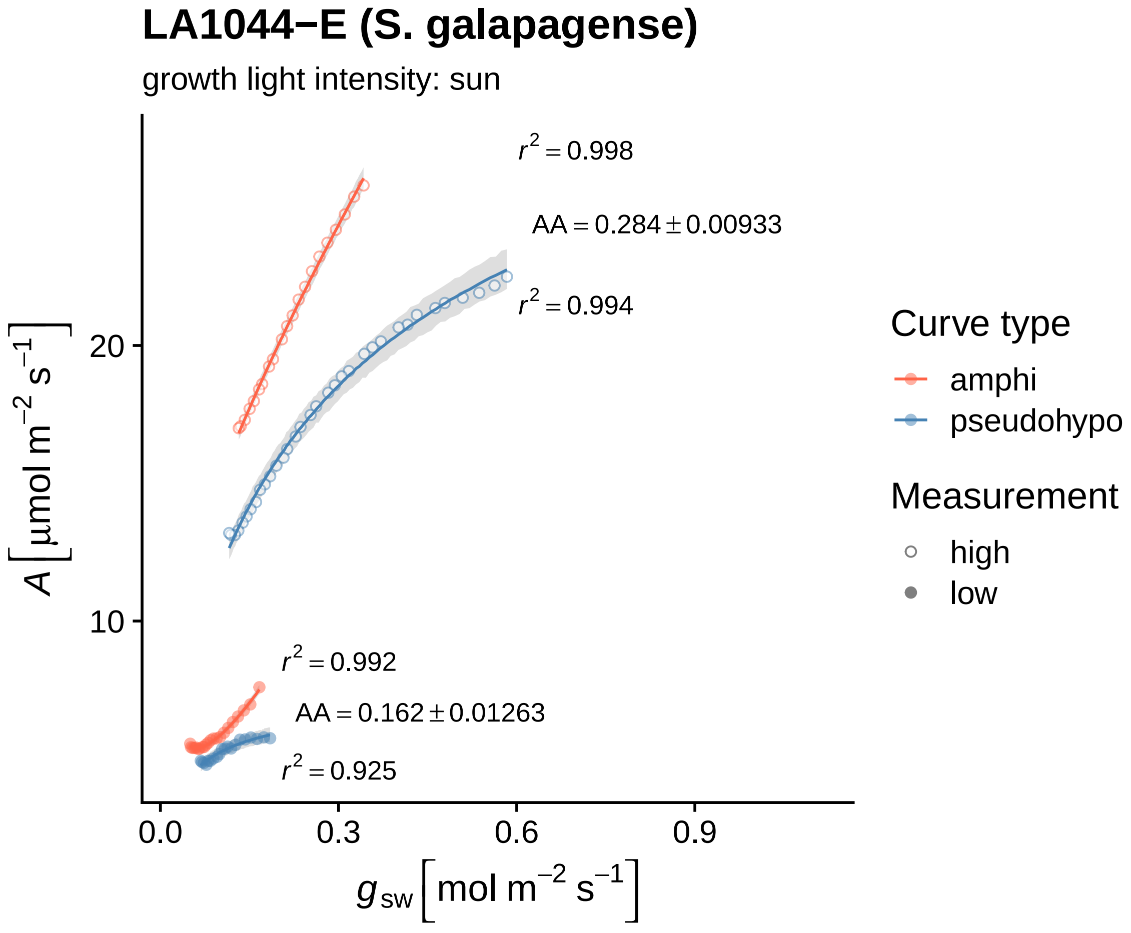
LA1044−E (S. galapagense)
growth light intensity: sun
$r^2 = 0.998$
$AA = 0.284 \pm 0.00933$
$r^2 = 0.994$
$r^2 = 0.992$
$AA = 0.162 \pm 0.01263$
$r^2 = 0.925$
20
10
0.0
0.3
0.6
0.9
$A\ [\mu mol\ m^{-2}\ s^{-1}]$
$g_{sw}\ [mol\ m^{-2}\ s^{-1}]$
Curve type
amphi
pseudohypo
Measurement
high
low

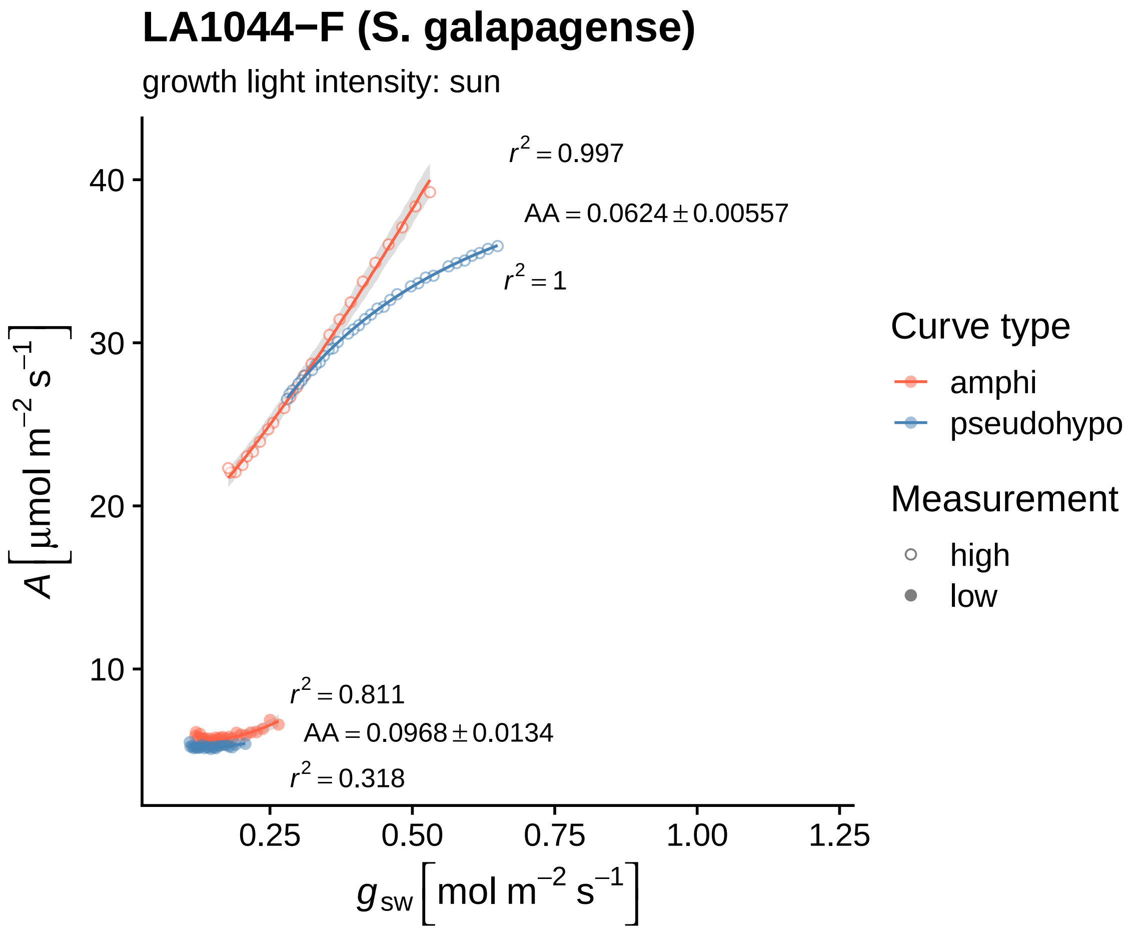
LA1044−F (S. galapagense)
growth light intensity: sun
$r^2 = 0.997$
$AA = 0.0624 \pm 0.00557$
$r^2 = 1$
$r^2 = 0.811$
$AA = 0.0968 \pm 0.0134$
$r^2 = 0.318$
40
30
20
10
0.25
0.50
0.75
1.00
1.25
$A\ [\mu mol\ m^{-2}\ s^{-1}]$
$g_{sw}\ [mol\ m^{-2}\ s^{-1}]$
Curve type
amphi
pseudohypo
Measurement
high
low

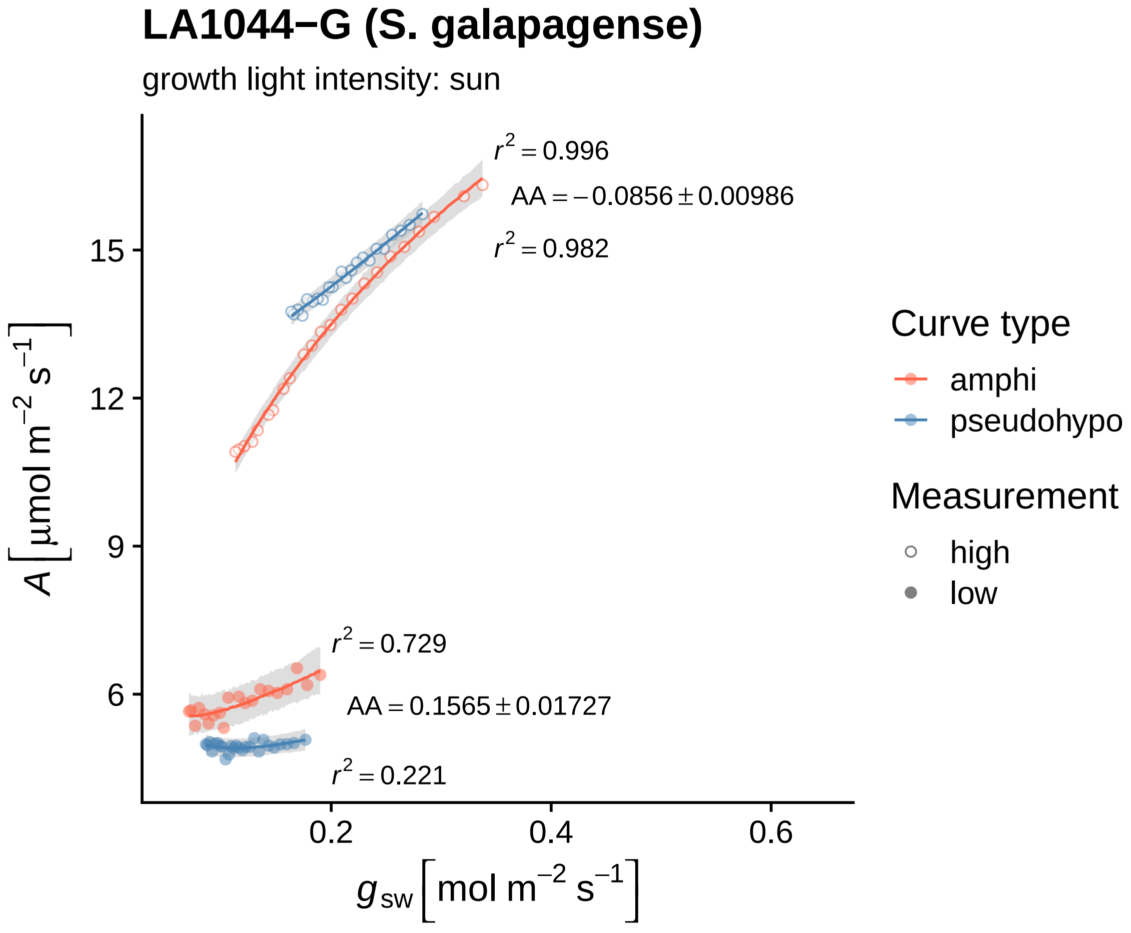

LA1044−G (S. galapagense)
growth light intensity: sun
$r^2 = 0.996$
$AA = -0.0856 \pm 0.00986$
$r^2 = 0.982$
$r^2 = 0.729$
$AA = 0.1565 \pm 0.01727$
$r^2 = 0.221$
$A\left[\mu mol\ m^{-2}\ s^{-1}\right]$
15
12
9
6
0.2
0.4
0.6
$g_{sw}\left[mol\ m^{-2}\ s^{-1}\right]$
Curve type
amphi
pseudohypo
Measurement
high
low

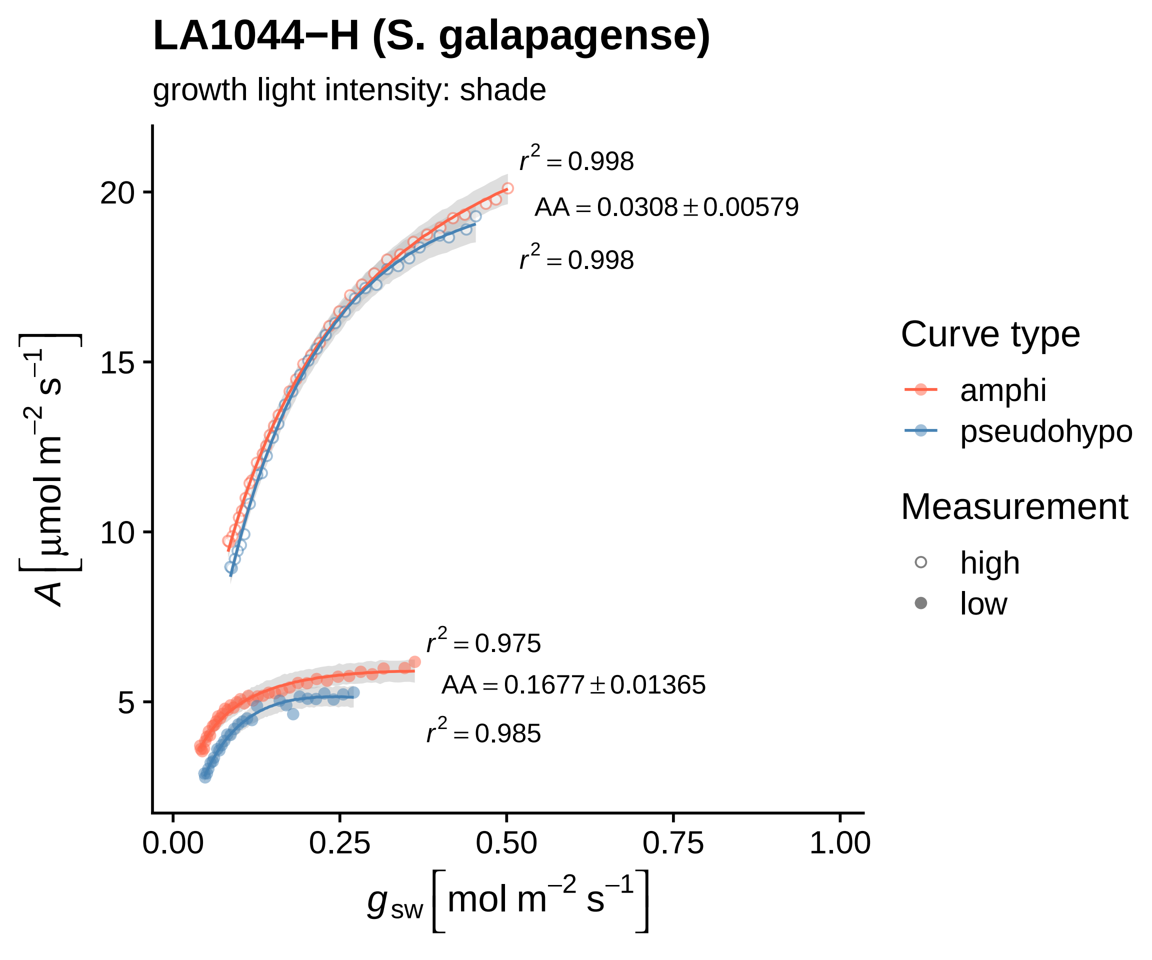

LA1044−H (S. galapagense)
growth light intensity: shade
$r^2 = 0.998$
$AA = 0.0308 \pm 0.00579$
$r^2 = 0.998$
$r^2 = 0.975$
$AA = 0.1677 \pm 0.01365$
$r^2 = 0.985$
$A\left[\mu mol\ m^{-2}\ s^{-1}\right]$
20
15
10
5
0.00
0.25
0.50
0.75
1.00
$g_{sw}\left[mol\ m^{-2}\ s^{-1}\right]$
Curve type
amphi
pseudohypo
Measurement
high
low

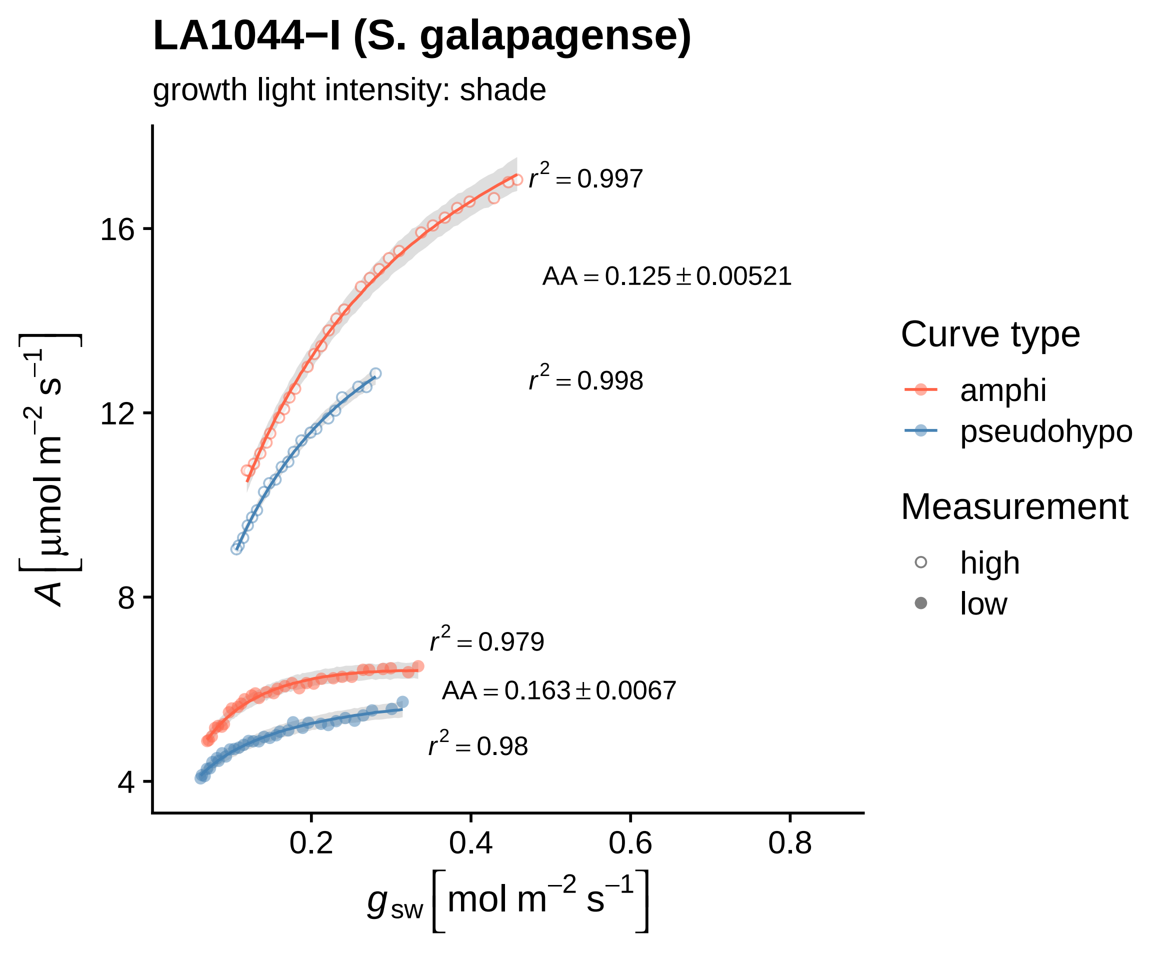
LA1044−I (S. galapagense)
growth light intensity: shade
$r^2 = 0.997$
$AA = 0.125 \pm 0.00521$
$r^2 = 0.998$
$r^2 = 0.979$
$AA = 0.163 \pm 0.0067$
$r^2 = 0.98$
16
12
8
4
0.2
0.4
0.6
0.8
$A\ [\mu mol\ m^{-2}\ s^{-1}]$
$g_{sw}\ [mol\ m^{-2}\ s^{-1}]$
Curve type
amphi
pseudohypo
Measurement
high
low

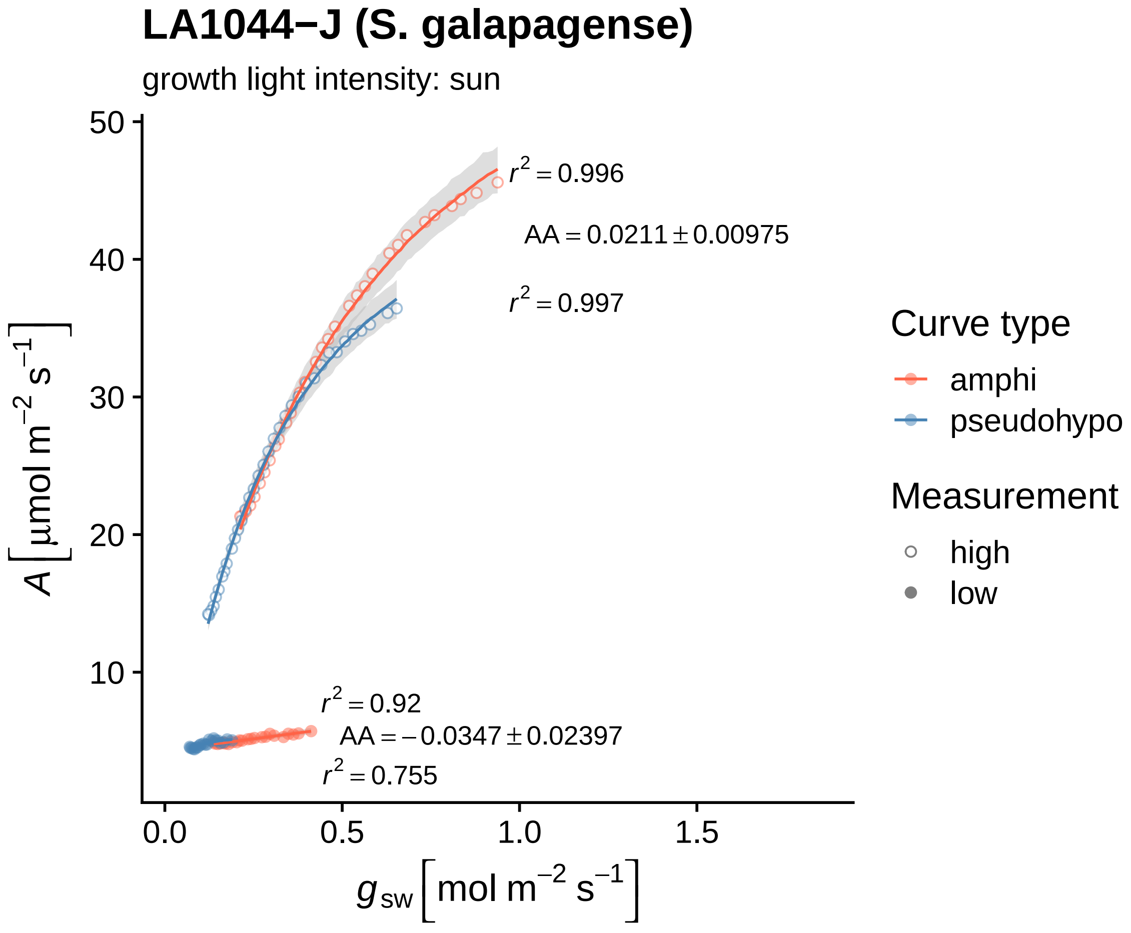

LA1044−J (S. galapagense)
growth light intensity: sun
$r^2 = 0.996$
$AA = 0.0211 \pm 0.00975$
$r^2 = 0.997$
$r^2 = 0.92$
$AA = -0.0347 \pm 0.02397$
$r^2 = 0.755$
$A\left[\mu mol\ m^{-2}\ s^{-1}\right]$
$g_{sw}\left[mol\ m^{-2}\ s^{-1}\right]$
50
40
30
20
10
0.0
0.5
1.0
1.5
Curve type
amphi
pseudohypo
Measurement
high
low

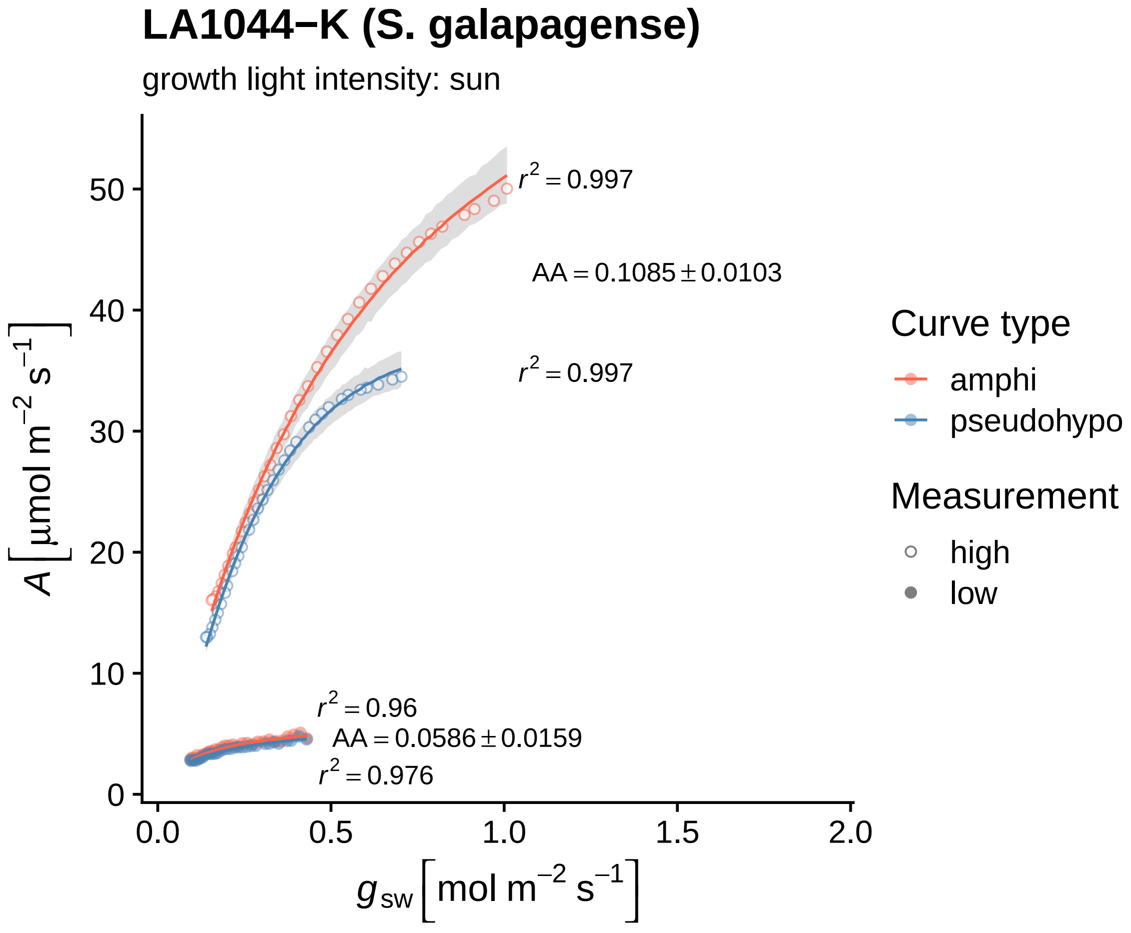

LA1044−K (S. galapagense)
growth light intensity: sun
$r^2 = 0.997$
AA = 0.1085 ± 0.0103
$r^2 = 0.997$
$r^2 = 0.96$
AA = 0.0586 ± 0.0159
$r^2 = 0.976$
$A$ [µmol m$^{-2}$ s$^{-1}$]
$g_{sw}$ [mol m$^{-2}$ s$^{-1}$]
0
10
20
30
40
50
0.0
0.5
1.0
1.5
2.0
Curve type
amphi
pseudohypo
Measurement
high
low

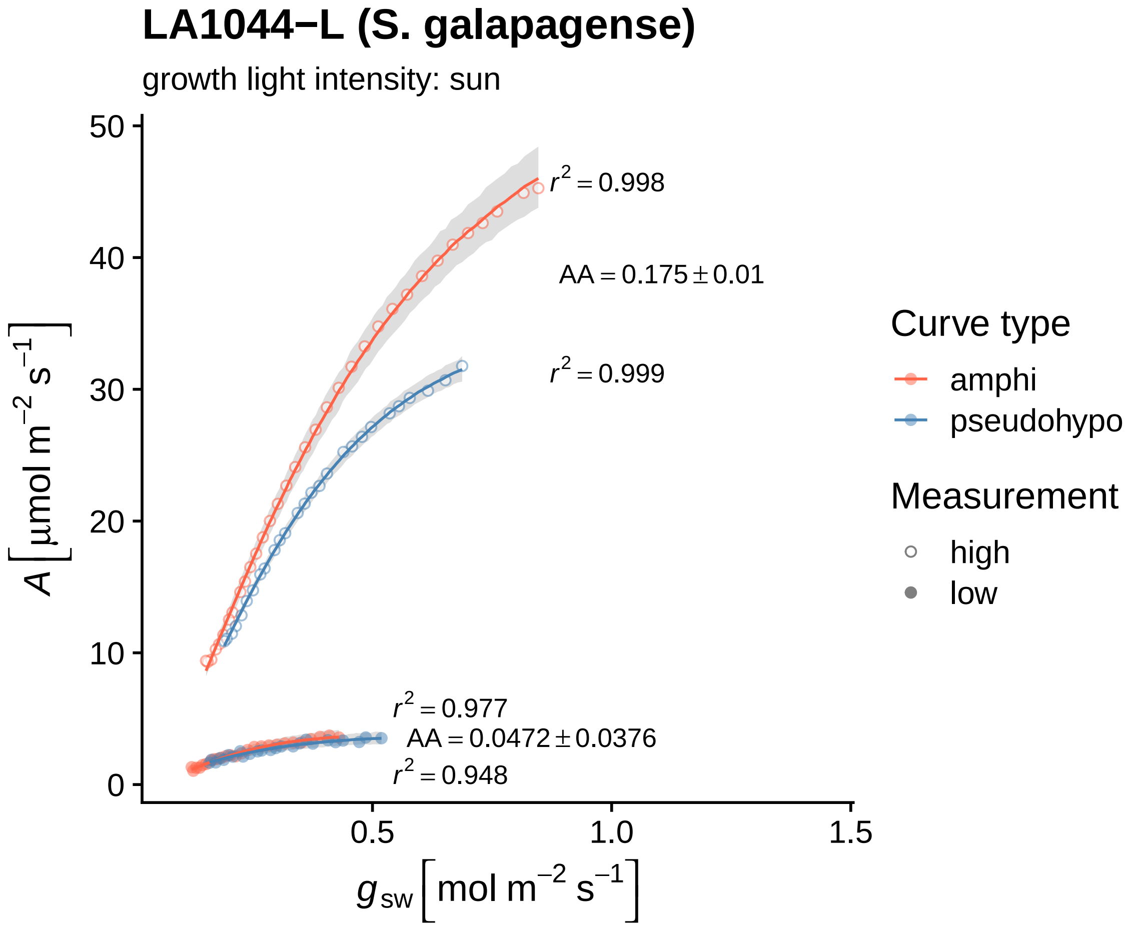
LA1044−L (S. galapagense)
growth light intensity: sun
$r^2 = 0.998$
$AA = 0.175 \pm 0.01$
$r^2 = 0.999$
$r^2 = 0.977$
$AA = 0.0472 \pm 0.0376$
$r^2 = 0.948$
$A\left[\mu mol\ m^{-2}\ s^{-1}\right]$
$g_{sw}\left[mol\ m^{-2}\ s^{-1}\right]$
0
10
20
30
40
50
0.5
1.0
1.5
Curve type
amphi
pseudohypo
Measurement
high
low

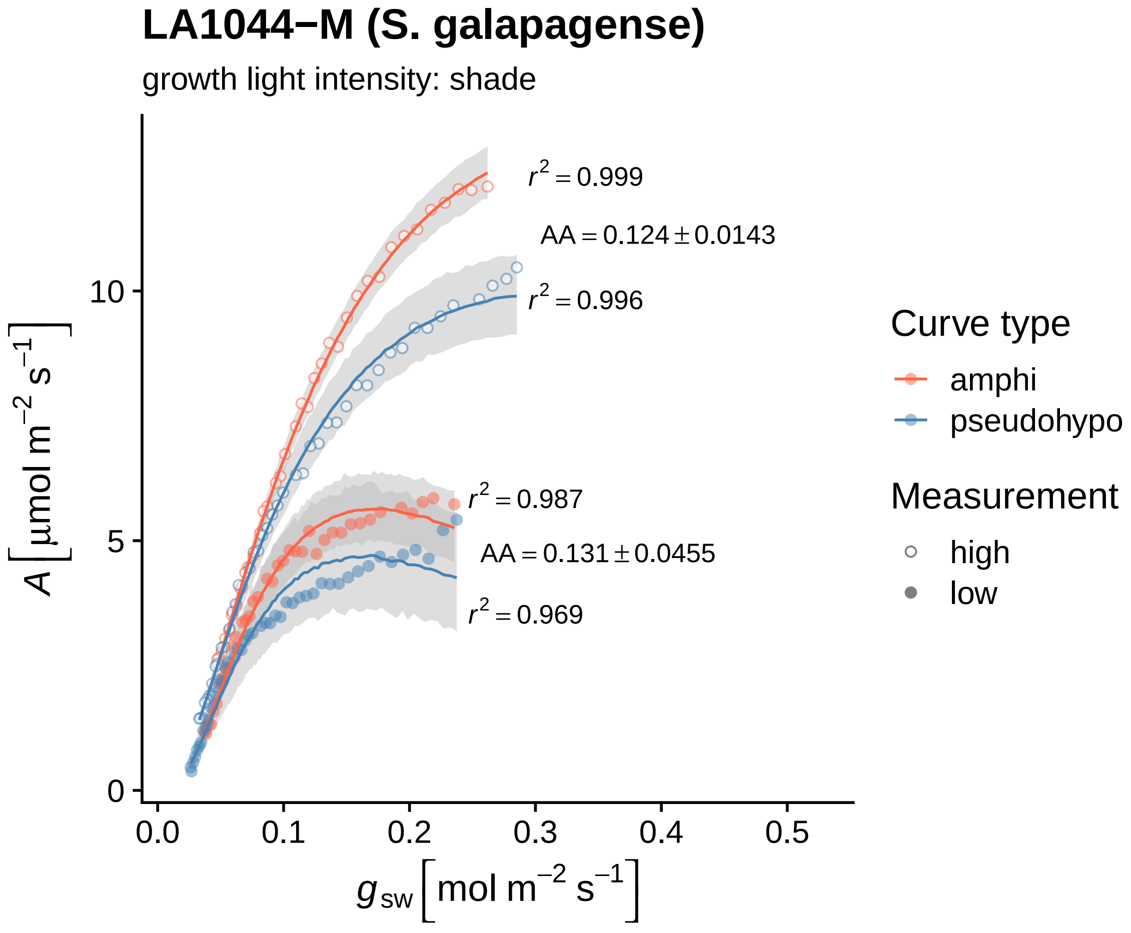

LA1044−M (S. galapagense)
growth light intensity: shade
$r^2 = 0.999$
$AA = 0.124 \pm 0.0143$
$r^2 = 0.996$
$r^2 = 0.987$
$AA = 0.131 \pm 0.0455$
$r^2 = 0.969$
0
5
10
0.0
0.1
0.2
0.3
0.4
0.5
$A\,[\mu mol\ m^{-2}\ s^{-1}]$
$g_{sw}\,[mol\ m^{-2}\ s^{-1}]$
Curve type
amphi
pseudohypo
Measurement
high
low

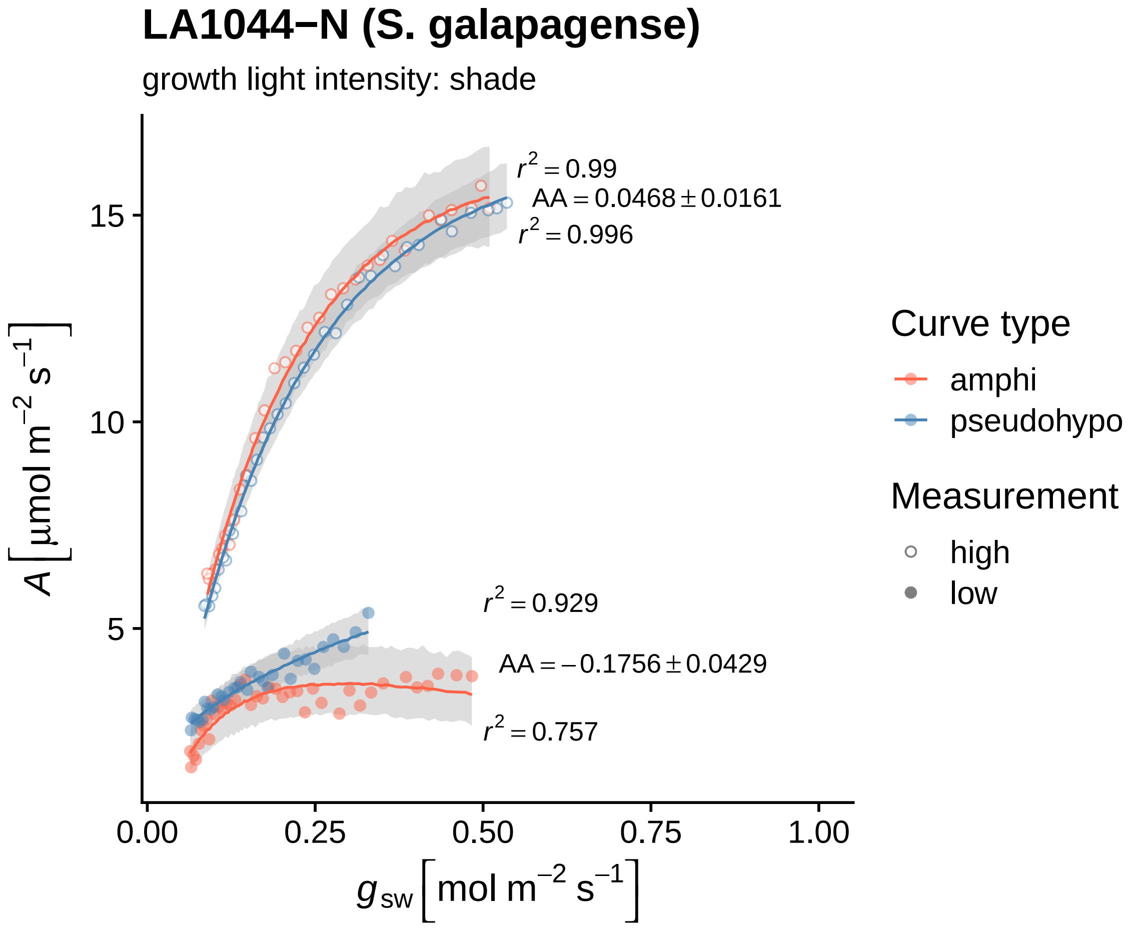
LA1044−N (S. galapagense)
growth light intensity: shade
r² = 0.99
AA = 0.0468 ± 0.0161
r² = 0.996
r² = 0.929
AA = −0.1756 ± 0.0429
r² = 0.757
A [µmol m⁻² s⁻¹]
15
10
5
0.00
0.25
0.50
0.75
1.00
g_sw [mol m⁻² s⁻¹]
Curve type
amphi
pseudohypo
Measurement
high
low

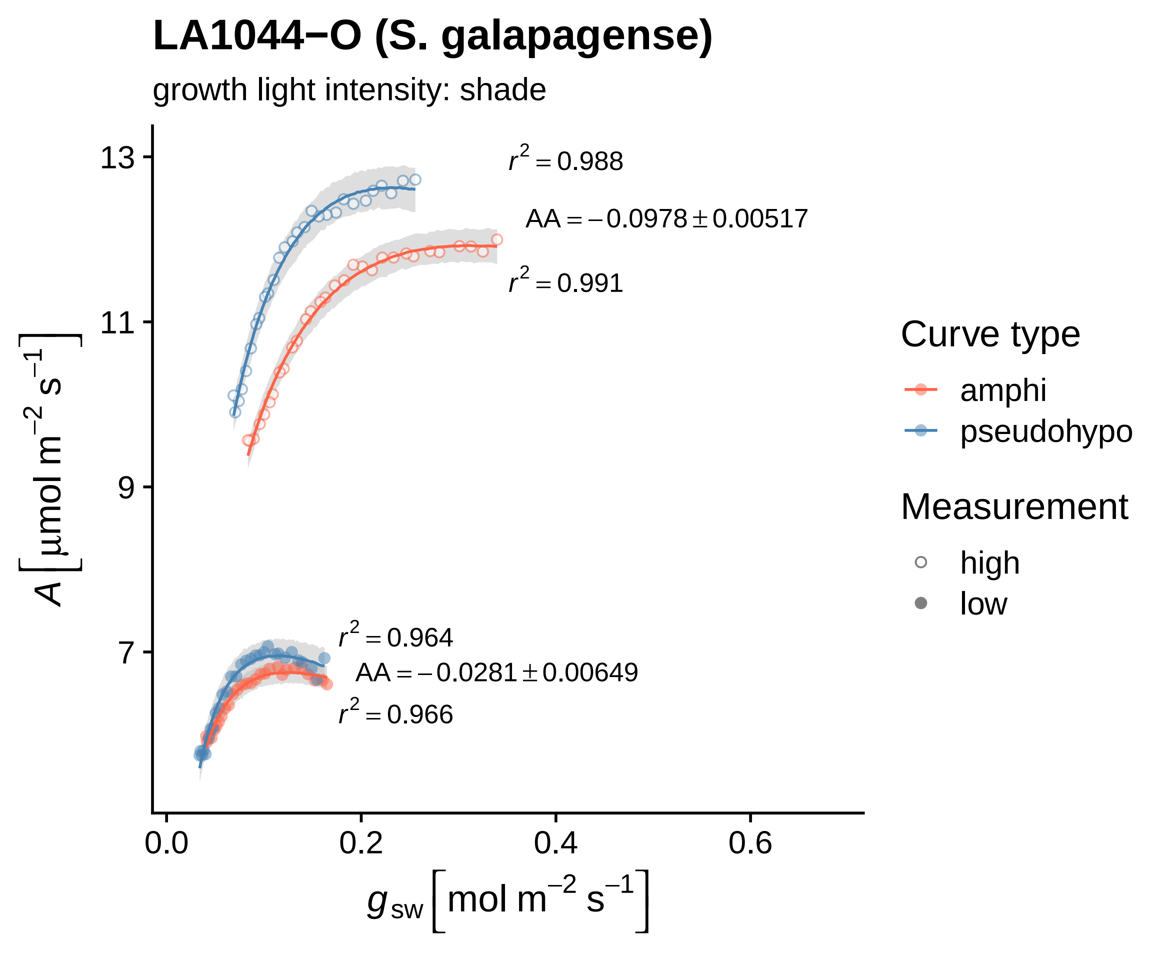
LA1044−O (S. galapagense)
growth light intensity: shade
$r^2 = 0.988$
$AA = -0.0978 \pm 0.00517$
$r^2 = 0.991$
$r^2 = 0.964$
$AA = -0.0281 \pm 0.00649$
$r^2 = 0.966$
13
11
9
7
0.0
0.2
0.4
0.6
$A\,[\mu mol\ m^{-2}\ s^{-1}]$
$g_{sw}\,[mol\ m^{-2}\ s^{-1}]$
Curve type
amphi
pseudohypo
Measurement
high
low

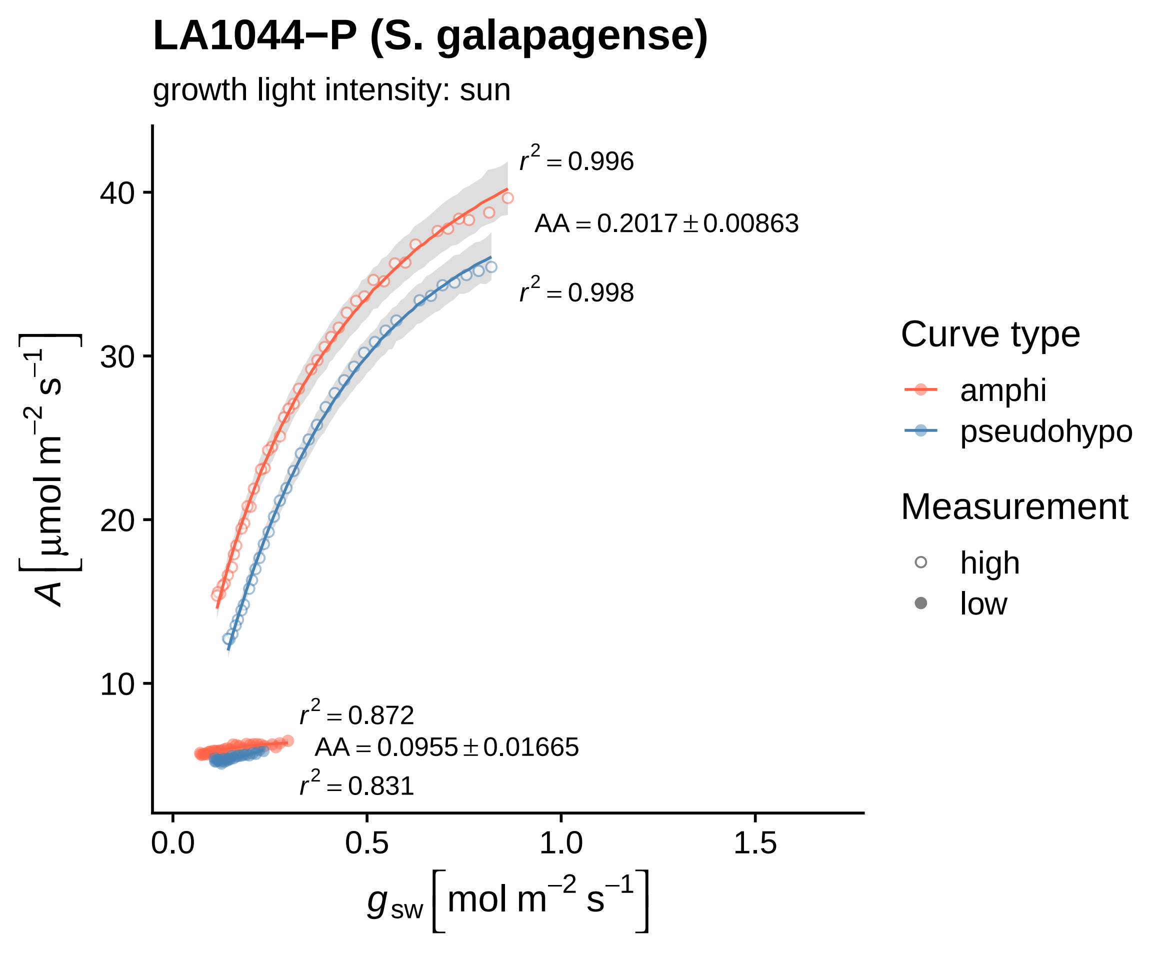
LA1044−P (S. galapagense)
growth light intensity: sun
r² = 0.996
AA = 0.2017 ± 0.00863
r² = 0.998
r² = 0.872
AA = 0.0955 ± 0.01665
r² = 0.831
A [µmol m⁻² s⁻¹]
g_sw [mol m⁻² s⁻¹]
0.0
0.5
1.0
1.5
10
20
30
40
Curve type
amphi
pseudohypo
Measurement
high
low

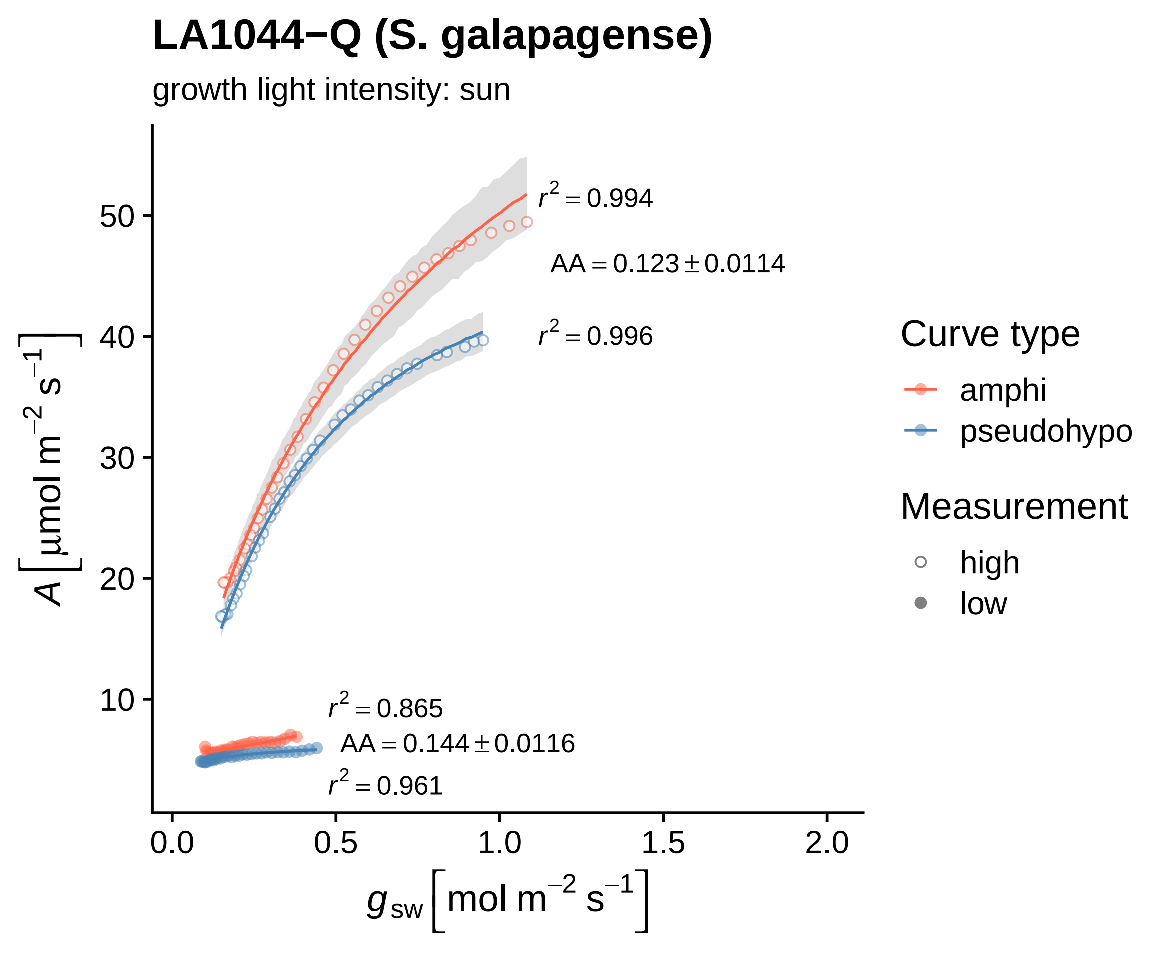

LA1044−Q (S. galapagense)
growth light intensity: sun
r2 = 0.994
AA = 0.123 ± 0.0114
r2 = 0.996
r2 = 0.865
AA = 0.144 ± 0.0116
r2 = 0.961
A [µmol m−2 s−1]
gsw [mol m−2 s−1]
50
40
30
20
10
0.0
0.5
1.0
1.5
2.0
Curve type
amphi
pseudohypo
Measurement
high
low

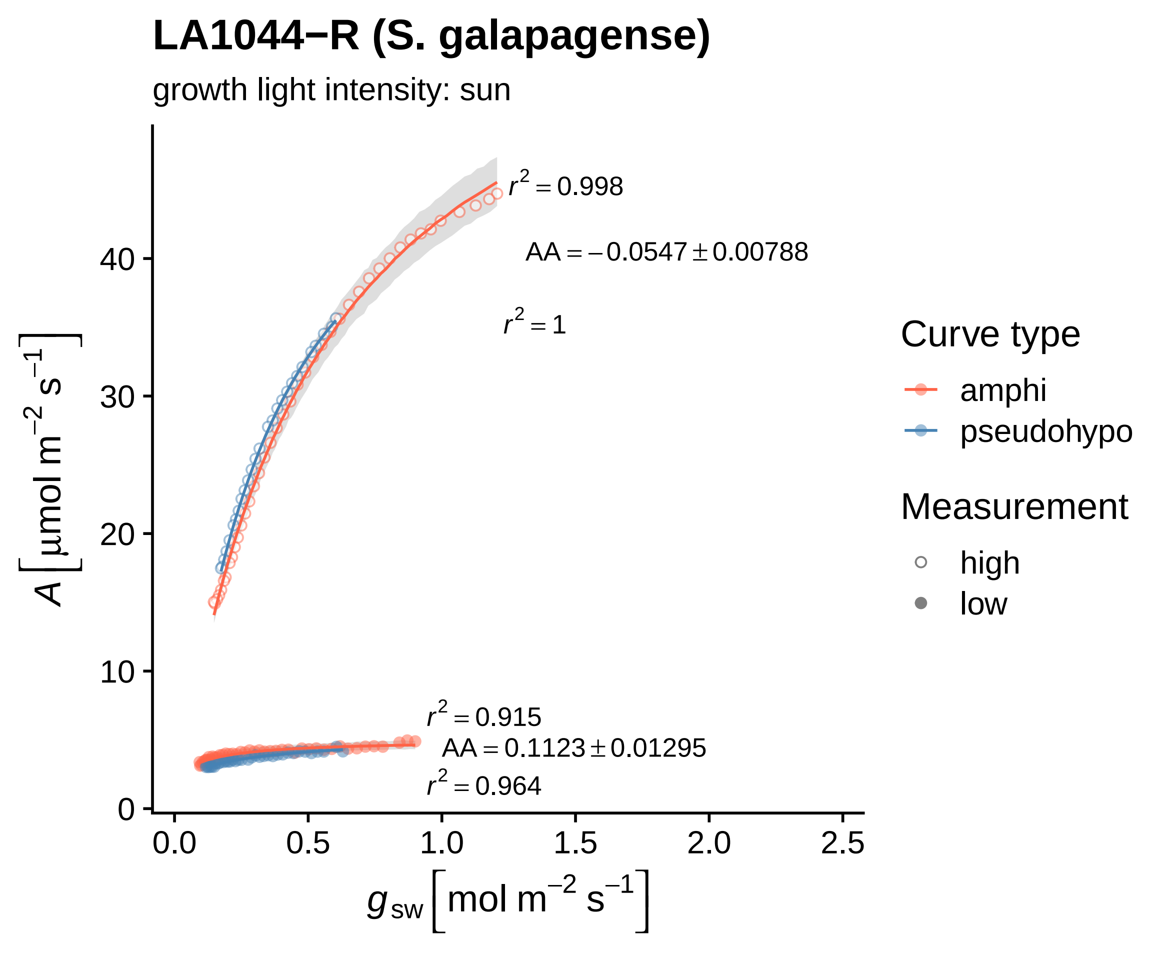
LA1044−R (S. galapagense)
growth light intensity: sun
$r^2 = 0.998$
$AA = -0.0547 \pm 0.00788$
$r^2 = 1$
$r^2 = 0.915$
$AA = 0.1123 \pm 0.01295$
$r^2 = 0.964$
$A\ [\mu mol\ m^{-2}\ s^{-1}]$
0
10
20
30
40
0.0
0.5
1.0
1.5
2.0
2.5
$g_{sw}\ [mol\ m^{-2}\ s^{-1}]$
Curve type
amphi
pseudohypo
Measurement
high
low

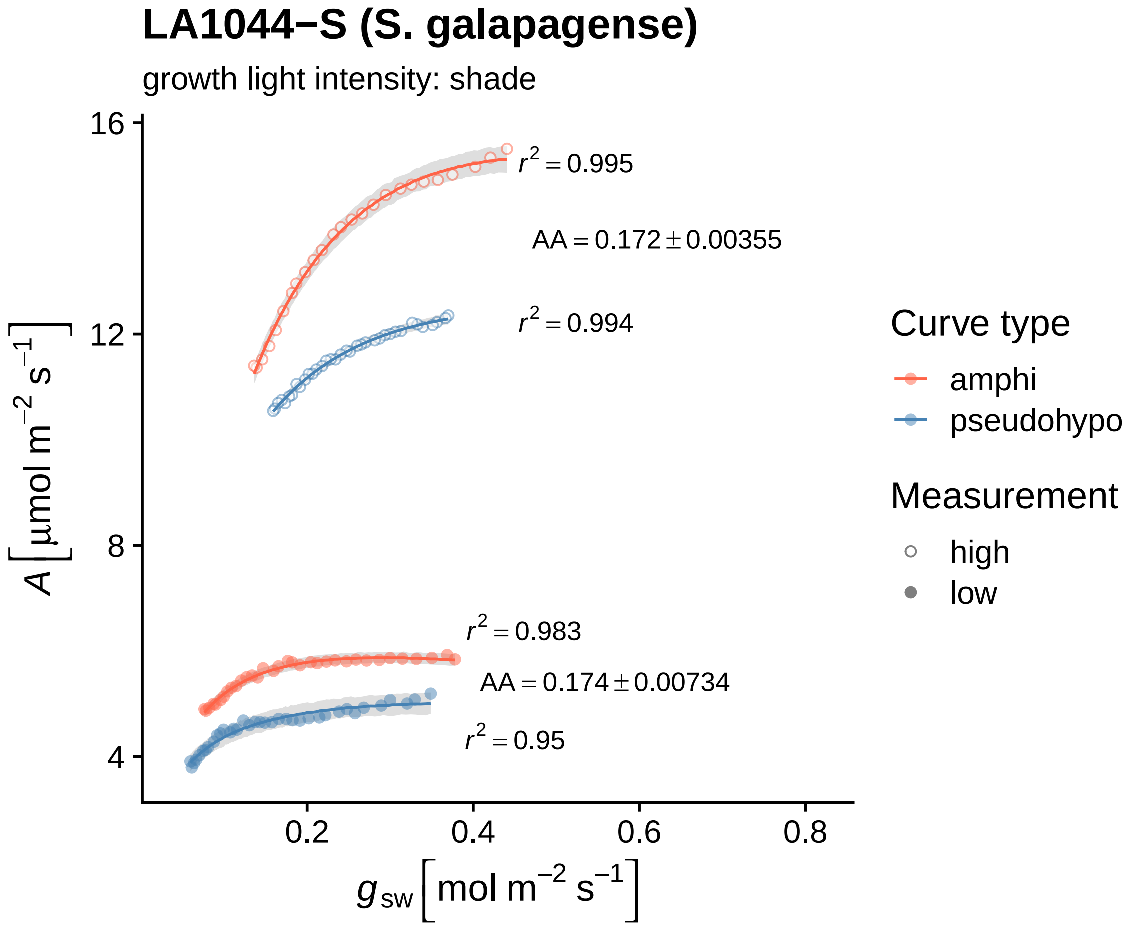
LA1044−S (S. galapagense)
growth light intensity: shade
r² = 0.995
AA = 0.172 ± 0.00355
r² = 0.994
r² = 0.983
AA = 0.174 ± 0.00734
r² = 0.95
16
12
8
4
0.2
0.4
0.6
0.8
A [µmol m⁻² s⁻¹]
g sw [mol m⁻² s⁻¹]
Curve type
amphi
pseudohypo
Measurement
high
low

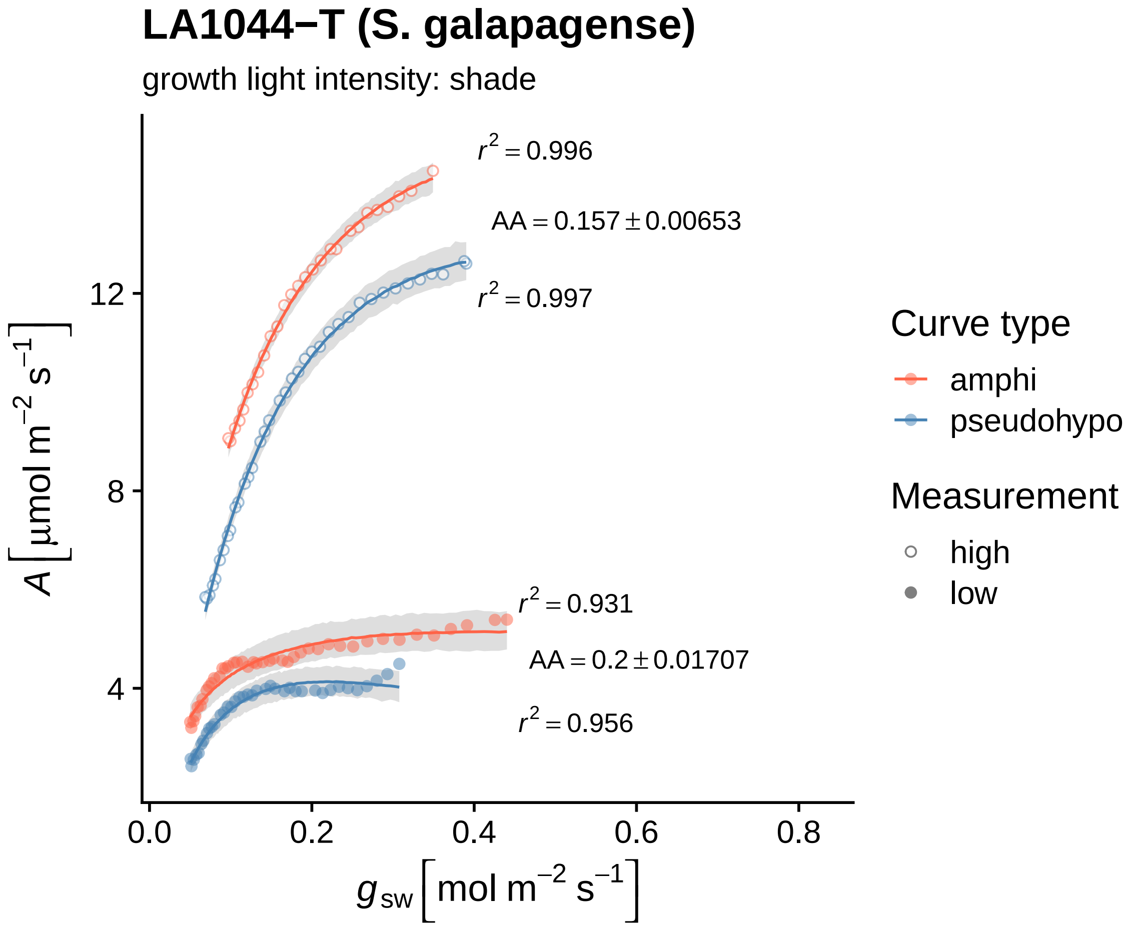

LA1044−T (S. galapagense)
growth light intensity: shade
$r^2 = 0.996$
AA = 0.157 ± 0.00653
$r^2 = 0.997$
$r^2 = 0.931$
AA = 0.2 ± 0.01707
$r^2 = 0.956$
12
8
4
0.0
0.2
0.4
0.6
0.8
$A\ [\mu mol\ m^{-2}\ s^{-1}]$
$g_{sw}\ [mol\ m^{-2}\ s^{-1}]$
Curve type
amphi
pseudohypo
Measurement
high
low

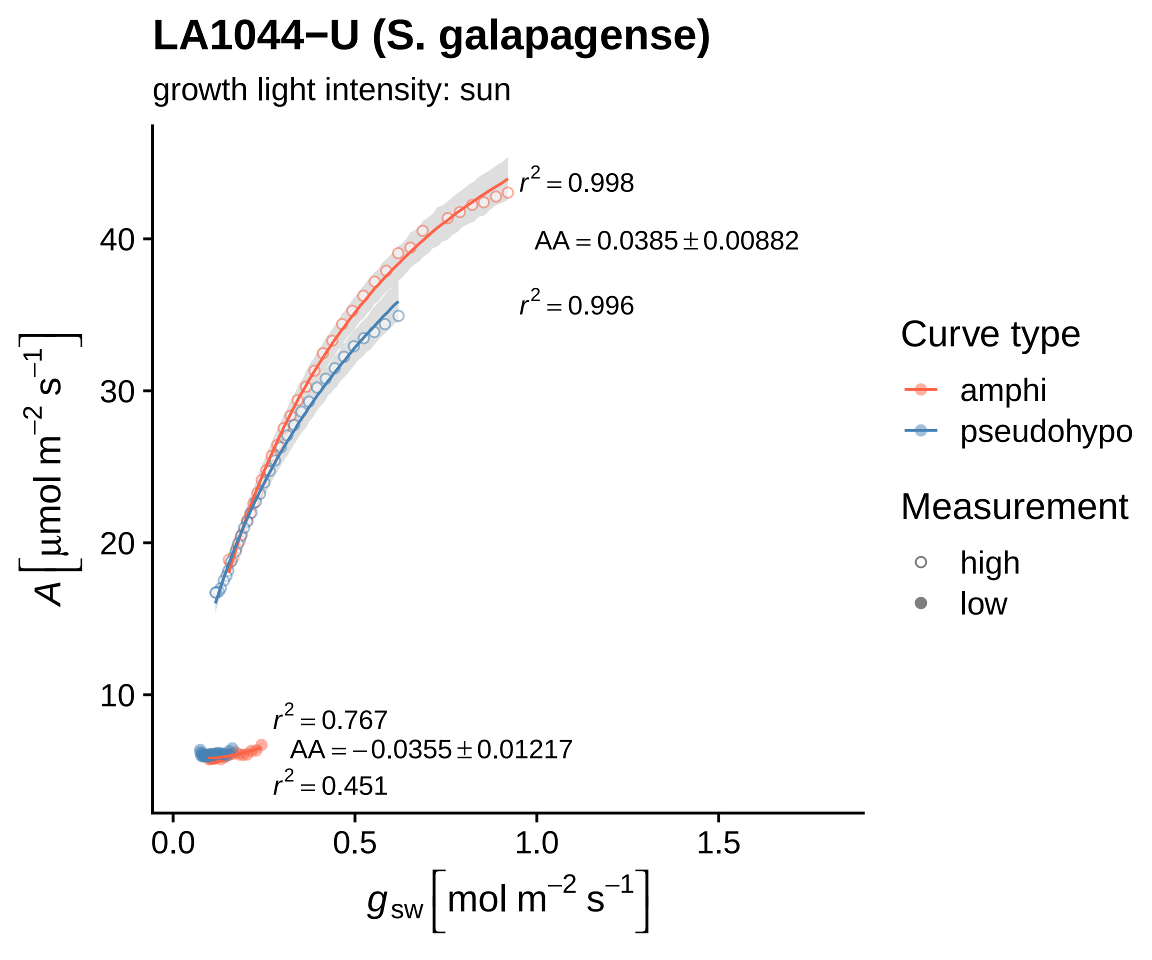
LA1044−U (S. galapagense)
growth light intensity: sun
$r^2 = 0.998$
$AA = 0.0385 \pm 0.00882$
$r^2 = 0.996$
$r^2 = 0.767$
$AA = -0.0355 \pm 0.01217$
$r^2 = 0.451$
$A\,[\mu mol\ m^{-2}\ s^{-1}]$
$g_{sw}\,[mol\ m^{-2}\ s^{-1}]$
10
20
30
40
0.0
0.5
1.0
1.5
Curve type
amphi
pseudohypo
Measurement
high
low

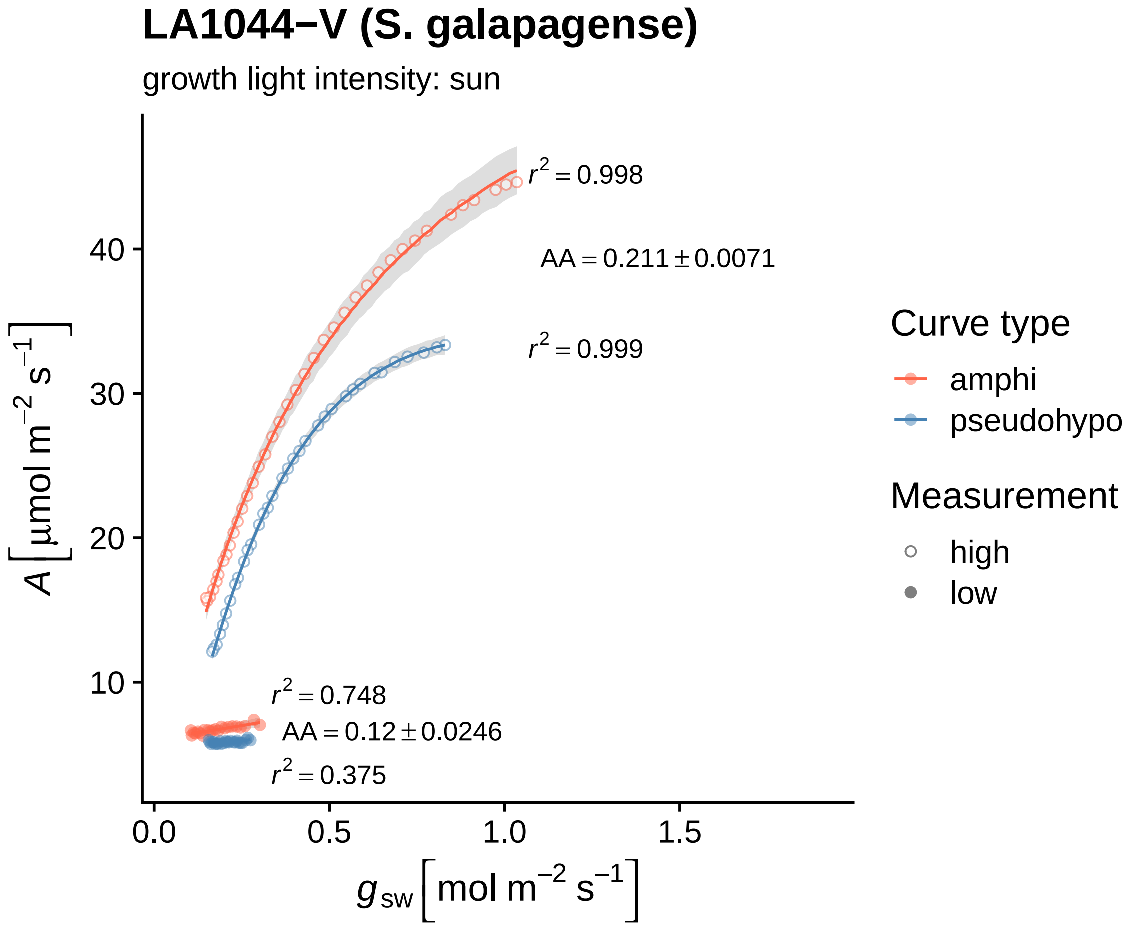

LA1044−V (S. galapagense)
growth light intensity: sun
$r^2 = 0.998$
$AA = 0.211 \pm 0.0071$
$r^2 = 0.999$
$r^2 = 0.748$
$AA = 0.12 \pm 0.0246$
$r^2 = 0.375$
$A\left[\mu mol\ m^{-2}\ s^{-1}\right]$
10
20
30
40
0.0
0.5
1.0
1.5
$g_{sw}\left[mol\ m^{-2}\ s^{-1}\right]$
Curve type
amphi
pseudohypo
Measurement
high
low

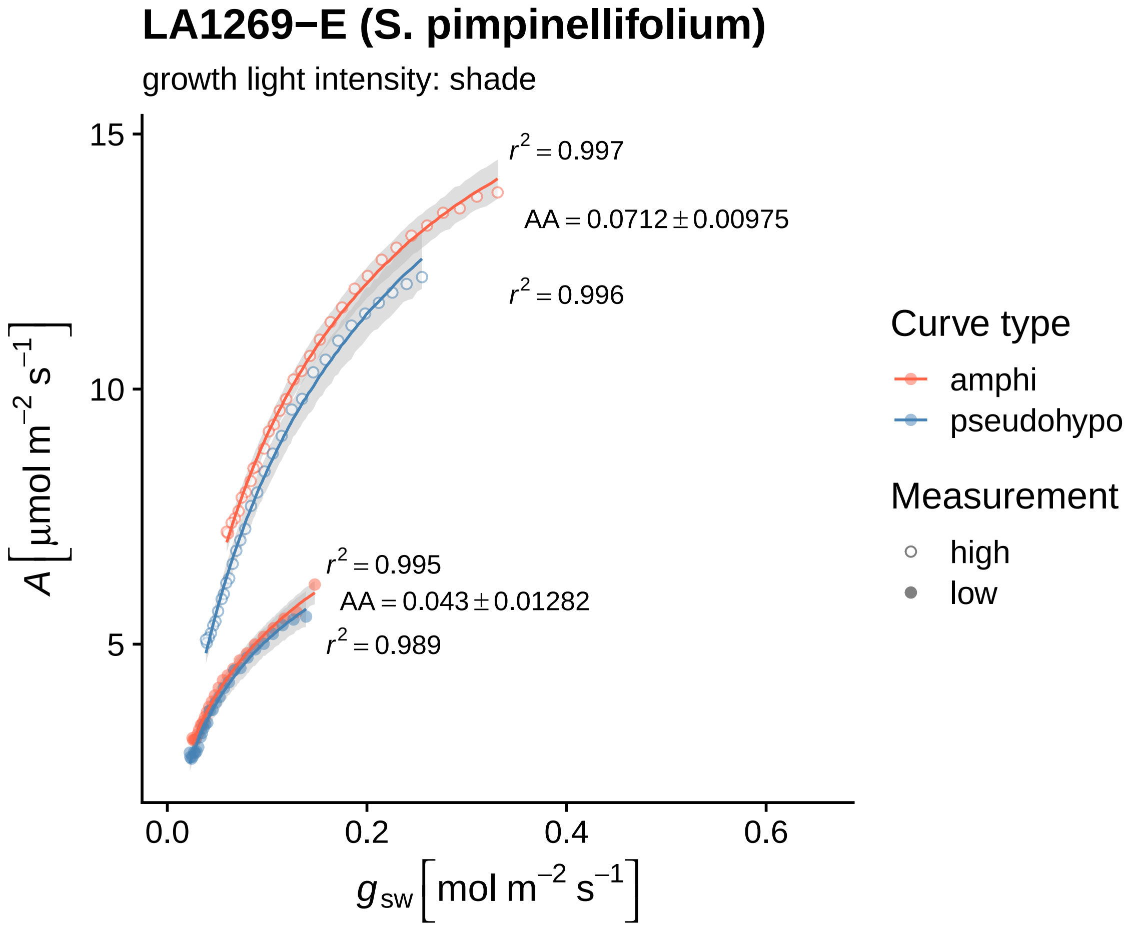

LA1269−E (S. pimpinellifolium)
growth light intensity: shade
$r^2 = 0.997$
$AA = 0.0712 \pm 0.00975$
$r^2 = 0.996$
$r^2 = 0.995$
$AA = 0.043 \pm 0.01282$
$r^2 = 0.989$
15
10
5
0.0
0.2
0.4
0.6
$A\left[\mu mol\ m^{-2}\ s^{-1}\right]$
$g_{sw}\left[mol\ m^{-2}\ s^{-1}\right]$
Curve type
amphi
pseudohypo
Measurement
high
low

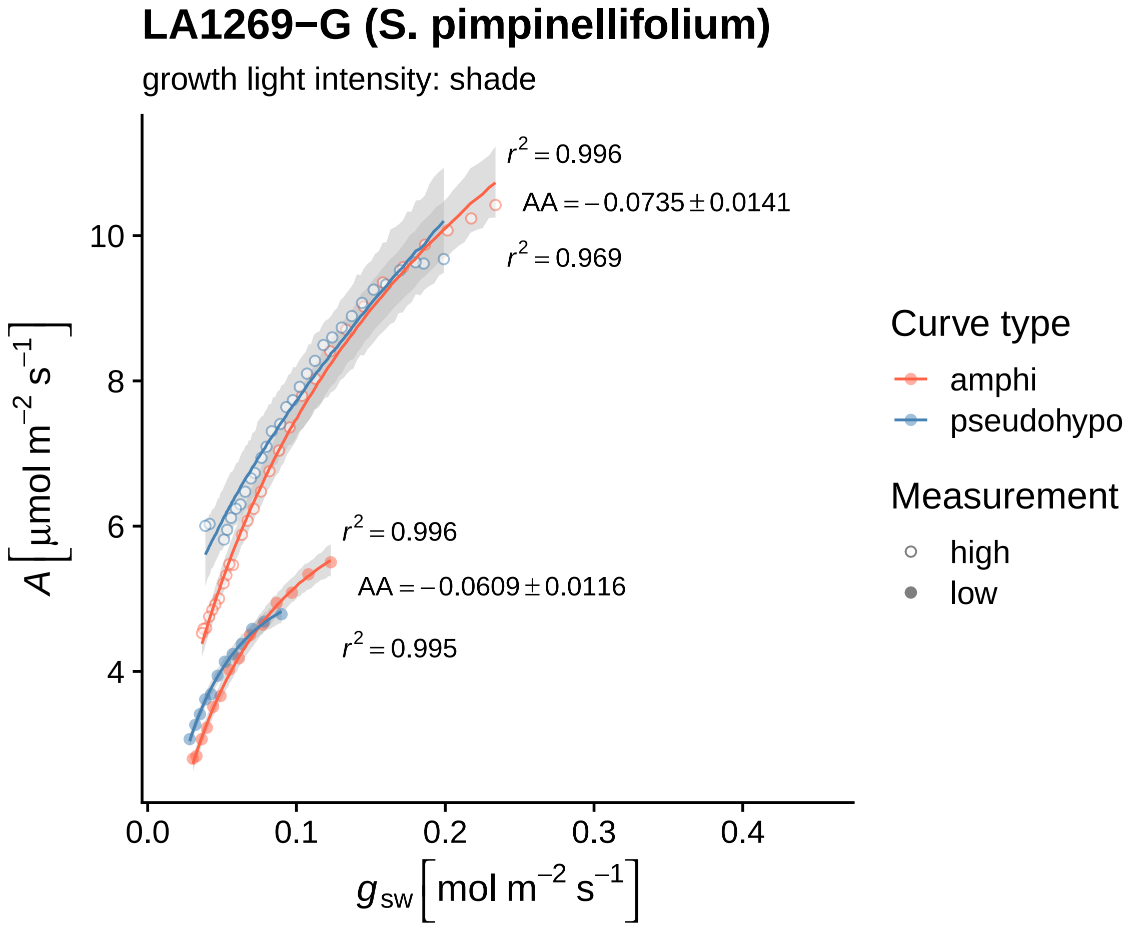

LA1269−G (S. pimpinellifolium)
growth light intensity: shade
r² = 0.996
AA = −0.0735 ± 0.0141
r² = 0.969
r² = 0.996
AA = −0.0609 ± 0.0116
r² = 0.995
A [μmol m⁻² s⁻¹]
10
8
6
4
0.0
0.1
0.2
0.3
0.4
gsw [mol m⁻² s⁻¹]
Curve type
amphi
pseudohypo
Measurement
high
low

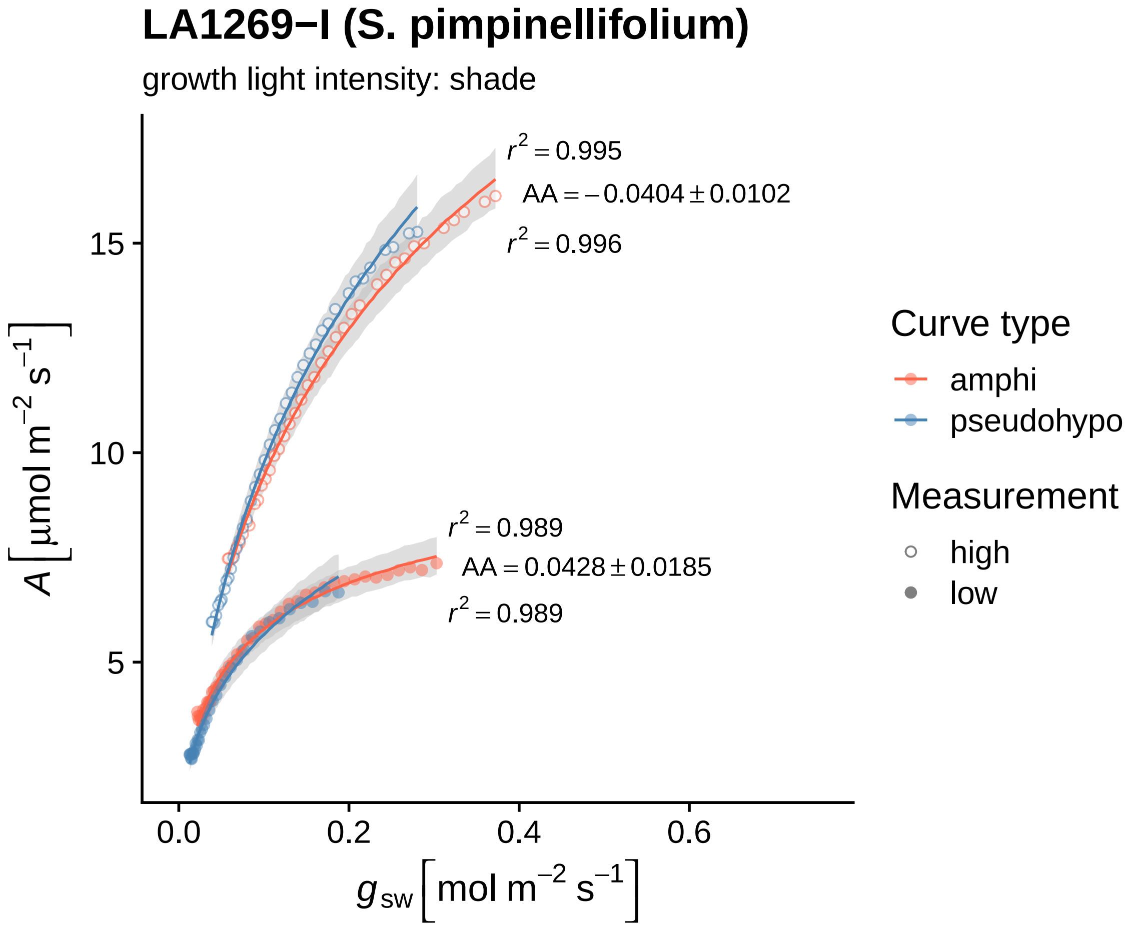
LA1269−I (S. pimpinellifolium)
growth light intensity: shade
$r^2 = 0.995$
$AA = -0.0404 \pm 0.0102$
$r^2 = 0.996$
$r^2 = 0.989$
$AA = 0.0428 \pm 0.0185$
$r^2 = 0.989$
15
10
5
0.0
0.2
0.4
0.6
$A\ [\mu mol\ m^{-2}\ s^{-1}]$
$g_{sw}\ [mol\ m^{-2}\ s^{-1}]$
Curve type
amphi
pseudohypo
Measurement
high
low

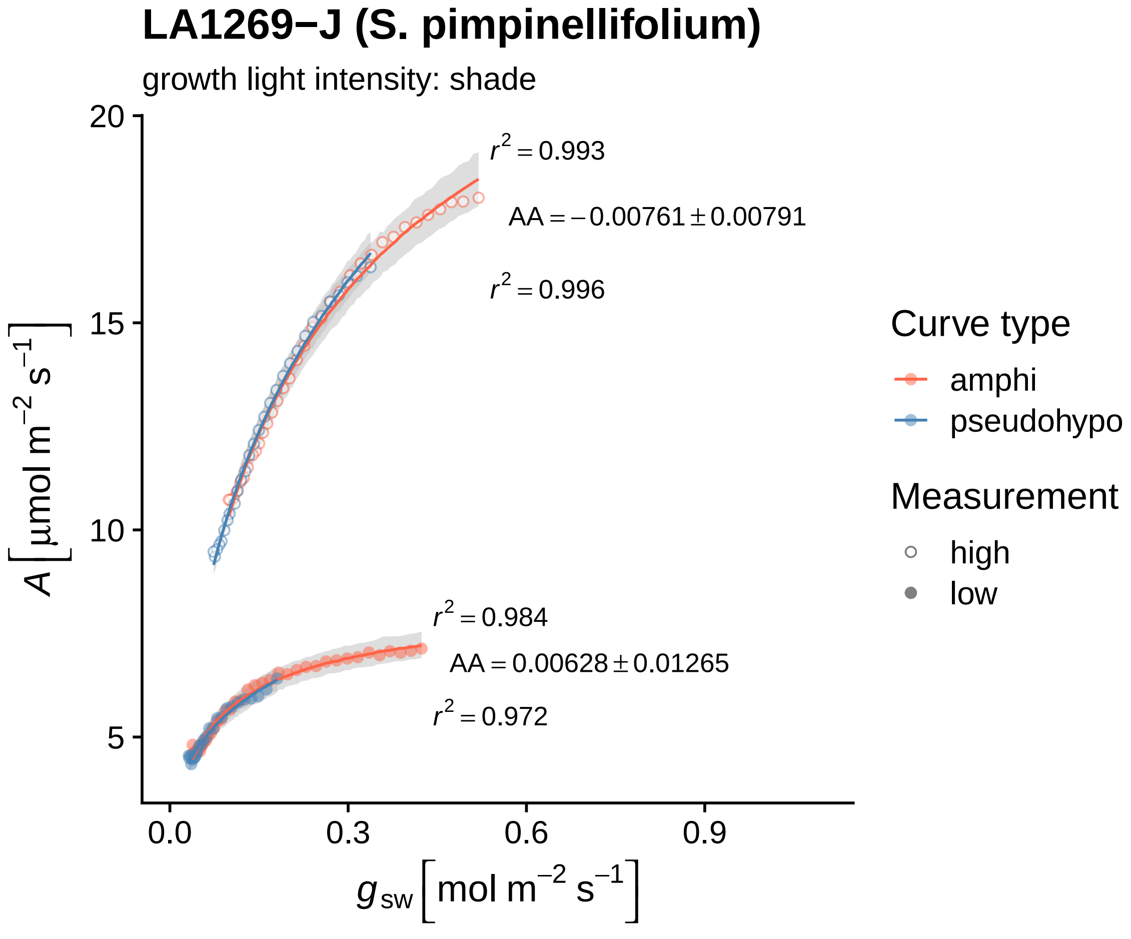

LA1269−J (S. pimpinellifolium)
growth light intensity: shade
$r^2 = 0.993$
$AA = -0.00761 \pm 0.00791$
$r^2 = 0.996$
$r^2 = 0.984$
$AA = 0.00628 \pm 0.01265$
$r^2 = 0.972$
20
15
10
5
0.0
0.3
0.6
0.9
$A\left[\mu mol\ m^{-2}\ s^{-1}\right]$
$g_{sw}\left[mol\ m^{-2}\ s^{-1}\right]$
Curve type
amphi
pseudohypo
Measurement
high
low

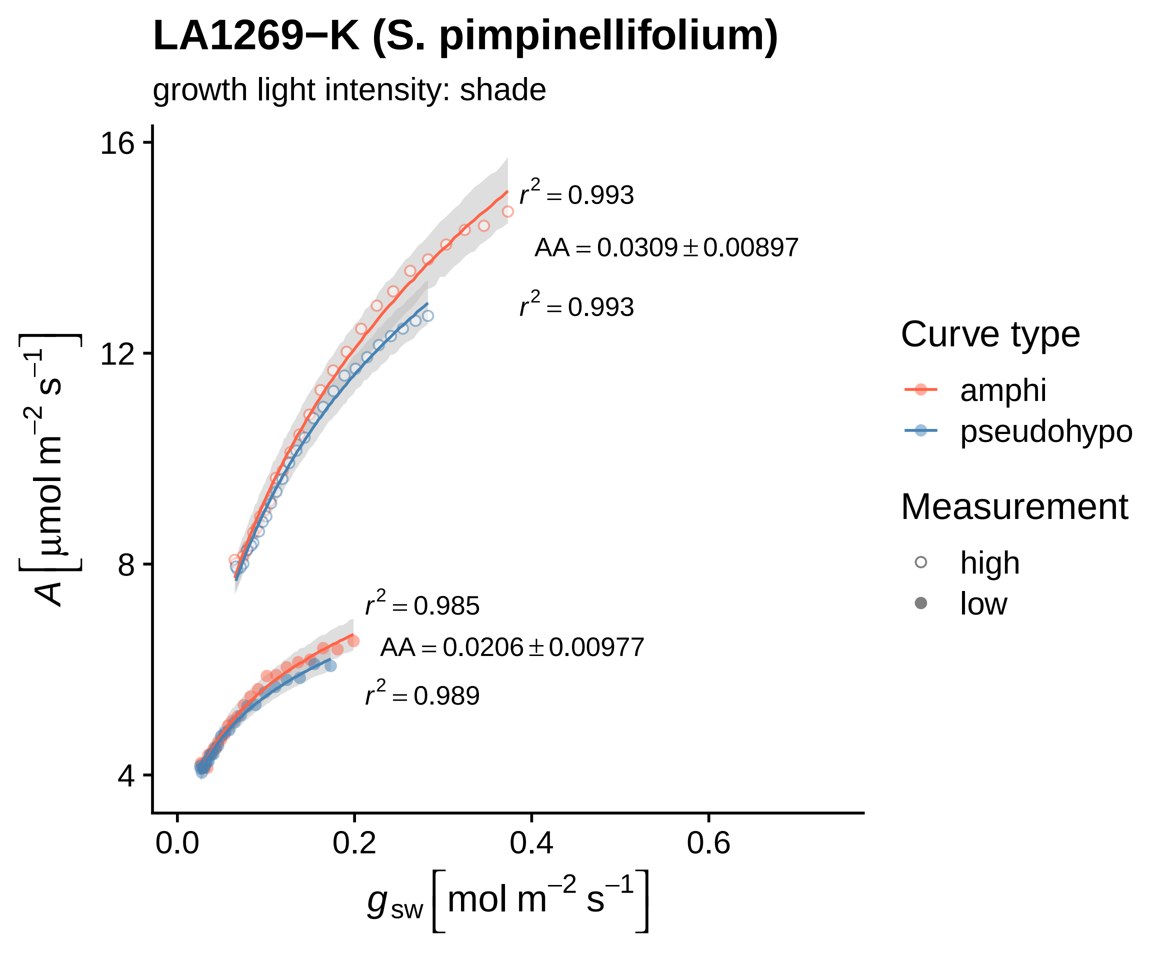

LA1269−K (S. pimpinellifolium)
growth light intensity: shade
$r^2 = 0.993$
$AA = 0.0309 \pm 0.00897$
$r^2 = 0.993$
$r^2 = 0.985$
$AA = 0.0206 \pm 0.00977$
$r^2 = 0.989$
16
12
8
4
0.0
0.2
0.4
0.6
$A\left[\mu mol\ m^{-2}\ s^{-1}\right]$
$g_{sw}\left[mol\ m^{-2}\ s^{-1}\right]$
Curve type
amphi
pseudohypo
Measurement
high
low

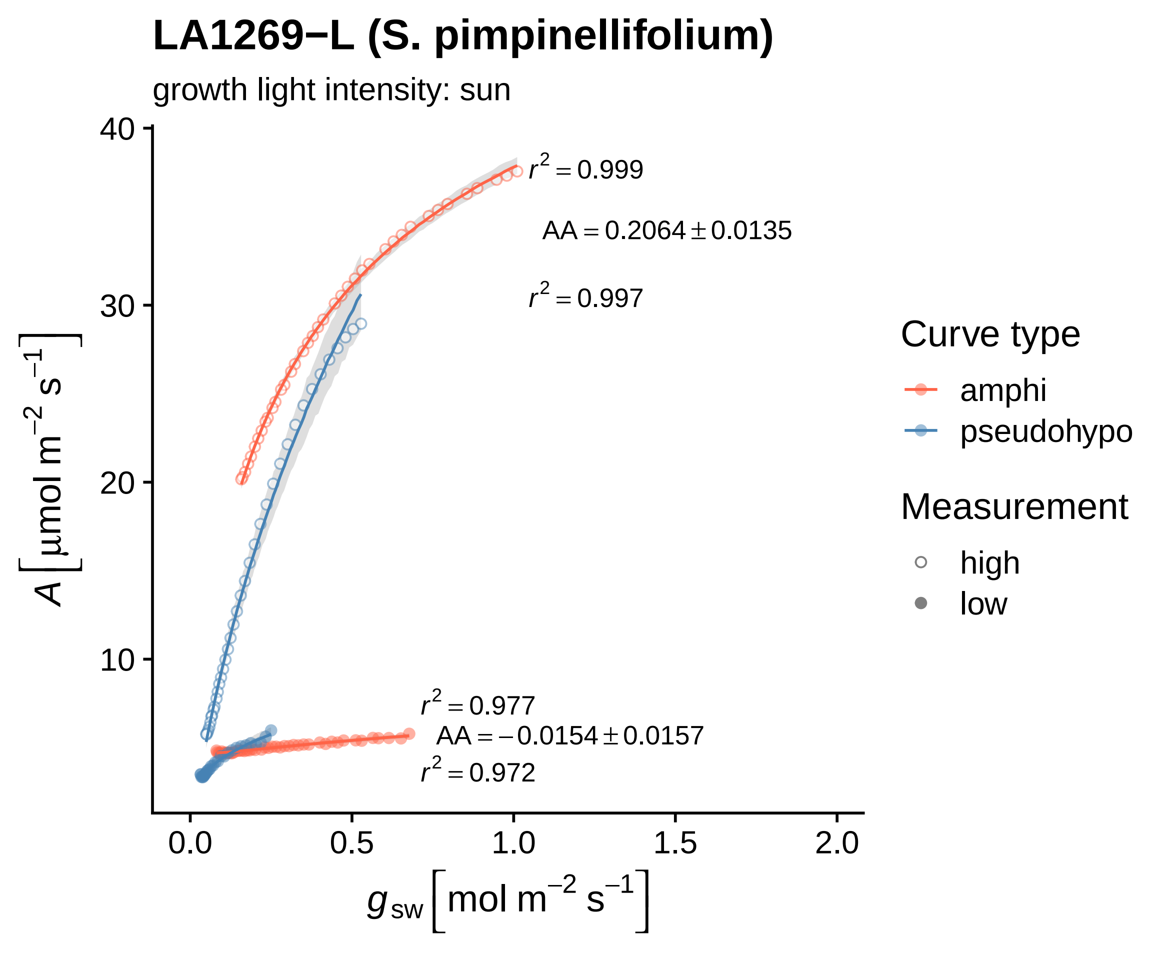
LA1269−L (S. pimpinellifolium)
growth light intensity: sun
$r^2 = 0.999$
AA = 0.2064 ± 0.0135
$r^2 = 0.997$
$r^2 = 0.977$
AA = −0.0154 ± 0.0157
$r^2 = 0.972$
40
30
20
10
0.0
0.5
1.0
1.5
2.0
$A \left[\mu mol\ m^{-2}\ s^{-1}\right]$
$g_{sw} \left[mol\ m^{-2}\ s^{-1}\right]$
Curve type
amphi
pseudohypo
Measurement
high
low

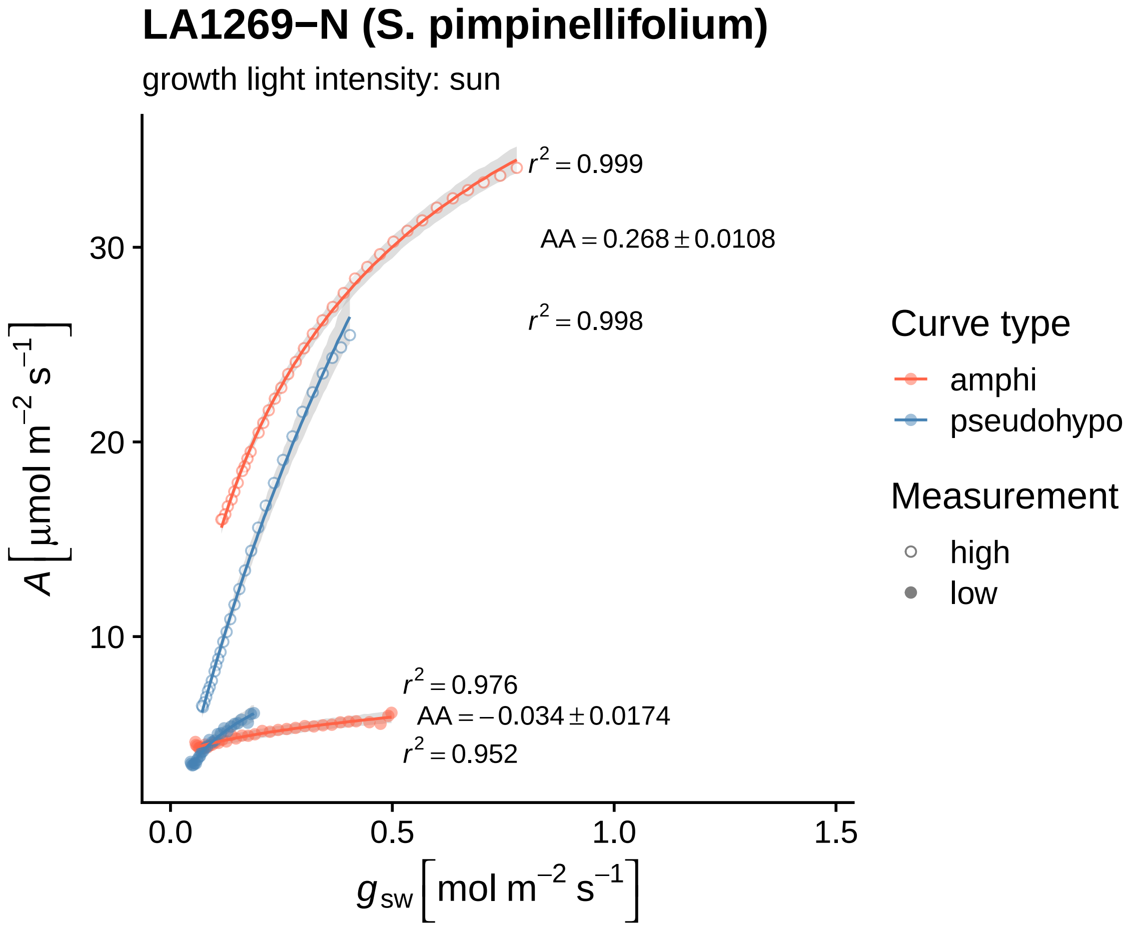
LA1269−N (S. pimpinellifolium)
growth light intensity: sun
r² = 0.999
AA = 0.268 ± 0.0108
r² = 0.998
r² = 0.976
AA = −0.034 ± 0.0174
r² = 0.952
30
20
10
0.0
0.5
1.0
1.5
A [µmol m⁻² s⁻¹]
g sw [mol m⁻² s⁻¹]
Curve type
amphi
pseudohypo
Measurement
high
low

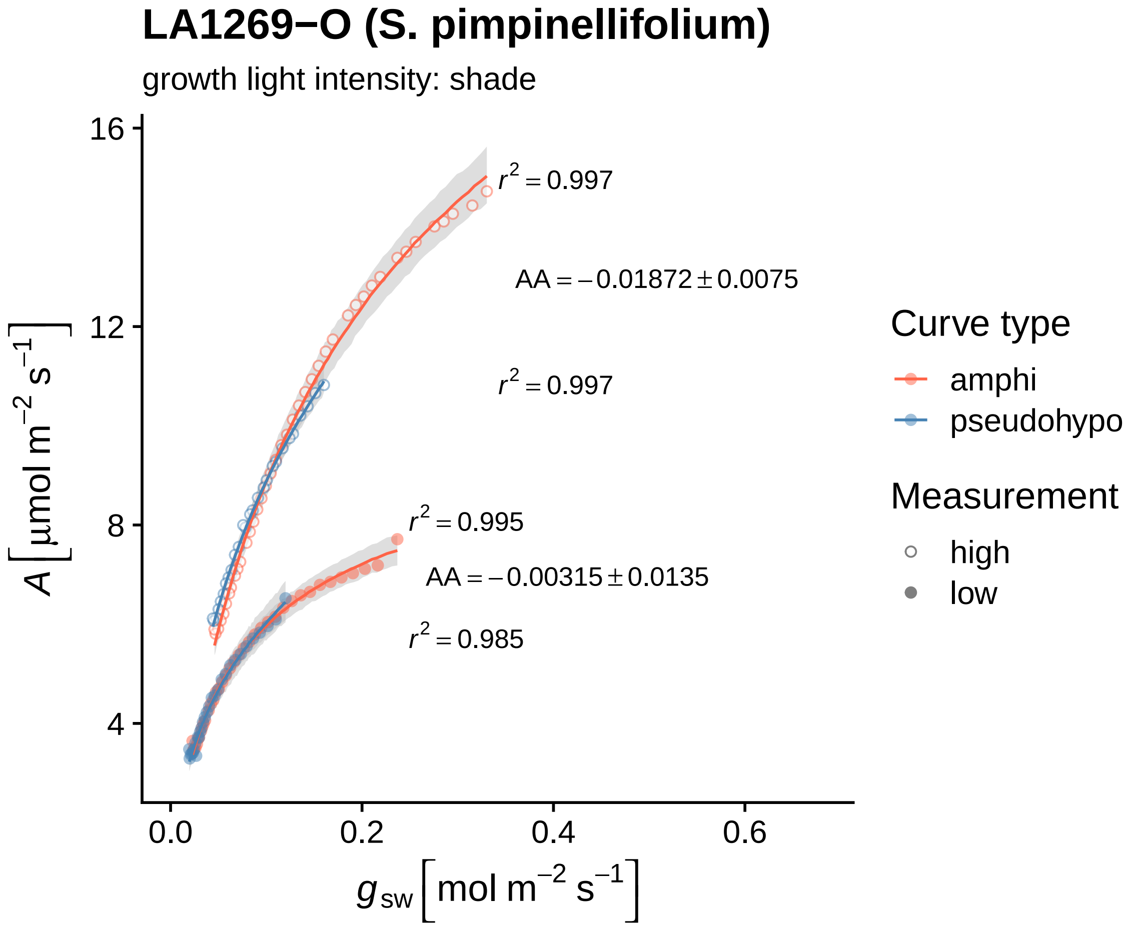

LA1269−O (S. pimpinellifolium)
growth light intensity: shade
$r^2 = 0.997$
AA = −0.01872 ± 0.0075
$r^2 = 0.997$
$r^2 = 0.995$
AA = −0.00315 ± 0.0135
$r^2 = 0.985$
16
12
8
4
0.0
0.2
0.4
0.6
$A\,[\mu mol\ m^{-2}\ s^{-1}]$
$g_{sw}\,[mol\ m^{-2}\ s^{-1}]$
Curve type
amphi
pseudohypo
Measurement
high
low

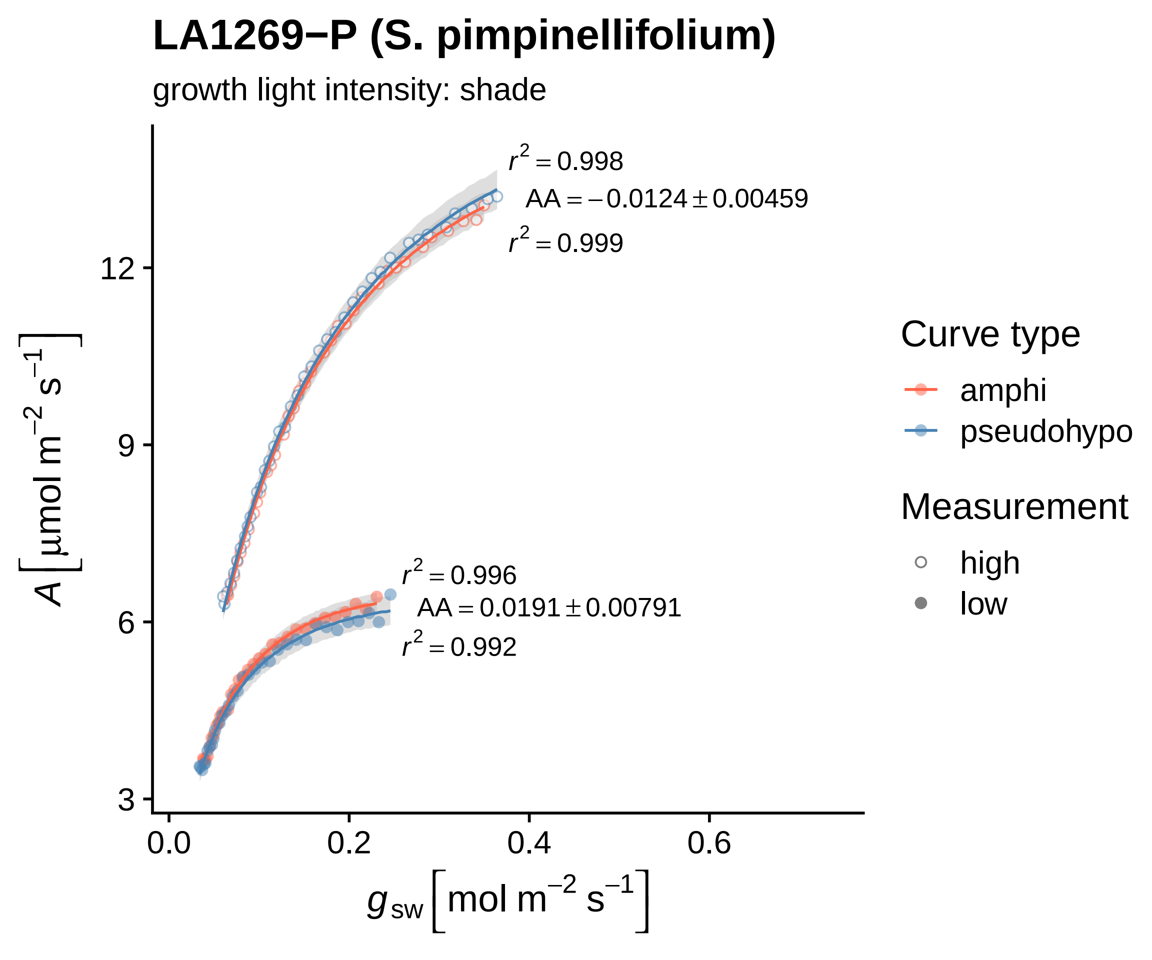

LA1269−P (S. pimpinellifolium)
growth light intensity: shade
$r^2 = 0.998$
$AA = -0.0124 \pm 0.00459$
$r^2 = 0.999$
$r^2 = 0.996$
$AA = 0.0191 \pm 0.00791$
$r^2 = 0.992$
12
9
6
3
0.0
0.2
0.4
0.6
$A\ [\mu mol\ m^{-2}\ s^{-1}]$
$g_{sw}\ [mol\ m^{-2}\ s^{-1}]$
Curve type
amphi
pseudohypo
Measurement
high
low

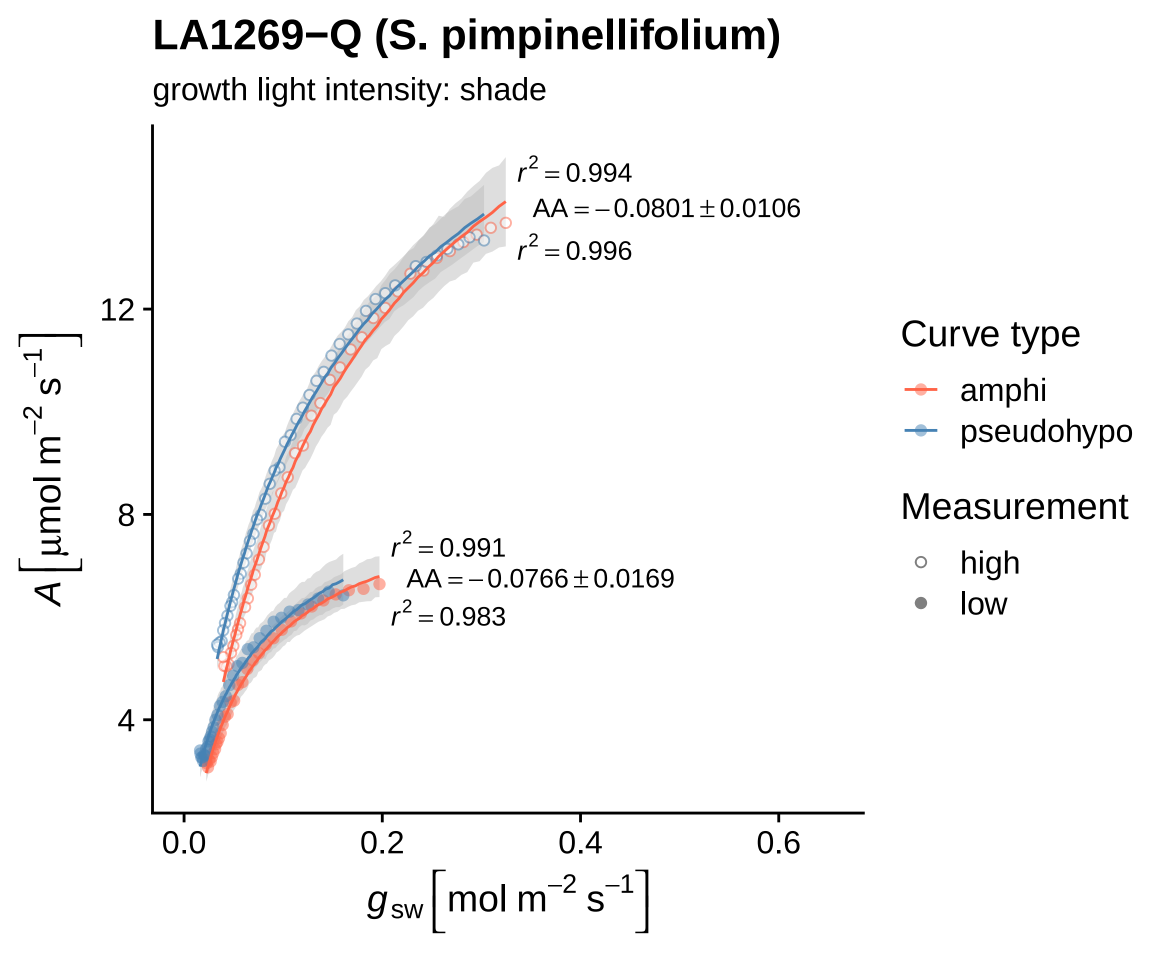
LA1269−Q (S. pimpinellifolium)
growth light intensity: shade
r² = 0.994
AA = −0.0801 ± 0.0106
r² = 0.996
r² = 0.991
AA = −0.0766 ± 0.0169
r² = 0.983
A [µmol m⁻² s⁻¹]
12
8
4
0.0
0.2
0.4
0.6
g_sw [mol m⁻² s⁻¹]
Curve type
amphi
pseudohypo
Measurement
high
low

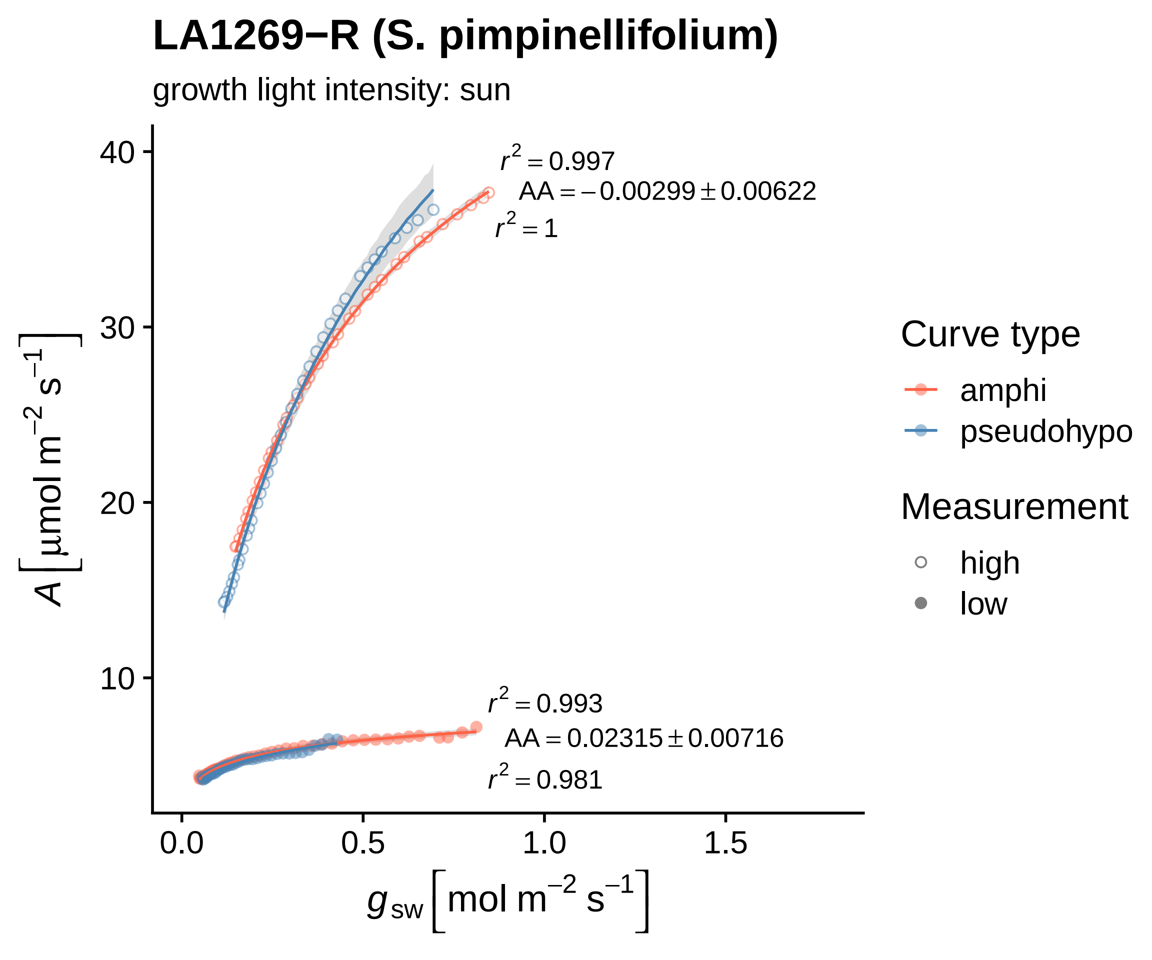
LA1269−R (S. pimpinellifolium)
growth light intensity: sun
$r^2 = 0.997$
$AA = -0.00299 \pm 0.00622$
$r^2 = 1$
$r^2 = 0.993$
$AA = 0.02315 \pm 0.00716$
$r^2 = 0.981$
40
30
20
10
0.0
0.5
1.0
1.5
$A\left[\mu mol\ m^{-2}\ s^{-1}\right]$
$g_{sw}\left[mol\ m^{-2}\ s^{-1}\right]$
Curve type
amphi
pseudohypo
Measurement
high
low

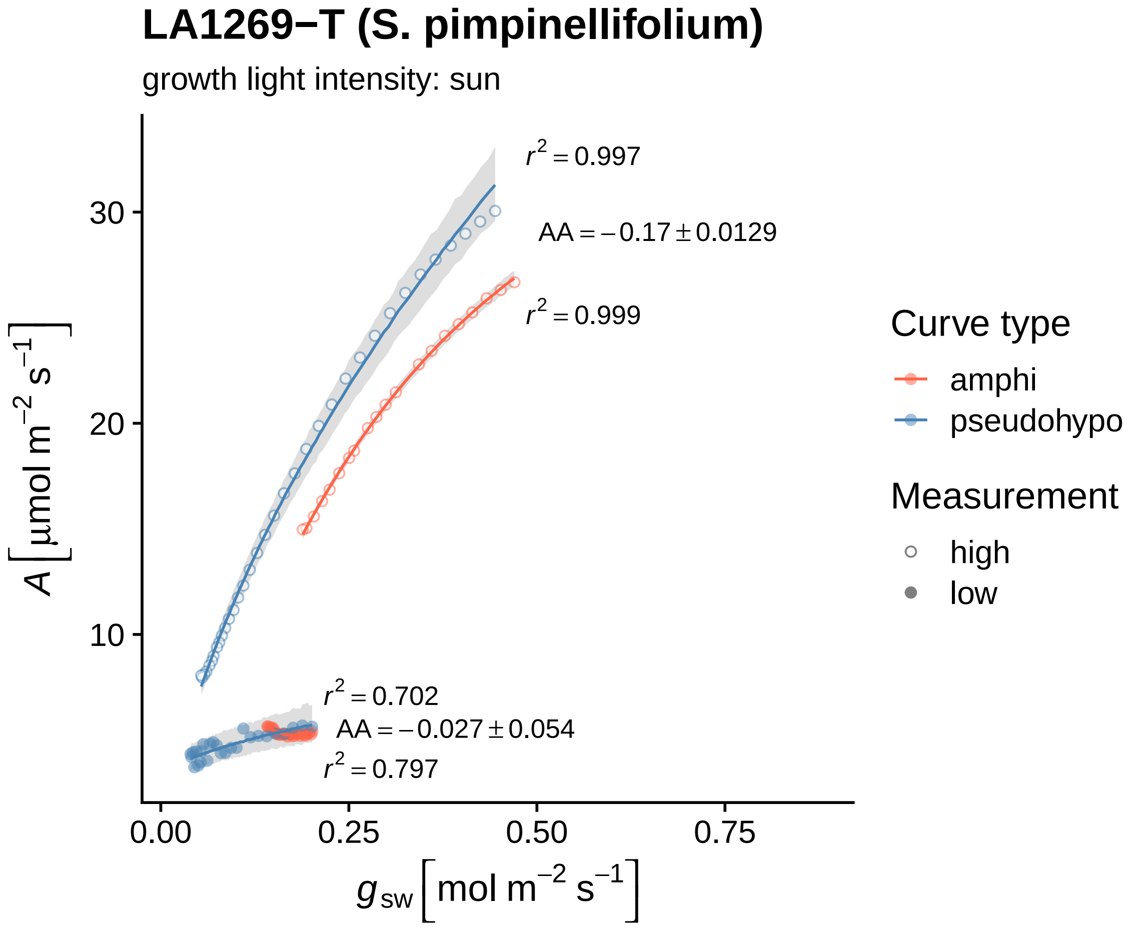
LA1269−T (S. pimpinellifolium)
growth light intensity: sun
$r^2 = 0.997$
$AA = -0.17 \pm 0.0129$
$r^2 = 0.999$
$r^2 = 0.702$
$AA = -0.027 \pm 0.054$
$r^2 = 0.797$
30
20
10
0.00
0.25
0.50
0.75
$A\left[\mu mol\ m^{-2}\ s^{-1}\right]$
$g_{sw}\left[mol\ m^{-2}\ s^{-1}\right]$
Curve type
amphi
pseudohypo
Measurement
high
low

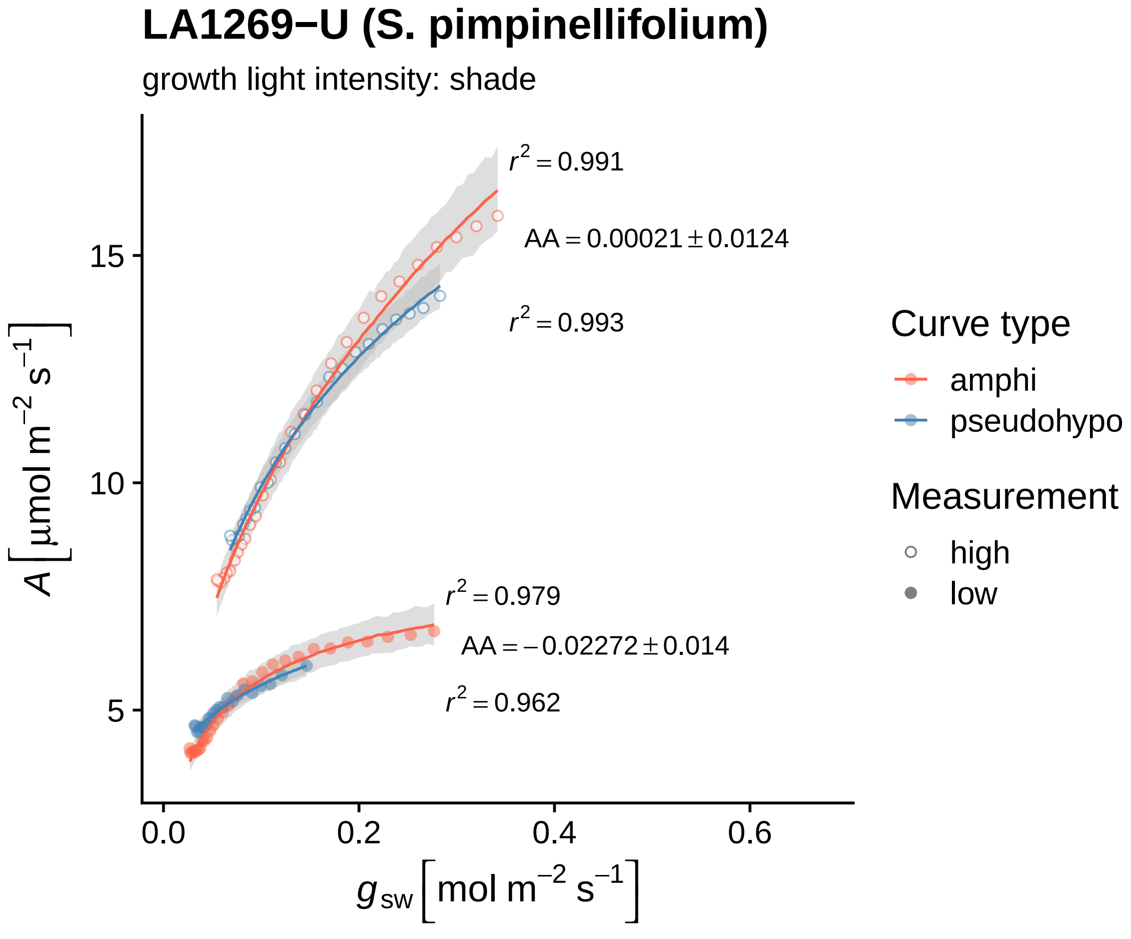

LA1269−U (S. pimpinellifolium)
growth light intensity: shade
r² = 0.991
AA = 0.00021 ± 0.0124
r² = 0.993
r² = 0.979
AA = −0.02272 ± 0.014
r² = 0.962
15
10
5
0.0
0.2
0.4
0.6
A [µmol m⁻² s⁻¹]
g_sw [mol m⁻² s⁻¹]
Curve type
amphi
pseudohypo
Measurement
high
low

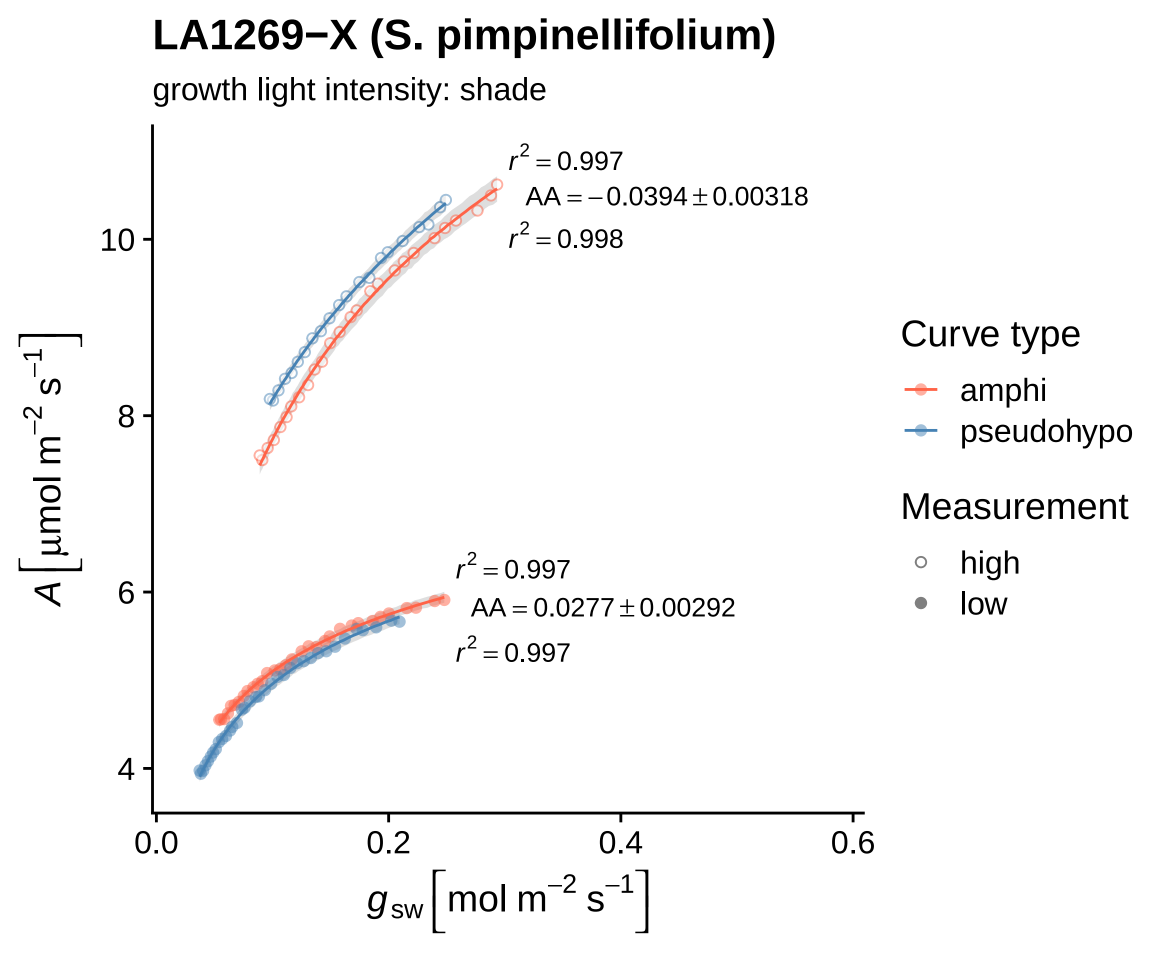
LA1269−X (S. pimpinellifolium)
growth light intensity: shade
$r^2 = 0.997$
$AA = -0.0394 \pm 0.00318$
$r^2 = 0.998$
$r^2 = 0.997$
$AA = 0.0277 \pm 0.00292$
$r^2 = 0.997$
$A\left[\mu mol\ m^{-2}\ s^{-1}\right]$
$g_{sw}\left[mol\ m^{-2}\ s^{-1}\right]$
4
6
8
10
0.0
0.2
0.4
0.6
Curve type
amphi
pseudohypo
Measurement
high
low

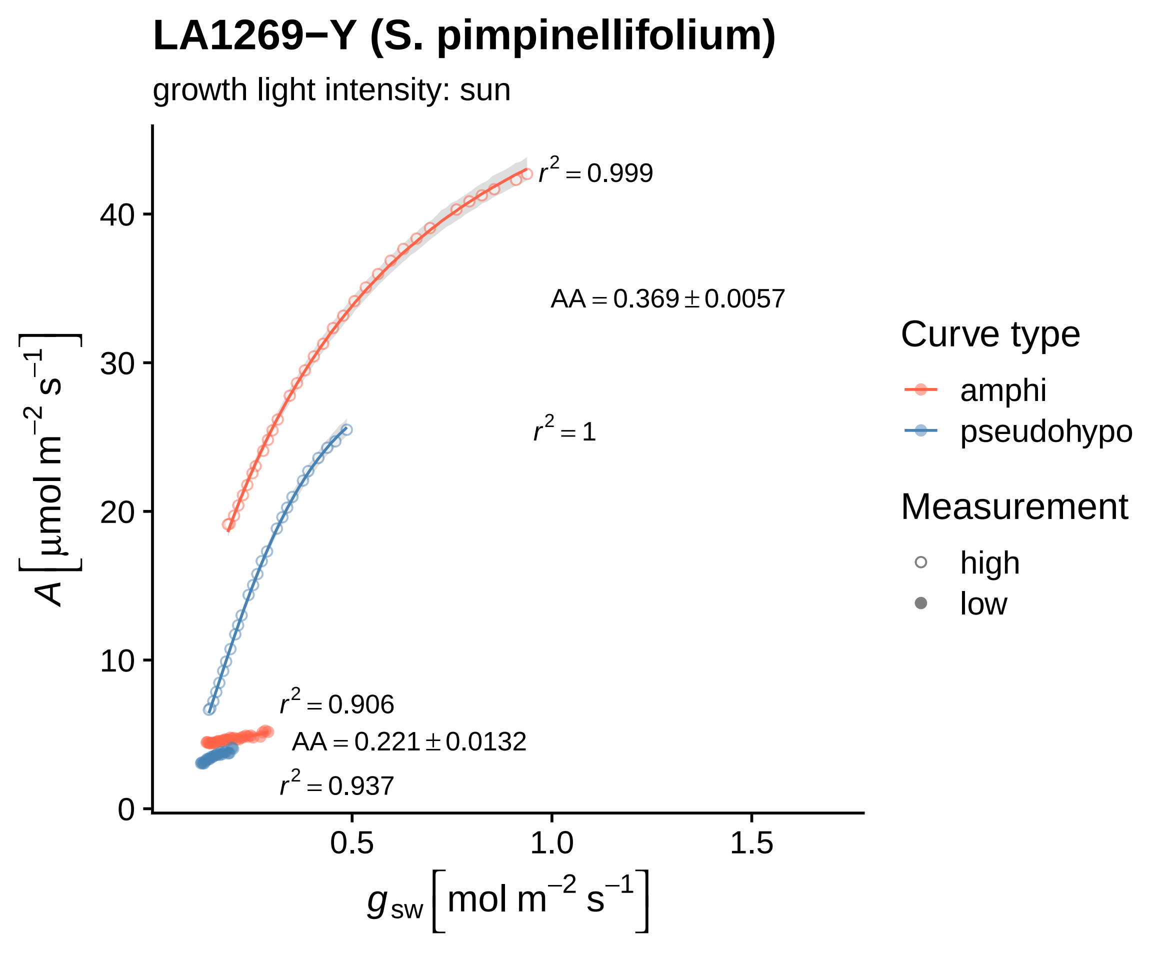
LA1269−Y (S. pimpinellifolium)
growth light intensity: sun
$r^2 = 0.999$
$AA = 0.369 \pm 0.0057$
$r^2 = 1$
$r^2 = 0.906$
$AA = 0.221 \pm 0.0132$
$r^2 = 0.937$
$A\left[\mu mol\ m^{-2}\ s^{-1}\right]$
0
10
20
30
40
0.5
1.0
1.5
$g_{sw}\left[mol\ m^{-2}\ s^{-1}\right]$
Curve type
amphi
pseudohypo
Measurement
high
low

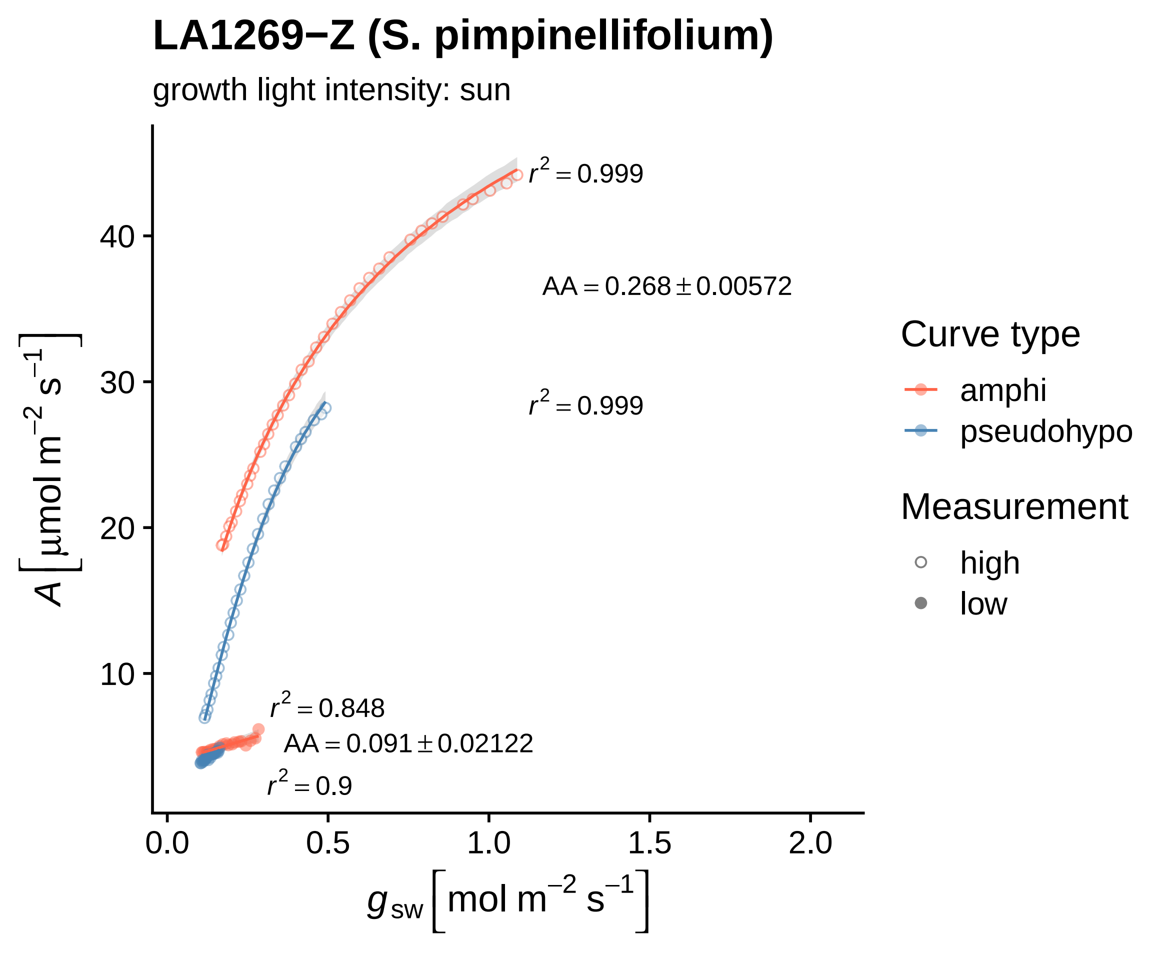
LA1269−Z (S. pimpinellifolium)
growth light intensity: sun
r² = 0.999
AA = 0.268 ± 0.00572
r² = 0.999
r² = 0.848
AA = 0.091 ± 0.02122
r² = 0.9
40
30
20
10
0.0
0.5
1.0
1.5
2.0
A [μmol m⁻² s⁻¹]
g sw [mol m⁻² s⁻¹]
Curve type
amphi
pseudohypo
Measurement
high
low

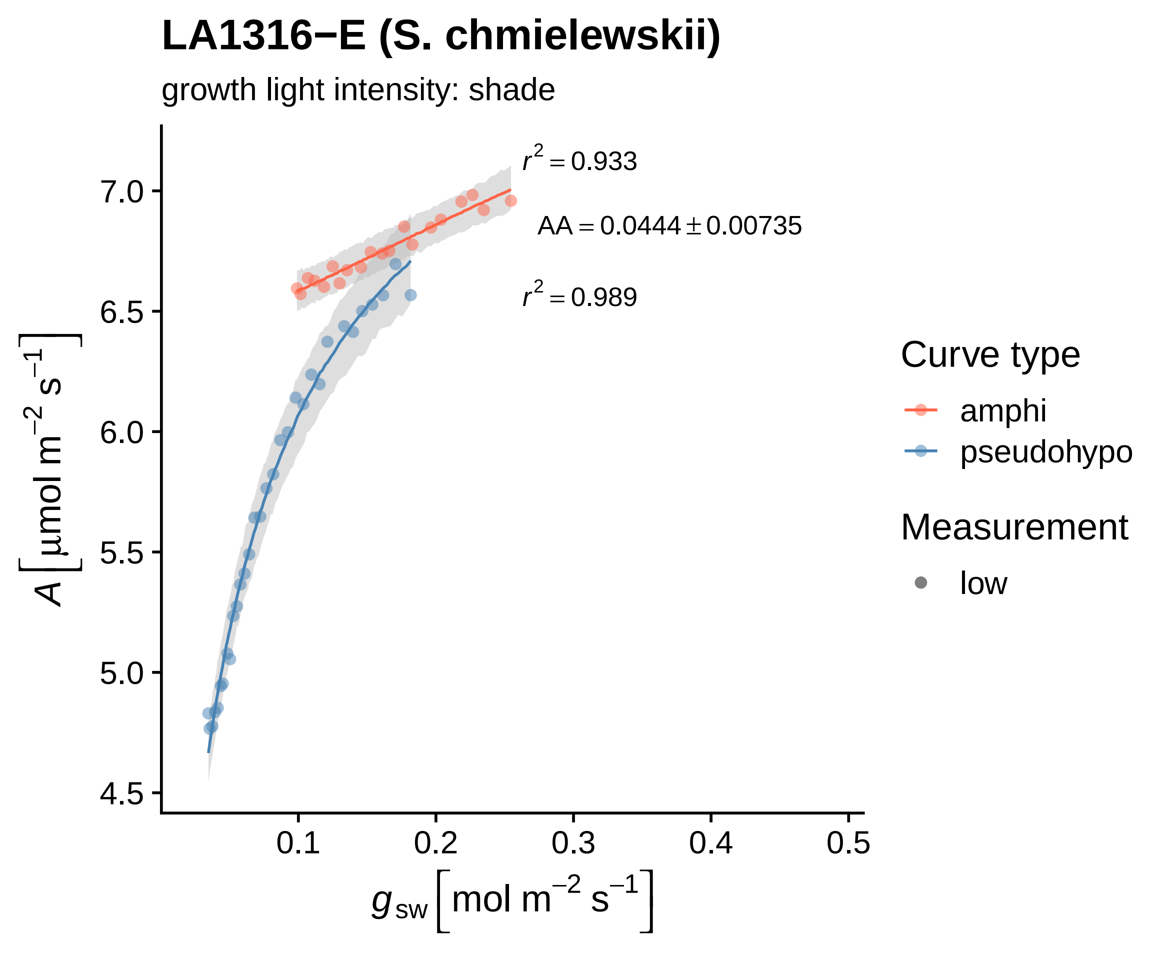

LA1316−E (S. chmielewskii)
growth light intensity: shade
$r^2 = 0.933$
$AA = 0.0444 \pm 0.00735$
$r^2 = 0.989$
7.0
6.5
6.0
5.5
5.0
4.5
$A\left[\mu mol\ m^{-2}\ s^{-1}\right]$
0.1
0.2
0.3
0.4
0.5
$g_{sw}\left[mol\ m^{-2}\ s^{-1}\right]$
Curve type
amphi
pseudohypo
Measurement
low

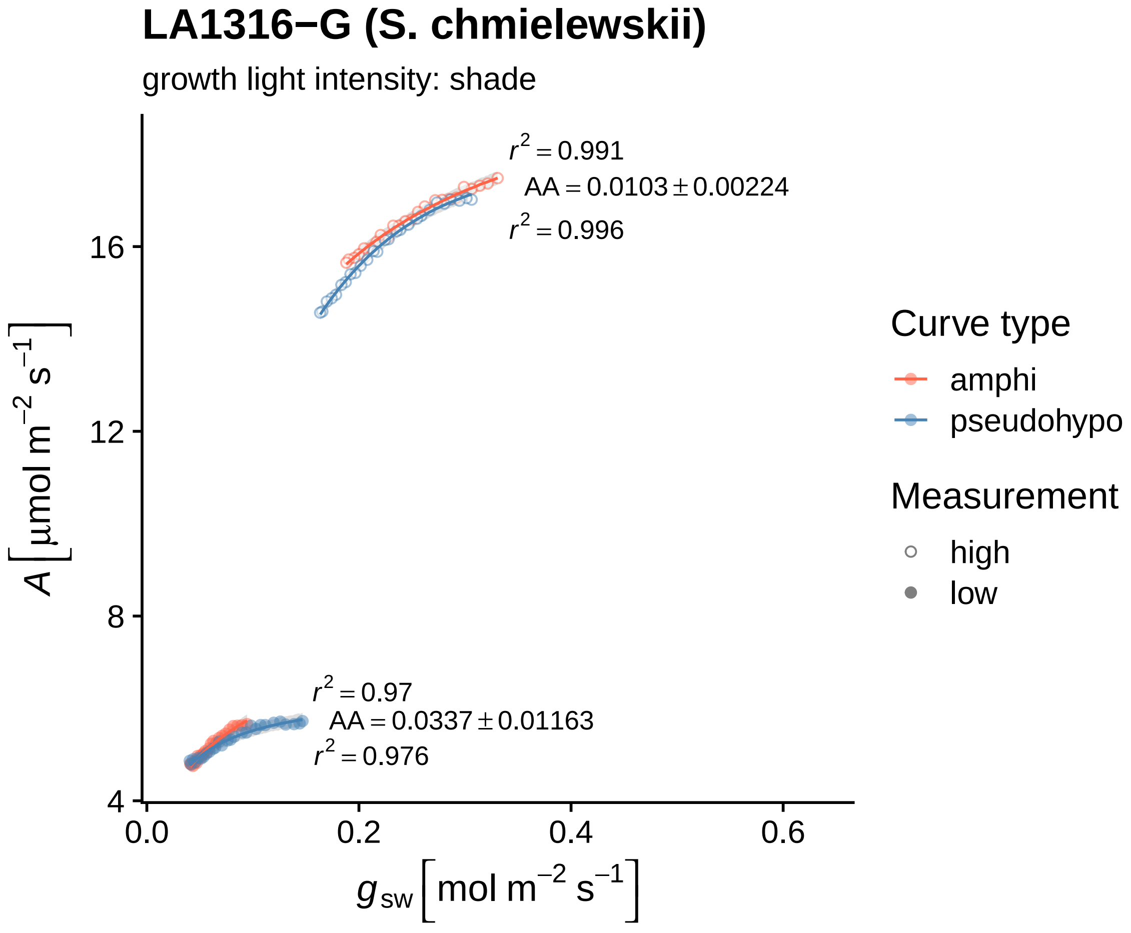
LA1316−G (S. chmielewskii)
growth light intensity: shade
r² = 0.991
AA = 0.0103 ± 0.00224
r² = 0.996
r² = 0.97
AA = 0.0337 ± 0.01163
r² = 0.976
A [μmol m⁻² s⁻¹]
16
12
8
4
0.0
0.2
0.4
0.6
g_sw [mol m⁻² s⁻¹]
Curve type
amphi
pseudohypo
Measurement
high
low

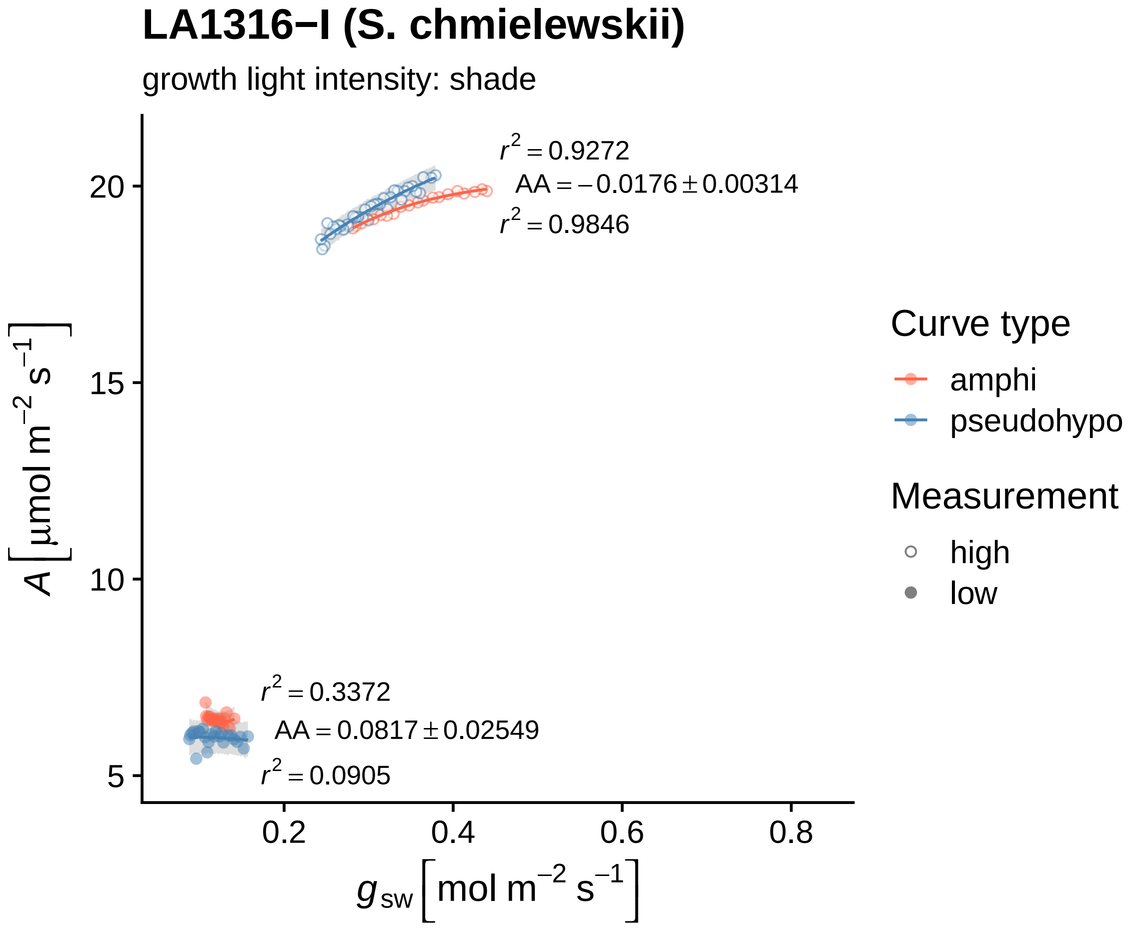
LA1316−I (S. chmielewskii)
growth light intensity: shade
$r^2 = 0.9272$
$AA = -0.0176 \pm 0.00314$
$r^2 = 0.9846$
$r^2 = 0.3372$
$AA = 0.0817 \pm 0.02549$
$r^2 = 0.0905$
20
15
10
5
0.2
0.4
0.6
0.8
$A\left[\mu mol\ m^{-2}\ s^{-1}\right]$
$g_{sw}\left[mol\ m^{-2}\ s^{-1}\right]$
Curve type
amphi
pseudohypo
Measurement
high
low

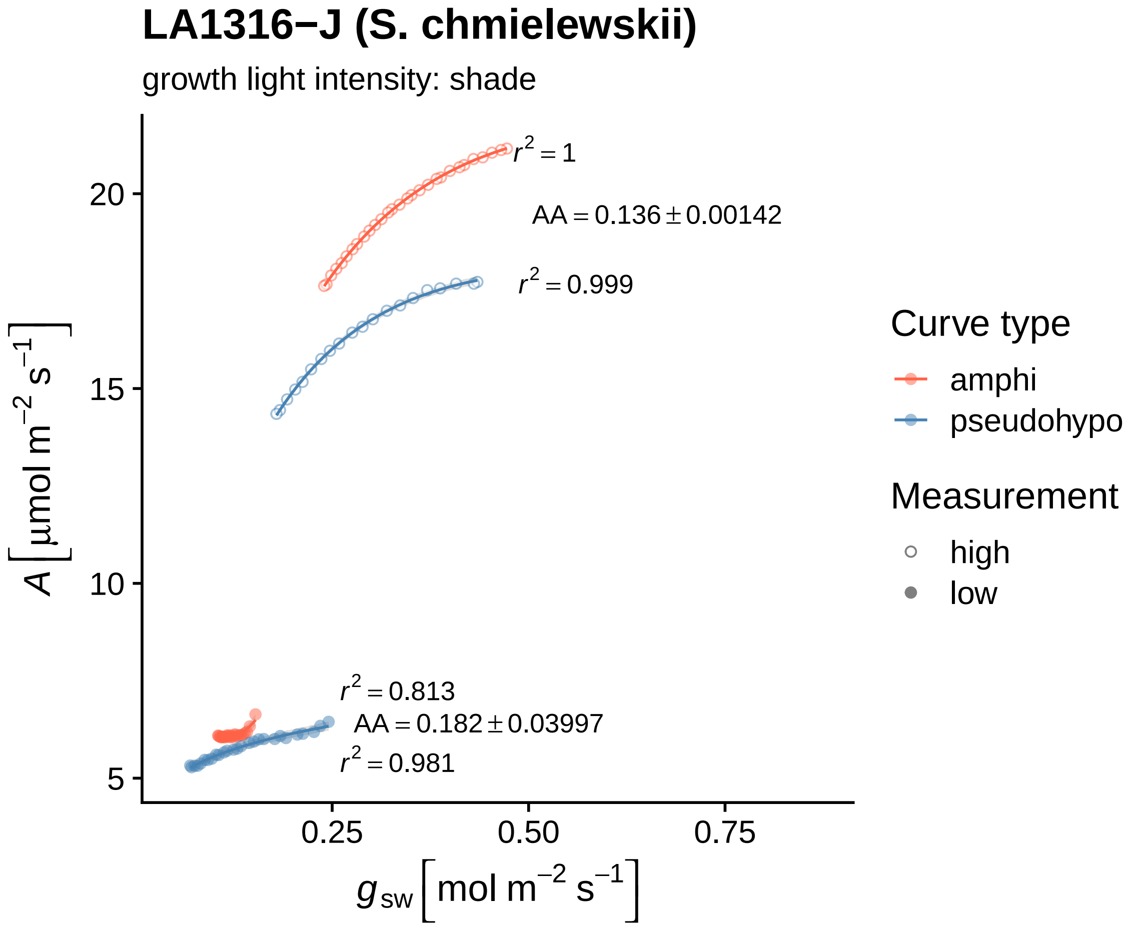

LA1316−J (S. chmielewskii)
growth light intensity: shade
$r^2 = 1$
$AA = 0.136 \pm 0.00142$
$r^2 = 0.999$
$r^2 = 0.813$
$AA = 0.182 \pm 0.03997$
$r^2 = 0.981$
20
15
10
5
0.25
0.50
0.75
$A\left[\mu mol\ m^{-2}\ s^{-1}\right]$
$g_{sw}\left[mol\ m^{-2}\ s^{-1}\right]$
Curve type
amphi
pseudohypo
Measurement
high
low

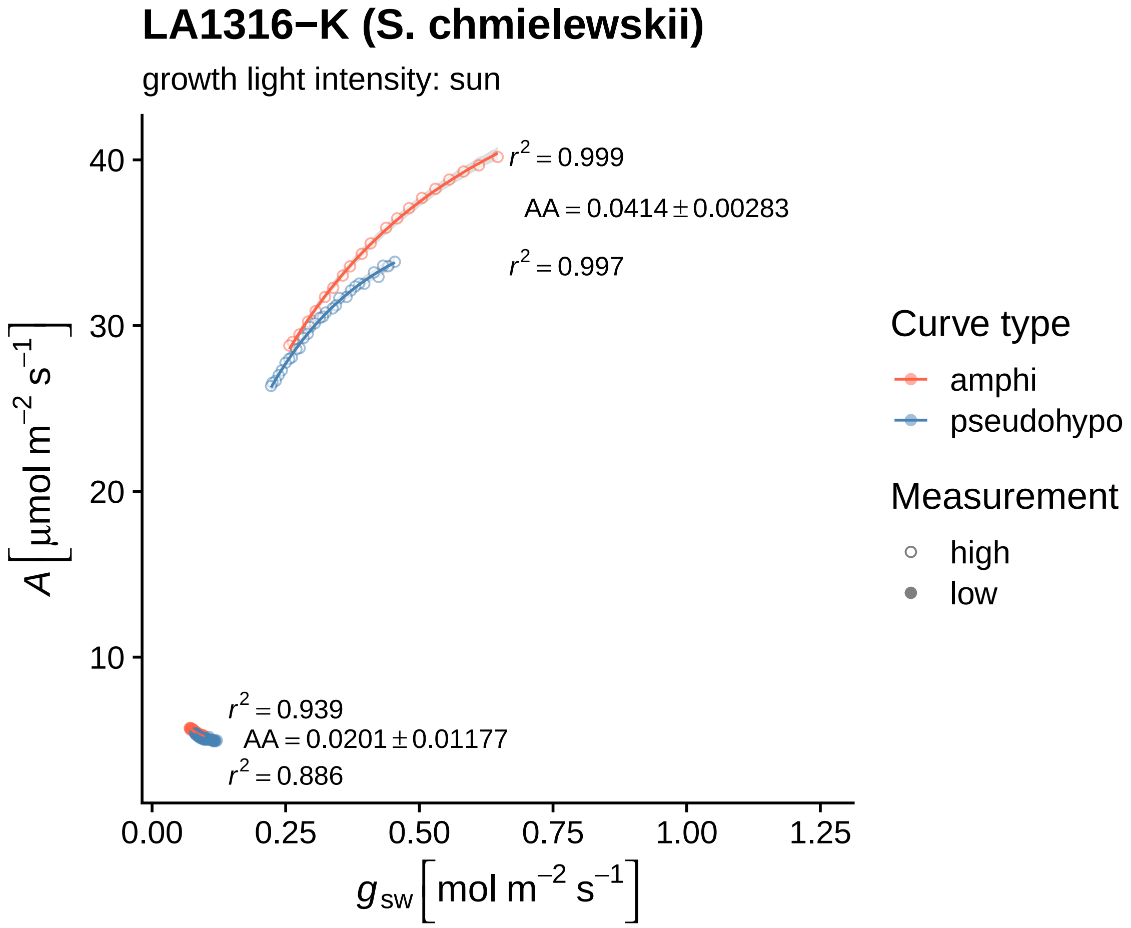
LA1316−K (S. chmielewskii)
growth light intensity: sun
r² = 0.999
AA = 0.0414 ± 0.00283
r² = 0.997
r² = 0.939
AA = 0.0201 ± 0.01177
r² = 0.886
A [µmol m⁻² s⁻¹]
40
30
20
10
0.00
0.25
0.50
0.75
1.00
1.25
g_sw [mol m⁻² s⁻¹]
Curve type
amphi
pseudohypo
Measurement
high
low

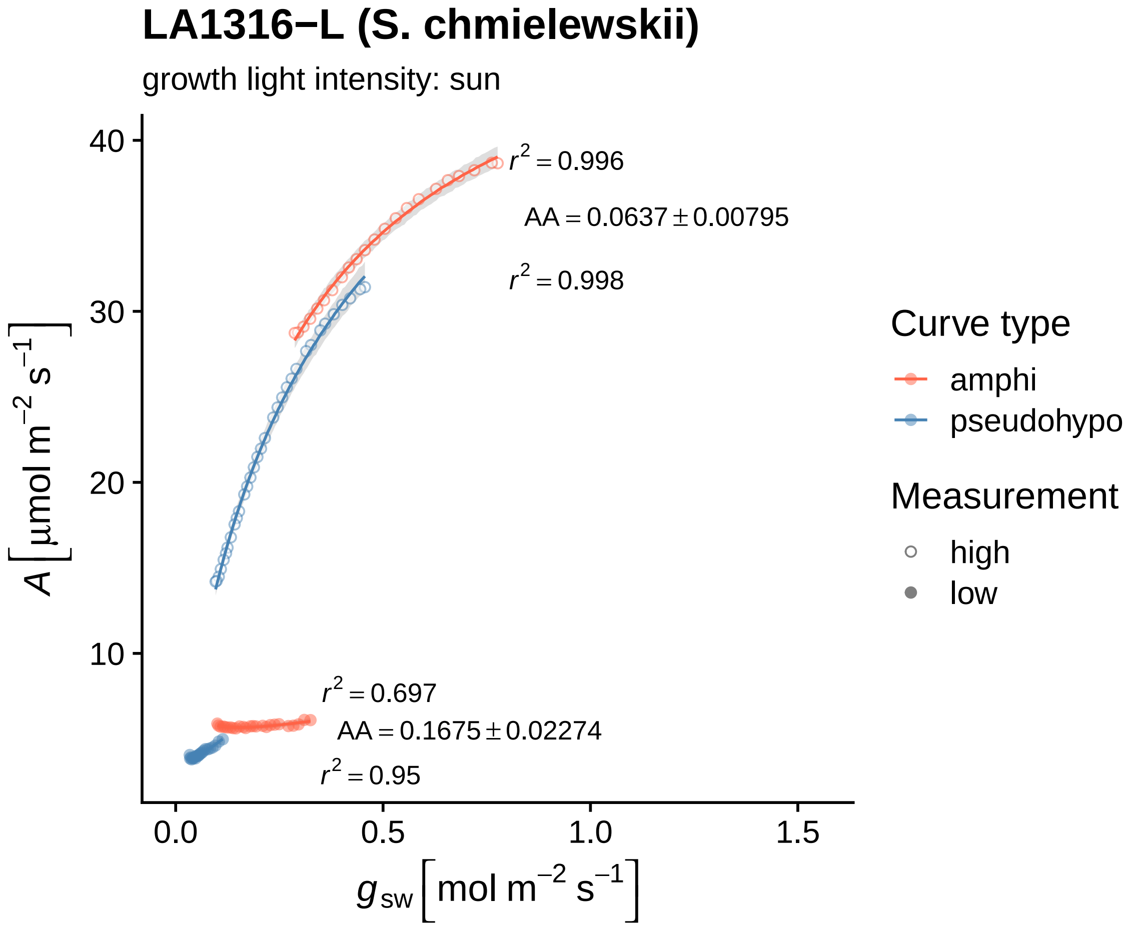
LA1316−L (S. chmielewskii)
growth light intensity: sun
$r^2 = 0.996$
$AA = 0.0637 \pm 0.00795$
$r^2 = 0.998$
$r^2 = 0.697$
$AA = 0.1675 \pm 0.02274$
$r^2 = 0.95$
40
30
20
10
0.0
0.5
1.0
1.5
$A\ [\mu mol\ m^{-2}\ s^{-1}]$
$g_{sw}\ [mol\ m^{-2}\ s^{-1}]$
Curve type
amphi
pseudohypo
Measurement
high
low

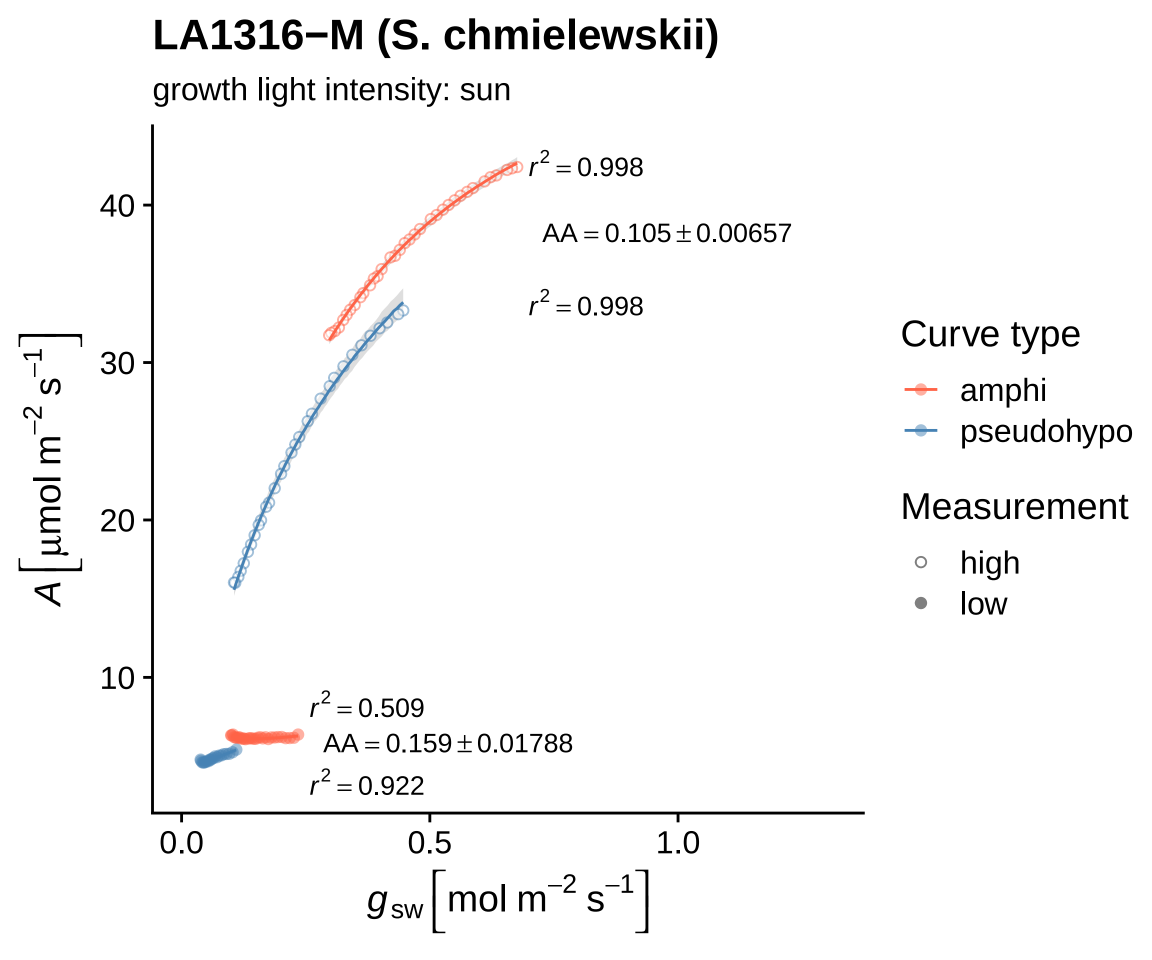

LA1316−M (S. chmielewskii)
growth light intensity: sun
$r^2 = 0.998$
AA = 0.105 ± 0.00657
$r^2 = 0.998$
$r^2 = 0.509$
AA = 0.159 ± 0.01788
$r^2 = 0.922$
40
30
20
10
0.0
0.5
1.0
A [μmol m⁻² s⁻¹]
$g_{sw}$ [mol m⁻² s⁻¹]
Curve type
amphi
pseudohypo
Measurement
high
low

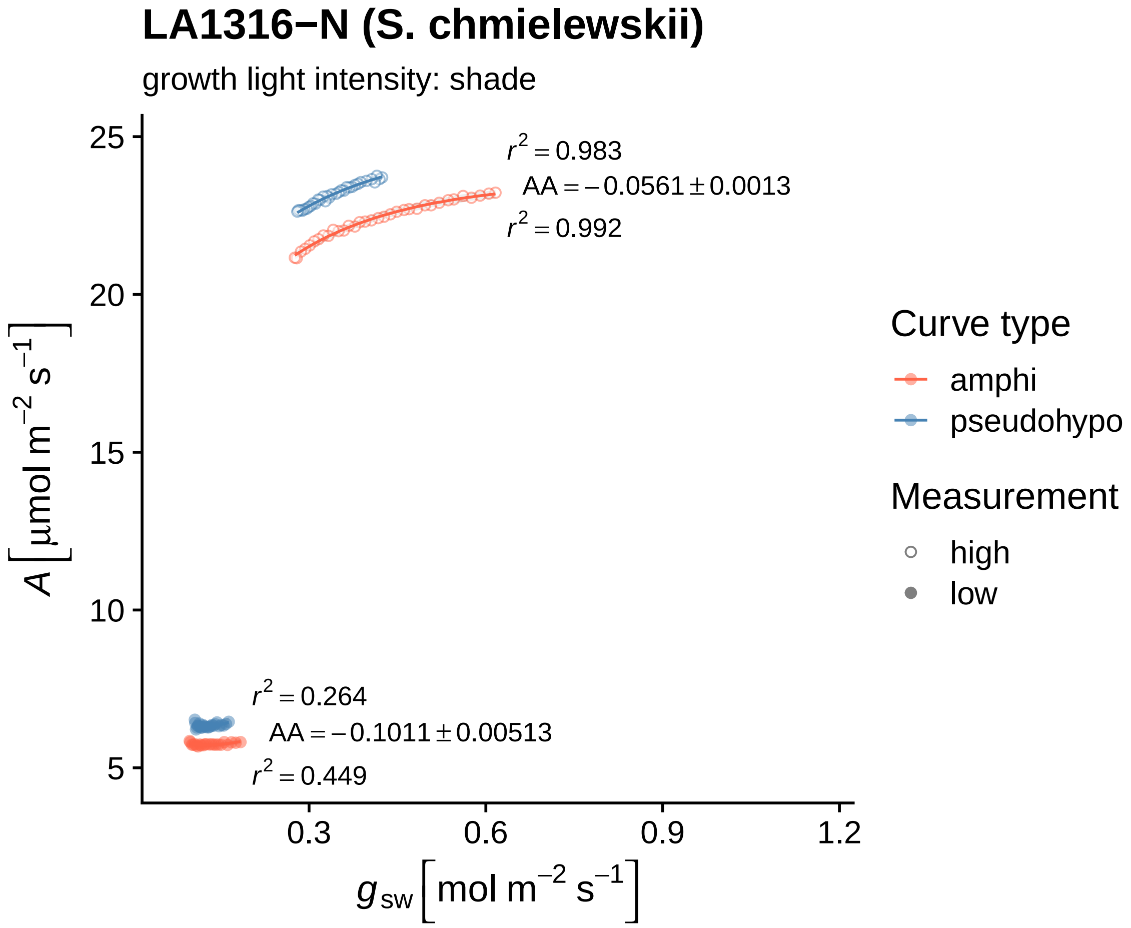

LA1316−N (S. chmielewskii)
growth light intensity: shade
$r^2 = 0.983$
$AA = -0.0561 \pm 0.0013$
$r^2 = 0.992$
$r^2 = 0.264$
$AA = -0.1011 \pm 0.00513$
$r^2 = 0.449$
25
20
15
10
5
0.3
0.6
0.9
1.2
$A\ [\mu mol\ m^{-2}\ s^{-1}]$
$g_{sw}\ [mol\ m^{-2}\ s^{-1}]$
Curve type
amphi
pseudohypo
Measurement
high
low

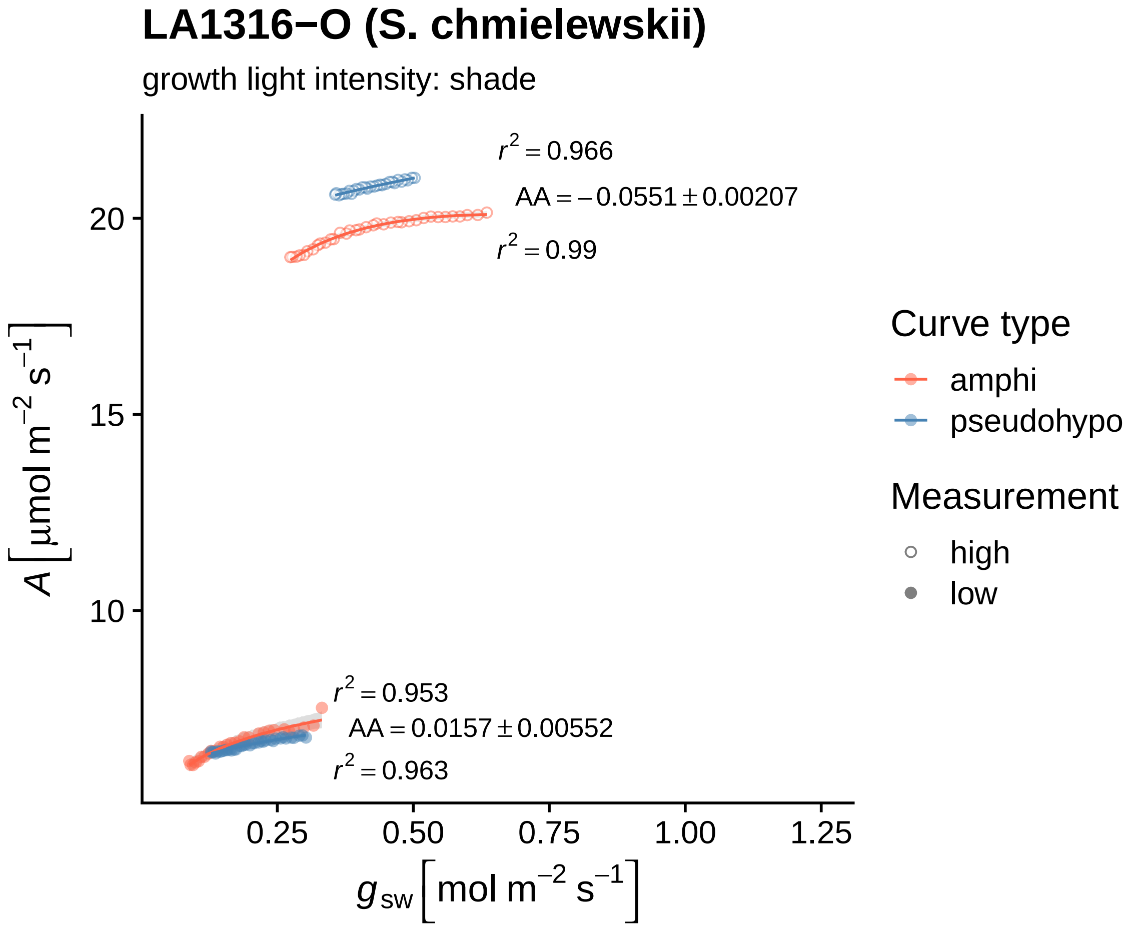

LA1316−O (S. chmielewskii)
growth light intensity: shade
$r^2 = 0.966$
$AA = -0.0551 \pm 0.00207$
$r^2 = 0.99$
$r^2 = 0.953$
$AA = 0.0157 \pm 0.00552$
$r^2 = 0.963$
20
15
10
0.25
0.50
0.75
1.00
1.25
$A\,[\mu mol\ m^{-2}\ s^{-1}]$
$g_{sw}\,[mol\ m^{-2}\ s^{-1}]$
Curve type
amphi
pseudohypo
Measurement
high
low

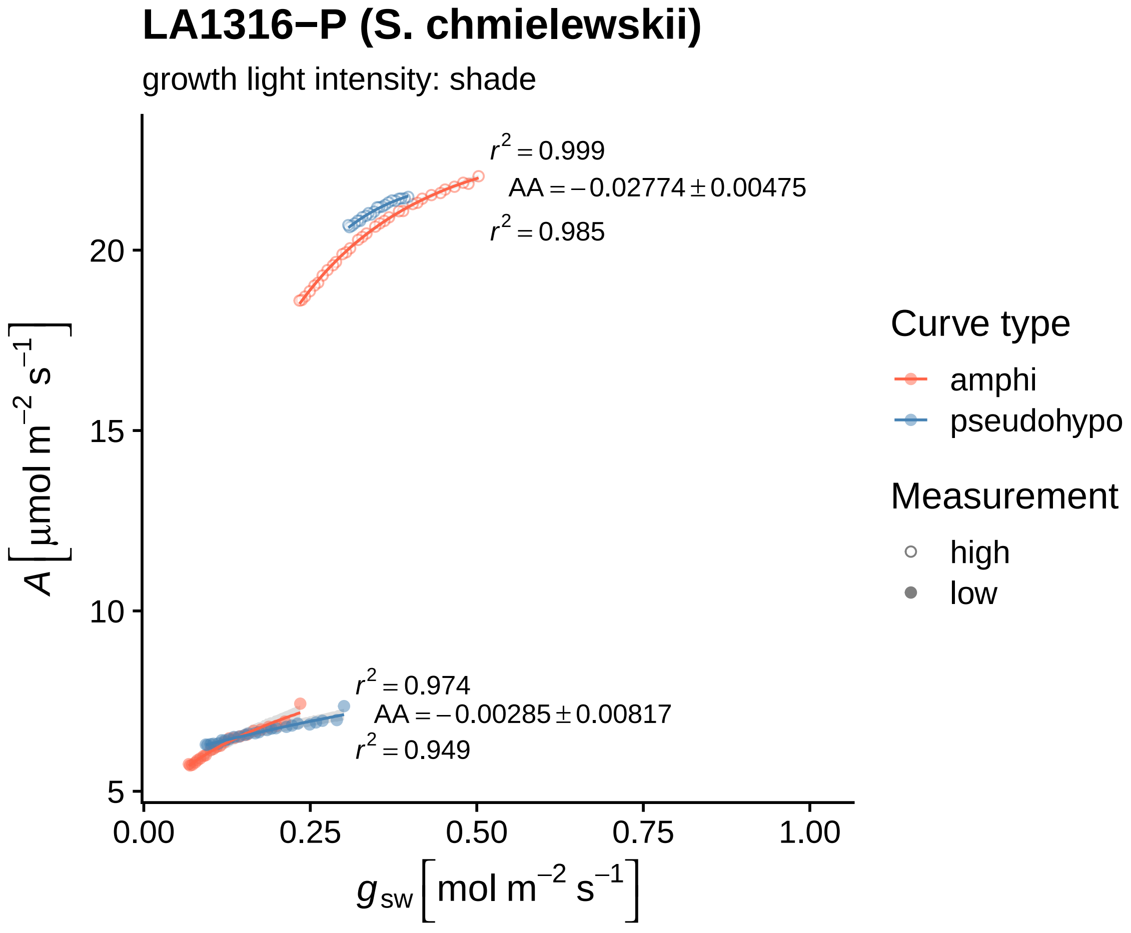

LA1316−P (S. chmielewskii)
growth light intensity: shade
$r^2 = 0.999$
$AA = -0.02774 \pm 0.00475$
$r^2 = 0.985$
$r^2 = 0.974$
$AA = -0.00285 \pm 0.00817$
$r^2 = 0.949$
20
15
10
5
0.00
0.25
0.50
0.75
1.00
$A\ [\mu mol\ m^{-2}\ s^{-1}]$
$g_{sw}\ [mol\ m^{-2}\ s^{-1}]$
Curve type
amphi
pseudohypo
Measurement
high
low

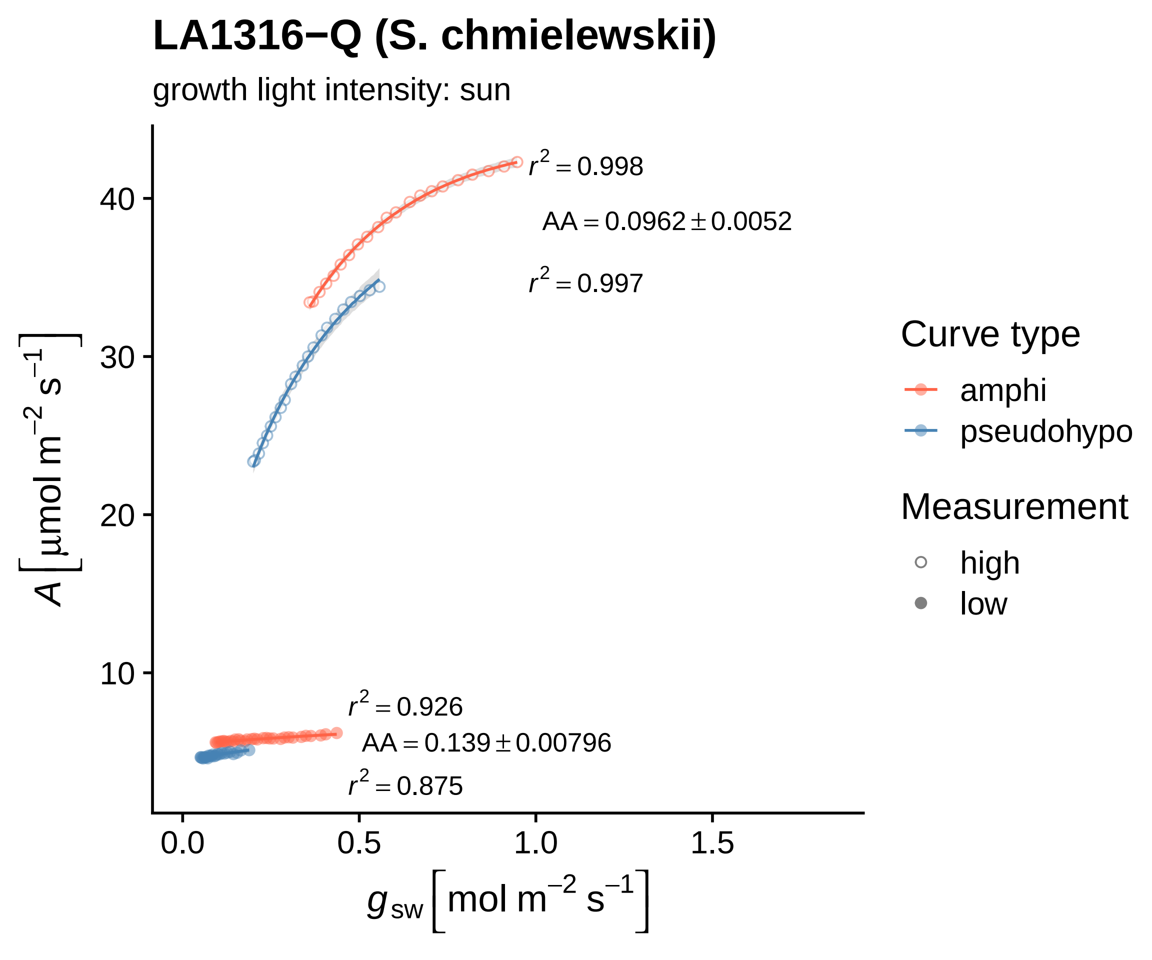

LA1316−Q (S. chmielewskii)
growth light intensity: sun
$r^2 = 0.998$
$AA = 0.0962 \pm 0.0052$
$r^2 = 0.997$
$r^2 = 0.926$
$AA = 0.139 \pm 0.00796$
$r^2 = 0.875$
40
30
20
10
0.0
0.5
1.0
1.5
$A\left[\mu mol\ m^{-2}\ s^{-1}\right]$
$g_{sw}\left[mol\ m^{-2}\ s^{-1}\right]$
Curve type
amphi
pseudohypo
Measurement
high
low

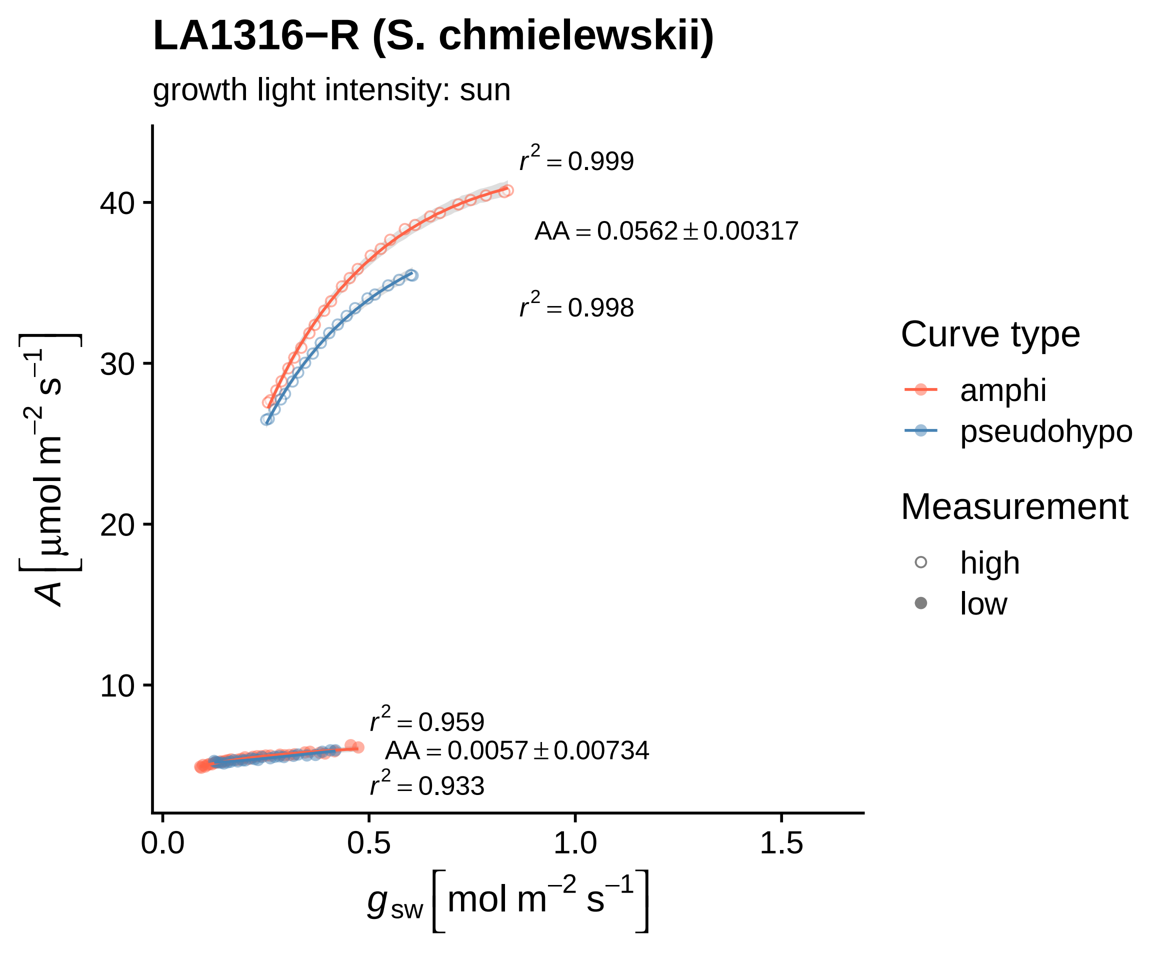
LA1316−R (S. chmielewskii)
growth light intensity: sun
$r^2 = 0.999$
$AA = 0.0562 \pm 0.00317$
$r^2 = 0.998$
$r^2 = 0.959$
$AA = 0.0057 \pm 0.00734$
$r^2 = 0.933$
40
30
20
10
0.0
0.5
1.0
1.5
$A\left[\mu mol\ m^{-2}\ s^{-1}\right]$
$g_{sw}\left[mol\ m^{-2}\ s^{-1}\right]$
Curve type
amphi
pseudohypo
Measurement
high
low

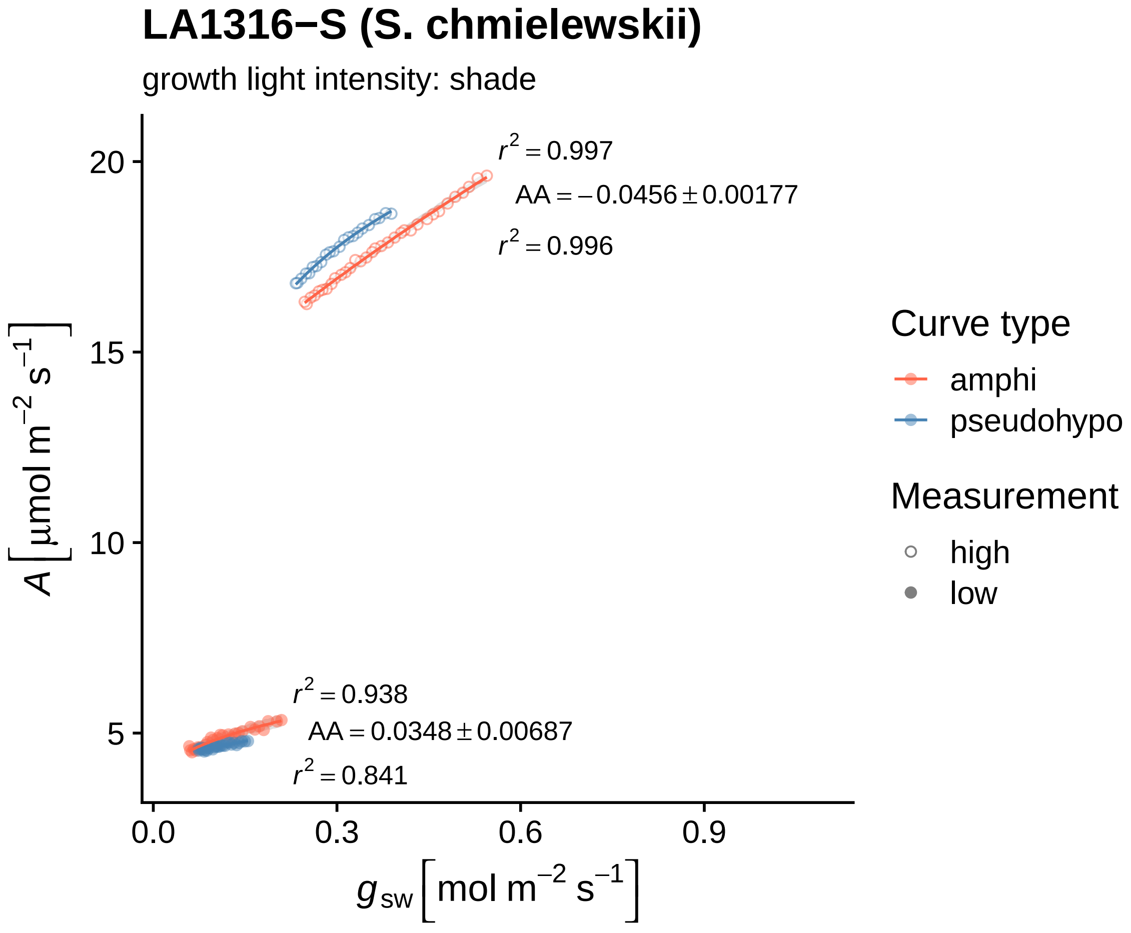
LA1316−S (S. chmielewskii)
growth light intensity: shade
$r^2 = 0.997$
$AA = -0.0456 \pm 0.00177$
$r^2 = 0.996$
$r^2 = 0.938$
$AA = 0.0348 \pm 0.00687$
$r^2 = 0.841$
20
15
10
5
0.0
0.3
0.6
0.9
$A\,[\mu mol\ m^{-2}\ s^{-1}]$
$g_{sw}\,[mol\ m^{-2}\ s^{-1}]$
Curve type
amphi
pseudohypo
Measurement
high
low

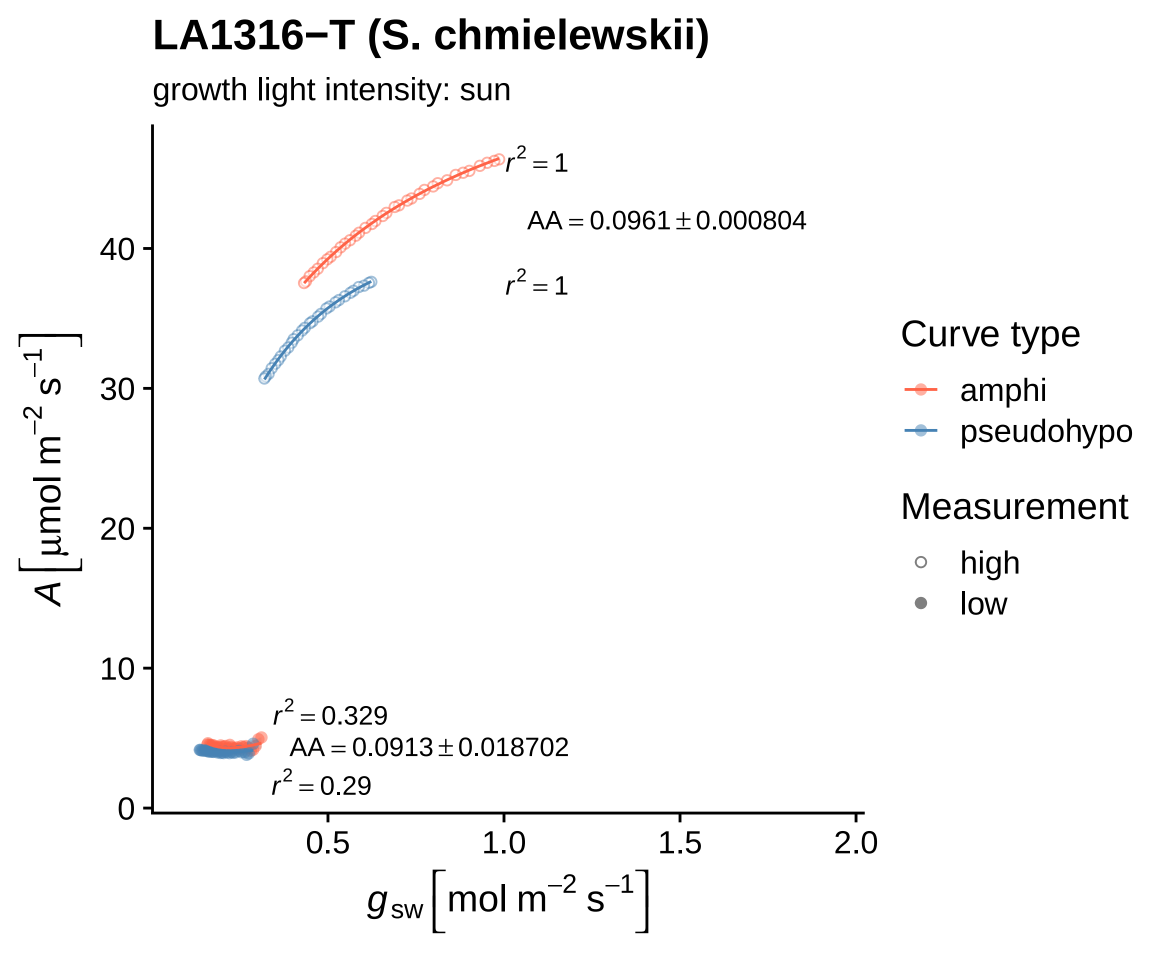

LA1316−T (S. chmielewskii)
growth light intensity: sun
r² = 1
AA = 0.0961 ± 0.000804
r² = 1
r² = 0.329
AA = 0.0913 ± 0.018702
r² = 0.29
A [μmol m⁻² s⁻¹]
g_sw [mol m⁻² s⁻¹]
0
10
20
30
40
0.5
1.0
1.5
2.0
Curve type
amphi
pseudohypo
Measurement
high
low

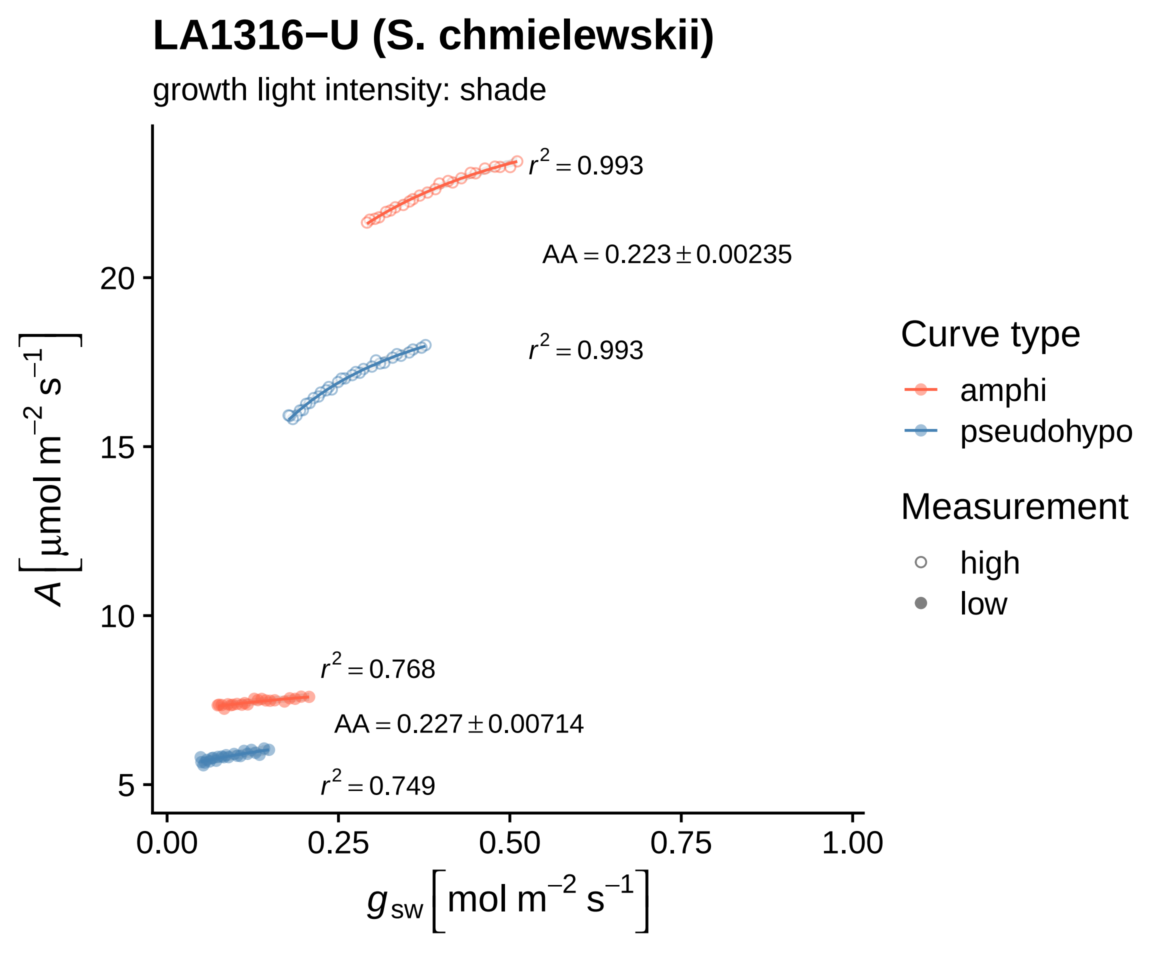

LA1316−U (S. chmielewskii)
growth light intensity: shade
$r^2 = 0.993$
$AA = 0.223 \pm 0.00235$
$r^2 = 0.993$
$r^2 = 0.768$
$AA = 0.227 \pm 0.00714$
$r^2 = 0.749$
20
15
10
5
0.00
0.25
0.50
0.75
1.00
$A\ [\mu mol\ m^{-2}\ s^{-1}]$
$g_{sw}\ [mol\ m^{-2}\ s^{-1}]$
Curve type
amphi
pseudohypo
Measurement
high
low

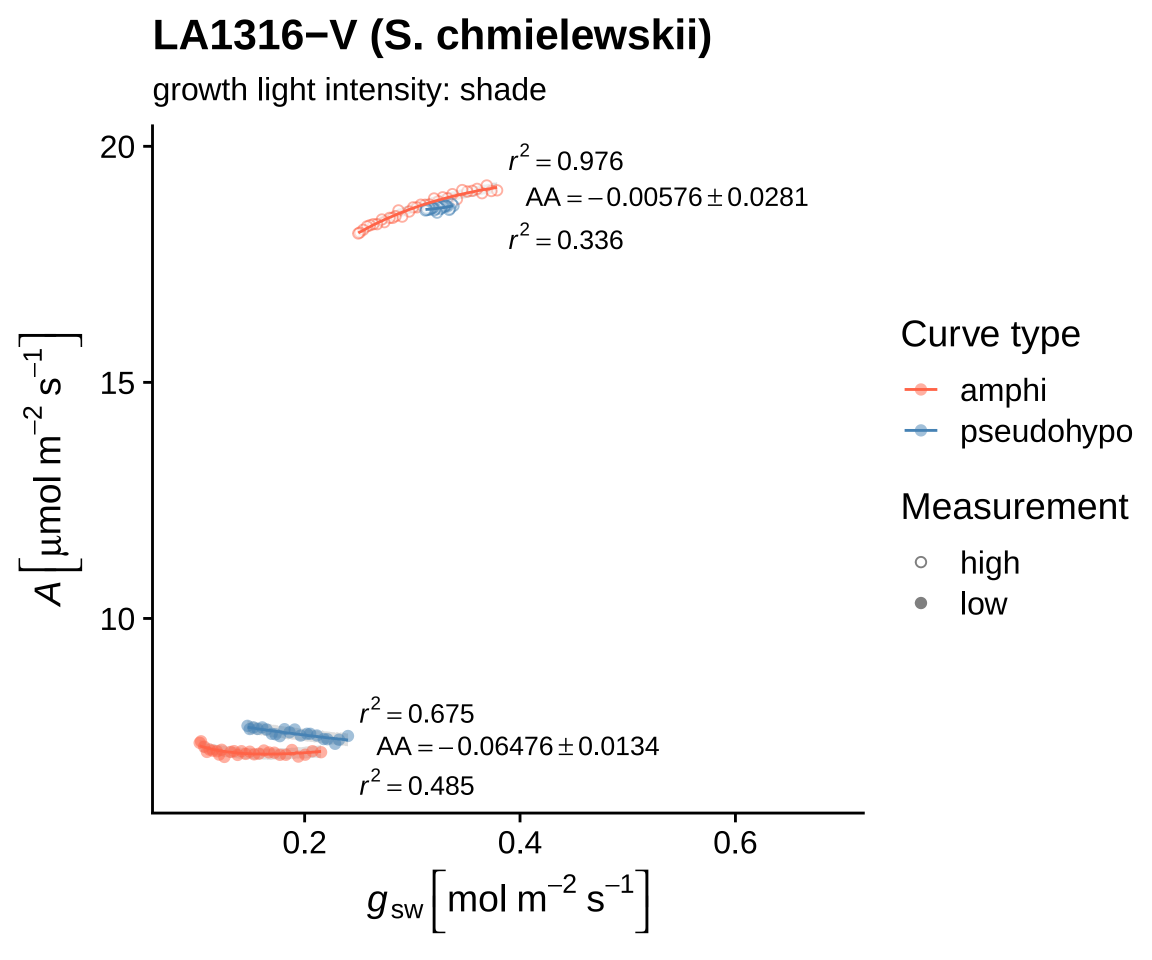

LA1316−V (S. chmielewskii)
growth light intensity: shade
$r^2 = 0.976$
$AA = -0.00576 \pm 0.0281$
$r^2 = 0.336$
$r^2 = 0.675$
$AA = -0.06476 \pm 0.0134$
$r^2 = 0.485$
20
15
10
0.2
0.4
0.6
$A\ [\mu mol\ m^{-2}\ s^{-1}]$
$g_{sw}\ [mol\ m^{-2}\ s^{-1}]$
Curve type
amphi
pseudohypo
Measurement
high
low

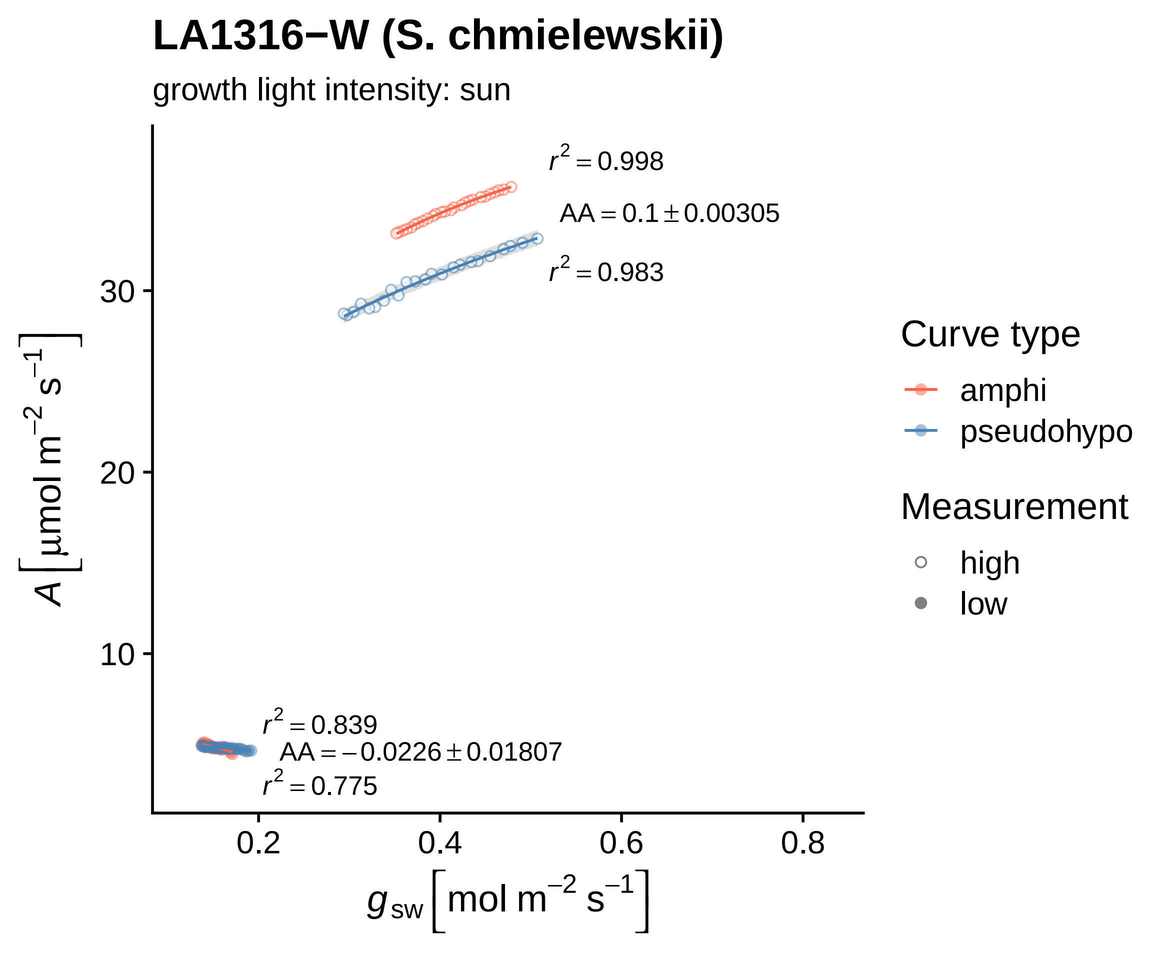
LA1316−W (S. chmielewskii)
growth light intensity: sun
$r^2 = 0.998$
$AA = 0.1 \pm 0.00305$
$r^2 = 0.983$
$r^2 = 0.839$
$AA = -0.0226 \pm 0.01807$
$r^2 = 0.775$
30
20
10
0.2
0.4
0.6
0.8
$A\,[\mu mol\ m^{-2}\ s^{-1}]$
$g_{sw}\,[mol\ m^{-2}\ s^{-1}]$
Curve type
amphi
pseudohypo
Measurement
high
low

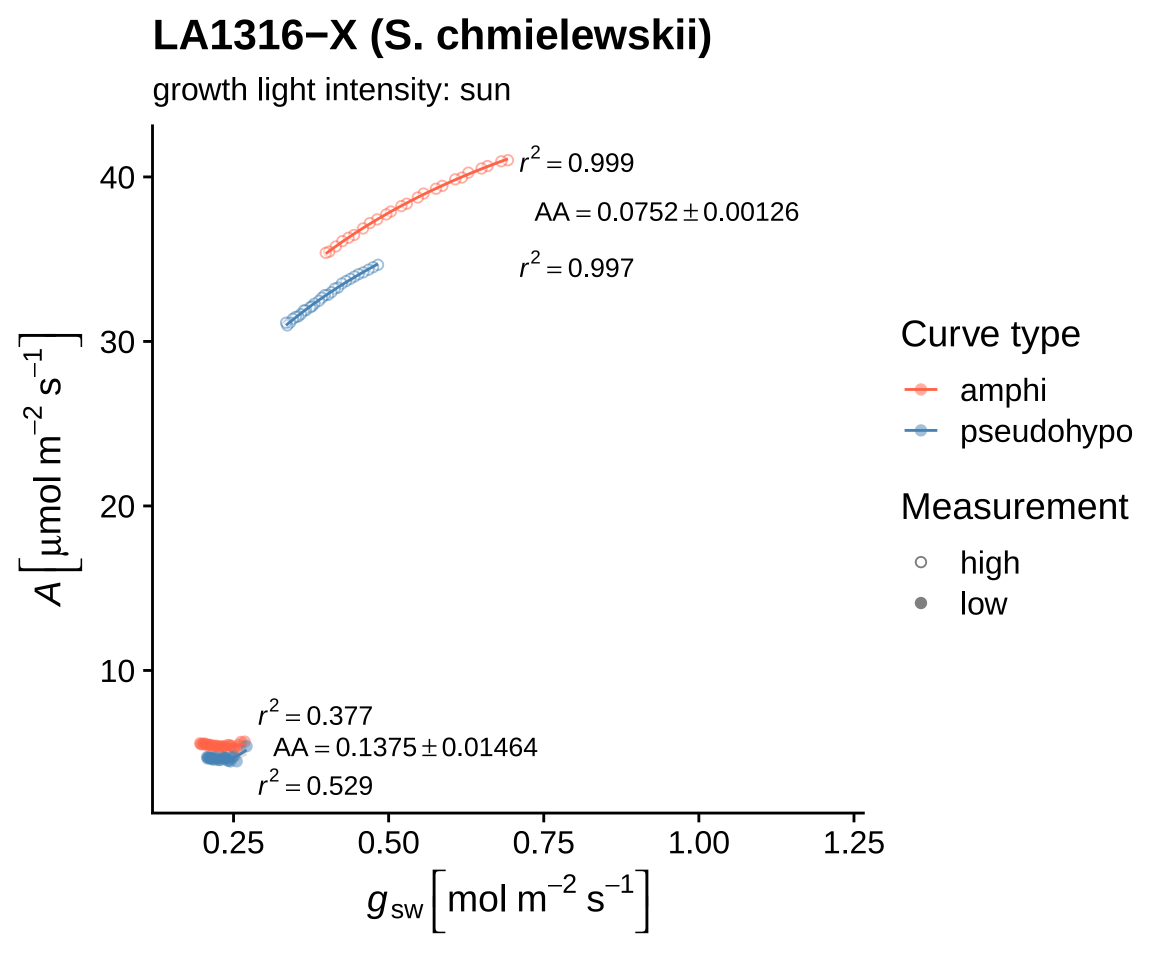

LA1316−X (S. chmielewskii)
growth light intensity: sun
$r^2 = 0.999$
$AA = 0.0752 \pm 0.00126$
$r^2 = 0.997$
$r^2 = 0.377$
$AA = 0.1375 \pm 0.01464$
$r^2 = 0.529$
40
30
20
10
0.25
0.50
0.75
1.00
1.25
$A\ [\mu mol\ m^{-2}\ s^{-1}]$
$g_{sw}\ [mol\ m^{-2}\ s^{-1}]$
Curve type
amphi
pseudohypo
Measurement
high
low

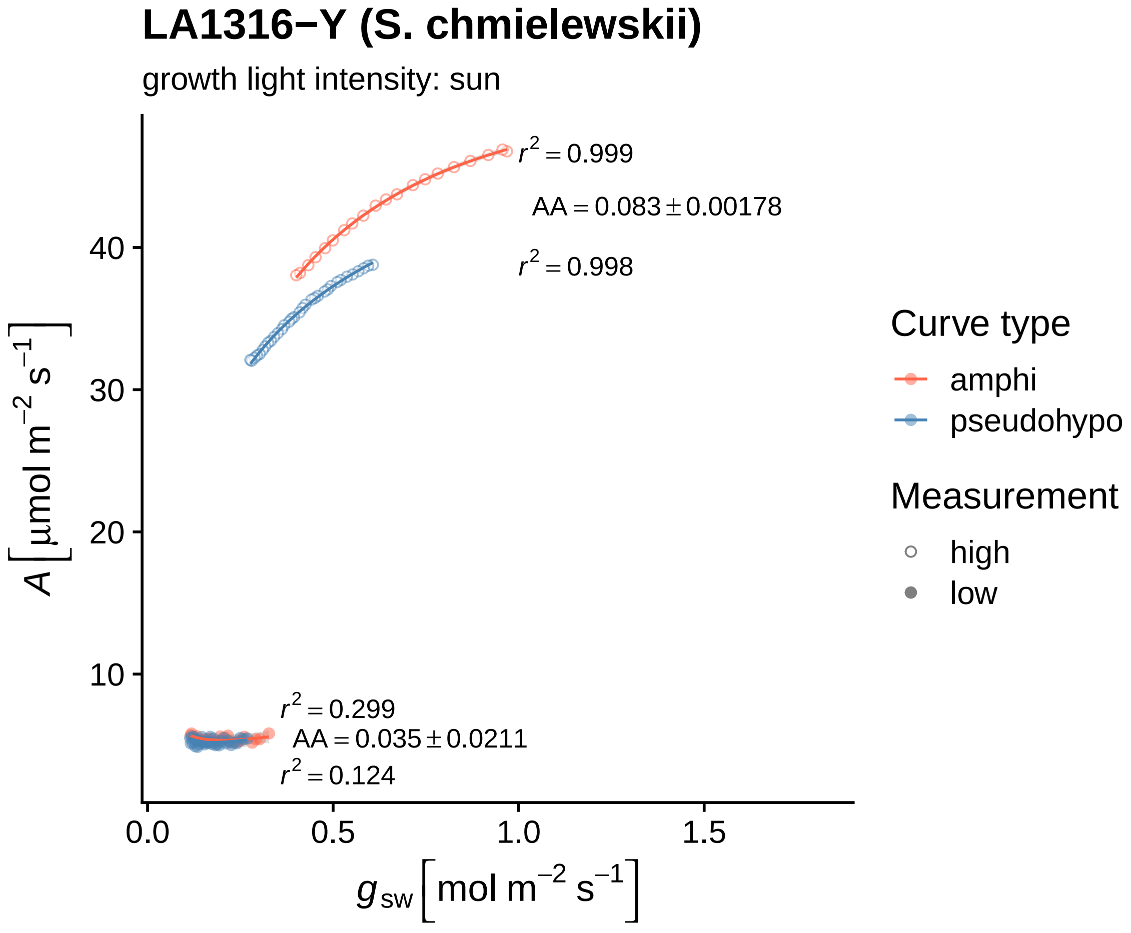
LA1316−Y (S. chmielewskii)
growth light intensity: sun
$r^2 = 0.999$
$AA = 0.083 \pm 0.00178$
$r^2 = 0.998$
$r^2 = 0.299$
$AA = 0.035 \pm 0.0211$
$r^2 = 0.124$
40
30
20
10
0.0
0.5
1.0
1.5
$A\ [\mu mol\ m^{-2}\ s^{-1}]$
$g_{sw}\ [mol\ m^{-2}\ s^{-1}]$
Curve type
amphi
pseudohypo
Measurement
high
low

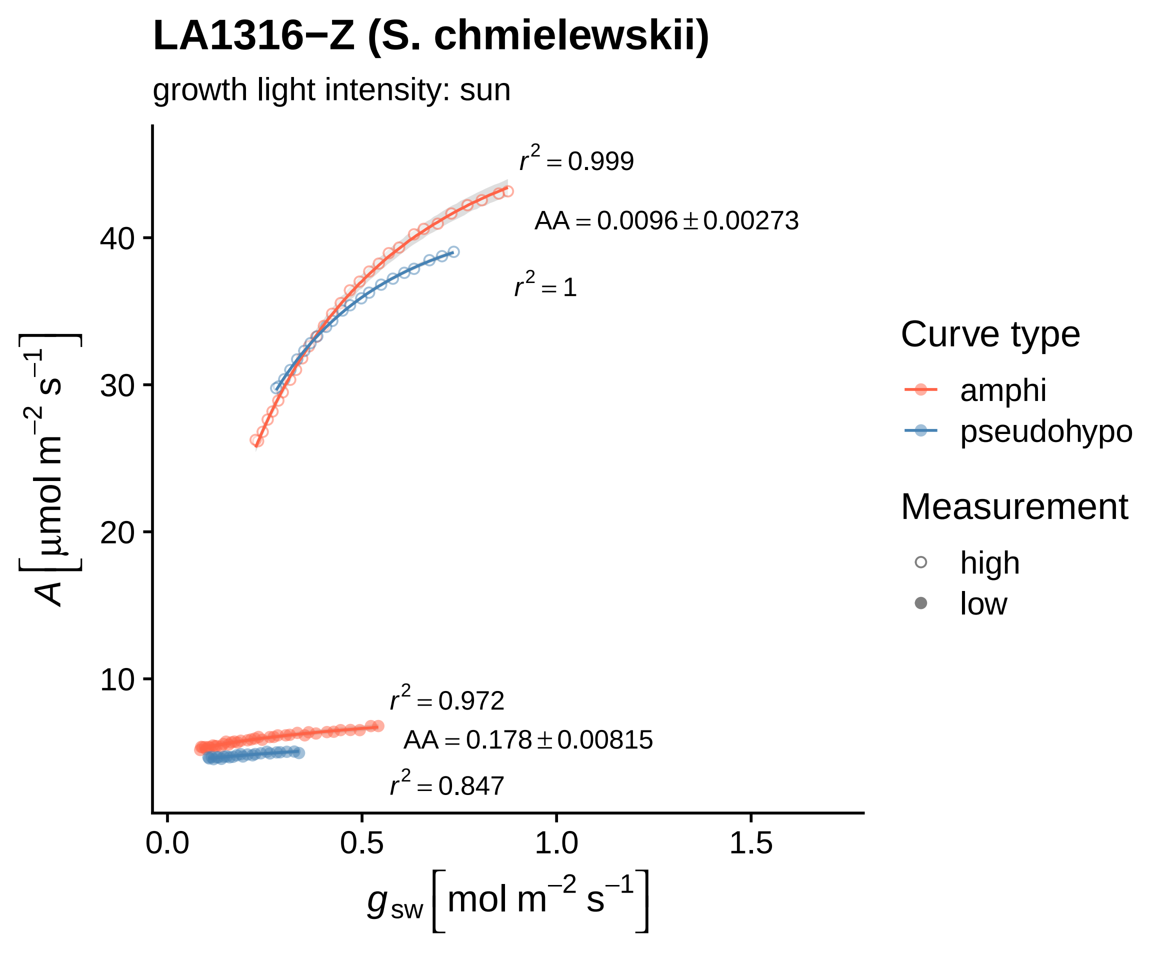

LA1316−Z (S. chmielewskii)
growth light intensity: sun
$r^2 = 0.999$
$AA = 0.0096 \pm 0.00273$
$r^2 = 1$
$r^2 = 0.972$
$AA = 0.178 \pm 0.00815$
$r^2 = 0.847$
40
30
20
10
0.0
0.5
1.0
1.5
$A\ [\mu mol\ m^{-2}\ s^{-1}]$
$g_{sw}\ [mol\ m^{-2}\ s^{-1}]$
Curve type
amphi
pseudohypo
Measurement
high
low

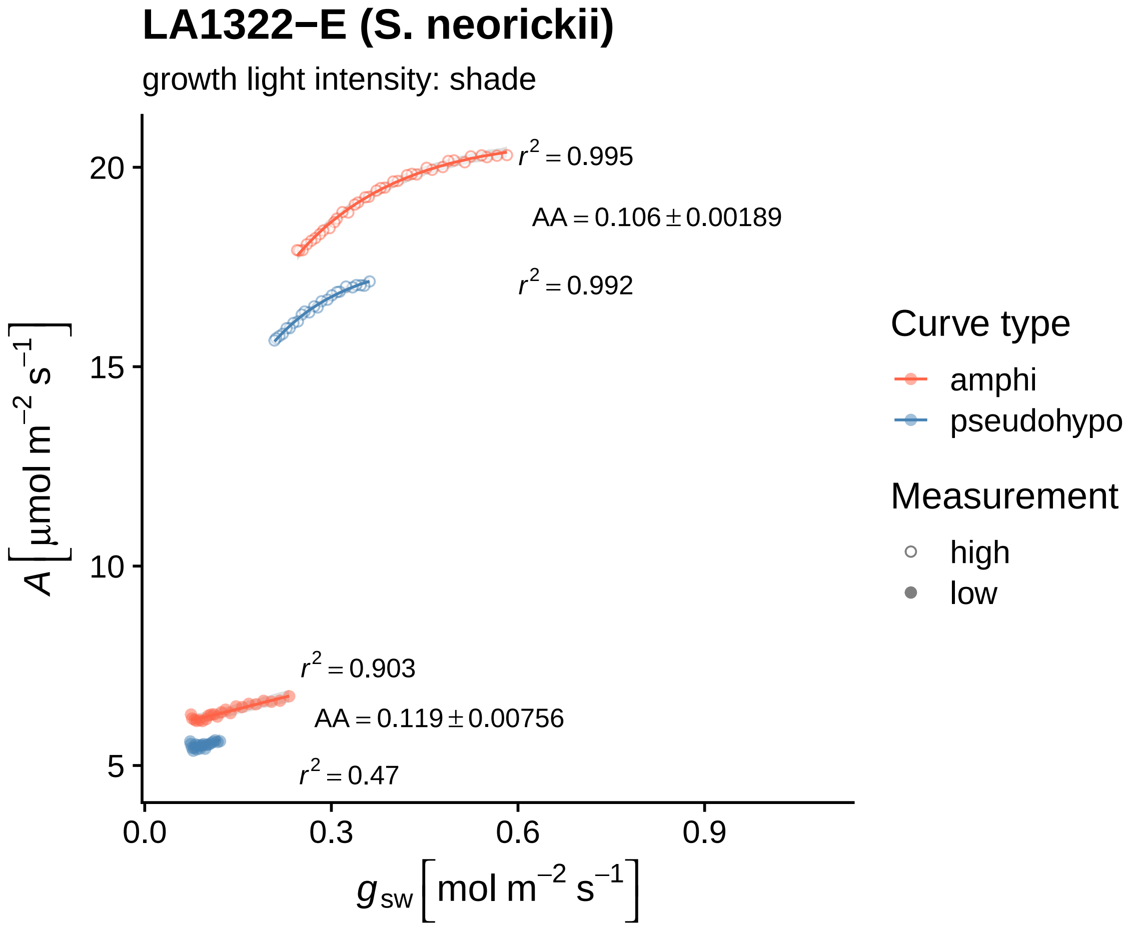

LA1322−E (S. neorickii)
growth light intensity: shade
$r^2 = 0.995$
$AA = 0.106 \pm 0.00189$
$r^2 = 0.992$
$r^2 = 0.903$
$AA = 0.119 \pm 0.00756$
$r^2 = 0.47$
20
15
10
5
0.0
0.3
0.6
0.9
$A\ [\mu mol\ m^{-2}\ s^{-1}]$
$g_{sw}\ [mol\ m^{-2}\ s^{-1}]$
Curve type
amphi
pseudohypo
Measurement
high
low

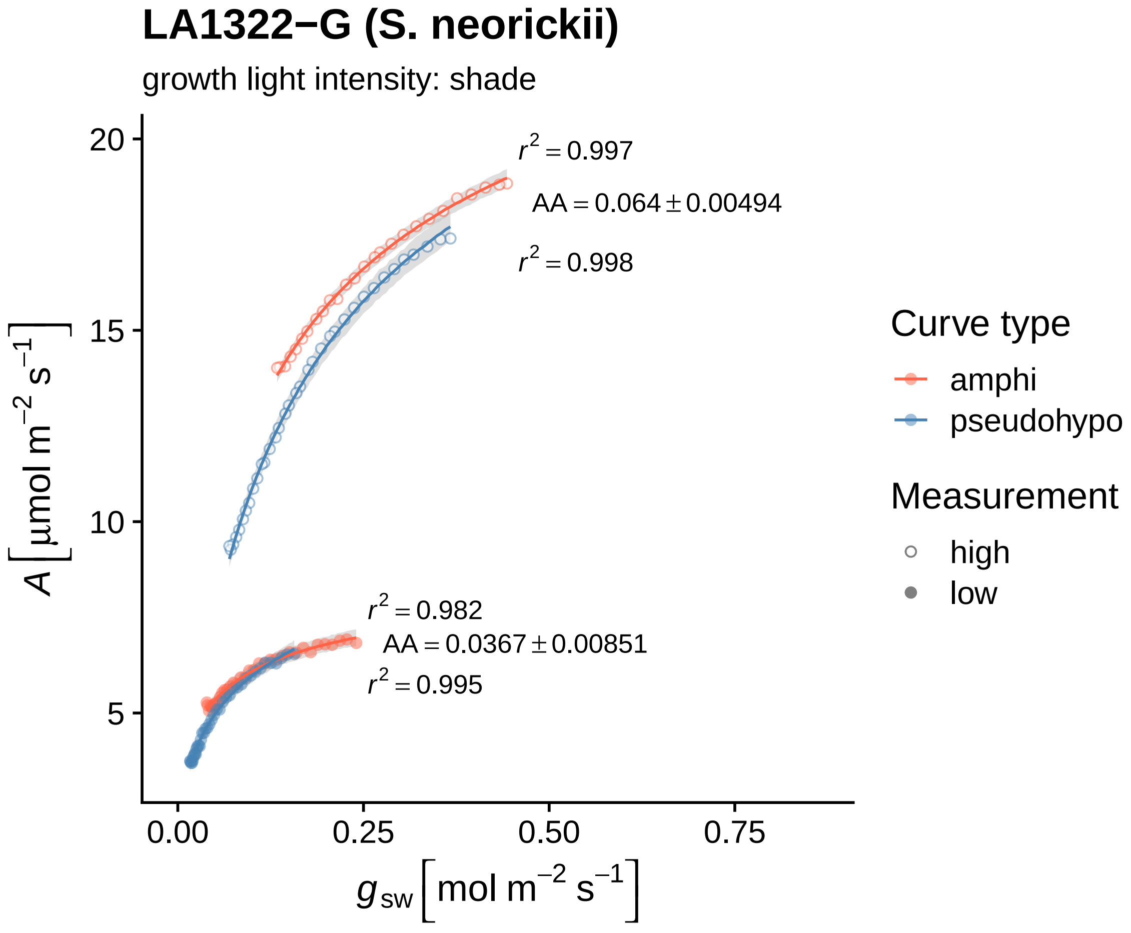
LA1322−G (S. neorickii)
growth light intensity: shade
$r^2 = 0.997$
$AA = 0.064 \pm 0.00494$
$r^2 = 0.998$
$r^2 = 0.982$
$AA = 0.0367 \pm 0.00851$
$r^2 = 0.995$
20
15
10
5
0.00
0.25
0.50
0.75
$A\,[\mu mol\ m^{-2}\ s^{-1}]$
$g_{sw}\,[mol\ m^{-2}\ s^{-1}]$
Curve type
amphi
pseudohypo
Measurement
high
low

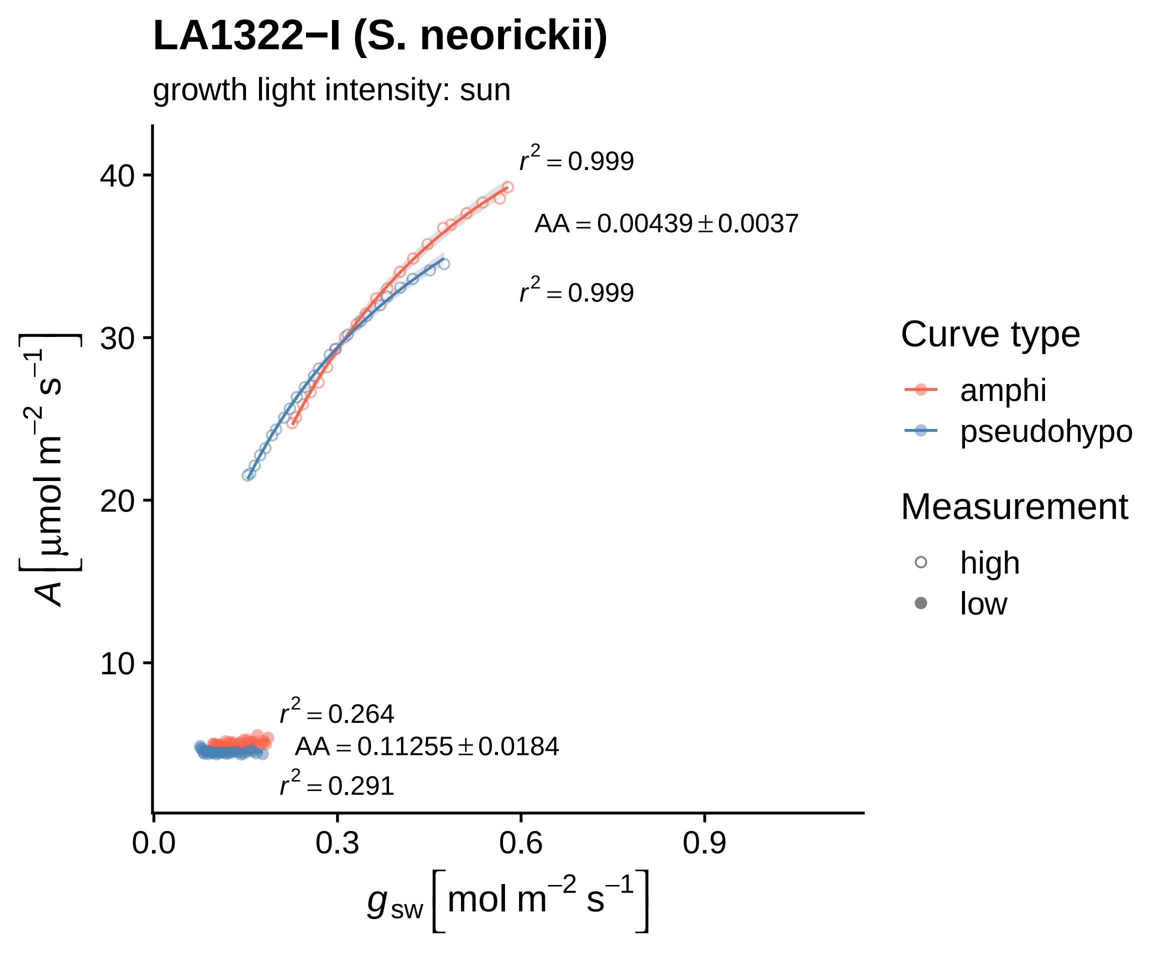

LA1322−I (S. neorickii)
growth light intensity: sun
$r^2 = 0.999$
AA = 0.00439 ± 0.0037
$r^2 = 0.999$
$r^2 = 0.264$
AA = 0.11255 ± 0.0184
$r^2 = 0.291$
40
30
20
10
0.0
0.3
0.6
0.9
$A\ [\mu mol\ m^{-2}\ s^{-1}]$
$g_{sw}\ [mol\ m^{-2}\ s^{-1}]$
Curve type
amphi
pseudohypo
Measurement
high
low

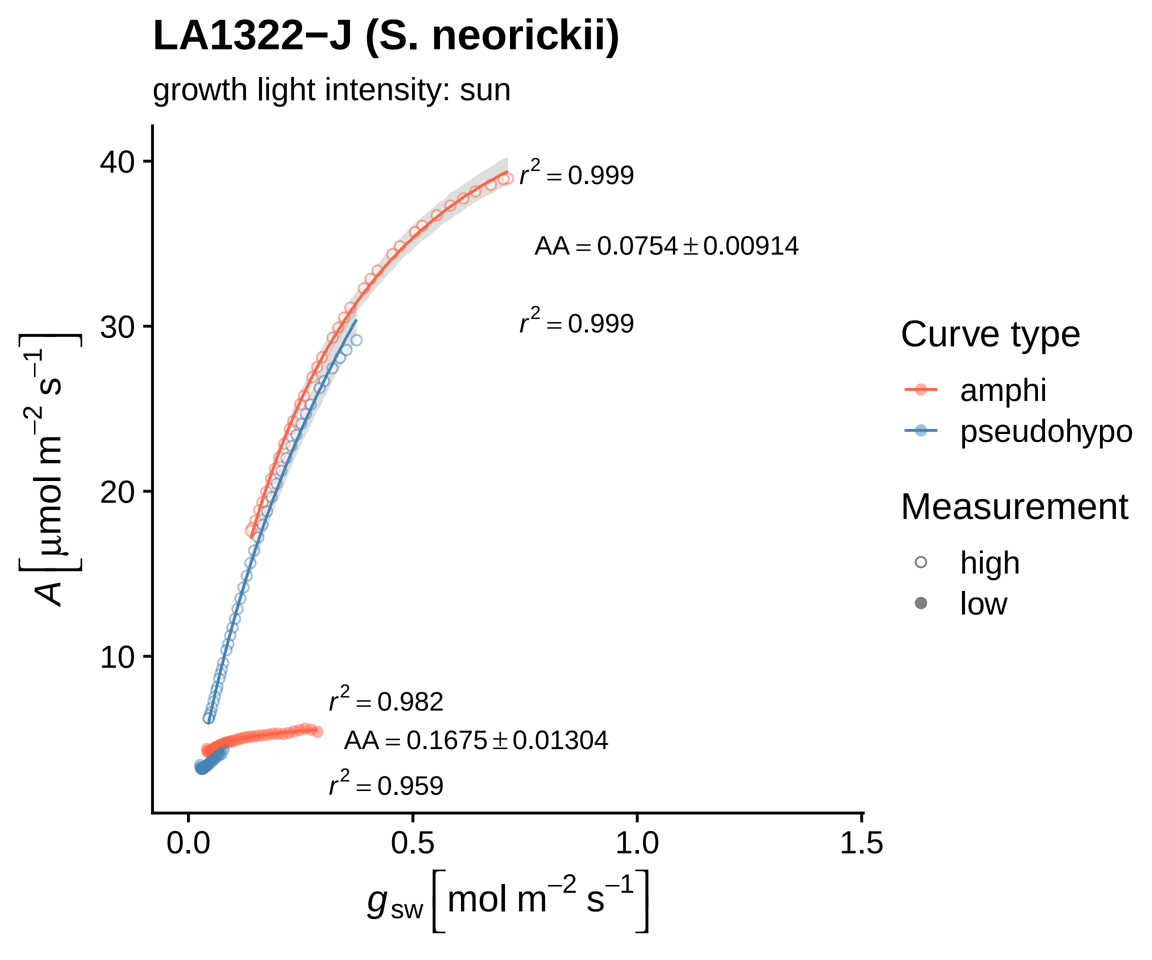

LA1322−J (S. neorickii)
growth light intensity: sun
r² = 0.999
AA = 0.0754 ± 0.00914
r² = 0.999
r² = 0.982
AA = 0.1675 ± 0.01304
r² = 0.959
40
30
20
10
0.0
0.5
1.0
1.5
A [μmol m⁻² s⁻¹]
g sw [mol m⁻² s⁻¹]
Curve type
amphi
pseudohypo
Measurement
high
low

### LA1322−K (S. neorickii)

growth light intensity: sun

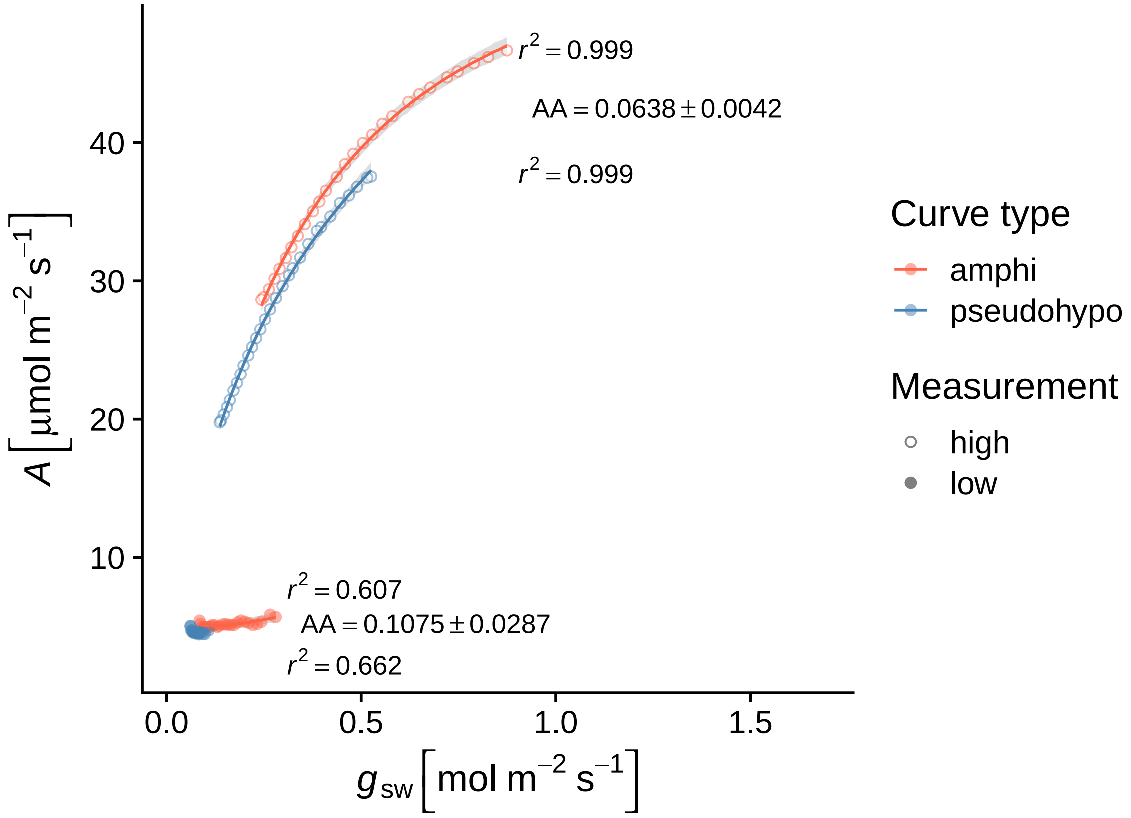

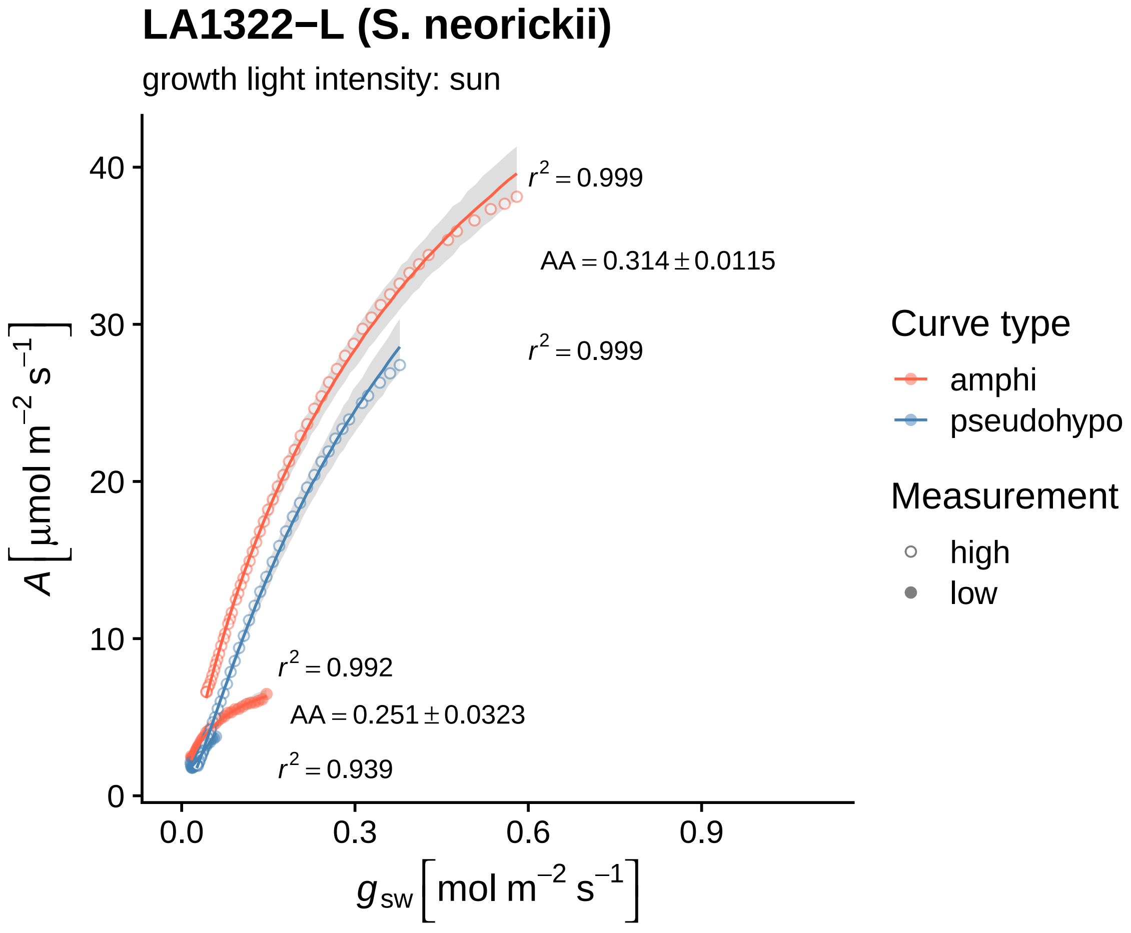
LA1322−L (S. neorickii)
growth light intensity: sun
$r^2 = 0.999$
AA = 0.314 ± 0.0115
$r^2 = 0.999$
$r^2 = 0.992$
AA = 0.251 ± 0.0323
$r^2 = 0.939$
$A\ [\mu mol\ m^{-2}\ s^{-1}]$
$g_{sw}\ [mol\ m^{-2}\ s^{-1}]$
0
10
20
30
40
0.0
0.3
0.6
0.9
Curve type
amphi
pseudohypo
Measurement
high
low

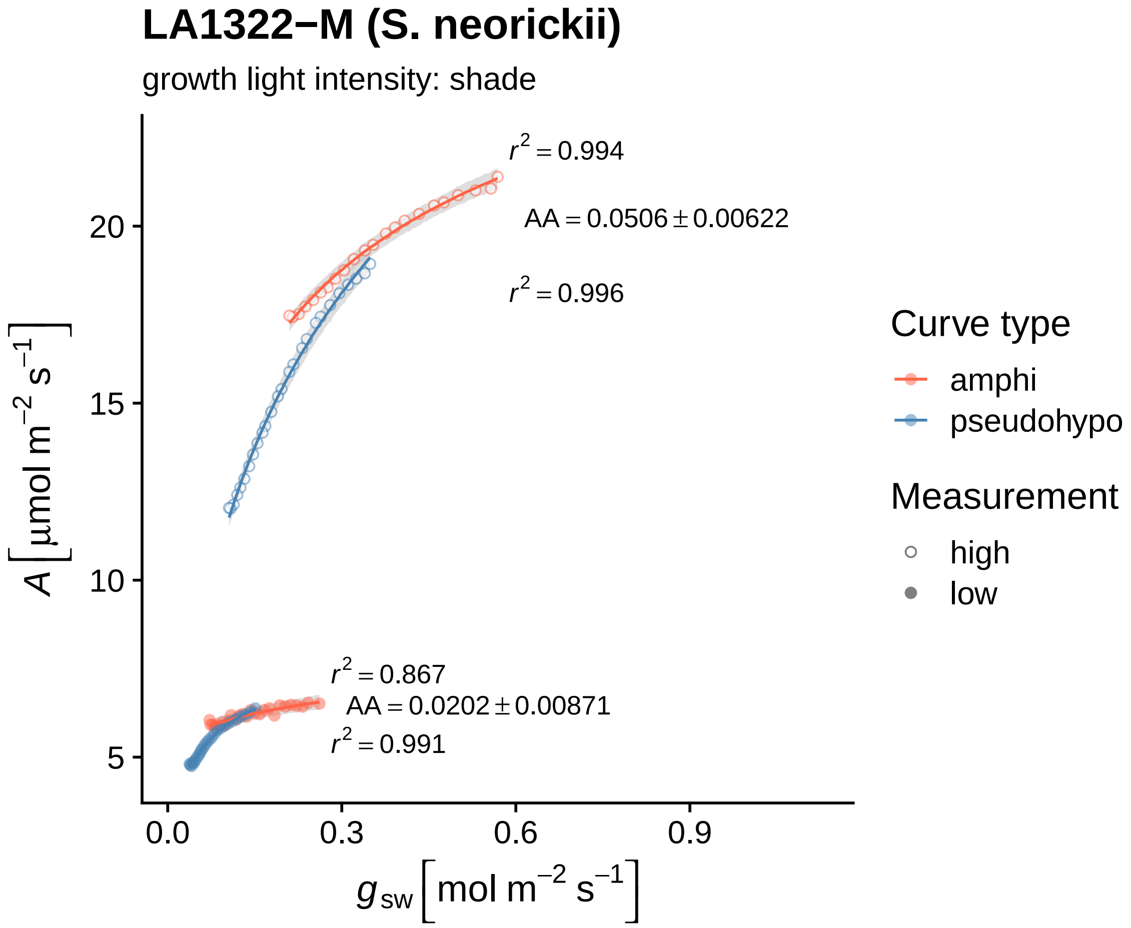

LA1322−M (S. neorickii)
growth light intensity: shade
r² = 0.994
AA = 0.0506 ± 0.00622
r² = 0.996
r² = 0.867
AA = 0.0202 ± 0.00871
r² = 0.991
20
15
10
5
0.0
0.3
0.6
0.9
A [µmol m⁻² s⁻¹]
g_sw [mol m⁻² s⁻¹]
Curve type
amphi
pseudohypo
Measurement
high
low

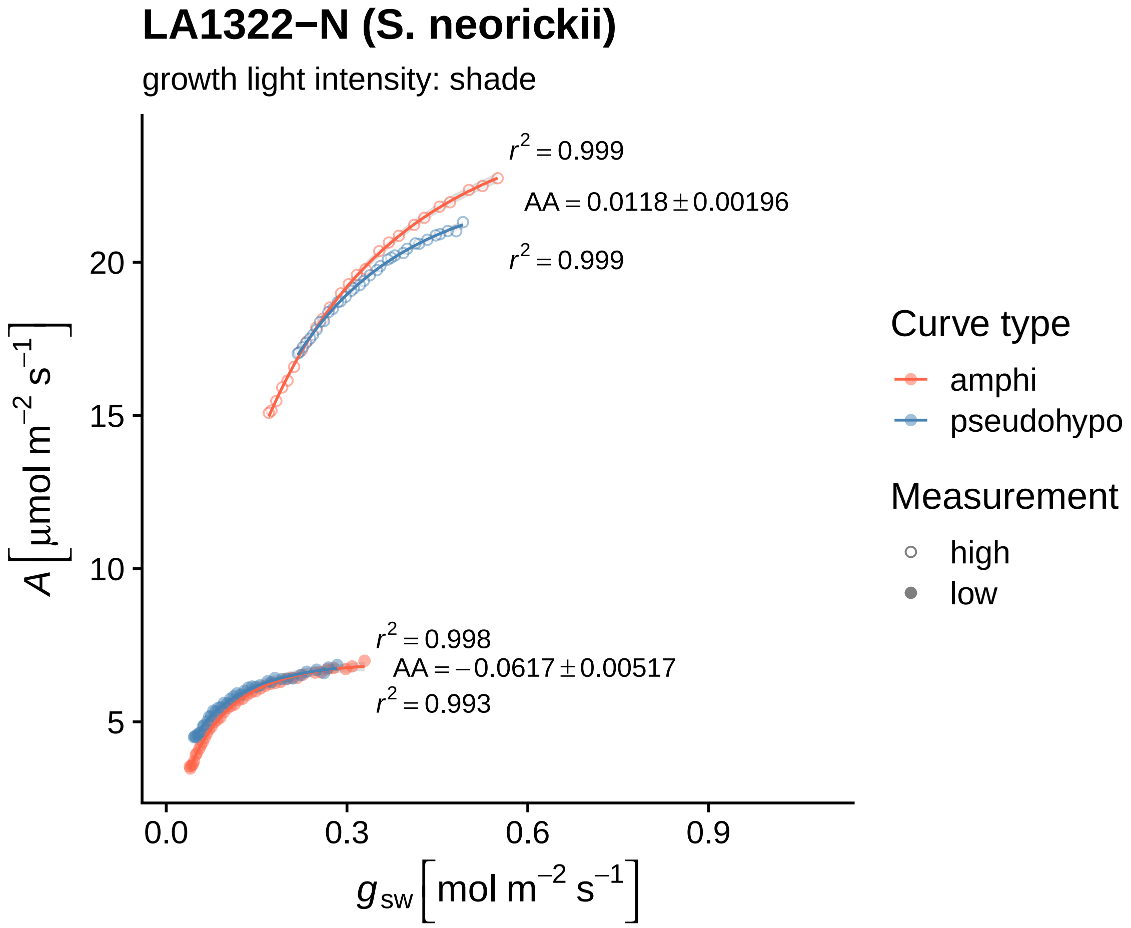

LA1322−N (S. neorickii)
growth light intensity: shade
$r^2 = 0.999$
$AA = 0.0118 \pm 0.00196$
$r^2 = 0.999$
$r^2 = 0.998$
$AA = -0.0617 \pm 0.00517$
$r^2 = 0.993$
20
15
10
5
0.0
0.3
0.6
0.9
$A\ [\mu mol\ m^{-2}\ s^{-1}]$
$g_{sw}\ [mol\ m^{-2}\ s^{-1}]$
Curve type
amphi
pseudohypo
Measurement
high
low

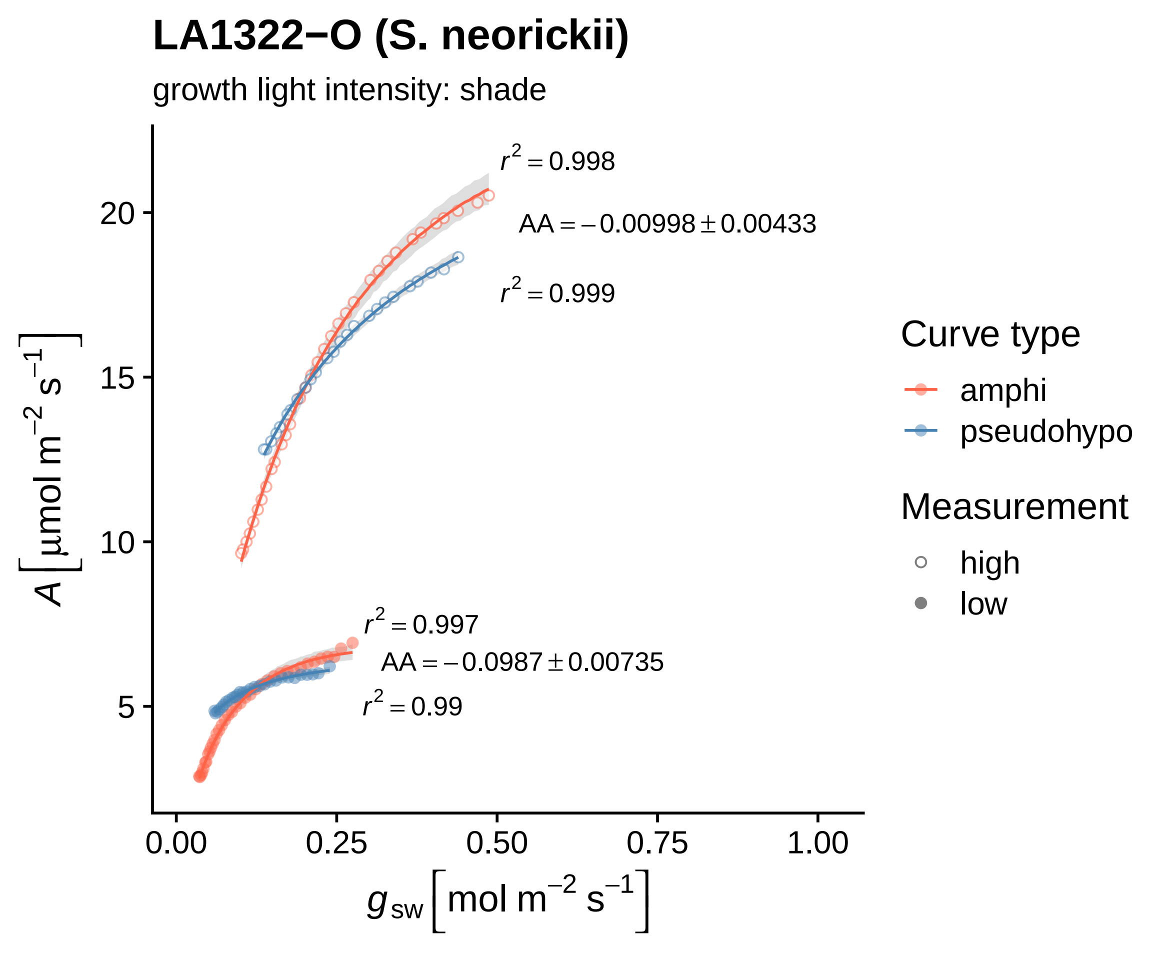
LA1322−O (S. neorickii)
growth light intensity: shade
$r^2 = 0.998$
$AA = -0.00998 \pm 0.00433$
$r^2 = 0.999$
$r^2 = 0.997$
$AA = -0.0987 \pm 0.00735$
$r^2 = 0.99$
$A\ [\mu mol\ m^{-2}\ s^{-1}]$
20
15
10
5
0.00
0.25
0.50
0.75
1.00
$g_{sw}\ [mol\ m^{-2}\ s^{-1}]$
Curve type
amphi
pseudohypo
Measurement
high
low

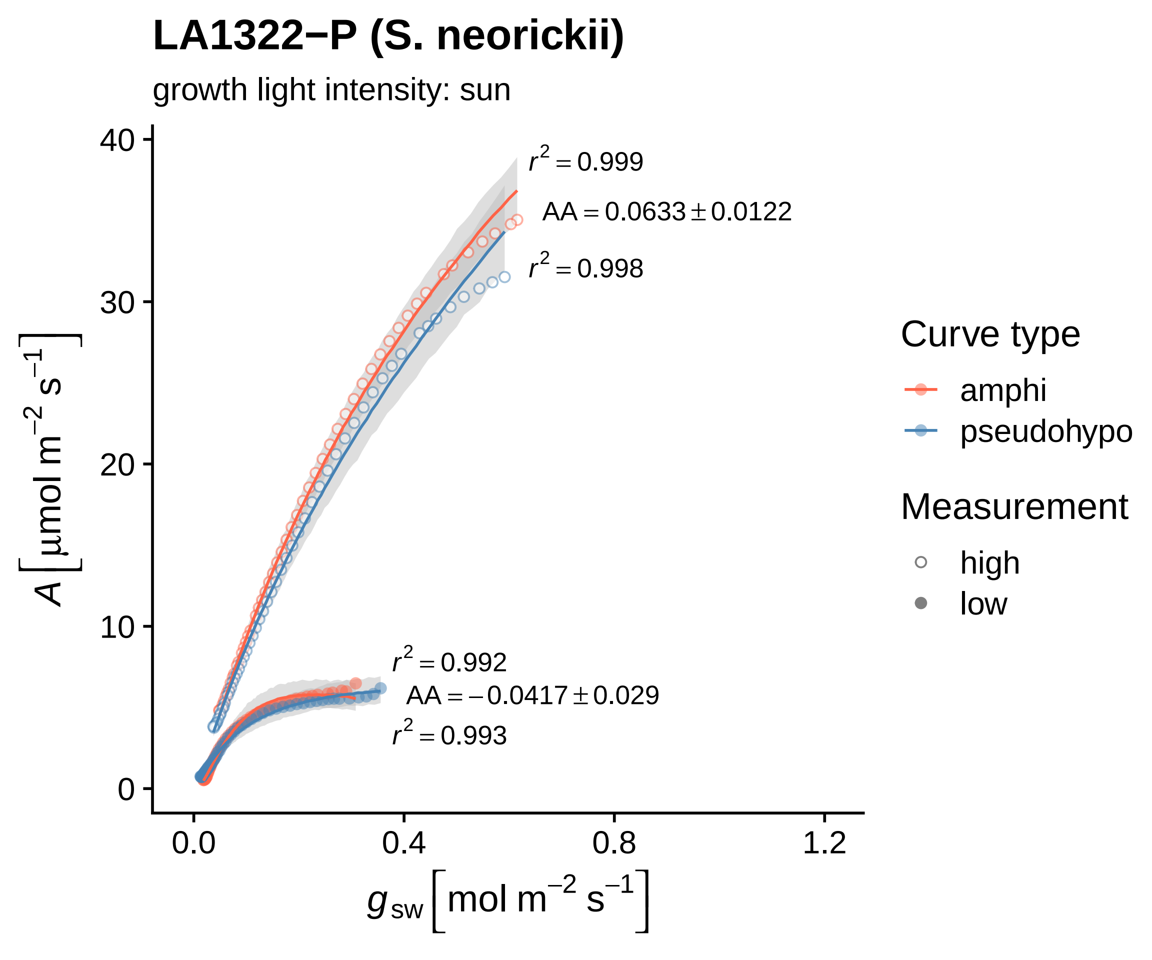
LA1322−P (S. neorickii)
growth light intensity: sun
$r^2 = 0.999$
$AA = 0.0633 \pm 0.0122$
$r^2 = 0.998$
$r^2 = 0.992$
$AA = -0.0417 \pm 0.029$
$r^2 = 0.993$
$A\ [\mu mol\ m^{-2}\ s^{-1}]$
$g_{sw}\ [mol\ m^{-2}\ s^{-1}]$
0
10
20
30
40
0.0
0.4
0.8
1.2
Curve type
amphi
pseudohypo
Measurement
high
low

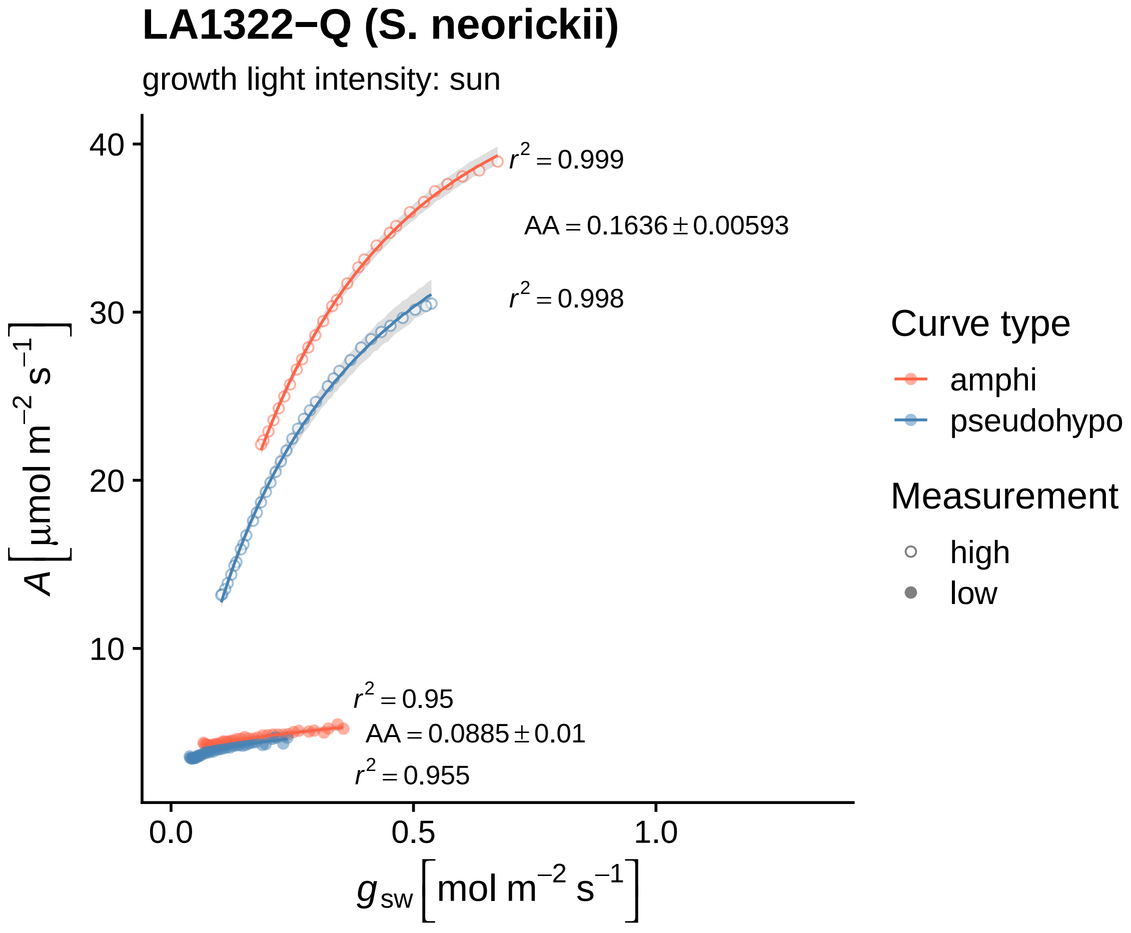
LA1322−Q (S. neorickii)
growth light intensity: sun
A [µmol m−2 s−1]
gsw [mol m−2 s−1]
40
30
20
10
0.0
0.5
1.0
r2 = 0.999
AA = 0.1636 ± 0.00593
r2 = 0.998
r2 = 0.95
AA = 0.0885 ± 0.01
r2 = 0.955
Curve type
amphi
pseudohypo
Measurement
high
low

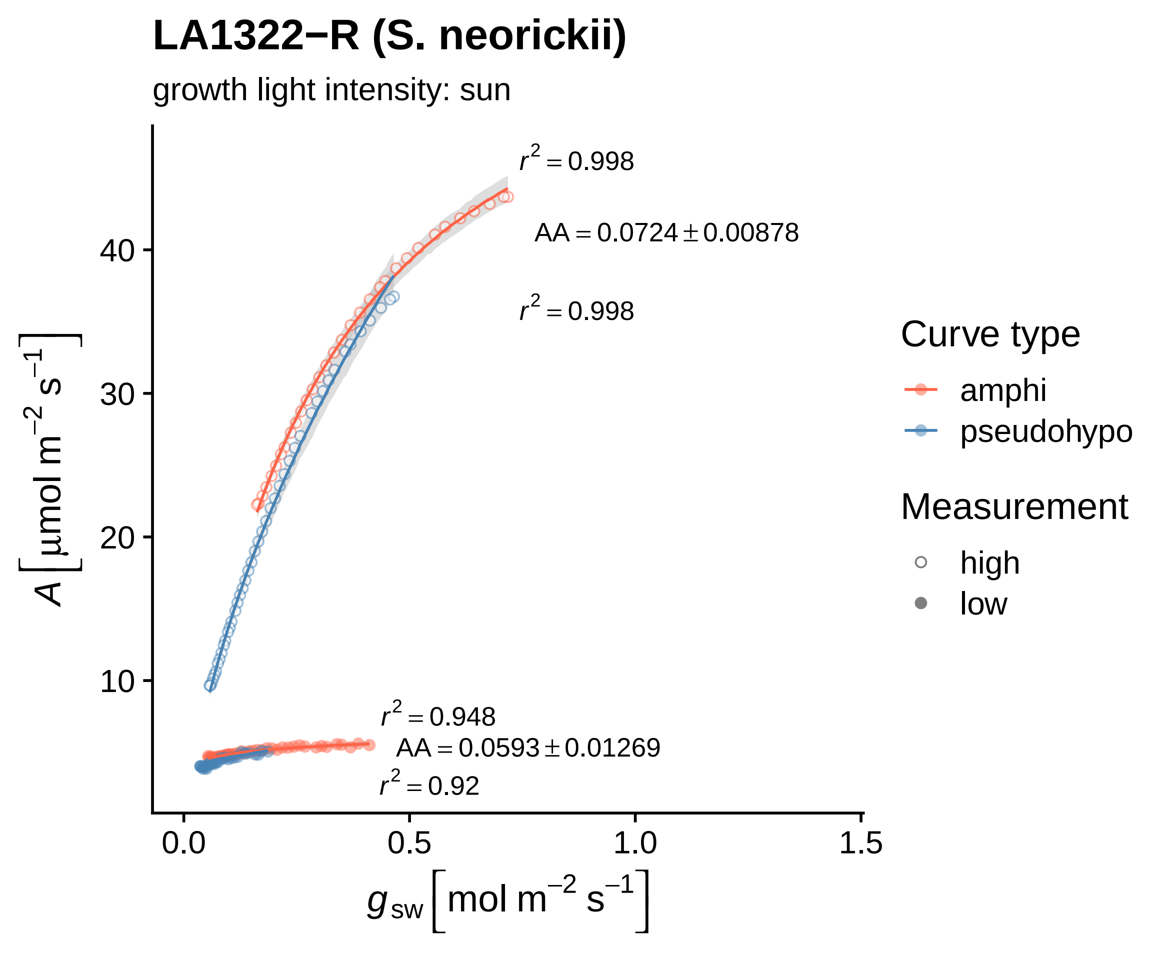

LA1322−R (S. neorickii)
growth light intensity: sun
$r^2 = 0.998$
AA = 0.0724 ± 0.00878
$r^2 = 0.998$
$r^2 = 0.948$
AA = 0.0593 ± 0.01269
$r^2 = 0.92$
10
20
30
40
0.0
0.5
1.0
1.5
$A\ [\mu mol\ m^{-2}\ s^{-1}]$
$g_{sw}\ [mol\ m^{-2}\ s^{-1}]$
Curve type
amphi
pseudohypo
Measurement
high
low

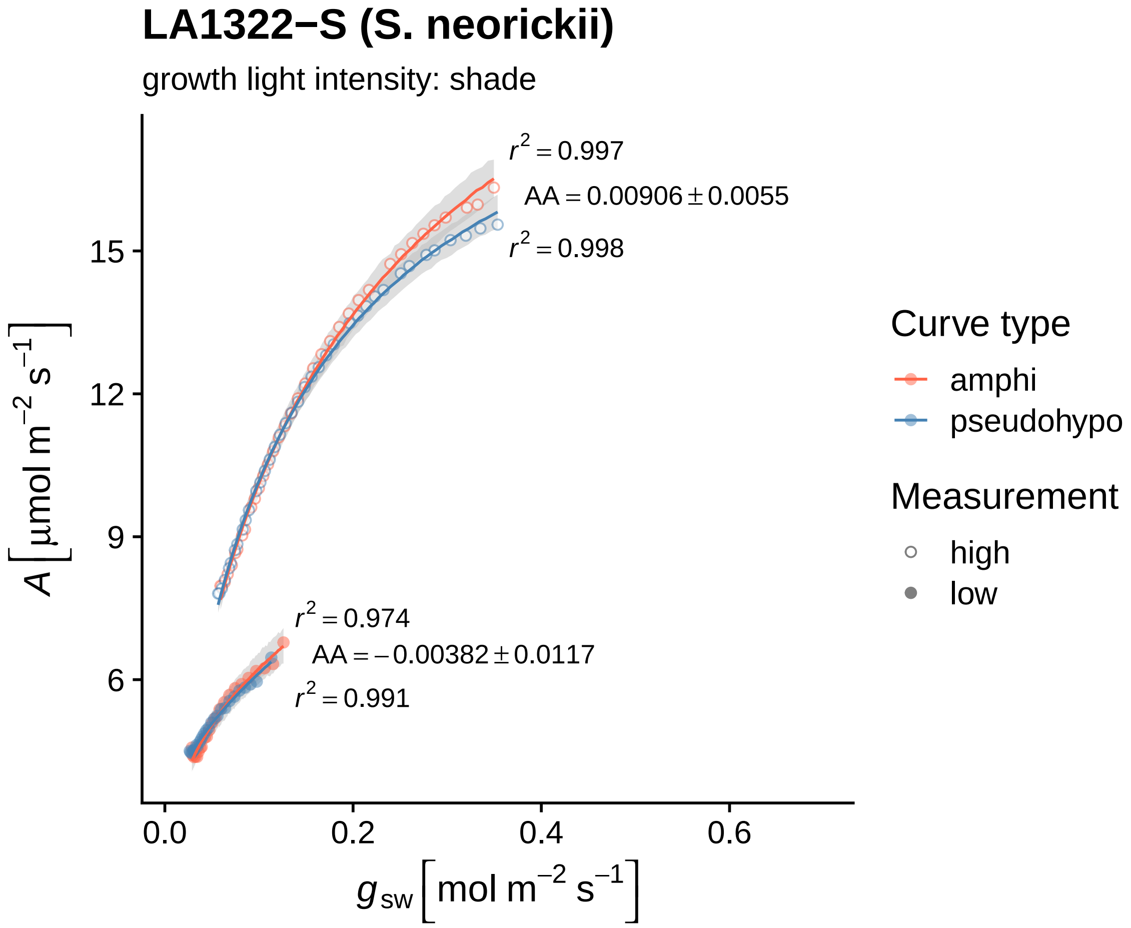
LA1322−S (S. neorickii)
growth light intensity: shade
r² = 0.997
AA = 0.00906 ± 0.0055
r² = 0.998
r² = 0.974
AA = −0.00382 ± 0.0117
r² = 0.991
15
12
9
6
0.0
0.2
0.4
0.6
A [μmol m⁻² s⁻¹]
g_sw [mol m⁻² s⁻¹]
Curve type
amphi
pseudohypo
Measurement
high
low

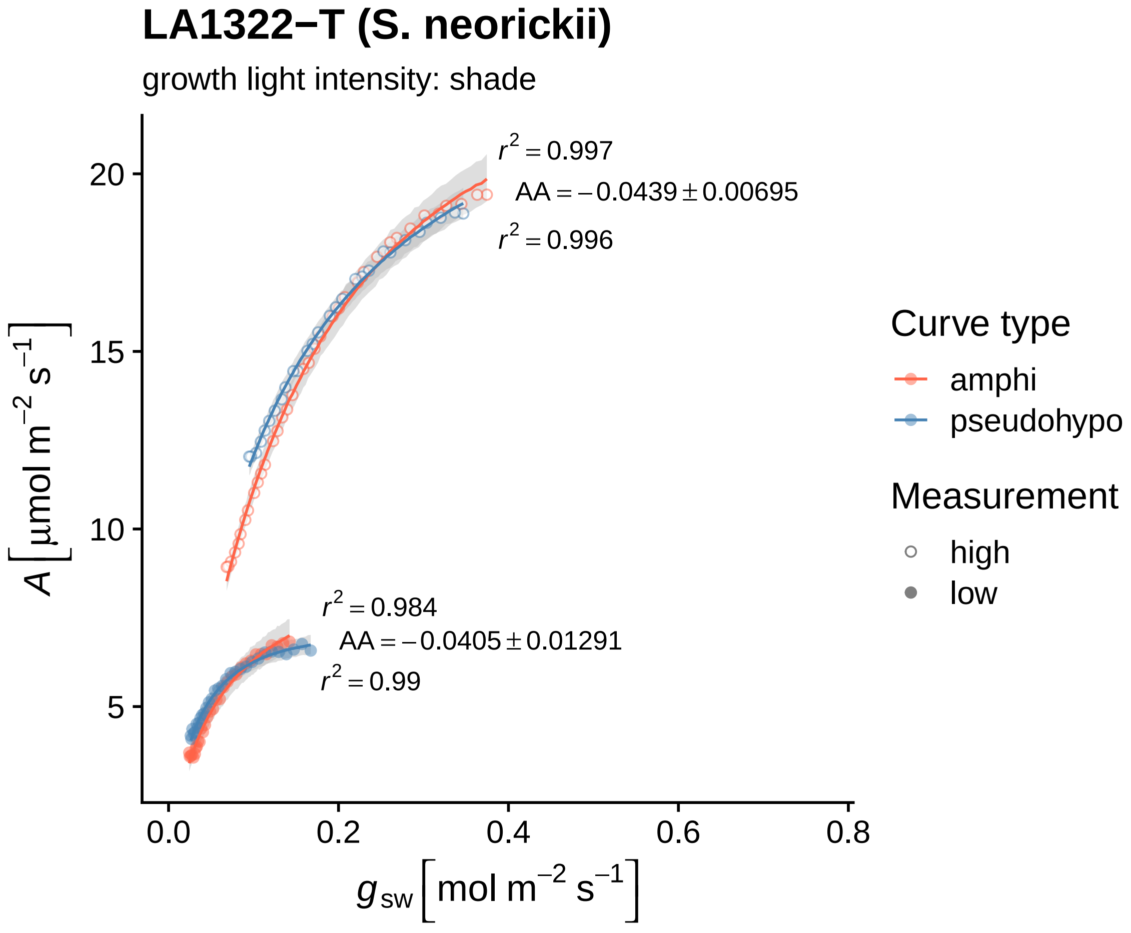

LA1322−T (S. neorickii)
growth light intensity: shade
$r^2 = 0.997$
$AA = -0.0439 \pm 0.00695$
$r^2 = 0.996$
$r^2 = 0.984$
$AA = -0.0405 \pm 0.01291$
$r^2 = 0.99$
20
15
10
5
0.0
0.2
0.4
0.6
0.8
$A\left[\mu mol\ m^{-2}\ s^{-1}\right]$
$g_{sw}\left[mol\ m^{-2}\ s^{-1}\right]$
Curve type
amphi
pseudohypo
Measurement
high
low

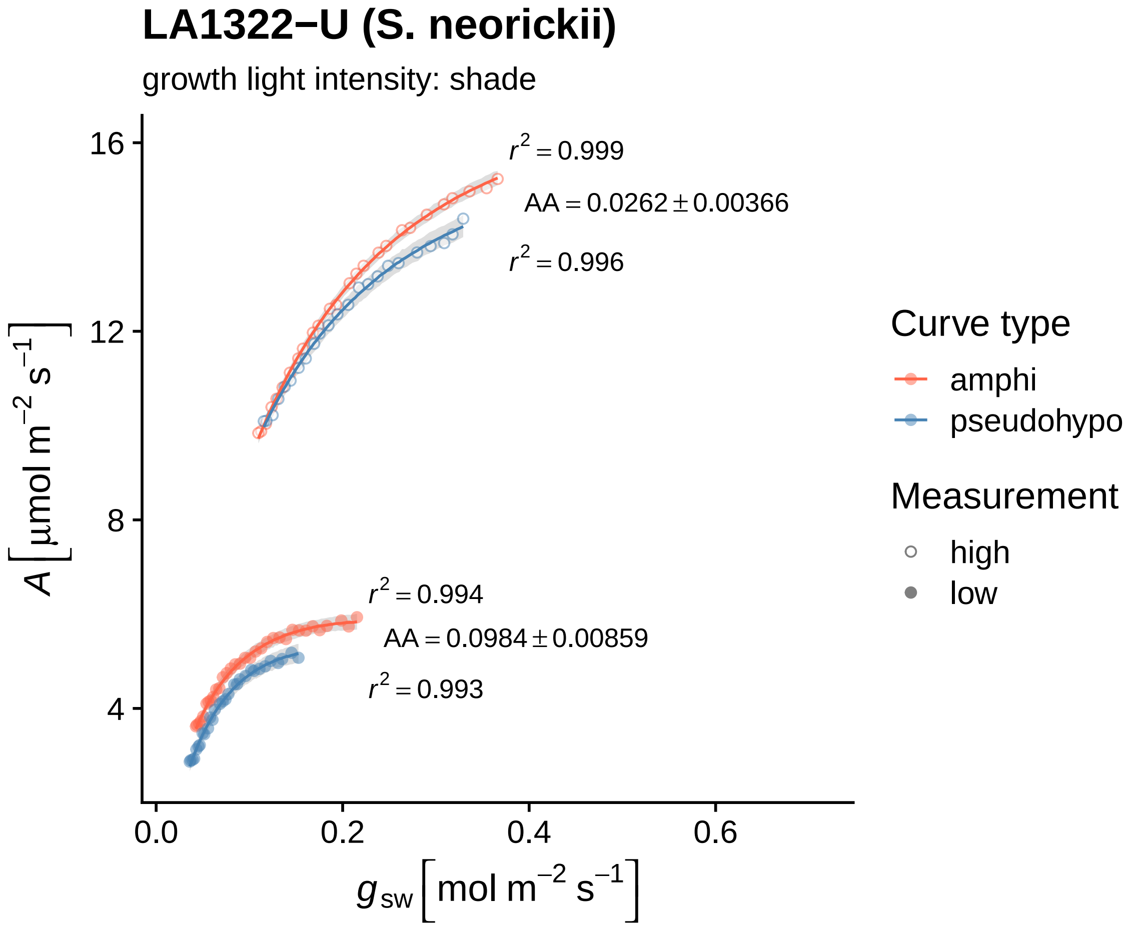

LA1322−U (S. neorickii)
growth light intensity: shade
$r^2 = 0.999$
$AA = 0.0262 \pm 0.00366$
$r^2 = 0.996$
$r^2 = 0.994$
$AA = 0.0984 \pm 0.00859$
$r^2 = 0.993$
16
12
8
4
0.0
0.2
0.4
0.6
$A\ [\mu mol\ m^{-2}\ s^{-1}]$
$g_{sw}\ [mol\ m^{-2}\ s^{-1}]$
Curve type
amphi
pseudohypo
Measurement
high
low

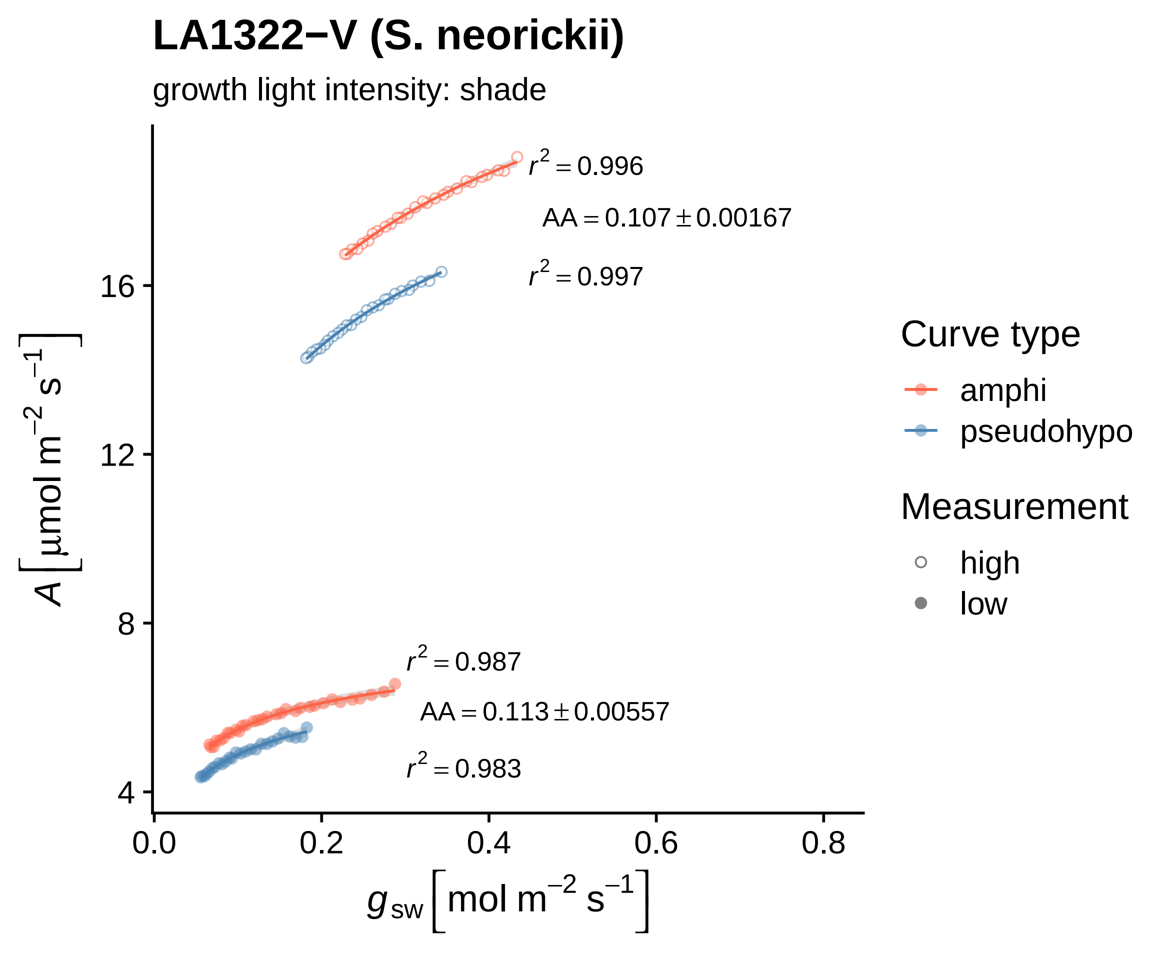
LA1322−V (S. neorickii)
growth light intensity: shade
$r^2 = 0.996$
$AA = 0.107 \pm 0.00167$
$r^2 = 0.997$
$r^2 = 0.987$
$AA = 0.113 \pm 0.00557$
$r^2 = 0.983$
16
12
8
4
0.0
0.2
0.4
0.6
0.8
$A\ [\mu mol\ m^{-2}\ s^{-1}]$
$g_{sw}\ [mol\ m^{-2}\ s^{-1}]$
Curve type
amphi
pseudohypo
Measurement
high
low

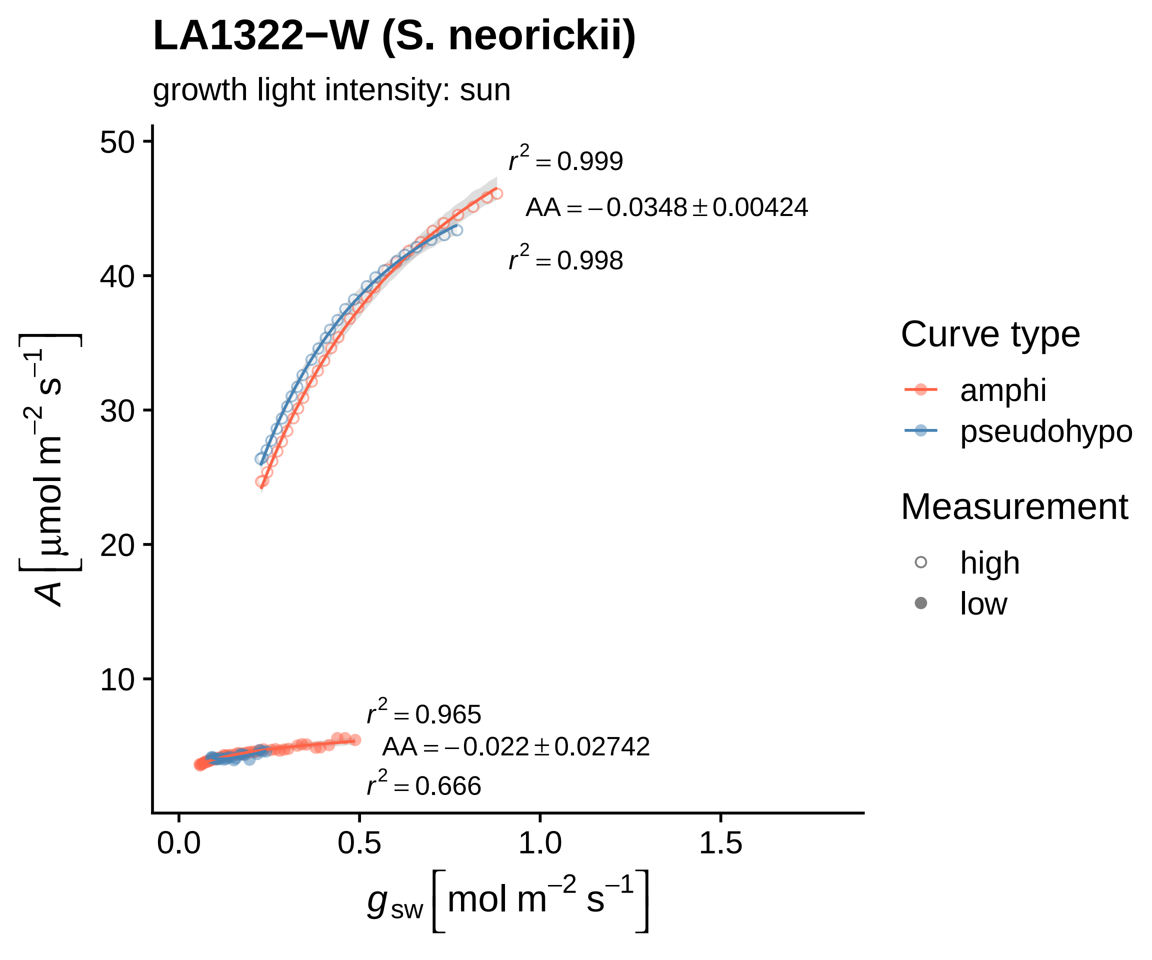

LA1322−W (S. neorickii)
growth light intensity: sun
$r^2 = 0.999$
$AA = -0.0348 \pm 0.00424$
$r^2 = 0.998$
$r^2 = 0.965$
$AA = -0.022 \pm 0.02742$
$r^2 = 0.666$
$A\ [\mu mol\ m^{-2}\ s^{-1}]$
$g_{sw}\ [mol\ m^{-2}\ s^{-1}]$
0.0
0.5
1.0
1.5
10
20
30
40
50
Curve type
amphi
pseudohypo
Measurement
high
low

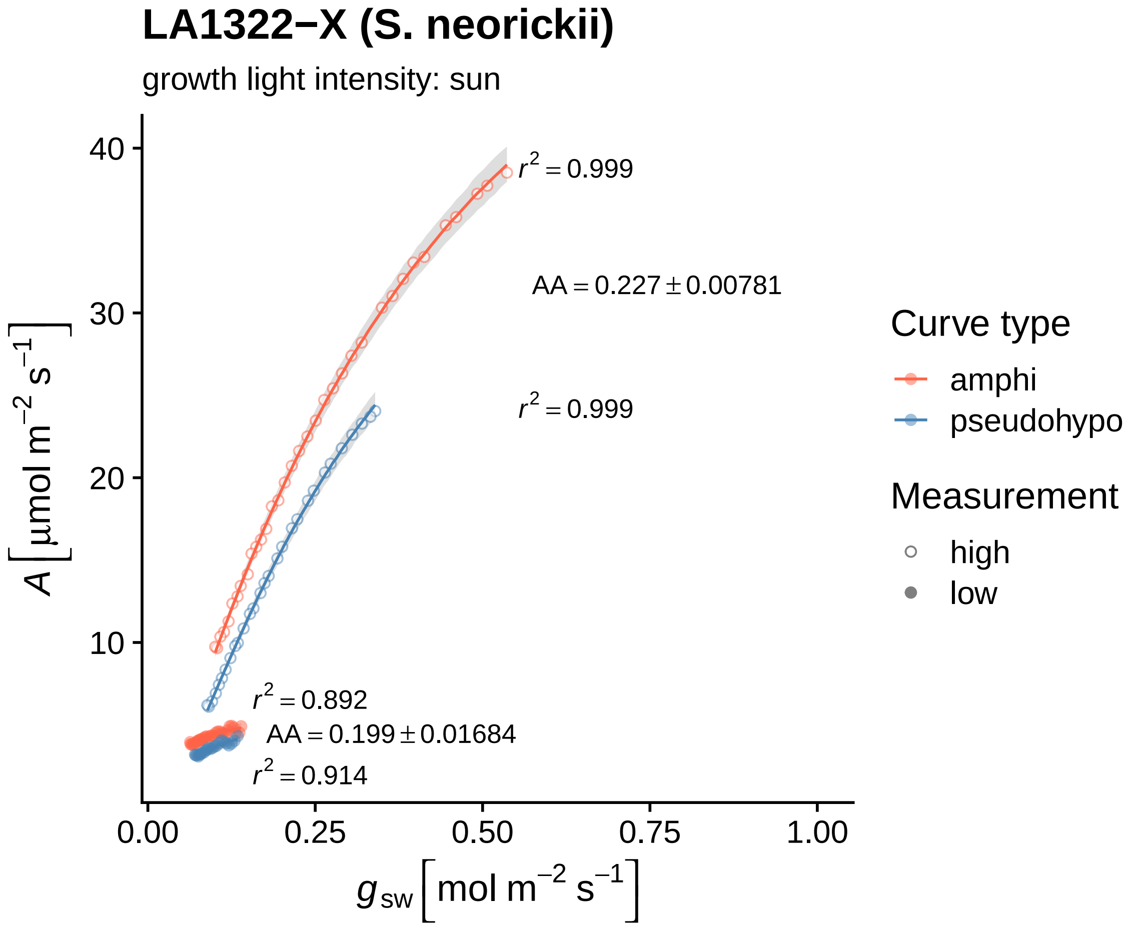
LA1322−X (S. neorickii)
growth light intensity: sun
$r^2 = 0.999$
AA = 0.227 ± 0.00781
$r^2 = 0.999$
$r^2 = 0.892$
AA = 0.199 ± 0.01684
$r^2 = 0.914$
$A$ [μmol m$^{-2}$ s$^{-1}$]
40
30
20
10
0.00
0.25
0.50
0.75
1.00
$g_{sw}$ [mol m$^{-2}$ s$^{-1}$]
Curve type
amphi
pseudohypo
Measurement
high
low

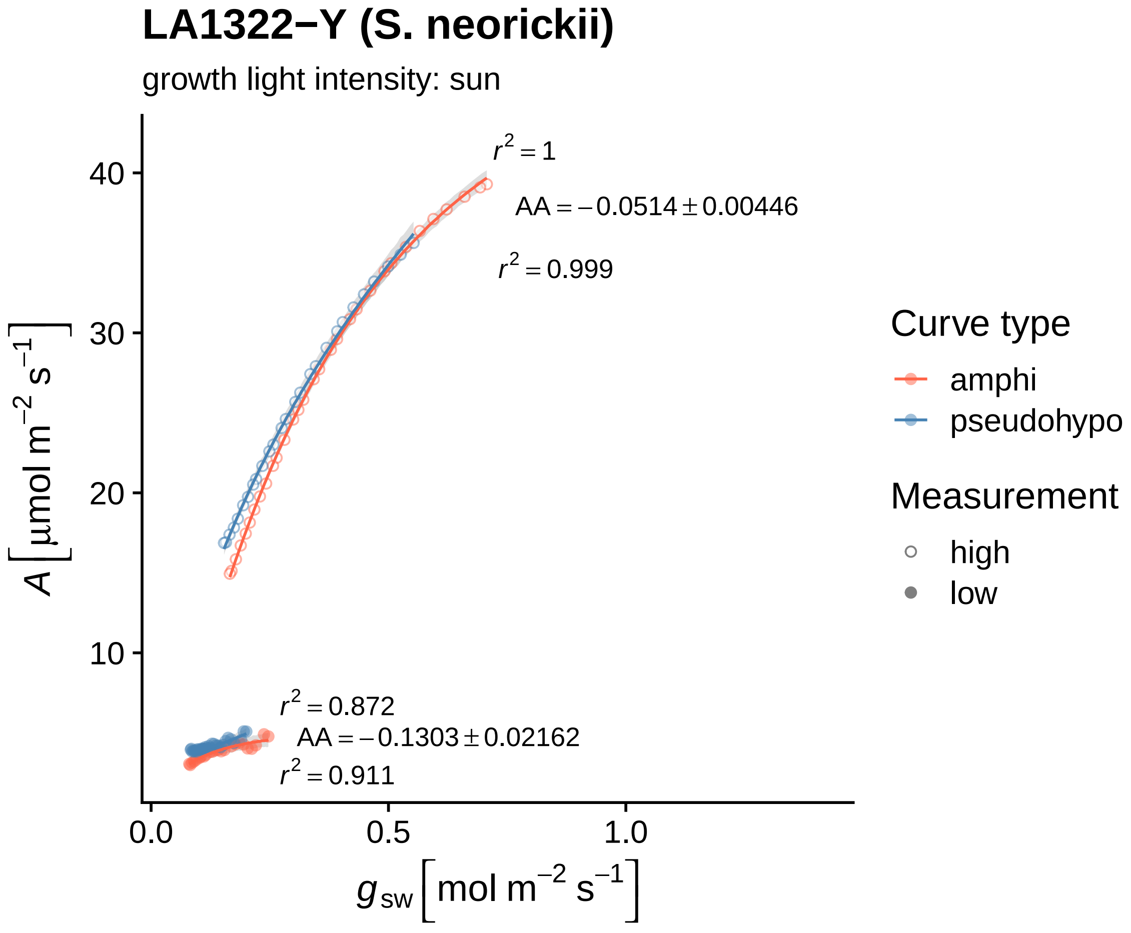

LA1322−Y (S. neorickii)
growth light intensity: sun
$r^2 = 1$
$AA = -0.0514 \pm 0.00446$
$r^2 = 0.999$
$r^2 = 0.872$
$AA = -0.1303 \pm 0.02162$
$r^2 = 0.911$
40
30
20
10
0.0
0.5
1.0
$A\left[\mu mol\ m^{-2}\ s^{-1}\right]$
$g_{sw}\left[mol\ m^{-2}\ s^{-1}\right]$
Curve type
amphi
pseudohypo
Measurement
high
low

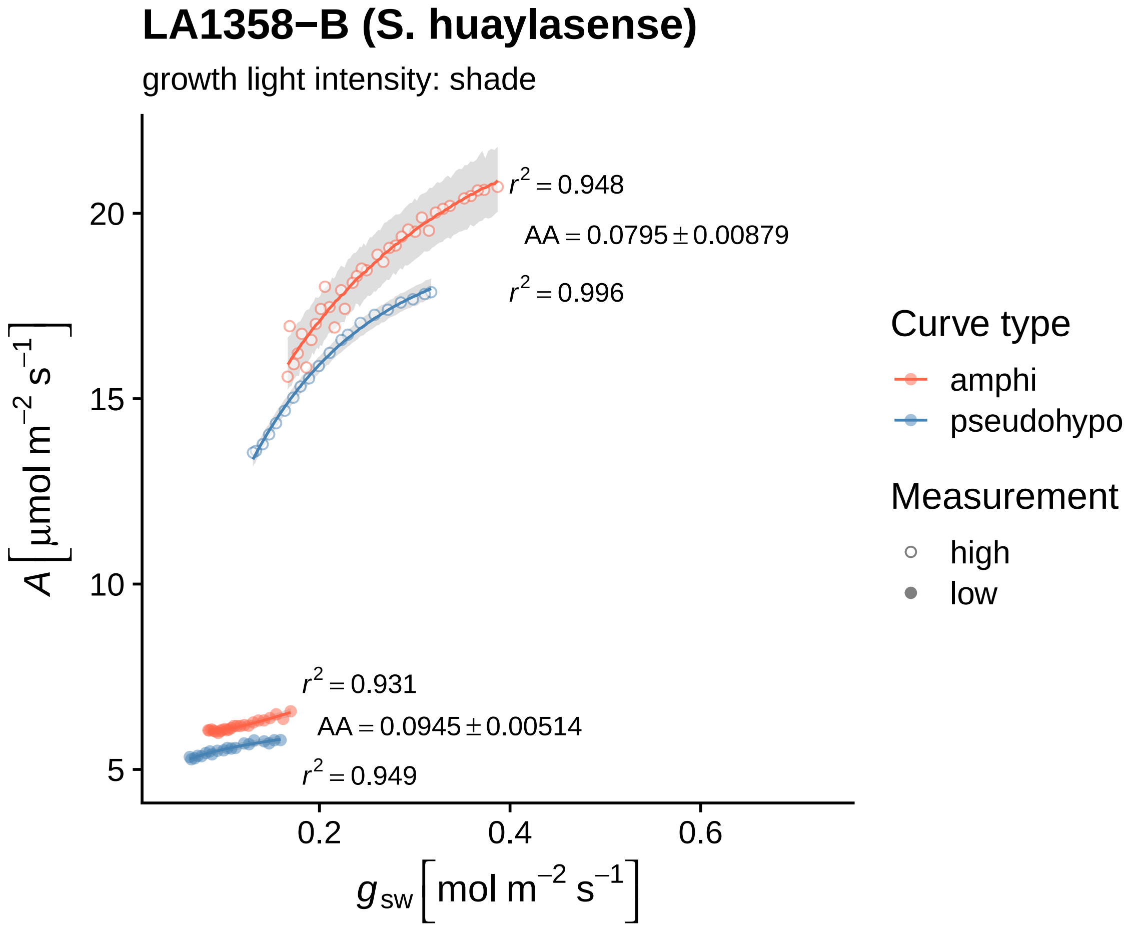
LA1358−B (S. huaylasense)
growth light intensity: shade
r² = 0.948
AA = 0.0795 ± 0.00879
r² = 0.996
r² = 0.931
AA = 0.0945 ± 0.00514
r² = 0.949
20
15
10
5
0.2
0.4
0.6
A [μmol m⁻² s⁻¹]
g sw [mol m⁻² s⁻¹]
Curve type
amphi
pseudohypo
Measurement
high
low

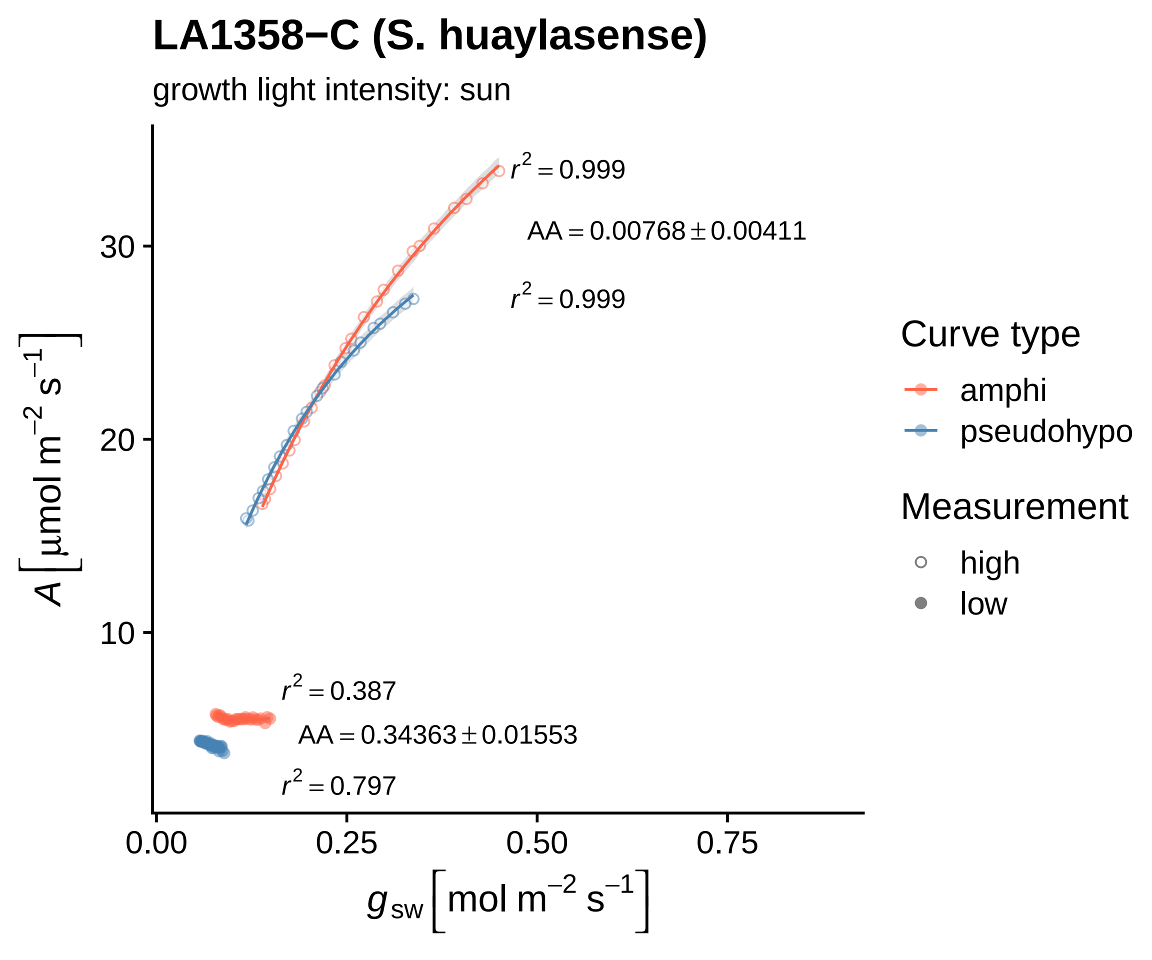

LA1358−C (S. huaylasense)
growth light intensity: sun
r² = 0.999
AA = 0.00768 ± 0.00411
r² = 0.999
r² = 0.387
AA = 0.34363 ± 0.01553
r² = 0.797
30
20
10
0.00
0.25
0.50
0.75
A [µmol m⁻² s⁻¹]
g_sw [mol m⁻² s⁻¹]
Curve type
amphi
pseudohypo
Measurement
high
low

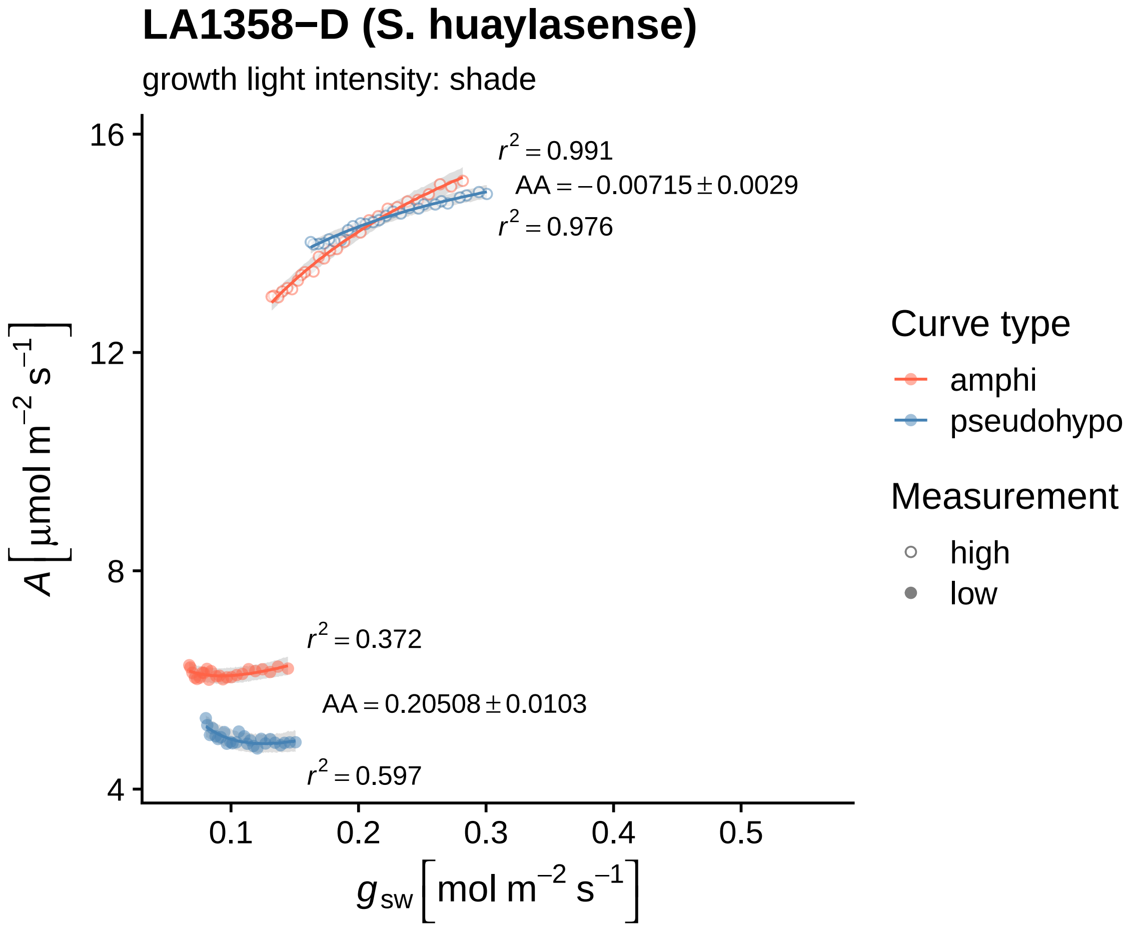

LA1358−D (S. huaylasense)
growth light intensity: shade
$r^2 = 0.991$
$AA = -0.00715 \pm 0.0029$
$r^2 = 0.976$
$r^2 = 0.372$
$AA = 0.20508 \pm 0.0103$
$r^2 = 0.597$
$A\,[\mu mol\ m^{-2}\ s^{-1}]$
$g_{sw}\,[mol\ m^{-2}\ s^{-1}]$
16
12
8
4
0.1
0.2
0.3
0.4
0.5
Curve type
amphi
pseudohypo
Measurement
high
low

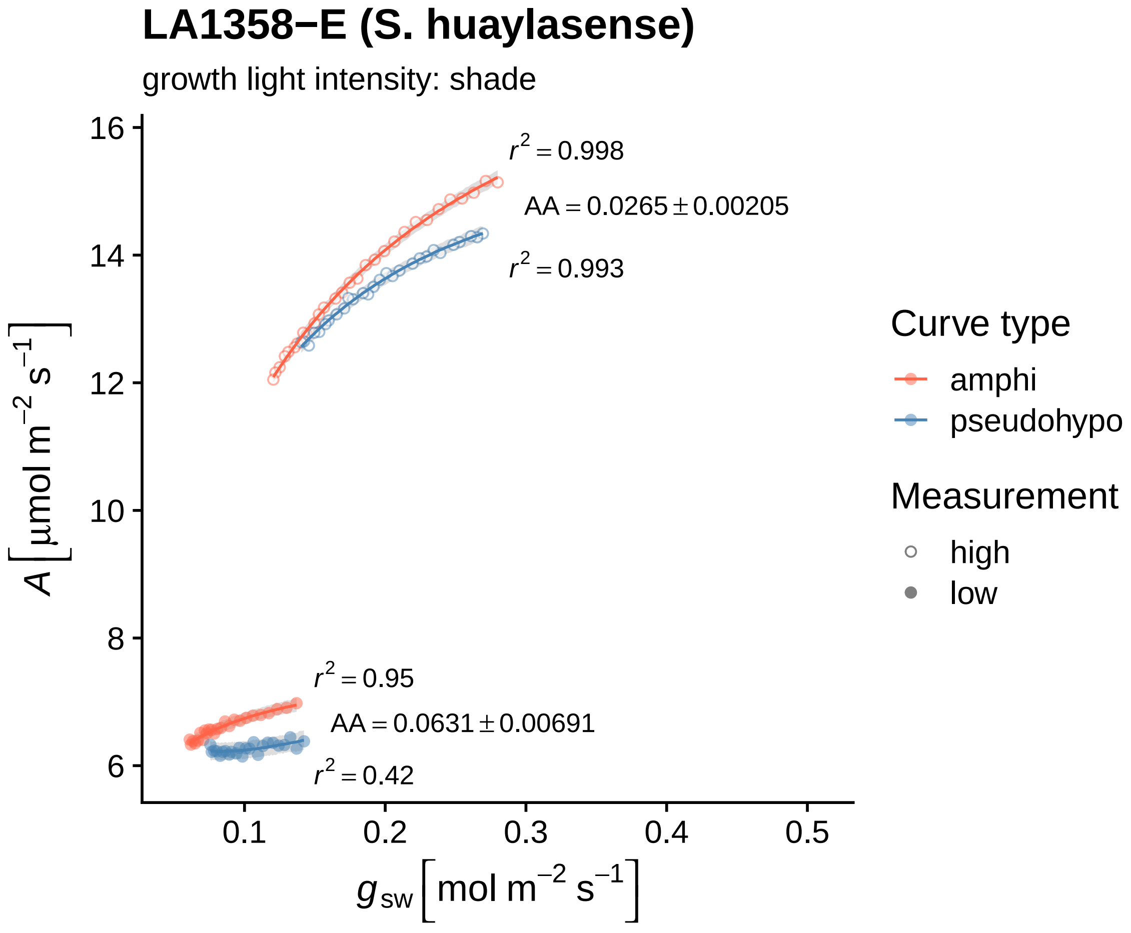

LA1358−E (S. huaylasense)
growth light intensity: shade
$r^2 = 0.998$
$AA = 0.0265 \pm 0.00205$
$r^2 = 0.993$
$r^2 = 0.95$
$AA = 0.0631 \pm 0.00691$
$r^2 = 0.42$
16
14
12
10
8
6
0.1
0.2
0.3
0.4
0.5
$A\ [\mu mol\ m^{-2}\ s^{-1}]$
$g_{sw}\ [mol\ m^{-2}\ s^{-1}]$
Curve type
amphi
pseudohypo
Measurement
high
low

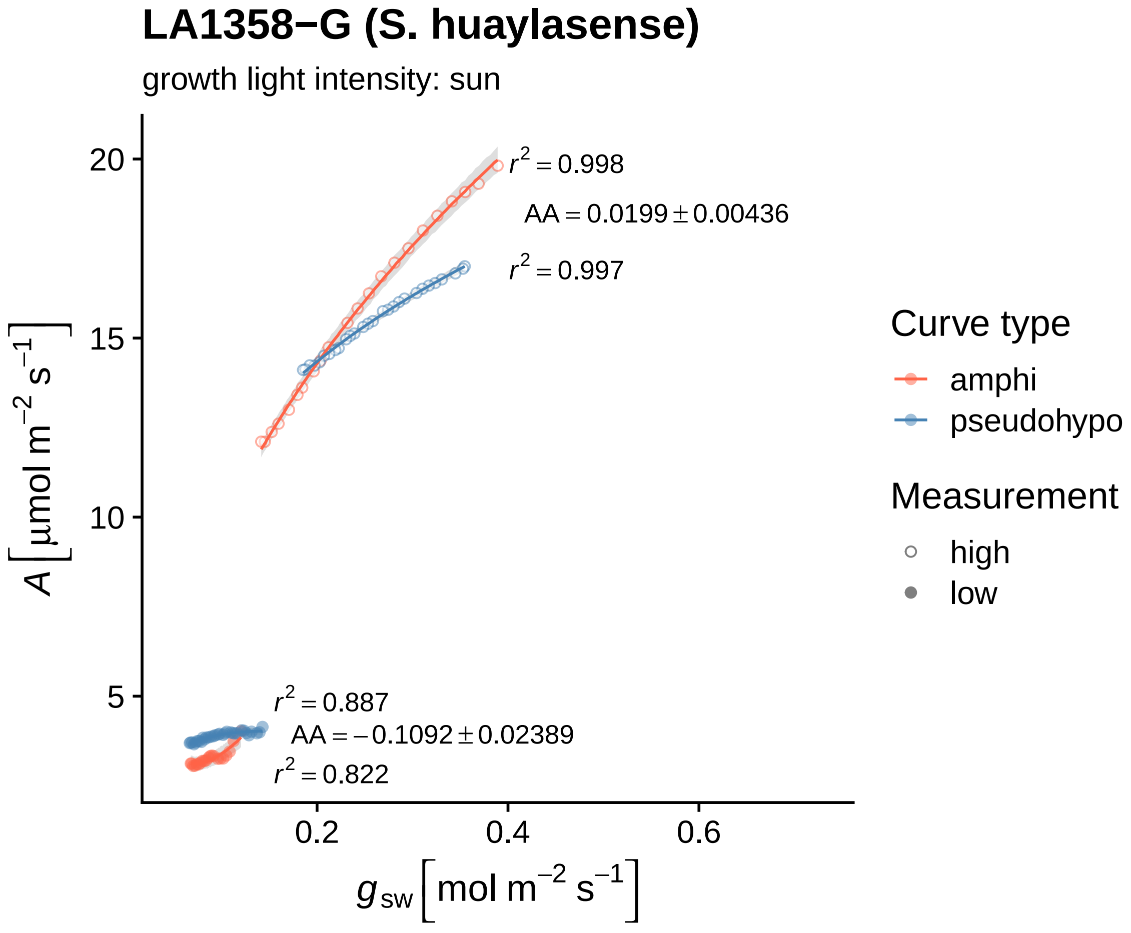

LA1358−G (S. huaylasense)
growth light intensity: sun
$r^2 = 0.998$
$AA = 0.0199 \pm 0.00436$
$r^2 = 0.997$
$r^2 = 0.887$
$AA = -0.1092 \pm 0.02389$
$r^2 = 0.822$
20
15
10
5
0.2
0.4
0.6
$A\ [\mu mol\ m^{-2}\ s^{-1}]$
$g_{sw}\ [mol\ m^{-2}\ s^{-1}]$
Curve type
amphi
pseudohypo
Measurement
high
low

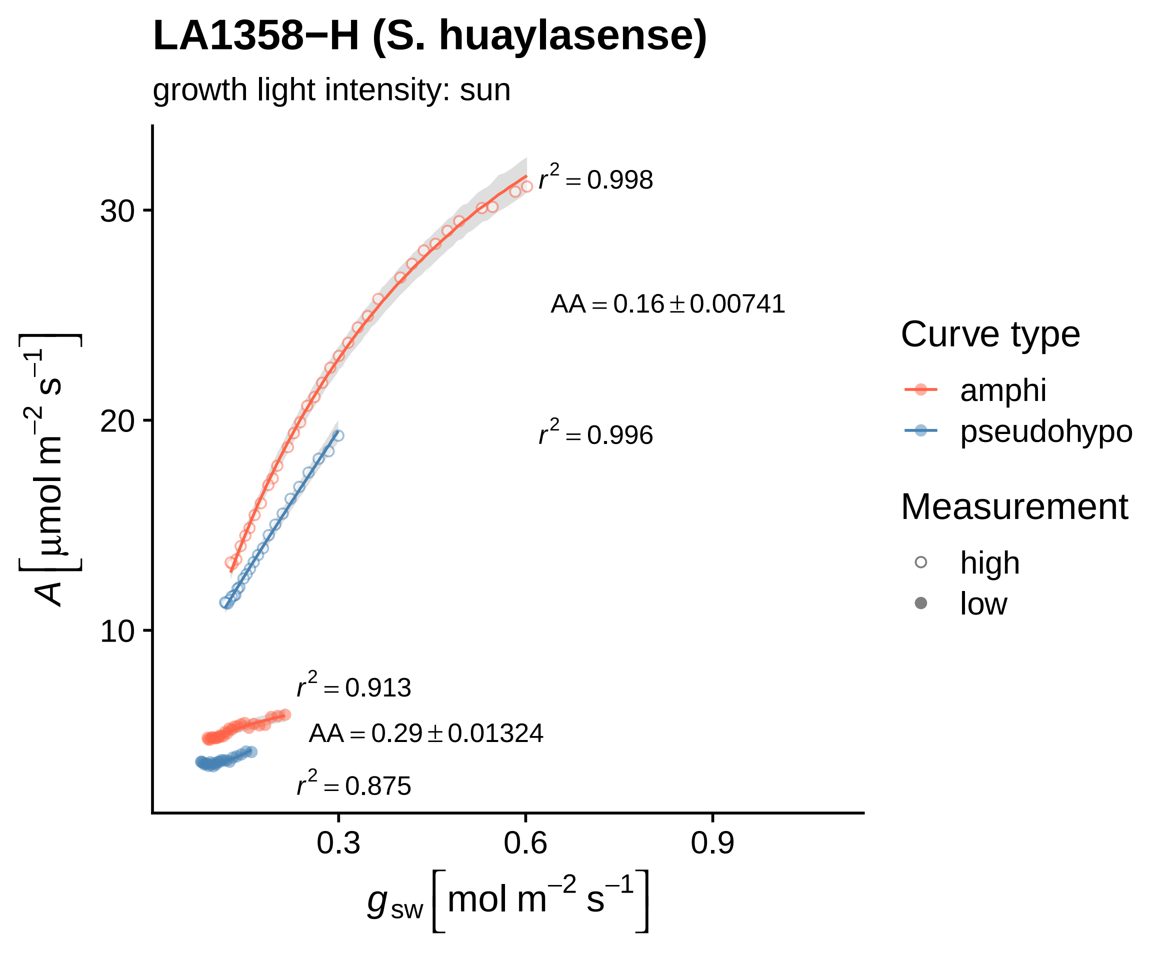
LA1358−H (S. huaylasense)
growth light intensity: sun
$r^2 = 0.998$
$AA = 0.16 \pm 0.00741$
$r^2 = 0.996$
$r^2 = 0.913$
$AA = 0.29 \pm 0.01324$
$r^2 = 0.875$
30
20
10
0.3
0.6
0.9
$A\ [\mu mol\ m^{-2}\ s^{-1}]$
$g_{sw}\ [mol\ m^{-2}\ s^{-1}]$
Curve type
amphi
pseudohypo
Measurement
high
low

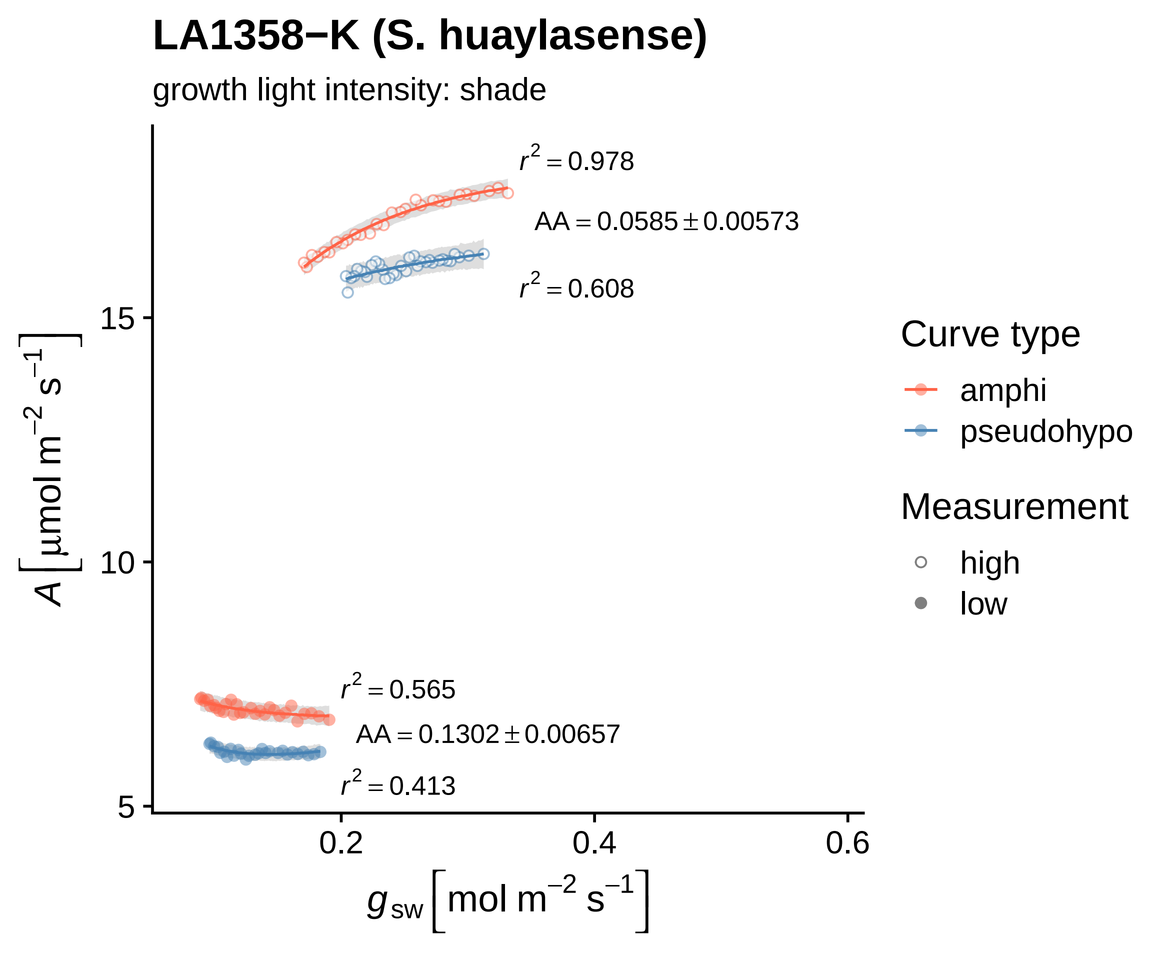

LA1358−K (S. huaylasense)
growth light intensity: shade
$r^2 = 0.978$
$AA = 0.0585 \pm 0.00573$
$r^2 = 0.608$
$r^2 = 0.565$
$AA = 0.1302 \pm 0.00657$
$r^2 = 0.413$
15
10
5
0.2
0.4
0.6
$A\left[\mu mol\ m^{-2}\ s^{-1}\right]$
$g_{sw}\left[mol\ m^{-2}\ s^{-1}\right]$
Curve type
amphi
pseudohypo
Measurement
high
low

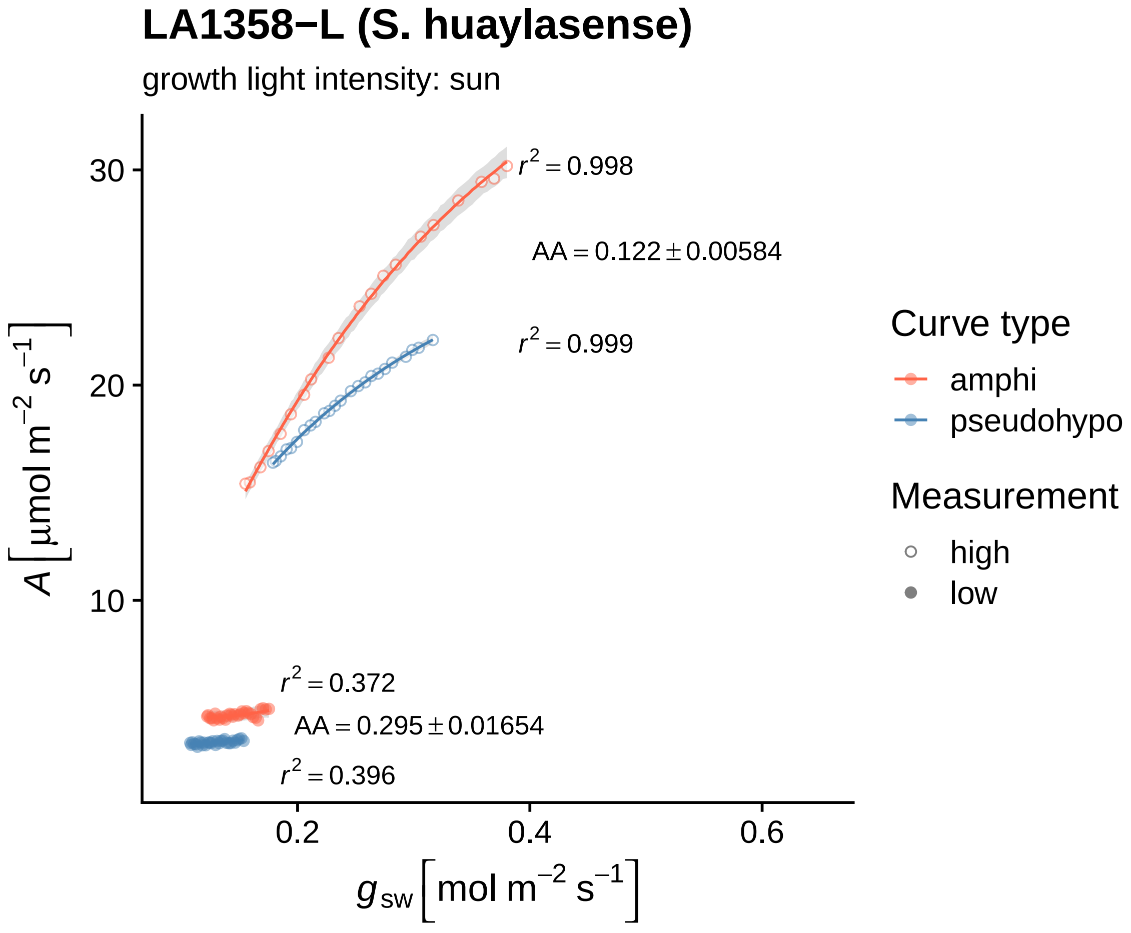

LA1358−L (S. huaylasense)
growth light intensity: sun
$r^2 = 0.998$
AA = 0.122 ± 0.00584
$r^2 = 0.999$
$r^2 = 0.372$
AA = 0.295 ± 0.01654
$r^2 = 0.396$
30
20
10
0.2
0.4
0.6
$A\ [\mu mol\ m^{-2}\ s^{-1}]$
$g_{sw}\ [mol\ m^{-2}\ s^{-1}]$
Curve type
amphi
pseudohypo
Measurement
high
low

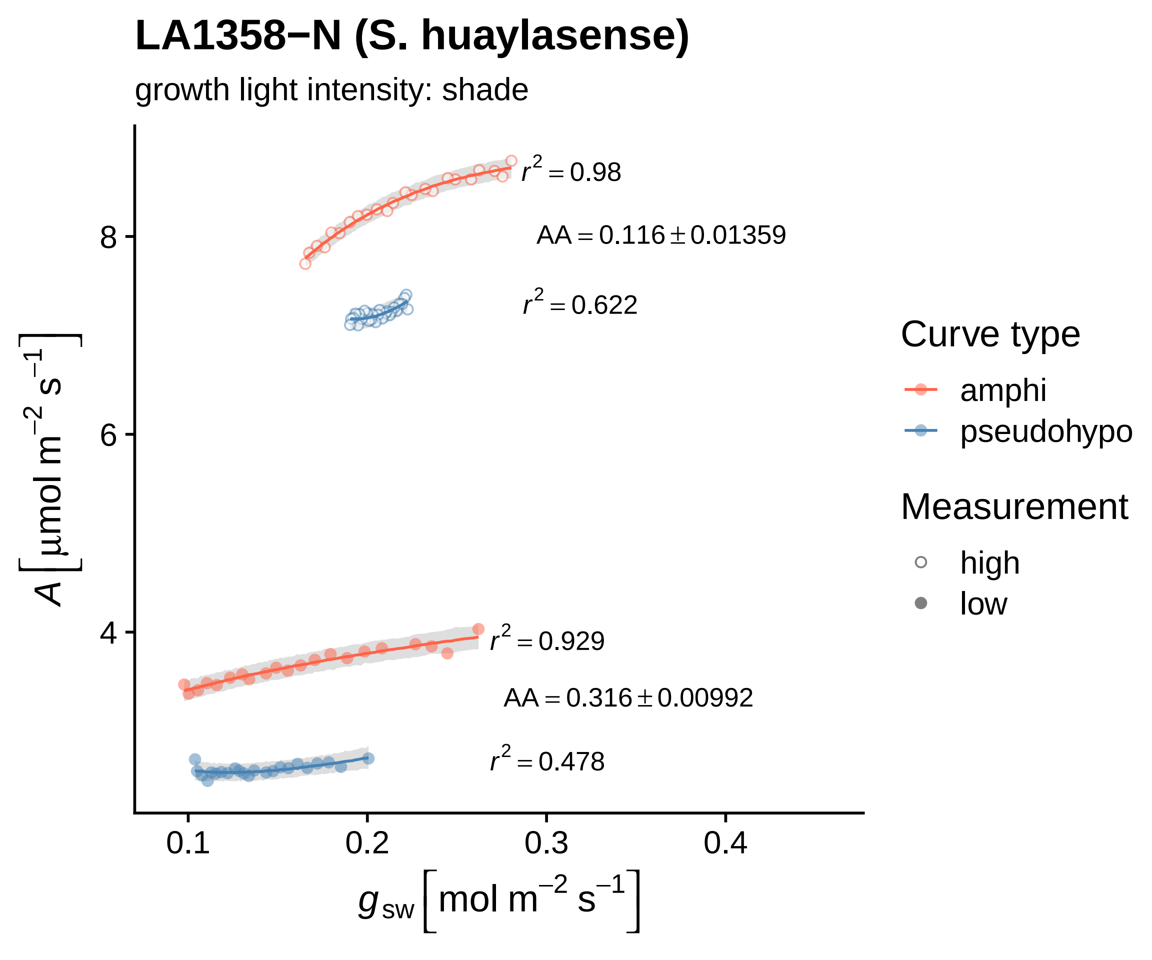
LA1358−N (S. huaylasense)
growth light intensity: shade
$r^2 = 0.98$
$AA = 0.116 \pm 0.01359$
$r^2 = 0.622$
$r^2 = 0.929$
$AA = 0.316 \pm 0.00992$
$r^2 = 0.478$
$A\ [\mu mol\ m^{-2}\ s^{-1}]$
8
6
4
0.1
0.2
0.3
0.4
$g_{sw}\ [mol\ m^{-2}\ s^{-1}]$
Curve type
amphi
pseudohypo
Measurement
high
low

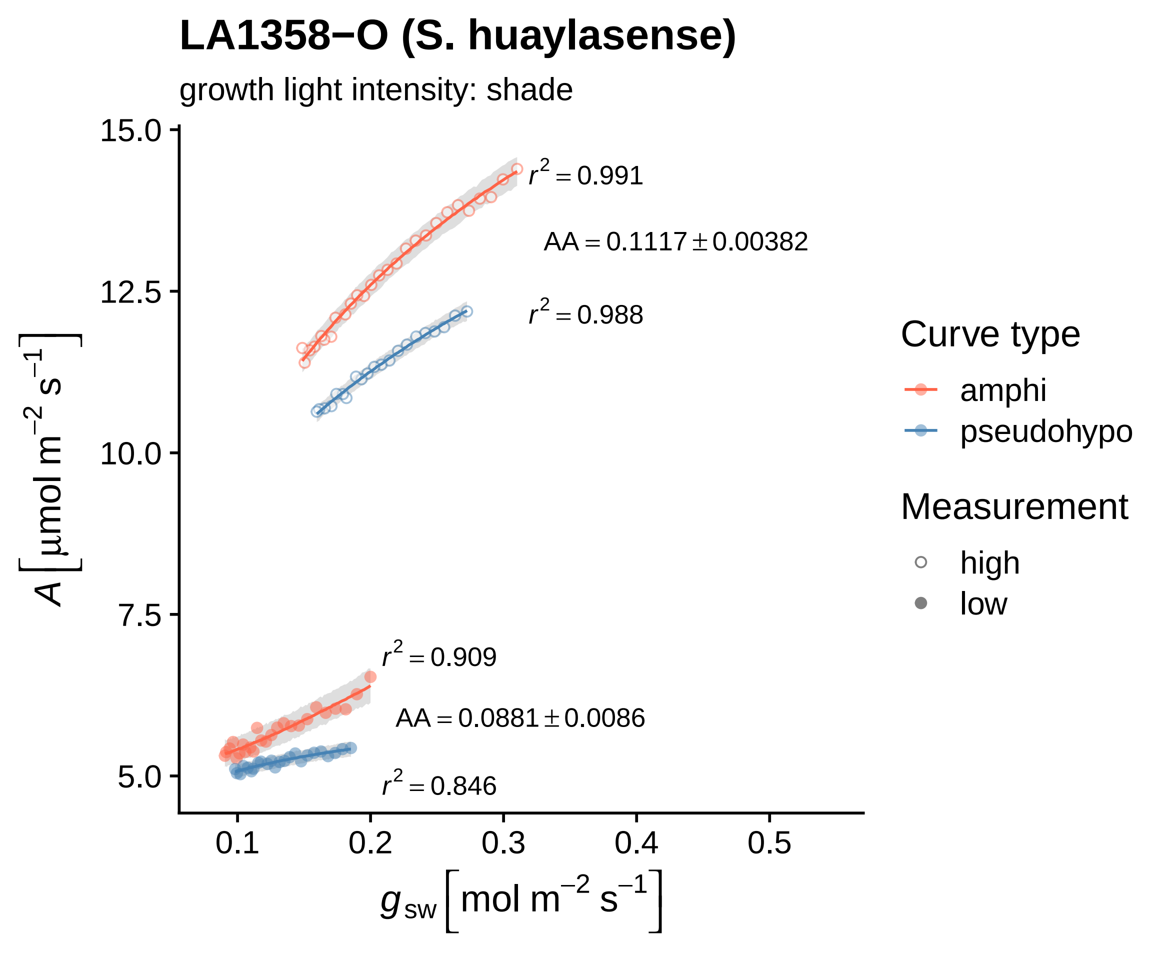

LA1358−O (S. huaylasense)
growth light intensity: shade
15.0
12.5
10.0
7.5
5.0
0.1
0.2
0.3
0.4
0.5
$A\ [\mu mol\ m^{-2}\ s^{-1}]$
$g_{sw}\ [mol\ m^{-2}\ s^{-1}]$
$r^2 = 0.991$
$AA = 0.1117 \pm 0.00382$
$r^2 = 0.988$
$r^2 = 0.909$
$AA = 0.0881 \pm 0.0086$
$r^2 = 0.846$
Curve type
amphi
pseudohypo
Measurement
high
low

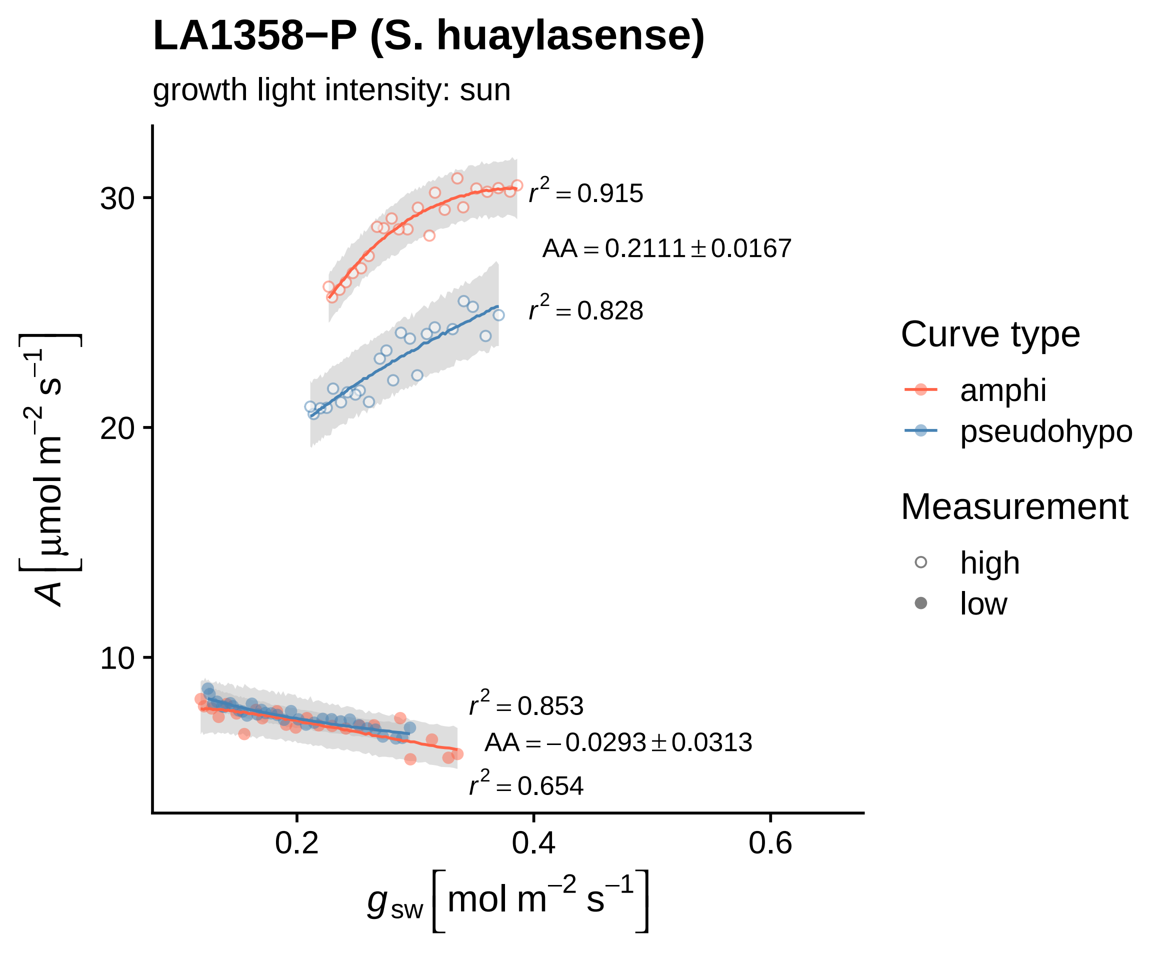

LA1358−P (S. huaylasense)
growth light intensity: sun
$r^2 = 0.915$
$AA = 0.2111 \pm 0.0167$
$r^2 = 0.828$
$r^2 = 0.853$
$AA = -0.0293 \pm 0.0313$
$r^2 = 0.654$
30
20
10
0.2
0.4
0.6
$A\ [\mu mol\ m^{-2}\ s^{-1}]$
$g_{sw}\ [mol\ m^{-2}\ s^{-1}]$
Curve type
amphi
pseudohypo
Measurement
high
low

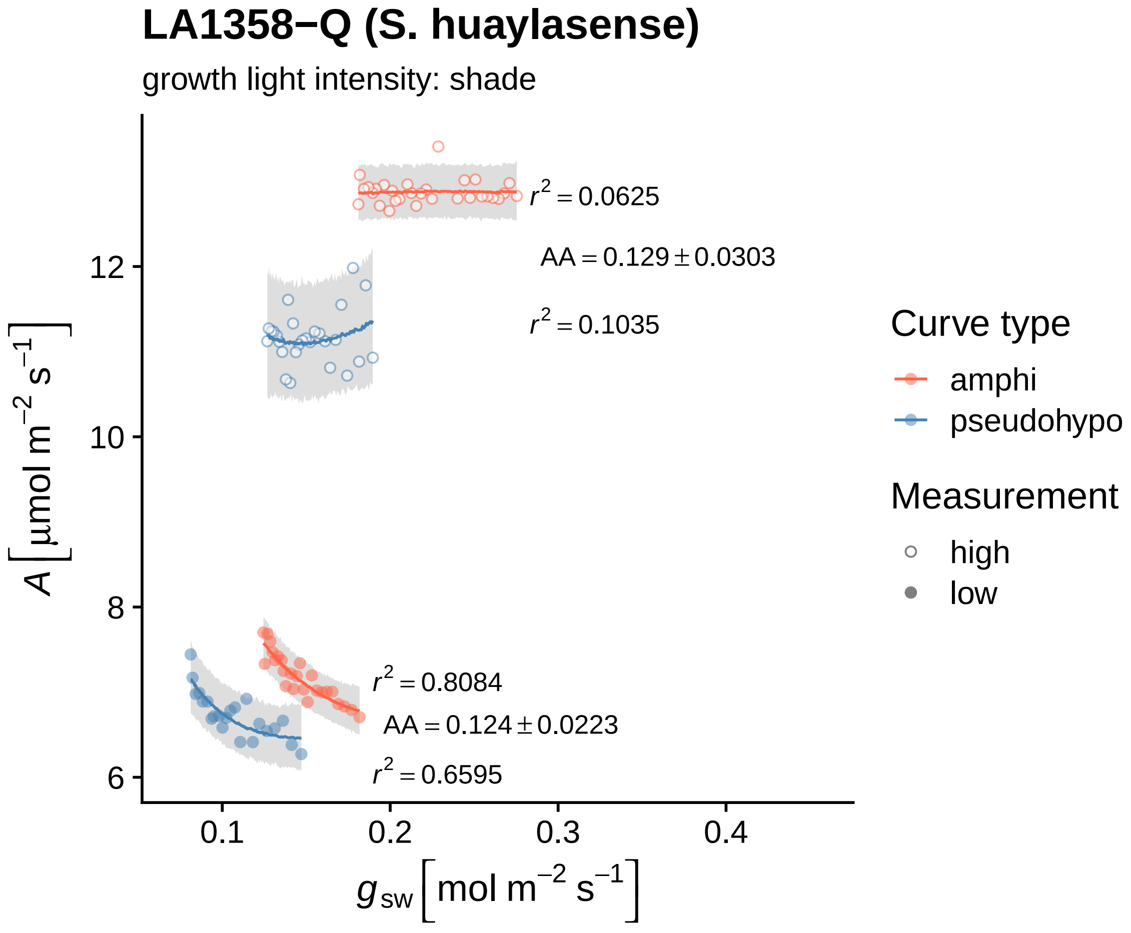

LA1358−Q (S. huaylasense)
growth light intensity: shade
$r^2 = 0.0625$
$AA = 0.129 \pm 0.0303$
$r^2 = 0.1035$
$r^2 = 0.8084$
$AA = 0.124 \pm 0.0223$
$r^2 = 0.6595$
12
10
8
6
0.1
0.2
0.3
0.4
$A\ [\mu mol\ m^{-2}\ s^{-1}]$
$g_{sw}\ [mol\ m^{-2}\ s^{-1}]$
Curve type
amphi
pseudohypo
Measurement
high
low

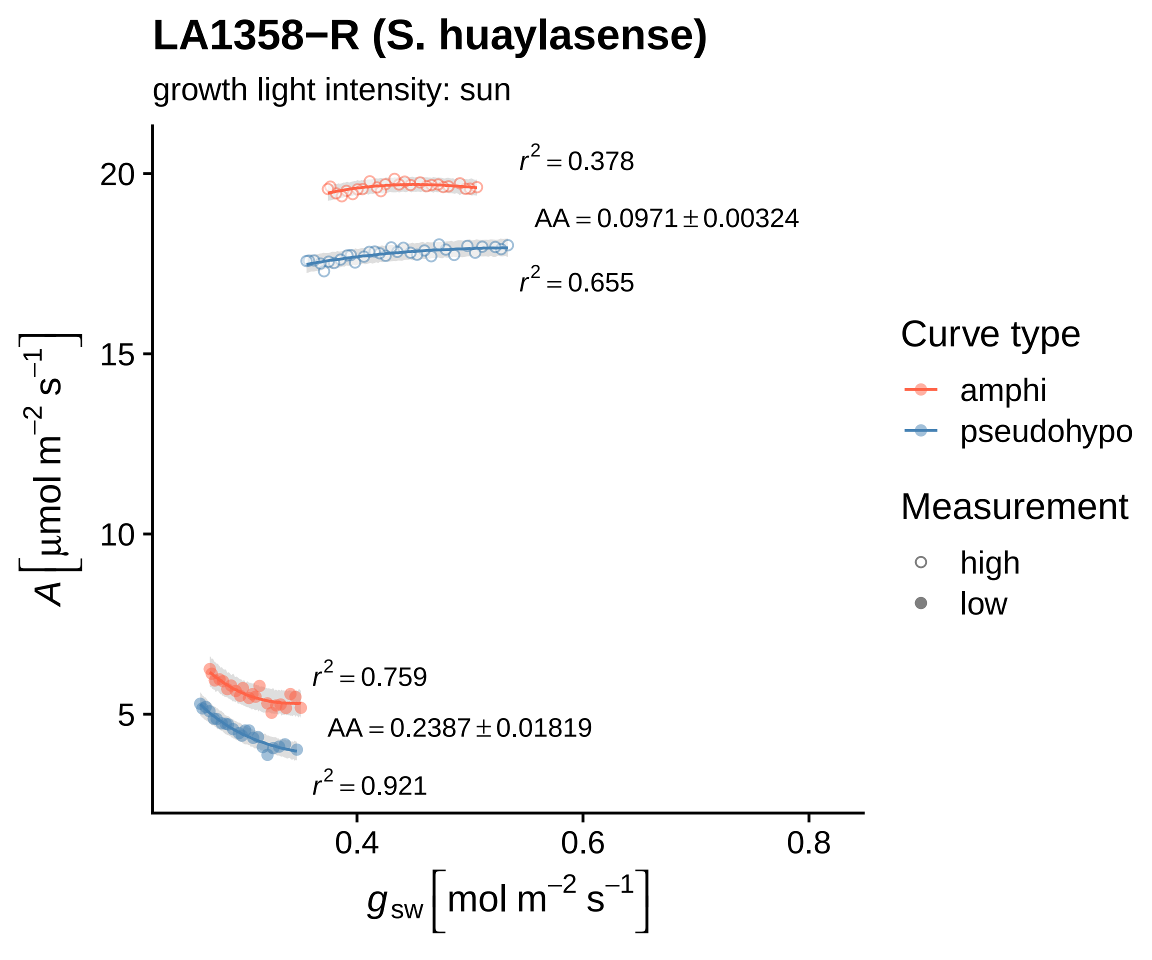

LA1358−R (S. huaylasense)
growth light intensity: sun
r² = 0.378
AA = 0.0971 ± 0.00324
r² = 0.655
r² = 0.759
AA = 0.2387 ± 0.01819
r² = 0.921
A [µmol m⁻² s⁻¹]
g_sw [mol m⁻² s⁻¹]
20
15
10
5
0.4
0.6
0.8
Curve type
amphi
pseudohypo
Measurement
high
low

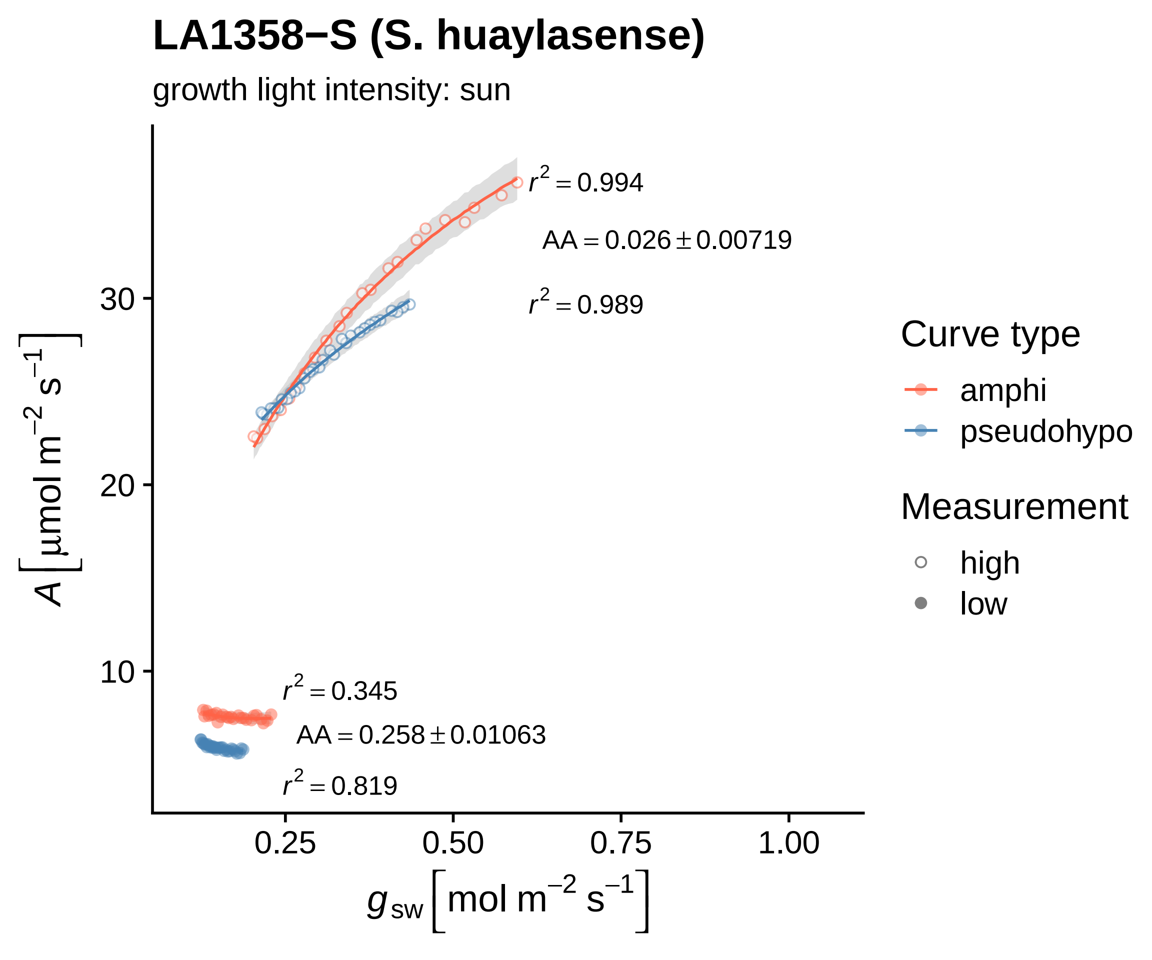

LA1358−S (S. huaylasense)
growth light intensity: sun
$r^2 = 0.994$
$AA = 0.026 \pm 0.00719$
$r^2 = 0.989$
$r^2 = 0.345$
$AA = 0.258 \pm 0.01063$
$r^2 = 0.819$
30
20
10
0.25
0.50
0.75
1.00
$A\ [\mu mol\ m^{-2}\ s^{-1}]$
$g_{sw}\ [mol\ m^{-2}\ s^{-1}]$
Curve type
amphi
pseudohypo
Measurement
high
low

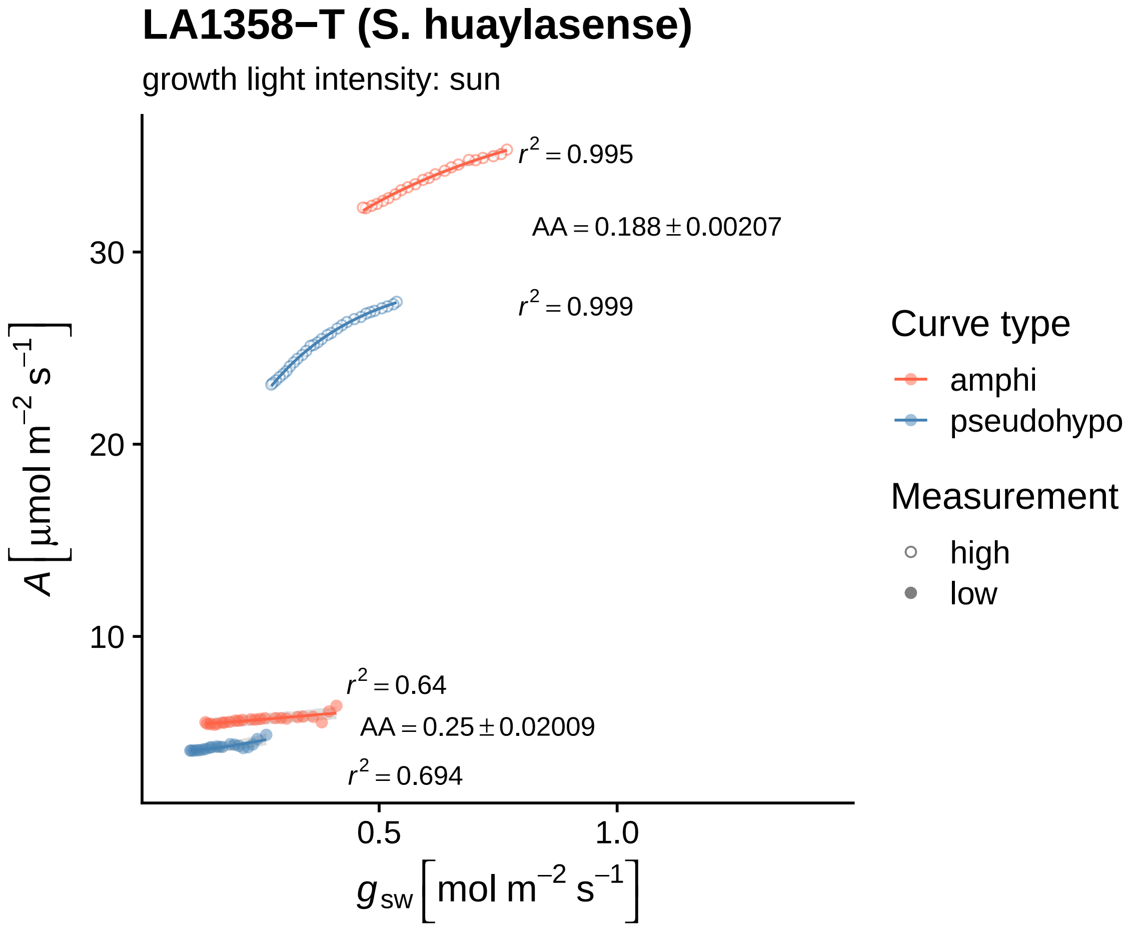
LA1358−T (S. huaylasense)
growth light intensity: sun
$r^2 = 0.995$
$AA = 0.188 \pm 0.00207$
$r^2 = 0.999$
$r^2 = 0.64$
$AA = 0.25 \pm 0.02009$
$r^2 = 0.694$
30
20
10
0.5
1.0
$A\,[\mu mol\,m^{-2}\,s^{-1}]$
$g_{sw}\,[mol\,m^{-2}\,s^{-1}]$
Curve type
amphi
pseudohypo
Measurement
high
low

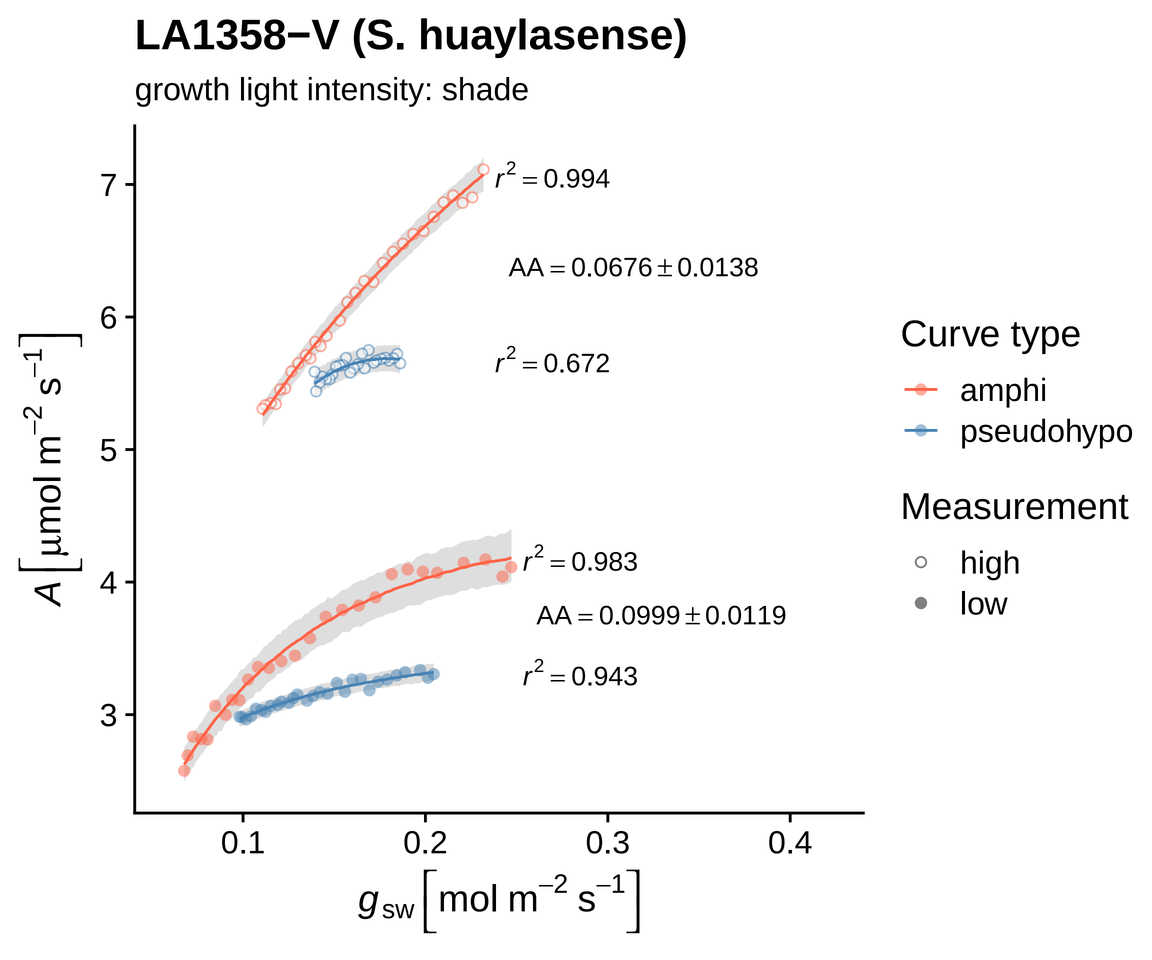

LA1358−V (S. huaylasense)
growth light intensity: shade
$r^2 = 0.994$
$AA = 0.0676 \pm 0.0138$
$r^2 = 0.672$
$r^2 = 0.983$
$AA = 0.0999 \pm 0.0119$
$r^2 = 0.943$
7
6
5
4
3
0.1
0.2
0.3
0.4
$A\ [\mu mol\ m^{-2}\ s^{-1}]$
$g_{sw}\ [mol\ m^{-2}\ s^{-1}]$
Curve type
amphi
pseudohypo
Measurement
high
low

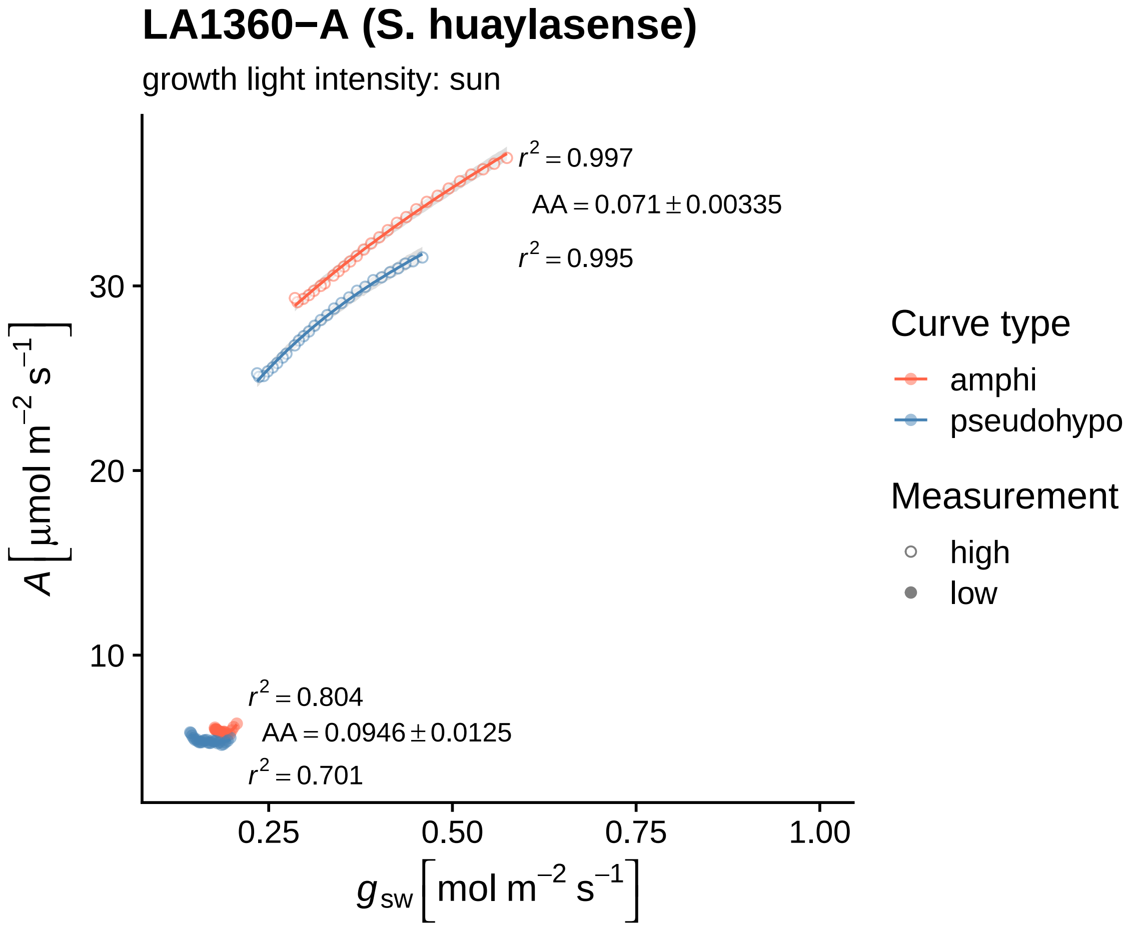

LA1360−A (S. huaylasense)
growth light intensity: sun
$r^2 = 0.997$
$AA = 0.071 \pm 0.00335$
$r^2 = 0.995$
$r^2 = 0.804$
$AA = 0.0946 \pm 0.0125$
$r^2 = 0.701$
30
20
10
0.25
0.50
0.75
1.00
$A\left[\mu mol\ m^{-2}\ s^{-1}\right]$
$g_{sw}\left[mol\ m^{-2}\ s^{-1}\right]$
Curve type
amphi
pseudohypo
Measurement
high
low

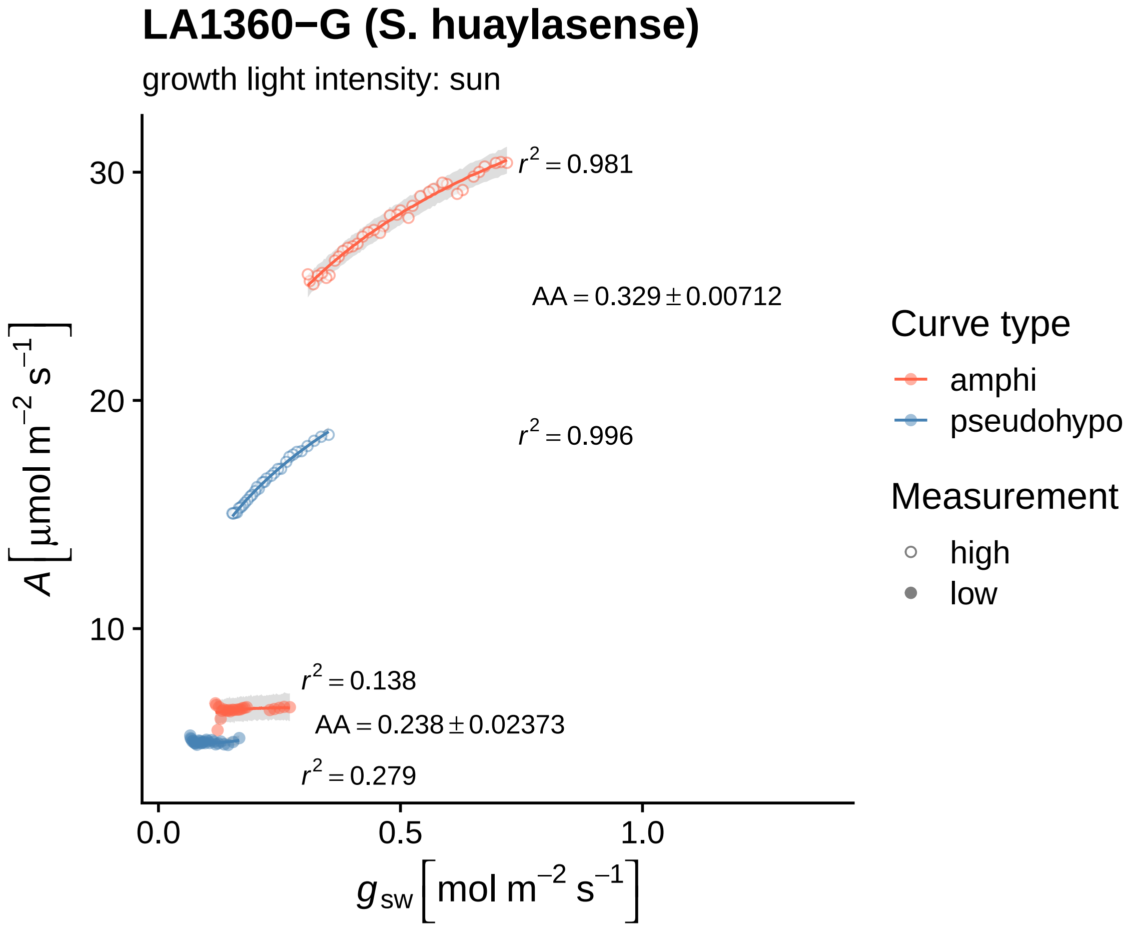
LA1360−G (S. huaylasense)
growth light intensity: sun
A [μmol m−2 s−1]
30
20
10
0.0
0.5
1.0
gsw [mol m−2 s−1]
r2 = 0.981
AA = 0.329 ± 0.00712
r2 = 0.996
r2 = 0.138
AA = 0.238 ± 0.02373
r2 = 0.279
Curve type
amphi
pseudohypo
Measurement
high
low

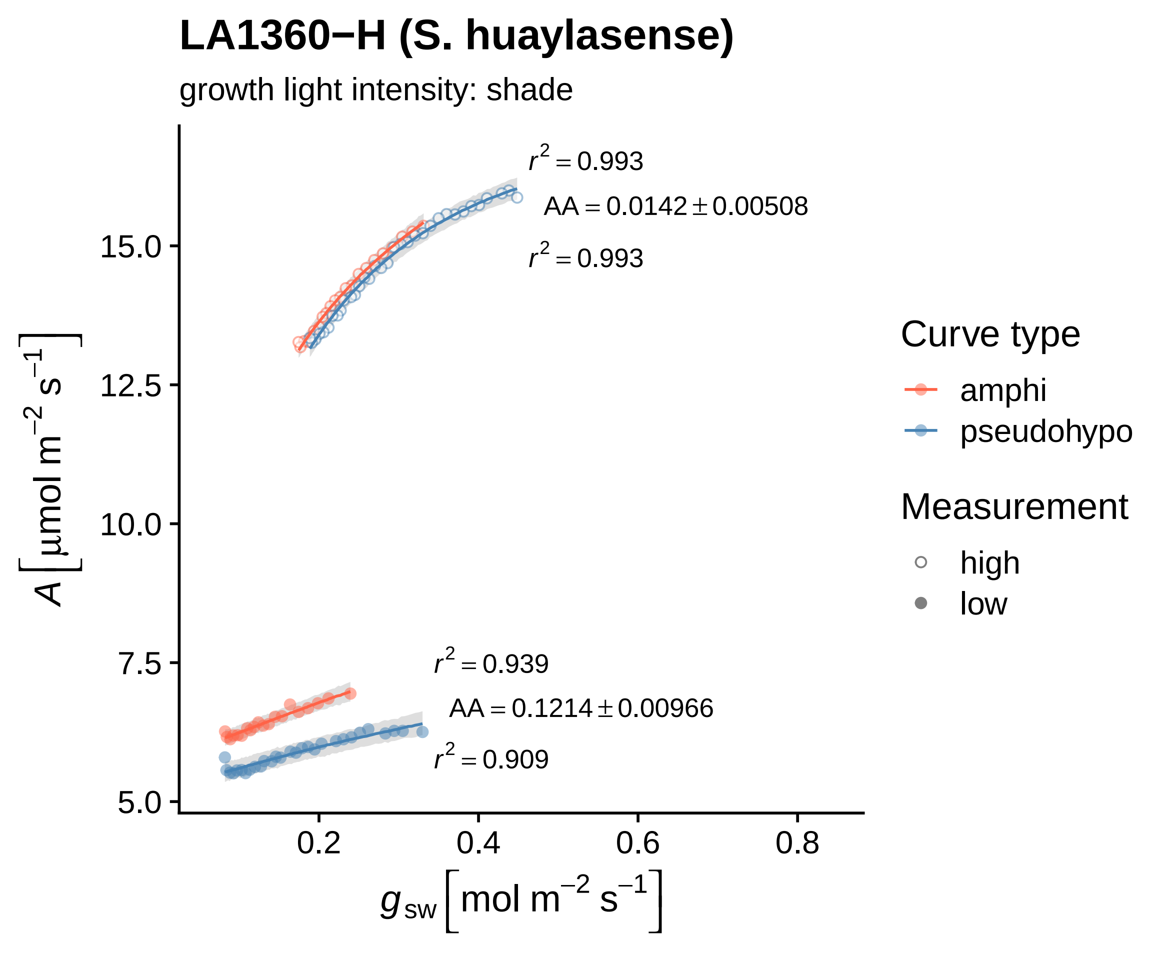
LA1360−H (S. huaylasense)
growth light intensity: shade
$r^2 = 0.993$
$AA = 0.0142 \pm 0.00508$
$r^2 = 0.993$
$r^2 = 0.939$
$AA = 0.1214 \pm 0.00966$
$r^2 = 0.909$
15.0
12.5
10.0
7.5
5.0
0.2
0.4
0.6
0.8
$A\left[\mu mol\ m^{-2}\ s^{-1}\right]$
$g_{sw}\left[mol\ m^{-2}\ s^{-1}\right]$
Curve type
amphi
pseudohypo
Measurement
high
low

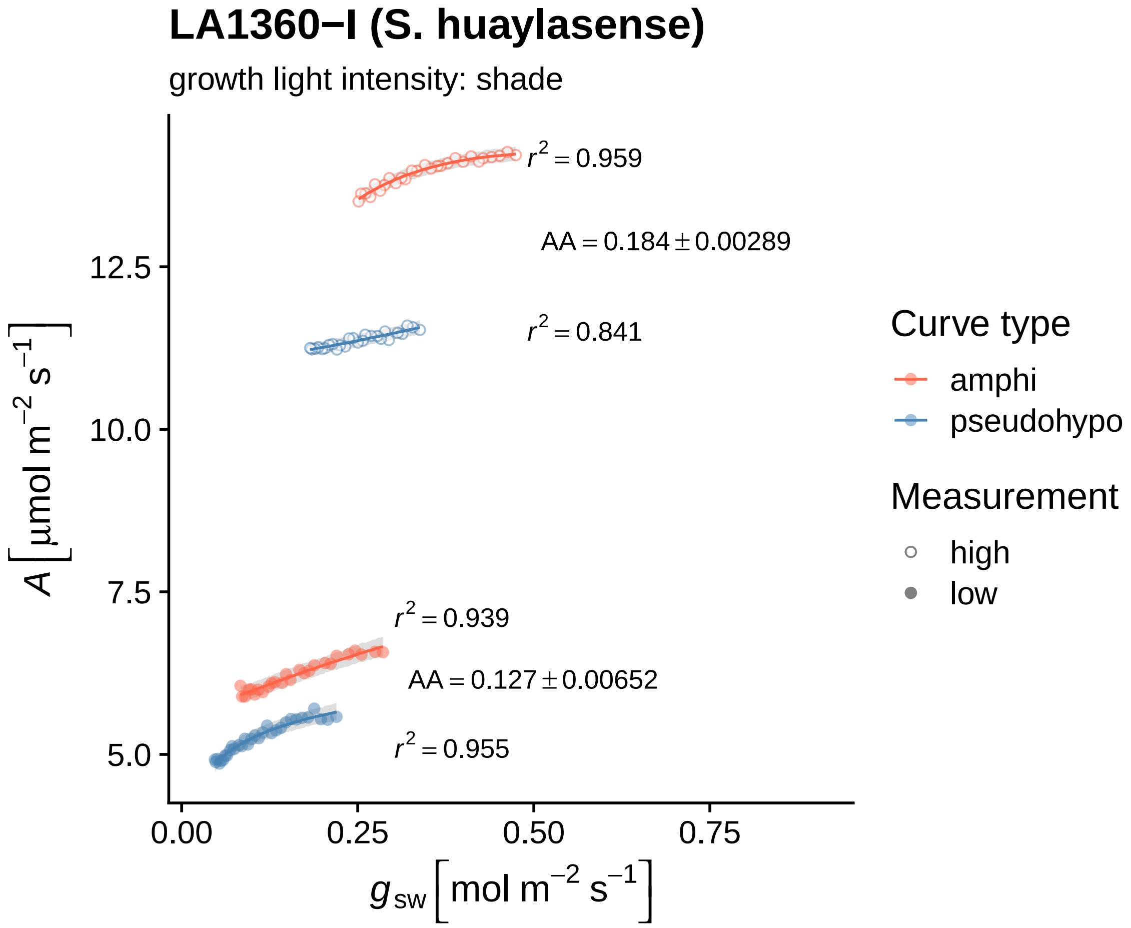

LA1360−I (S. huaylasense)
growth light intensity: shade
$r^2 = 0.959$
$AA = 0.184 \pm 0.00289$
$r^2 = 0.841$
$r^2 = 0.939$
$AA = 0.127 \pm 0.00652$
$r^2 = 0.955$
12.5
10.0
7.5
5.0
0.00
0.25
0.50
0.75
$A\ [\mu mol\ m^{-2}\ s^{-1}]$
$g_{sw}\ [mol\ m^{-2}\ s^{-1}]$
Curve type
amphi
pseudohypo
Measurement
high
low

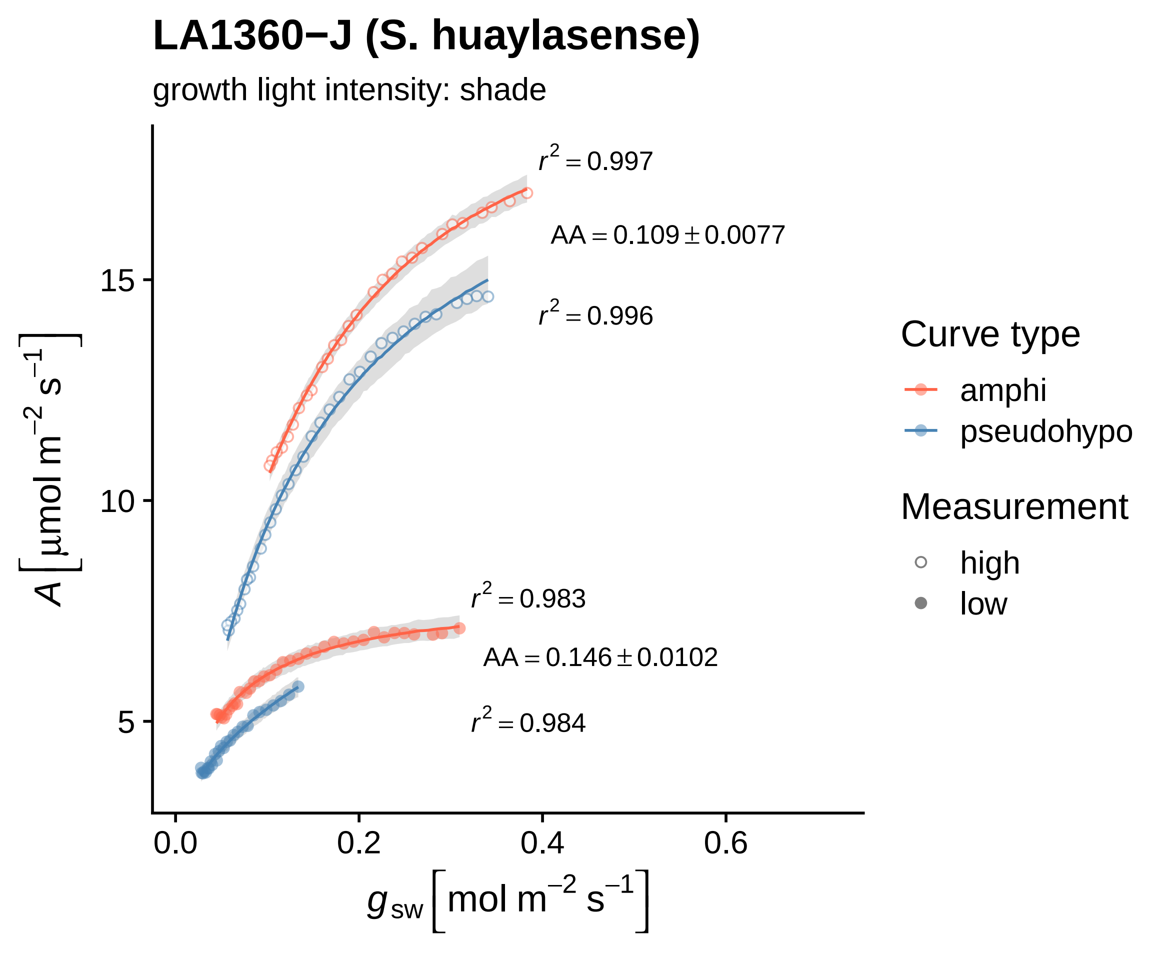

LA1360−J (S. huaylasense)
growth light intensity: shade
$r^2 = 0.997$
$AA = 0.109 \pm 0.0077$
$r^2 = 0.996$
$r^2 = 0.983$
$AA = 0.146 \pm 0.0102$
$r^2 = 0.984$
15
10
5
0.0
0.2
0.4
0.6
$A\ [\mu mol\ m^{-2}\ s^{-1}]$
$g_{sw}\ [mol\ m^{-2}\ s^{-1}]$
Curve type
amphi
pseudohypo
Measurement
high
low

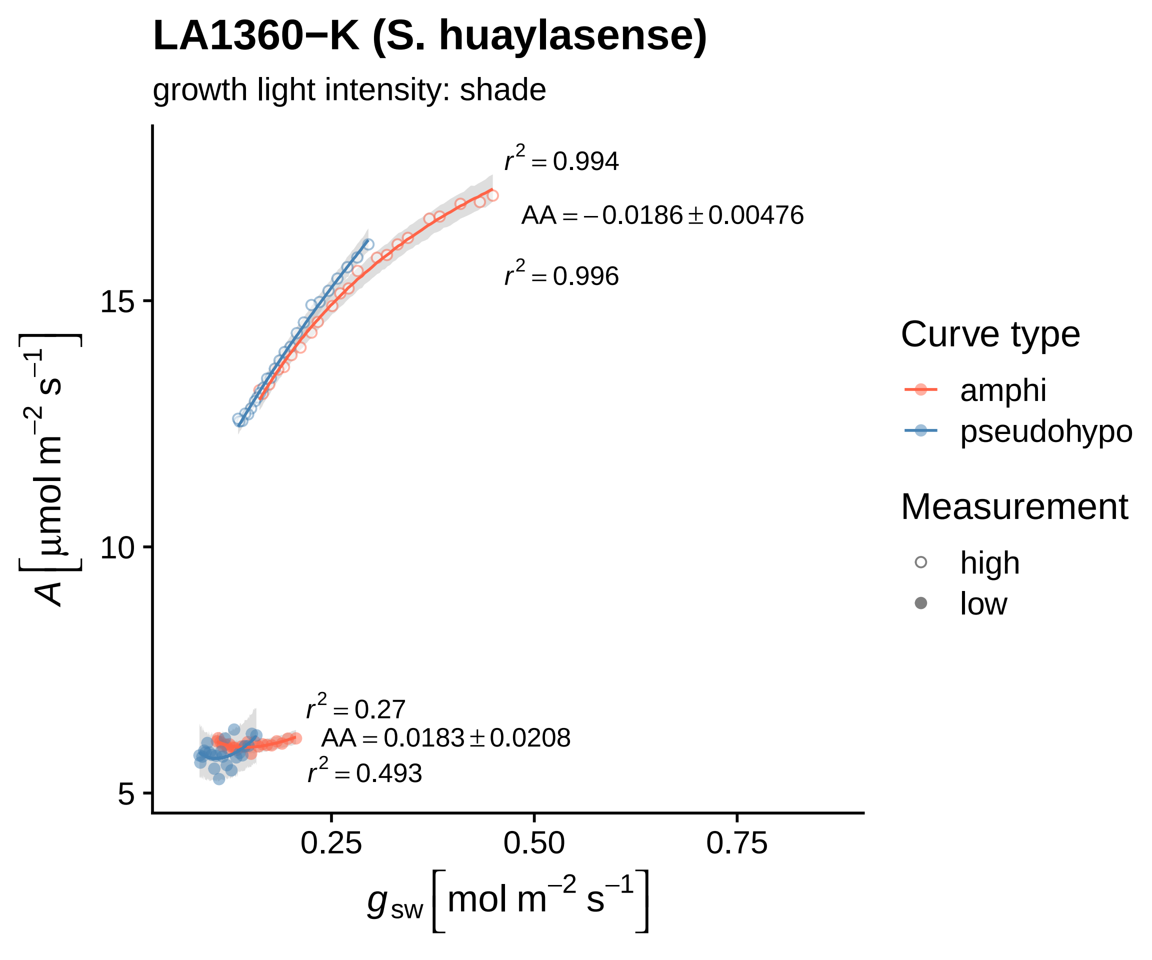

LA1360−K (S. huaylasense)
growth light intensity: shade
$r^2 = 0.994$
$AA = -0.0186 \pm 0.00476$
$r^2 = 0.996$
$r^2 = 0.27$
$AA = 0.0183 \pm 0.0208$
$r^2 = 0.493$
15
10
5
0.25
0.50
0.75
$A\left[\mu mol\ m^{-2}\ s^{-1}\right]$
$g_{sw}\left[mol\ m^{-2}\ s^{-1}\right]$
Curve type
amphi
pseudohypo
Measurement
high
low

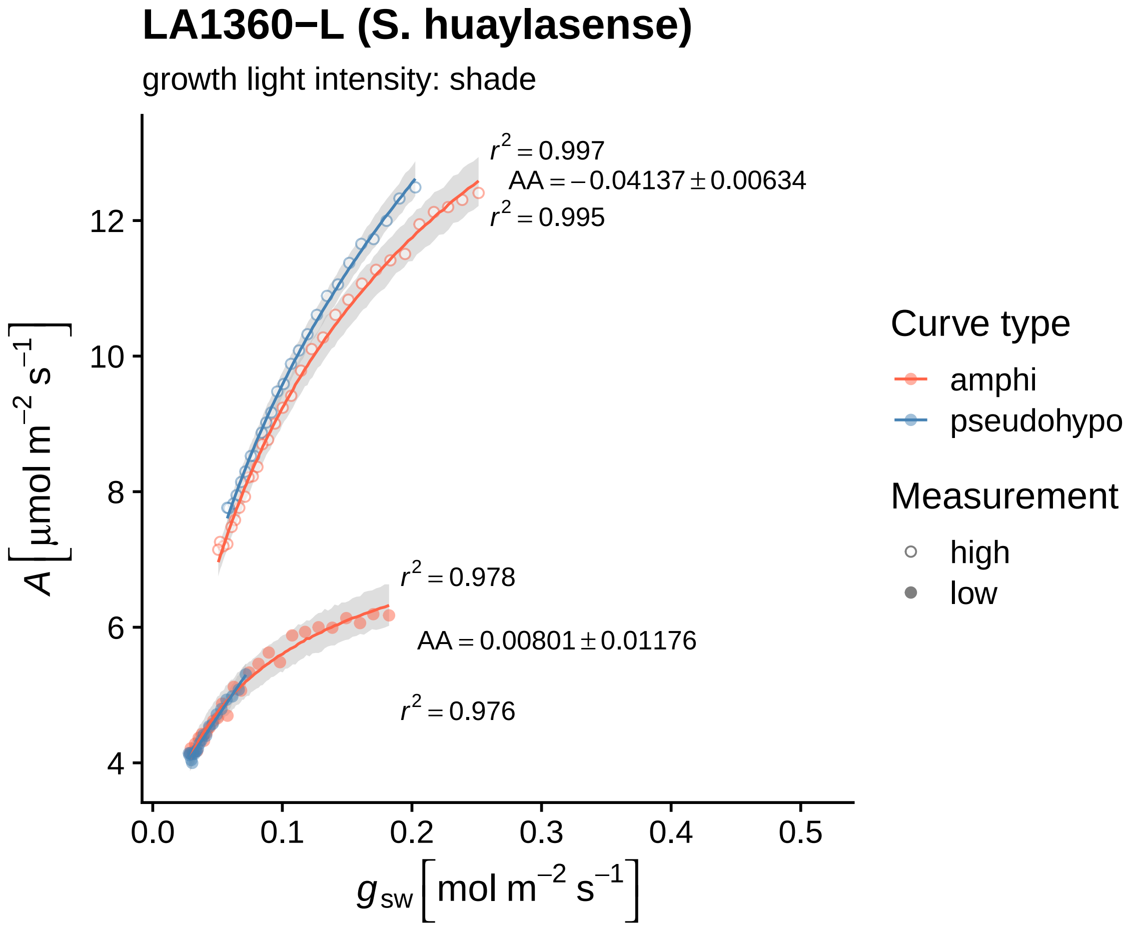

LA1360−L (S. huaylasense)
growth light intensity: shade
$r^2 = 0.997$
$AA = -0.04137 \pm 0.00634$
$r^2 = 0.995$
$r^2 = 0.978$
$AA = 0.00801 \pm 0.01176$
$r^2 = 0.976$
12
10
8
6
4
$A\left[\mu mol\ m^{-2}\ s^{-1}\right]$
0.0
0.1
0.2
0.3
0.4
0.5
$g_{sw}\left[mol\ m^{-2}\ s^{-1}\right]$
Curve type
amphi
pseudohypo
Measurement
high
low

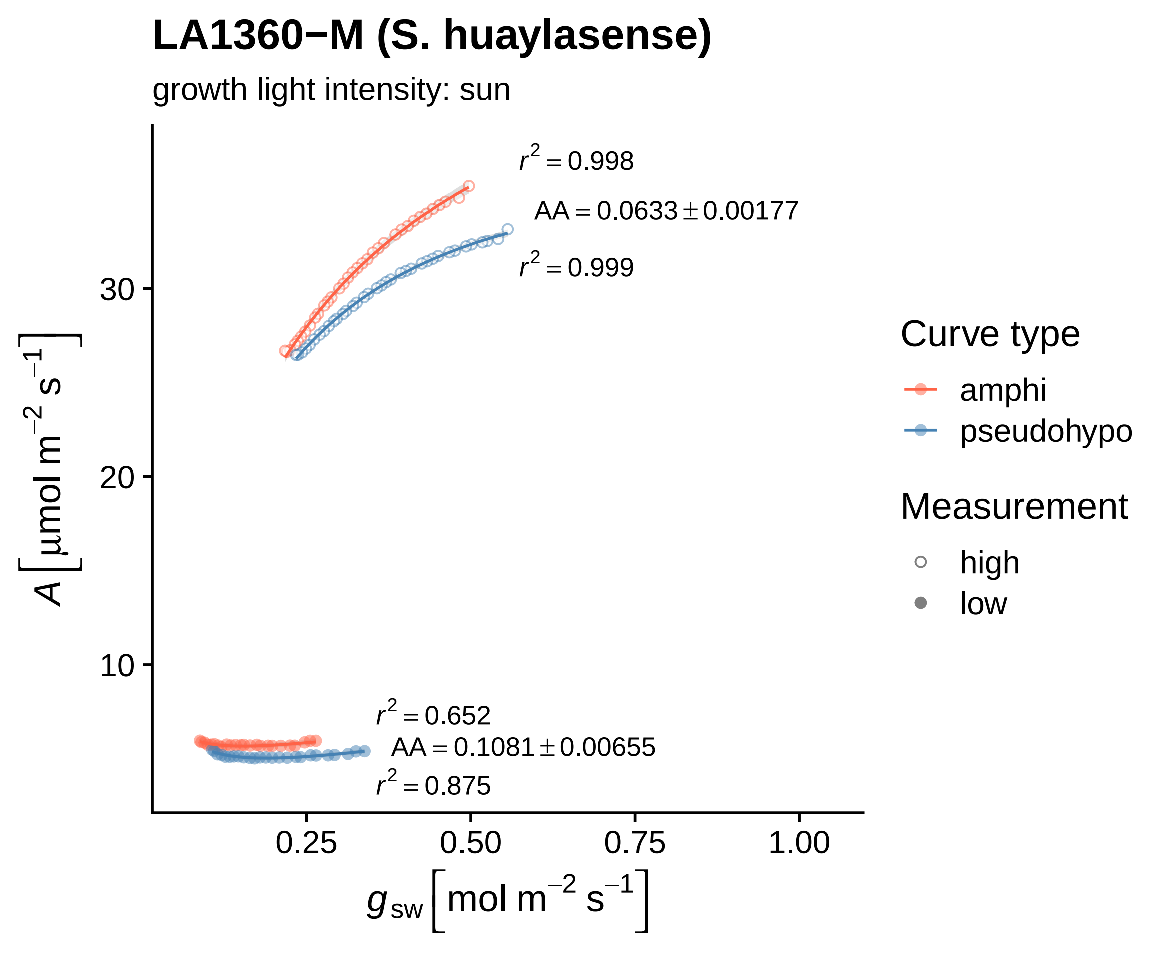
LA1360−M (S. huaylasense)
growth light intensity: sun
r² = 0.998
AA = 0.0633 ± 0.00177
r² = 0.999
r² = 0.652
AA = 0.1081 ± 0.00655
r² = 0.875
30
20
10
0.25
0.50
0.75
1.00
A [µmol m⁻² s⁻¹]
g_sw [mol m⁻² s⁻¹]
Curve type
amphi
pseudohypo
Measurement
high
low

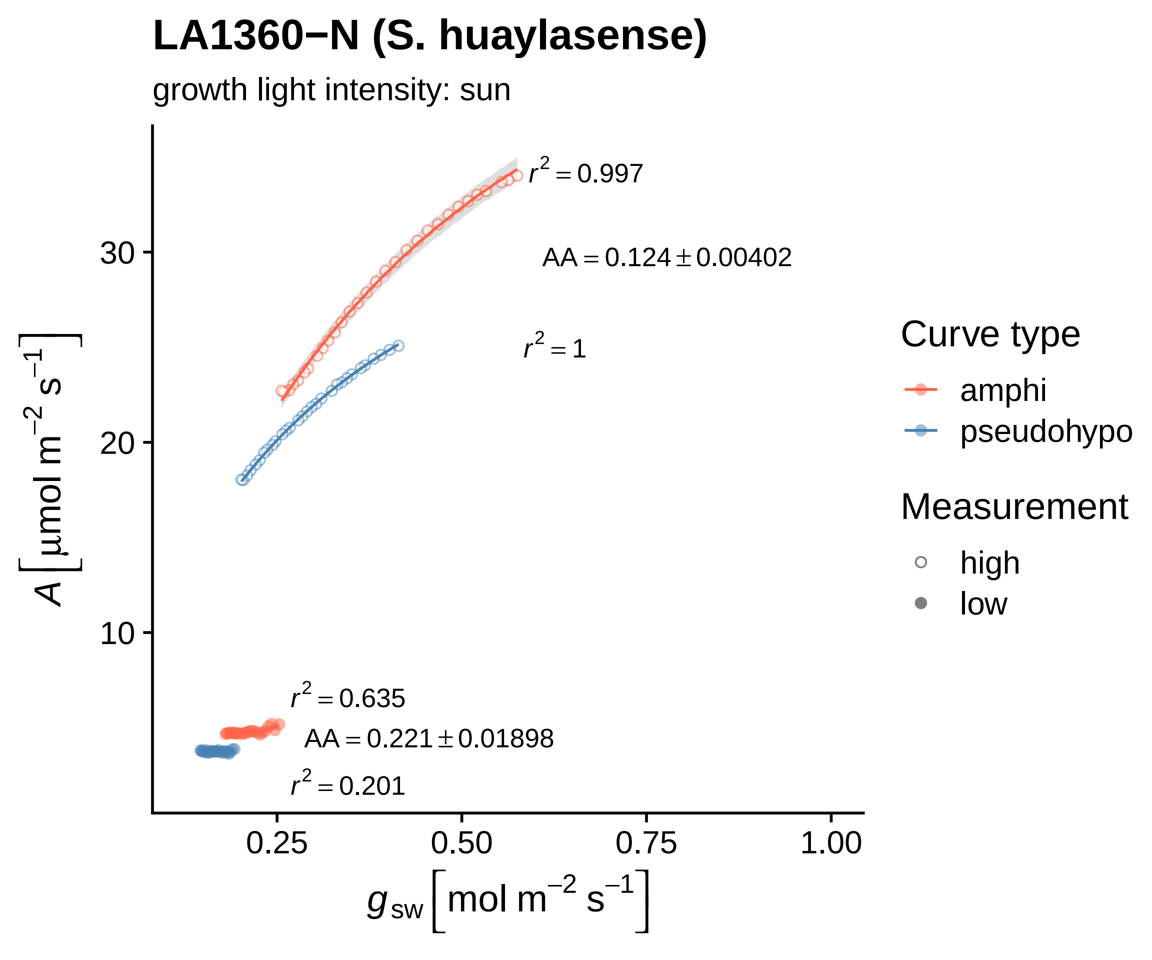

LA1360−N (S. huaylasense)
growth light intensity: sun
r² = 0.997
AA = 0.124 ± 0.00402
r² = 1
r² = 0.635
AA = 0.221 ± 0.01898
r² = 0.201
30
20
10
0.25
0.50
0.75
1.00
A [µmol m⁻² s⁻¹]
g_sw [mol m⁻² s⁻¹]
Curve type
amphi
pseudohypo
Measurement
high
low

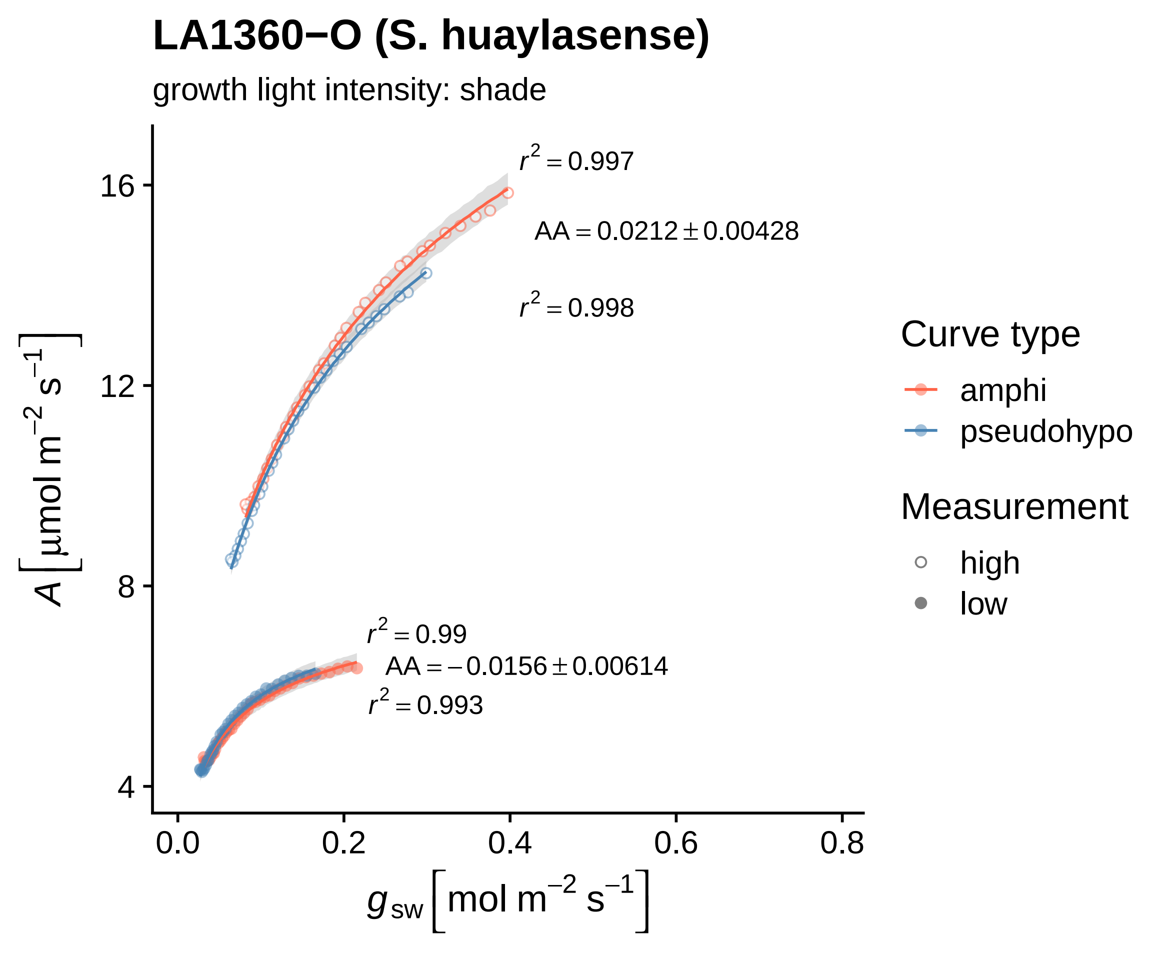
LA1360−O (S. huaylasense)
growth light intensity: shade
r² = 0.997
AA = 0.0212 ± 0.00428
r² = 0.998
r² = 0.99
AA = −0.0156 ± 0.00614
r² = 0.993
A [µmol m⁻² s⁻¹]
g_sw [mol m⁻² s⁻¹]
16
12
8
4
0.0
0.2
0.4
0.6
0.8
Curve type
amphi
pseudohypo
Measurement
high
low

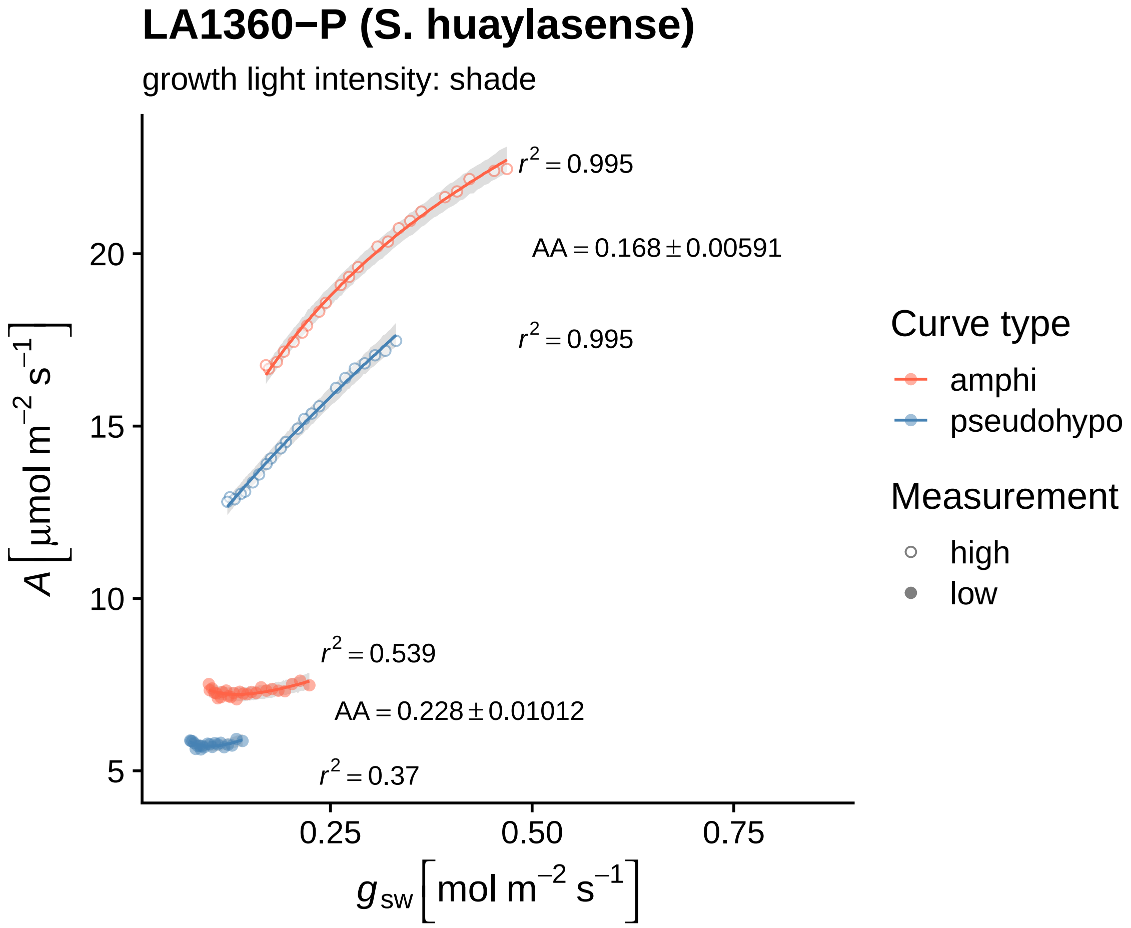

LA1360−P (S. huaylasense)
growth light intensity: shade
$r^2 = 0.995$
AA = 0.168 ± 0.00591
$r^2 = 0.995$
$r^2 = 0.539$
AA = 0.228 ± 0.01012
$r^2 = 0.37$
20
15
10
5
0.25
0.50
0.75
$A\ [\mu mol\ m^{-2}\ s^{-1}]$
$g_{sw}\ [mol\ m^{-2}\ s^{-1}]$
Curve type
amphi
pseudohypo
Measurement
high
low

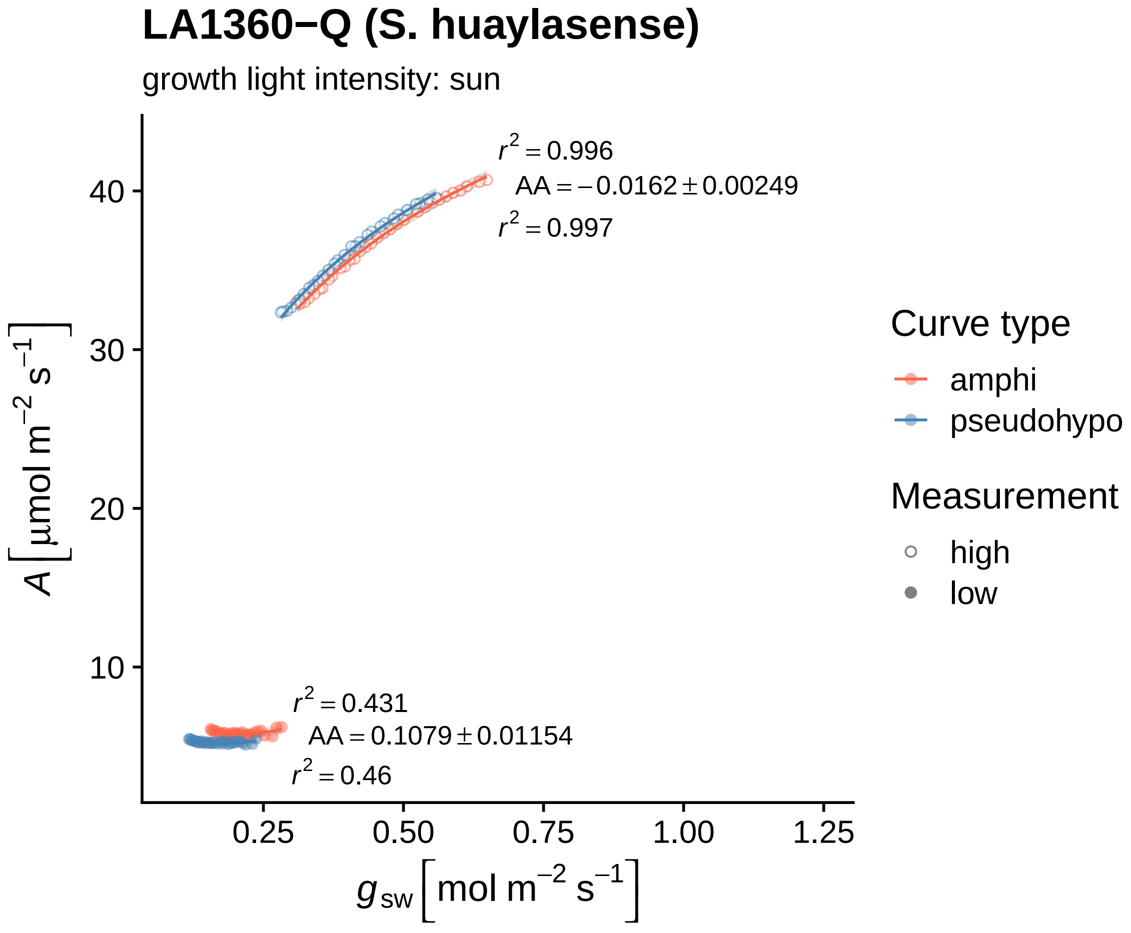

LA1360−Q (S. huaylasense)
growth light intensity: sun
$r^2 = 0.996$
$AA = -0.0162 \pm 0.00249$
$r^2 = 0.997$
$r^2 = 0.431$
$AA = 0.1079 \pm 0.01154$
$r^2 = 0.46$
$A\left[\mu mol\ m^{-2}\ s^{-1}\right]$
40
30
20
10
0.25
0.50
0.75
1.00
1.25
$g_{sw}\left[mol\ m^{-2}\ s^{-1}\right]$
Curve type
amphi
pseudohypo
Measurement
high
low

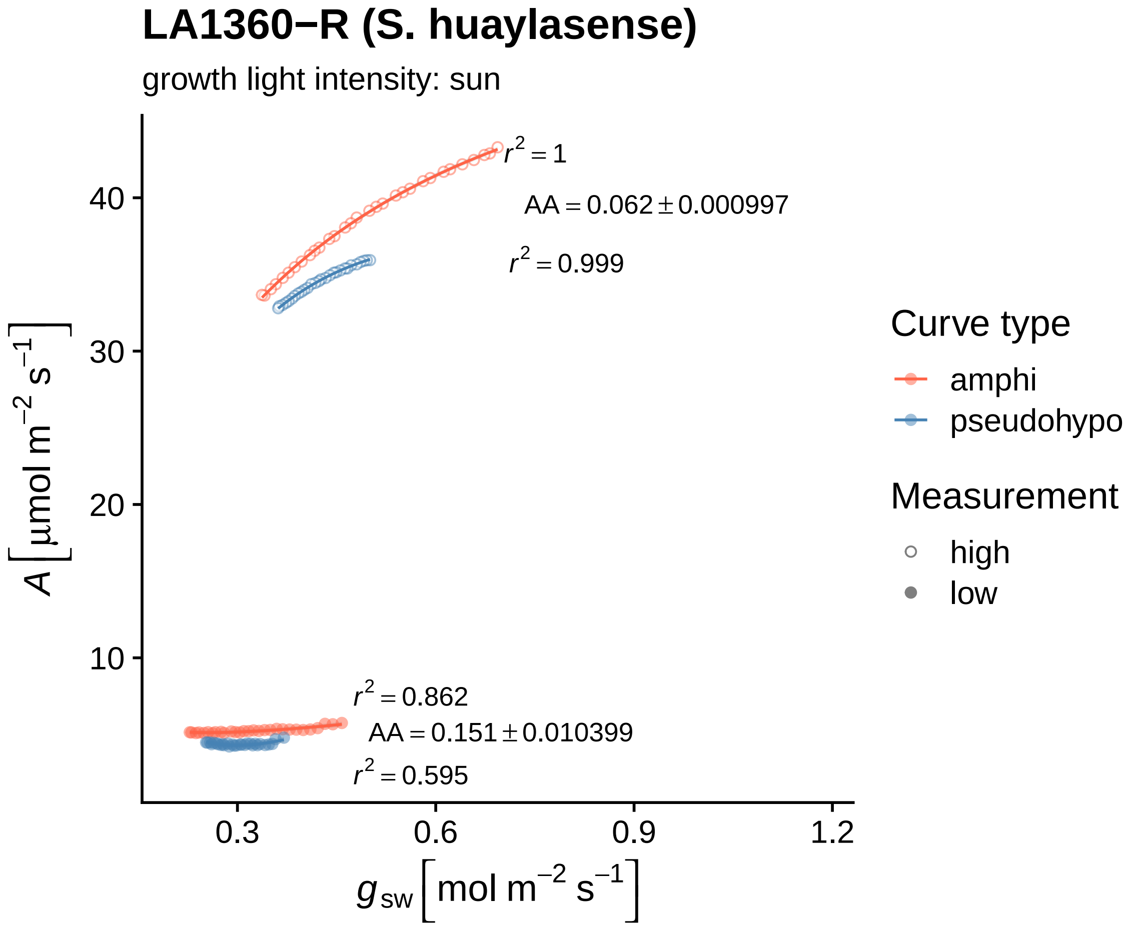

LA1360−R (S. huaylasense)
growth light intensity: sun
r² = 1
AA = 0.062 ± 0.000997
r² = 0.999
r² = 0.862
AA = 0.151 ± 0.010399
r² = 0.595
A [μmol m⁻² s⁻¹]
g_sw [mol m⁻² s⁻¹]
10
20
30
40
0.3
0.6
0.9
1.2
Curve type
amphi
pseudohypo
Measurement
high
low

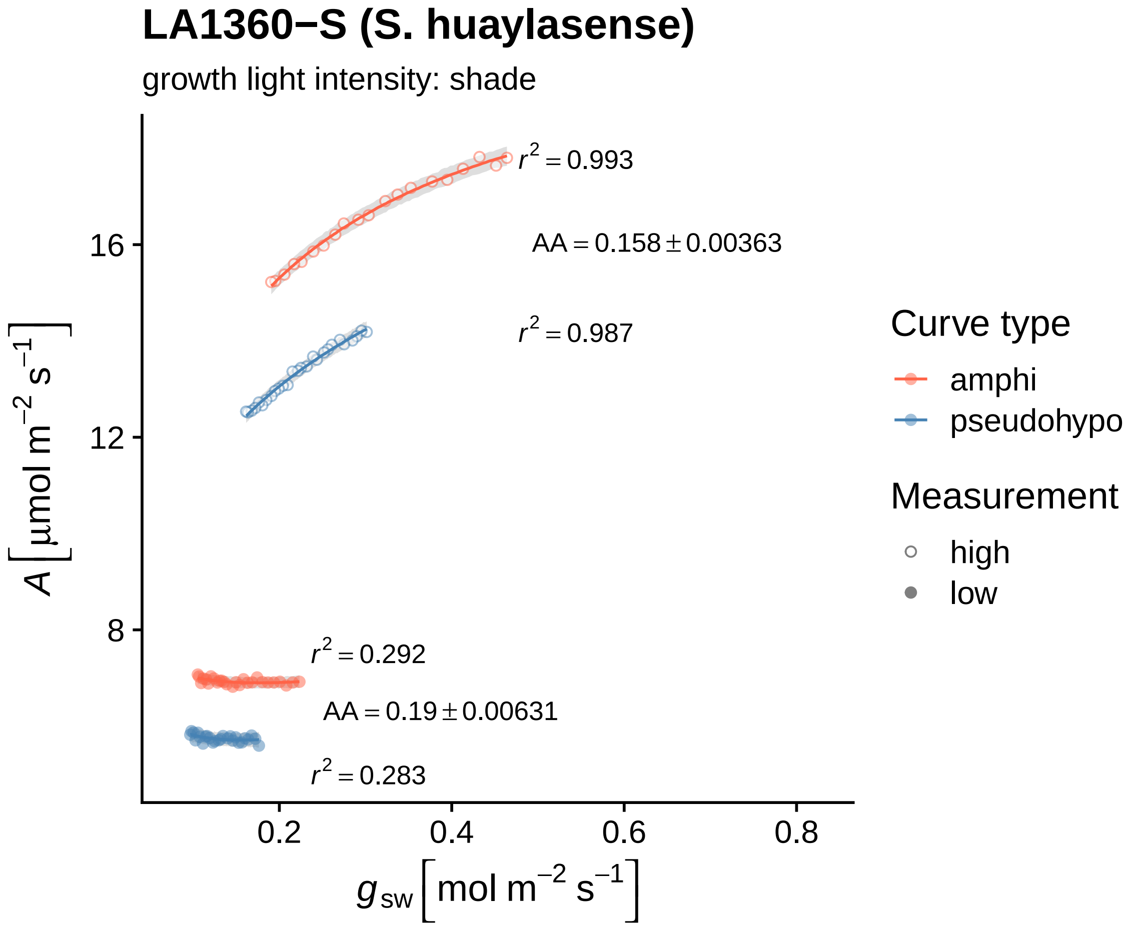
LA1360−S (S. huaylasense)
growth light intensity: shade
$r^2 = 0.993$
$AA = 0.158 \pm 0.00363$
$r^2 = 0.987$
$r^2 = 0.292$
$AA = 0.19 \pm 0.00631$
$r^2 = 0.283$
16
12
8
0.2
0.4
0.6
0.8
$A\ [\mu mol\ m^{-2}\ s^{-1}]$
$g_{sw}\ [mol\ m^{-2}\ s^{-1}]$
Curve type
amphi
pseudohypo
Measurement
high
low

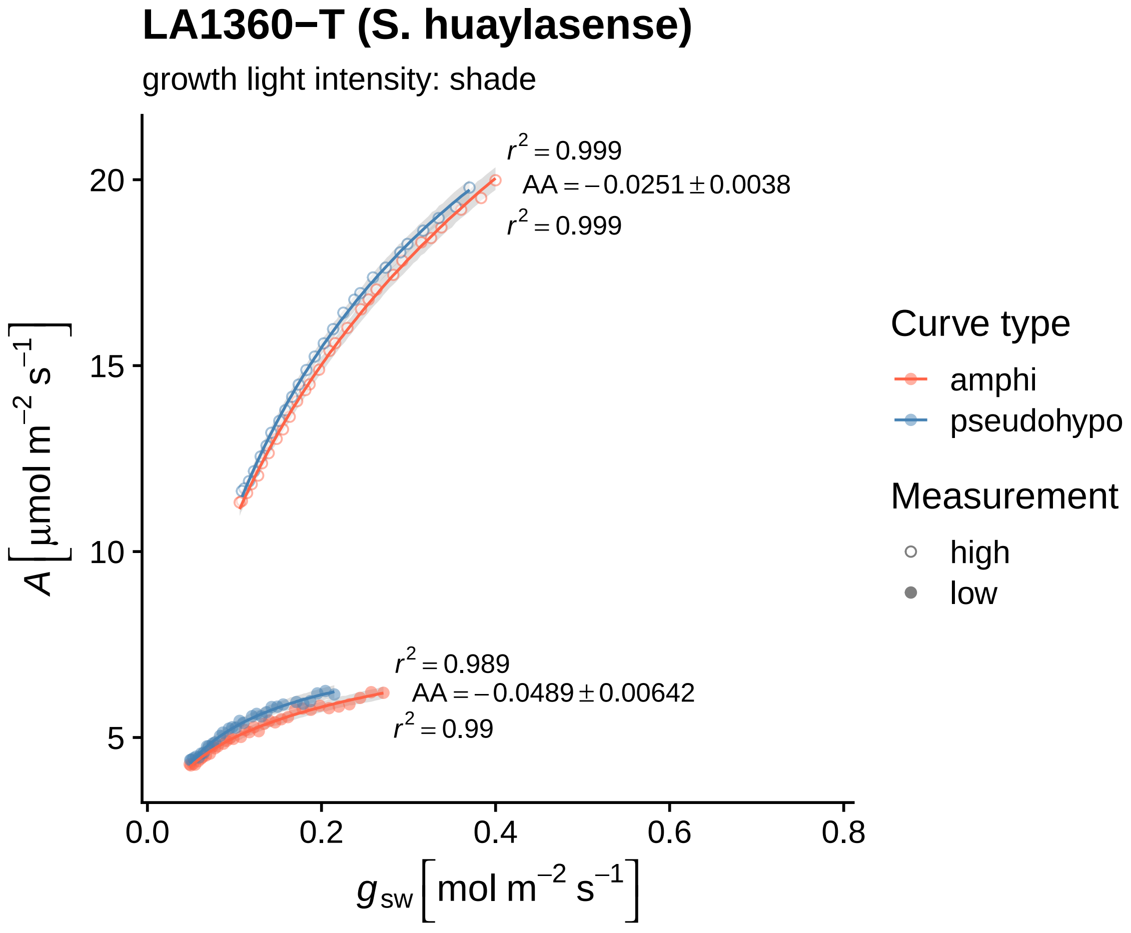
LA1360−T (S. huaylasense)
growth light intensity: shade
$r^2 = 0.999$
$AA = -0.0251 \pm 0.0038$
$r^2 = 0.999$
$r^2 = 0.989$
$AA = -0.0489 \pm 0.00642$
$r^2 = 0.99$
20
15
10
5
0.0
0.2
0.4
0.6
0.8
$A\ [\mu mol\ m^{-2}\ s^{-1}]$
$g_{sw}\ [mol\ m^{-2}\ s^{-1}]$
Curve type
amphi
pseudohypo
Measurement
high
low

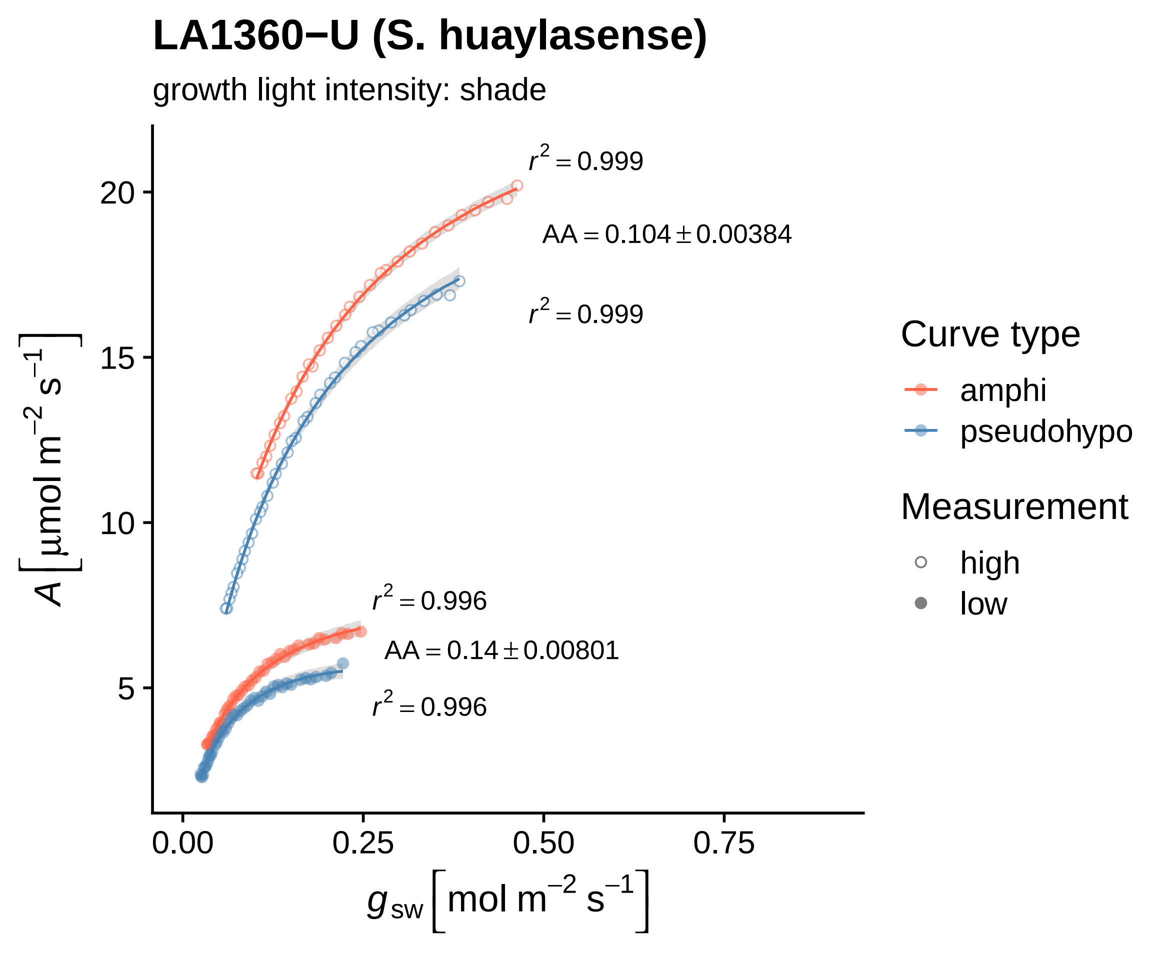

LA1360−U (S. huaylasense)
growth light intensity: shade
$r^2 = 0.999$
$AA = 0.104 \pm 0.00384$
$r^2 = 0.999$
$r^2 = 0.996$
$AA = 0.14 \pm 0.00801$
$r^2 = 0.996$
20
15
10
5
0.00
0.25
0.50
0.75
$A\left[\mu mol\ m^{-2}\ s^{-1}\right]$
$g_{sw}\left[mol\ m^{-2}\ s^{-1}\right]$
Curve type
amphi
pseudohypo
Measurement
high
low

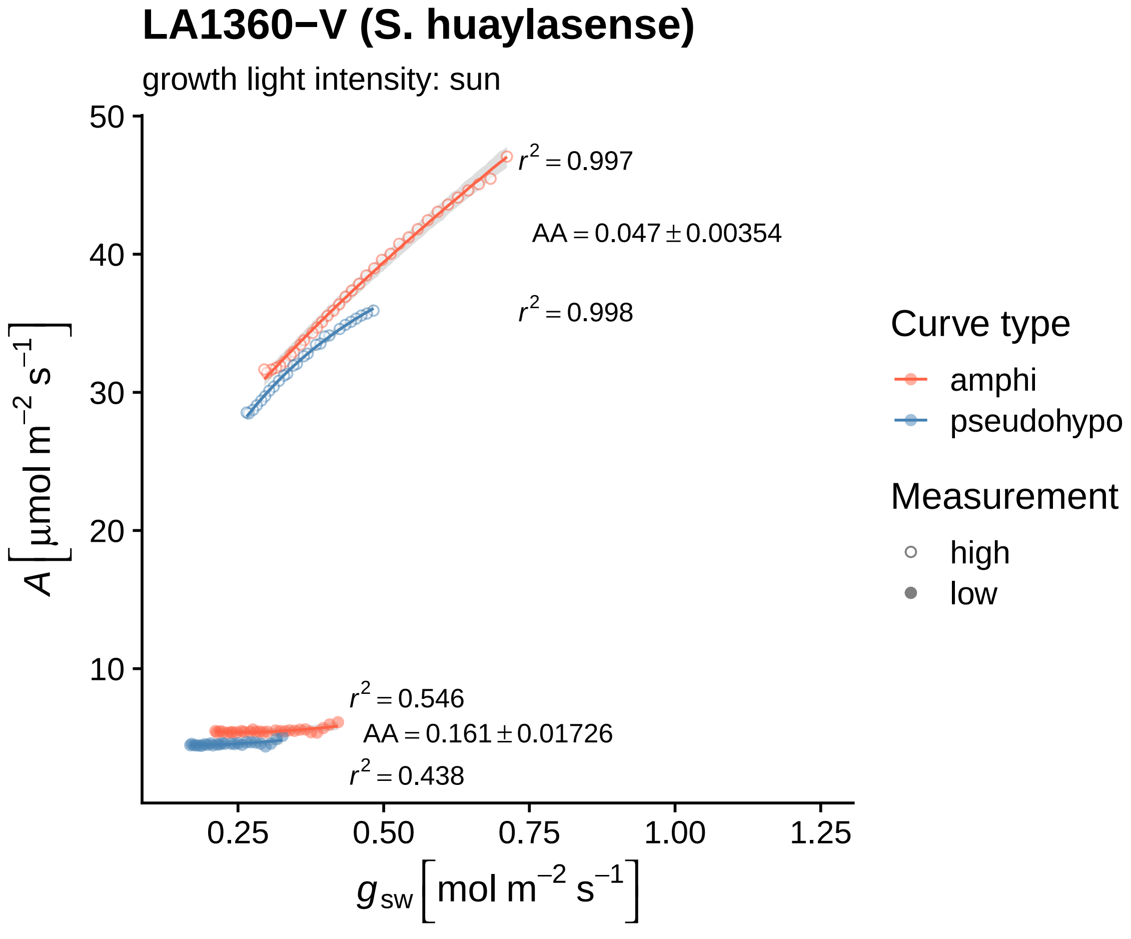

LA1360−V (S. huaylasense)
growth light intensity: sun
$r^2 = 0.997$
$AA = 0.047 \pm 0.00354$
$r^2 = 0.998$
$r^2 = 0.546$
$AA = 0.161 \pm 0.01726$
$r^2 = 0.438$
50
40
30
20
10
0.25
0.50
0.75
1.00
1.25
$A\ [\mu mol\ m^{-2}\ s^{-1}]$
$g_{sw}\ [mol\ m^{-2}\ s^{-1}]$
Curve type
amphi
pseudohypo
Measurement
high
low

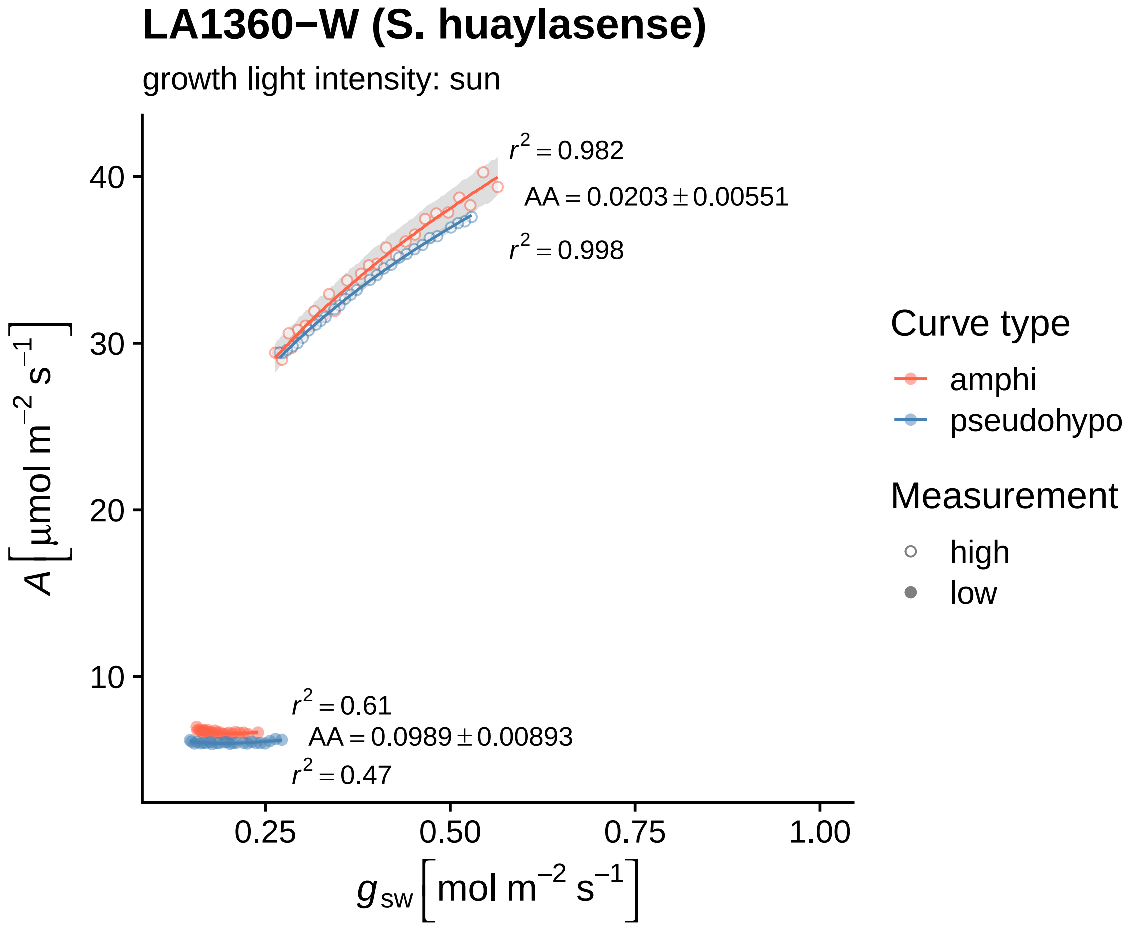

LA1360−W (S. huaylasense)
growth light intensity: sun
$r^2 = 0.982$
$AA = 0.0203 \pm 0.00551$
$r^2 = 0.998$
$r^2 = 0.61$
$AA = 0.0989 \pm 0.00893$
$r^2 = 0.47$
40
30
20
10
0.25
0.50
0.75
1.00
$A\left[\mu mol\ m^{-2}\ s^{-1}\right]$
$g_{sw}\left[mol\ m^{-2}\ s^{-1}\right]$
Curve type
amphi
pseudohypo
Measurement
high
low

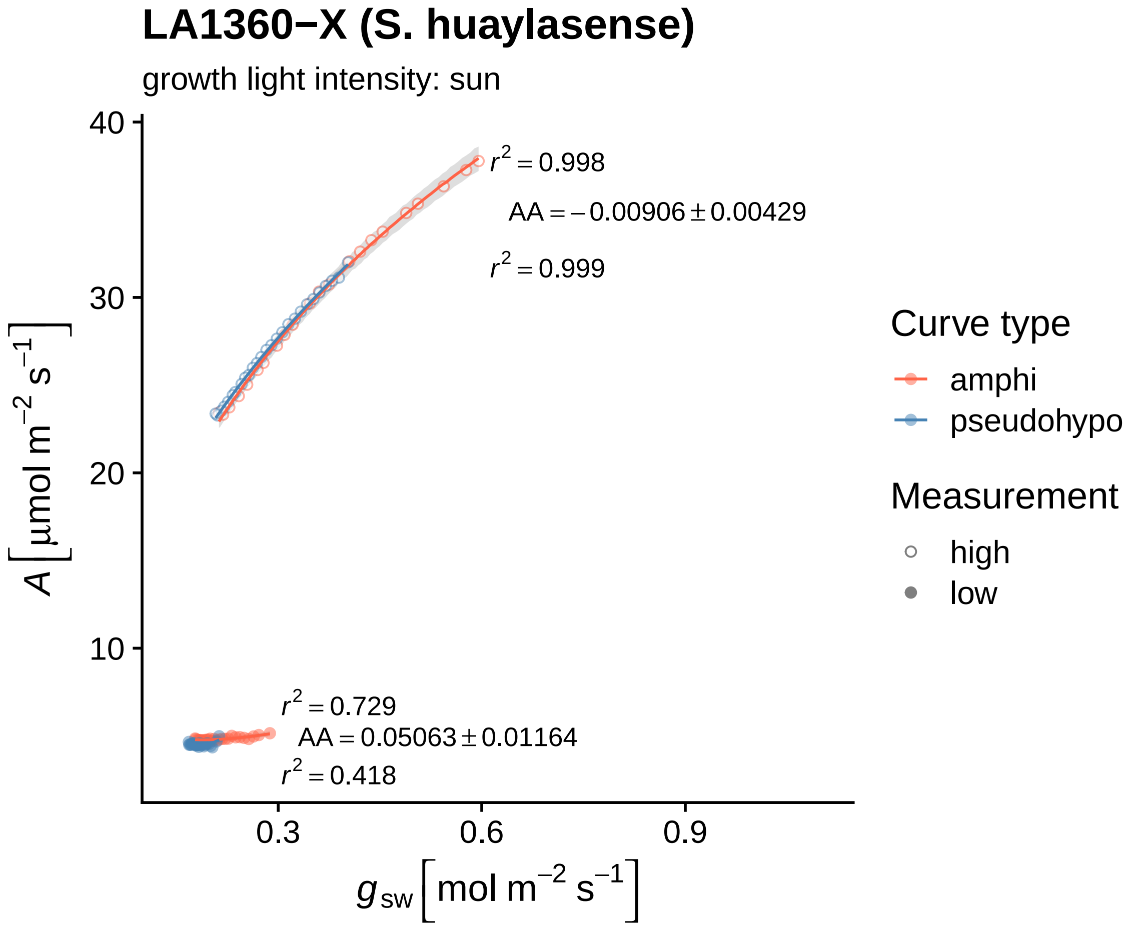

LA1360−X (S. huaylasense)
growth light intensity: sun
$r^2 = 0.998$
$AA = -0.00906 \pm 0.00429$
$r^2 = 0.999$
$r^2 = 0.729$
$AA = 0.05063 \pm 0.01164$
$r^2 = 0.418$
40
30
20
10
0.3
0.6
0.9
$A\ [\mu mol\ m^{-2}\ s^{-1}]$
$g_{sw}\ [mol\ m^{-2}\ s^{-1}]$
Curve type
amphi
pseudohypo
Measurement
high
low

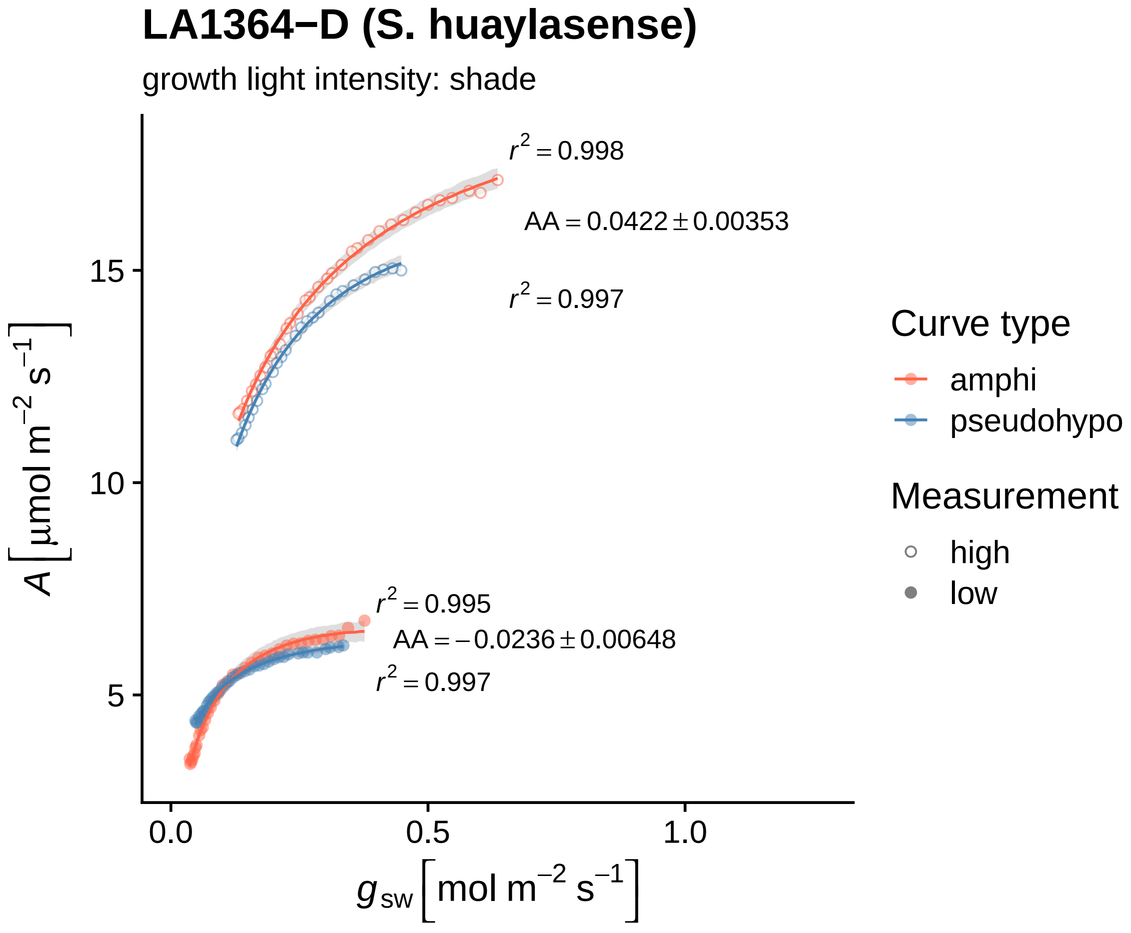

LA1364−D (S. huaylasense)
growth light intensity: shade
$r^2 = 0.998$
$AA = 0.0422 \pm 0.00353$
$r^2 = 0.997$
$r^2 = 0.995$
$AA = -0.0236 \pm 0.00648$
$r^2 = 0.997$
15
10
5
0.0
0.5
1.0
$A$ [µmol m$^{-2}$ s$^{-1}$]
$g_{sw}$ [mol m$^{-2}$ s$^{-1}$]
Curve type
amphi
pseudohypo
Measurement
high
low

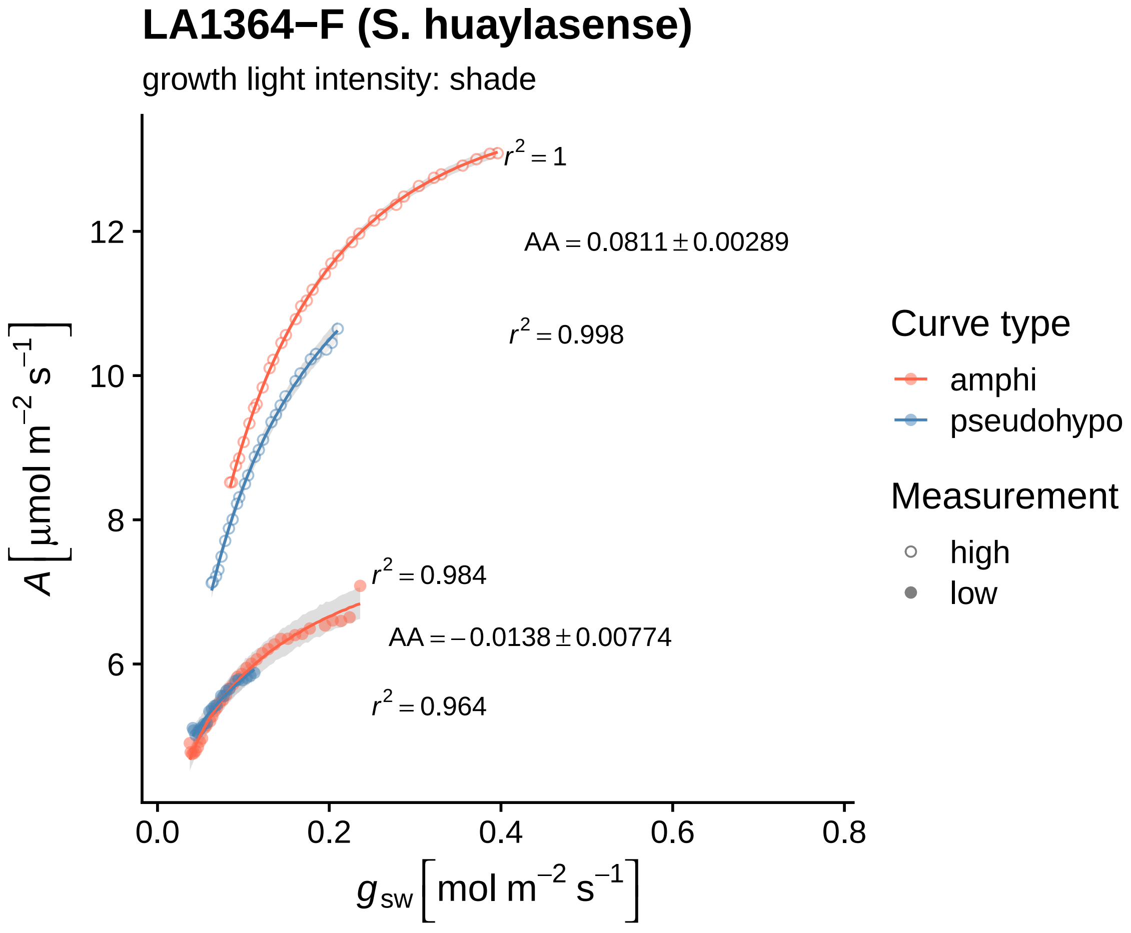

LA1364−F (S. huaylasense)
growth light intensity: shade
$r^2 = 1$
$AA = 0.0811 \pm 0.00289$
$r^2 = 0.998$
$r^2 = 0.984$
$AA = -0.0138 \pm 0.00774$
$r^2 = 0.964$
12
10
8
6
0.0
0.2
0.4
0.6
0.8
$A\,[\mu mol\ m^{-2}\ s^{-1}]$
$g_{sw}\,[mol\ m^{-2}\ s^{-1}]$
Curve type
amphi
pseudohypo
Measurement
high
low

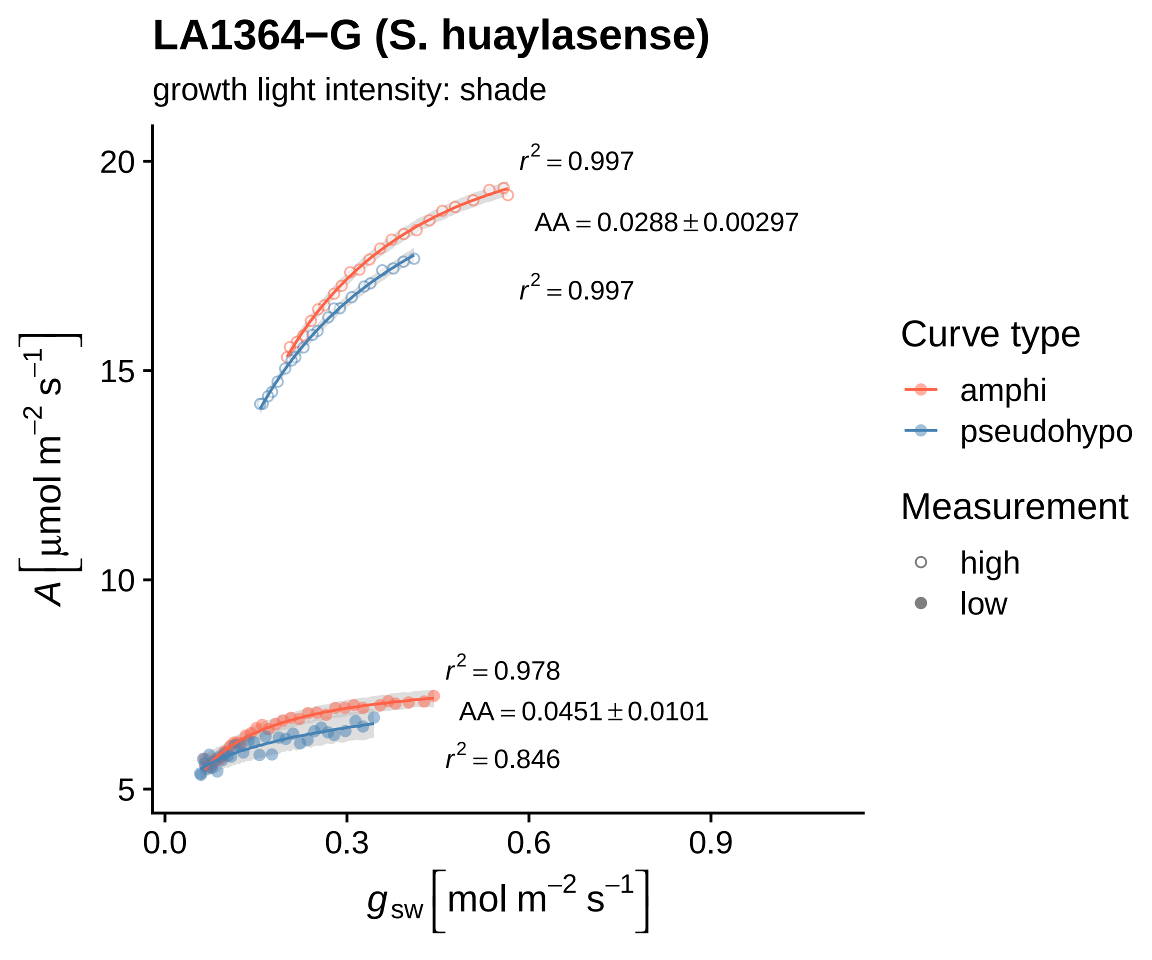

LA1364−G (S. huaylasense)
growth light intensity: shade
$r^2 = 0.997$
$AA = 0.0288 \pm 0.00297$
$r^2 = 0.997$
$r^2 = 0.978$
$AA = 0.0451 \pm 0.0101$
$r^2 = 0.846$
20
15
10
5
0.0
0.3
0.6
0.9
$A\ [\mu mol\ m^{-2}\ s^{-1}]$
$g_{sw}\ [mol\ m^{-2}\ s^{-1}]$
Curve type
amphi
pseudohypo
Measurement
high
low

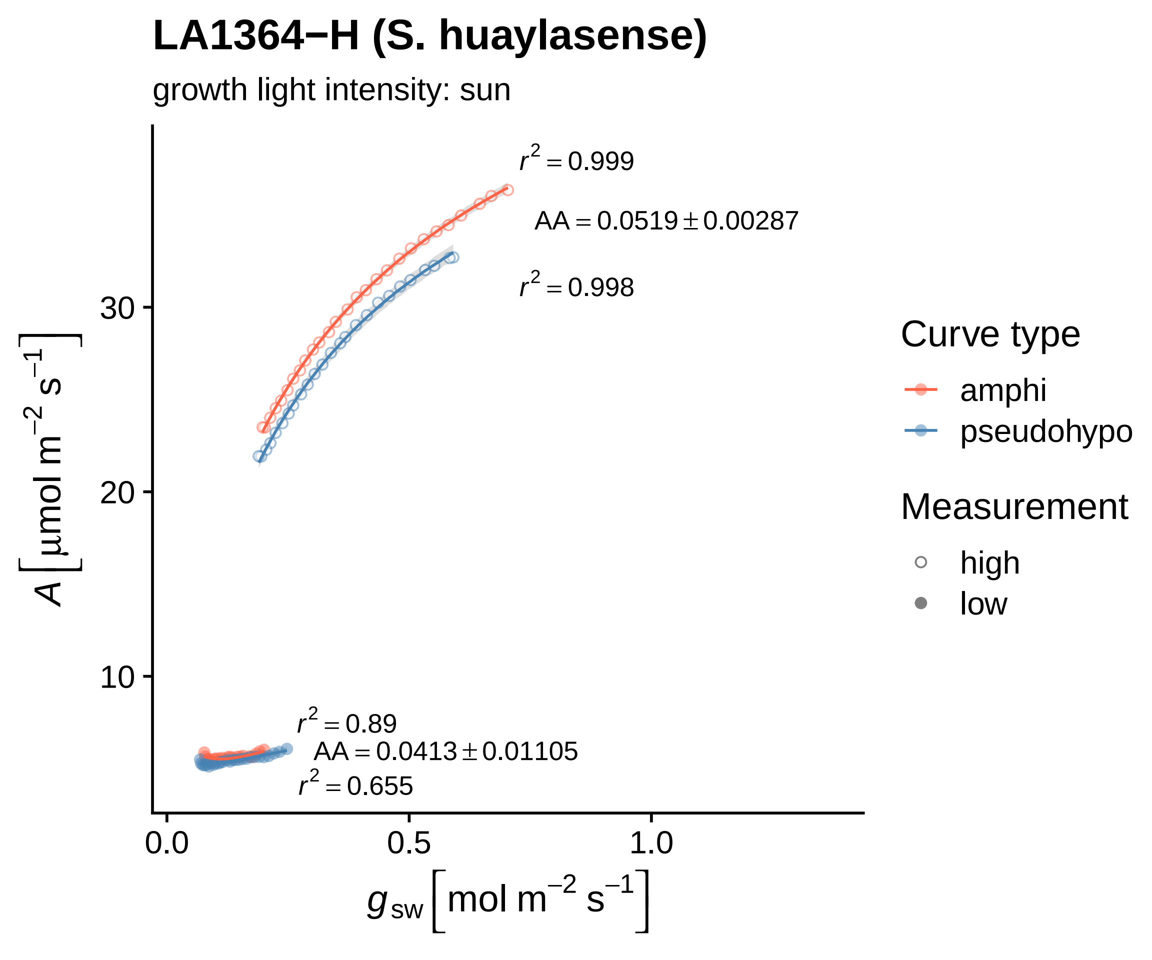
LA1364−H (S. huaylasense)
growth light intensity: sun
$r^2 = 0.999$
$AA = 0.0519 \pm 0.00287$
$r^2 = 0.998$
$r^2 = 0.89$
$AA = 0.0413 \pm 0.01105$
$r^2 = 0.655$
30
20
10
0.0
0.5
1.0
$A\left[\mu mol\ m^{-2}\ s^{-1}\right]$
$g_{sw}\left[mol\ m^{-2}\ s^{-1}\right]$
Curve type
amphi
pseudohypo
Measurement
high
low

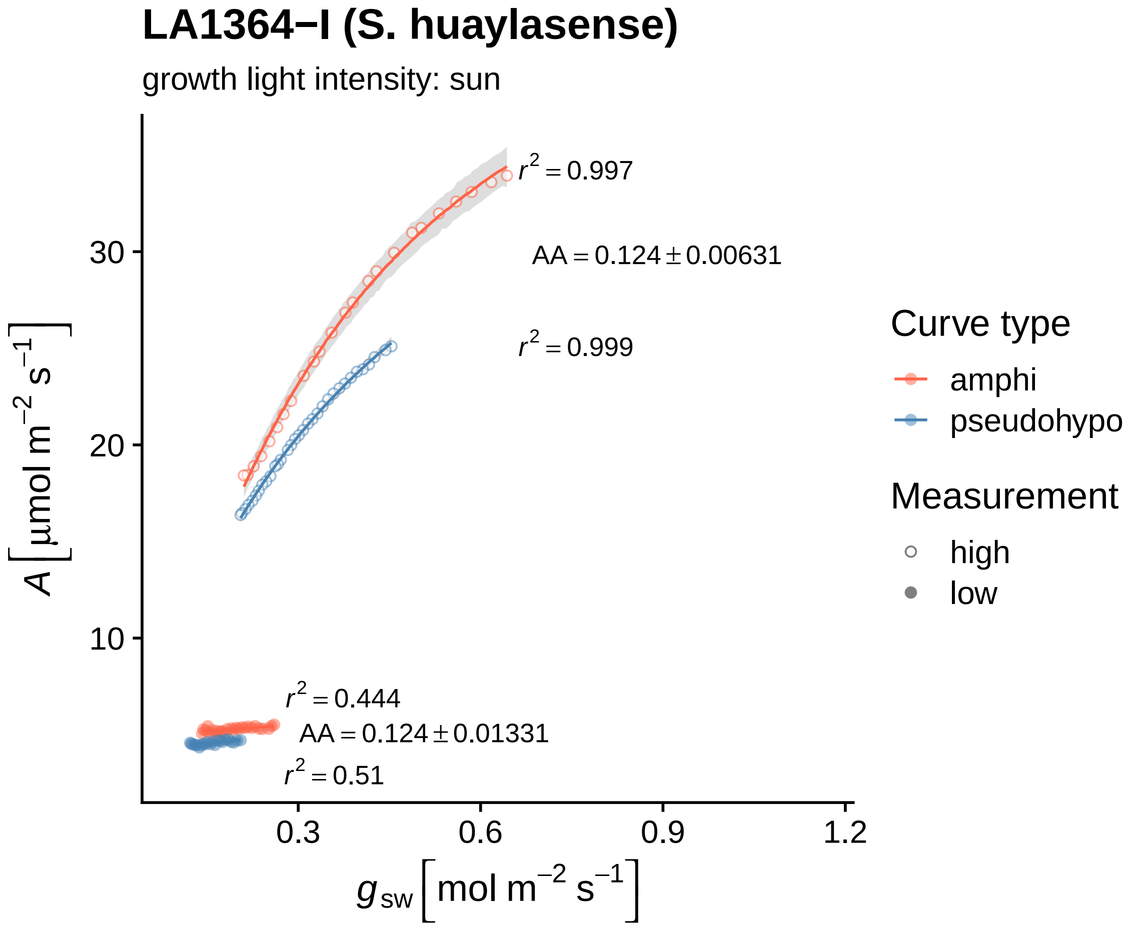
LA1364−I (S. huaylasense)
growth light intensity: sun
$r^2 = 0.997$
$AA = 0.124 \pm 0.00631$
$r^2 = 0.999$
$r^2 = 0.444$
$AA = 0.124 \pm 0.01331$
$r^2 = 0.51$
30
20
10
0.3
0.6
0.9
1.2
$A\ [\mu mol\ m^{-2}\ s^{-1}]$
$g_{sw}\ [mol\ m^{-2}\ s^{-1}]$
Curve type
amphi
pseudohypo
Measurement
high
low

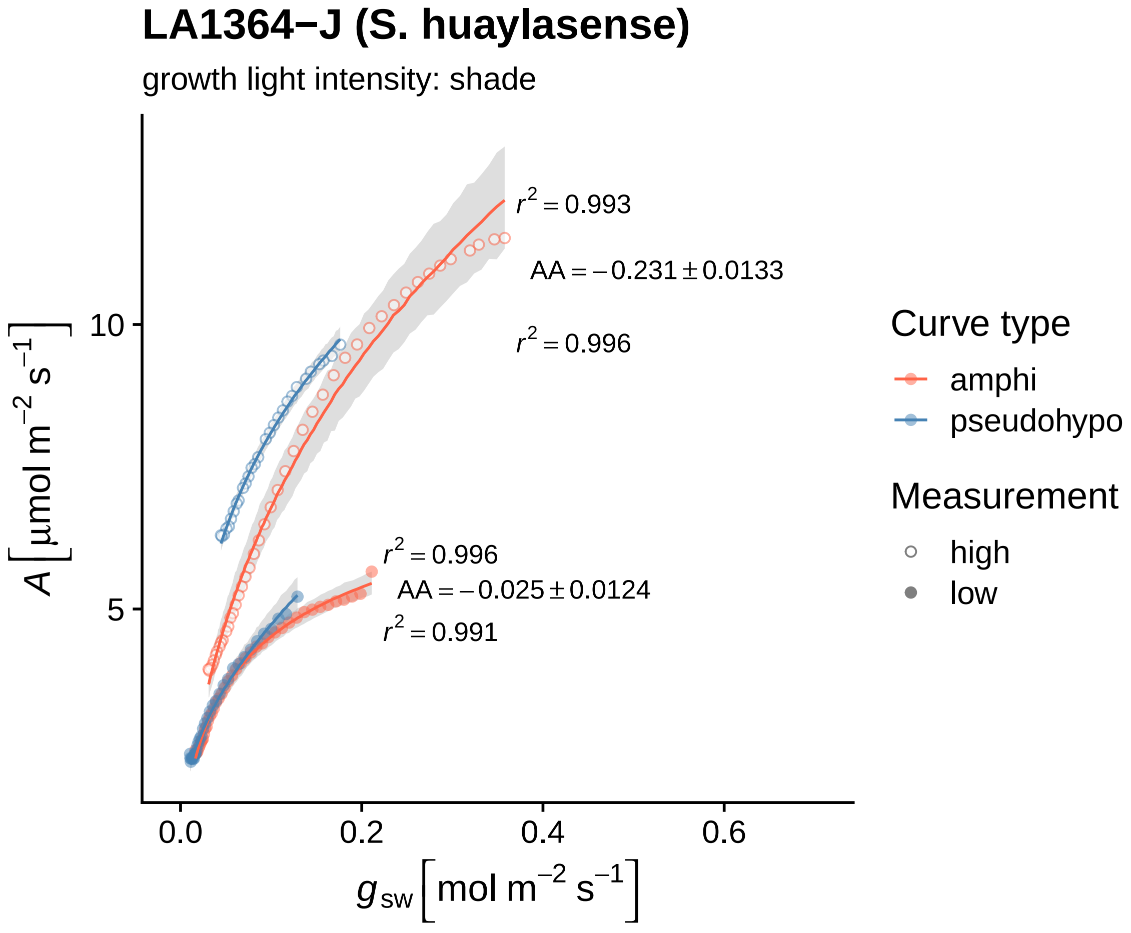

LA1364−J (S. huaylasense)
growth light intensity: shade
r² = 0.993
AA = −0.231 ± 0.0133
r² = 0.996
r² = 0.996
AA = −0.025 ± 0.0124
r² = 0.991
10
5
0.0
0.2
0.4
0.6
A [μmol m⁻² s⁻¹]
g_sw [mol m⁻² s⁻¹]
Curve type
amphi
pseudohypo
Measurement
high
low

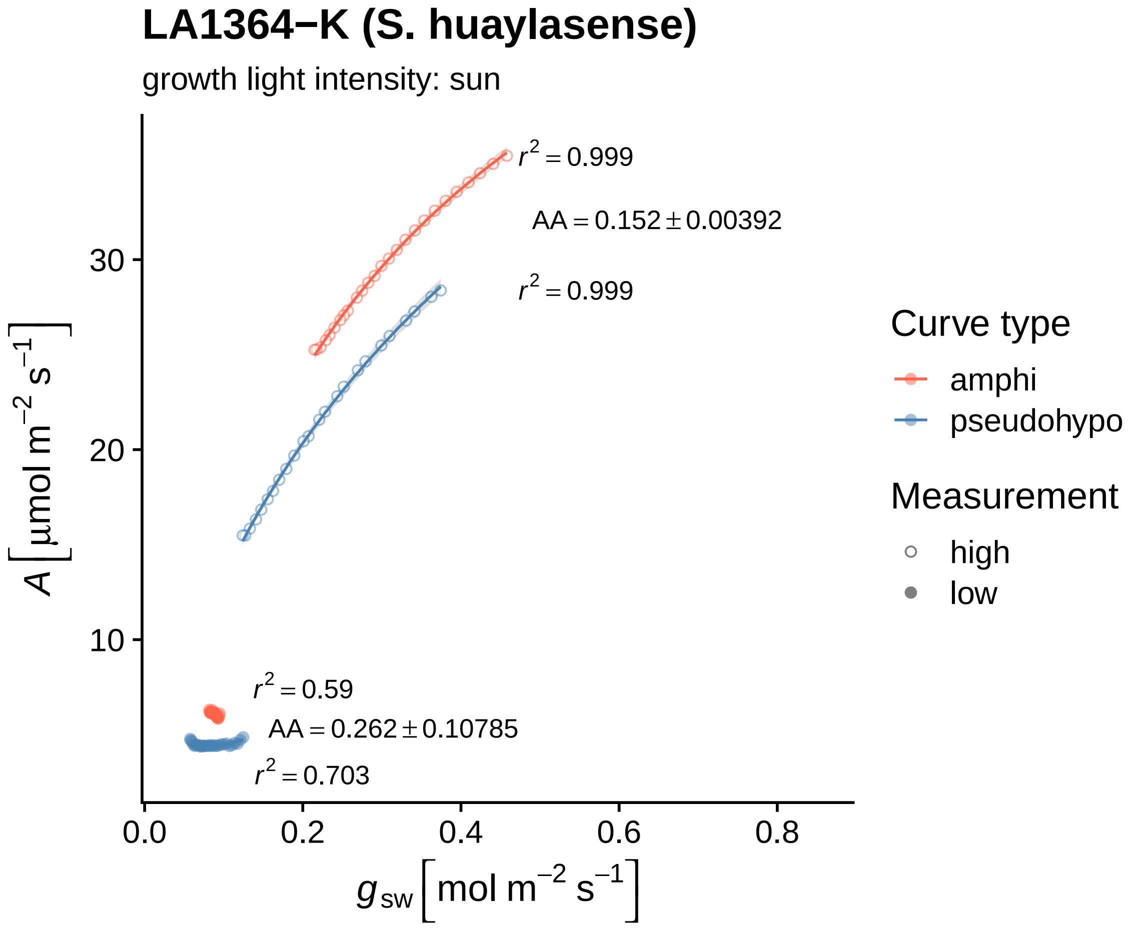

LA1364−K (S. huaylasense)
growth light intensity: sun
$r^2 = 0.999$
$AA = 0.152 \pm 0.00392$
$r^2 = 0.999$
$r^2 = 0.59$
$AA = 0.262 \pm 0.10785$
$r^2 = 0.703$
30
20
10
0.0
0.2
0.4
0.6
0.8
$A\ [\mu mol\ m^{-2}\ s^{-1}]$
$g_{sw}\ [mol\ m^{-2}\ s^{-1}]$
Curve type
amphi
pseudohypo
Measurement
high
low

### LA1364−L (S. huaylasense)

growth light intensity: sun

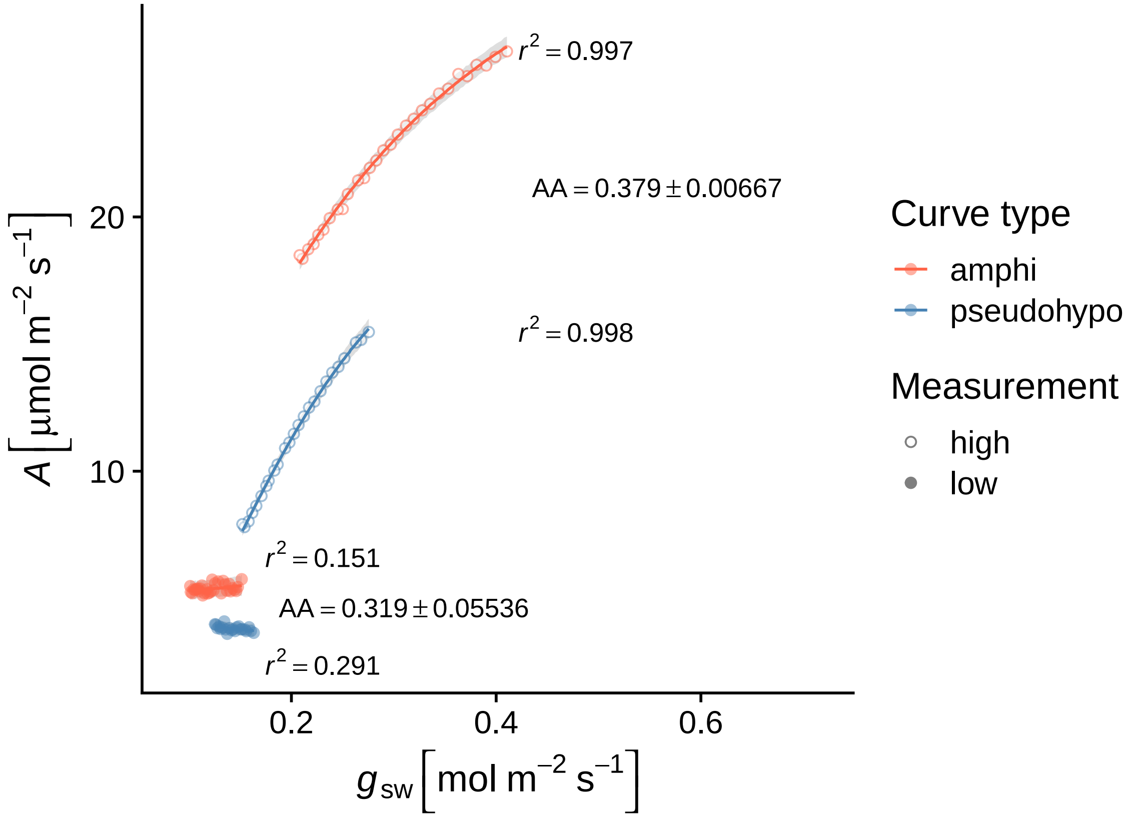

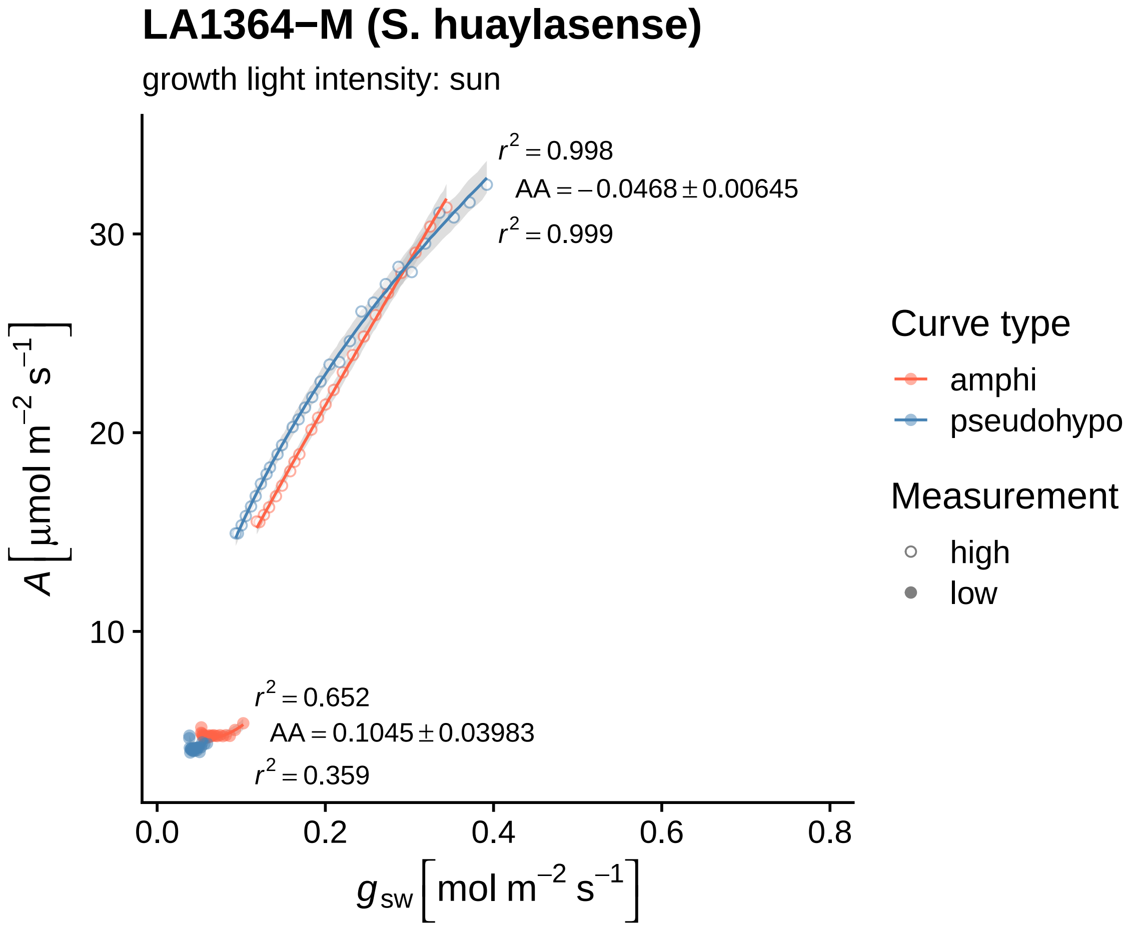
LA1364−M (S. huaylasense)
growth light intensity: sun
r² = 0.998
AA = −0.0468 ± 0.00645
r² = 0.999
r² = 0.652
AA = 0.1045 ± 0.03983
r² = 0.359
A [μmol m⁻² s⁻¹]
30
20
10
0.0
0.2
0.4
0.6
0.8
g_sw [mol m⁻² s⁻¹]
Curve type
amphi
pseudohypo
Measurement
high
low

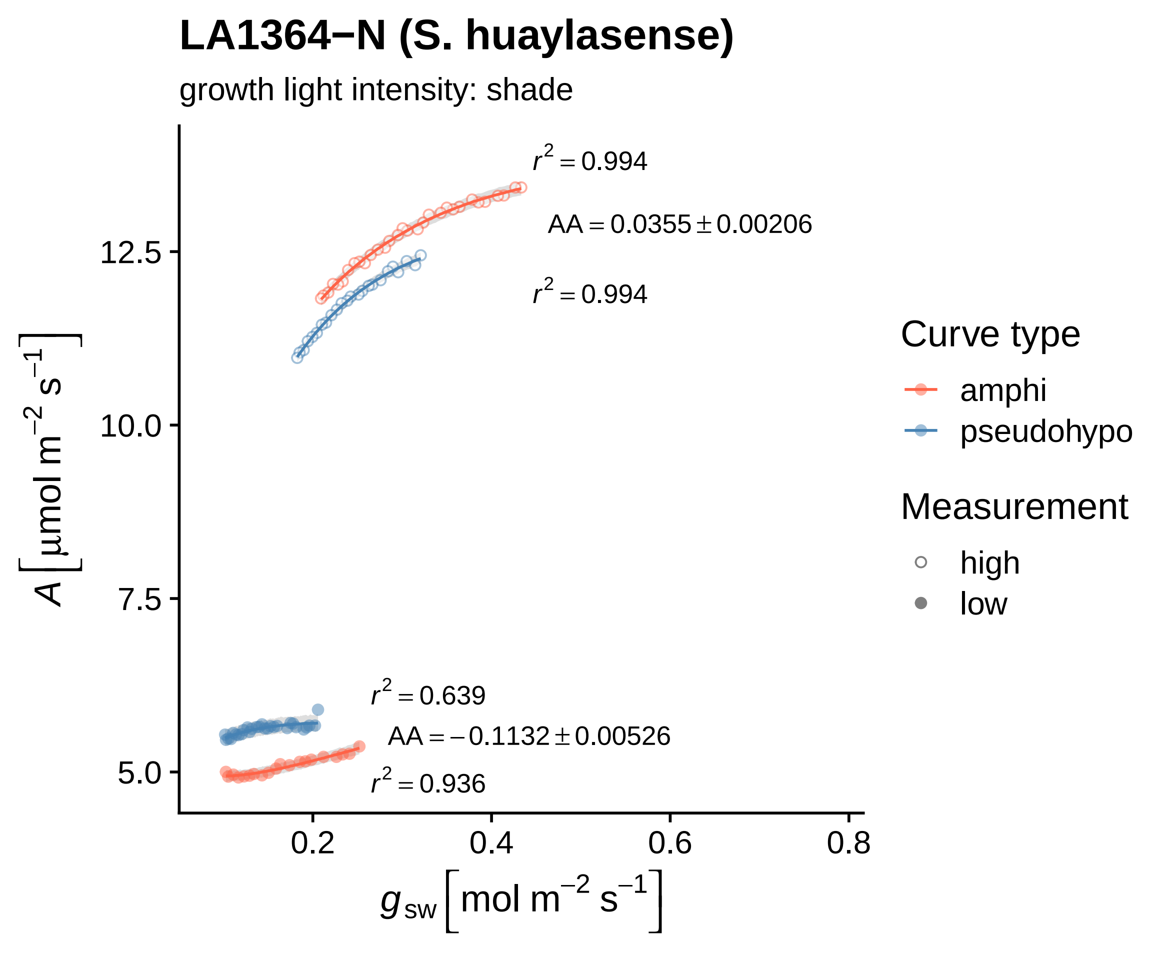
LA1364−N (S. huaylasense)
growth light intensity: shade
$r^2 = 0.994$
$AA = 0.0355 \pm 0.00206$
$r^2 = 0.994$
$r^2 = 0.639$
$AA = -0.1132 \pm 0.00526$
$r^2 = 0.936$
12.5
10.0
7.5
5.0
0.2
0.4
0.6
0.8
$A\,[\mu mol\ m^{-2}\ s^{-1}]$
$g_{sw}\,[mol\ m^{-2}\ s^{-1}]$
Curve type
amphi
pseudohypo
Measurement
high
low

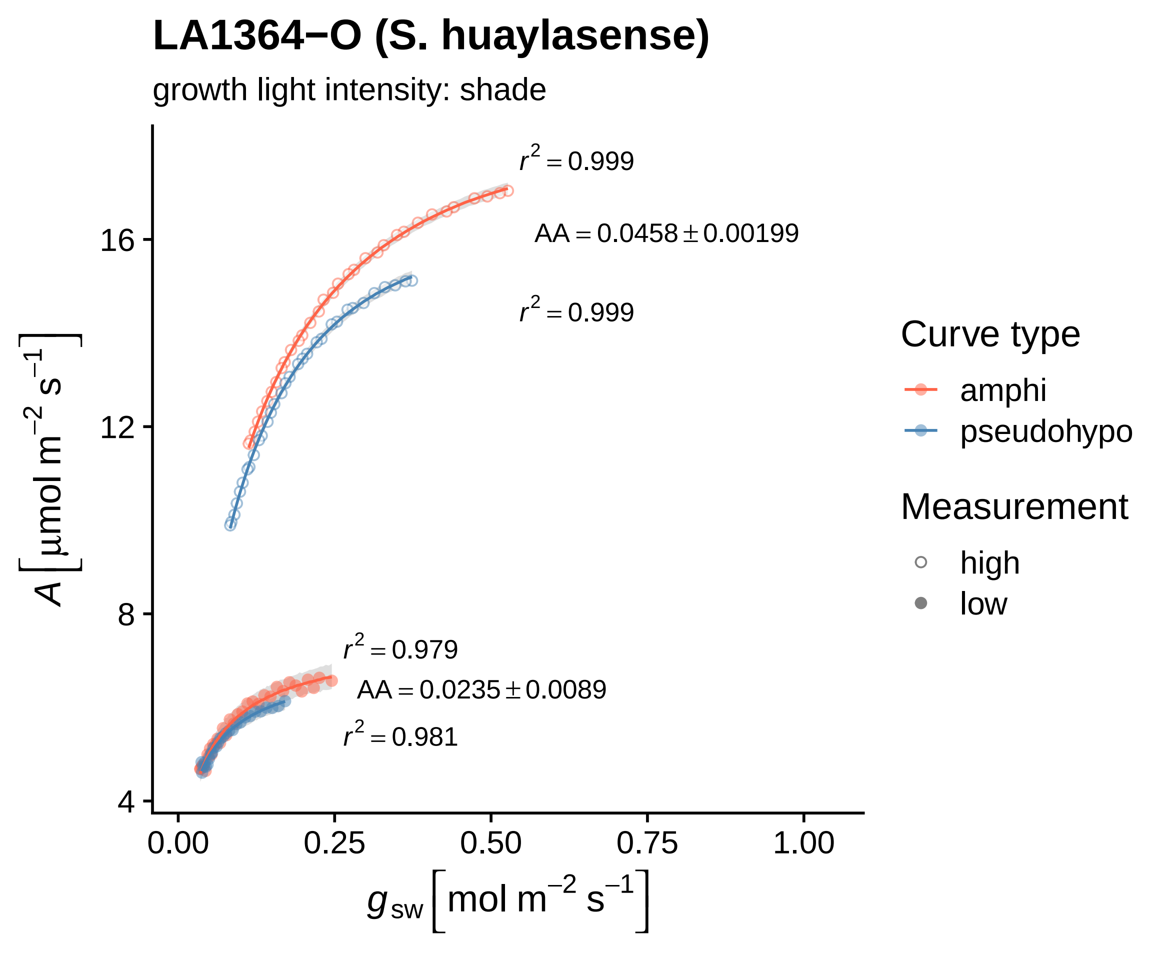

LA1364−O (S. huaylasense)
growth light intensity: shade
$r^2 = 0.999$
$AA = 0.0458 \pm 0.00199$
$r^2 = 0.999$
$r^2 = 0.979$
$AA = 0.0235 \pm 0.0089$
$r^2 = 0.981$
16
12
8
4
0.00
0.25
0.50
0.75
1.00
$A\ [\mu mol\ m^{-2}\ s^{-1}]$
$g_{sw}\ [mol\ m^{-2}\ s^{-1}]$
Curve type
amphi
pseudohypo
Measurement
high
low

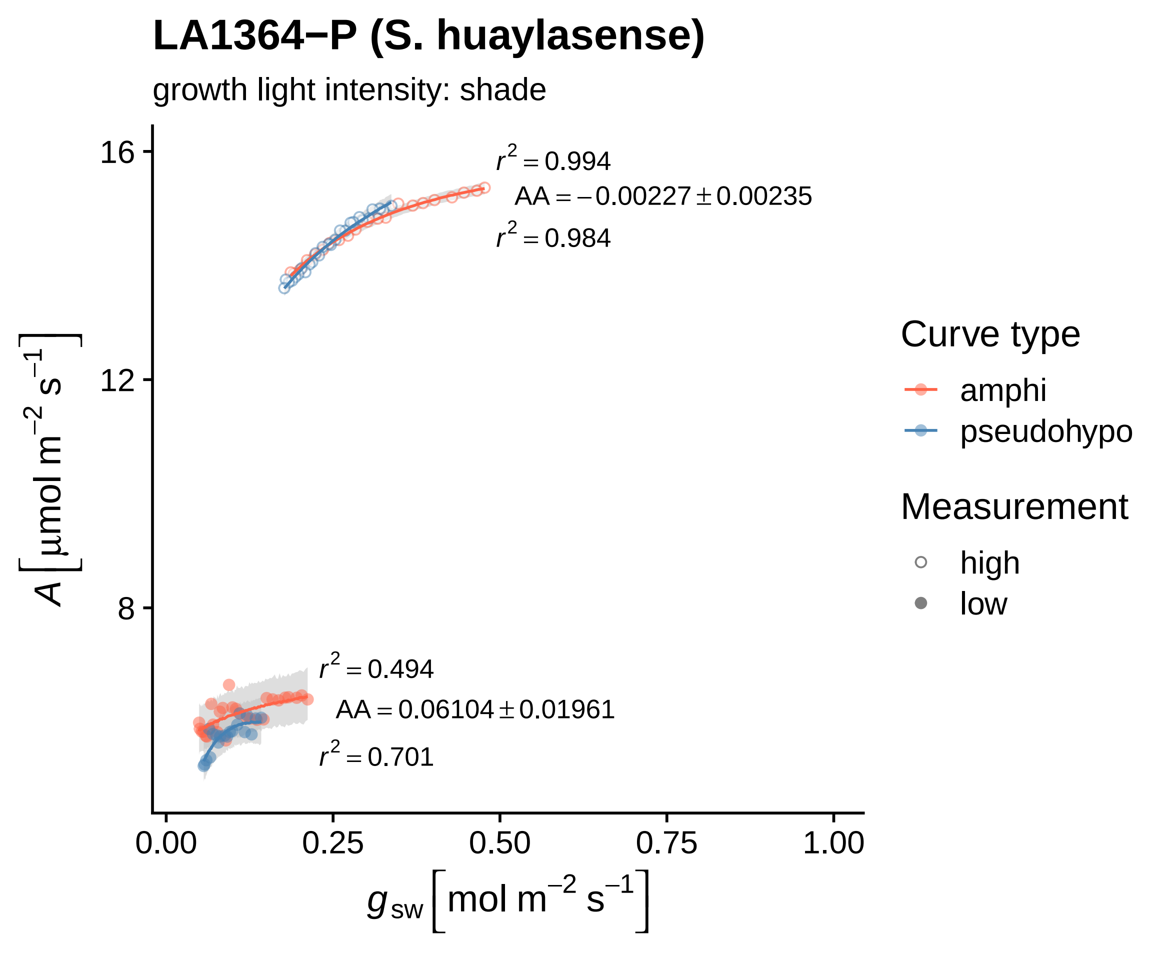
LA1364−P (S. huaylasense)
growth light intensity: shade
r² = 0.994
AA = −0.00227 ± 0.00235
r² = 0.984
r² = 0.494
AA = 0.06104 ± 0.01961
r² = 0.701
16
12
8
0.00
0.25
0.50
0.75
1.00
A [μmol m⁻² s⁻¹]
g_sw [mol m⁻² s⁻¹]
Curve type
amphi
pseudohypo
Measurement
high
low

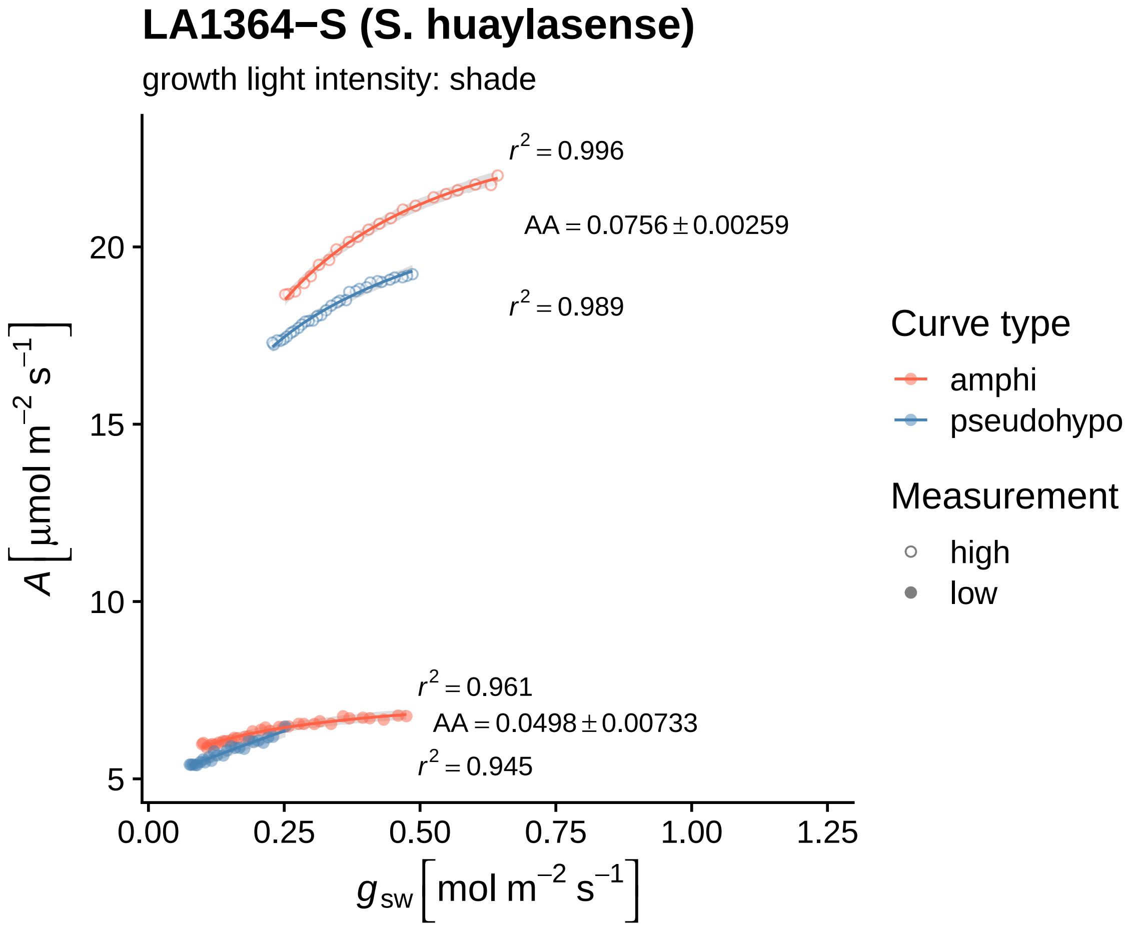
LA1364−S (S. huaylasense)
growth light intensity: shade
$r^2 = 0.996$
$AA = 0.0756 \pm 0.00259$
$r^2 = 0.989$
$r^2 = 0.961$
$AA = 0.0498 \pm 0.00733$
$r^2 = 0.945$
$A\left[\mu mol\ m^{-2}\ s^{-1}\right]$
20
15
10
5
0.00
0.25
0.50
0.75
1.00
1.25
$g_{sw}\left[mol\ m^{-2}\ s^{-1}\right]$
Curve type
amphi
pseudohypo
Measurement
high
low

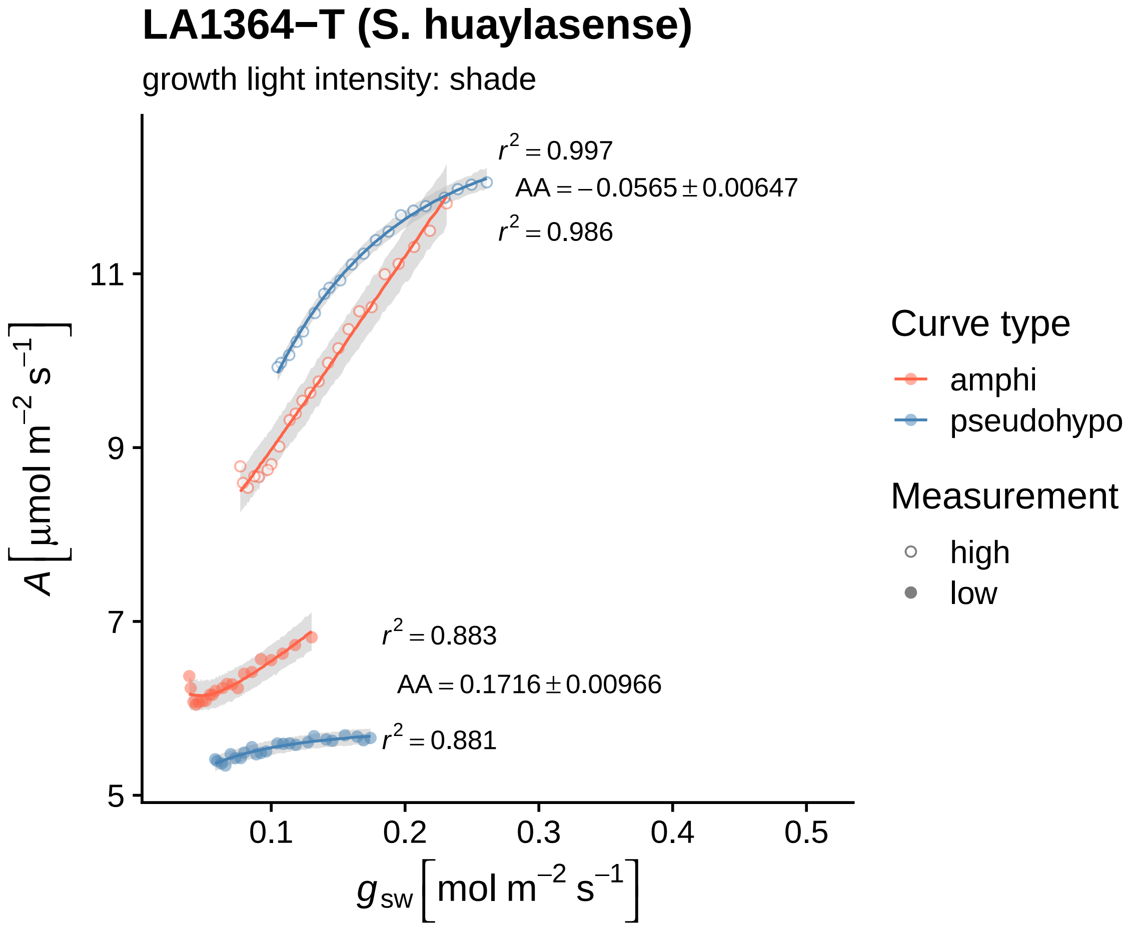

LA1364−T (S. huaylasense)
growth light intensity: shade
$r^2 = 0.997$
$AA = -0.0565 \pm 0.00647$
$r^2 = 0.986$
$r^2 = 0.883$
$AA = 0.1716 \pm 0.00966$
$r^2 = 0.881$
11
9
7
5
0.1
0.2
0.3
0.4
0.5
$A\,[\mu mol\ m^{-2}\ s^{-1}]$
$g_{sw}\,[mol\ m^{-2}\ s^{-1}]$
Curve type
amphi
pseudohypo
Measurement
high
low

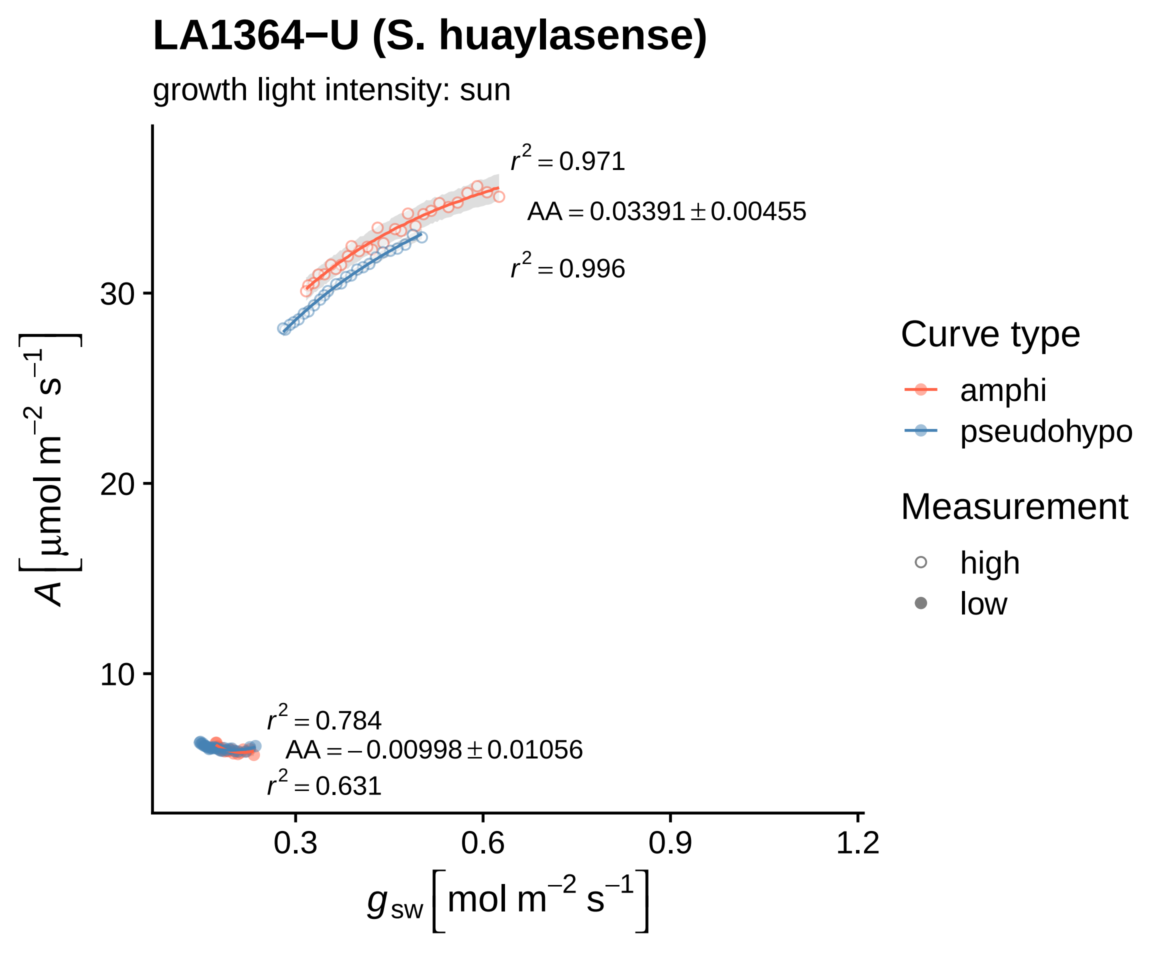

LA1364−U (S. huaylasense)
growth light intensity: sun
r² = 0.971
AA = 0.03391 ± 0.00455
r² = 0.996
r² = 0.784
AA = −0.00998 ± 0.01056
r² = 0.631
30
20
10
0.3
0.6
0.9
1.2
A [μmol m⁻² s⁻¹]
g_sw [mol m⁻² s⁻¹]
Curve type
amphi
pseudohypo
Measurement
high
low

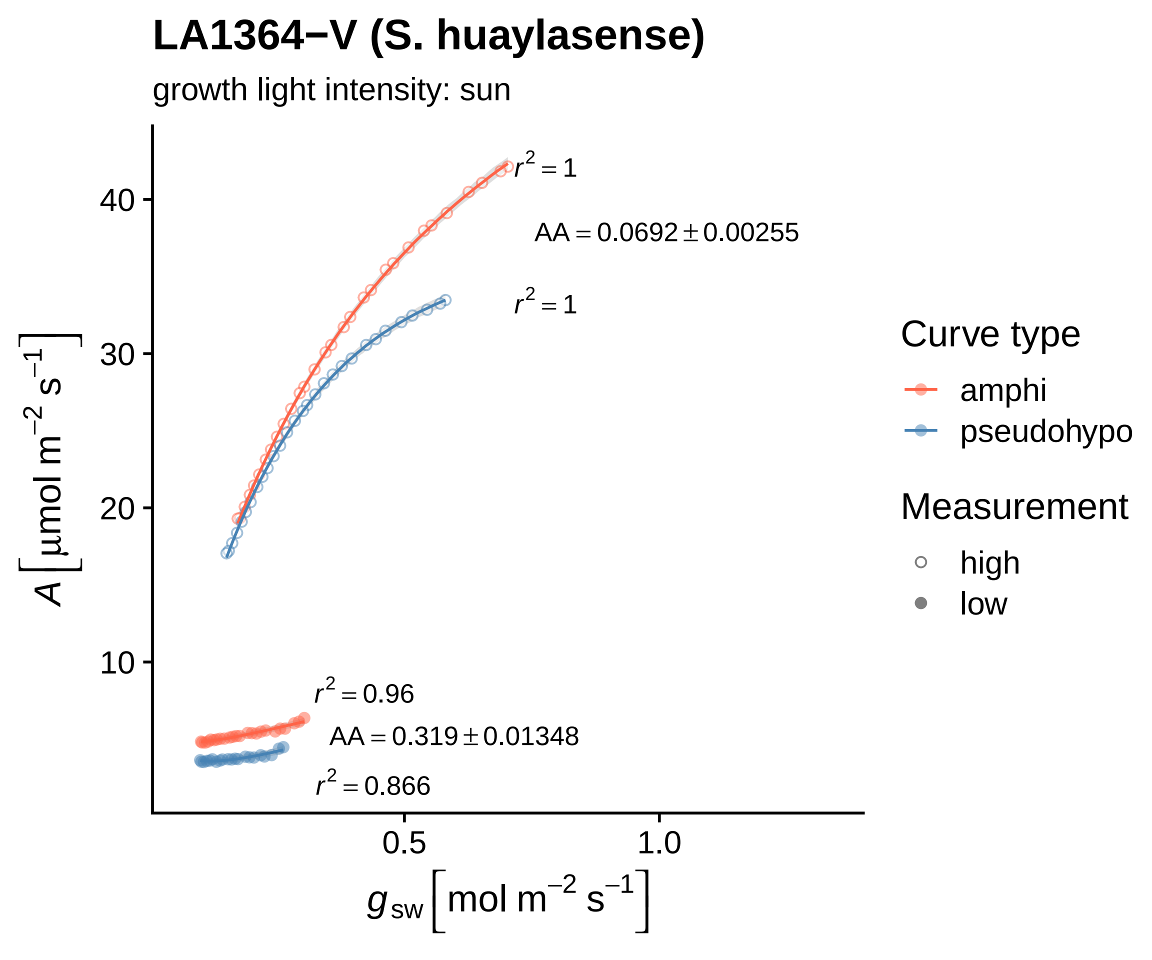

LA1364−V (S. huaylasense)
growth light intensity: sun
r² = 1
AA = 0.0692 ± 0.00255
r² = 1
r² = 0.96
AA = 0.319 ± 0.01348
r² = 0.866
10
20
30
40
0.5
1.0
A [μmol m⁻² s⁻¹]
g_sw [mol m⁻² s⁻¹]
Curve type
amphi
pseudohypo
Measurement
high
low

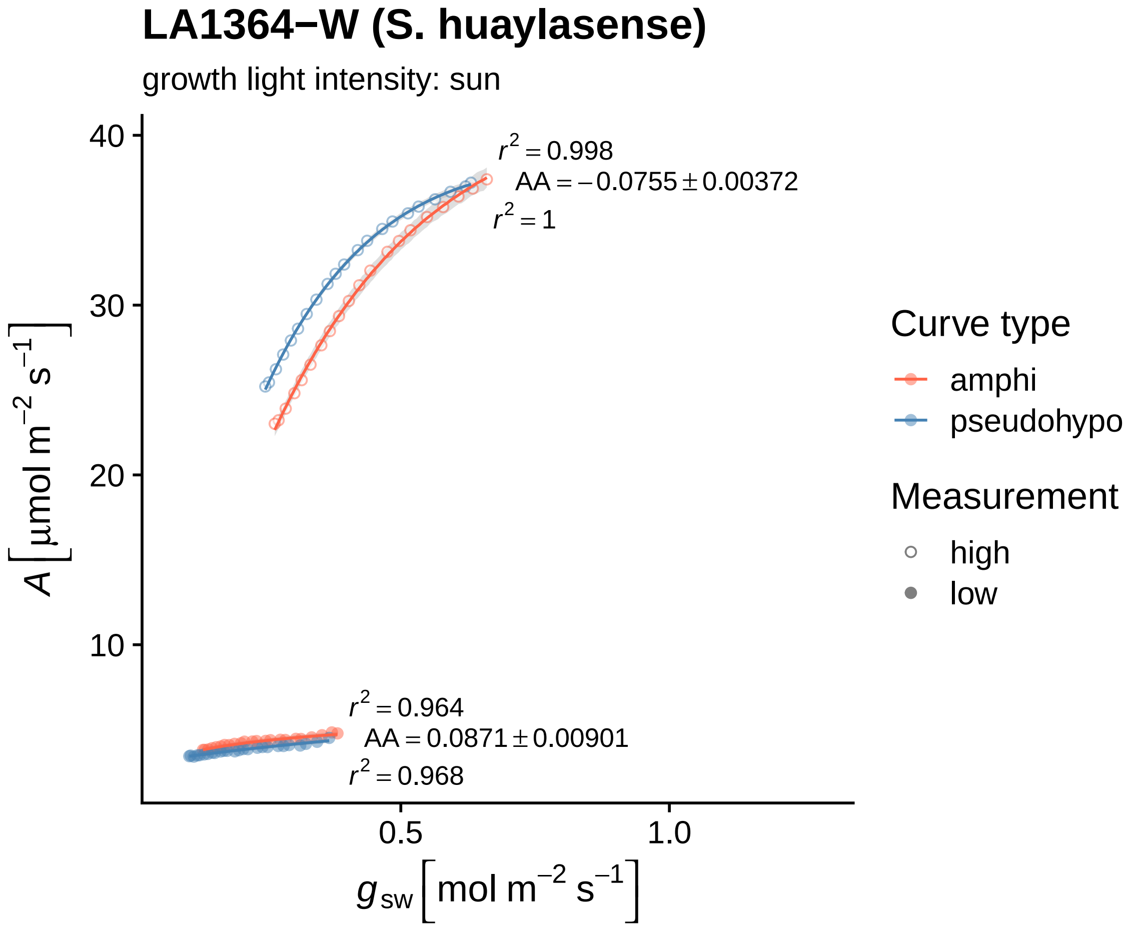
LA1364−W (S. huaylasense)
growth light intensity: sun
$r^2 = 0.998$
$AA = -0.0755 \pm 0.00372$
$r^2 = 1$
$r^2 = 0.964$
$AA = 0.0871 \pm 0.00901$
$r^2 = 0.968$
40
30
20
10
0.5
1.0
$A\ [\mu mol\ m^{-2}\ s^{-1}]$
$g_{sw}\ [mol\ m^{-2}\ s^{-1}]$
Curve type
amphi
pseudohypo
Measurement
high
low

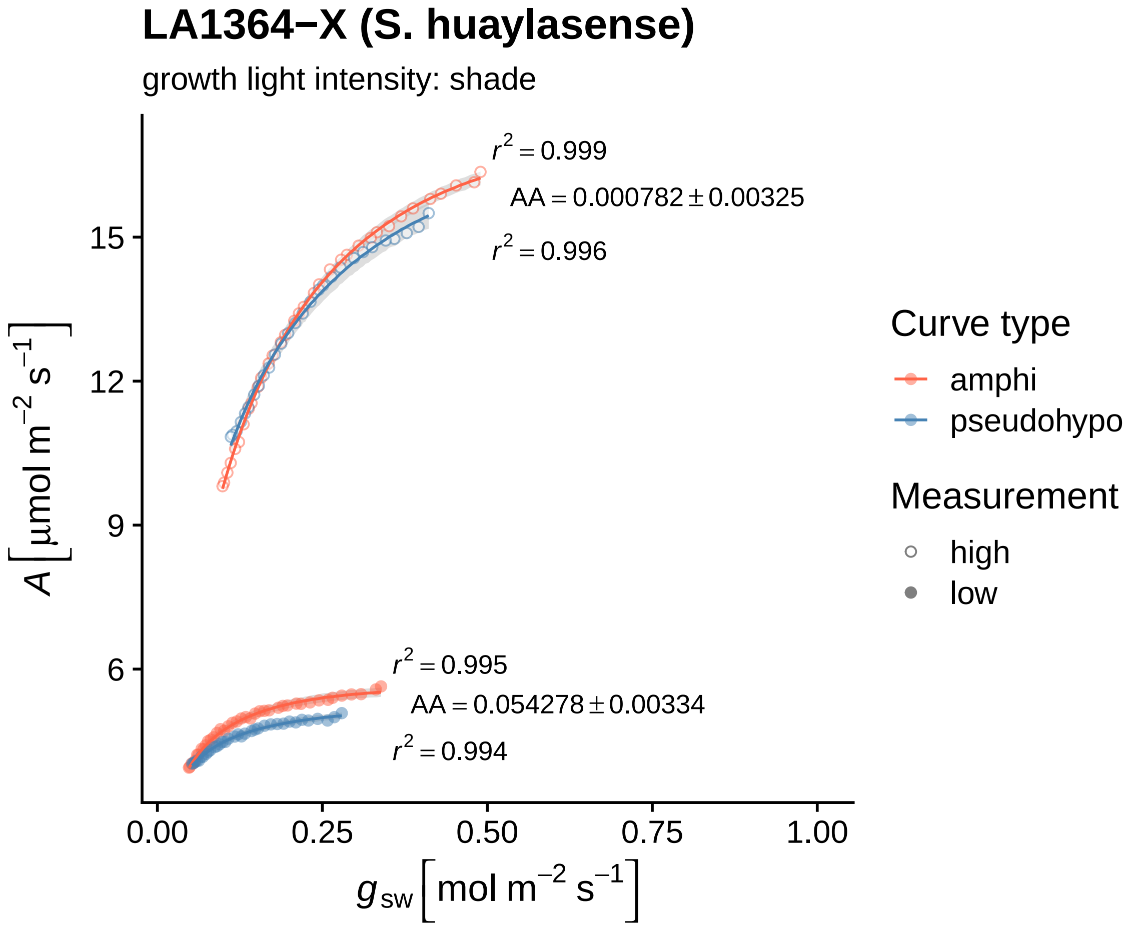

LA1364−X (S. huaylasense)
growth light intensity: shade
$r^2 = 0.999$
$AA = 0.000782 \pm 0.00325$
$r^2 = 0.996$
$r^2 = 0.995$
$AA = 0.054278 \pm 0.00334$
$r^2 = 0.994$
15
12
9
6
0.00
0.25
0.50
0.75
1.00
$A\ [\mu mol\ m^{-2}\ s^{-1}]$
$g_{sw}\ [mol\ m^{-2}\ s^{-1}]$
Curve type
amphi
pseudohypo
Measurement
high
low

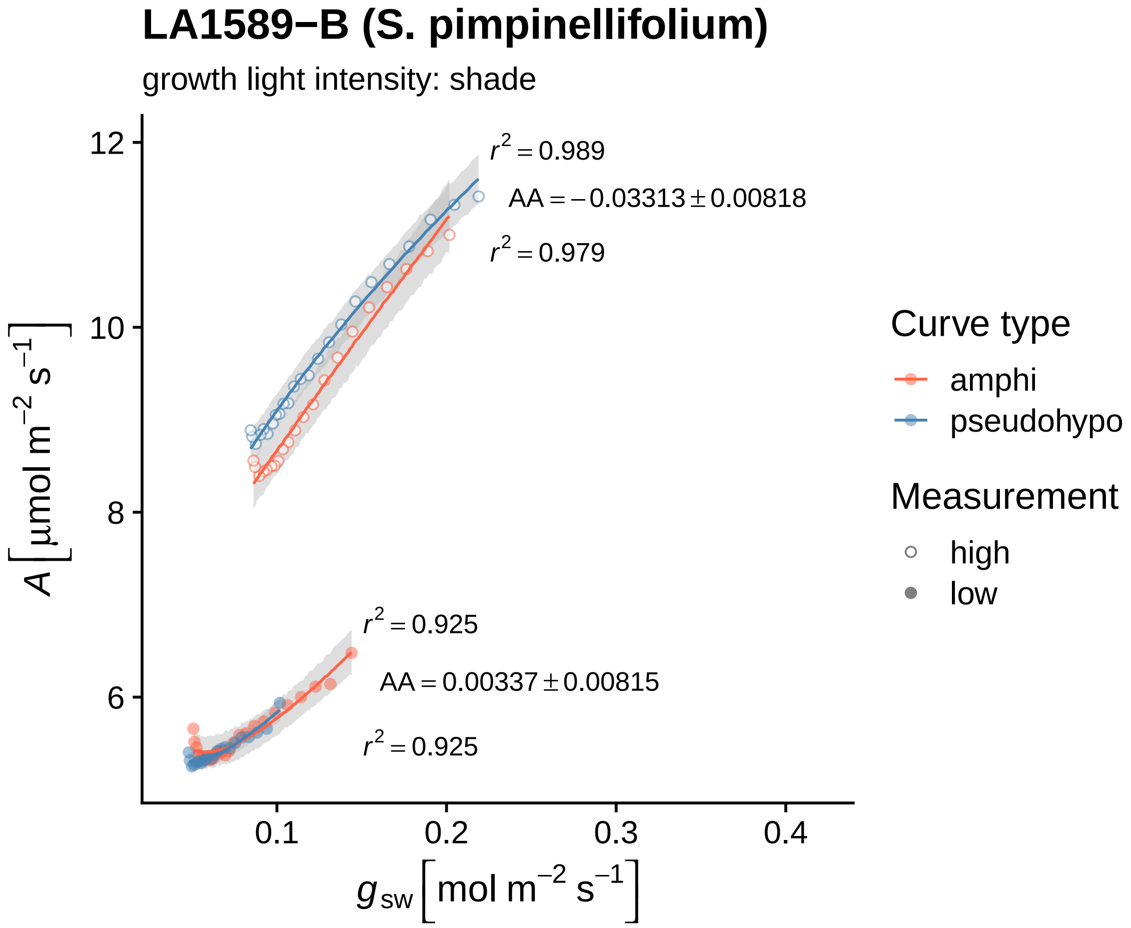

LA1589−B (S. pimpinellifolium)
growth light intensity: shade
r² = 0.989
AA = −0.03313 ± 0.00818
r² = 0.979
r² = 0.925
AA = 0.00337 ± 0.00815
r² = 0.925
12
10
8
6
0.1
0.2
0.3
0.4
A [μmol m⁻² s⁻¹]
g_sw [mol m⁻² s⁻¹]
Curve type
amphi
pseudohypo
Measurement
high
low

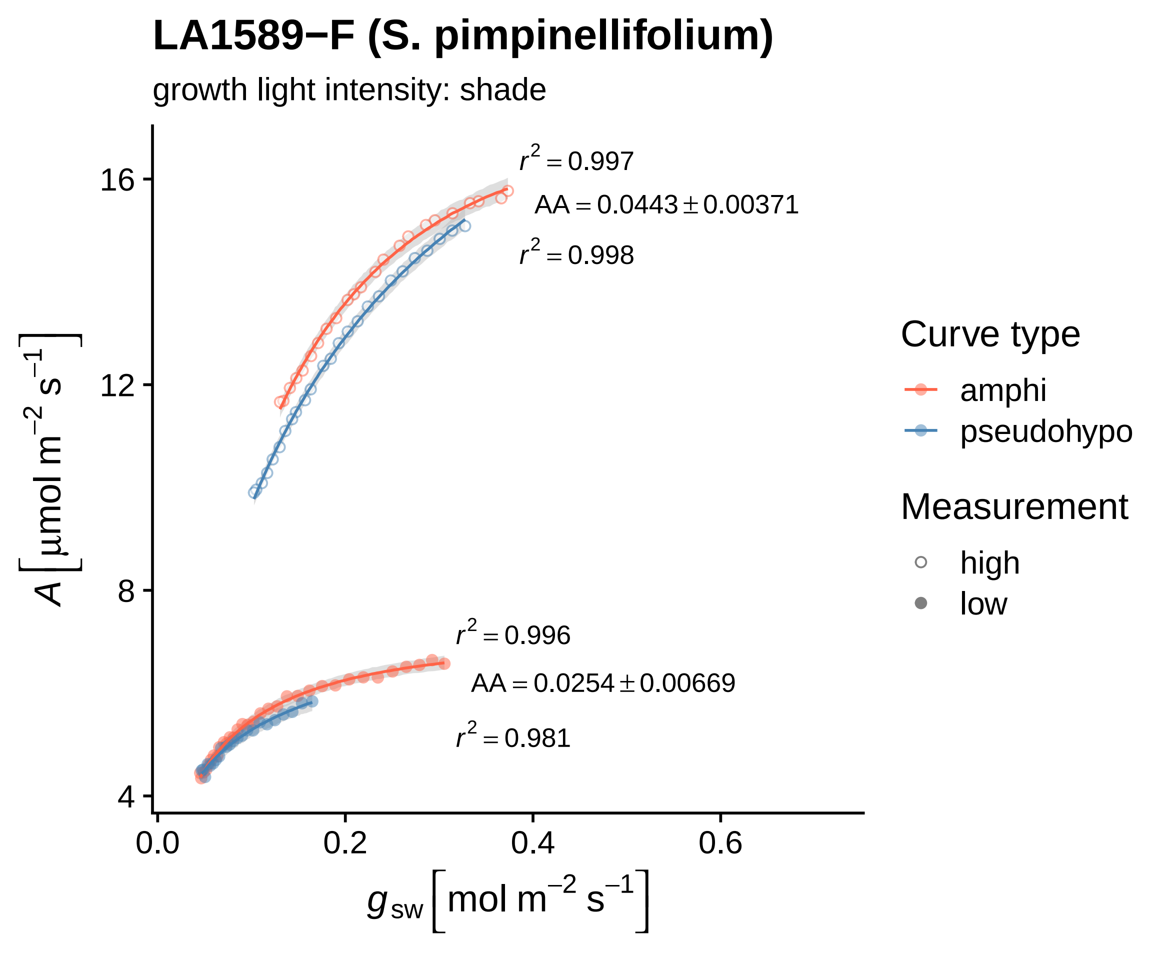
LA1589−F (S. pimpinellifolium)
growth light intensity: shade
$r^2 = 0.997$
$AA = 0.0443 \pm 0.00371$
$r^2 = 0.998$
$r^2 = 0.996$
$AA = 0.0254 \pm 0.00669$
$r^2 = 0.981$
16
12
8
4
0.0
0.2
0.4
0.6
$A\ [\mu mol\ m^{-2}\ s^{-1}]$
$g_{sw}\ [mol\ m^{-2}\ s^{-1}]$
Curve type
amphi
pseudohypo
Measurement
high
low

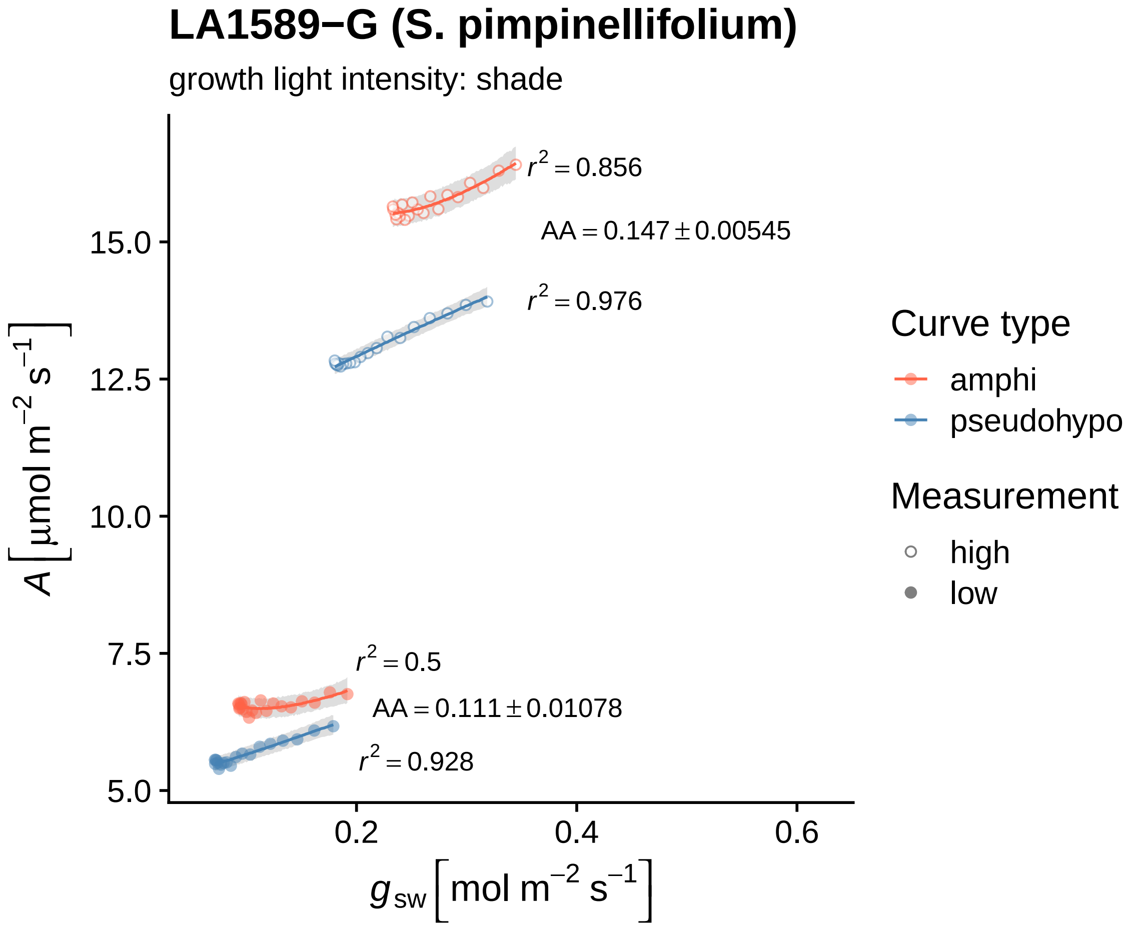
LA1589−G (S. pimpinellifolium)
growth light intensity: shade
$r^2 = 0.856$
$AA = 0.147 \pm 0.00545$
$r^2 = 0.976$
$r^2 = 0.5$
$AA = 0.111 \pm 0.01078$
$r^2 = 0.928$
15.0
12.5
10.0
7.5
5.0
0.2
0.4
0.6
$A\ [\mu mol\ m^{-2}\ s^{-1}]$
$g_{sw}\ [mol\ m^{-2}\ s^{-1}]$
Curve type
amphi
pseudohypo
Measurement
high
low

### LA1589−I (S. pimpinellifolium)

growth light intensity: sun

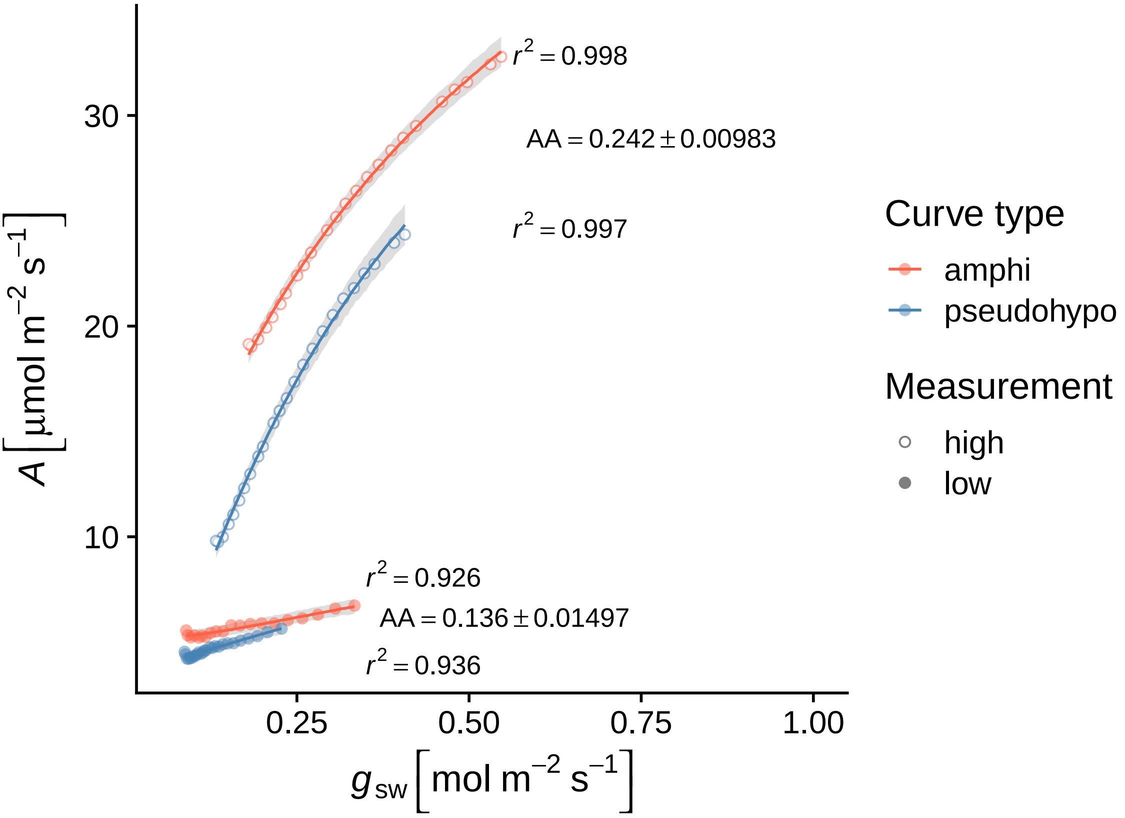

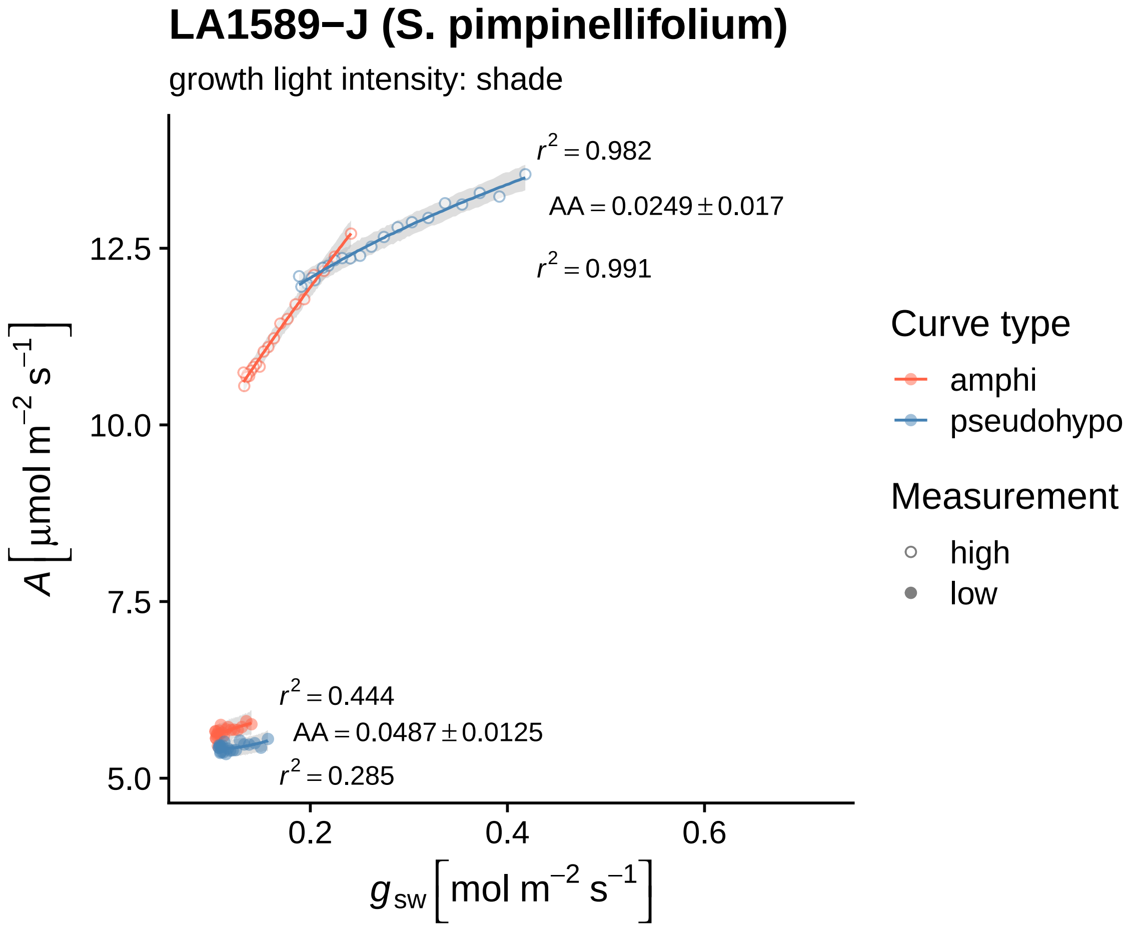
LA1589−J (S. pimpinellifolium)
growth light intensity: shade
$r^2 = 0.982$
$AA = 0.0249 \pm 0.017$
$r^2 = 0.991$
$r^2 = 0.444$
$AA = 0.0487 \pm 0.0125$
$r^2 = 0.285$
12.5
10.0
7.5
5.0
0.2
0.4
0.6
$A\ [\mu mol\ m^{-2}\ s^{-1}]$
$g_{sw}\ [mol\ m^{-2}\ s^{-1}]$
Curve type
amphi
pseudohypo
Measurement
high
low

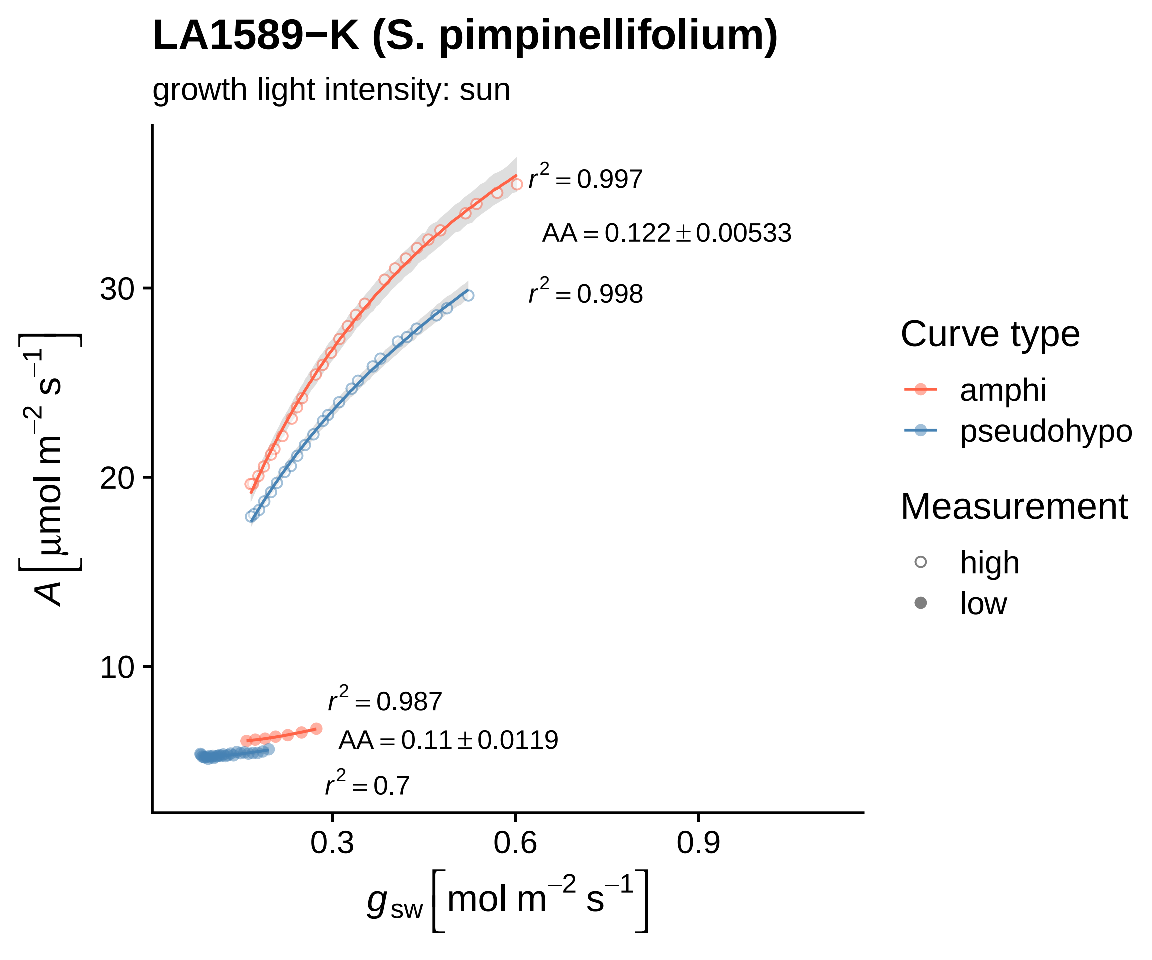

LA1589−K (S. pimpinellifolium)
growth light intensity: sun
r2 = 0.997
AA = 0.122 ± 0.00533
r2 = 0.998
r2 = 0.987
AA = 0.11 ± 0.0119
r2 = 0.7
10
20
30
0.3
0.6
0.9
A [μmol m−2 s−1]
gsw [mol m−2 s−1]
Curve type
amphi
pseudohypo
Measurement
high
low

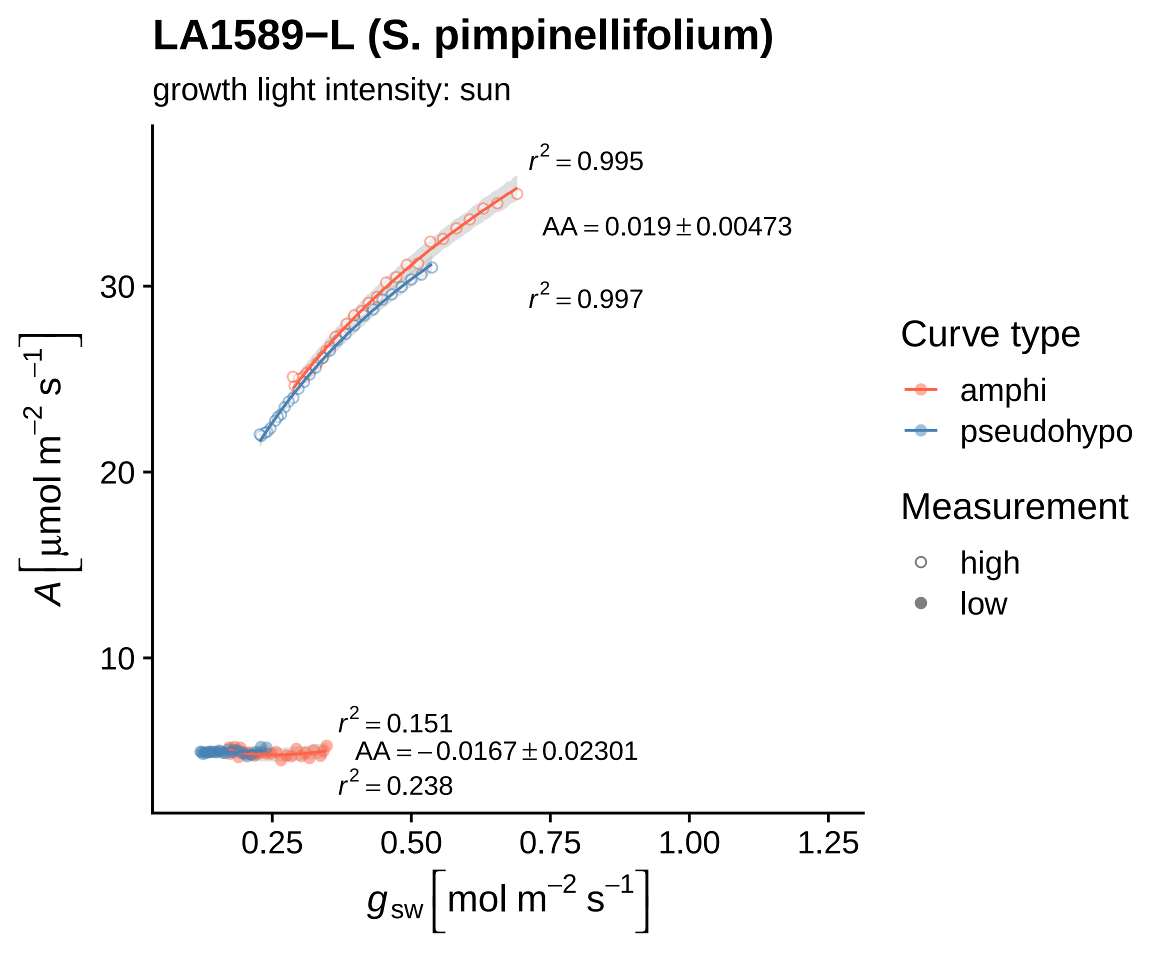

LA1589−L (S. pimpinellifolium)
growth light intensity: sun
r² = 0.995
AA = 0.019 ± 0.00473
r² = 0.997
r² = 0.151
AA = −0.0167 ± 0.02301
r² = 0.238
30
20
10
0.25
0.50
0.75
1.00
1.25
A [µmol m⁻² s⁻¹]
g_sw [mol m⁻² s⁻¹]
Curve type
amphi
pseudohypo
Measurement
high
low

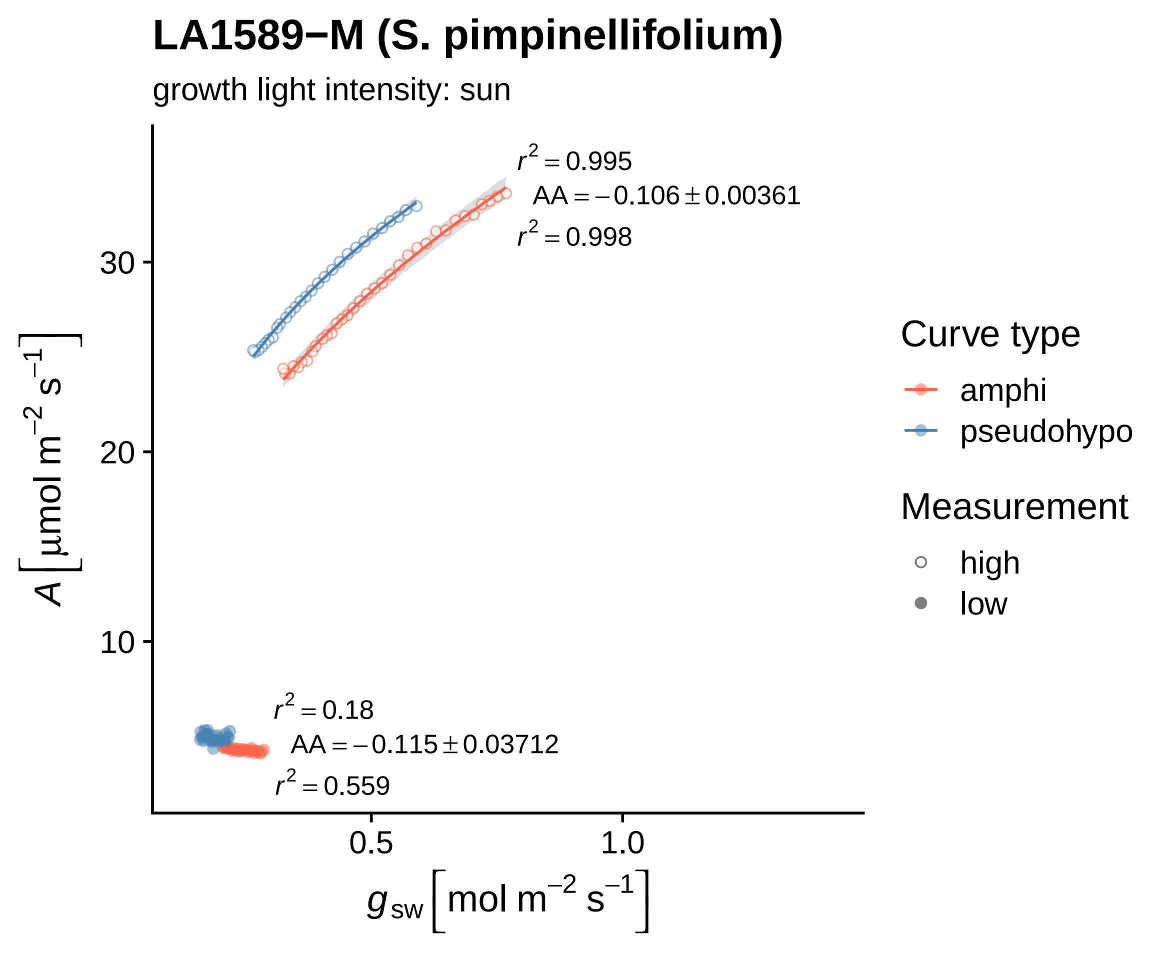

LA1589−M (S. pimpinellifolium)
growth light intensity: sun
$r^2 = 0.995$
$AA = -0.106 \pm 0.00361$
$r^2 = 0.998$
$r^2 = 0.18$
$AA = -0.115 \pm 0.03712$
$r^2 = 0.559$
30
20
10
0.5
1.0
$A\left[\mu mol\ m^{-2}\ s^{-1}\right]$
$g_{sw}\left[mol\ m^{-2}\ s^{-1}\right]$
Curve type
amphi
pseudohypo
Measurement
high
low

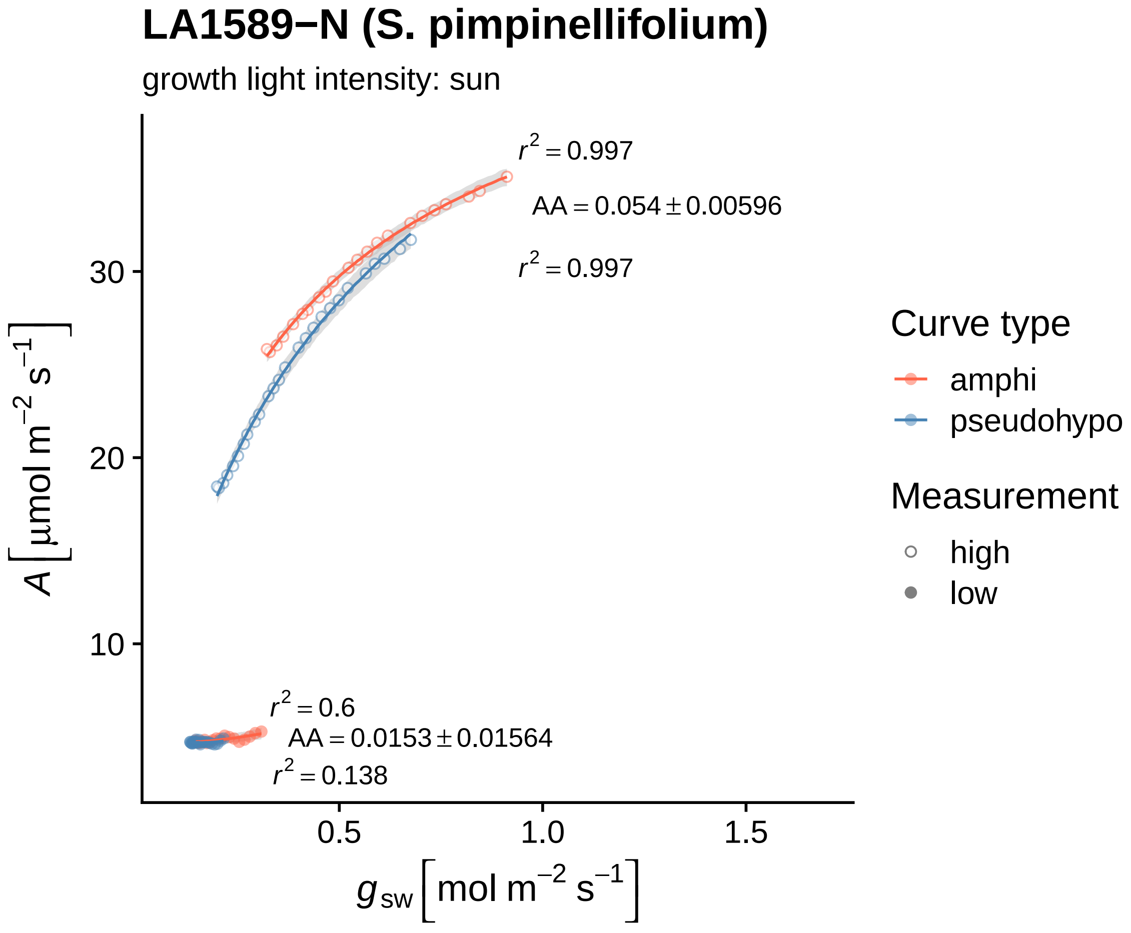
LA1589−N (S. pimpinellifolium)
growth light intensity: sun
r² = 0.997
AA = 0.054 ± 0.00596
r² = 0.997
r² = 0.6
AA = 0.0153 ± 0.01564
r² = 0.138
A [μmol m⁻² s⁻¹]
30
20
10
0.5
1.0
1.5
g_sw [mol m⁻² s⁻¹]
Curve type
amphi
pseudohypo
Measurement
high
low

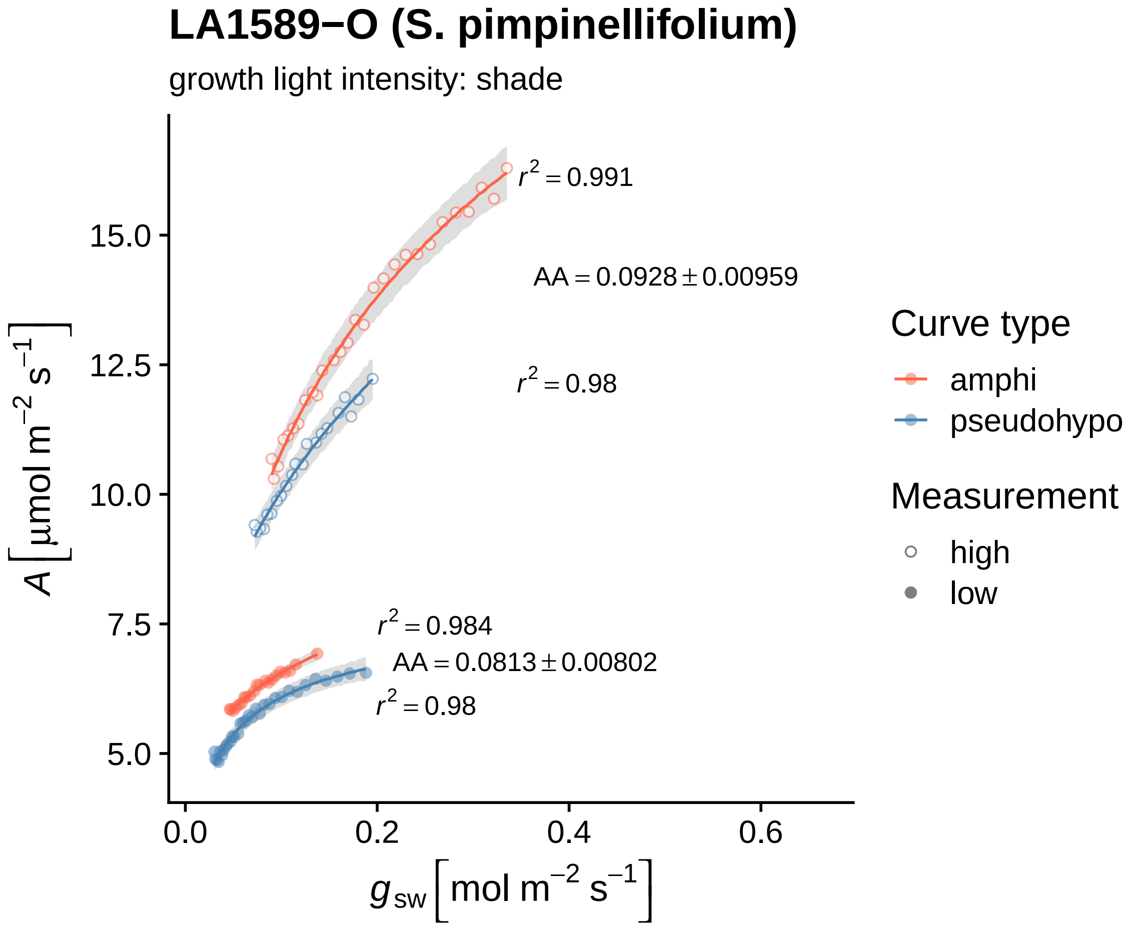
LA1589−O (S. pimpinellifolium)
growth light intensity: shade
$r^2 = 0.991$
$AA = 0.0928 \pm 0.00959$
$r^2 = 0.98$
$r^2 = 0.984$
$AA = 0.0813 \pm 0.00802$
$r^2 = 0.98$
15.0
12.5
10.0
7.5
5.0
0.0
0.2
0.4
0.6
$A\left[\mu mol\ m^{-2}\ s^{-1}\right]$
$g_{sw}\left[mol\ m^{-2}\ s^{-1}\right]$
Curve type
amphi
pseudohypo
Measurement
high
low

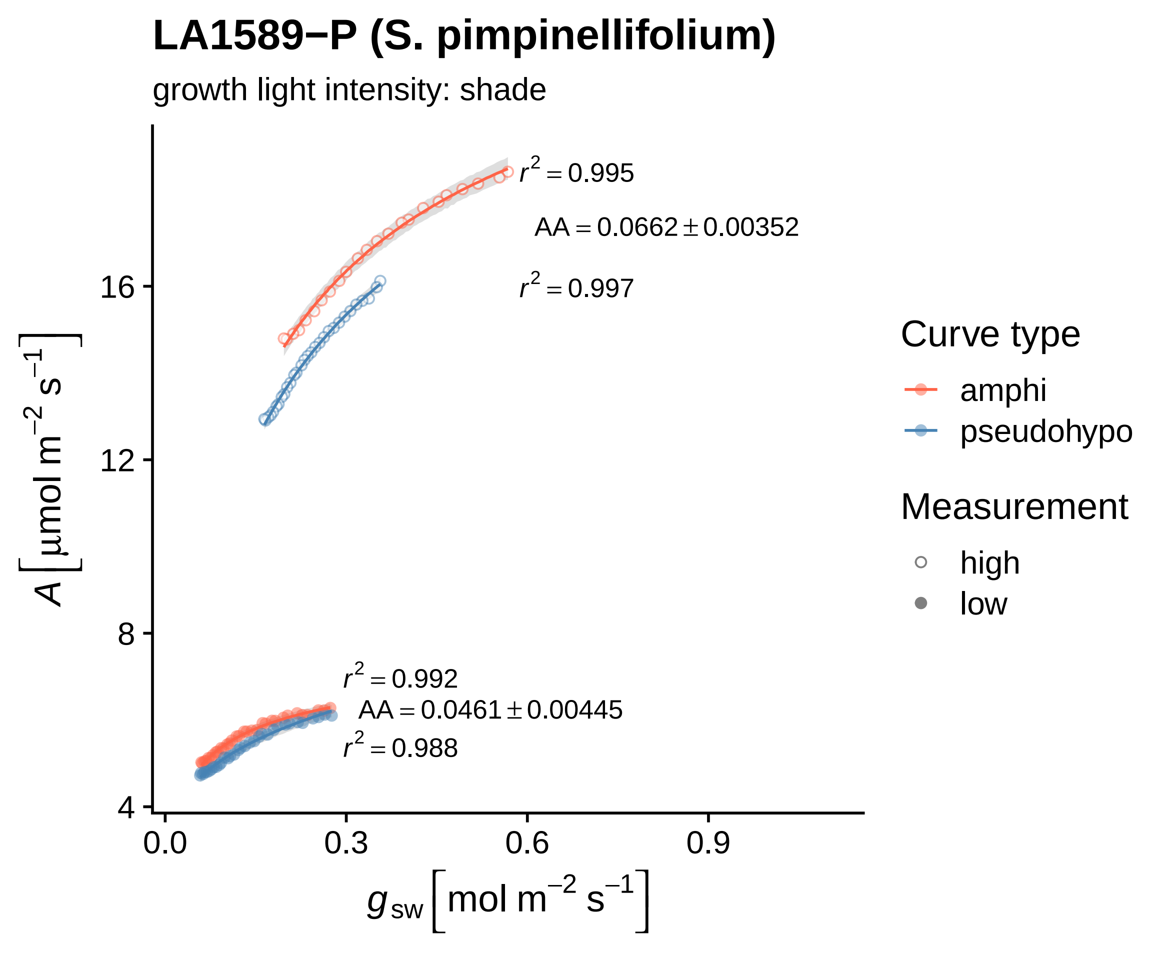

LA1589−P (S. pimpinellifolium)
growth light intensity: shade
$r^2 = 0.995$
$AA = 0.0662 \pm 0.00352$
$r^2 = 0.997$
$r^2 = 0.992$
$AA = 0.0461 \pm 0.00445$
$r^2 = 0.988$
16
12
8
4
0.0
0.3
0.6
0.9
$A \left[\mu mol\ m^{-2}\ s^{-1}\right]$
$g_{sw} \left[mol\ m^{-2}\ s^{-1}\right]$
Curve type
amphi
pseudohypo
Measurement
high
low

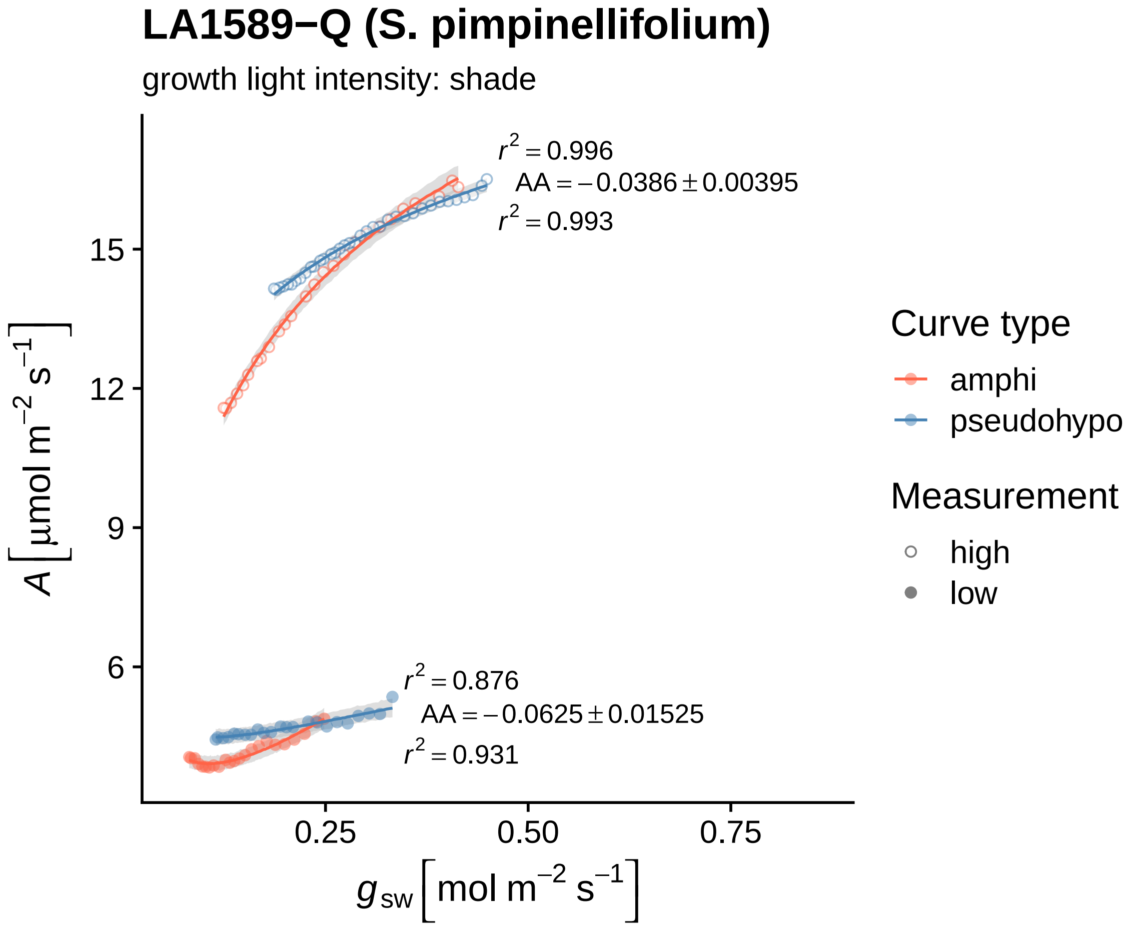
LA1589−Q (S. pimpinellifolium)
growth light intensity: shade
r^2 = 0.996
AA = −0.0386 ± 0.00395
r^2 = 0.993
r^2 = 0.876
AA = −0.0625 ± 0.01525
r^2 = 0.931
15
12
9
6
0.25
0.50
0.75
A [µmol m^−2 s^−1]
g_sw [mol m^−2 s^−1]
Curve type
amphi
pseudohypo
Measurement
high
low

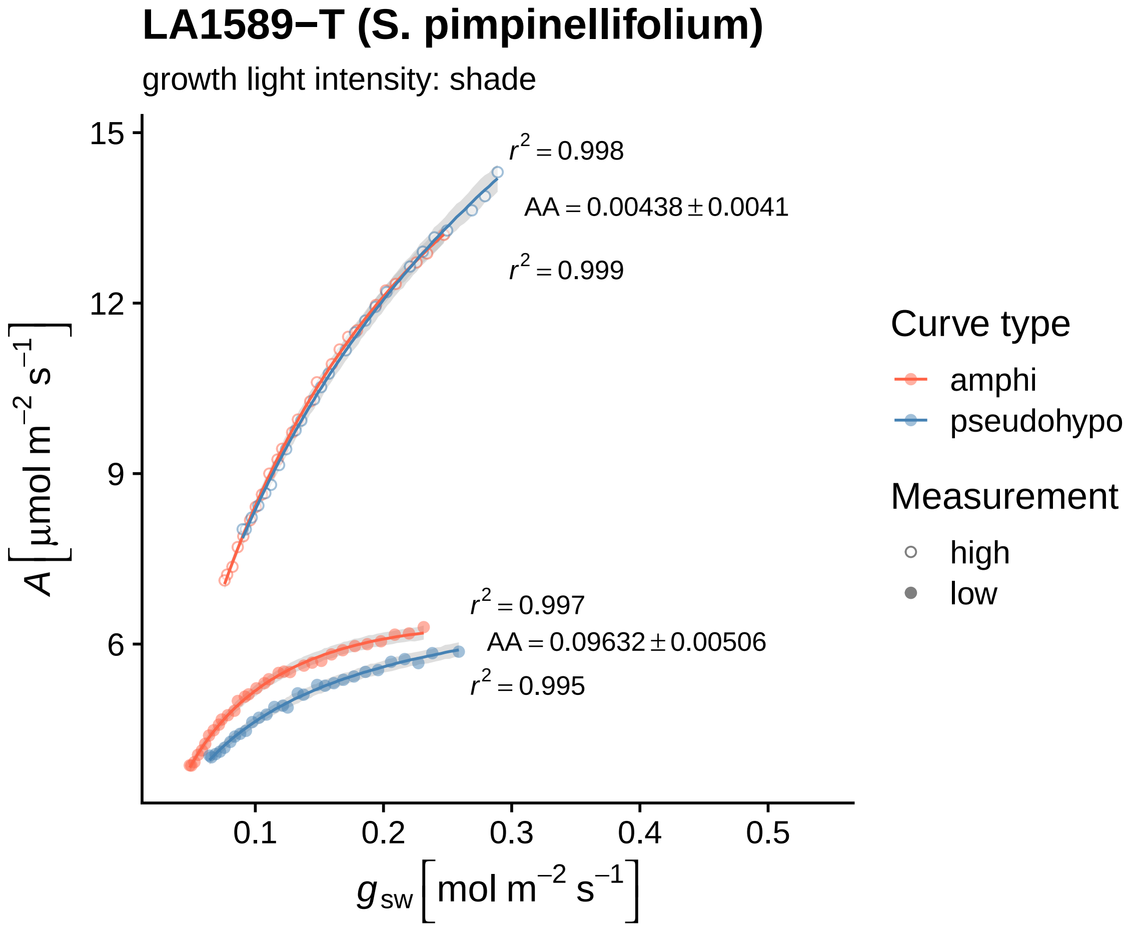
LA1589−T (S. pimpinellifolium)
growth light intensity: shade
r² = 0.998
AA = 0.00438 ± 0.0041
r² = 0.999
r² = 0.997
AA = 0.09632 ± 0.00506
r² = 0.995
15
12
9
6
0.1
0.2
0.3
0.4
0.5
A [μmol m⁻² s⁻¹]
g sw [mol m⁻² s⁻¹]
Curve type
amphi
pseudohypo
Measurement
high
low

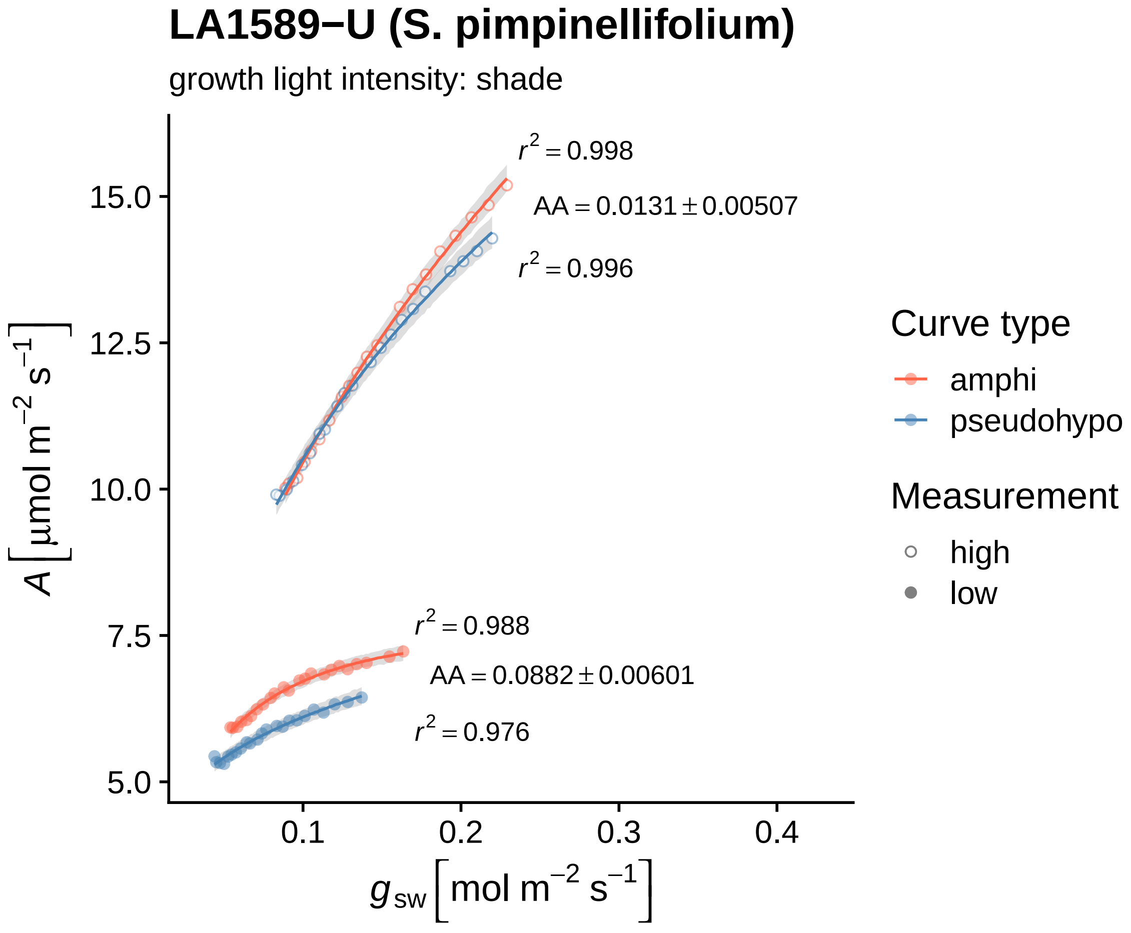

LA1589−U (S. pimpinellifolium)
growth light intensity: shade
$r^2 = 0.998$
$AA = 0.0131 \pm 0.00507$
$r^2 = 0.996$
$r^2 = 0.988$
$AA = 0.0882 \pm 0.00601$
$r^2 = 0.976$
15.0
12.5
10.0
7.5
5.0
0.1
0.2
0.3
0.4
$A\left[\mu mol\ m^{-2}\ s^{-1}\right]$
$g_{sw}\left[mol\ m^{-2}\ s^{-1}\right]$
Curve type
amphi
pseudohypo
Measurement
high
low

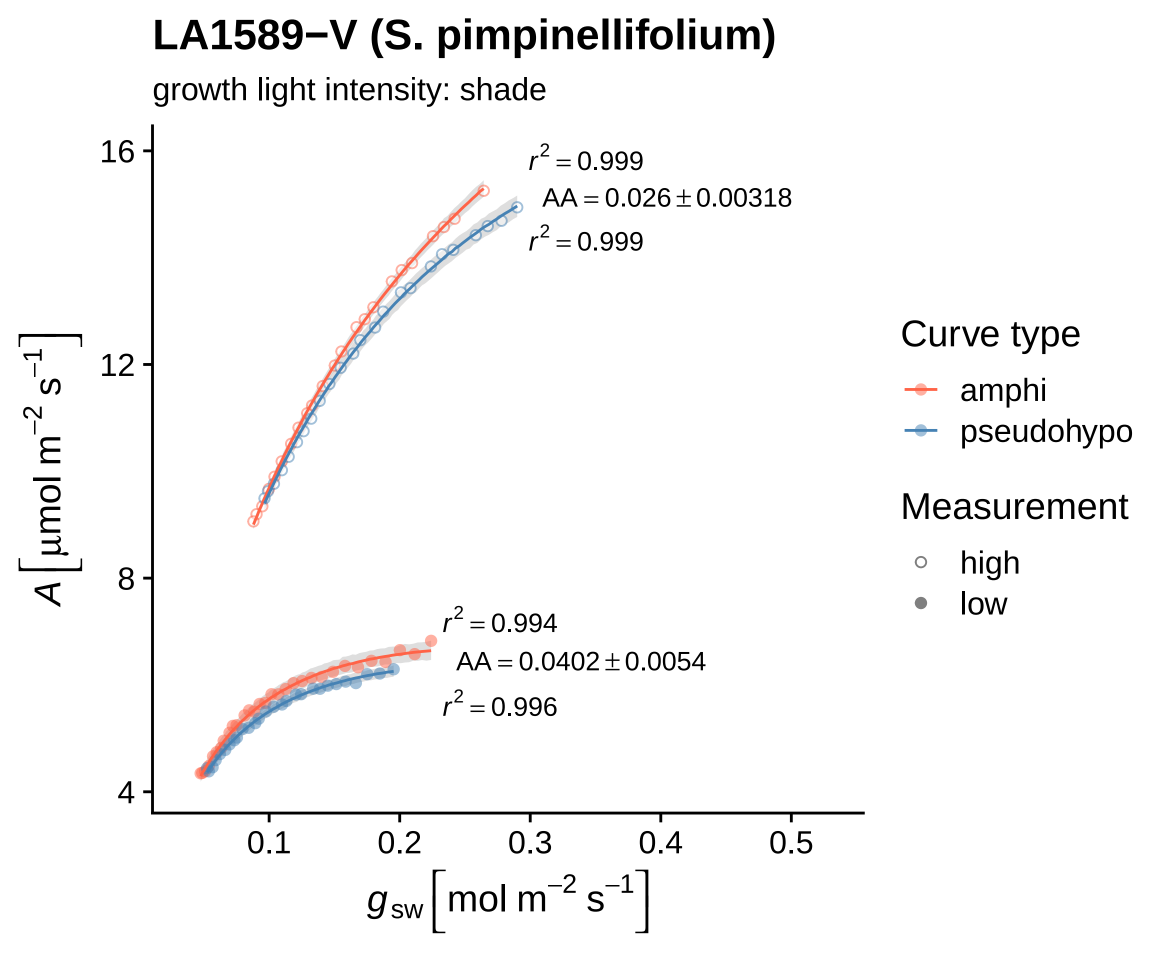

LA1589−V (S. pimpinellifolium)
growth light intensity: shade
r² = 0.999
AA = 0.026 ± 0.00318
r² = 0.999
r² = 0.994
AA = 0.0402 ± 0.0054
r² = 0.996
16
12
8
4
0.1
0.2
0.3
0.4
0.5
A [μmol m⁻² s⁻¹]
gsw [mol m⁻² s⁻¹]
Curve type
amphi
pseudohypo
Measurement
high
low

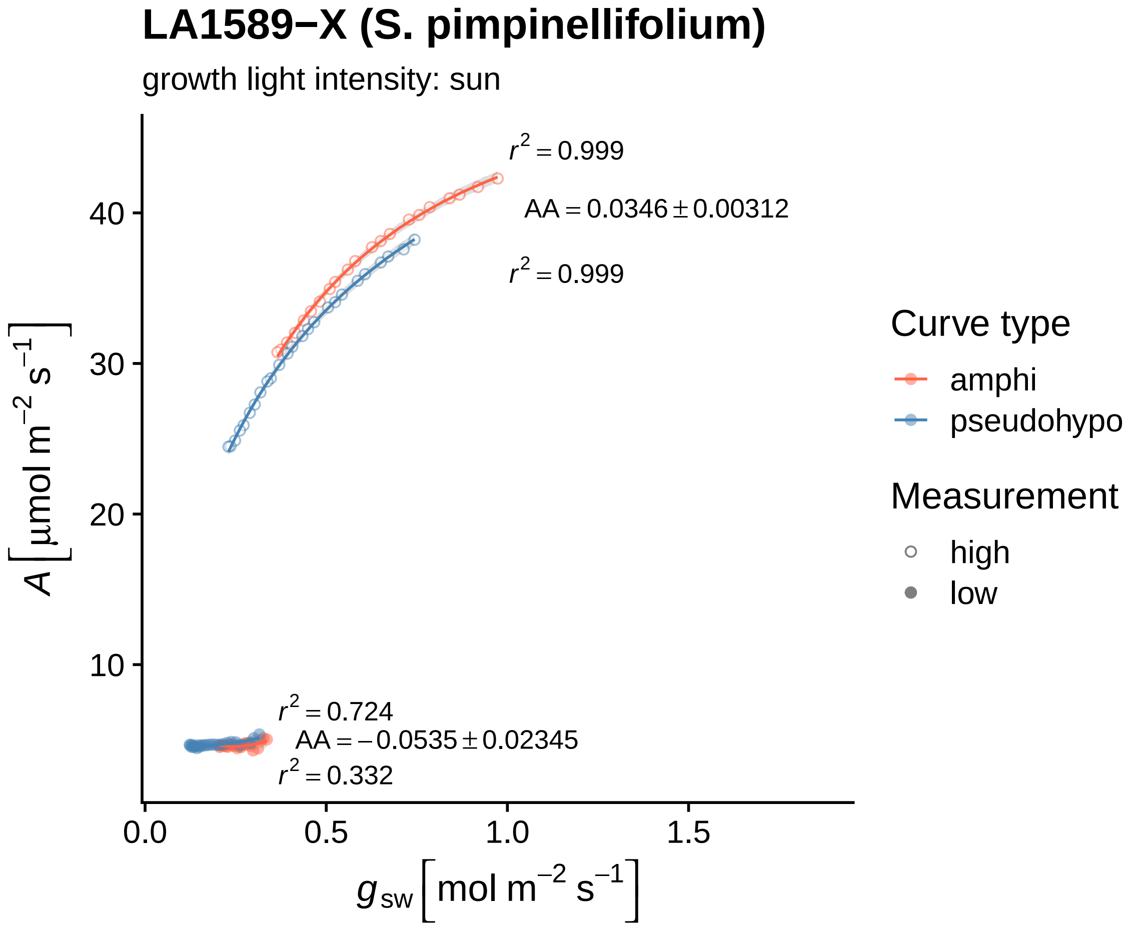

LA1589−X (S. pimpinellifolium)
growth light intensity: sun
$r^2 = 0.999$
$AA = 0.0346 \pm 0.00312$
$r^2 = 0.999$
$r^2 = 0.724$
$AA = -0.0535 \pm 0.02345$
$r^2 = 0.332$
40
30
20
10
0.0
0.5
1.0
1.5
$A\ [\mu mol\ m^{-2}\ s^{-1}]$
$g_{sw}\ [mol\ m^{-2}\ s^{-1}]$
Curve type
amphi
pseudohypo
Measurement
high
low

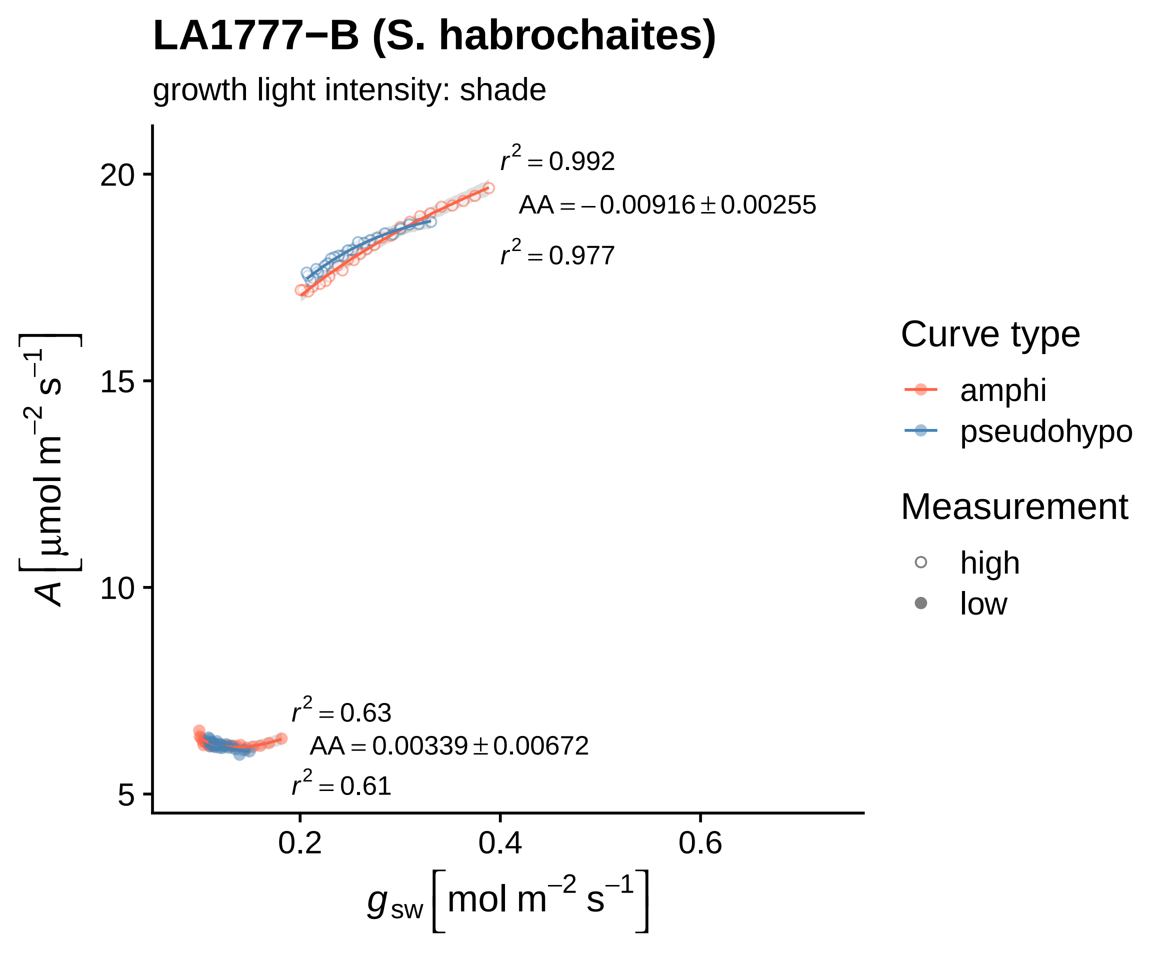

LA1777−B (S. habrochaites)
growth light intensity: shade
$r^2 = 0.992$
$AA = -0.00916 \pm 0.00255$
$r^2 = 0.977$
$r^2 = 0.63$
$AA = 0.00339 \pm 0.00672$
$r^2 = 0.61$
20
15
10
5
0.2
0.4
0.6
$A\ [\mu mol\ m^{-2}\ s^{-1}]$
$g_{sw}\ [mol\ m^{-2}\ s^{-1}]$
Curve type
amphi
pseudohypo
Measurement
high
low

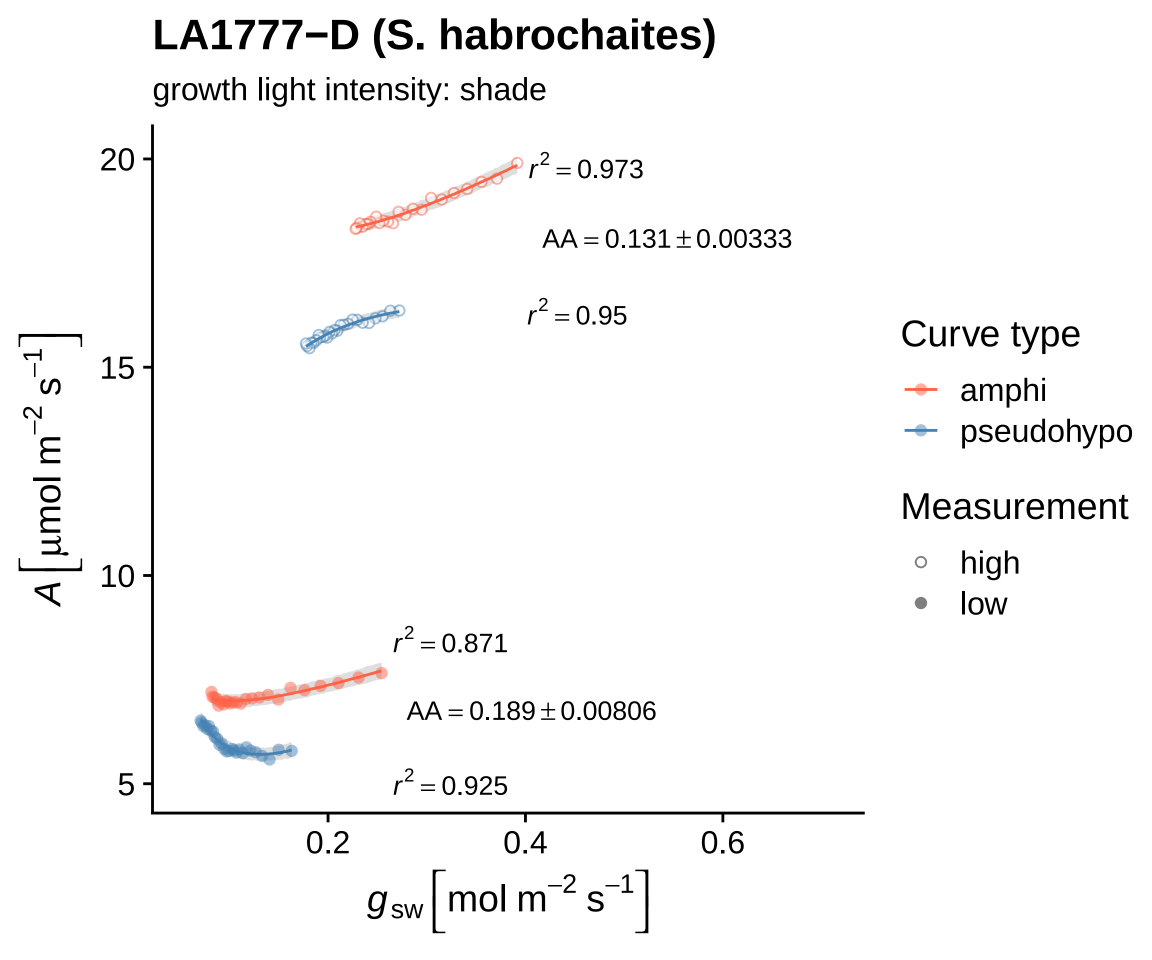
LA1777−D (S. habrochaites)
growth light intensity: shade
$r^2 = 0.973$
$AA = 0.131 \pm 0.00333$
$r^2 = 0.95$
$r^2 = 0.871$
$AA = 0.189 \pm 0.00806$
$r^2 = 0.925$
20
15
10
5
0.2
0.4
0.6
$A\ [\mu mol\ m^{-2}\ s^{-1}]$
$g_{sw}\ [mol\ m^{-2}\ s^{-1}]$
Curve type
amphi
pseudohypo
Measurement
high
low

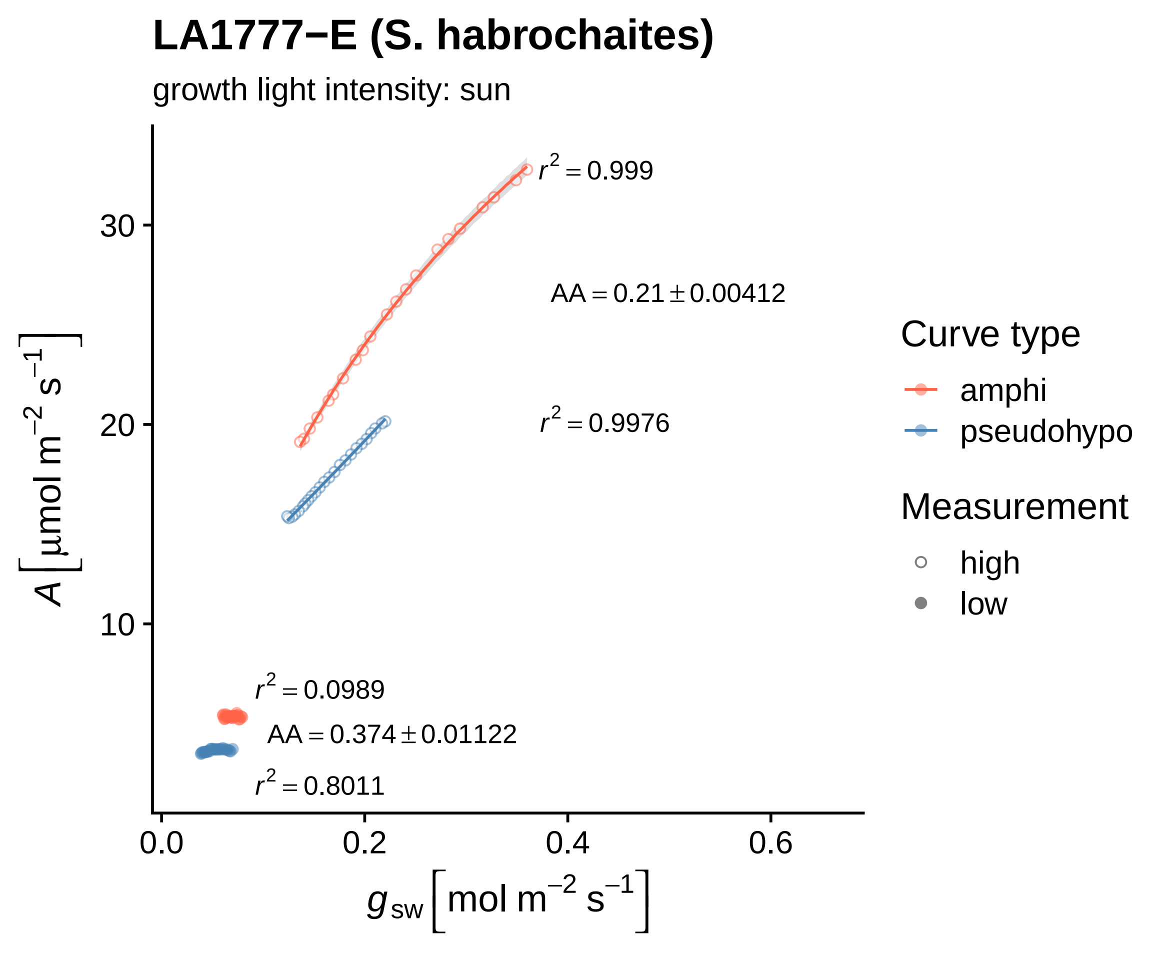

LA1777−E (S. habrochaites)
growth light intensity: sun
$r^2 = 0.999$
$AA = 0.21 \pm 0.00412$
$r^2 = 0.9976$
$r^2 = 0.0989$
$AA = 0.374 \pm 0.01122$
$r^2 = 0.8011$
30
20
10
0.0
0.2
0.4
0.6
$A\left[\mu mol\ m^{-2}\ s^{-1}\right]$
$g_{sw}\left[mol\ m^{-2}\ s^{-1}\right]$
Curve type
amphi
pseudohypo
Measurement
high
low

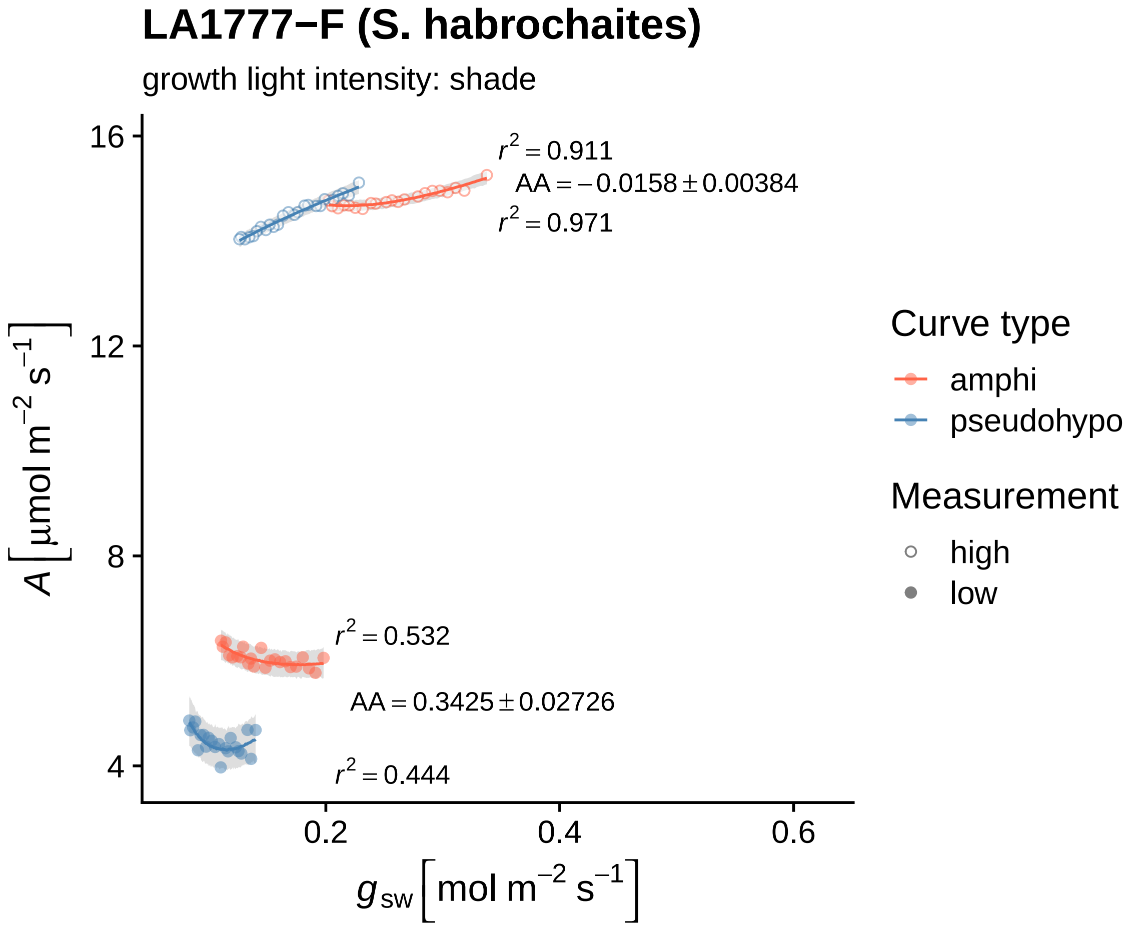

LA1777−F (S. habrochaites)
growth light intensity: shade
$r^2 = 0.911$
$AA = -0.0158 \pm 0.00384$
$r^2 = 0.971$
$r^2 = 0.532$
$AA = 0.3425 \pm 0.02726$
$r^2 = 0.444$
$A\ [\mu mol\ m^{-2}\ s^{-1}]$
16
12
8
4
0.2
0.4
0.6
$g_{sw}\ [mol\ m^{-2}\ s^{-1}]$
Curve type
amphi
pseudohypo
Measurement
high
low

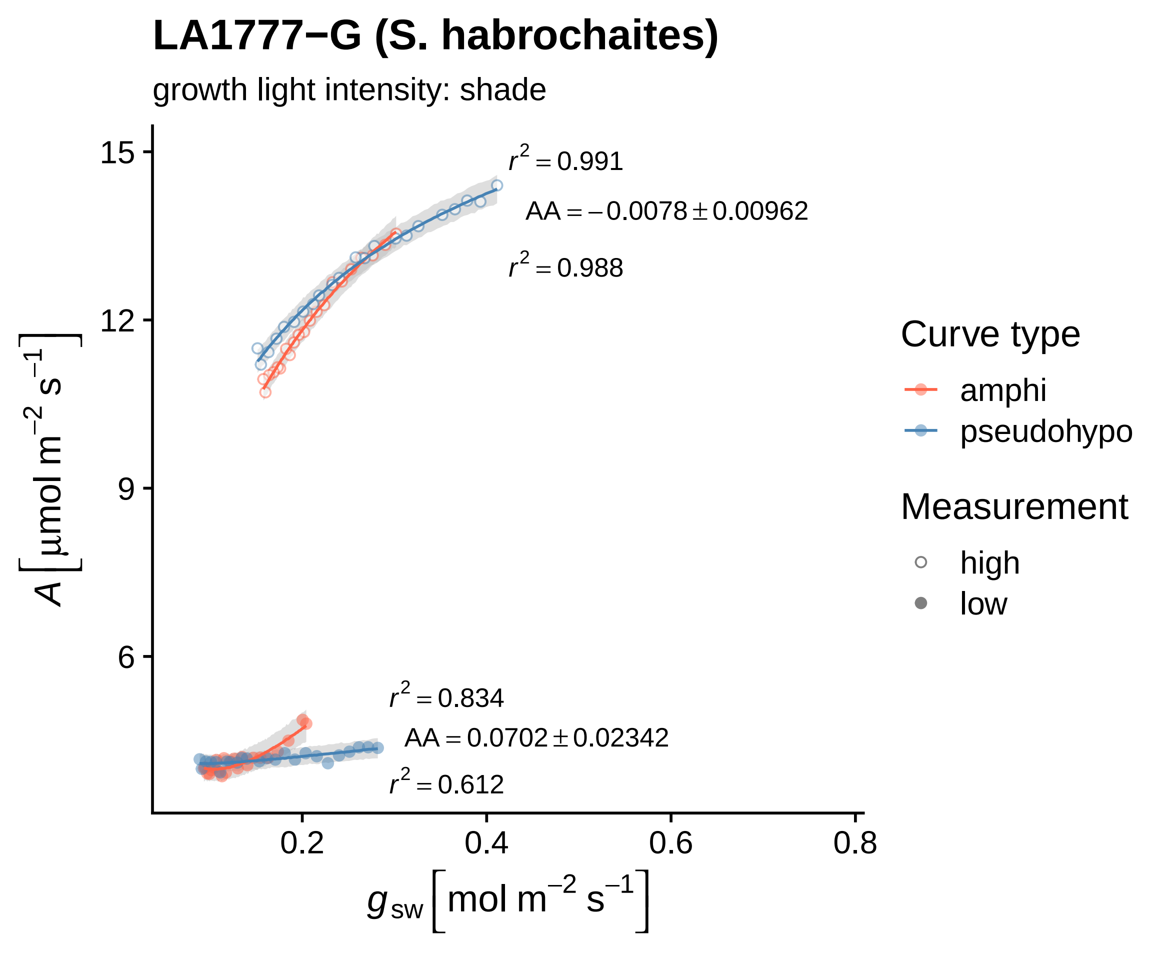
LA1777−G (S. habrochaites)
growth light intensity: shade
$r^2 = 0.991$
$AA = -0.0078 \pm 0.00962$
$r^2 = 0.988$
$r^2 = 0.834$
$AA = 0.0702 \pm 0.02342$
$r^2 = 0.612$
15
12
9
6
0.2
0.4
0.6
0.8
$A\ [\mu mol\ m^{-2}\ s^{-1}]$
$g_{sw}\ [mol\ m^{-2}\ s^{-1}]$
Curve type
amphi
pseudohypo
Measurement
high
low

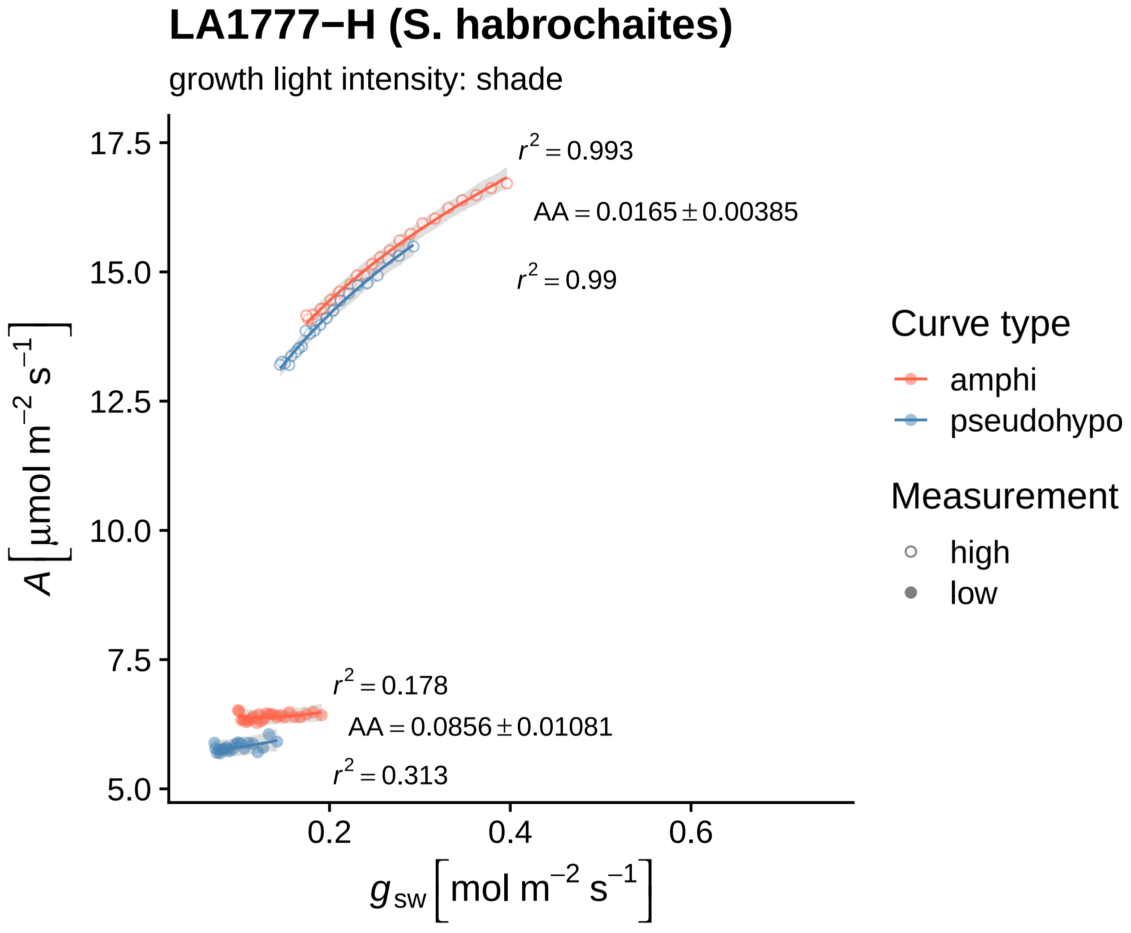

LA1777−H (S. habrochaites)
growth light intensity: shade
r^2 = 0.993
AA = 0.0165 ± 0.00385
r^2 = 0.99
r^2 = 0.178
AA = 0.0856 ± 0.01081
r^2 = 0.313
17.5
15.0
12.5
10.0
7.5
5.0
0.2
0.4
0.6
A [μmol m^−2 s^−1]
g_sw [mol m^−2 s^−1]
Curve type
amphi
pseudohypo
Measurement
high
low

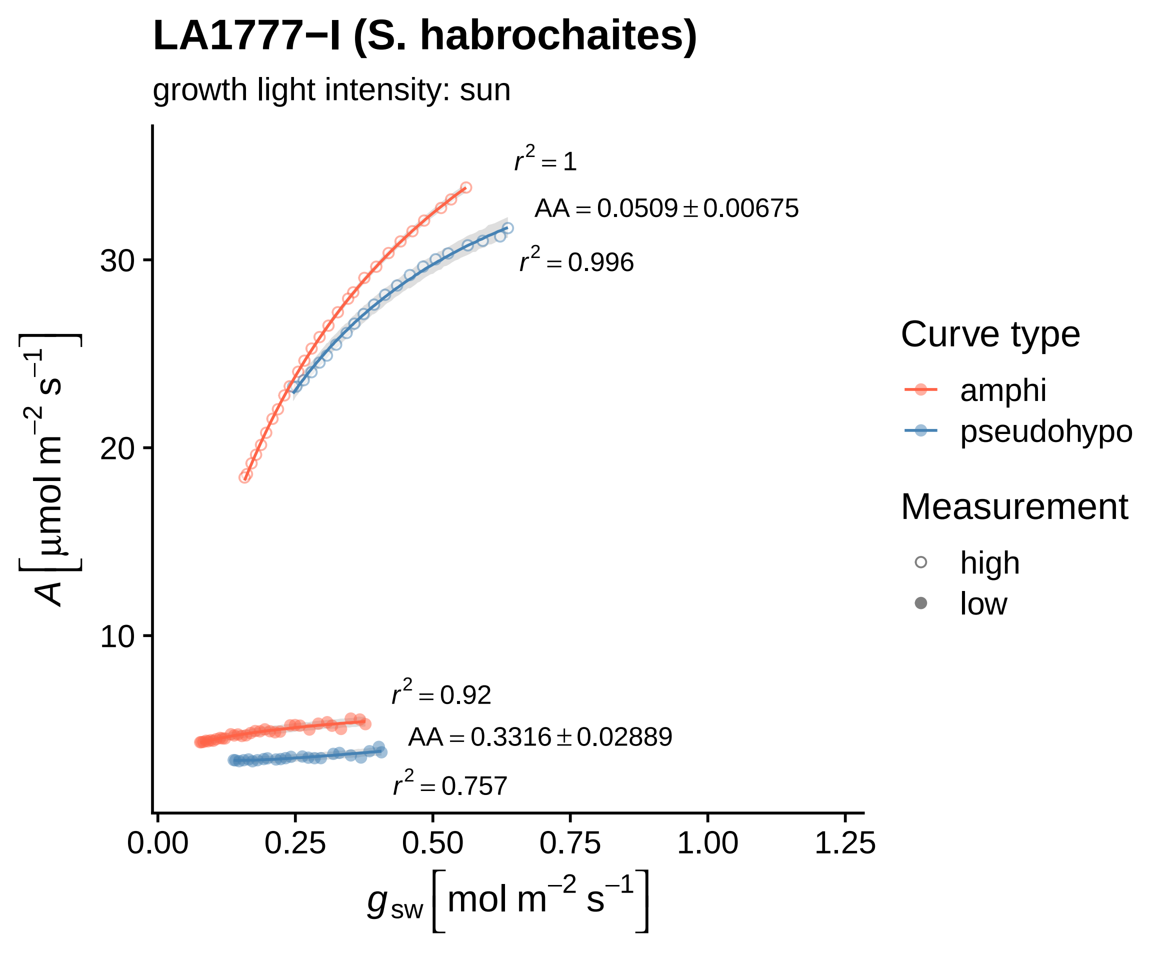

LA1777−I (S. habrochaites)
growth light intensity: sun
r² = 1
AA = 0.0509 ± 0.00675
r² = 0.996
r² = 0.92
AA = 0.3316 ± 0.02889
r² = 0.757
30
20
10
0.00
0.25
0.50
0.75
1.00
1.25
A [µmol m⁻² s⁻¹]
g_sw [mol m⁻² s⁻¹]
Curve type
amphi
pseudohypo
Measurement
high
low

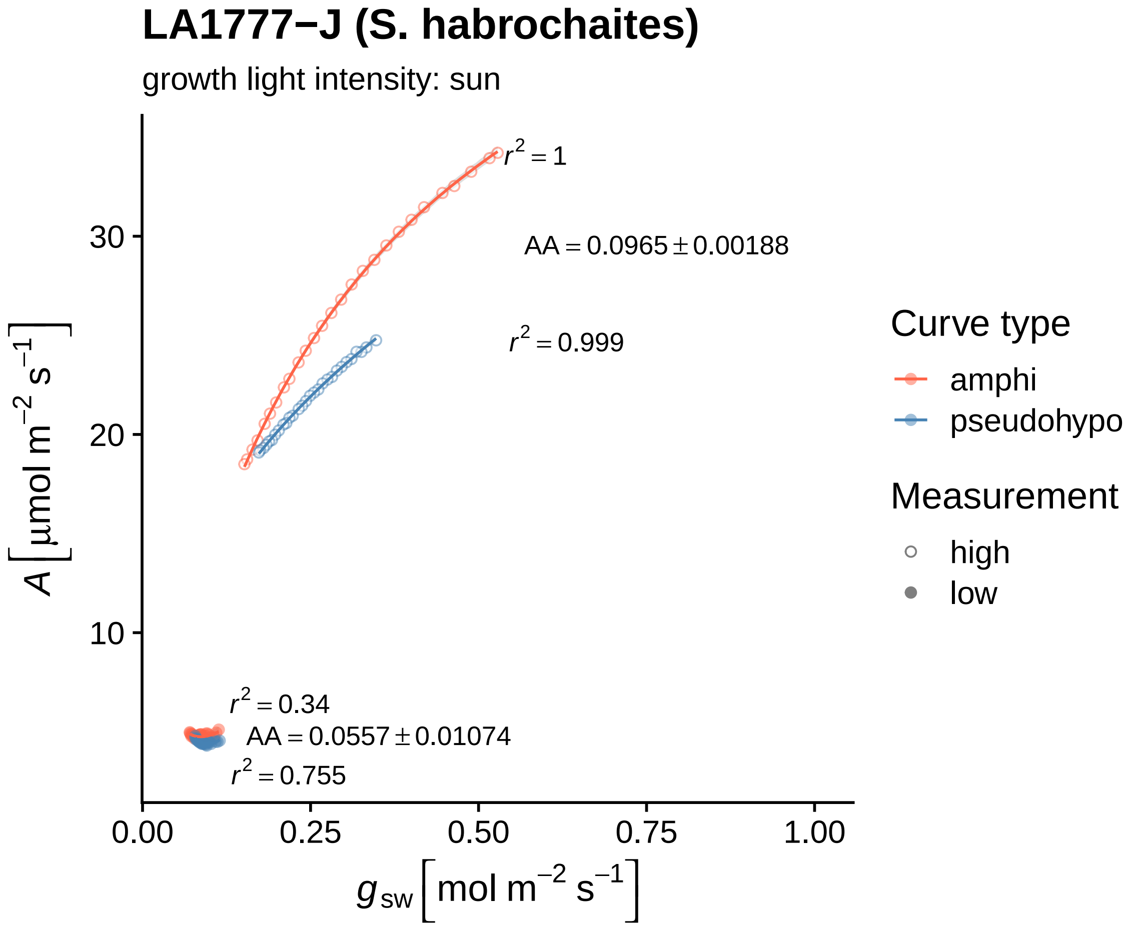

LA1777−J (S. habrochaites)
growth light intensity: sun
$r^2 = 1$
AA = 0.0965 ± 0.00188
$r^2 = 0.999$
$r^2 = 0.34$
AA = 0.0557 ± 0.01074
$r^2 = 0.755$
30
20
10
0.00
0.25
0.50
0.75
1.00
$A\ [\mu mol\ m^{-2}\ s^{-1}]$
$g_{sw}\ [mol\ m^{-2}\ s^{-1}]$
Curve type
amphi
pseudohypo
Measurement
high
low

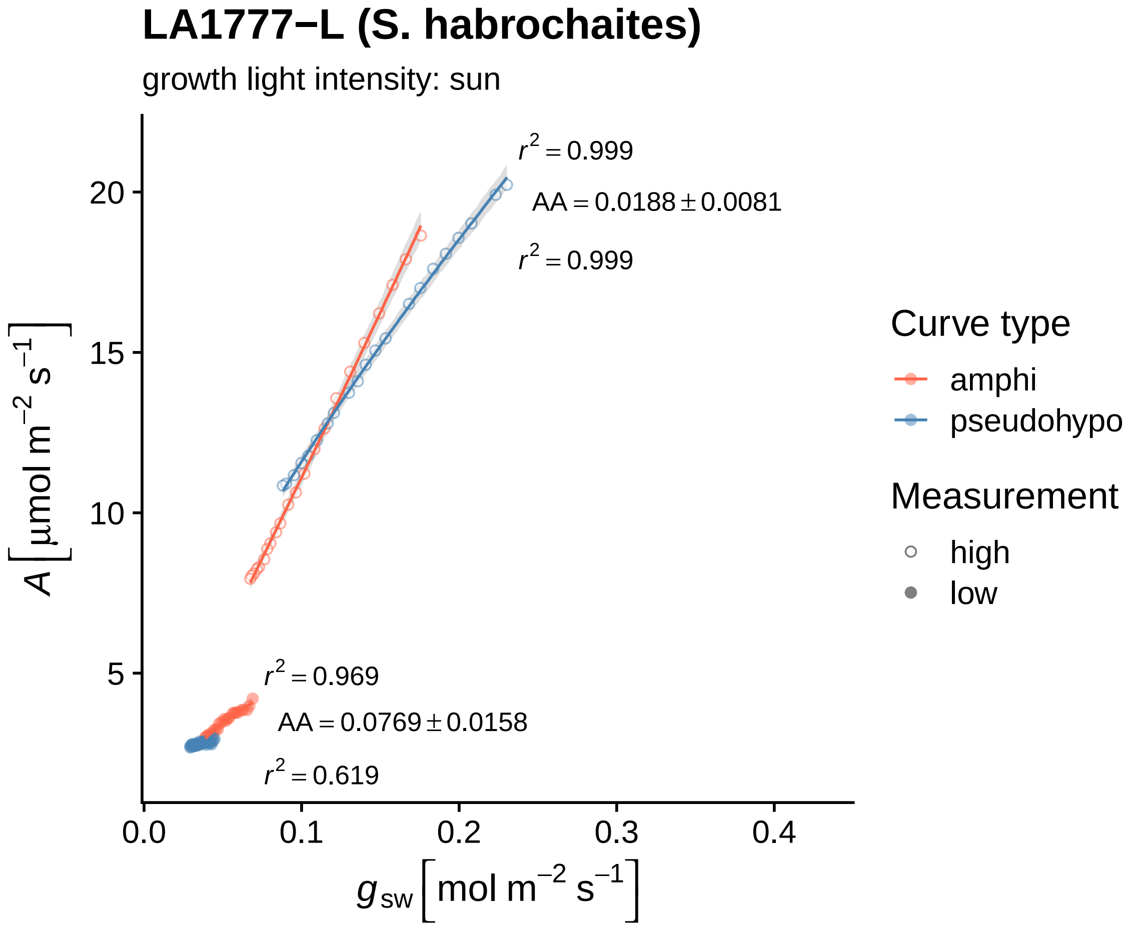

LA1777−L (S. habrochaites)
growth light intensity: sun
$r^2 = 0.999$
AA = 0.0188 ± 0.0081
$r^2 = 0.999$
$r^2 = 0.969$
AA = 0.0769 ± 0.0158
$r^2 = 0.619$
20
15
10
5
0.0
0.1
0.2
0.3
0.4
$A\ [\mu mol\ m^{-2}\ s^{-1}]$
$g_{sw}\ [mol\ m^{-2}\ s^{-1}]$
Curve type
amphi
pseudohypo
Measurement
high
low

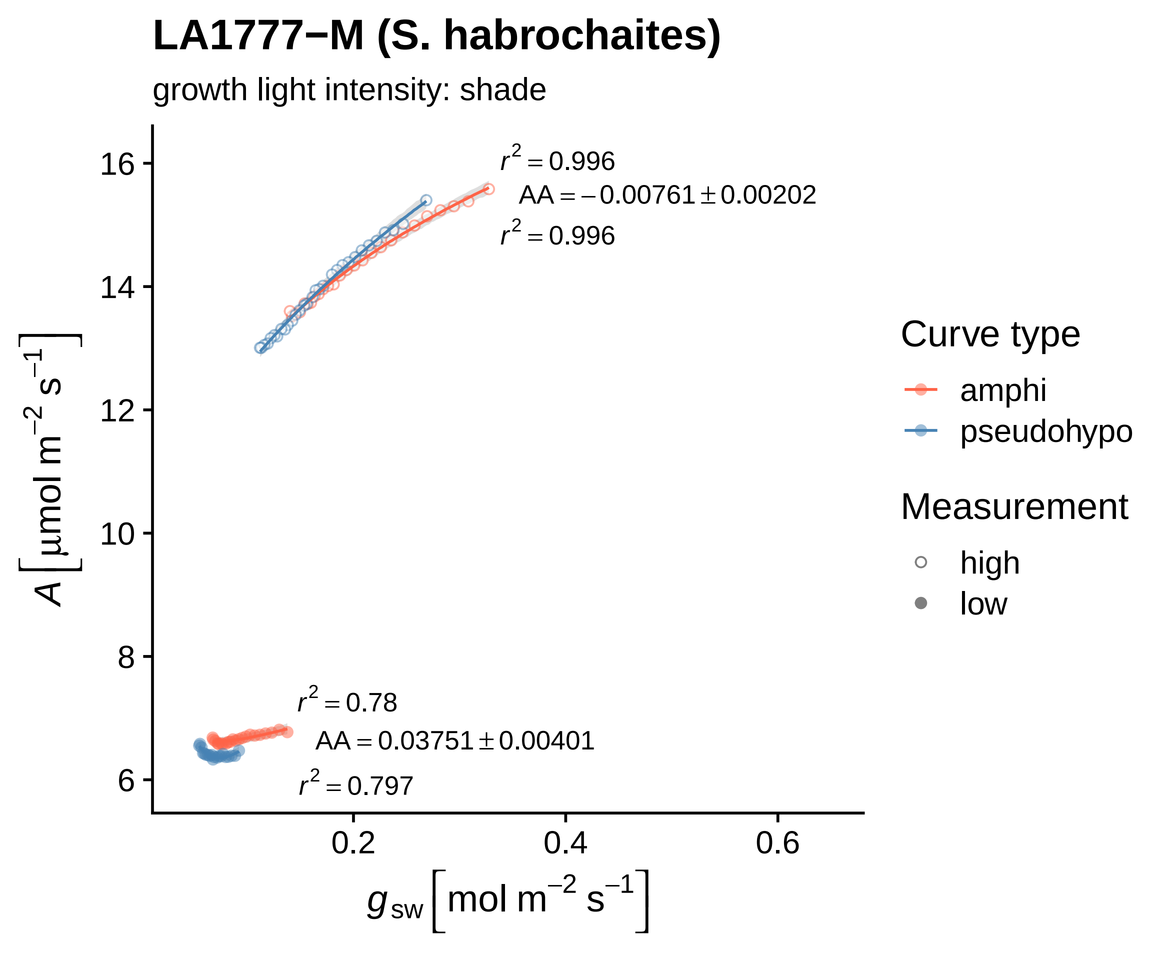

LA1777−M (S. habrochaites)
growth light intensity: shade
$r^2 = 0.996$
$AA = -0.00761 \pm 0.00202$
$r^2 = 0.996$
$r^2 = 0.78$
$AA = 0.03751 \pm 0.00401$
$r^2 = 0.797$
$A\left[\mu mol\ m^{-2}\ s^{-1}\right]$
$g_{sw}\left[mol\ m^{-2}\ s^{-1}\right]$
16
14
12
10
8
6
0.2
0.4
0.6
Curve type
amphi
pseudohypo
Measurement
high
low

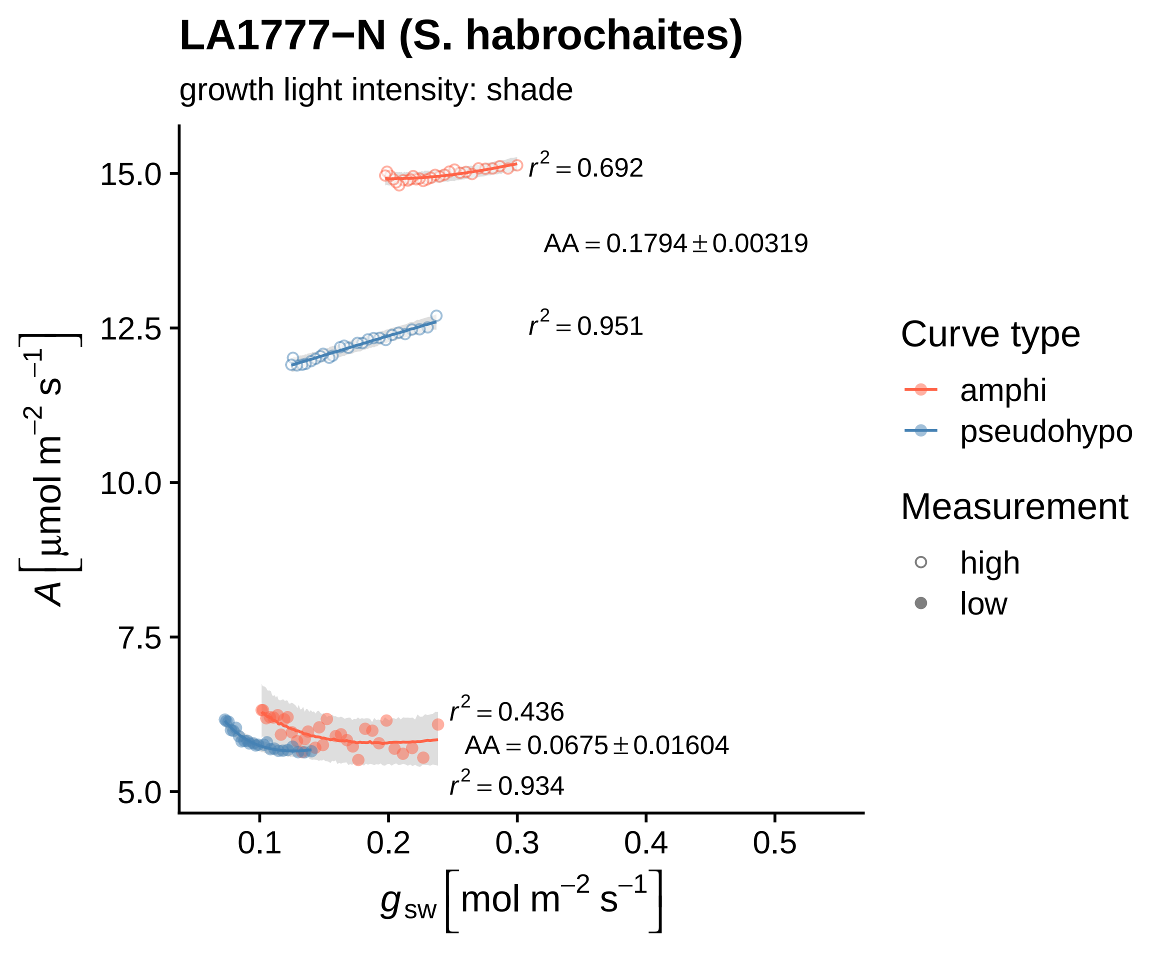
LA1777−N (S. habrochaites)
growth light intensity: shade
$r^2 = 0.692$
$AA = 0.1794 \pm 0.00319$
$r^2 = 0.951$
$r^2 = 0.436$
$AA = 0.0675 \pm 0.01604$
$r^2 = 0.934$
15.0
12.5
10.0
7.5
5.0
0.1
0.2
0.3
0.4
0.5
$A\ [\mu mol\ m^{-2}\ s^{-1}]$
$g_{sw}\ [mol\ m^{-2}\ s^{-1}]$
Curve type
amphi
pseudohypo
Measurement
high
low

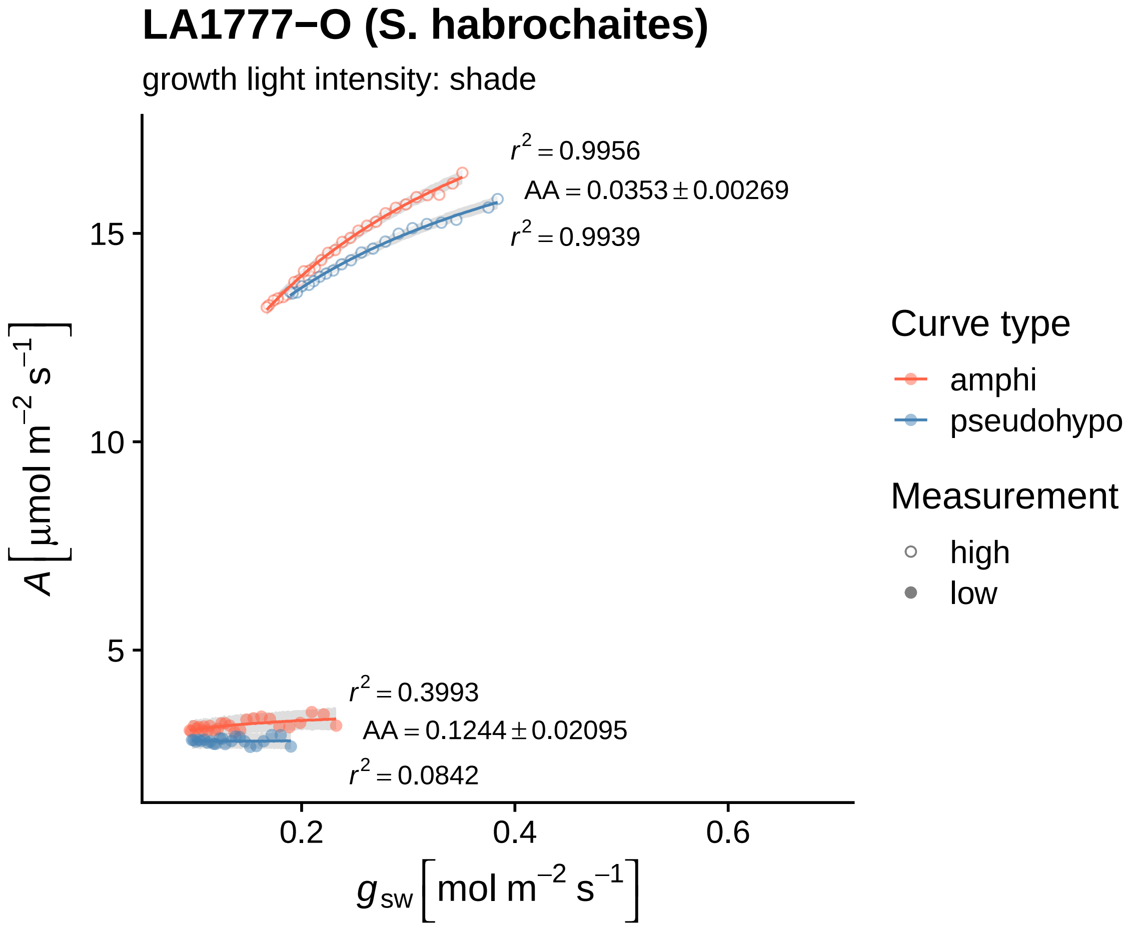

LA1777−O (S. habrochaites)
growth light intensity: shade
r² = 0.9956
AA = 0.0353 ± 0.00269
r² = 0.9939
r² = 0.3993
AA = 0.1244 ± 0.02095
r² = 0.0842
15
10
5
0.2
0.4
0.6
A [μmol m⁻² s⁻¹]
g sw [mol m⁻² s⁻¹]
Curve type
amphi
pseudohypo
Measurement
high
low

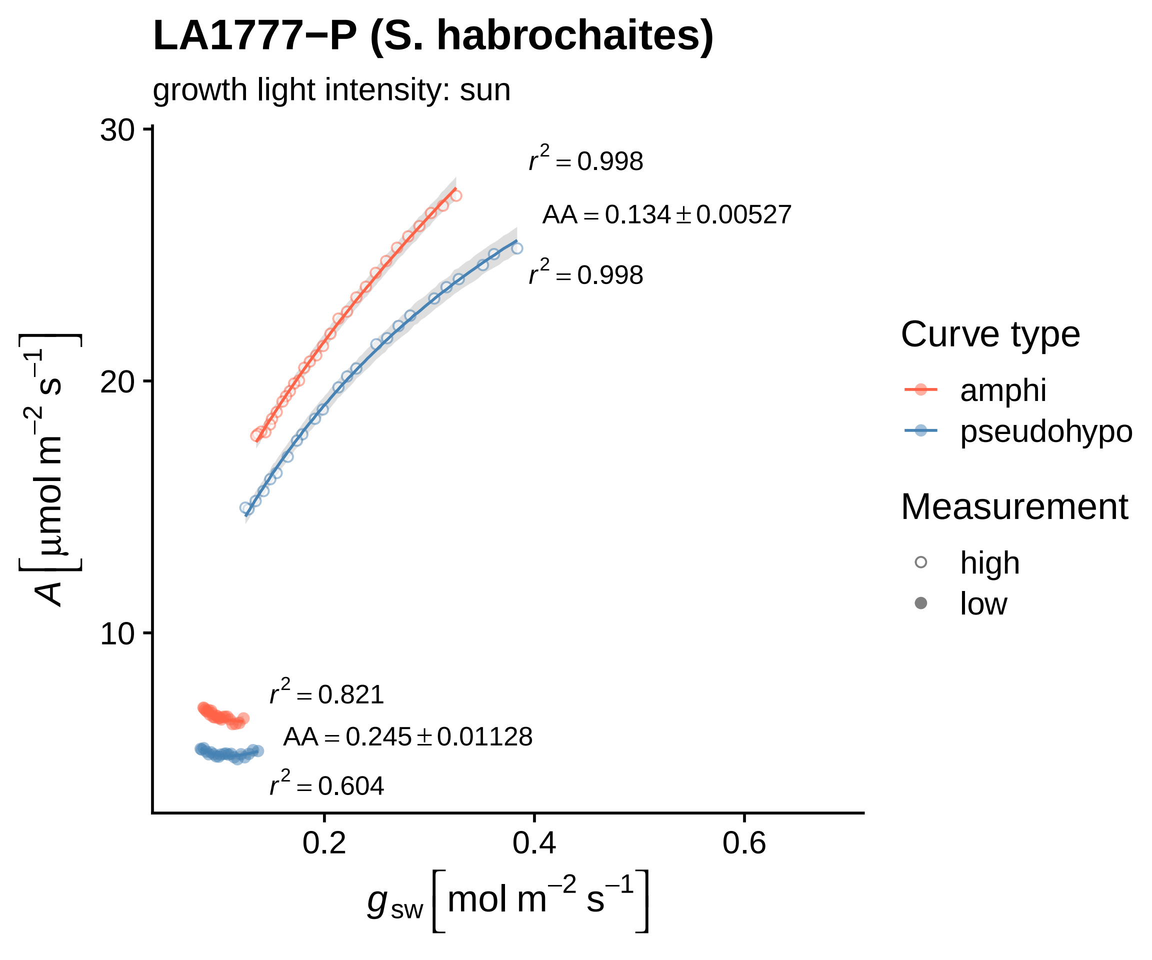

LA1777−P (S. habrochaites)
growth light intensity: sun
30
20
10
0.2
0.4
0.6
$A\left[\mu mol\ m^{-2}\ s^{-1}\right]$
$g_{sw}\left[mol\ m^{-2}\ s^{-1}\right]$
$r^2 = 0.998$
$AA = 0.134 \pm 0.00527$
$r^2 = 0.998$
$r^2 = 0.821$
$AA = 0.245 \pm 0.01128$
$r^2 = 0.604$
Curve type
amphi
pseudohypo
Measurement
high
low

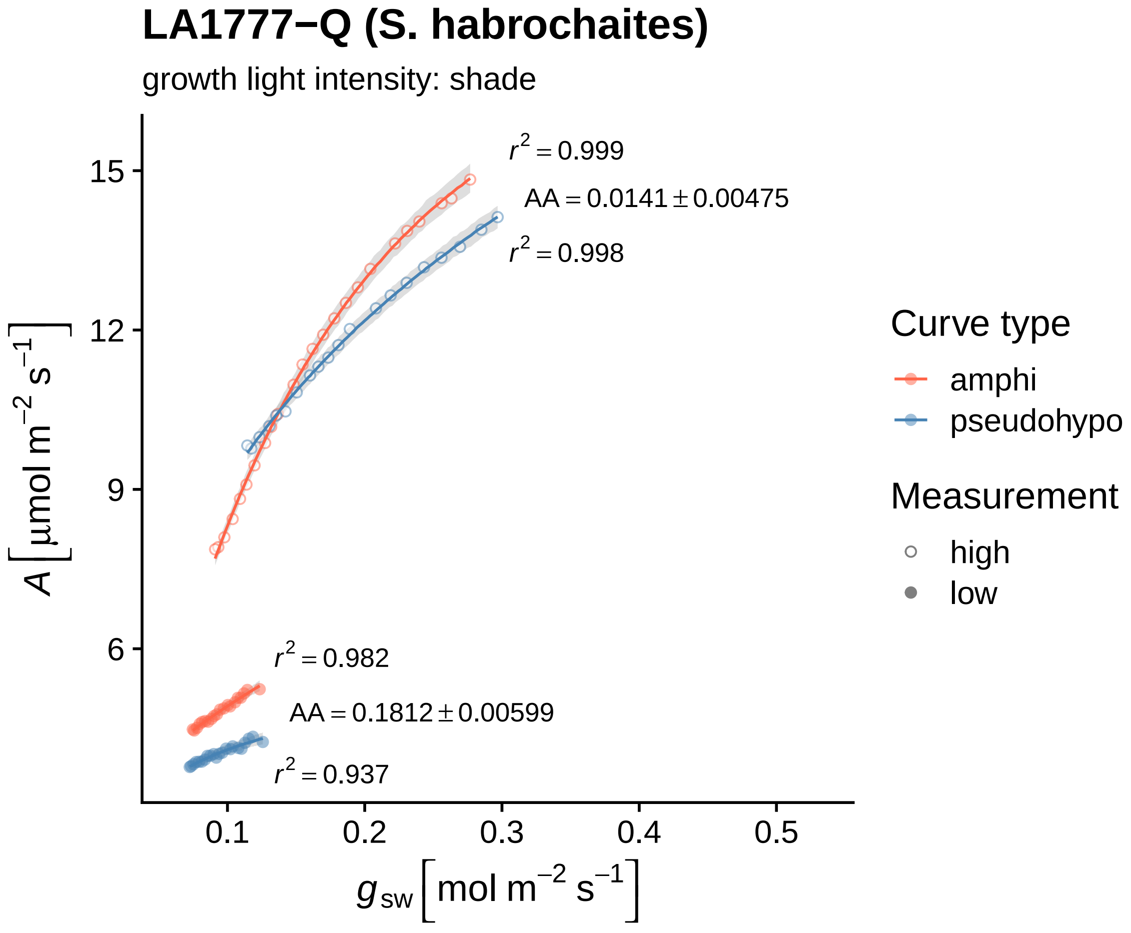
LA1777−Q (S. habrochaites)
growth light intensity: shade
$r^2 = 0.999$
$AA = 0.0141 \pm 0.00475$
$r^2 = 0.998$
$r^2 = 0.982$
$AA = 0.1812 \pm 0.00599$
$r^2 = 0.937$
15
12
9
6
0.1
0.2
0.3
0.4
0.5
$A\ [\mu mol\ m^{-2}\ s^{-1}]$
$g_{sw}\ [mol\ m^{-2}\ s^{-1}]$
Curve type
amphi
pseudohypo
Measurement
high
low

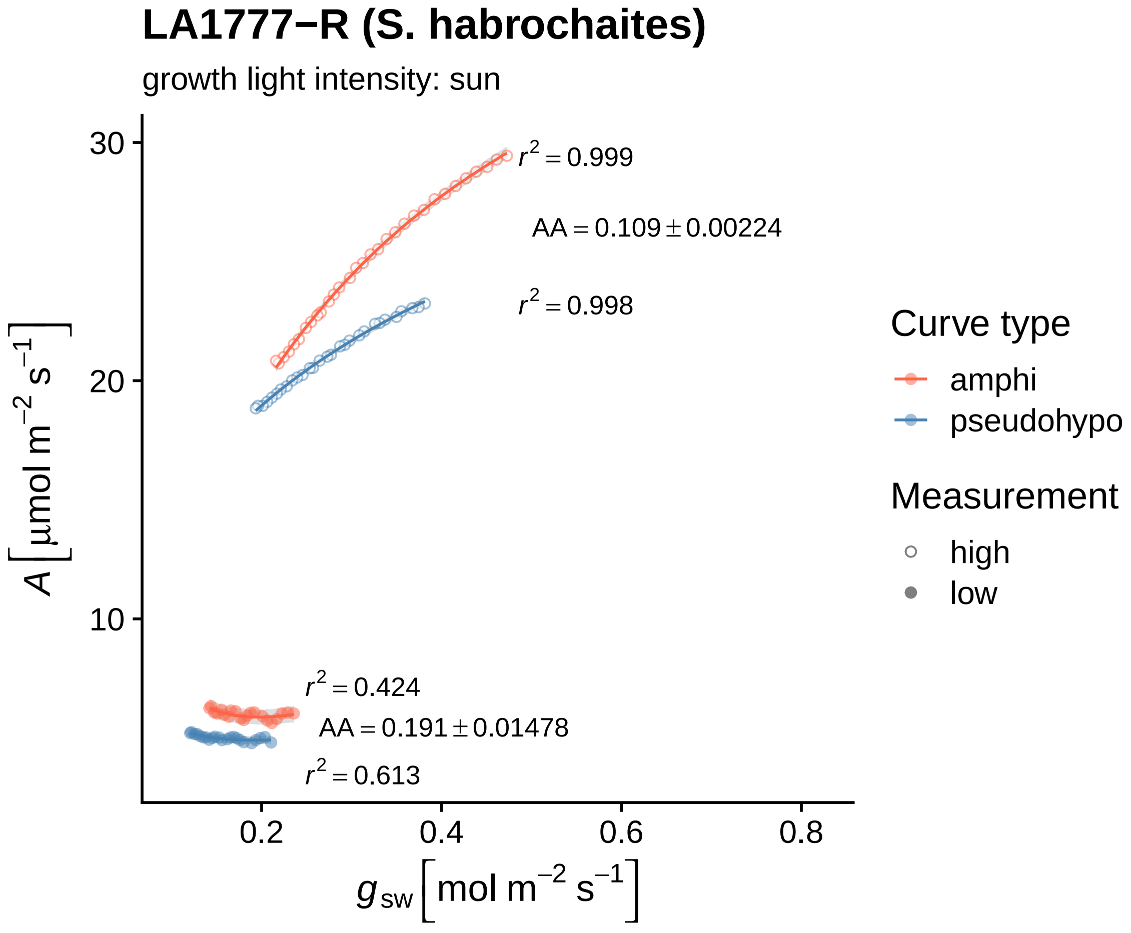

LA1777−R (S. habrochaites)
growth light intensity: sun
$r^2 = 0.999$
$AA = 0.109 \pm 0.00224$
$r^2 = 0.998$
$r^2 = 0.424$
$AA = 0.191 \pm 0.01478$
$r^2 = 0.613$
30
20
10
0.2
0.4
0.6
0.8
$A\ [\mu mol\ m^{-2}\ s^{-1}]$
$g_{sw}\ [mol\ m^{-2}\ s^{-1}]$
Curve type
amphi
pseudohypo
Measurement
high
low

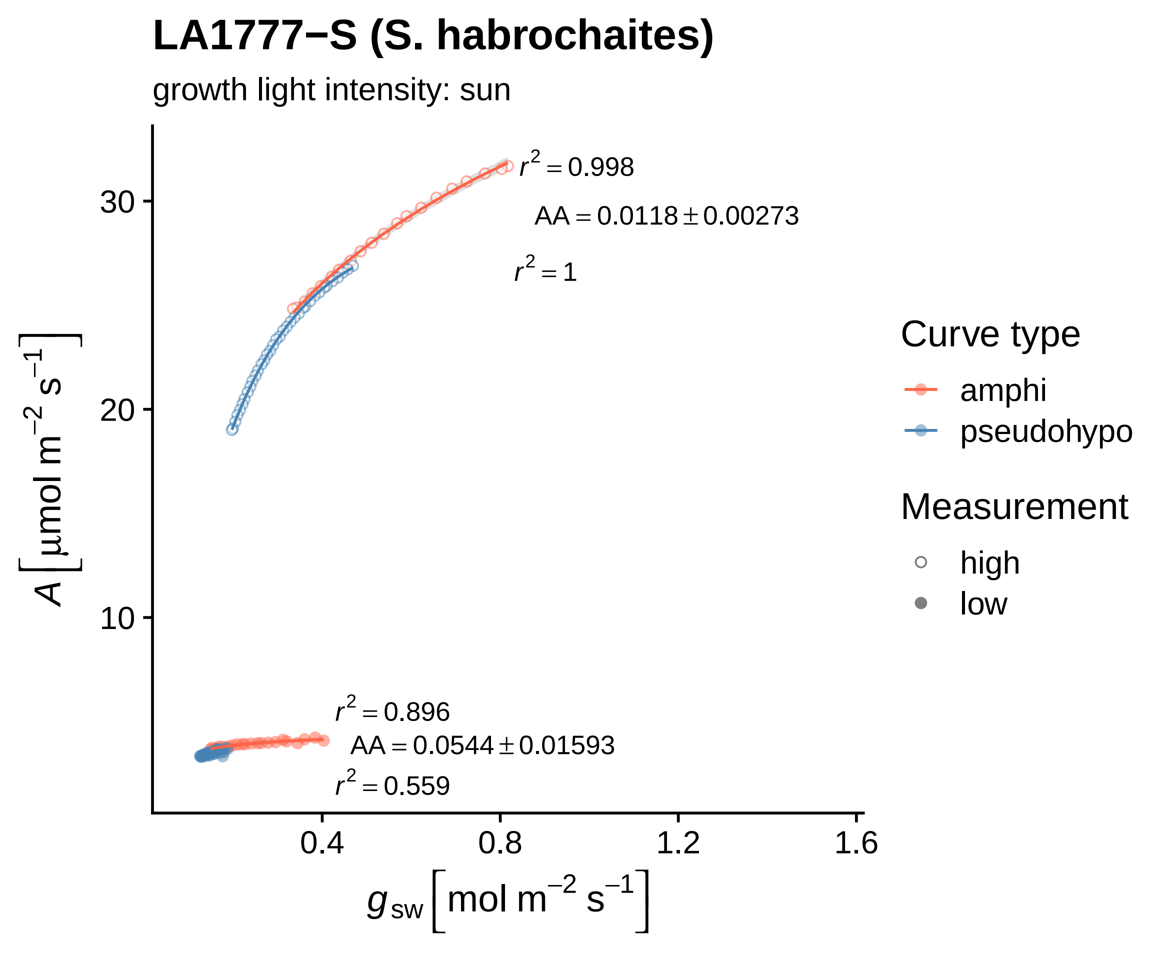

LA1777−S (S. habrochaites)
growth light intensity: sun
$r^2 = 0.998$
$AA = 0.0118 \pm 0.00273$
$r^2 = 1$
$r^2 = 0.896$
$AA = 0.0544 \pm 0.01593$
$r^2 = 0.559$
30
20
10
0.4
0.8
1.2
1.6
$A\left[\mu mol\ m^{-2}\ s^{-1}\right]$
$g_{sw}\left[mol\ m^{-2}\ s^{-1}\right]$
Curve type
amphi
pseudohypo
Measurement
high
low

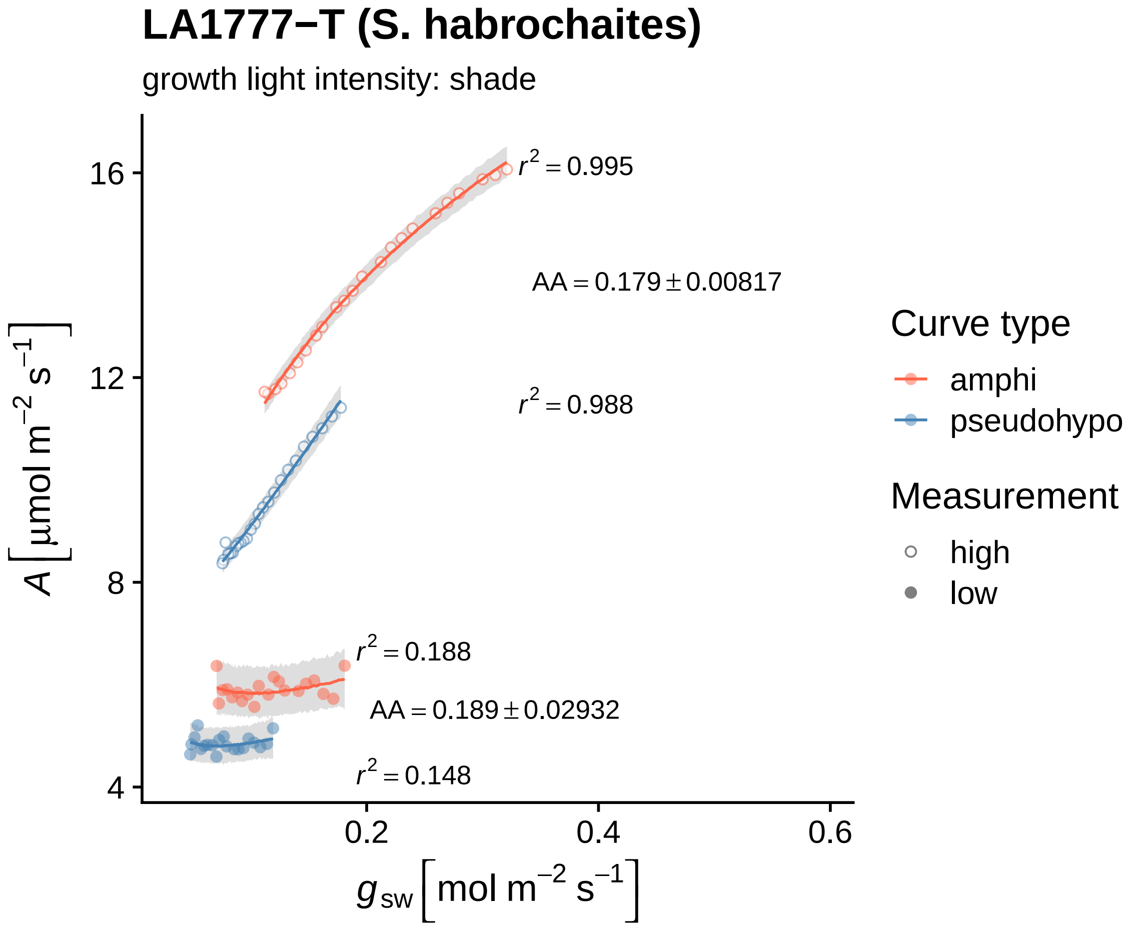

LA1777−T (S. habrochaites)
growth light intensity: shade
$r^2 = 0.995$
$AA = 0.179 \pm 0.00817$
$r^2 = 0.988$
$r^2 = 0.188$
$AA = 0.189 \pm 0.02932$
$r^2 = 0.148$
16
12
8
4
0.2
0.4
0.6
$A\left[\mu mol\ m^{-2}\ s^{-1}\right]$
$g_{sw}\left[mol\ m^{-2}\ s^{-1}\right]$
Curve type
amphi
pseudohypo
Measurement
high
low

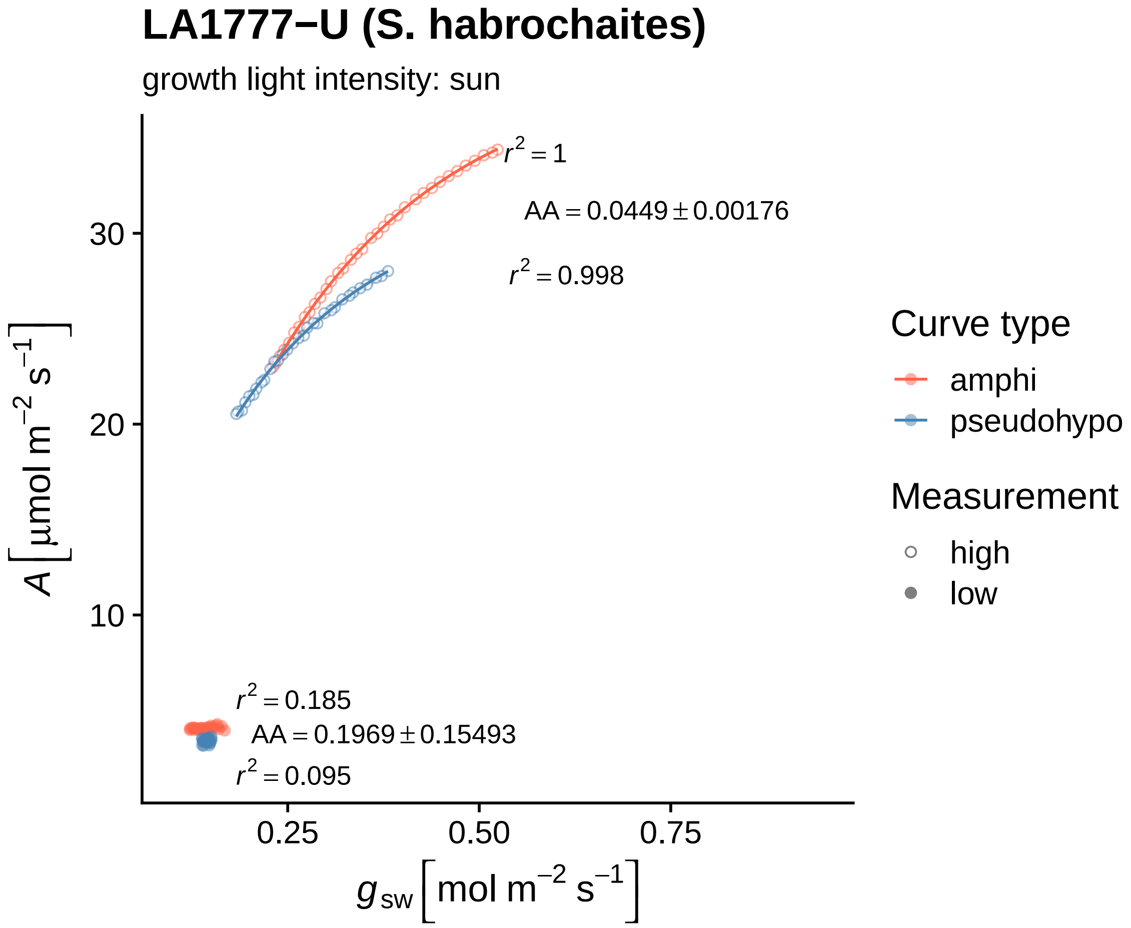

LA1777−U (S. habrochaites)
growth light intensity: sun
$A\left[\mu mol\ m^{-2}\ s^{-1}\right]$
30
20
10
$r^2 = 1$
$AA = 0.0449 \pm 0.00176$
$r^2 = 0.998$
$r^2 = 0.185$
$AA = 0.1969 \pm 0.15493$
$r^2 = 0.095$
0.25
0.50
0.75
$g_{sw}\left[mol\ m^{-2}\ s^{-1}\right]$
Curve type
amphi
pseudohypo
Measurement
high
low

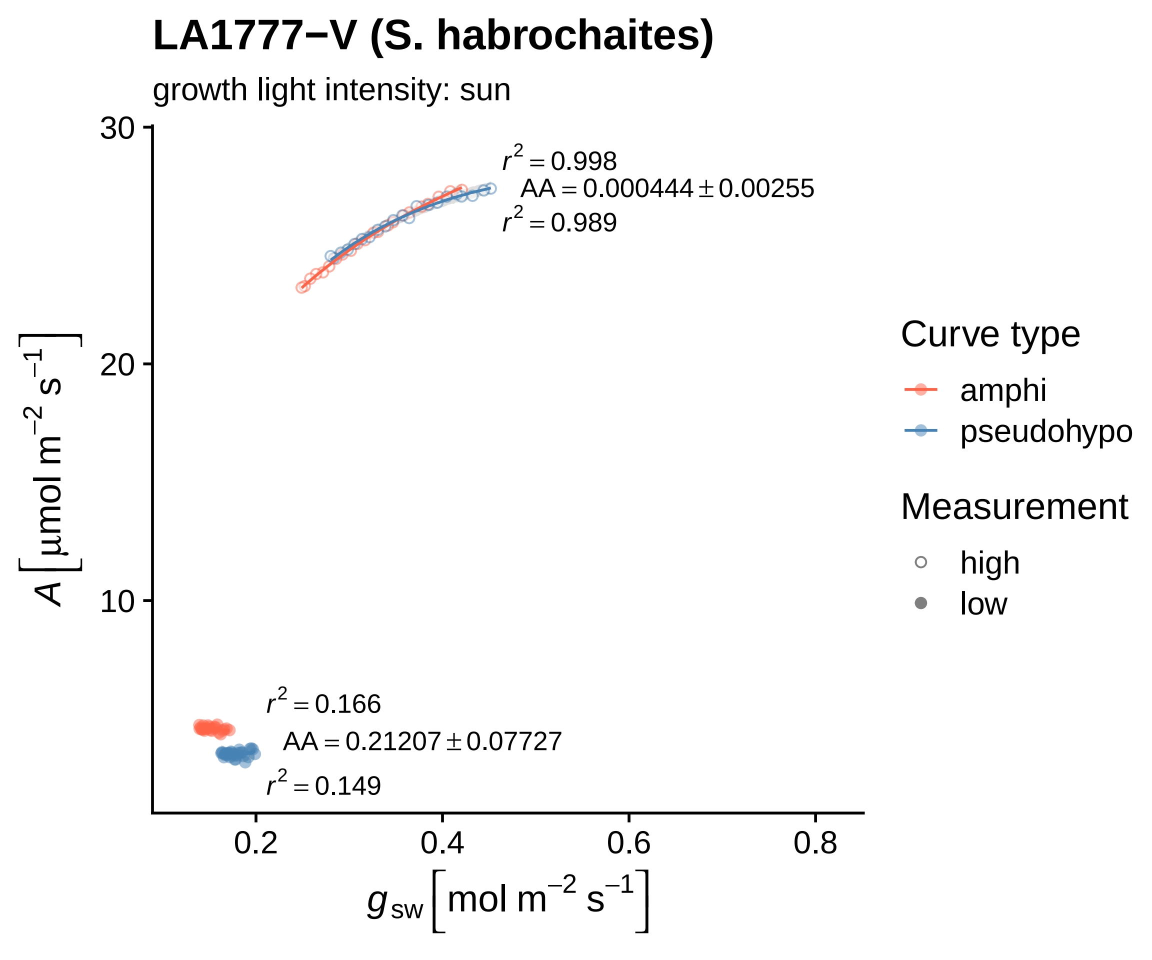

LA1777−V (S. habrochaites)
growth light intensity: sun
$r^2 = 0.998$
AA = 0.000444 ± 0.00255
$r^2 = 0.989$
$r^2 = 0.166$
AA = 0.21207 ± 0.07727
$r^2 = 0.149$
30
20
10
0.2
0.4
0.6
0.8
$A\ [\mu mol\ m^{-2}\ s^{-1}]$
$g_{sw}\ [mol\ m^{-2}\ s^{-1}]$
Curve type
amphi
pseudohypo
Measurement
high
low

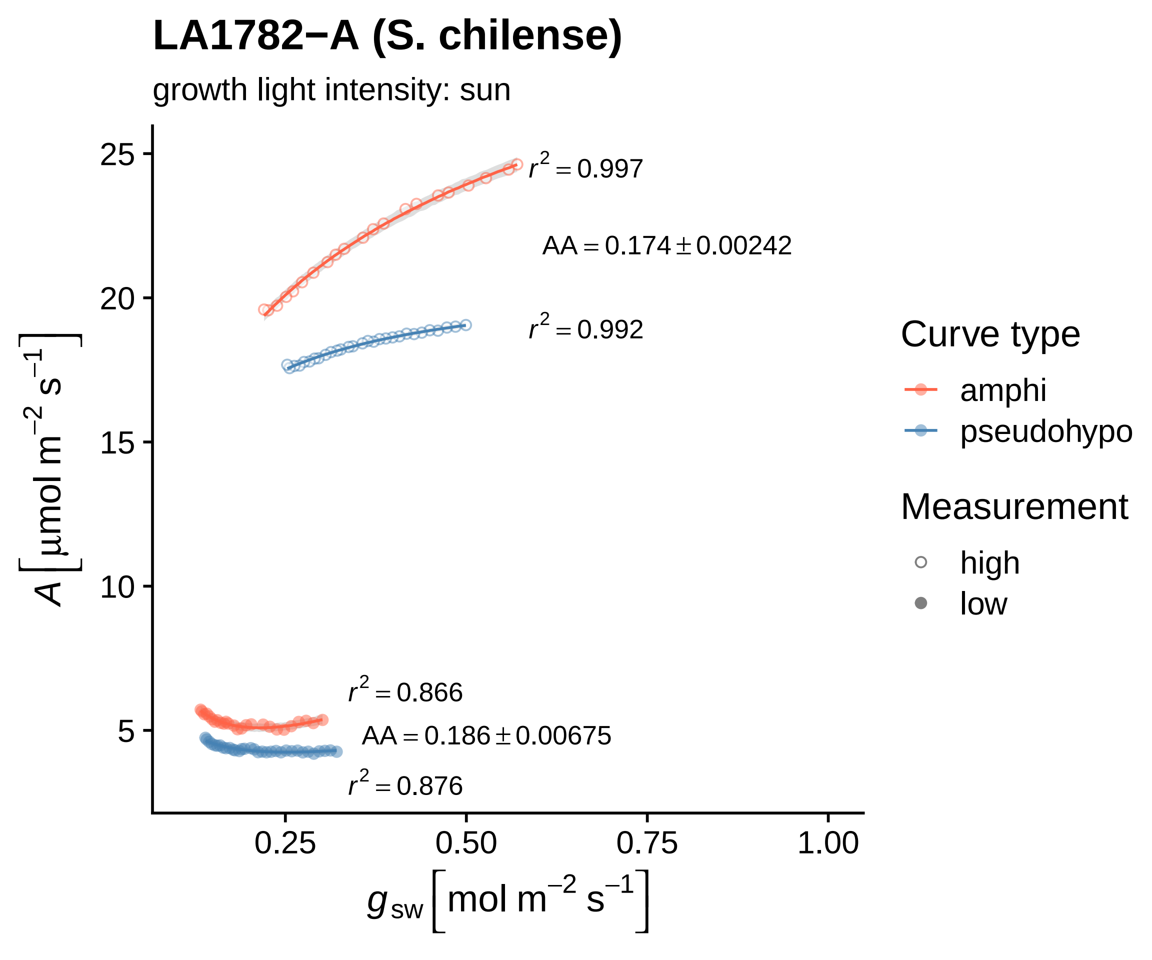
LA1782−A (S. chilense)
growth light intensity: sun
r² = 0.997
AA = 0.174 ± 0.00242
r² = 0.992
r² = 0.866
AA = 0.186 ± 0.00675
r² = 0.876
25
20
15
10
5
0.25
0.50
0.75
1.00
A [μmol m⁻² s⁻¹]
g sw [mol m⁻² s⁻¹]
Curve type
amphi
pseudohypo
Measurement
high
low

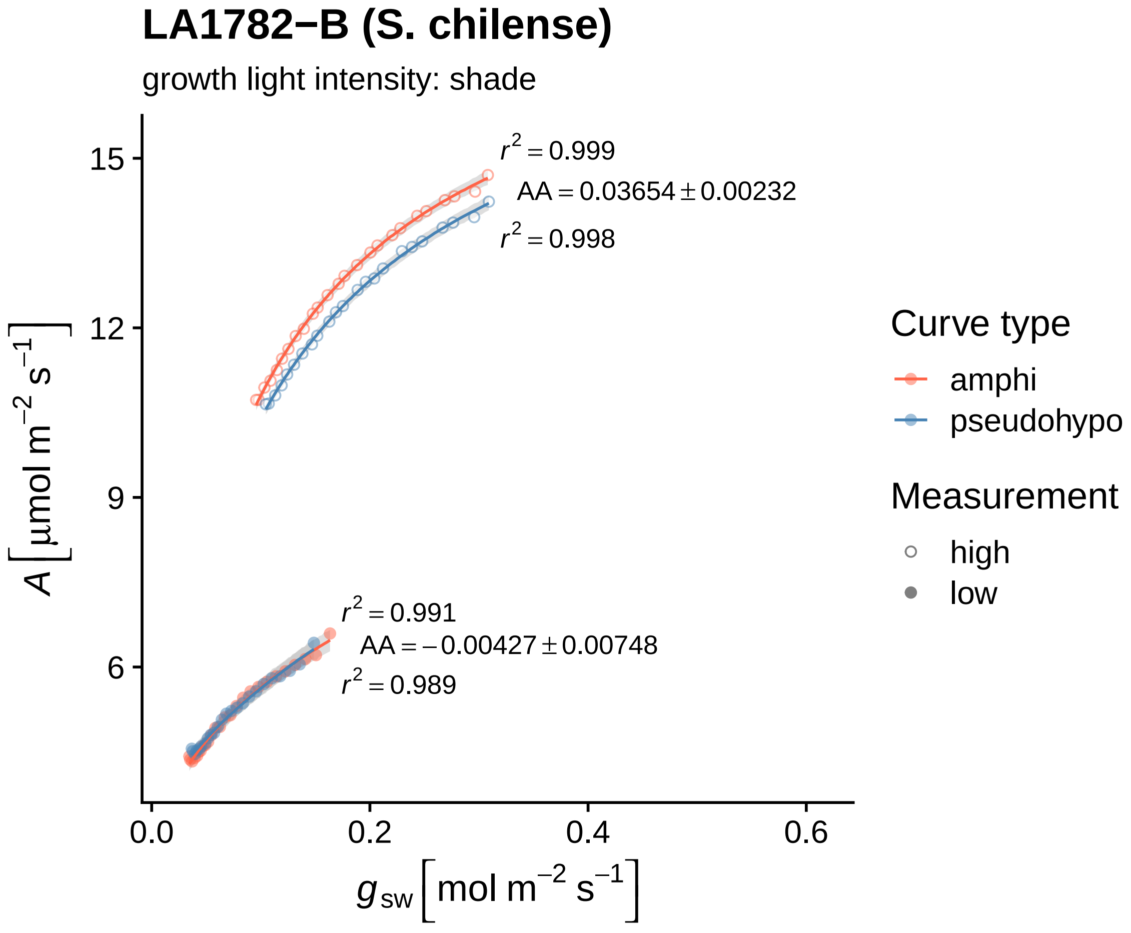

LA1782−B (S. chilense)
growth light intensity: shade
$r^2 = 0.999$
$AA = 0.03654 \pm 0.00232$
$r^2 = 0.998$
$r^2 = 0.991$
$AA = -0.00427 \pm 0.00748$
$r^2 = 0.989$
15
12
9
6
0.0
0.2
0.4
0.6
$A\left[\mu mol\ m^{-2}\ s^{-1}\right]$
$g_{sw}\left[mol\ m^{-2}\ s^{-1}\right]$
Curve type
amphi
pseudohypo
Measurement
high
low

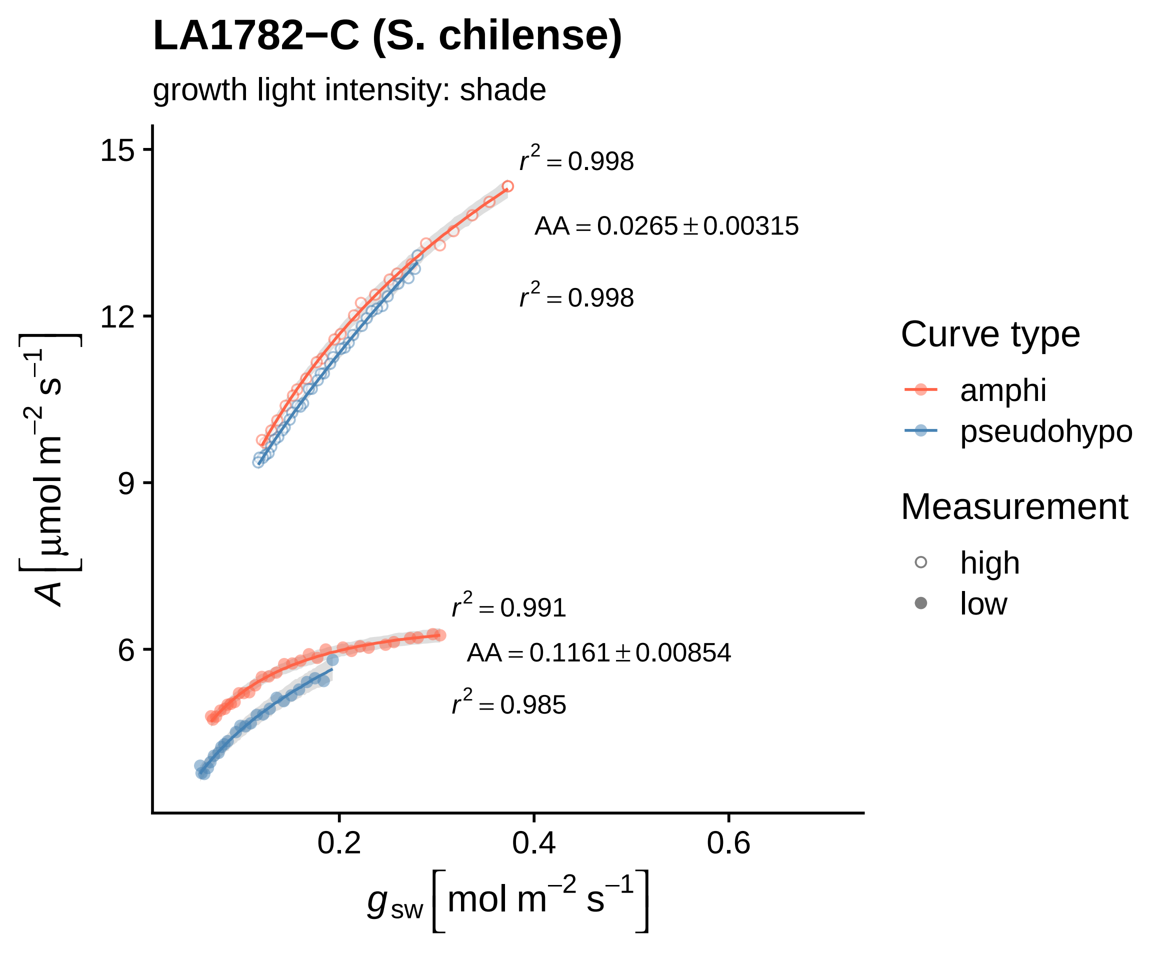

LA1782−C (S. chilense)
growth light intensity: shade
$r^2 = 0.998$
$AA = 0.0265 \pm 0.00315$
$r^2 = 0.998$
$r^2 = 0.991$
$AA = 0.1161 \pm 0.00854$
$r^2 = 0.985$
15
12
9
6
0.2
0.4
0.6
$A\ [\mu mol\ m^{-2}\ s^{-1}]$
$g_{sw}\ [mol\ m^{-2}\ s^{-1}]$
Curve type
amphi
pseudohypo
Measurement
high
low

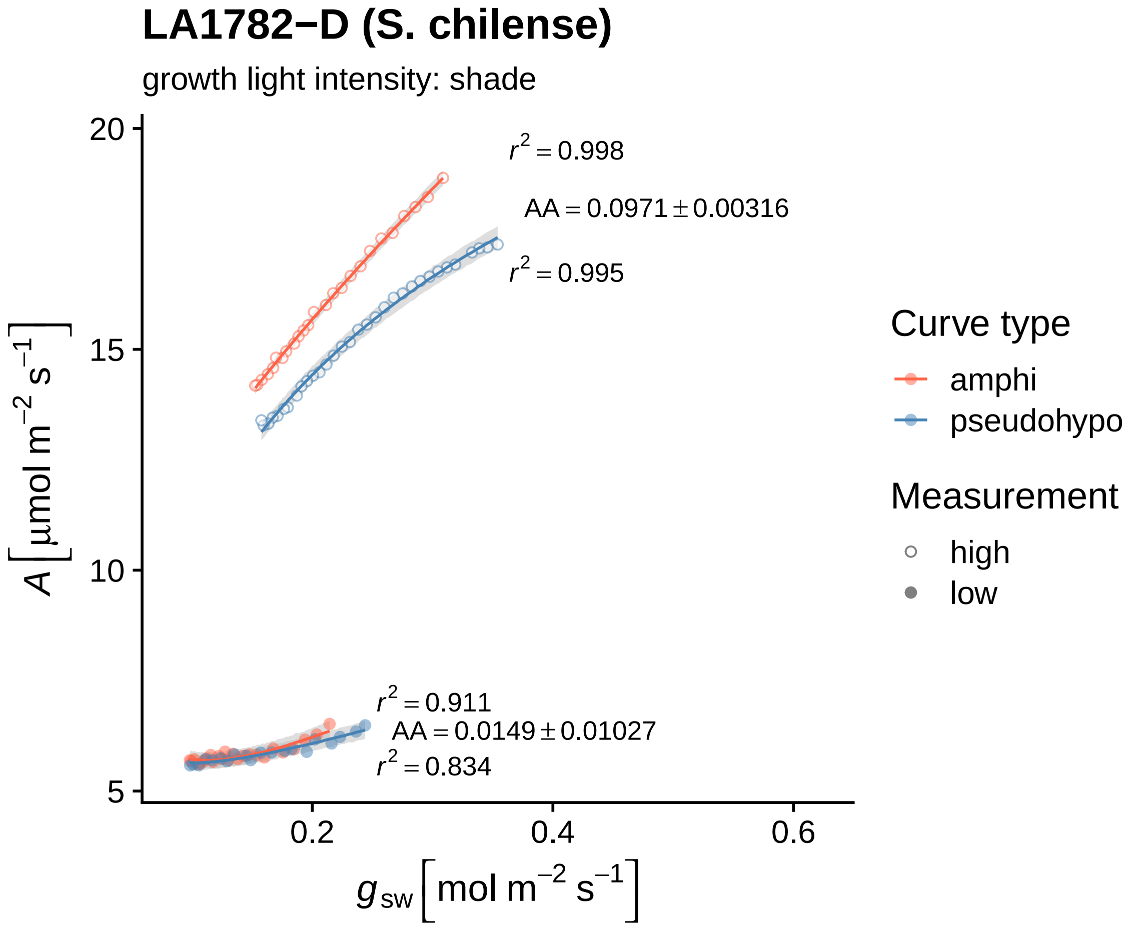
LA1782−D (S. chilense)
growth light intensity: shade
$r^2 = 0.998$
$AA = 0.0971 \pm 0.00316$
$r^2 = 0.995$
$r^2 = 0.911$
$AA = 0.0149 \pm 0.01027$
$r^2 = 0.834$
20
15
10
5
0.2
0.4
0.6
$A\left[\mu mol\ m^{-2}\ s^{-1}\right]$
$g_{sw}\left[mol\ m^{-2}\ s^{-1}\right]$
Curve type
amphi
pseudohypo
Measurement
high
low

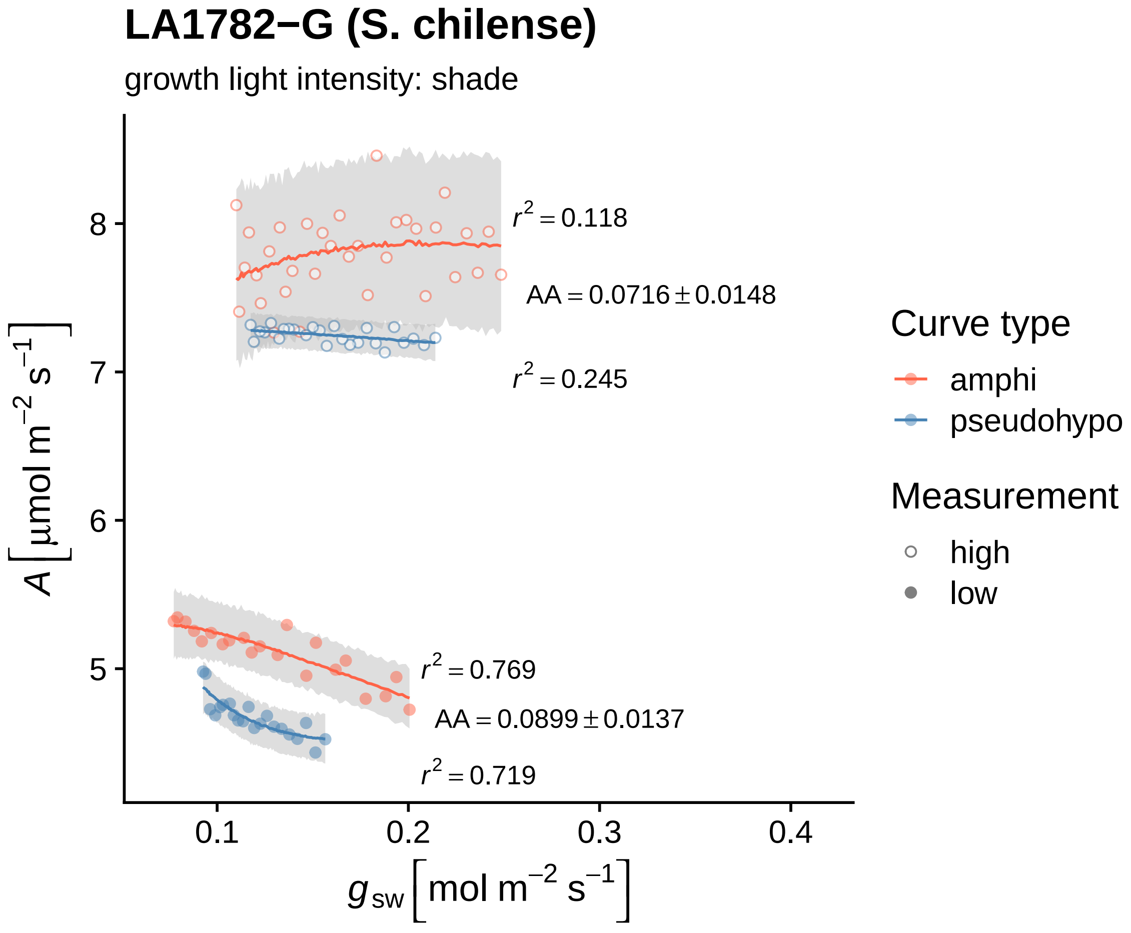

LA1782−G (S. chilense)
growth light intensity: shade
$r^2 = 0.118$
$AA = 0.0716 \pm 0.0148$
$r^2 = 0.245$
$r^2 = 0.769$
$AA = 0.0899 \pm 0.0137$
$r^2 = 0.719$
8
7
6
5
0.1
0.2
0.3
0.4
$A\ [\mu mol\ m^{-2}\ s^{-1}]$
$g_{sw}\ [mol\ m^{-2}\ s^{-1}]$
Curve type
amphi
pseudohypo
Measurement
high
low

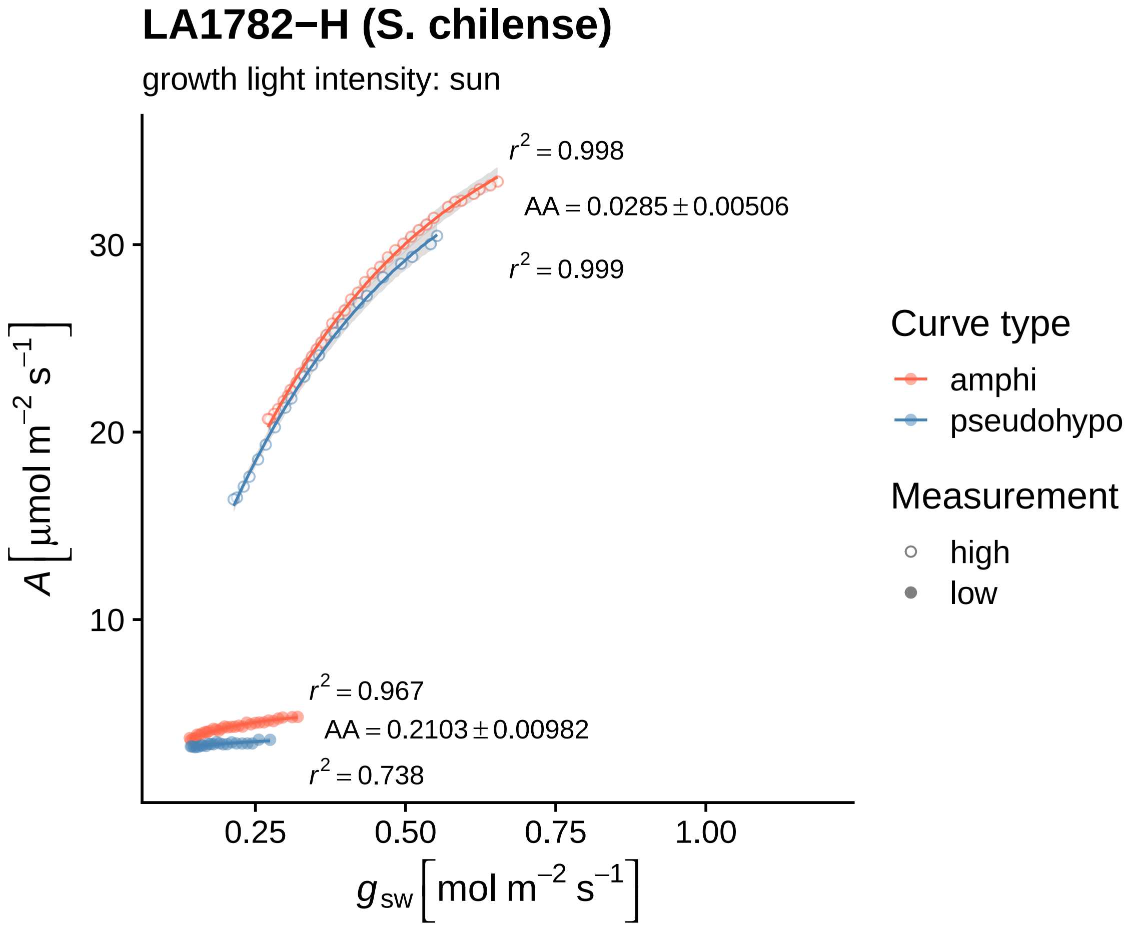

LA1782−H (S. chilense)
growth light intensity: sun
r² = 0.998
AA = 0.0285 ± 0.00506
r² = 0.999
r² = 0.967
AA = 0.2103 ± 0.00982
r² = 0.738
30
20
10
0.25
0.50
0.75
1.00
A [μmol m⁻² s⁻¹]
g sw [mol m⁻² s⁻¹]
Curve type
amphi
pseudohypo
Measurement
high
low

#### LA1782−I (S. chilense)

growth light intensity: shade

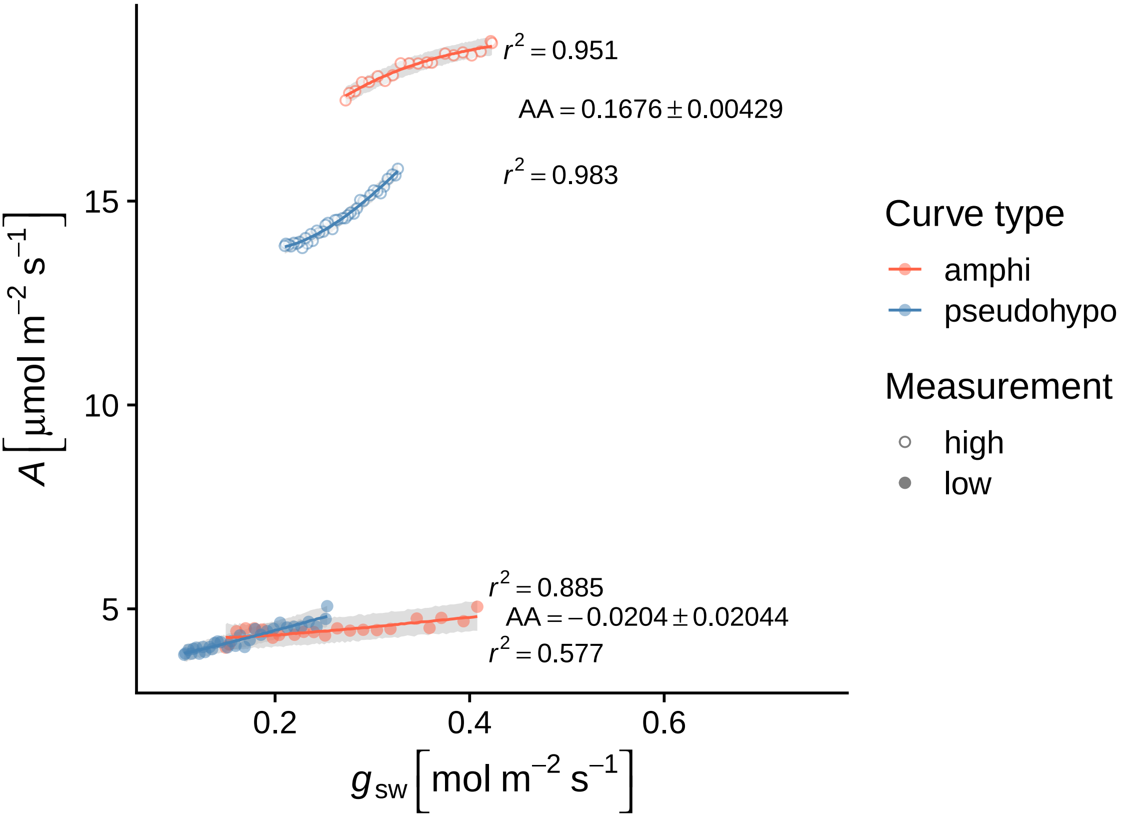

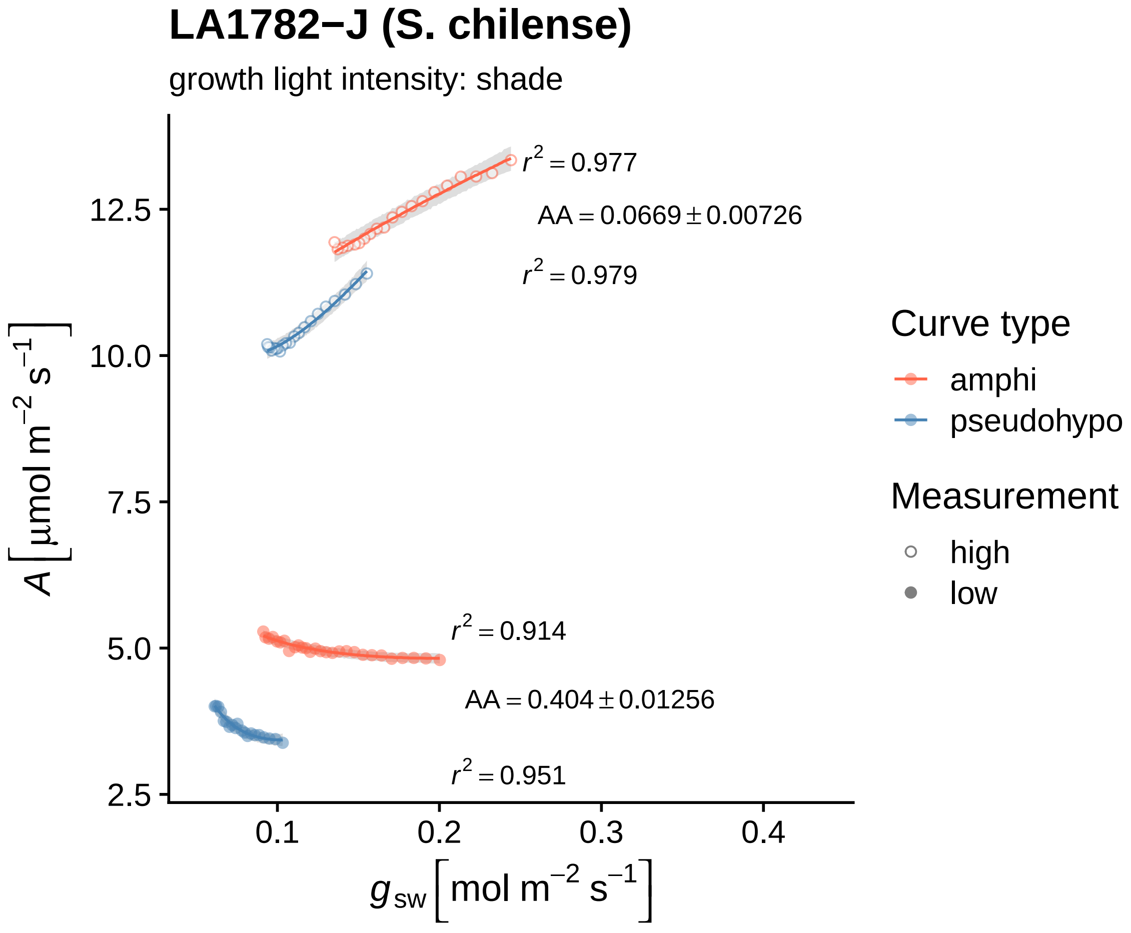

LA1782−J (S. chilense)
growth light intensity: shade
$r^2 = 0.977$
$AA = 0.0669 \pm 0.00726$
$r^2 = 0.979$
$r^2 = 0.914$
$AA = 0.404 \pm 0.01256$
$r^2 = 0.951$
$A\ [\mu mol\ m^{-2}\ s^{-1}]$
12.5
10.0
7.5
5.0
2.5
0.1
0.2
0.3
0.4
$g_{sw}\ [mol\ m^{-2}\ s^{-1}]$
Curve type
amphi
pseudohypo
Measurement
high
low

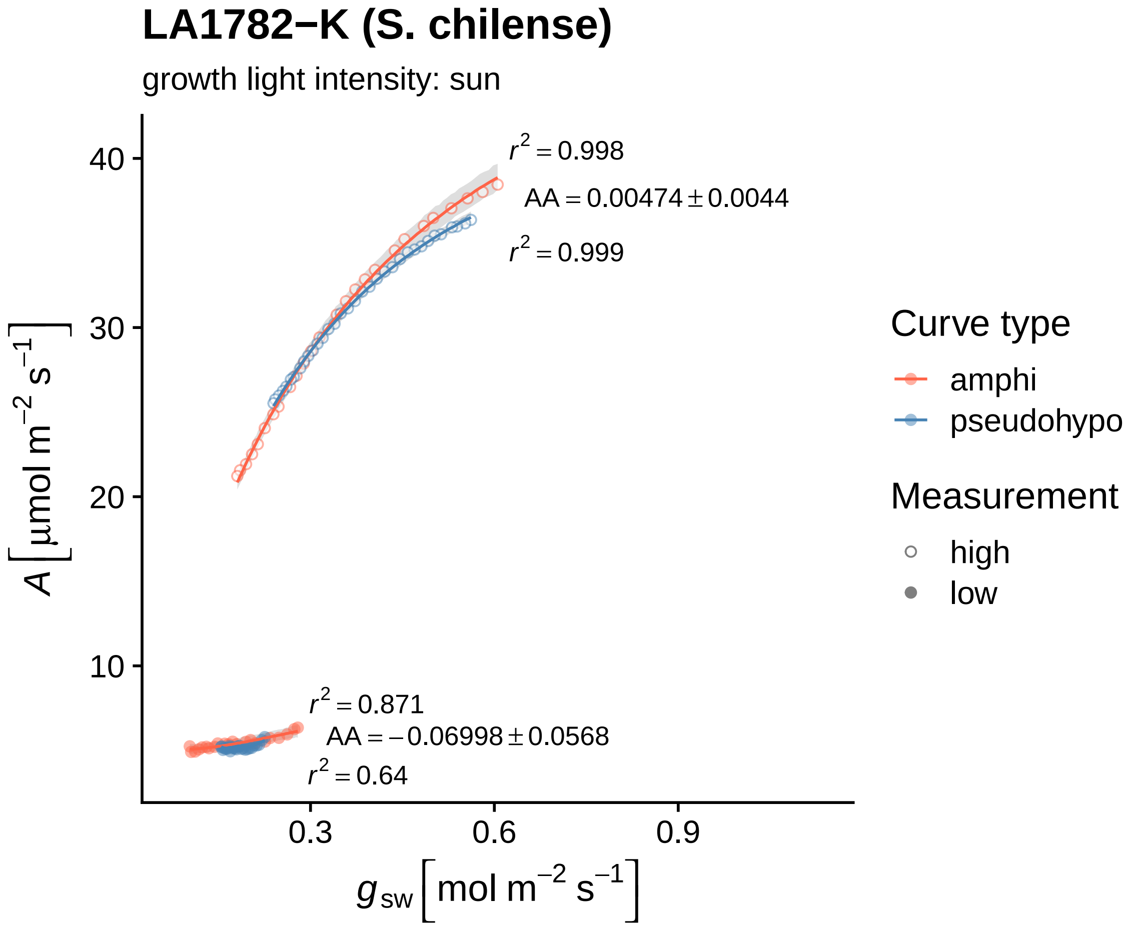
LA1782−K (S. chilense)
growth light intensity: sun
$r^2 = 0.998$
$AA = 0.00474 \pm 0.0044$
$r^2 = 0.999$
$r^2 = 0.871$
$AA = -0.06998 \pm 0.0568$
$r^2 = 0.64$
40
30
20
10
0.3
0.6
0.9
$A\ [\mu mol\ m^{-2}\ s^{-1}]$
$g_{sw}\ [mol\ m^{-2}\ s^{-1}]$
Curve type
amphi
pseudohypo
Measurement
high
low

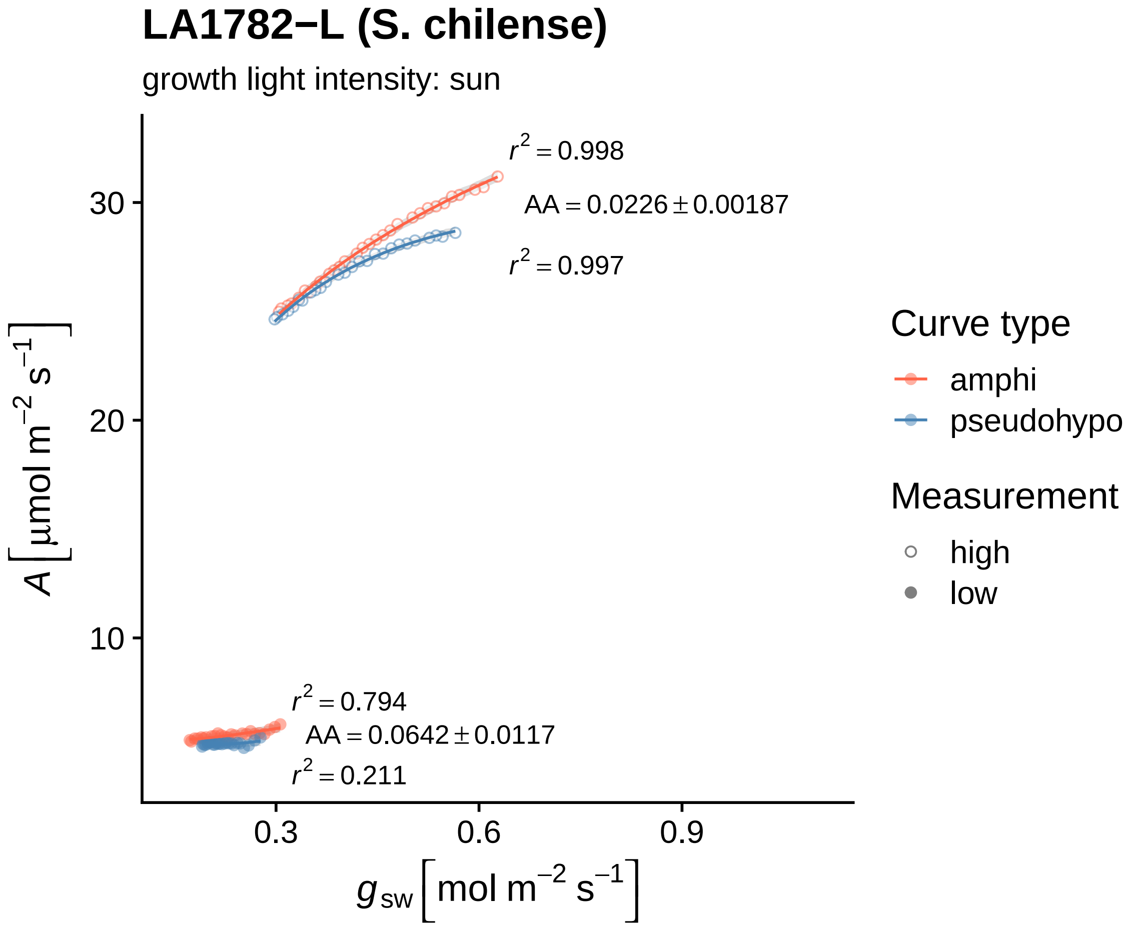

LA1782−L (S. chilense)
growth light intensity: sun
$r^2 = 0.998$
$AA = 0.0226 \pm 0.00187$
$r^2 = 0.997$
$r^2 = 0.794$
$AA = 0.0642 \pm 0.0117$
$r^2 = 0.211$
30
20
10
0.3
0.6
0.9
$A\ [\mu mol\ m^{-2}\ s^{-1}]$
$g_{sw}\ [mol\ m^{-2}\ s^{-1}]$
Curve type
amphi
pseudohypo
Measurement
high
low

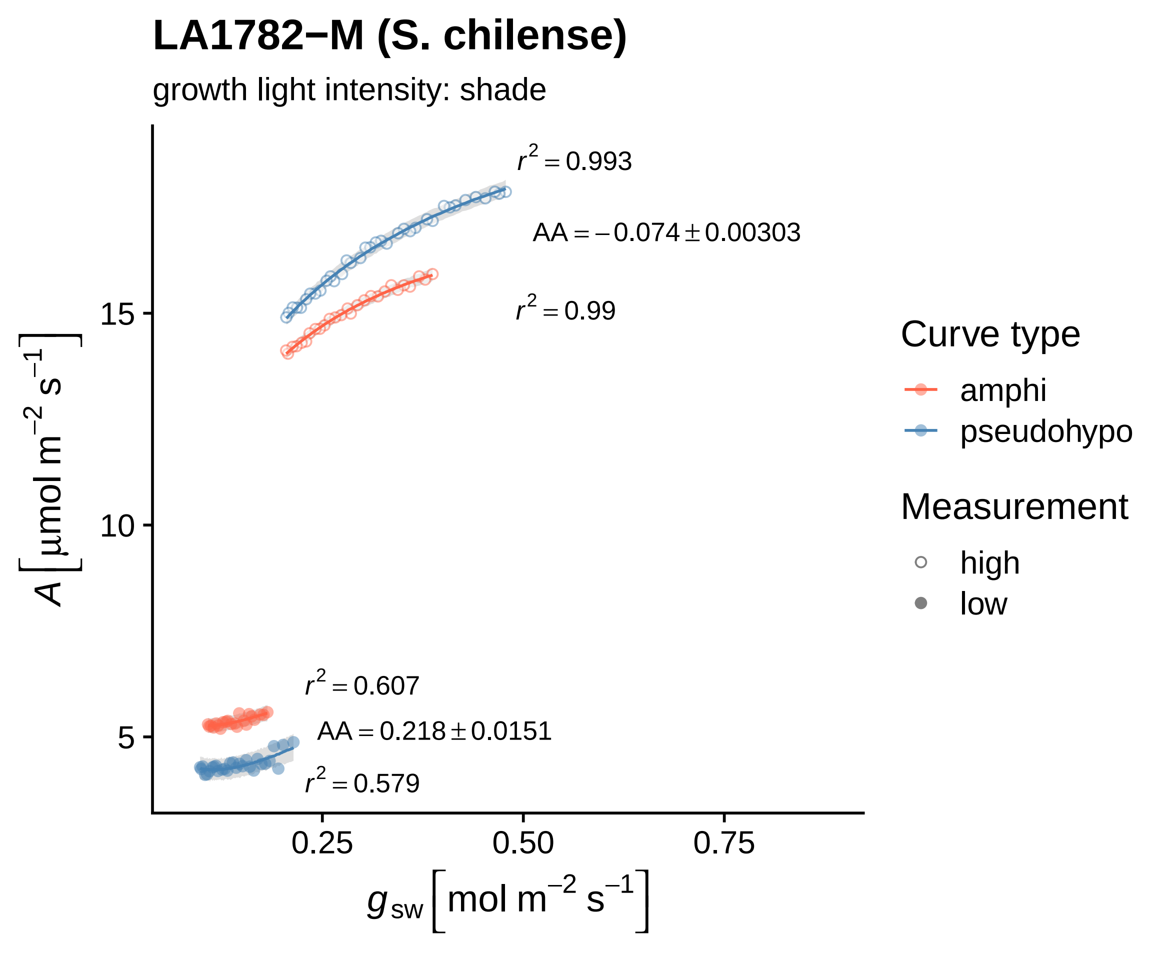

LA1782−M (S. chilense)
growth light intensity: shade
$r^2 = 0.993$
$AA = -0.074 \pm 0.00303$
$r^2 = 0.99$
$r^2 = 0.607$
$AA = 0.218 \pm 0.0151$
$r^2 = 0.579$
15
10
5
0.25
0.50
0.75
$A\left[\mu mol\ m^{-2}\ s^{-1}\right]$
$g_{sw}\left[mol\ m^{-2}\ s^{-1}\right]$
Curve type
amphi
pseudohypo
Measurement
high
low

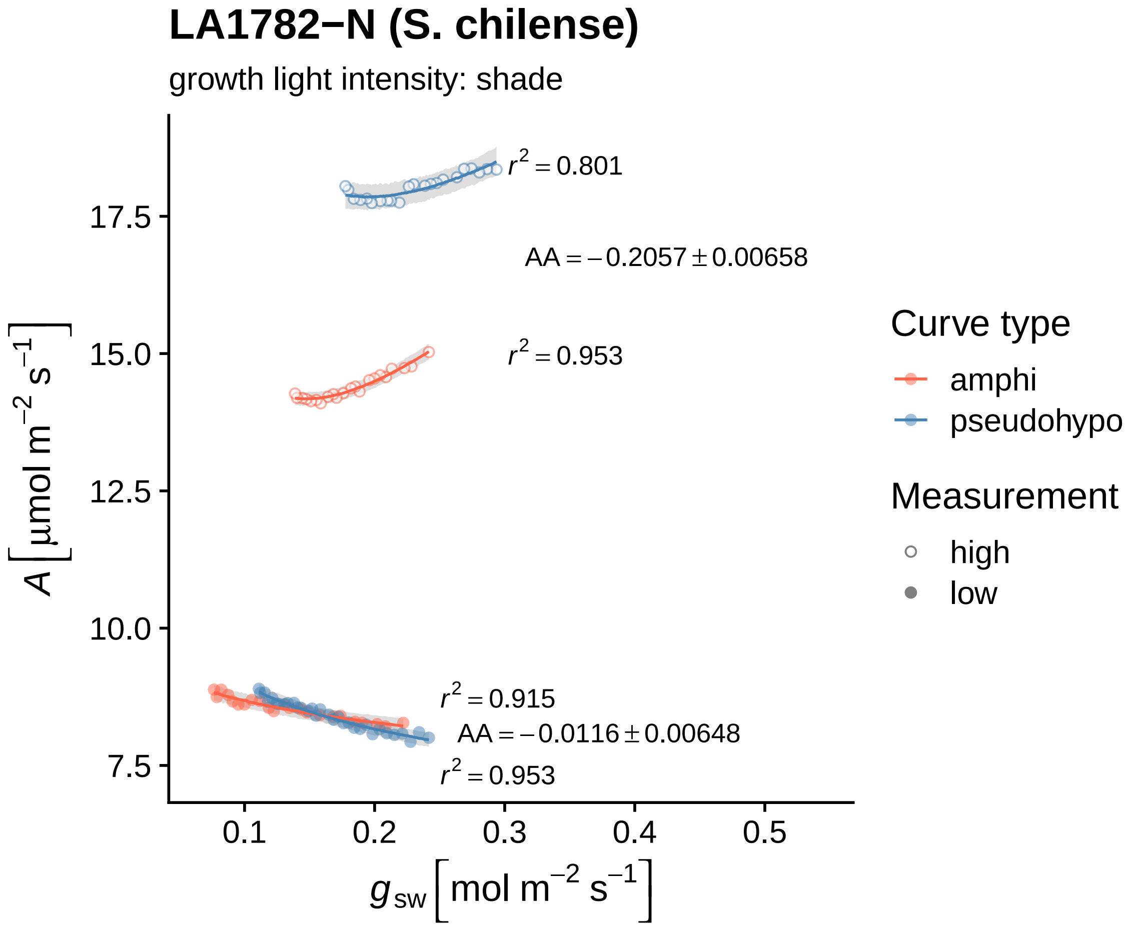

LA1782−N (S. chilense)
growth light intensity: shade
$r^2 = 0.801$
$AA = -0.2057 \pm 0.00658$
$r^2 = 0.953$
$r^2 = 0.915$
$AA = -0.0116 \pm 0.00648$
$r^2 = 0.953$
17.5
15.0
12.5
10.0
7.5
0.1
0.2
0.3
0.4
0.5
$A\ [\mu mol\ m^{-2}\ s^{-1}]$
$g_{sw}\ [mol\ m^{-2}\ s^{-1}]$
Curve type
amphi
pseudohypo
Measurement
high
low

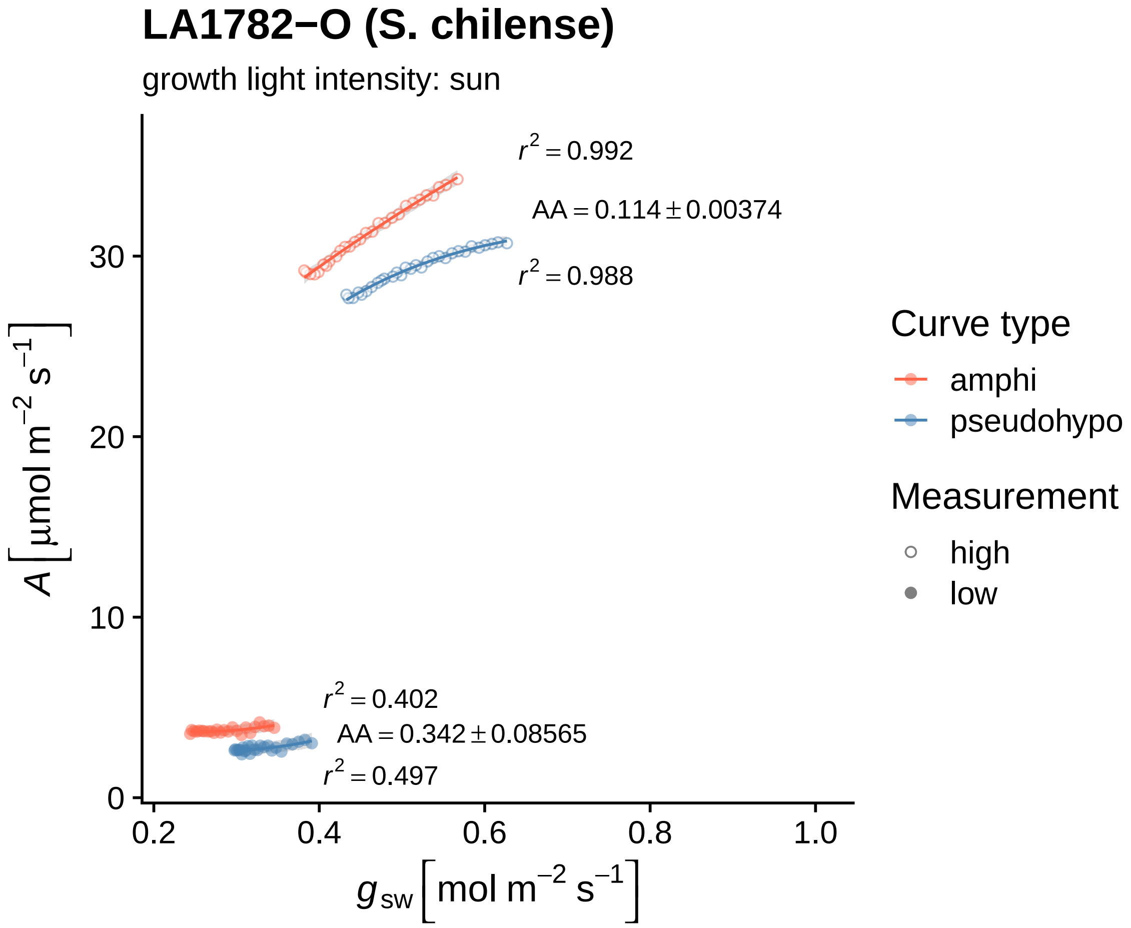

LA1782−O (S. chilense)
growth light intensity: sun
r² = 0.992
AA = 0.114 ± 0.00374
r² = 0.988
r² = 0.402
AA = 0.342 ± 0.08565
r² = 0.497
0
10
20
30
0.2
0.4
0.6
0.8
1.0
A [µmol m⁻² s⁻¹]
g sw [mol m⁻² s⁻¹]
Curve type
amphi
pseudohypo
Measurement
high
low

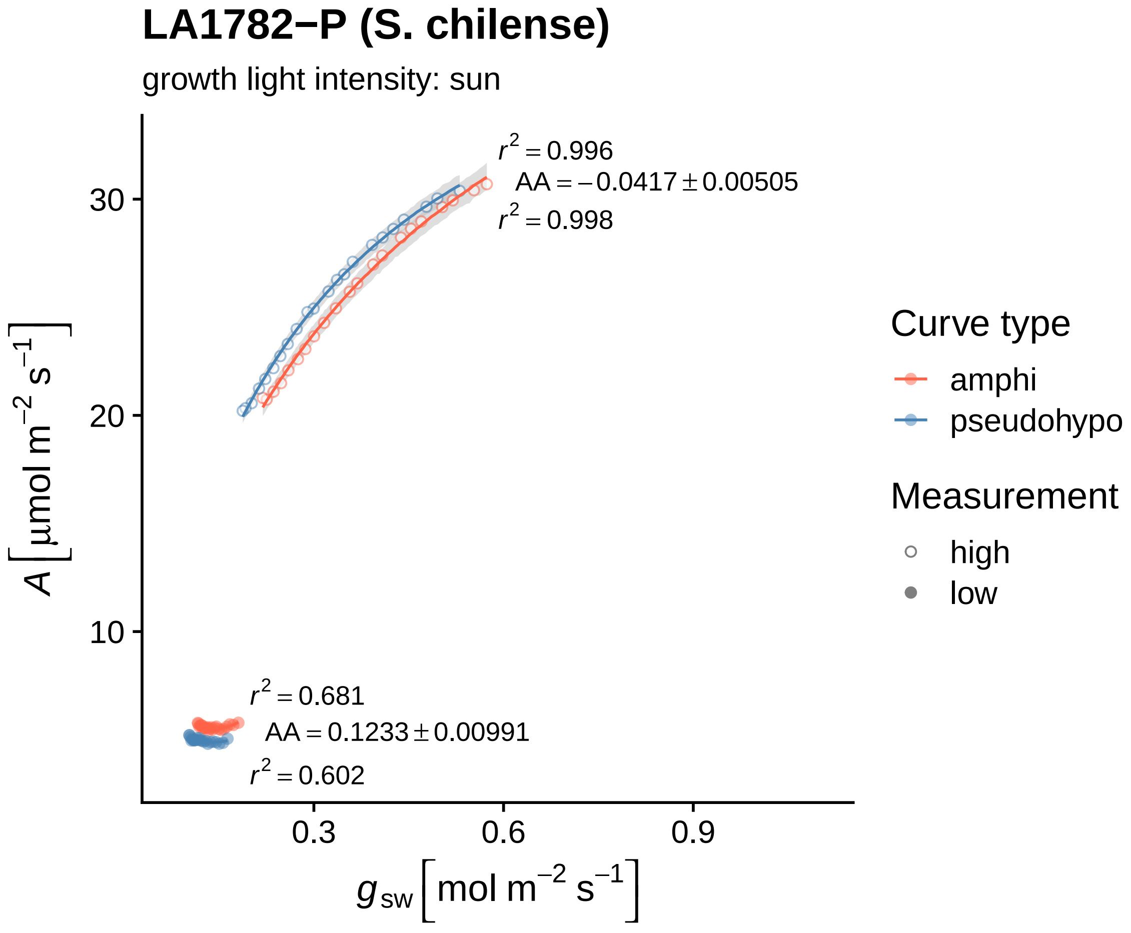

LA1782−P (S. chilense)
growth light intensity: sun
r² = 0.996
AA = −0.0417 ± 0.00505
r² = 0.998
r² = 0.681
AA = 0.1233 ± 0.00991
r² = 0.602
30
20
10
0.3
0.6
0.9
A [μmol m⁻² s⁻¹]
g_sw [mol m⁻² s⁻¹]
Curve type
amphi
pseudohypo
Measurement
high
low

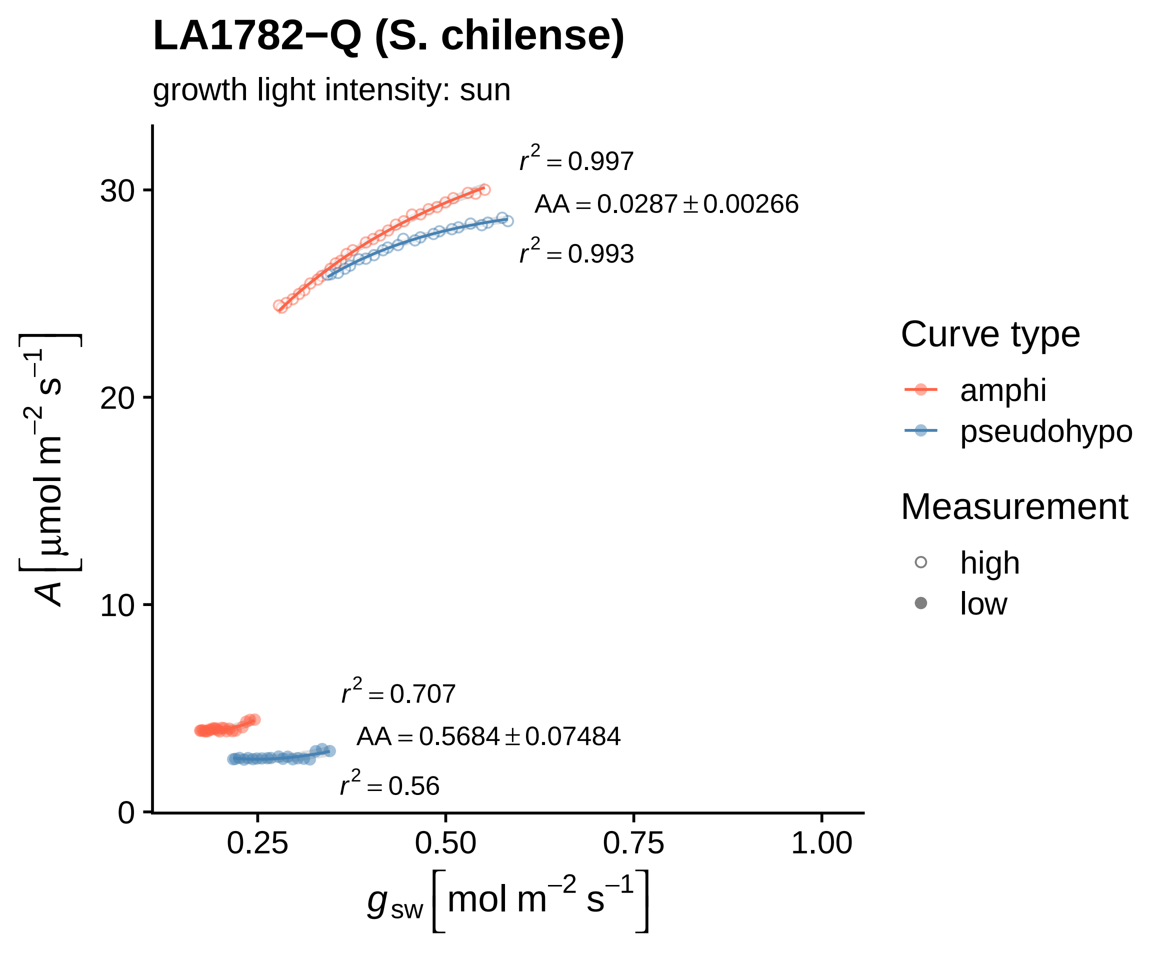
LA1782−Q (S. chilense)
growth light intensity: sun
$r^2 = 0.997$
$AA = 0.0287 \pm 0.00266$
$r^2 = 0.993$
$r^2 = 0.707$
$AA = 0.5684 \pm 0.07484$
$r^2 = 0.56$
$A\ [\mu mol\ m^{-2}\ s^{-1}]$
30
20
10
0
0.25
0.50
0.75
1.00
$g_{sw}\ [mol\ m^{-2}\ s^{-1}]$
Curve type
amphi
pseudohypo
Measurement
high
low

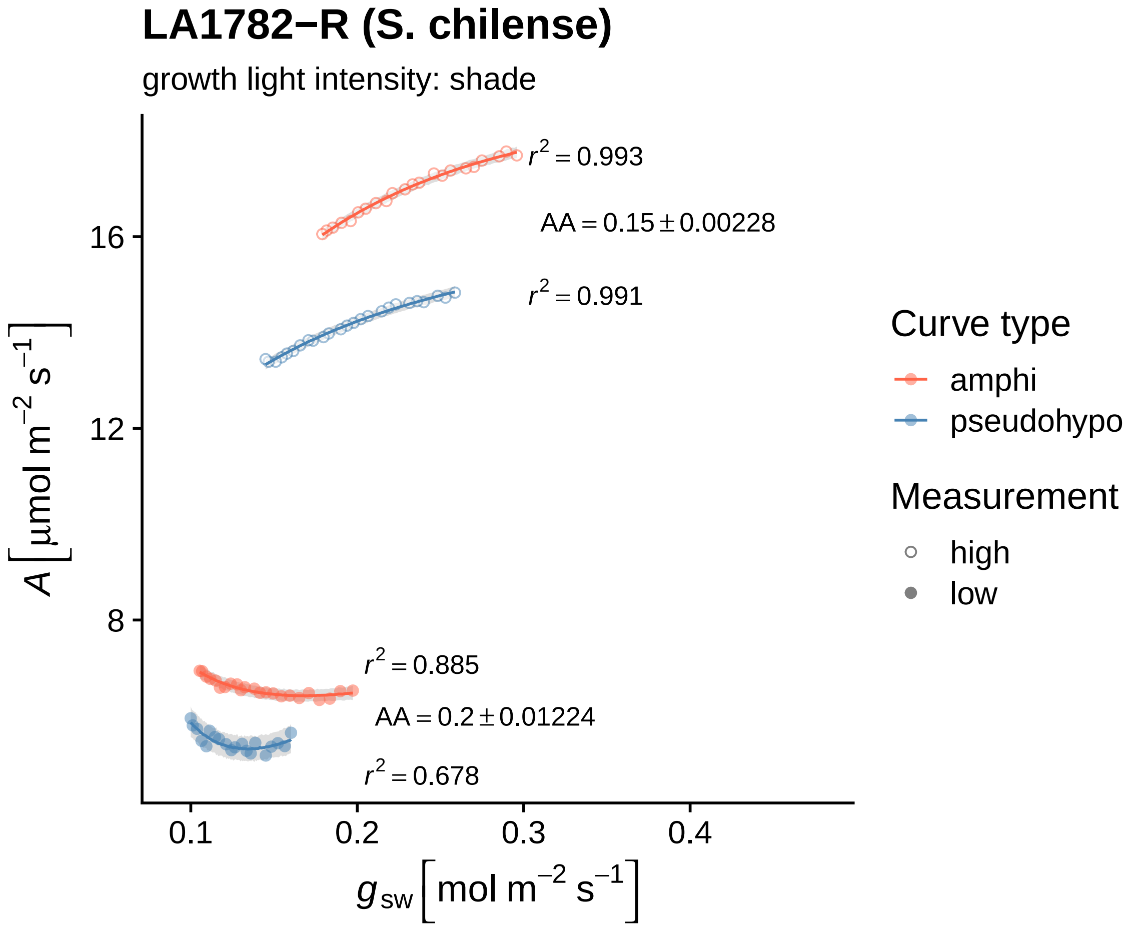

LA1782−R (S. chilense)
growth light intensity: shade
r² = 0.993
AA = 0.15 ± 0.00228
r² = 0.991
r² = 0.885
AA = 0.2 ± 0.01224
r² = 0.678
16
12
8
0.1
0.2
0.3
0.4
A [μmol m⁻² s⁻¹]
g_sw [mol m⁻² s⁻¹]
Curve type
amphi
pseudohypo
Measurement
high
low

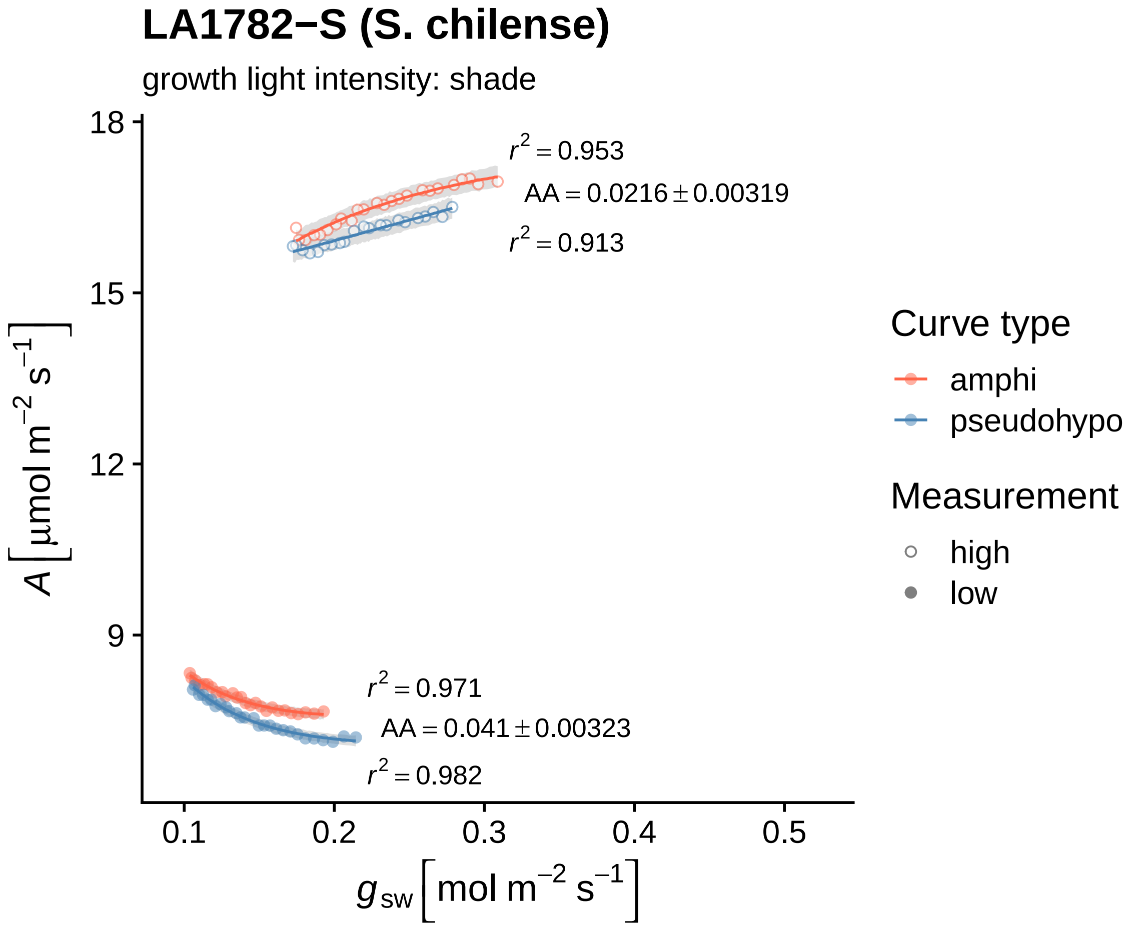

LA1782−S (S. chilense)
growth light intensity: shade
18
15
12
9
$r^2 = 0.953$
$AA = 0.0216 \pm 0.00319$
$r^2 = 0.913$
$r^2 = 0.971$
$AA = 0.041 \pm 0.00323$
$r^2 = 0.982$
0.1
0.2
0.3
0.4
0.5
$g_{sw}$ [mol m$^{-2}$ s$^{-1}$]
$A$ [µmol m$^{-2}$ s$^{-1}$]
Curve type
amphi
pseudohypo
Measurement
high
low

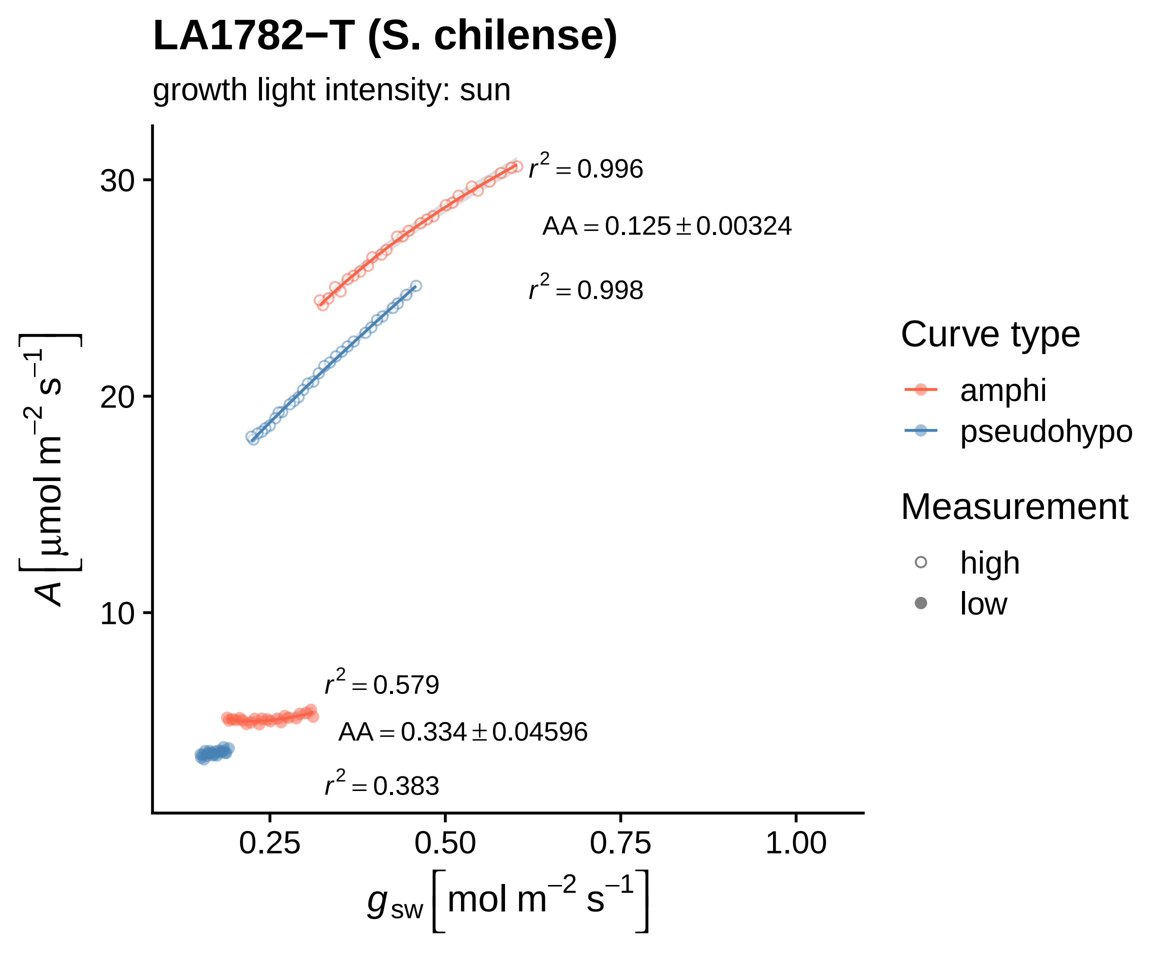

LA1782−T (S. chilense)
growth light intensity: sun
$r^2 = 0.996$
$AA = 0.125 \pm 0.00324$
$r^2 = 0.998$
$r^2 = 0.579$
$AA = 0.334 \pm 0.04596$
$r^2 = 0.383$
30
20
10
0.25
0.50
0.75
1.00
$A\ [\mu mol\ m^{-2}\ s^{-1}]$
$g_{sw}\ [mol\ m^{-2}\ s^{-1}]$
Curve type
amphi
pseudohypo
Measurement
high
low

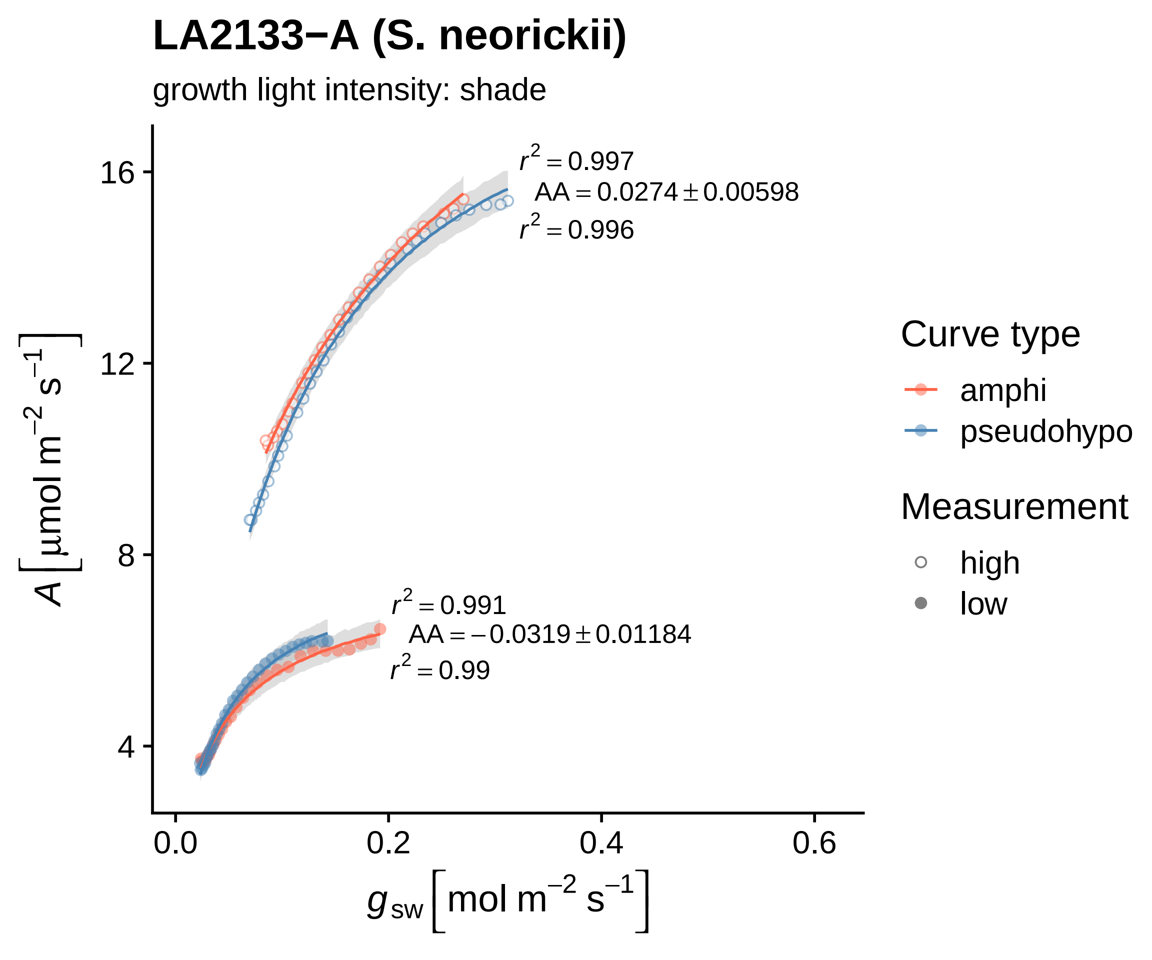
LA2133−A (S. neorickii)
growth light intensity: shade
$r^2 = 0.997$
$AA = 0.0274 \pm 0.00598$
$r^2 = 0.996$
$r^2 = 0.991$
$AA = -0.0319 \pm 0.01184$
$r^2 = 0.99$
16
12
8
4
0.0
0.2
0.4
0.6
$A\ [\mu mol\ m^{-2}\ s^{-1}]$
$g_{sw}\ [mol\ m^{-2}\ s^{-1}]$
Curve type
amphi
pseudohypo
Measurement
high
low

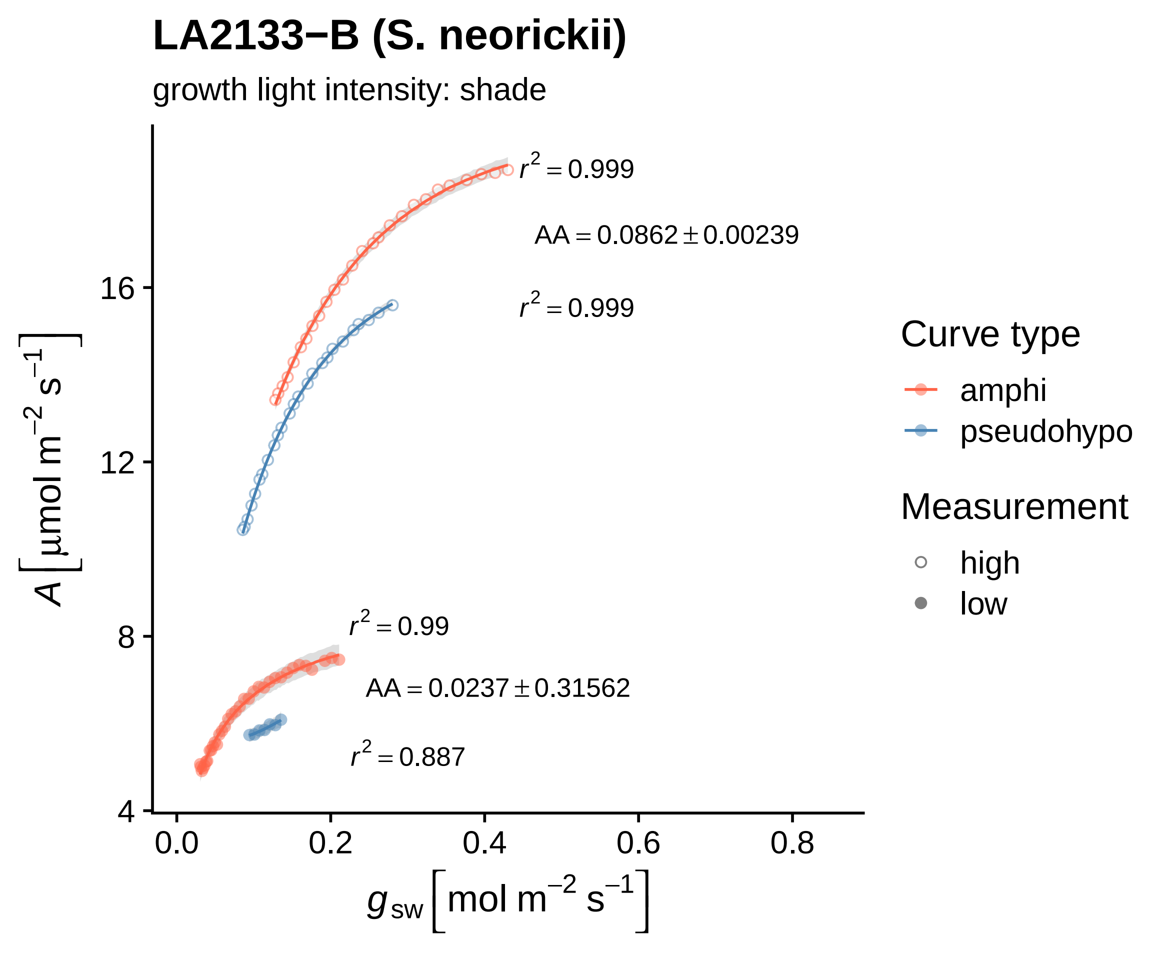

LA2133−B (S. neorickii)
growth light intensity: shade
$r^2 = 0.999$
$AA = 0.0862 \pm 0.00239$
$r^2 = 0.999$
$r^2 = 0.99$
$AA = 0.0237 \pm 0.31562$
$r^2 = 0.887$
16
12
8
4
0.0
0.2
0.4
0.6
0.8
$A \left[\mu mol\ m^{-2}\ s^{-1}\right]$
$g_{sw} \left[mol\ m^{-2}\ s^{-1}\right]$
Curve type
amphi
pseudohypo
Measurement
high
low

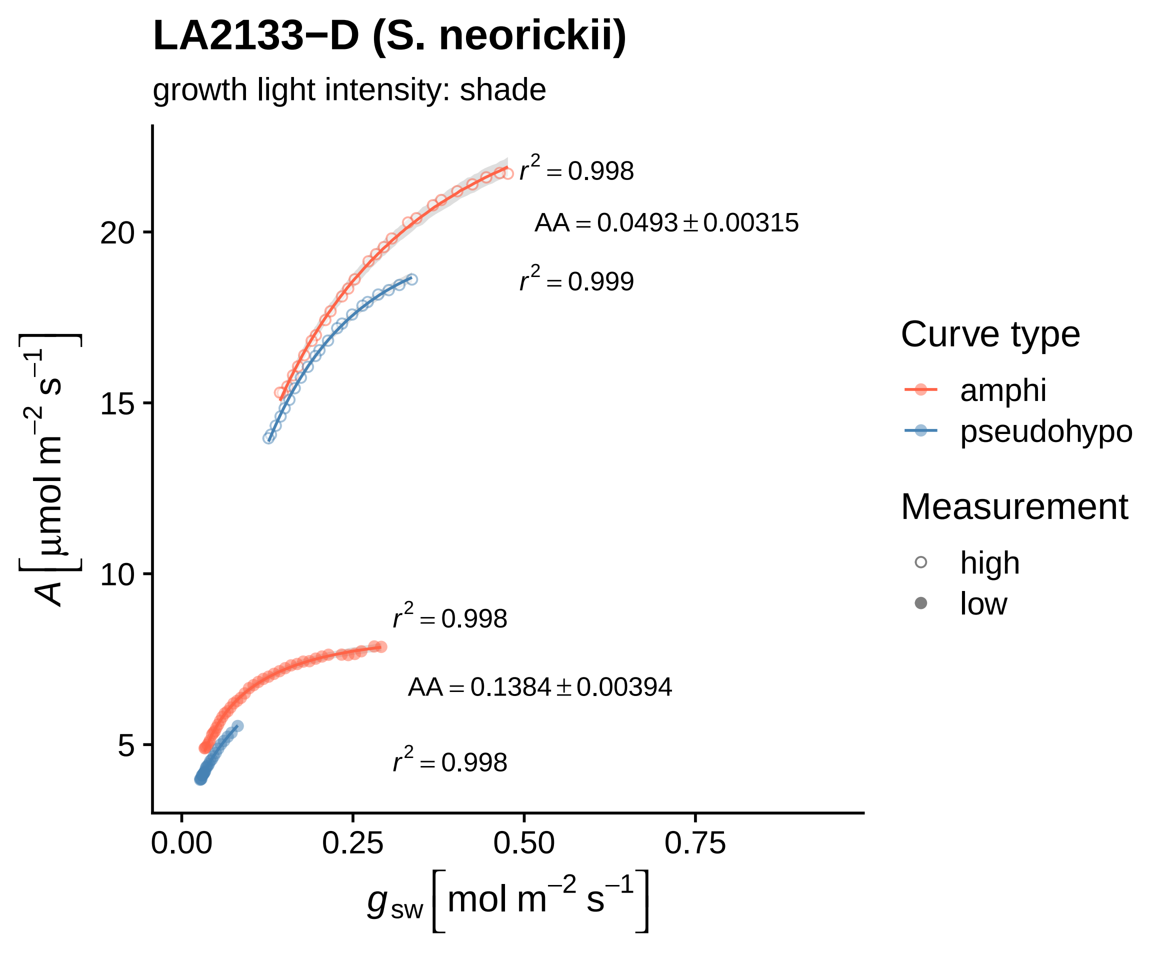

LA2133−D (S. neorickii)
growth light intensity: shade
$r^2 = 0.998$
$AA = 0.0493 \pm 0.00315$
$r^2 = 0.999$
$r^2 = 0.998$
$AA = 0.1384 \pm 0.00394$
$r^2 = 0.998$
20
15
10
5
0.00
0.25
0.50
0.75
$A \left[\mu mol\ m^{-2}\ s^{-1}\right]$
$g_{sw} \left[mol\ m^{-2}\ s^{-1}\right]$
Curve type
amphi
pseudohypo
Measurement
high
low

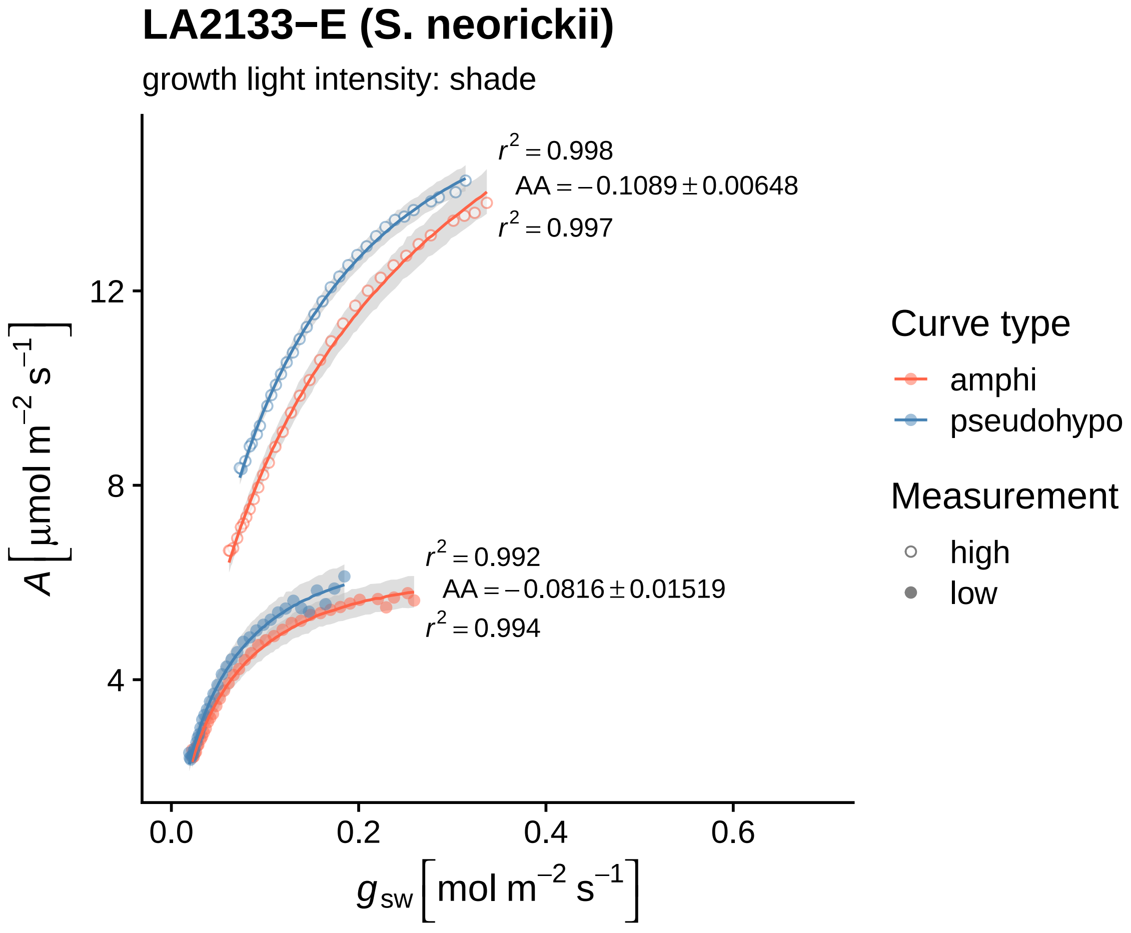

LA2133−E (S. neorickii)
growth light intensity: shade
r² = 0.998
AA = −0.1089 ± 0.00648
r² = 0.997
r² = 0.992
AA = −0.0816 ± 0.01519
r² = 0.994
12
8
4
0.0
0.2
0.4
0.6
A [μmol m⁻² s⁻¹]
g_sw [mol m⁻² s⁻¹]
Curve type
amphi
pseudohypo
Measurement
high
low

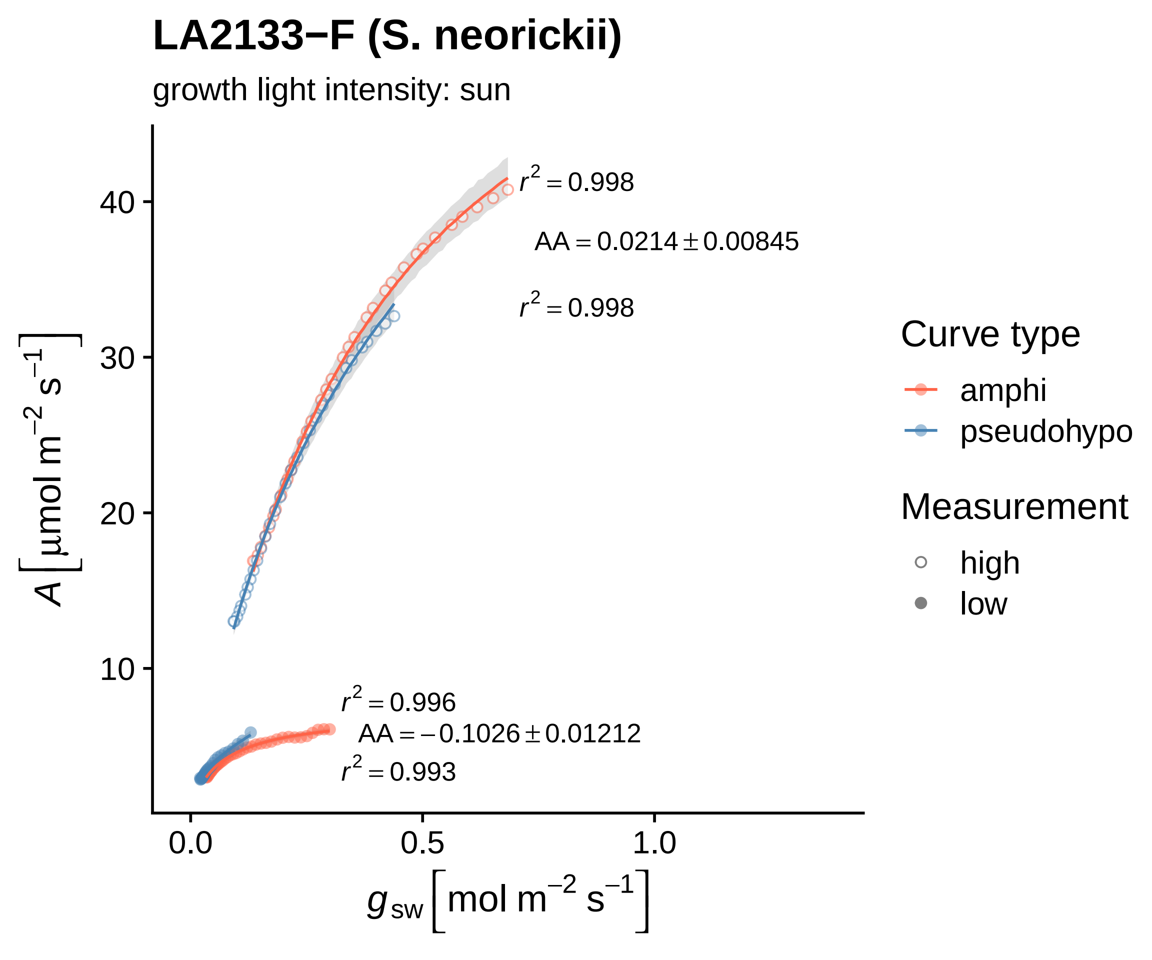
LA2133−F (S. neorickii)
growth light intensity: sun
$r^2 = 0.998$
$AA = 0.0214 \pm 0.00845$
$r^2 = 0.998$
$r^2 = 0.996$
$AA = -0.1026 \pm 0.01212$
$r^2 = 0.993$
40
30
20
10
0.0
0.5
1.0
$A\ [\mu mol\ m^{-2}\ s^{-1}]$
$g_{sw}\ [mol\ m^{-2}\ s^{-1}]$
Curve type
amphi
pseudohypo
Measurement
high
low

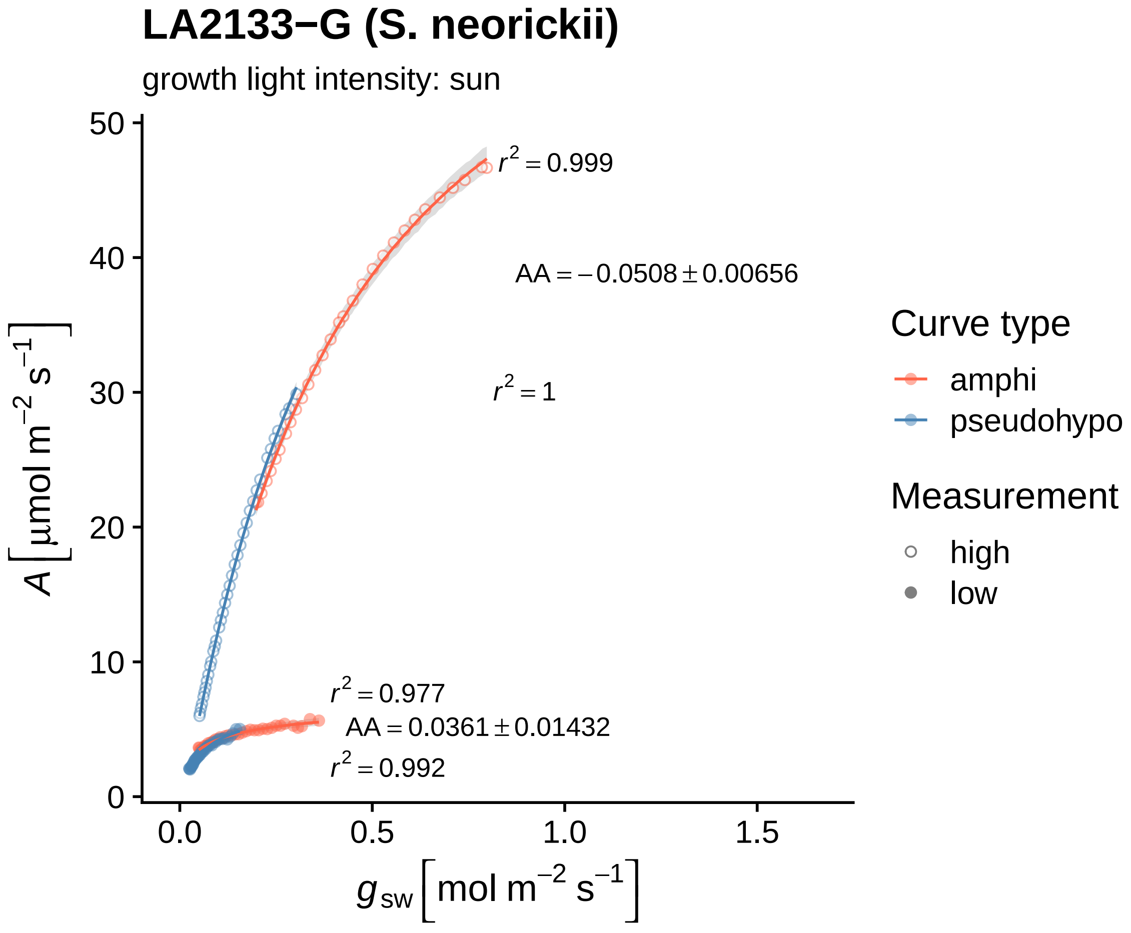

LA2133−G (S. neorickii)
growth light intensity: sun
$r^2 = 0.999$
$AA = -0.0508 \pm 0.00656$
$r^2 = 1$
$r^2 = 0.977$
$AA = 0.0361 \pm 0.01432$
$r^2 = 0.992$
$A\left[\mu mol\ m^{-2}\ s^{-1}\right]$
$g_{sw}\left[mol\ m^{-2}\ s^{-1}\right]$
0
10
20
30
40
50
0.0
0.5
1.0
1.5
Curve type
amphi
pseudohypo
Measurement
high
low

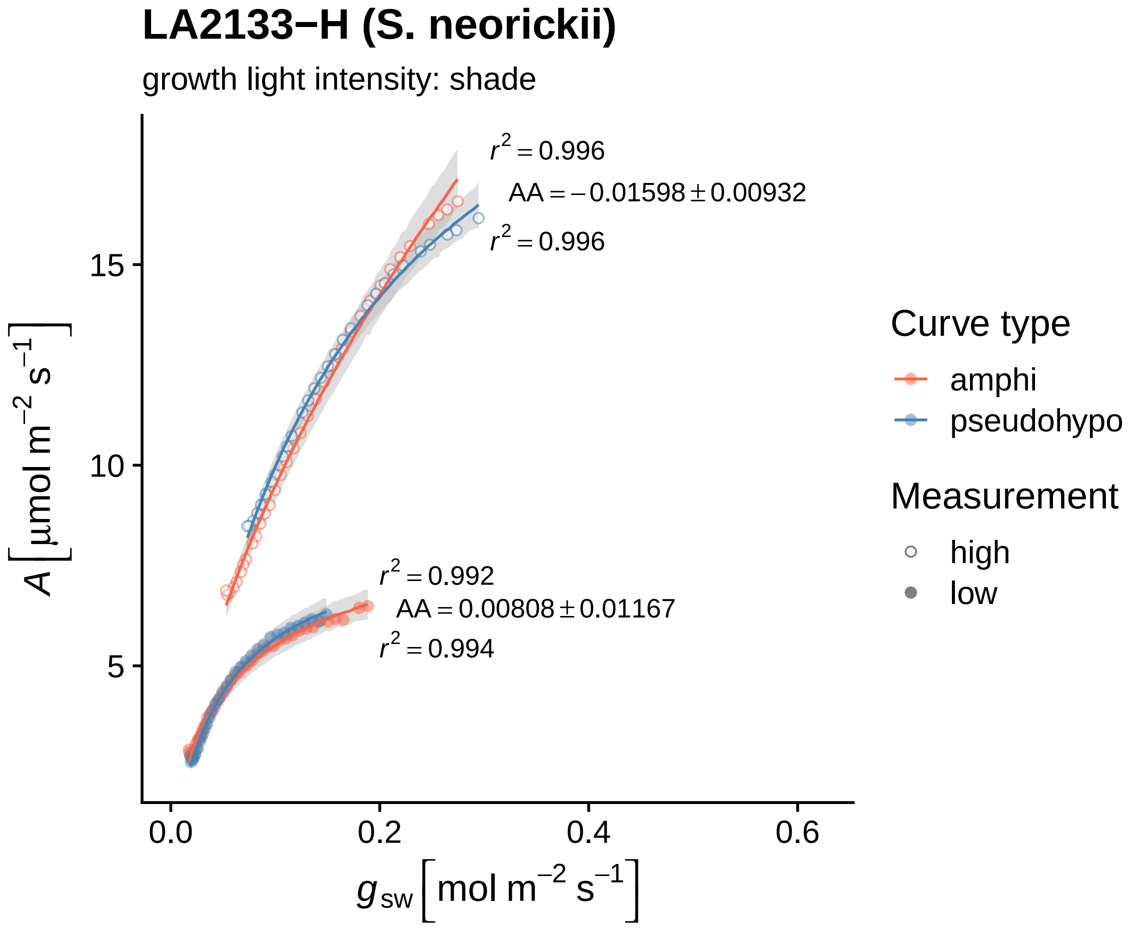

LA2133−H (S. neorickii)
growth light intensity: shade
r² = 0.996
AA = −0.01598 ± 0.00932
r² = 0.996
r² = 0.992
AA = 0.00808 ± 0.01167
r² = 0.994
15
10
5
0.0
0.2
0.4
0.6
A [μmol m⁻² s⁻¹]
g_sw [mol m⁻² s⁻¹]
Curve type
amphi
pseudohypo
Measurement
high
low

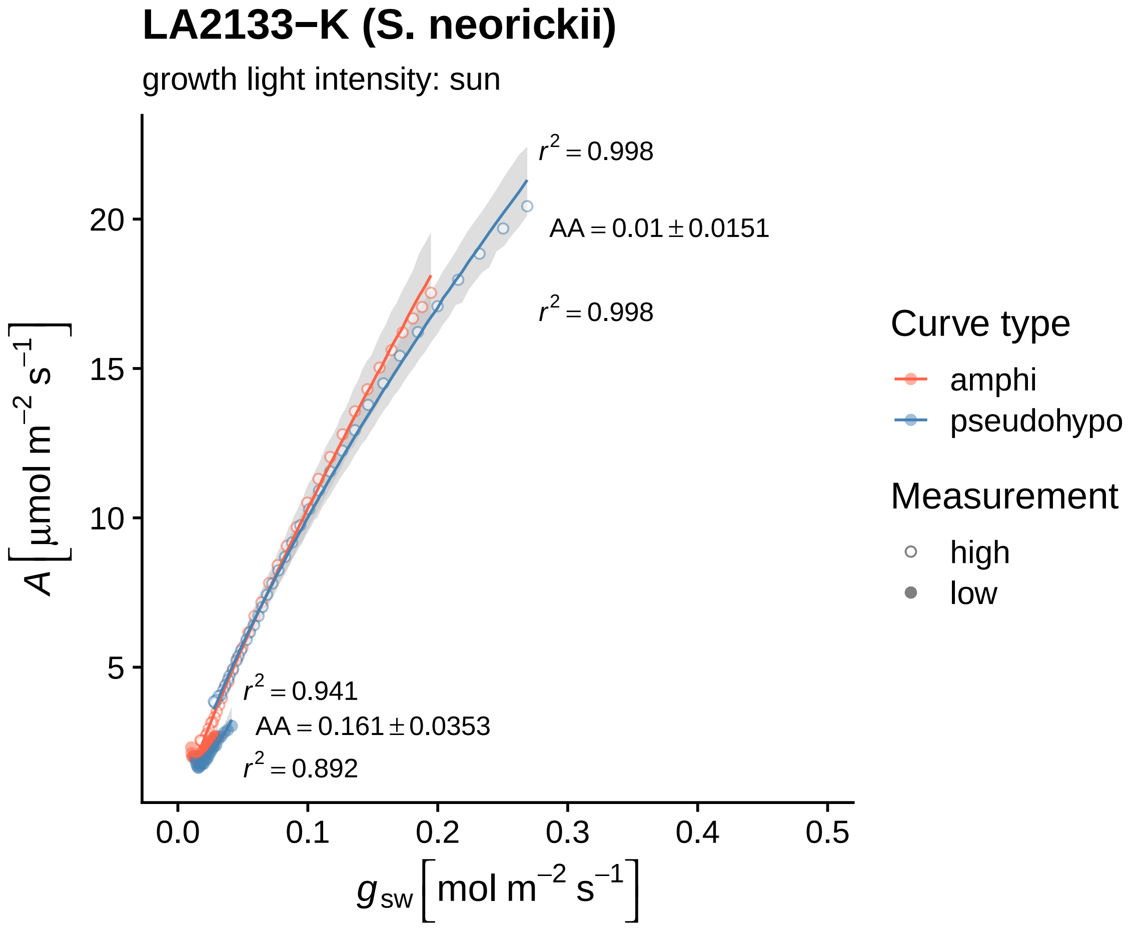

LA2133−K (S. neorickii)
growth light intensity: sun
r² = 0.998
AA = 0.01 ± 0.0151
r² = 0.998
r² = 0.941
AA = 0.161 ± 0.0353
r² = 0.892
A [μmol m⁻² s⁻¹]
g_sw [mol m⁻² s⁻¹]
20
15
10
5
0.0
0.1
0.2
0.3
0.4
0.5
Curve type
amphi
pseudohypo
Measurement
high
low

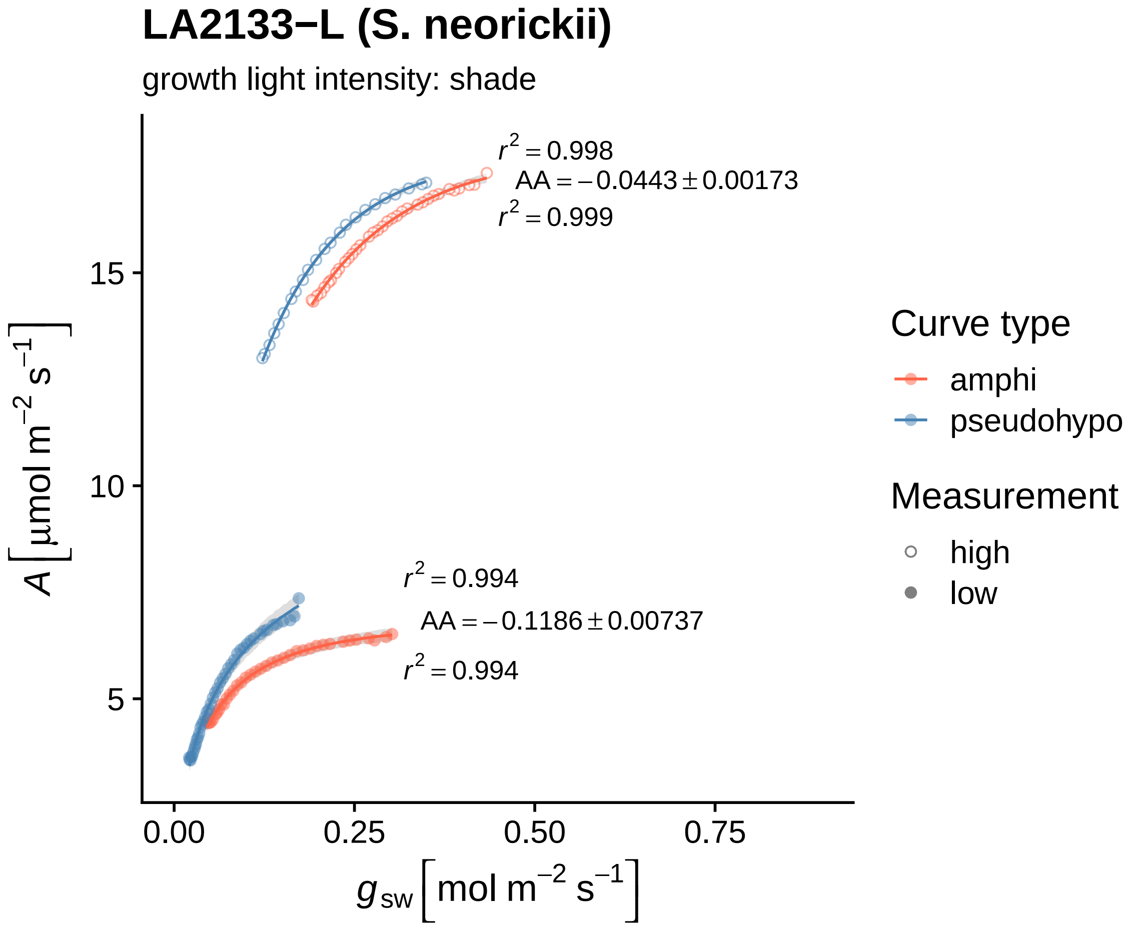
LA2133−L (S. neorickii)
growth light intensity: shade
$r^2 = 0.998$
$AA = -0.0443 \pm 0.00173$
$r^2 = 0.999$
$r^2 = 0.994$
$AA = -0.1186 \pm 0.00737$
$r^2 = 0.994$
15
10
5
0.00
0.25
0.50
0.75
$A\ [\mu mol\ m^{-2}\ s^{-1}]$
$g_{sw}\ [mol\ m^{-2}\ s^{-1}]$
Curve type
amphi
pseudohypo
Measurement
high
low

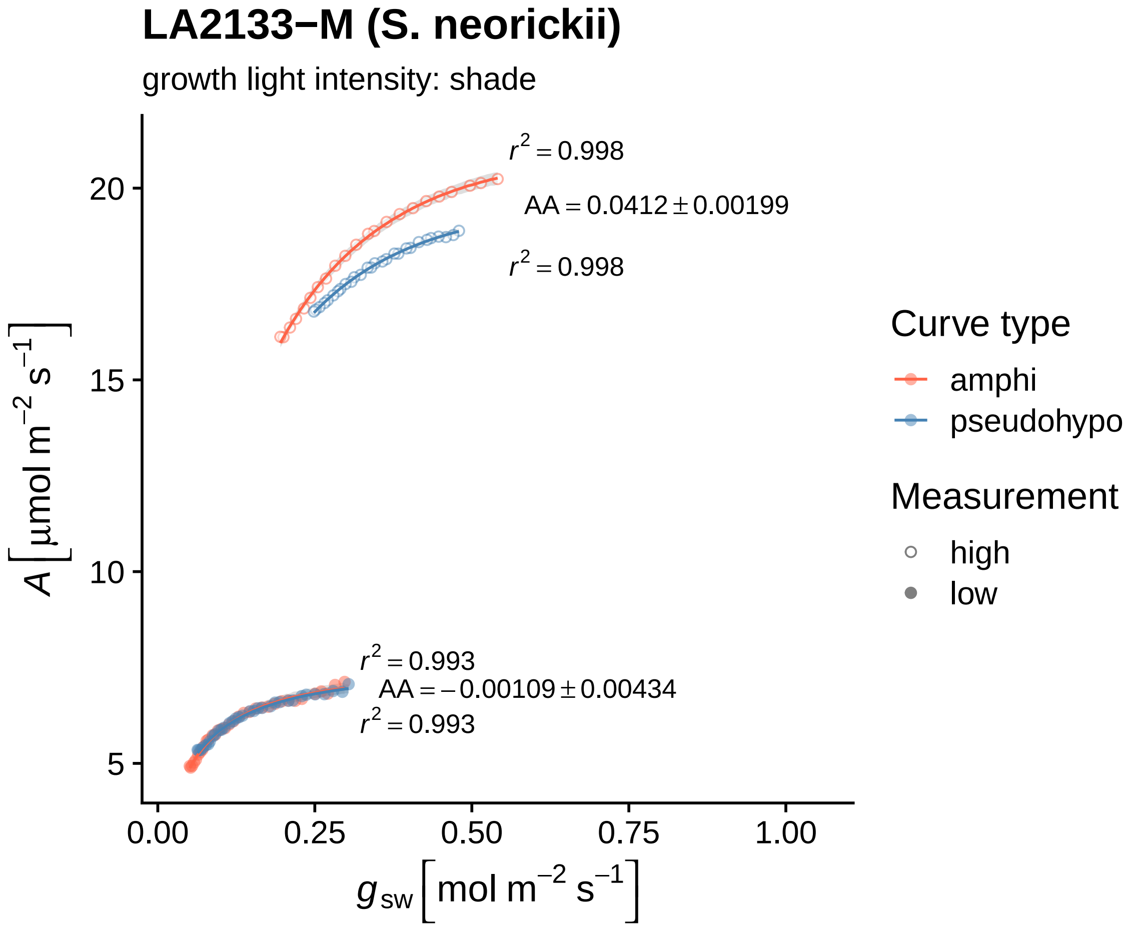

LA2133−M (S. neorickii)
growth light intensity: shade
r² = 0.998
AA = 0.0412 ± 0.00199
r² = 0.998
r² = 0.993
AA = −0.00109 ± 0.00434
r² = 0.993
20
15
10
5
0.00
0.25
0.50
0.75
1.00
A [μmol m⁻² s⁻¹]
g_sw [mol m⁻² s⁻¹]
Curve type
amphi
pseudohypo
Measurement
high
low

**LA2133−N (S. neorickii)**

growth light intensity: shade

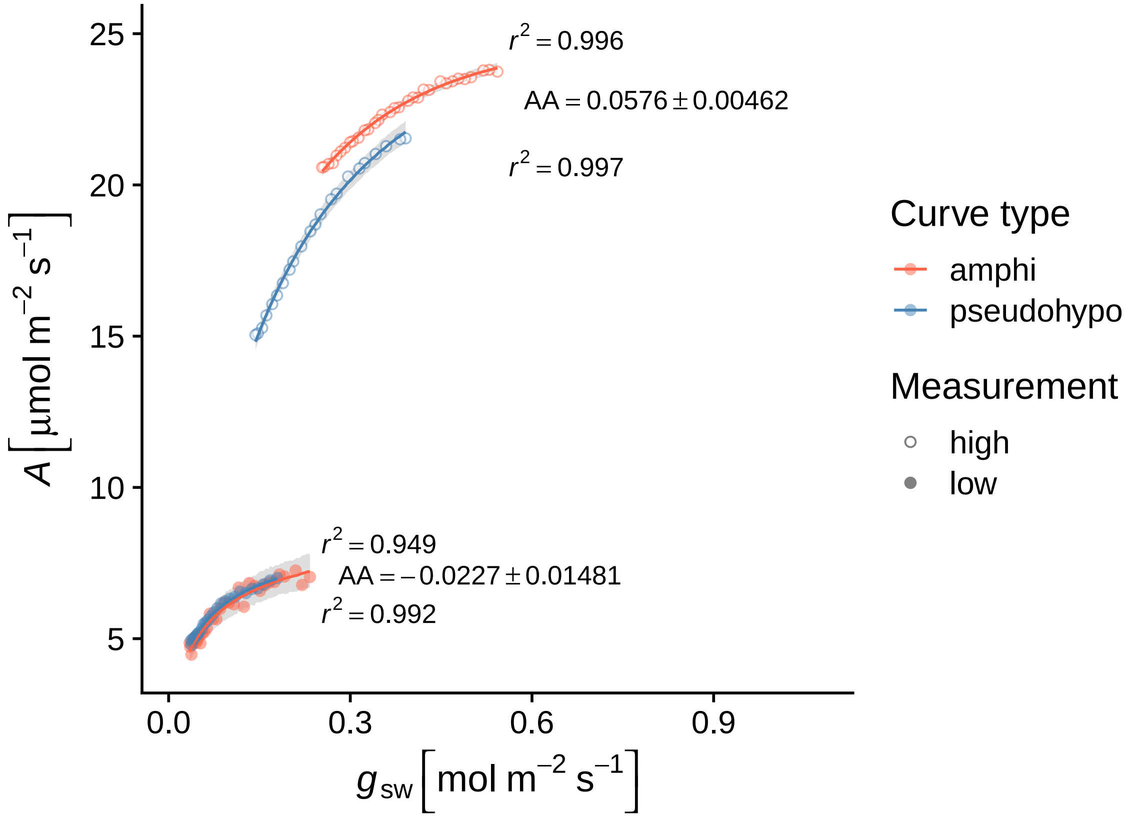

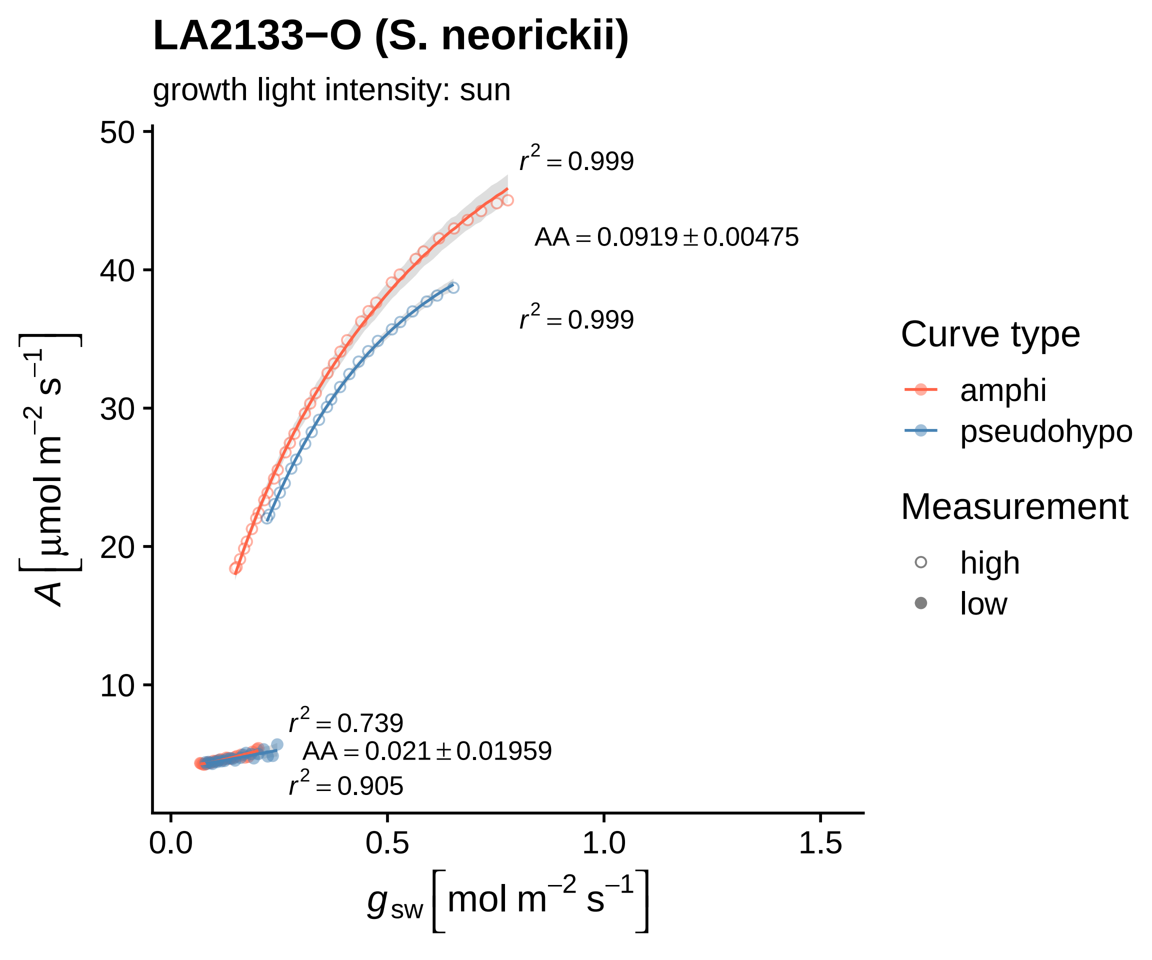

LA2133−O (S. neorickii)
growth light intensity: sun
$r^2 = 0.999$
$AA = 0.0919 \pm 0.00475$
$r^2 = 0.999$
$r^2 = 0.739$
$AA = 0.021 \pm 0.01959$
$r^2 = 0.905$
50
40
30
20
10
0.0
0.5
1.0
1.5
$A\left[\mu mol\ m^{-2}\ s^{-1}\right]$
$g_{sw}\left[mol\ m^{-2}\ s^{-1}\right]$
Curve type
amphi
pseudohypo
Measurement
high
low

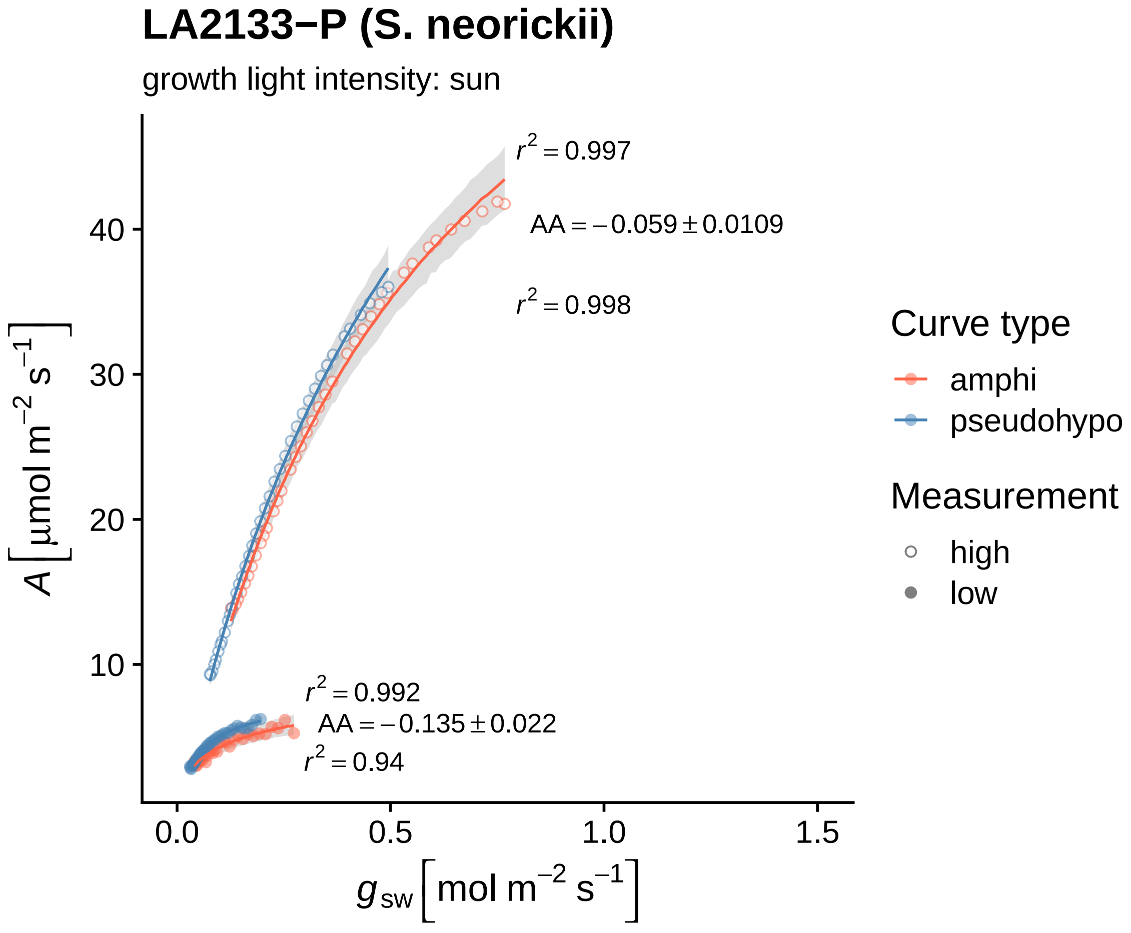
LA2133−P (S. neorickii)
growth light intensity: sun
$r^2 = 0.997$
$AA = -0.059 \pm 0.0109$
$r^2 = 0.998$
$r^2 = 0.992$
$AA = -0.135 \pm 0.022$
$r^2 = 0.94$
$A\ [\mu mol\ m^{-2}\ s^{-1}]$
$g_{sw}\ [mol\ m^{-2}\ s^{-1}]$
10
20
30
40
0.0
0.5
1.0
1.5
Curve type
amphi
pseudohypo
Measurement
high
low

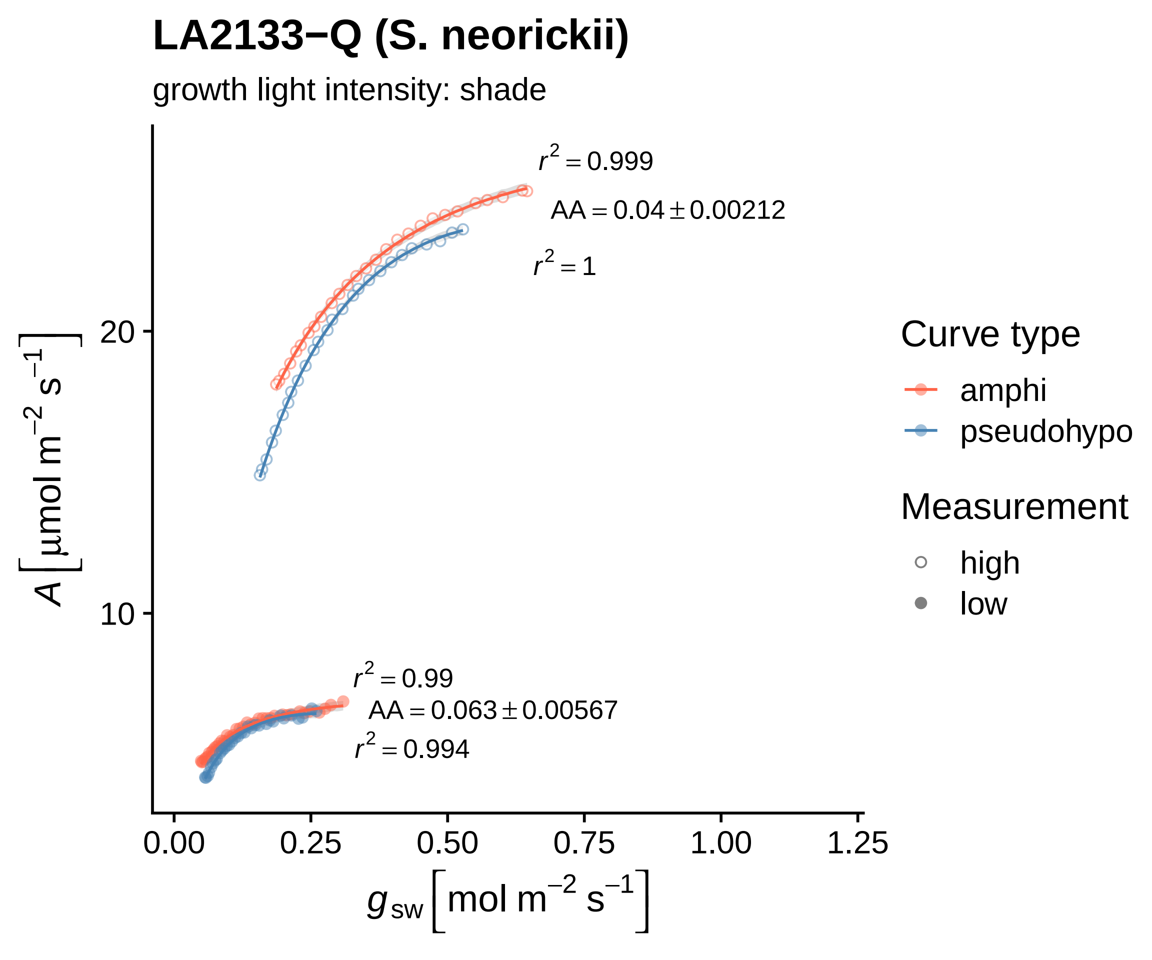

LA2133−Q (S. neorickii)
growth light intensity: shade
r2 = 0.999
AA = 0.04 ± 0.00212
r2 = 1
r2 = 0.99
AA = 0.063 ± 0.00567
r2 = 0.994
20
10
0.00
0.25
0.50
0.75
1.00
1.25
A [μmol m−2 s−1]
gsw [mol m−2 s−1]
Curve type
amphi
pseudohypo
Measurement
high
low

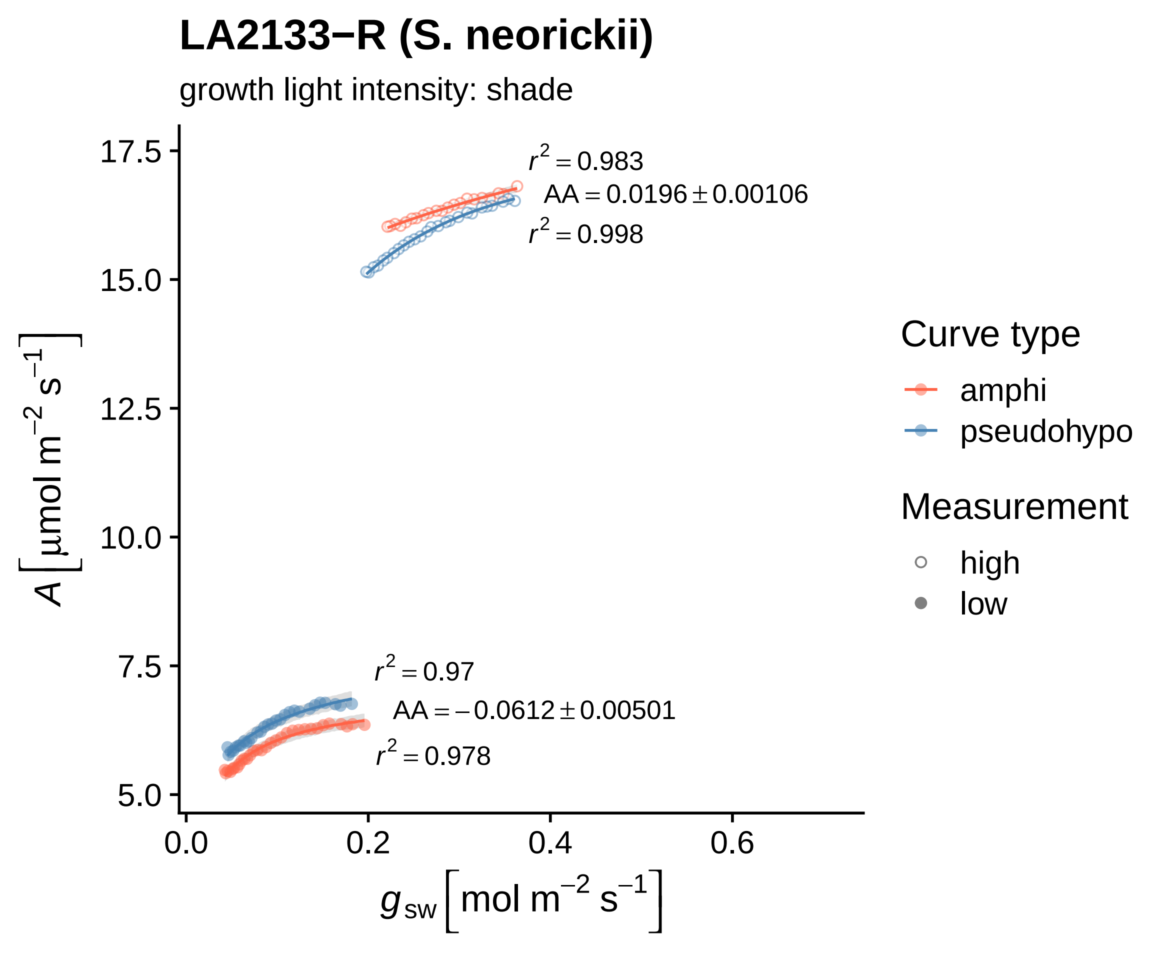

LA2133−R (S. neorickii)
growth light intensity: shade
$r^2 = 0.983$
$AA = 0.0196 \pm 0.00106$
$r^2 = 0.998$
$r^2 = 0.97$
$AA = -0.0612 \pm 0.00501$
$r^2 = 0.978$
17.5
15.0
12.5
10.0
7.5
5.0
0.0
0.2
0.4
0.6
$A\ [\mu mol\ m^{-2}\ s^{-1}]$
$g_{sw}\ [mol\ m^{-2}\ s^{-1}]$
Curve type
amphi
pseudohypo
Measurement
high
low

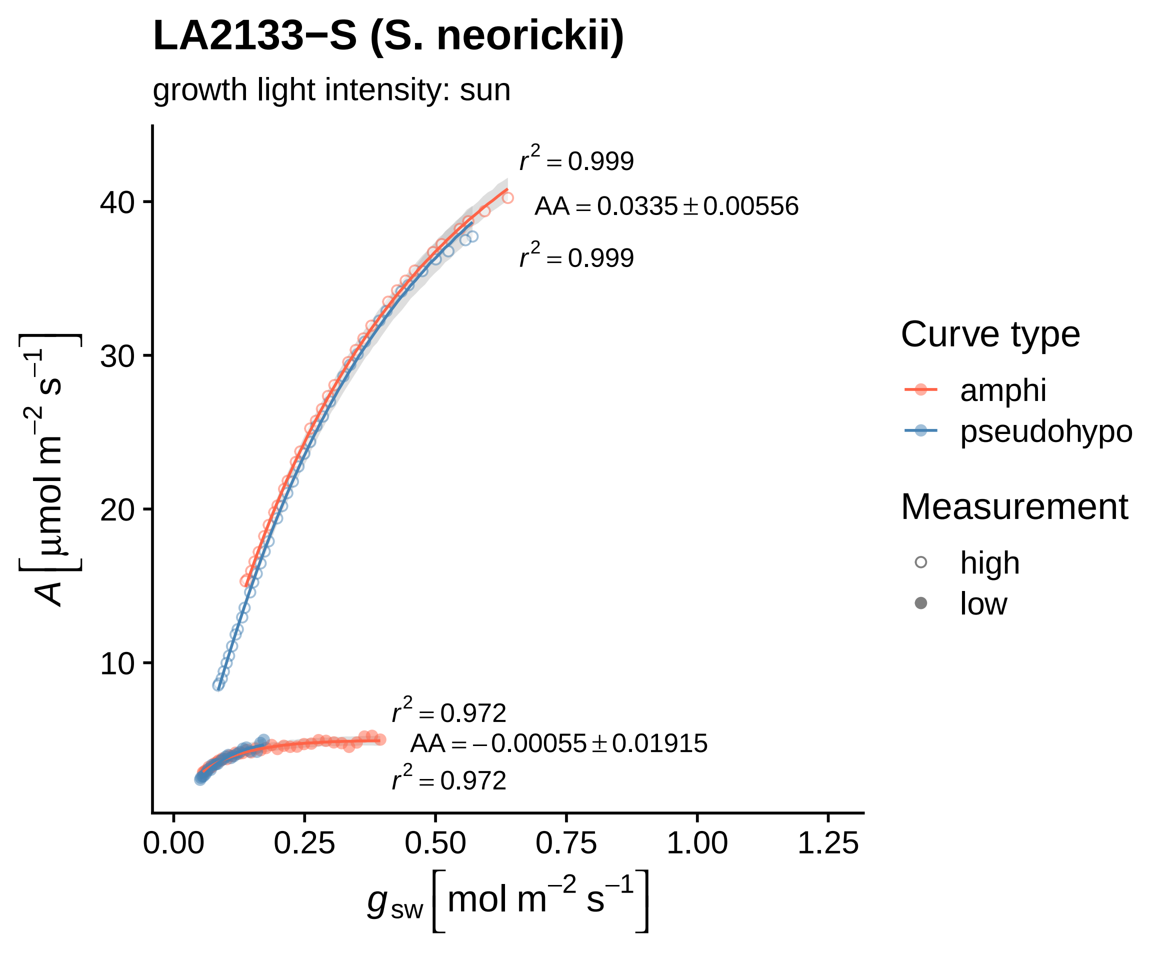
LA2133−S (S. neorickii)
growth light intensity: sun
$r^2 = 0.999$
$AA = 0.0335 \pm 0.00556$
$r^2 = 0.999$
$r^2 = 0.972$
$AA = -0.00055 \pm 0.01915$
$r^2 = 0.972$
10
20
30
40
0.00
0.25
0.50
0.75
1.00
1.25
$A\left[\mu mol\ m^{-2}\ s^{-1}\right]$
$g_{sw}\left[mol\ m^{-2}\ s^{-1}\right]$
Curve type
amphi
pseudohypo
Measurement
high
low

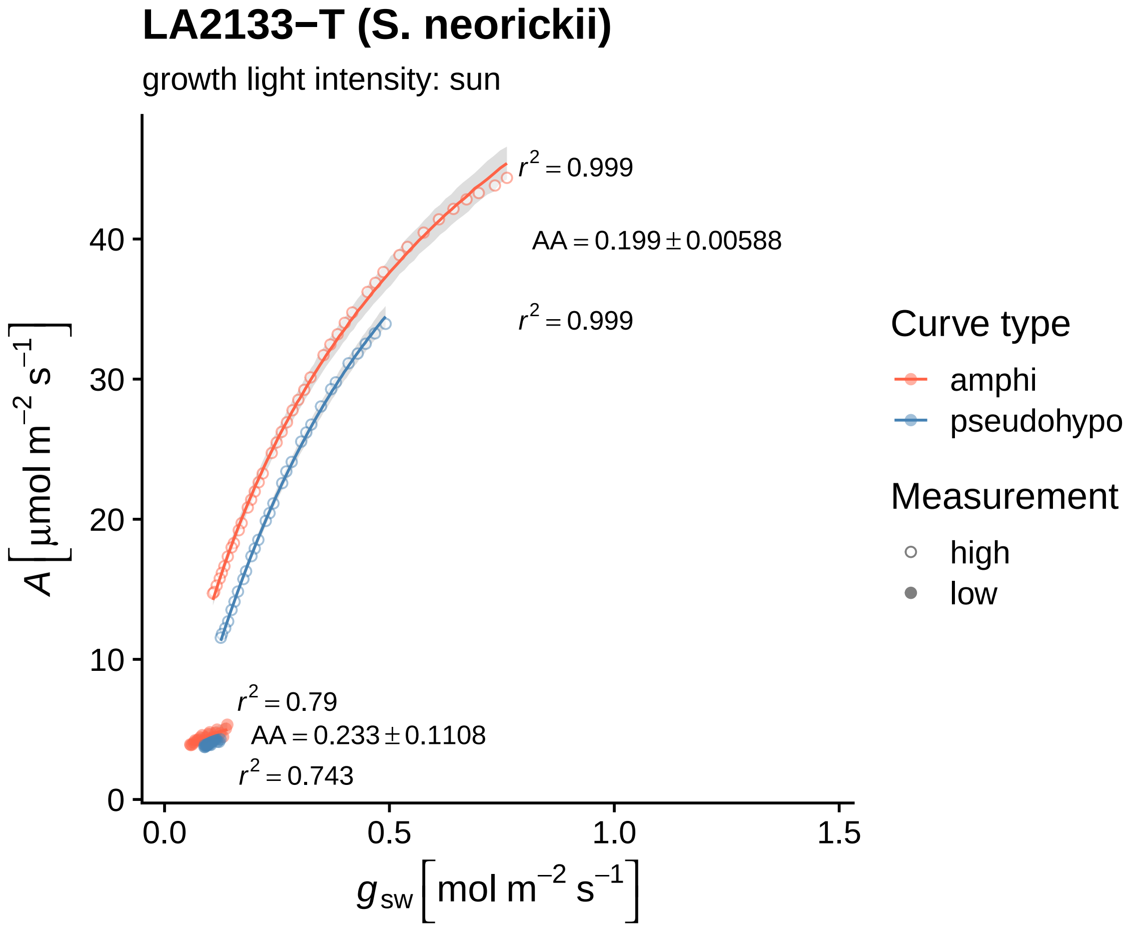
LA2133−T (S. neorickii)
growth light intensity: sun
$r^2 = 0.999$
$AA = 0.199 \pm 0.00588$
$r^2 = 0.999$
$r^2 = 0.79$
$AA = 0.233 \pm 0.1108$
$r^2 = 0.743$
40
30
20
10
0
0.0
0.5
1.0
1.5
$A\ [\mu mol\ m^{-2}\ s^{-1}]$
$g_{sw}\ [mol\ m^{-2}\ s^{-1}]$
Curve type
amphi
pseudohypo
Measurement
high
low

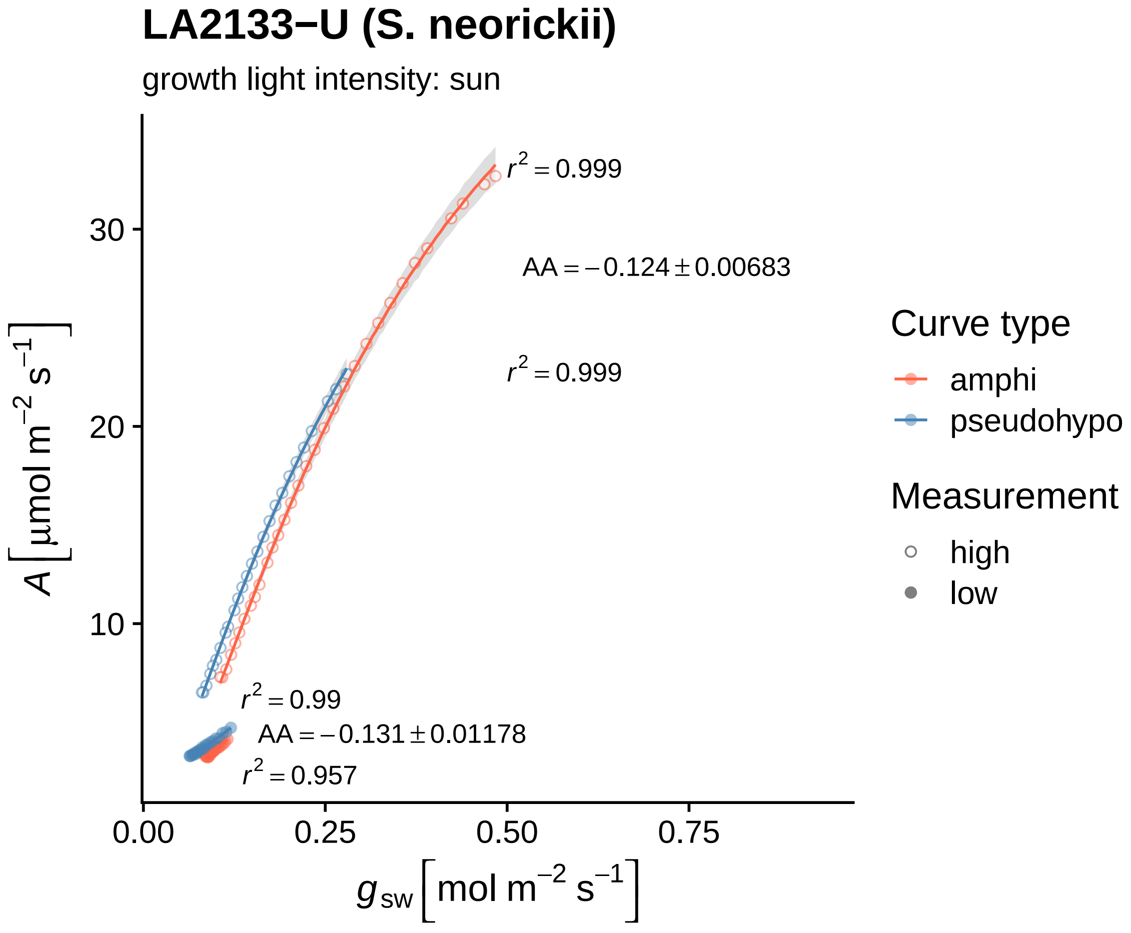
LA2133−U (S. neorickii)
growth light intensity: sun
$r^2 = 0.999$
$AA = -0.124 \pm 0.00683$
$r^2 = 0.999$
$r^2 = 0.99$
$AA = -0.131 \pm 0.01178$
$r^2 = 0.957$
30
20
10
0.00
0.25
0.50
0.75
$A\left[\mu mol\ m^{-2}\ s^{-1}\right]$
$g_{sw}\left[mol\ m^{-2}\ s^{-1}\right]$
Curve type
amphi
pseudohypo
Measurement
high
low

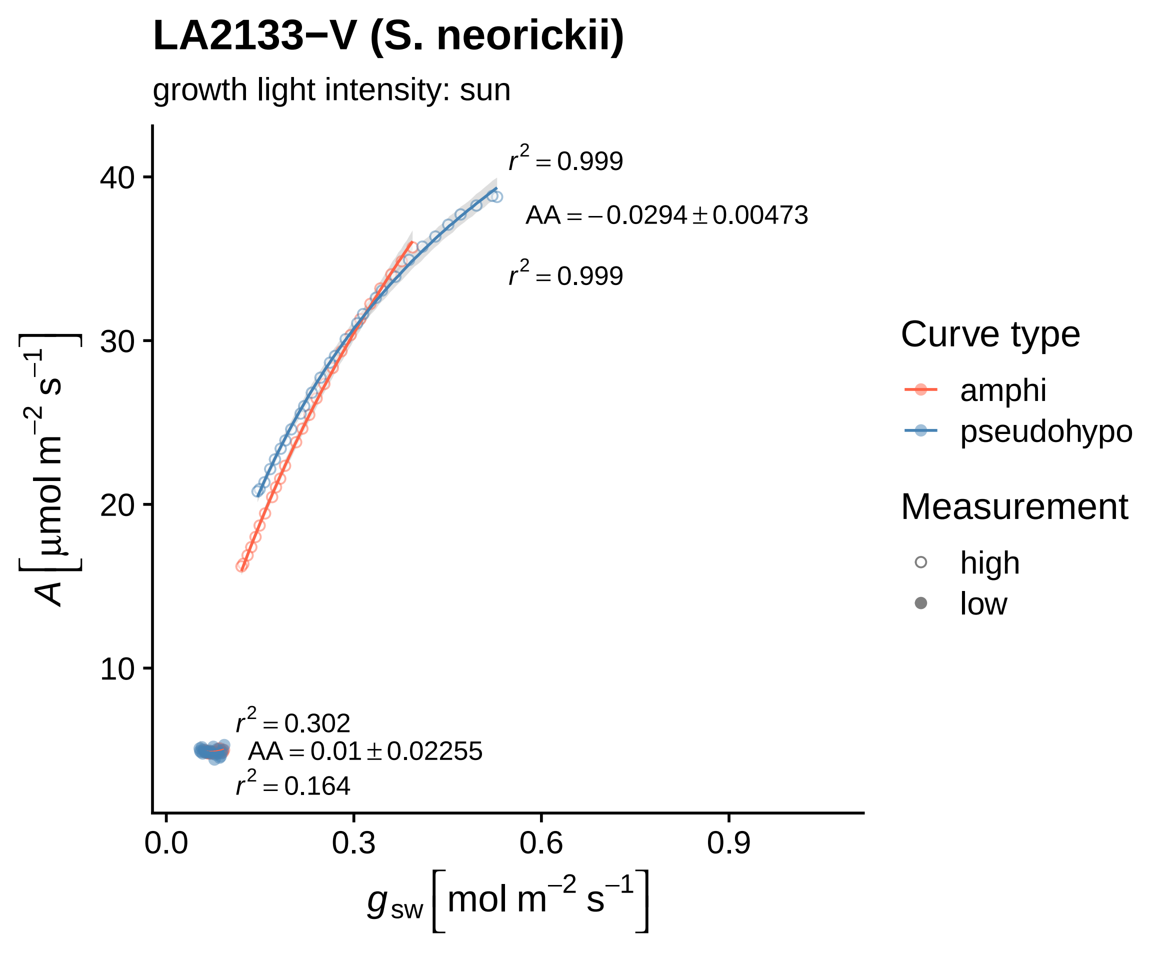

LA2133−V (S. neorickii)
growth light intensity: sun
r² = 0.999
AA = −0.0294 ± 0.00473
r² = 0.999
r² = 0.302
AA = 0.01 ± 0.02255
r² = 0.164
40
30
20
10
0.0
0.3
0.6
0.9
A [μmol m⁻² s⁻¹]
g_sw [mol m⁻² s⁻¹]
Curve type
amphi
pseudohypo
Measurement
high
low

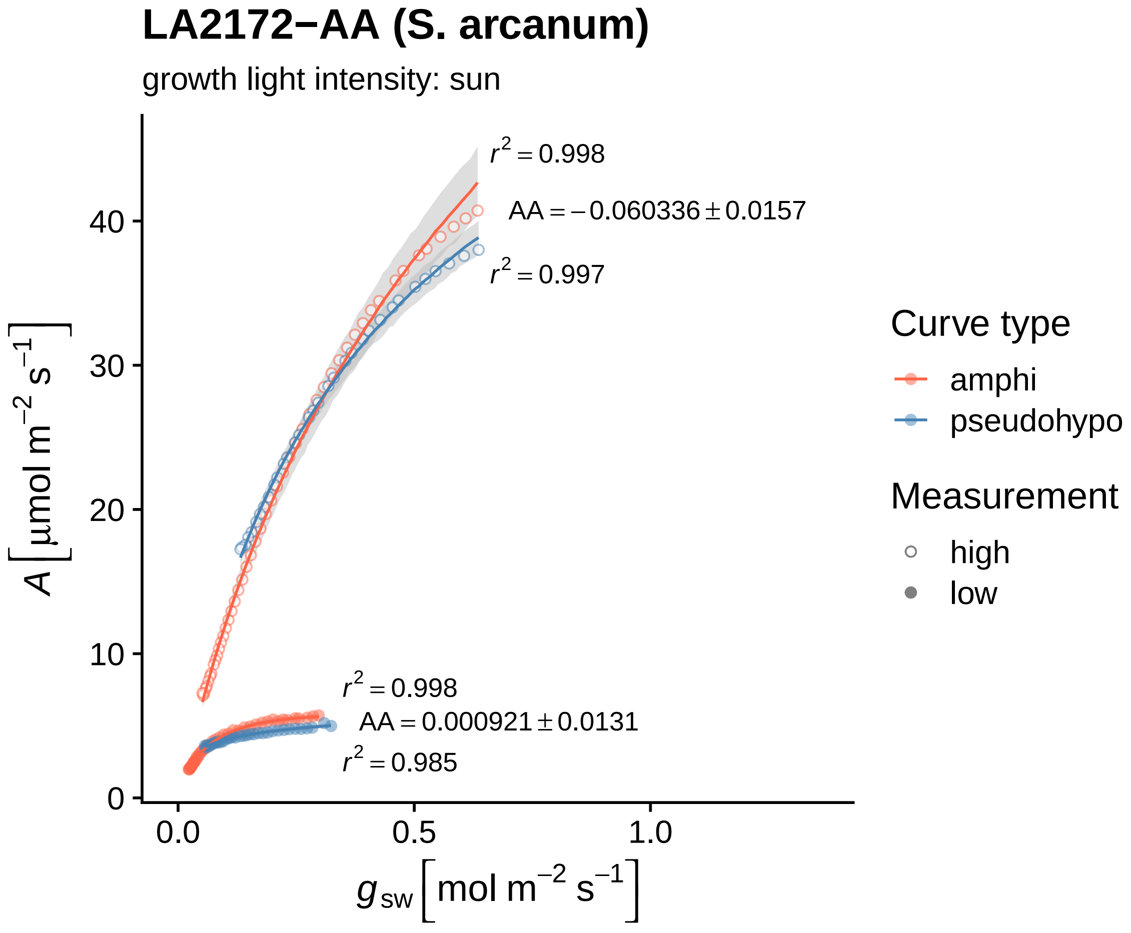

LA2172−AA (S. arcanum)
growth light intensity: sun
$r^2 = 0.998$
$AA = -0.060336 \pm 0.0157$
$r^2 = 0.997$
$r^2 = 0.998$
$AA = 0.000921 \pm 0.0131$
$r^2 = 0.985$
$A\ [\mu mol\ m^{-2}\ s^{-1}]$
$g_{sw}\ [mol\ m^{-2}\ s^{-1}]$
0
10
20
30
40
0.0
0.5
1.0
Curve type
amphi
pseudohypo
Measurement
high
low

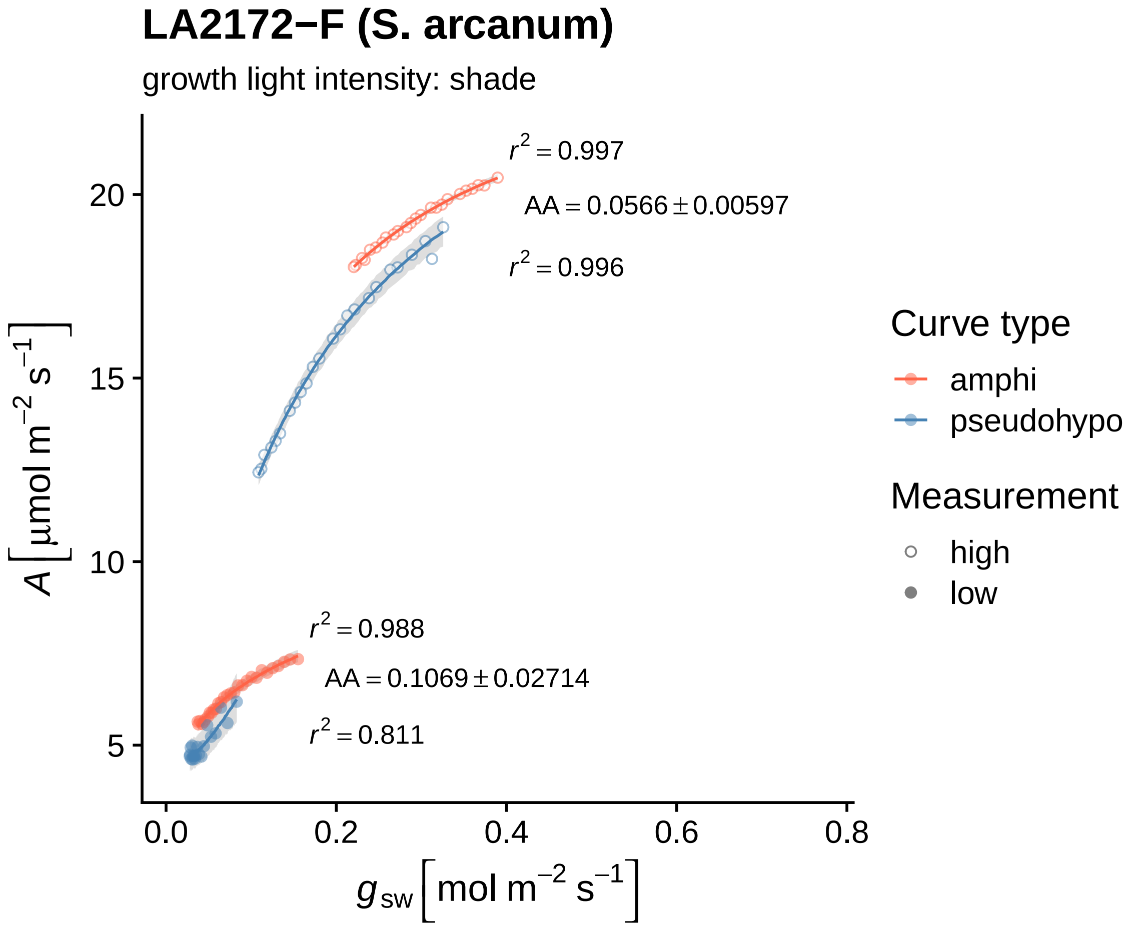

LA2172−F (S. arcanum)
growth light intensity: shade
$r^2 = 0.997$
$AA = 0.0566 \pm 0.00597$
$r^2 = 0.996$
$r^2 = 0.988$
$AA = 0.1069 \pm 0.02714$
$r^2 = 0.811$
$A\left[\mu mol\ m^{-2}\ s^{-1}\right]$
$g_{sw}\left[mol\ m^{-2}\ s^{-1}\right]$
20
15
10
5
0.0
0.2
0.4
0.6
0.8
Curve type
amphi
pseudohypo
Measurement
high
low

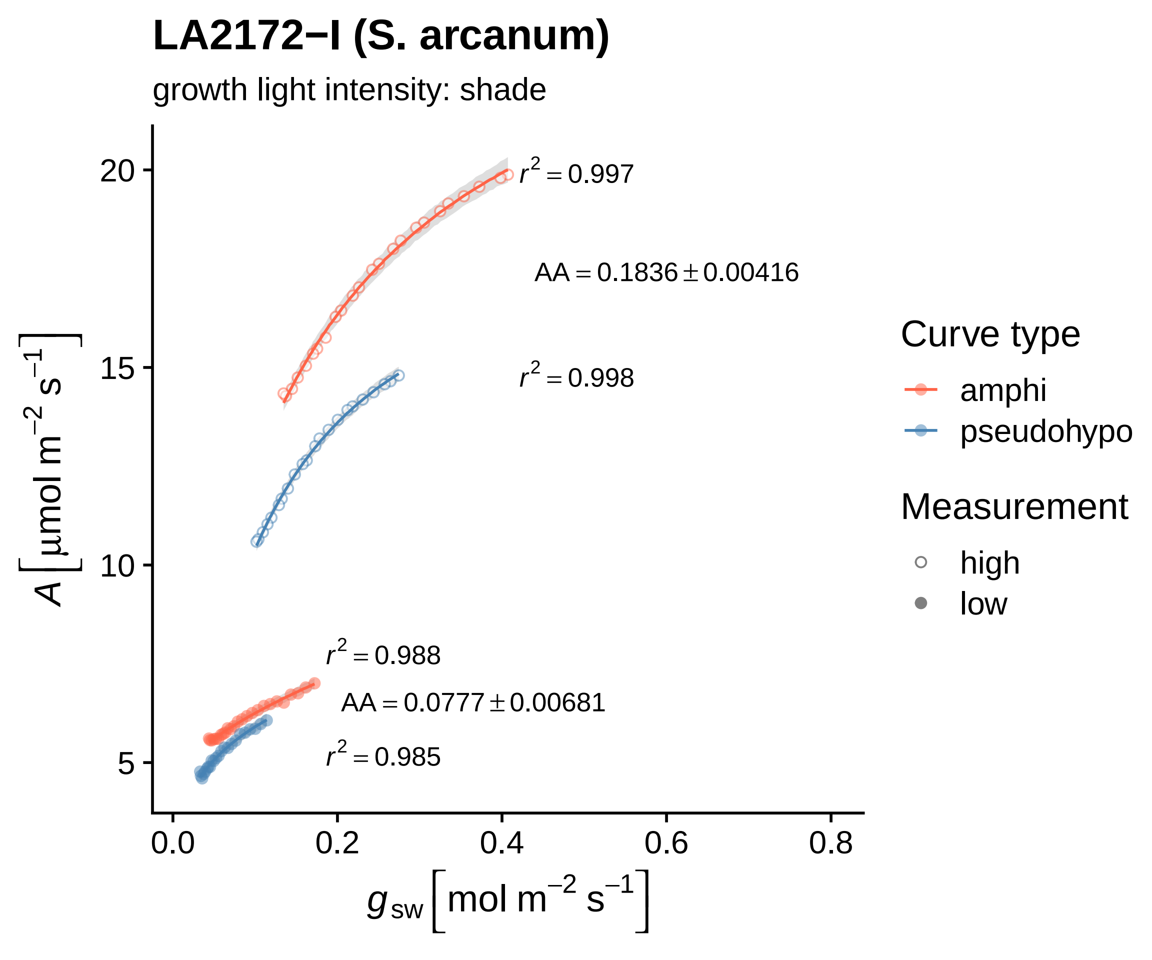

LA2172−I (S. arcanum)
growth light intensity: shade
$r^2 = 0.997$
AA = 0.1836 ± 0.00416
$r^2 = 0.998$
$r^2 = 0.988$
AA = 0.0777 ± 0.00681
$r^2 = 0.985$
20
15
10
5
0.0
0.2
0.4
0.6
0.8
$A\ [\mu mol\ m^{-2}\ s^{-1}]$
$g_{sw}\ [mol\ m^{-2}\ s^{-1}]$
Curve type
amphi
pseudohypo
Measurement
high
low

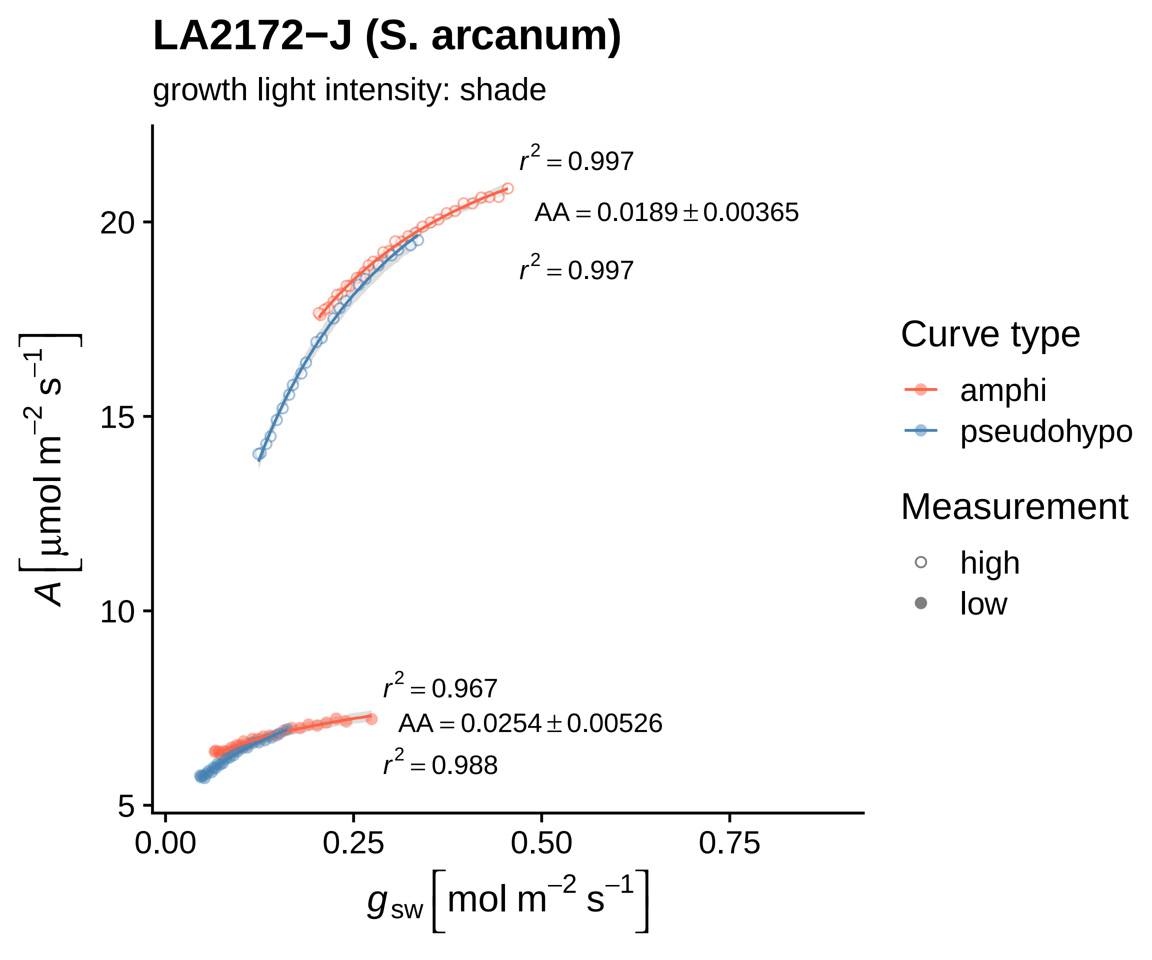

LA2172−J (S. arcanum)
growth light intensity: shade
$r^2 = 0.997$
$AA = 0.0189 \pm 0.00365$
$r^2 = 0.997$
$r^2 = 0.967$
$AA = 0.0254 \pm 0.00526$
$r^2 = 0.988$
$A\left[\mu mol\ m^{-2}\ s^{-1}\right]$
20
15
10
5
0.00
0.25
0.50
0.75
$g_{sw}\left[mol\ m^{-2}\ s^{-1}\right]$
Curve type
amphi
pseudohypo
Measurement
high
low

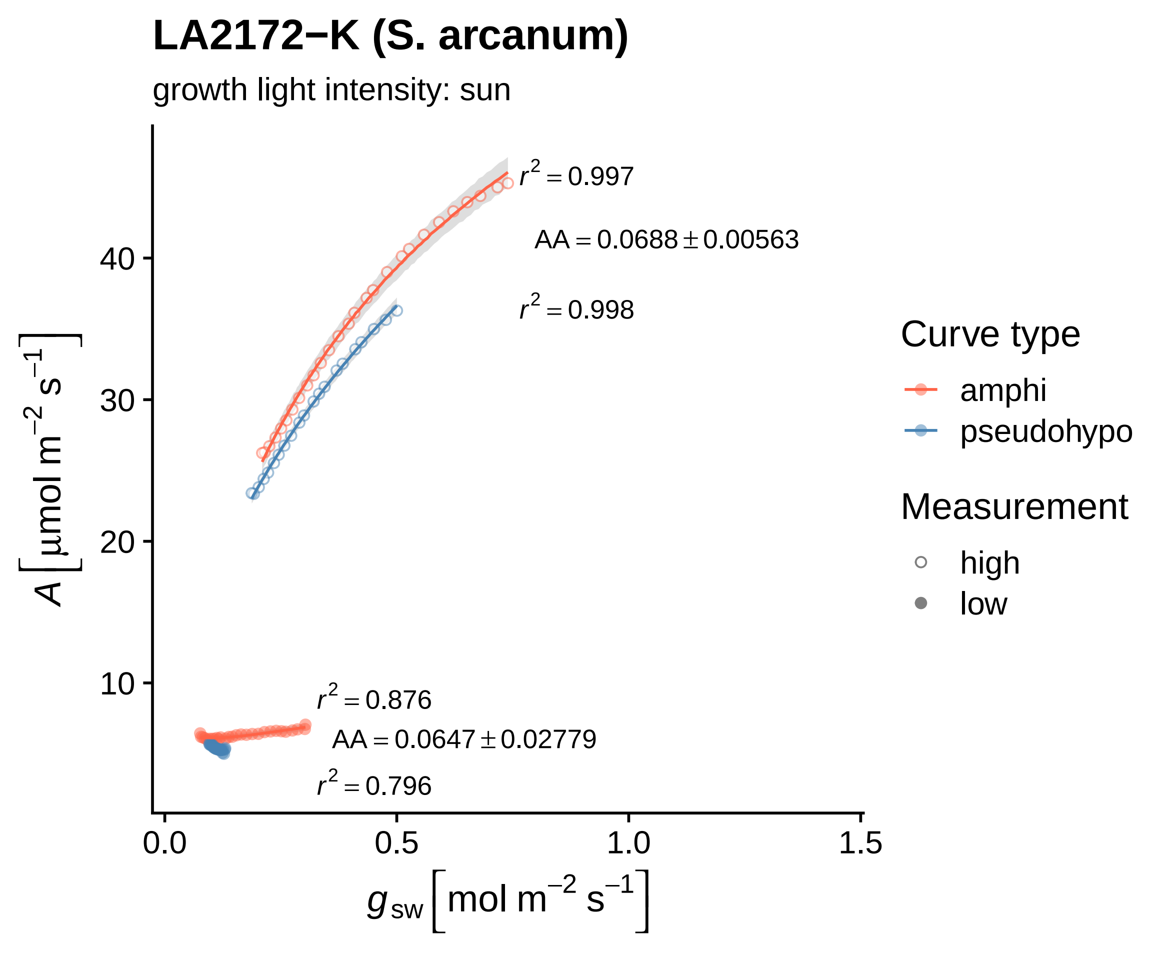

LA2172−K (S. arcanum)
growth light intensity: sun
$r^2 = 0.997$
$AA = 0.0688 \pm 0.00563$
$r^2 = 0.998$
$r^2 = 0.876$
$AA = 0.0647 \pm 0.02779$
$r^2 = 0.796$
40
30
20
10
0.0
0.5
1.0
1.5
$A\left[\mu mol\ m^{-2}\ s^{-1}\right]$
$g_{sw}\left[mol\ m^{-2}\ s^{-1}\right]$
Curve type
amphi
pseudohypo
Measurement
high
low

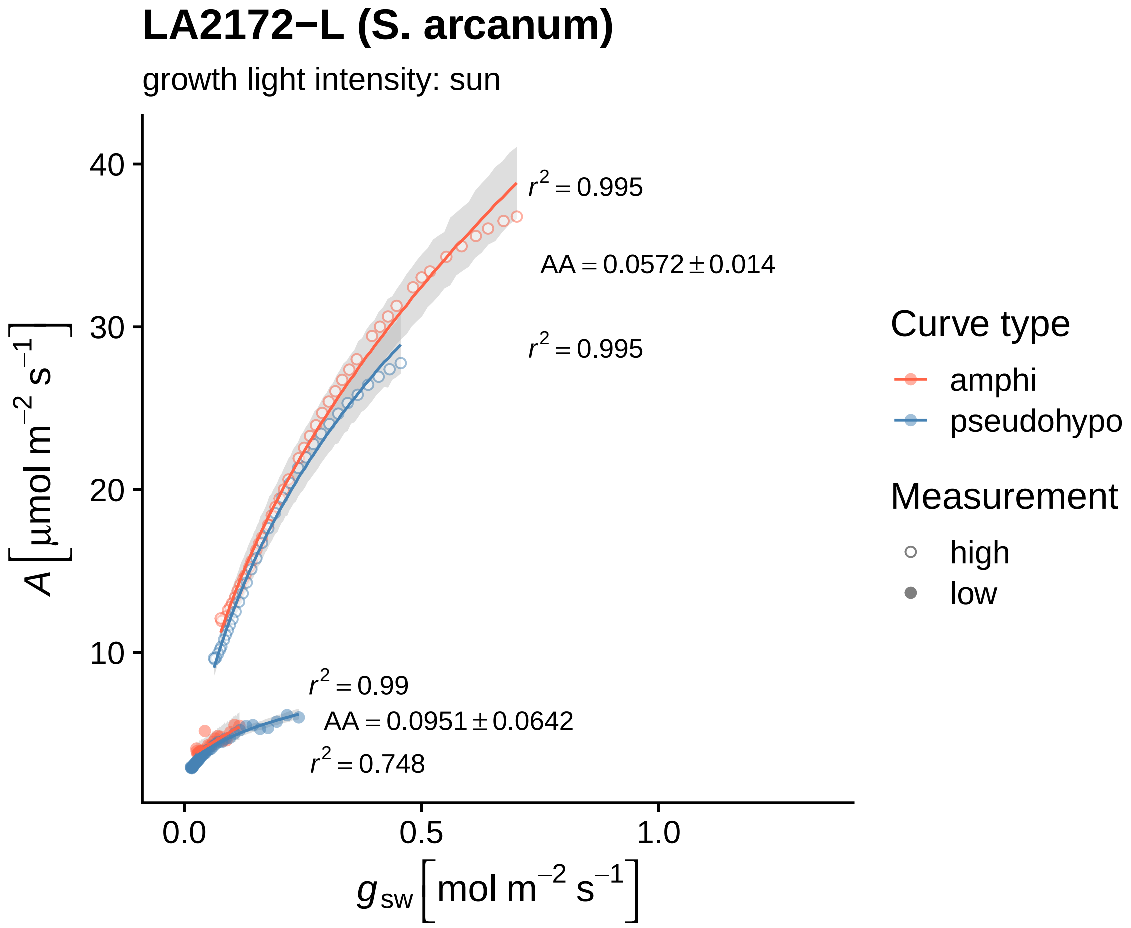

LA2172−L (S. arcanum)
growth light intensity: sun
$r^2 = 0.995$
AA = 0.0572 ± 0.014
$r^2 = 0.995$
$r^2 = 0.99$
AA = 0.0951 ± 0.0642
$r^2 = 0.748$
40
30
20
10
0.0
0.5
1.0
$A\ [\mu mol\ m^{-2}\ s^{-1}]$
$g_{sw}\ [mol\ m^{-2}\ s^{-1}]$
Curve type
amphi
pseudohypo
Measurement
high
low

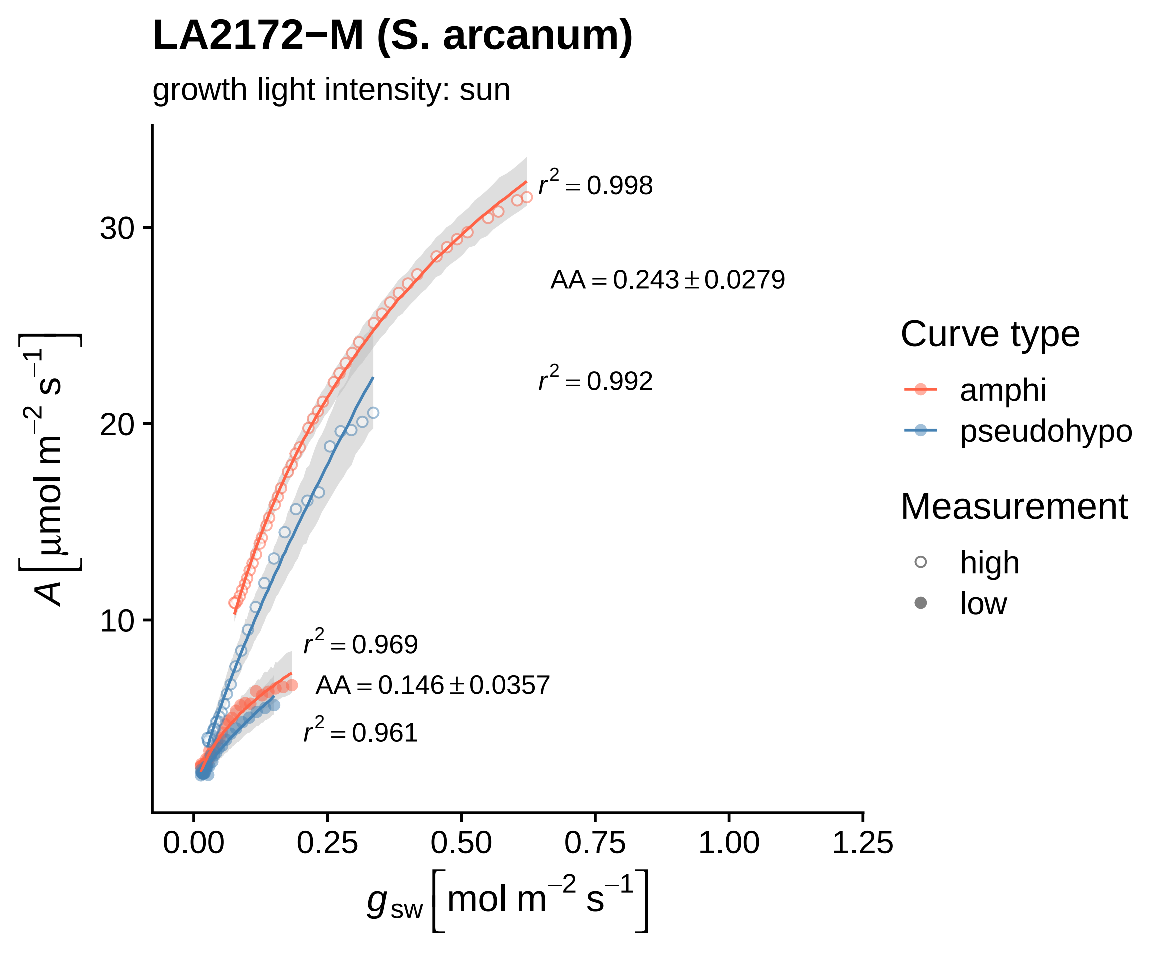

LA2172−M (S. arcanum)
growth light intensity: sun
$r^2 = 0.998$
$AA = 0.243 \pm 0.0279$
$r^2 = 0.992$
$r^2 = 0.969$
$AA = 0.146 \pm 0.0357$
$r^2 = 0.961$
30
20
10
0.00
0.25
0.50
0.75
1.00
1.25
$A\ [\mu mol\ m^{-2}\ s^{-1}]$
$g_{sw}\ [mol\ m^{-2}\ s^{-1}]$
Curve type
amphi
pseudohypo
Measurement
high
low

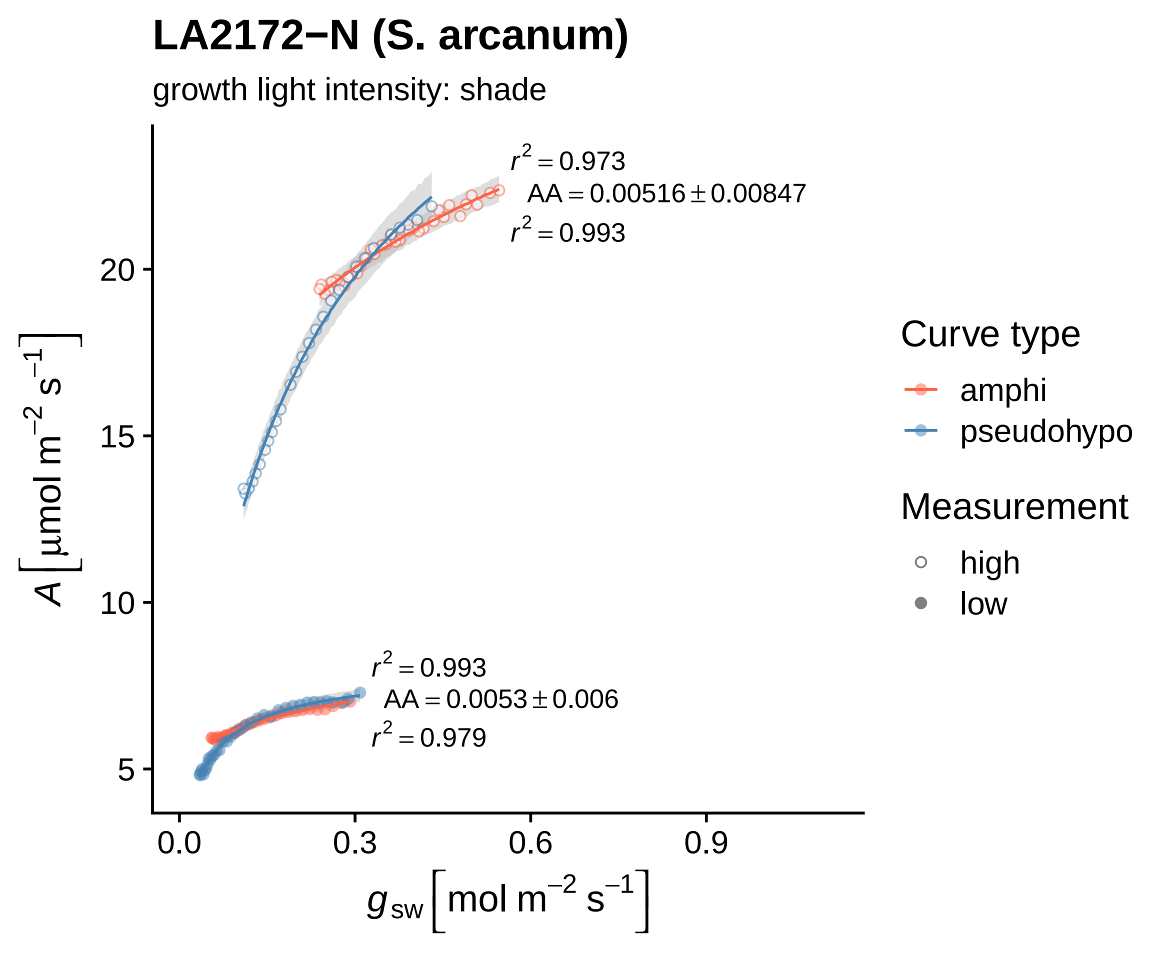

LA2172−N (S. arcanum)
growth light intensity: shade
$r^2 = 0.973$
AA = 0.00516 ± 0.00847
$r^2 = 0.993$
$r^2 = 0.993$
AA = 0.0053 ± 0.006
$r^2 = 0.979$
$A$ [µmol m$^{-2}$ s$^{-1}$]
$g_{sw}$ [mol m$^{-2}$ s$^{-1}$]
0.0
0.3
0.6
0.9
5
10
15
20
Curve type
amphi
pseudohypo
Measurement
high
low

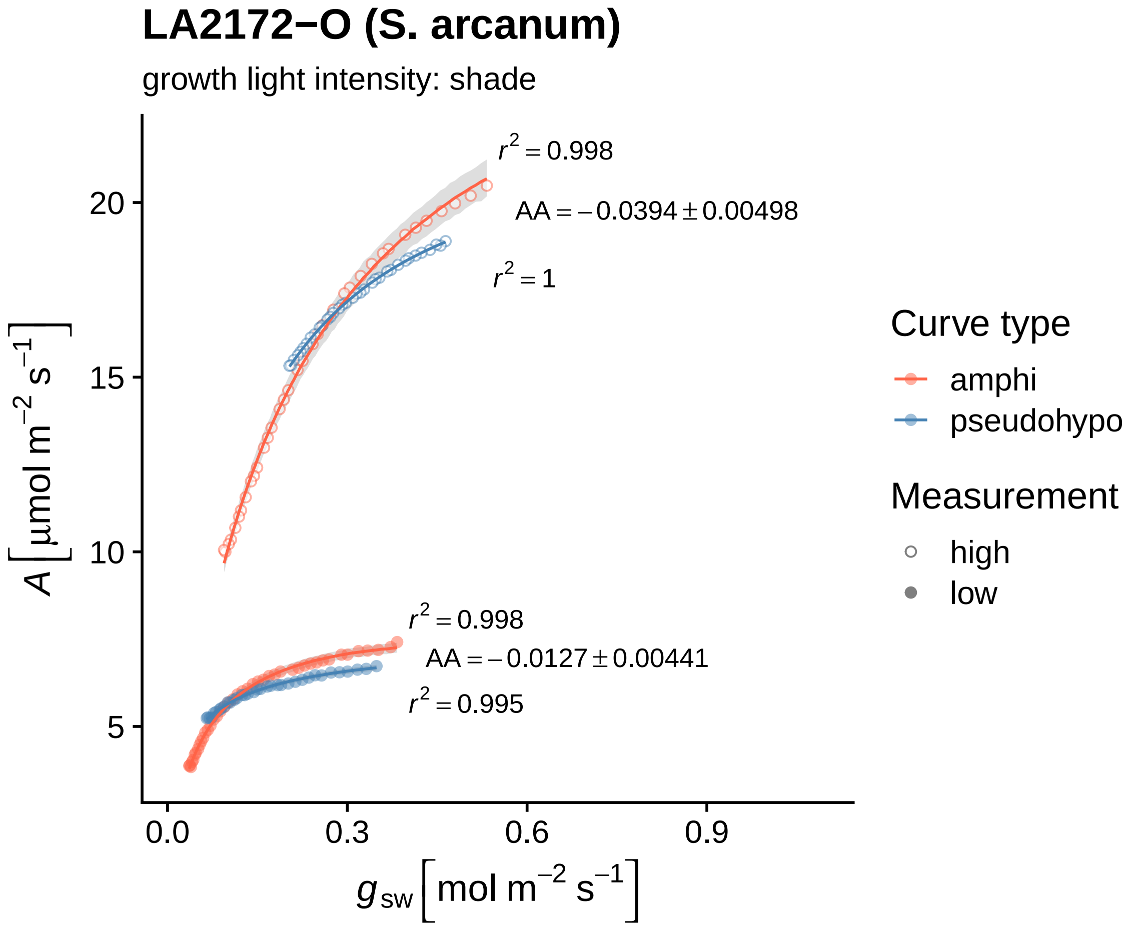
LA2172−O (S. arcanum)
growth light intensity: shade
$r^2 = 0.998$
$AA = -0.0394 \pm 0.00498$
$r^2 = 1$
$r^2 = 0.998$
$AA = -0.0127 \pm 0.00441$
$r^2 = 0.995$
$A\left[\mu mol\ m^{-2}\ s^{-1}\right]$
$g_{sw}\left[mol\ m^{-2}\ s^{-1}\right]$
20
15
10
5
0.0
0.3
0.6
0.9
Curve type
amphi
pseudohypo
Measurement
high
low

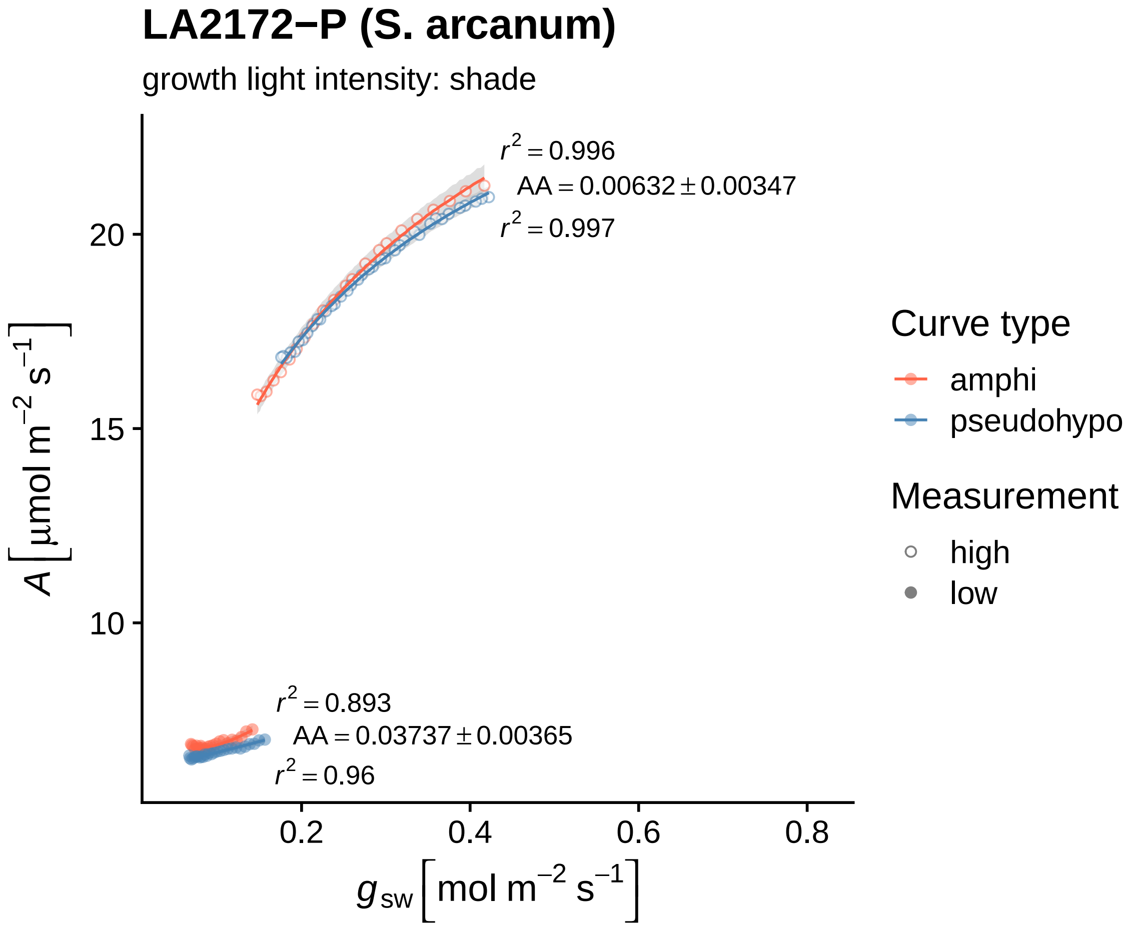

LA2172−P (S. arcanum)
growth light intensity: shade
$r^2 = 0.996$
$AA = 0.00632 \pm 0.00347$
$r^2 = 0.997$
$r^2 = 0.893$
$AA = 0.03737 \pm 0.00365$
$r^2 = 0.96$
20
15
10
0.2
0.4
0.6
0.8
$A\left[\mu mol\ m^{-2}\ s^{-1}\right]$
$g_{sw}\left[mol\ m^{-2}\ s^{-1}\right]$
Curve type
amphi
pseudohypo
Measurement
high
low

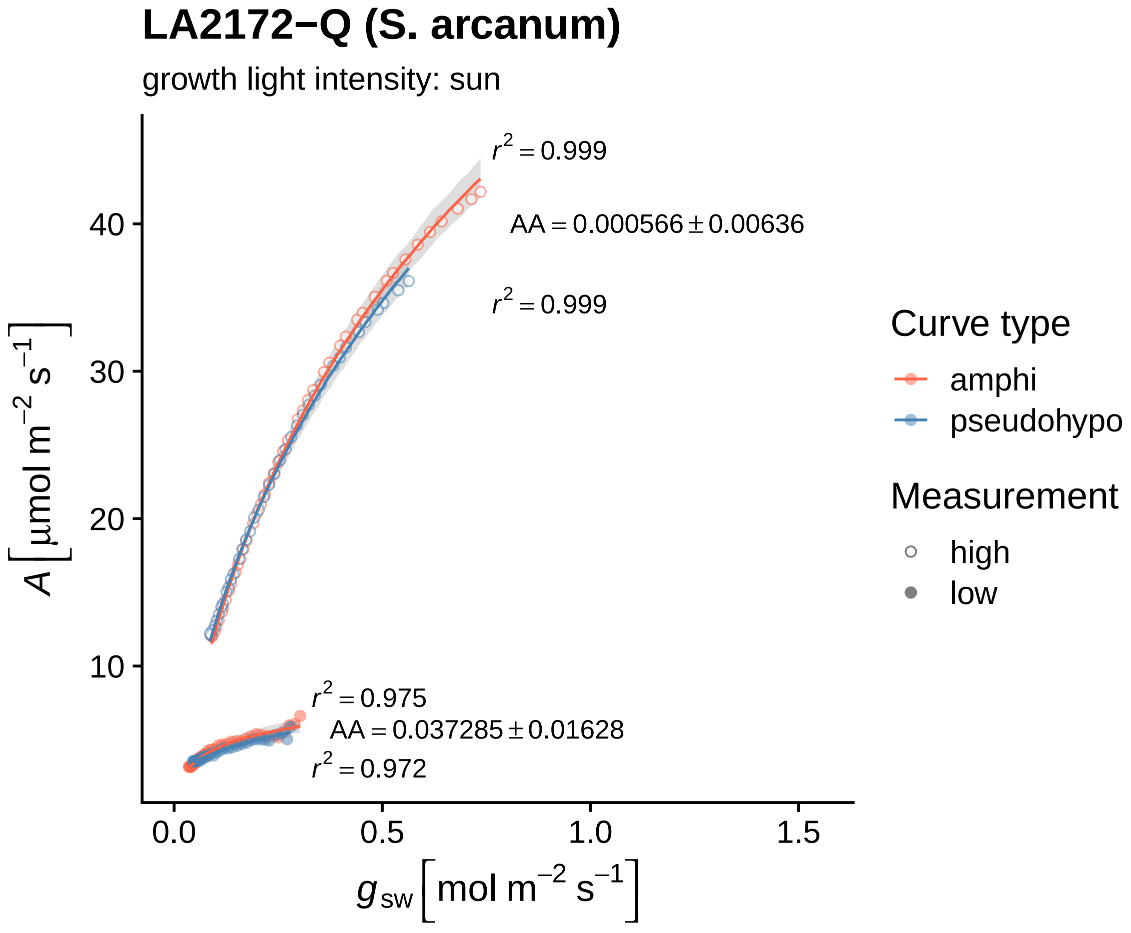

LA2172−Q (S. arcanum)
growth light intensity: sun
$r^2 = 0.999$
AA = 0.000566 ± 0.00636
$r^2 = 0.999$
$r^2 = 0.975$
AA = 0.037285 ± 0.01628
$r^2 = 0.972$
40
30
20
10
0.0
0.5
1.0
1.5
$A\ [\mu mol\ m^{-2}\ s^{-1}]$
$g_{sw}\ [mol\ m^{-2}\ s^{-1}]$
Curve type
amphi
pseudohypo
Measurement
high
low

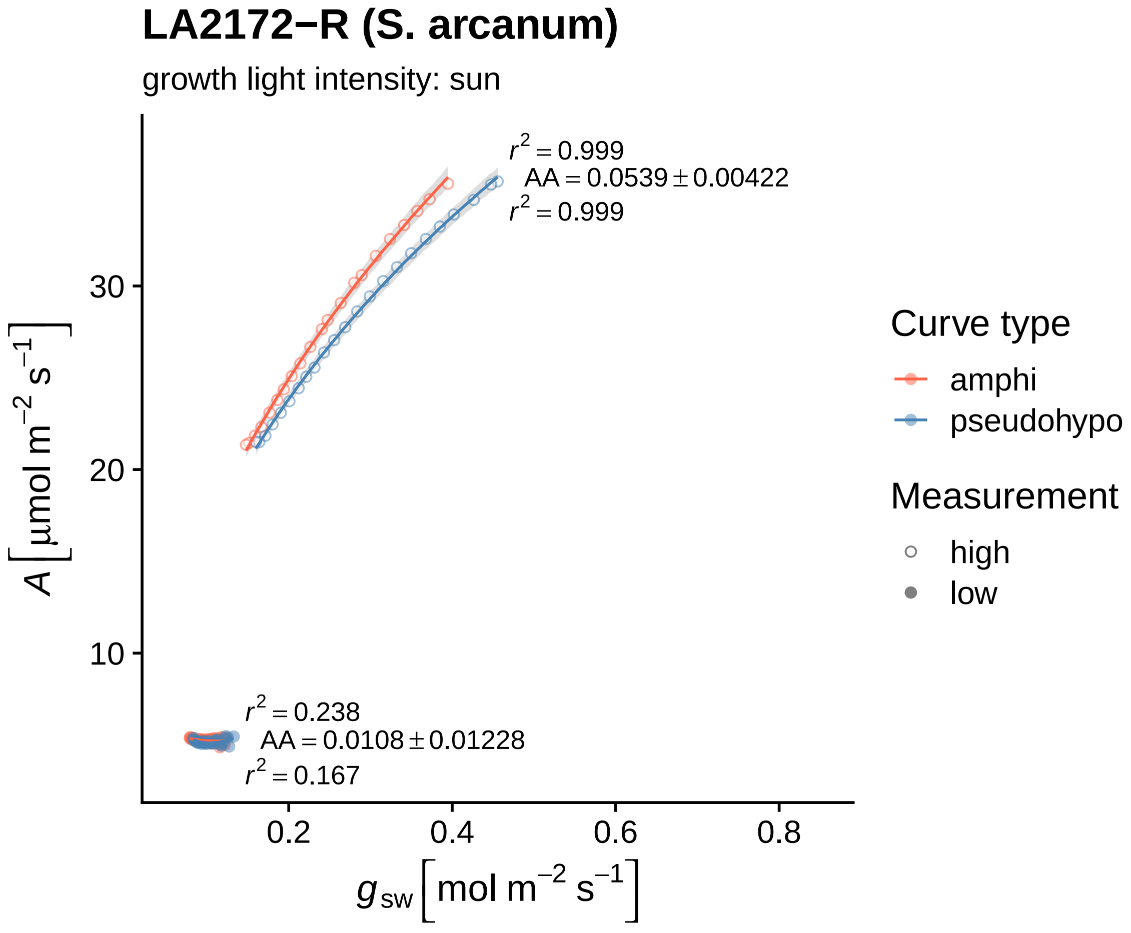

LA2172−R (S. arcanum)
growth light intensity: sun
$r^2 = 0.999$
$AA = 0.0539 \pm 0.00422$
$r^2 = 0.999$
$r^2 = 0.238$
$AA = 0.0108 \pm 0.01228$
$r^2 = 0.167$
30
20
10
0.2
0.4
0.6
0.8
$A \left[\mu mol\ m^{-2}\ s^{-1}\right]$
$g_{sw} \left[mol\ m^{-2}\ s^{-1}\right]$
Curve type
amphi
pseudohypo
Measurement
high
low

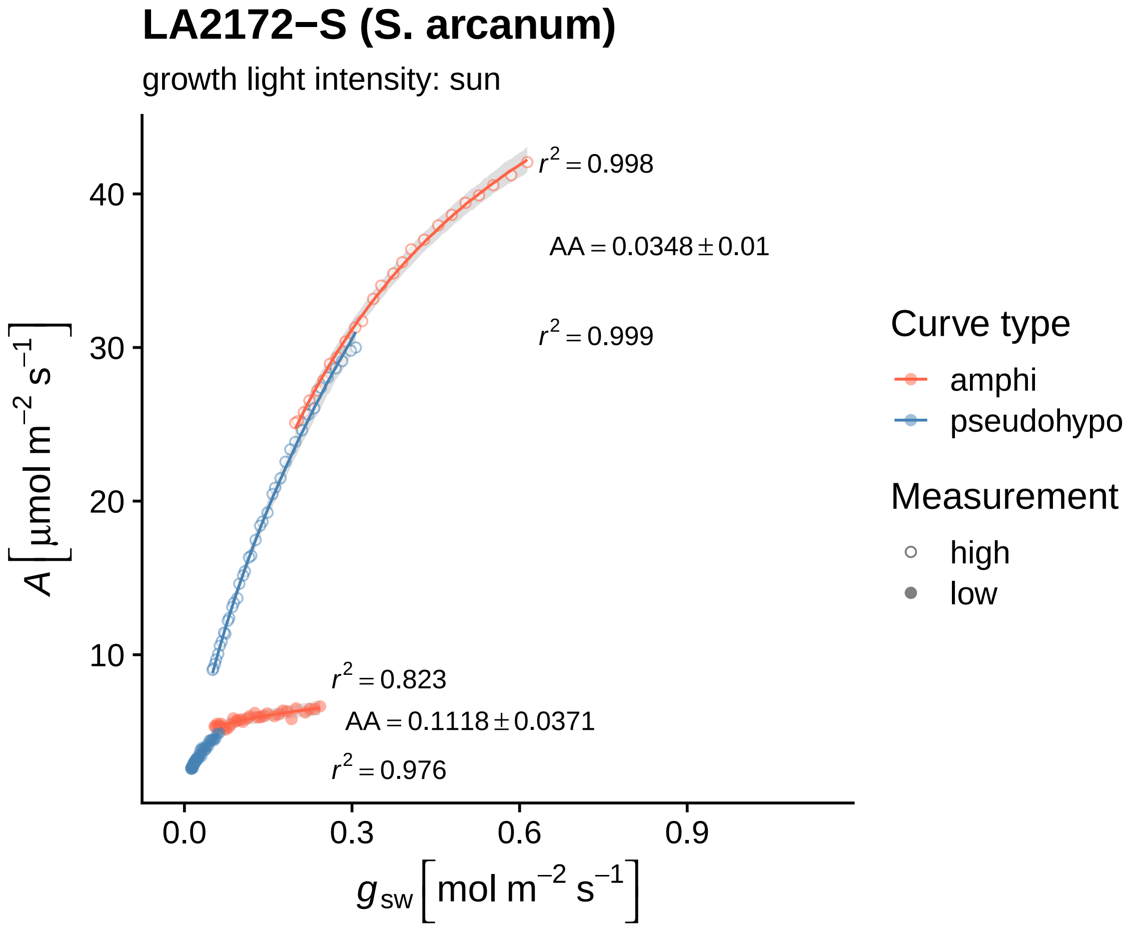
LA2172−S (S. arcanum)
growth light intensity: sun
$r^2 = 0.998$
$AA = 0.0348 \pm 0.01$
$r^2 = 0.999$
$r^2 = 0.823$
$AA = 0.1118 \pm 0.0371$
$r^2 = 0.976$
40
30
20
10
0.0
0.3
0.6
0.9
$A\,[\mu mol\ m^{-2}\ s^{-1}]$
$g_{sw}\,[mol\ m^{-2}\ s^{-1}]$
Curve type
amphi
pseudohypo
Measurement
high
low

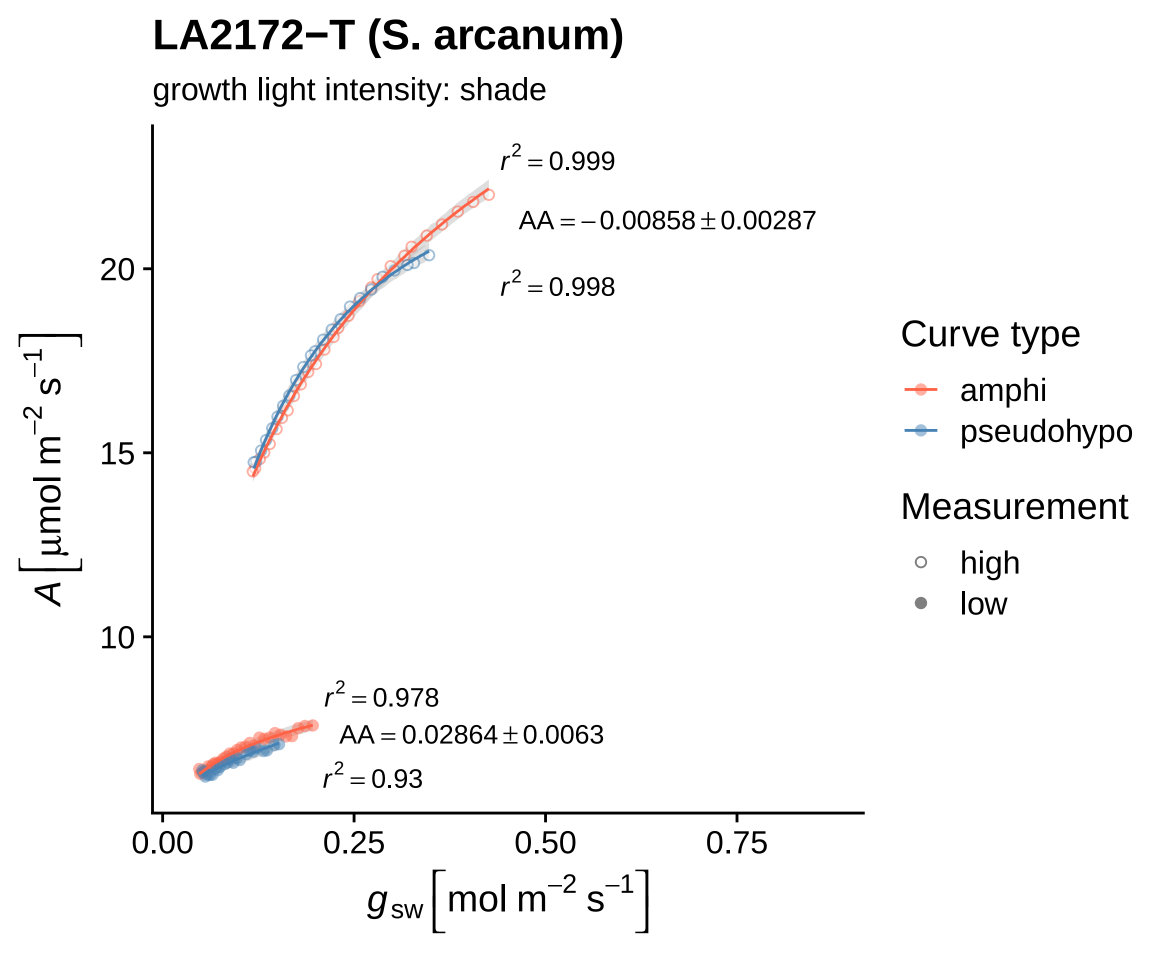

LA2172−T (S. arcanum)
growth light intensity: shade
$r^2 = 0.999$
$AA = -0.00858 \pm 0.00287$
$r^2 = 0.998$
$r^2 = 0.978$
$AA = 0.02864 \pm 0.0063$
$r^2 = 0.93$
20
15
10
0.00
0.25
0.50
0.75
$A\ [\mu mol\ m^{-2}\ s^{-1}]$
$g_{sw}\ [mol\ m^{-2}\ s^{-1}]$
Curve type
amphi
pseudohypo
Measurement
high
low

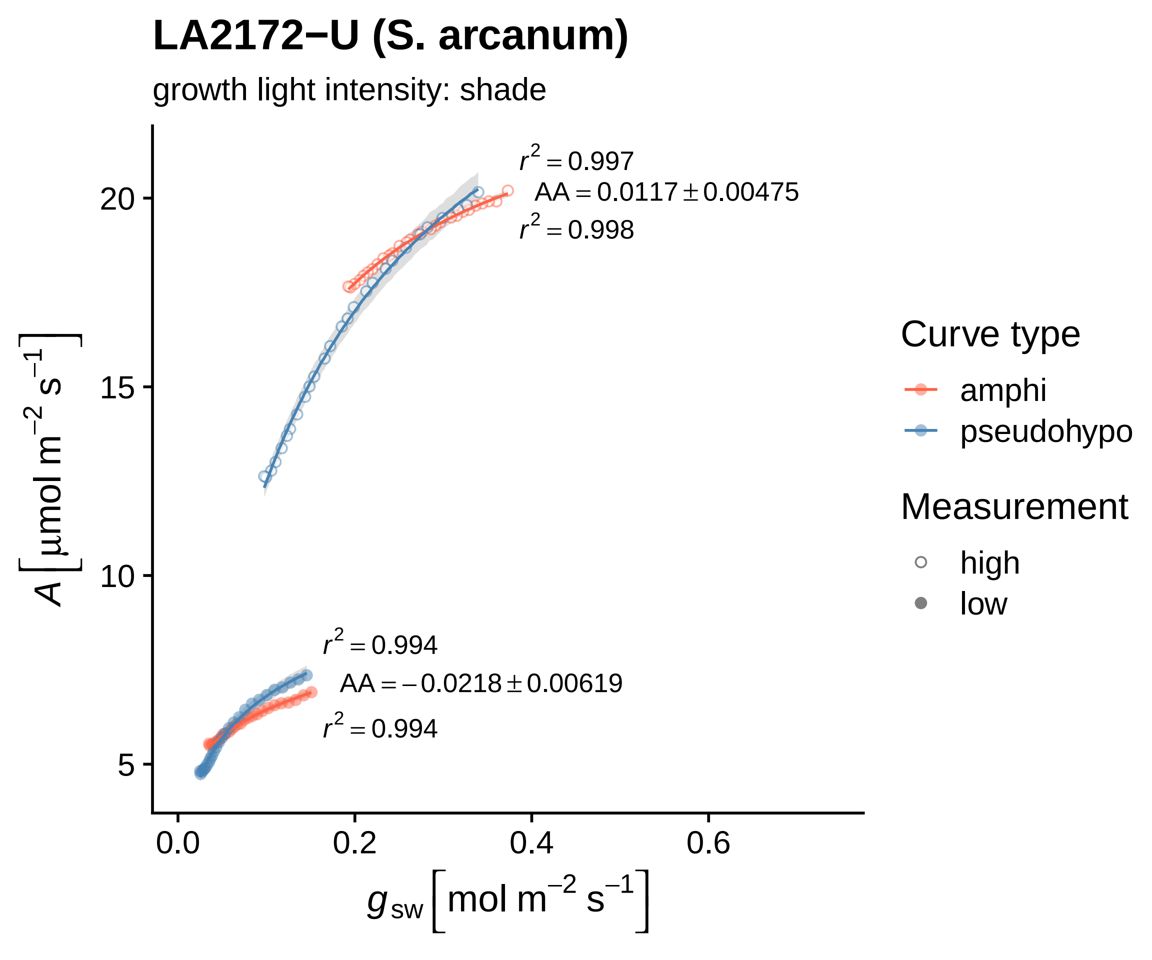

LA2172−U (S. arcanum)
growth light intensity: shade
$r^2 = 0.997$
$AA = 0.0117 \pm 0.00475$
$r^2 = 0.998$
$r^2 = 0.994$
$AA = -0.0218 \pm 0.00619$
$r^2 = 0.994$
20
15
10
5
0.0
0.2
0.4
0.6
$A \left[\mu mol\ m^{-2}\ s^{-1}\right]$
$g_{sw} \left[mol\ m^{-2}\ s^{-1}\right]$
Curve type
amphi
pseudohypo
Measurement
high
low

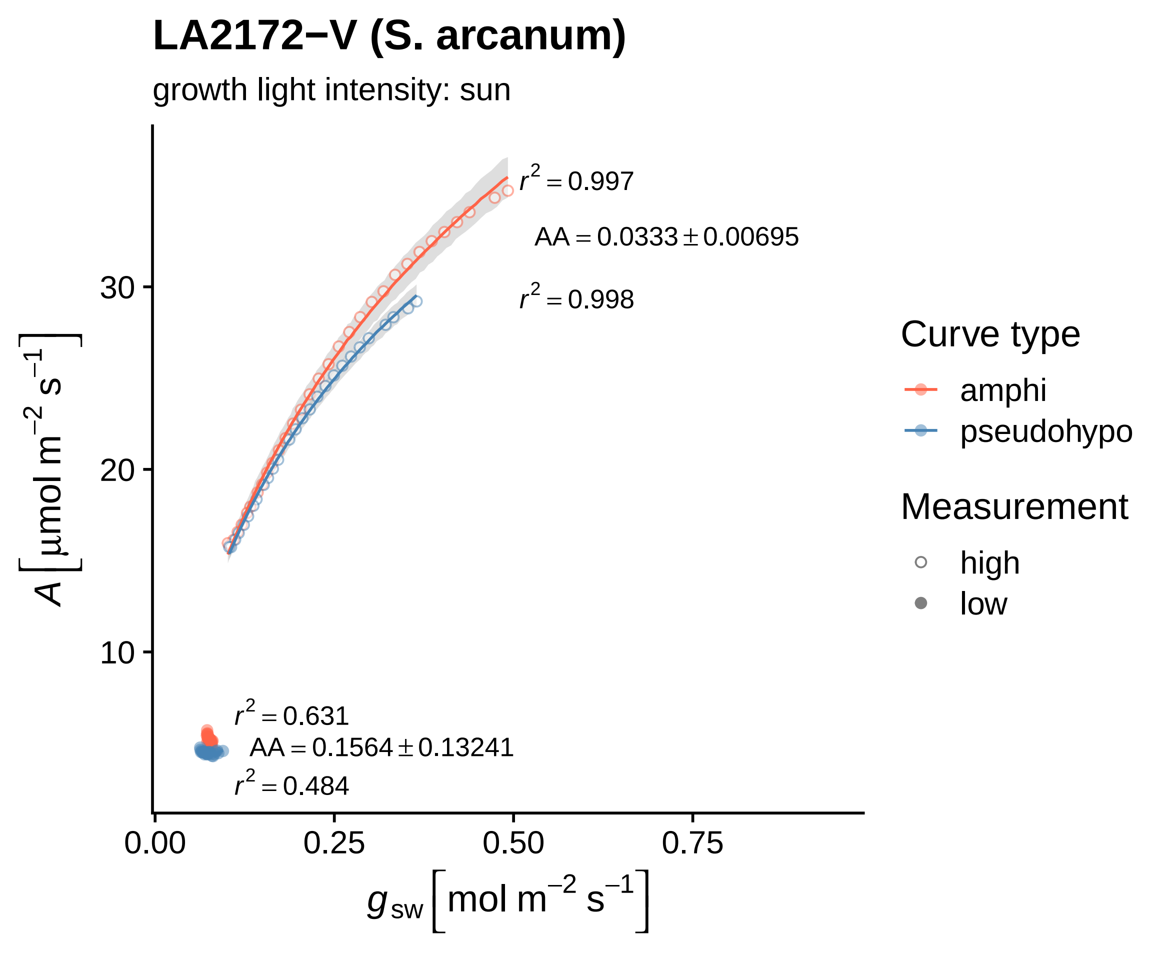

LA2172−V (S. arcanum)
growth light intensity: sun
$r^2 = 0.997$
$AA = 0.0333 \pm 0.00695$
$r^2 = 0.998$
$r^2 = 0.631$
$AA = 0.1564 \pm 0.13241$
$r^2 = 0.484$
30
20
10
0.00
0.25
0.50
0.75
$A\left[\mu mol\ m^{-2}\ s^{-1}\right]$
$g_{sw}\left[mol\ m^{-2}\ s^{-1}\right]$
Curve type
amphi
pseudohypo
Measurement
high
low

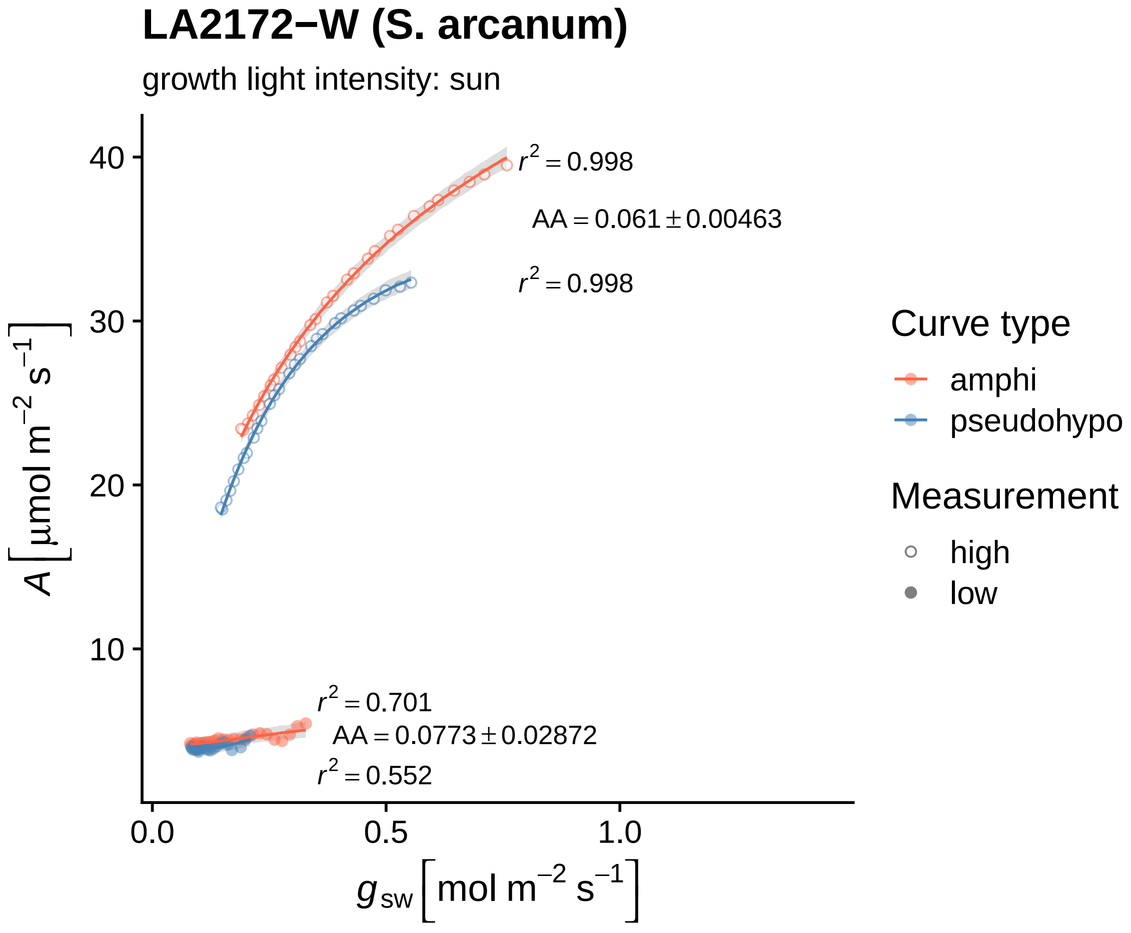

LA2172−W (S. arcanum)
growth light intensity: sun
$r^2 = 0.998$
$AA = 0.061 \pm 0.00463$
$r^2 = 0.998$
$r^2 = 0.701$
$AA = 0.0773 \pm 0.02872$
$r^2 = 0.552$
40
30
20
10
0.0
0.5
1.0
$A\ [\mu mol\ m^{-2}\ s^{-1}]$
$g_{sw}\ [mol\ m^{-2}\ s^{-1}]$
Curve type
amphi
pseudohypo
Measurement
high
low

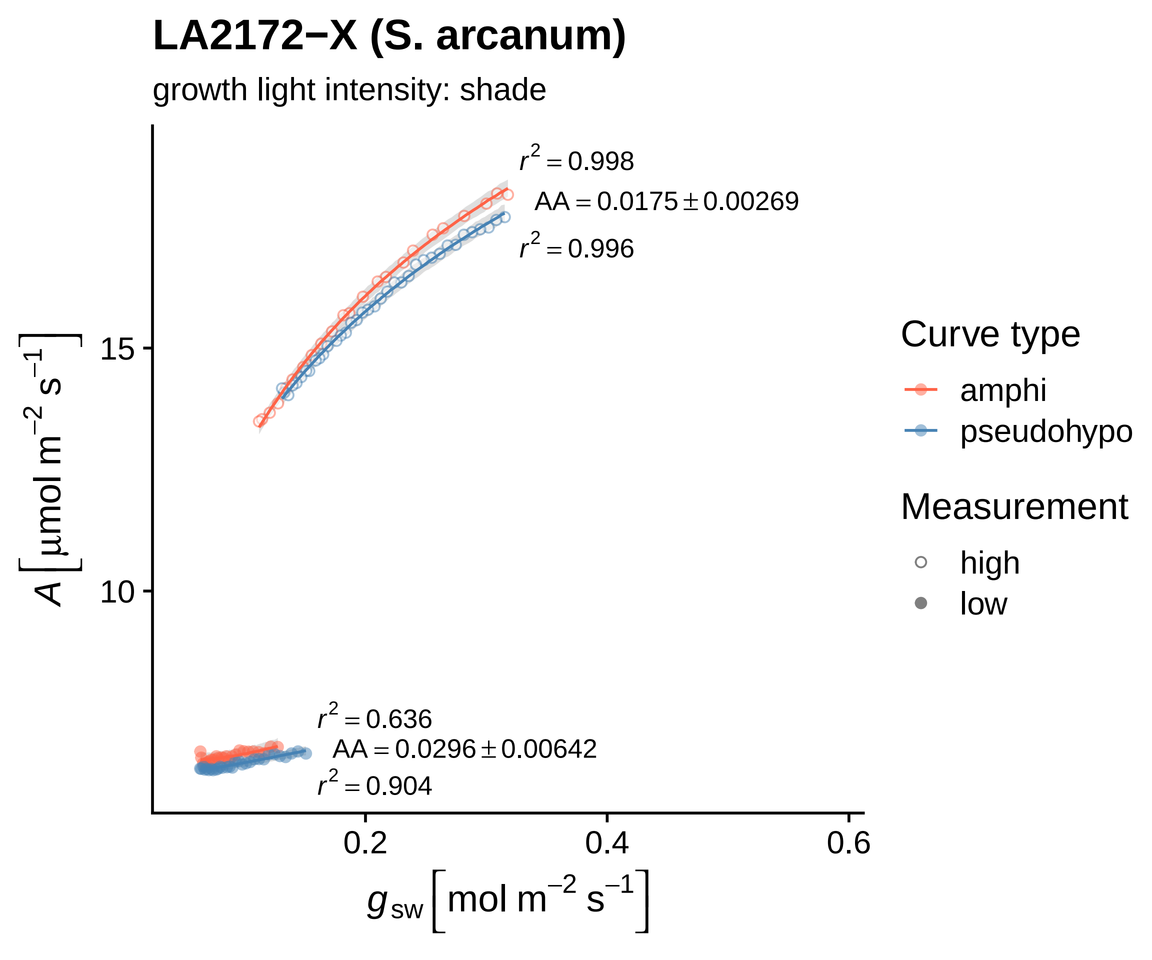

LA2172−X (S. arcanum)
growth light intensity: shade
$r^2 = 0.998$
$AA = 0.0175 \pm 0.00269$
$r^2 = 0.996$
$r^2 = 0.636$
$AA = 0.0296 \pm 0.00642$
$r^2 = 0.904$
15
10
0.2
0.4
0.6
$A\ [\mu mol\ m^{-2}\ s^{-1}]$
$g_{sw}\ [mol\ m^{-2}\ s^{-1}]$
Curve type
amphi
pseudohypo
Measurement
high
low

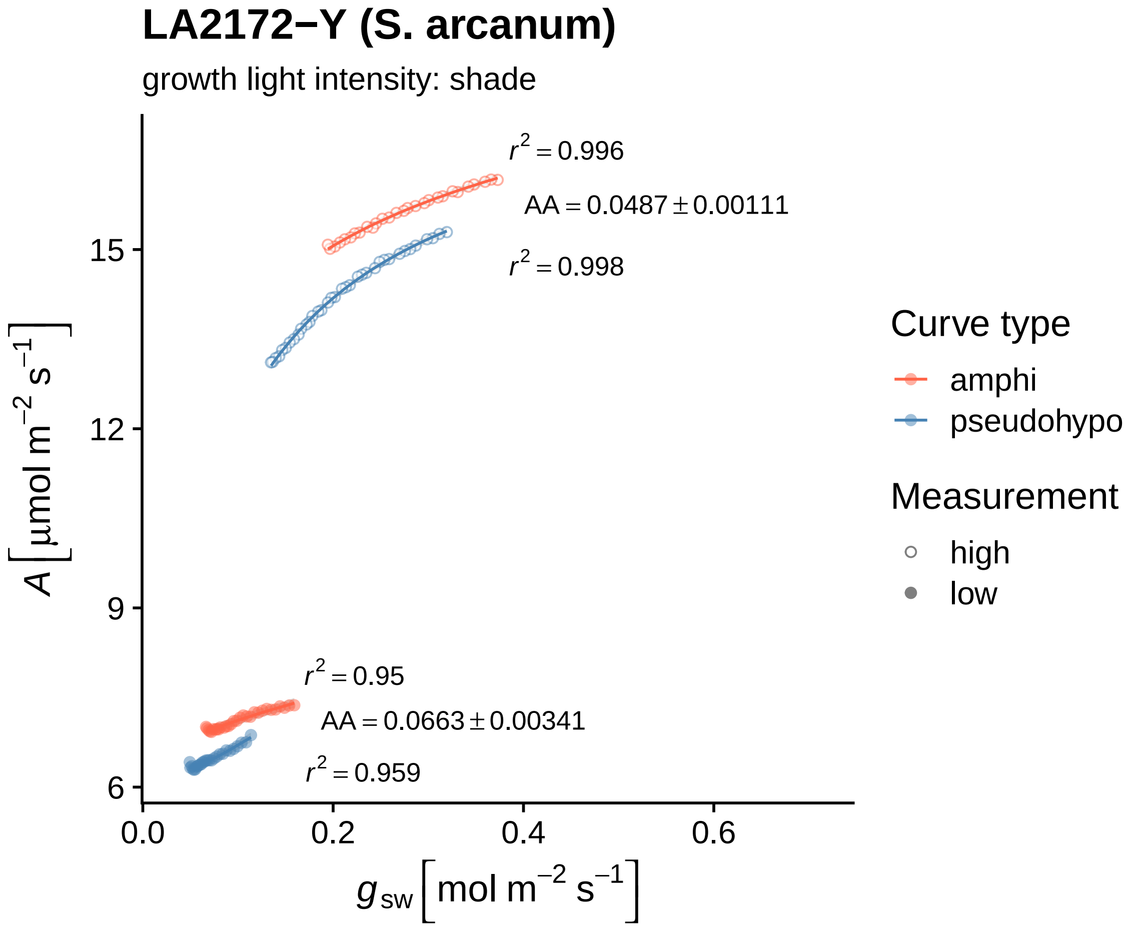

LA2172−Y (S. arcanum)
growth light intensity: shade
$r^2 = 0.996$
$AA = 0.0487 \pm 0.00111$
$r^2 = 0.998$
$r^2 = 0.95$
$AA = 0.0663 \pm 0.00341$
$r^2 = 0.959$
15
12
9
6
0.0
0.2
0.4
0.6
$A\left[\mu mol\ m^{-2}\ s^{-1}\right]$
$g_{sw}\left[mol\ m^{-2}\ s^{-1}\right]$
Curve type
amphi
pseudohypo
Measurement
high
low

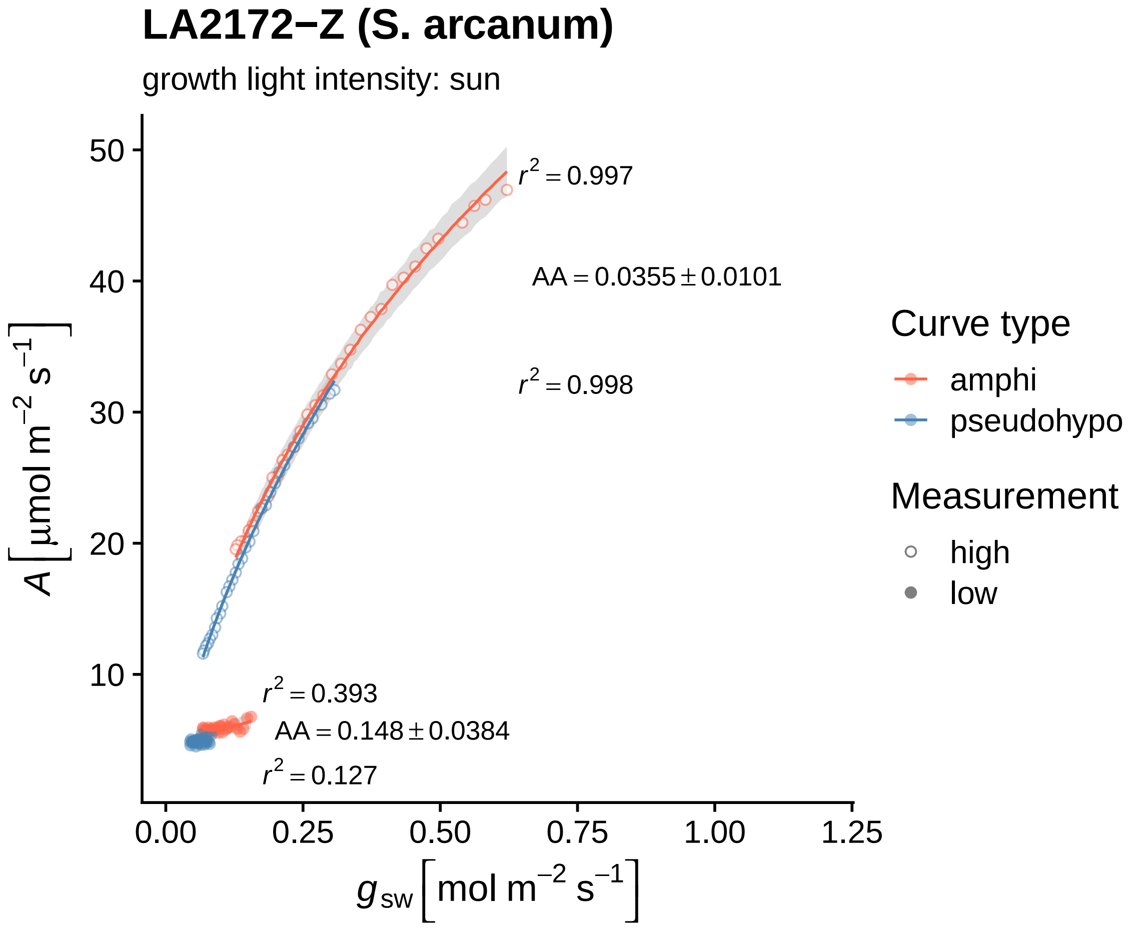

LA2172−Z (S. arcanum)
growth light intensity: sun
$r^2 = 0.997$
$AA = 0.0355 \pm 0.0101$
$r^2 = 0.998$
$r^2 = 0.393$
$AA = 0.148 \pm 0.0384$
$r^2 = 0.127$
50
40
30
20
10
0.00
0.25
0.50
0.75
1.00
1.25
$A\ [\mu mol\ m^{-2}\ s^{-1}]$
$g_{sw}\ [mol\ m^{-2}\ s^{-1}]$
Curve type
amphi
pseudohypo
Measurement
high
low

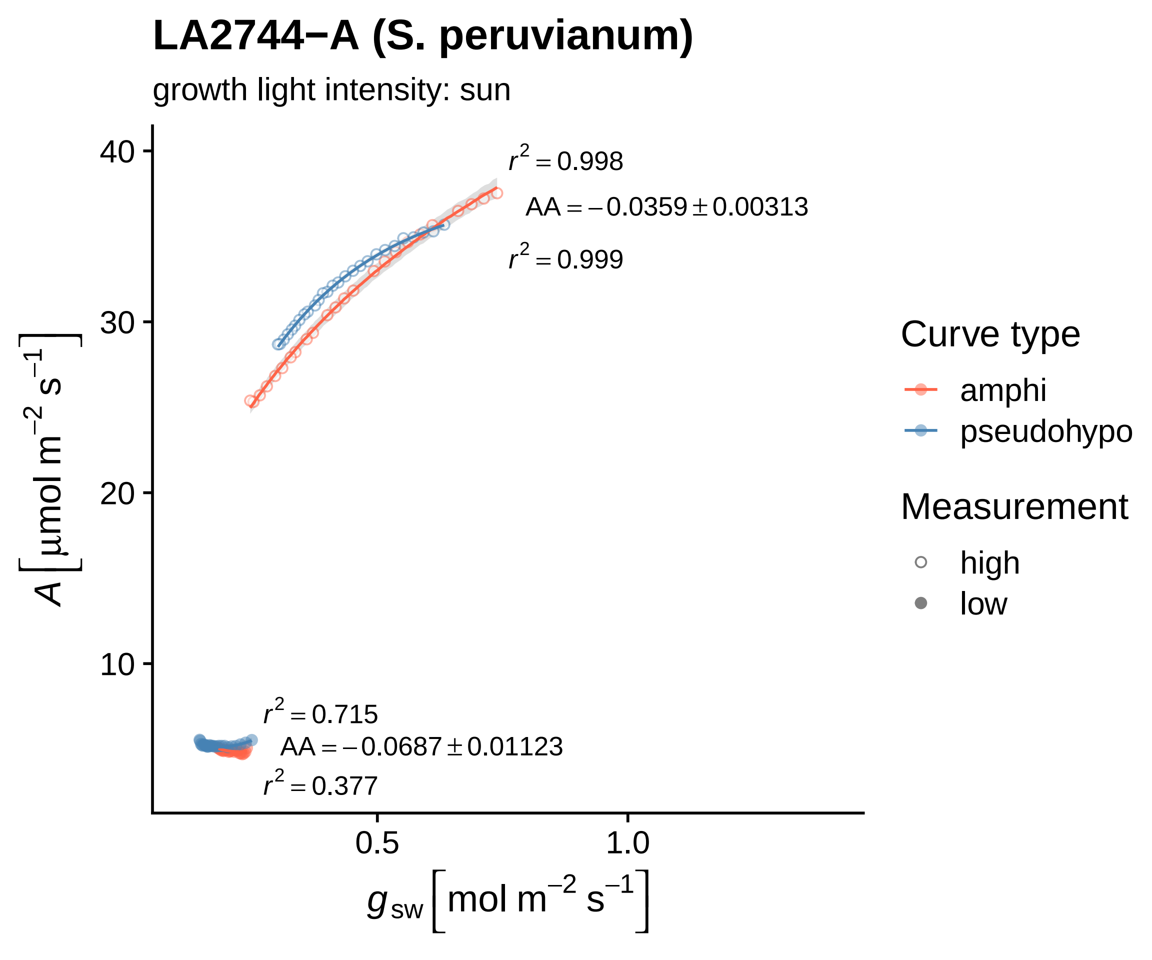

LA2744−A (S. peruvianum)
growth light intensity: sun
r² = 0.998
AA = −0.0359 ± 0.00313
r² = 0.999
r² = 0.715
AA = −0.0687 ± 0.01123
r² = 0.377
40
30
20
10
0.5
1.0
A [µmol m⁻² s⁻¹]
g sw [mol m⁻² s⁻¹]
Curve type
amphi
pseudohypo
Measurement
high
low

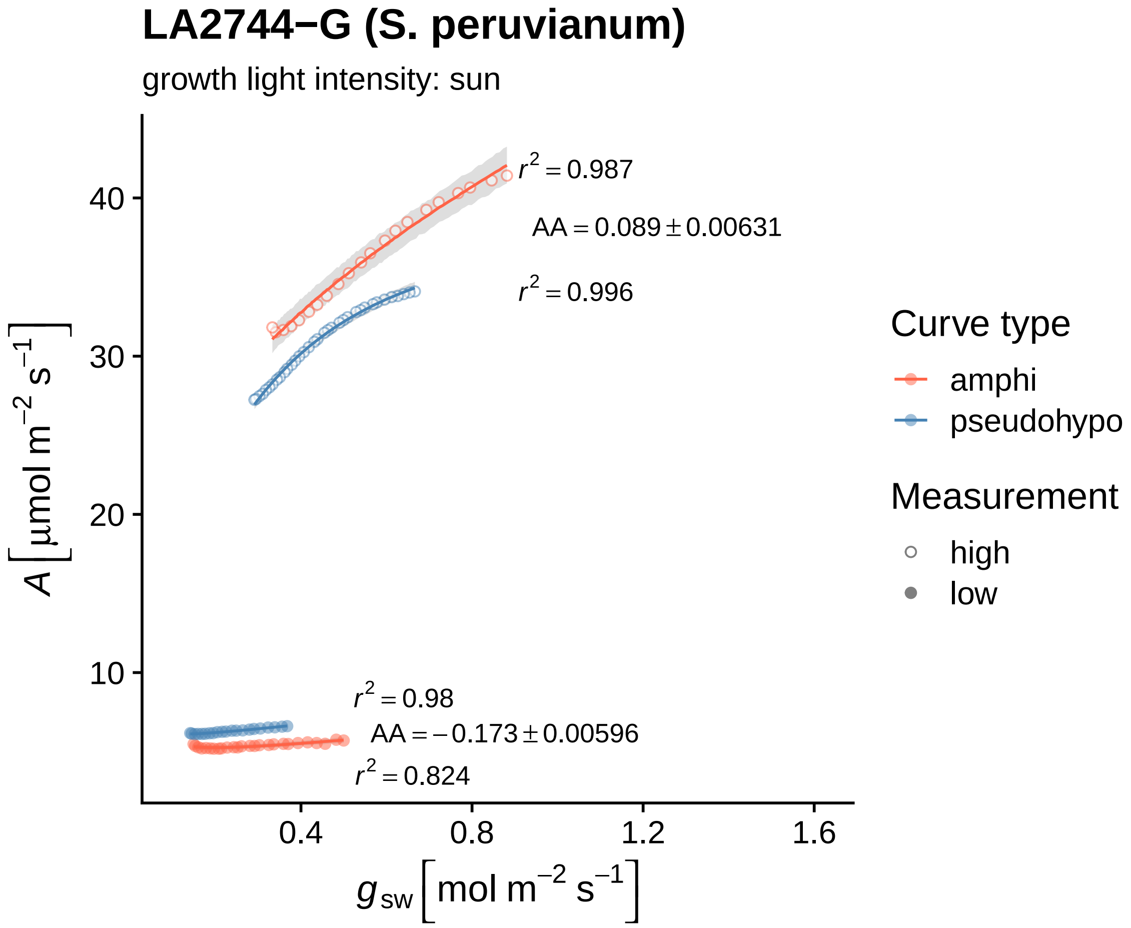
LA2744−G (S. peruvianum)
growth light intensity: sun
$r^2 = 0.987$
$AA = 0.089 \pm 0.00631$
$r^2 = 0.996$
$r^2 = 0.98$
$AA = -0.173 \pm 0.00596$
$r^2 = 0.824$
40
30
20
10
0.4
0.8
1.2
1.6
$A\,[\mu mol\ m^{-2}\ s^{-1}]$
$g_{sw}\,[mol\ m^{-2}\ s^{-1}]$
Curve type
amphi
pseudohypo
Measurement
high
low

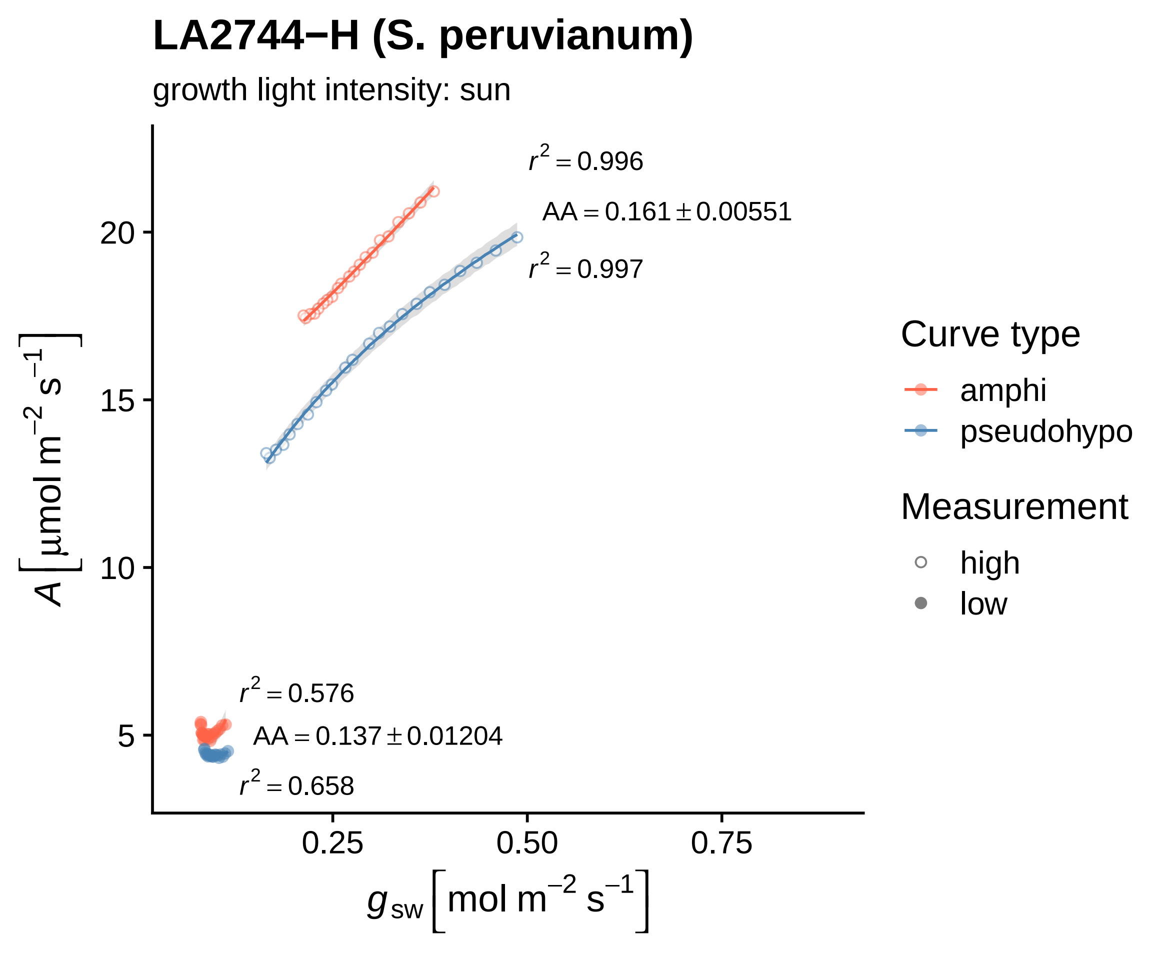

LA2744−H (S. peruvianum)
growth light intensity: sun
$r^2 = 0.996$
$AA = 0.161 \pm 0.00551$
$r^2 = 0.997$
$r^2 = 0.576$
$AA = 0.137 \pm 0.01204$
$r^2 = 0.658$
20
15
10
5
0.25
0.50
0.75
$A\ [\mu mol\ m^{-2}\ s^{-1}]$
$g_{sw}\ [mol\ m^{-2}\ s^{-1}]$
Curve type
amphi
pseudohypo
Measurement
high
low

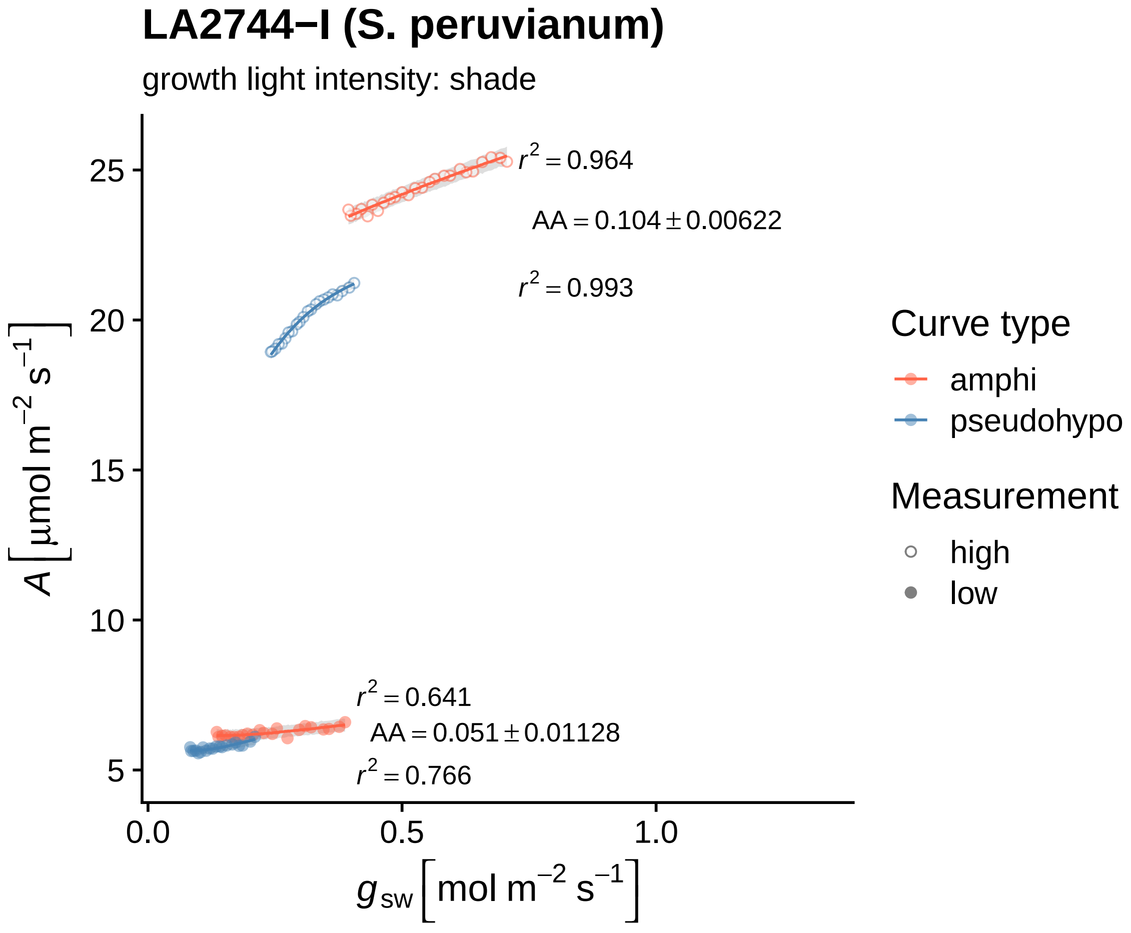

LA2744−I (S. peruvianum)
growth light intensity: shade
$r^2 = 0.964$
$AA = 0.104 \pm 0.00622$
$r^2 = 0.993$
$r^2 = 0.641$
$AA = 0.051 \pm 0.01128$
$r^2 = 0.766$
25
20
15
10
5
0.0
0.5
1.0
$A\left[\mu mol\ m^{-2}\ s^{-1}\right]$
$g_{sw}\left[mol\ m^{-2}\ s^{-1}\right]$
Curve type
amphi
pseudohypo
Measurement
high
low

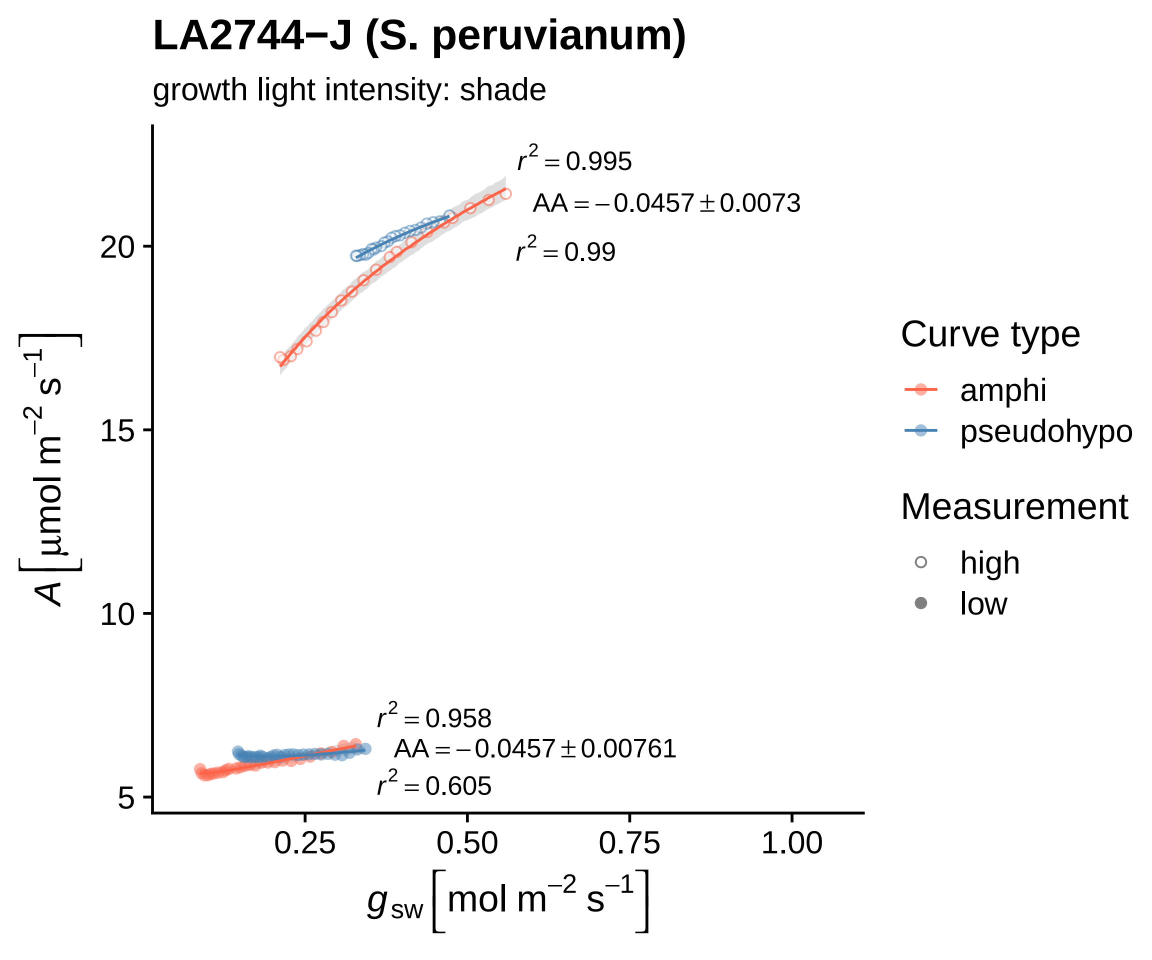

LA2744−J (S. peruvianum)
growth light intensity: shade
$r^2 = 0.995$
$AA = -0.0457 \pm 0.0073$
$r^2 = 0.99$
$r^2 = 0.958$
$AA = -0.0457 \pm 0.00761$
$r^2 = 0.605$
$A\ [\mu mol\ m^{-2}\ s^{-1}]$
20
15
10
5
0.25
0.50
0.75
1.00
$g_{sw}\ [mol\ m^{-2}\ s^{-1}]$
Curve type
amphi
pseudohypo
Measurement
high
low

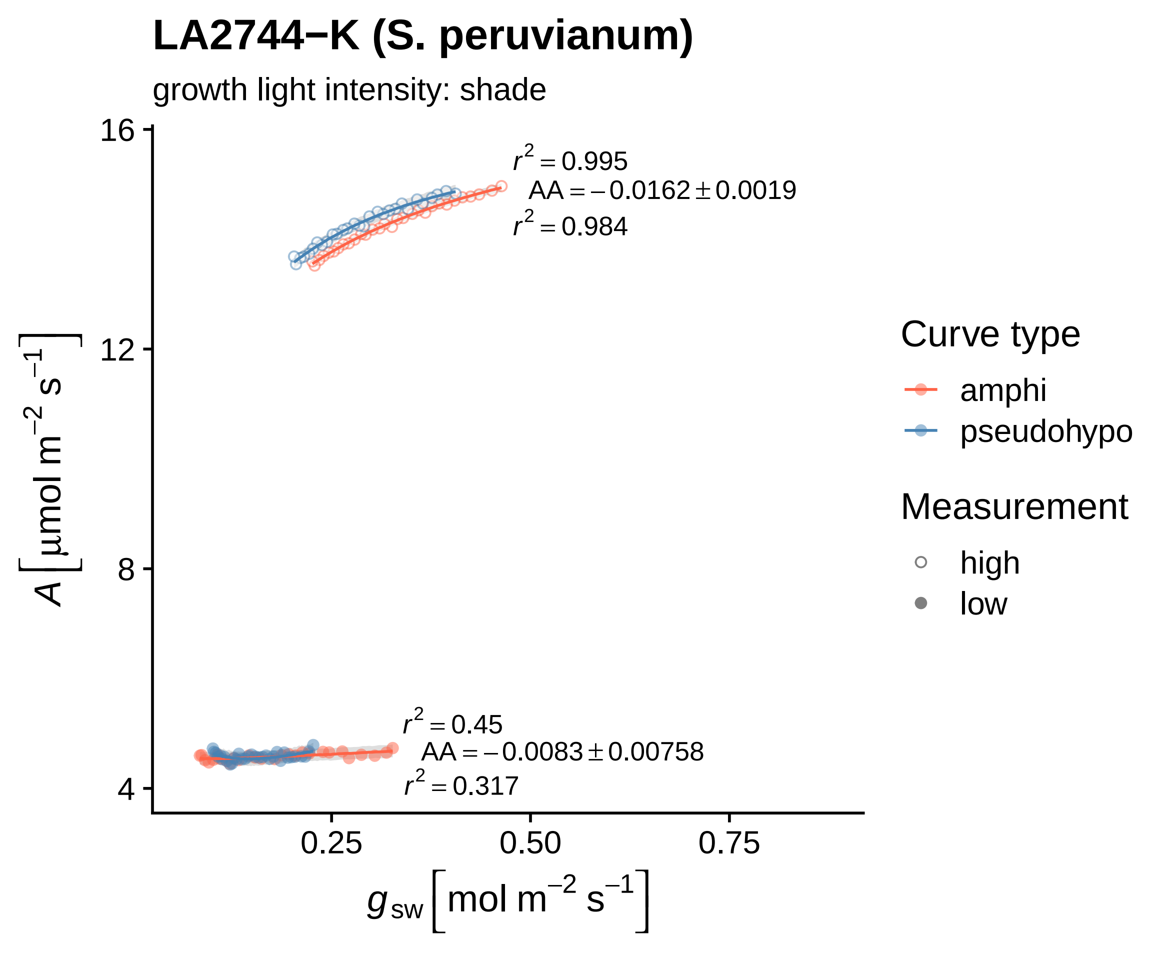

LA2744−K (S. peruvianum)
growth light intensity: shade
$r^2 = 0.995$
$AA = -0.0162 \pm 0.0019$
$r^2 = 0.984$
$r^2 = 0.45$
$AA = -0.0083 \pm 0.00758$
$r^2 = 0.317$
16
12
8
4
0.25
0.50
0.75
$A\ [\mu mol\ m^{-2}\ s^{-1}]$
$g_{sw}\ [mol\ m^{-2}\ s^{-1}]$
Curve type
amphi
pseudohypo
Measurement
high
low

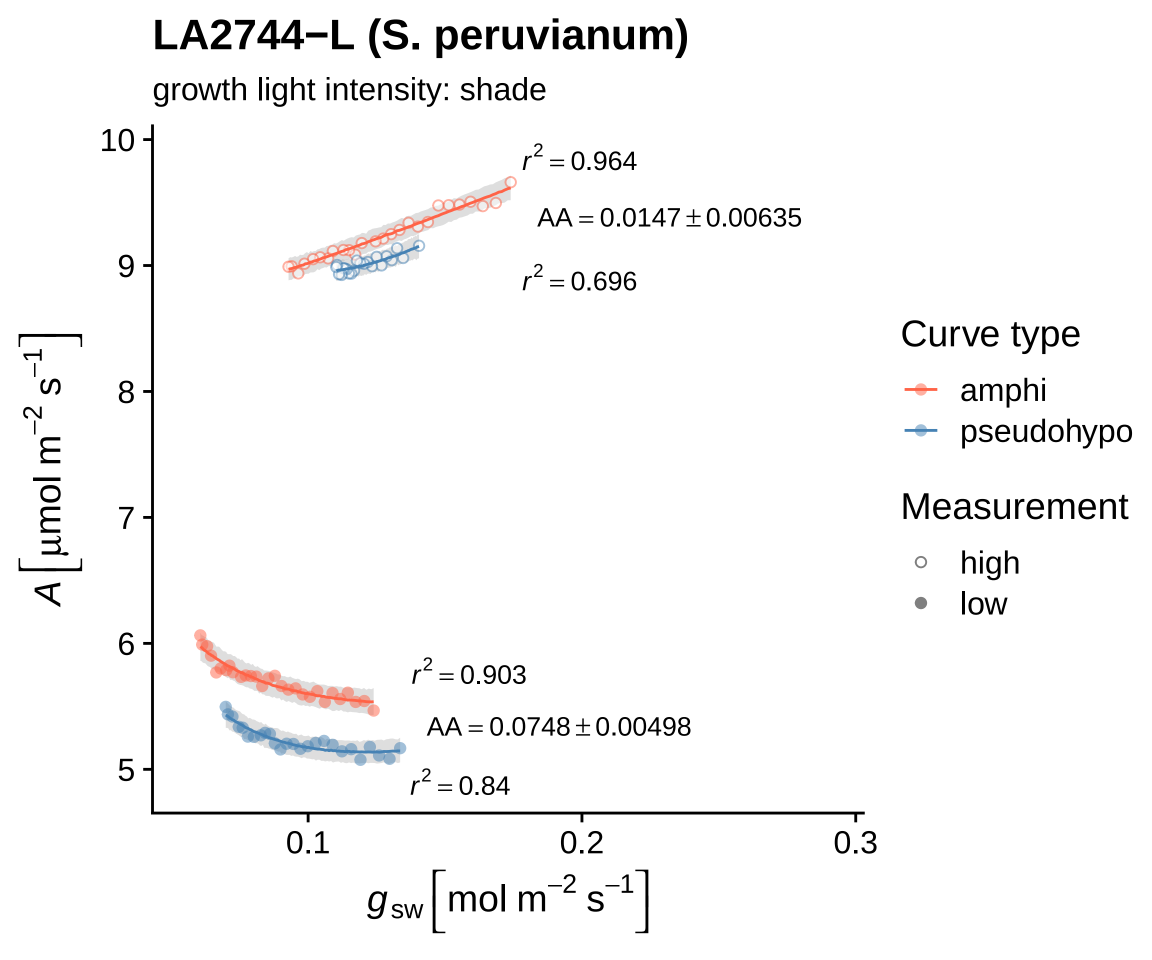
LA2744−L (S. peruvianum)
growth light intensity: shade
$r^2 = 0.964$
$AA = 0.0147 \pm 0.00635$
$r^2 = 0.696$
$r^2 = 0.903$
$AA = 0.0748 \pm 0.00498$
$r^2 = 0.84$
$A\left[\mu mol\ m^{-2}\ s^{-1}\right]$
$g_{sw}\left[mol\ m^{-2}\ s^{-1}\right]$
10
9
8
7
6
5
0.1
0.2
0.3
Curve type
amphi
pseudohypo
Measurement
high
low

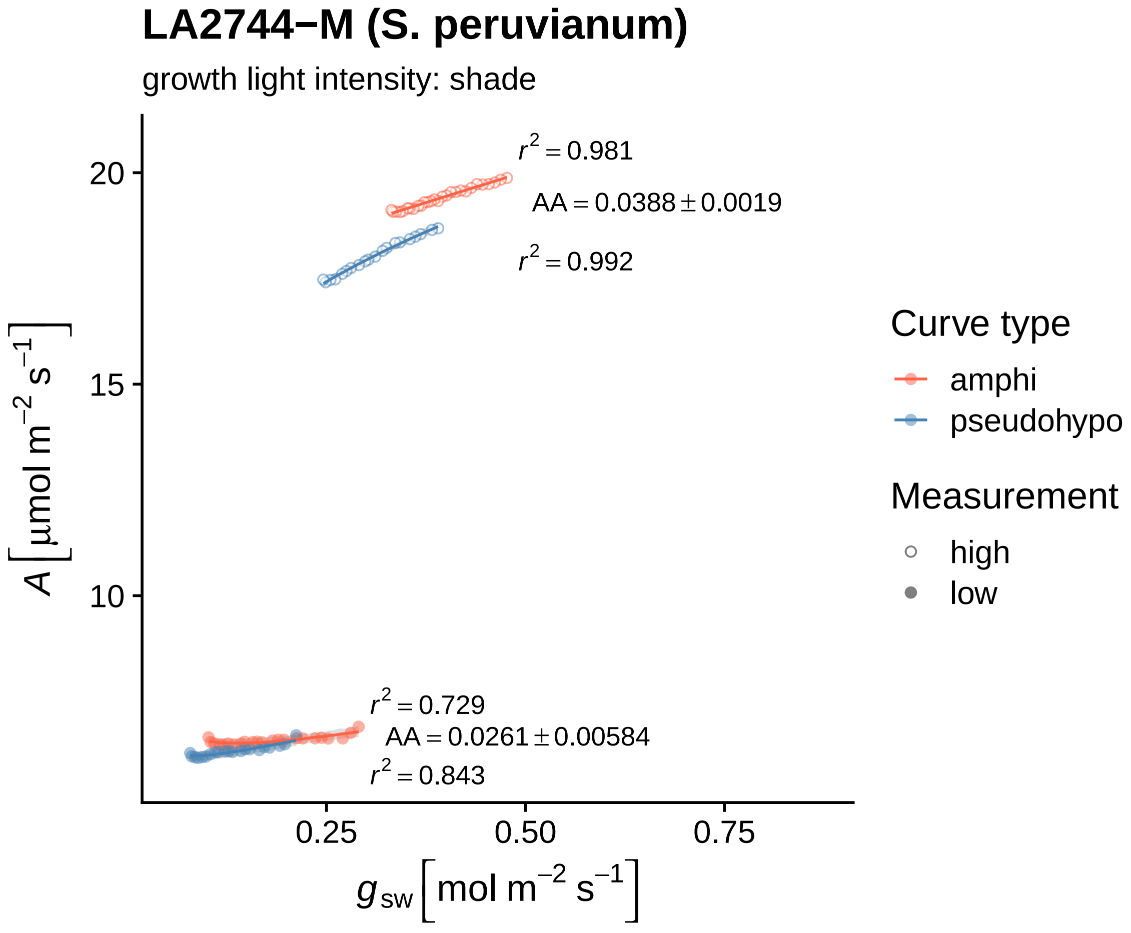
LA2744−M (S. peruvianum)
growth light intensity: shade
$r^2 = 0.981$
$AA = 0.0388 \pm 0.0019$
$r^2 = 0.992$
$r^2 = 0.729$
$AA = 0.0261 \pm 0.00584$
$r^2 = 0.843$
20
15
10
0.25
0.50
0.75
$A\ [\mu mol\ m^{-2}\ s^{-1}]$
$g_{sw}\ [mol\ m^{-2}\ s^{-1}]$
Curve type
amphi
pseudohypo
Measurement
high
low

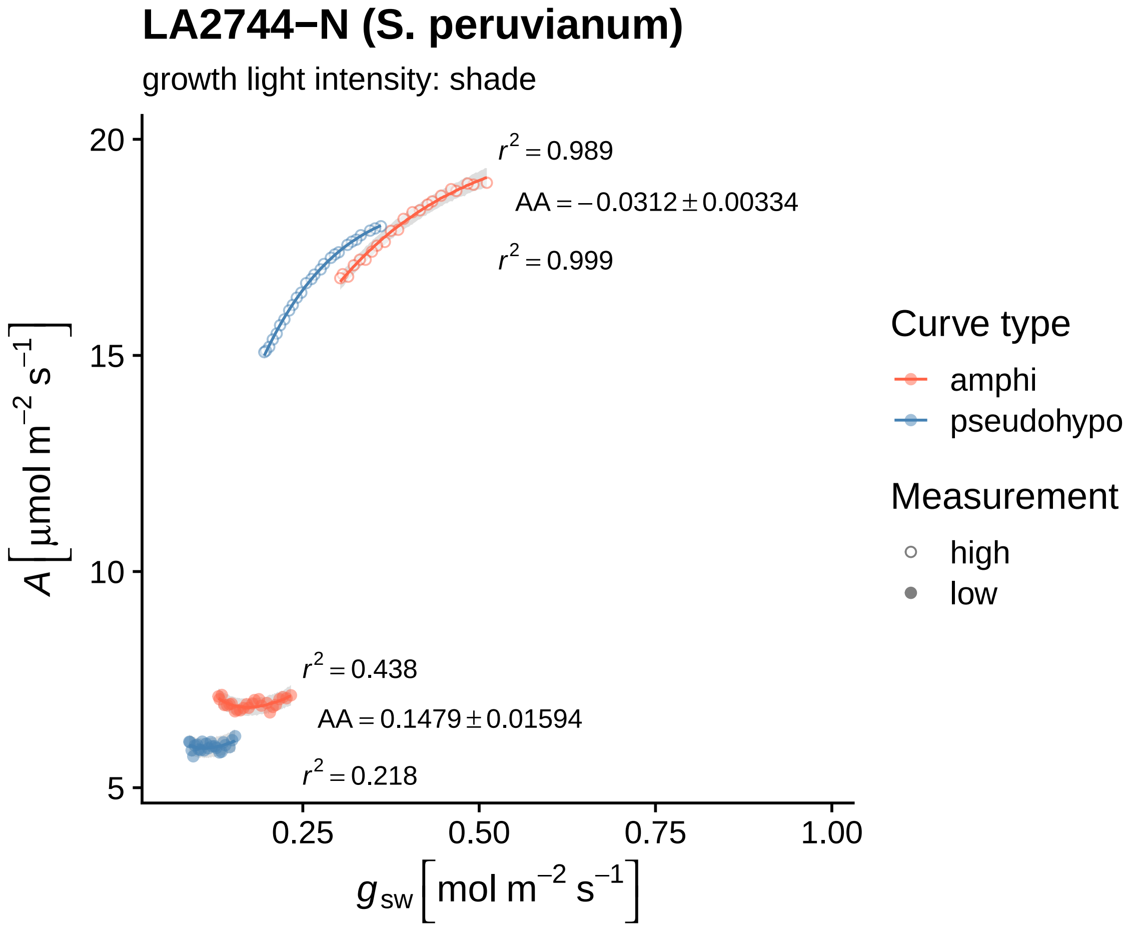

LA2744−N (S. peruvianum)
growth light intensity: shade
r² = 0.989
AA = −0.0312 ± 0.00334
r² = 0.999
r² = 0.438
AA = 0.1479 ± 0.01594
r² = 0.218
20
15
10
5
0.25
0.50
0.75
1.00
A [µmol m⁻² s⁻¹]
g sw [mol m⁻² s⁻¹]
Curve type
amphi
pseudohypo
Measurement
high
low

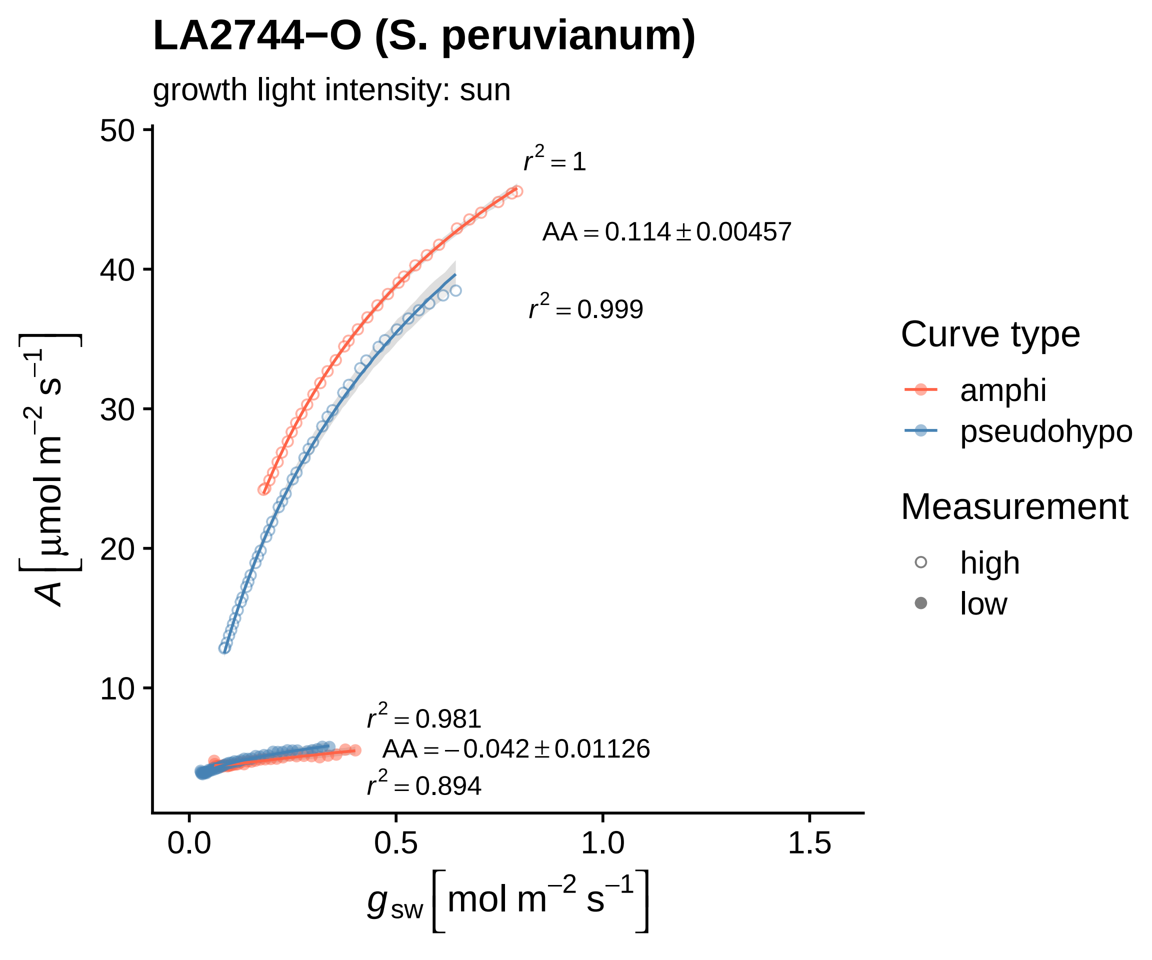

LA2744−O (S. peruvianum)
growth light intensity: sun
$r^2 = 1$
$AA = 0.114 \pm 0.00457$
$r^2 = 0.999$
$r^2 = 0.981$
$AA = -0.042 \pm 0.01126$
$r^2 = 0.894$
50
40
30
20
10
0.0
0.5
1.0
1.5
$A\left[\mu\text{mol m}^{-2}\text{ s}^{-1}\right]$
$g_{sw}\left[\text{mol m}^{-2}\text{ s}^{-1}\right]$
Curve type
amphi
pseudohypo
Measurement
high
low

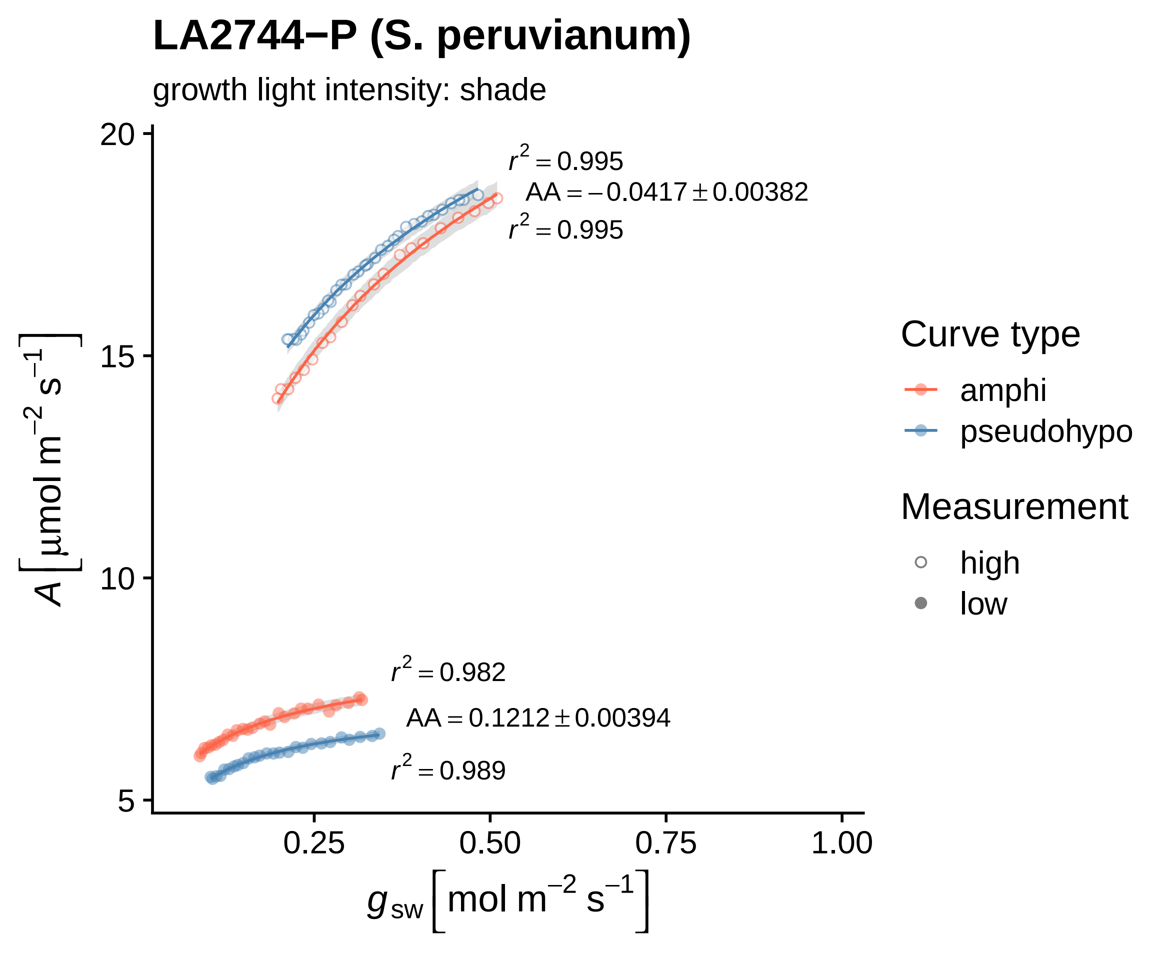

LA2744−P (S. peruvianum)
growth light intensity: shade
r² = 0.995
AA = −0.0417 ± 0.00382
r² = 0.995
r² = 0.982
AA = 0.1212 ± 0.00394
r² = 0.989
20
15
10
5
0.25
0.50
0.75
1.00
A [µmol m⁻² s⁻¹]
g sw [mol m⁻² s⁻¹]
Curve type
amphi
pseudohypo
Measurement
high
low

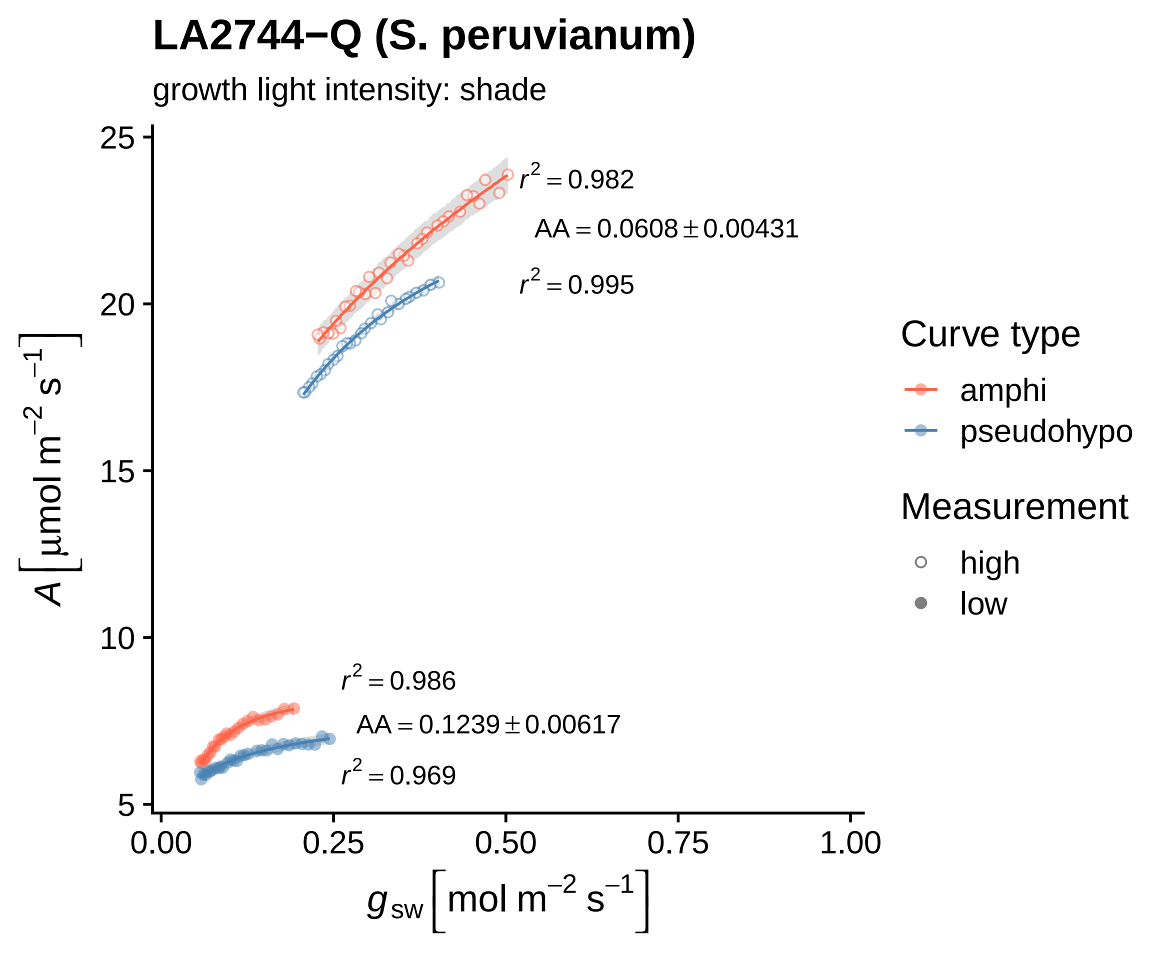

LA2744−Q (S. peruvianum)
growth light intensity: shade
r² = 0.982
AA = 0.0608 ± 0.00431
r² = 0.995
r² = 0.986
AA = 0.1239 ± 0.00617
r² = 0.969
A [μmol m⁻² s⁻¹]
25
20
15
10
5
0.00
0.25
0.50
0.75
1.00
g sw [mol m⁻² s⁻¹]
Curve type
amphi
pseudohypo
Measurement
high
low

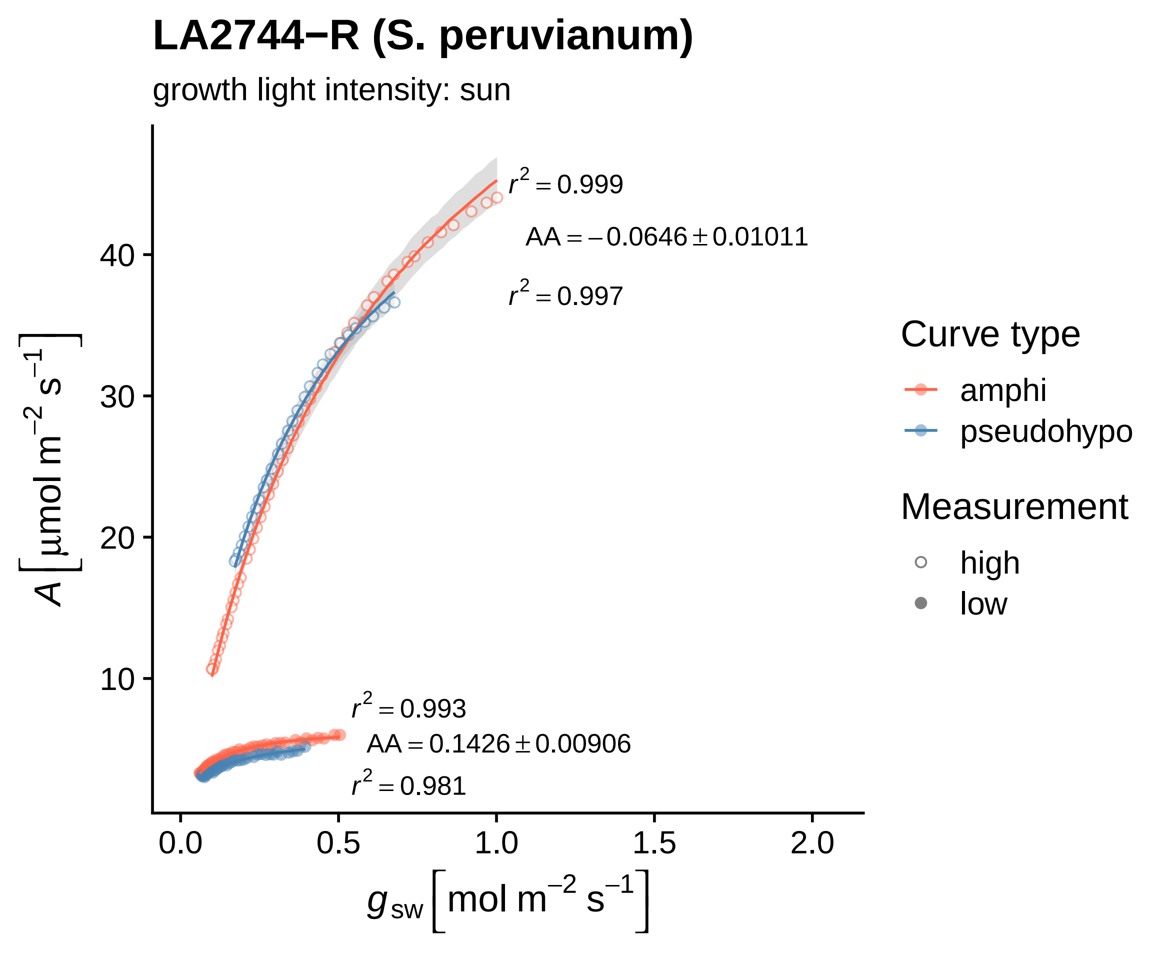
LA2744−R (S. peruvianum)
growth light intensity: sun
r² = 0.999
AA = −0.0646 ± 0.01011
r² = 0.997
r² = 0.993
AA = 0.1426 ± 0.00906
r² = 0.981
A [μmol m⁻² s⁻¹]
g_sw [mol m⁻² s⁻¹]
0.0
0.5
1.0
1.5
2.0
10
20
30
40
Curve type
amphi
pseudohypo
Measurement
high
low

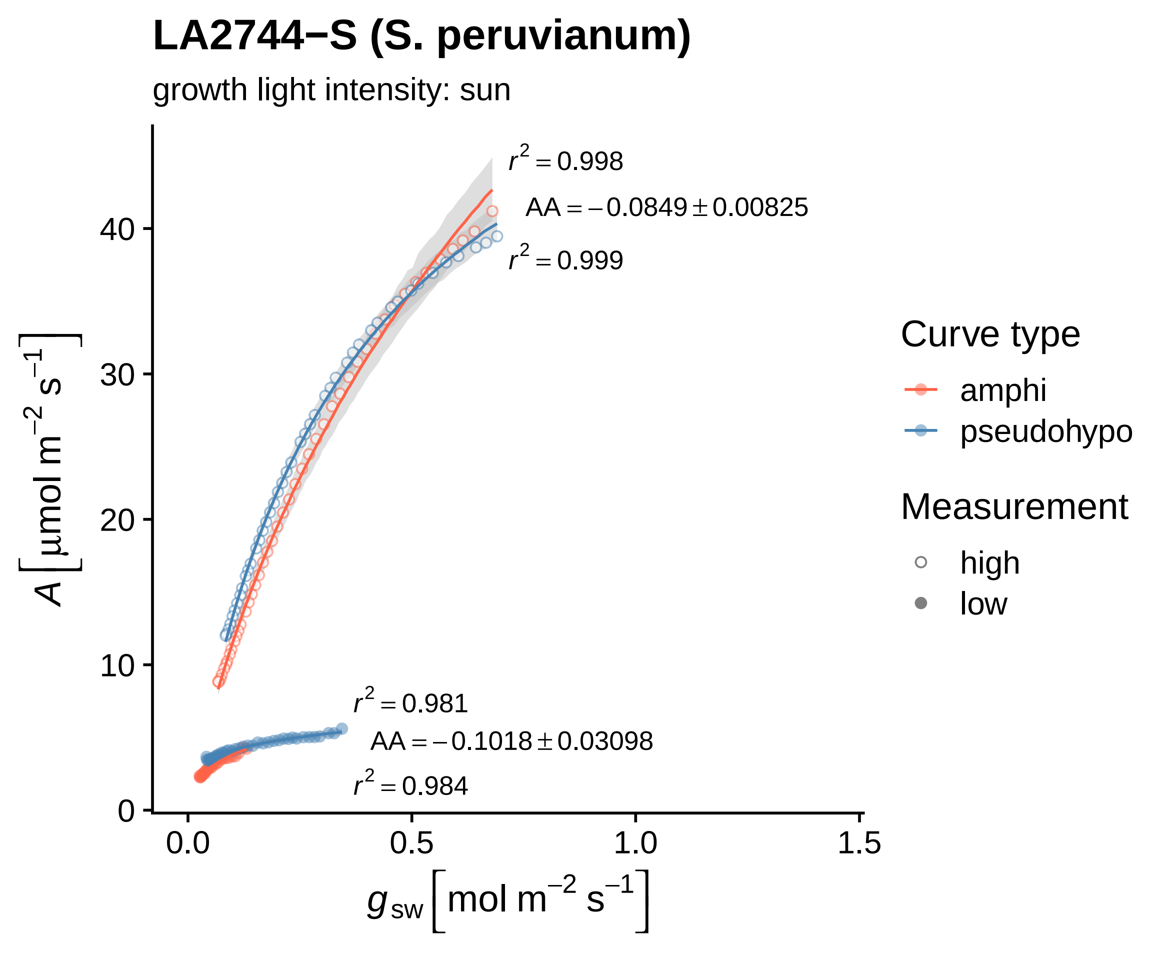
LA2744−S (S. peruvianum)
growth light intensity: sun
$r^2 = 0.998$
$AA = -0.0849 \pm 0.00825$
$r^2 = 0.999$
$r^2 = 0.981$
$AA = -0.1018 \pm 0.03098$
$r^2 = 0.984$
$A\ [\mu mol\ m^{-2}\ s^{-1}]$
0
10
20
30
40
0.0
0.5
1.0
1.5
$g_{sw}\ [mol\ m^{-2}\ s^{-1}]$
Curve type
amphi
pseudohypo
Measurement
high
low

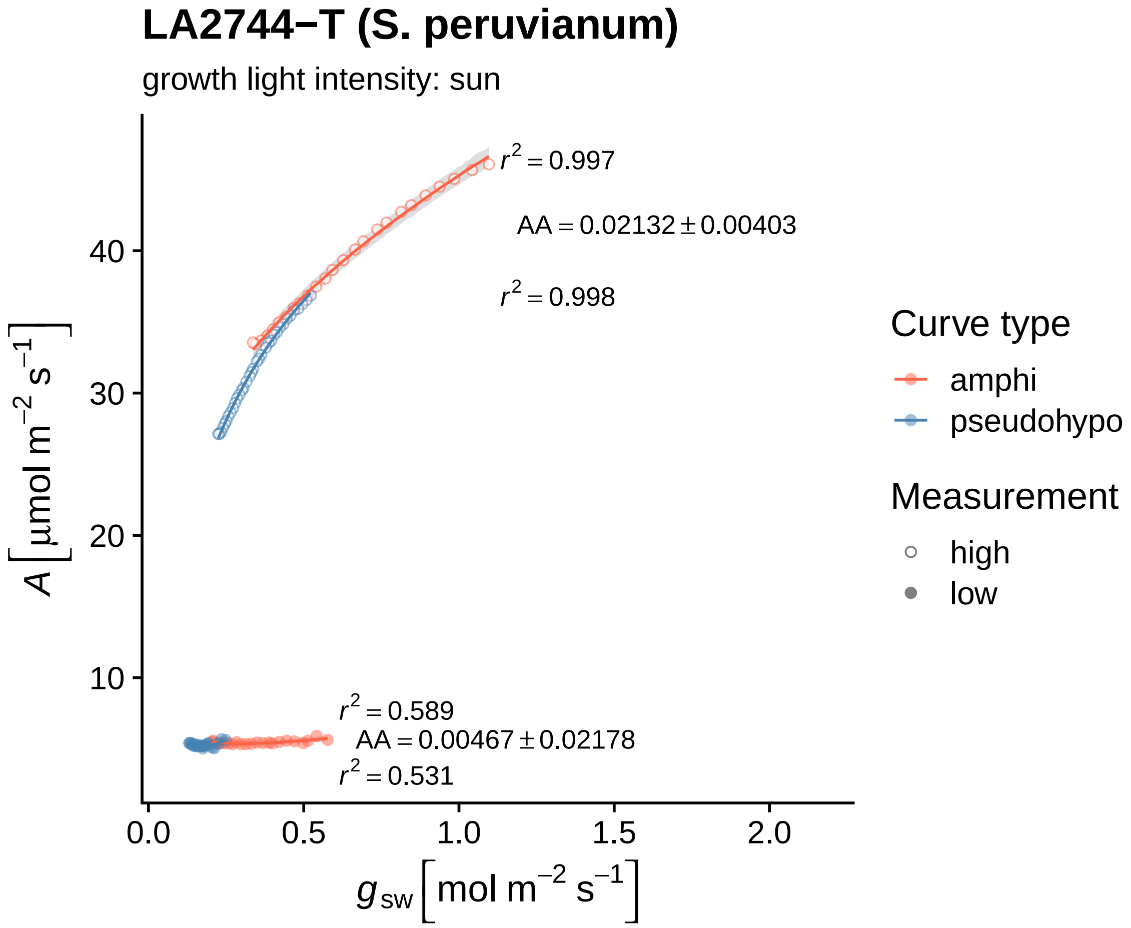

LA2744−T (S. peruvianum)
growth light intensity: sun
r2 = 0.997
AA = 0.02132 ± 0.00403
r2 = 0.998
r2 = 0.589
AA = 0.00467 ± 0.02178
r2 = 0.531
A [μmol m−2 s−1]
gsw [mol m−2 s−1]
40
30
20
10
0.0
0.5
1.0
1.5
2.0
Curve type
amphi
pseudohypo
Measurement
high
low

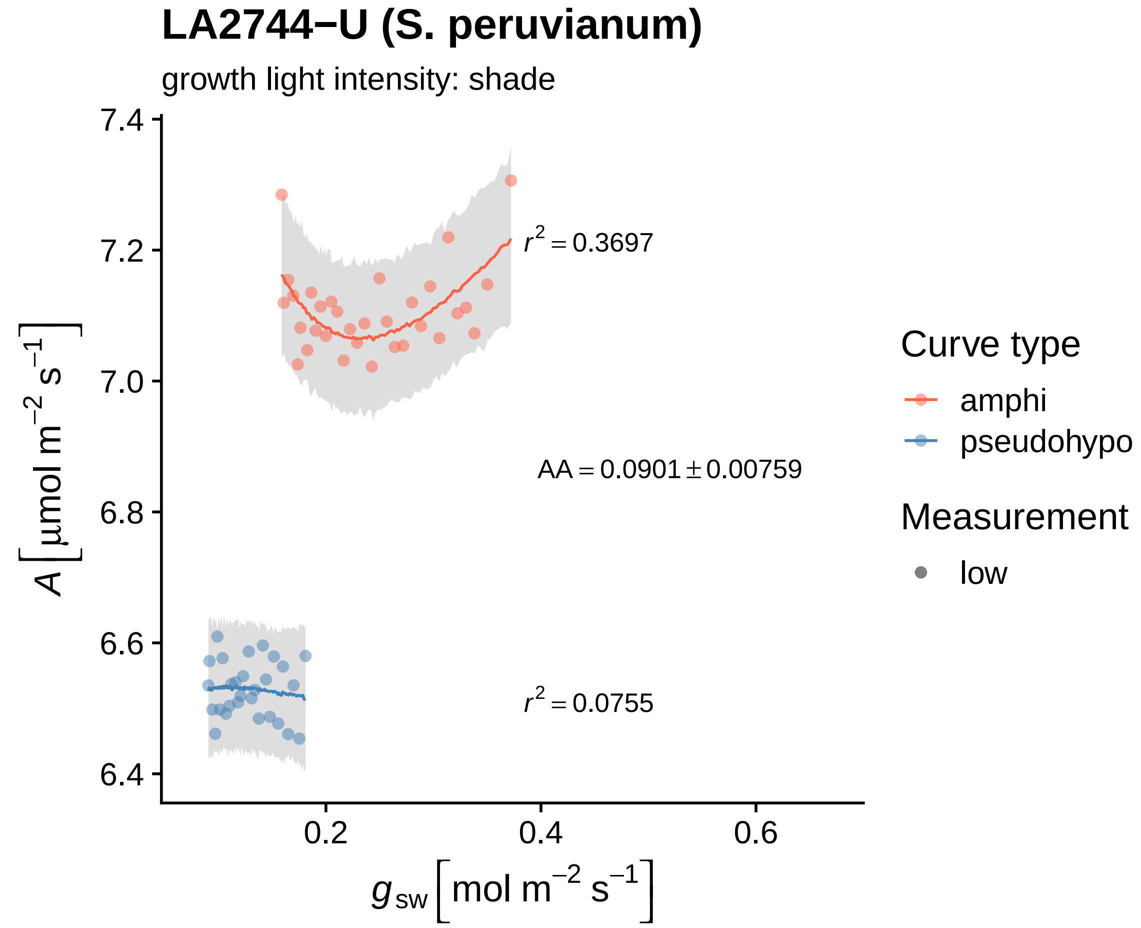

LA2744−U (S. peruvianum)
growth light intensity: shade
7.4
7.2
7.0
6.8
6.6
6.4
0.2
0.4
0.6
$r^2 = 0.3697$
$AA = 0.0901 \pm 0.00759$
$r^2 = 0.0755$
$A\left[\mu mol\ m^{-2}\ s^{-1}\right]$
$g_{sw}\left[mol\ m^{-2}\ s^{-1}\right]$
Curve type
amphi
pseudohypo
Measurement
low

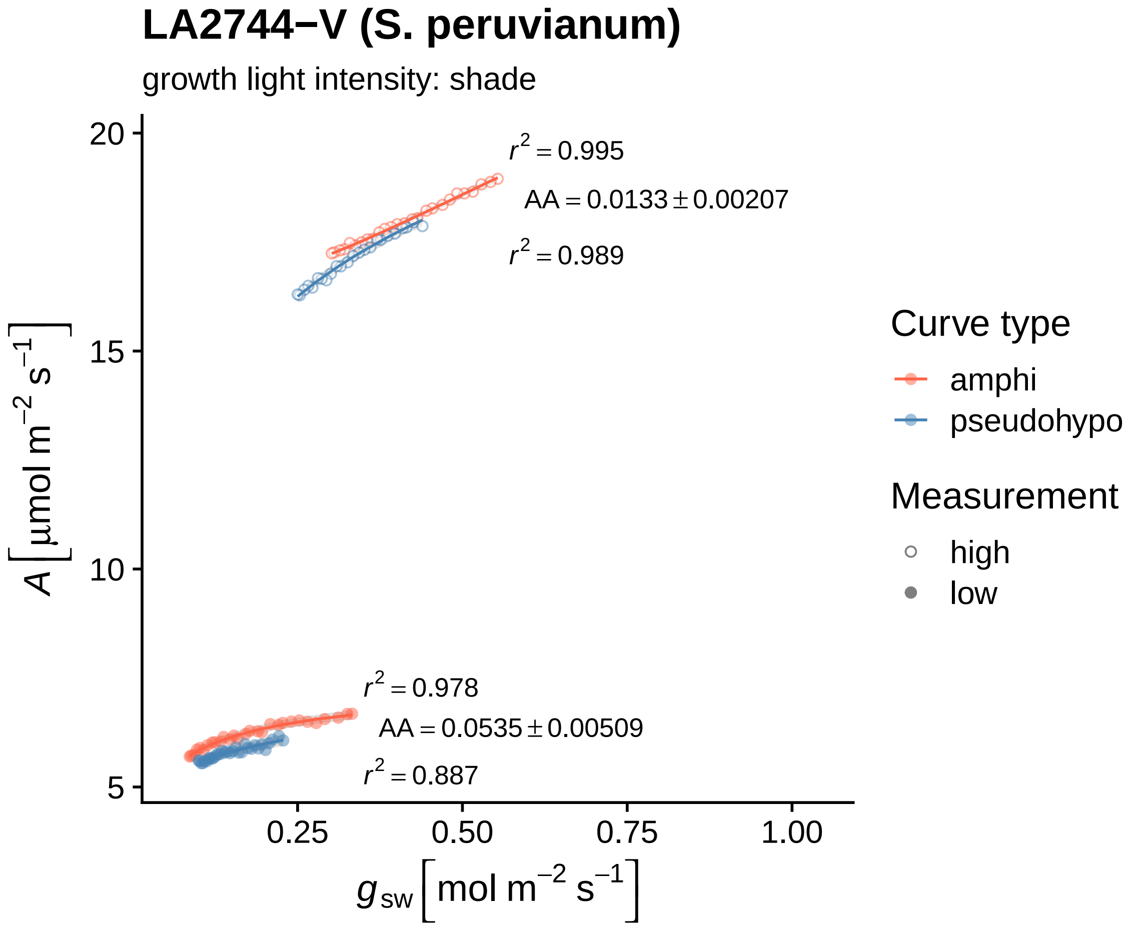

LA2744−V (S. peruvianum)
growth light intensity: shade
$r^2 = 0.995$
$AA = 0.0133 \pm 0.00207$
$r^2 = 0.989$
$r^2 = 0.978$
$AA = 0.0535 \pm 0.00509$
$r^2 = 0.887$
20
15
10
5
0.25
0.50
0.75
1.00
$A\ [\mu mol\ m^{-2}\ s^{-1}]$
$g_{sw}\ [mol\ m^{-2}\ s^{-1}]$
Curve type
amphi
pseudohypo
Measurement
high
low

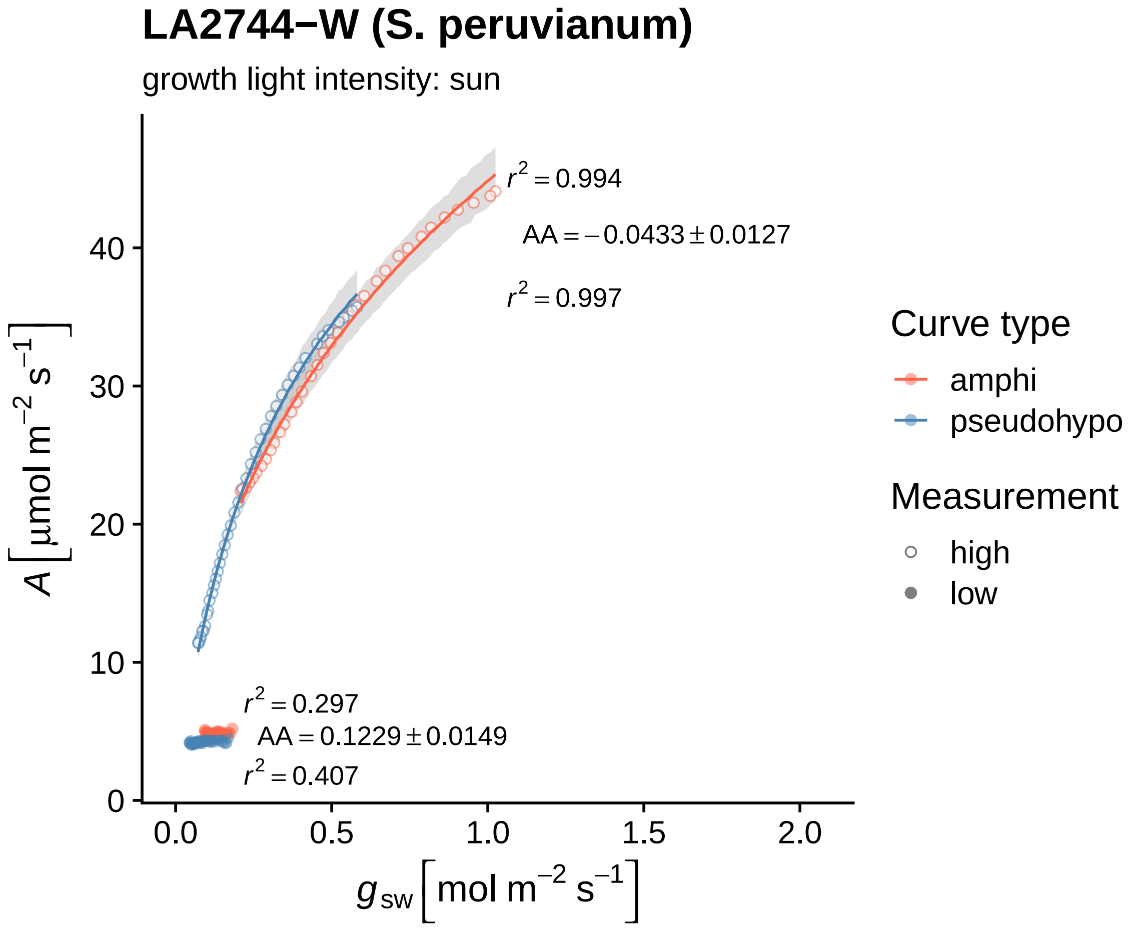

LA2744−W (S. peruvianum)
growth light intensity: sun
$r^2 = 0.994$
$AA = -0.0433 \pm 0.0127$
$r^2 = 0.997$
$r^2 = 0.297$
$AA = 0.1229 \pm 0.0149$
$r^2 = 0.407$
0
10
20
30
40
0.0
0.5
1.0
1.5
2.0
$A\,[\mu mol\ m^{-2}\ s^{-1}]$
$g_{sw}\,[mol\ m^{-2}\ s^{-1}]$
Curve type
amphi
pseudohypo
Measurement
high
low

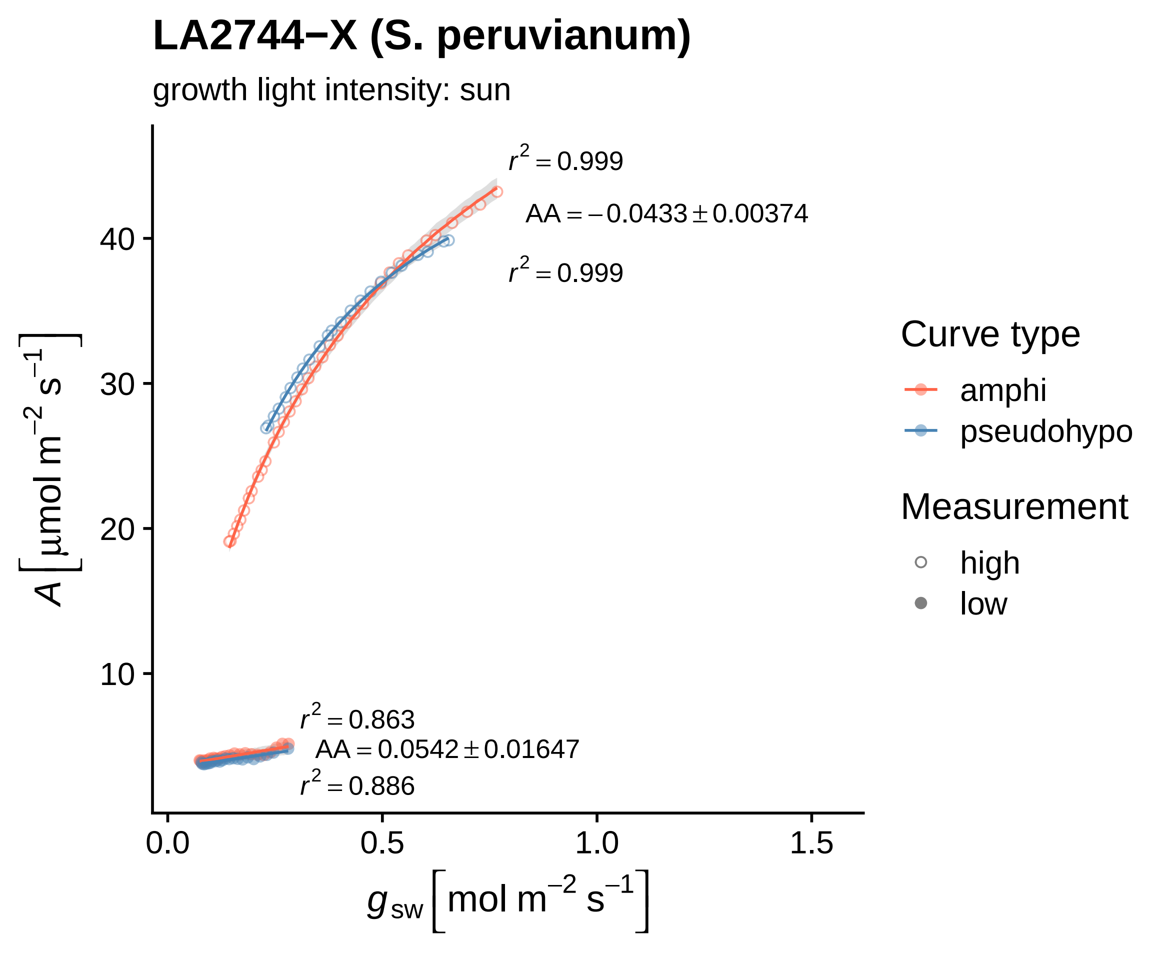
LA2744−X (S. peruvianum)
growth light intensity: sun
$r^2 = 0.999$
$AA = -0.0433 \pm 0.00374$
$r^2 = 0.999$
$r^2 = 0.863$
$AA = 0.0542 \pm 0.01647$
$r^2 = 0.886$
$A\ [\mu mol\ m^{-2}\ s^{-1}]$
$g_{sw}\ [mol\ m^{-2}\ s^{-1}]$
10
20
30
40
0.0
0.5
1.0
1.5
Curve type
amphi
pseudohypo
Measurement
high
low

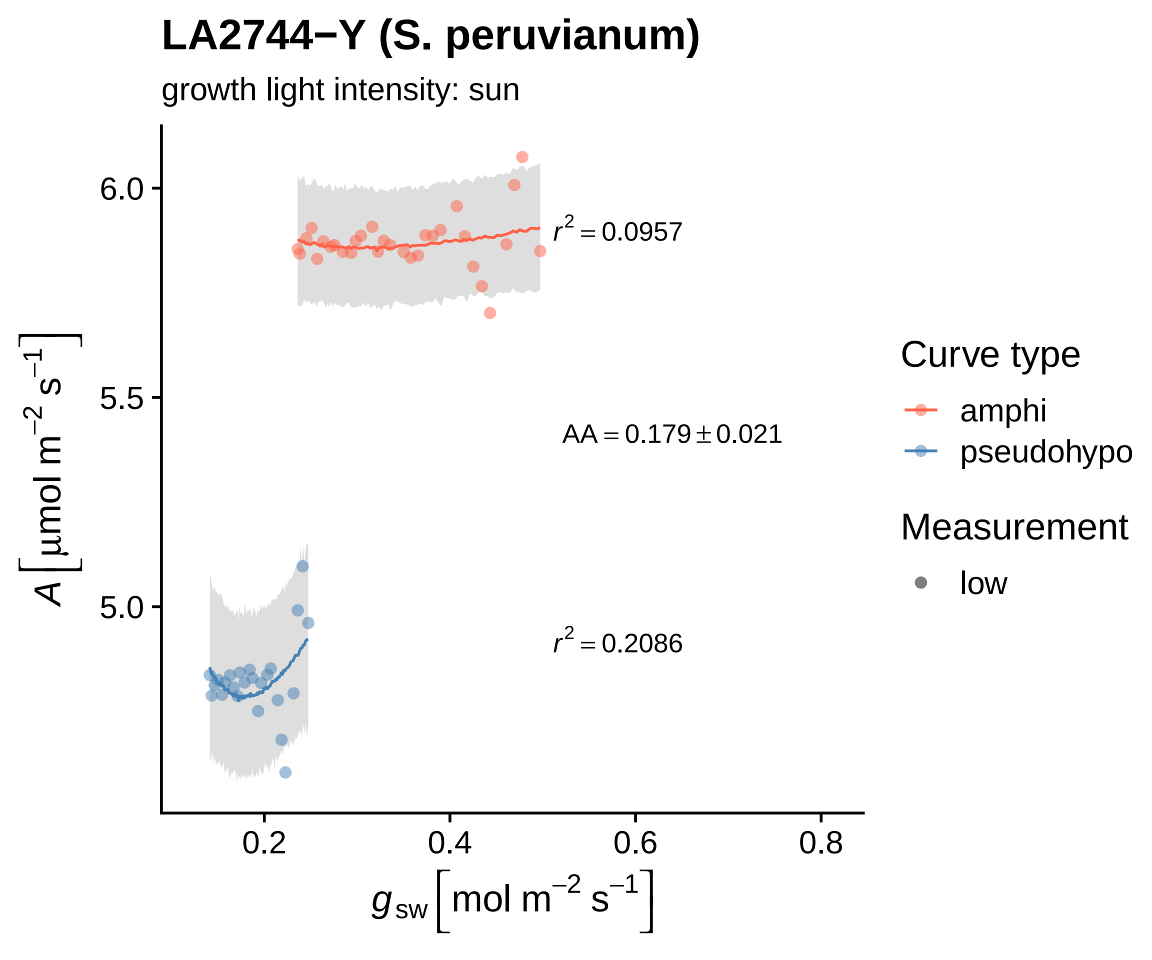
LA2744−Y (S. peruvianum)
growth light intensity: sun
$A\left[\mu mol\ m^{-2}\ s^{-1}\right]$
6.0
5.5
5.0
0.2
0.4
0.6
0.8
$g_{sw}\left[mol\ m^{-2}\ s^{-1}\right]$
$r^2 = 0.0957$
$AA = 0.179 \pm 0.021$
$r^2 = 0.2086$
Curve type
amphi
pseudohypo
Measurement
low

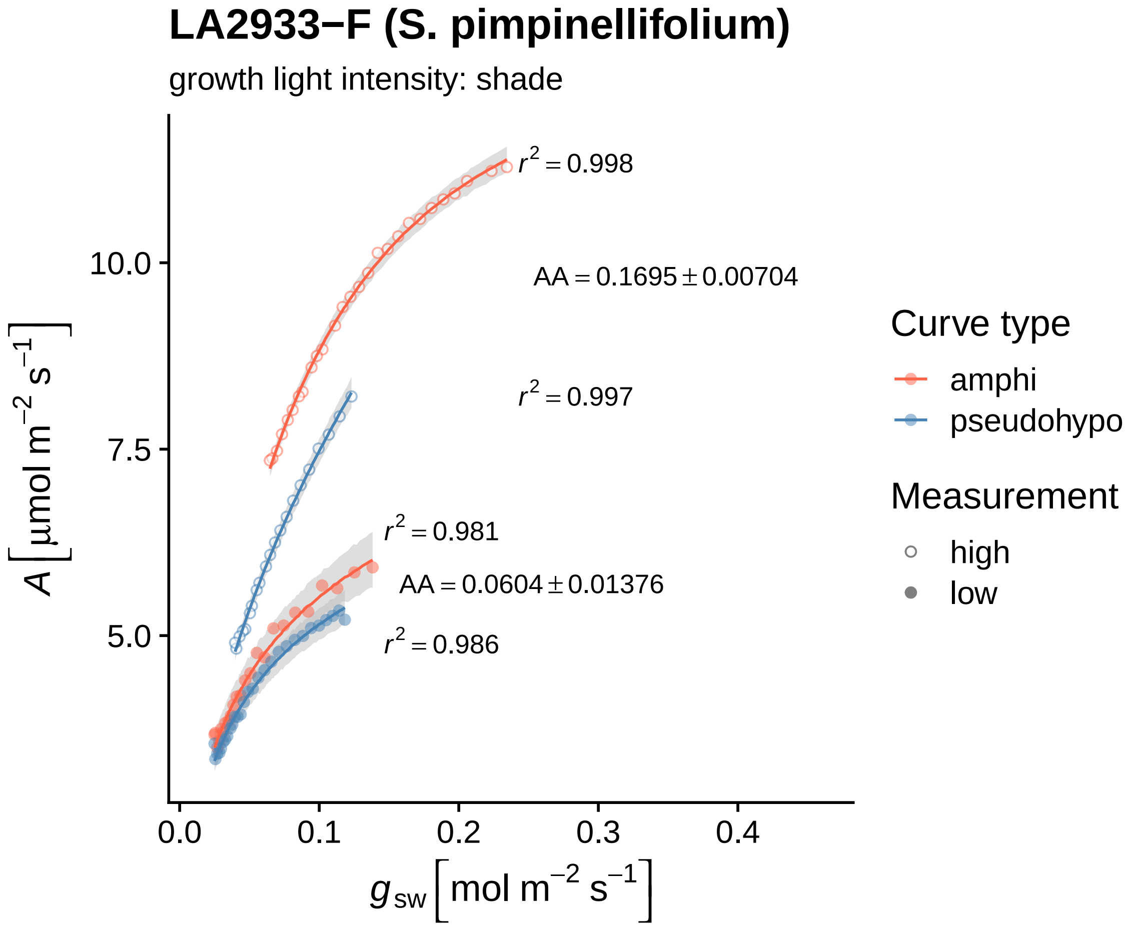

LA2933−F (S. pimpinellifolium)
growth light intensity: shade
$r^2 = 0.998$
AA = 0.1695 ± 0.00704
$r^2 = 0.997$
$r^2 = 0.981$
AA = 0.0604 ± 0.01376
$r^2 = 0.986$
10.0
7.5
5.0
0.0
0.1
0.2
0.3
0.4
$A\,[\mu mol\ m^{-2}\ s^{-1}]$
$g_{sw}\,[mol\ m^{-2}\ s^{-1}]$
Curve type
amphi
pseudohypo
Measurement
high
low

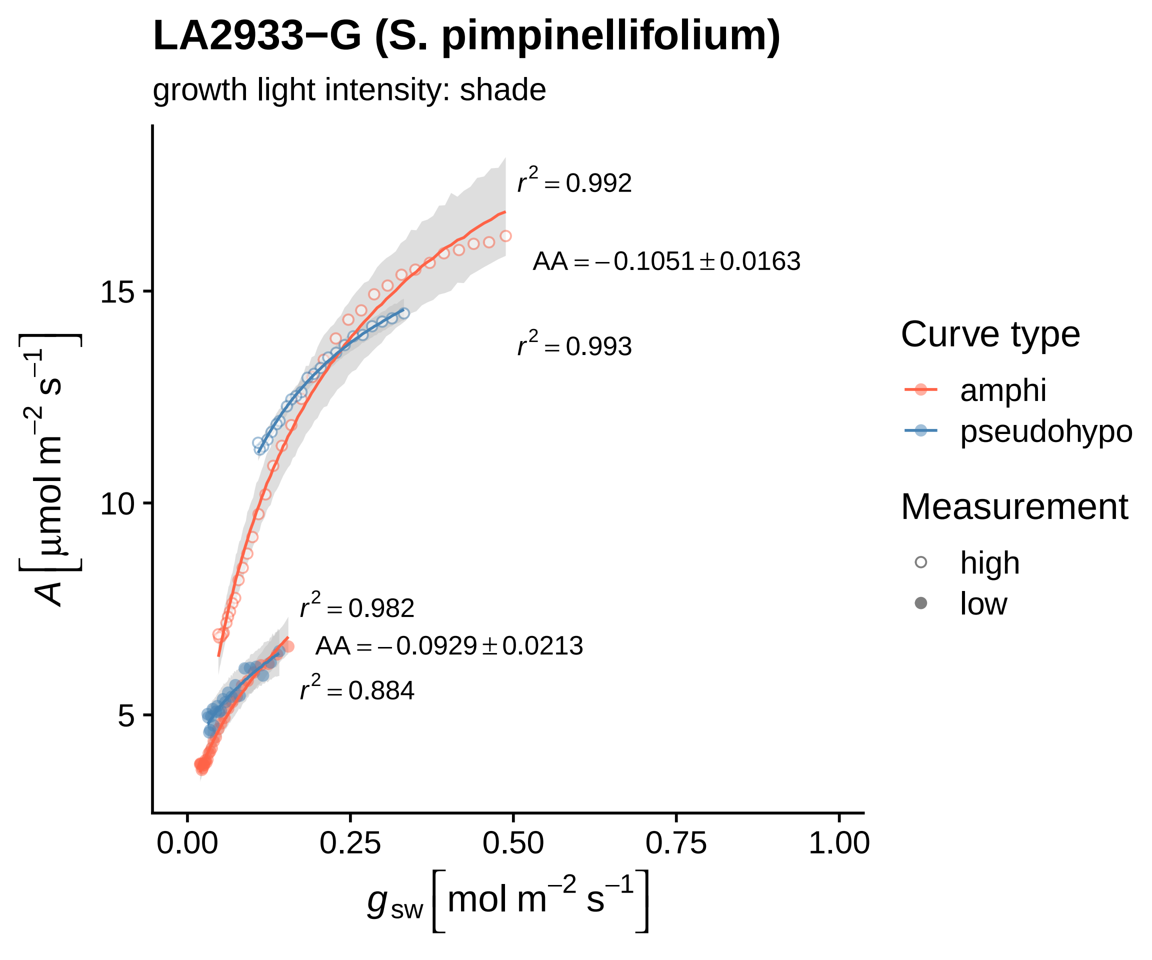
LA2933−G (S. pimpinellifolium)
growth light intensity: shade
r^2 = 0.992
AA = −0.1051 ± 0.0163
r^2 = 0.993
r^2 = 0.982
AA = −0.0929 ± 0.0213
r^2 = 0.884
15
10
5
0.00
0.25
0.50
0.75
1.00
A [μmol m^−2 s^−1]
g_sw [mol m^−2 s^−1]
Curve type
amphi
pseudohypo
Measurement
high
low

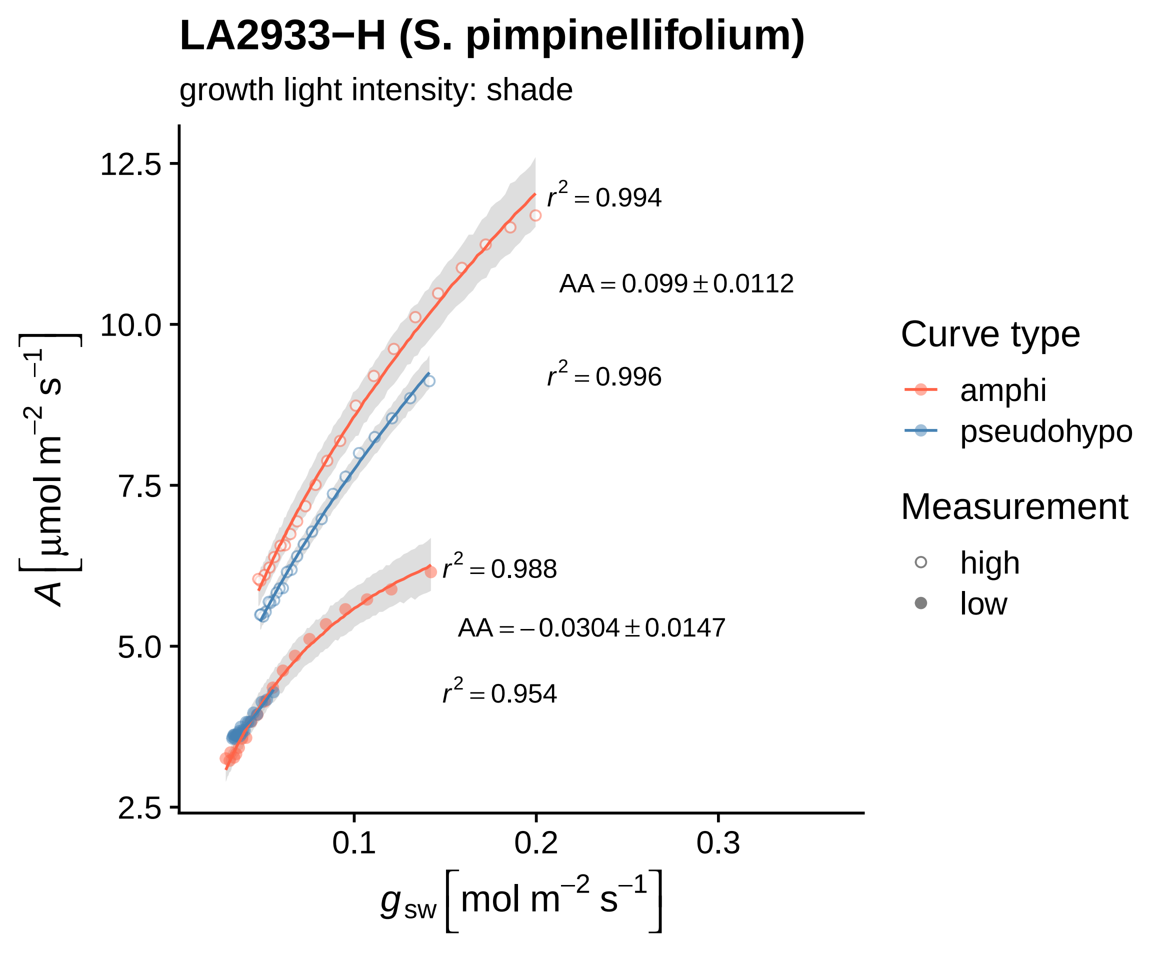
LA2933−H (S. pimpinellifolium)
growth light intensity: shade
$r^2 = 0.994$
$AA = 0.099 \pm 0.0112$
$r^2 = 0.996$
$r^2 = 0.988$
$AA = -0.0304 \pm 0.0147$
$r^2 = 0.954$
12.5
10.0
7.5
5.0
2.5
0.1
0.2
0.3
$A\,[\mu mol\ m^{-2}\ s^{-1}]$
$g_{sw}\,[mol\ m^{-2}\ s^{-1}]$
Curve type
amphi
pseudohypo
Measurement
high
low

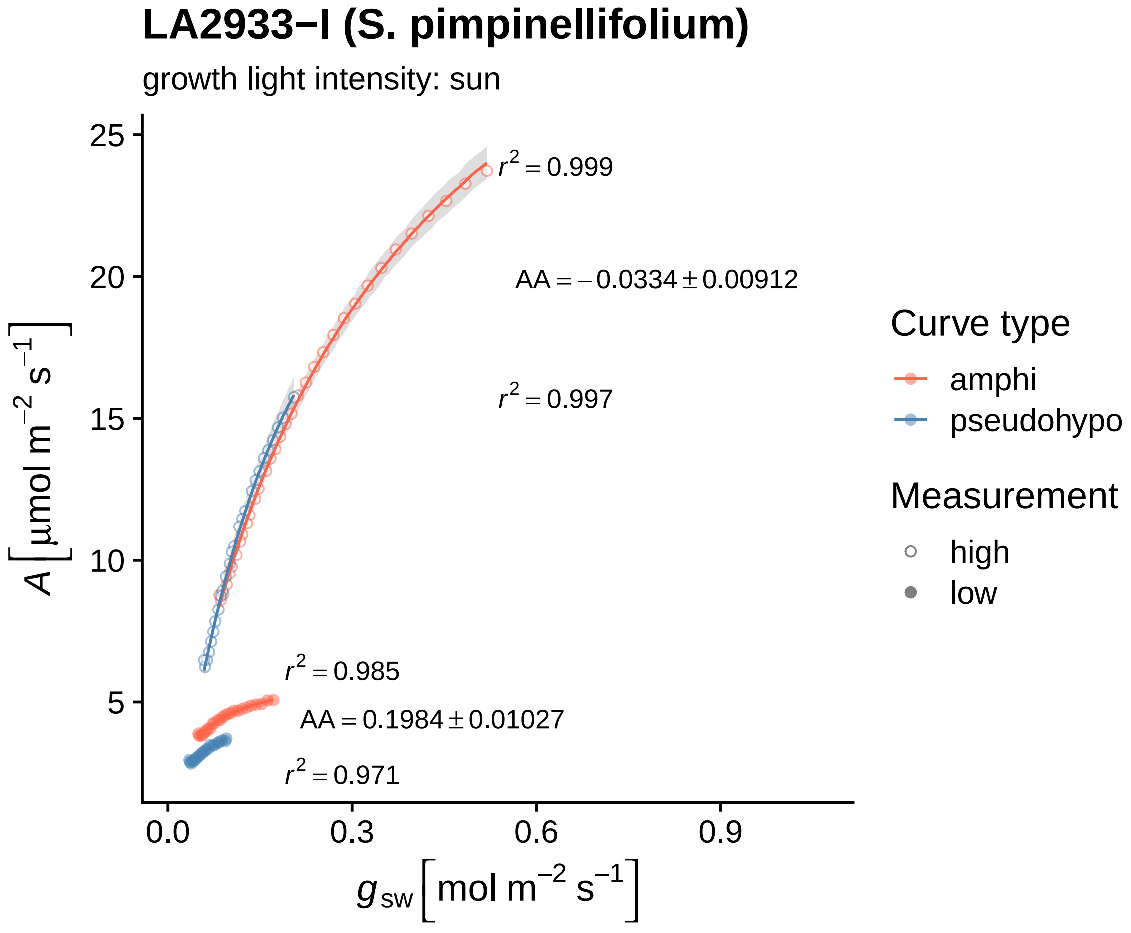

LA2933−I (S. pimpinellifolium)
growth light intensity: sun
$r^2 = 0.999$
$AA = -0.0334 \pm 0.00912$
$r^2 = 0.997$
$r^2 = 0.985$
$AA = 0.1984 \pm 0.01027$
$r^2 = 0.971$
25
20
15
10
5
0.0
0.3
0.6
0.9
$A\ [\mu mol\ m^{-2}\ s^{-1}]$
$g_{sw}\ [mol\ m^{-2}\ s^{-1}]$
Curve type
amphi
pseudohypo
Measurement
high
low

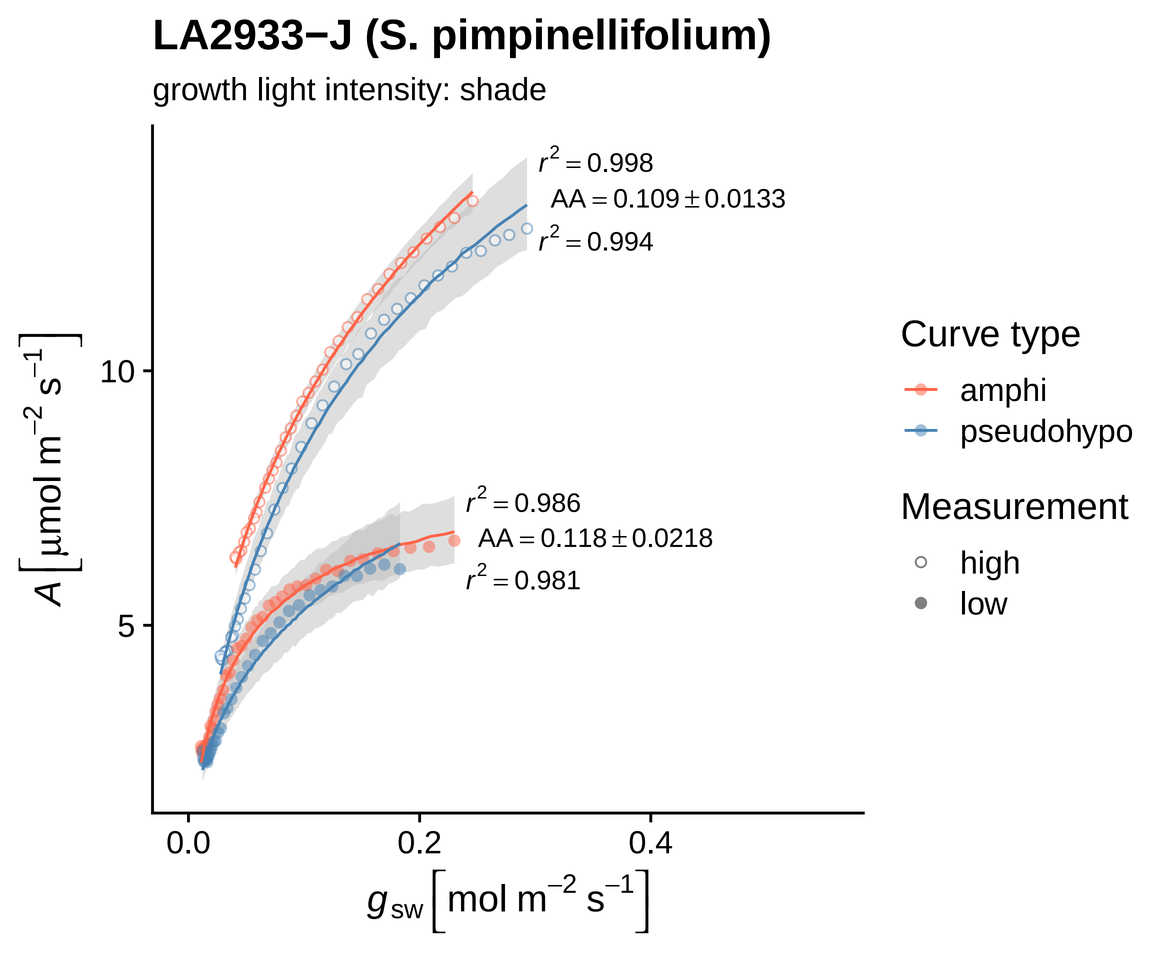
LA2933−J (S. pimpinellifolium)
growth light intensity: shade
$r^2 = 0.998$
$AA = 0.109 \pm 0.0133$
$r^2 = 0.994$
$r^2 = 0.986$
$AA = 0.118 \pm 0.0218$
$r^2 = 0.981$
$A\left[\mu mol\ m^{-2}\ s^{-1}\right]$
10
5
0.0
0.2
0.4
$g_{sw}\left[mol\ m^{-2}\ s^{-1}\right]$
Curve type
amphi
pseudohypo
Measurement
high
low

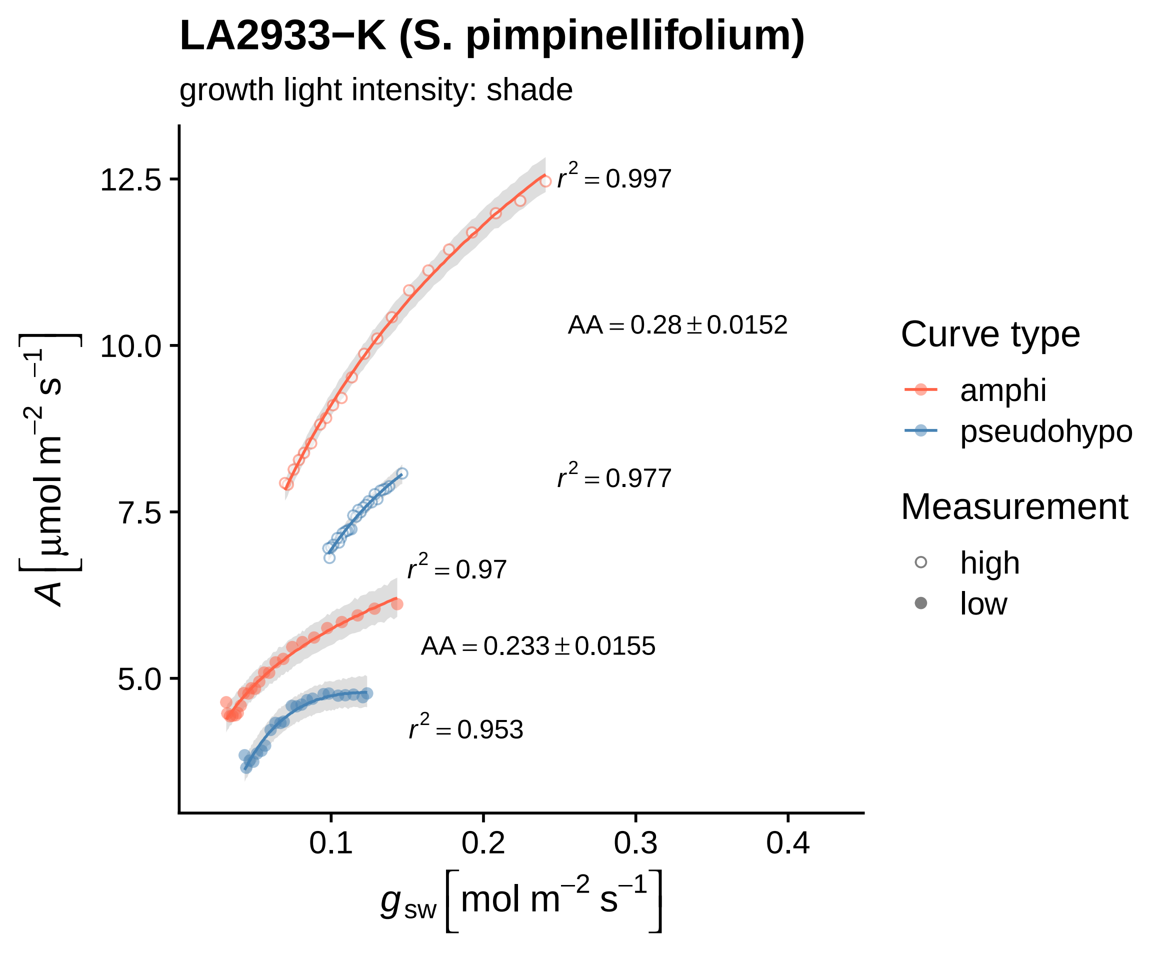
LA2933−K (S. pimpinellifolium)
growth light intensity: shade
$r^2 = 0.997$
AA = 0.28 ± 0.0152
$r^2 = 0.977$
$r^2 = 0.97$
AA = 0.233 ± 0.0155
$r^2 = 0.953$
12.5
10.0
7.5
5.0
0.1
0.2
0.3
0.4
$A\ [\mu mol\ m^{-2}\ s^{-1}]$
$g_{sw}\ [mol\ m^{-2}\ s^{-1}]$
Curve type
amphi
pseudohypo
Measurement
high
low

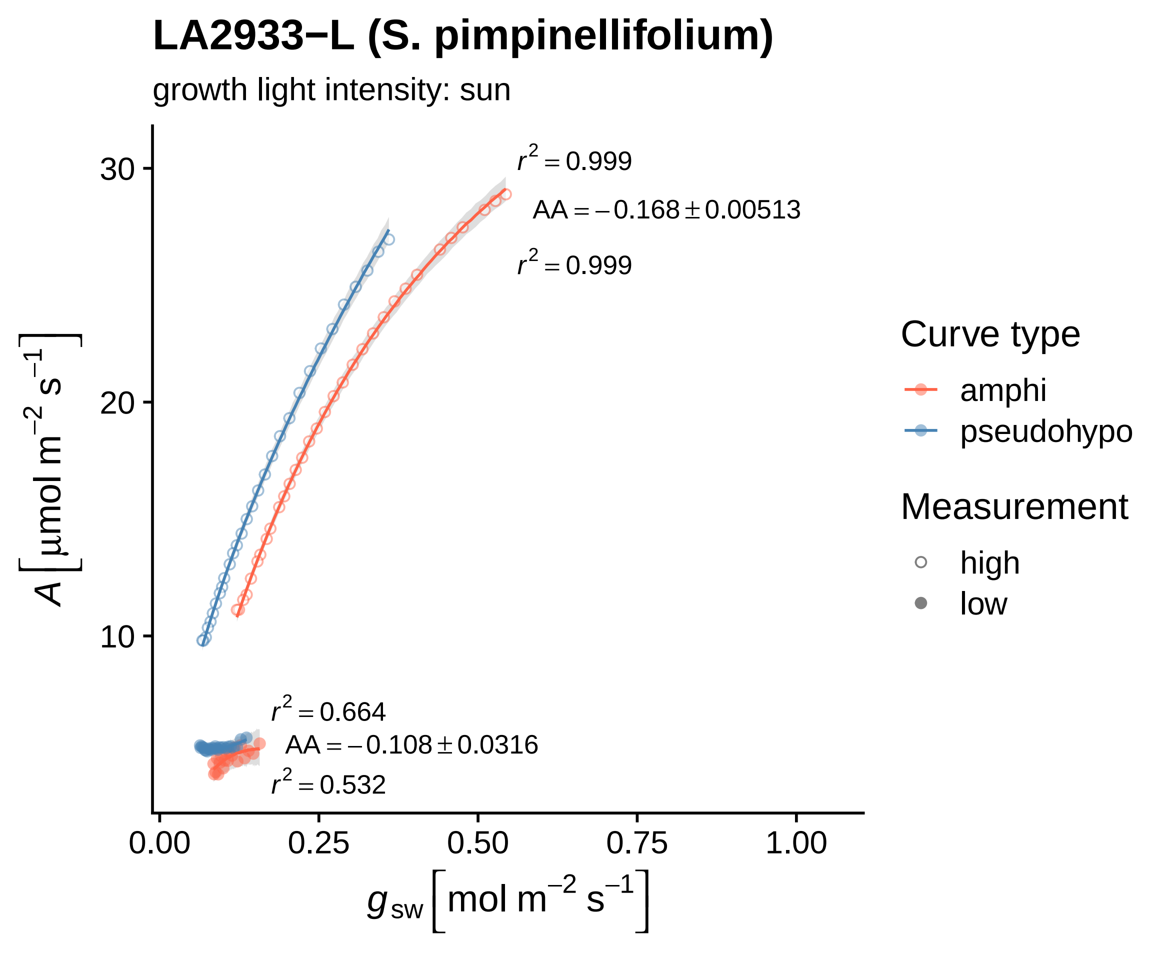
LA2933−L (S. pimpinellifolium)
growth light intensity: sun
r² = 0.999
AA = −0.168 ± 0.00513
r² = 0.999
r² = 0.664
AA = −0.108 ± 0.0316
r² = 0.532
30
20
10
0.00
0.25
0.50
0.75
1.00
A [μmol m⁻² s⁻¹]
g sw [mol m⁻² s⁻¹]
Curve type
amphi
pseudohypo
Measurement
high
low

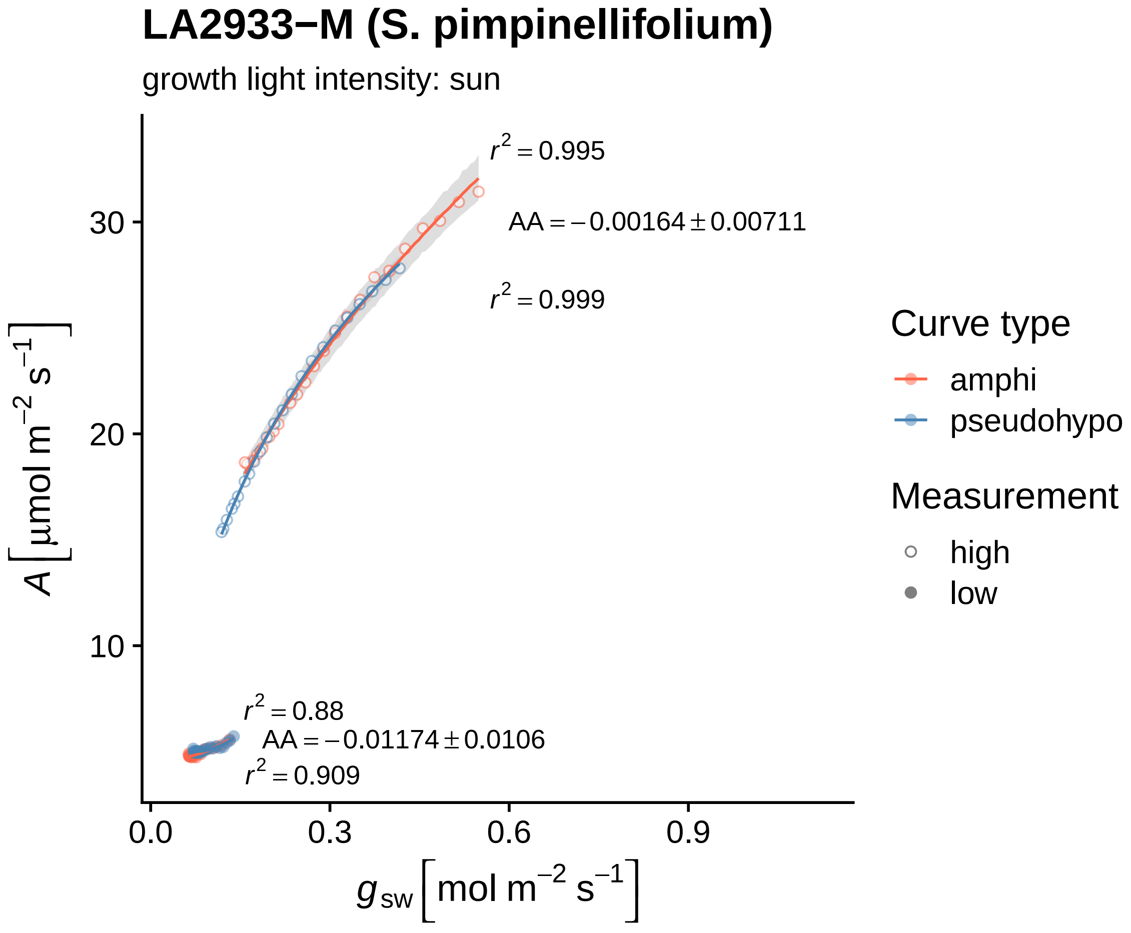

LA2933−M (S. pimpinellifolium)
growth light intensity: sun
r² = 0.995
AA = −0.00164 ± 0.00711
r² = 0.999
r² = 0.88
AA = −0.01174 ± 0.0106
r² = 0.909
30
20
10
0.0
0.3
0.6
0.9
A [µmol m⁻² s⁻¹]
gsw [mol m⁻² s⁻¹]
Curve type
amphi
pseudohypo
Measurement
high
low

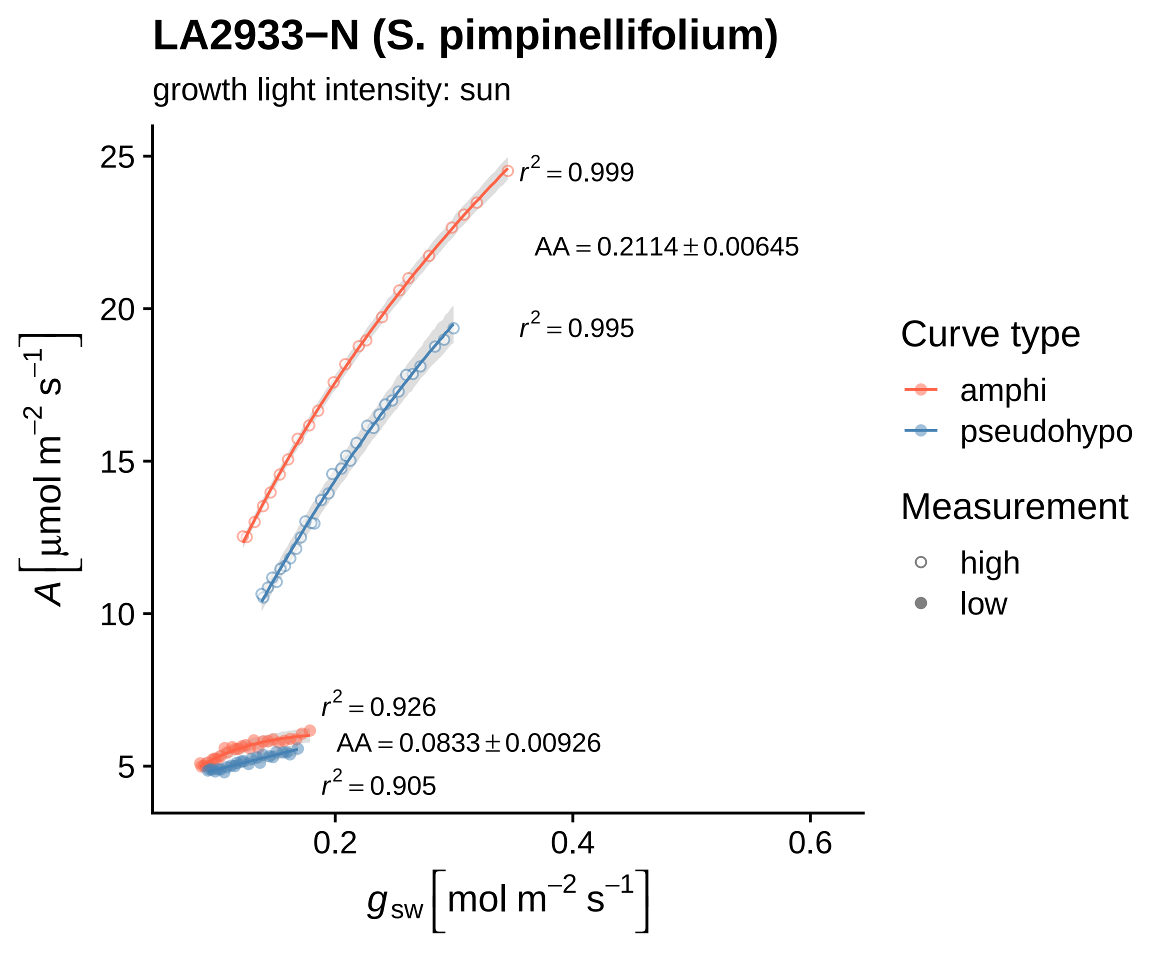
LA2933−N (S. pimpinellifolium)
growth light intensity: sun
$r^2 = 0.999$
$AA = 0.2114 \pm 0.00645$
$r^2 = 0.995$
$r^2 = 0.926$
$AA = 0.0833 \pm 0.00926$
$r^2 = 0.905$
25
20
15
10
5
0.2
0.4
0.6
$A\left[\mu mol\ m^{-2}\ s^{-1}\right]$
$g_{sw}\left[mol\ m^{-2}\ s^{-1}\right]$
Curve type
amphi
pseudohypo
Measurement
high
low

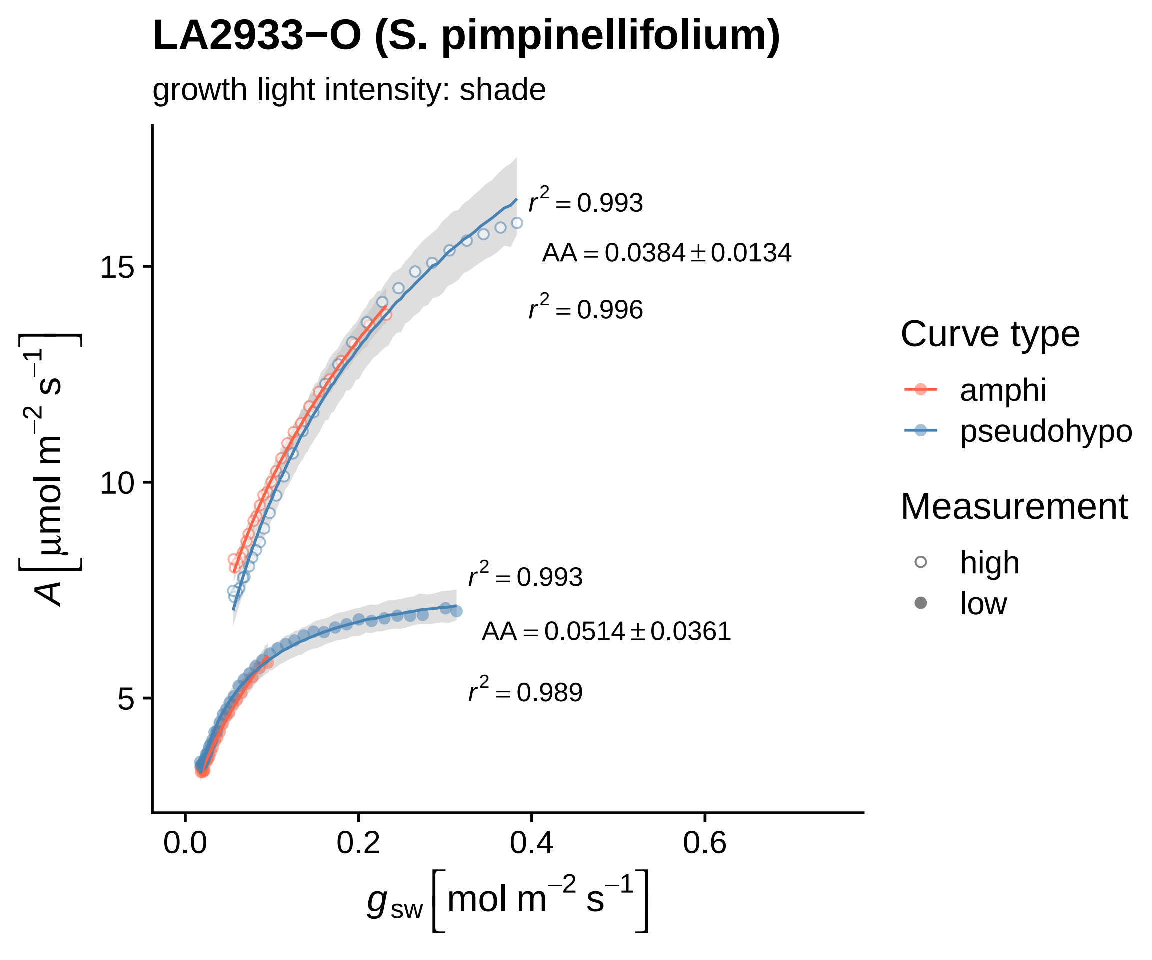

LA2933−O (S. pimpinellifolium)
growth light intensity: shade
$r^2 = 0.993$
$AA = 0.0384 \pm 0.0134$
$r^2 = 0.996$
$r^2 = 0.993$
$AA = 0.0514 \pm 0.0361$
$r^2 = 0.989$
15
10
5
0.0
0.2
0.4
0.6
$A\ [\mu mol\ m^{-2}\ s^{-1}]$
$g_{sw}\ [mol\ m^{-2}\ s^{-1}]$
Curve type
amphi
pseudohypo
Measurement
high
low

### LA2933−P (S. pimpinellifolium)

growth light intensity: shade

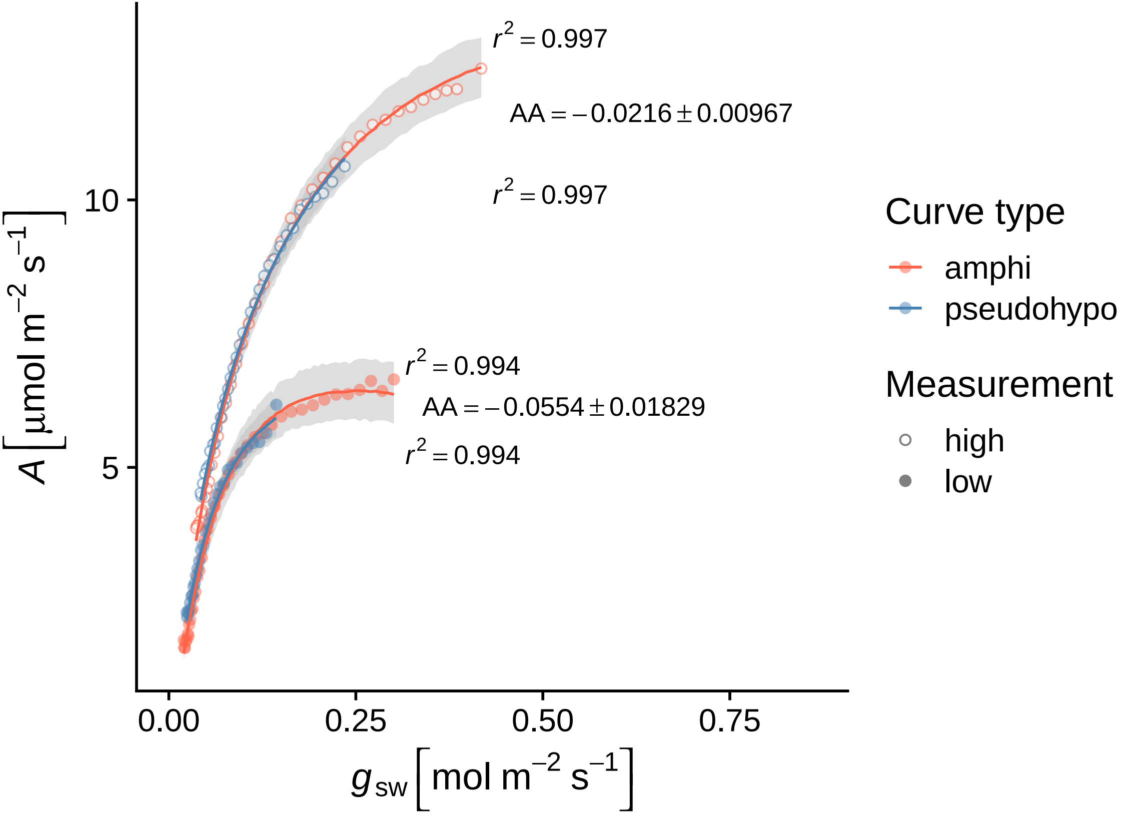

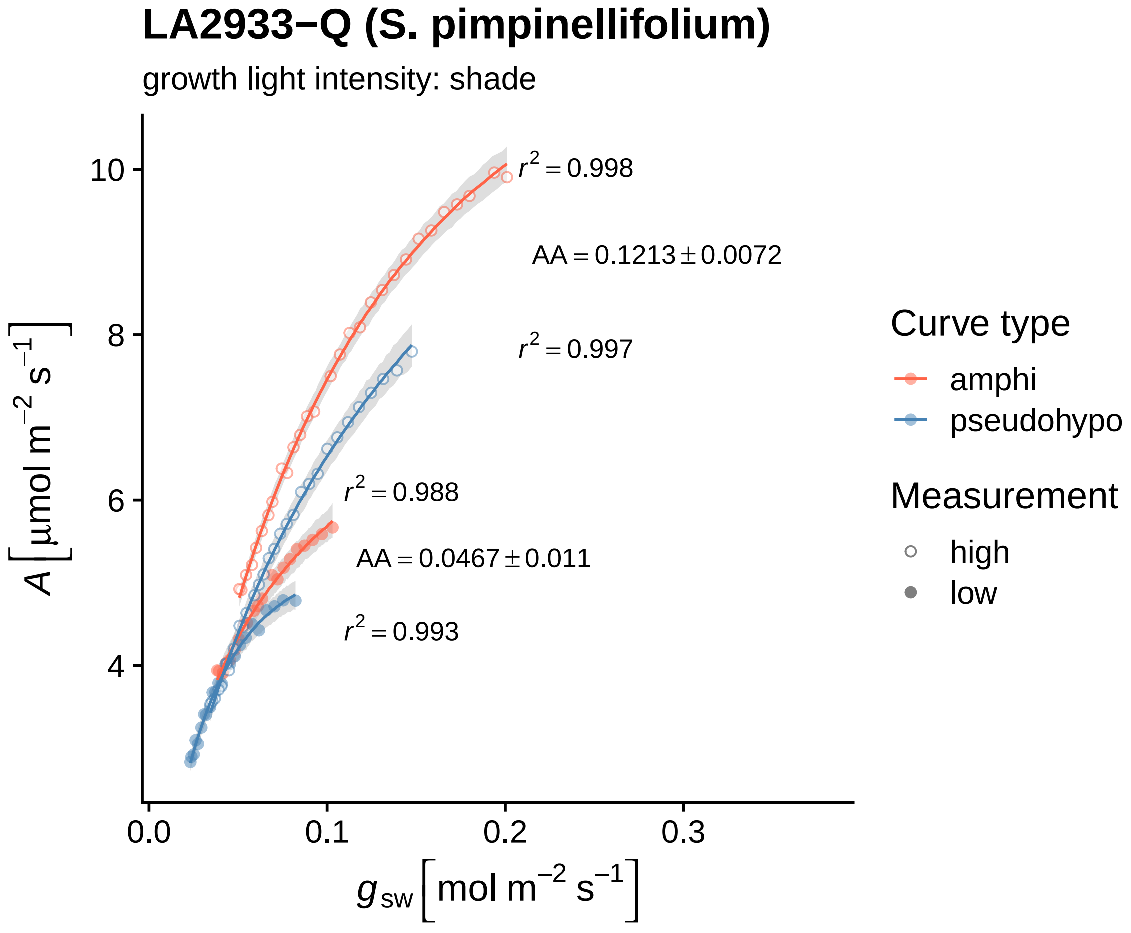

LA2933−Q (S. pimpinellifolium)
growth light intensity: shade
$r^2 = 0.998$
AA = 0.1213 ± 0.0072
$r^2 = 0.997$
$r^2 = 0.988$
AA = 0.0467 ± 0.011
$r^2 = 0.993$
10
8
6
4
0.0
0.1
0.2
0.3
$A\ [\mu mol\ m^{-2}\ s^{-1}]$
$g_{sw}\ [mol\ m^{-2}\ s^{-1}]$
Curve type
amphi
pseudohypo
Measurement
high
low

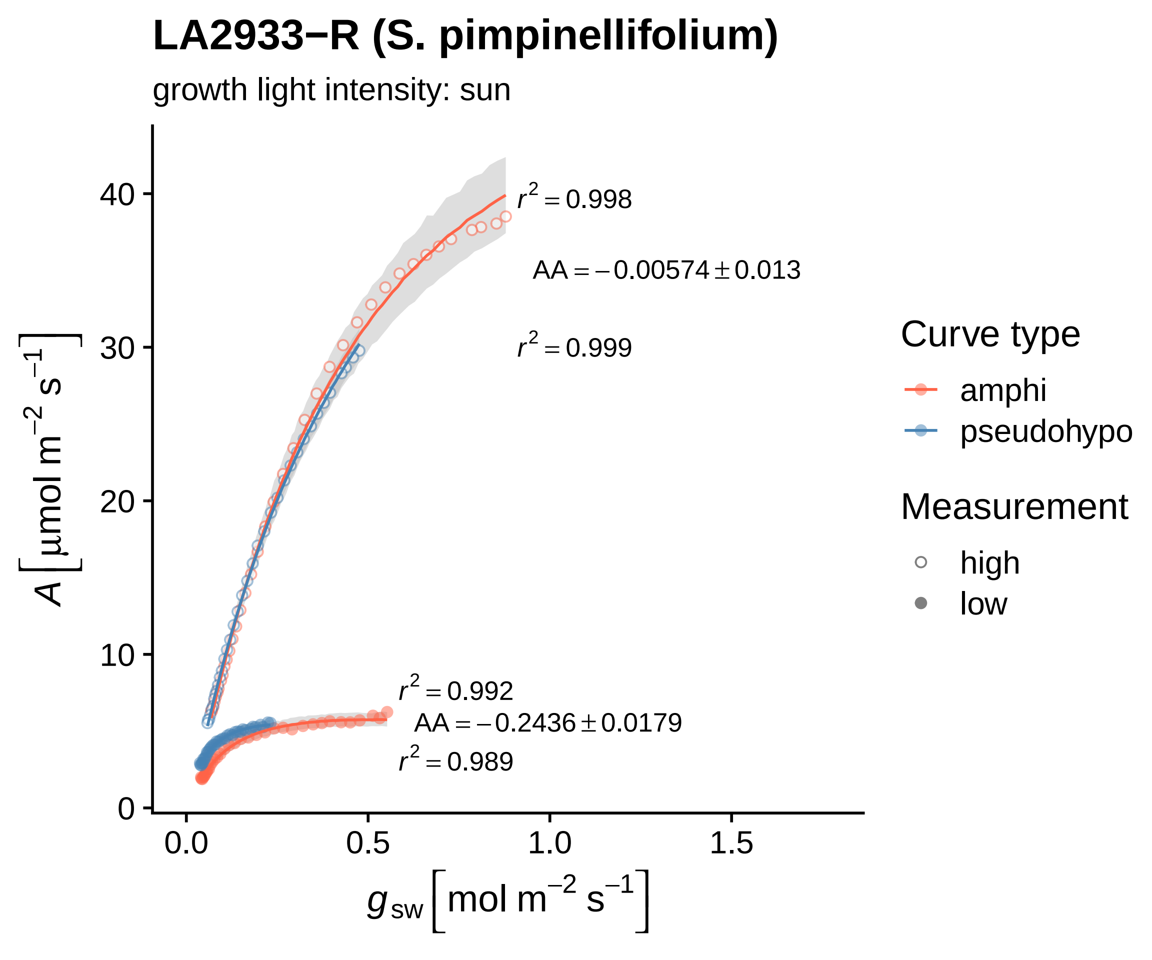

LA2933−R (S. pimpinellifolium)
growth light intensity: sun
$r^2 = 0.998$
$AA = -0.00574 \pm 0.013$
$r^2 = 0.999$
$r^2 = 0.992$
$AA = -0.2436 \pm 0.0179$
$r^2 = 0.989$
40
30
20
10
0
0.0
0.5
1.0
1.5
$A \left[ \mu mol\ m^{-2}\ s^{-1} \right]$
$g_{sw} \left[ mol\ m^{-2}\ s^{-1} \right]$
Curve type
amphi
pseudohypo
Measurement
high
low

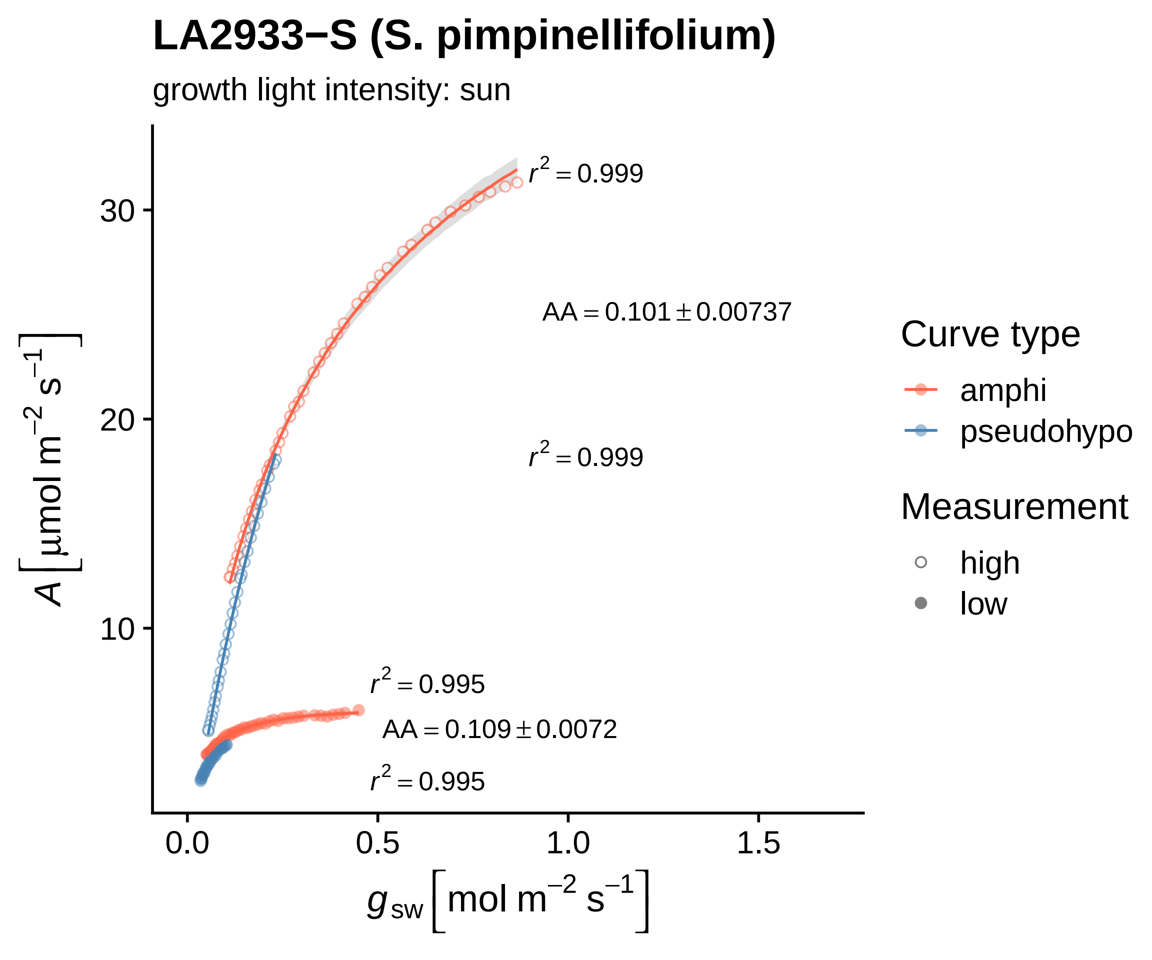
LA2933−S (S. pimpinellifolium)
growth light intensity: sun
$r^2 = 0.999$
$AA = 0.101 \pm 0.00737$
$r^2 = 0.999$
$r^2 = 0.995$
$AA = 0.109 \pm 0.0072$
$r^2 = 0.995$
30
20
10
0.0
0.5
1.0
1.5
$A\ [\mu mol\ m^{-2}\ s^{-1}]$
$g_{sw}\ [mol\ m^{-2}\ s^{-1}]$
Curve type
amphi
pseudohypo
Measurement
high
low

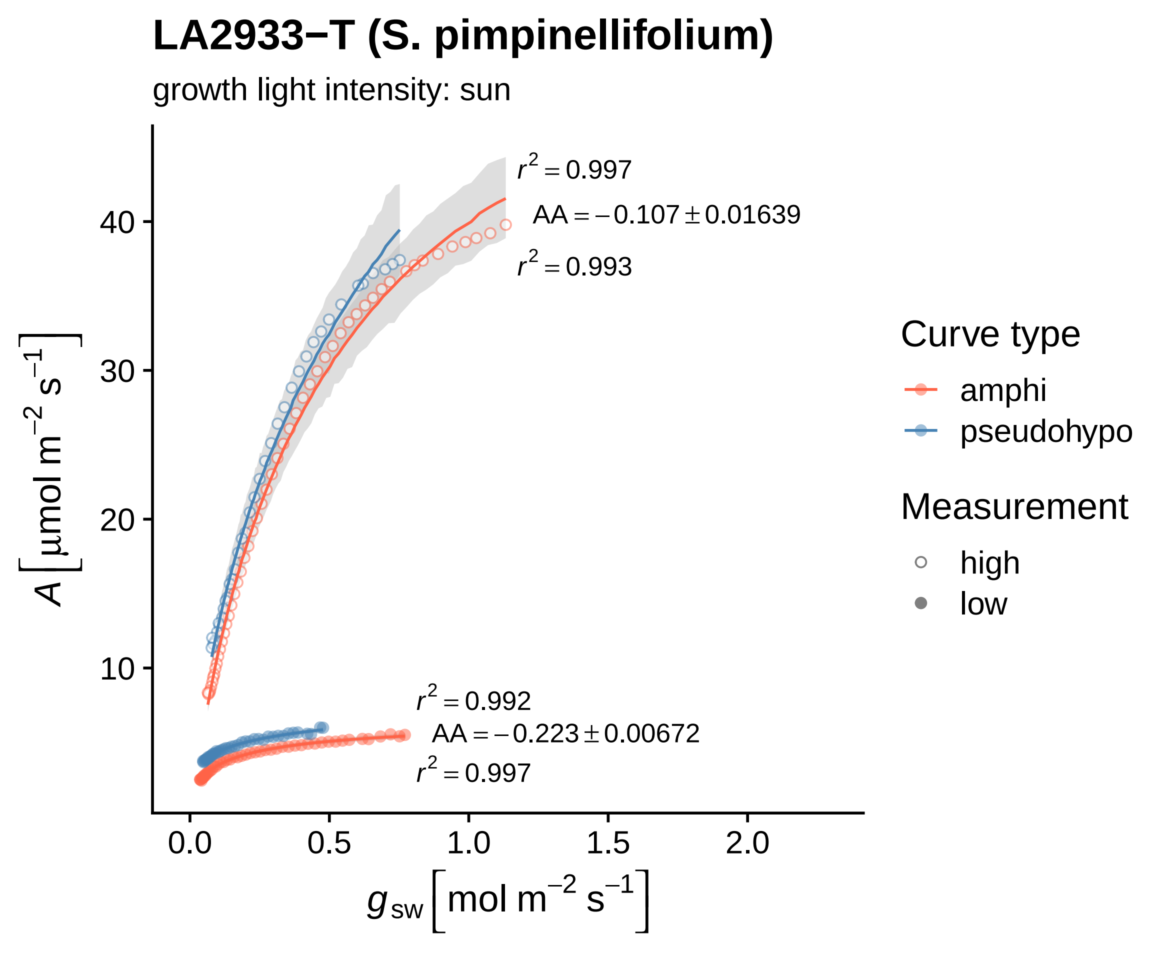

LA2933−T (S. pimpinellifolium)
growth light intensity: sun
r² = 0.997
AA = −0.107 ± 0.01639
r² = 0.993
r² = 0.992
AA = −0.223 ± 0.00672
r² = 0.997
A [µmol m⁻² s⁻¹]
g_sw [mol m⁻² s⁻¹]
10
20
30
40
0.0
0.5
1.0
1.5
2.0
Curve type
amphi
pseudohypo
Measurement
high
low

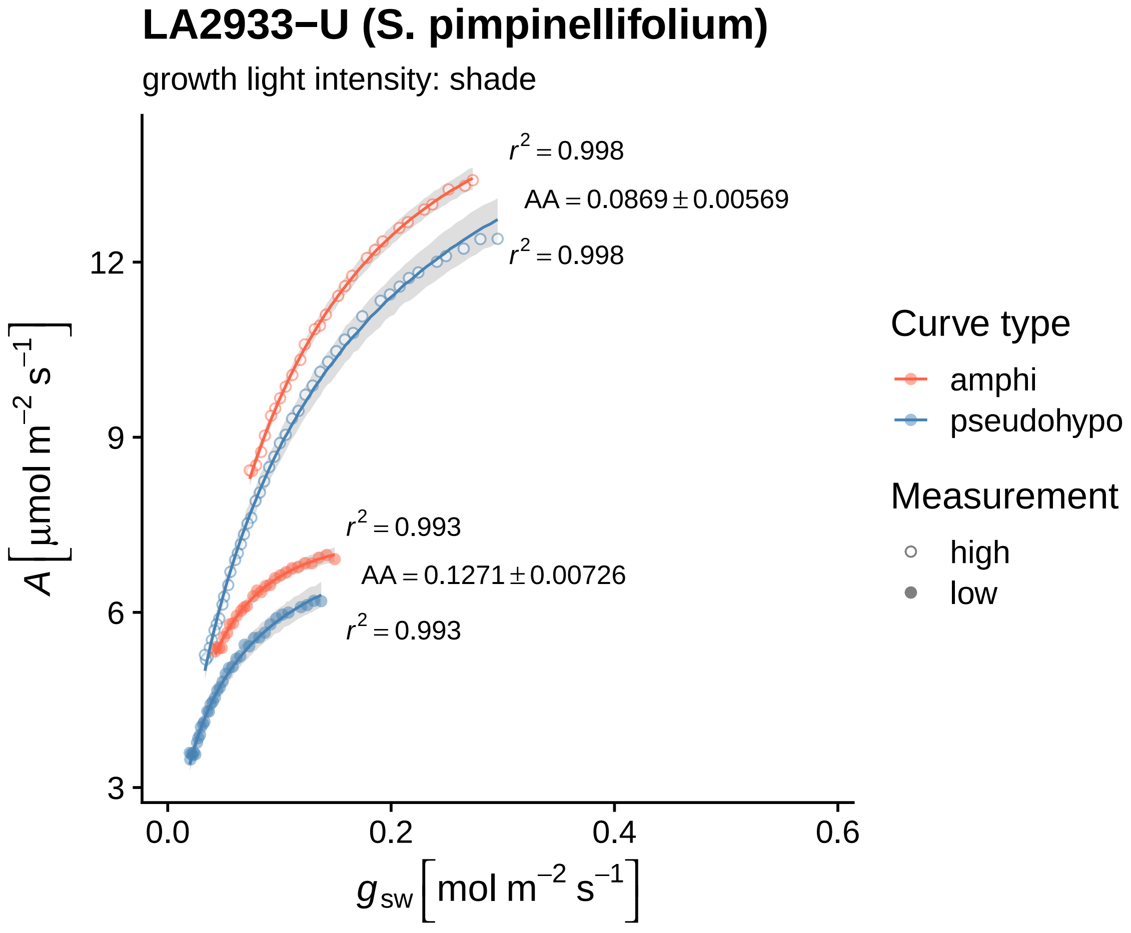

LA2933−U (S. pimpinellifolium)
growth light intensity: shade
$r^2 = 0.998$
$AA = 0.0869 \pm 0.00569$
$r^2 = 0.998$
$r^2 = 0.993$
$AA = 0.1271 \pm 0.00726$
$r^2 = 0.993$
12
9
6
3
0.0
0.2
0.4
0.6
$A\ [\mu mol\ m^{-2}\ s^{-1}]$
$g_{sw}\ [mol\ m^{-2}\ s^{-1}]$
Curve type
amphi
pseudohypo
Measurement
high
low

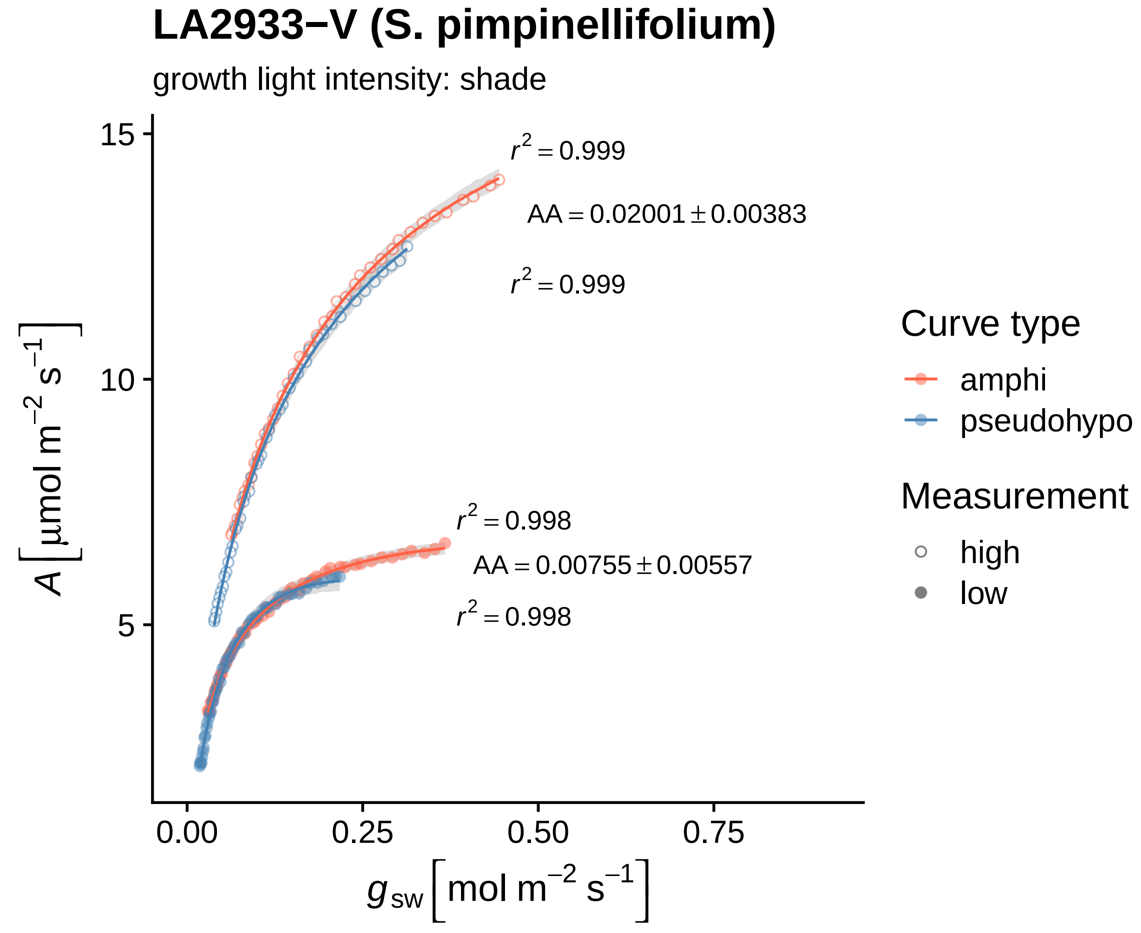

LA2933−V (S. pimpinellifolium)
growth light intensity: shade
$r^2 = 0.999$
AA = 0.02001 ± 0.00383
$r^2 = 0.999$
$r^2 = 0.998$
AA = 0.00755 ± 0.00557
$r^2 = 0.998$
15
10
5
0.00
0.25
0.50
0.75
$A\left[\mu mol\ m^{-2}\ s^{-1}\right]$
$g_{sw}\left[mol\ m^{-2}\ s^{-1}\right]$
Curve type
amphi
pseudohypo
Measurement
high
low

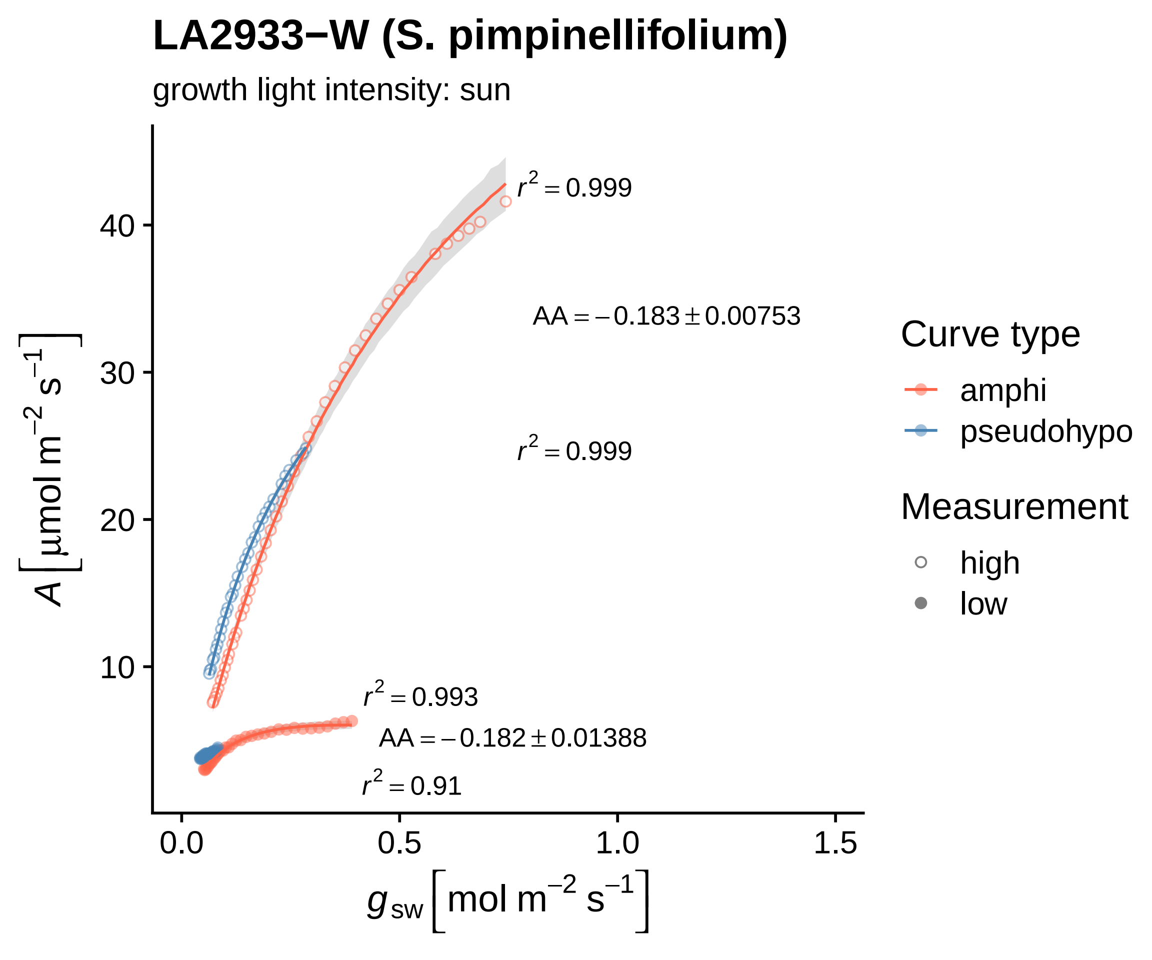

LA2933−W (S. pimpinellifolium)
growth light intensity: sun
r² = 0.999
AA = −0.183 ± 0.00753
r² = 0.999
r² = 0.993
AA = −0.182 ± 0.01388
r² = 0.91
A [µmol m⁻² s⁻¹]
g sw [mol m⁻² s⁻¹]
0.0
0.5
1.0
1.5
10
20
30
40
Curve type
amphi
pseudohypo
Measurement
high
low

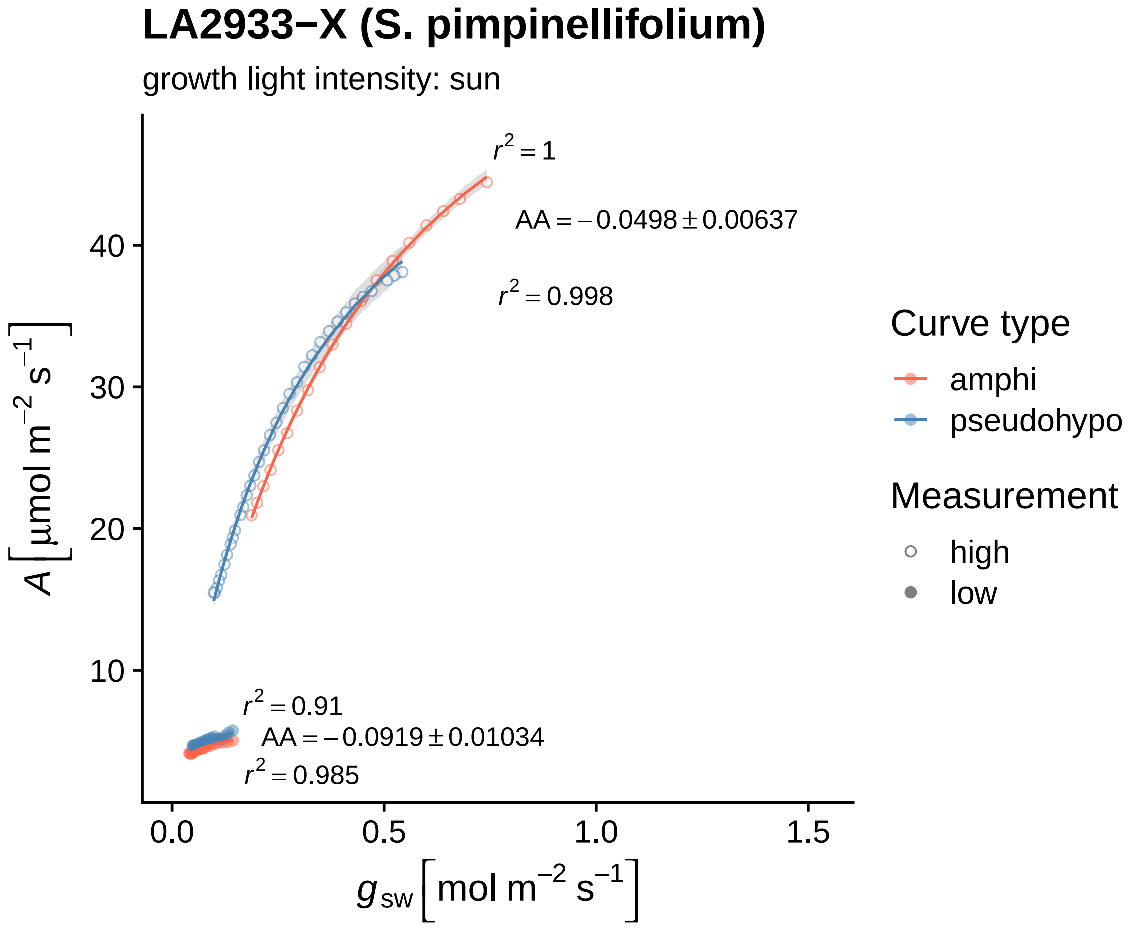
LA2933−X (S. pimpinellifolium)
growth light intensity: sun
$r^2 = 1$
$AA = -0.0498 \pm 0.00637$
$r^2 = 0.998$
$r^2 = 0.91$
$AA = -0.0919 \pm 0.01034$
$r^2 = 0.985$
10
20
30
40
0.0
0.5
1.0
1.5
$A\ [\mu mol\ m^{-2}\ s^{-1}]$
$g_{sw}\ [mol\ m^{-2}\ s^{-1}]$
Curve type
amphi
pseudohypo
Measurement
high
low

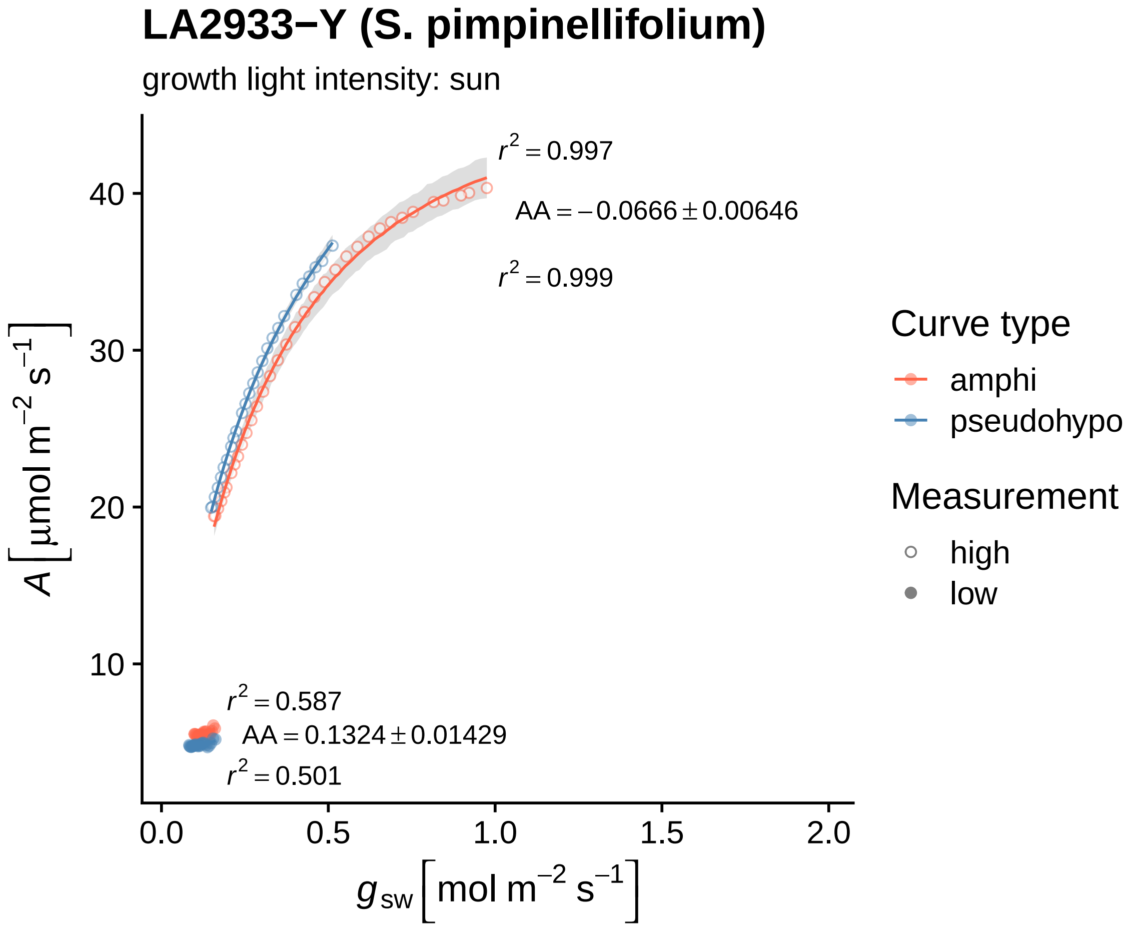
LA2933−Y (S. pimpinellifolium)
growth light intensity: sun
$r^2 = 0.997$
$AA = -0.0666 \pm 0.00646$
$r^2 = 0.999$
$r^2 = 0.587$
$AA = 0.1324 \pm 0.01429$
$r^2 = 0.501$
40
30
20
10
0.0
0.5
1.0
1.5
2.0
$A\left[\mu mol\ m^{-2}\ s^{-1}\right]$
$g_{sw}\left[mol\ m^{-2}\ s^{-1}\right]$
Curve type
amphi
pseudohypo
Measurement
high
low

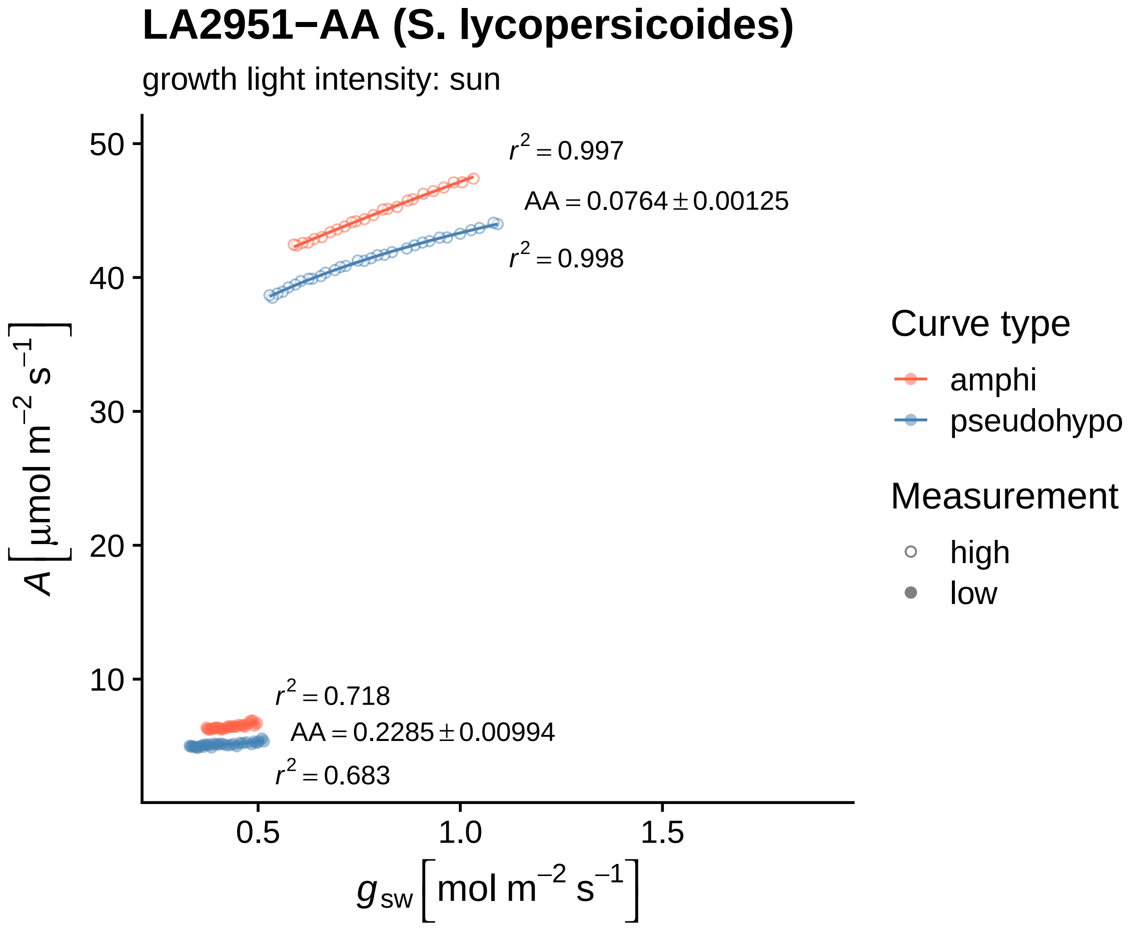
LA2951−AA (S. lycopersicoides)
growth light intensity: sun
$r^2 = 0.997$
$AA = 0.0764 \pm 0.00125$
$r^2 = 0.998$
$r^2 = 0.718$
$AA = 0.2285 \pm 0.00994$
$r^2 = 0.683$
50
40
30
20
10
0.5
1.0
1.5
$A\ [\mu mol\ m^{-2}\ s^{-1}]$
$g_{sw}\ [mol\ m^{-2}\ s^{-1}]$
Curve type
amphi
pseudohypo
Measurement
high
low

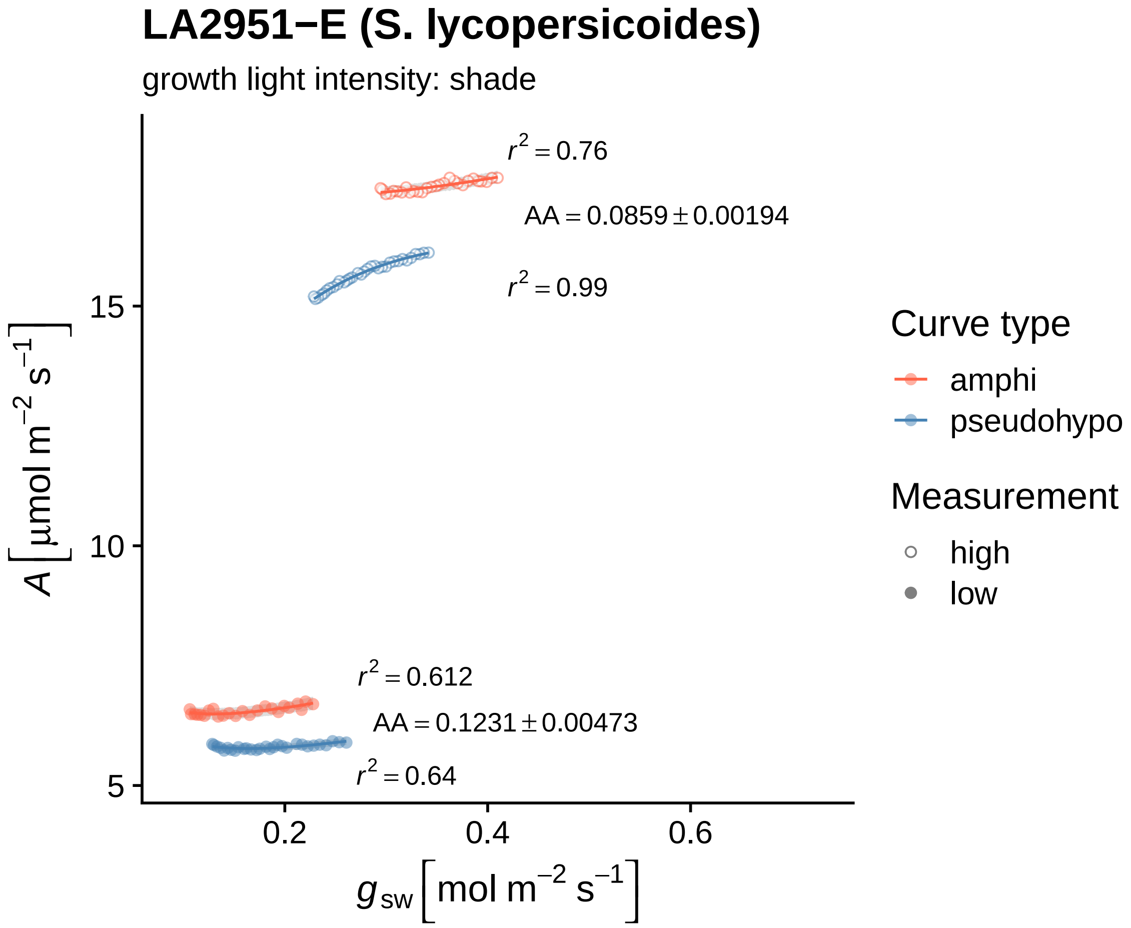

LA2951−E (S. lycopersicoides)
growth light intensity: shade
$r^2 = 0.76$
$AA = 0.0859 \pm 0.00194$
$r^2 = 0.99$
$r^2 = 0.612$
$AA = 0.1231 \pm 0.00473$
$r^2 = 0.64$
15
10
5
0.2
0.4
0.6
$A\ [\mu mol\ m^{-2}\ s^{-1}]$
$g_{sw}\ [mol\ m^{-2}\ s^{-1}]$
Curve type
amphi
pseudohypo
Measurement
high
low

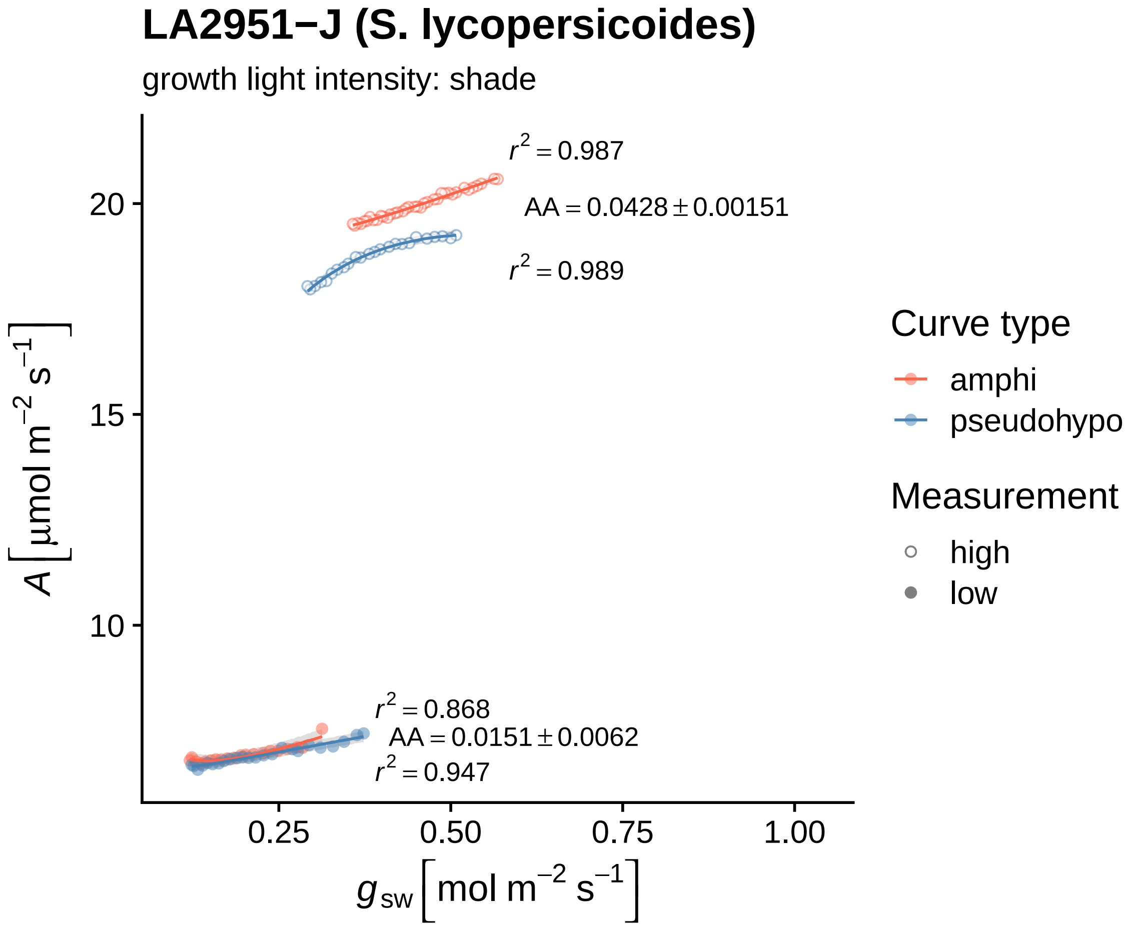

LA2951−J (S. lycopersicoides)
growth light intensity: shade
$r^2 = 0.987$
AA = 0.0428 ± 0.00151
$r^2 = 0.989$
$r^2 = 0.868$
AA = 0.0151 ± 0.0062
$r^2 = 0.947$
20
15
10
0.25
0.50
0.75
1.00
$A\ [\mu mol\ m^{-2}\ s^{-1}]$
$g_{sw}\ [mol\ m^{-2}\ s^{-1}]$
Curve type
amphi
pseudohypo
Measurement
high
low

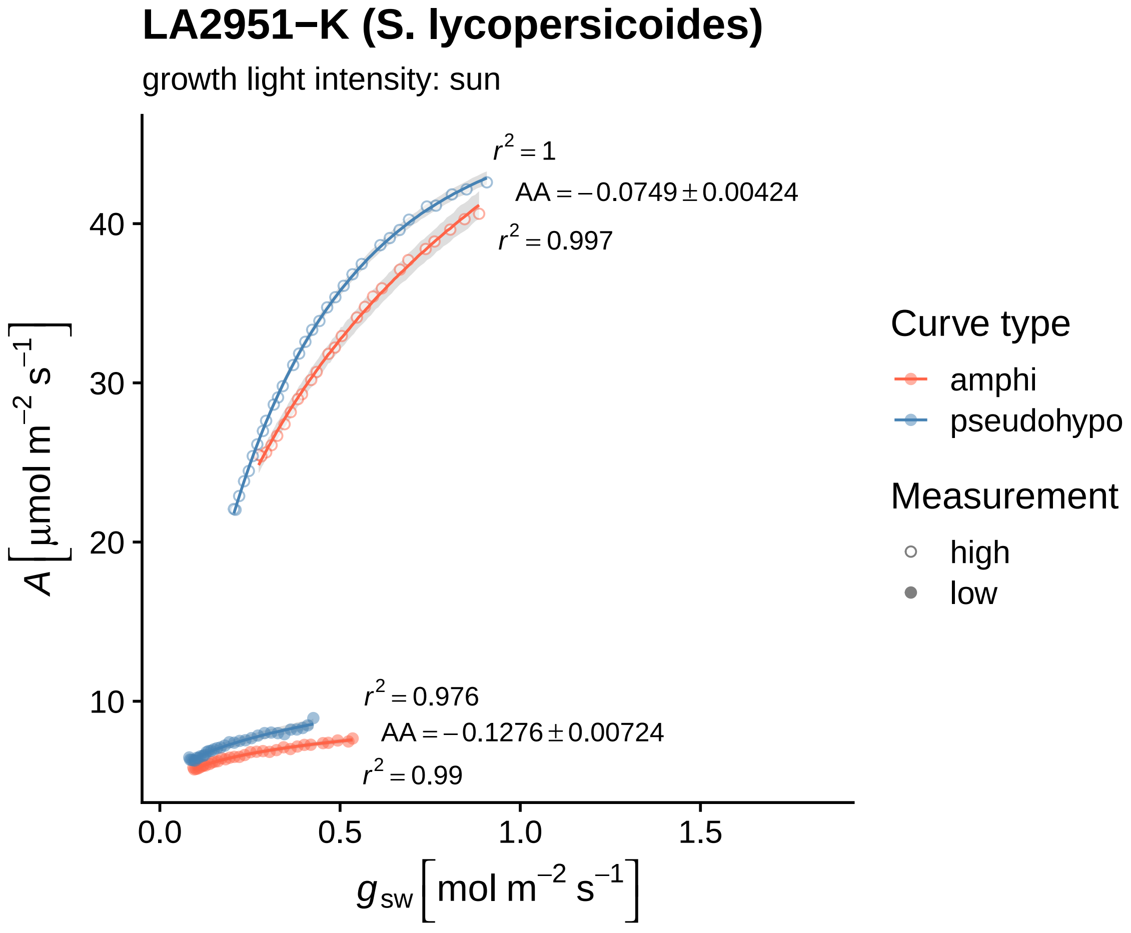

LA2951−K (S. lycopersicoides)
growth light intensity: sun
$r^2 = 1$
$AA = -0.0749 \pm 0.00424$
$r^2 = 0.997$
$r^2 = 0.976$
$AA = -0.1276 \pm 0.00724$
$r^2 = 0.99$
10
20
30
40
0.0
0.5
1.0
1.5
$A\ [\mu mol\ m^{-2}\ s^{-1}]$
$g_{sw}\ [mol\ m^{-2}\ s^{-1}]$
Curve type
amphi
pseudohypo
Measurement
high
low

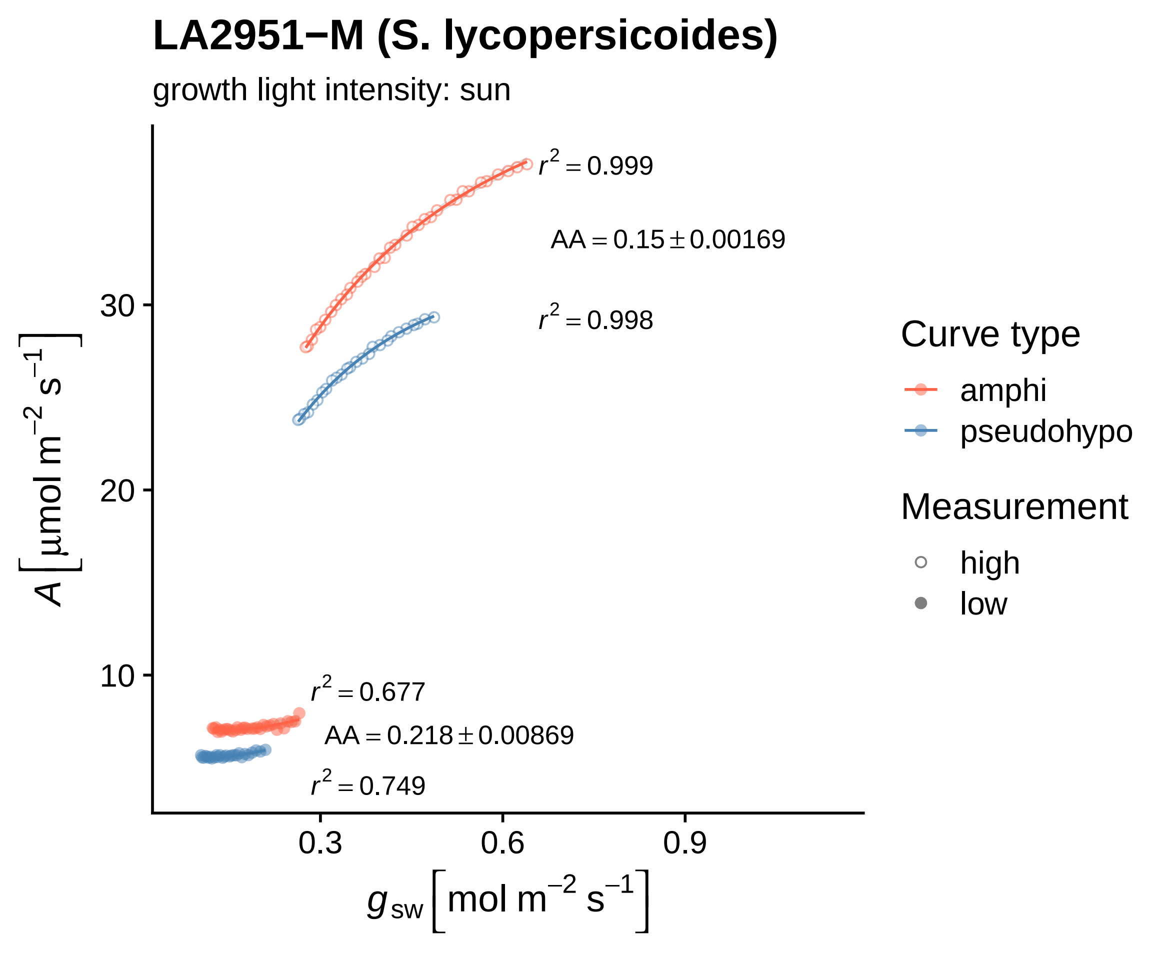
LA2951−M (S. lycopersicoides)
growth light intensity: sun
r² = 0.999
AA = 0.15 ± 0.00169
r² = 0.998
r² = 0.677
AA = 0.218 ± 0.00869
r² = 0.749
30
20
10
0.3
0.6
0.9
A [μmol m⁻² s⁻¹]
g_sw [mol m⁻² s⁻¹]
Curve type
amphi
pseudohypo
Measurement
high
low

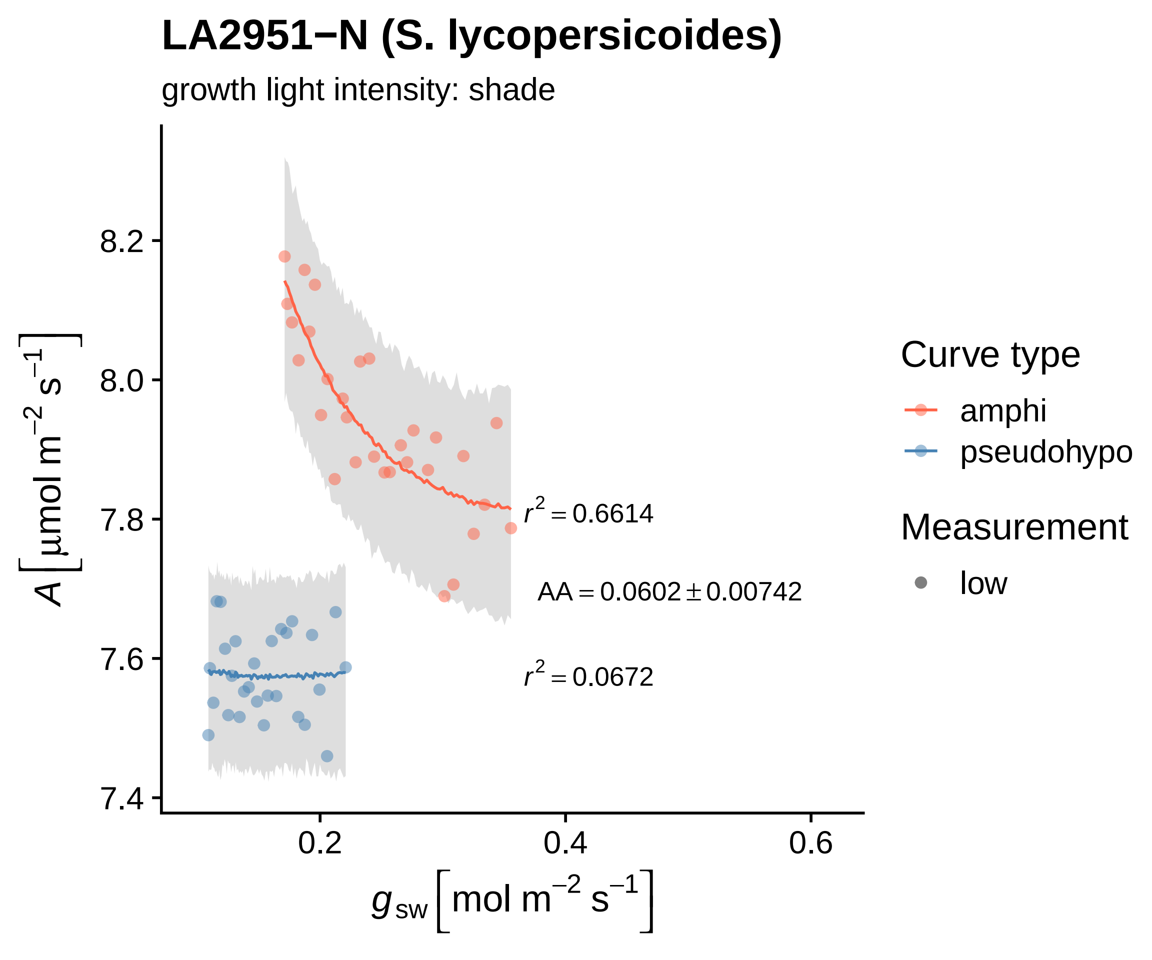
LA2951−N (S. lycopersicoides)
growth light intensity: shade
$r^2 = 0.6614$
$AA = 0.0602 \pm 0.00742$
$r^2 = 0.0672$
Curve type
amphi
pseudohypo
Measurement
low
8.2
8.0
7.8
7.6
7.4
0.2
0.4
0.6
$A\left[\mu mol\ m^{-2}\ s^{-1}\right]$
$g_{sw}\left[mol\ m^{-2}\ s^{-1}\right]$

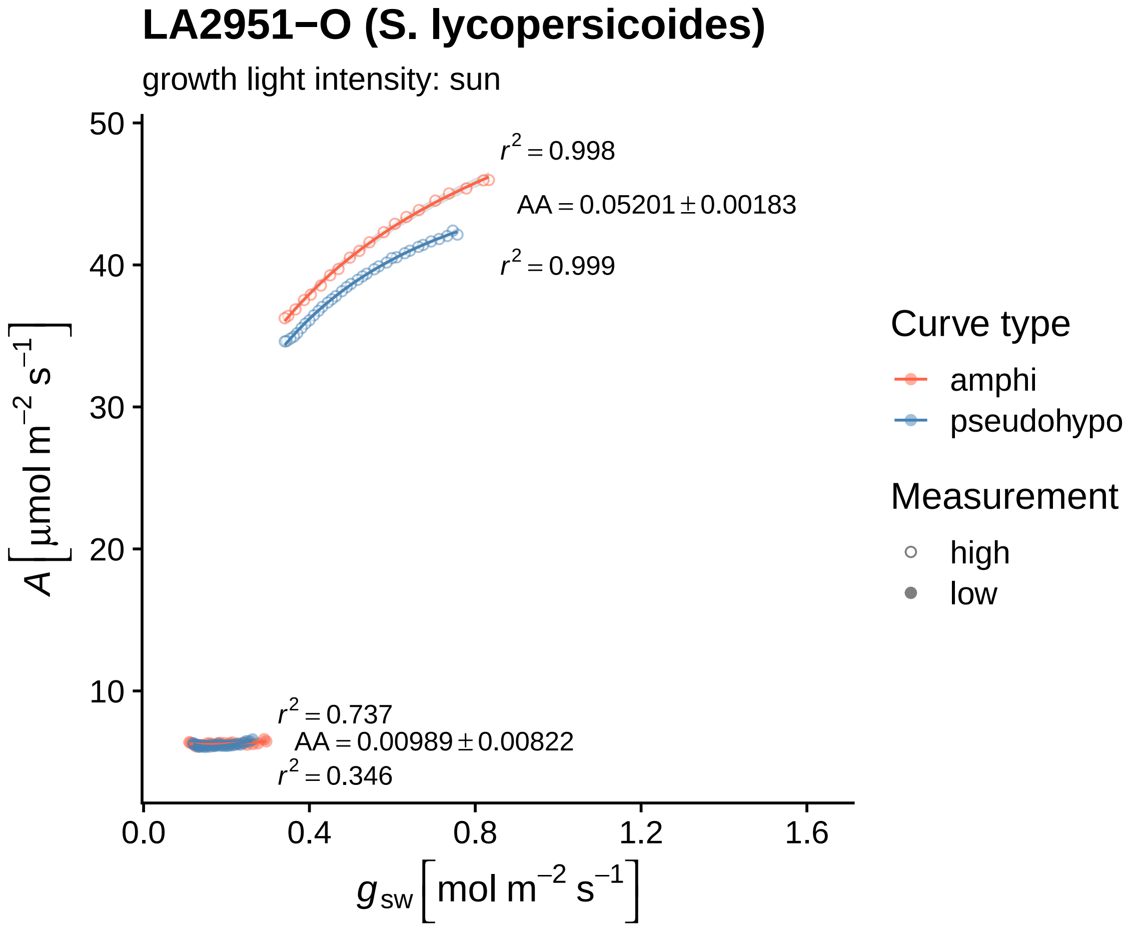

LA2951−O (S. lycopersicoides)
growth light intensity: sun
r² = 0.998
AA = 0.05201 ± 0.00183
r² = 0.999
r² = 0.737
AA = 0.00989 ± 0.00822
r² = 0.346
A [µmol m⁻² s⁻¹]
50
40
30
20
10
0.0
0.4
0.8
1.2
1.6
gsw [mol m⁻² s⁻¹]
Curve type
amphi
pseudohypo
Measurement
high
low

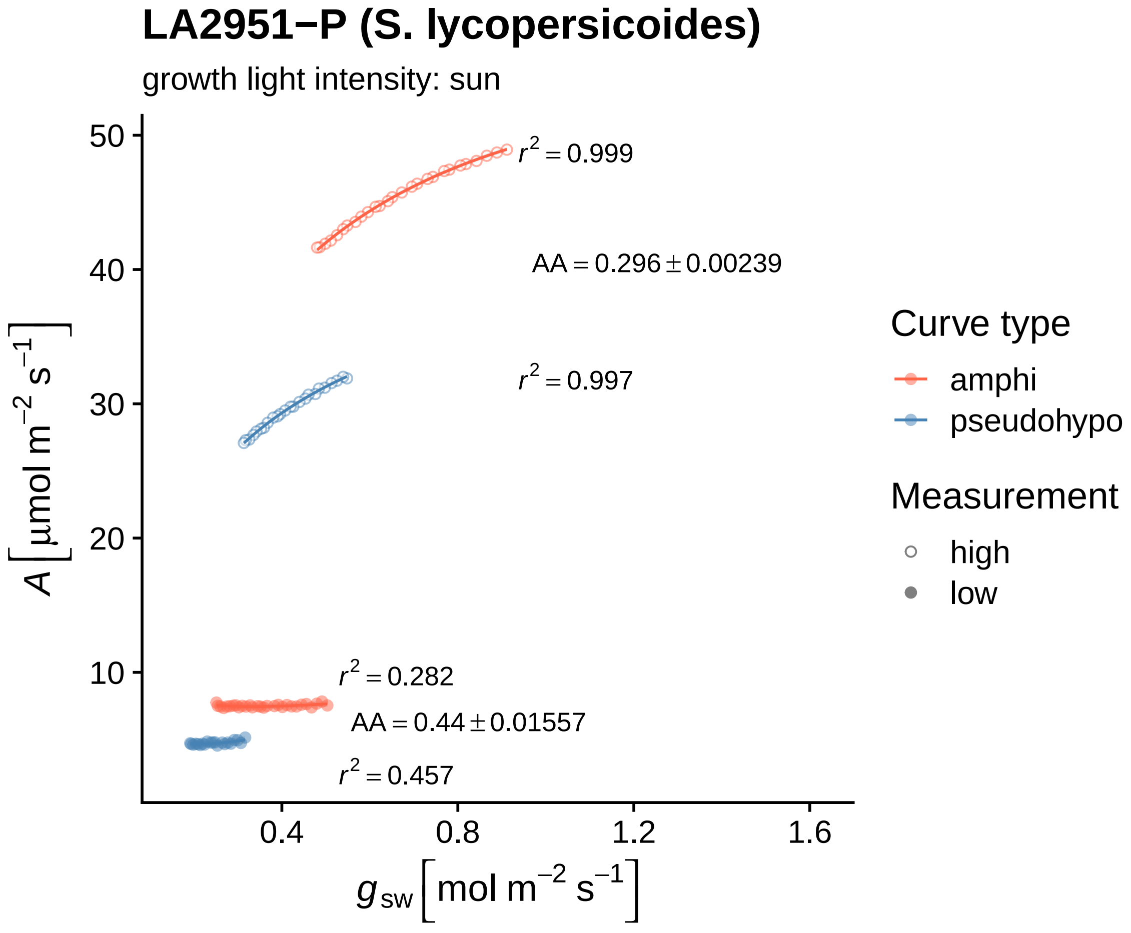

LA2951−P (S. lycopersicoides)
growth light intensity: sun
$r^2 = 0.999$
$AA = 0.296 \pm 0.00239$
$r^2 = 0.997$
$r^2 = 0.282$
$AA = 0.44 \pm 0.01557$
$r^2 = 0.457$
50
40
30
20
10
0.4
0.8
1.2
1.6
$A\ [\mu mol\ m^{-2}\ s^{-1}]$
$g_{sw}\ [mol\ m^{-2}\ s^{-1}]$
Curve type
amphi
pseudohypo
Measurement
high
low

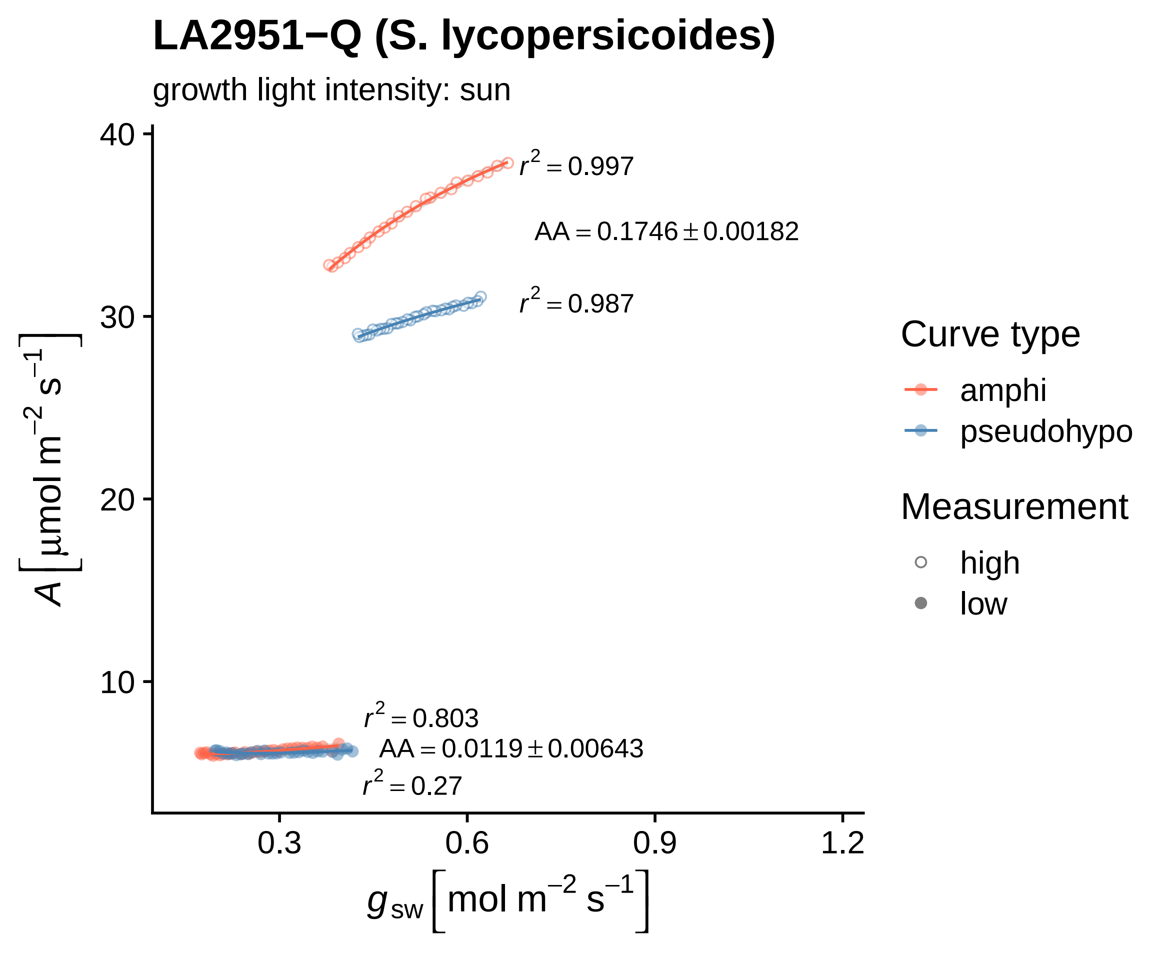

LA2951−Q (S. lycopersicoides)
growth light intensity: sun
$r^2 = 0.997$
$AA = 0.1746 \pm 0.00182$
$r^2 = 0.987$
$r^2 = 0.803$
$AA = 0.0119 \pm 0.00643$
$r^2 = 0.27$
$A\ [\mu mol\ m^{-2}\ s^{-1}]$
$g_{sw}\ [mol\ m^{-2}\ s^{-1}]$
40
30
20
10
0.3
0.6
0.9
1.2
Curve type
amphi
pseudohypo
Measurement
high
low

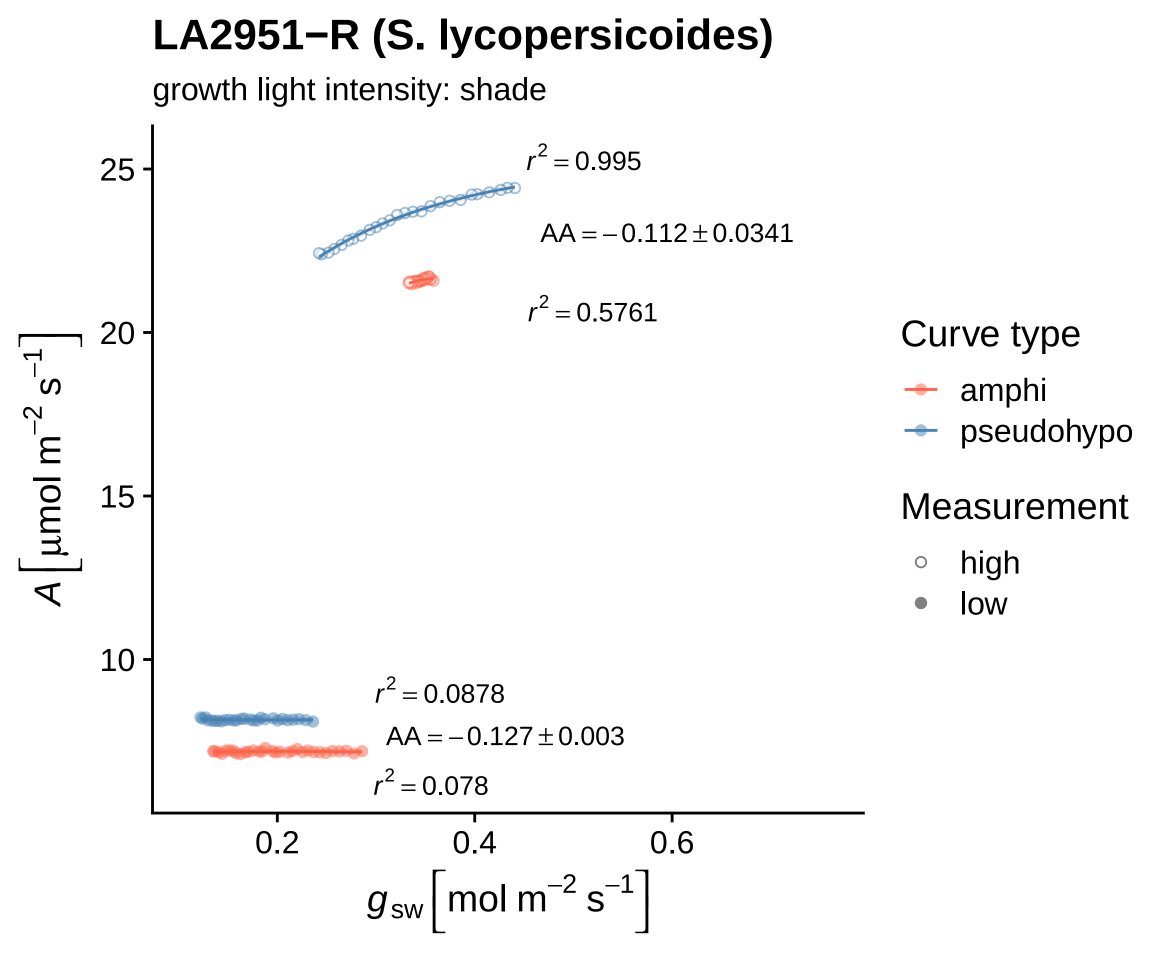

LA2951−R (S. lycopersicoides)
growth light intensity: shade
r² = 0.995
AA = −0.112 ± 0.0341
r² = 0.5761
r² = 0.0878
AA = −0.127 ± 0.003
r² = 0.078
25
20
15
10
0.2
0.4
0.6
A [µmol m⁻² s⁻¹]
g_sw [mol m⁻² s⁻¹]
Curve type
amphi
pseudohypo
Measurement
high
low

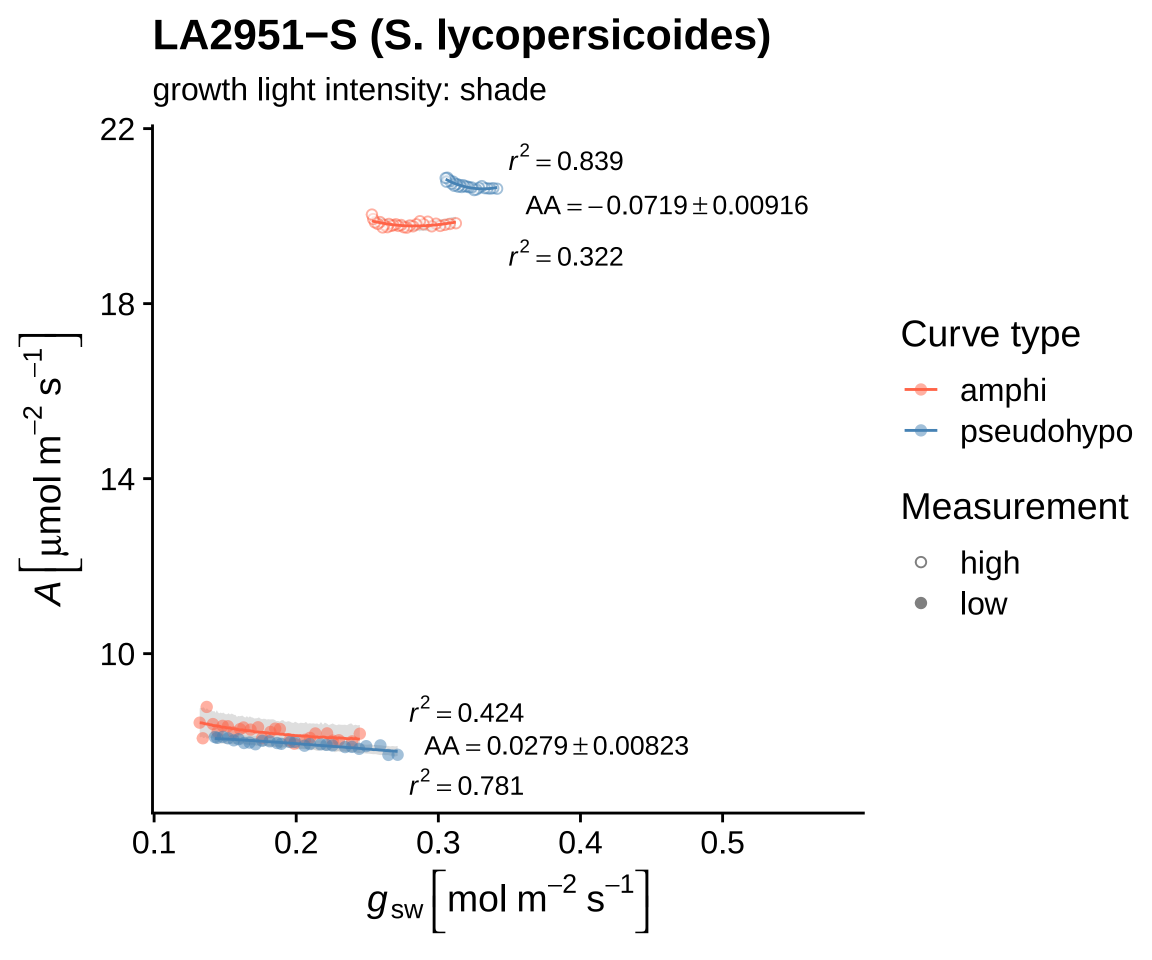

LA2951−S (S. lycopersicoides)
growth light intensity: shade
r² = 0.839
AA = −0.0719 ± 0.00916
r² = 0.322
r² = 0.424
AA = 0.0279 ± 0.00823
r² = 0.781
A [µmol m⁻² s⁻¹]
g_sw [mol m⁻² s⁻¹]
22
18
14
10
0.1
0.2
0.3
0.4
0.5
Curve type
amphi
pseudohypo
Measurement
high
low

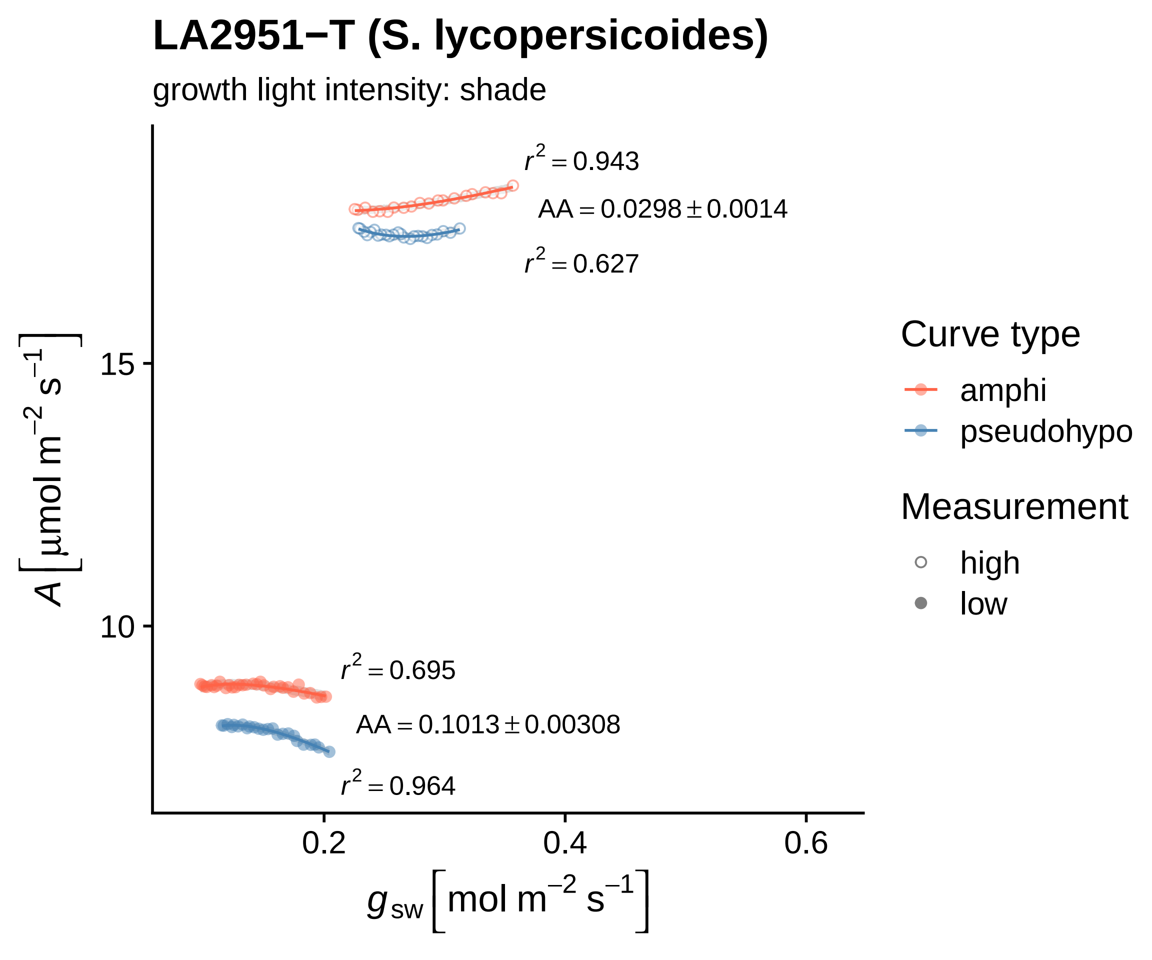

LA2951−T (S. lycopersicoides)
growth light intensity: shade
r² = 0.943
AA = 0.0298 ± 0.0014
r² = 0.627
r² = 0.695
AA = 0.1013 ± 0.00308
r² = 0.964
15
10
0.2
0.4
0.6
A [µmol m⁻² s⁻¹]
gsw [mol m⁻² s⁻¹]
Curve type
amphi
pseudohypo
Measurement
high
low

### LA2951−U (S. lycopersicoides)

growth light intensity: sun

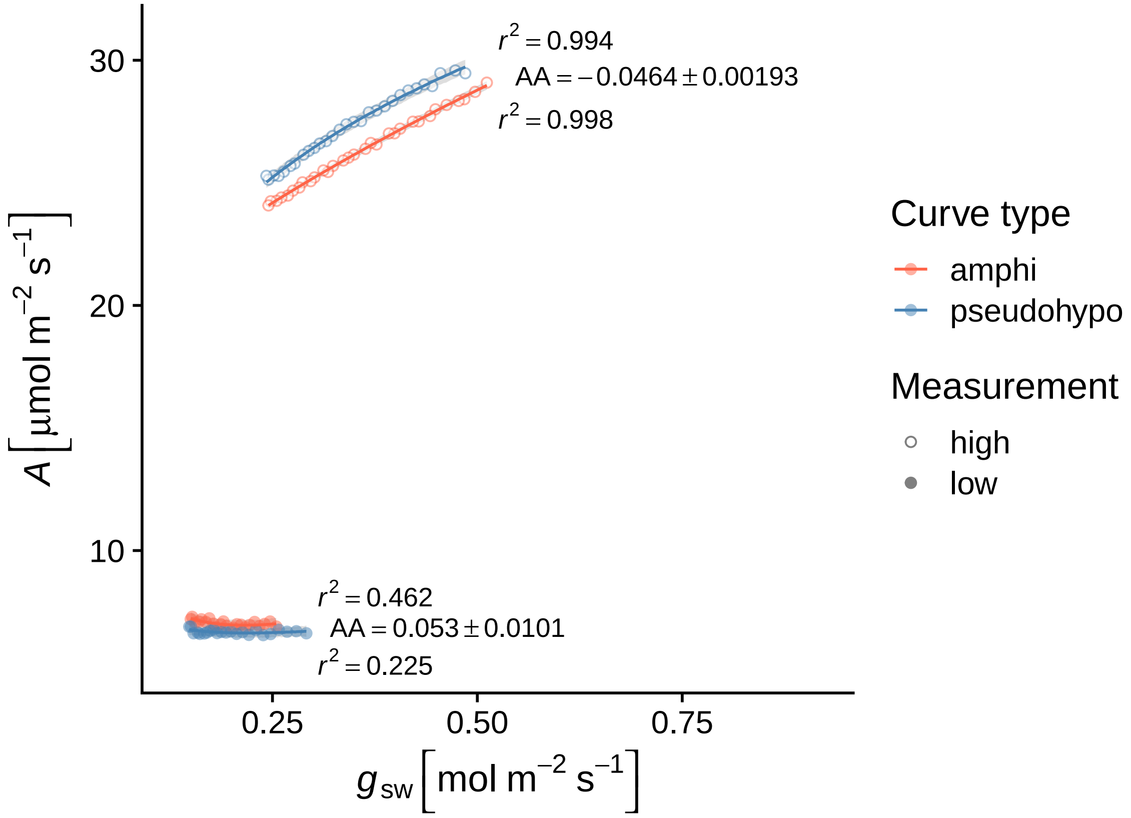

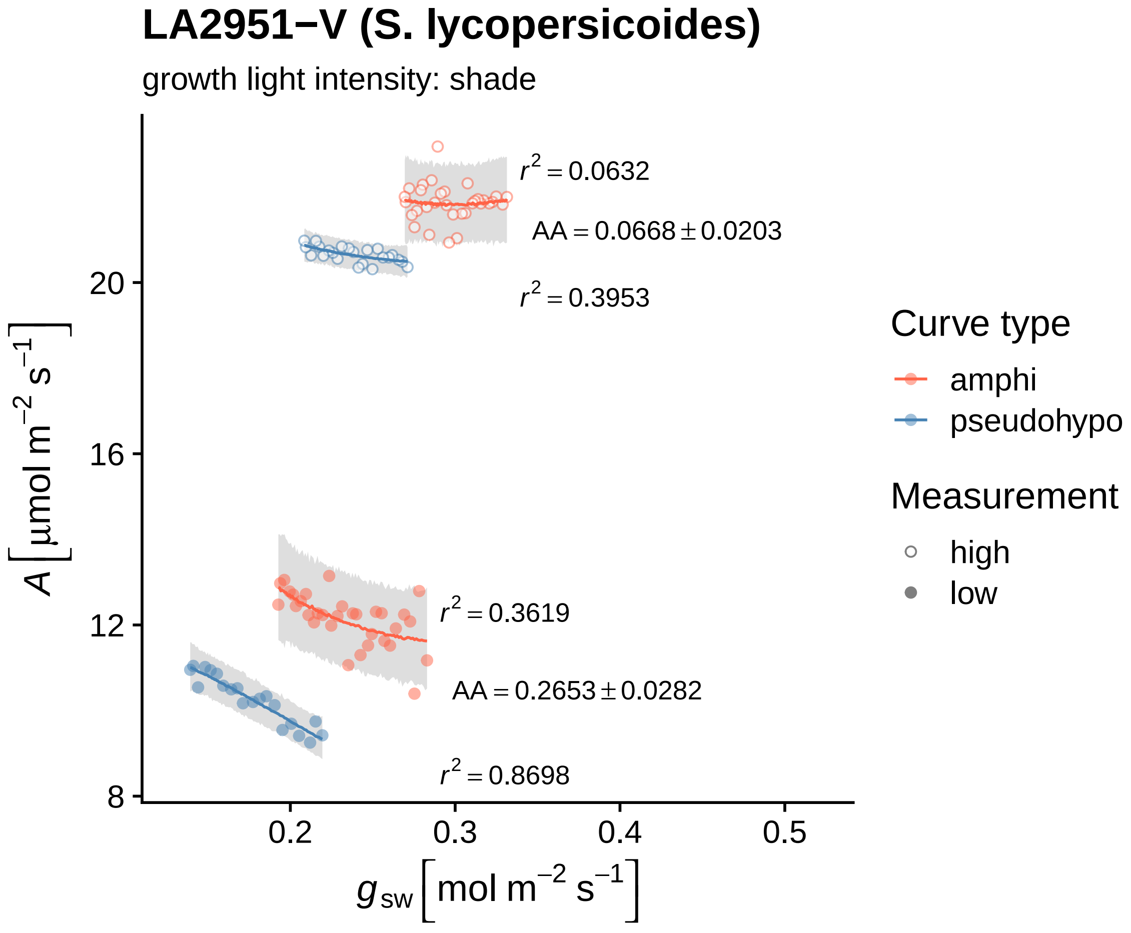
LA2951−V (S. lycopersicoides)
growth light intensity: shade
$r^2 = 0.0632$
$AA = 0.0668 \pm 0.0203$
$r^2 = 0.3953$
$r^2 = 0.3619$
$AA = 0.2653 \pm 0.0282$
$r^2 = 0.8698$
20
16
12
8
0.2
0.3
0.4
0.5
$A\ [\mu mol\ m^{-2}\ s^{-1}]$
$g_{sw}\ [mol\ m^{-2}\ s^{-1}]$
Curve type
amphi
pseudohypo
Measurement
high
low

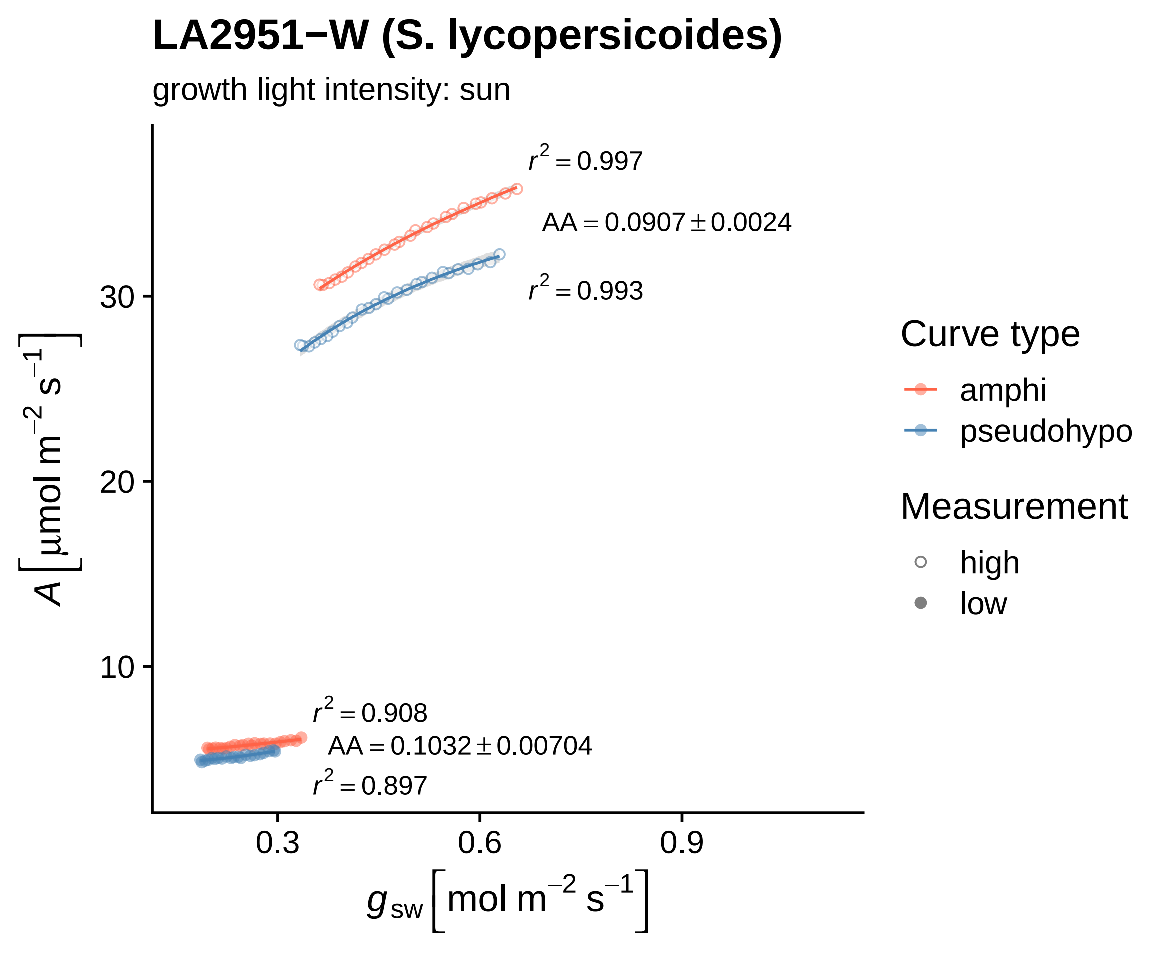

LA2951−W (S. lycopersicoides)
growth light intensity: sun
r² = 0.997
AA = 0.0907 ± 0.0024
r² = 0.993
r² = 0.908
AA = 0.1032 ± 0.00704
r² = 0.897
30
20
10
0.3
0.6
0.9
A [µmol m⁻² s⁻¹]
g_sw [mol m⁻² s⁻¹]
Curve type
amphi
pseudohypo
Measurement
high
low

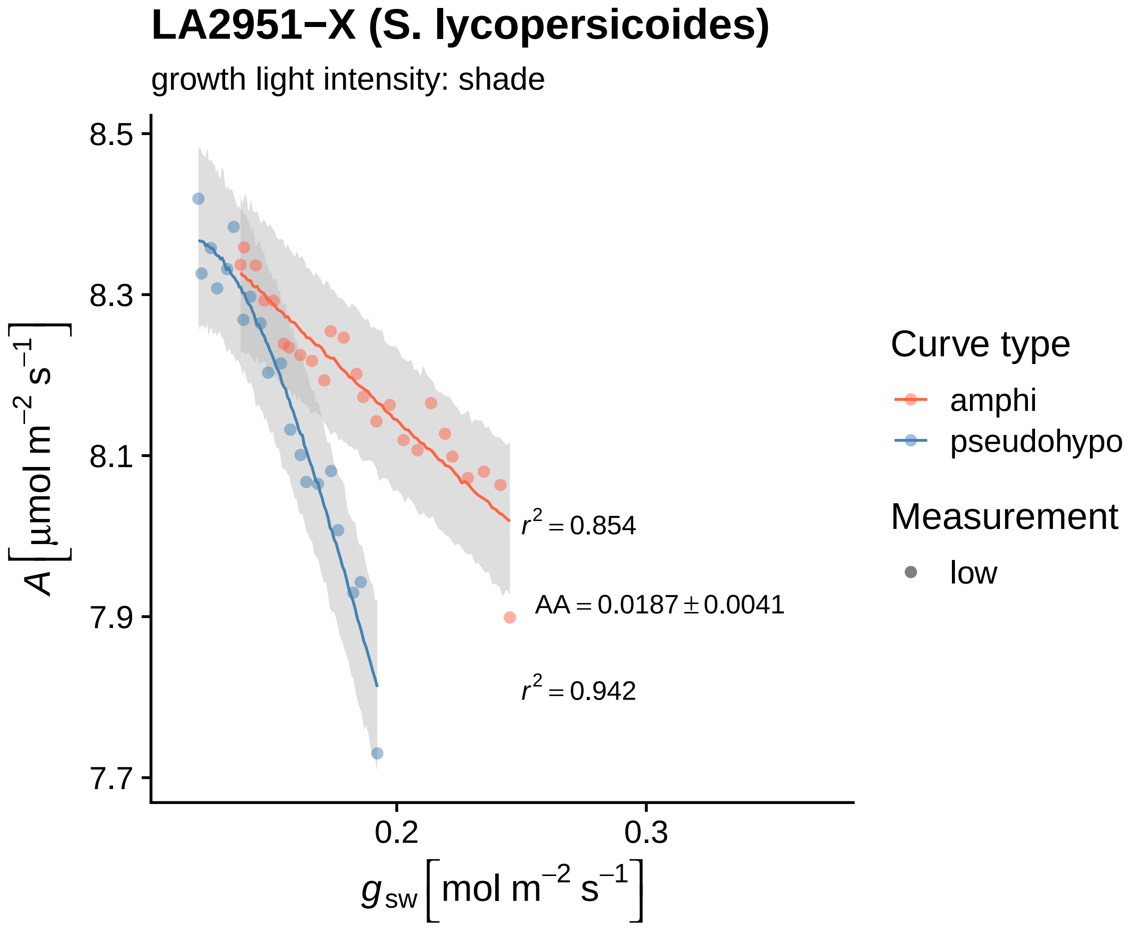
LA2951−X (S. lycopersicoides)
growth light intensity: shade
8.5
8.3
8.1
7.9
7.7
A [μmol m−2 s−1]
0.2
0.3
gsw [mol m−2 s−1]
r2 = 0.854
AA = 0.0187 ± 0.0041
r2 = 0.942
Curve type
amphi
pseudohypo
Measurement
low

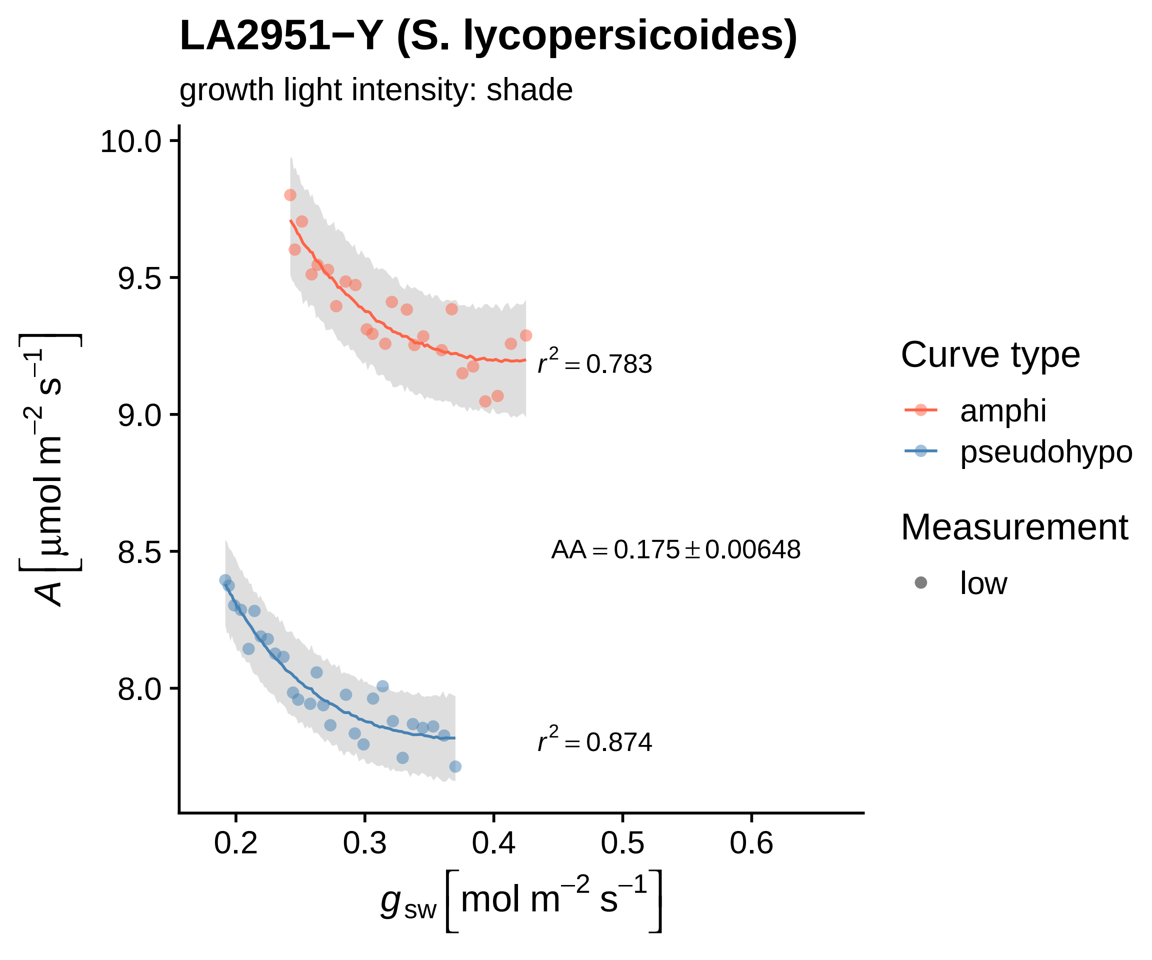
LA2951−Y (S. lycopersicoides)
growth light intensity: shade
10.0
9.5
9.0
8.5
8.0
0.2
0.3
0.4
0.5
0.6
A [μmol m−2 s−1]
gsw [mol m−2 s−1]
r2 = 0.783
AA = 0.175 ± 0.00648
r2 = 0.874
Curve type
amphi
pseudohypo
Measurement
low

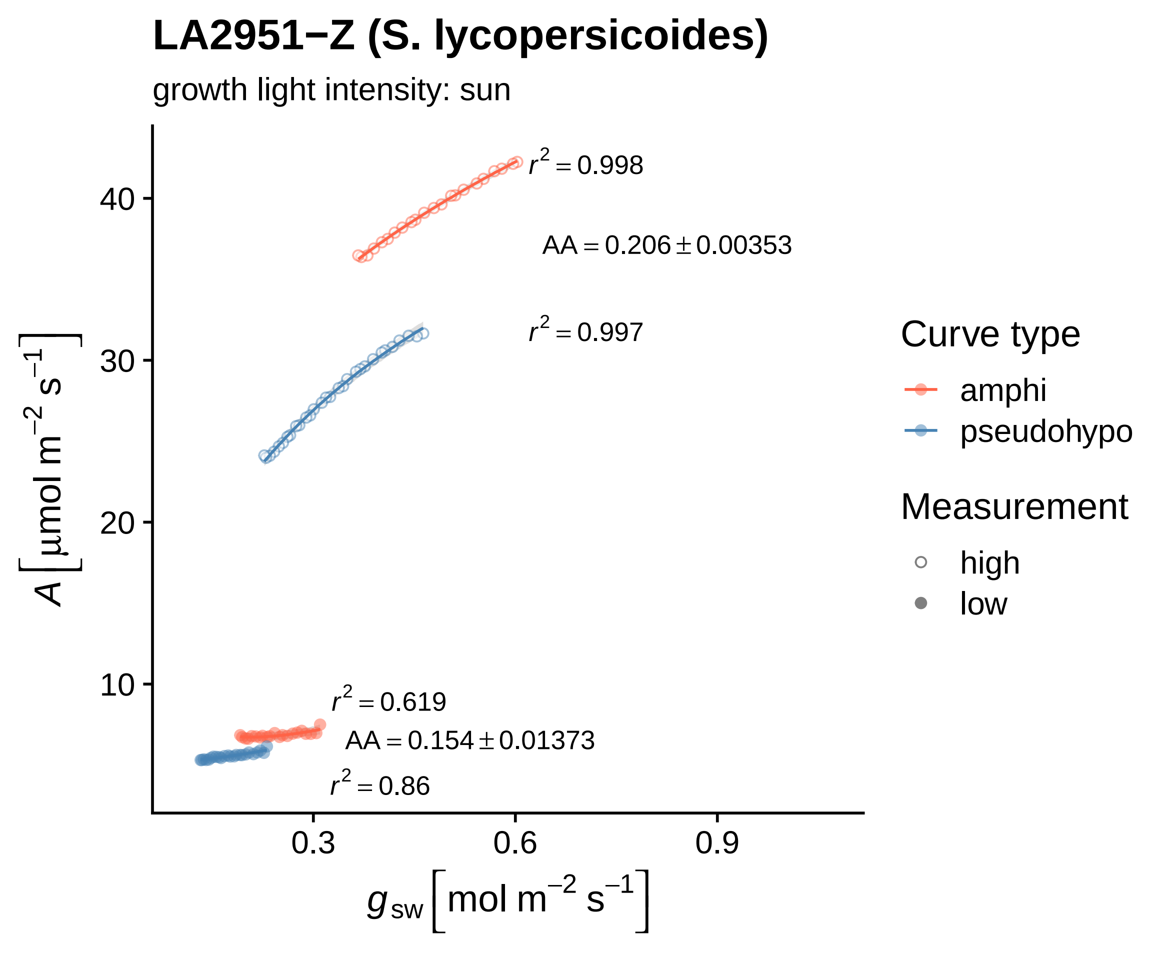

LA2951−Z (S. lycopersicoides)
growth light intensity: sun
r² = 0.998
AA = 0.206 ± 0.00353
r² = 0.997
r² = 0.619
AA = 0.154 ± 0.01373
r² = 0.86
40
30
20
10
0.3
0.6
0.9
A [µmol m⁻² s⁻¹]
gsw [mol m⁻² s⁻¹]
Curve type
amphi
pseudohypo
Measurement
high
low

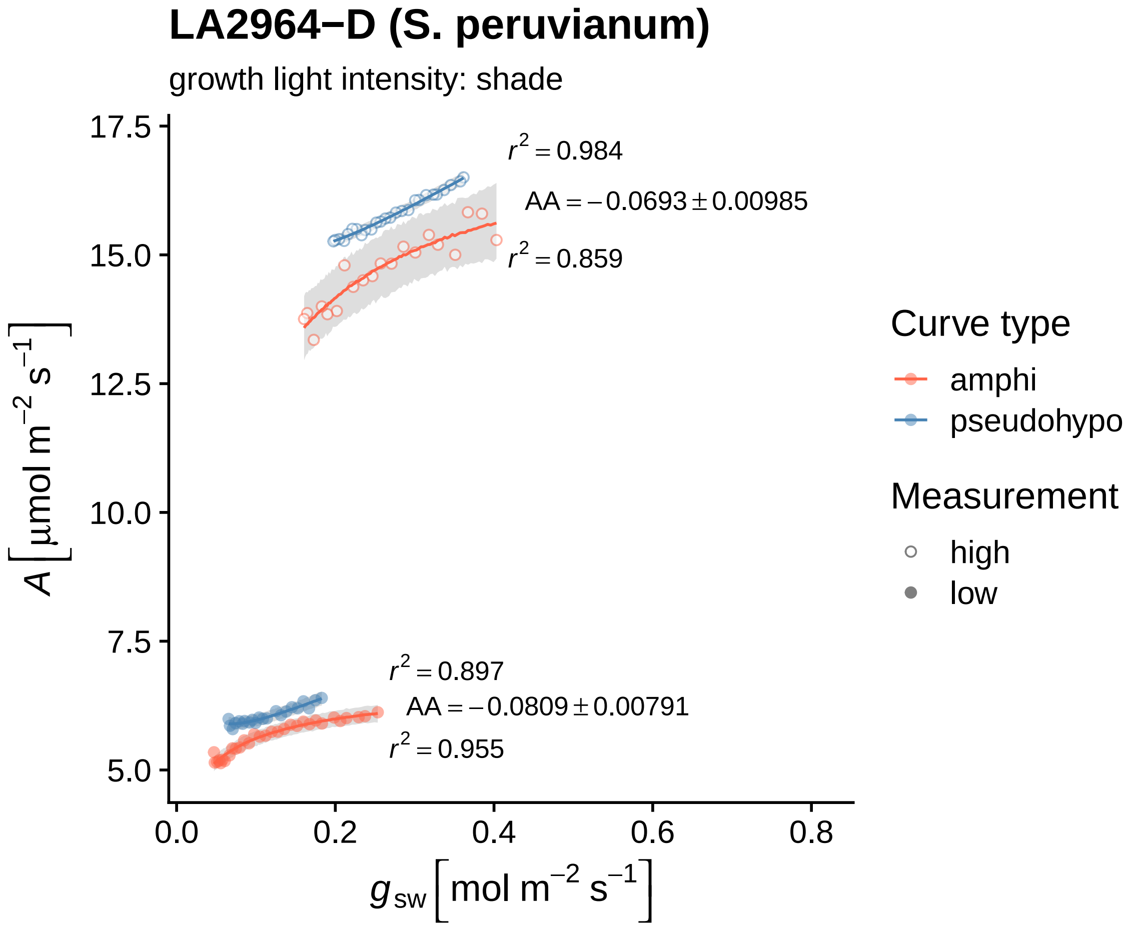

LA2964−D (S. peruvianum)
growth light intensity: shade
$r^2 = 0.984$
$AA = -0.0693 \pm 0.00985$
$r^2 = 0.859$
$r^2 = 0.897$
$AA = -0.0809 \pm 0.00791$
$r^2 = 0.955$
17.5
15.0
12.5
10.0
7.5
5.0
0.0
0.2
0.4
0.6
0.8
$A\ [\mu mol\ m^{-2}\ s^{-1}]$
$g_{sw}\ [mol\ m^{-2}\ s^{-1}]$
Curve type
amphi
pseudohypo
Measurement
high
low

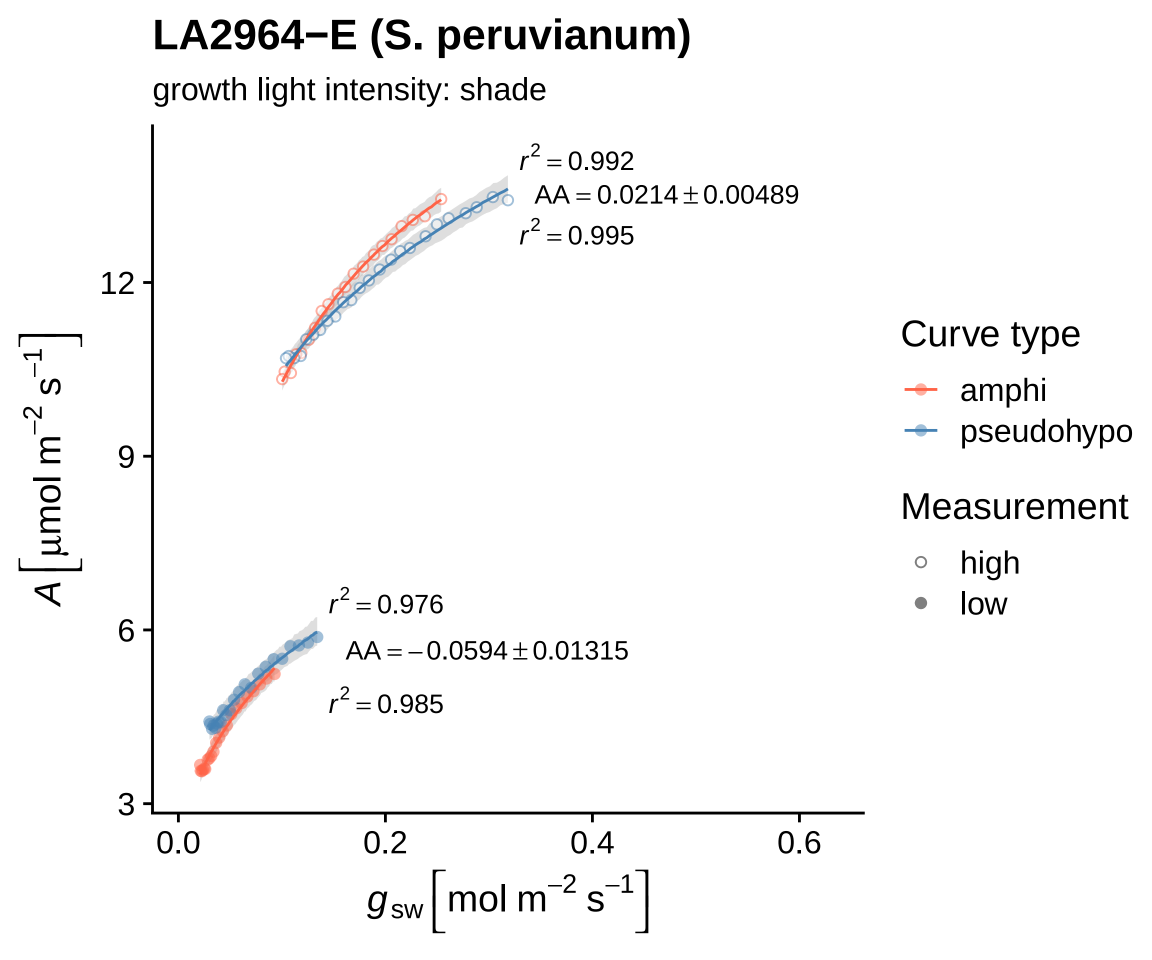

LA2964−E (S. peruvianum)
growth light intensity: shade
$r^2 = 0.992$
$AA = 0.0214 \pm 0.00489$
$r^2 = 0.995$
$r^2 = 0.976$
$AA = -0.0594 \pm 0.01315$
$r^2 = 0.985$
12
9
6
3
0.0
0.2
0.4
0.6
$A\ [\mu mol\ m^{-2}\ s^{-1}]$
$g_{sw}\ [mol\ m^{-2}\ s^{-1}]$
Curve type
amphi
pseudohypo
Measurement
high
low

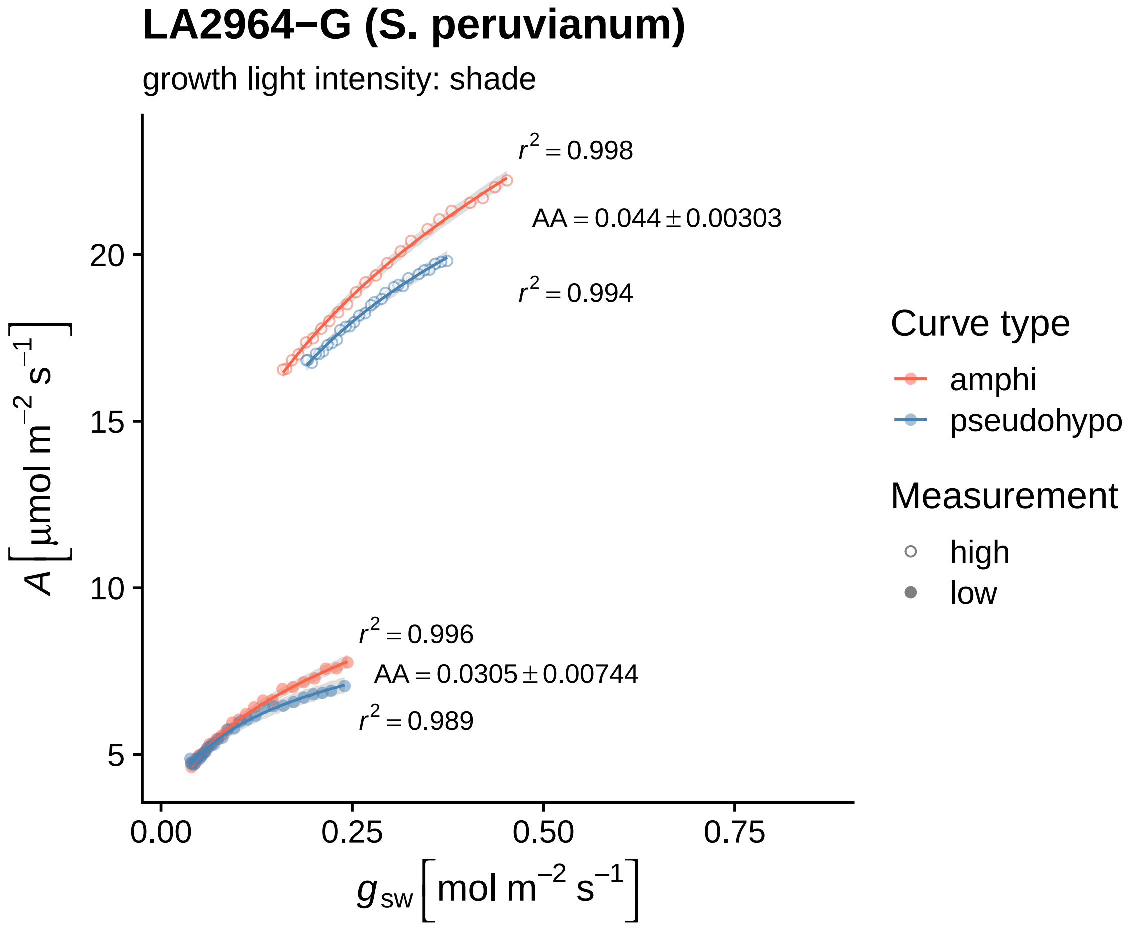

LA2964−G (S. peruvianum)
growth light intensity: shade
$r^2 = 0.998$
AA = 0.044 ± 0.00303
$r^2 = 0.994$
$r^2 = 0.996$
AA = 0.0305 ± 0.00744
$r^2 = 0.989$
20
15
10
5
0.00
0.25
0.50
0.75
$A\ [\mu mol\ m^{-2}\ s^{-1}]$
$g_{sw}\ [mol\ m^{-2}\ s^{-1}]$
Curve type
amphi
pseudohypo
Measurement
high
low

### LA2964−H (S. peruvianum)

growth light intensity: shade

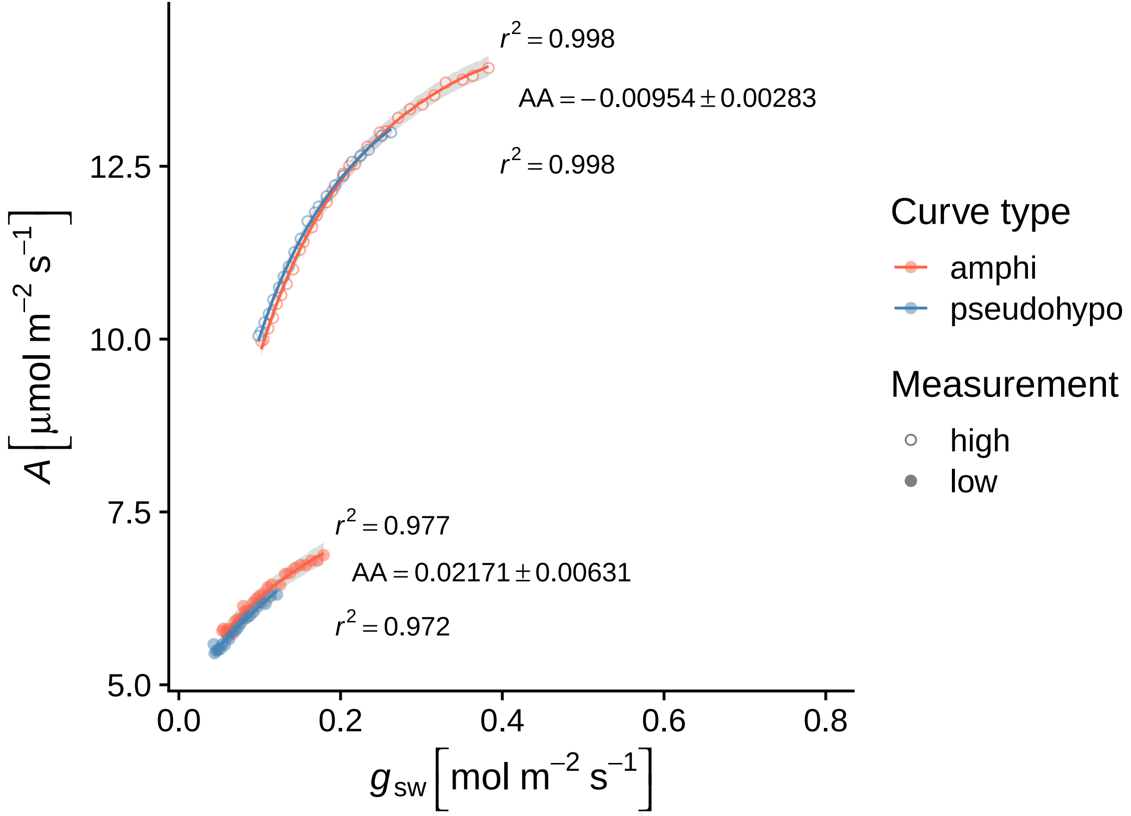

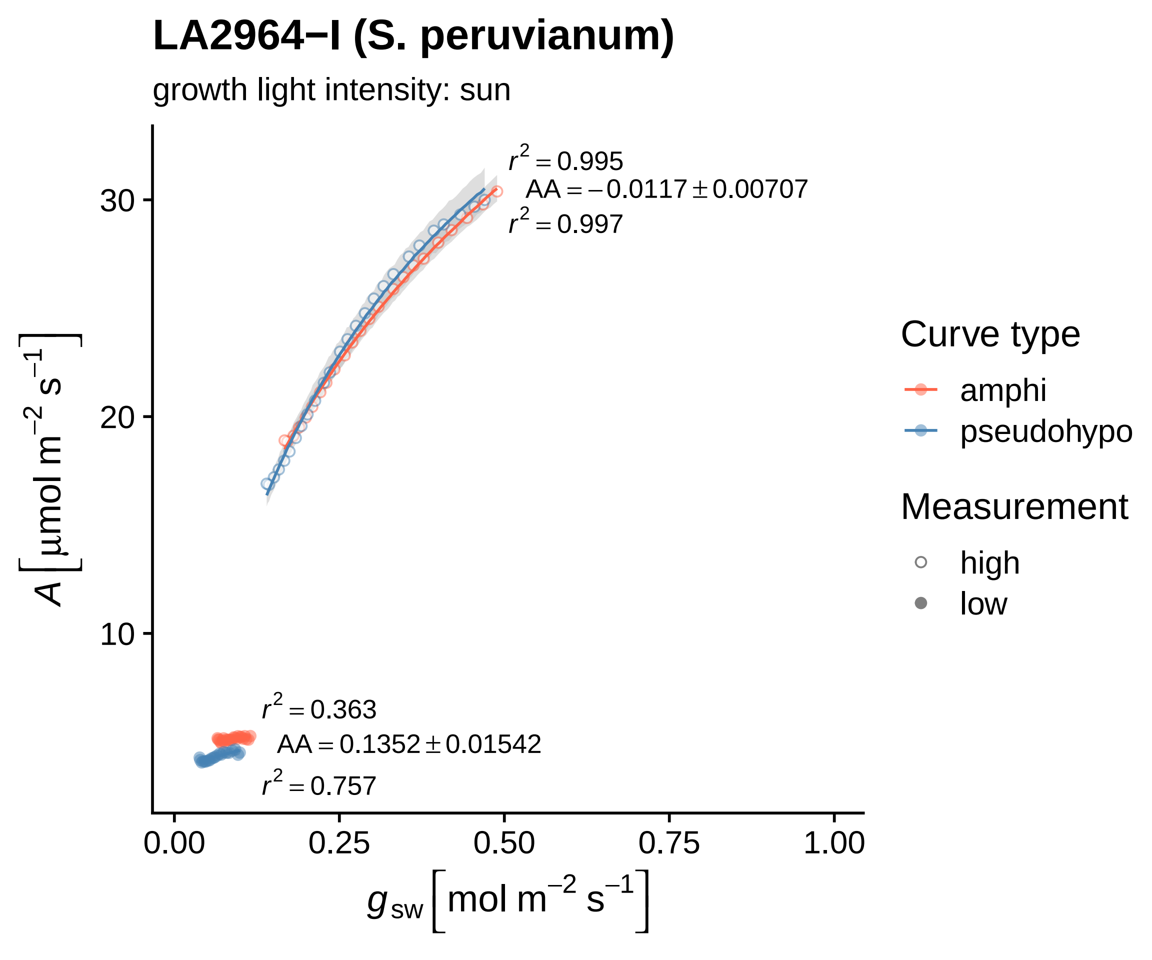

LA2964−I (S. peruvianum)
growth light intensity: sun
$r^2 = 0.995$
$AA = -0.0117 \pm 0.00707$
$r^2 = 0.997$
$r^2 = 0.363$
$AA = 0.1352 \pm 0.01542$
$r^2 = 0.757$
30
20
10
0.00
0.25
0.50
0.75
1.00
$A\,[\mu mol\ m^{-2}\ s^{-1}]$
$g_{sw}\,[mol\ m^{-2}\ s^{-1}]$
Curve type
amphi
pseudohypo
Measurement
high
low

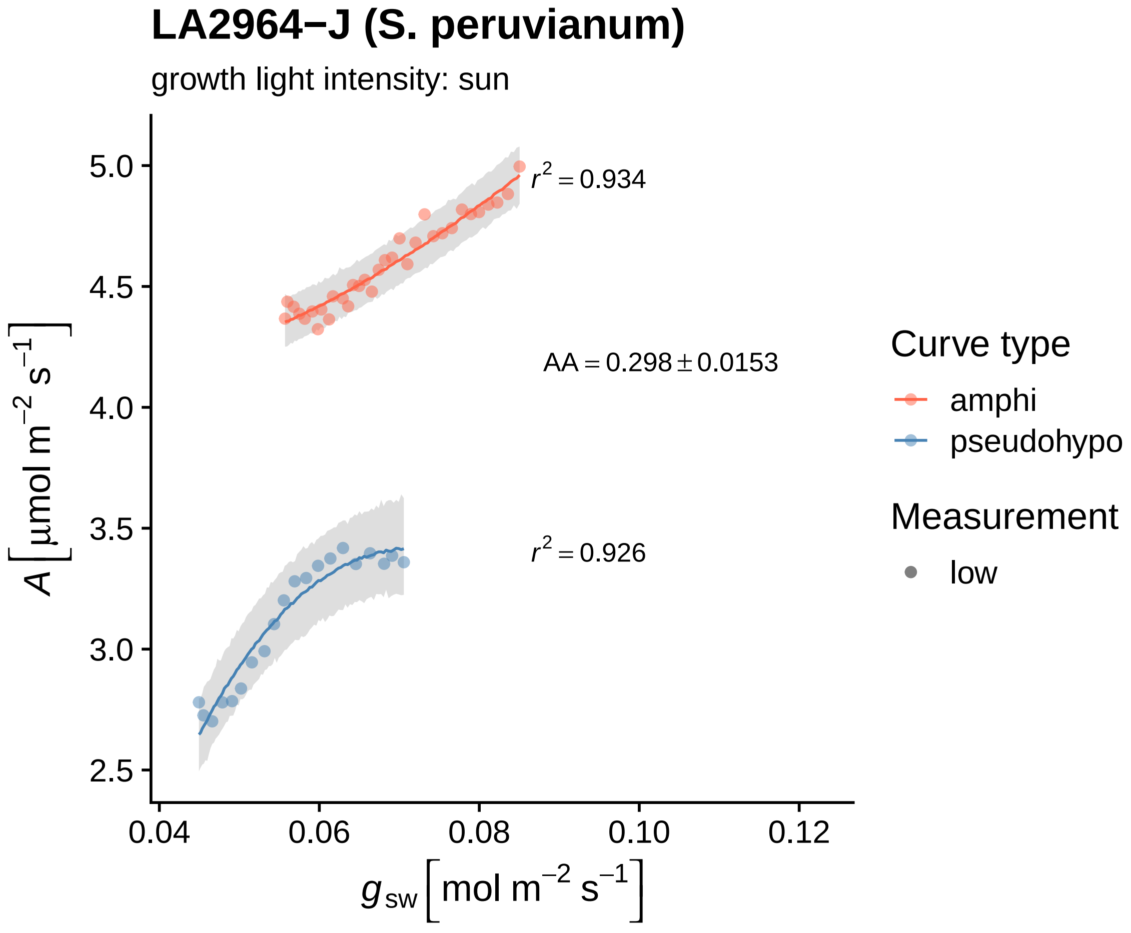

LA2964−J (S. peruvianum)
growth light intensity: sun
$r^2 = 0.934$
AA = 0.298 ± 0.0153
$r^2 = 0.926$
5.0
4.5
4.0
3.5
3.0
2.5
0.04
0.06
0.08
0.10
0.12
$A\left[\mu mol\ m^{-2}\ s^{-1}\right]$
$g_{sw}\left[mol\ m^{-2}\ s^{-1}\right]$
Curve type
amphi
pseudohypo
Measurement
low

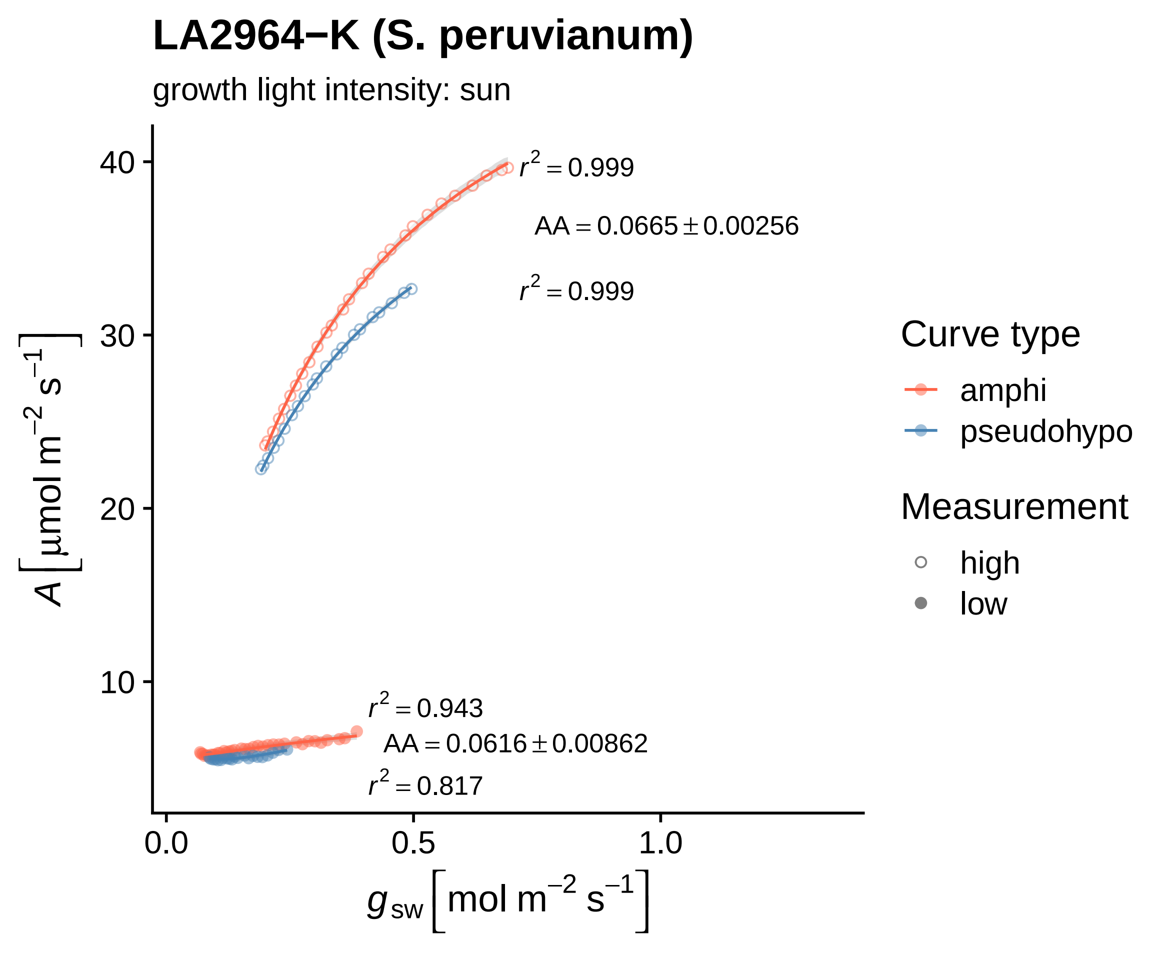

LA2964−K (S. peruvianum)
growth light intensity: sun
$r^2 = 0.999$
$AA = 0.0665 \pm 0.00256$
$r^2 = 0.999$
$r^2 = 0.943$
$AA = 0.0616 \pm 0.00862$
$r^2 = 0.817$
40
30
20
10
0.0
0.5
1.0
$A\ [\mu mol\ m^{-2}\ s^{-1}]$
$g_{sw}\ [mol\ m^{-2}\ s^{-1}]$
Curve type
amphi
pseudohypo
Measurement
high
low

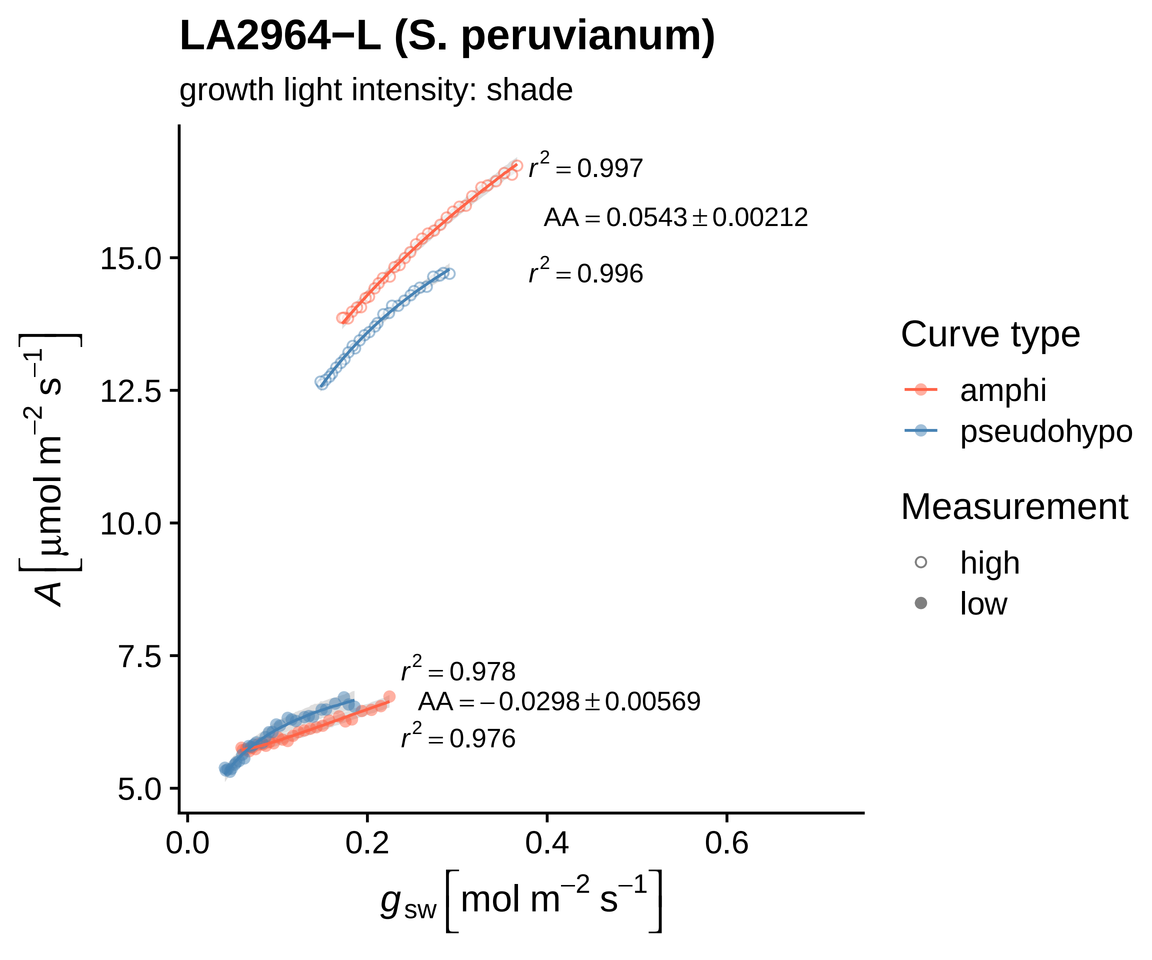

LA2964−L (S. peruvianum)
growth light intensity: shade
$r^2 = 0.997$
$AA = 0.0543 \pm 0.00212$
$r^2 = 0.996$
$r^2 = 0.978$
$AA = -0.0298 \pm 0.00569$
$r^2 = 0.976$
15.0
12.5
10.0
7.5
5.0
0.0
0.2
0.4
0.6
$A\left[\mu mol\ m^{-2}\ s^{-1}\right]$
$g_{sw}\left[mol\ m^{-2}\ s^{-1}\right]$
Curve type
amphi
pseudohypo
Measurement
high
low

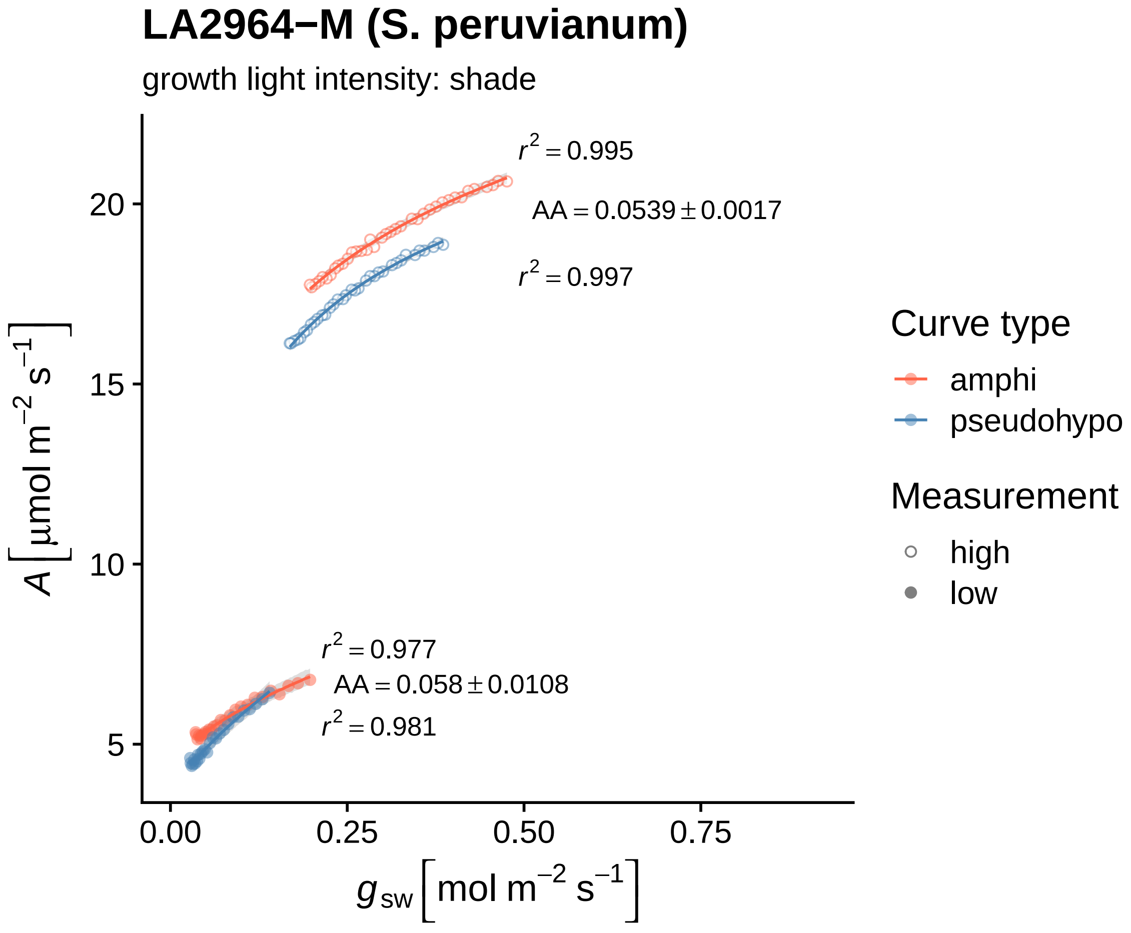

LA2964−M (S. peruvianum)
growth light intensity: shade
$r^2 = 0.995$
$AA = 0.0539 \pm 0.0017$
$r^2 = 0.997$
$r^2 = 0.977$
$AA = 0.058 \pm 0.0108$
$r^2 = 0.981$
20
15
10
5
0.00
0.25
0.50
0.75
$A\ [\mu mol\ m^{-2}\ s^{-1}]$
$g_{sw}\ [mol\ m^{-2}\ s^{-1}]$
Curve type
amphi
pseudohypo
Measurement
high
low

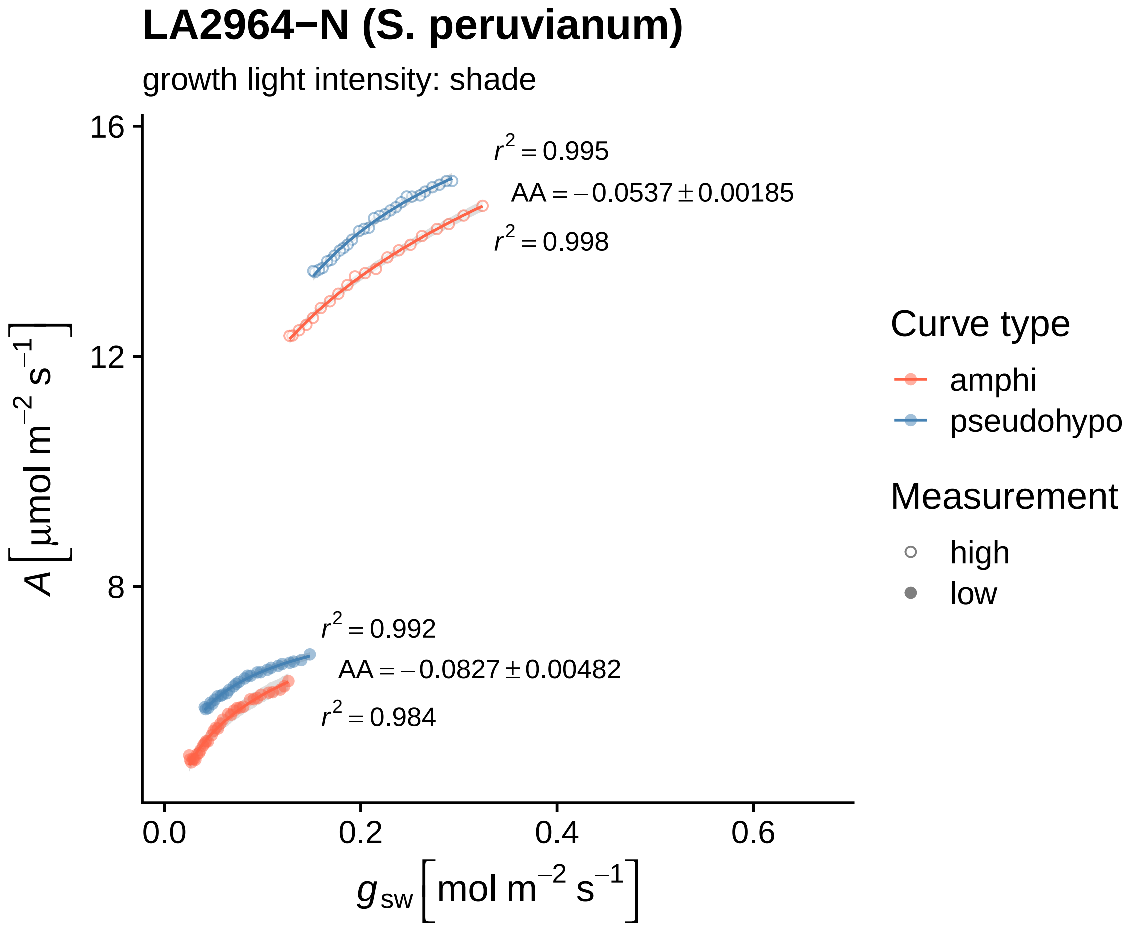

LA2964−N (S. peruvianum)
growth light intensity: shade
r² = 0.995
AA = −0.0537 ± 0.00185
r² = 0.998
r² = 0.992
AA = −0.0827 ± 0.00482
r² = 0.984
16
12
8
0.0
0.2
0.4
0.6
A [µmol m⁻² s⁻¹]
g_sw [mol m⁻² s⁻¹]
Curve type
amphi
pseudohypo
Measurement
high
low

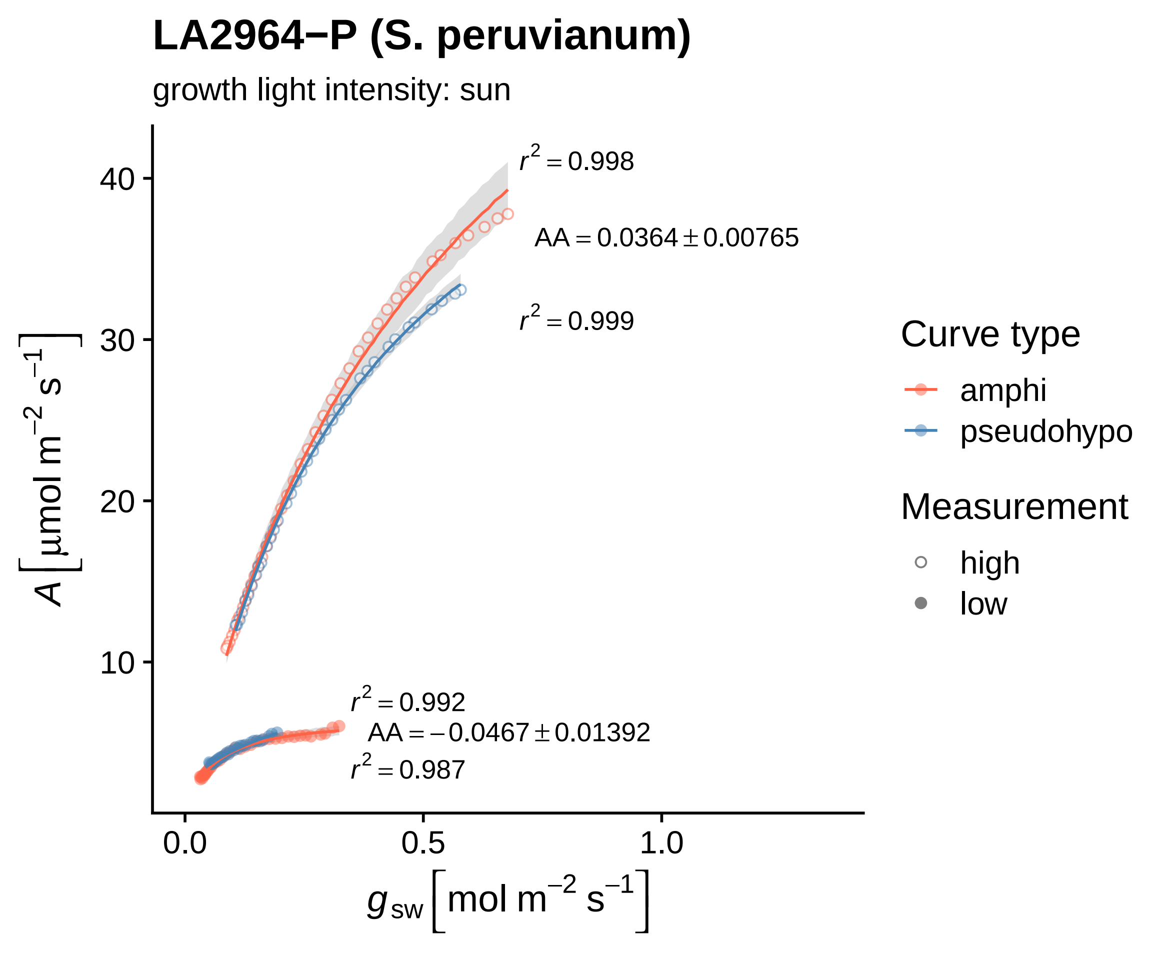
LA2964−P (S. peruvianum)
growth light intensity: sun
$r^2 = 0.998$
$AA = 0.0364 \pm 0.00765$
$r^2 = 0.999$
$r^2 = 0.992$
$AA = -0.0467 \pm 0.01392$
$r^2 = 0.987$
40
30
20
10
0.0
0.5
1.0
$A\ [\mu mol\ m^{-2}\ s^{-1}]$
$g_{sw}\ [mol\ m^{-2}\ s^{-1}]$
Curve type
amphi
pseudohypo
Measurement
high
low

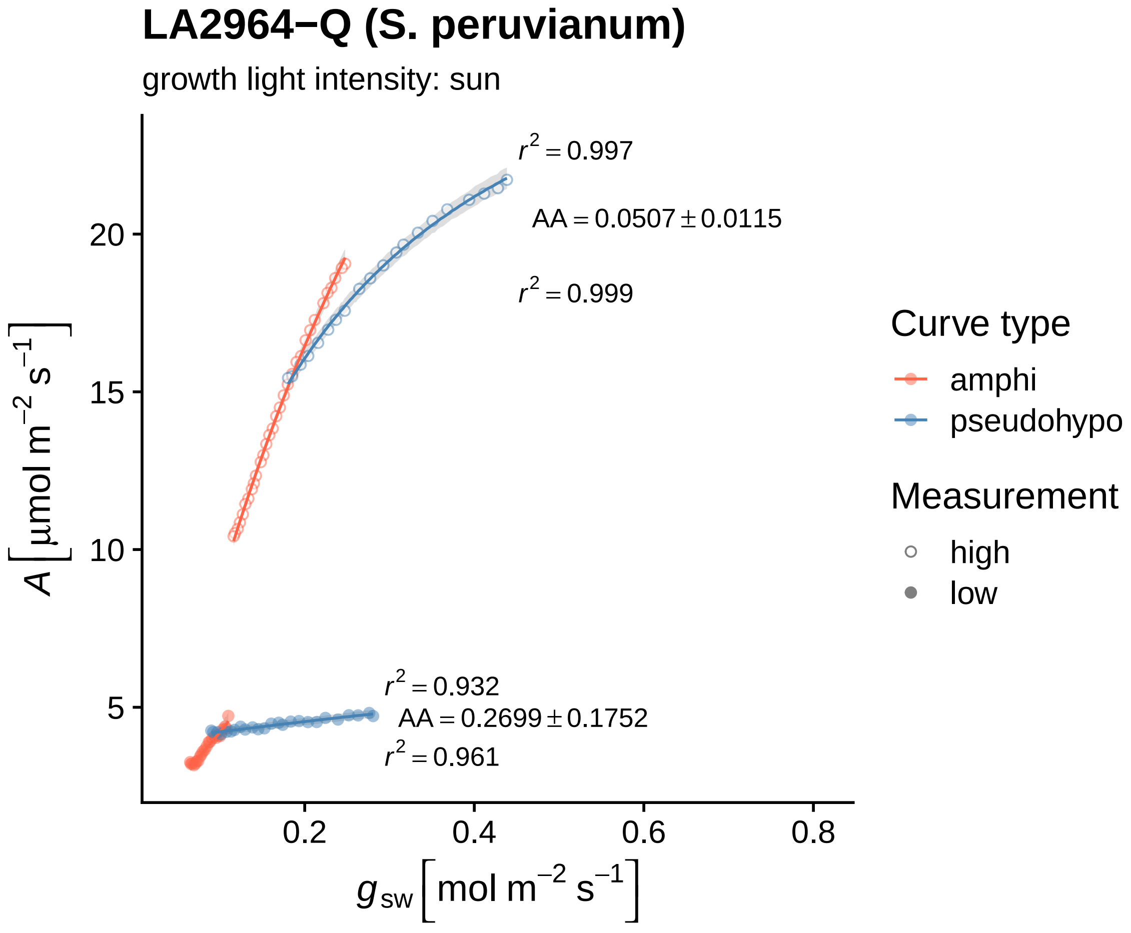

LA2964−Q (S. peruvianum)
growth light intensity: sun
r² = 0.997
AA = 0.0507 ± 0.0115
r² = 0.999
r² = 0.932
AA = 0.2699 ± 0.1752
r² = 0.961
20
15
10
5
0.2
0.4
0.6
0.8
A [μmol m⁻² s⁻¹]
g sw [mol m⁻² s⁻¹]
Curve type
amphi
pseudohypo
Measurement
high
low

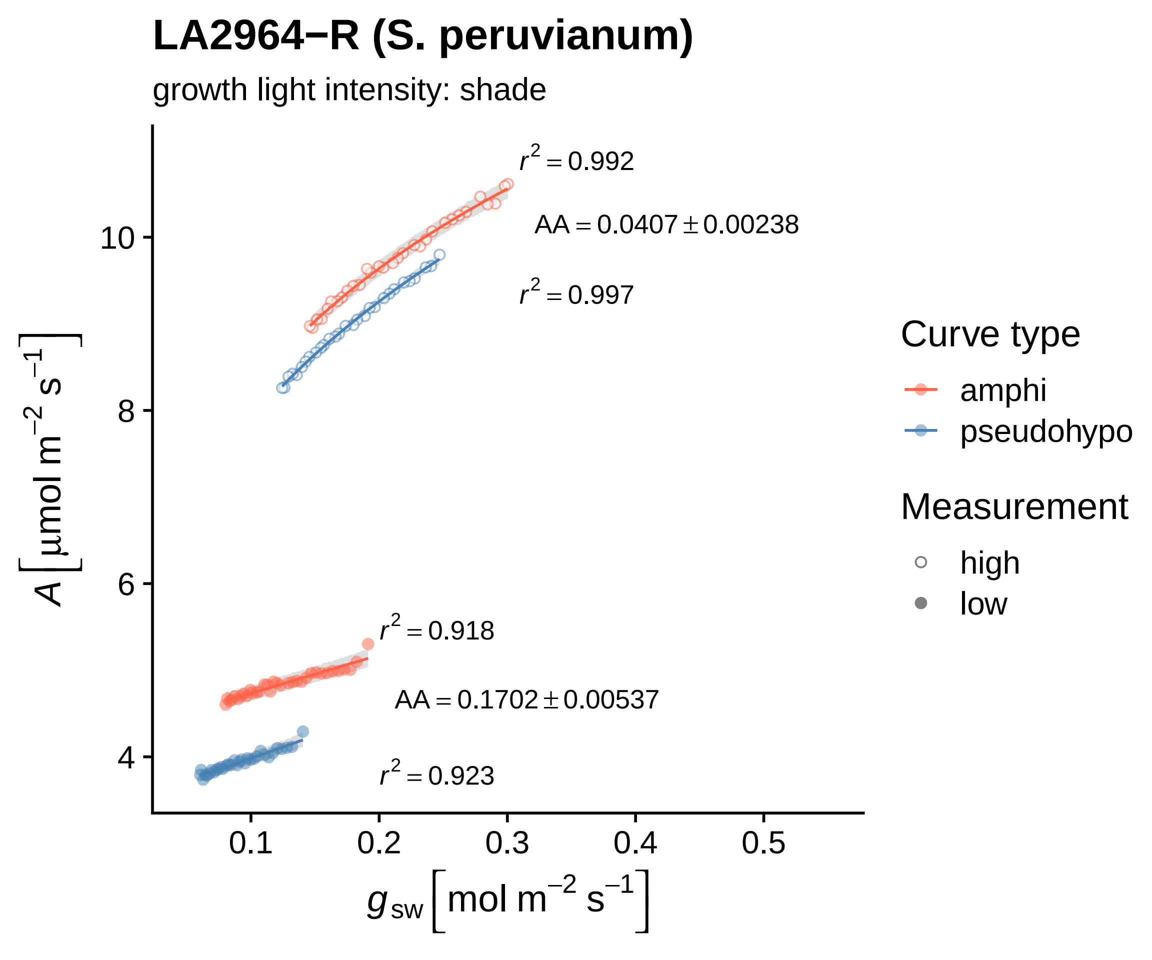

LA2964−R (S. peruvianum)
growth light intensity: shade
r² = 0.992
AA = 0.0407 ± 0.00238
r² = 0.997
r² = 0.918
AA = 0.1702 ± 0.00537
r² = 0.923
10
8
6
4
0.1
0.2
0.3
0.4
0.5
A [μmol m⁻² s⁻¹]
g_sw [mol m⁻² s⁻¹]
Curve type
amphi
pseudohypo
Measurement
high
low

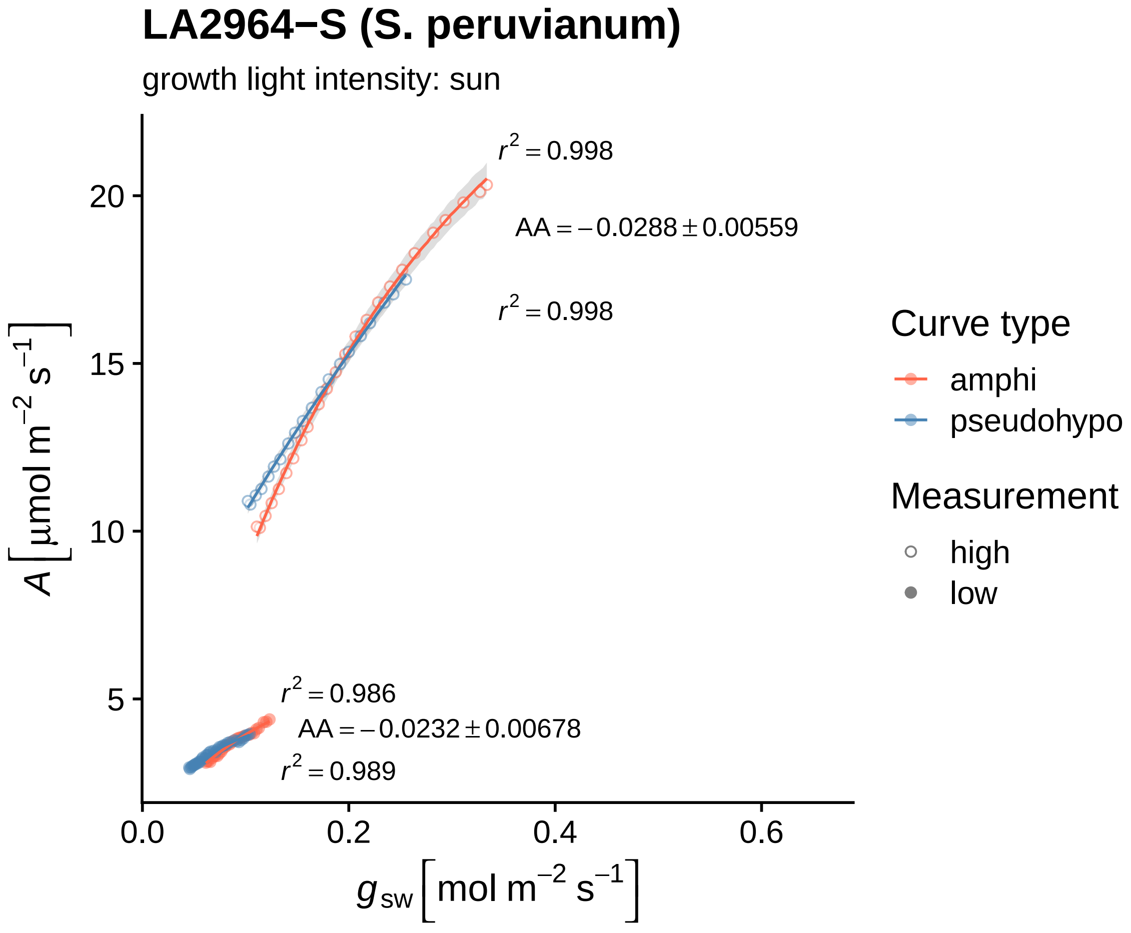

LA2964−S (S. peruvianum)
growth light intensity: sun
r² = 0.998
AA = −0.0288 ± 0.00559
r² = 0.998
r² = 0.986
AA = −0.0232 ± 0.00678
r² = 0.989
A [μmol m⁻² s⁻¹]
g sw [mol m⁻² s⁻¹]
0.0
0.2
0.4
0.6
5
10
15
20
Curve type
amphi
pseudohypo
Measurement
high
low

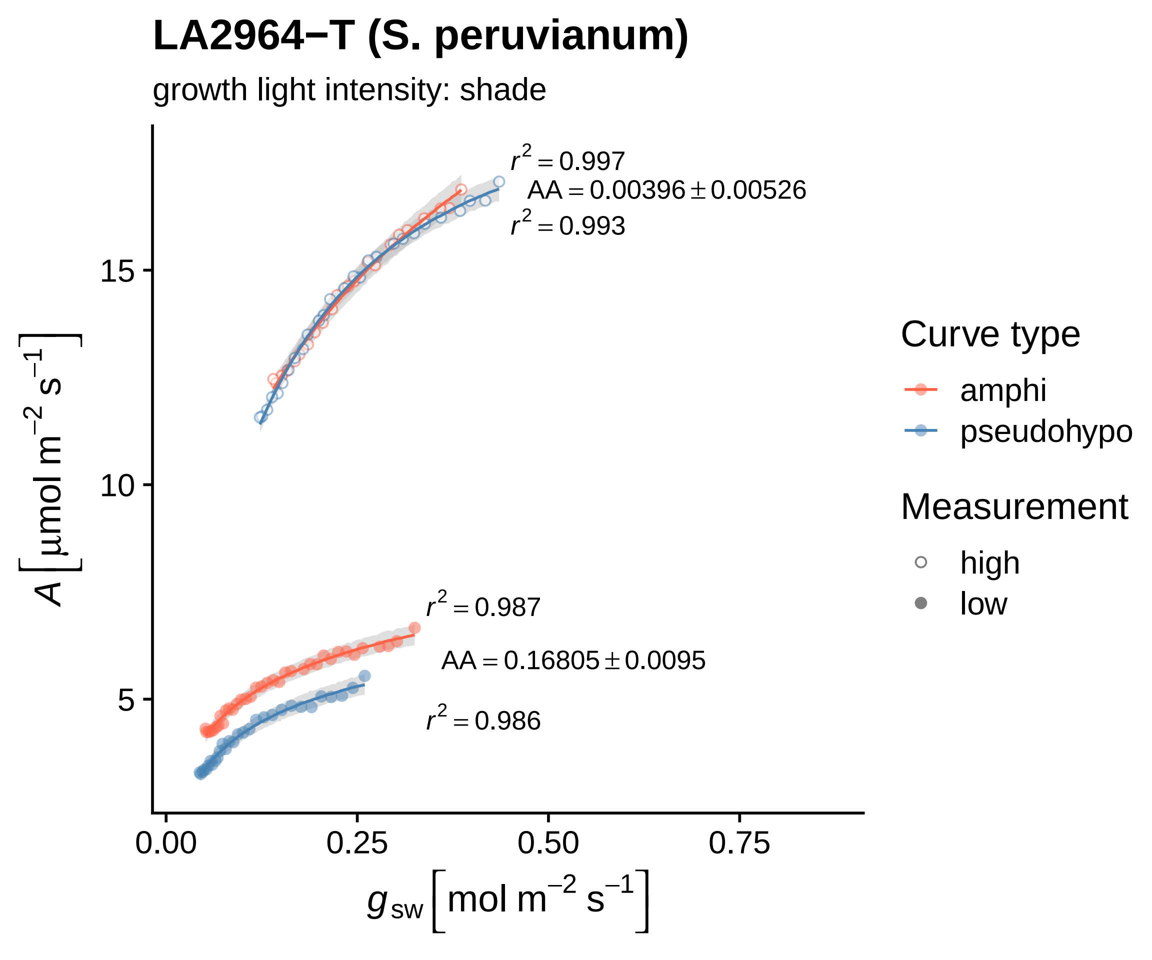

LA2964−T (S. peruvianum)
growth light intensity: shade
$r^2 = 0.997$
AA = 0.00396 ± 0.00526
$r^2 = 0.993$
$r^2 = 0.987$
AA = 0.16805 ± 0.0095
$r^2 = 0.986$
15
10
5
0.00
0.25
0.50
0.75
$A\ [\mu mol\ m^{-2}\ s^{-1}]$
$g_{sw}\ [mol\ m^{-2}\ s^{-1}]$
Curve type
amphi
pseudohypo
Measurement
high
low

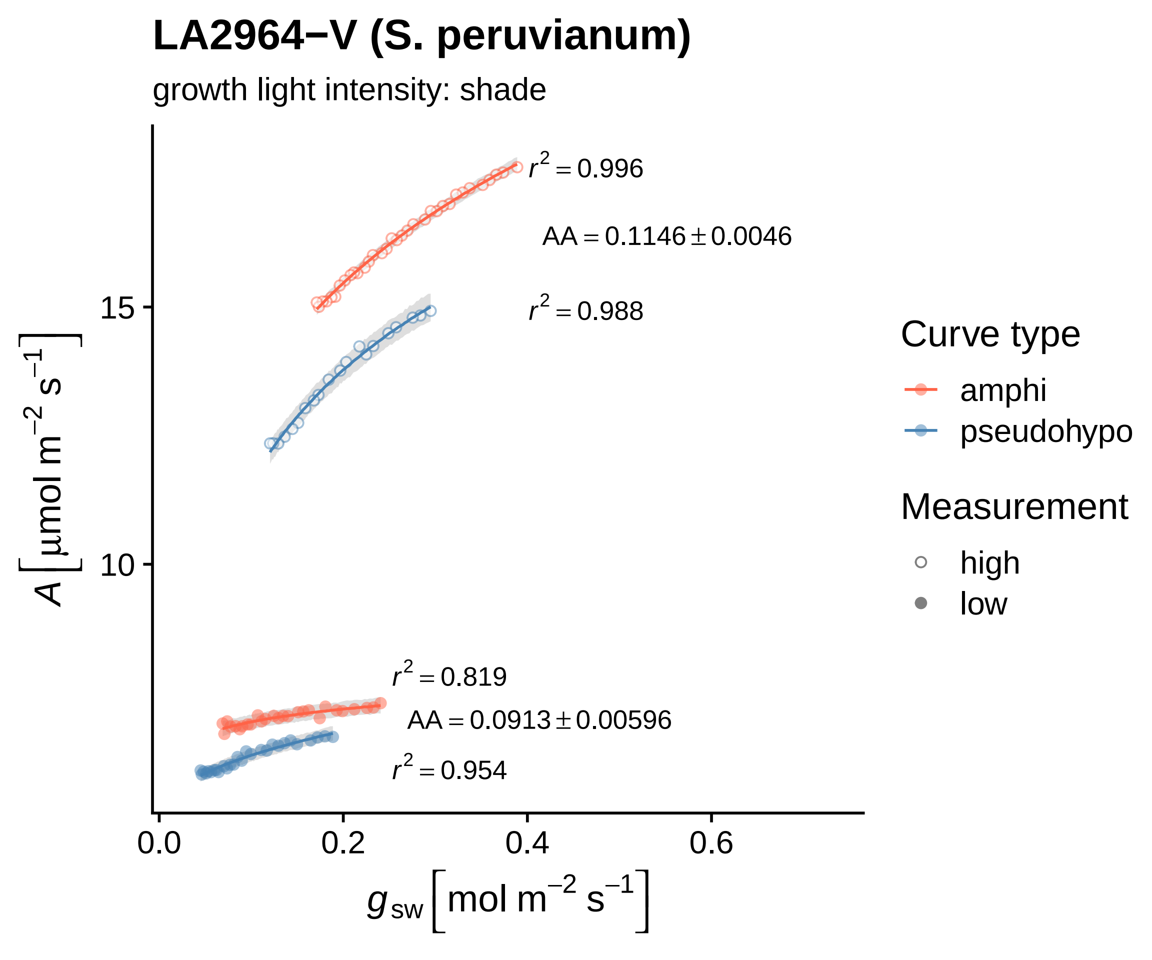

LA2964−V (S. peruvianum)
growth light intensity: shade
$r^2 = 0.996$
$AA = 0.1146 \pm 0.0046$
$r^2 = 0.988$
$r^2 = 0.819$
$AA = 0.0913 \pm 0.00596$
$r^2 = 0.954$
15
10
0.0
0.2
0.4
0.6
$A\ [\mu mol\ m^{-2}\ s^{-1}]$
$g_{sw}\ [mol\ m^{-2}\ s^{-1}]$
Curve type
amphi
pseudohypo
Measurement
high
low

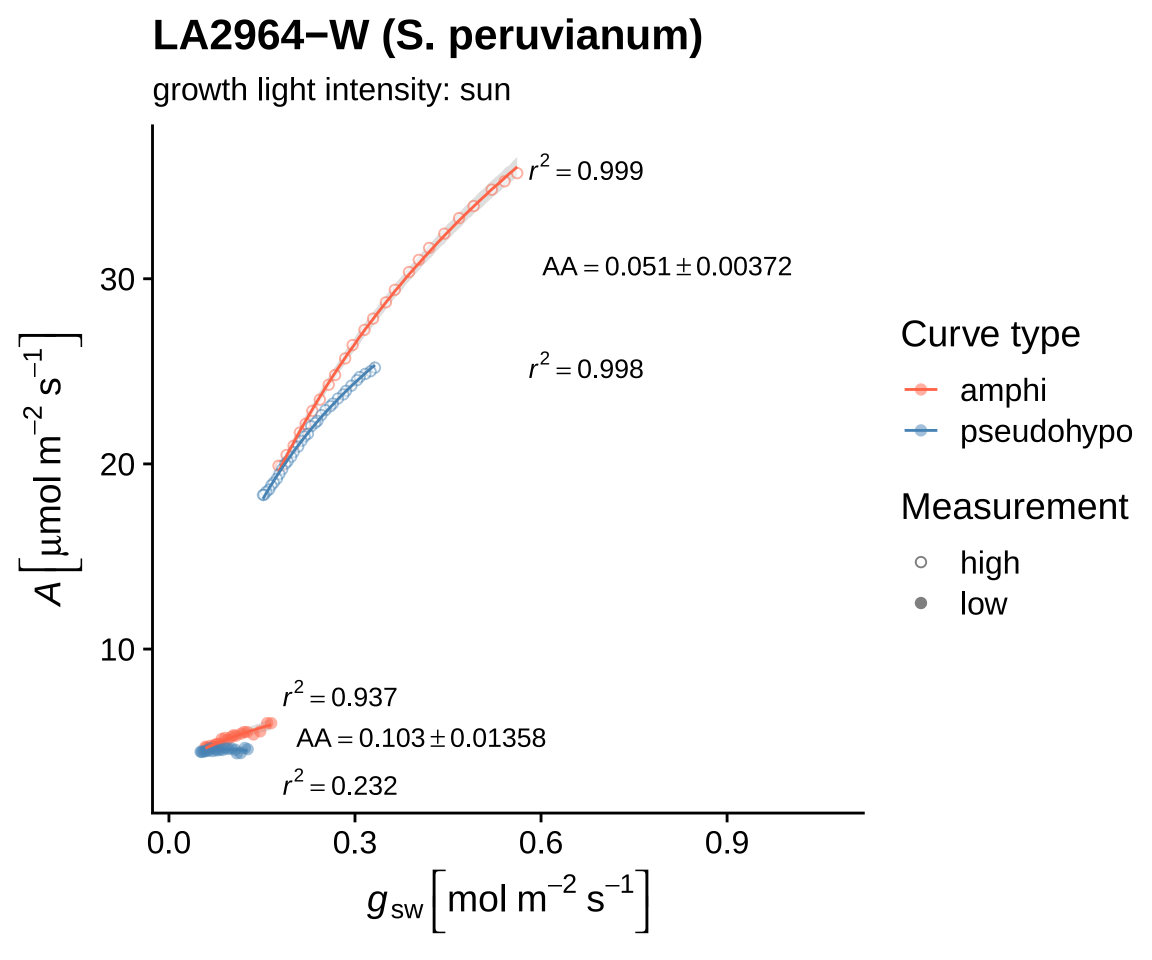

LA2964−W (S. peruvianum)
growth light intensity: sun
$r^2 = 0.999$
$AA = 0.051 \pm 0.00372$
$r^2 = 0.998$
$r^2 = 0.937$
$AA = 0.103 \pm 0.01358$
$r^2 = 0.232$
$A\,[\mu mol\ m^{-2}\ s^{-1}]$
30
20
10
0.0
0.3
0.6
0.9
$g_{sw}\,[mol\ m^{-2}\ s^{-1}]$
Curve type
amphi
pseudohypo
Measurement
high
low

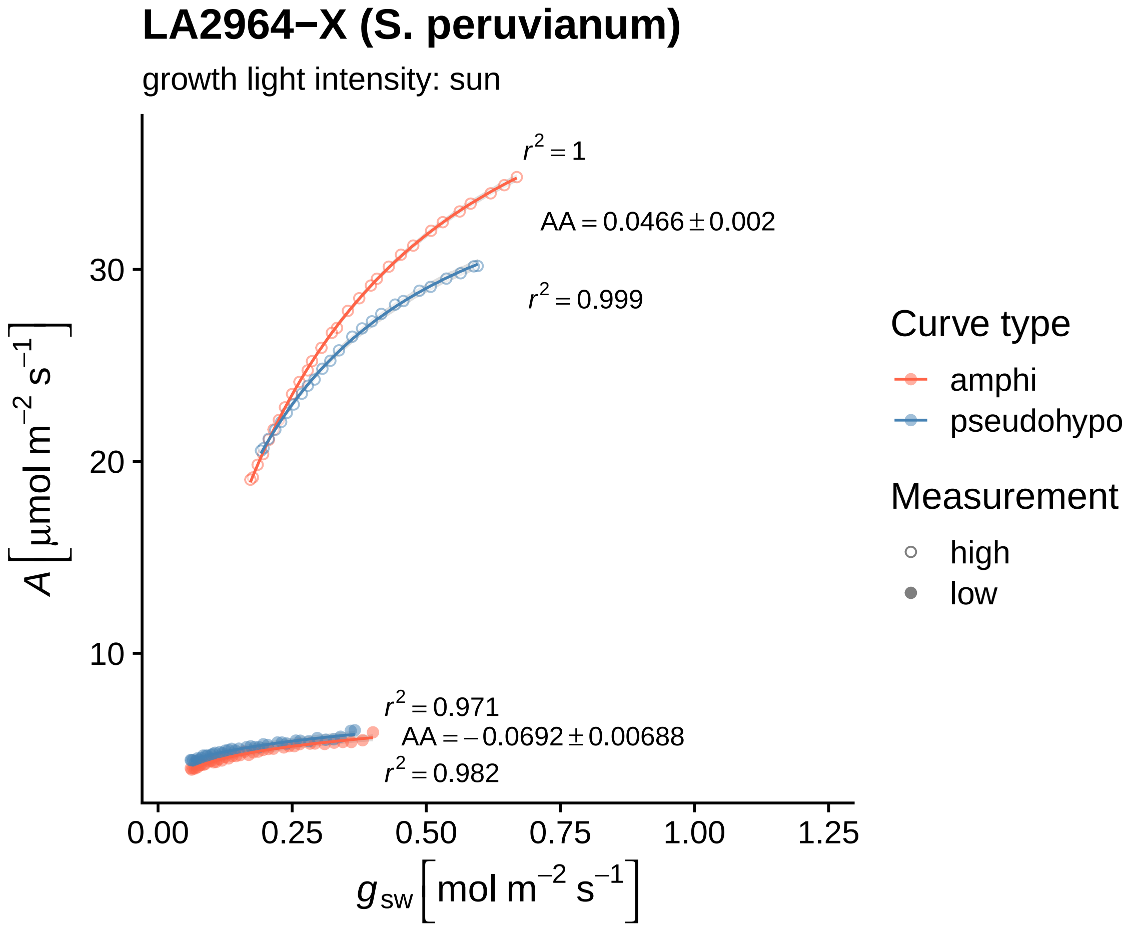

LA2964−X (S. peruvianum)
growth light intensity: sun
$r^2 = 1$
$AA = 0.0466 \pm 0.002$
$r^2 = 0.999$
$r^2 = 0.971$
$AA = -0.0692 \pm 0.00688$
$r^2 = 0.982$
30
20
10
0.00
0.25
0.50
0.75
1.00
1.25
$A\ [\mu mol\ m^{-2}\ s^{-1}]$
$g_{sw}\ [mol\ m^{-2}\ s^{-1}]$
Curve type
amphi
pseudohypo
Measurement
high
low

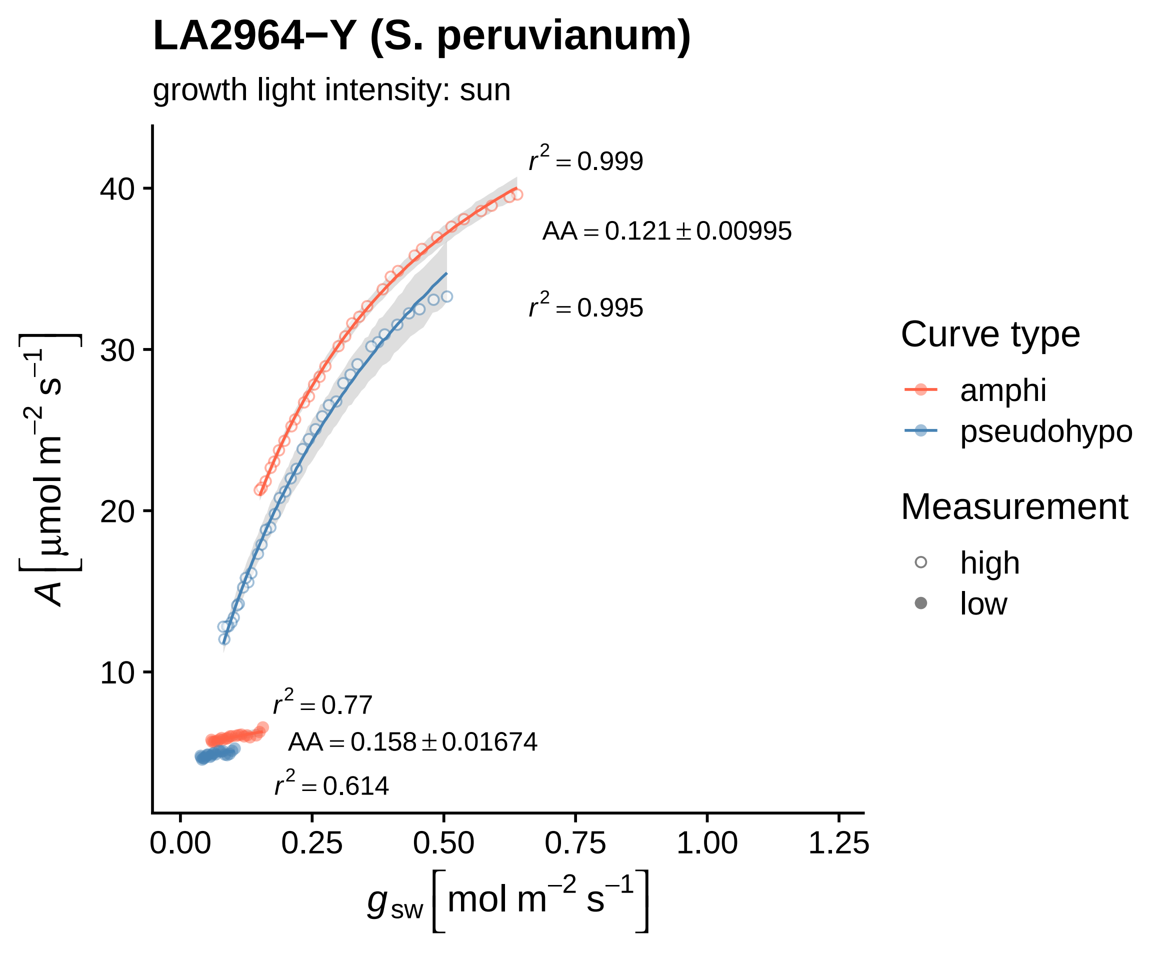

LA2964−Y (S. peruvianum)
growth light intensity: sun
r² = 0.999
AA = 0.121 ± 0.00995
r² = 0.995
r² = 0.77
AA = 0.158 ± 0.01674
r² = 0.614
40
30
20
10
0.00
0.25
0.50
0.75
1.00
1.25
A [µmol m⁻² s⁻¹]
gsw [mol m⁻² s⁻¹]
Curve type
amphi
pseudohypo
Measurement
high
low

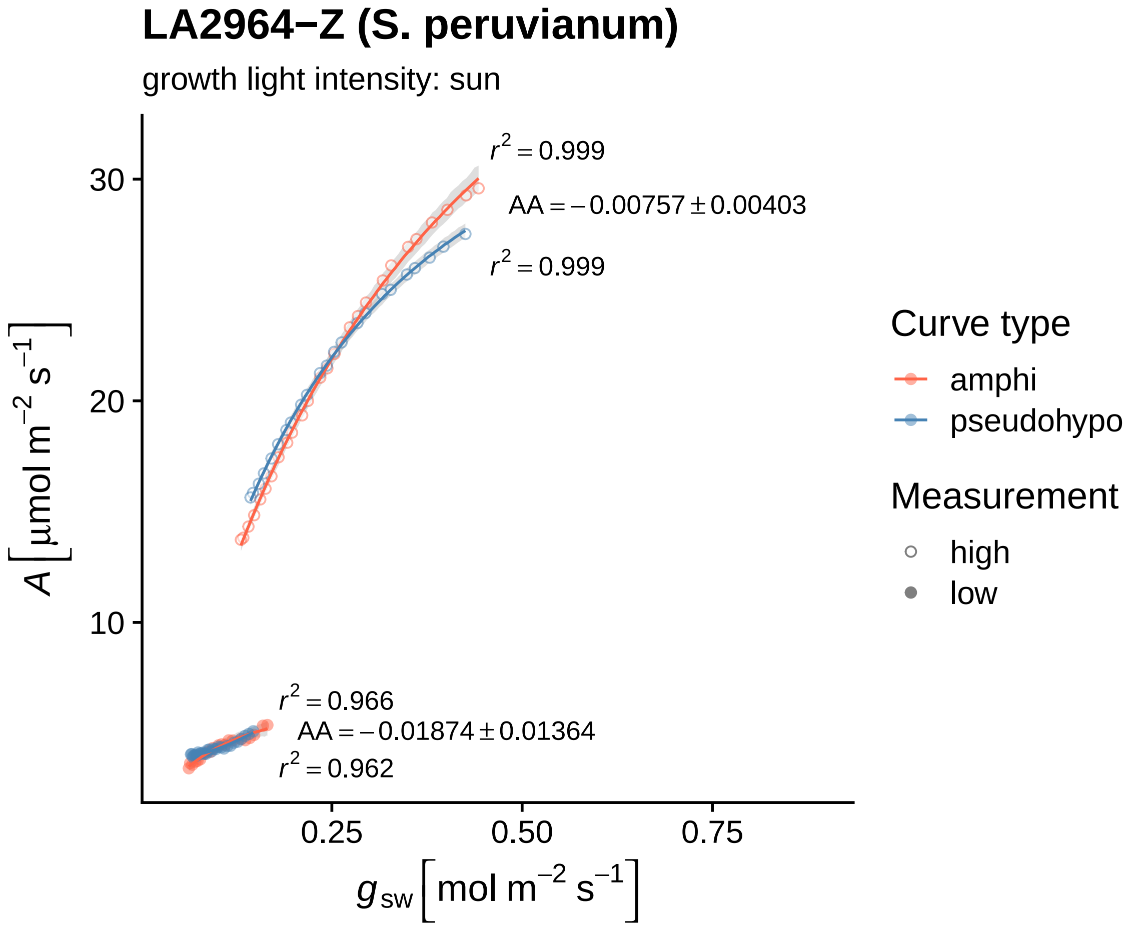
LA2964−Z (S. peruvianum)
growth light intensity: sun
$r^2 = 0.999$
$AA = -0.00757 \pm 0.00403$
$r^2 = 0.999$
$r^2 = 0.966$
$AA = -0.01874 \pm 0.01364$
$r^2 = 0.962$
30
20
10
0.25
0.50
0.75
$A\ [\mu mol\ m^{-2}\ s^{-1}]$
$g_{sw}\ [mol\ m^{-2}\ s^{-1}]$
Curve type
amphi
pseudohypo
Measurement
high
low

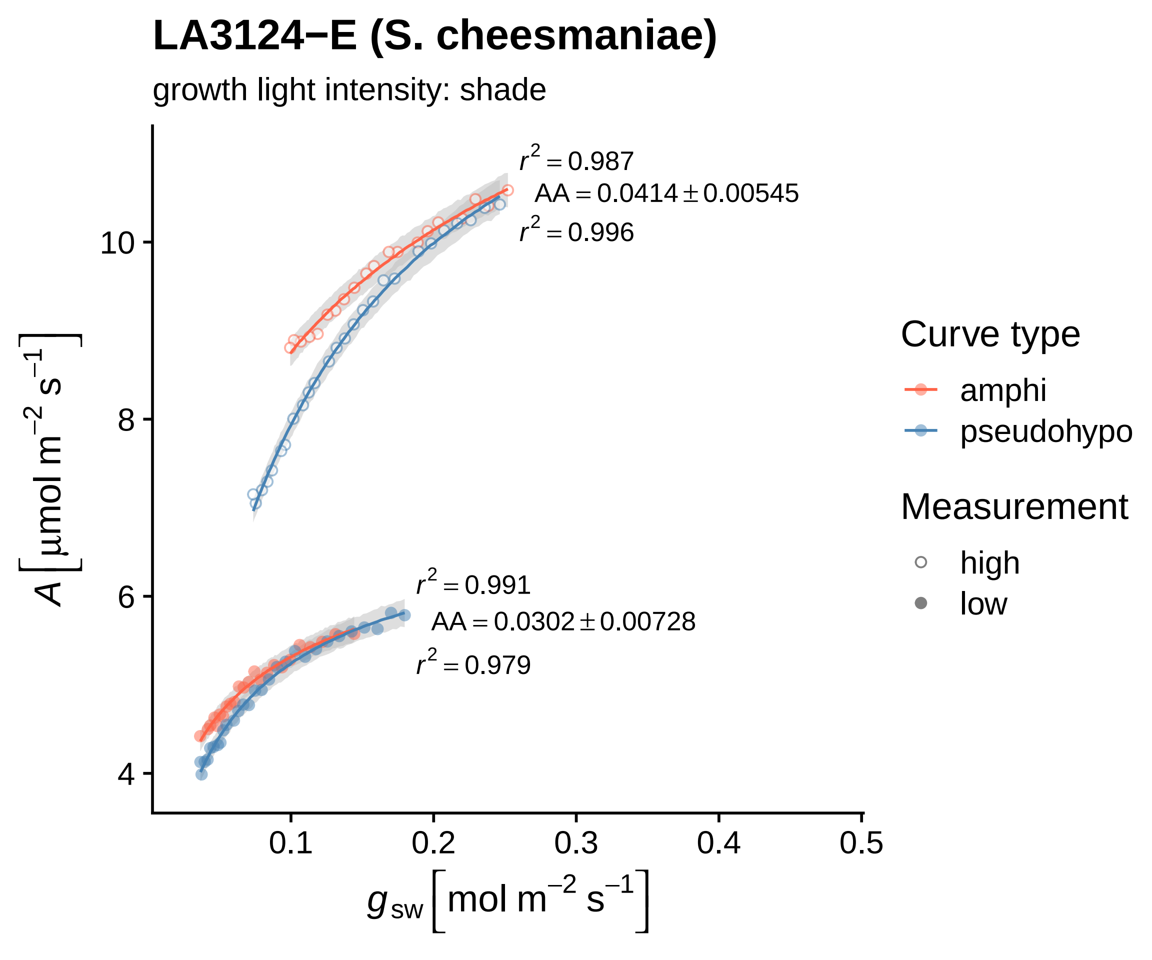

LA3124−E (S. cheesmaniae)
growth light intensity: shade
$r^2 = 0.987$
$AA = 0.0414 \pm 0.00545$
$r^2 = 0.996$
$r^2 = 0.991$
$AA = 0.0302 \pm 0.00728$
$r^2 = 0.979$
10
8
6
4
0.1
0.2
0.3
0.4
0.5
$A\ [\mu mol\ m^{-2}\ s^{-1}]$
$g_{sw}\ [mol\ m^{-2}\ s^{-1}]$
Curve type
amphi
pseudohypo
Measurement
high
low

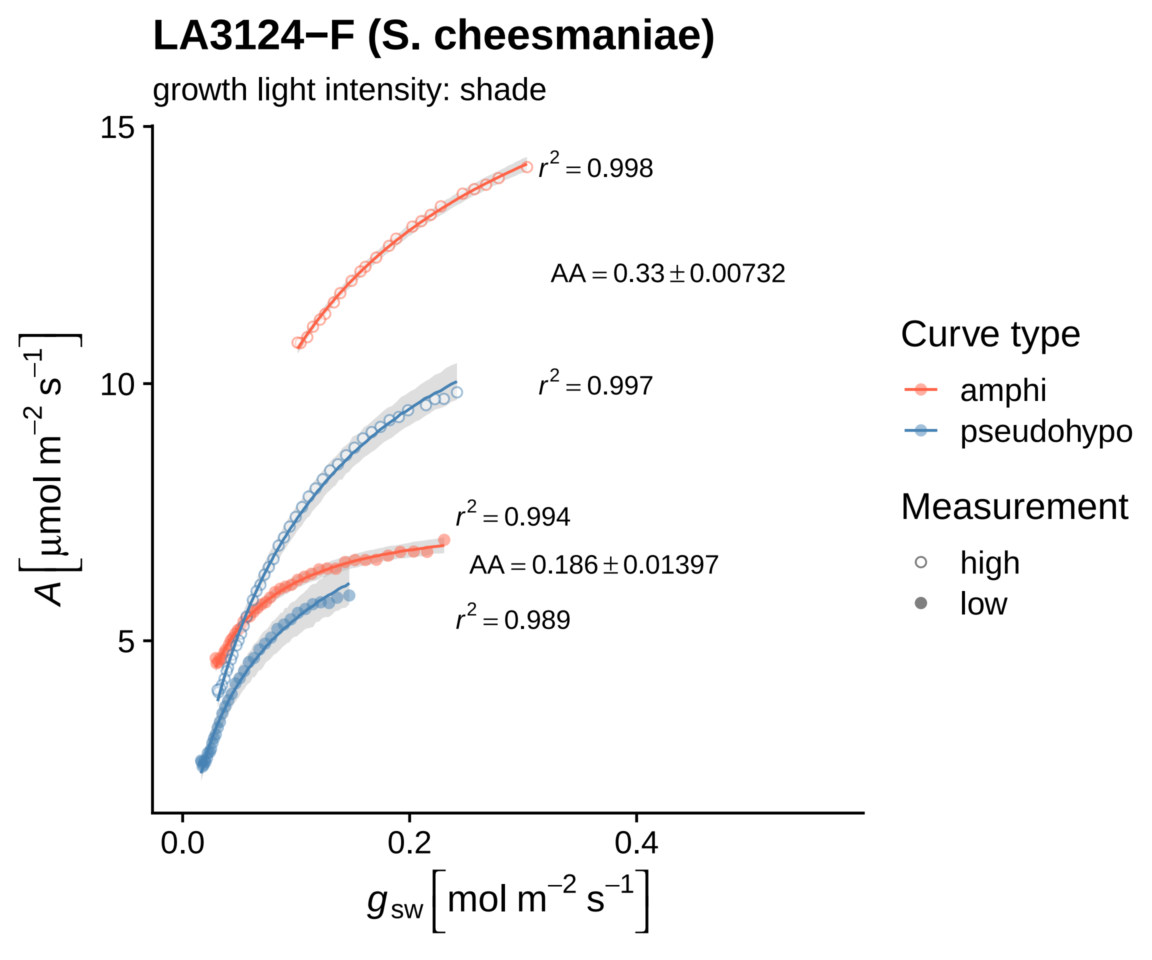
LA3124−F (S. cheesmaniae)
growth light intensity: shade
$r^2 = 0.998$
$AA = 0.33 \pm 0.00732$
$r^2 = 0.997$
$r^2 = 0.994$
$AA = 0.186 \pm 0.01397$
$r^2 = 0.989$
15
10
5
0.0
0.2
0.4
$A\left[\mu mol\ m^{-2}\ s^{-1}\right]$
$g_{sw}\left[mol\ m^{-2}\ s^{-1}\right]$
Curve type
amphi
pseudohypo
Measurement
high
low

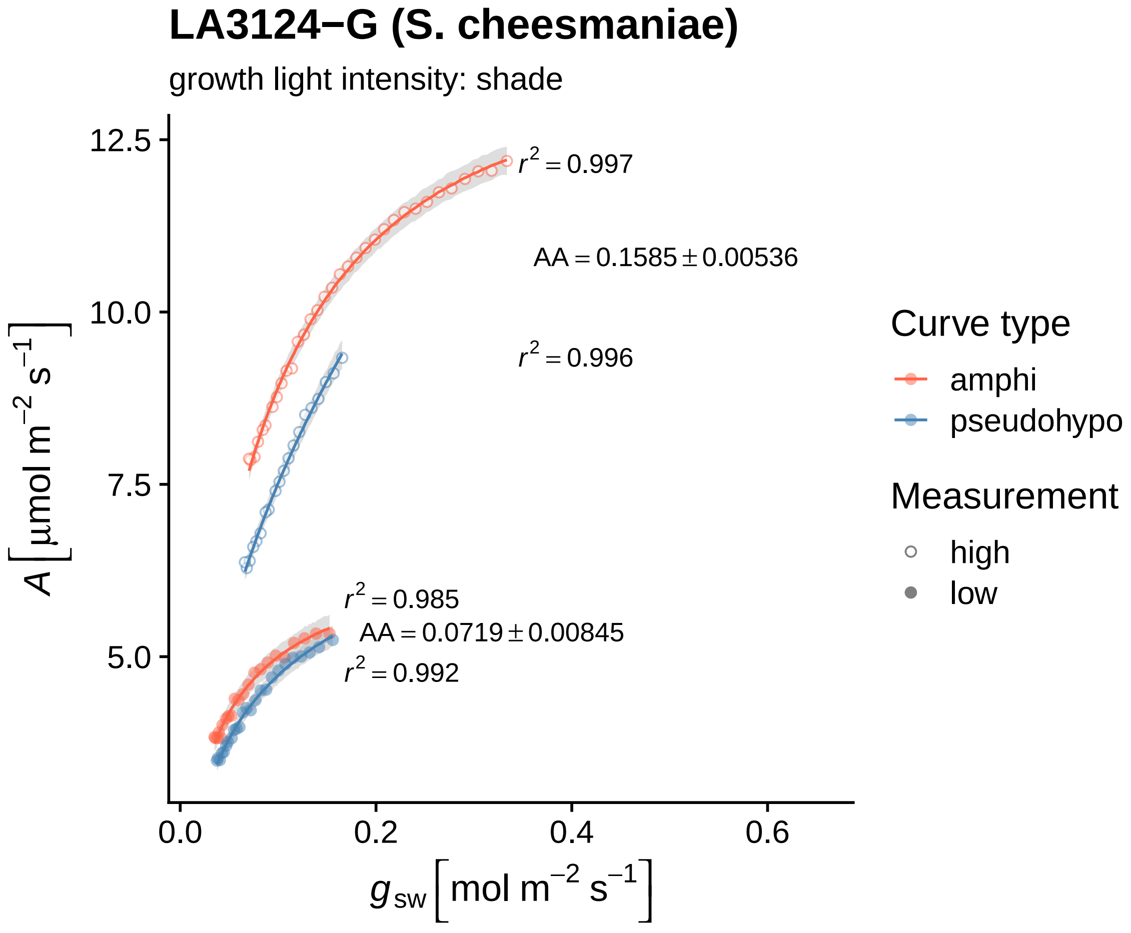

LA3124−G (S. cheesmaniae)
growth light intensity: shade
12.5
10.0
7.5
5.0
0.0
0.2
0.4
0.6
$A\left[\mu mol\ m^{-2}\ s^{-1}\right]$
$g_{sw}\left[mol\ m^{-2}\ s^{-1}\right]$
$r^2 = 0.997$
$AA = 0.1585 \pm 0.00536$
$r^2 = 0.996$
$r^2 = 0.985$
$AA = 0.0719 \pm 0.00845$
$r^2 = 0.992$
Curve type
amphi
pseudohypo
Measurement
high
low

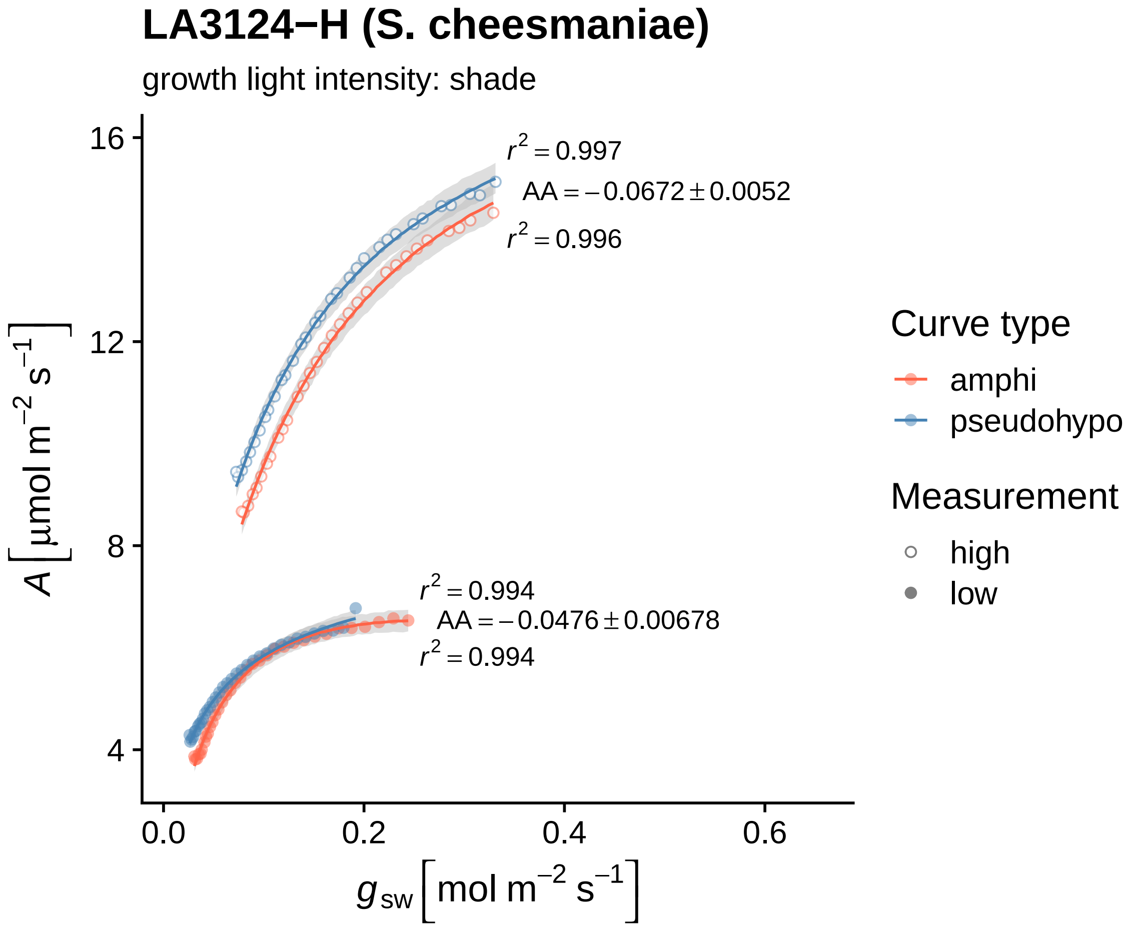

LA3124−H (S. cheesmaniae)
growth light intensity: shade
$r^2 = 0.997$
$AA = -0.0672 \pm 0.0052$
$r^2 = 0.996$
$r^2 = 0.994$
$AA = -0.0476 \pm 0.00678$
$r^2 = 0.994$
16
12
8
4
0.0
0.2
0.4
0.6
$A\left[\mu mol\ m^{-2}\ s^{-1}\right]$
$g_{sw}\left[mol\ m^{-2}\ s^{-1}\right]$
Curve type
amphi
pseudohypo
Measurement
high
low

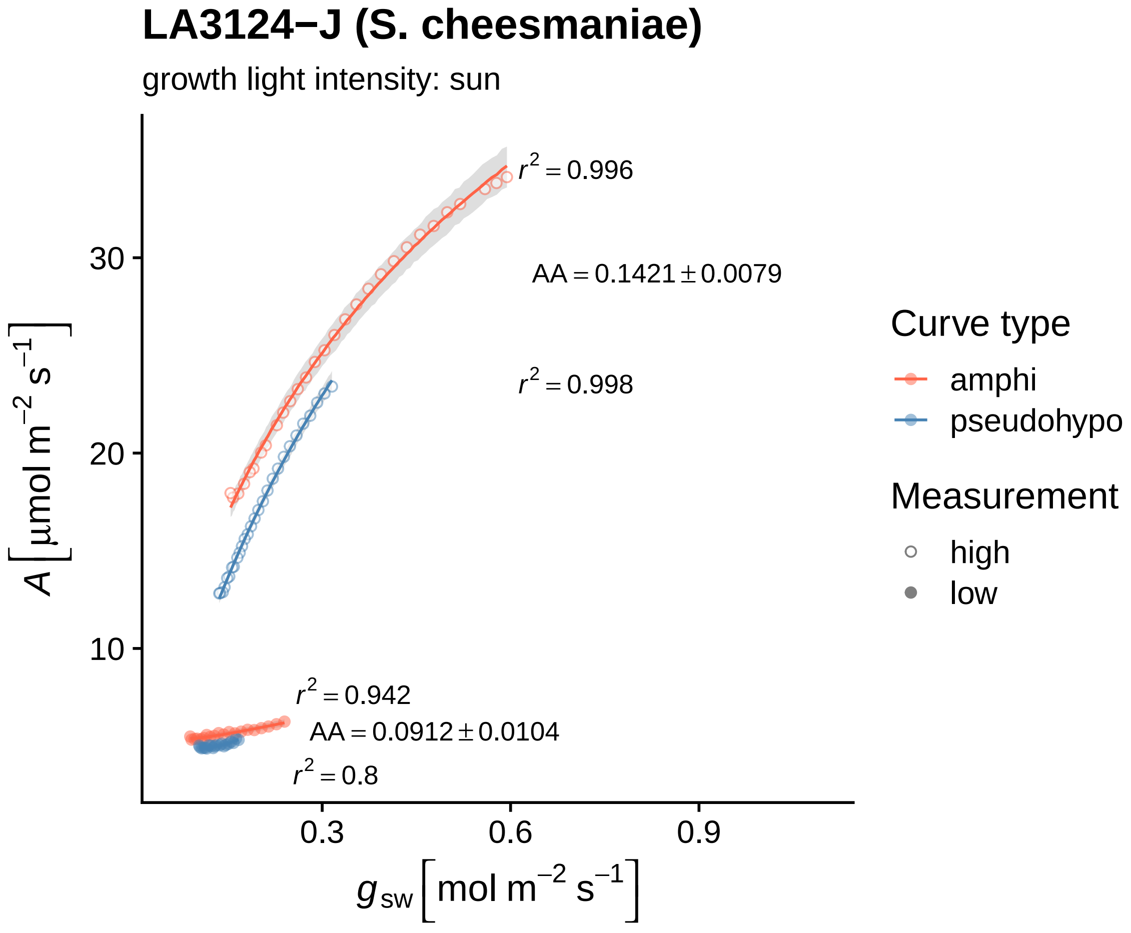

LA3124−J (S. cheesmaniae)
growth light intensity: sun
$r^2 = 0.996$
$AA = 0.1421 \pm 0.0079$
$r^2 = 0.998$
$r^2 = 0.942$
$AA = 0.0912 \pm 0.0104$
$r^2 = 0.8$
30
20
10
0.3
0.6
0.9
$A\left[\mu mol\ m^{-2}\ s^{-1}\right]$
$g_{sw}\left[mol\ m^{-2}\ s^{-1}\right]$
Curve type
amphi
pseudohypo
Measurement
high
low

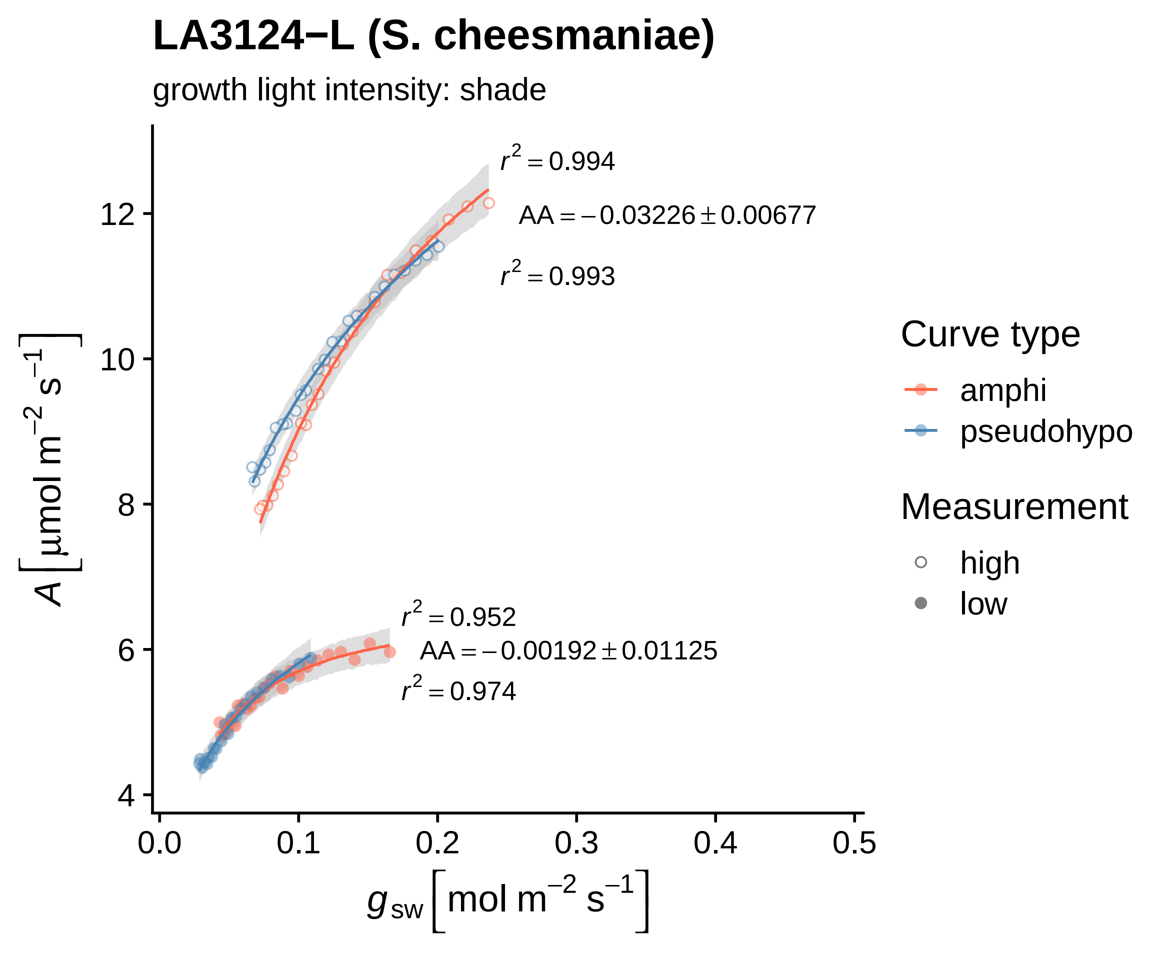

LA3124−L (S. cheesmaniae)
growth light intensity: shade
r² = 0.994
AA = −0.03226 ± 0.00677
r² = 0.993
r² = 0.952
AA = −0.00192 ± 0.01125
r² = 0.974
12
10
8
6
4
0.0
0.1
0.2
0.3
0.4
0.5
A [μmol m⁻² s⁻¹]
g sw [mol m⁻² s⁻¹]
Curve type
amphi
pseudohypo
Measurement
high
low

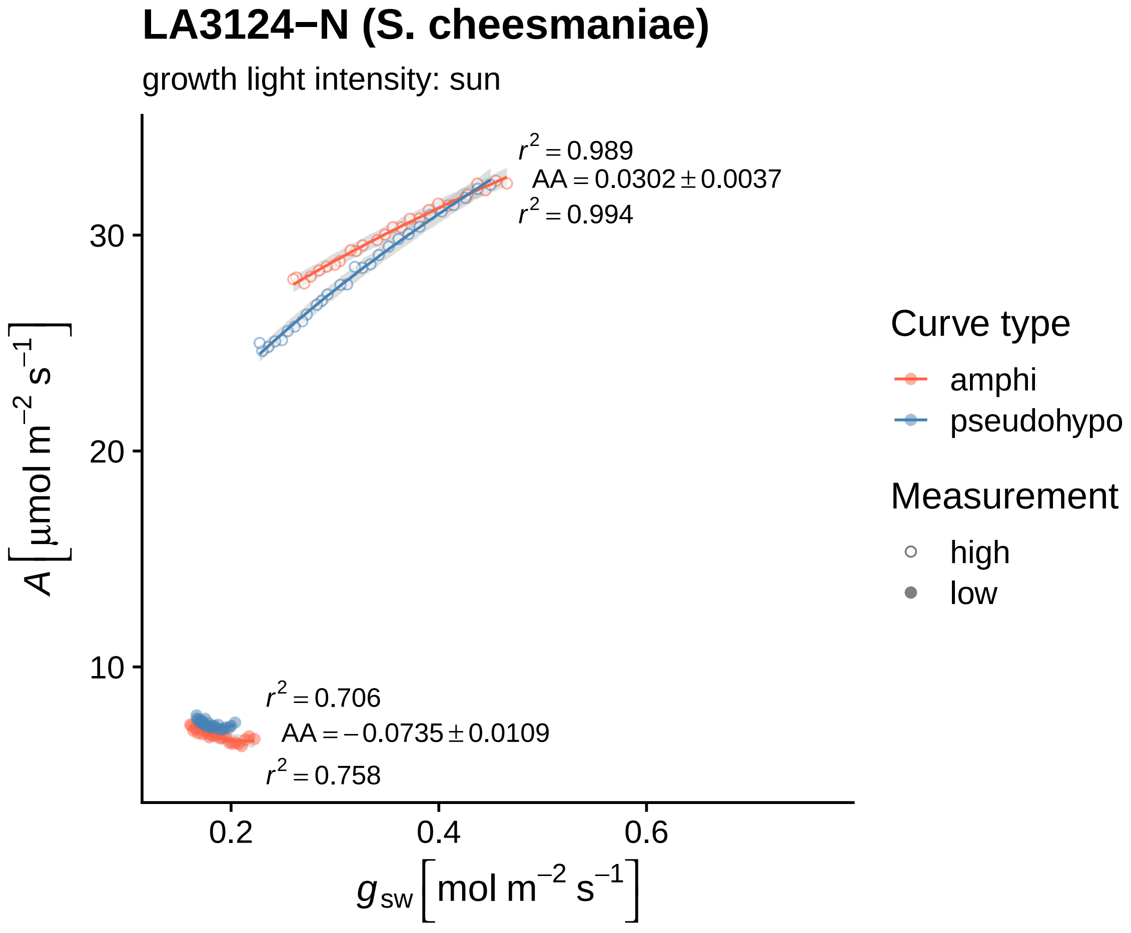
LA3124−N (S. cheesmaniae)
growth light intensity: sun
r² = 0.989
AA = 0.0302 ± 0.0037
r² = 0.994
r² = 0.706
AA = −0.0735 ± 0.0109
r² = 0.758
30
20
10
0.2
0.4
0.6
A [μmol m⁻² s⁻¹]
g sw [mol m⁻² s⁻¹]
Curve type
amphi
pseudohypo
Measurement
high
low

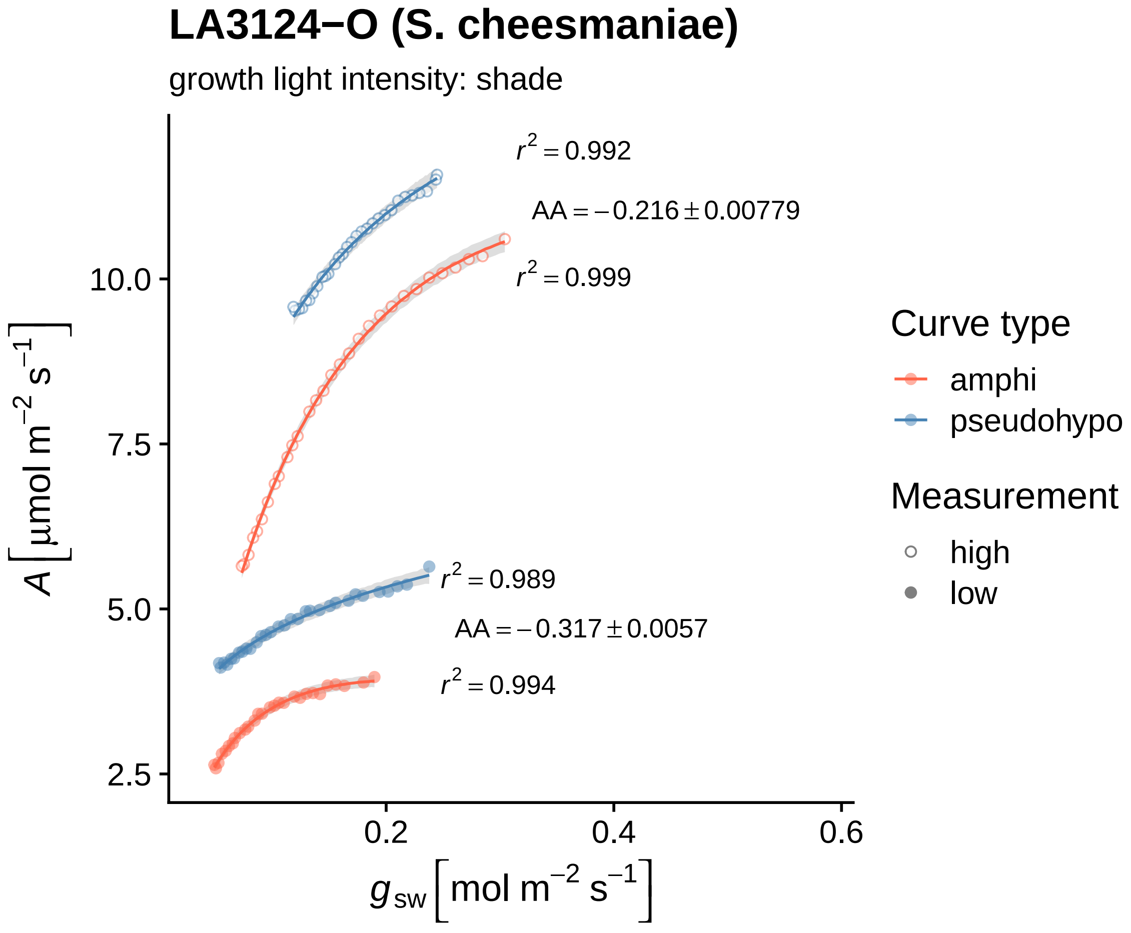
LA3124−O (S. cheesmaniae)
growth light intensity: shade
$r^2 = 0.992$
$AA = -0.216 \pm 0.00779$
$r^2 = 0.999$
$r^2 = 0.989$
$AA = -0.317 \pm 0.0057$
$r^2 = 0.994$
10.0
7.5
5.0
2.5
0.2
0.4
0.6
$A\,[\mu mol\ m^{-2}\ s^{-1}]$
$g_{sw}\,[mol\ m^{-2}\ s^{-1}]$
Curve type
amphi
pseudohypo
Measurement
high
low

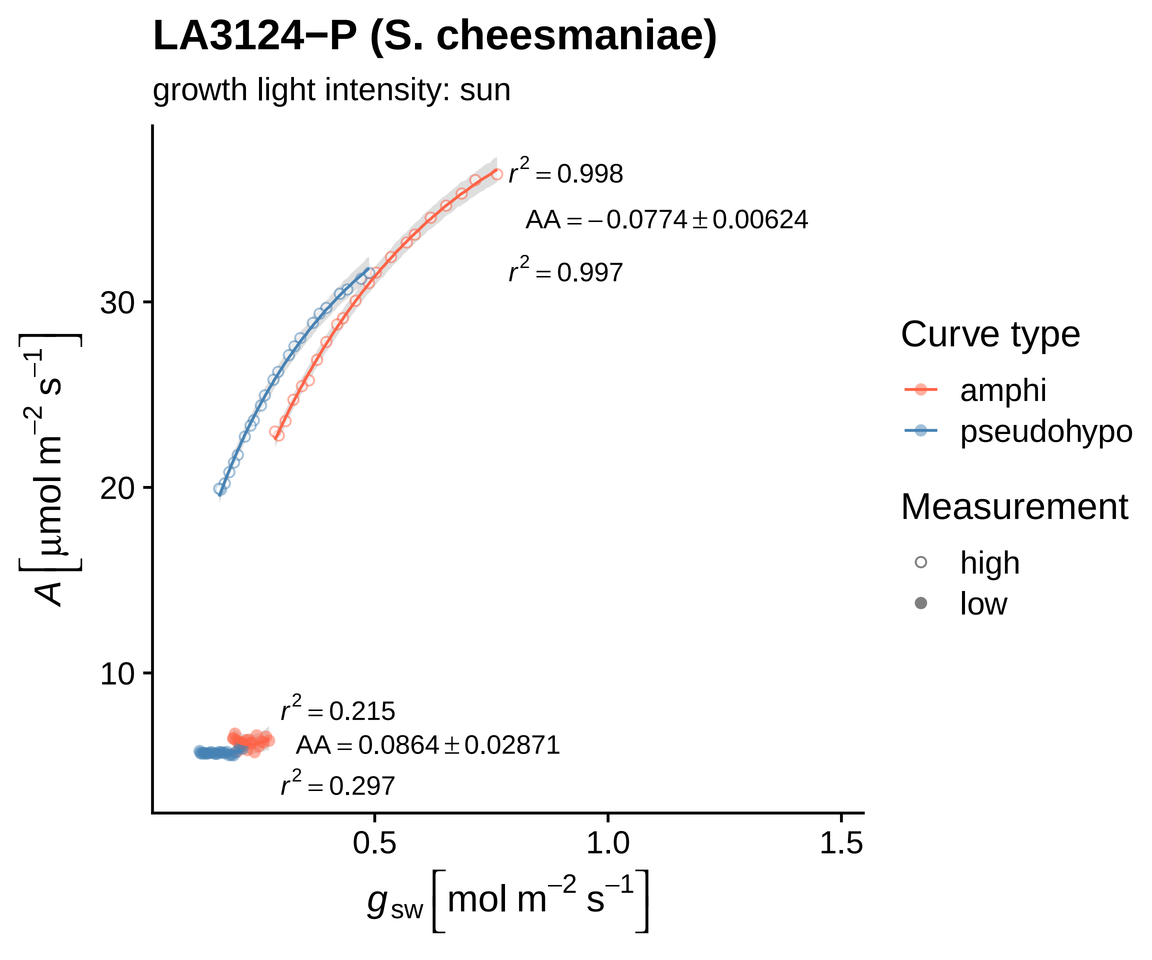

LA3124−P (S. cheesmaniae)
growth light intensity: sun
$r^2 = 0.998$
$AA = -0.0774 \pm 0.00624$
$r^2 = 0.997$
$r^2 = 0.215$
$AA = 0.0864 \pm 0.02871$
$r^2 = 0.297$
30
20
10
0.5
1.0
1.5
$A\left[\mu mol\ m^{-2}\ s^{-1}\right]$
$g_{sw}\left[mol\ m^{-2}\ s^{-1}\right]$
Curve type
amphi
pseudohypo
Measurement
high
low

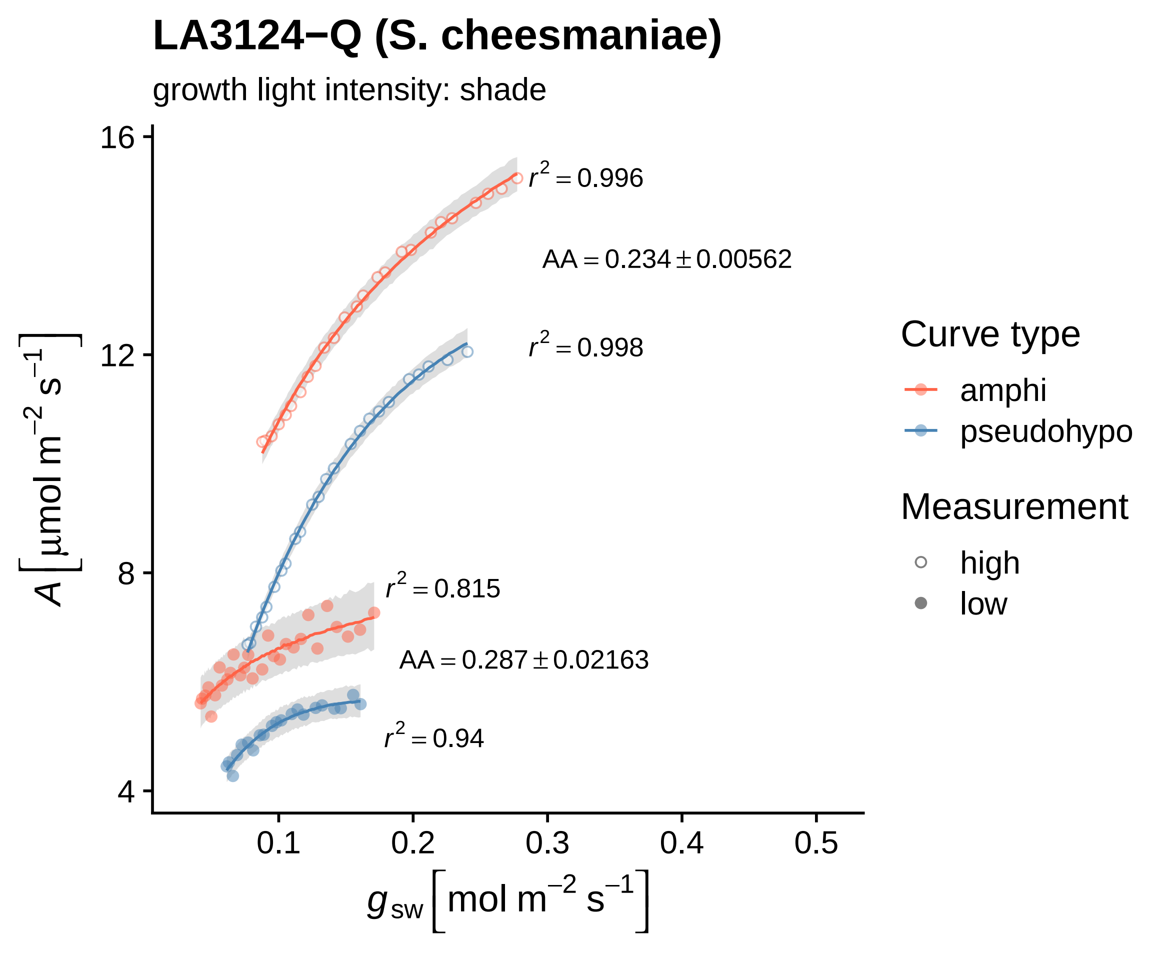

LA3124−Q (S. cheesmaniae)
growth light intensity: shade
$A\left[\mu mol\ m^{-2}\ s^{-1}\right]$
16
12
8
4
0.1
0.2
0.3
0.4
0.5
$g_{sw}\left[mol\ m^{-2}\ s^{-1}\right]$
$r^2 = 0.996$
AA = 0.234 ± 0.00562
$r^2 = 0.998$
$r^2 = 0.815$
AA = 0.287 ± 0.02163
$r^2 = 0.94$
Curve type
amphi
pseudohypo
Measurement
high
low

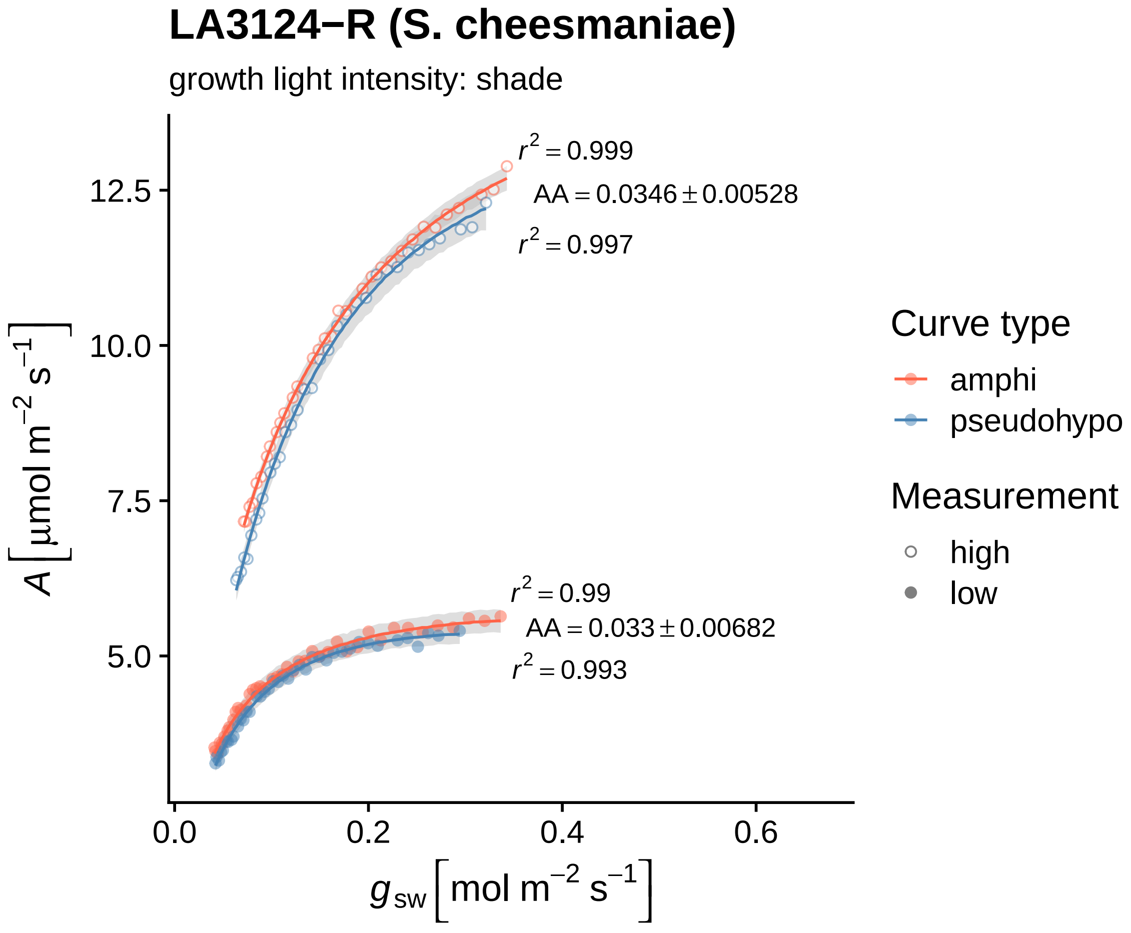

LA3124−R (S. cheesmaniae)
growth light intensity: shade
r² = 0.999
AA = 0.0346 ± 0.00528
r² = 0.997
r² = 0.99
AA = 0.033 ± 0.00682
r² = 0.993
12.5
10.0
7.5
5.0
0.0
0.2
0.4
0.6
A [μmol m⁻² s⁻¹]
g_sw [mol m⁻² s⁻¹]
Curve type
amphi
pseudohypo
Measurement
high
low

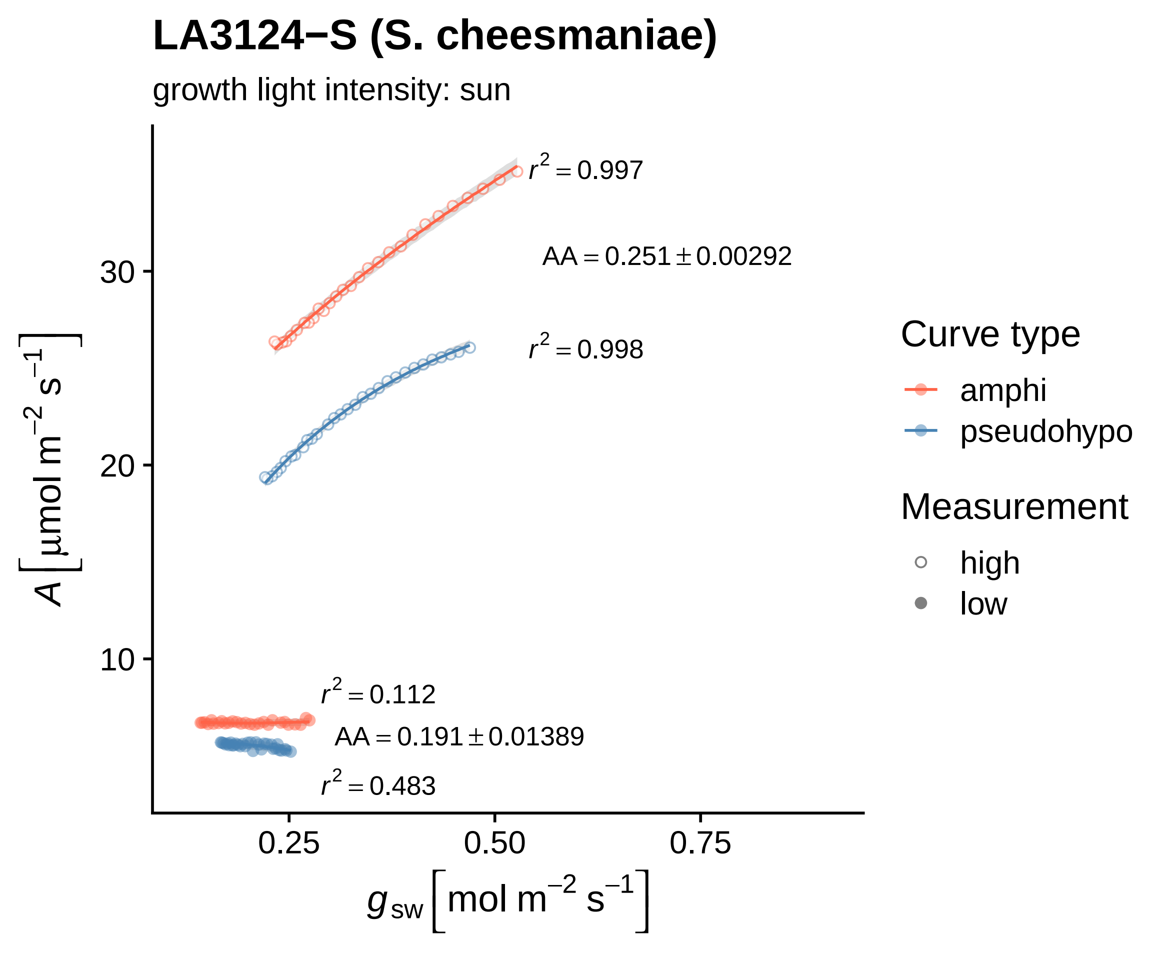

LA3124−S (S. cheesmaniae)
growth light intensity: sun
$r^2 = 0.997$
AA = 0.251 ± 0.00292
$r^2 = 0.998$
$r^2 = 0.112$
AA = 0.191 ± 0.01389
$r^2 = 0.483$
30
20
10
0.25
0.50
0.75
$A\left[\mu mol\ m^{-2}\ s^{-1}\right]$
$g_{sw}\left[mol\ m^{-2}\ s^{-1}\right]$
Curve type
amphi
pseudohypo
Measurement
high
low

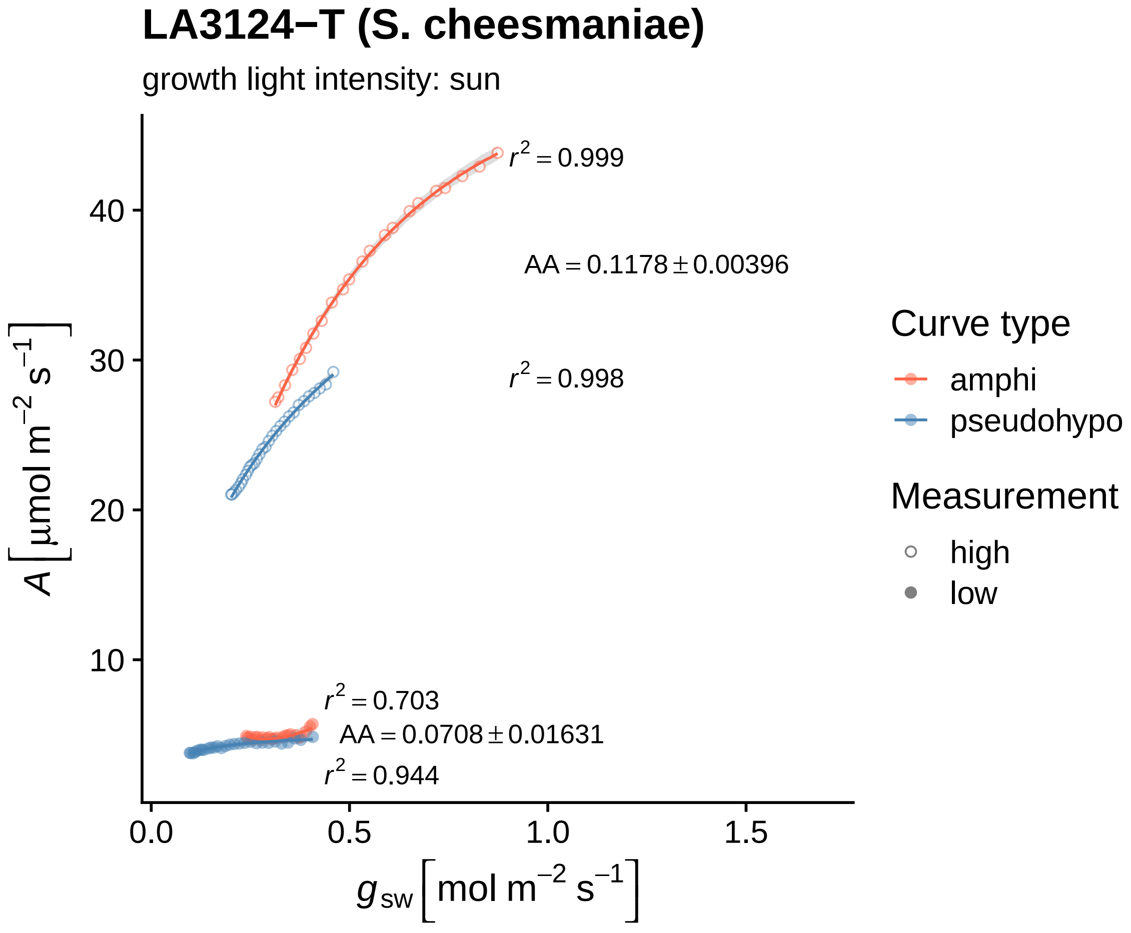

LA3124−T (S. cheesmaniae)
growth light intensity: sun
$r^2 = 0.999$
AA = 0.1178 ± 0.00396
$r^2 = 0.998$
$r^2 = 0.703$
AA = 0.0708 ± 0.01631
$r^2 = 0.944$
40
30
20
10
0.0
0.5
1.0
1.5
$A\ [\mu mol\ m^{-2}\ s^{-1}]$
$g_{sw}\ [mol\ m^{-2}\ s^{-1}]$
Curve type
amphi
pseudohypo
Measurement
high
low

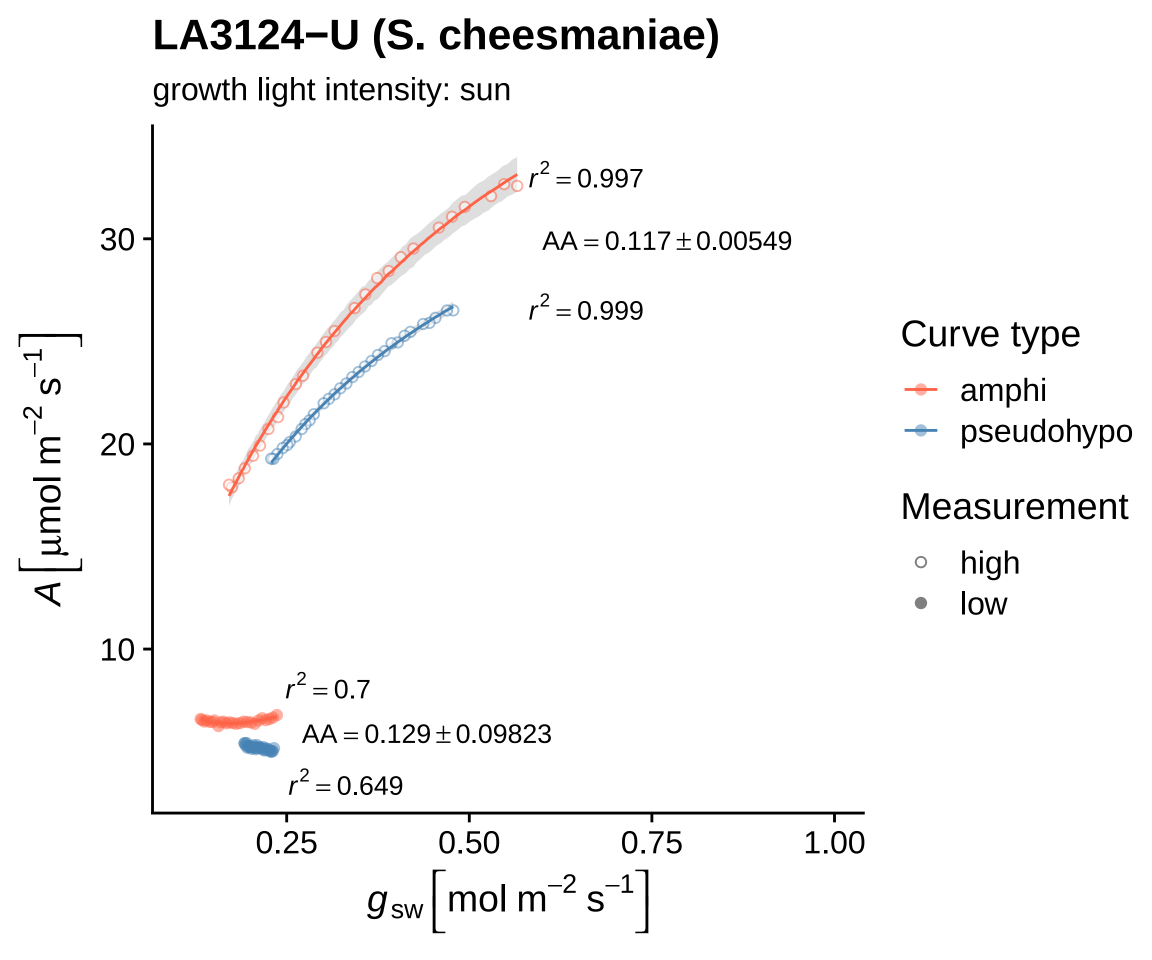

LA3124−U (S. cheesmaniae)
growth light intensity: sun
$r^2 = 0.997$
$AA = 0.117 \pm 0.00549$
$r^2 = 0.999$
$r^2 = 0.7$
$AA = 0.129 \pm 0.09823$
$r^2 = 0.649$
30
20
10
0.25
0.50
0.75
1.00
$A\ [\mu mol\ m^{-2}\ s^{-1}]$
$g_{sw}\ [mol\ m^{-2}\ s^{-1}]$
Curve type
amphi
pseudohypo
Measurement
high
low

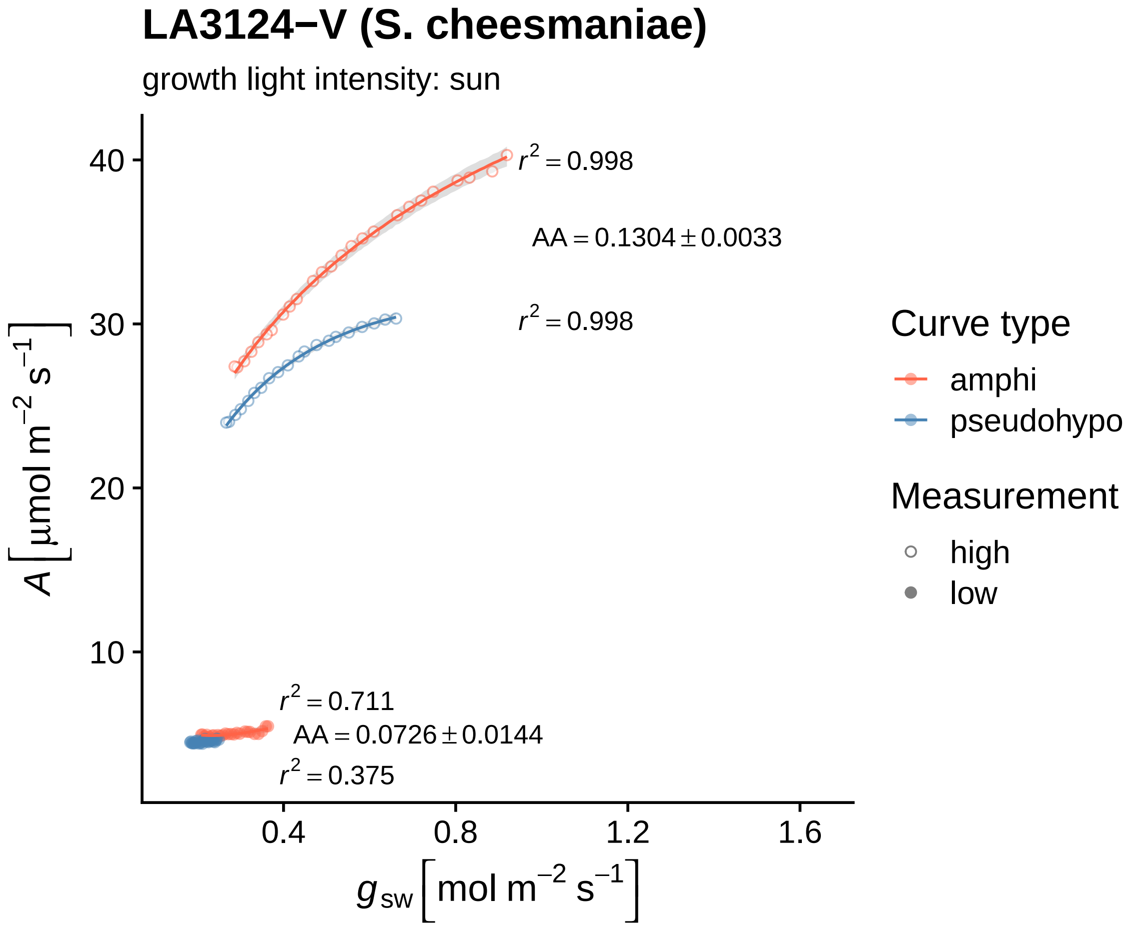

LA3124−V (S. cheesmaniae)
growth light intensity: sun
$r^2 = 0.998$
$AA = 0.1304 \pm 0.0033$
$r^2 = 0.998$
$r^2 = 0.711$
$AA = 0.0726 \pm 0.0144$
$r^2 = 0.375$
40
30
20
10
0.4
0.8
1.2
1.6
$A\left[\mu mol\ m^{-2}\ s^{-1}\right]$
$g_{sw}\left[mol\ m^{-2}\ s^{-1}\right]$
Curve type
amphi
pseudohypo
Measurement
high
low

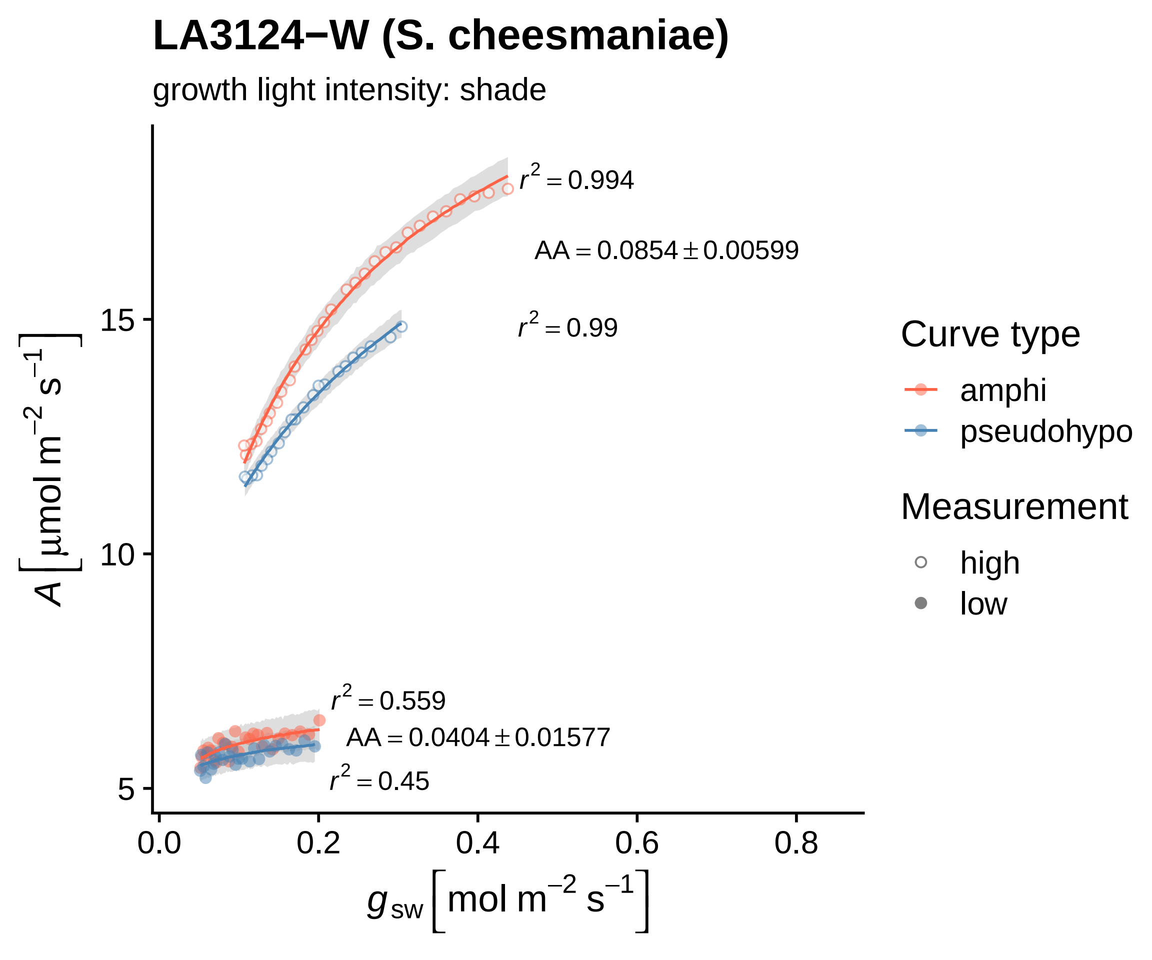

LA3124−W (S. cheesmaniae)
growth light intensity: shade
$r^2 = 0.994$
$AA = 0.0854 \pm 0.00599$
$r^2 = 0.99$
$r^2 = 0.559$
$AA = 0.0404 \pm 0.01577$
$r^2 = 0.45$
$A\ [\mu mol\ m^{-2}\ s^{-1}]$
15
10
5
0.0
0.2
0.4
0.6
0.8
$g_{sw}\ [mol\ m^{-2}\ s^{-1}]$
Curve type
amphi
pseudohypo
Measurement
high
low

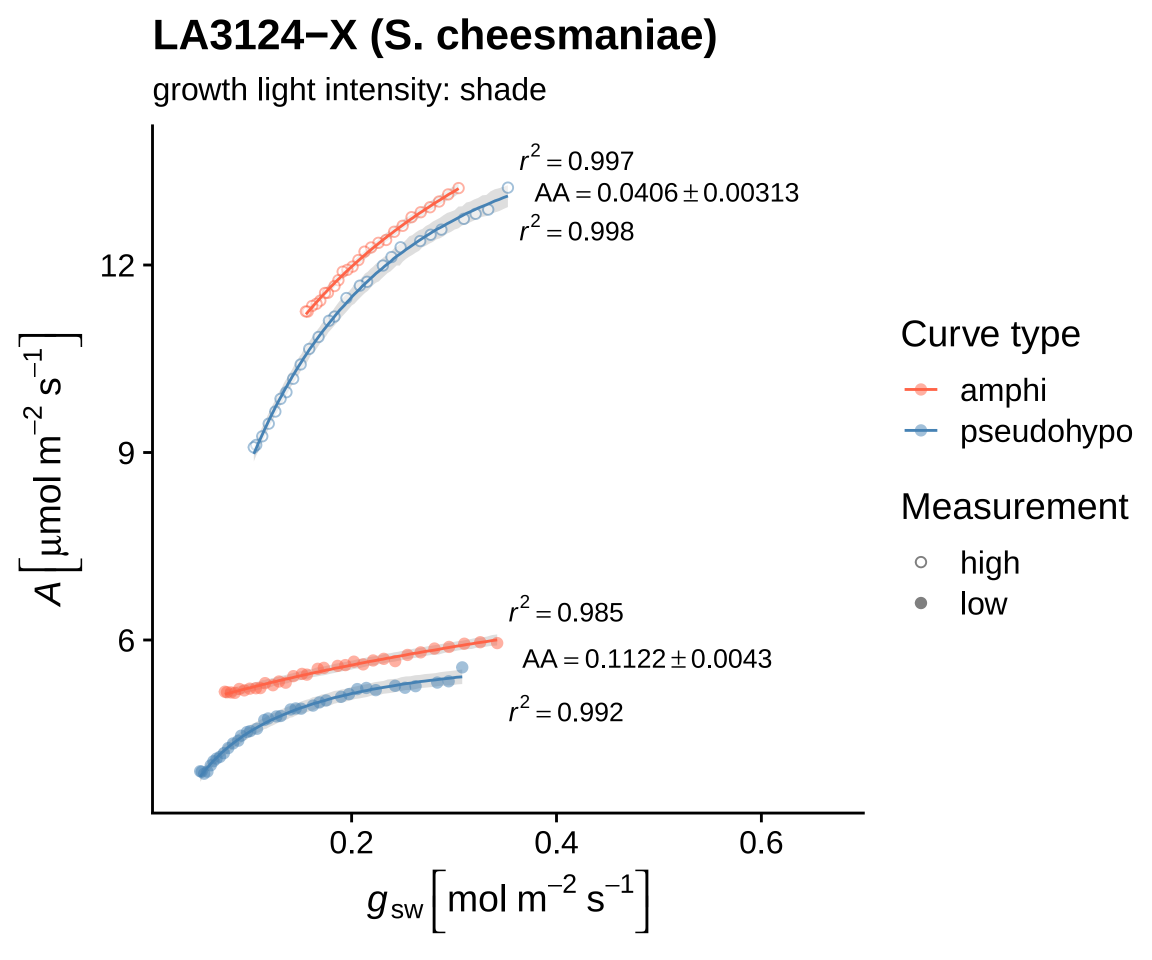
LA3124−X (S. cheesmaniae)
growth light intensity: shade
$r^2 = 0.997$
$AA = 0.0406 \pm 0.00313$
$r^2 = 0.998$
$r^2 = 0.985$
$AA = 0.1122 \pm 0.0043$
$r^2 = 0.992$
12
9
6
0.2
0.4
0.6
$A\left[\mu mol\ m^{-2}\ s^{-1}\right]$
$g_{sw}\left[mol\ m^{-2}\ s^{-1}\right]$
Curve type
amphi
pseudohypo
Measurement
high
low

#### LA3124−Y (S. cheesmaniae)

growth light intensity: sun

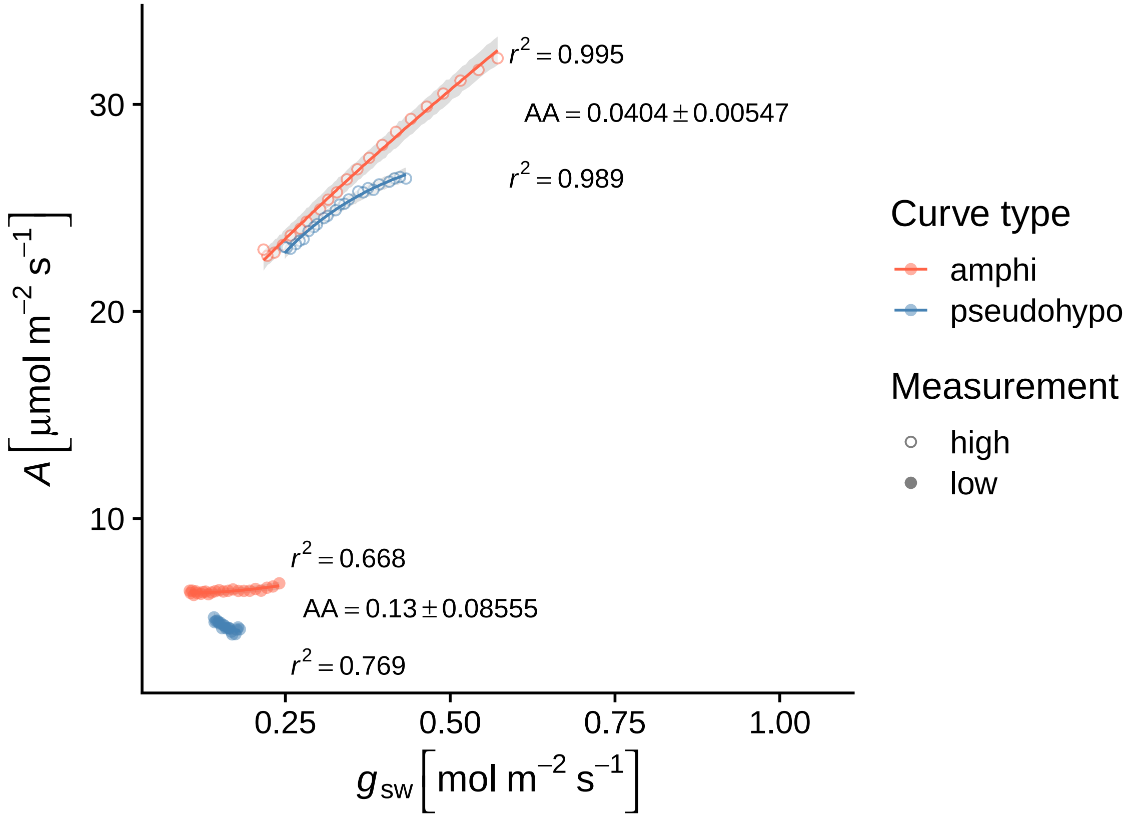

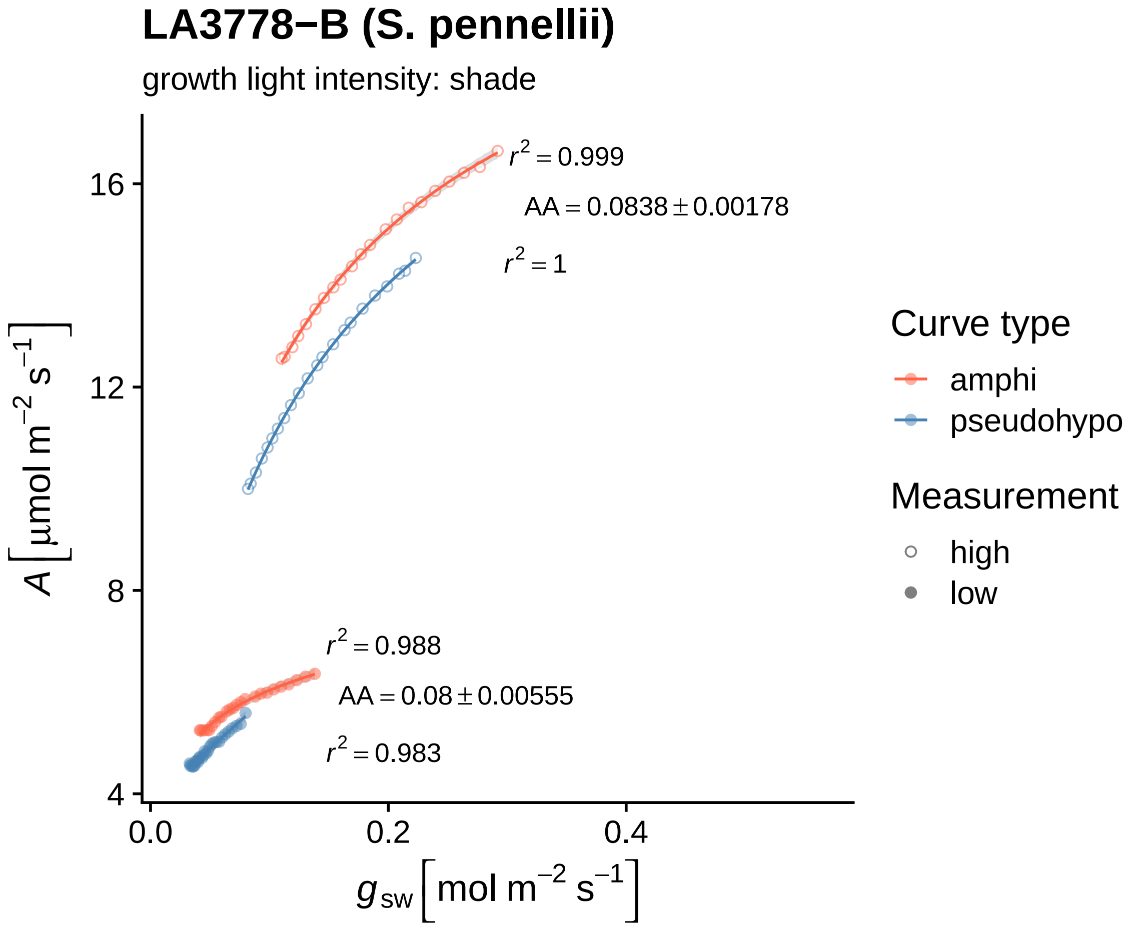
LA3778−B (S. pennellii)
growth light intensity: shade
$r^2 = 0.999$
$AA = 0.0838 \pm 0.00178$
$r^2 = 1$
$r^2 = 0.988$
$AA = 0.08 \pm 0.00555$
$r^2 = 0.983$
16
12
8
4
0.0
0.2
0.4
$A\left[\mu mol\ m^{-2}\ s^{-1}\right]$
$g_{sw}\left[mol\ m^{-2}\ s^{-1}\right]$
Curve type
amphi
pseudohypo
Measurement
high
low

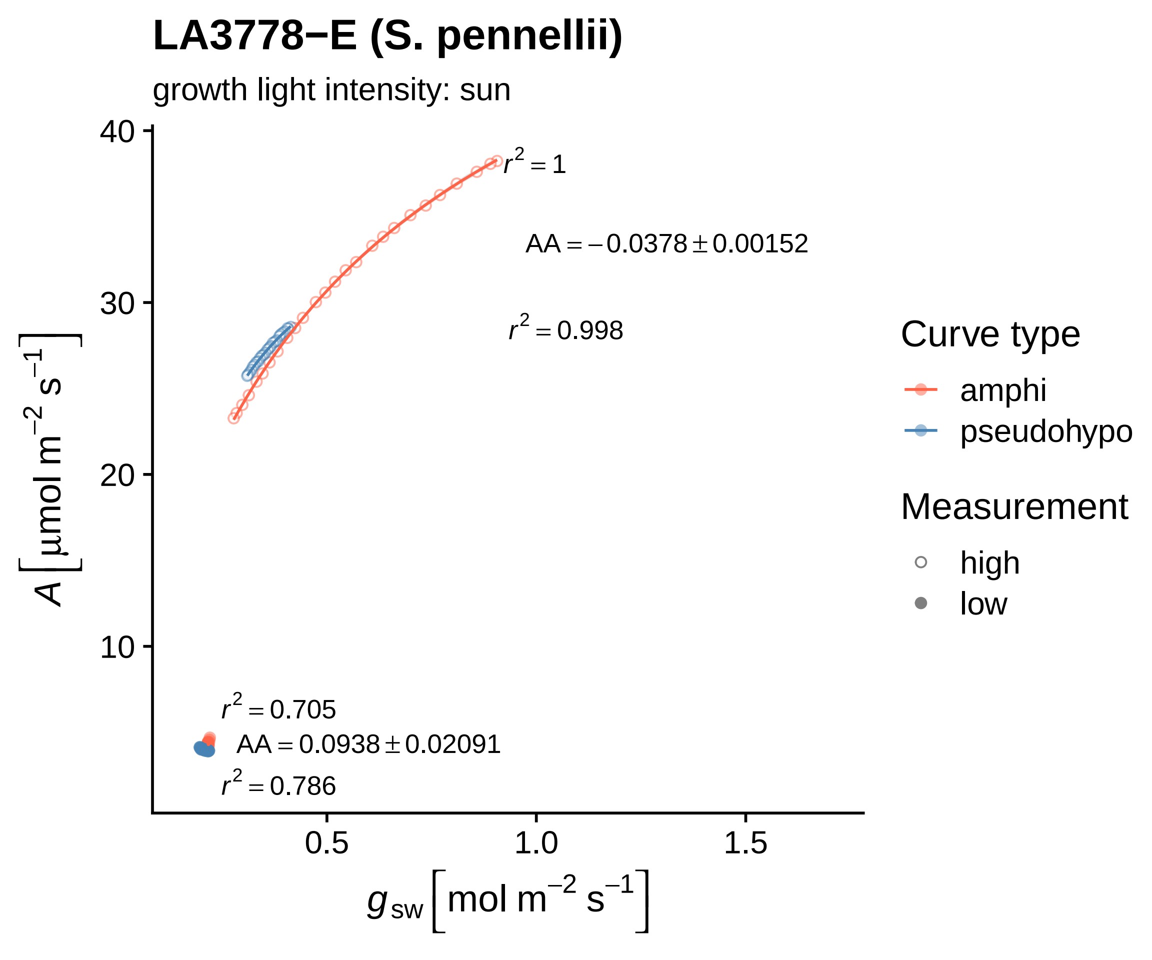
LA3778−E (S. pennellii)
growth light intensity: sun
40
30
20
10
0.5
1.0
1.5
$r^2 = 1$
$AA = -0.0378 \pm 0.00152$
$r^2 = 0.998$
$r^2 = 0.705$
$AA = 0.0938 \pm 0.02091$
$r^2 = 0.786$
$A\left[\mu mol\ m^{-2}\ s^{-1}\right]$
$g_{sw}\left[mol\ m^{-2}\ s^{-1}\right]$
Curve type
amphi
pseudohypo
Measurement
high
low

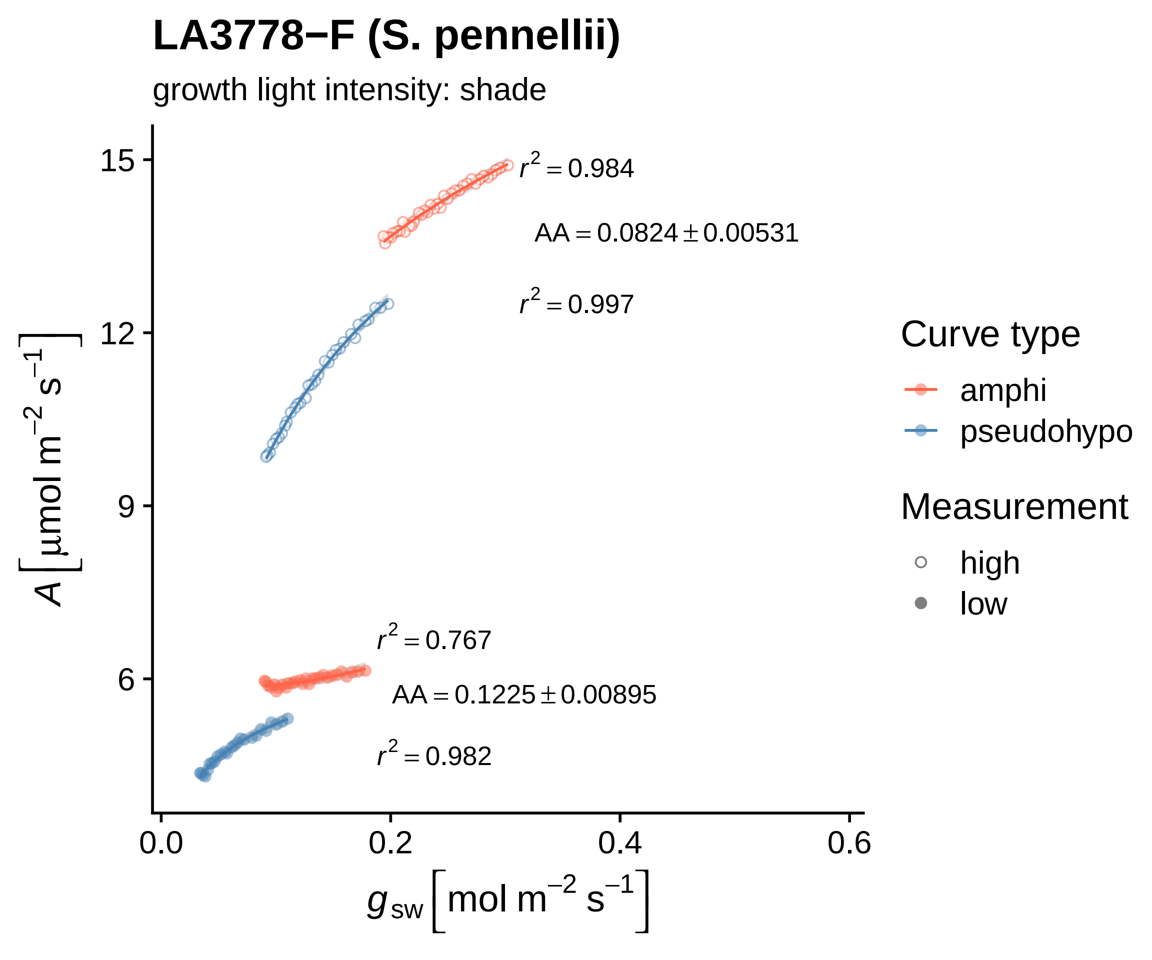

LA3778−F (S. pennellii)
growth light intensity: shade
$r^2 = 0.984$
$AA = 0.0824 \pm 0.00531$
$r^2 = 0.997$
$r^2 = 0.767$
$AA = 0.1225 \pm 0.00895$
$r^2 = 0.982$
15
12
9
6
0.0
0.2
0.4
0.6
$A\ [\mu mol\ m^{-2}\ s^{-1}]$
$g_{sw}\ [mol\ m^{-2}\ s^{-1}]$
Curve type
amphi
pseudohypo
Measurement
high
low

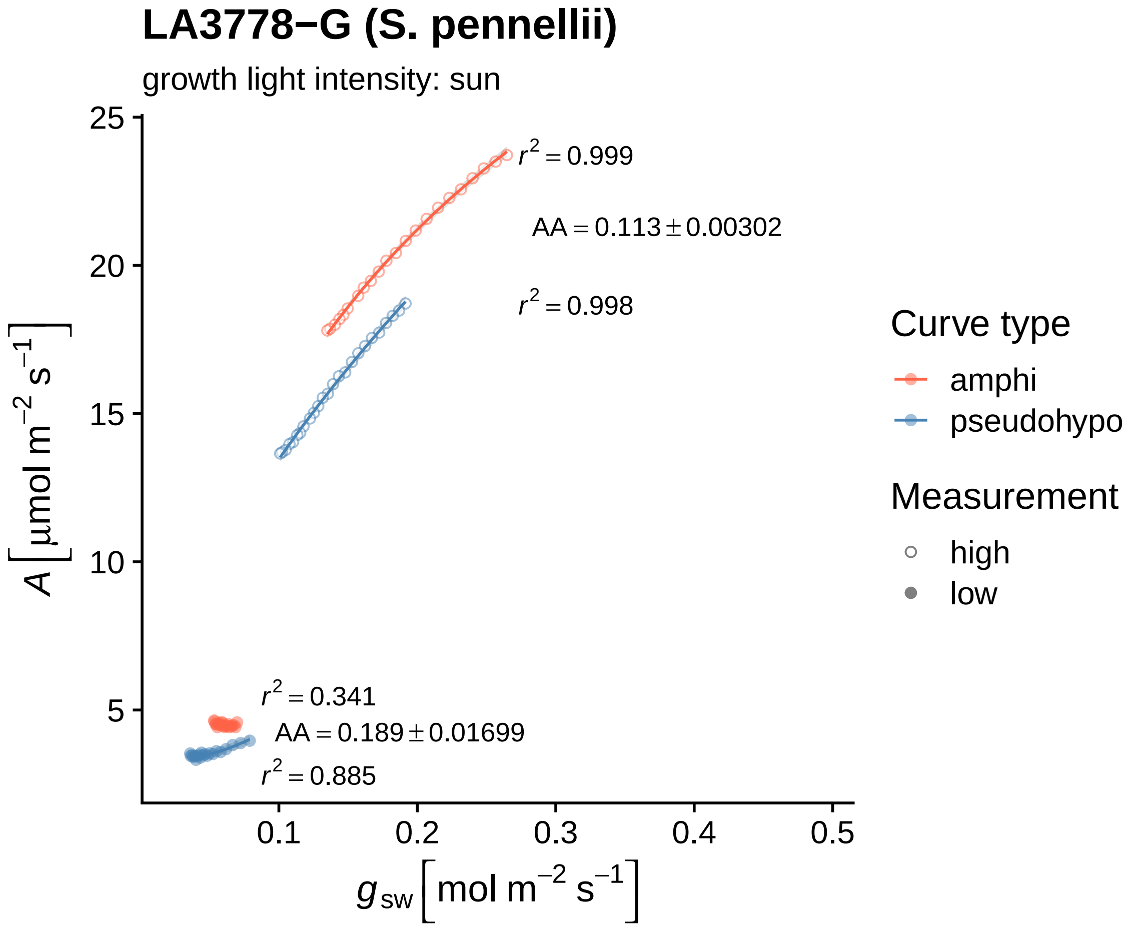

LA3778−G (S. pennellii)
growth light intensity: sun
$r^2 = 0.999$
$AA = 0.113 \pm 0.00302$
$r^2 = 0.998$
$r^2 = 0.341$
$AA = 0.189 \pm 0.01699$
$r^2 = 0.885$
25
20
15
10
5
0.1
0.2
0.3
0.4
0.5
$A\ [\mu mol\ m^{-2}\ s^{-1}]$
$g_{sw}\ [mol\ m^{-2}\ s^{-1}]$
Curve type
amphi
pseudohypo
Measurement
high
low

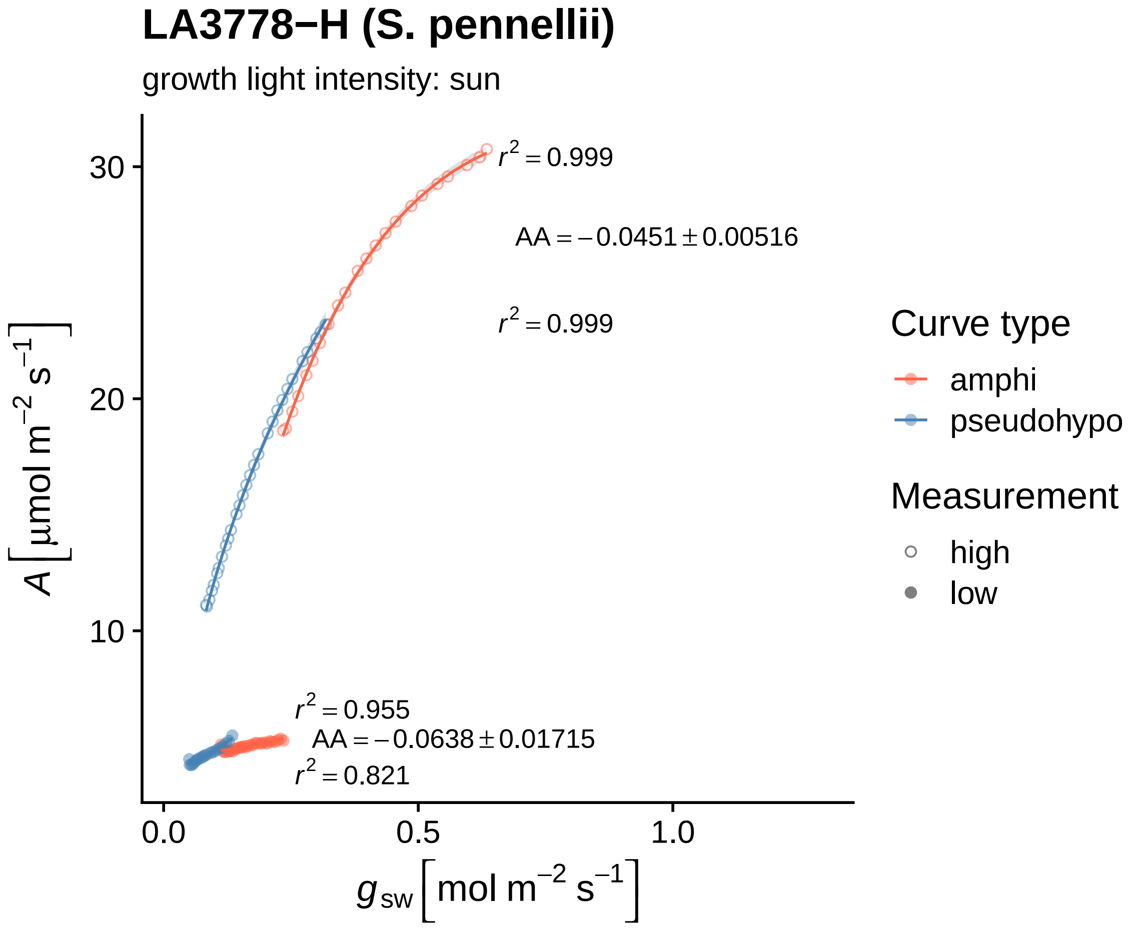

LA3778−H (S. pennellii)
growth light intensity: sun
$r^2 = 0.999$
AA = −0.0451 ± 0.00516
$r^2 = 0.999$
$r^2 = 0.955$
AA = −0.0638 ± 0.01715
$r^2 = 0.821$
30
20
10
0.0
0.5
1.0
$A\ [\mu mol\ m^{-2}\ s^{-1}]$
$g_{sw}\ [mol\ m^{-2}\ s^{-1}]$
Curve type
amphi
pseudohypo
Measurement
high
low

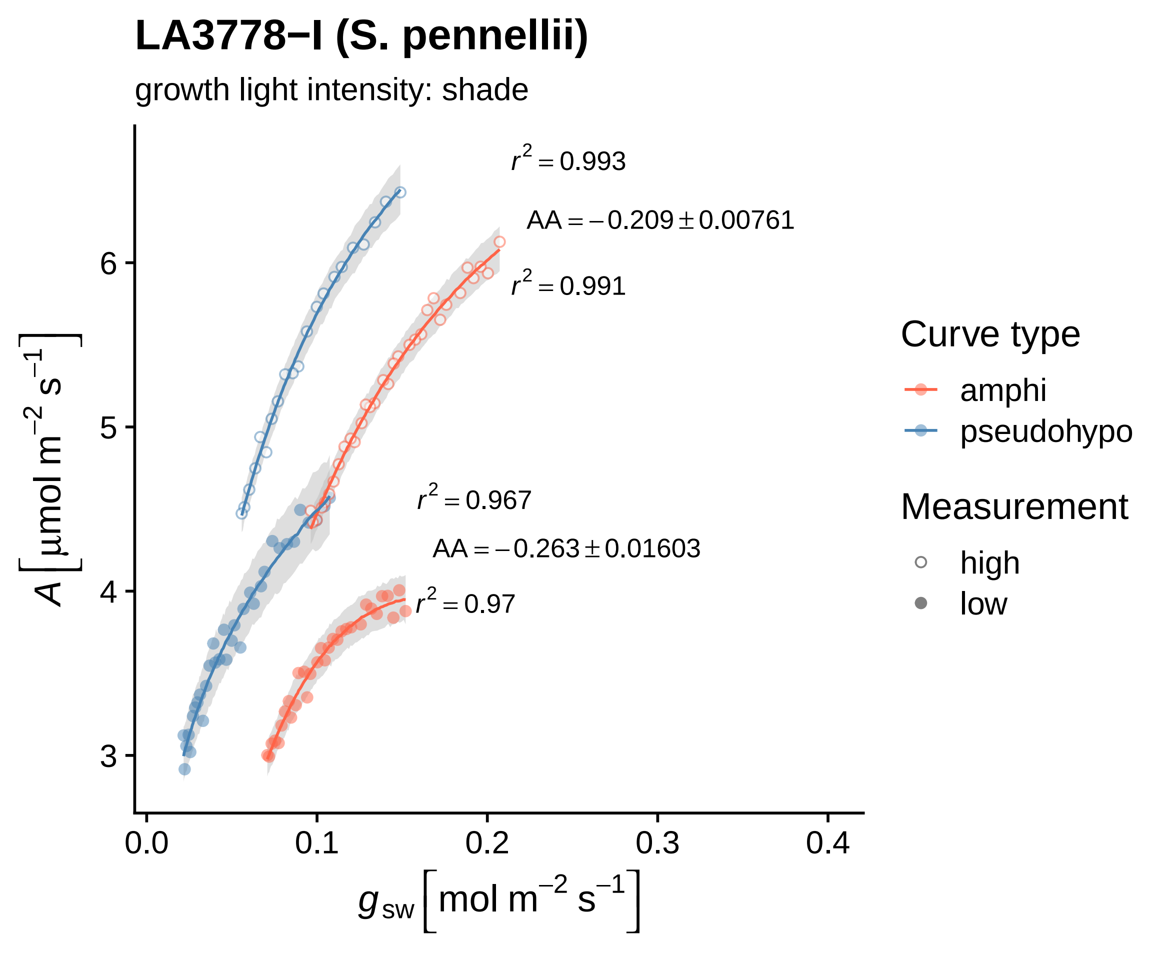
LA3778−I (S. pennellii)
growth light intensity: shade
$r^2 = 0.993$
$AA = -0.209 \pm 0.00761$
$r^2 = 0.991$
$r^2 = 0.967$
$AA = -0.263 \pm 0.01603$
$r^2 = 0.97$
Curve type
amphi
pseudohypo
Measurement
high
low
$A\ [\mu mol\ m^{-2}\ s^{-1}]$
$g_{sw}\ [mol\ m^{-2}\ s^{-1}]$
3
4
5
6
0.0
0.1
0.2
0.3
0.4

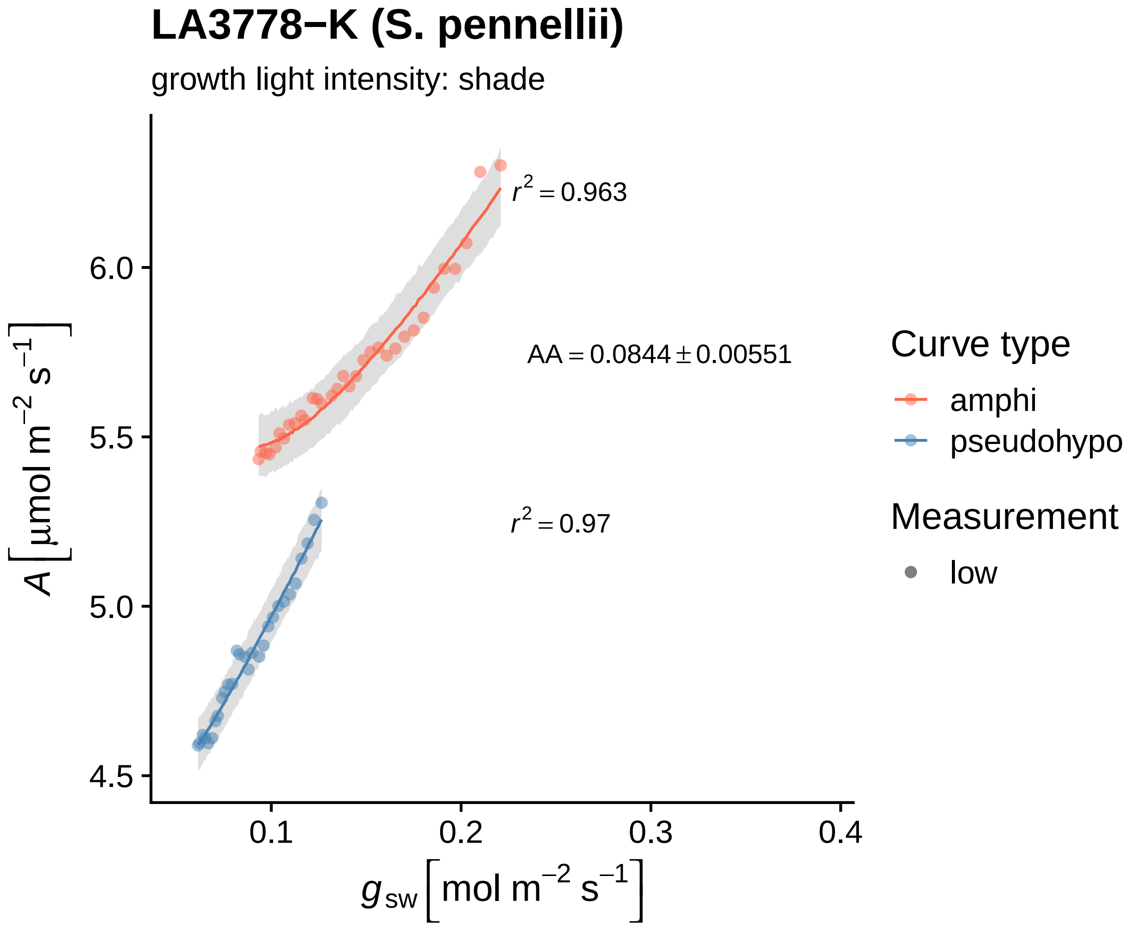

LA3778−K (S. pennellii)
growth light intensity: shade
r² = 0.963
AA = 0.0844 ± 0.00551
r² = 0.97
6.0
5.5
5.0
4.5
0.1
0.2
0.3
0.4
A [μmol m⁻² s⁻¹]
g_sw [mol m⁻² s⁻¹]
Curve type
amphi
pseudohypo
Measurement
low

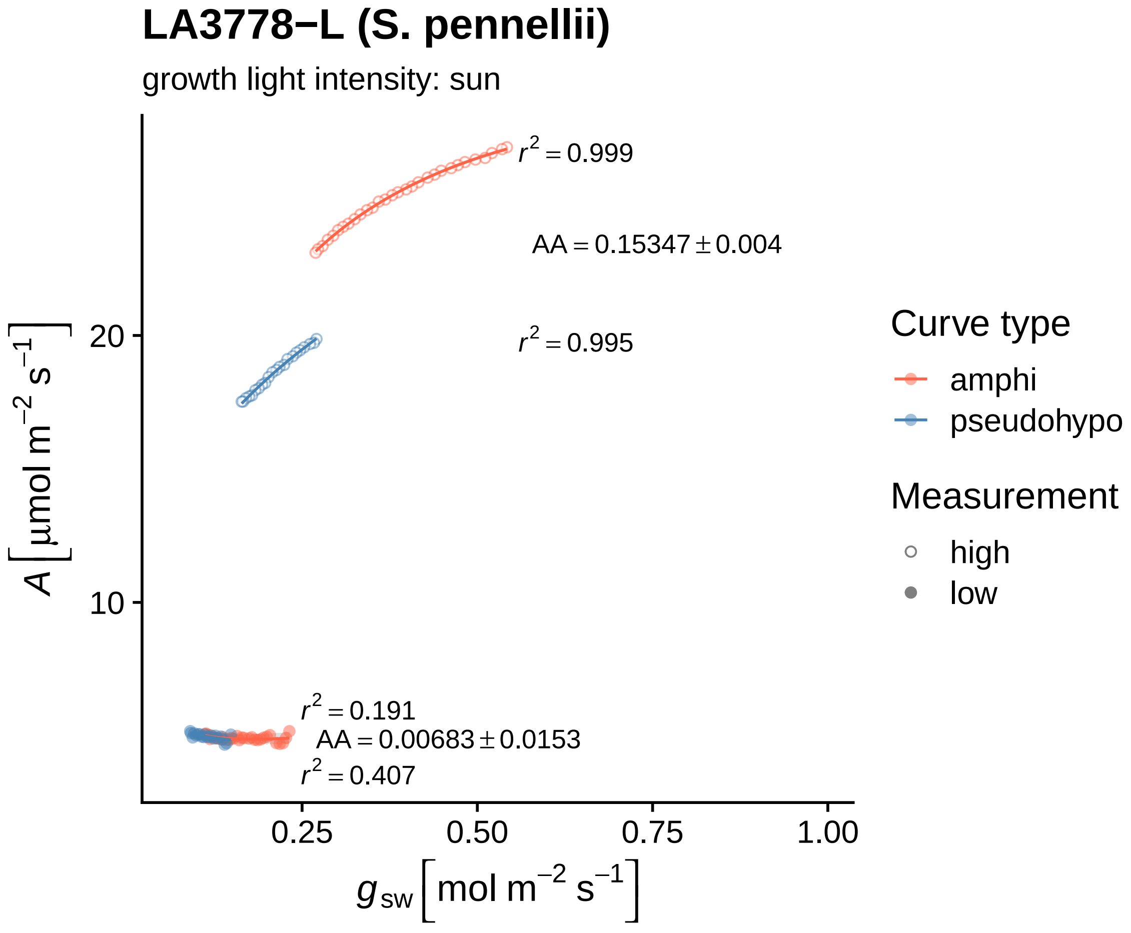

LA3778−L (S. pennellii)
growth light intensity: sun
$r^2 = 0.999$
$AA = 0.15347 \pm 0.004$
$r^2 = 0.995$
$r^2 = 0.191$
$AA = 0.00683 \pm 0.0153$
$r^2 = 0.407$
20
10
0.25
0.50
0.75
1.00
$A\left[\mu mol\ m^{-2}\ s^{-1}\right]$
$g_{sw}\left[mol\ m^{-2}\ s^{-1}\right]$
Curve type
amphi
pseudohypo
Measurement
high
low

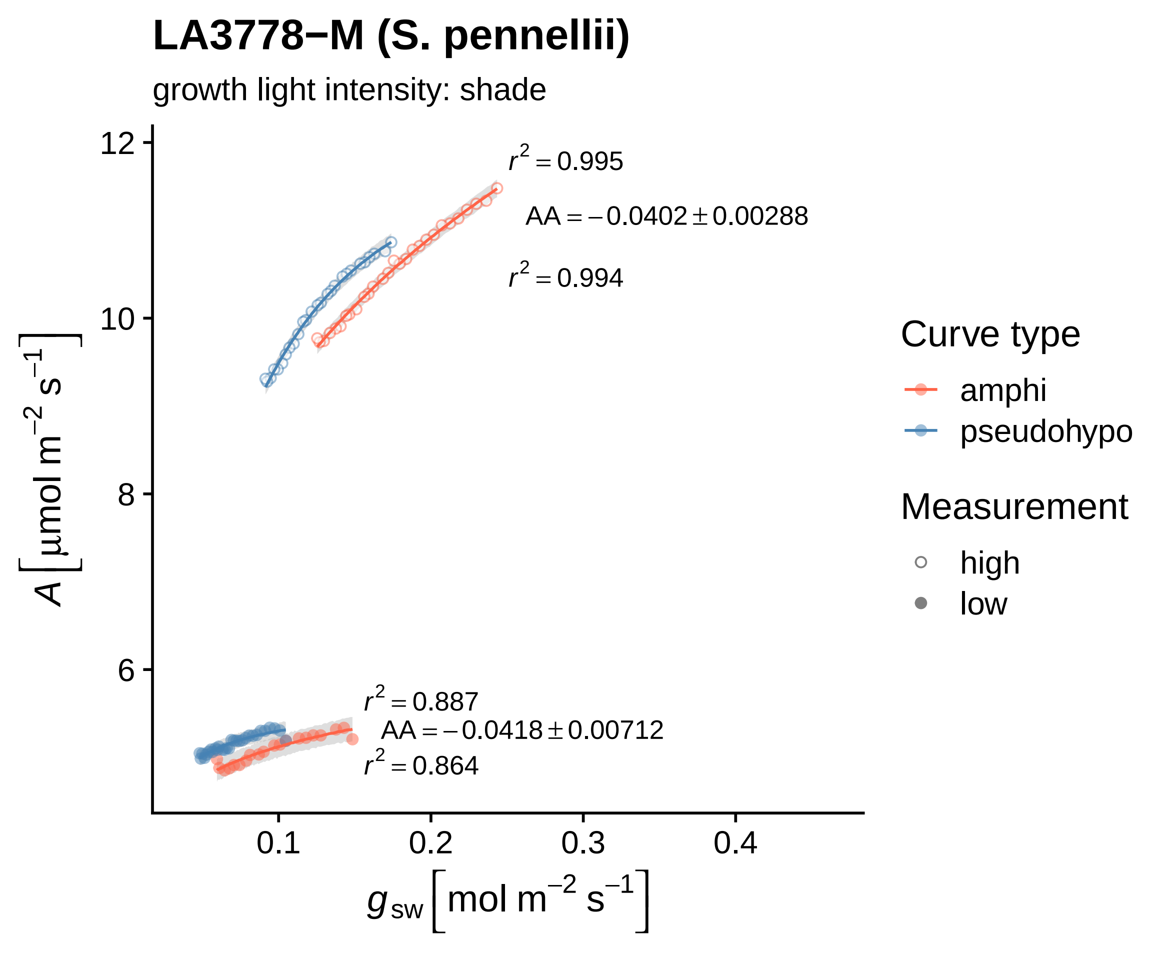

LA3778−M (S. pennellii)
growth light intensity: shade
$r^2 = 0.995$
$AA = -0.0402 \pm 0.00288$
$r^2 = 0.994$
$r^2 = 0.887$
$AA = -0.0418 \pm 0.00712$
$r^2 = 0.864$
12
10
8
6
0.1
0.2
0.3
0.4
$A\ [\mu mol\ m^{-2}\ s^{-1}]$
$g_{sw}\ [mol\ m^{-2}\ s^{-1}]$
Curve type
amphi
pseudohypo
Measurement
high
low

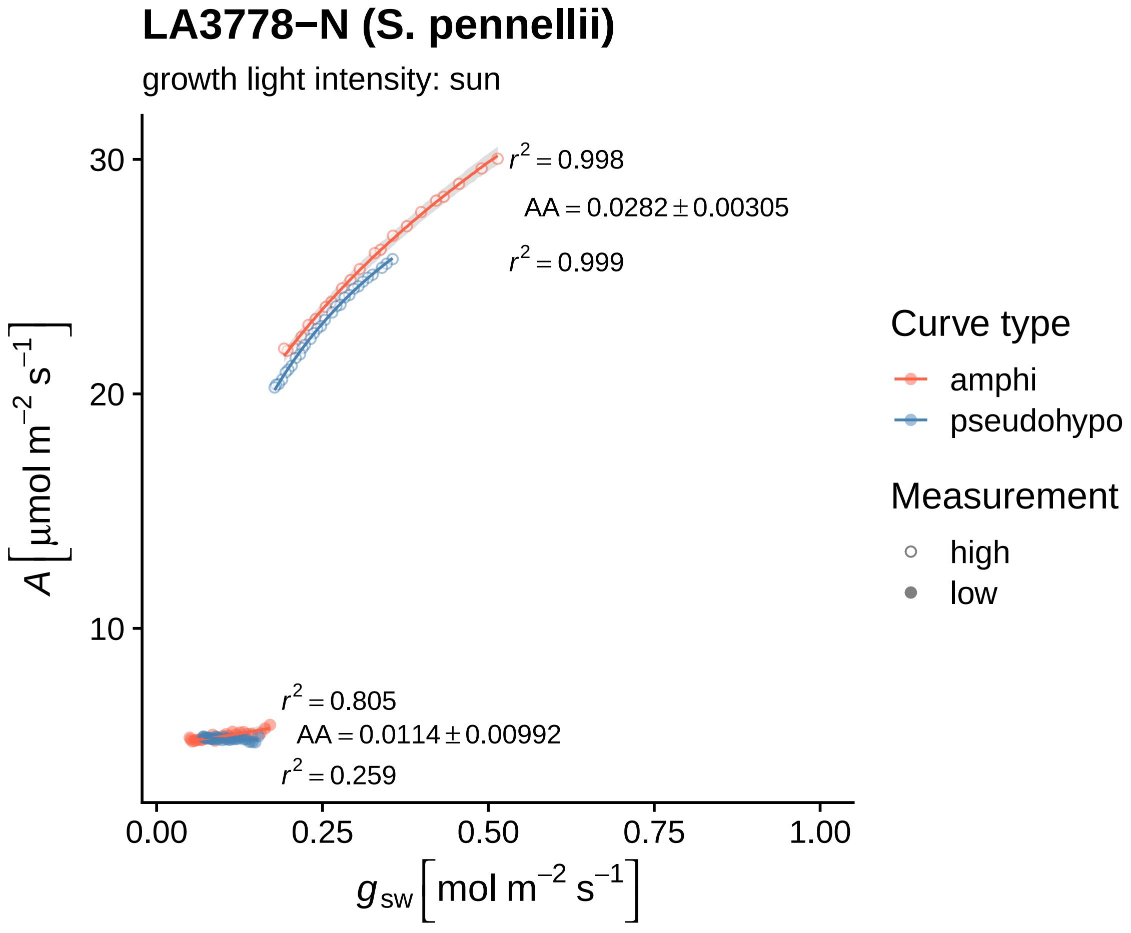
LA3778−N (S. pennellii)
growth light intensity: sun
$r^2 = 0.998$
$AA = 0.0282 \pm 0.00305$
$r^2 = 0.999$
$r^2 = 0.805$
$AA = 0.0114 \pm 0.00992$
$r^2 = 0.259$
30
20
10
0.00
0.25
0.50
0.75
1.00
$A\ [\mu mol\ m^{-2}\ s^{-1}]$
$g_{sw}\ [mol\ m^{-2}\ s^{-1}]$
Curve type
amphi
pseudohypo
Measurement
high
low

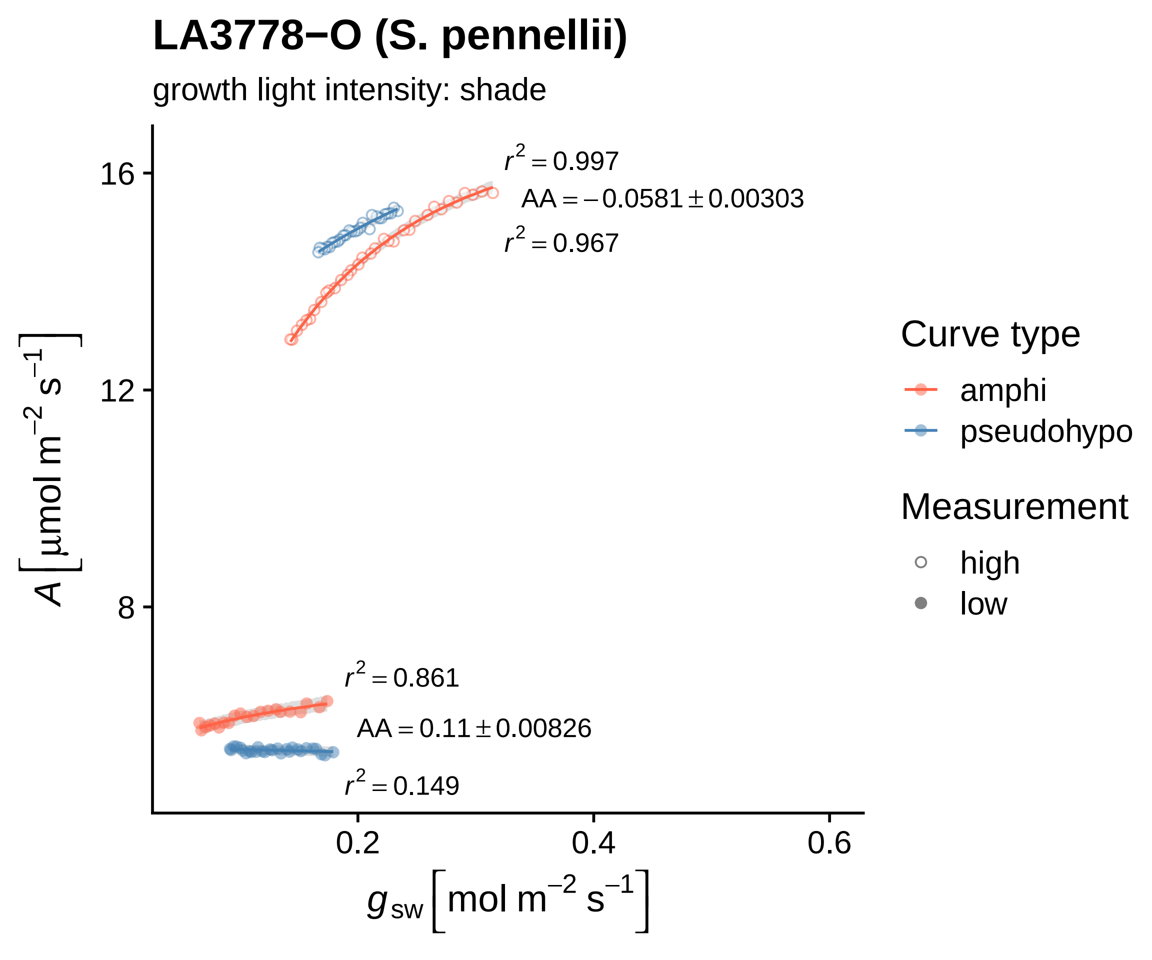

LA3778−O (S. pennellii)
growth light intensity: shade
$r^2 = 0.997$
$AA = -0.0581 \pm 0.00303$
$r^2 = 0.967$
$r^2 = 0.861$
$AA = 0.11 \pm 0.00826$
$r^2 = 0.149$
16
12
8
0.2
0.4
0.6
$A\ [\mu mol\ m^{-2}\ s^{-1}]$
$g_{sw}\ [mol\ m^{-2}\ s^{-1}]$
Curve type
amphi
pseudohypo
Measurement
high
low

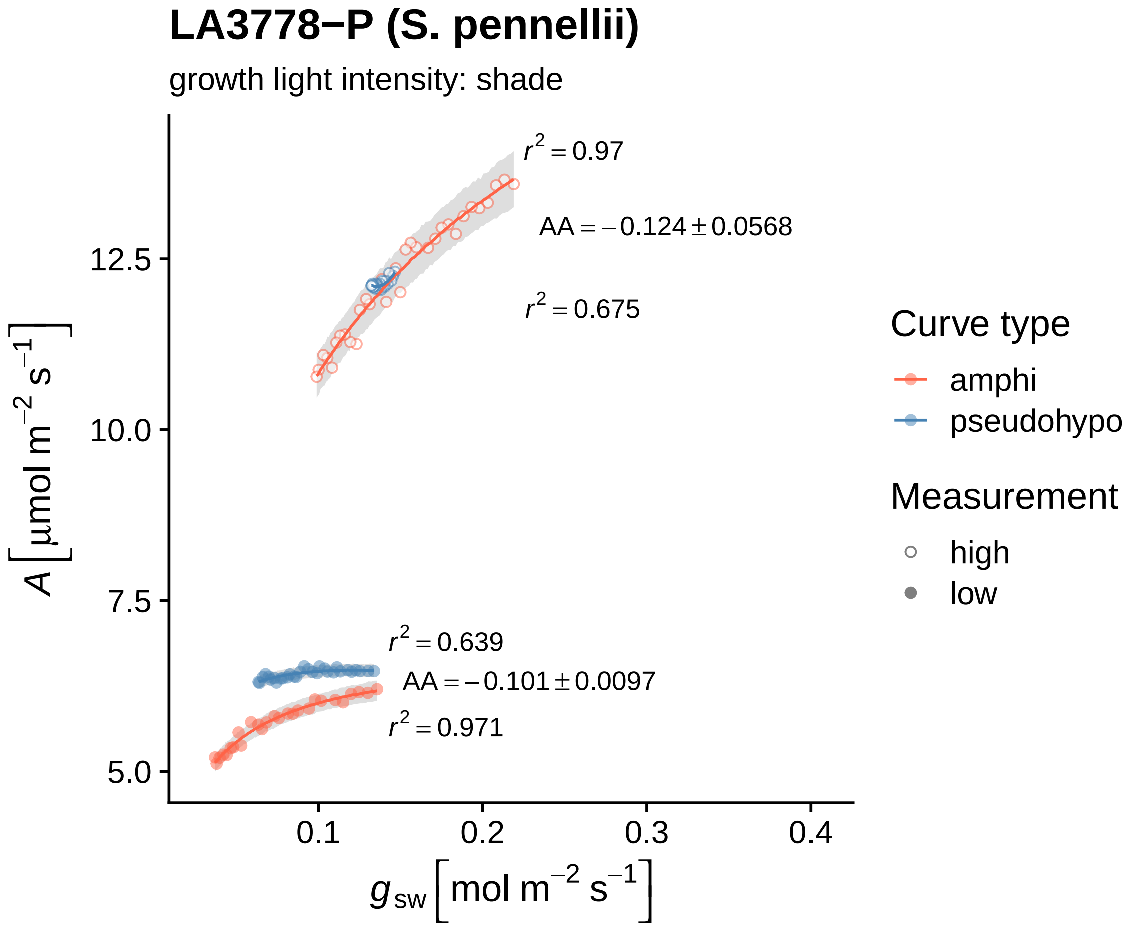
LA3778−P (S. pennellii)
growth light intensity: shade
$r^2 = 0.97$
$AA = -0.124 \pm 0.0568$
$r^2 = 0.675$
$r^2 = 0.639$
$AA = -0.101 \pm 0.0097$
$r^2 = 0.971$
12.5
10.0
7.5
5.0
0.1
0.2
0.3
0.4
$A\,[\mu mol\ m^{-2}\ s^{-1}]$
$g_{sw}\,[mol\ m^{-2}\ s^{-1}]$
Curve type
amphi
pseudohypo
Measurement
high
low

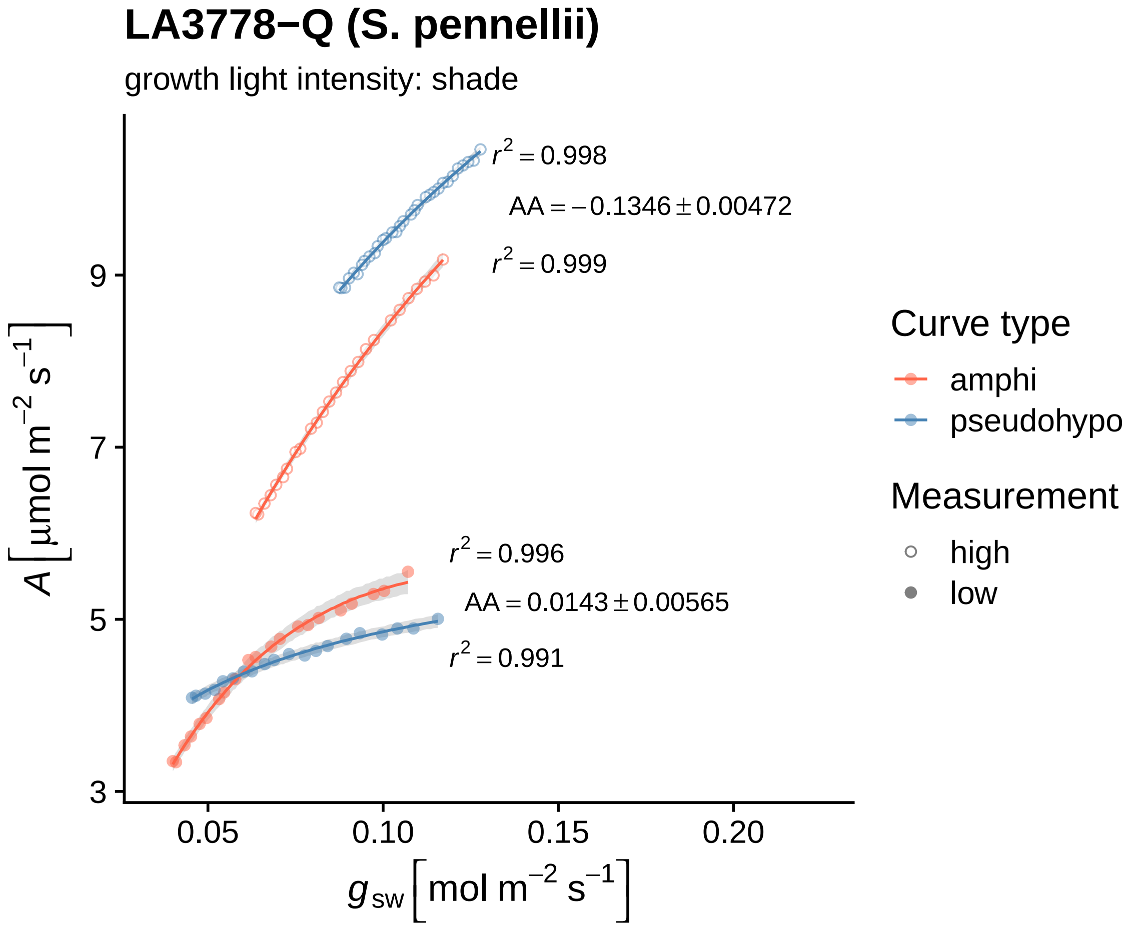
LA3778−Q (S. pennellii)
growth light intensity: shade
$r^2 = 0.998$
$AA = -0.1346 \pm 0.00472$
$r^2 = 0.999$
$r^2 = 0.996$
$AA = 0.0143 \pm 0.00565$
$r^2 = 0.991$
$A\,[\mu mol\ m^{-2}\ s^{-1}]$
$g_{sw}\,[mol\ m^{-2}\ s^{-1}]$
3
5
7
9
0.05
0.10
0.15
0.20
Curve type
amphi
pseudohypo
Measurement
high
low

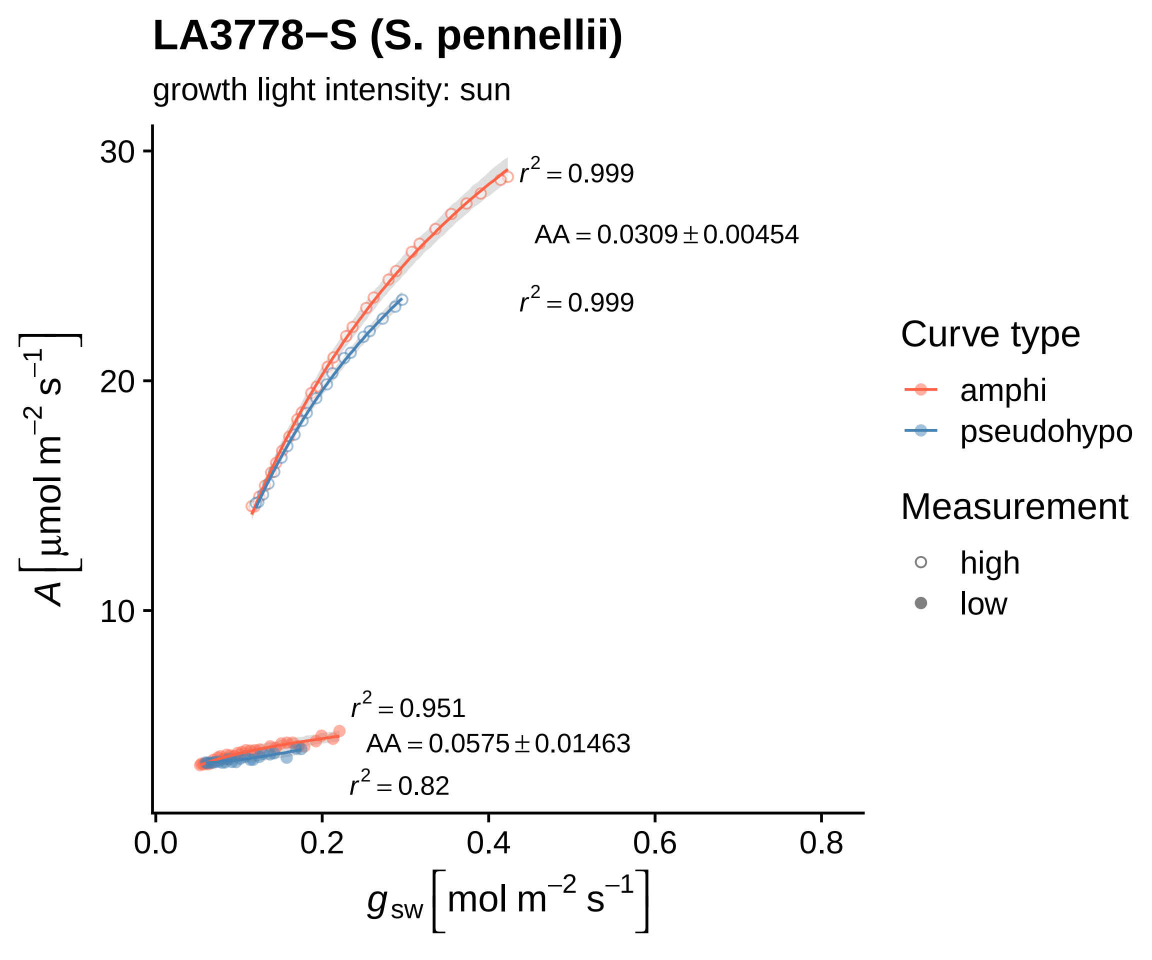

LA3778−S (S. pennellii)
growth light intensity: sun
$r^2 = 0.999$
$AA = 0.0309 \pm 0.00454$
$r^2 = 0.999$
$r^2 = 0.951$
$AA = 0.0575 \pm 0.01463$
$r^2 = 0.82$
30
20
10
0.0
0.2
0.4
0.6
0.8
$A\ [\mu mol\ m^{-2}\ s^{-1}]$
$g_{sw}\ [mol\ m^{-2}\ s^{-1}]$
Curve type
amphi
pseudohypo
Measurement
high
low

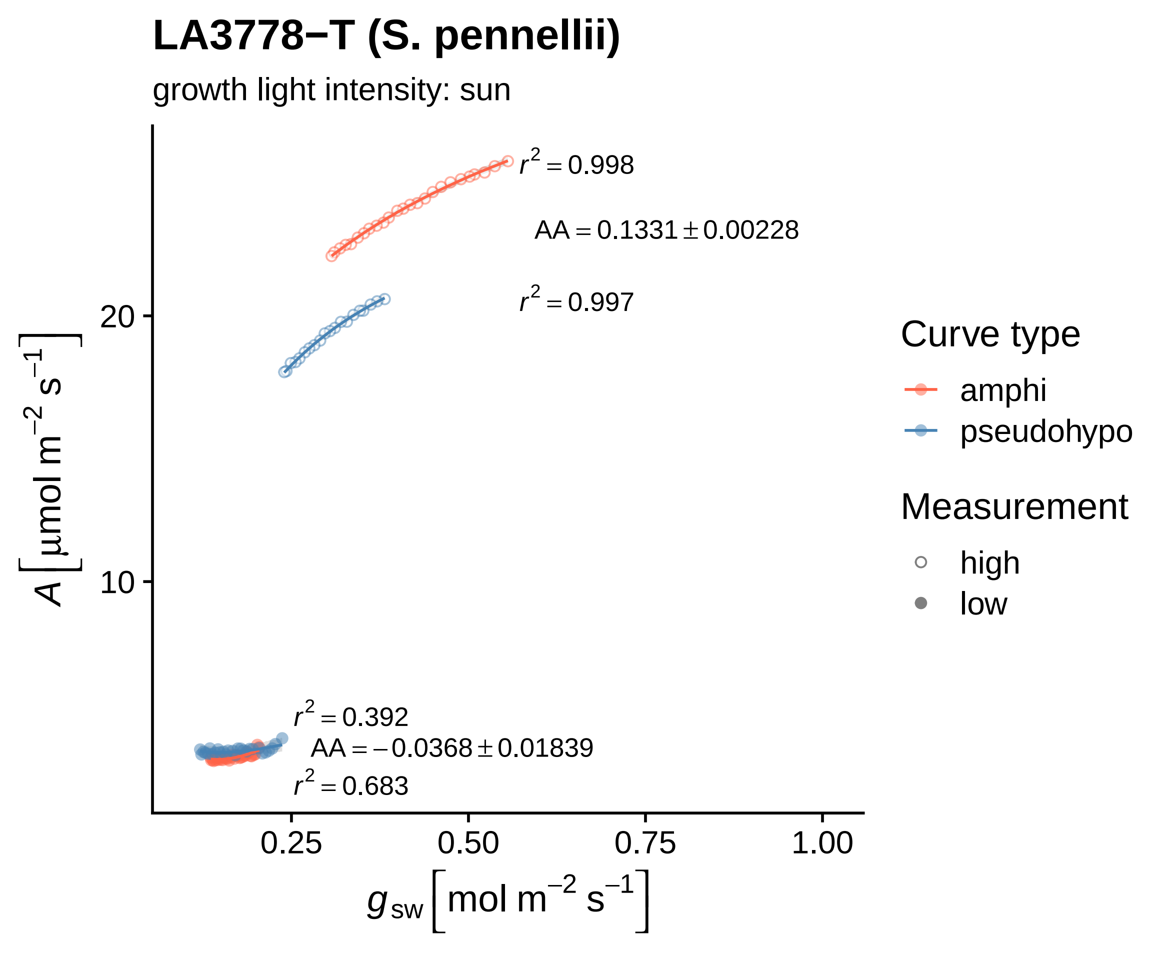

LA3778−T (S. pennellii)
growth light intensity: sun
r2 = 0.998
AA = 0.1331 ± 0.00228
r2 = 0.997
r2 = 0.392
AA = −0.0368 ± 0.01839
r2 = 0.683
20
10
0.25
0.50
0.75
1.00
A [µmol m−2 s−1]
gsw [mol m−2 s−1]
Curve type
amphi
pseudohypo
Measurement
high
low

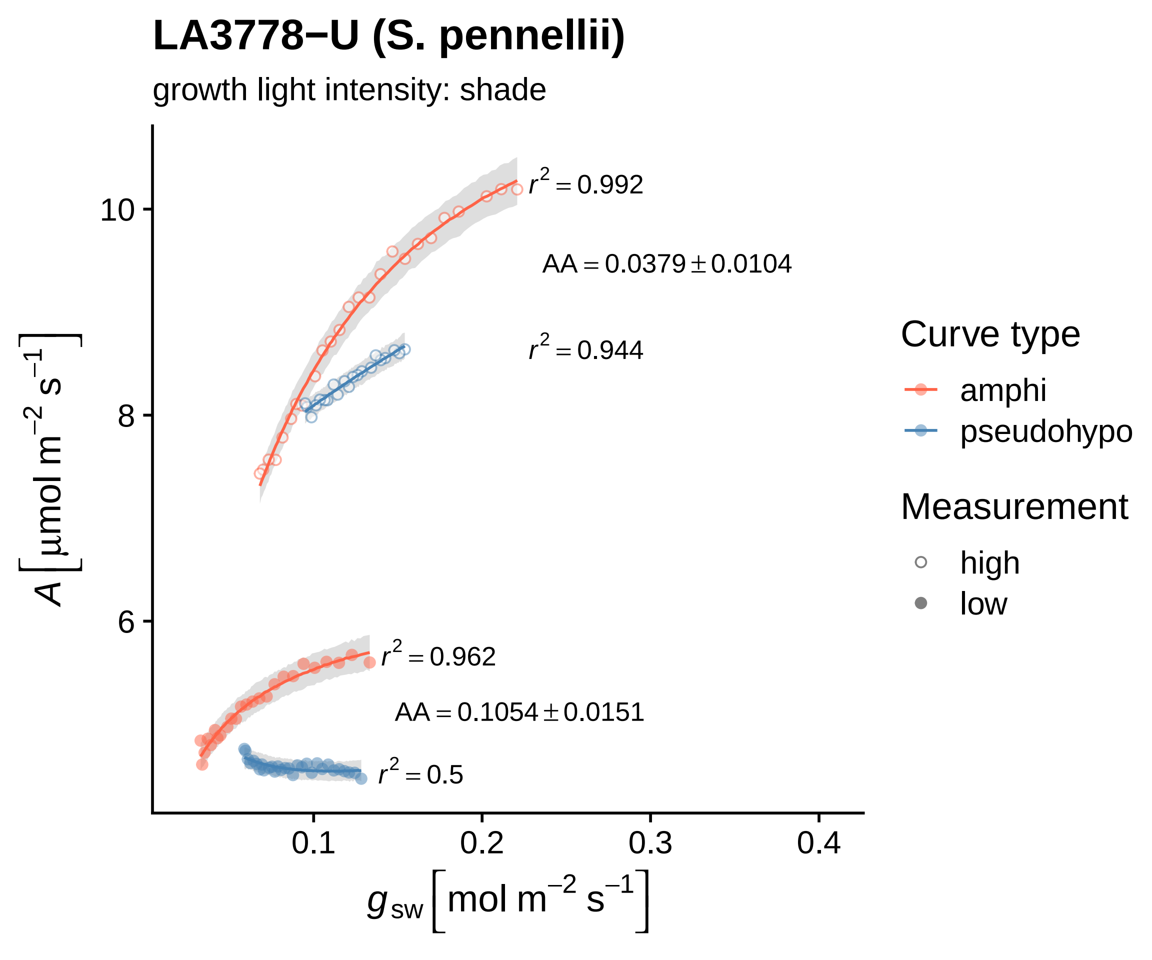

LA3778−U (S. pennellii)
growth light intensity: shade
$r^2 = 0.992$
$AA = 0.0379 \pm 0.0104$
$r^2 = 0.944$
$r^2 = 0.962$
$AA = 0.1054 \pm 0.0151$
$r^2 = 0.5$
10
8
6
0.1
0.2
0.3
0.4
$A\left[\mu mol\ m^{-2}\ s^{-1}\right]$
$g_{sw}\left[mol\ m^{-2}\ s^{-1}\right]$
Curve type
amphi
pseudohypo
Measurement
high
low

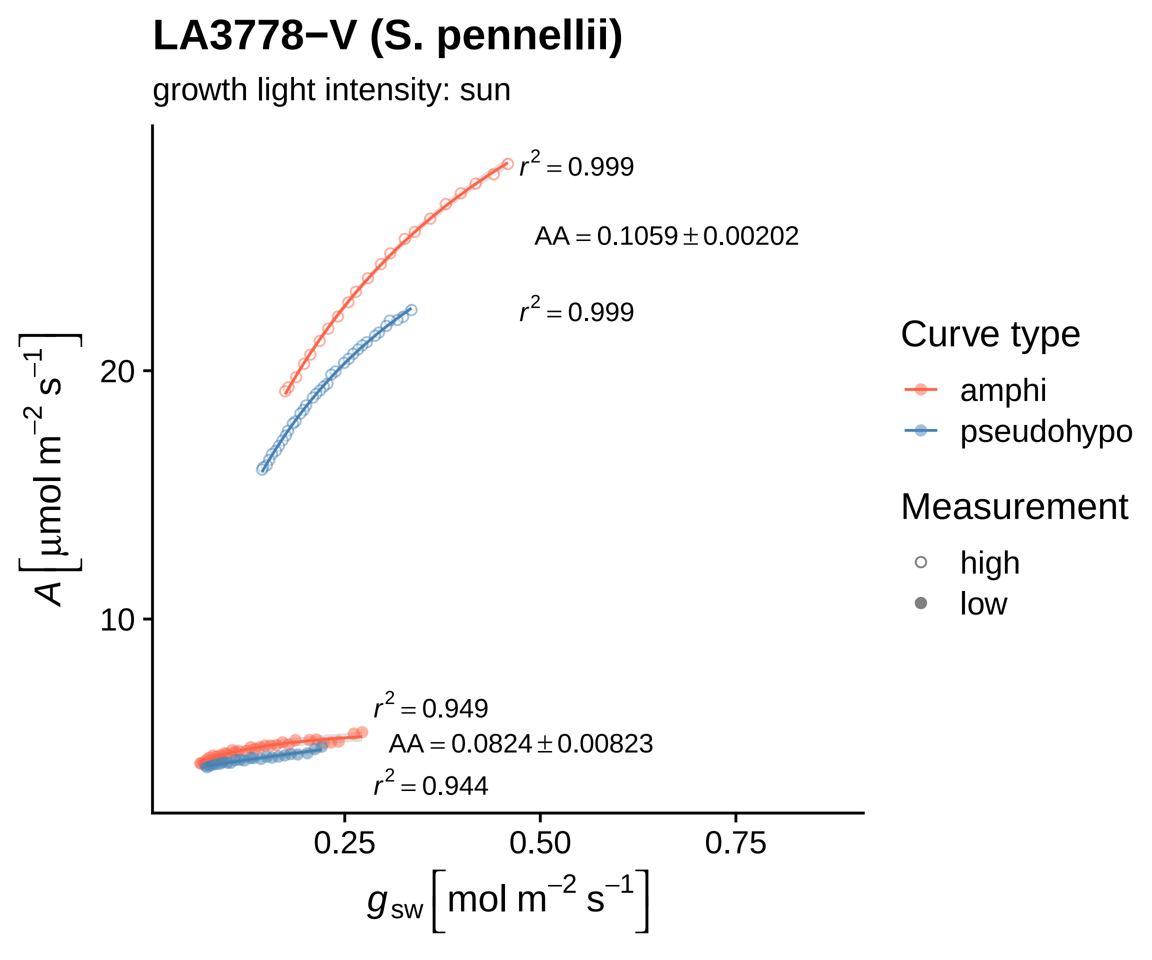

LA3778−V (S. pennellii)
growth light intensity: sun
$r^2 = 0.999$
$AA = 0.1059 \pm 0.00202$
$r^2 = 0.999$
$r^2 = 0.949$
$AA = 0.0824 \pm 0.00823$
$r^2 = 0.944$
20
10
0.25
0.50
0.75
$A\ [\mu mol\ m^{-2}\ s^{-1}]$
$g_{sw}\ [mol\ m^{-2}\ s^{-1}]$
Curve type
amphi
pseudohypo
Measurement
high
low

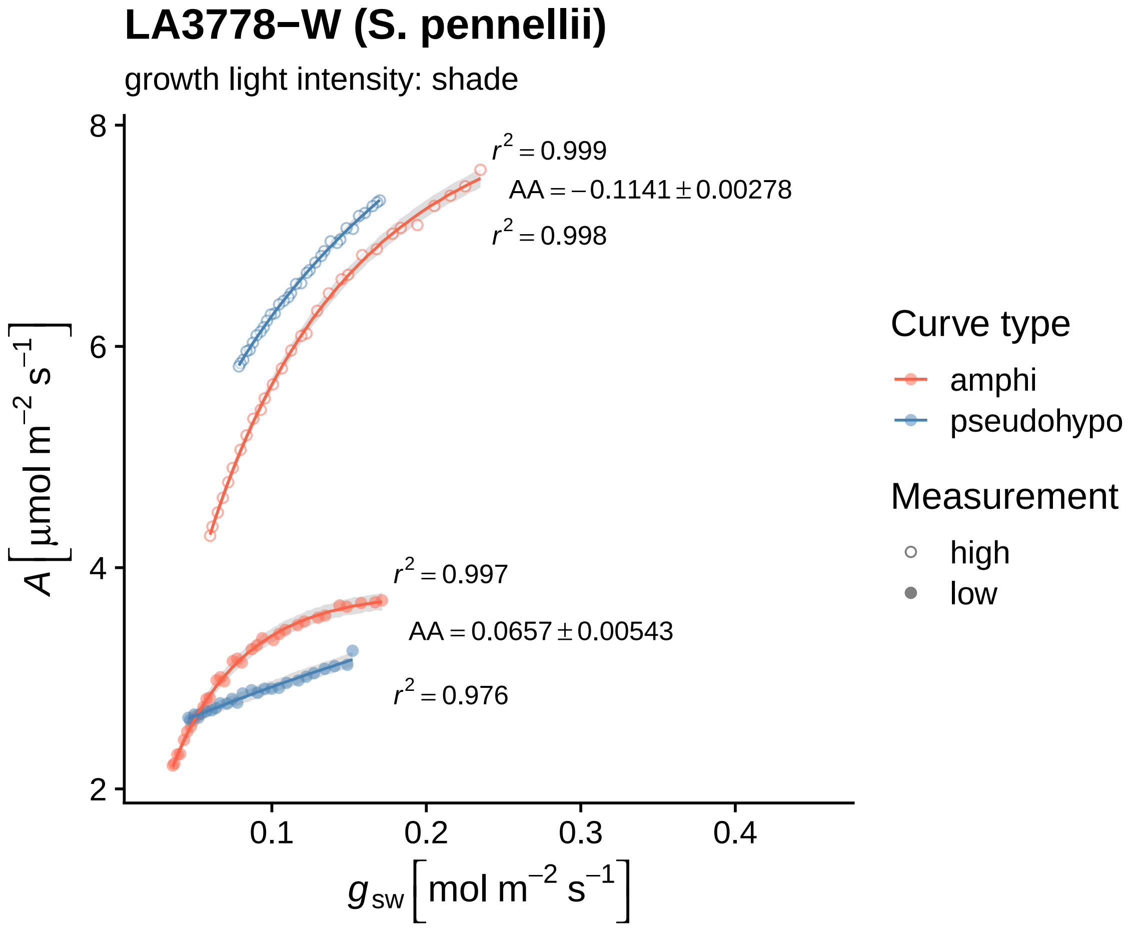
LA3778−W (S. pennellii)
growth light intensity: shade
$r^2 = 0.999$
$AA = -0.1141 \pm 0.00278$
$r^2 = 0.998$
$r^2 = 0.997$
$AA = 0.0657 \pm 0.00543$
$r^2 = 0.976$
$A \left[\mu mol\ m^{-2}\ s^{-1}\right]$
8
6
4
2
0.1
0.2
0.3
0.4
$g_{sw} \left[mol\ m^{-2}\ s^{-1}\right]$
Curve type
amphi
pseudohypo
Measurement
high
low

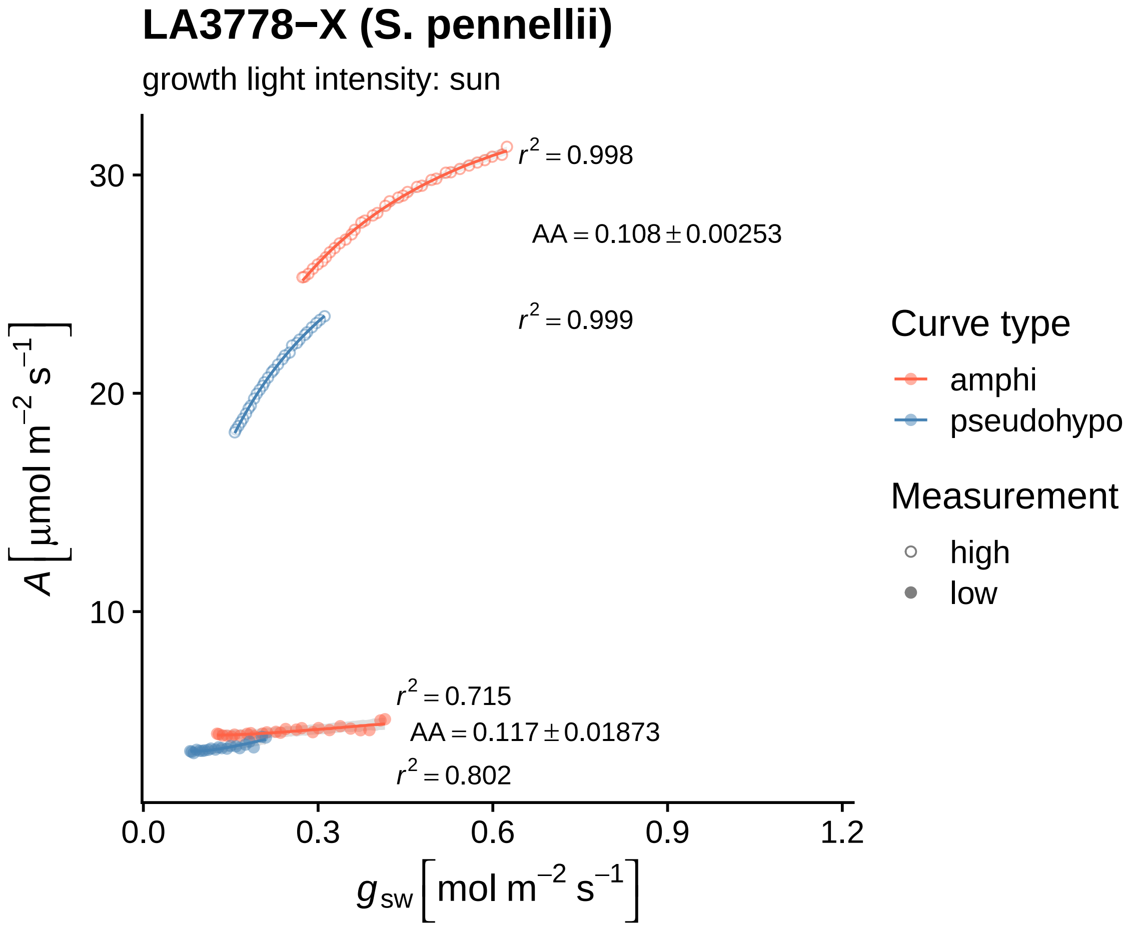
LA3778−X (S. pennellii)
growth light intensity: sun
$r^2 = 0.998$
$AA = 0.108 \pm 0.00253$
$r^2 = 0.999$
$r^2 = 0.715$
$AA = 0.117 \pm 0.01873$
$r^2 = 0.802$
30
20
10
0.0
0.3
0.6
0.9
1.2
$A\left[\mu mol\ m^{-2}\ s^{-1}\right]$
$g_{sw}\left[mol\ m^{-2}\ s^{-1}\right]$
Curve type
amphi
pseudohypo
Measurement
high
low

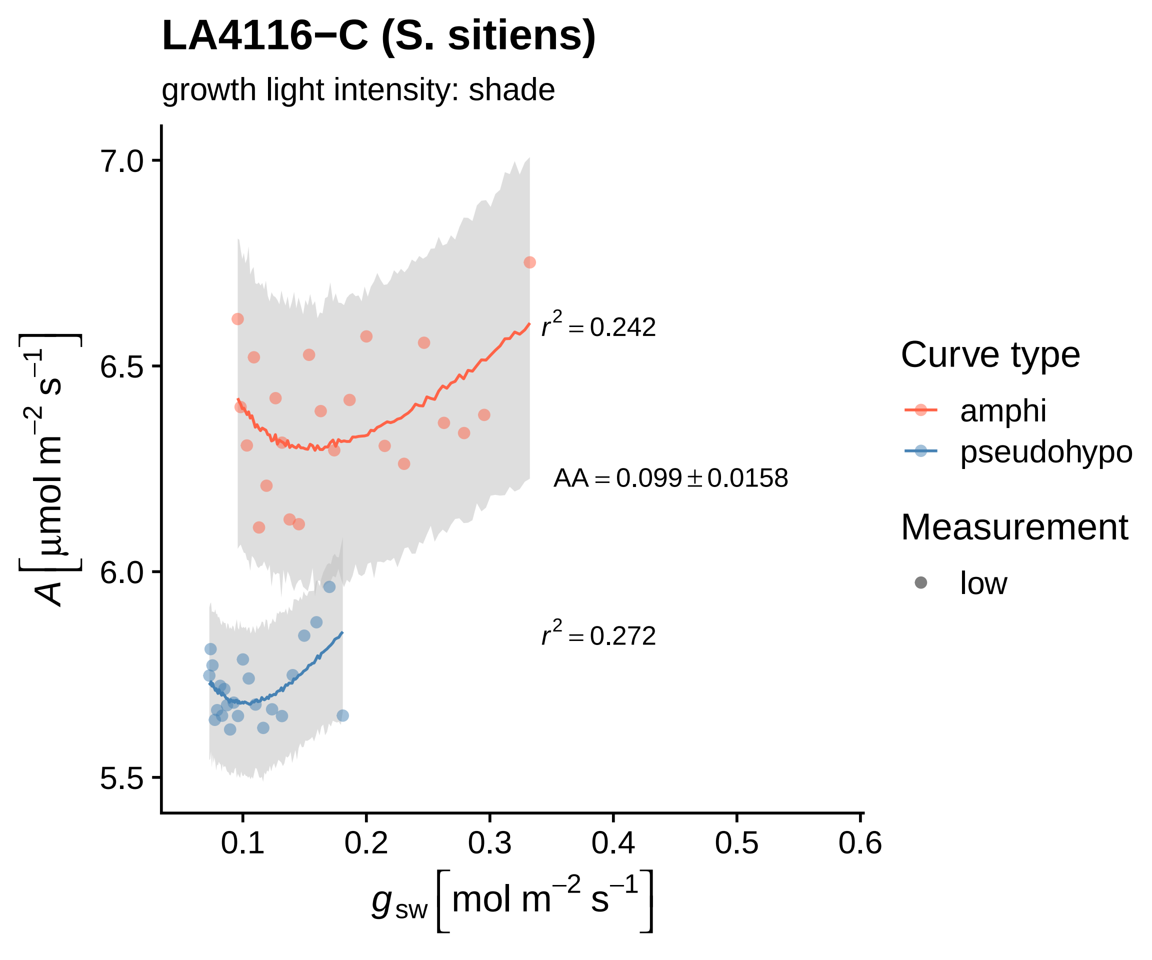

LA4116−C (S. sitiens)
growth light intensity: shade
7.0
6.5
6.0
5.5
A [μmol m−2 s−1]
0.1
0.2
0.3
0.4
0.5
0.6
gsw [mol m−2 s−1]
r2 = 0.242
AA = 0.099 ± 0.0158
r2 = 0.272
Curve type
amphi
pseudohypo
Measurement
low

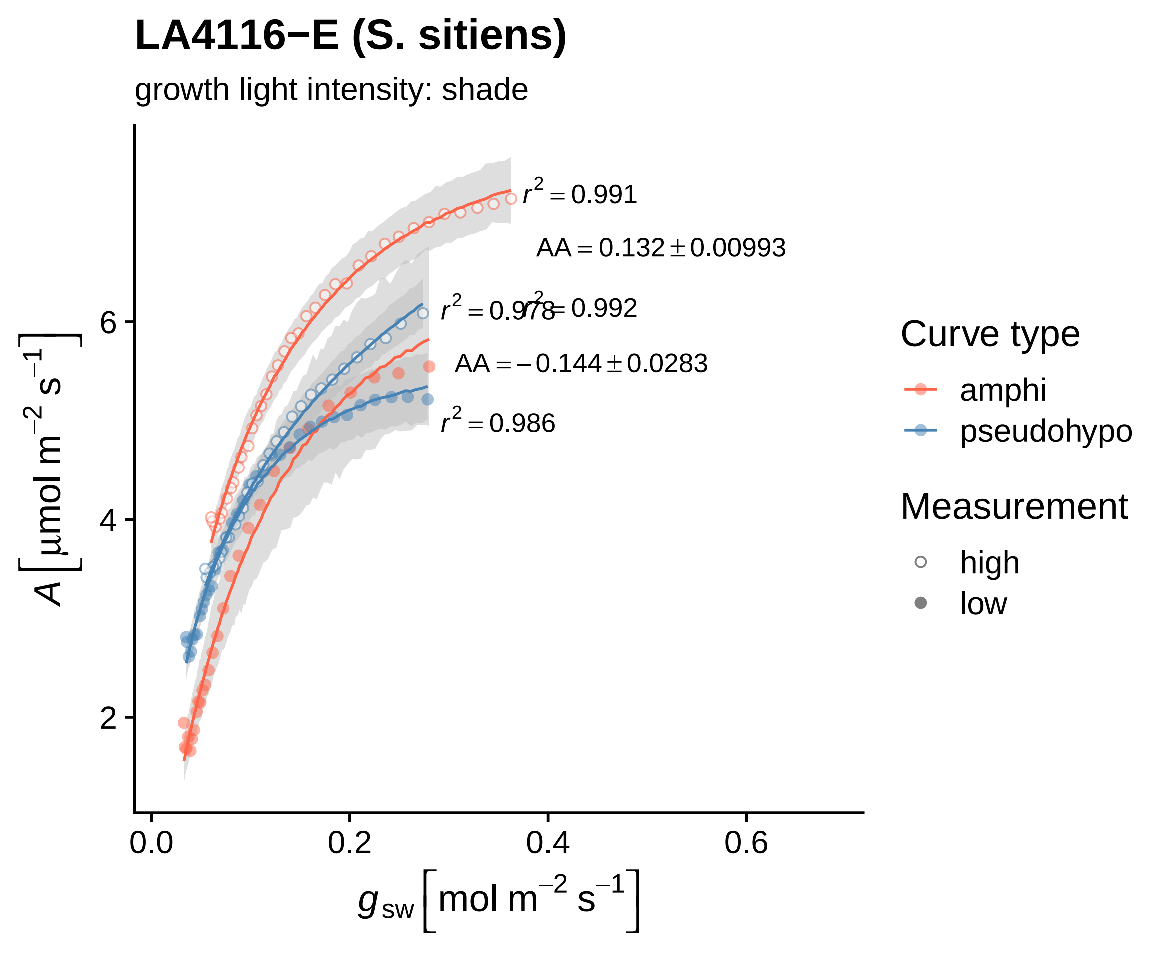

LA4116−E (S. sitiens)
growth light intensity: shade
r² = 0.991
AA = 0.132 ± 0.00993
r² = 0.978
r² = 0.992
AA = −0.144 ± 0.0283
r² = 0.986
2
4
6
0.0
0.2
0.4
0.6
A [μmol m⁻² s⁻¹]
g_sw [mol m⁻² s⁻¹]
Curve type
amphi
pseudohypo
Measurement
high
low

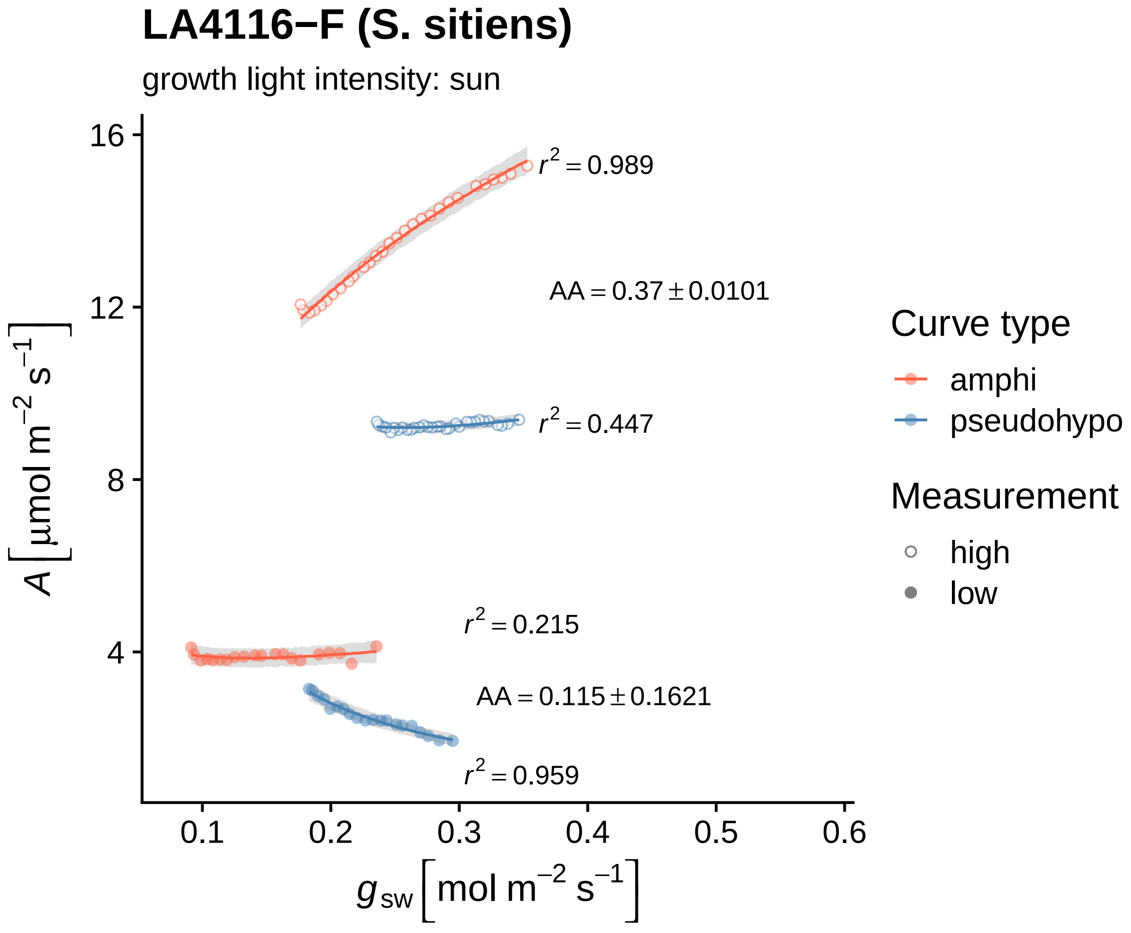
LA4116−F (S. sitiens)
growth light intensity: sun
$r^2 = 0.989$
AA = 0.37 ± 0.0101
$r^2 = 0.447$
$r^2 = 0.215$
AA = 0.115 ± 0.1621
$r^2 = 0.959$
16
12
8
4
0.1
0.2
0.3
0.4
0.5
0.6
$A\ [\mu mol\ m^{-2}\ s^{-1}]$
$g_{sw}\ [mol\ m^{-2}\ s^{-1}]$
Curve type
amphi
pseudohypo
Measurement
high
low

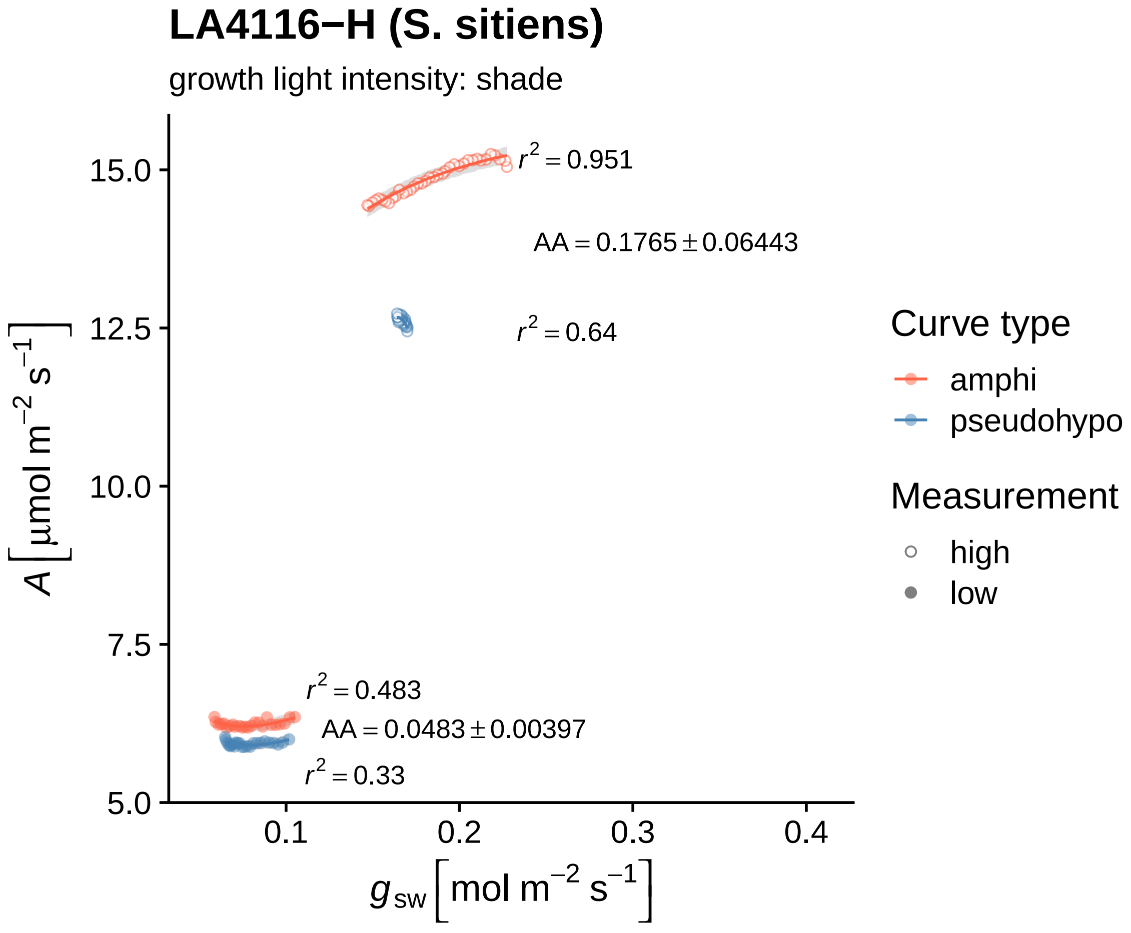

LA4116−H (S. sitiens)
growth light intensity: shade
r^2 = 0.951
AA = 0.1765 ± 0.06443
r^2 = 0.64
r^2 = 0.483
AA = 0.0483 ± 0.00397
r^2 = 0.33
15.0
12.5
10.0
7.5
5.0
0.1
0.2
0.3
0.4
A [µmol m^−2 s^−1]
g_sw [mol m^−2 s^−1]
Curve type
amphi
pseudohypo
Measurement
high
low

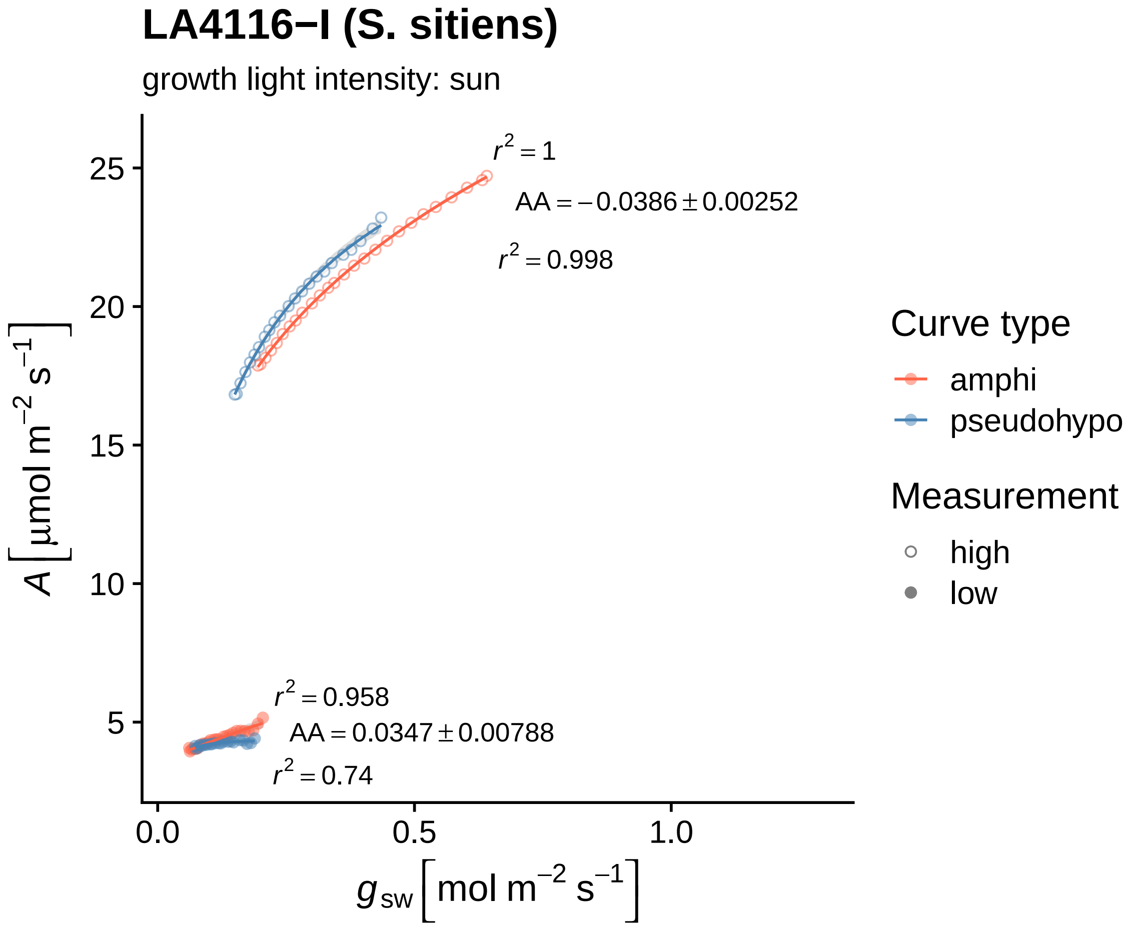
LA4116−I (S. sitiens)
growth light intensity: sun
$r^2 = 1$
$AA = -0.0386 \pm 0.00252$
$r^2 = 0.998$
$r^2 = 0.958$
$AA = 0.0347 \pm 0.00788$
$r^2 = 0.74$
25
20
15
10
5
0.0
0.5
1.0
$A\left[\mu mol\ m^{-2}\ s^{-1}\right]$
$g_{sw}\left[mol\ m^{-2}\ s^{-1}\right]$
Curve type
amphi
pseudohypo
Measurement
high
low

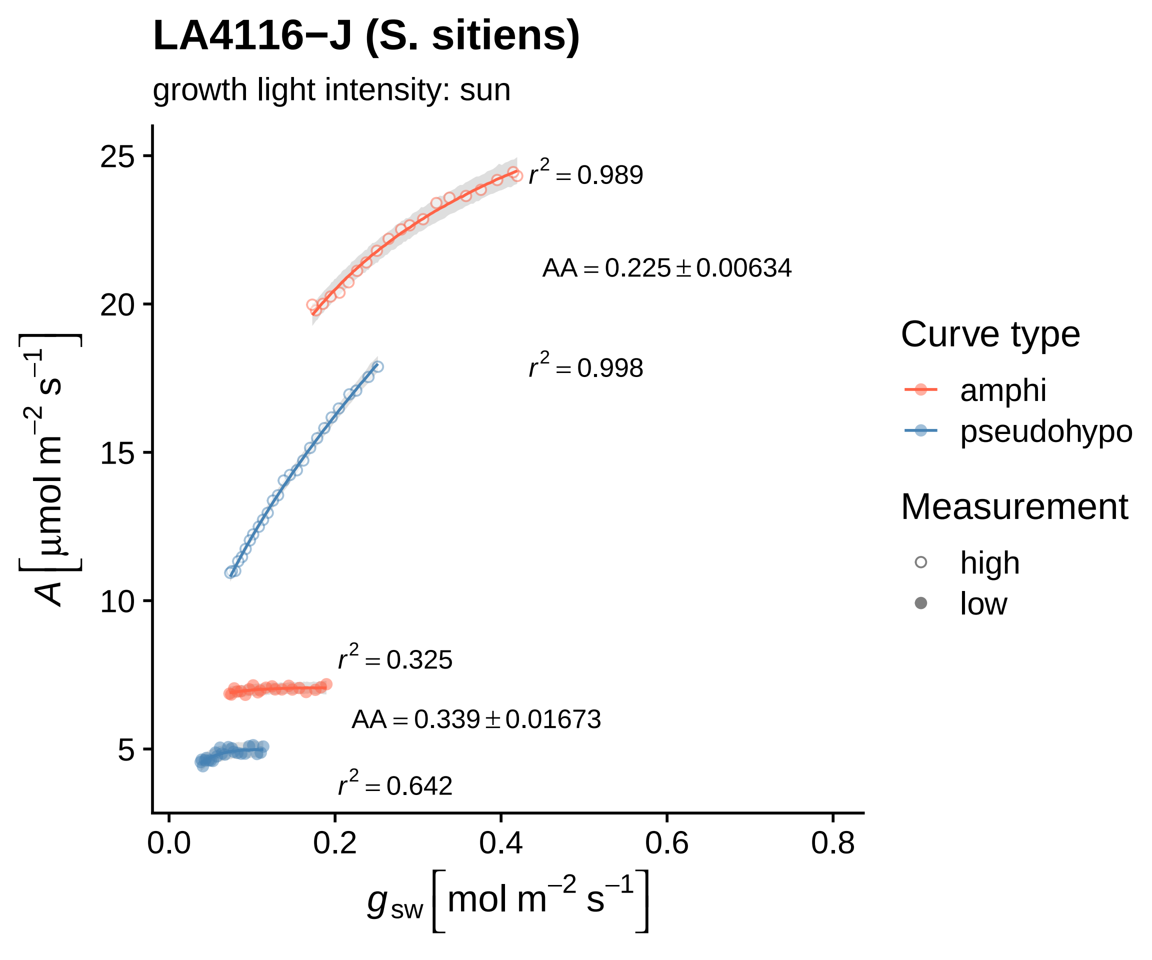

LA4116−J (S. sitiens)
growth light intensity: sun
$r^2 = 0.989$
$AA = 0.225 \pm 0.00634$
$r^2 = 0.998$
$r^2 = 0.325$
$AA = 0.339 \pm 0.01673$
$r^2 = 0.642$
$A\ [\mu mol\ m^{-2}\ s^{-1}]$
25
20
15
10
5
0.0
0.2
0.4
0.6
0.8
$g_{sw}\ [mol\ m^{-2}\ s^{-1}]$
Curve type
amphi
pseudohypo
Measurement
high
low

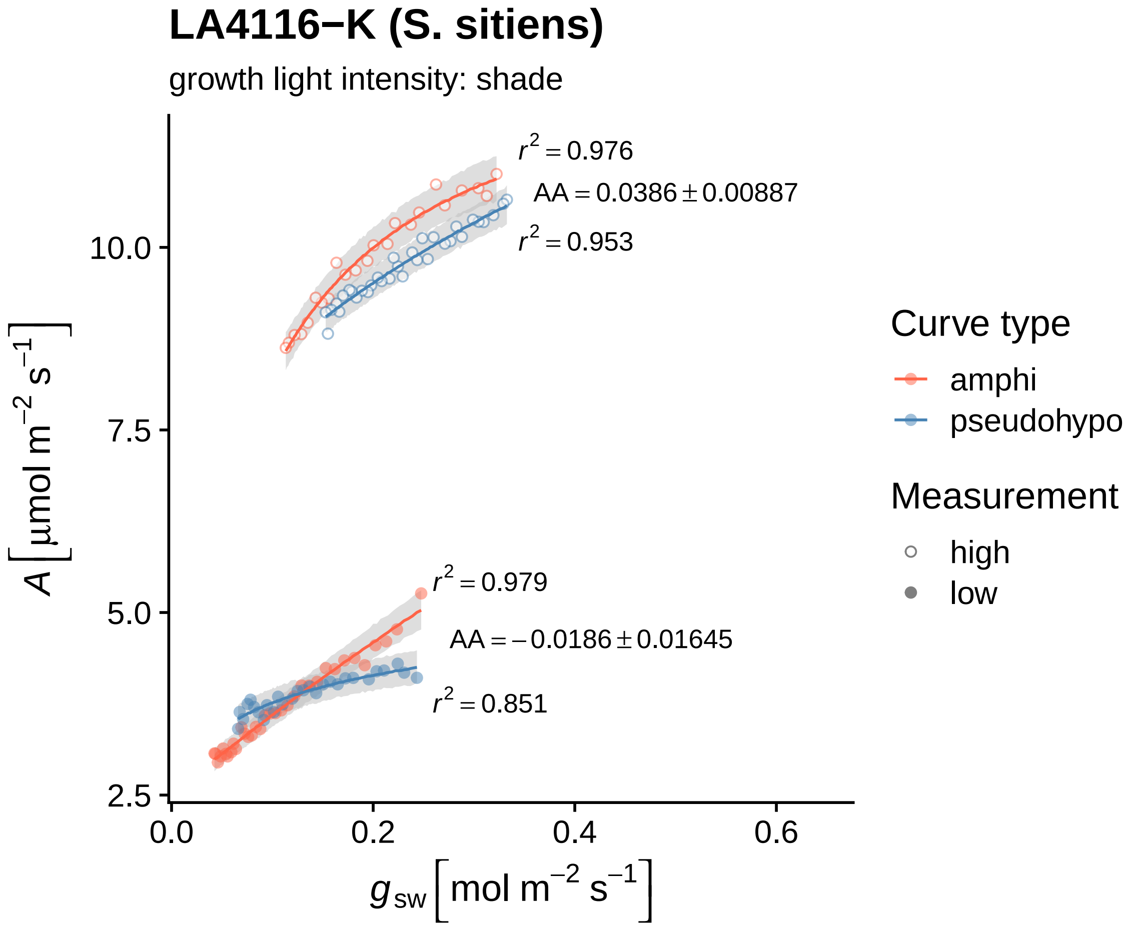

LA4116−K (S. sitiens)
growth light intensity: shade
$r^2 = 0.976$
$AA = 0.0386 \pm 0.00887$
$r^2 = 0.953$
$r^2 = 0.979$
$AA = -0.0186 \pm 0.01645$
$r^2 = 0.851$
10.0
7.5
5.0
2.5
0.0
0.2
0.4
0.6
$A\left[\mu mol\ m^{-2}\ s^{-1}\right]$
$g_{sw}\left[mol\ m^{-2}\ s^{-1}\right]$
Curve type
amphi
pseudohypo
Measurement
high
low

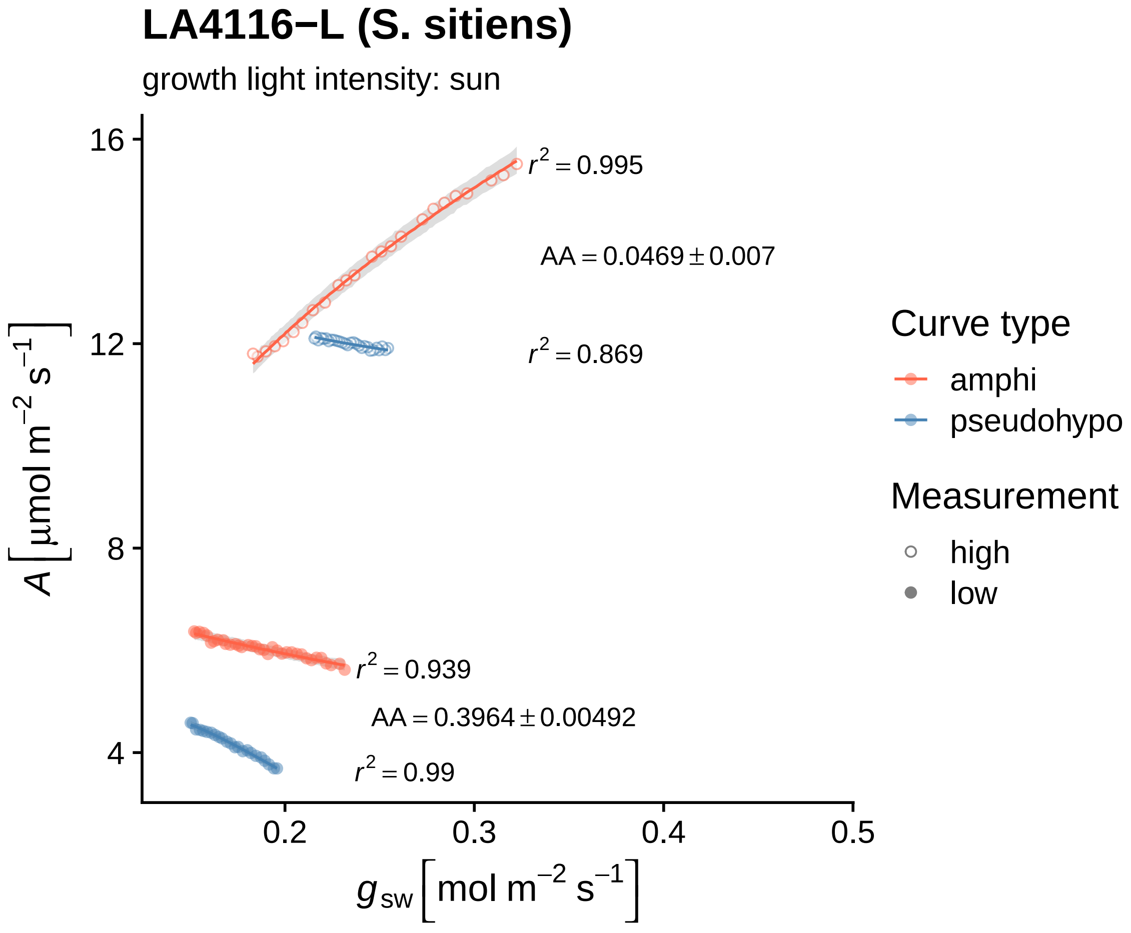
LA4116−L (S. sitiens)
growth light intensity: sun
$r^2 = 0.995$
$AA = 0.0469 \pm 0.007$
$r^2 = 0.869$
$r^2 = 0.939$
$AA = 0.3964 \pm 0.00492$
$r^2 = 0.99$
16
12
8
4
0.2
0.3
0.4
0.5
$A\,[\mu mol\ m^{-2}\ s^{-1}]$
$g_{sw}\,[mol\ m^{-2}\ s^{-1}]$
Curve type
amphi
pseudohypo
Measurement
high
low

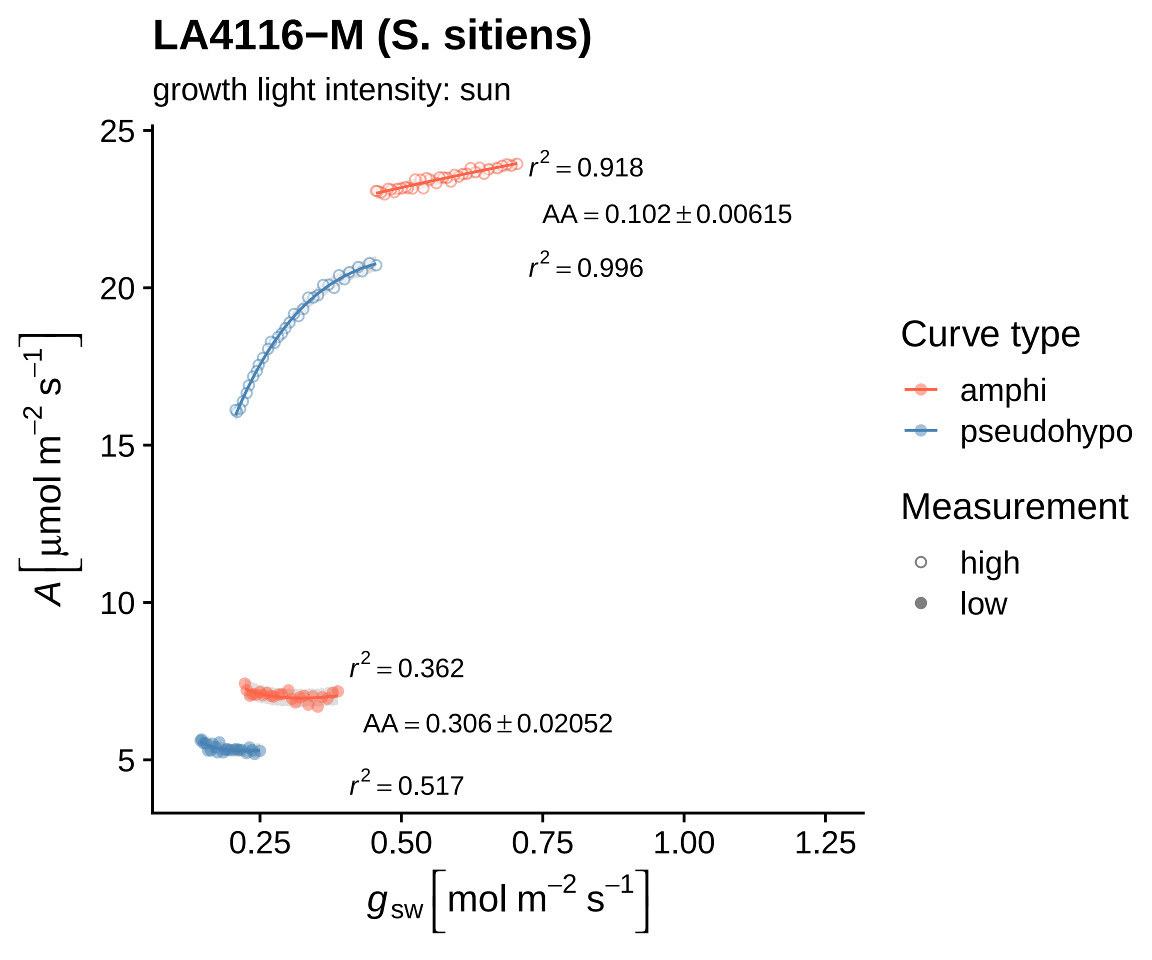
LA4116−M (S. sitiens)
growth light intensity: sun
r² = 0.918
AA = 0.102 ± 0.00615
r² = 0.996
r² = 0.362
AA = 0.306 ± 0.02052
r² = 0.517
25
20
15
10
5
0.25
0.50
0.75
1.00
1.25
A [µmol m⁻² s⁻¹]
g_sw [mol m⁻² s⁻¹]
Curve type
amphi
pseudohypo
Measurement
high
low

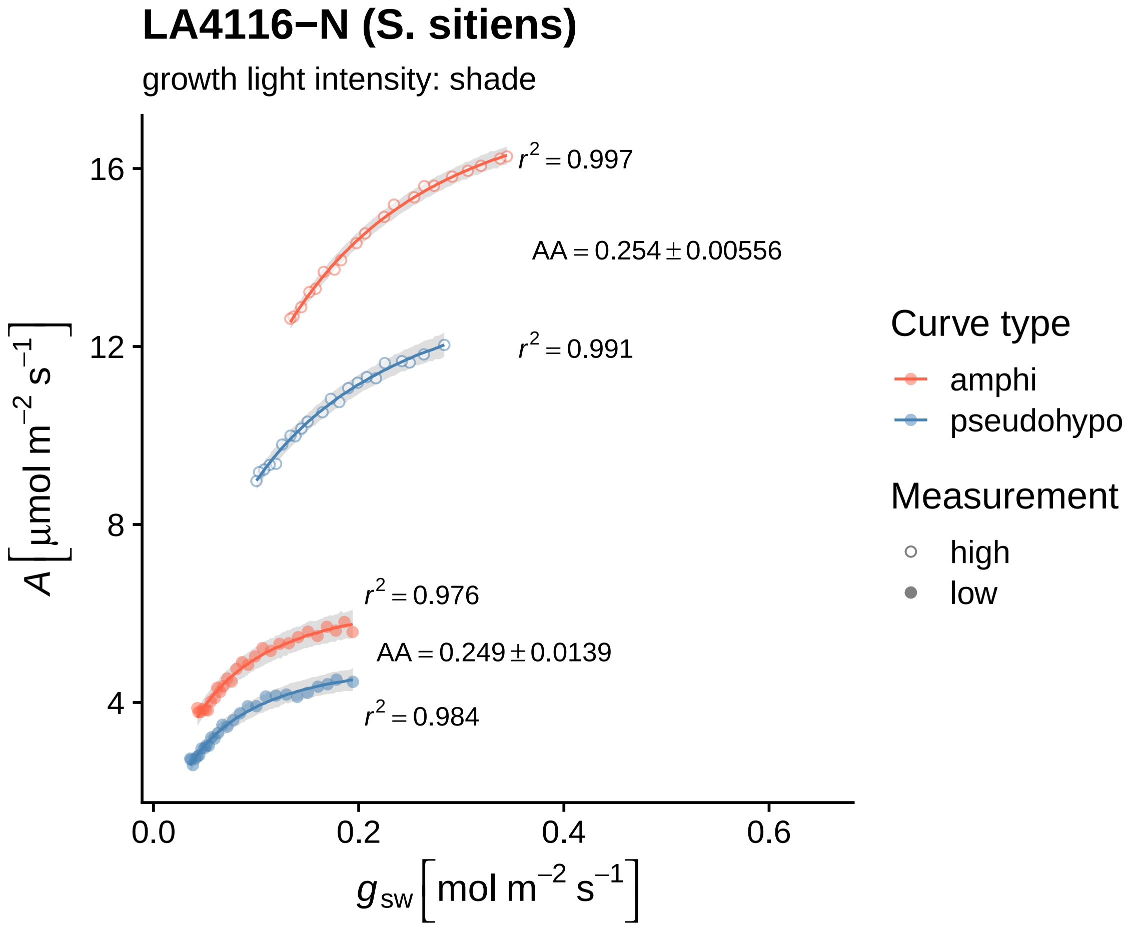
LA4116−N (S. sitiens)
growth light intensity: shade
A [μmol m⁻² s⁻¹]
16
12
8
4
0.0
0.2
0.4
0.6
g_sw [mol m⁻² s⁻¹]
r² = 0.997
AA = 0.254 ± 0.00556
r² = 0.991
r² = 0.976
AA = 0.249 ± 0.0139
r² = 0.984
Curve type
amphi
pseudohypo
Measurement
high
low

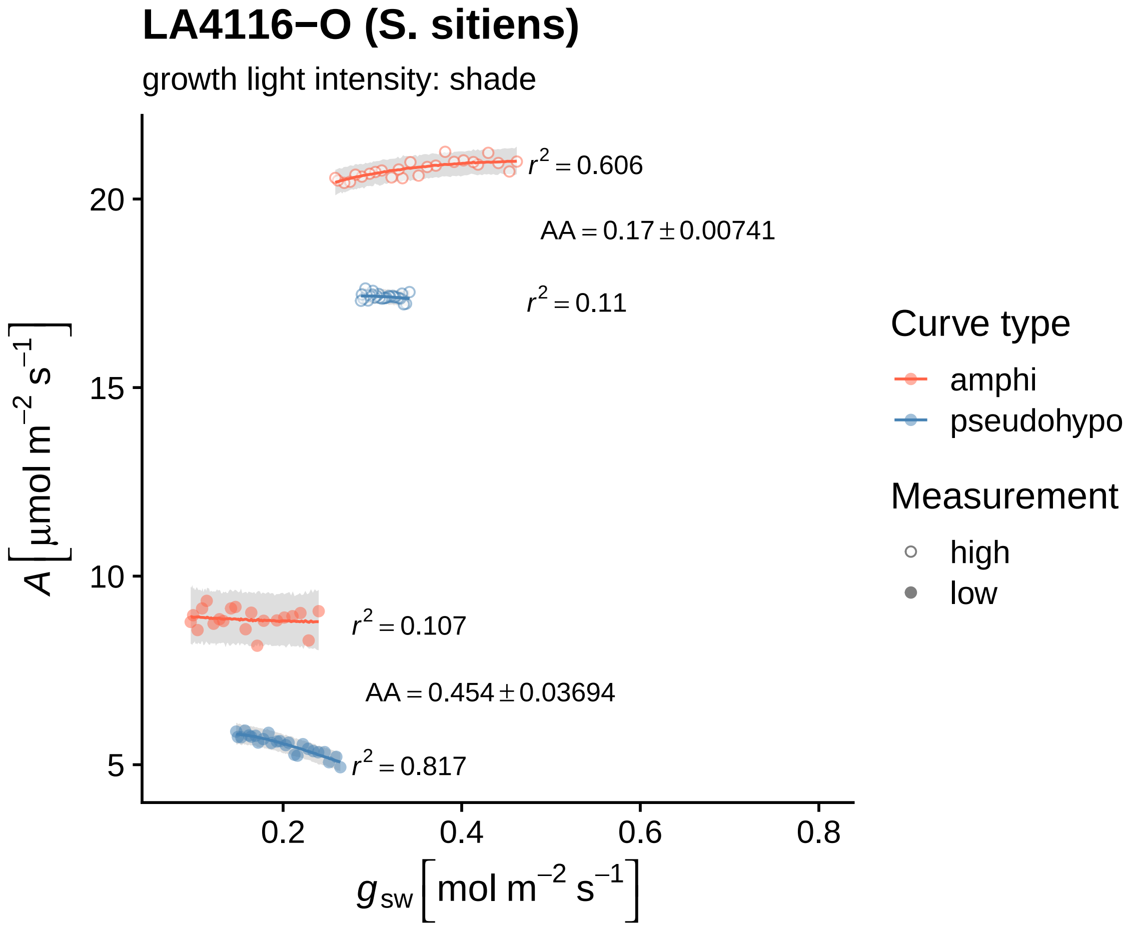
LA4116−O (S. sitiens)
growth light intensity: shade
A [µmol m−2 s−1]
20
15
10
5
0.2
0.4
0.6
0.8
gsw [mol m−2 s−1]
r2 = 0.606
AA = 0.17 ± 0.00741
r2 = 0.11
r2 = 0.107
AA = 0.454 ± 0.03694
r2 = 0.817
Curve type
amphi
pseudohypo
Measurement
high
low

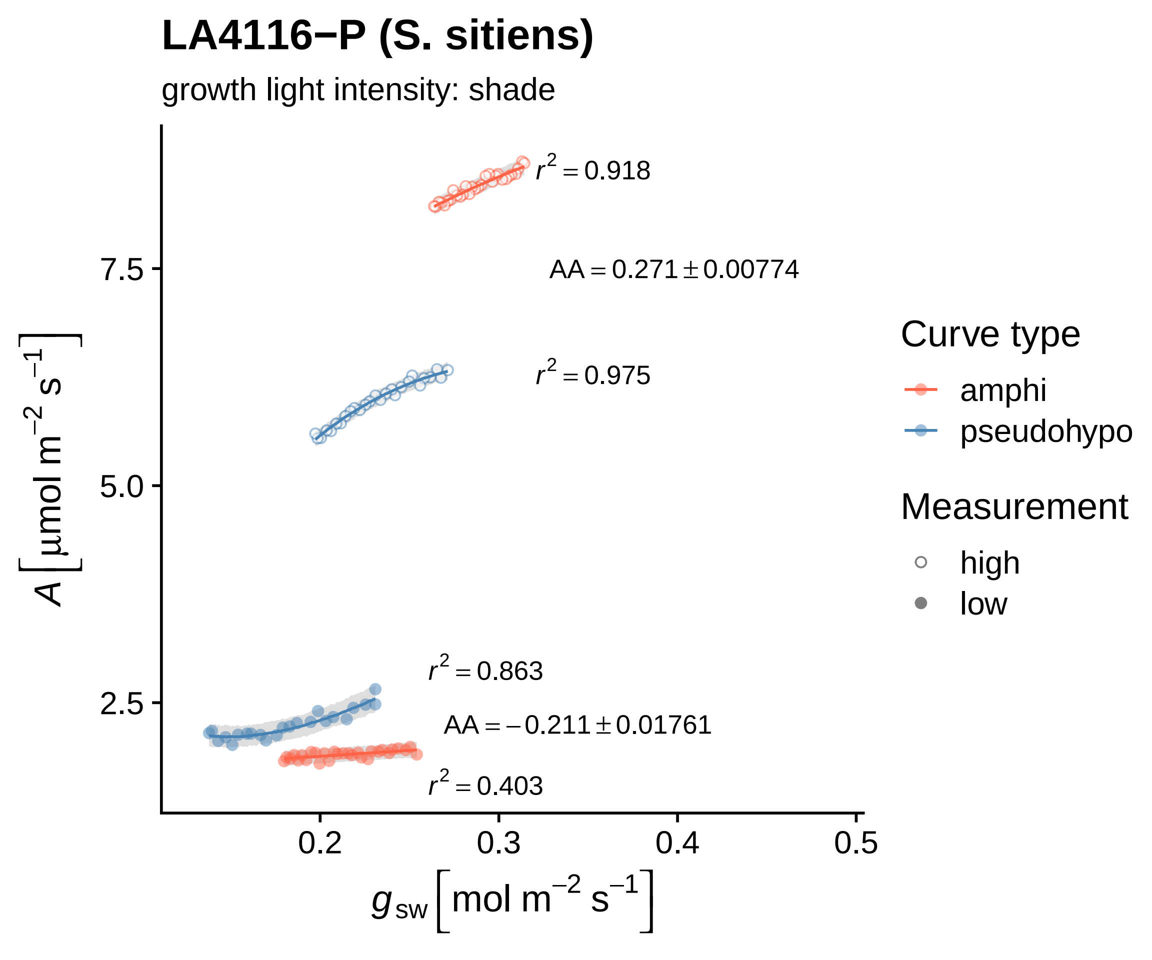

LA4116−P (S. sitiens)
growth light intensity: shade
$r^2 = 0.918$
$AA = 0.271 \pm 0.00774$
$r^2 = 0.975$
$r^2 = 0.863$
$AA = -0.211 \pm 0.01761$
$r^2 = 0.403$
7.5
5.0
2.5
0.2
0.3
0.4
0.5
$A\left[\mu mol\ m^{-2}\ s^{-1}\right]$
$g_{sw}\left[mol\ m^{-2}\ s^{-1}\right]$
Curve type
amphi
pseudohypo
Measurement
high
low

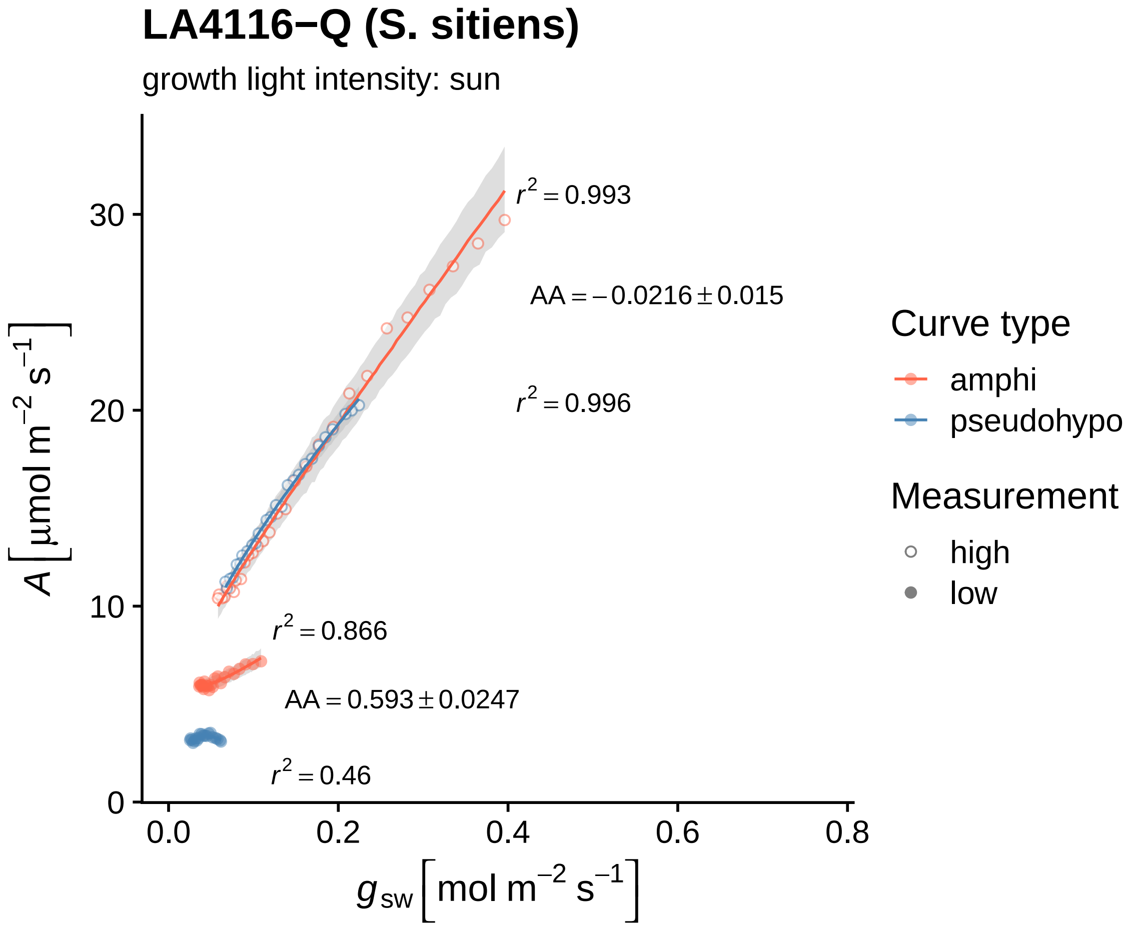
LA4116−Q (S. sitiens)
growth light intensity: sun
$r^2 = 0.993$
$AA = -0.0216 \pm 0.015$
$r^2 = 0.996$
$r^2 = 0.866$
$AA = 0.593 \pm 0.0247$
$r^2 = 0.46$
30
20
10
0
0.0
0.2
0.4
0.6
0.8
$A\left[\mu mol\ m^{-2}\ s^{-1}\right]$
$g_{sw}\left[mol\ m^{-2}\ s^{-1}\right]$
Curve type
amphi
pseudohypo
Measurement
high
low

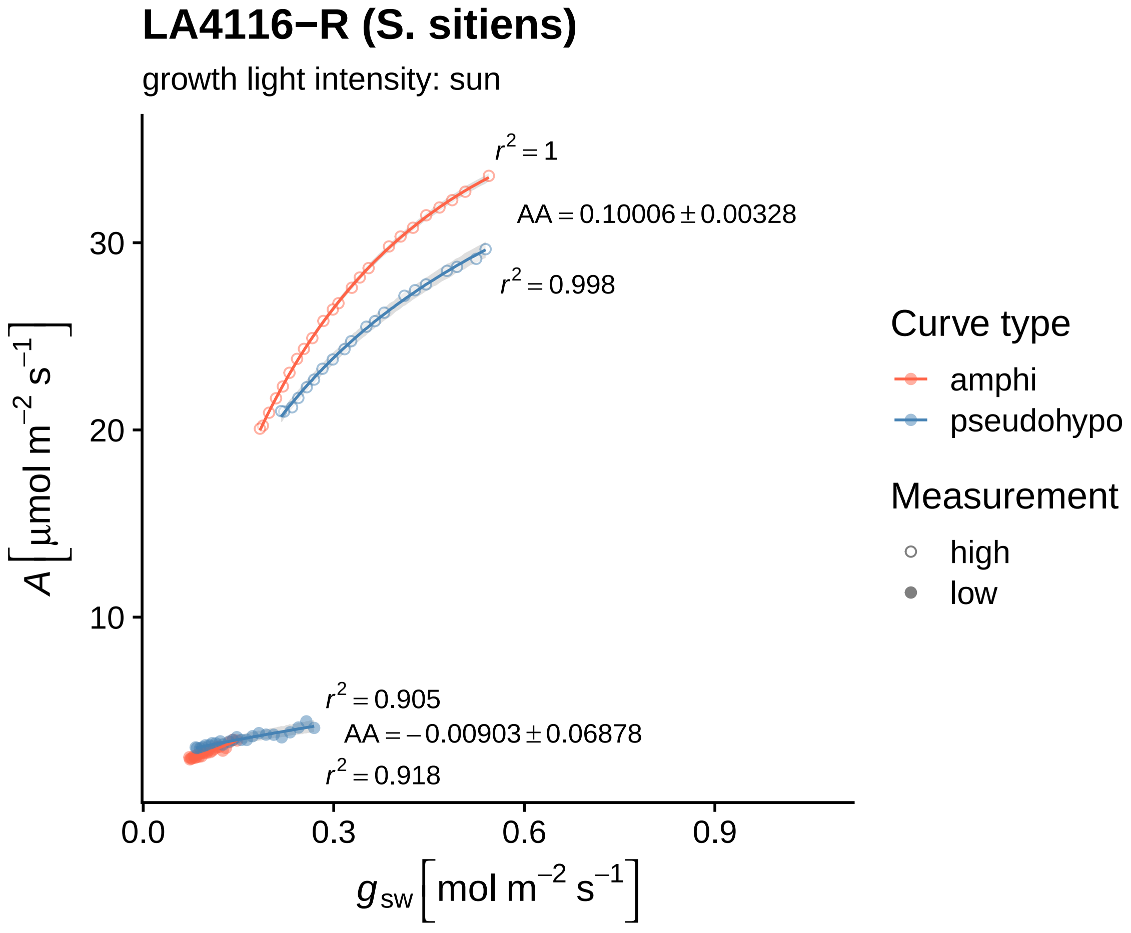
LA4116−R (S. sitiens)
growth light intensity: sun
$r^2 = 1$
AA = 0.10006 ± 0.00328
$r^2 = 0.998$
$r^2 = 0.905$
AA = −0.00903 ± 0.06878
$r^2 = 0.918$
30
20
10
0.0
0.3
0.6
0.9
$A\,[\mu mol\ m^{-2}\ s^{-1}]$
$g_{sw}\,[mol\ m^{-2}\ s^{-1}]$
Curve type
amphi
pseudohypo
Measurement
high
low

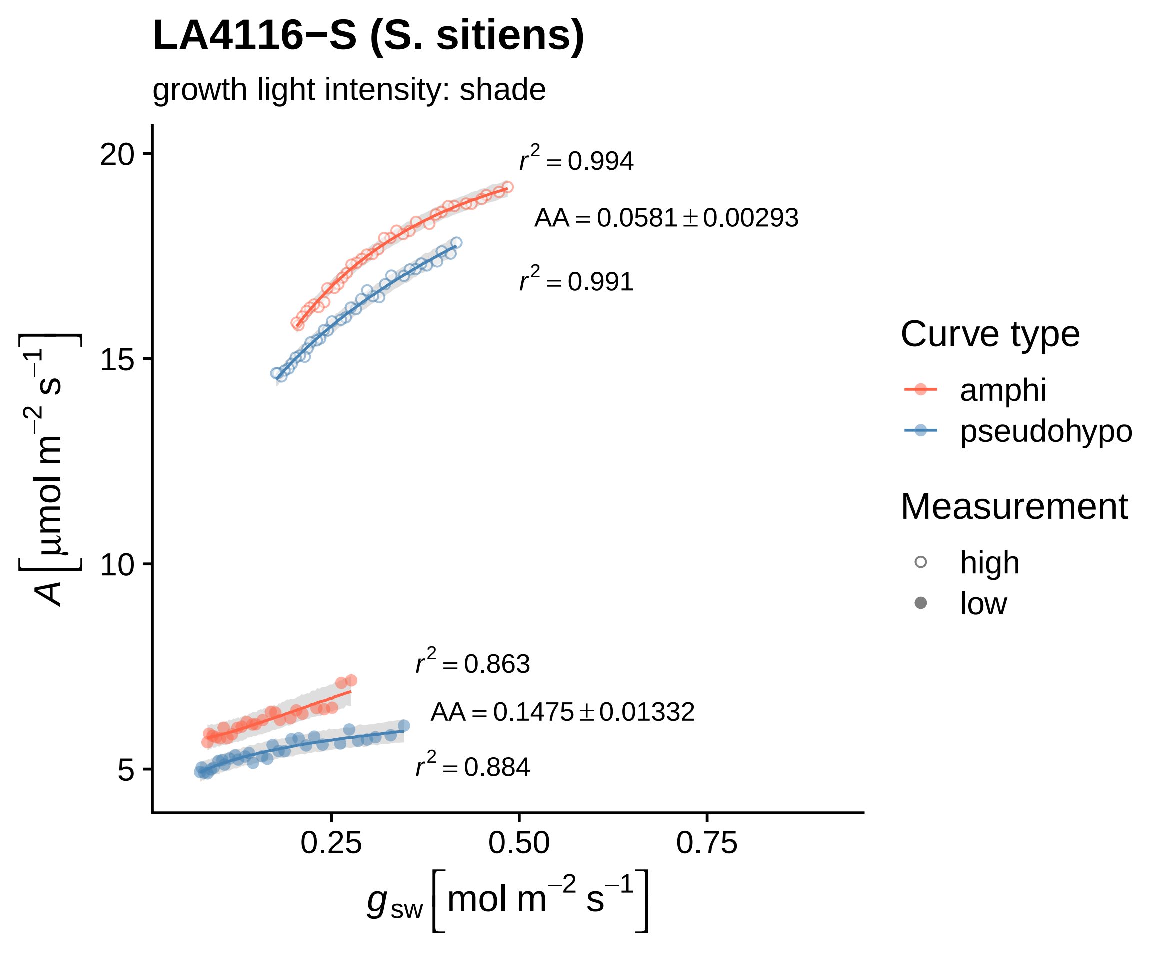

LA4116−S (S. sitiens)
growth light intensity: shade
$r^2 = 0.994$
$AA = 0.0581 \pm 0.00293$
$r^2 = 0.991$
$r^2 = 0.863$
$AA = 0.1475 \pm 0.01332$
$r^2 = 0.884$
20
15
10
5
0.25
0.50
0.75
$A\left[\mu mol\ m^{-2}\ s^{-1}\right]$
$g_{sw}\left[mol\ m^{-2}\ s^{-1}\right]$
Curve type
amphi
pseudohypo
Measurement
high
low

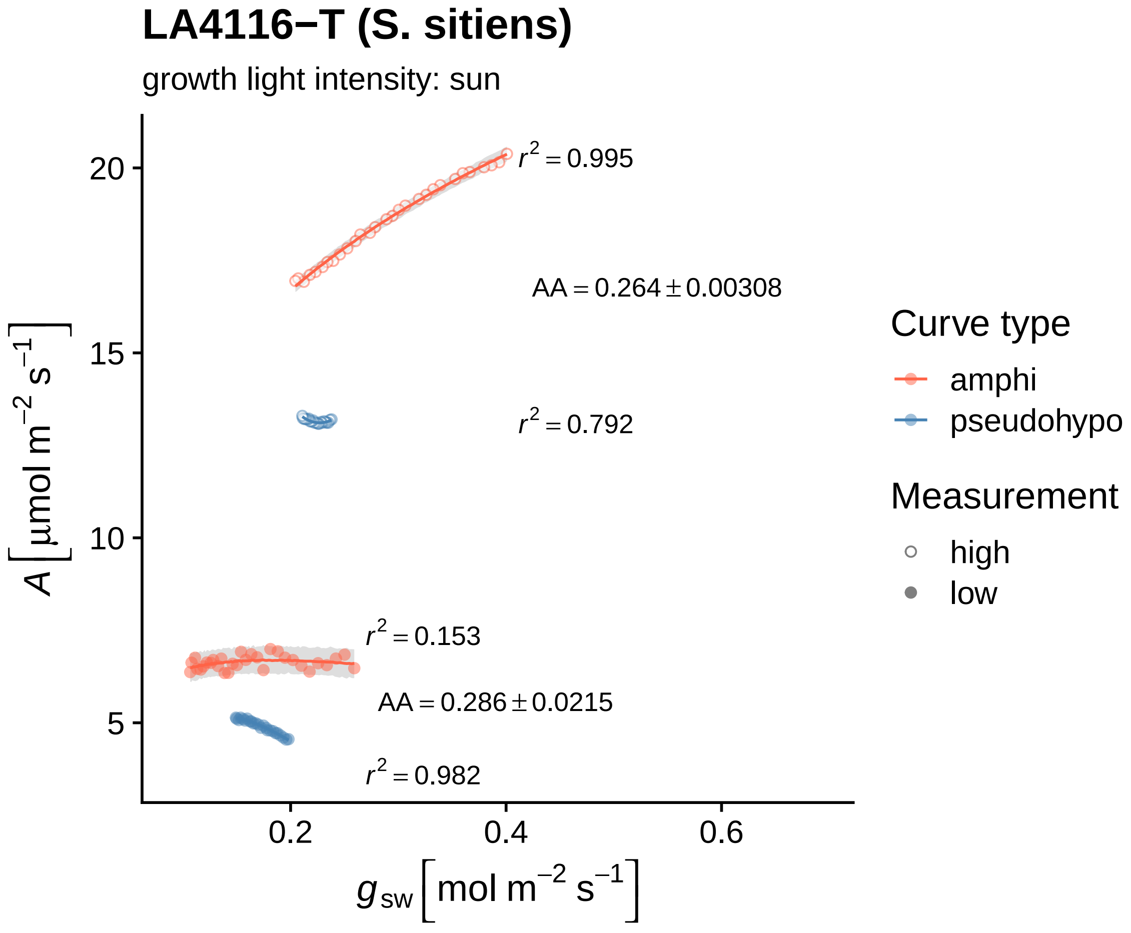

LA4116−T (S. sitiens)
growth light intensity: sun
r² = 0.995
AA = 0.264 ± 0.00308
r² = 0.792
r² = 0.153
AA = 0.286 ± 0.0215
r² = 0.982
20
15
10
5
0.2
0.4
0.6
A [µmol m⁻² s⁻¹]
g_sw [mol m⁻² s⁻¹]
Curve type
amphi
pseudohypo
Measurement
high
low

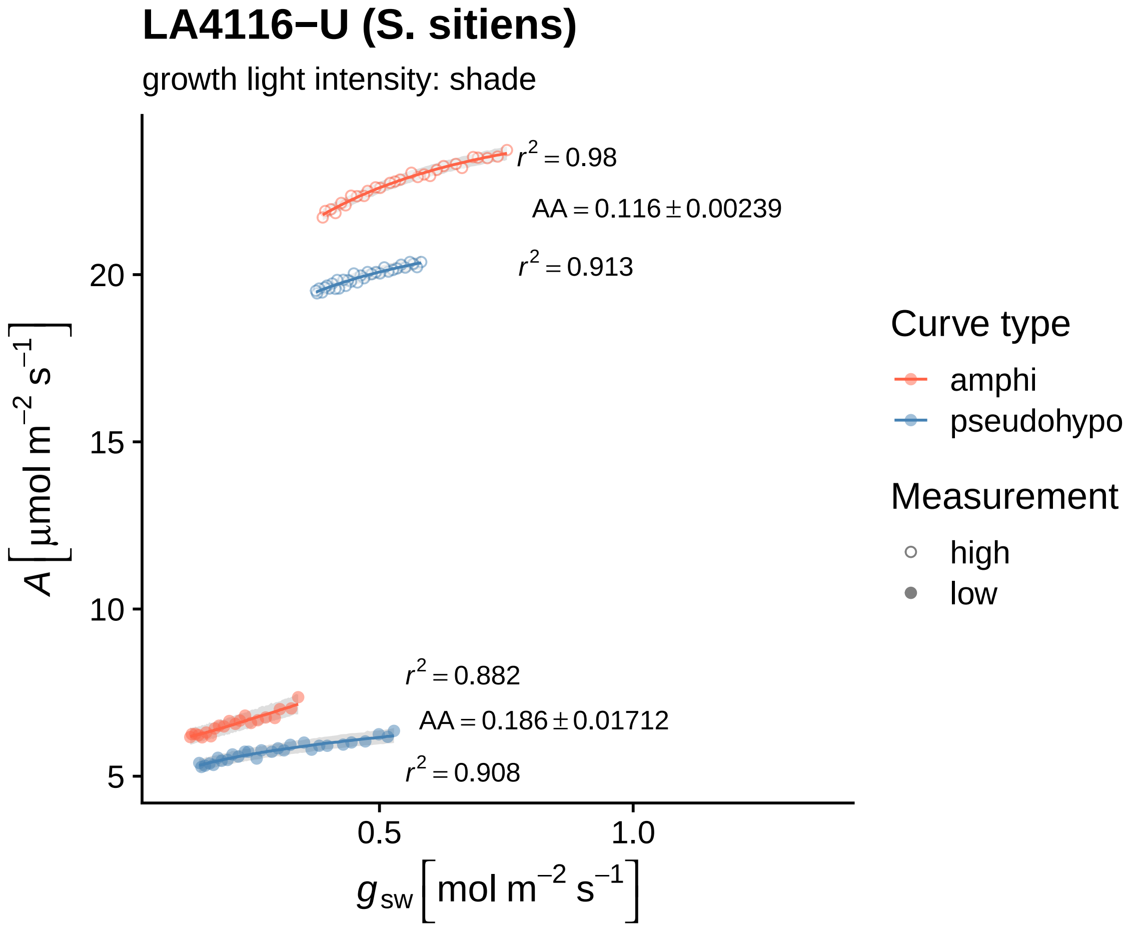
LA4116−U (S. sitiens)
growth light intensity: shade
r² = 0.98
AA = 0.116 ± 0.00239
r² = 0.913
r² = 0.882
AA = 0.186 ± 0.01712
r² = 0.908
A [µmol m⁻² s⁻¹]
20
15
10
5
0.5
1.0
g_sw [mol m⁻² s⁻¹]
Curve type
amphi
pseudohypo
Measurement
high
low

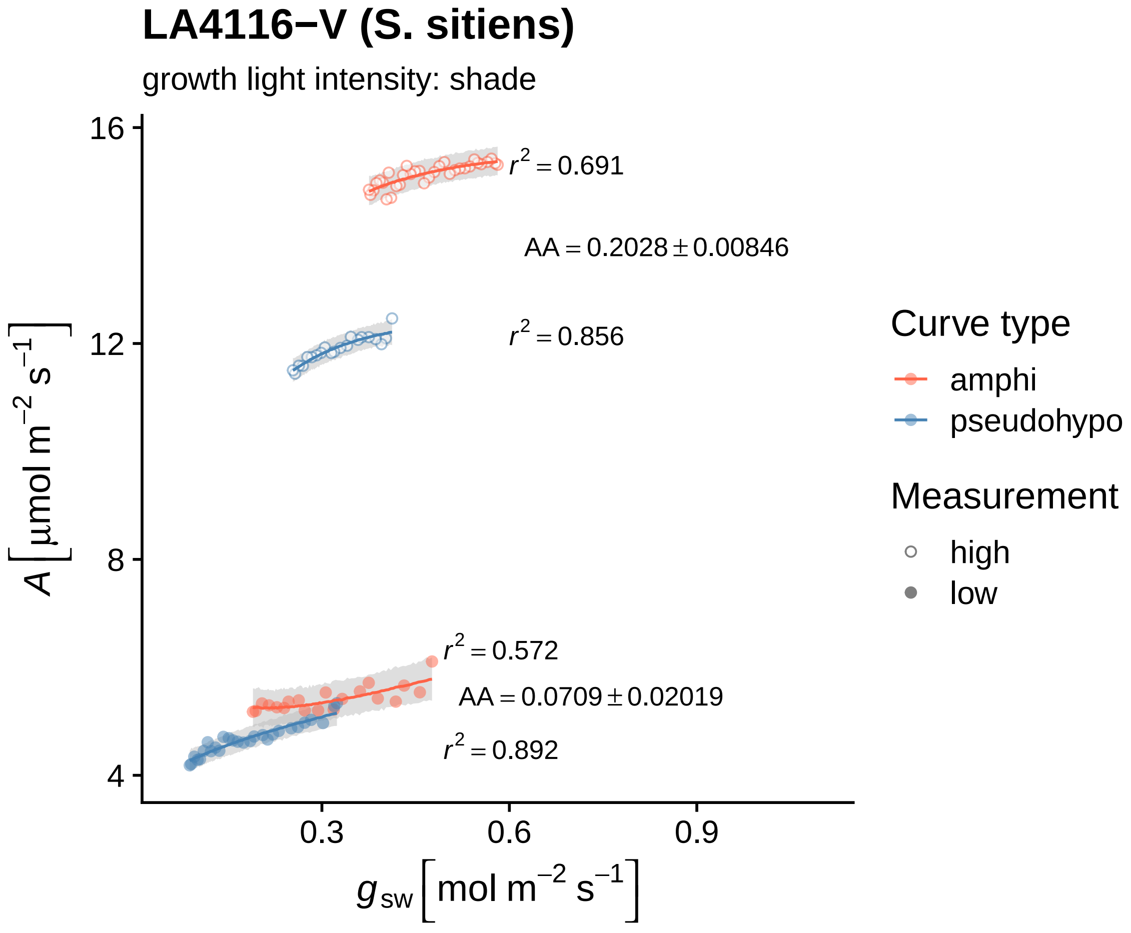

LA4116−V (S. sitiens)
growth light intensity: shade
$r^2 = 0.691$
AA = 0.2028 ± 0.00846
$r^2 = 0.856$
$r^2 = 0.572$
AA = 0.0709 ± 0.02019
$r^2 = 0.892$
16
12
8
4
0.3
0.6
0.9
A [μmol m⁻² s⁻¹]
$g_{sw}$ [mol m⁻² s⁻¹]
Curve type
amphi
pseudohypo
Measurement
high
low

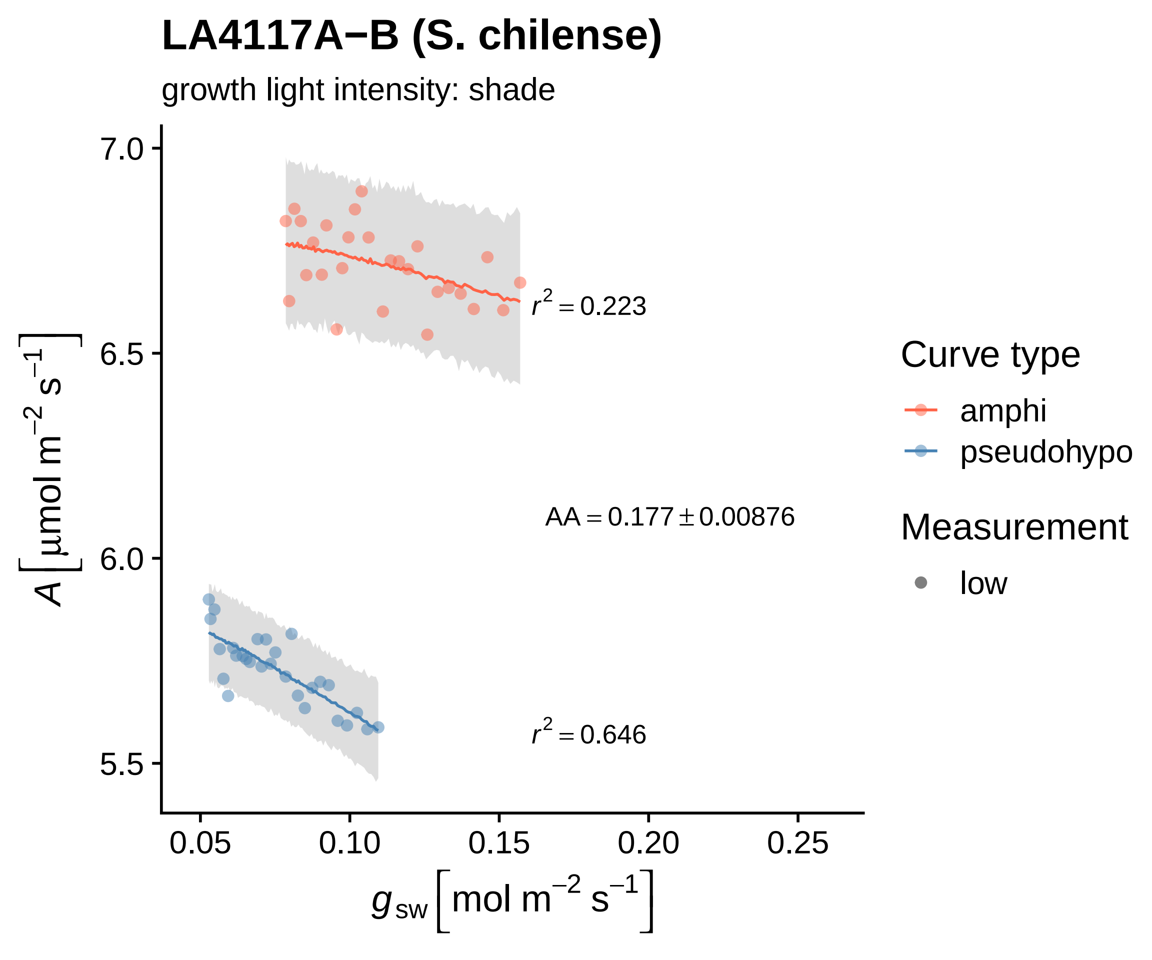

LA4117A−B (S. chilense)
growth light intensity: shade
7.0
6.5
6.0
5.5
0.05
0.10
0.15
0.20
0.25
A [μmol m−2 s−1]
gsw [mol m−2 s−1]
r2 = 0.223
AA = 0.177 ± 0.00876
r2 = 0.646
Curve type
amphi
pseudohypo
Measurement
low

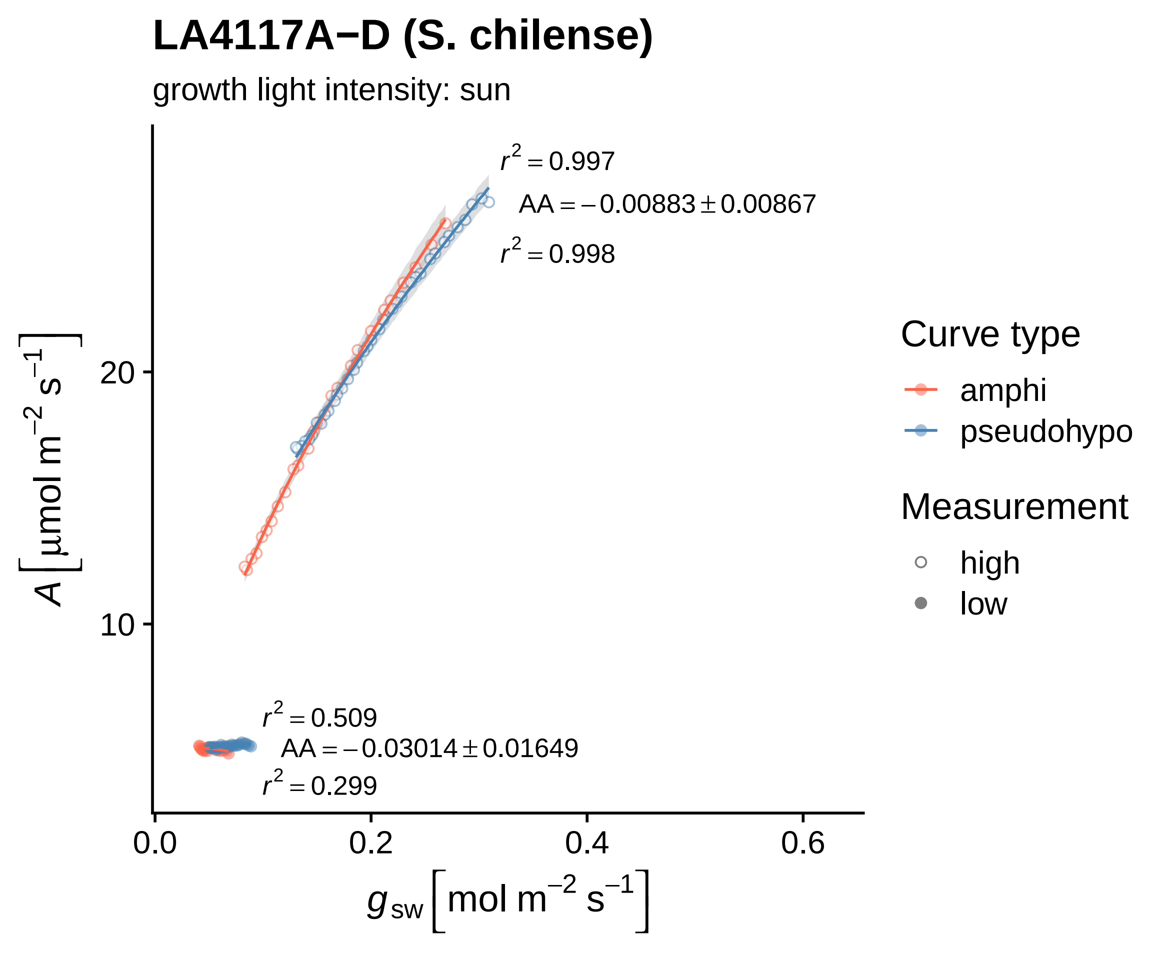

LA4117A−D (S. chilense)
growth light intensity: sun
r² = 0.997
AA = −0.00883 ± 0.00867
r² = 0.998
r² = 0.509
AA = −0.03014 ± 0.01649
r² = 0.299
20
10
0.0
0.2
0.4
0.6
A [µmol m⁻² s⁻¹]
g sw [mol m⁻² s⁻¹]
Curve type
amphi
pseudohypo
Measurement
high
low

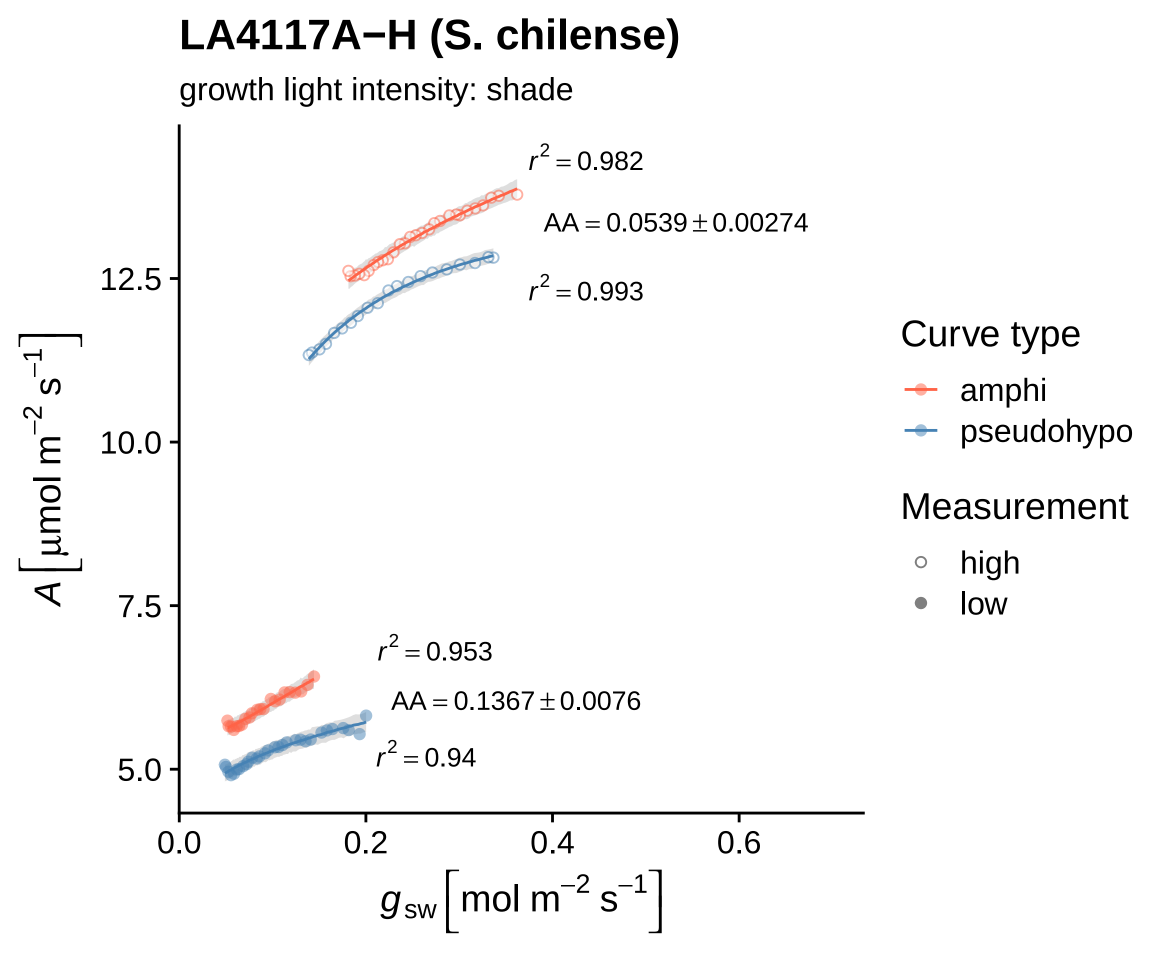

LA4117A−H (S. chilense)
growth light intensity: shade
$r^2 = 0.982$
$AA = 0.0539 \pm 0.00274$
$r^2 = 0.993$
$r^2 = 0.953$
$AA = 0.1367 \pm 0.0076$
$r^2 = 0.94$
12.5
10.0
7.5
5.0
0.0
0.2
0.4
0.6
$A \left[\mu mol\ m^{-2}\ s^{-1}\right]$
$g_{sw} \left[mol\ m^{-2}\ s^{-1}\right]$
Curve type
amphi
pseudohypo
Measurement
high
low

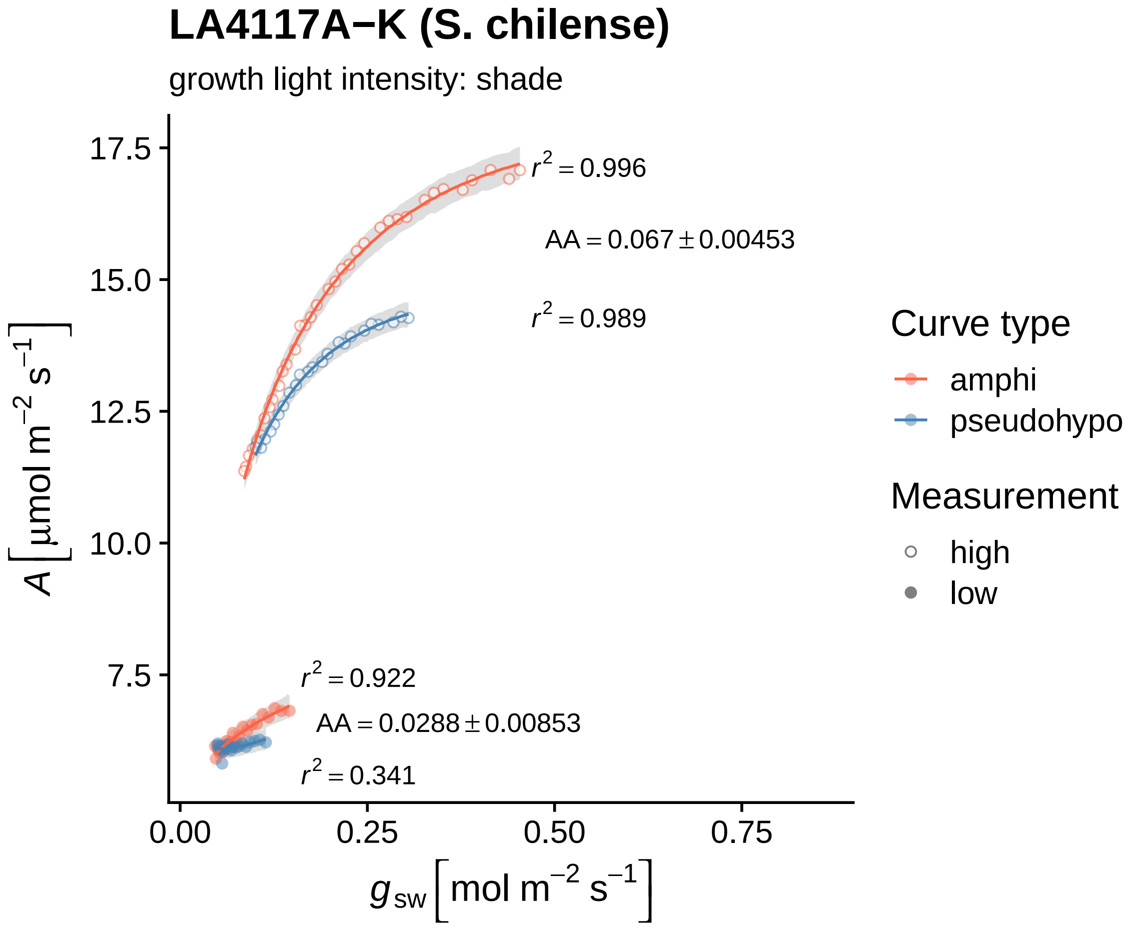

LA4117A−K (S. chilense)
growth light intensity: shade
$r^2 = 0.996$
$AA = 0.067 \pm 0.00453$
$r^2 = 0.989$
$r^2 = 0.922$
$AA = 0.0288 \pm 0.00853$
$r^2 = 0.341$
17.5
15.0
12.5
10.0
7.5
0.00
0.25
0.50
0.75
$A\ [\mu mol\ m^{-2}\ s^{-1}]$
$g_{sw}\ [mol\ m^{-2}\ s^{-1}]$
Curve type
amphi
pseudohypo
Measurement
high
low

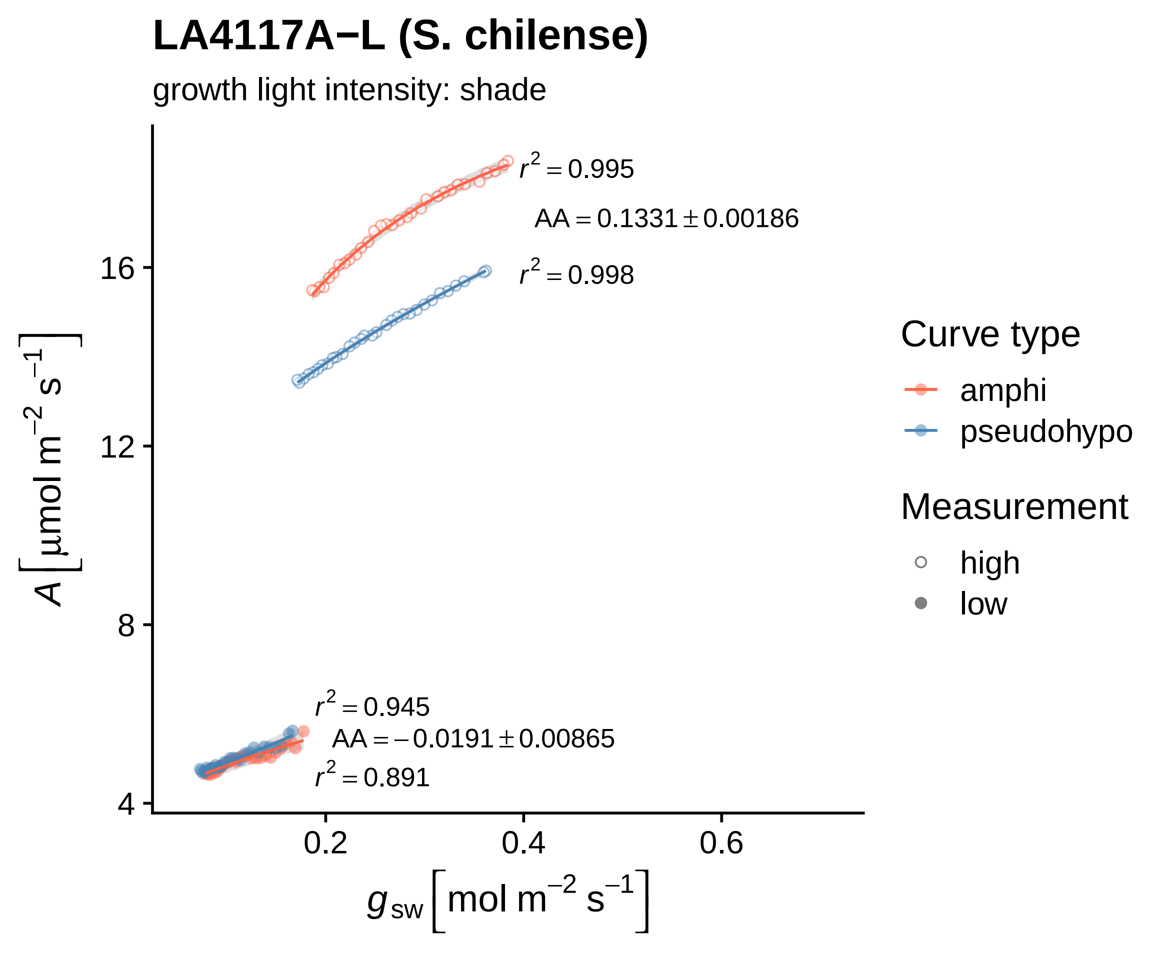
LA4117A−L (S. chilense)
growth light intensity: shade
$r^2 = 0.995$
$AA = 0.1331 \pm 0.00186$
$r^2 = 0.998$
$r^2 = 0.945$
$AA = -0.0191 \pm 0.00865$
$r^2 = 0.891$
16
12
8
4
0.2
0.4
0.6
$A\ [\mu mol\ m^{-2}\ s^{-1}]$
$g_{sw}\ [mol\ m^{-2}\ s^{-1}]$
Curve type
amphi
pseudohypo
Measurement
high
low

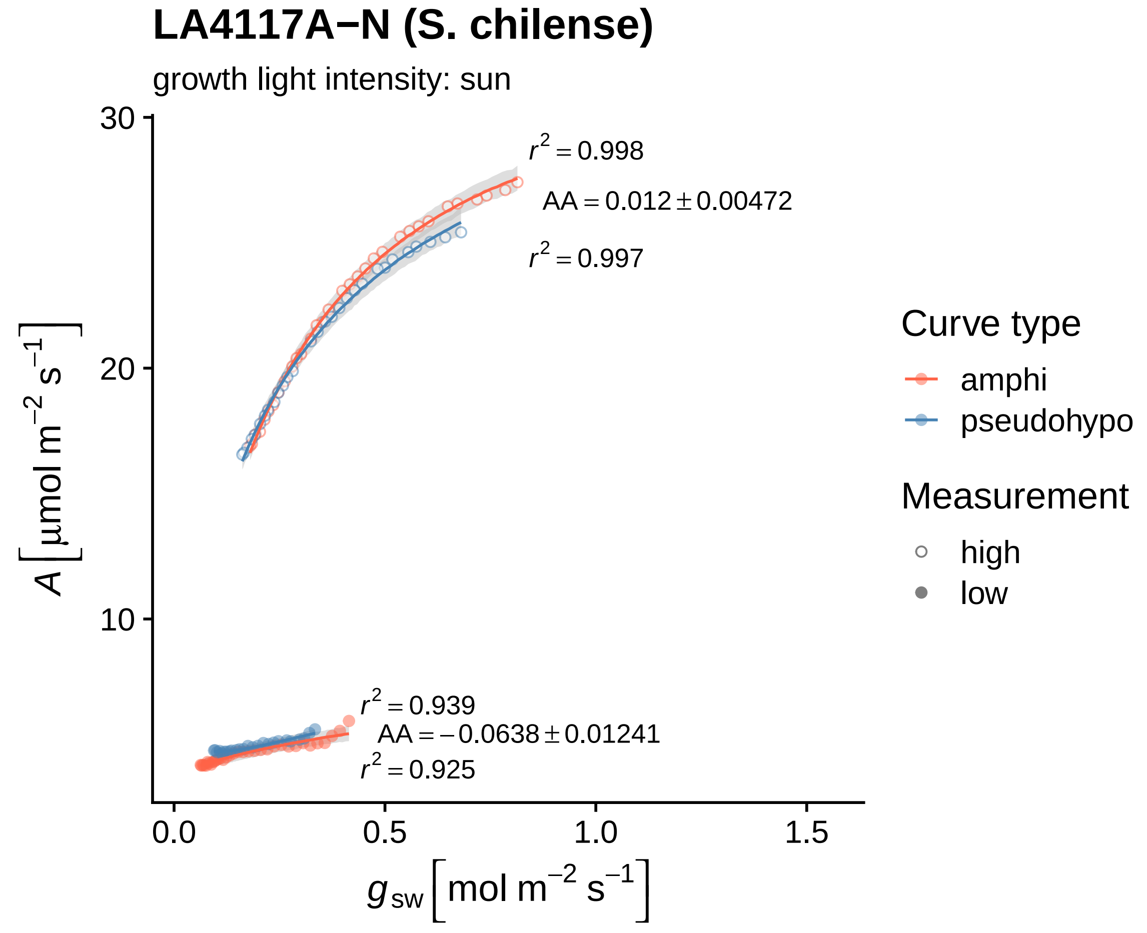

LA4117A−N (S. chilense)
growth light intensity: sun
$r^2 = 0.998$
$AA = 0.012 \pm 0.00472$
$r^2 = 0.997$
$r^2 = 0.939$
$AA = -0.0638 \pm 0.01241$
$r^2 = 0.925$
30
20
10
0.0
0.5
1.0
1.5
$A\left[\mu mol\ m^{-2}\ s^{-1}\right]$
$g_{sw}\left[mol\ m^{-2}\ s^{-1}\right]$
Curve type
amphi
pseudohypo
Measurement
high
low

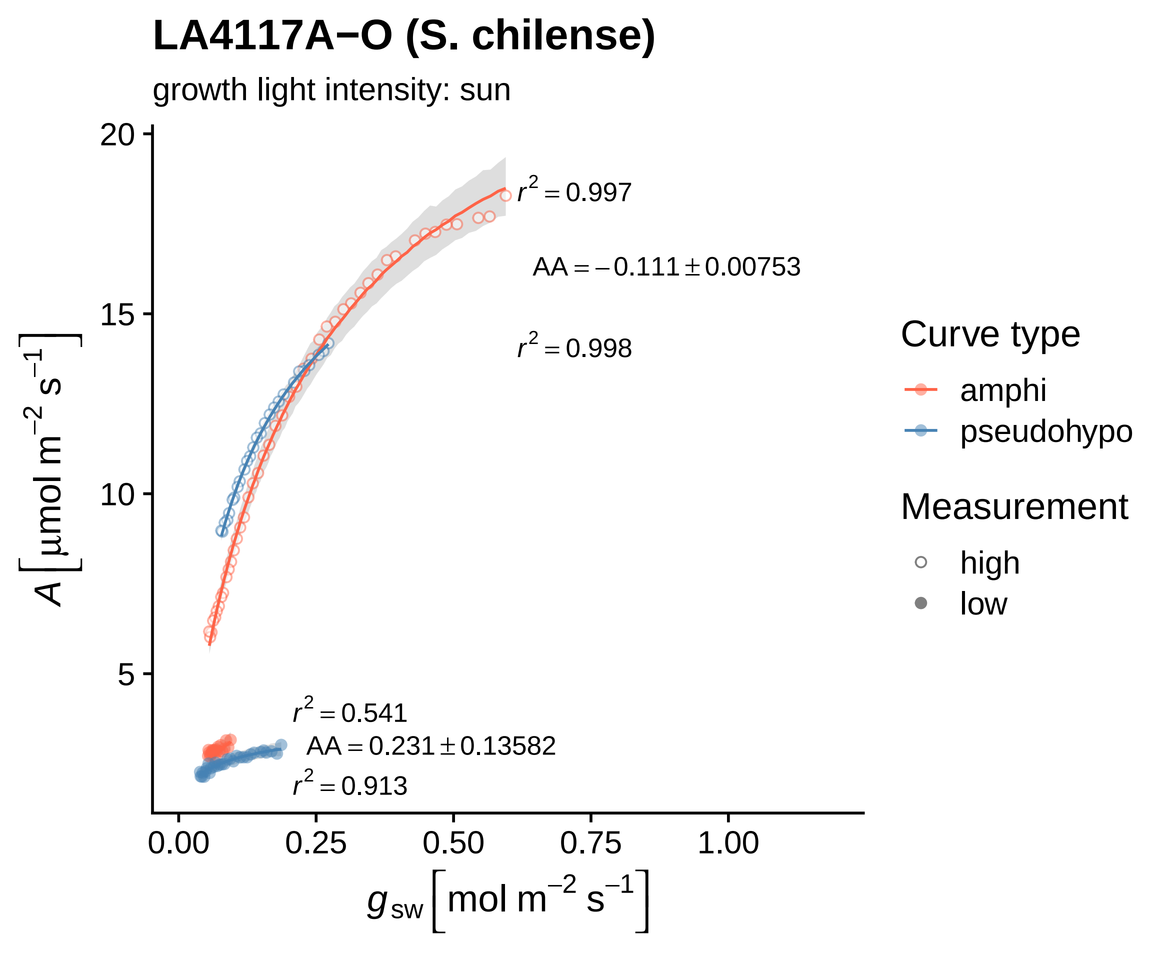

LA4117A−O (S. chilense)
growth light intensity: sun
$r^2 = 0.997$
$AA = -0.111 \pm 0.00753$
$r^2 = 0.998$
$r^2 = 0.541$
$AA = 0.231 \pm 0.13582$
$r^2 = 0.913$
20
15
10
5
0.00
0.25
0.50
0.75
1.00
$A\,[\mu mol\ m^{-2}\ s^{-1}]$
$g_{sw}\,[mol\ m^{-2}\ s^{-1}]$
Curve type
amphi
pseudohypo
Measurement
high
low

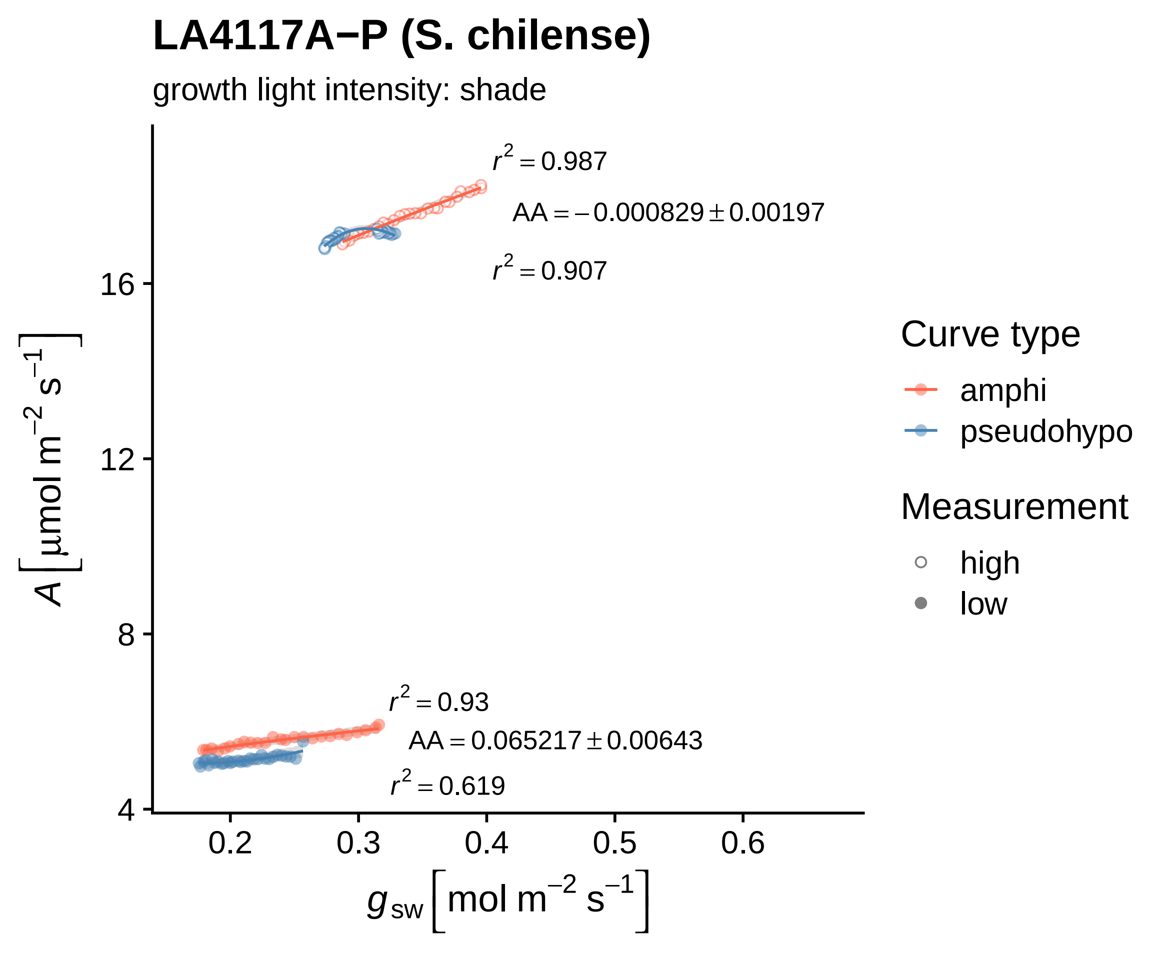

LA4117A−P (S. chilense)
growth light intensity: shade
$r^2 = 0.987$
$AA = -0.000829 \pm 0.00197$
$r^2 = 0.907$
$r^2 = 0.93$
$AA = 0.065217 \pm 0.00643$
$r^2 = 0.619$
16
12
8
4
0.2
0.3
0.4
0.5
0.6
$A\left[\mu mol\ m^{-2}\ s^{-1}\right]$
$g_{sw}\left[mol\ m^{-2}\ s^{-1}\right]$
Curve type
amphi
pseudohypo
Measurement
high
low

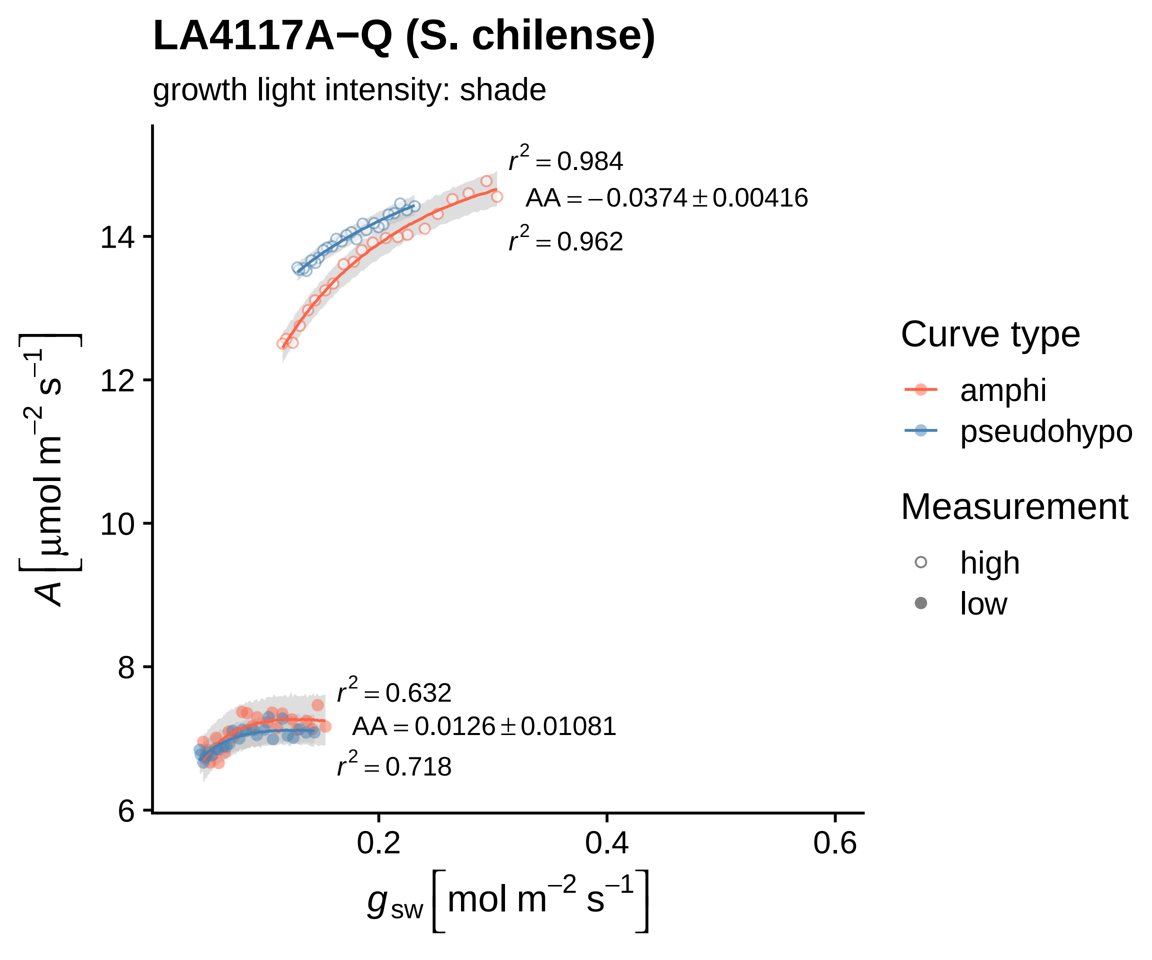

LA4117A−Q (S. chilense)
growth light intensity: shade
r² = 0.984
AA = −0.0374 ± 0.00416
r² = 0.962
r² = 0.632
AA = 0.0126 ± 0.01081
r² = 0.718
A [µmol m⁻² s⁻¹]
14
12
10
8
6
0.2
0.4
0.6
g_sw [mol m⁻² s⁻¹]
Curve type
amphi
pseudohypo
Measurement
high
low

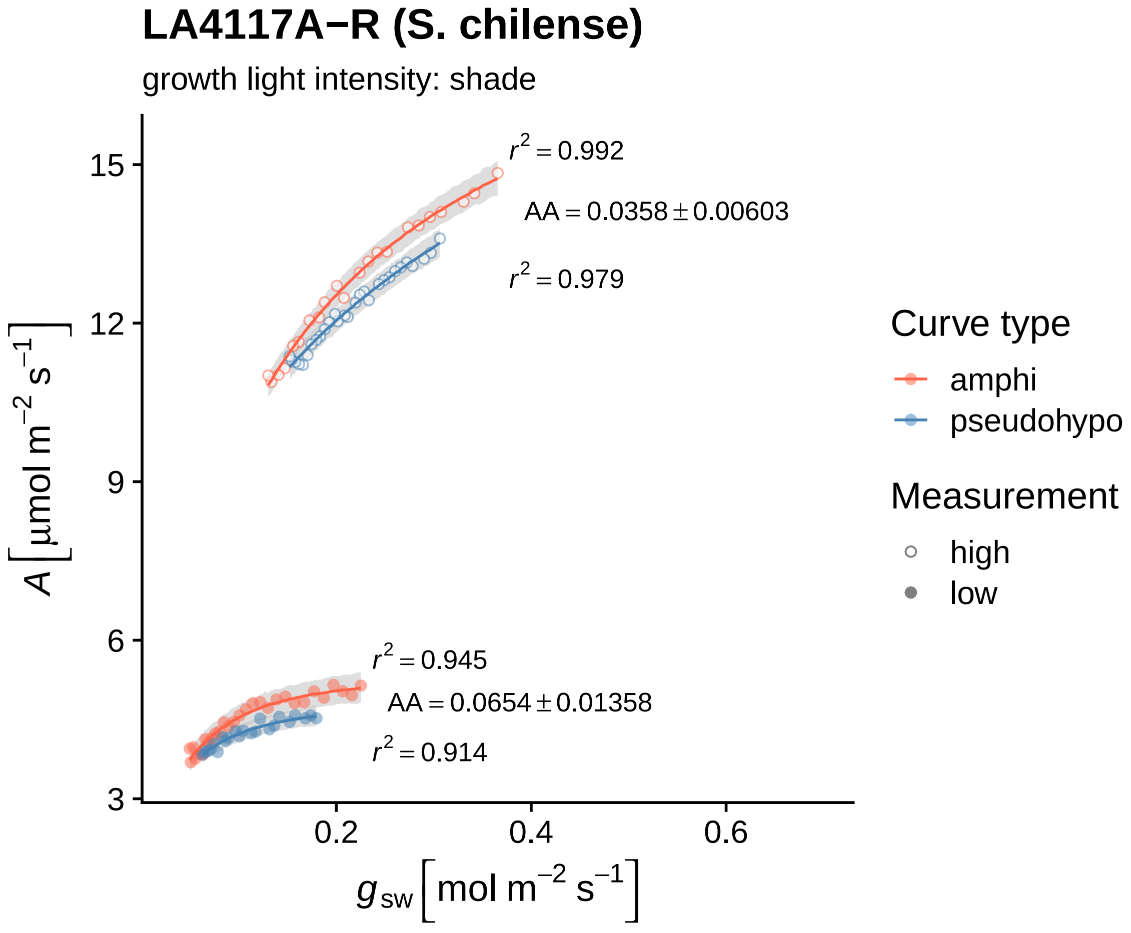

LA4117A−R (S. chilense)
growth light intensity: shade
$r^2 = 0.992$
$AA = 0.0358 \pm 0.00603$
$r^2 = 0.979$
$r^2 = 0.945$
$AA = 0.0654 \pm 0.01358$
$r^2 = 0.914$
15
12
9
6
3
0.2
0.4
0.6
$A\ [\mu mol\ m^{-2}\ s^{-1}]$
$g_{sw}\ [mol\ m^{-2}\ s^{-1}]$
Curve type
amphi
pseudohypo
Measurement
high
low

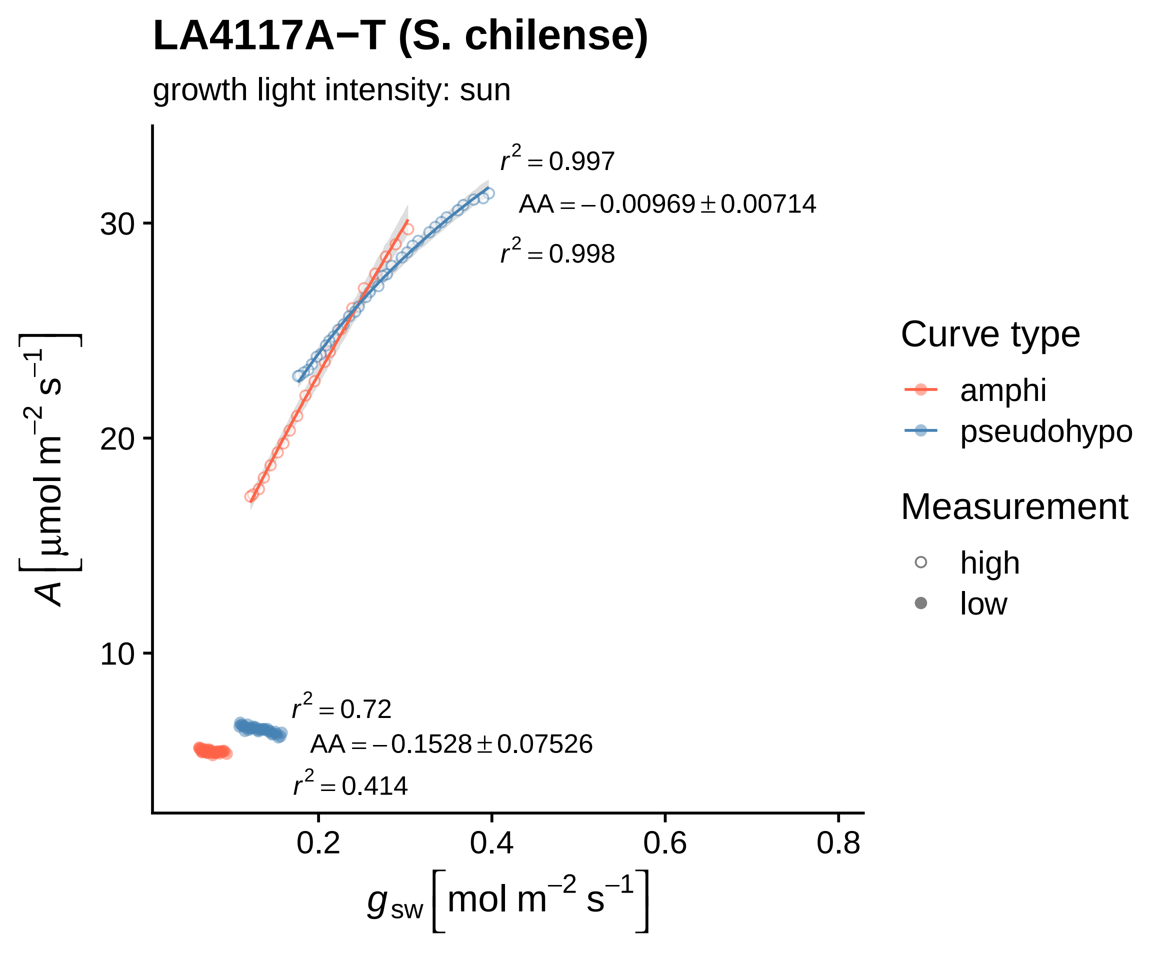

LA4117A−T (S. chilense)
growth light intensity: sun
r² = 0.997
AA = −0.00969 ± 0.00714
r² = 0.998
r² = 0.72
AA = −0.1528 ± 0.07526
r² = 0.414
A [µmol m⁻² s⁻¹]
30
20
10
0.2
0.4
0.6
0.8
g_sw [mol m⁻² s⁻¹]
Curve type
amphi
pseudohypo
Measurement
high
low

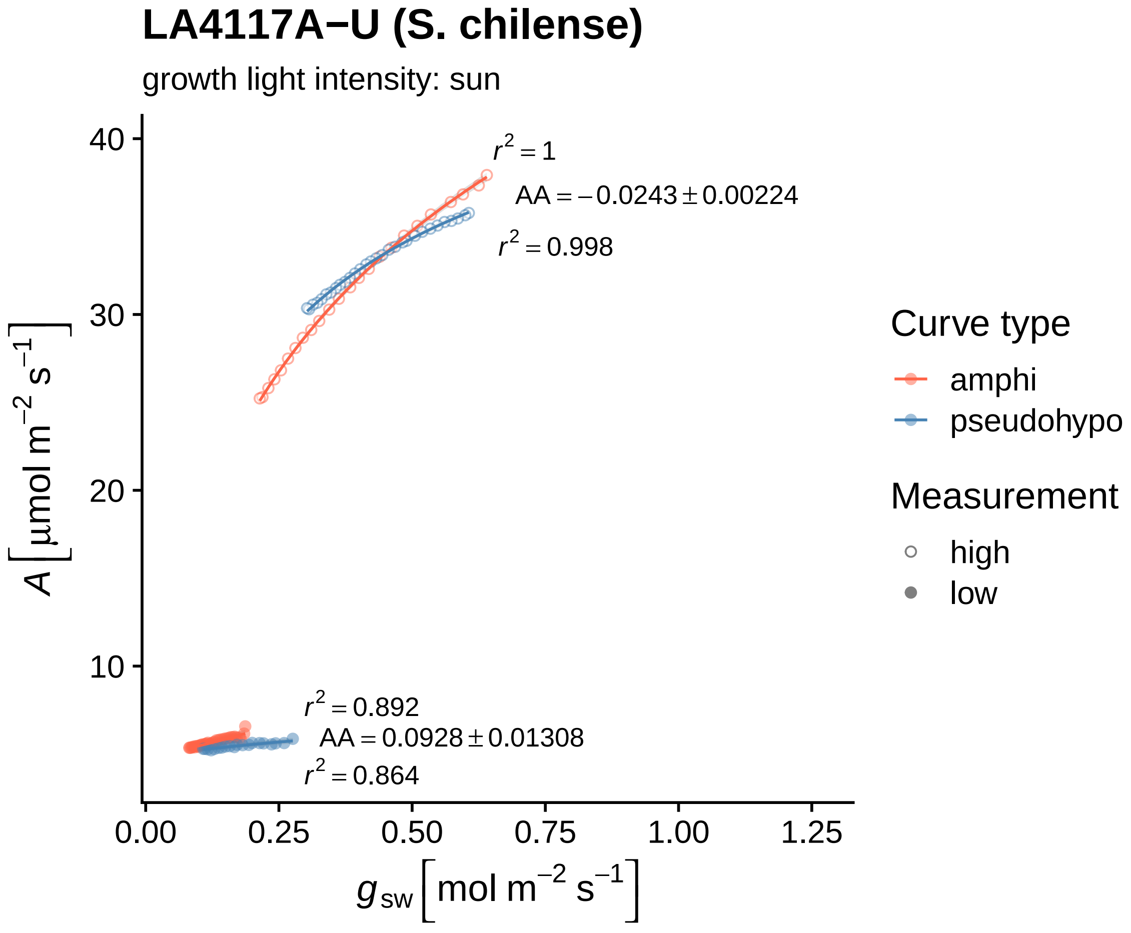

LA4117A−U (S. chilense)
growth light intensity: sun
$r^2 = 1$
$AA = -0.0243 \pm 0.00224$
$r^2 = 0.998$
$r^2 = 0.892$
$AA = 0.0928 \pm 0.01308$
$r^2 = 0.864$
40
30
20
10
0.00
0.25
0.50
0.75
1.00
1.25
$A\,[\mu mol\ m^{-2}\ s^{-1}]$
$g_{sw}\,[mol\ m^{-2}\ s^{-1}]$
Curve type
amphi
pseudohypo
Measurement
high
low

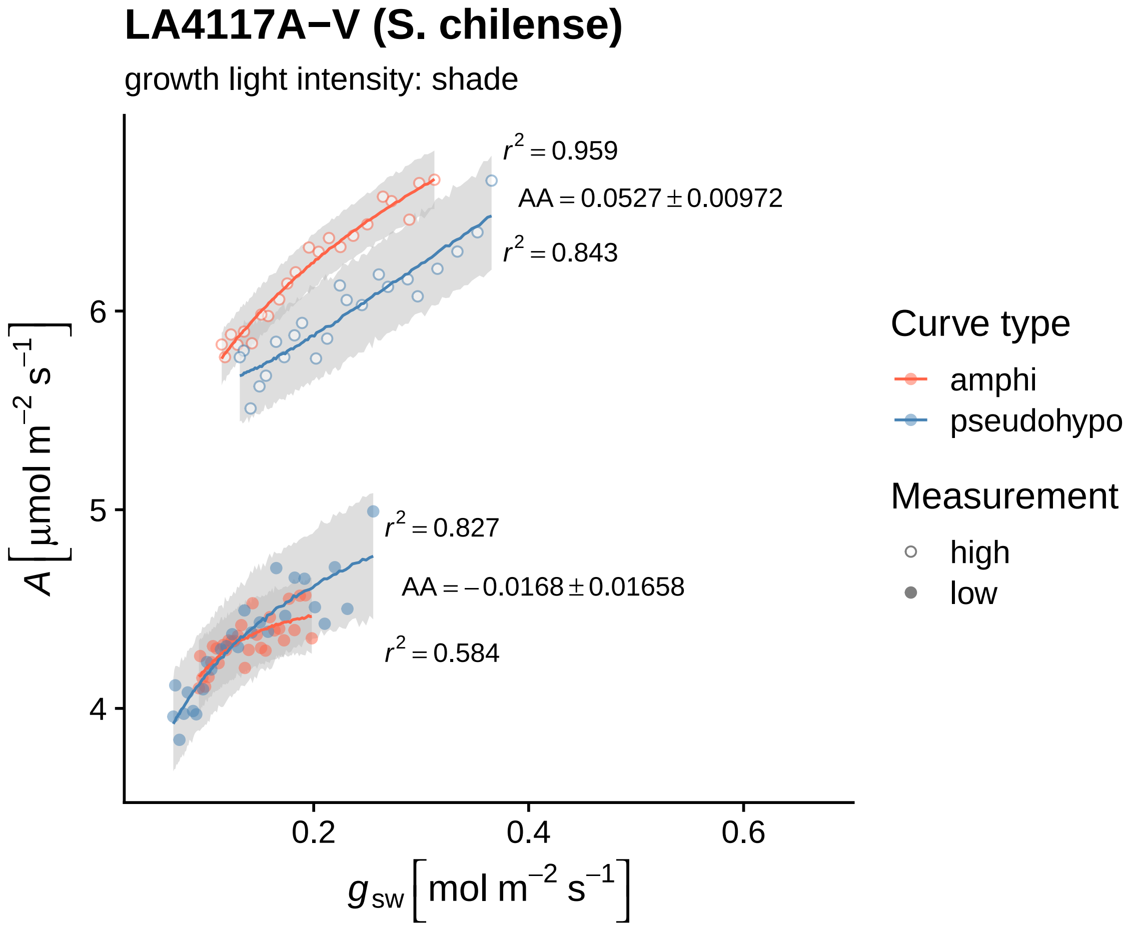
LA4117A−V (S. chilense)
growth light intensity: shade
r² = 0.959
AA = 0.0527 ± 0.00972
r² = 0.843
r² = 0.827
AA = −0.0168 ± 0.01658
r² = 0.584
6
5
4
0.2
0.4
0.6
A [μmol m⁻² s⁻¹]
g_sw [mol m⁻² s⁻¹]
Curve type
amphi
pseudohypo
Measurement
high
low

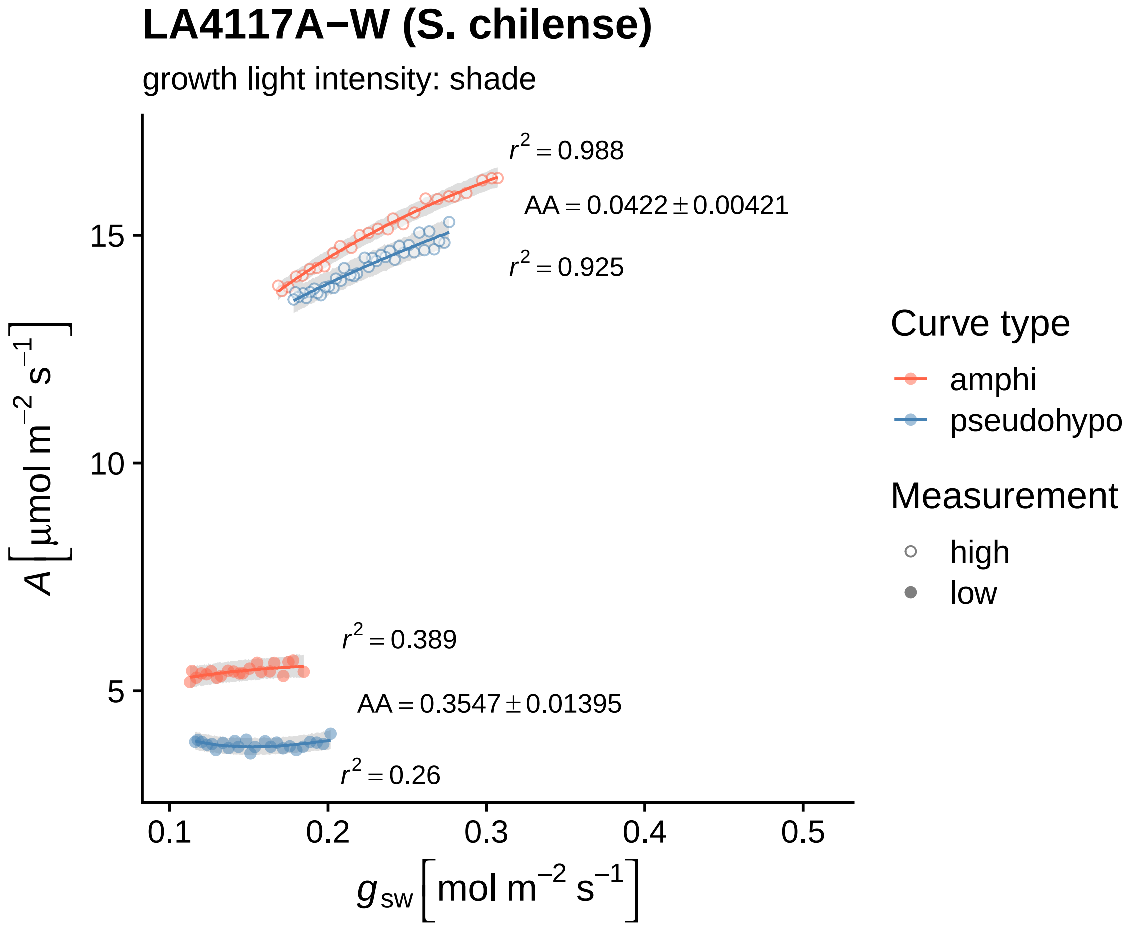

LA4117A−W (S. chilense)
growth light intensity: shade
$r^2 = 0.988$
$AA = 0.0422 \pm 0.00421$
$r^2 = 0.925$
$r^2 = 0.389$
$AA = 0.3547 \pm 0.01395$
$r^2 = 0.26$
15
10
5
0.1
0.2
0.3
0.4
0.5
$A\ [\mu mol\ m^{-2}\ s^{-1}]$
$g_{sw}\ [mol\ m^{-2}\ s^{-1}]$
Curve type
amphi
pseudohypo
Measurement
high
low

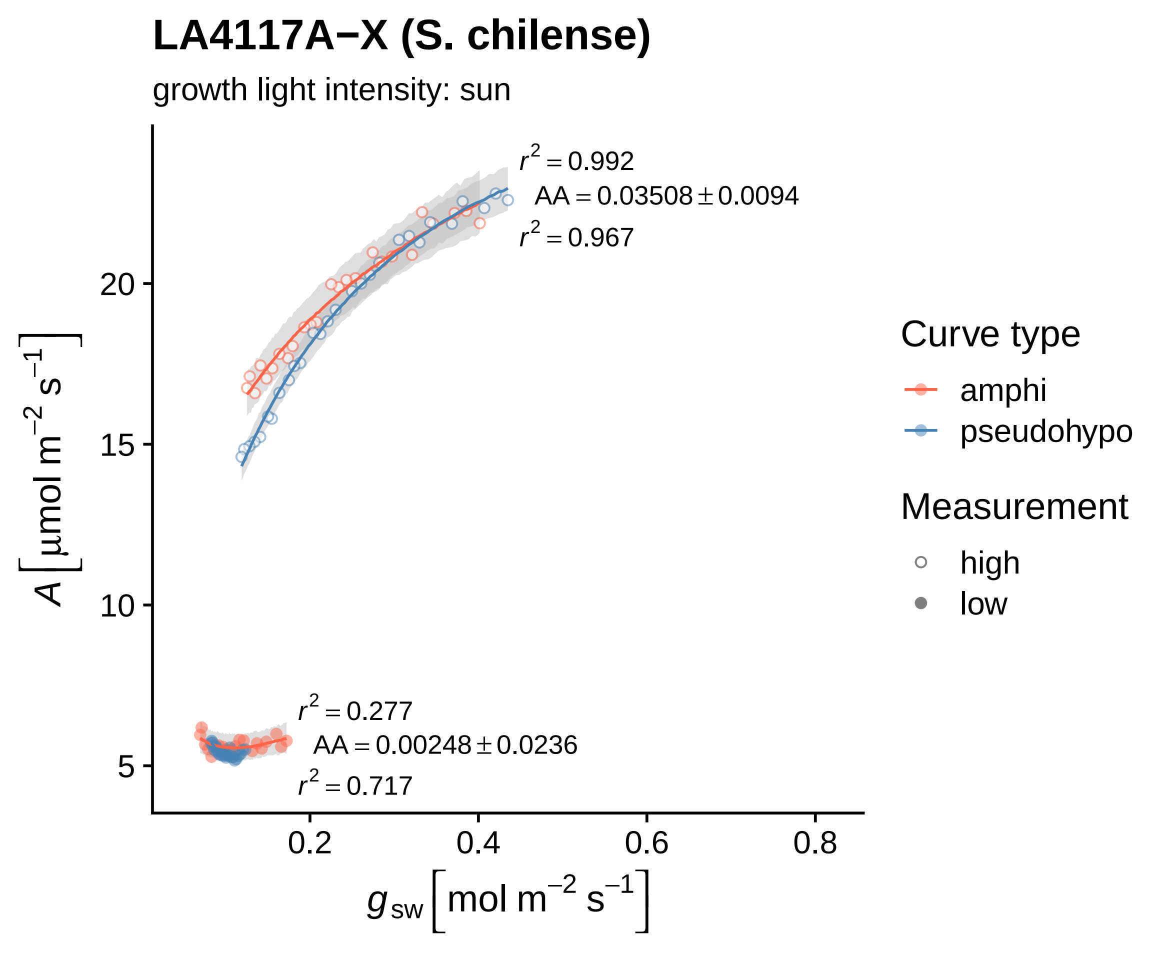
LA4117A−X (S. chilense)
growth light intensity: sun
r² = 0.992
AA = 0.03508 ± 0.0094
r² = 0.967
r² = 0.277
AA = 0.00248 ± 0.0236
r² = 0.717
A [µmol m⁻² s⁻¹]
g_sw [mol m⁻² s⁻¹]
20
15
10
5
0.2
0.4
0.6
0.8
Curve type
amphi
pseudohypo
Measurement
high
low

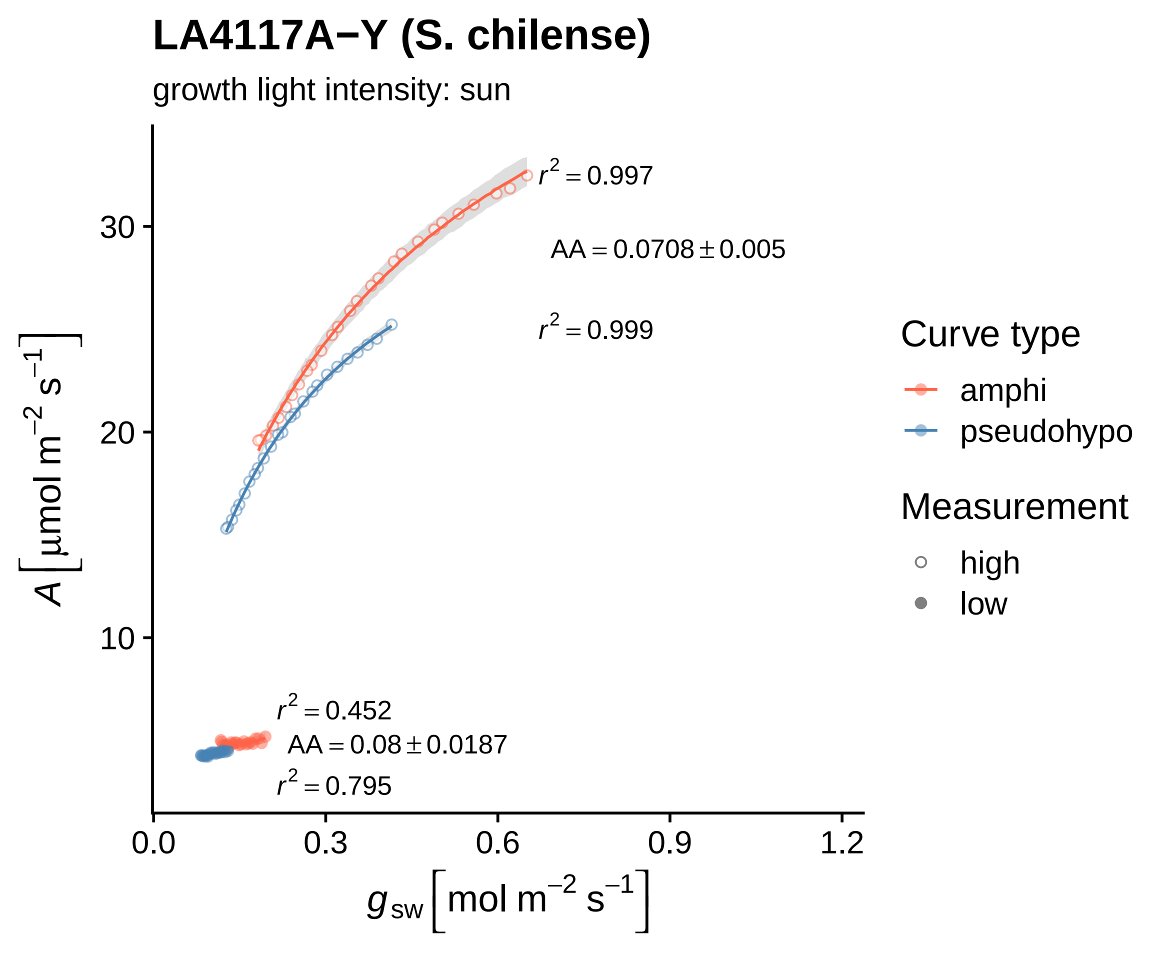

LA4117A−Y (S. chilense)
growth light intensity: sun
A [μmol m−2 s−1]
30
20
10
0.0
0.3
0.6
0.9
1.2
gsw [mol m−2 s−1]
r2 = 0.997
AA = 0.0708 ± 0.005
r2 = 0.999
r2 = 0.452
AA = 0.08 ± 0.0187
r2 = 0.795
Curve type
amphi
pseudohypo
Measurement
high
low

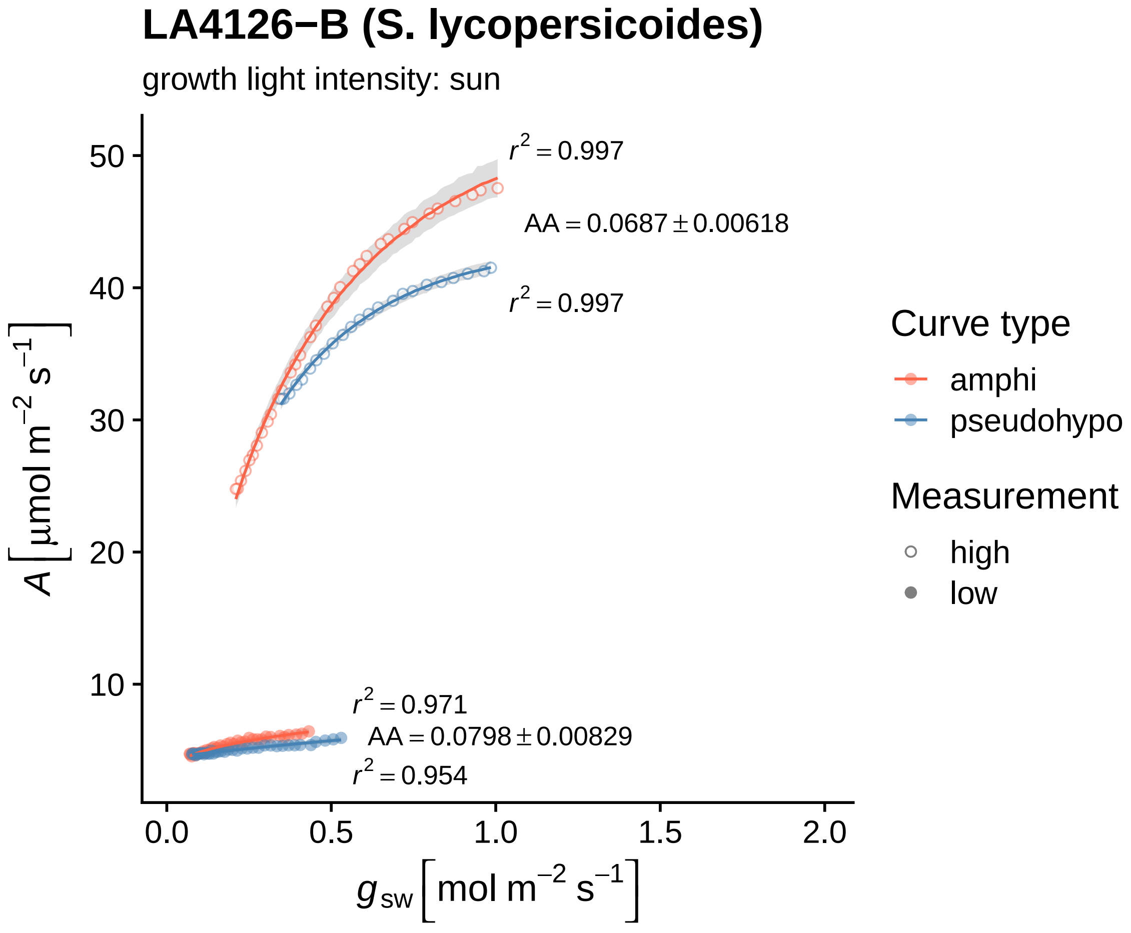

LA4126−B (S. lycopersicoides)
growth light intensity: sun
$r^2 = 0.997$
$AA = 0.0687 \pm 0.00618$
$r^2 = 0.997$
$r^2 = 0.971$
$AA = 0.0798 \pm 0.00829$
$r^2 = 0.954$
50
40
30
20
10
0.0
0.5
1.0
1.5
2.0
$A\ [\mu mol\ m^{-2}\ s^{-1}]$
$g_{sw}\ [mol\ m^{-2}\ s^{-1}]$
Curve type
amphi
pseudohypo
Measurement
high
low

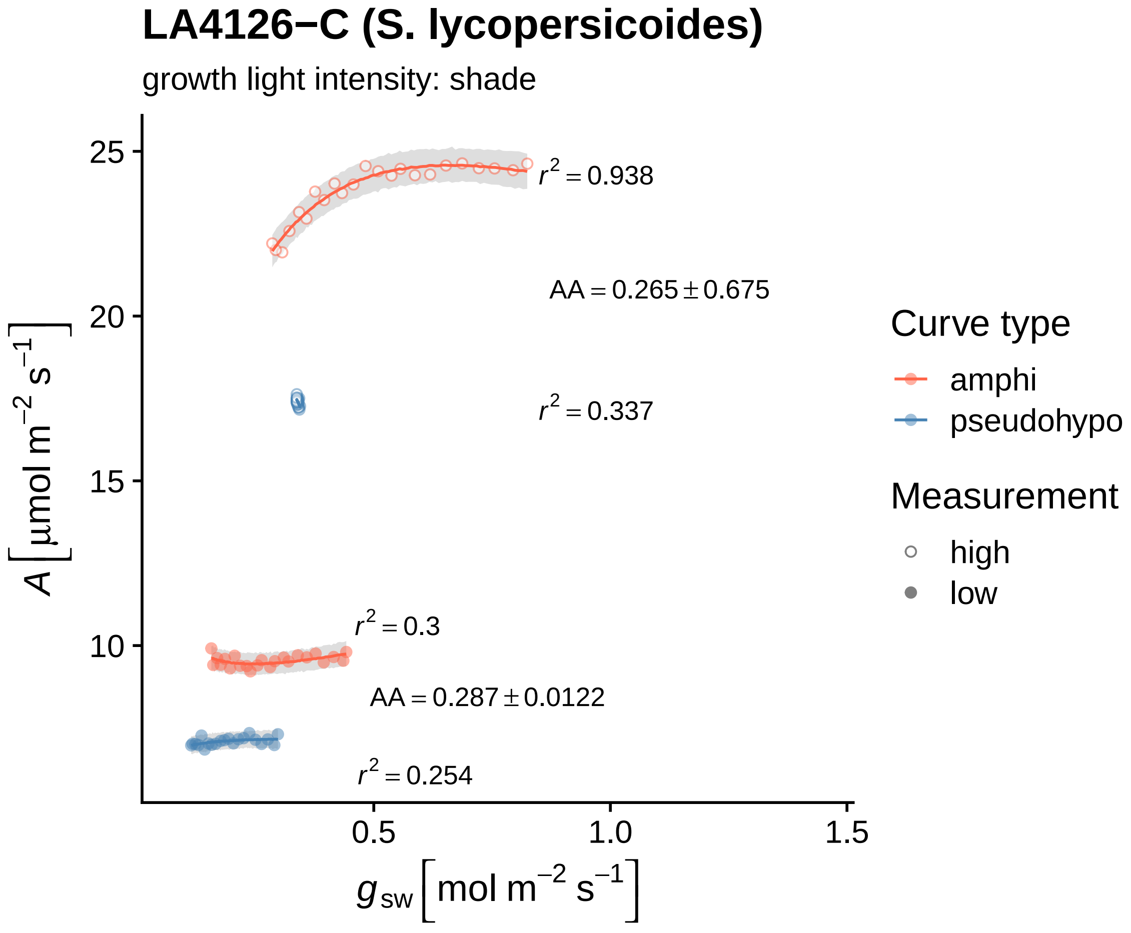

LA4126−C (S. lycopersicoides)
growth light intensity: shade
$r^2 = 0.938$
$AA = 0.265 \pm 0.675$
$r^2 = 0.337$
$r^2 = 0.3$
$AA = 0.287 \pm 0.0122$
$r^2 = 0.254$
25
20
15
10
0.5
1.0
1.5
$A\ [\mu mol\ m^{-2}\ s^{-1}]$
$g_{sw}\ [mol\ m^{-2}\ s^{-1}]$
Curve type
amphi
pseudohypo
Measurement
high
low

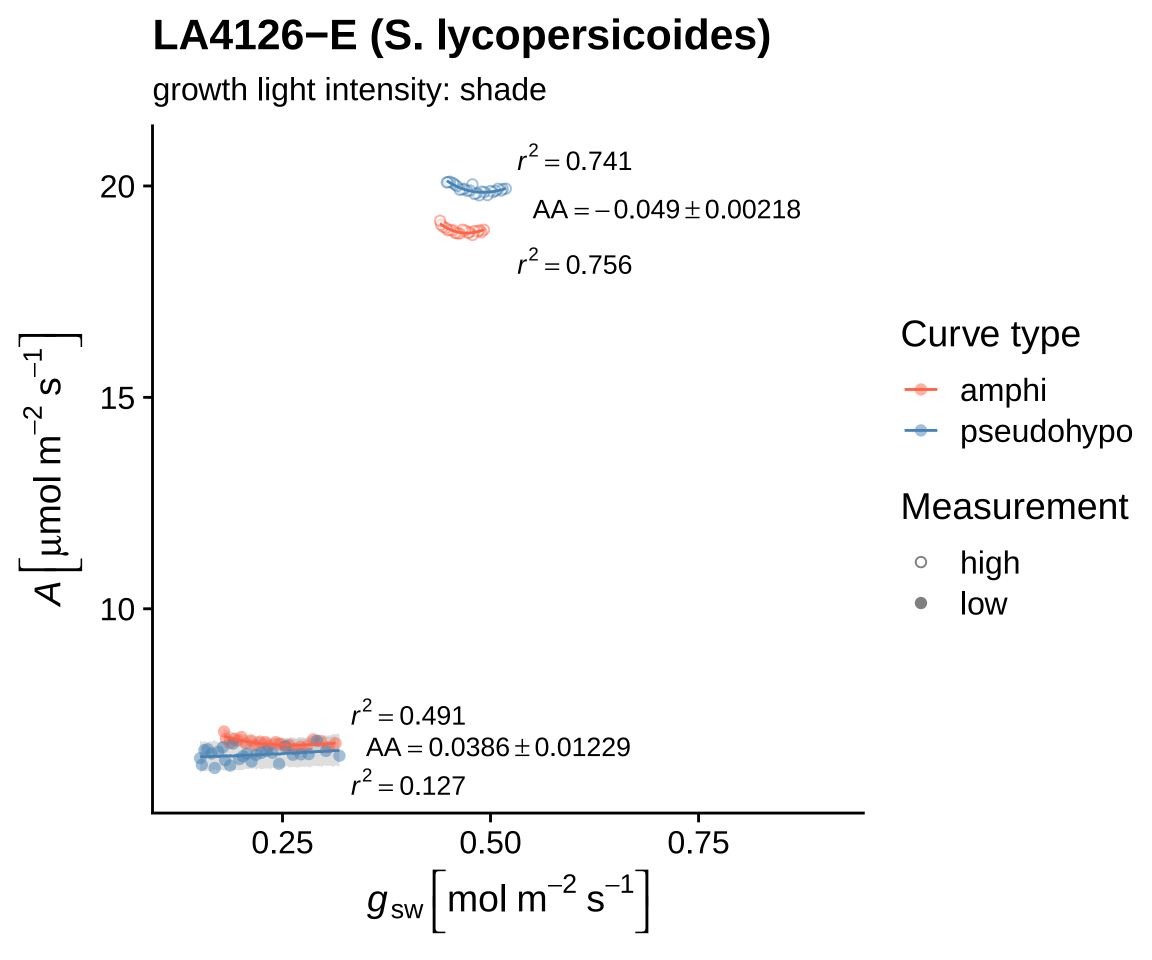
LA4126−E (S. lycopersicoides)
growth light intensity: shade
$r^2 = 0.741$
$AA = -0.049 \pm 0.00218$
$r^2 = 0.756$
$r^2 = 0.491$
$AA = 0.0386 \pm 0.01229$
$r^2 = 0.127$
20
15
10
0.25
0.50
0.75
$A$ [µmol m$^{-2}$ s$^{-1}$]
$g_{sw}$ [mol m$^{-2}$ s$^{-1}$]
Curve type
amphi
pseudohypo
Measurement
high
low

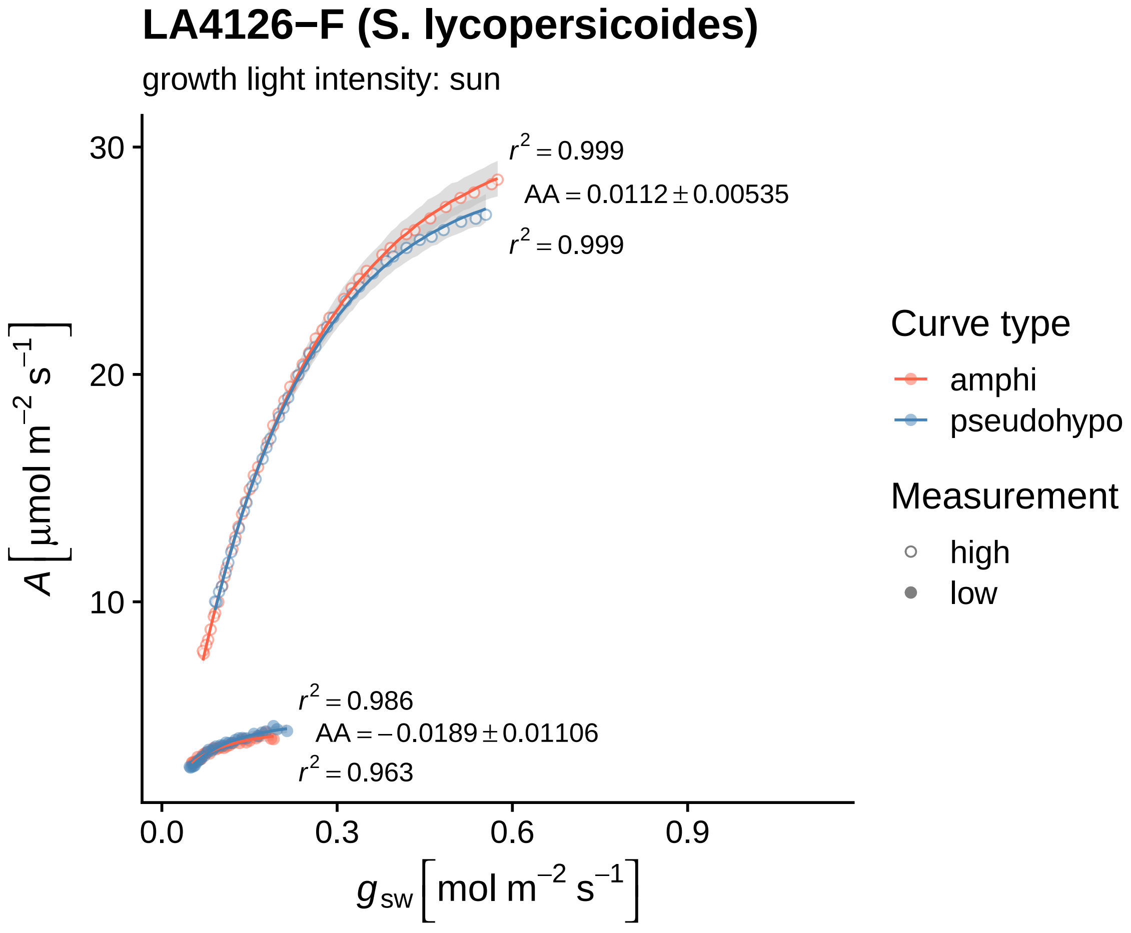

LA4126−F (S. lycopersicoides)
growth light intensity: sun
r² = 0.999
AA = 0.0112 ± 0.00535
r² = 0.999
r² = 0.986
AA = −0.0189 ± 0.01106
r² = 0.963
30
20
10
0.0
0.3
0.6
0.9
A [µmol m⁻² s⁻¹]
g_sw [mol m⁻² s⁻¹]
Curve type
amphi
pseudohypo
Measurement
high
low

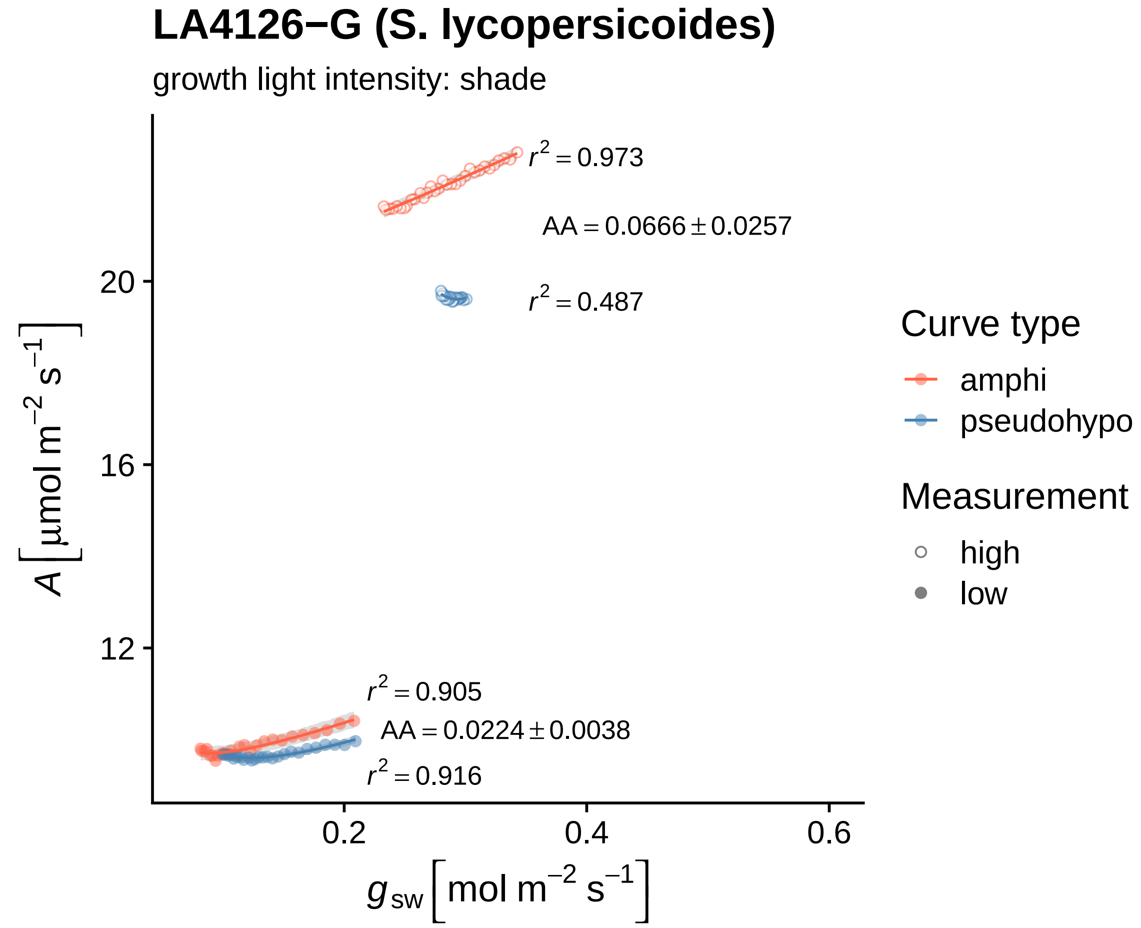

LA4126−G (S. lycopersicoides)
growth light intensity: shade
r² = 0.973
AA = 0.0666 ± 0.0257
r² = 0.487
r² = 0.905
AA = 0.0224 ± 0.0038
r² = 0.916
20
16
12
0.2
0.4
0.6
A [μmol m⁻² s⁻¹]
g_sw [mol m⁻² s⁻¹]
Curve type
amphi
pseudohypo
Measurement
high
low

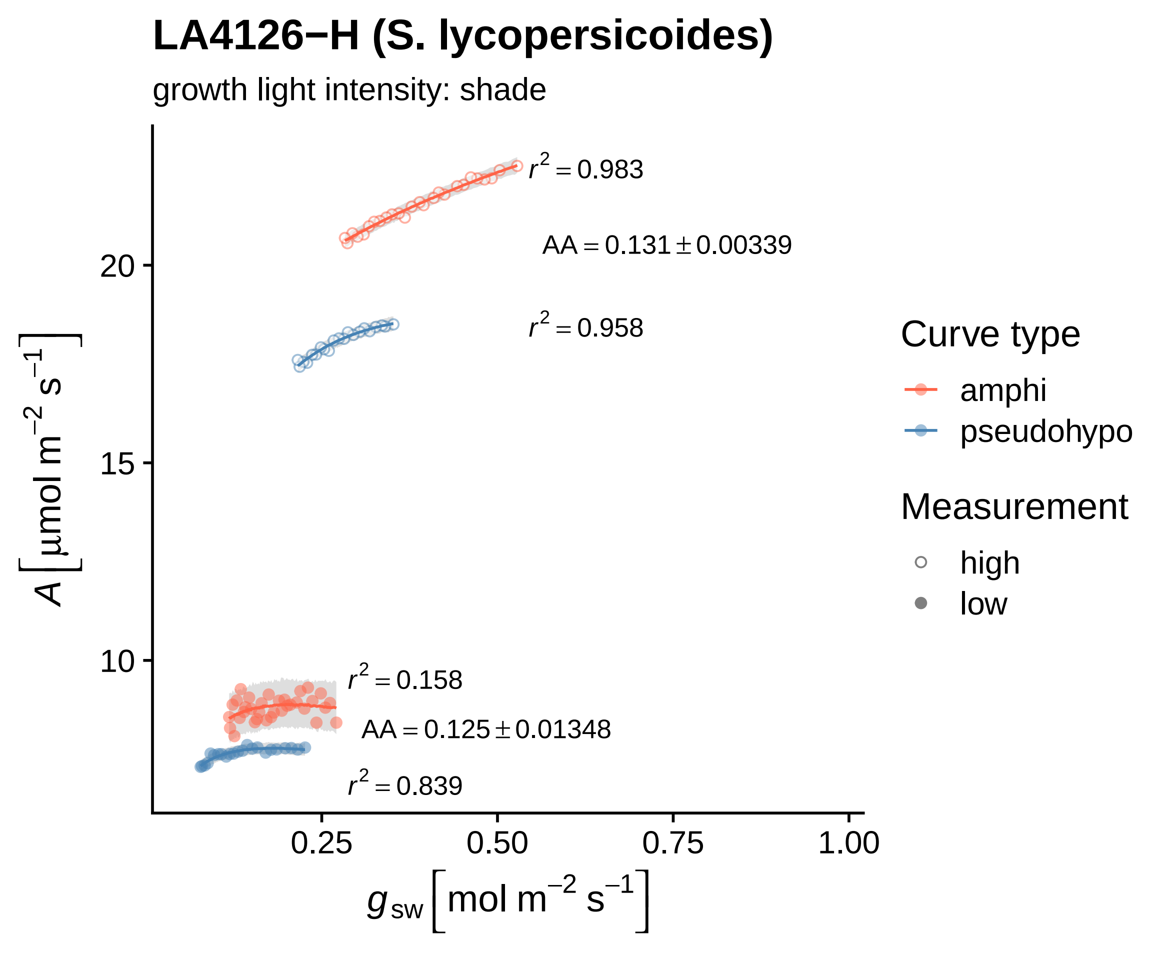
LA4126−H (S. lycopersicoides)
growth light intensity: shade
$r^2 = 0.983$
$AA = 0.131 \pm 0.00339$
$r^2 = 0.958$
$r^2 = 0.158$
$AA = 0.125 \pm 0.01348$
$r^2 = 0.839$
20
15
10
0.25
0.50
0.75
1.00
$A\ [\mu mol\ m^{-2}\ s^{-1}]$
$g_{sw}\ [mol\ m^{-2}\ s^{-1}]$
Curve type
amphi
pseudohypo
Measurement
high
low

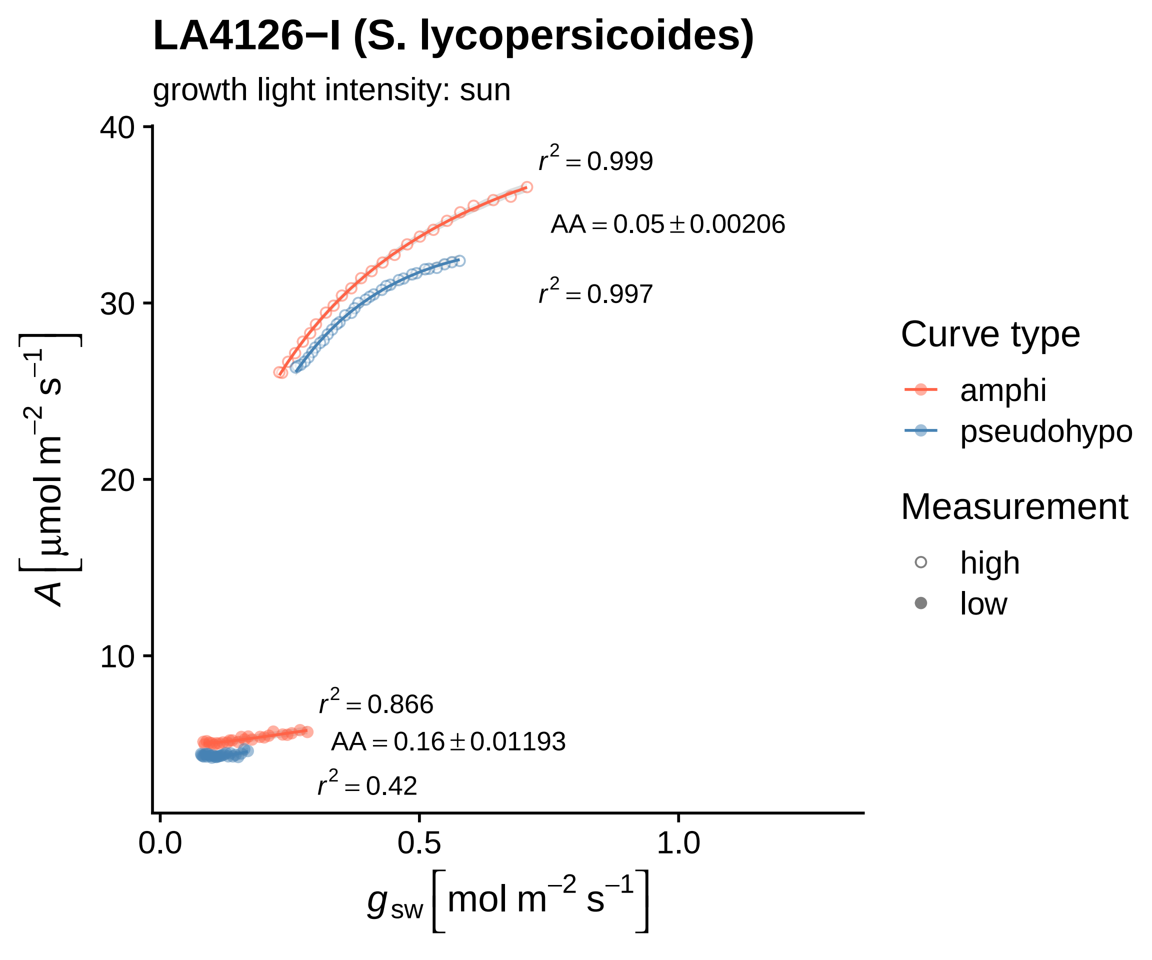
LA4126−l (S. lycopersicoides)
growth light intensity: sun
40
30
20
10
0.0
0.5
1.0
r² = 0.999
AA = 0.05 ± 0.00206
r² = 0.997
r² = 0.866
AA = 0.16 ± 0.01193
r² = 0.42
A [μmol m⁻² s⁻¹]
g_sw [mol m⁻² s⁻¹]
Curve type
amphi
pseudohypo
Measurement
high
low

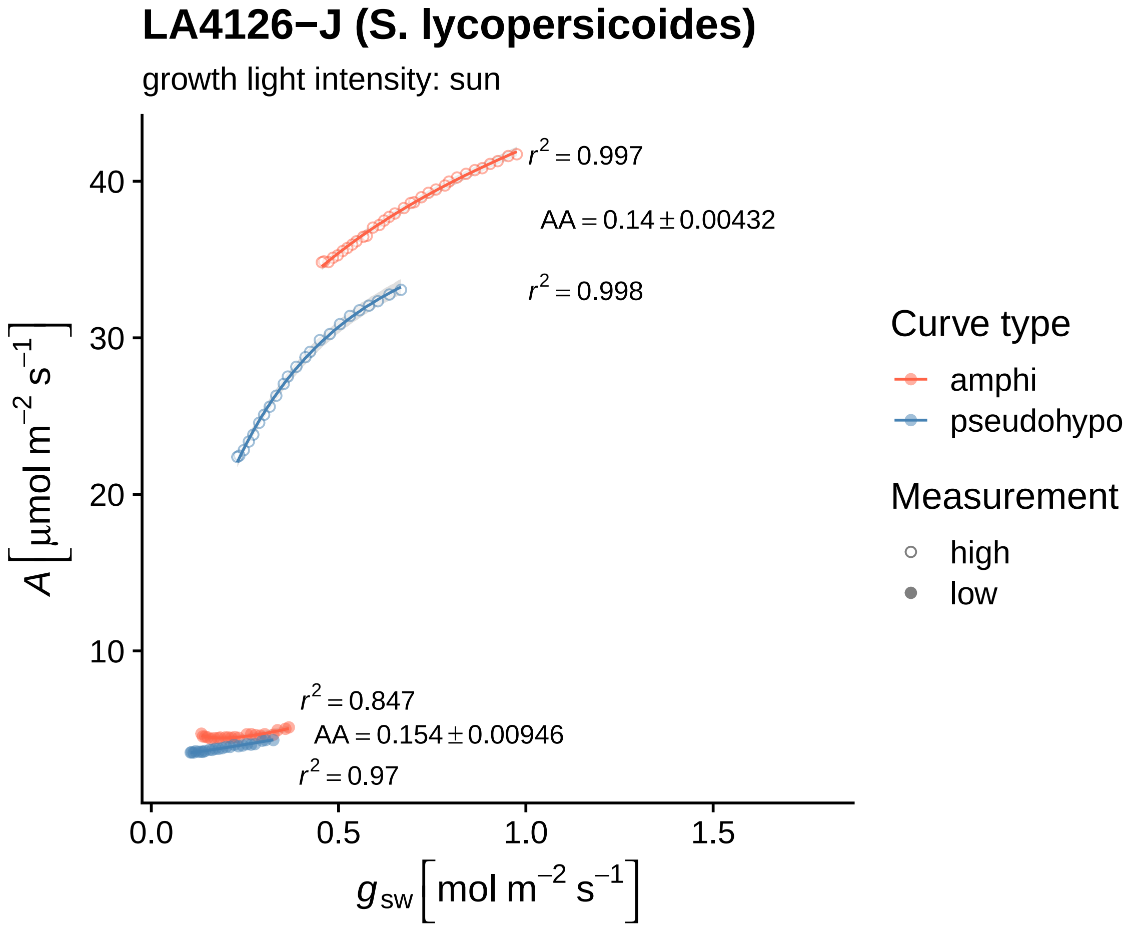

LA4126−J (S. lycopersicoides)
growth light intensity: sun
$r^2 = 0.997$
$AA = 0.14 \pm 0.00432$
$r^2 = 0.998$
$r^2 = 0.847$
$AA = 0.154 \pm 0.00946$
$r^2 = 0.97$
$A\ [\mu mol\ m^{-2}\ s^{-1}]$
$g_{sw}\ [mol\ m^{-2}\ s^{-1}]$
40
30
20
10
0.0
0.5
1.0
1.5
Curve type
amphi
pseudohypo
Measurement
high
low

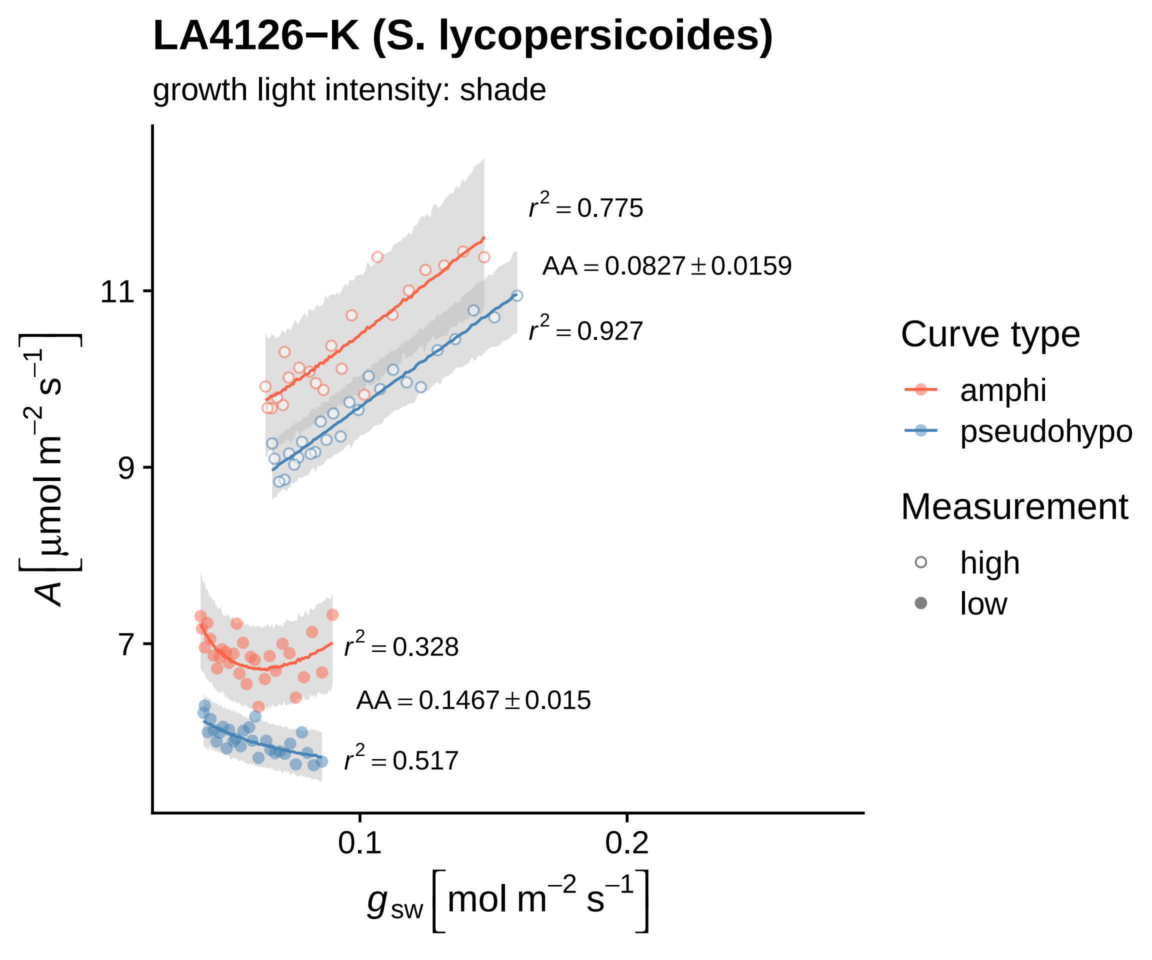

LA4126−K (S. lycopersicoides)
growth light intensity: shade
$r^2 = 0.775$
$AA = 0.0827 \pm 0.0159$
$r^2 = 0.927$
$r^2 = 0.328$
$AA = 0.1467 \pm 0.015$
$r^2 = 0.517$
11
9
7
0.1
0.2
$A\,[\mu mol\ m^{-2}\ s^{-1}]$
$g_{sw}\,[mol\ m^{-2}\ s^{-1}]$
Curve type
amphi
pseudohypo
Measurement
high
low

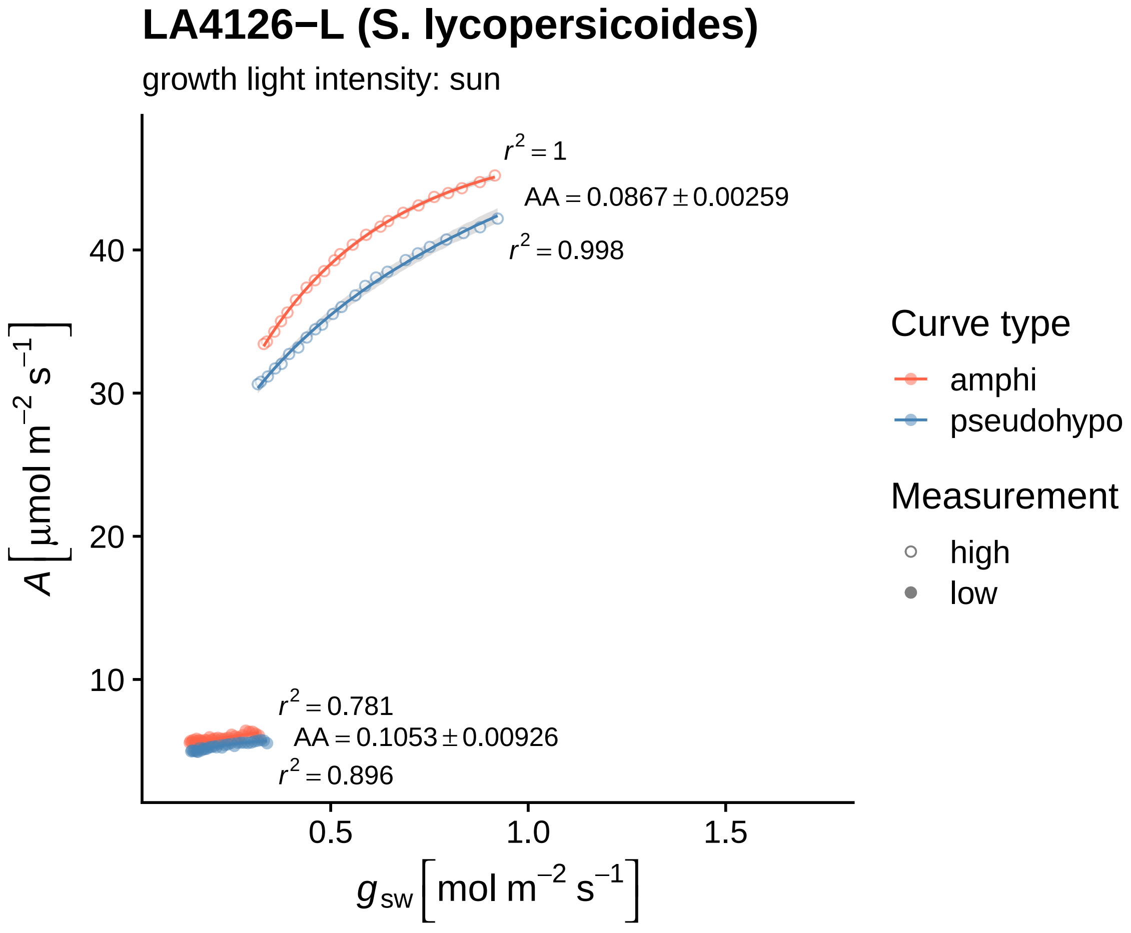
LA4126−L (S. lycopersicoides)
growth light intensity: sun
$r^2 = 1$
AA = 0.0867 ± 0.00259
$r^2 = 0.998$
$r^2 = 0.781$
AA = 0.1053 ± 0.00926
$r^2 = 0.896$
40
30
20
10
0.5
1.0
1.5
$A$ [µmol m$^{-2}$ s$^{-1}$]
$g_{sw}$ [mol m$^{-2}$ s$^{-1}$]
Curve type
amphi
pseudohypo
Measurement
high
low

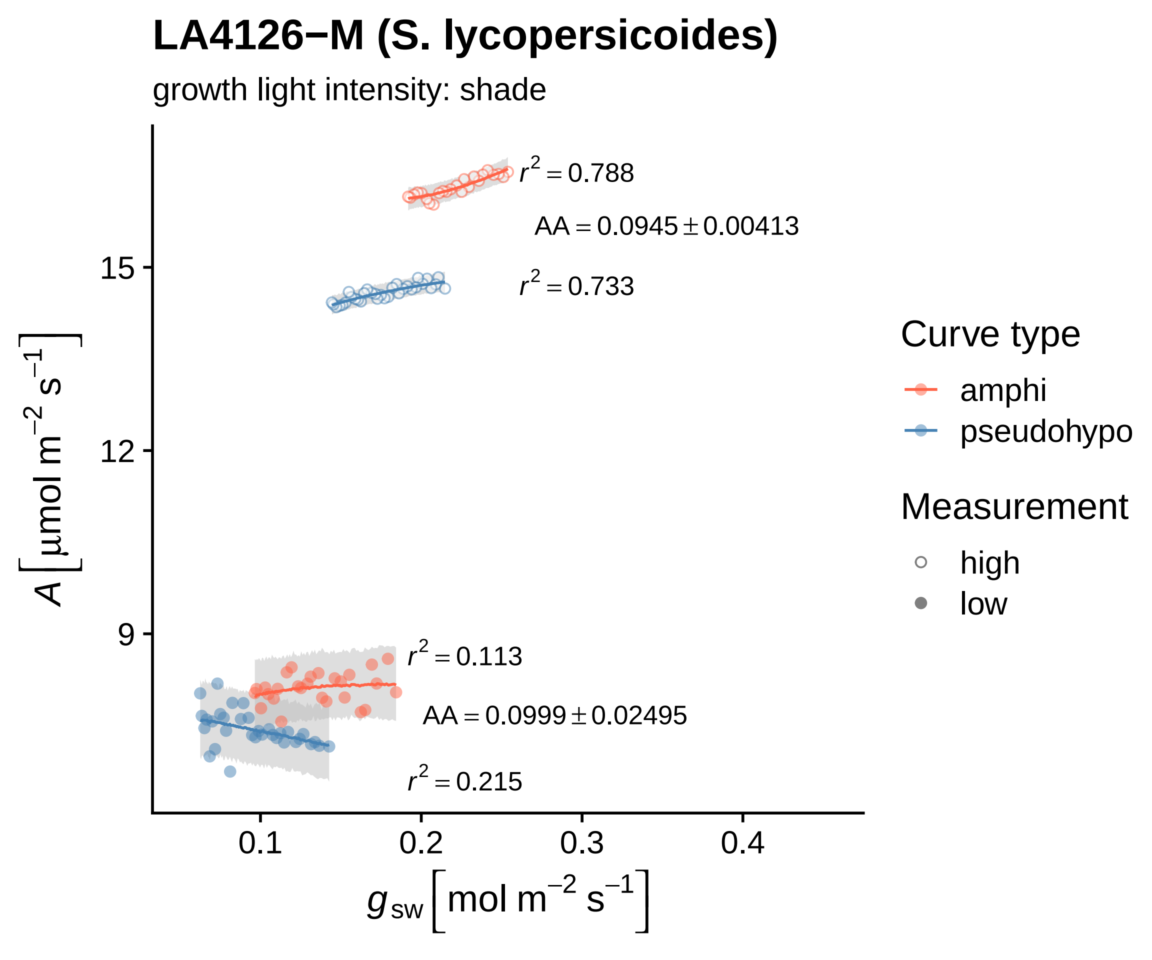
LA4126−M (S. lycopersicoides)
growth light intensity: shade
$r^2 = 0.788$
$AA = 0.0945 \pm 0.00413$
$r^2 = 0.733$
$r^2 = 0.113$
$AA = 0.0999 \pm 0.02495$
$r^2 = 0.215$
15
12
9
0.1
0.2
0.3
0.4
$A\ [\mu mol\ m^{-2}\ s^{-1}]$
$g_{sw}\ [mol\ m^{-2}\ s^{-1}]$
Curve type
amphi
pseudohypo
Measurement
high
low

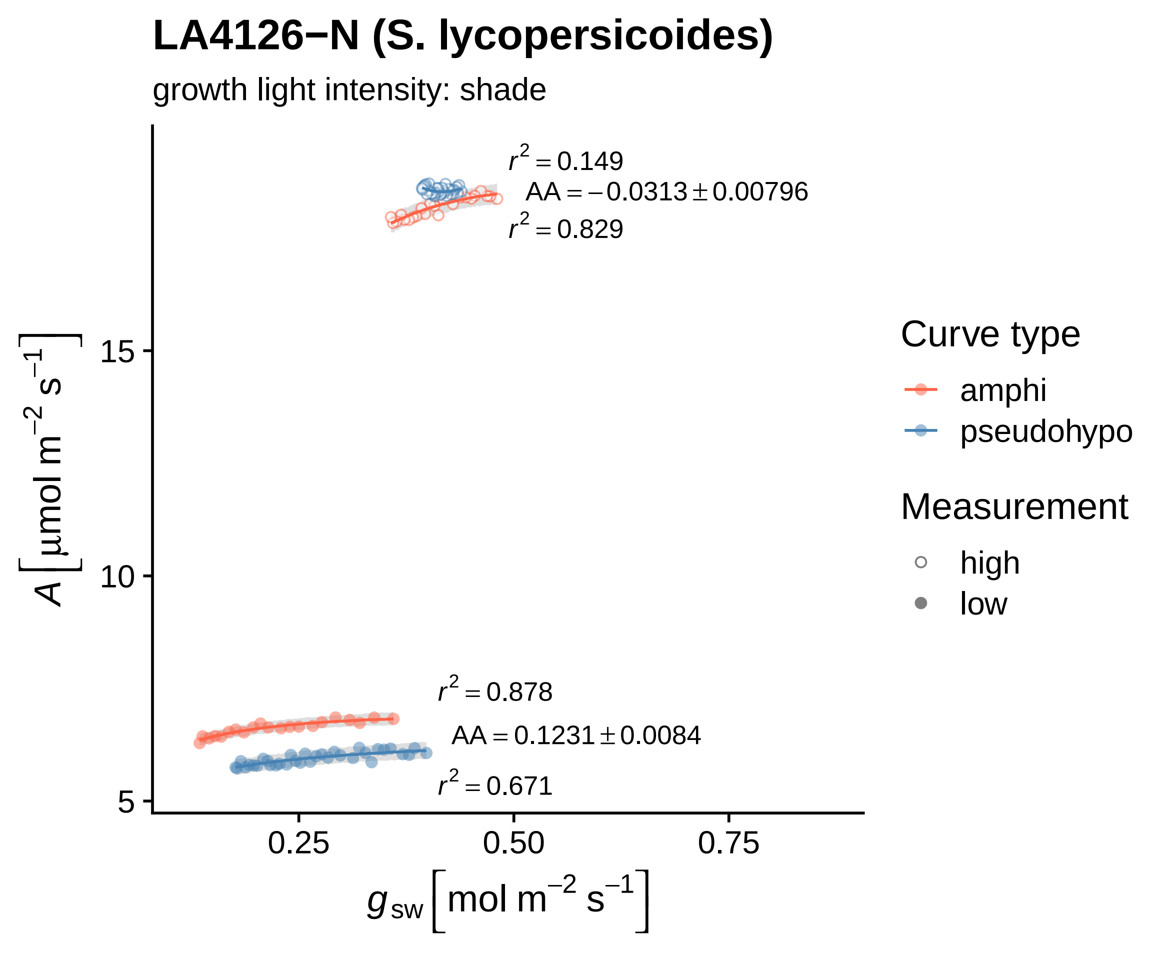
LA4126−N (S. lycopersicoides)
growth light intensity: shade
r² = 0.149
AA = −0.0313 ± 0.00796
r² = 0.829
r² = 0.878
AA = 0.1231 ± 0.0084
r² = 0.671
A [µmol m⁻² s⁻¹]
15
10
5
0.25
0.50
0.75
g sw [mol m⁻² s⁻¹]
Curve type
amphi
pseudohypo
Measurement
high
low

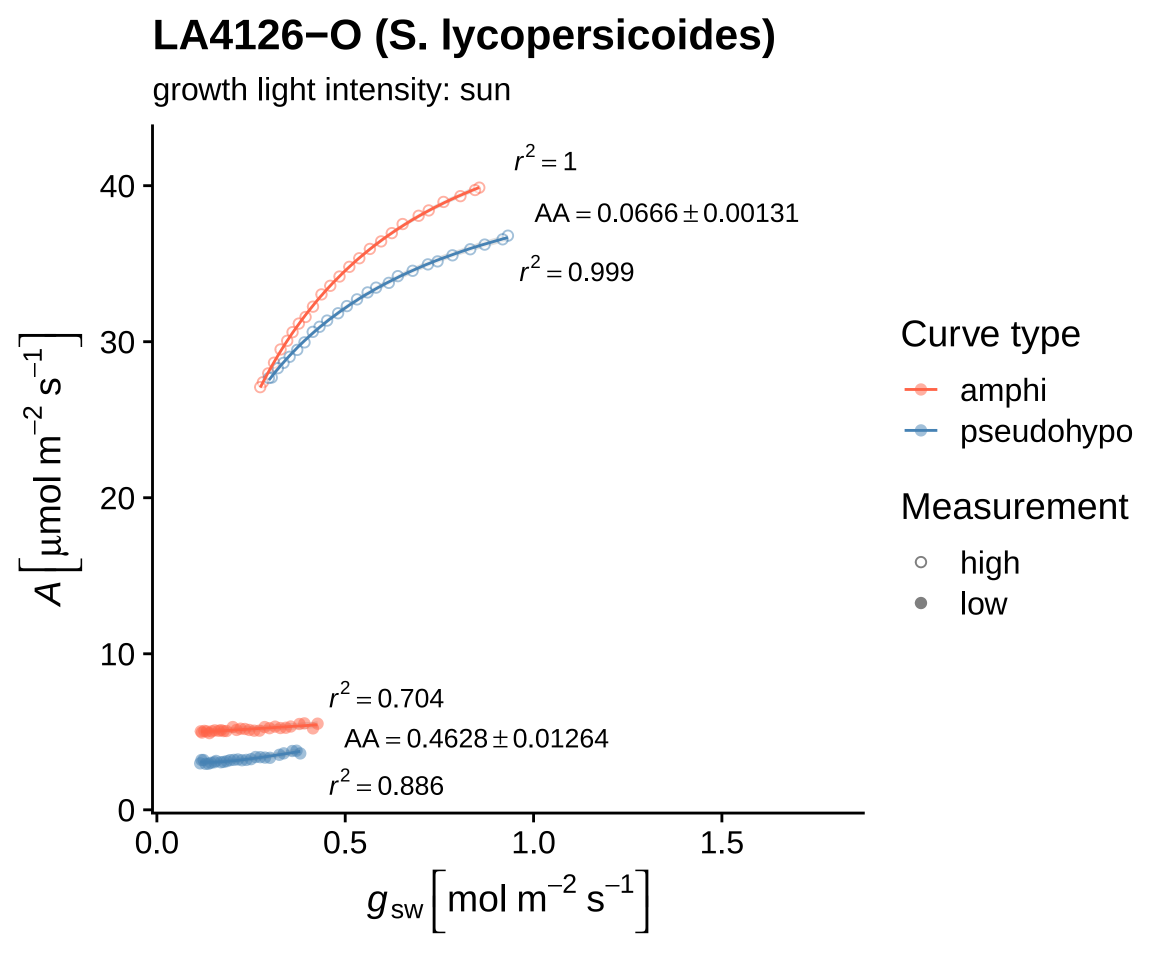
LA4126−O (S. lycopersicoides)
growth light intensity: sun
$r^2 = 1$
$AA = 0.0666 \pm 0.00131$
$r^2 = 0.999$
$r^2 = 0.704$
$AA = 0.4628 \pm 0.01264$
$r^2 = 0.886$
40
30
20
10
0
0.0
0.5
1.0
1.5
$A\,[\mu mol\ m^{-2}\ s^{-1}]$
$g_{sw}\,[mol\ m^{-2}\ s^{-1}]$
Curve type
amphi
pseudohypo
Measurement
high
low

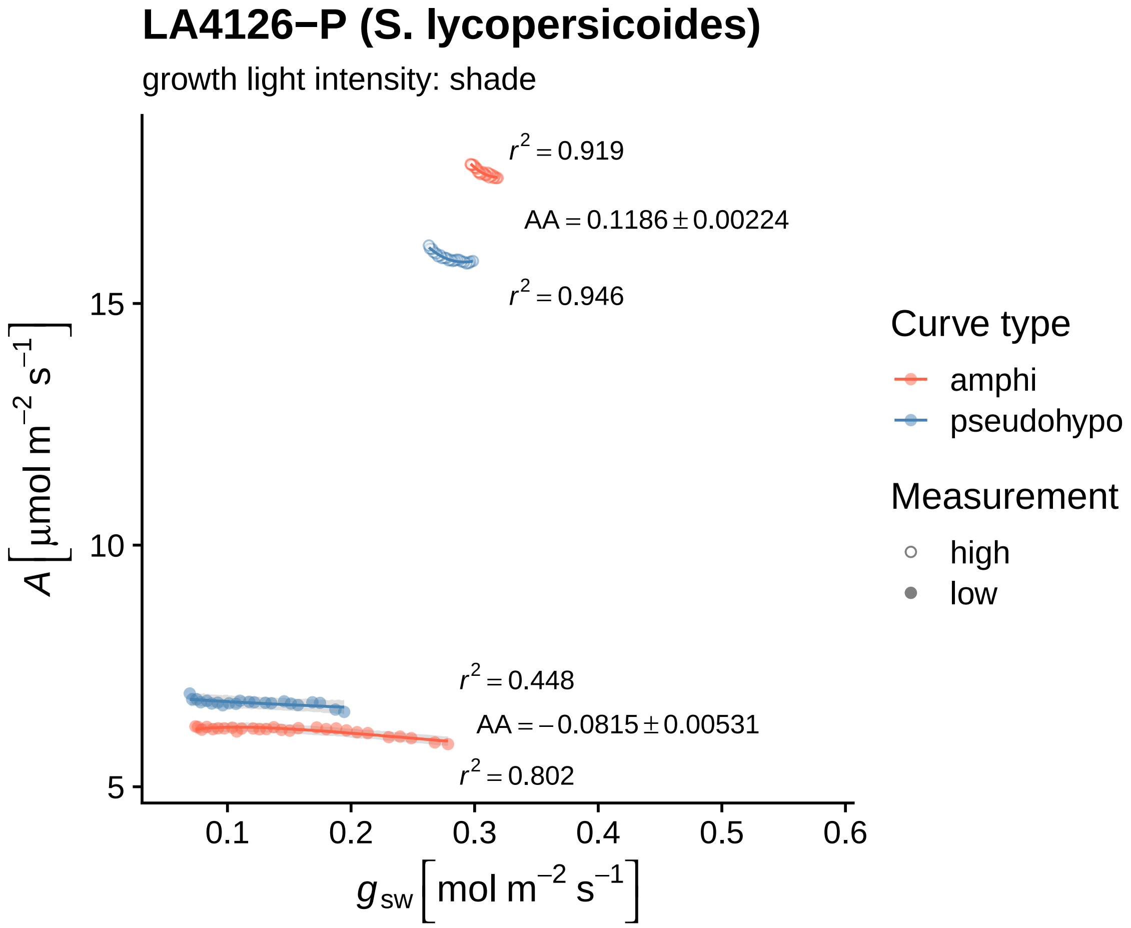

LA4126−P (S. lycopersicoides)
growth light intensity: shade
r² = 0.919
AA = 0.1186 ± 0.00224
r² = 0.946
r² = 0.448
AA = −0.0815 ± 0.00531
r² = 0.802
15
10
5
0.1
0.2
0.3
0.4
0.5
0.6
A [μmol m⁻² s⁻¹]
g_sw [mol m⁻² s⁻¹]
Curve type
amphi
pseudohypo
Measurement
high
low

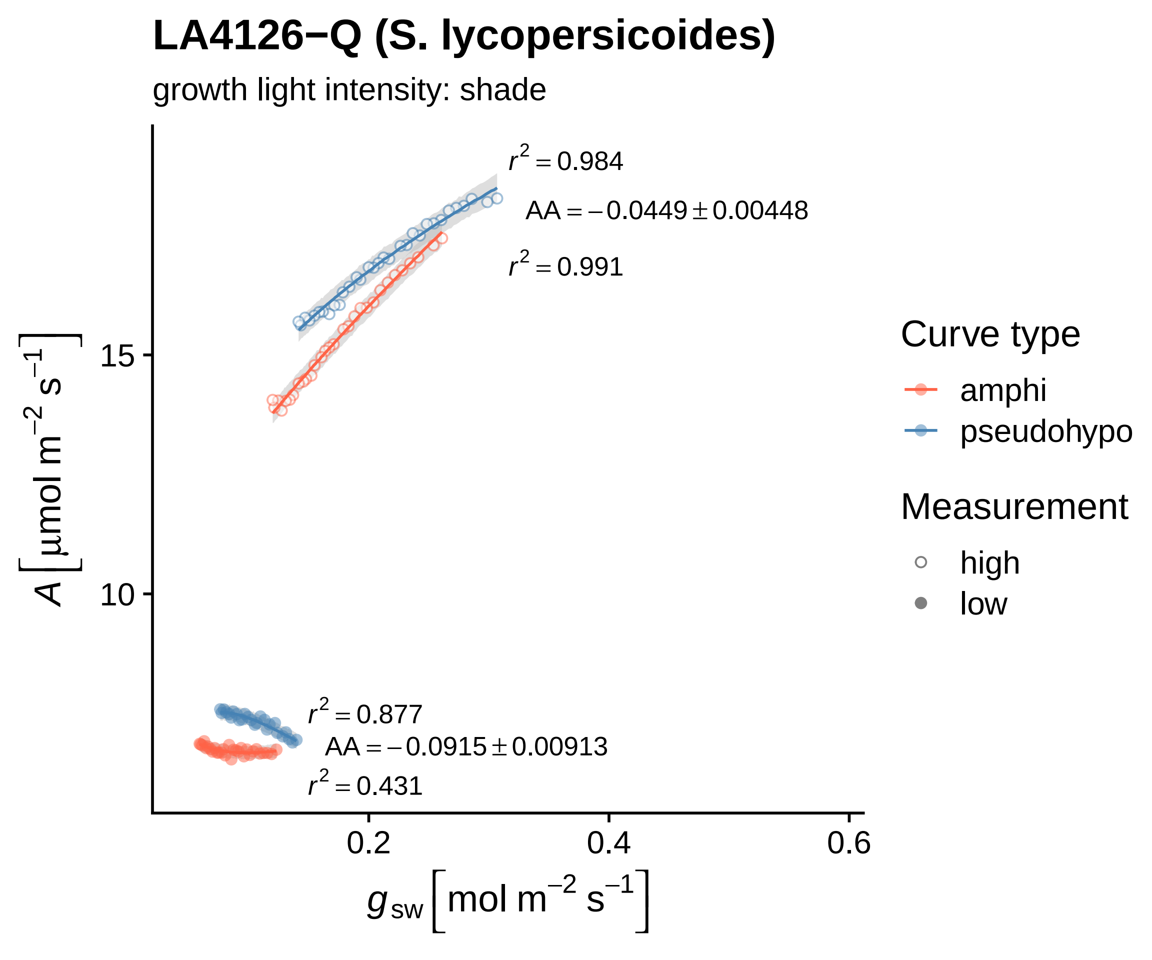
LA4126−Q (S. lycopersicoides)
growth light intensity: shade
$r^2 = 0.984$
$AA = -0.0449 \pm 0.00448$
$r^2 = 0.991$
$r^2 = 0.877$
$AA = -0.0915 \pm 0.00913$
$r^2 = 0.431$
15
10
0.2
0.4
0.6
$A\,[\mu mol\ m^{-2}\ s^{-1}]$
$g_{sw}\,[mol\ m^{-2}\ s^{-1}]$
Curve type
amphi
pseudohypo
Measurement
high
low

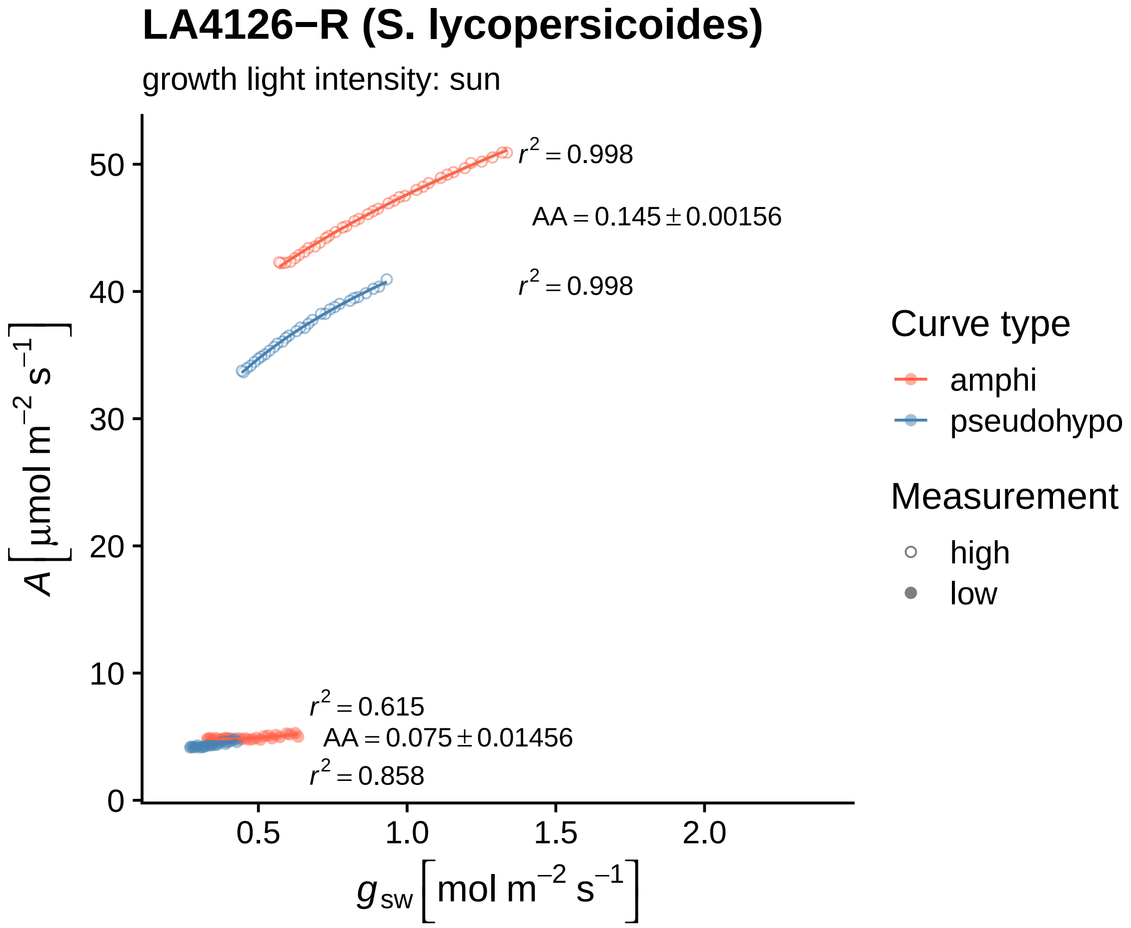

LA4126−R (S. lycopersicoides)
growth light intensity: sun
r² = 0.998
AA = 0.145 ± 0.00156
r² = 0.998
r² = 0.615
AA = 0.075 ± 0.01456
r² = 0.858
A [µmol m⁻² s⁻¹]
0
10
20
30
40
50
0.5
1.0
1.5
2.0
g_sw [mol m⁻² s⁻¹]
Curve type
amphi
pseudohypo
Measurement
high
low

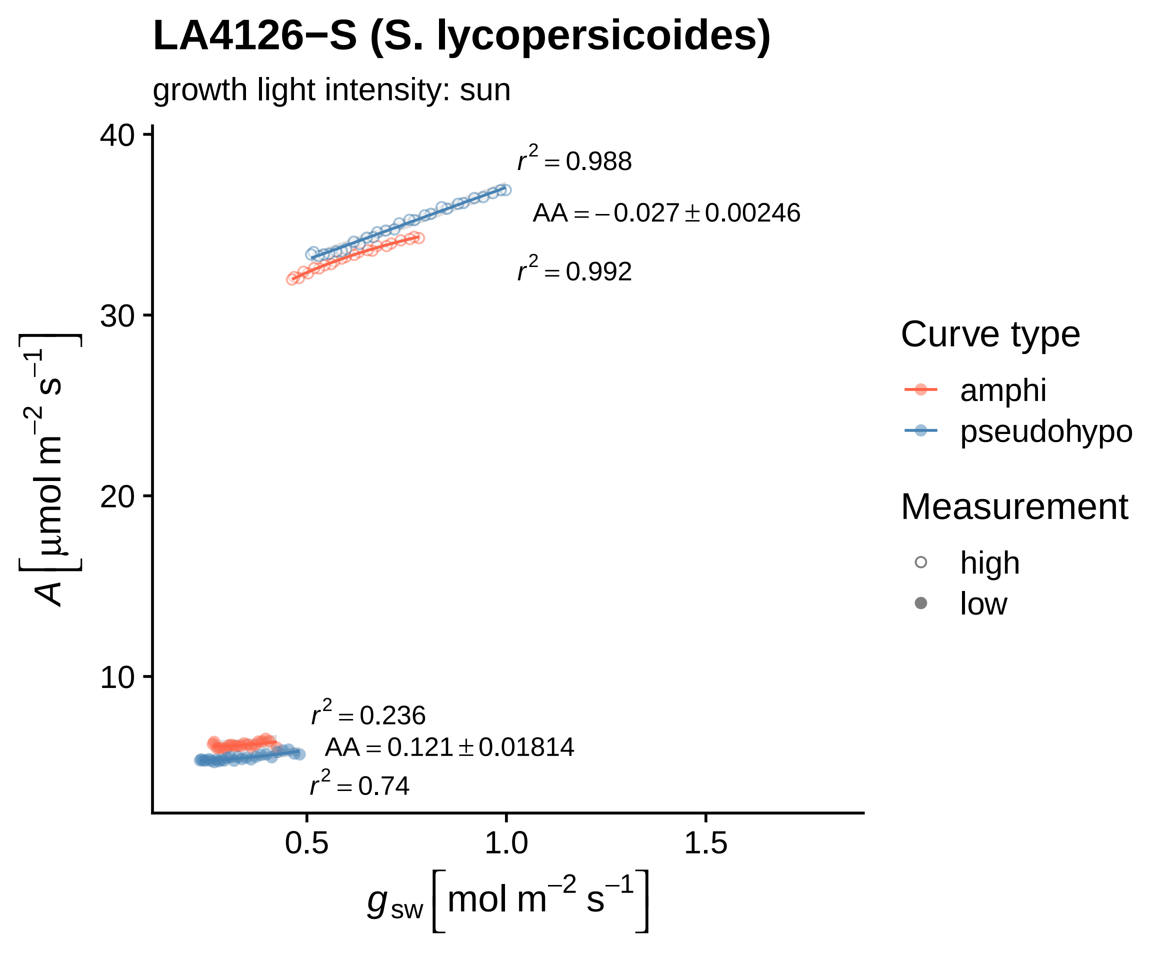
LA4126−S (S. lycopersicoides)
growth light intensity: sun
$r^2 = 0.988$
$AA = -0.027 \pm 0.00246$
$r^2 = 0.992$
$r^2 = 0.236$
$AA = 0.121 \pm 0.01814$
$r^2 = 0.74$
40
30
20
10
0.5
1.0
1.5
$A\left[\mu mol\ m^{-2}\ s^{-1}\right]$
$g_{sw}\left[mol\ m^{-2}\ s^{-1}\right]$
Curve type
amphi
pseudohypo
Measurement
high
low

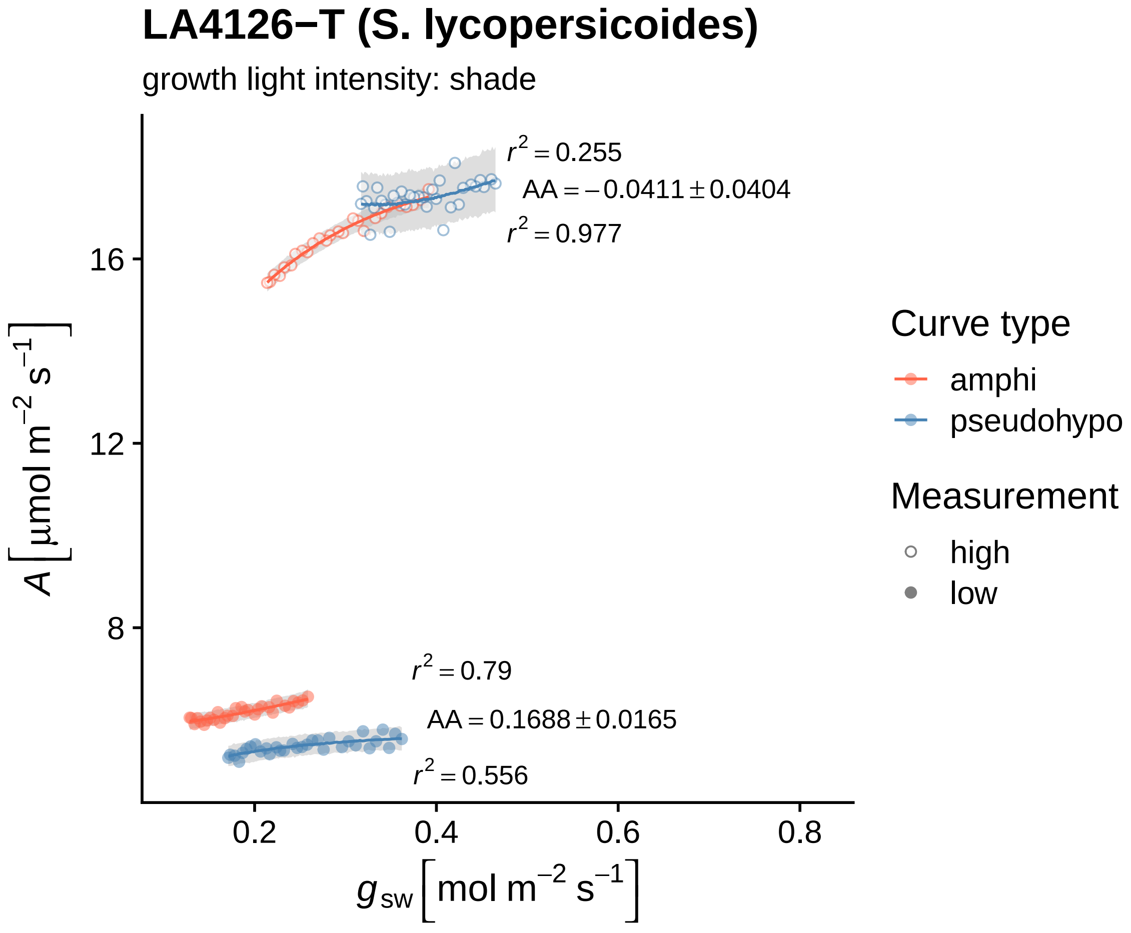
LA4126−T (S. lycopersicoides)
growth light intensity: shade
$r^2 = 0.255$
$AA = -0.0411 \pm 0.0404$
$r^2 = 0.977$
$r^2 = 0.79$
$AA = 0.1688 \pm 0.0165$
$r^2 = 0.556$
16
12
8
$A\,[\mu mol\ m^{-2}\ s^{-1}]$
0.2
0.4
0.6
0.8
$g_{sw}\,[mol\ m^{-2}\ s^{-1}]$
Curve type
amphi
pseudohypo
Measurement
high
low

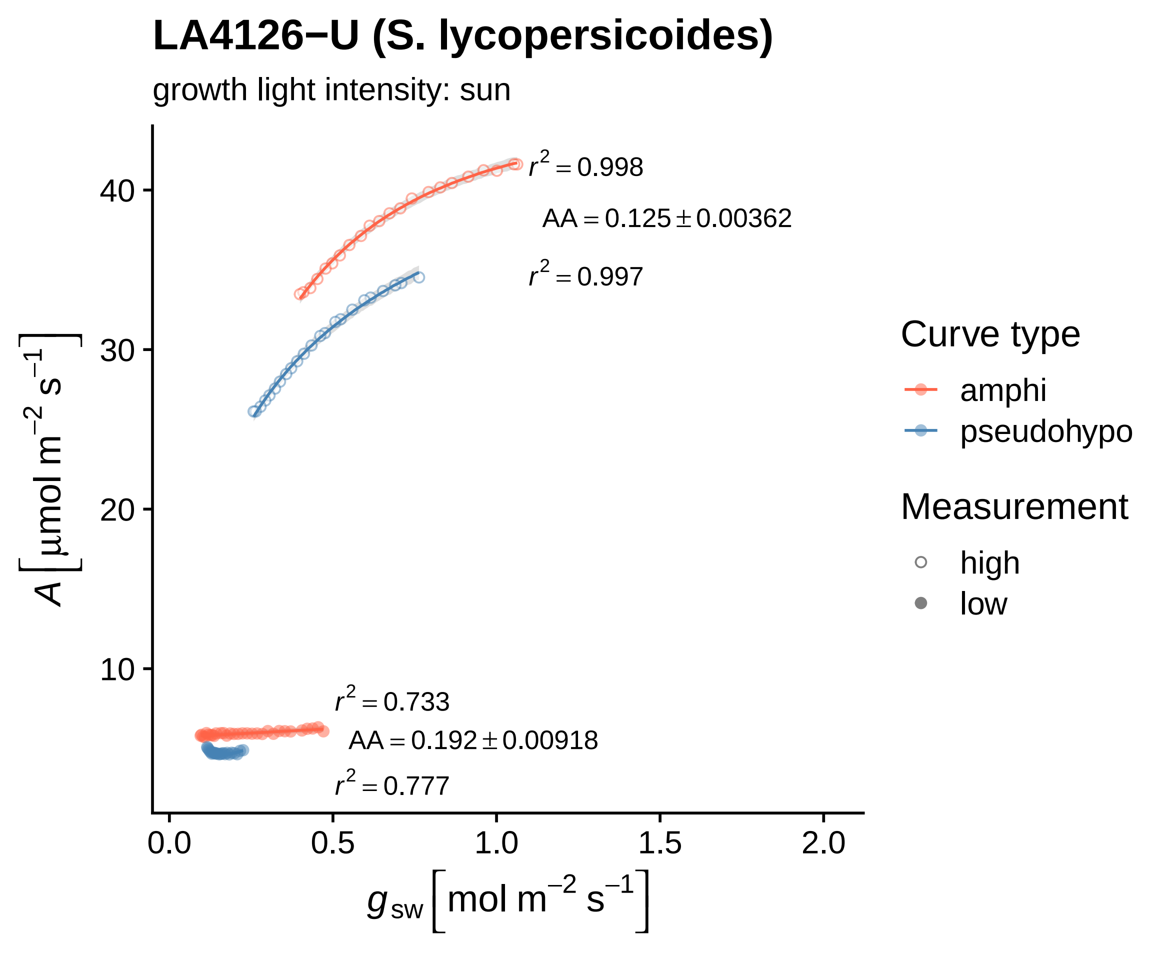
LA4126−U (S. lycopersicoides)
growth light intensity: sun
$r^2 = 0.998$
$AA = 0.125 \pm 0.00362$
$r^2 = 0.997$
$r^2 = 0.733$
$AA = 0.192 \pm 0.00918$
$r^2 = 0.777$
40
30
20
10
0.0
0.5
1.0
1.5
2.0
$A\left[\mu mol\ m^{-2}\ s^{-1}\right]$
$g_{sw}\left[mol\ m^{-2}\ s^{-1}\right]$
Curve type
amphi
pseudohypo
Measurement
high
low

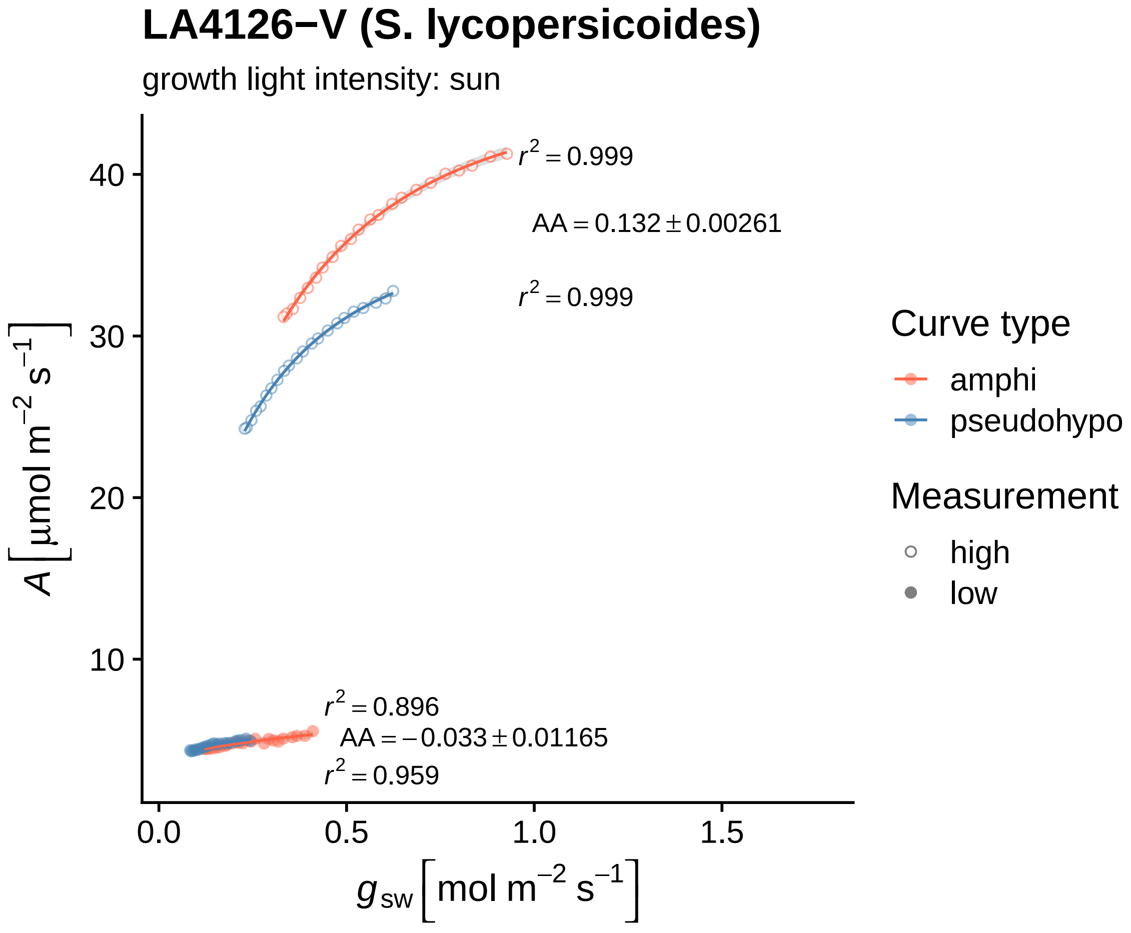

LA4126−V (S. lycopersicoides)
growth light intensity: sun
$r^2 = 0.999$
$AA = 0.132 \pm 0.00261$
$r^2 = 0.999$
$r^2 = 0.896$
$AA = -0.033 \pm 0.01165$
$r^2 = 0.959$
$A$ [μmol m$^{-2}$ s$^{-1}$]
40
30
20
10
0.0
0.5
1.0
1.5
$g_{sw}$ [mol m$^{-2}$ s$^{-1}$]
Curve type
amphi
pseudohypo
Measurement
high
low

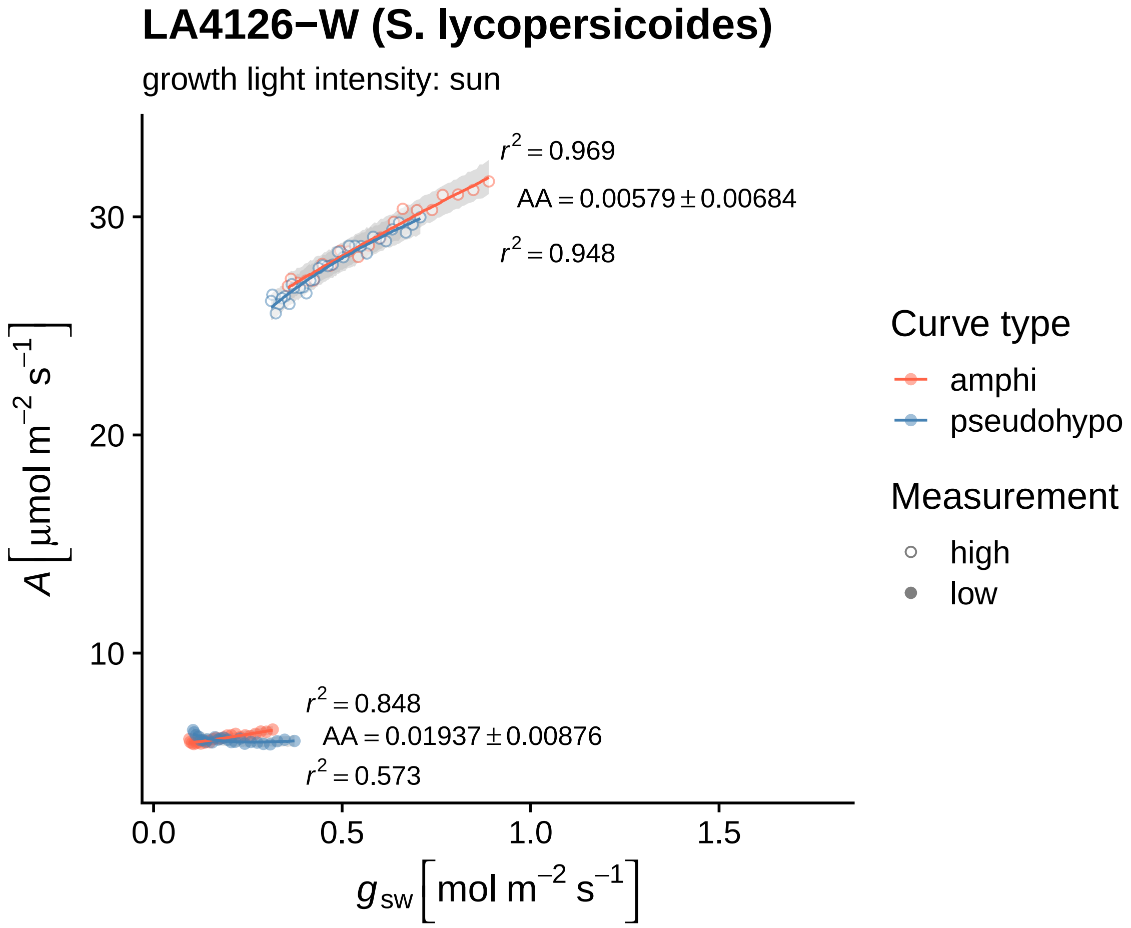

LA4126−W (S. lycopersicoides)
growth light intensity: sun
r² = 0.969
AA = 0.00579 ± 0.00684
r² = 0.948
r² = 0.848
AA = 0.01937 ± 0.00876
r² = 0.573
30
20
10
0.0
0.5
1.0
1.5
A [µmol m⁻² s⁻¹]
g sw [mol m⁻² s⁻¹]
Curve type
amphi
pseudohypo
Measurement
high
low

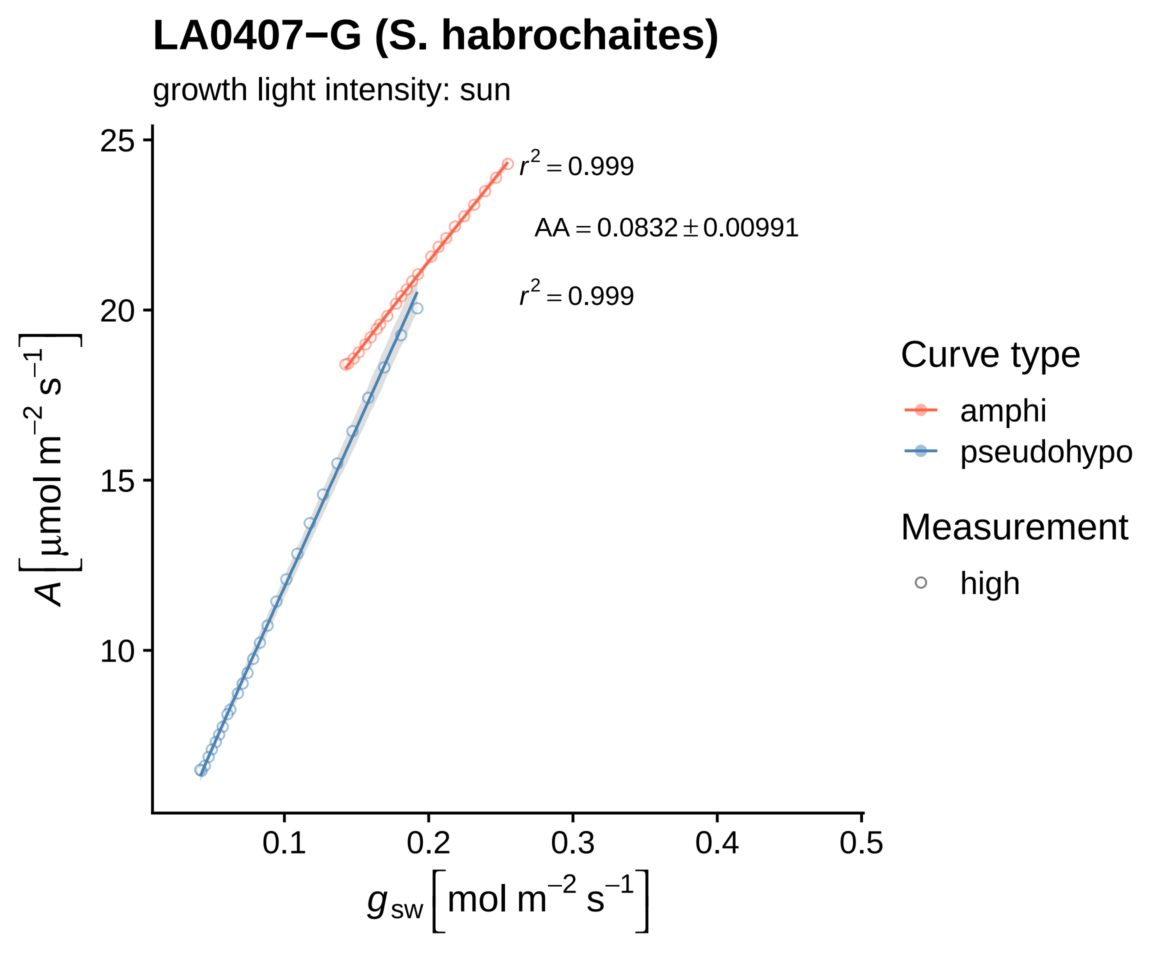

LA0407−G (S. habrochaites)
growth light intensity: sun
$r^2 = 0.999$
$AA = 0.0832 \pm 0.00991$
$r^2 = 0.999$
25
20
15
10
$A\left[\mu mol\ m^{-2}\ s^{-1}\right]$
0.1
0.2
0.3
0.4
0.5
$g_{sw}\left[mol\ m^{-2}\ s^{-1}\right]$
Curve type
amphi
pseudohypo
Measurement
high

### LA0750−E (S. pennellii)

growth light intensity: sun

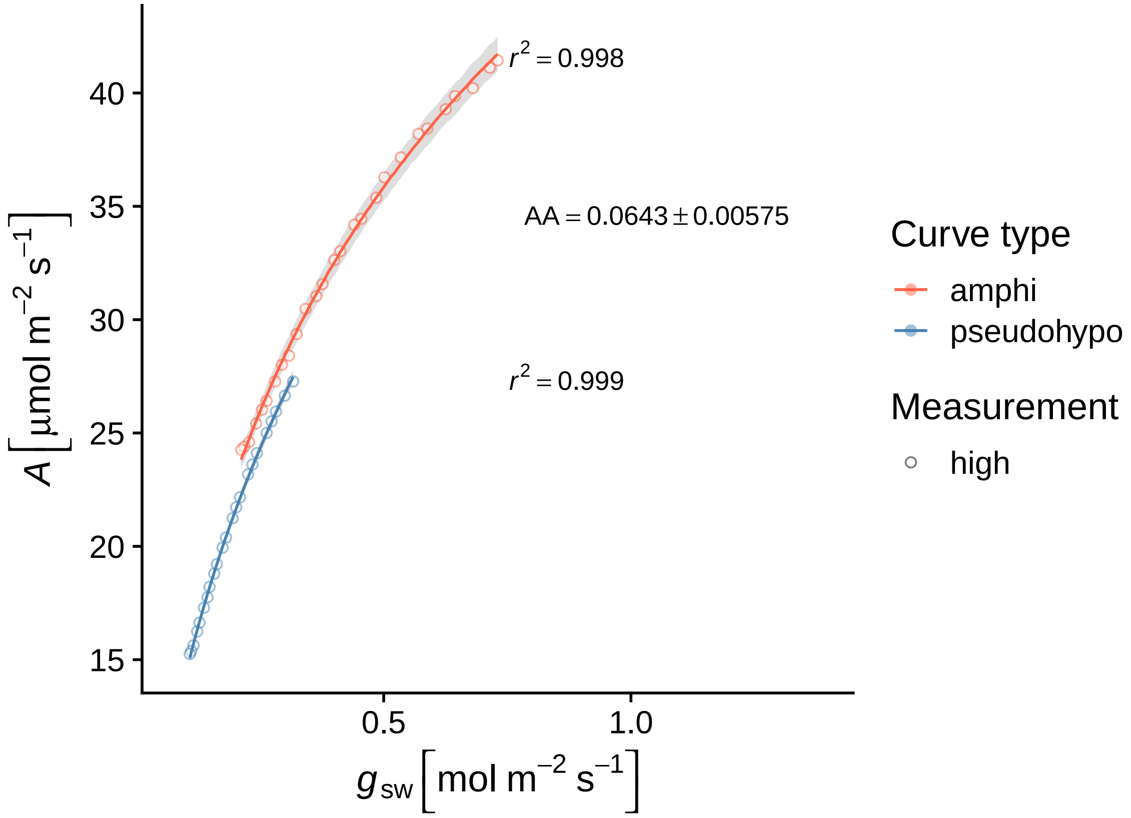

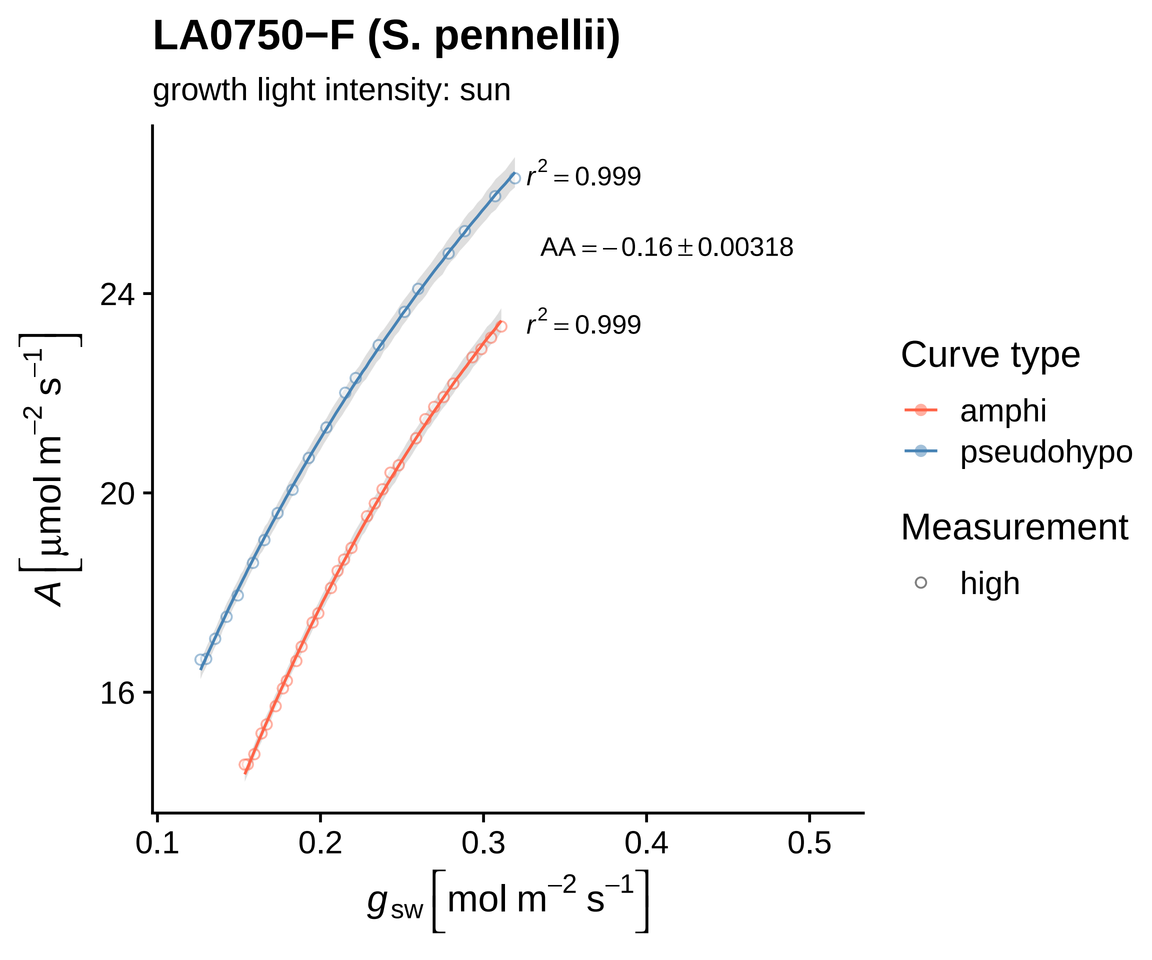
LA0750−F (S. pennellii)
growth light intensity: sun
$r^2 = 0.999$
$AA = -0.16 \pm 0.00318$
$r^2 = 0.999$
24
20
16
0.1
0.2
0.3
0.4
0.5
$A\ [\mu mol\ m^{-2}\ s^{-1}]$
$g_{sw}\ [mol\ m^{-2}\ s^{-1}]$
Curve type
amphi
pseudohypo
Measurement
high

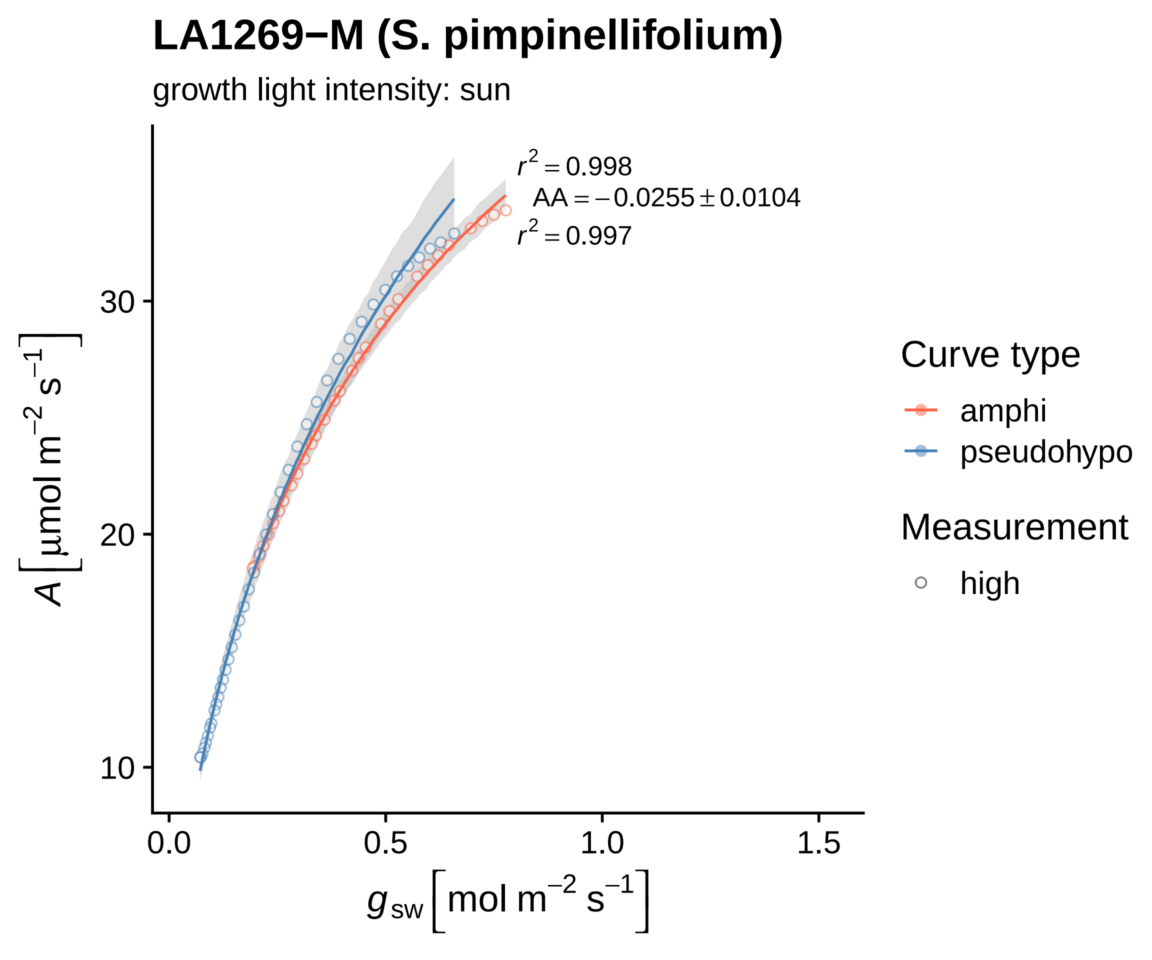

LA1269−M (S. pimpinellifolium)
growth light intensity: sun
$r^2 = 0.998$
$AA = -0.0255 \pm 0.0104$
$r^2 = 0.997$
A [µmol m−2 s−1]
30
20
10
0.0
0.5
1.0
1.5
gsw [mol m−2 s−1]
Curve type
amphi
pseudohypo
Measurement
high

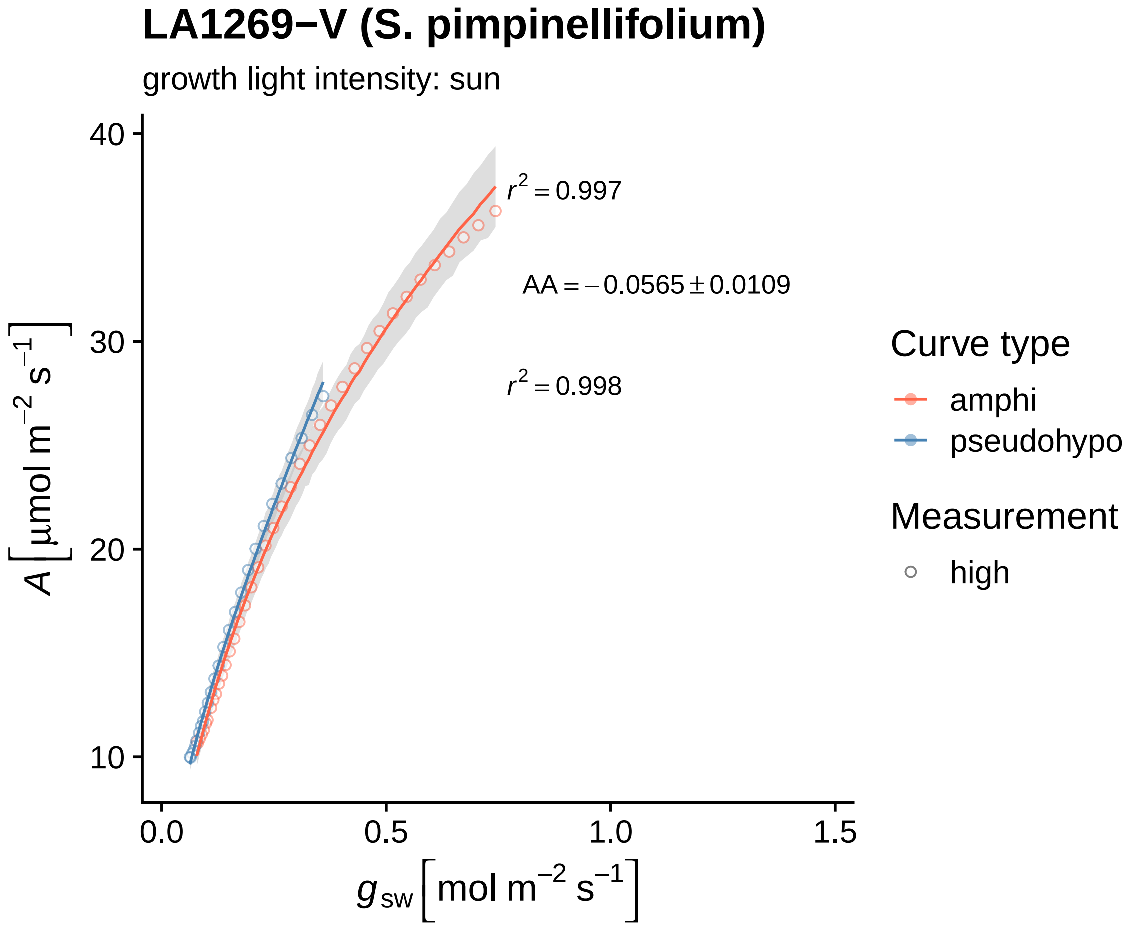
LA1269−V (S. pimpinellifolium)
growth light intensity: sun
$r^2 = 0.997$
AA = −0.0565 ± 0.0109
$r^2 = 0.998$
Curve type
amphi
pseudohypo
Measurement
high
$A\ [\mu mol\ m^{-2}\ s^{-1}]$
$g_{sw}\ [mol\ m^{-2}\ s^{-1}]$
10
20
30
40
0.0
0.5
1.0
1.5

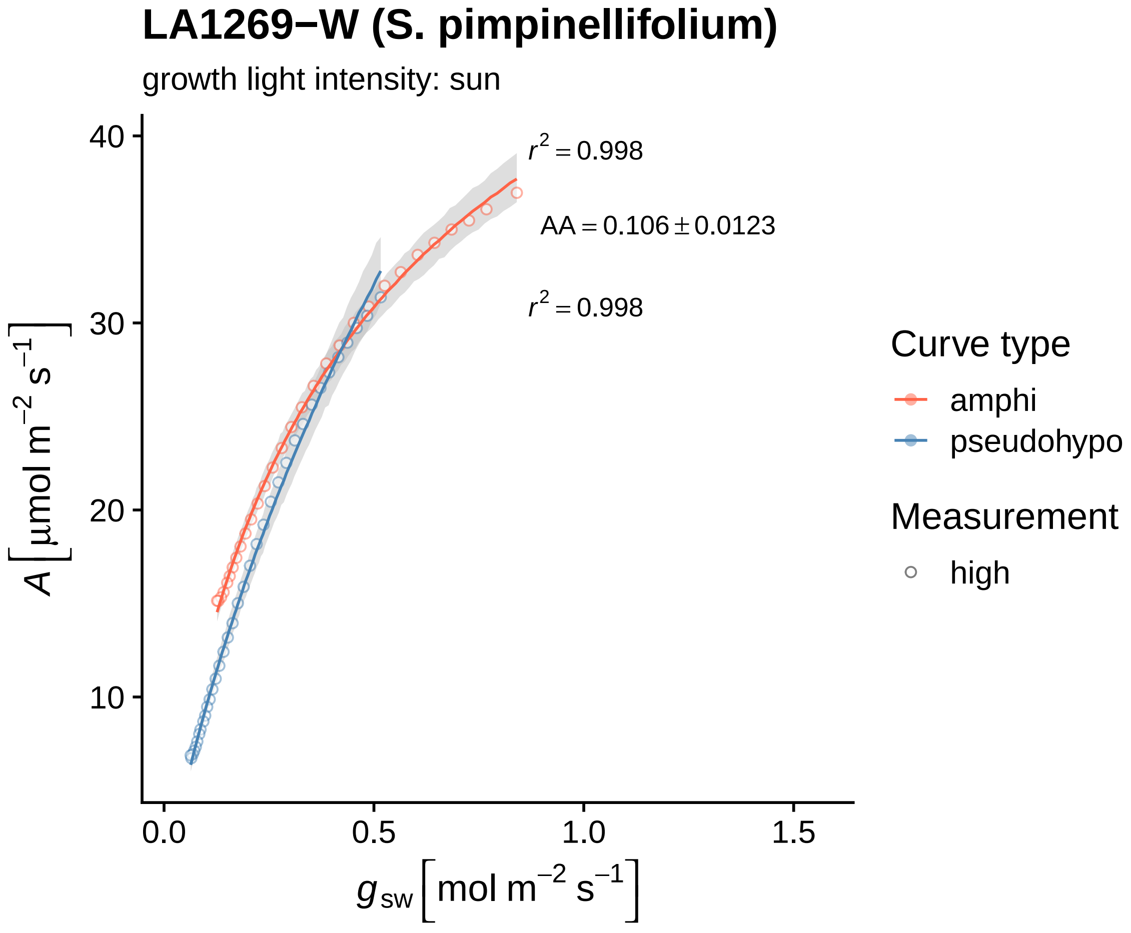

LA1269−W (S. pimpinellifolium)
growth light intensity: sun
$r^2 = 0.998$
$AA = 0.106 \pm 0.0123$
$r^2 = 0.998$
$A \left[ \mu mol\ m^{-2}\ s^{-1} \right]$
40
30
20
10
0.0
0.5
1.0
1.5
$g_{sw} \left[ mol\ m^{-2}\ s^{-1} \right]$
Curve type
amphi
pseudohypo
Measurement
high

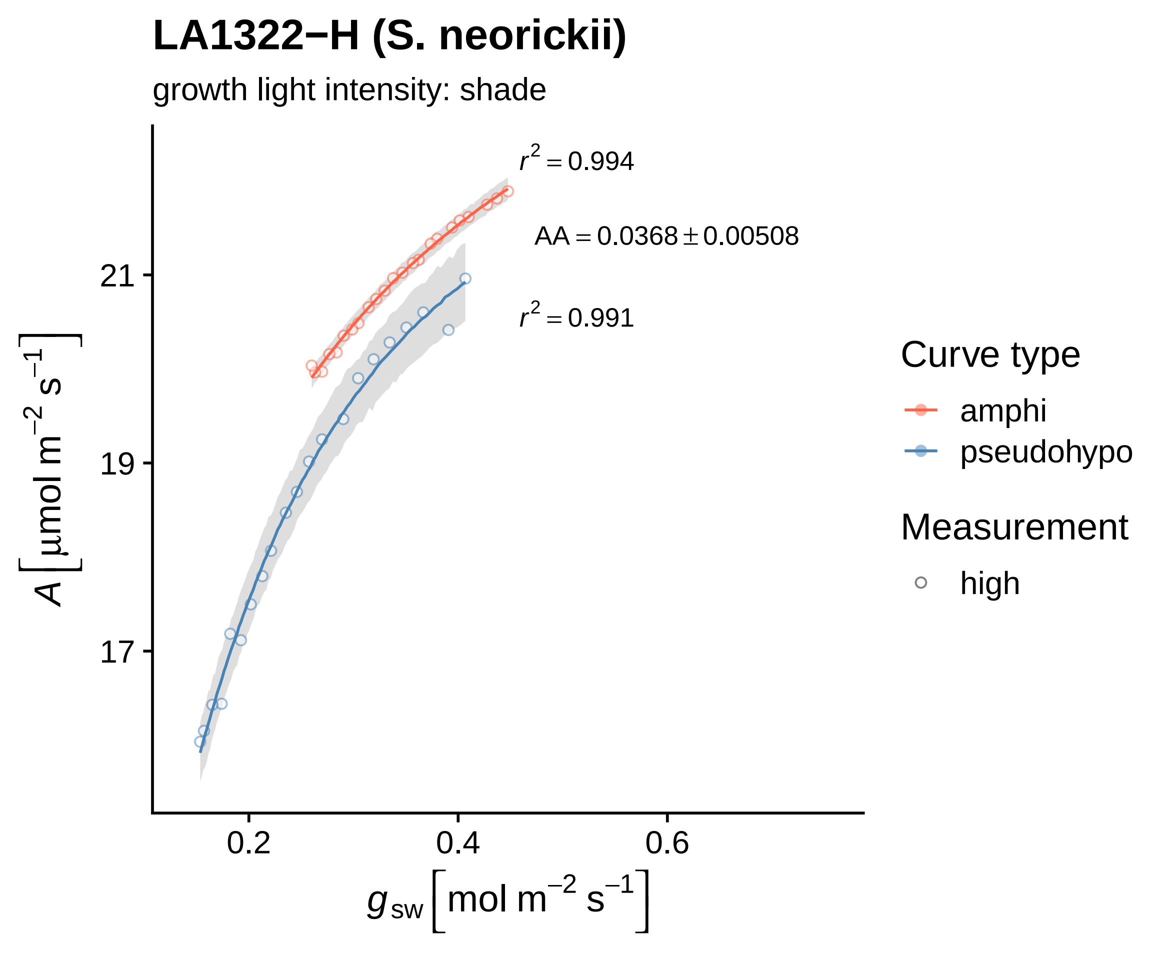
LA1322−H (S. neorickii)
growth light intensity: shade
$r^2 = 0.994$
$AA = 0.0368 \pm 0.00508$
$r^2 = 0.991$
21
19
17
0.2
0.4
0.6
$A\ [\mu mol\ m^{-2}\ s^{-1}]$
$g_{sw}\ [mol\ m^{-2}\ s^{-1}]$
Curve type
amphi
pseudohypo
Measurement
high

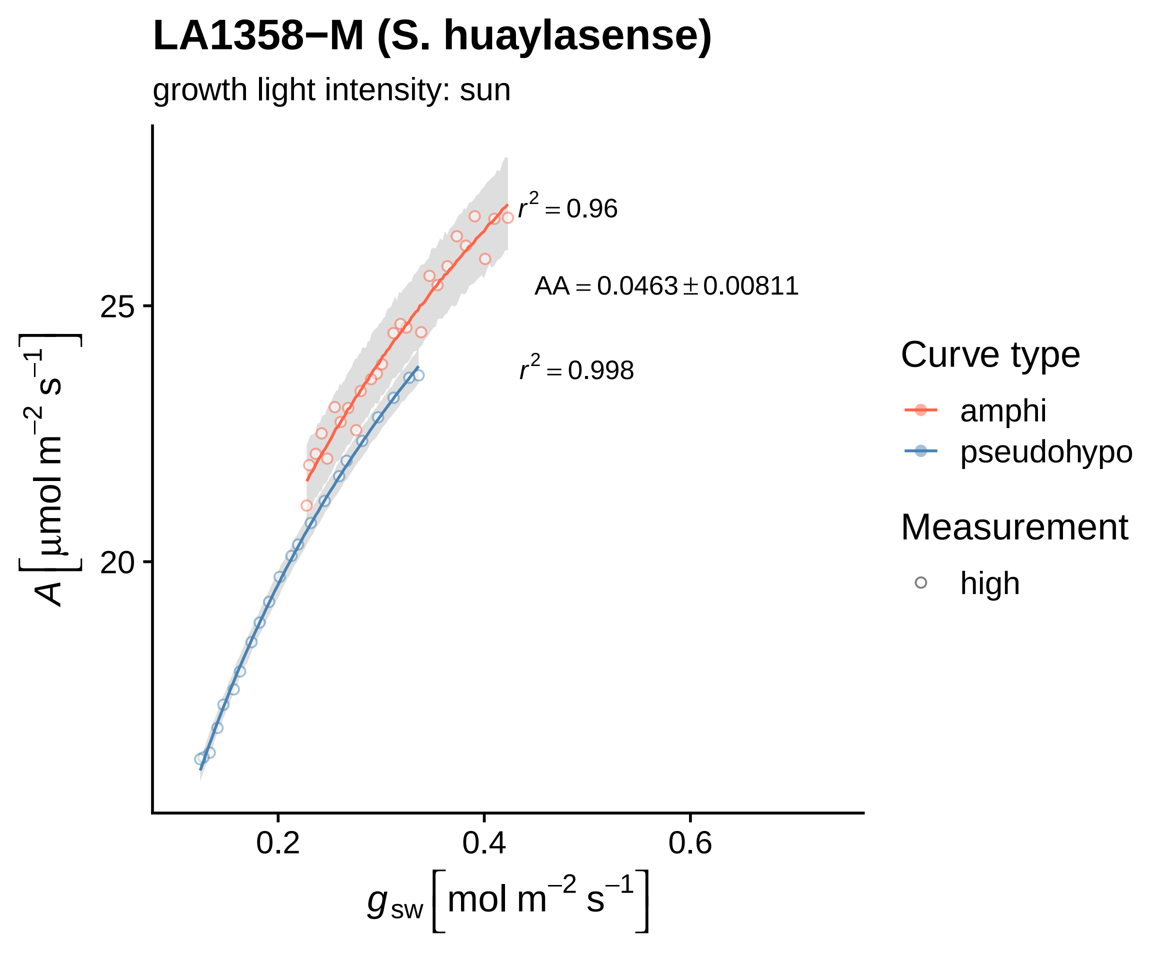

LA1358−M (S. huaylasense)
growth light intensity: sun
r2 = 0.96
AA = 0.0463 ± 0.00811
r2 = 0.998
25
20
0.2
0.4
0.6
A [μmol m−2 s−1]
gsw [mol m−2 s−1]
Curve type
amphi
pseudohypo
Measurement
high

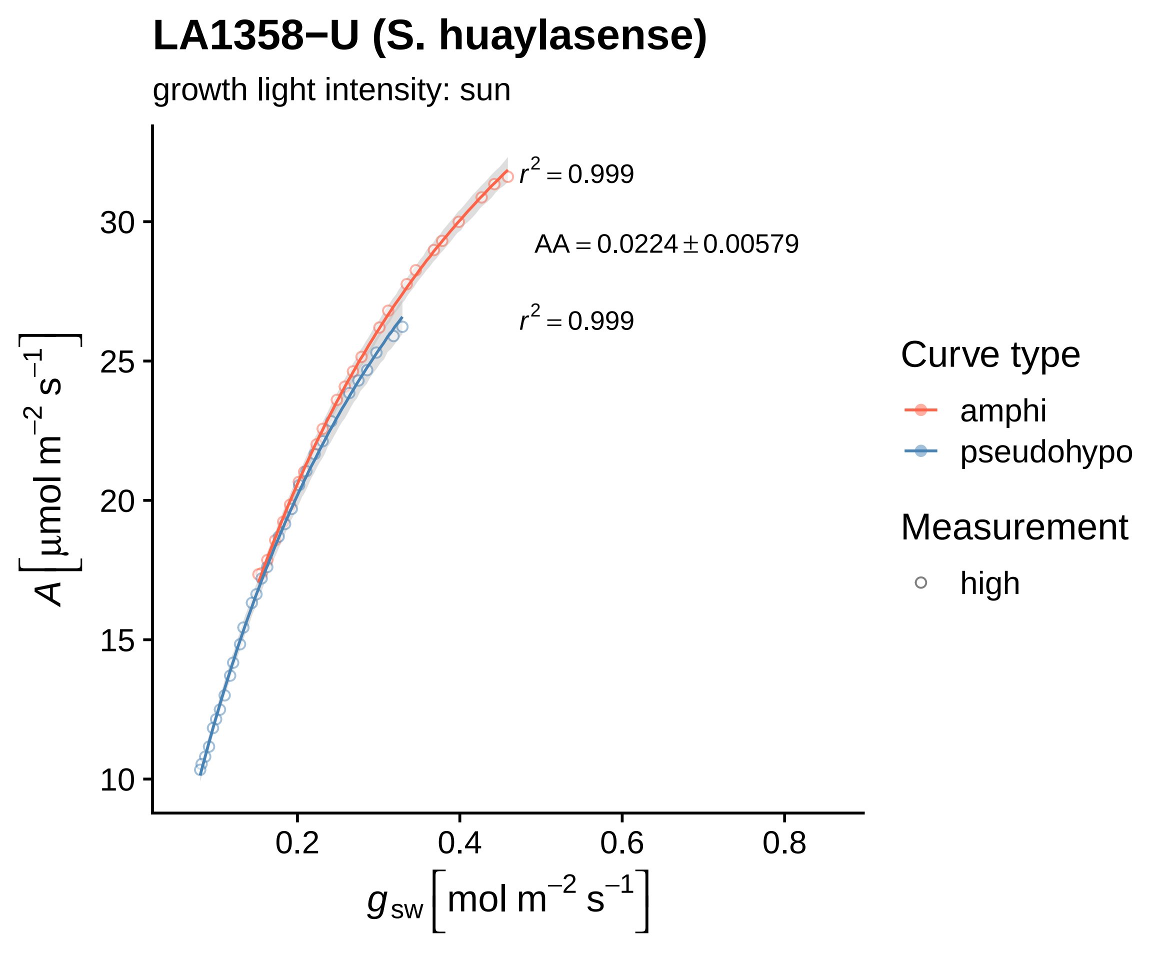

LA1358−U (S. huaylasense)
growth light intensity: sun
r^2 = 0.999
AA = 0.0224 ± 0.00579
r^2 = 0.999
A [μmol m^−2 s^−1]
g_sw [mol m^−2 s^−1]
10
15
20
25
30
0.2
0.4
0.6
0.8
Curve type
amphi
pseudohypo
Measurement
high

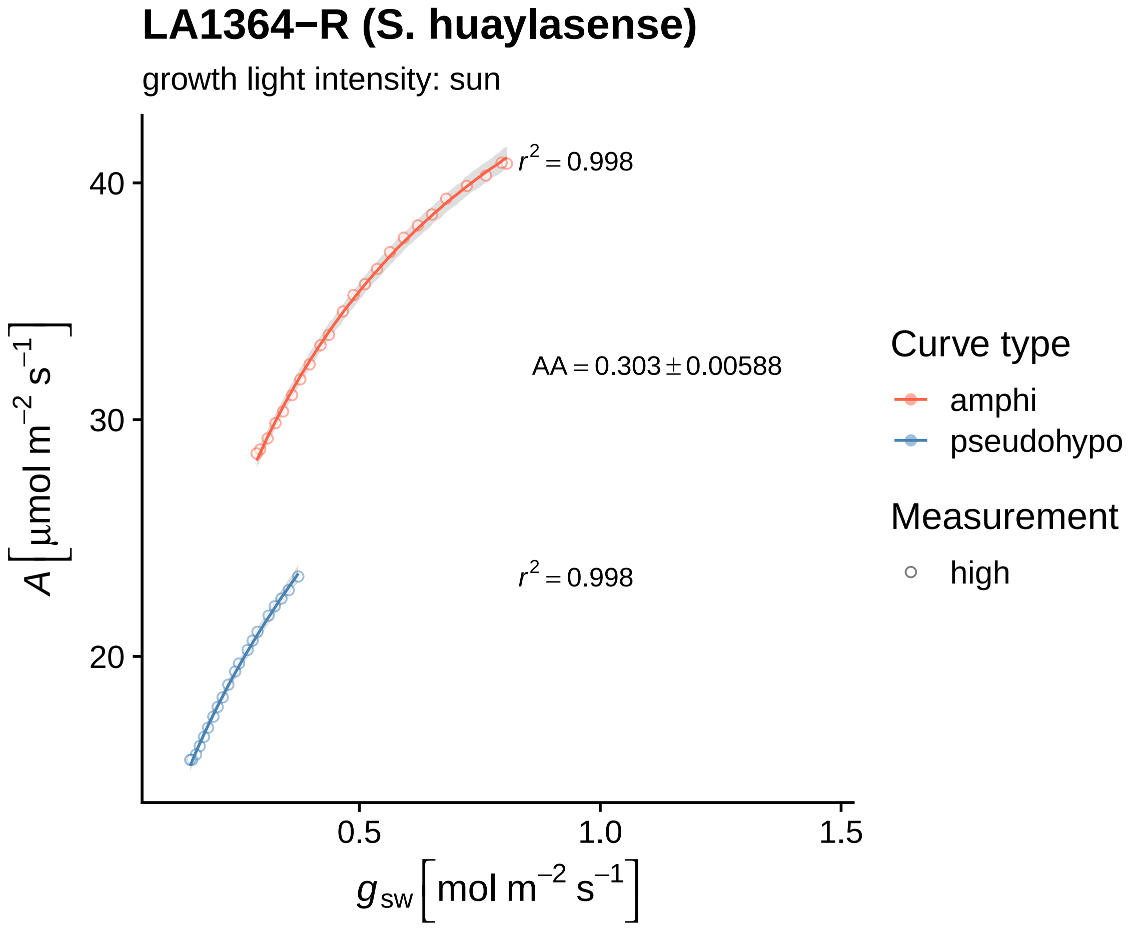
LA1364−R (S. huaylasense)
growth light intensity: sun
r2 = 0.998
AA = 0.303 ± 0.00588
r2 = 0.998
40
30
20
0.5
1.0
1.5
A [µmol m−2 s−1]
gsw [mol m−2 s−1]
Curve type
amphi
pseudohypo
Measurement
high

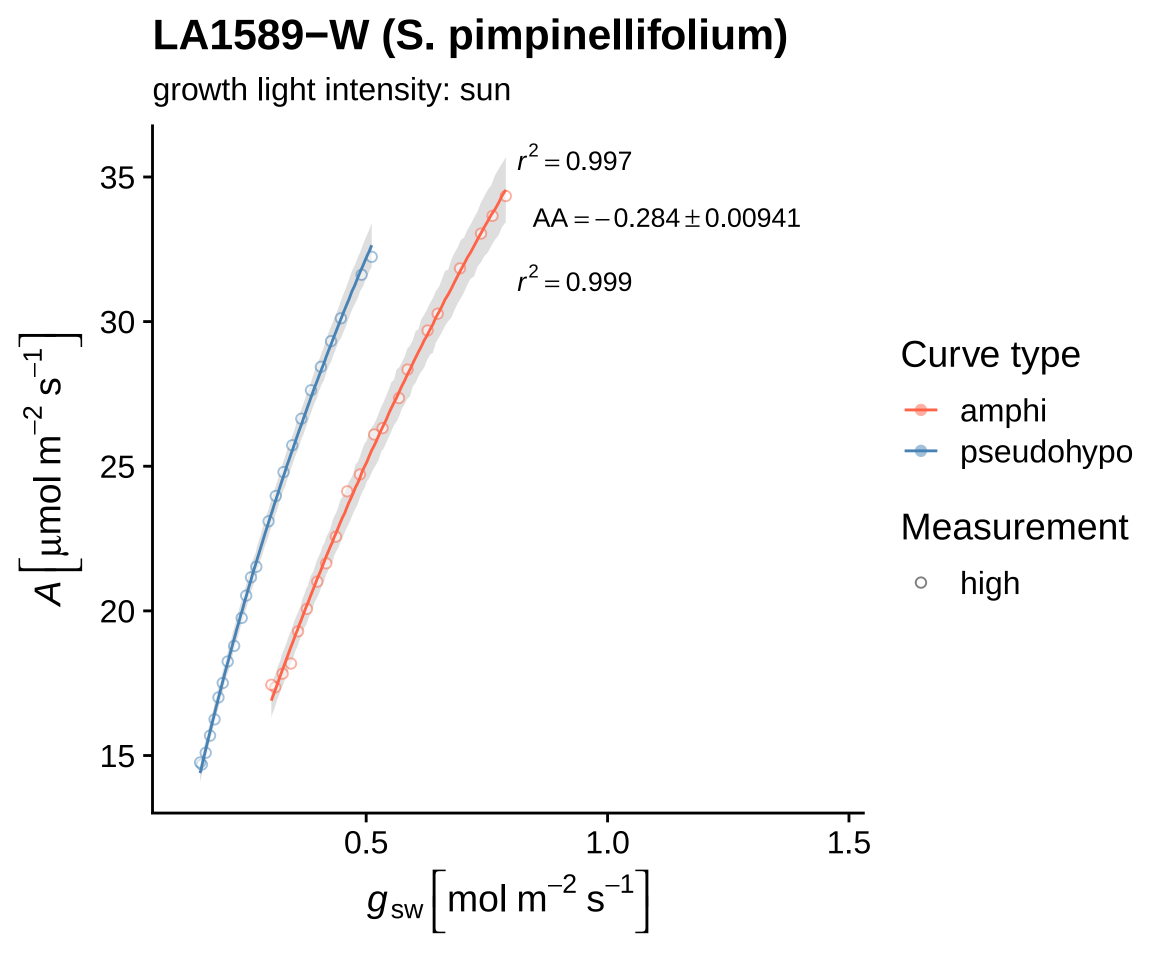
LA1589−W (S. pimpinellifolium)
growth light intensity: sun
r² = 0.997
AA = −0.284 ± 0.00941
r² = 0.999
A [μmol m⁻² s⁻¹]
35
30
25
20
15
0.5
1.0
1.5
g sw [mol m⁻² s⁻¹]
Curve type
amphi
pseudohypo
Measurement
high

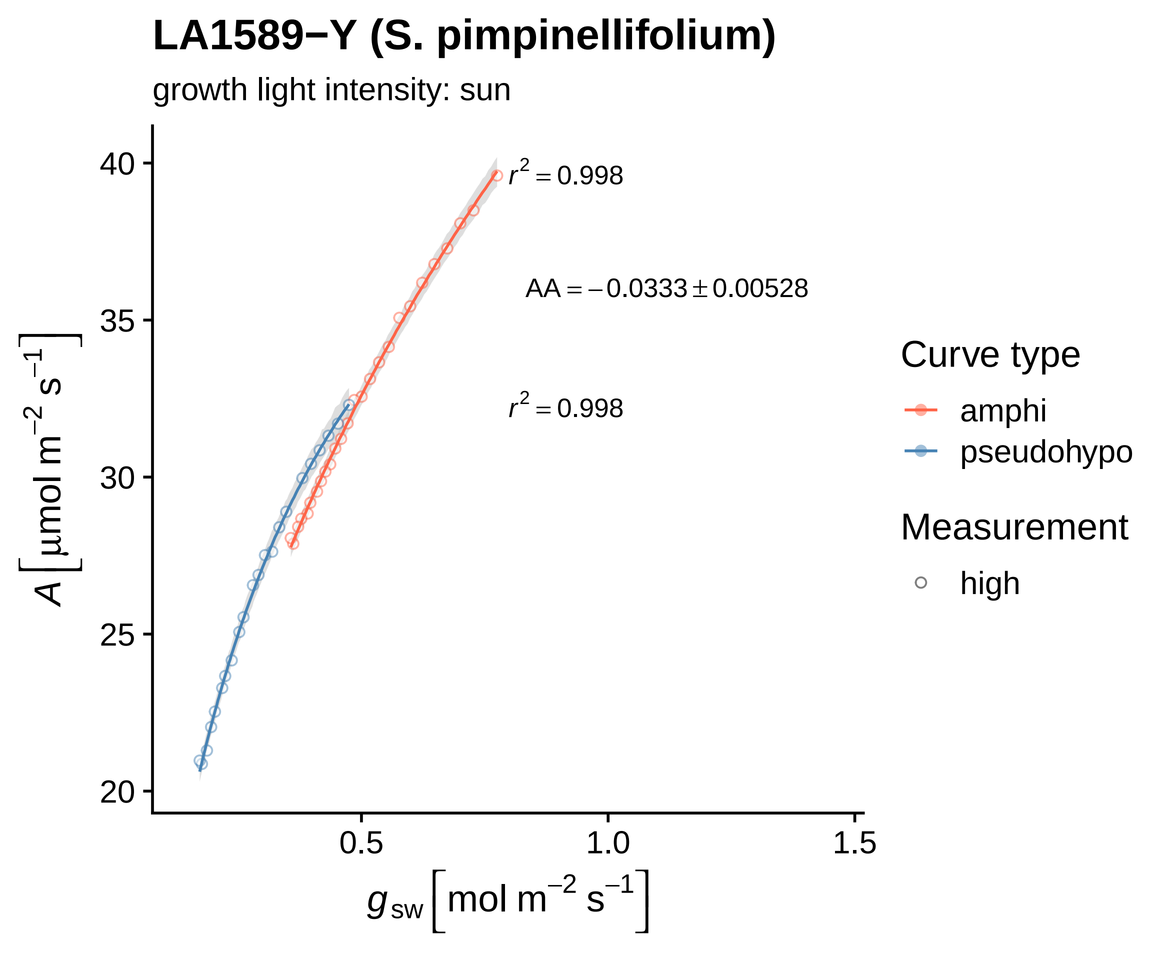

LA1589−Y (S. pimpinellifolium)
growth light intensity: sun
$r^2 = 0.998$
AA = −0.0333 ± 0.00528
$r^2 = 0.998$
40
35
30
25
20
0.5
1.0
1.5
$A$ [μmol m$^{-2}$ s$^{-1}$]
$g_{sw}$ [mol m$^{-2}$ s$^{-1}$]
Curve type
amphi
pseudohypo
Measurement
high

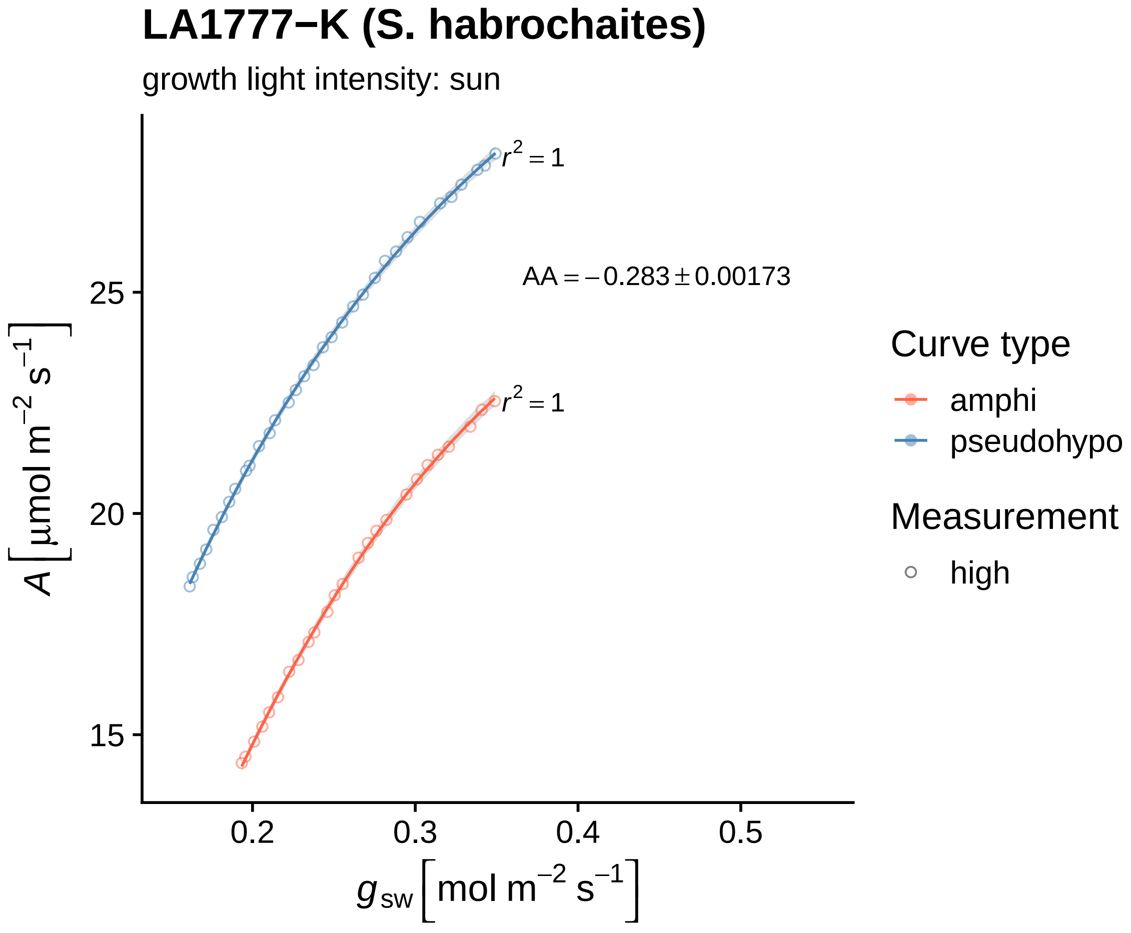

LA1777−K (S. habrochaites)
growth light intensity: sun
$r^2 = 1$
AA = −0.283 ± 0.00173
$r^2 = 1$
25
20
15
0.2
0.3
0.4
0.5
$A$ [µmol m$^{-2}$ s$^{-1}$]
$g_{sw}$ [mol m$^{-2}$ s$^{-1}$]
Curve type
amphi
pseudohypo
Measurement
high

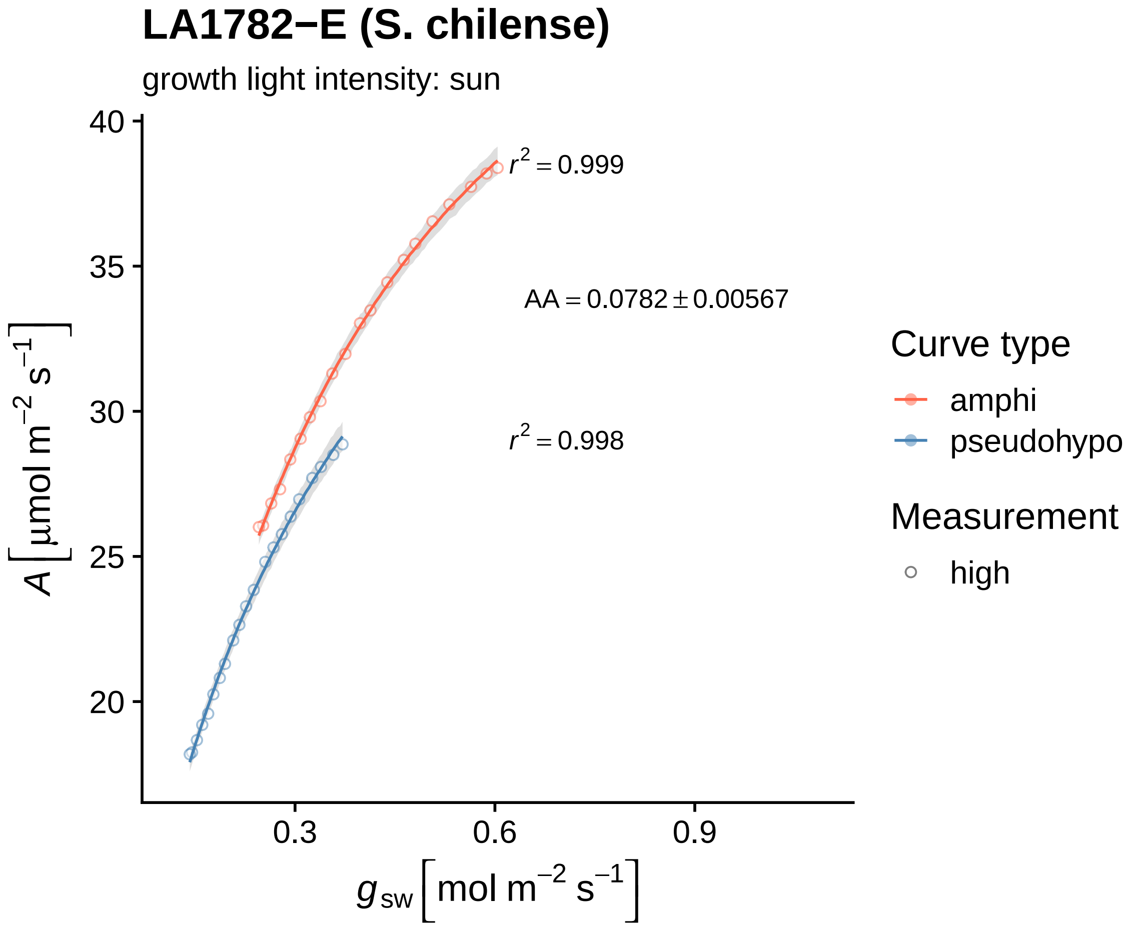
LA1782−E (S. chilense)
growth light intensity: sun
$r^2 = 0.999$
$AA = 0.0782 \pm 0.00567$
$r^2 = 0.998$
$A\left[\mu mol\ m^{-2}\ s^{-1}\right]$
$g_{sw}\left[mol\ m^{-2}\ s^{-1}\right]$
40
35
30
25
20
0.3
0.6
0.9
Curve type
amphi
pseudohypo
Measurement
high

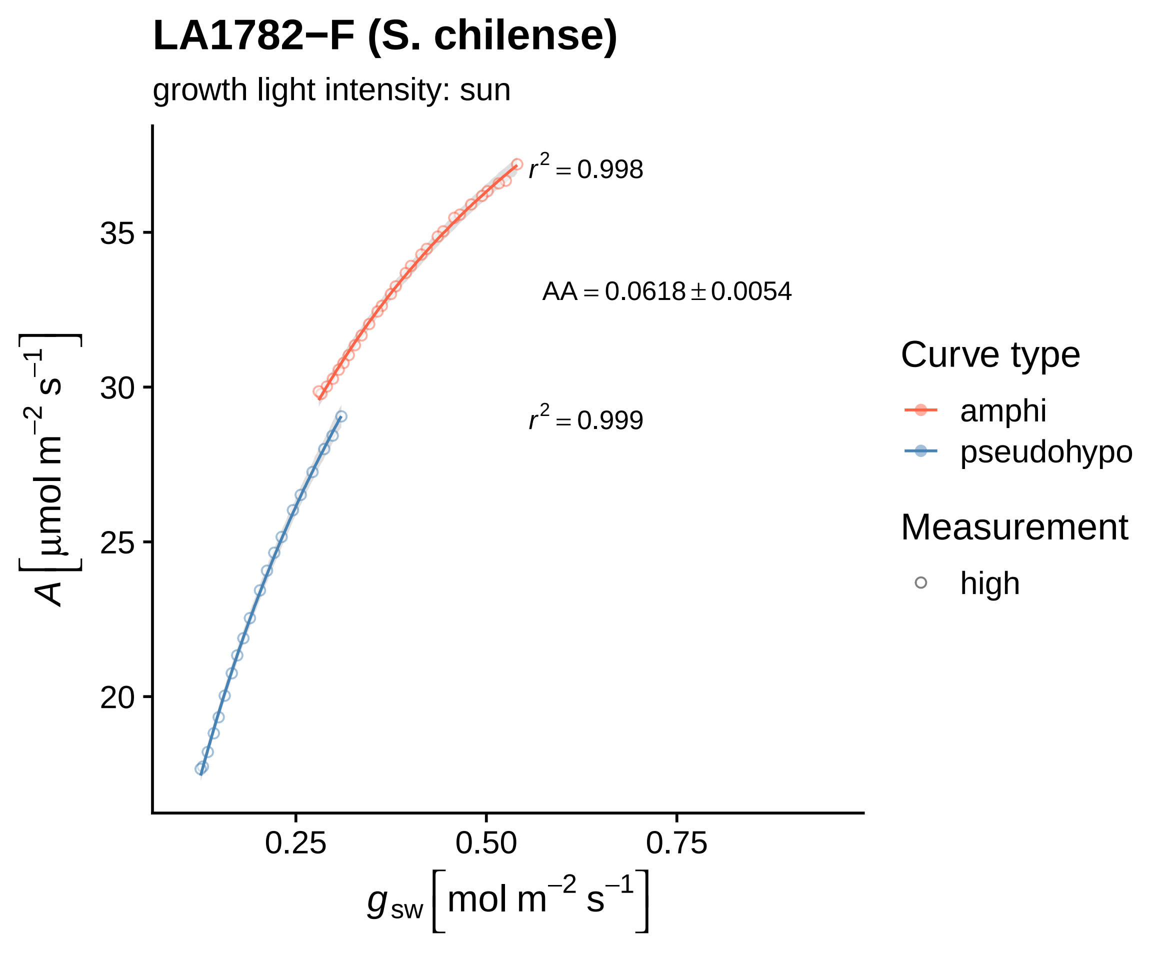
LA1782−F (S. chilense)
growth light intensity: sun
$r^2 = 0.998$
AA = 0.0618 ± 0.0054
$r^2 = 0.999$
35
30
25
20
0.25
0.50
0.75
$A\left[\mu mol\ m^{-2}\ s^{-1}\right]$
$g_{sw}\left[mol\ m^{-2}\ s^{-1}\right]$
Curve type
amphi
pseudohypo
Measurement
high

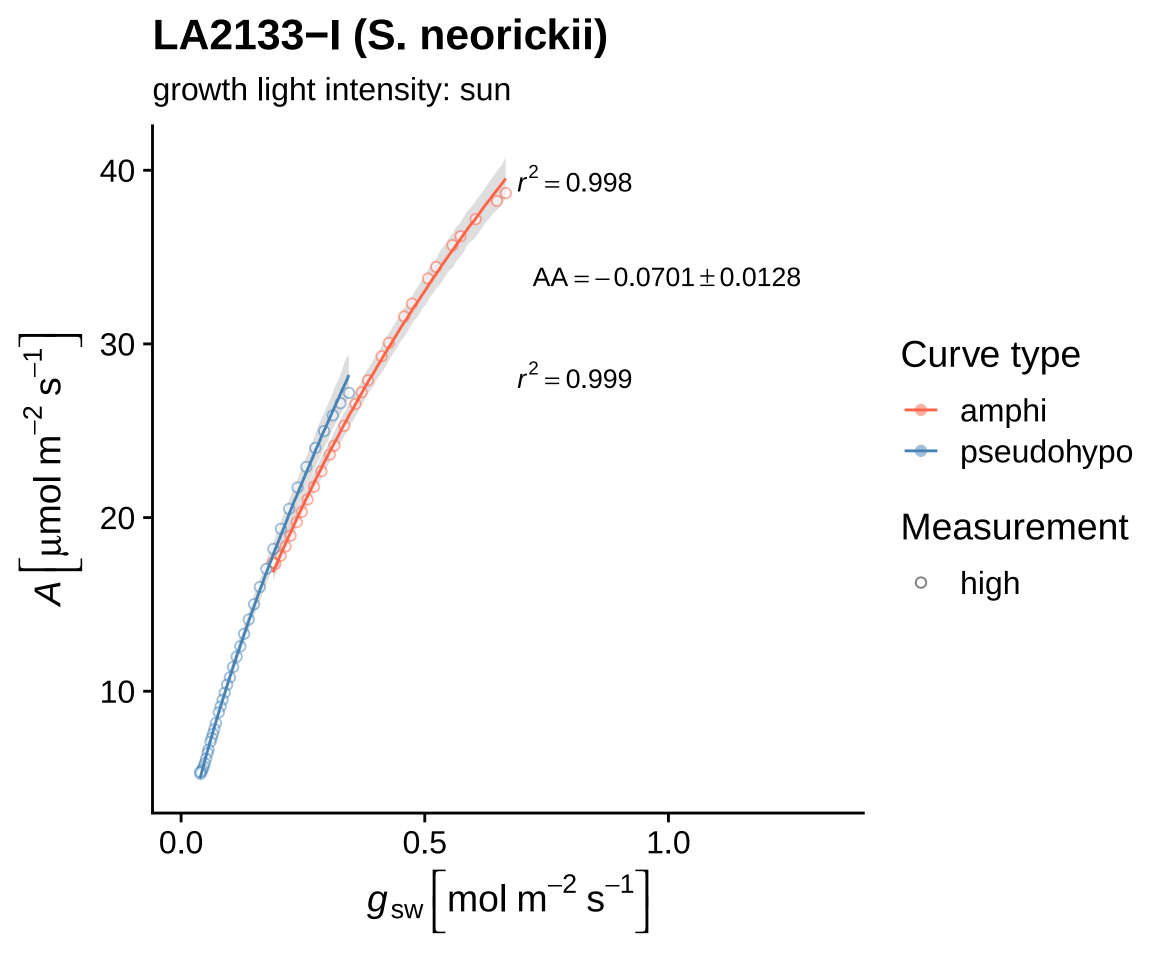
LA2133−I (S. neorickii)
growth light intensity: sun
$r^2 = 0.998$
$AA = -0.0701 \pm 0.0128$
$r^2 = 0.999$
40
30
20
10
0.0
0.5
1.0
$A\left[\mu mol\ m^{-2}\ s^{-1}\right]$
$g_{sw}\left[mol\ m^{-2}\ s^{-1}\right]$
Curve type
amphi
pseudohypo
Measurement
high

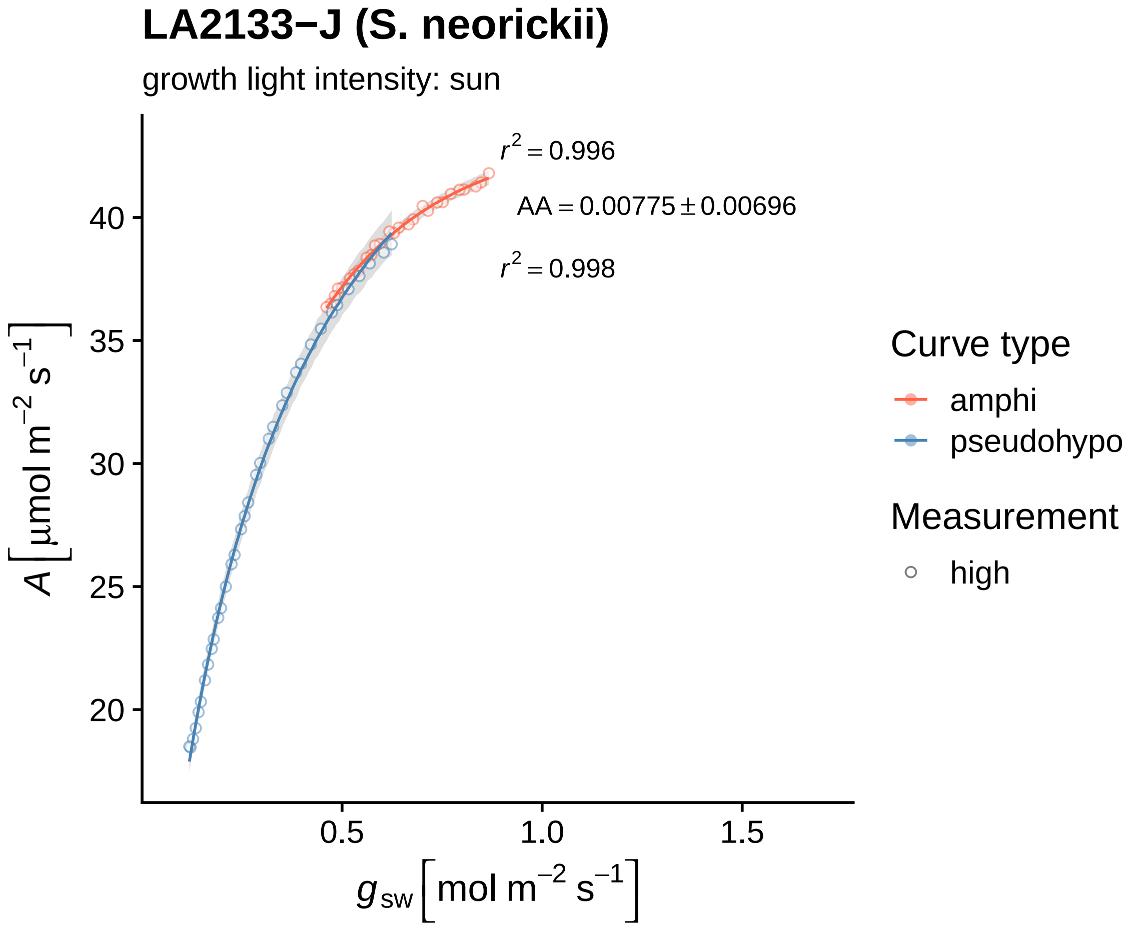
LA2133−J (S. neorickii)
growth light intensity: sun
$r^2 = 0.996$
$AA = 0.00775 \pm 0.00696$
$r^2 = 0.998$
$A\left[\mu mol\ m^{-2}\ s^{-1}\right]$
40
35
30
25
20
0.5
1.0
1.5
$g_{sw}\left[mol\ m^{-2}\ s^{-1}\right]$
Curve type
amphi
pseudohypo
Measurement
high

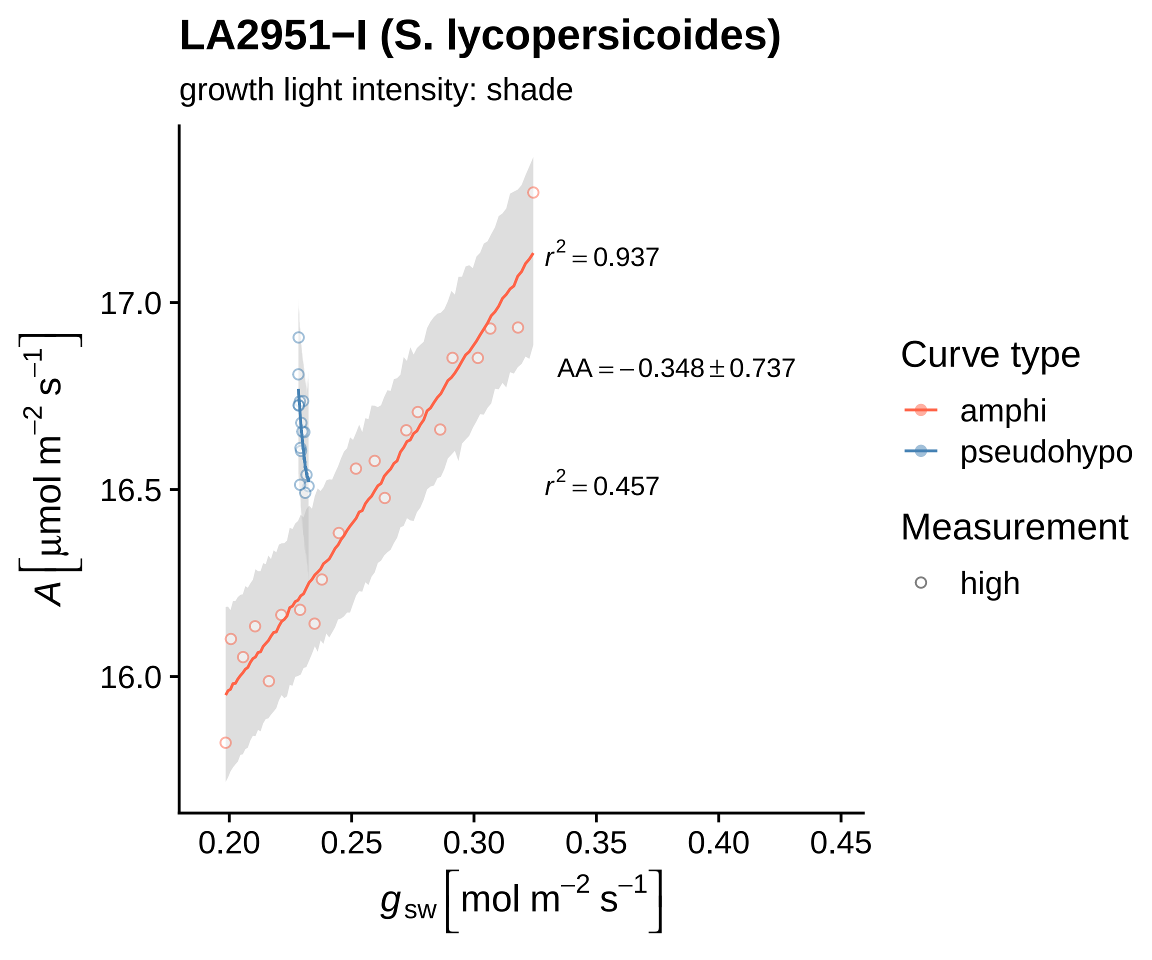

LA2951−I (S. lycopersicoides)
growth light intensity: shade
$r^2 = 0.937$
$AA = -0.348 \pm 0.737$
$r^2 = 0.457$
17.0
16.5
16.0
0.20
0.25
0.30
0.35
0.40
0.45
$A\left[\mu mol\ m^{-2}\ s^{-1}\right]$
$g_{sw}\left[mol\ m^{-2}\ s^{-1}\right]$
Curve type
amphi
pseudohypo
Measurement
high

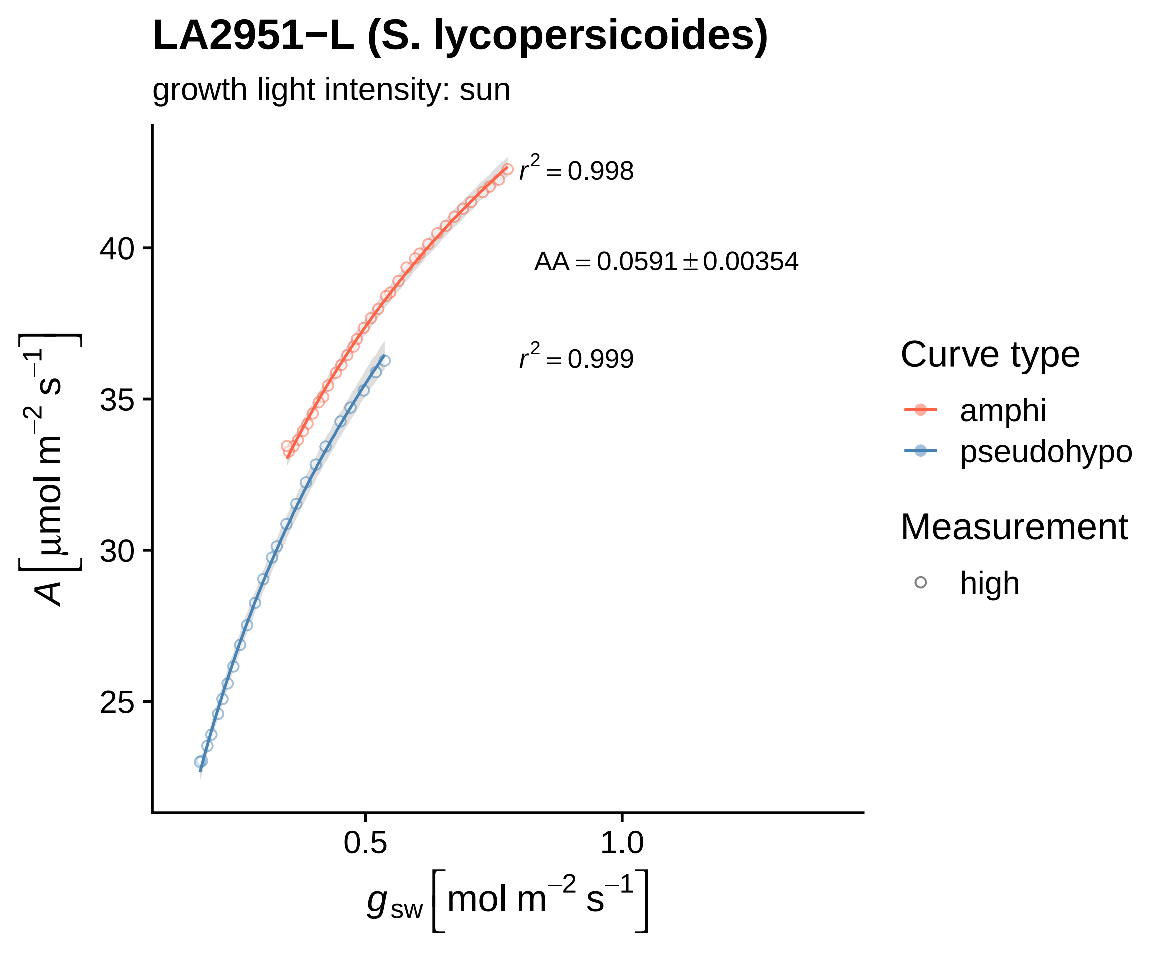

LA2951−L (S. lycopersicoides)
growth light intensity: sun
r² = 0.998
AA = 0.0591 ± 0.00354
r² = 0.999
A [µmol m⁻² s⁻¹]
40
35
30
25
0.5
1.0
g sw [mol m⁻² s⁻¹]
Curve type
amphi
pseudohypo
Measurement
high

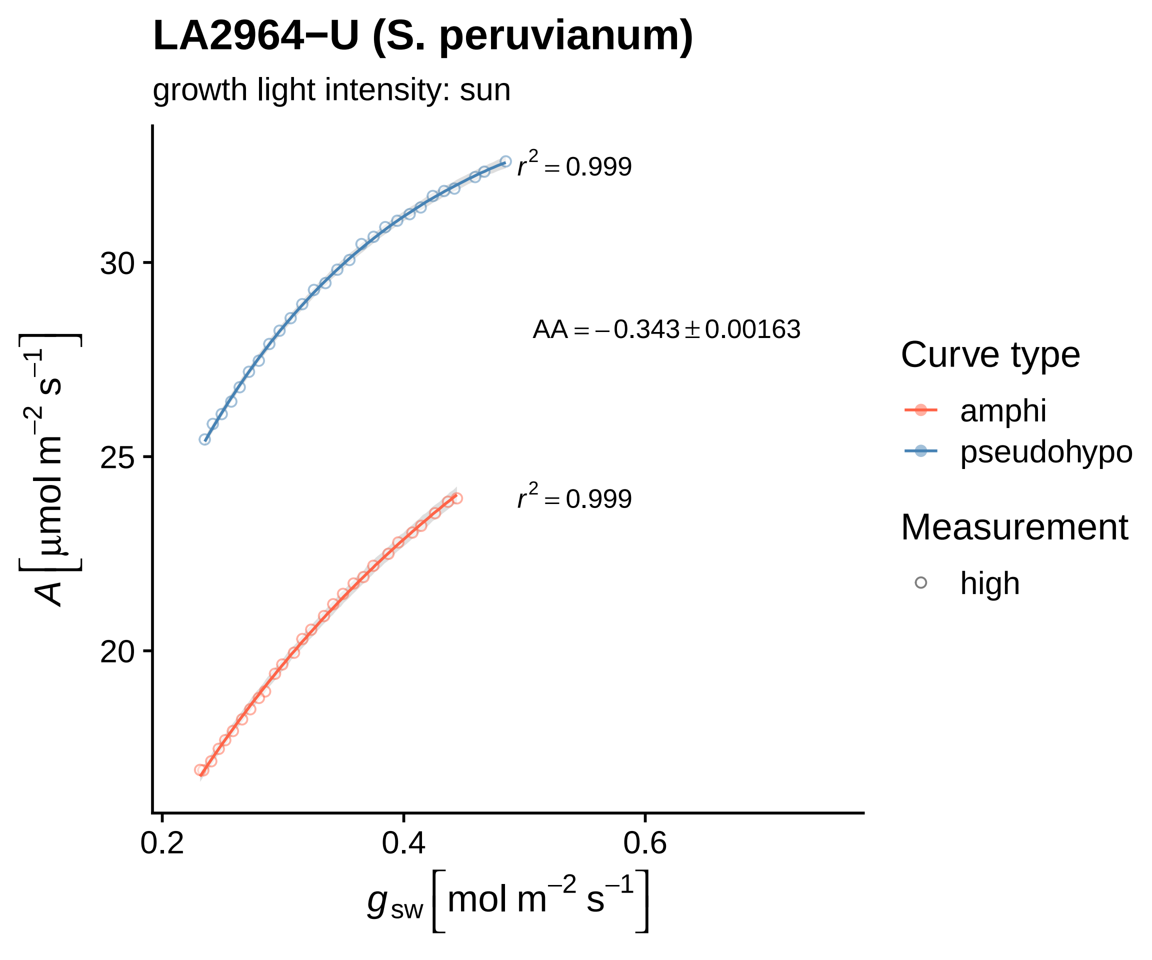
LA2964−U (S. peruvianum)
growth light intensity: sun
$r^2 = 0.999$
$AA = -0.343 \pm 0.00163$
$r^2 = 0.999$
30
25
20
0.2
0.4
0.6
$A\,[\mu mol\ m^{-2}\ s^{-1}]$
$g_{sw}\,[mol\ m^{-2}\ s^{-1}]$
Curve type
amphi
pseudohypo
Measurement
high

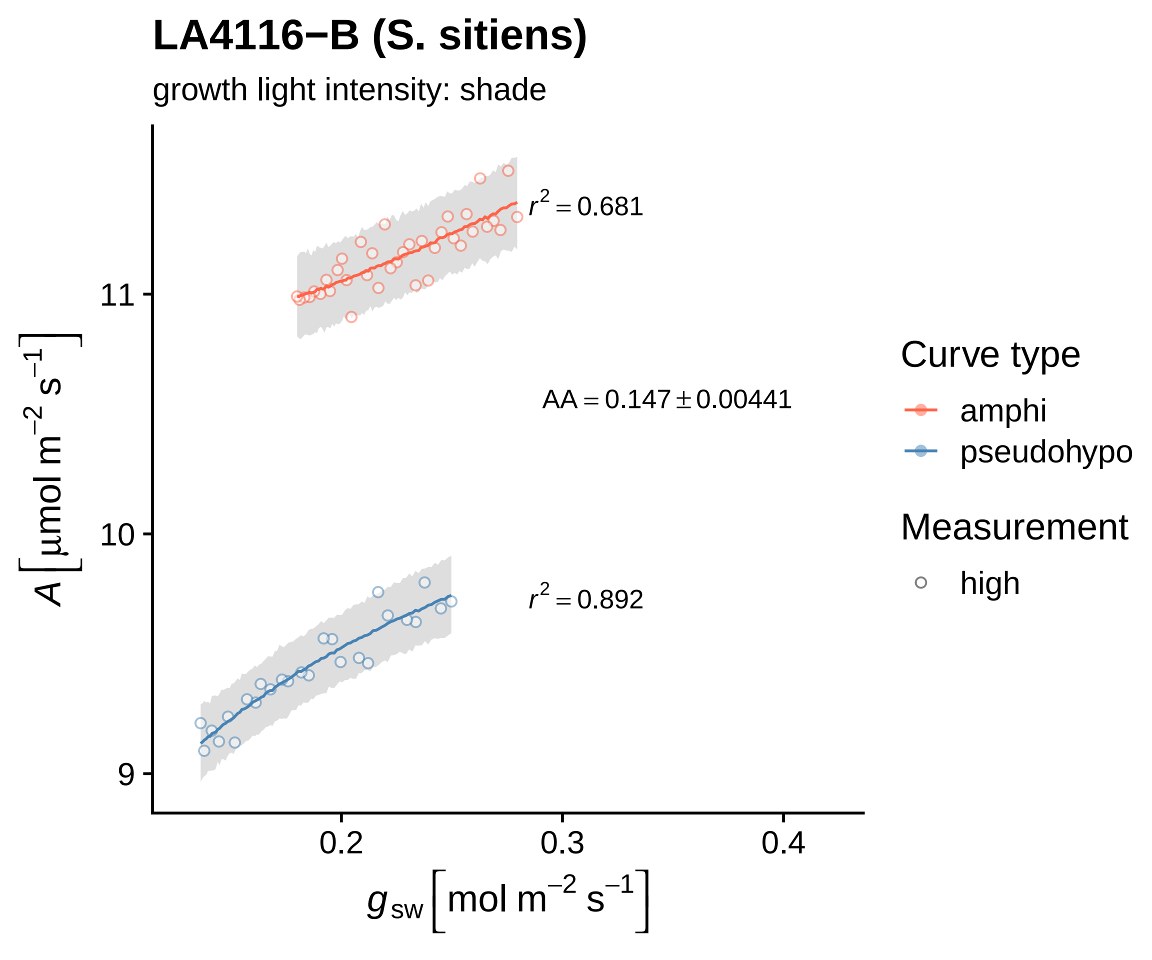
LA4116−B (S. sitiens)
growth light intensity: shade
$r^2 = 0.681$
$AA = 0.147 \pm 0.00441$
$r^2 = 0.892$
11
10
9
0.2
0.3
0.4
$A\,[\mu mol\ m^{-2}\ s^{-1}]$
$g_{sw}\,[mol\ m^{-2}\ s^{-1}]$
Curve type
amphi
pseudohypo
Measurement
high

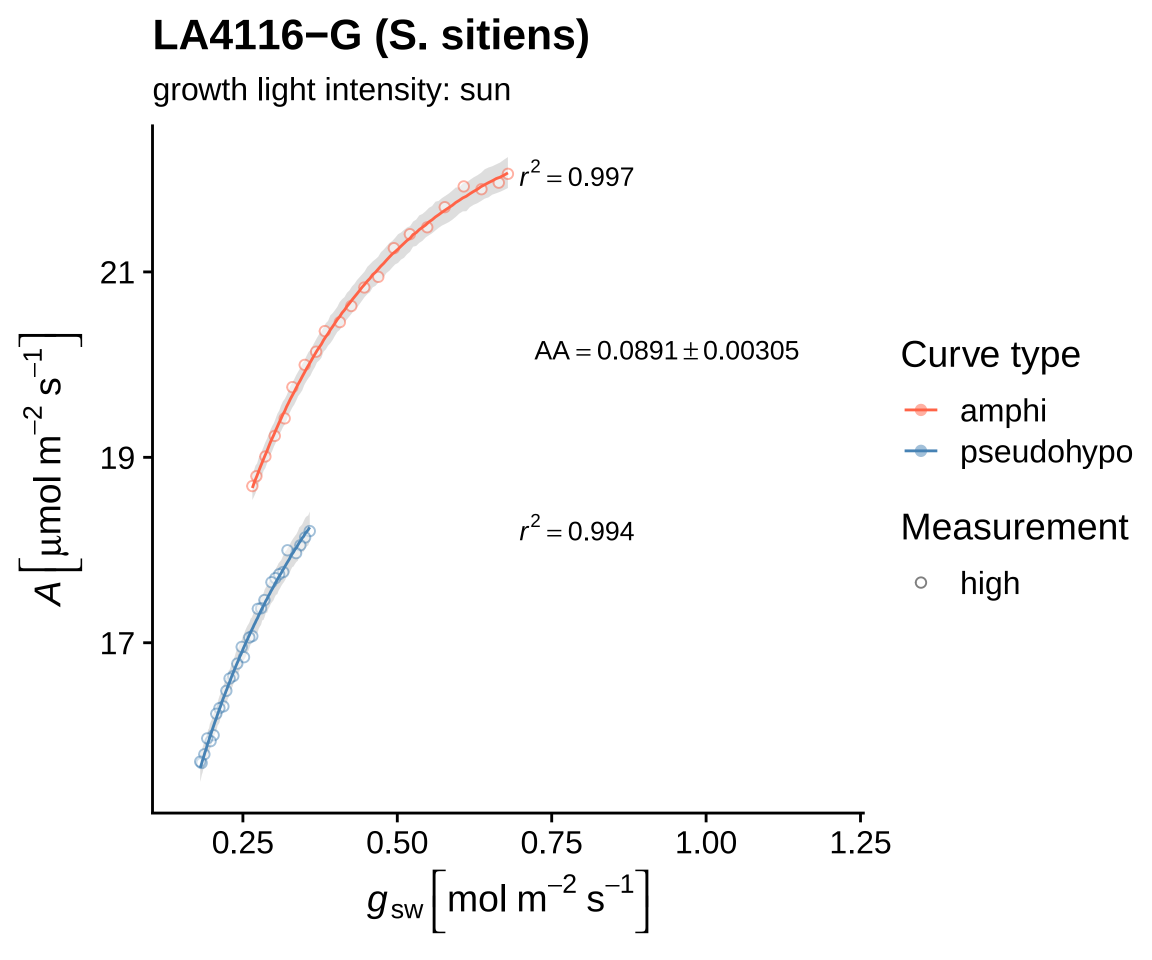
LA4116−G (S. sitiens)
growth light intensity: sun
$r^2 = 0.997$
$AA = 0.0891 \pm 0.00305$
$r^2 = 0.994$
21
19
17
$A\left[\mu mol\ m^{-2}\ s^{-1}\right]$
0.25
0.50
0.75
1.00
1.25
$g_{sw}\left[mol\ m^{-2}\ s^{-1}\right]$
Curve type
amphi
pseudohypo
Measurement
high

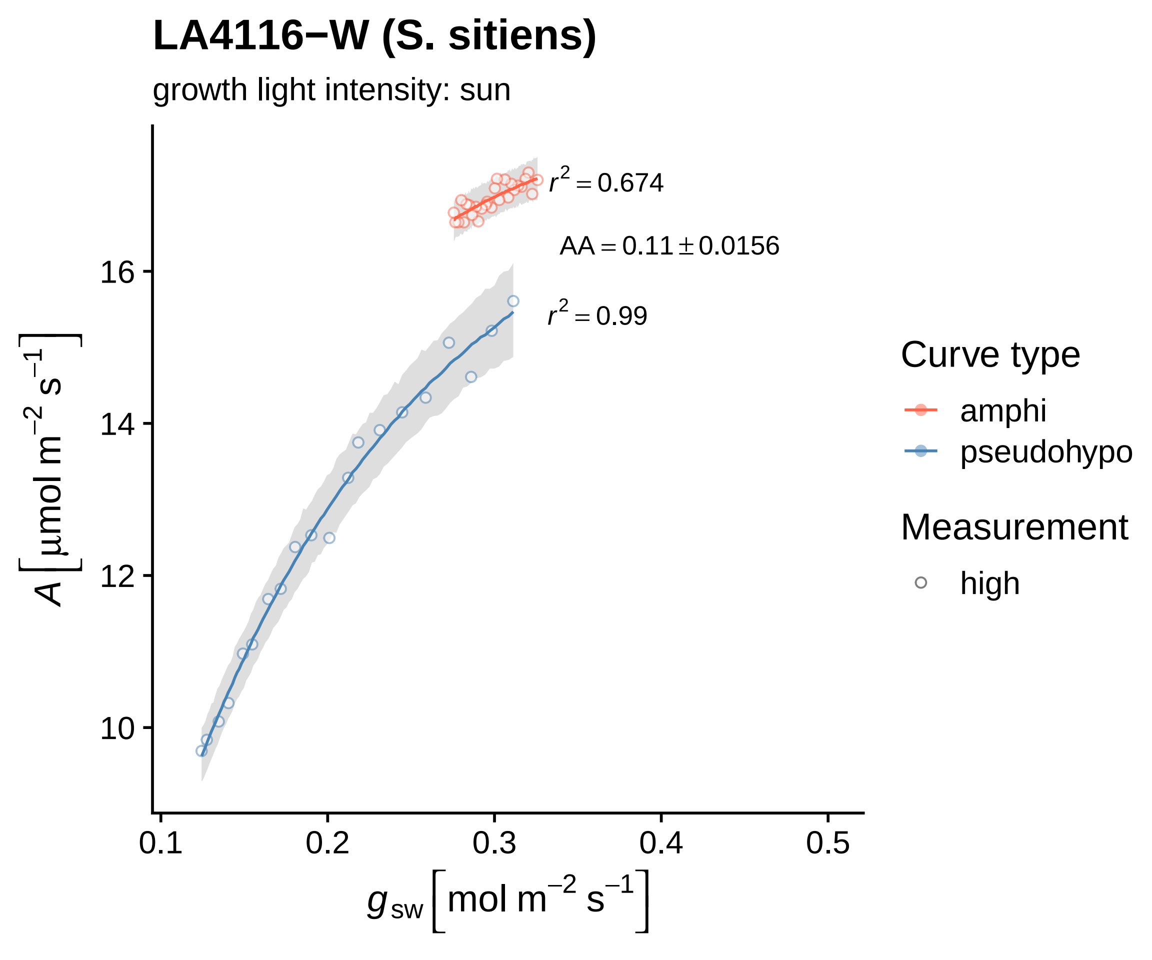

LA4116−W (S. sitiens)
growth light intensity: sun
$r^2 = 0.674$
$AA = 0.11 \pm 0.0156$
$r^2 = 0.99$
16
14
12
10
$A \left[ \mu mol\ m^{-2}\ s^{-1} \right]$
0.1
0.2
0.3
0.4
0.5
$g_{sw} \left[ mol\ m^{-2}\ s^{-1} \right]$
Curve type
amphi
pseudohypo
Measurement
high

### LA4117A−I (S. chilense)

growth light intensity: sun

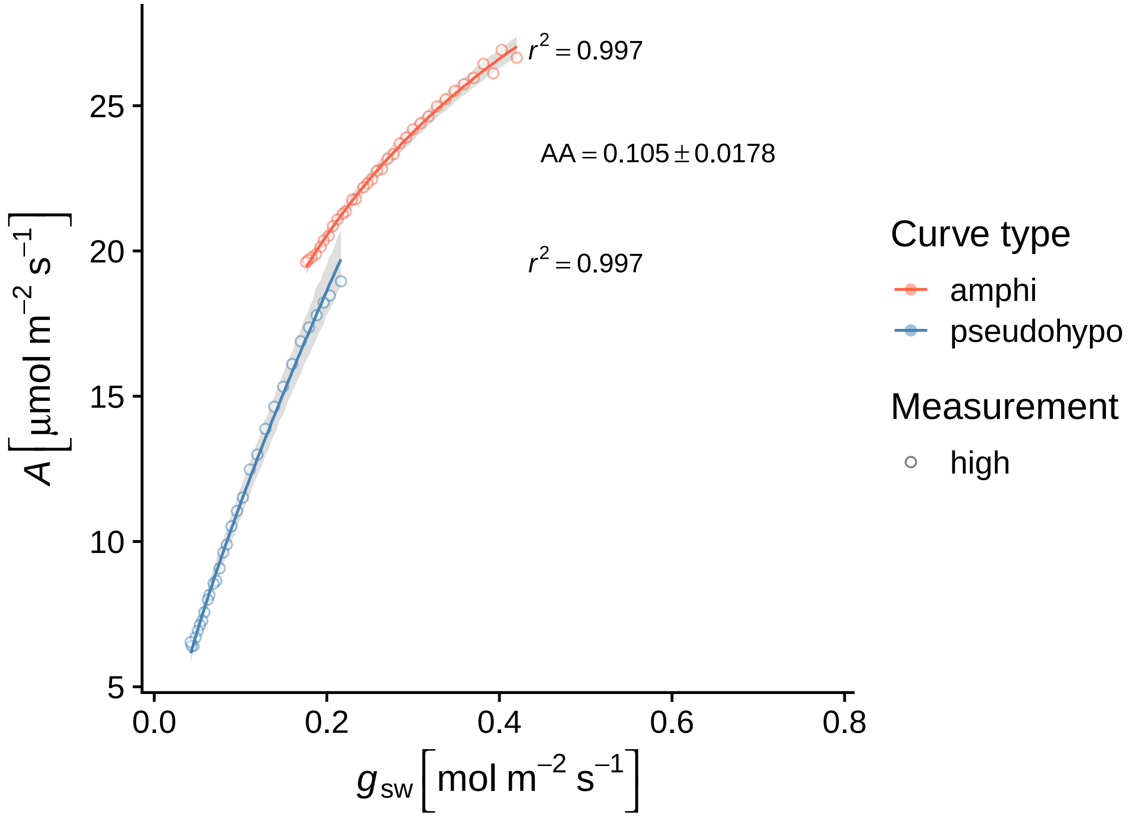

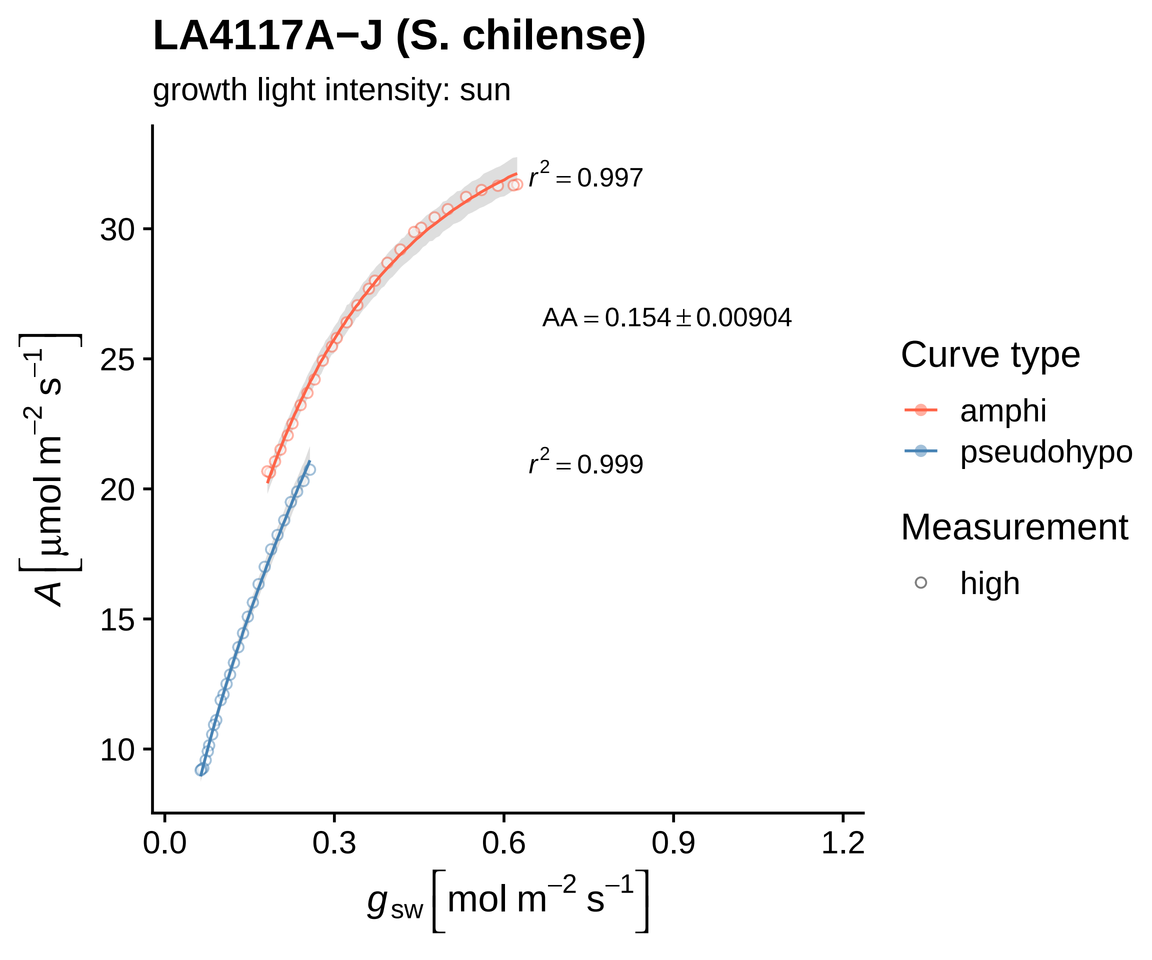
LA4117A−J (S. chilense)
growth light intensity: sun
$r^2 = 0.997$
$AA = 0.154 \pm 0.00904$
$r^2 = 0.999$
Curve type
amphi
pseudohypo
Measurement
high
30
25
20
15
10
0.0
0.3
0.6
0.9
1.2
$A\left[\mu mol\ m^{-2}\ s^{-1}\right]$
$g_{sw}\left[mol\ m^{-2}\ s^{-1}\right]$
